# Detection and Phylogenetic Analysis of Highly Pathogenic A/H5N1 Avian Influenza Clade 2.3.4.4b Virus in Chile, 2022

**DOI:** 10.1101/2023.02.01.526205

**Authors:** Pedro Jimenez-Bluhm, Jurre Y. Siegers, Shaoyuan Tan, Bridgett Sharp, Pamela Freiden, Katherinne Orozco, Soledad Ruiz, Cecilia Baumberger, Pablo Galdames, Maria Antonieta Gonzalez, Camila Rojas, Erik A. Karlsson, Christopher Hamilton-West, Stacey Schultz-Cherry

**Author notes:** **Address for Correspondence**, Pedro Jiménez-Bluhm, Stacey Schultz-Cherry.

## Abstract

Since the emergence of the Highly Pathogenic Avian Influenza H5N1 goose/Guangdong (Gs/GD) lineage in China in 1996, these viruses have spread globally. This includes the recent introduction of the Gs/GD lineage 2.3.4.4b in the Americas in 2021, which has affected both wild and domestic bird populations causing high mortality. Since November 2022, reports of mortality in wild birds in numerous South American countries along the Pacific Migratory Flyway have been attributed to this lineage. Through an ongoing longitudinal avian influenza virus (AIV) surveillance, we determined that AIV prevalence remained below 1% in the Lluta river wetland from August through October 2022. However, there was a significant increase in prevalence to 2.6% and 11.7% in November and December respectively, coinciding with the arrival of migratory birds from the Northern Hemisphere. Of the AIV RT-qPCR positive environmental feces samples 7 were identified as A/H5. Sequencing of the COI gene demonstrated that the H5 positive samples were obtained from Peruvian pelican (n=1), Franklin’s gull (n=1), Gray gull (n=1), Elegant tern (n=2) and Black skimmer (n=2). Full genomes were obtained from 3 samples and the putative cleavage site composition of the HA was polybasic REKRRKR/GLF for all. Phylogenetic analysis of the samples revealed them to belong to the H5 2.3.4.4b clade, closely related to isolates obtained in Peru in late November. The emergence of H5 clade 2.3.4.4b in South America has an immediate impact on public health, animal production and wildlife, hence increased active surveillance is warranted.

## MAIN TEXT

Highly pathogenic avian influenza (HPAI) viruses remain a major threat to both animal and human health. Since the emergence of the HPAI A/H5N1 goose/Guangdong (Gs/GD) lineage in 1996, these viruses have spread globally. This includes the recent introduction and reassortment of lineage 2.3.4.4b into the Americas in 2021, impacting both wild and domestic bird populations with high mortality [1]. Since November 2022, reports of mortality in wild birds in numerous South American countries along the Pacific Migratory Flyway have been attributed to HPAI A/H5Nx viruses. These same countries have also reported infections in domestic bird populations. However, genomic data for the viruses entering South American remains sparse. Here we report an initial genomic characterization of A/H5N1 Clade 2.3.4.4b viruses detected in wild birds in Chile.

On 7^th^ December 2022, the Agricultural and Livestock Service of Chile (SAG) reported the first confirmed case of HPAI A/H5N1 infection in Peruvian pelicans (*Pelecanus thagus*) at the Lluta River wetland located 10 km south of the Chilean-Peruvian border. We have been conducting active, longitudinal avian influenza (AIV) surveillance in Chile since 2015, with a focus on wild birds and high-risk interfaces with domestic poultry and humans. In anticipation of 2.3.4.4 encroachment into South America, we intensified our ongoing AIV surveillance in the Lluta river estuary (S 18° 24’ 59.306”; W 70° 19’ 20.668) to biweekly sampling since August 2022. AIV prevalence (as measured by M gene specific qRT-PCR) remained below 1% in August and September 2022. However, prevalence jumped to 2.6% and 11.7% in November and December, respectively, coinciding with the arrival of migratory birds from the Northern Hemisphere. Of the 69 AIV positive environmental samples from November and December (total samples: 2023), 7 (10.1%) were identified as hemagglutinin (HA) subtype A/H5: 1 obtained in late November, and 6 in December. As per regulations, original samples and RNA were submitted to SAG (submission #230014). Cytochrome Oxidase I speciation [2] shows the A/H5 positive samples come from several species: Peruvian pelican (*Pelecanus thagus*) (n=1), Franklin’s gull (*Larus pipixcan*) (n=1), Gray gull (*Leucophaeus modestus*) (n=1), Elegant tern (*Thalasseus elegans*) (n=2) and Black skimmer (*Rynchops niger*) (n=2).

Full genome sequencing of 3 of the samples yielded 2 complete genomes (A/Grey gull/Chile/61947/2022 and A/Black skimmer/Chile/61962/2022) and one partial genome (A/Peruvian pelican/Chile/61740/2022) missing segments 2 and 4 due to low viral load. The A/H5 HA genes were closest to Peruvian pelican and chicken samples (EPI_ISL_16249730, EPI_ISL_16249681 and EPI_ISL_16249274) detected in Peru around the same period, and both Peruvian and Chilean strains derive from A/H5N1 viruses causing widespread outbreaks in poultry and wild birds across North America [3] (Figure 1). Chilean A/H5N1 HA sequences have 99.9%/99.5% similarity in nucleotide/amino acid identity to each other, respectively, and 98.1%/98.9% nucleotide/amino acid similarity relative to the A/H5N1 clade 2.3.4.4b candidate vaccine strain, A/Astrakhan/3212/2020 (Supplemental Table 1). HA mutations 101M, 112Q, 200A and 486V (H3 numbering) were shared across both Chilean A/H5N1 HA segments (Supplemental Table 1). These mutations do not correlate with any previously reported phenotypic traits associated with increased risk [4]. Consistent with 2.3.4.4b HA genes, Chilean A/H5N1 viruses contain the HPAI polybasic cleavage site motif, REKRRKR|GLF. Likewise, N1 neuraminidase (NA) gene segments were closest to Peruvian pelican strains. The NA sequences do not contain any stalk deletions or markers of antiviral drug resistance (Supplemental Figure 1).

**Figure 1.**
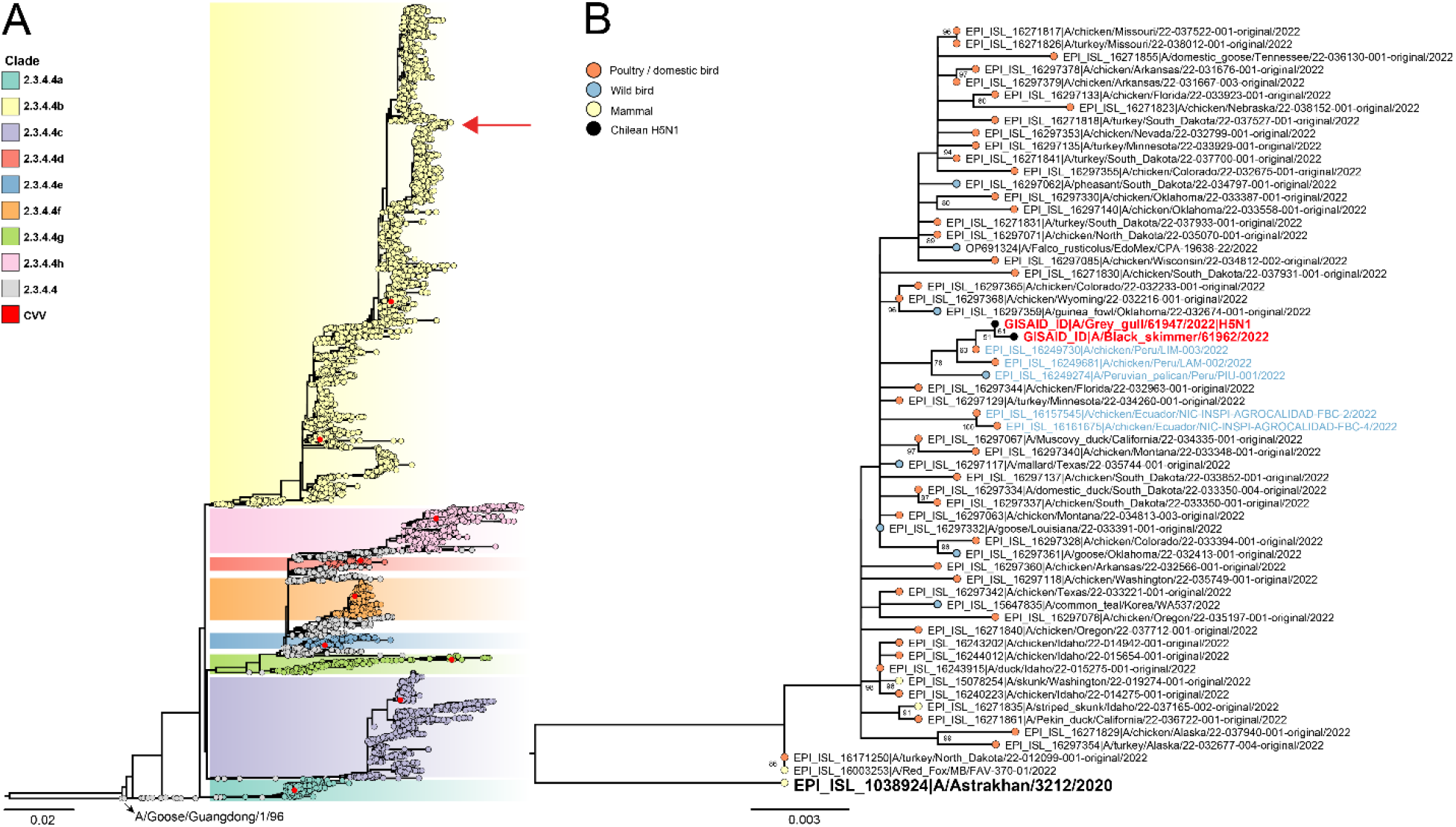
Maximum likelihood phylogeny of the HA genes of H5N1 viruses detected in Chile, 2022. **A**, Full tree of subclades of A/H5Nx clade 2.3.4.4. Candidate vaccine viruses are shown in red circles. Tip colors indicate subclades of 2.3.4.4 avian H5Nx viruses. Location of Chile strains is indicated with a red arrow. **B**, A/H5N1 phylogeny of 2.3.4.4b clade containing Chilean A/H5N1 viruses and closest relatives. Chilean viruses are highlighted in red with black tip labels. Strains from Central/South America are highlighted with light blue text. Candidate vaccine virus is shown in bold. Tip colors indicate cases of mammalian (pale yellow), wild bird (light blue), and poultry (pale red) infection with avian H5Nx viruses.

Emergence of A/H5 2.3.4.4b in South America poses a major risk, as exemplified by the catastrophic impacts to the domestic poultry and wildlife sectors in Europe and North America since 2021 [5]. There is limited experience in South America on management and containment of AIV, possibly due to minimal HPAI in the region (Chile, 2002)[6]. For the moment, regional A/H5 outbreaks/detections are limited to shorebirds, backyard poultry, and a few commercial farms. However, prevalence is likely grossly underestimated, constrained by the sheer size of the affected territory, economic factors, a dearth of exhaustive AIV surveillance, diagnostics, and adequately trained veterinary personnel to identify the disease. There is also a lack of incentive to report due to insufficient economic compensation mechanisms for culling of exposed poultry throughout the region. Considering the large, socio-economically vulnerable population in South America, mass culling of exposed poultry could also greatly reduce access to critical animal protein sources and promote food insecurity, further complicating eradication plans. Aside from domestic poultry, A/H5 2.3.4.4b in South America will also likely affect entire populations of already vulnerable wild bird species. Over 22,000 wild birds were lost in Peru to A/H5 infection between late November and early December 2022, many of them already considered endangered in the country [3].

Enhanced surveillance in both domestic and wild birds is critical in Chile and across South America. HPAI A/H5Nx viruses in South America have only been detected in shorebirds and Charadriiformes hosts so far, likely due to passive detection systems. Active, longitudinal surveillance, especially in known AIV hotspots [7,8] should also focus on Anseriformes, given year long AIV circulation and genetic diversity in these species in Chile [8]. Further sequencing of A/H5Nx and non-A/H5 positive samples is critical to identify possible reassortment events with locally circulating AIV strains.

## ACKNOWLEDGMENTS

We would like to thank the Municipality of Arica for facilitating sampling at the Lluta river wetland Municipal natural reserve. The investigators thank all the authors and laboratories who contribute genetic sequence data via GISAID. A complete list of GISAID acknowledgements is provided in Supplementary Data 1.

## APPEDICIES

Supplemental Figure 1 - Neuraminidase phylogeny

Supplemental Table 1 - Percent homology of nucleotides and amino acids

Supplemental Table 2 - Amino acid mutations in hemagglutinin

Supplemental Data 1 – GISAID/NCBI acknowledgement tables

**Supplemental Figure 1.**
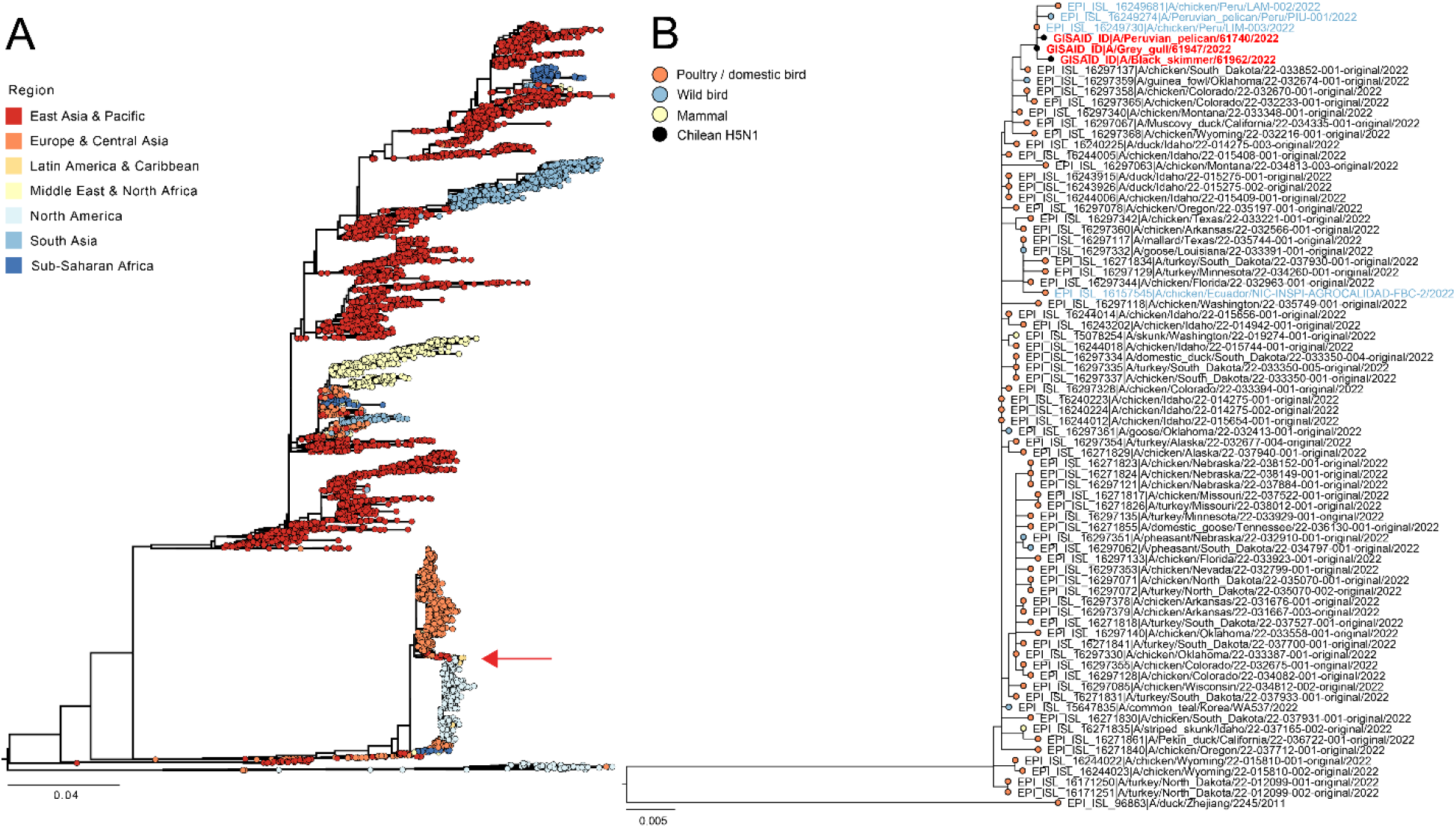
Maximum likelihood phylogeny of the NA genes of H5N1 viruses detected in Chile, 2022. **A**, Full tree of H5N1 NA sequences. Tip colors indicate the geographic region of H5N1 viruses. Location of Chile strains is indicated with a red arrow. **B**, A/H5N1 NA phylogeny containing Chilean A/H5N1 viruses and closest relatives. Chilean viruses are highlighted in red with black tip labels. Strains from Central/South America are highlighted with light blue text. Tip colors indicate cases of mammalian (pale yellow), wild bird (light blue), and poultry (pale red) infection with avian H5Nx viruses.

**Supplemental Table 1.** Percent homology of nucleotides and amino acids between CVV, Chilean and closely related Peruvian H5N1 strains.

| **Percent nucleotide identity HA** |  |  |  |  |  |
| --- | --- | --- | --- | --- | --- |
|  | A/Astrakhan/3212/2020 | A/chicken/Peru/LA M-002/2022 | A/chicken/Peru/LI M-003/2022 | A/Grey_gull/Chile/61947/2022 | A/Black_skimmer/Chile/61962/2022 |
| A/Astrakhan/3212/2020 |  | **97,9** | **97,9** | **98** | **98,1** |
| A/chicken/Peru/LA M-002/2022 | **97,9** |  | **99,8** | **99,8** | **99,7** |
| A/chicken/Peru/LI M-003/2022 | **97,9** | **99,8** |  | **99,9** | **99,9** |
| A/Grey_gull/Chile/61947/2022 | **98** | **99,8** | **99,9** |  | **99,9** |
| A/Black_skimmer/Chile/61962/2022 | **98,1** | **99,7** | **99,9** | **99,9** |  |
| **Percent amino acid identity HA** |  |  |  |  |  |
|  | A/Astrakhan/3212/2020 | A/chicken/Peru/LA M-002/2022 | A/chicken/Peru/LI M-003/2022 | A/Grey_gull/Chile/61947/2022 | A/Black_skimmer/Chile/61962/2022 |
| A/Astrakhan/3212/2020 |  | **99,3** | **99,3** | **98,8** | **98,9** |
| A/chicken/Peru/LA M-002/2022 | **99,3** |  | **100** | **99,5** | **99,7** |
| A/chicken/Peru/LI M-003/2022 | **99,3** | **100** |  | **99,5** | **99,7** |
| A/Grey_gull/Chile/61947/2022 | **98,8** | **99,5** | **99,5** |  | **99,5** |
| A/Black_skimmer/Chile/61962/2022 | **98,9** | **99,7** | **99,7** | **99,5** |  |

**Supplemental Table 2.** Amino acid mutations in hemagglutinin relative to the reference strain A/Astrakhan/3212/2020 in clade 2.3.4.4b avian influenza A/H5N1 viruses detected in Chile, 2022.

| **H5 clade 2.3.4.4b strain** | **HA amino acid position†** |  |  |  |
| --- | --- | --- | --- | --- |
|  | 101[104] | 112[115] | 200[206] | 486[497] |
| A/Astrakhan/3212/2020 (CVV)‡ | L | L | V | I |
| A/Black_skimmer/Chile/61962/2022 | M | Q | A | V |
| A/Grey_gull/Chile/61947/2022 | M | Q | A | V |

HA, hemagglutinin.
†Amino acids were numbered according to the hemagglutinin H3 and [H5] numbering system.
‡Candidate vaccine virus reference strain (GISAID accession no. EPI\_ISL\_1038924; https://www.gisaid.org).

## REFERENCES

1. Bevins SN, Shriner SA, Cumbee Jr JC, et al. Intercontinental movement of highly pathogenic avian influenza A (H5N1) clade 2.3. 4.4 virus to the United States, 2021. Emerging infectious diseases. 2022;28(5):1006.

2. Cheung PP, Leung YC, Chow C-K, et al. Identifying the species-origin of faecal droppings used for avian influenza virus surveillance in wild-birds. Journal of clinical virology. 2009;46(1):90–93.

3. Gamarra-Toledo V, Plaza PI, Gutiérrez R, et al. Avian flu threatens Neotropical birds. Science. 2023;379(6629):246–246.

4. Suttie A, Deng Y-M, Greenhill AR, et al. Inventory of molecular markers affecting biological characteristics of avian influenza A viruses. Virus Genes. 2019;55:739–768.

5. Stokstad E. Deadly flu spreads through North American birds. Science. 2022;376(6592):441–442.

6. Spackman E, McCracken KG, Winker K, et al. H7N3 avian influenza virus found in a South American wild duck is related to the Chilean 2002 poultry outbreak, contains genes from equine and North American wild bird lineages, and is adapted to domestic turkeys. Journal of virology. 2006;80(15):7760–7764.

7. Ruiz S, Jimenez-Bluhm P, Di Pillo F, et al. Temporal dynamics and the influence of environmental variables on the prevalence of avian influenza virus in main wetlands in central Chile. Transboundary and emerging diseases. 2021;68(3):1601–1614.

8. Jiménez-Bluhm P, Karlsson EA, Freiden P, et al. Wild birds in Chile Harbor diverse avian influenza A viruses. Emerging Microbes & Infections. 2018;7(1):1–4.

