## Supplemental Data 1 for "Detection and Phylogenetic Analysis of Highly Pathogenic A/H5N1 Avian Influenza Clade 2.3.4.4b Virus in Chile, 2022"

### Supplemental Data 1. NCBI and GISAD acknowledgement

### NCBI:

| accession | length | host | segment | serotype | country | region | date | name | mutations | age | gender | lineage | vac_strain | fulllength_plus |
|---|---|---|---|---|---|---|---|---|---|---|---|---|---|---|
| AB932556 | 1742 | Avian | 4 (HA) | H5N8 | Japan | N | 2014/04/13 | Influenza A virus (A/chicken/kumamoto/1-7/2014(H5N8)) | | | | | | c |
| AB979455 | 1742 | Avian | 4 (HA) | H5N1 | Viet Nam | T | 2014/04/15 | Influenza A virus (A/muscovy duck/Vietnam/LBM631/2014(H5N1)) | | | | | | c |
| AB979463 | 1742 | Avian | 4 (HA) | H5N1 | Viet Nam | T | 2014/04/15 | Influenza A virus (A/duck/Vietnam/LBM632/2014(H5N1)) | | | | | | c |
| AB979471 | 1742 | Avian | 4 (HA) | H5N1 | Viet Nam | T | 2014/04/15 | Influenza A virus (A/duck/Vietnam/LBM633/2014(H5N1)) | | | | | | c |
| AB979479 | 1742 | Avian | 4 (HA) | H5N1 | Viet Nam | T | 2014/04/15 | Influenza A virus (A/muscovy duck/Vietnam/LBM634/2014(H5N1)) | | | | | | c |
| AB979487 | 1734 | Avian | 4 (HA) | H5N1 | Viet Nam | T | 2014/04/15 | Influenza A virus (A/muscovy duck/Vietnam/LBM635/2014(H5N1)) | | | | | | c |
| AB979495 | 1741 | Avian | 4 (HA) | H5N1 | Viet Nam | T | 2014/04/15 | Influenza A virus (A/muscovy duck/Vietnam/LBM636/2014(H5N1)) | | | | | | c |
| AB979503 | 1740 | Avian | 4 (HA) | H5N1 | Viet Nam | T | 2014/04/15 | Influenza A virus (A/duck/Vietnam/LBM638/2014(H5N1)) | | | | | | c |
| AB979511 | 1742 | Avian | 4 (HA) | H5N1 | Viet Nam | T | 2014/04/15 | Influenza A virus (A/duck/Vietnam/LBM639/2014(H5N1)) | | | | | | c |
| KJ508916 | 1742 | Avian | 4 (HA) | H5N8 | South Korea | N | 2014/01/19 | Influenza A virus (A/broiler duck/Korea/H32/2014(H5N8)) | | | | | | c |
| KP090439 | 1776 | Avian | 4 (HA) | H5N6 | China | N | 2014/06/29 | Influenza A virus (A/chicken/Jiangxi/NCDZT1123/2014(H5N6)) | | | | | | c |
| KP090447 | 1776 | Avian | 4 (HA) | H5N6 | China | N | 2014/06/29 | Influenza A virus (A/chicken/Jiangxi/NCDZT1126/2014(H5N6)) | | | | | | c |
| KP714479 | 1776 | Avian | 4 (HA) | H5N8 | Taiwan | N | 2015/01/09 | Influenza A virus (A/goose/Taiwan/01003/2015(H5N8)) | | | | | | c |
| KP714480 | 1776 | Avian | 4 (HA) | H5N2 | Taiwan | N | 2015/01/09 | Influenza A virus (A/goose/Taiwan/01004/2015(H5N2)) | | | | | | c |
| KP714481 | 1776 | Avian | 4 (HA) | H5N3 | Taiwan | N | 2015/01/12 | Influenza A virus (A/goose/Taiwan/01042/2015(H5N3)) | | | | | | c |
| KP732644 | 1704 | Avian | 4 (HA) | H5N6 | China | N | 2014/03/22 | Influenza A virus (A/duck/Eastern China/S0322/2014(H5N6)) | | | | | | c |
| KP851843 | 1743 | Avian | 4 (HA) | H5N8 | South Korea | N | 2014/02/15 | Influenza A virus (A/baikal teal/Korea/K14-E016/2014(H5N8)) | | | | | | c |
| KT280932 | 1744 | Avian | 4 (HA) | H5N2 | USA | N | 2015/04/27 | Influenza A virus (A/chicken/Iowa/21981/2015(H5N2)) | | | | | | c |
| KT327395 | 1776 | Avian | 4 (HA) | H5N2 | Taiwan | N | 2015/01/19 | Influenza A virus (A/night heron/Taiwan/A289/2015(H5N2)) | | | | | | c |
| KT327403 | 1776 | Avian | 4 (HA) | H5N2 | Taiwan | N | 2015/01/17 | Influenza A virus (A/thrush/Taiwan/A234/2015(H5N2)) | | | | | | c |
| KT327411 | 1776 | Avian | 4 (HA) | H5N3 | Taiwan | N | 2015/01/15 | Influenza A virus (A/Bulbul/Taiwan/01156/2015(H5N3)) | | | | | | c |
| KT370058 | 1704 | Environment | 4 (HA) | H5N6 | China | N | 2014/05/06 | Influenza A virus (A/environment/Guangdong/QY208/2014(H5N6)) | | | | | | c |
| KT370059 | 1704 | Environment | 4 (HA) | H5N6 | China | N | 2013/04/10 | Influenza A virus (A/environment/Guangdong/QY025/2013(H5N6)) | | | | | | c |
| KT370060 | 1704 | Environment | 4 (HA) | H5N6 | China | N | 2014/05/06 | Influenza A virus (A/environment/Guangdong/QY197/2014(H5N6)) | | | | | | c |
| KT370061 | 1704 | Environment | 4 (HA) | H5N6 | China | N | 2014/07/17 | Influenza A virus (A/environment/Guangdong/ZS356/2014(H5N6)) | | | | | | c |
| KT370062 | 1704 | Environment | 4 (HA) | H5N6 | China | N | 2014/12/20 | Influenza A virus (A/environment/Guangdong/PY955/2014(H5N6)) | | | | | | c |
| KT370063 | 1704 | Environment | 4 (HA) | H5N6 | China | N | 2015/02/23 | Influenza A virus (A/environment/Guangdong/HY243/2015(H5N6)) | | | | | | c |
| KT370064 | 1704 | Environment | 4 (HA) | H5N6 | China | N | 2014/03/02 | Influenza A virus (A/environment/Guangdong/JY137/2014(H5N6)) | | | | | | c |
| KT936689 | 1619 | Avian | 4 (HA) | H5N8 | China | N | 2014/12/15 | Influenza A virus (A/Eurasian curlew/Shanghai/DT1215-1/2014(H5N8)) | | | | | | p |
| KT936690 | 1587 | Avian | 4 (HA) | H5N8 | China | N | 2014/12/31 | Influenza A virus (A/Eurasian curlew/Shanghai/DT1231-1/2014(H5N8)) | | | | | | p |
| KT936691 | 1585 | Avian | 4 (HA) | H5N8 | China | N | 2014/12/31 | Influenza A virus (A/Eurasian curlew/Shanghai/DT1231-2/2014(H5N8)) | | | | | | p |
| KT936692 | 1598 | Avian | 4 (HA) | H5N8 | China | N | 2014/12/31 | Influenza A virus (A/Eurasian curlew/Shanghai/DT1231-3/2014(H5N8)) | | | | | | p |
| KT936693 | 1586 | Avian | 4 (HA) | H5N8 | China | N | 2014/12/31 | Influenza A virus (A/Eurasian curlew/Shanghai/DT1231-4/2014(H5N8)) | | | | | | p |
| KT936694 | 1598 | Avian | 4 (HA) | H5N8 | China | N | 2014/12/31 | Influenza A virus (A/Eurasian curlew/Shanghai/DT1231-5/2014(H5N8)) | | | | | | p |
| KT936695 | 1590 | Avian | 4 (HA) | H5N8 | China | N | 2014/12/31 | Influenza A virus (A/Eurasian curlew/Shanghai/DT1231-6/2014(H5N8)) | | | | | | p |
| KX121198 | 1704 | Avian | 4 (HA) | H5N6 | China | N | 2014/11/12 | Influenza A virus (A/goose/Hunan/118/2014(H5N6)) | | | | | | c |
| KX121206 | 1704 | Avian | 4 (HA) | H5N6 | China | N | 2014/11/13 | Influenza A virus (A/duck/Hunan/144/2014(H5N6)) | | | | | | c |
| KX297862 | 1704 | Environment | 4 (HA) | H5N8 | South Korea | N | 2014/02/06 | Influenza A virus (A/environment/Korea/W454/2014(H5N8)) | | | | | | c |
| KX297863 | 1704 | Avian | 4 (HA) | H5N8 | South Korea | N | 2014/02/14 | Influenza A virus (A/mallard duck/Korea/W456/2014(H5N8)) | | | | | | c |
| KX297864 | 1704 | Avian | 4 (HA) | H5N8 | South Korea | N | 2014/02/14 | Influenza A virus (A/mallard duck/Korea/W457/2014(H5N8)) | | | | | | c |
| KX297865 | 1704 | Environment | 4 (HA) | H5N8 | South Korea | N | 2014/02/18 | Influenza A virus (A/environment/Korea/W458/2014(H5N8)) | | | | | | c |
| KX297866 | 1704 | Environment | 4 (HA) | H5N8 | South Korea | N | 2014/12/21 | Influenza A virus (A/environment/Korea/W464/2014(H5N8)) | | | | | | c |
| KX297867 | 1704 | Environment | 4 (HA) | H5N8 | South Korea | N | 2014/12/21 | Influenza A virus (A/environment/Korea/W465/2014(H5N8)) | | | | | | c |
| KX297868 | 1704 | Environment | 4 (HA) | H5N8 | South Korea | N | 2014/12/21 | Influenza A virus (A/environment/Korea/W466/2014(H5N8)) | | | | | | c |
| KX297869 | 1704 | Environment | 4 (HA) | H5N8 | South Korea | N | 2014/12/21 | Influenza A virus (A/environment/Korea/W467/2014(H5N8)) | | | | | | c |
| KX297870 | 1704 | Environment | 4 (HA) | H5N8 | South Korea | N | 2014/12/21 | Influenza A virus (A/environment/Korea/W468/2014(H5N8)) | | | | | | c |
| KX297871 | 1704 | Environment | 4 (HA) | H5N8 | South Korea | N | 2014/12/21 | Influenza A virus (A/environment/Korea/W469/2014(H5N8)) | | | | | | c |
| KX297872 | 1704 | Environment | 4 (HA) | H5N8 | South Korea | N | 2014/12/21 | Influenza A virus (A/environment/Korea/W470/2014(H5N8)) | | | | | | c |
| KX297873 | 1704 | Environment | 4 (HA) | H5N8 | South Korea | N | 2014/12/21 | Influenza A virus (A/environment/Korea/W471/2014(H5N8)) | | | | | | c |
| KX297874 | 1704 | Environment | 4 (HA) | H5N8 | South Korea | N | 2014/12/21 | Influenza A virus (A/environment/Korea/W472/2014(H5N8)) | | | | | | c |
| KX297875 | 1704 | Environment | 4 (HA) | H5N8 | South Korea | N | 2014/12/21 | Influenza A virus (A/environment/Korea/W473/2014(H5N8)) | | | | | | c |
| KX297876 | 1704 | Environment | 4 (HA) | H5N8 | South Korea | N | 2014/12/21 | Influenza A virus (A/environment/Korea/W474/2014(H5N8)) | | | | | | c |
| KX297877 | 1704 | Environment | 4 (HA) | H5N8 | South Korea | N | 2014/12/21 | Influenza A virus (A/environment/Korea/W475/2014(H5N8)) | | | | | | c |
| KX297878 | 1704 | Environment | 4 (HA) | H5N8 | South Korea | N | 2014/12/21 | Influenza A virus (A/environment/Korea/W476/2014(H5N8)) | | | | | | c |
| KX297879 | 1704 | Environment | 4 (HA) | H5N8 | South Korea | N | 2014/12/21 | Influenza A virus (A/environment/Korea/W477/2014(H5N8)) | | | | | | c |
| KX297880 | 1704 | Environment | 4 (HA) | H5N8 | South Korea | N | 2015/01/14 | Influenza A virus (A/environment/Korea/W482/2015(H5N8)) | | | | | | c |
| KX297881 | 1704 | Environment | 4 (HA) | H5N8 | South Korea | N | 2015/01/14 | Influenza A virus (A/environment/Korea/W483/2015(H5N8)) | | | | | | c |
| KX297882 | 1704 | Environment | 4 (HA) | H5N8 | South Korea | N | 2015/01/21 | Influenza A virus (A/environment/Korea/W486/2015(H5N8)) | | | | | | c |
| KX297883 | 1704 | Environment | 4 (HA) | H5N8 | South Korea | N | 2015/01/21 | Influenza A virus (A/environment/Korea/W487/2015(H5N8)) | | | | | | c |
| KX297884 | 1704 | Environment | 4 (HA) | H5N8 | South Korea | N | 2015/01/21 | Influenza A virus (A/environment/Korea/W488/2015(H5N8)) | | | | | | c |
| KX297885 | 1704 | Environment | 4 (HA) | H5N8 | South Korea | N | 2015/01/21 | Influenza A virus (A/environment/Korea/W490/2015(H5N8)) | | | | | | c |
| KX297886 | 1704 | Environment | 4 (HA) | H5N8 | South Korea | N | 2015/02/27 | Influenza A virus (A/environment/Korea/W492/2015(H5N8)) | | | | | | c |
| KX960164 | 1701 | Avian | 4 (HA) | H5N6 | China | N | 2015/01/07 | Influenza A virus (A/wild bird/Jiangxi/P560/2015(H5N6)) | | | | | | c |
| KX960166 | 1704 | Avian | 4 (HA) | H5N1 | China | N | 2015/01/07 | Influenza A virus (A/wild bird/Jiangxi/P410/2015(H5N1)) | | | | | | c |
| KX960168 | 1701 | Avian | 4 (HA) | H5N6 | China | N | 2015/01/07 | Influenza A virus (A/wild bird/Jiangxi/P38/2015(H5N6)) | | | | | | c |
| KX960170 | 1701 | Avian | 4 (HA) | H5N6 | China | N | 2015/01/07 | Influenza A virus (A/wild bird/Jiangxi/P31/2015(H5N6)) | | | | | | c |
| KX960172 | 1701 | Avian | 4 (HA) | H5N6 | China | N | 2015/01/07 | Influenza A virus (A/wild bird/Jiangxi/P21/2015(H5N6)) | | | | | | c |
| KX960174 | 1701 | Avian | 4 (HA) | H5N6 | China | N | 2015/01/07 | Influenza A virus (A/wild bird/Jiangxi/P6/2015(H5N6)) | | | | | | c |
| KX960176 | 1701 | Avian | 4 (HA) | H5N6 | China | N | 2015/01/07 | Influenza A virus (A/wild bird/Jiangxi/P5/2015(H5N6)) | | | | | | c |
| KX960178 | 1701 | Avian | 4 (HA) | H5N6 | China | N | 2015/01/07 | Influenza A virus (A/wild bird/Jiangxi/P562/2015(H5N6)) | | | | | | c |
| KX960180 | 1701 | Avian | 4 (HA) | H5N6 | China | N | 2015/01/07 | Influenza A virus (A/wild bird/Jiangxi/P469/2015(H5N6)) | | | | | | c |
| KX960182 | 1701 | Avian | 4 (HA) | H5N6 | China | N | 2015/01/07 | Influenza A virus (A/wild bird/Jiangxi/P237/2015(H5N6)) | | | | | | c |
| KX960184 | 1704 | Avian | 4 (HA) | H5N1 | China | N | 2015/01/07 | Influenza A virus (A/wild bird/Jiangxi/P126/2015(H5N1)) | | | | | | c |
| KY316512 | 1701 | Avian | 4 (HA) | H5N6 | South Korea | N | 2016/11/25 | Influenza A virus (A/waterfowl/Korea/S57/2016(H5N6)) | | | | | | c |
| LC041308 | 1734 | Avian | 4 (HA) | H5N6 | Viet Nam | T | 2014/11/18 | Influenza A virus (A/duck/Vietnam/HU1-1152/2014(H5N6)) | | | | | | c |
| LC041313 | 1710 | Avian | 4 (HA) | H5N6 | Viet Nam | T | 2014/09/03 | Influenza A virus (A/duck/Vietnam/HU1-1151/2014(H5N6)) | | | | | | p |

| | | | | | | | | | |
|---|---|---|---|---|---|---|---|---|---|
| LC041317 | 1734 | Avian | 4 (HA) | H5N6 | Viet Nam | T | 2014/11/18 | Influenza A virus (A/duck/Vietnam/HU1-1507/2014(H5N6)) | c |
| LC041319 | 1738 | Avian | 4 (HA) | H5N6 | Viet Nam | T | 2014/11/18 | Influenza A virus (A/duck/Vietnam/HU1-1511/2014(H5N6)) | c |
| LC041321 | 1736 | Avian | 4 (HA) | H5N6 | Viet Nam | T | 2014/11/18 | Influenza A virus (A/duck/Vietnam/HU1-1434/2014(H5N6)) | c |
| LC041323 | 1736 | Avian | 4 (HA) | H5N6 | Viet Nam | T | 2014/11/18 | Influenza A virus (A/duck/Vietnam/HU1-1144/2014(H5N6)) | c |
| LC275037 | 1739 | Avian | 4 (HA) | H5N6 | Japan | N | 2017/01/15 | Influenza A virus (A/bean goose/Ishikawa/1701A012/2017(H5N6)) | c |
| LC275045 | 1739 | Avian | 4 (HA) | H5N6 | Japan | N | 2017/01/29 | Influenza A virus (A/bean goose/Ishikawa/1701A014/2017(H5N6)) | c |
| LC279816 | 1742 | Avian | 4 (HA) | H5N6 | Viet Nam | T | 2015/12/08 | Influenza A virus (A/duck/Vietnam/LBM837/2015(H5N6)) | c |
| LC279824 | 1734 | Avian | 4 (HA) | H5N6 | Viet Nam | T | 2015/11/14 | Influenza A virus (A/chicken/Nha Trang/122/2015(H5N6)) | c |
| LC279832 | 1721 | Avian | 4 (HA) | H5N6 | Viet Nam | T | 2015/11/14 | Influenza A virus (A/chicken/Nha Trang/124/2015(H5N6)) | c |
| LC279840 | 1742 | Avian | 4 (HA) | H5N6 | Viet Nam | T | 2015/11/14 | Influenza A virus (A/chicken/Nha Trang/127/2015(H5N6)) | c |
| LC279848 | 1727 | Avian | 4 (HA) | H5N6 | Viet Nam | T | 2015/11/14 | Influenza A virus (A/chicken/Nha Trang/128/2015(H5N6)) | c |
| LC279856 | 1728 | Avian | 4 (HA) | H5N6 | Viet Nam | T | 2015/11/14 | Influenza A virus (A/chicken/Nha Trang/129/2015(H5N6)) | c |
| LC279864 | 1742 | Avian | 4 (HA) | H5N6 | Viet Nam | T | 2016/03/04 | Influenza A virus (A/muscovy duck/Quang Ninh/254/2016(H5N6)) | c |
| LC318856 | 1760 | Avian | 4 (HA) | H5N6 | Japan | N | 2017/01/16 | Influenza A virus (A/whooper swan/Iwate/17/2017(H5N6)) | c |
| LC348834 | 1739 | Environment | 4 (HA) | H5N6 | Japan | N | 2016/12/17 | Influenza A virus (A/water/Aichi/C3/2016(H5N6)) | c |
| LC348842 | 1739 | Environment | 4 (HA) | H5N6 | Japan | N | 2016/12/17 | Influenza A virus (A/water/Aichi/W4/2016(H5N6)) | c |
| LC420034 | 1742 | Environment | 4 (HA) | H5N8 | Japan | N | 2014/12/29 | Influenza A virus (A/environment/Yamaguchi/8/2014) | c |
| LC420042 | 1742 | Environment | 4 (HA) | H5N8 | Japan | N | 2014/12/14 | Influenza A virus (A/environment/Miyazaki/11/2014) | c |
| LC536298 | 1742 | Avian | 4 (HA) | H5N6 | Viet Nam | T | 2017/12/27 | Influenza A virus (A/duck/Vietnam/LBM1042/2017) | c |
| LC536322 | 1742 | Avian | 4 (HA) | H5N6 | Viet Nam | T | 2017/10/27 | Influenza A virus (A/chicken/Nha Trang/247/2017) | c |
| LC718210 | 1773 | Avian | 4 (HA) | H5N1 | Japan | N | 2022/01/27 | Influenza A virus (A/common buzzard/Kyoto/2601B013/2022) | c |
| LC718218 | 1766 | Avian | 4 (HA) | H5N1 | Japan | N | 2022/02/08 | Influenza A virus (A/whooper swan/Iwate/0302I001T/2022) | c |
| LC718226 | 1772 | Avian | 4 (HA) | H5N1 | Japan | N | 2022/02/12 | Influenza A virus (A/white-fronted goose/Iwate/TU16-74/2022) | c |
| LC718234 | 1771 | Avian | 4 (HA) | H5N1 | Japan | N | 2022/03/04 | Influenza A virus (A/jungle crow/Iwate/0303I003/2022) | c |
| LC718242 | 1768 | Avian | 4 (HA) | H5N1 | Japan | N | 2022/03/18 | Influenza A virus (A/whooper swan/Iwate/0303B006/2022) | c |
| LC718250 | 1770 | Avian | 4 (HA) | H5N1 | Japan | N | 2022/04/06 | Influenza A virus (A/jungle crow/Iwate/0304I001/2022) | c |
| LC718258 | 1767 | Avian | 4 (HA) | H5N1 | Japan | N | 2022/04/09 | Influenza A virus (A/jungle crow/Hokkaido/0104B085/2022) | c |
| LC718266 | 1776 | Avian | 4 (HA) | H5N1 | Japan | N | 2022/04/19 | Influenza A virus (A/jungle crow/Akita/0504F001/2022) | c |
| LC718274 | 1767 | Avian | 4 (HA) | H5N1 | Japan | N | 2022/04/18 | Influenza A virus (A/jungle crow/Hokkaido/0104B087/2022) | c |
| LC718346 | 1771 | Avian | 4 (HA) | H5N1 | Japan | N | 2022/04/18 | Influenza A virus (A/chicken/Akita/TU22-31/2022) | c |
| LC718362 | 1765 | Avian | 4 (HA) | H5N1 | Japan | N | 2022/05/14 | Influenza A virus (A/chicken/Hokkaido/TU25-3/2022) | c |
| LC723835 | 1766 | Avian | 4 (HA) | H5N8 | Japan | N | 2020 | Influenza A virus (A/Unidentified_ducks/Kagoshima/NIES229/2020) | p |
| MF399540 | 1704 | Avian | 4 (HA) | H5N6 | China | N | 2015/11/11 | Influenza A virus (A/Ferruginous Pochard/Ningxia/473-8/2015(H5N6)) | c |
| MF399556 | 1704 | Avian | 4 (HA) | H5N6 | China | N | 2015/11/11 | Influenza A virus (A/Northern Shoveler/Ningxia/475-11/2015(H5N6)) | c |
| MF399564 | 1704 | Avian | 4 (HA) | H5N6 | China | N | 2015/11/11 | Influenza A virus A/Eurasian Wigeon/Ningxia/476-12/2015(H5N6) | c |
| MF399588 | 1704 | Avian | 4 (HA) | H5N6 | China | N | 2015/11/11 | Influenza A virus (A/Ferruginous Pochard/Ningxia/479-16/2015(H5N6)) | c |
| MF399596 | 1704 | Avian | 4 (HA) | H5N6 | China | N | 2015/11/11 | Influenza A virus (A/Ferruginous Pochard/Ningxia/480-17/2015(H5N6)) | c |
| MF399604 | 1704 | Avian | 4 (HA) | H5N6 | China | N | 2015/11/11 | Influenza A virus (A/Northern Shoveler/Ningxia/481-21/2015(H5N6)) | c |
| MF399620 | 1704 | Avian | 4 (HA) | H5N6 | China | N | 2015/11/11 | Influenza A virus (A/Northern Shoveler/Ningxia/483-28/2015(H5N6)) | c |
| MF399668 | 1704 | Avian | 4 (HA) | H5N6 | China | N | 2015/11/16 | Influenza A virus (A/Northern Shoveler/Ningxia/488-53/2015(H5N6)) | c |
| MF926453 | 1742 | Avian | 4 (HA) | H5N8 | Russia | N | 2017/02/28 | Influenza A virus (A/chicken/Moscow/94/2017(H5N8)) | c |
| MF926461 | 1742 | Avian | 4 (HA) | H5N8 | Russia | N | 2017/01/08 | Influenza A virus (A/chicken/Chechnya/58/2017(H5N8)) | c |
| MF926469 | 1742 | Avian | 4 (HA) | H5N8 | Russia | N | 2017/01/05 | Influenza A virus (A/swan/Krasnodar/44/2017(H5N8)) | c |
| MF926477 | 1742 | Avian | 4 (HA) | H5N8 | Russia | N | 2017/01/03 | Influenza A virus (A/swan/Voronezh/2/2017(H5N8)) | c |
| MF926485 | 1742 | Avian | 4 (HA) | H5N8 | Russia | N | 2016/11/07 | Influenza A virus (A/goose/Kalmykia/813/2016(H5N8)) | c |
| MG029170 | 1701 | Avian | 4 (HA) | H5N6 | China | N | 2016/09 | Influenza A virus (A/poultry/China/XY165.4/2016(H5N6)) | c |
| MG029172 | 1704 | Avian | 4 (HA) | H5N6 | China | N | 2016/09 | Influenza A virus (A/poultry/China/XY01.4/2016(H5N6)) | c |
| MG220416 | 1751 | Environment | 4 (HA) | H5N6 | China | N | 2017/03/17 | Influenza A virus (A/environment/Hunankaifu/303/2017(H5N6)) | c |
| MH022716 | 1766 | Avian | 4 (HA) | H5N6 | China | N | 2016/11/29 | Influenza A virus (A/greenwing duck/Shanghai/SH1/2016) | c |
| MH156490 | 1748 | Environment | 4 (HA) | H5N6 | China | N | 2014/06/17 | Influenza A virus (A/environment/Chang Sha/308/2014) | c |
| MH156491 | 1748 | Environment | 4 (HA) | H5N6 | China | N | 2014/09/18 | Influenza A virus (A/environment/Chang Sha/399/2014) | c |
| MH156492 | 1748 | Environment | 4 (HA) | H5N6 | China | N | 2014/10/27 | Influenza A virus (A/environment/Chang Sha/441/2014) | c |
| MH156493 | 1748 | Environment | 4 (HA) | H5N6 | China | N | 2014/10/27 | Influenza A virus (A/environment/Chang Sha/443/2014) | c |
| MH156494 | 1748 | Environment | 4 (HA) | H5N6 | China | N | 2014/10/27 | Influenza A virus (A/environment/Chang Sha/445/2014) | c |
| MH156495 | 1748 | Environment | 4 (HA) | H5N6 | China | N | 2014/10/28 | Influenza A virus (A/environment/Chang Sha/448/2014) | c |
| MH156496 | 1748 | Environment | 4 (HA) | H5N6 | China | N | 2014/10/31 | Influenza A virus (A/environment/Chang Sha/455/2014) | c |
| MH156497 | 1748 | Environment | 4 (HA) | H5N6 | China | N | 2014/10/31 | Influenza A virus (A/environment/Chang Sha/458/2014) | c |
| MH156498 | 1748 | Environment | 4 (HA) | H5N6 | China | N | 2014/11/25 | Influenza A virus (A/environment/Chang Sha/488/2014) | c |
| MH156499 | 1748 | Environment | 4 (HA) | H5N6 | China | N | 2014/11/26 | Influenza A virus (A/environment/Chang Sha/499/2014) | c |
| MH283029 | 1704 | Avian | 4 (HA) | H5N6 | China | N | 2017/10/21 | Influenza A virus (A/common mallard/Ningxia/YG71/2017) | c |
| MH283037 | 1704 | Avian | 4 (HA) | H5N6 | China | N | 2017/10/21 | Influenza A virus (A/spot-billed duck/Ningxia/YG83/2017) | c |
| MH283045 | 1704 | Avian | 4 (HA) | H5N6 | China | N | 2017/10/21 | Influenza A virus (A/spot-billed duck/Ningxia/YG87/2017) | c |
| MH283053 | 1704 | Avian | 4 (HA) | H5N6 | China | N | 2017/10/21 | Influenza A virus (A/spot-billed duck/Ningxia/YG93/2017) | c |
| MH569637 | 1704 | Avian | 4 (HA) | H5N8 | Uganda | T | 2017/01 | Influenza A virus (A/white_winged_black_tern/Uganda/17RS115-3/2017) | c |
| MH988777 | 1735 | Environment | 4 (HA) | H5N8 | Ukraine | N | 2017/01/27 | Influenza A virus (A/Environmental fecal/AN/2/2017) | c |
| MK418658 | 1701 | Mink | 4 (HA) | H5N6 | China | N | 2018/01/30 | Influenza A virus (A/mink/Northern China/F0130m/2018) | c |
| MK631789 | 1762 | Avian | 4 (HA) | H5N8 | Democratic R | T | 2017/05/13 | Influenza A virus (A/Muscovy duck/Democratic Republic of the Congo/KAF1/2017) | c |
| MK636726 | 1761 | Avian | 4 (HA) | H5N8 | Democratic R | T | 2017/05/14 | Influenza A virus (A/Muscovy duck/Democratic Republic of the Congo/KAF4_HA/201 | c |
| MK636734 | 1762 | Avian | 4 (HA) | H5N8 | Democratic R | T | 2017/05/14 | Influenza A virus (A/Muscovy duck/Democratic Republic of the Congo/NYA4_HA/20 | c |
| MK636742 | 1762 | Avian | 4 (HA) | H5N8 | Democratic R | T | 2017/05/13 | Influenza A virus (A/Muscovy duck/Democratic Republic of the Congo/NYA14_HA/2 | c |
| MK636750 | 1762 | Avian | 4 (HA) | H5N8 | Democratic R | T | 2017/05/13 | Influenza A virus (A/Muscovy duck/Democratic Republic of the Congo/NYA15_HA/2 | c |
| MK636758 | 1762 | Avian | 4 (HA) | H5N8 | Democratic R | T | 2017/05/13 | Influenza A virus (A/Muscovy duck/Democratic Republic of the Congo/TCH4_HA/201 | c |
| MK636766 | 1762 | Avian | 4 (HA) | H5N8 | Democratic R | T | 2017/05/13 | Influenza A virus (A/Muscovy duck/Democratic Republic of the Congo/TCH5_HA/201 | c |
| MK636774 | 1760 | Avian | 4 (HA) | H5N8 | Democratic R | T | 2017/05/15 | Influenza A virus (A/Muscovy duck/Democratic Republic of the Congo/TCH6_HA/201 | c |
| MK734294 | 1704 | Avian | 4 (HA) | H5N8 | South Africa | S | 2018/01/29 | Influenza A virus (A/Sandwich tern/South Africa/18010369/2018) | c |
| MK734295 | 1704 | Penguin | 4 (HA) | H5N8 | South Africa | S | 2018/03/01 | Influenza A virus (A/African penguin/South Africa/476266/2018) | c |
| MK734296 | 1704 | Avian | 4 (HA) | H5N8 | South Africa | S | 2018/01/29 | Influenza A virus (A/Common tern/South Africa/18010371/2018) | c |
| MK734297 | 1704 | Avian | 4 (HA) | H5N8 | South Africa | S | 2018/01/03 | Influenza A virus (A/Swift tern/South Africa/18010027/2018) | c |
| MK734298 | 1704 | Avian | 4 (HA) | H5N8 | South Africa | S | 2018/01/03 | Influenza A virus (A/Swift tern/South Africa/18010028/2018) | c |
| MK734299 | 1704 | Avian | 4 (HA) | H5N8 | South Africa | S | 2018/01/29 | Influenza A virus (A/Swift tern/South Africa/18010370/2018) | c |
| MK734300 | 1704 | Avian | 4 (HA) | H5N8 | South Africa | S | 2018/01/31 | Influenza A virus (A/tern/South Africa/18010417/2018) | c |
| MK779131 | 1772 | Avian | 4 (HA) | H5N6 | China | N | 2018/08/11 | Influenza A virus (A/white-naped crane/China/ya3/2018) | c |
| MK942972 | 1749 | Avian | 4 (HA) | H5N6 | Viet Nam | T | 2015/10/25 | Influenza A virus (A/muscovy duck/Viet Nam/HN-2508/2015) | c |

| | | | | | | | | | |
|---|---|---|---|---|---|---|---|---|---|
| MK943406 | 1749 | Avian | 4 (HA) | H5N6 | Viet Nam | T | 2015/10/25 | Influenza A virus (A/muscovy duck/Viet Nam/HN-2501/2015) | c |
| MK943414 | 1749 | Avian | 4 (HA) | H5N6 | Viet Nam | T | 2015/10/25 | Influenza A virus (A/muscovy duck/Viet Nam/HN-2504/2015) | c |
| MK943422 | 1749 | Avian | 4 (HA) | H5N6 | Viet Nam | T | 2015/10/25 | Influenza A virus (A/muscovy duck/Viet Nam/HN-2506/2015) | c |
| MK943478 | 1749 | Avian | 4 (HA) | H5N6 | Viet Nam | T | 2016/01/05 | Influenza A virus (A/muscovy duck/Viet Nam/QN-2611/2016) | c |
| MK943486 | 1749 | Avian | 4 (HA) | H5N6 | Viet Nam | T | 2016/01/05 | Influenza A virus (A/muscovy duck/Viet Nam/QN-2612/2016) | c |
| MK943502 | 1749 | Avian | 4 (HA) | H5N6 | Viet Nam | T | 2016/01/05 | Influenza A virus (A/muscovy duck/Viet Nam/QN-2616/2016) | c |
| MK943522 | 1749 | Avian | 4 (HA) | H5N6 | Viet Nam | T | 2016/04/16 | Influenza A virus (A/muscovy duck/Viet Nam/QN-2706/2016) | c |
| MK963921 | 1749 | Avian | 4 (HA) | H5 | Viet Nam | T | 2014/08/04 | Influenza A virus (A/muscovy duck/Viet Nam/HN-1697/2014) | c |
| MK964973 | 1749 | Avian | 4 (HA) | H5N6 | Viet Nam | T | 2014/08/04 | Influenza A virus (A/muscovy duck/Viet Nam/HN-1699/2014) | c |
| MK975995 | 1532 | Avian | 4 (HA) | H5N8 | Egypt | N | 2018 | Influenza A virus (A/chicken/Egypt/AB2/2018) | p |
| MN025431 | 1766 | Avian | 4 (HA) | H5N8 | Denmark | N | 2016 | Influenza A virus (A/tufted duck/Denmark/11740/2016) | c |
| MN037415 | 1776 | Avian | 4 (HA) | H5N2 | USA | N | 2015/01/09 | Influenza A virus (A/cooper's hawk/Washington/2551/2015) | c |
| MN124544 | 1624 | Avian | 4 (HA) | H5N8 | South Africa | S | 2018/01/04 | Influenza A virus (A/Tern/South Africa/18010043/2018) | p |
| MN124552 | 1624 | Avian | 4 (HA) | H5N8 | South Africa | S | 2018/01/11 | Influenza A virus (A/Jackal buzzard/South Africa/18010106/2018) | p |
| MN124560 | 1624 | Avian | 4 (HA) | H5N8 | South Africa | S | 2018/01/08 | Influenza A virus (A/Tern/South Africa/18010107/2018) | p |
| MN124568 | 1624 | Avian | 4 (HA) | H5N8 | South Africa | S | 2018/01/18 | Influenza A virus (A/Common tern/South Africa/18010259/2018) | p |
| MN124576 | 1624 | Penguin | 4 (HA) | H5N8 | South Africa | S | 2018/01/31 | Influenza A virus (A/African penguin/South Africa/18010422/2018) | p |
| MN124584 | 1624 | Penguin | 4 (HA) | H5N8 | South Africa | S | 2018/01/31 | Influenza A virus (A/African penguin/South Africa/18010423/2018) | p |
| MN124592 | 1624 | Avian | 4 (HA) | H5N8 | South Africa | S | 2018/02/15 | Influenza A virus (A/Swift tern/South Africa/18020273/2018) | p |
| MN124600 | 1624 | Avian | 4 (HA) | H5N8 | South Africa | S | 2018/02/15 | Influenza A virus (A/Sandwich tern/South Africa/18020302/2018) | p |
| MN124608 | 1624 | Avian | 4 (HA) | H5N8 | South Africa | S | 2018/02/15 | Influenza A virus (A/Cape cormorant/South Africa/18020303/2018) | p |
| MN124616 | 1624 | Penguin | 4 (HA) | H5N8 | South Africa | S | 2018/02/15 | Influenza A virus (A/African penguin/South Africa/18020304/2018) | p |
| MN124624 | 1624 | Penguin | 4 (HA) | H5N8 | South Africa | S | 2018/02/22 | Influenza A virus (A/African penguin/South Africa/18020408/2018) | p |
| MN124632 | 1624 | Avian | 4 (HA) | H5N8 | South Africa | S | 2018/03/09 | Influenza A virus (A/Crowned cormorant/South Africa/18030213/2018) | p |
| MN124640 | 1624 | Avian | 4 (HA) | H5N8 | South Africa | S | 2018/03/09 | Influenza A virus (A/African oystercatcher/South Africa/18030214/2018) | p |
| MN124648 | 1624 | Avian | 4 (HA) | H5N8 | South Africa | S | 2018/03/27 | Influenza A virus (A/Swift tern/South Africa/18030478/2018) | p |
| MN124656 | 1624 | Avian | 4 (HA) | H5N8 | South Africa | S | 2018/04/12 | Influenza A virus (A/Hartlaubs gull/South Africa/18040224/2018) | p |
| MN124664 | 1624 | Avian | 4 (HA) | H5N8 | South Africa | S | 2018/04/17 | Influenza A virus (A/Swift tern/South Africa/18040275/2018) | p |
| MN124672 | 1624 | Avian | 4 (HA) | H5N8 | South Africa | S | 2018/04/20 | Influenza A virus (A/Hartlaubs gull/South Africa/18040367/2018) | p |
| MN124680 | 1624 | Penguin | 4 (HA) | H5N8 | South Africa | S | 2018/05/12 | Influenza A virus (A/African penguin/South Africa/18050256/2018) | p |
| MN128638 | 1704 | Avian | 4 (HA) | H5N6 | China | N | 2015/12/16 | Influenza A virus (A/goose/Guangdong/A-Goose-Guangdong-GS017-2015) | c |
| MN128651 | 1704 | Avian | 4 (HA) | H5N6 | China | N | 2015/12/16 | Influenza A virus (A/goose/Guangdong/A-Goose-Guangdong-GS114-2015) | c |
| MN128659 | 1704 | Avian | 4 (HA) | H5N6 | China | N | 2015/12/16 | Influenza A virus (A/goose/Guangdong/A-Goose-Guangdong-GS116-2015) | c |
| MN128667 | 1704 | Avian | 4 (HA) | H5N6 | China | N | 2015/12/16 | Influenza A virus (A/goose/Guangdong/A-Goose-Guangdong-GS120-2015) | c |
| MN128675 | 1704 | Avian | 4 (HA) | H5N6 | China | N | 2015/12/16 | Influenza A virus (A/goose/Guangdong/A-Goose-Guangdong-GS119-2015) | c |
| MN128680 | 1704 | Avian | 4 (HA) | H5N6 | China | N | 2015/12/16 | Influenza A virus (A/goose/Guangdong/A-Goose-Guangdong-GS144-2015) | c |
| MN128689 | 1704 | Avian | 4 (HA) | H5N6 | China | N | 2016/01/08 | Influenza A virus (A/goose/Guangdong/A-Goose-Guangdong-GS148-2016) | c |
| MN128870 | 1704 | Avian | 4 (HA) | H5N6 | China | N | 2015/12/16 | Influenza A virus (A/goose/Guangdong/A-Goose-Guangdong-GS018-2015) | c |
| MN556609 | 1704 | Avian | 4 (HA) | H5N6 | China | N | 2013/11/26 | Influenza A virus (A/duck/China/13087-4.seq/2013) | c |
| MN556681 | 1704 | Avian | 4 (HA) | H5N6 | China | N | 2014/01/06 | Influenza A virus (A/goose/China/14016HA-4.seq/2014) | c |
| MN556689 | 1704 | Avian | 4 (HA) | H5N6 | China | N | 2014/01/06 | Influenza A virus (A/duck/China/14017HA-4.seq/2014) | c |
| MN556725 | 1704 | Avian | 4 (HA) | H5N6 | China | N | 2014/03/15 | Influenza A virus (A/duck/China/14085HA-4.seq/2014) | c |
| MN559704 | 1695 | Avian | 4 (HA) | H5N8 | Egypt | N | 2017/01/22 | Influenza A virus (A/goose/Egypt/A_Geese_Egypt_MG3_2017/2017) | c |
| MN559705 | 1567 | Avian | 4 (HA) | H5N8 | Egypt | N | 2019/04/01 | Influenza A virus (A/goose/Egypt/A_Geese_Egypt_MG4_2019/2019) | p |
| MN565986 | 1701 | Avian | 4 (HA) | H5N6 | South Korea | N | 2017/12/12 | Influenza A virus (A/Anas platyrhynchos/South Korea/1702/2017) | c |
| MN566011 | 1701 | Avian | 4 (HA) | H5N6 | South Korea | N | 2017/12 | Influenza A virus (A/Anas platyrhynchos/South Korea/1703/2017) | c |
| MN566019 | 1701 | Avian | 4 (HA) | H5N6 | South Korea | N | 2017/12 | Influenza A virus (A/Wild bird(Anas platyrhynchos)/South Korea/1705/2017) | c |
| MN566028 | 1701 | Avian | 4 (HA) | H5N6 | South Korea | N | 2017/12 | Influenza A virus (A/Anas platyrhynchos/South Korea/1709/2017) | c |
| MN566036 | 1701 | Avian | 4 (HA) | H5N6 | South Korea | N | 2017/12 | Influenza A virus (A/wild bird(Anas platyrhynchos)/South Korea/1710/2017) | c |
| MN566053 | 1704 | Avian | 4 (HA) | H5N6 | South Korea | N | 2018/11 | Influenza A virus (A/wild duck/South Korea/1801/2018) | c |
| MN577280 | 1701 | Avian | 4 (HA) | H5N6 | South Korea | N | 2019/08 | Influenza A virus (A/wild duck/South Korea/1908/2019) | c |
| MN577314 | 1701 | Avian | 4 (HA) | H5N6 | South Korea | N | 2019/08 | Influenza A virus (A/wild duck/South Korea/1914/2019) | c |
| MN577323 | 1701 | Avian | 4 (HA) | H5N6 | South Korea | N | 2019/08 | Influenza A virus (A/wild duck/South Korea/1915/2019) | c |
| MN577334 | 1701 | Avian | 4 (HA) | H5N6 | South Korea | N | 2019/08 | Influenza A virus (A/wild duck/South Korea/1920/2019) | c |
| MN577345 | 1701 | Avian | 4 (HA) | H5N6 | South Korea | N | 2019/08 | Influenza A virus (A/wild duck/South Korea/1922/2019) | c |
| MN708201 | 1776 | Avian | 4 (HA) | H5N8 | Denmark | N | 2016/11/07 | Influenza A virus (A/Tufted duck/Denmark/11740-LWPL/2016) | c |
| MN759479 | 1704 | Avian | 4 (HA) | H5N8 | Nigeria | T | 2017/09 | Influenza A virus (A/duck/Nigeria/KNSB-11-13T_18RS1971-38/2017) | c |
| MN759480 | 1704 | Avian | 4 (HA) | H5N8 | Nigeria | T | 2017/09 | Influenza A virus (A/duck/Nigeria/KNSB-8-10T_18RS1971-37/2017) | c |
| MN759481 | 1704 | Avian | 4 (HA) | H5N8 | Nigeria | T | 2017/09 | Influenza A virus (A/duck/Nigeria/KNSB-20-23T_18RS1971-39/2017) | c |
| MN759482 | 1704 | Avian | 4 (HA) | H5N8 | Nigeria | T | 2017/09 | Influenza A virus (A/duck/Nigeria/KDIG-164-165T_18RS1971-40/2017) | c |
| MN759483 | 1704 | Avian | 4 (HA) | H5N8 | Nigeria | T | 2017/09 | Influenza A virus (A/duck/Nigeria/KDIG-171-172C_18RS1971-41/2017) | c |
| MN759484 | 1704 | Avian | 4 (HA) | H5N8 | Nigeria | T | 2017/09 | Influenza A virus (A/duck/Nigeria/KDIG-173-174_18RS1971-42/2017) | c |
| MN759485 | 1704 | Avian | 4 (HA) | H5N8 | Nigeria | T | 2017/02 | Influenza A virus (A/mallard_duck/Nigeria/KN-AR91-3T_18RS1971-45/2017) | c |
| MN759486 | 1704 | Avian | 4 (HA) | H5N8 | Nigeria | T | 2017/02 | Influenza A virus (A/muscovy_duck/Nigeria/KN-P31-33T_18RS1971-46/2017) | c |
| MN759487 | 1704 | Avian | 4 (HA) | H5N8 | Nigeria | T | 2017/02 | Influenza A virus (A/muscovy_duck/Nigeria/KN-P52-53T_18RS1971-47/2017) | c |
| MN759488 | 1704 | Avian | 4 (HA) | H5N8 | Nigeria | T | 2017/02 | Influenza A virus (A/muscovy_duck/Nigeria/KN-AR71-3T_18RS1971-49/2017) | c |
| MN759489 | 1704 | Avian | 4 (HA) | H5N8 | Nigeria | T | 2017/02 | Influenza A virus (A/duck/Nigeria/KN-K71-4C_18RS1971-50/2017) | c |
| MN759490 | 1704 | Avian | 4 (HA) | H5N8 | Nigeria | T | 2016/11 | Influenza A virus (A/chicken/Nigeria/VRD16-KN623_17RS737-43/2016) | c |
| MN809352 | 1704 | Avian | 4 (HA) | H5N6 | South Korea | N | 2018/12 | Influenza A virus (A/Wild duck/South Korea/1804/2018) | c |
| MN875108 | 1656 | Avian | 4 (HA) | H5N8 | France | N | 2016/12/28 | Influenza A virus (A/chicken/France/161585/2016) | p |
| MN875109 | 1656 | Avian | 4 (HA) | H5N8 | France | N | 2016/12/18 | Influenza A virus (A/eurasian wigeon/France/161323/2016) | p |
| MT027086 | 1704 | Avian | 4 (HA) | H5N8 | Slovakia | N | 2020/01/03 | Influenza A virus (A/layer hen/Slovakia/A-chicken-Slovakia-Pah_14-2020_H5N8_HA/ | c |
| MT200035 | 1762 | Avian | 4 (HA) | H5N6 | Viet Nam | T | 2019/12/30 | Influenza A virus (A/Domestic duck/Viet Nam/A_duck_Vietnam_HU12-1473_2019/2 | c |
| MT256068 | 1704 | Avian | 4 (HA) | H5N8 | Egypt | N | 2019/03/16 | Influenza A virus (A/Broiler chicken/Giza/VRLCU/2019) | c |
| MT256069 | 1704 | Avian | 4 (HA) | H5N8 | Egypt | N | 2019/05/10 | Influenza A virus (A/cattle egret/Monofiya/VRLCU/2019) | c |
| MT547570 | 1758 | Avian | 4 (HA) | H5N1 | Viet Nam | T | 2019/08 | Influenza A virus (A/chicken/Viet Nam/HU12-1328/2019) | c |
| MT781470 | 1704 | Avian | 4 (HA) | H5N6 | Denmark | N | 2018/09/03 | Influenza A virus (A/eider/Denmark/12292-1/2018) | c |
| MT781474 | 1702 | Avian | 4 (HA) | H5N6 | Denmark | N | 2018/09/03 | Influenza A virus (A/mute swan/Denmark/12293-1/2018) | nc |
| MT781490 | 1704 | Avian | 4 (HA) | H5N6 | Denmark | N | 2018/04/18 | Influenza A virus (A/common buzzard/Denmark/4907-1/2018) | c |
| MT781495 | 1704 | Avian | 4 (HA) | H5N6 | Denmark | N | 2019/01/04 | Influenza A virus (A/common buzzard/Denmark/613-1/2019) | c |
| MT781503 | 1704 | Avian | 4 (HA) | H5N6 | Denmark | N | 2018/07/11 | Influenza A virus (A/eider/Denmark/9885-1/2018) | c |
| MT781511 | 1704 | Avian | 4 (HA) | H5N6 | Denmark | N | 2018/08/12 | Influenza A virus (A/mute swan/Denmark/10771-1S/2018) | c |
| MT781519 | 1704 | Avian | 4 (HA) | H5N6 | Denmark | N | 2018/03/27 | Influenza A virus (A/mute swan/Denmark/4905-1/2018) | c |

| | | | | | | | | | |
|---|---|---|---|---|---|---|---|---|---|
| MT781535 | 1704 | Avian | 4 (HA) | H5N6 | Denmark | N | 2018/12/22 | Influenza A virus (A/white-tailed eagle/Denmark/0429-1/2018) | c |
| MT781546 | 1704 | Avian | 4 (HA) | H5N6 | Denmark | N | 2018/02/13 | Influenza A virus (A/white-tailed eagle/Denmark/3073-1/2018) | c |
| MT781554 | 1704 | Avian | 4 (HA) | H5N6 | Denmark | N | 2018/02/09 | Influenza A virus (A/white-tailed eagle/Denmark/4227-1/2018) | c |
| MT781562 | 1704 | Avian | 4 (HA) | H5N6 | Denmark | N | 2018/03/16 | Influenza A virus (A/white-tailed eagle/Denmark/4240-1/2018) | c |
| MT781570 | 1704 | Avian | 4 (HA) | H5N6 | Denmark | N | 2018/03/14 | Influenza A virus (A/white-tailed eagle/Denmark/4241-1/2018) | c |
| MT781578 | 1704 | Avian | 4 (HA) | H5N6 | Denmark | N | 2018/04/03 | Influenza A virus (A/white-tailed eagle/Denmark/4896-1/2018) | c |
| MT781586 | 1704 | Avian | 4 (HA) | H5N6 | Denmark | N | 2018/04/02 | Influenza A virus (A/white-tailed eagle/Denmark/4897-1/2018) | c |
| MT781591 | 1704 | Avian | 4 (HA) | H5N6 | Denmark | N | 2019/03/27 | Influenza A virus (A/white-tailed eagle/Denmark/4903-1/2019) | c |
| MT781602 | 1704 | Avian | 4 (HA) | H5N6 | Denmark | N | 2018/02/12 | Influenza A virus (A/white-tailed eagle/Denmark/5921-1/2018) | c |
| MT912721 | 1701 | Avian | 4 (HA) | H5N6 | China | N | 2017/12 | Influenza A virus (A/egret/Zhejiang/W18/2017) | c |
| MT912731 | 1704 | Avian | 4 (HA) | H5N6 | China | N | 2017/12 | Influenza A virus (A/egret/Zhejiang/W15/2017) | c |
| MW026072 | 1737 | Avian | 4 (HA) | H5N8 | Denmark | N | 2017/03/23 | Influenza A virus (A/common buzzard/Denmark/4079-1p1/2017) | c |
| MW026080 | 1737 | Avian | 4 (HA) | H5N8 | Denmark | N | 2017/03/03 | Influenza A virus (A/common buzzard/Denmark/3794-1p1/2017) | c |
| MW026088 | 1737 | Avian | 4 (HA) | H5N8 | Denmark | N | 2016/11/16 | Influenza A virus (A/common gull/Denmark/18577-1/2016) | c |
| MW026112 | 1737 | Avian | 4 (HA) | H5N8 | Denmark | N | 2016/11/16 | Influenza A virus (A/great black-backed gull/Denmark/19069-1/2016) | c |
| MW026120 | 1737 | Avian | 4 (HA) | H5N8 | Denmark | N | 2016/11/19 | Influenza A virus (A/mute swan/Denmark/19192-1/2016) | c |
| MW026128 | 1737 | Avian | 4 (HA) | H5N8 | Denmark | N | 2016/11/07 | Influenza A virus (A/tufted duck/Denmark/17740-1p1/2016) | c |
| MW097446 | 1704 | Environment | 4 (HA) | H5N6 | China | N | 2017/02/15 | Influenza A virus (A/environment/Yunnan/2.15_DQXC083-E/2017) | c |
| MW097447 | 1704 | Environment | 4 (HA) | H5N6 | China | N | 2017/03/02 | Influenza A virus (A/environment/Yunnan/3.20_DQSPZ024-E/2017) | c |
| MW097754 | 1745 | Avian | 4 (HA) | H5N6 | China | N | 2018/03/15 | Influenza A virus (A/goose/Fujian/3.15_FZHX0004-O/2018) | c |
| MW097755 | 1755 | Avian | 4 (HA) | H5N6 | China | N | 2018/03/15 | Influenza A virus (A/goose/Fujian/3.15_FZHX0006-O/2018) | c |
| MW097756 | 1757 | Avian | 4 (HA) | H5N6 | China | N | 2018/03/15 | Influenza A virus (A/goose/Fujian/3.15_FZHX0009-O/2018) | c |
| MW097768 | 1769 | Avian | 4 (HA) | H5N6 | China | N | 2018/12/26 | Influenza A virus (A/chicken/Shandong/12.26_TAWL023-O/2018) | c |
| MW097770 | 1770 | Environment | 4 (HA) | H5N6 | China | N | 2018/12/26 | Influenza A virus (A/environment/Shandong/12.26_TAWL002-E/2018) | c |
| MW097774 | 1769 | Avian | 4 (HA) | H5N6 | China | N | 2018/11/03 | Influenza A virus (A/chicken/Shanxi/11.30_TGRL018-O/2018) | c |
| MW097777 | 1773 | Avian | 4 (HA) | H5N6 | China | N | 2018/11/28 | Influenza A virus (A/chicken/Shandong/11.28_TAWL019-O/2018) | c |
| MW097778 | 1769 | Avian | 4 (HA) | H5N6 | China | N | 2018/11/28 | Influenza A virus (A/chicken/Shandong/11.28_TAWL021-O/2018) | c |
| MW097779 | 1769 | Avian | 4 (HA) | H5N6 | China | N | 2018/11/28 | Influenza A virus (A/chicken/Shandong/11.28_TAWL022-O/2018) | c |
| MW097780 | 1769 | Avian | 4 (HA) | H5N6 | China | N | 2018/11/28 | Influenza A virus (A/chicken/Shandong/11.28_TAWL024-O/2018) | c |
| MW097781 | 1769 | Environment | 4 (HA) | H5N6 | China | N | 2018/11/28 | Influenza A virus (A/environment/Shandong/11.28_TAWL002-E/2018) | c |
| MW097782 | 1769 | Avian | 4 (HA) | H5N6 | China | N | 2018/11/01 | Influenza A virus (A/chicken/Henan/11.01_XXHM013-C/2018) | c |
| MW097784 | 1773 | Avian | 4 (HA) | H5N6 | China | N | 2018/11/29 | Influenza A virus (A/chicken/Anhui/11.29_YHZGS007-O/2018) | c |
| MW097785 | 1773 | Avian | 4 (HA) | H5N6 | China | N | 2018/11/29 | Influenza A virus (A/chicken/Anhui/11.29_YHZGS009-O/2018) | c |
| MW097786 | 1769 | Avian | 4 (HA) | H5N6 | China | N | 2018/11/29 | Influenza A virus (A/chicken/Anhui/11.29_YHZGS010-O/2018) | c |
| MW097787 | 1769 | Avian | 4 (HA) | H5N6 | China | N | 2018/11/29 | Influenza A virus (A/chicken/Anhui/11.29_YHZGS011-O/2018) | c |
| MW097788 | 1769 | Avian | 4 (HA) | H5N6 | China | N | 2018/11/29 | Influenza A virus (A/chicken/Anhui/11.29_YHZGS012-O/2018) | c |
| MW097789 | 1772 | Avian | 4 (HA) | H5N6 | China | N | 2018/11/29 | Influenza A virus (A/chicken/Anhui/11.29_YHZGS013-O/2018) | c |
| MW097790 | 1769 | Avian | 4 (HA) | H5N6 | China | N | 2018/11/29 | Influenza A virus (A/chicken/Anhui/11.29_YHZGS014-O/2018) | c |
| MW097791 | 1771 | Avian | 4 (HA) | H5N6 | China | N | 2018/11/29 | Influenza A virus (A/chicken/Anhui/11.29_YHZGS015-O/2018) | c |
| MW097792 | 1769 | Avian | 4 (HA) | H5N6 | China | N | 2018/11/29 | Influenza A virus (A/chicken/Anhui/11.29_YHZGS016-O/2018) | c |
| MW097793 | 1769 | Avian | 4 (HA) | H5N6 | China | N | 2018/11/29 | Influenza A virus (A/chicken/Anhui/11.29_YHZGS017-O/2018) | c |
| MW097794 | 1769 | Avian | 4 (HA) | H5N6 | China | N | 2018/11/29 | Influenza A virus (A/chicken/Anhui/11.29_YHZGS019-O/2018) | c |
| MW097795 | 1772 | Avian | 4 (HA) | H5N6 | China | N | 2018/11/29 | Influenza A virus (A/chicken/Anhui/11.29_YHZGS020-O/2018) | c |
| MW097796 | 1769 | Avian | 4 (HA) | H5N6 | China | N | 2018/11/29 | Influenza A virus (A/chicken/Anhui/11.29_YHZGS022-O/2018) | c |
| MW097797 | 1769 | Avian | 4 (HA) | H5N6 | China | N | 2018/11/29 | Influenza A virus (A/chicken/Anhui/11.29_YHZGS023-O/2018) | c |
| MW097798 | 1769 | Avian | 4 (HA) | H5N6 | China | N | 2018/11/29 | Influenza A virus (A/chicken/Anhui/11.29_YHZGS024-O/2018) | c |
| MW097799 | 1769 | Avian | 4 (HA) | H5N6 | China | N | 2018/11/29 | Influenza A virus (A/chicken/Anhui/11.29_YHZGS025-O/2018) | c |
| MW097800 | 1772 | Avian | 4 (HA) | H5N6 | China | N | 2018/11/29 | Influenza A virus (A/chicken/Anhui/11.29_YHZGS001-C/2018) | c |
| MW097801 | 1771 | Avian | 4 (HA) | H5N6 | China | N | 2018/11/29 | Influenza A virus (A/chicken/Anhui/11.29_YHZGS004-C/2018) | c |
| MW097802 | 1769 | Avian | 4 (HA) | H5N6 | China | N | 2018/11/29 | Influenza A virus (A/chicken/Anhui/11.29_YHZGS006-C/2018) | c |
| MW097803 | 1773 | Avian | 4 (HA) | H5N6 | China | N | 2018/11/29 | Influenza A virus (A/chicken/Anhui/11.29_YHZGS009-C/2018) | c |
| MW097804 | 1773 | Avian | 4 (HA) | H5N6 | China | N | 2018/11/29 | Influenza A virus (A/chicken/Anhui/11.29_YHZGS019-C/2018) | c |
| MW097805 | 1773 | Avian | 4 (HA) | H5N6 | China | N | 2018/11/29 | Influenza A virus (A/chicken/Anhui/11.29_YHZGS021-C/2018) | c |
| MW097807 | 1773 | Environment | 4 (HA) | H5N6 | China | N | 2018/11/29 | Influenza A virus (A/environment/Anhui/11.29_YHZGS001-E/2018) | c |
| MW097810 | 1771 | Avian | 4 (HA) | H5N6 | China | N | 2018/10/28 | Influenza A virus (A/chicken/Chongqing/10.28_RCTYH003-O/2018) | c |
| MW097811 | 1771 | Avian | 4 (HA) | H5N6 | China | N | 2018/10/28 | Influenza A virus (A/chicken/Chongqing/10.28_RCTYH020-O/2018) | c |
| MW097812 | 1773 | Avian | 4 (HA) | H5N6 | China | N | 2018/11/29 | Influenza A virus (A/chicken/Henan/11.29_XXHM015-O/2018) | c |
| MW097813 | 1769 | Avian | 4 (HA) | H5N6 | China | N | 2018/11/29 | Influenza A virus (A/chicken/Henan/11.29_XXHM017-O/2018) | c |
| MW097815 | 1769 | Environment | 4 (HA) | H5N6 | China | N | 2018/11/29 | Influenza A virus (A/environment/Henan/11.29_XXHM001-E/2018) | c |
| MW097816 | 1773 | Environment | 4 (HA) | H5N6 | China | N | 2018/11/29 | Influenza A virus (A/environment/Henan/11.29_XXHM003-E/2018) | c |
| MW097817 | 1773 | Environment | 4 (HA) | H5N6 | China | N | 2018/11/29 | Influenza A virus (A/environment/Henan/11.29_XXHM004-E/2018) | c |
| MW097818 | 1769 | Avian | 4 (HA) | H5N6 | China | N | 2018/11/03 | Influenza A virus (A/chicken/Shanxi/11.30_TGRL002-O/2018) | c |
| MW097819 | 1770 | Avian | 4 (HA) | H5N6 | China | N | 2018/12/01 | Influenza A virus (A/chicken/Shanxi/12.01_TGRL005-O/2018) | c |
| MW097820 | 1770 | Avian | 4 (HA) | H5N6 | China | N | 2018/11/03 | Influenza A virus (A/chicken/Shanxi/11.30_TGRL007-O/2018) | c |
| MW097823 | 1773 | Avian | 4 (HA) | H5N2 | China | N | 2018/10/28 | Influenza A virus (A/chicken/Chongqing/10.28_RCTYH007-O/2018) | c |
| MW097824 | 1574 | Avian | 4 (HA) | H5N6 | China | N | 2018/08/28 | Influenza A virus (A/chicken/Anhui/8.28_YHZGS012-O/2018) | p |
| MW097827 | 1753 | Avian | 4 (HA) | H5N6 | China | N | 2017/05/26 | Influenza A virus (A/chicken/Jiangxi/5.26_NCNP13Q2-OC/2017) | c |
| MW097845 | 1755 | Environment | 4 (HA) | H5N2 | China | N | 2017/01/11 | Influenza A virus (A/environment/Jiangxi/1.11_NCNP65E2-E/2017) | c |
| MW097850 | 1754 | Environment | 4 (HA) | H5N6 | China | N | 2017/01/11 | Influenza A virus (A/environment/Jiangxi/1.11_NCDZT99F2-E/2017) | c |
| MW097861 | 1751 | Avian | 4 (HA) | H5N6 | China | N | 2017/10/03 | Influenza A virus (A/chicken/Jiangxi/10.30_NCDZT17B3-OC/2017) | c |
| MW097866 | 1747 | Avian | 4 (HA) | H5N6 | China | N | 2018/03/07 | Influenza A virus (A/duck/Hunan/03.07_YYGK31L3-OC/2018) | c |
| MW097868 | 1750 | Avian | 4 (HA) | H5N6 | China | N | 2018/03/07 | Influenza A virus (A/chicken/Hunan/03.07_YYGK40L3-OC/2018) | c |
| MW097948 | 1769 | Avian | 4 (HA) | H5N6 | China | N | 2018/08/28 | Influenza A virus (A/chicken/Henan/8.28_XXHM003-O/2018) | c |
| MW097949 | 1769 | Avian | 4 (HA) | H5N6 | China | N | 2018/08/28 | Influenza A virus (A/chicken/Henan/8.28_XXHM007-O/2018) | c |
| MW097950 | 1771 | Avian | 4 (HA) | H5N6 | China | N | 2018/08/28 | Influenza A virus (A/chicken/Henan/8.28_XXHM016-O/2018) | c |
| MW097951 | 1773 | Avian | 4 (HA) | H5N6 | China | N | 2018/08/28 | Influenza A virus (A/chicken/Henan/8.28_XXHM004-C/2018) | c |
| MW097952 | 1770 | Avian | 4 (HA) | H5N6 | China | N | 2018/08/28 | Influenza A virus (A/chicken/Shanxi/8.28_TGRL001-O/2018) | c |
| MW097953 | 1771 | Avian | 4 (HA) | H5N6 | China | N | 2018/08/28 | Influenza A virus (A/chicken/Shanxi/8.28_TGRL002-O/2018) | c |
| MW097954 | 1771 | Avian | 4 (HA) | H5N6 | China | N | 2018/08/28 | Influenza A virus (A/chicken/Shanxi/8.28_TGRL004-O/2018) | c |
| MW097955 | 1772 | Avian | 4 (HA) | H5N6 | China | N | 2018/08/28 | Influenza A virus (A/chicken/Shanxi/8.28_TGRL009-O/2018) | c |
| MW097956 | 1769 | Avian | 4 (HA) | H5N6 | China | N | 2018/08/28 | Influenza A virus (A/chicken/Shanxi/8.28_TGRL011-O/2018) | c |
| MW097957 | 1772 | Avian | 4 (HA) | H5N6 | China | N | 2018/08/28 | Influenza A virus (A/chicken/Shanxi/8.28_TGRL016-O/2018) | c |
| MW097958 | 1773 | Avian | 4 (HA) | H5N6 | China | N | 2018/08/28 | Influenza A virus (A/chicken/Shanxi/8.28_TGRL020-O/2018) | c |

| | | | | | | | | | |
|---|---|---|---|---|---|---|---|---|---|
| MW097959 | 1773 | Avian | 4 (HA) | H5N6 | China | N | 2018/08/28 | Influenza A virus (A/chicken/Shanxi/8.28_TGRL023-O/2018) | c |
| MW097960 | 1772 | Avian | 4 (HA) | H5N6 | China | N | 2018/08/28 | Influenza A virus (A/chicken/Shanxi/8.28_TGRL018-C/2018) | c |
| MW098006 | 1773 | Environment | 4 (HA) | H5N6 | China | N | 2019/01/23 | Influenza A virus (A/environment/Shanxi/01.23_TGRL004-E/2019) | c |
| MW098013 | 1773 | Avian | 4 (HA) | H5N6 | China | N | 2019/01/26 | Influenza A virus (A/chicken/Shandong/01.26_TAWL003-C/2019) | c |
| MW098017 | 1748 | Avian | 4 (HA) | H5N6 | China | N | 2019/01/26 | Influenza A virus (A/chicken/Shandong/01.26_TAWL017-C/2019) | c |
| MW098019 | 1769 | Environment | 4 (HA) | H5 | China | N | 2018/12/26 | Influenza A virus (A/environment/Shandong/12.26_TAWL001-E/2018) | c |
| MW098064 | 1749 | Avian | 4 (HA) | H5 | China | N | 2018/10/26 | Influenza A virus (A/duck/Zhejiang/10.26_HZBX001-C/2018) | c |
| MW098069 | 1767 | Avian | 4 (HA) | H5 | China | N | 2018/10/26 | Influenza A virus (A/duck/Zhejiang/10.26_HZBX004-C/2018) | c |
| MW098073 | 1742 | Avian | 4 (HA) | H5 | China | N | 2018/10/26 | Influenza A virus (A/duck/Zhejiang/10.26_HZBX012-C/2018) | c |
| MW098112 | 1745 | Environment | 4 (HA) | H5 | China | N | 2018/12/02 | Influenza A virus (A/environment/Inner_mongolia/12.02_EEDSWSQ013-E/2018) | c |
| MW098121 | 1769 | Avian | 4 (HA) | H5 | China | N | 2018/03/21 | Influenza A virus (A/chicken/Yunnan/3.21_DQXBL001-C/2018) | c |
| MW098127 | 1773 | Avian | 4 (HA) | H5 | China | N | 2017/03/08 | Influenza A virus (A/chicken/Jilin/3.08_CCNQM002-O/2017) | c |
| MW098134 | 1769 | Avian | 4 (HA) | H5 | China | N | 2018/12/24 | Influenza A virus (A/chicken/Xinjiang/12.24_WLMQXL023-O/2018) | c |
| MW098143 | 1748 | Avian | 4 (HA) | H5 | China | N | 2018/10/13 | Influenza A virus (A/chicken/Xinjiang/10.13_WLMQXL014-O/2018) | c |
| MW098144 | 1699 | Avian | 4 (HA) | H5 | China | N | 2018/10/13 | Influenza A virus (A/chicken/Xinjiang/10.13_WLMQXL018-O/2018) | p |
| MW098145 | 1747 | Avian | 4 (HA) | H5 | China | N | 2018/10/13 | Influenza A virus (A/duck/Xinjiang/10.13_WLMQXL005-O/2018) | nc |
| MW098147 | 1712 | Avian | 4 (HA) | H5 | China | N | 2018/09/19 | Influenza A virus (A/chicken/Xinjiang/09.19_WLMQXL002-O/2018) | p |
| MW098148 | 1728 | Avian | 4 (HA) | H5 | China | N | 2018/09/19 | Influenza A virus (A/chicken/Xinjiang/09.19_WLMQXL008-O/2018) | p |
| MW098156 | 1773 | Avian | 4 (HA) | H5N6 | China | N | 2018/08/26 | Influenza A virus (A/chicken/Fujian/8.26_FZHX0065-C/2018) | c |
| MW098164 | 1773 | Avian | 4 (HA) | H5 | China | N | 2018/01/25 | Influenza A virus (A/duck/Fujian/1.25_FZHX0043-O/2018) | c |
| MW098165 | 1773 | Avian | 4 (HA) | H5 | China | N | 2018/09/24 | Influenza A virus (A/chicken/Fujian/9.24_FZHX0087-C/2018) | c |
| MW098167 | 1770 | Avian | 4 (HA) | H5 | China | N | 2018/10/23 | Influenza A virus (A/chicken/Fujian/10.23_FZHX0106-O/2018) | c |
| MW098169 | 1772 | Avian | 4 (HA) | H5N6 | China | N | 2018/10/23 | Influenza A virus (A/duck/Fujian/10.23_FZHX0142-C/2018) | c |
| MW098170 | 1773 | Avian | 4 (HA) | H5N6 | China | N | 2018/11/26 | Influenza A virus (A/duck/Fujian/11.26_FZHX0143-O/2018) | c |
| MW098178 | 1773 | Avian | 4 (HA) | H5 | China | N | 2018/11/26 | Influenza A virus (A/duck/Fujian/11.26_FZHX0193-C/2018) | c |
| MW098250 | 1770 | Avian | 4 (HA) | H5N6 | China | N | 2018/03/23 | Influenza A virus (A/chicken/Jiangxi/3.23_NCDZT15M3-OC/2018) | c |
| MW098251 | 1769 | Avian | 4 (HA) | H5N6 | China | N | 2018/03/23 | Influenza A virus (A/chicken/Jiangxi/3.23_NCDZT57M3-OC/2018) | c |
| MW098252 | 1773 | Avian | 4 (HA) | H5N6 | China | N | 2018/03/23 | Influenza A virus (A/chicken/Jiangxi/3.23_NCDZT63M3-OC/2018) | c |
| MW098253 | 1770 | Avian | 4 (HA) | H5N6 | China | N | 2018/03/23 | Influenza A virus (A/duck/Jiangxi/3.23_NCNP87M3-OC/2018) | c |
| MW098254 | 1769 | Avian | 4 (HA) | H5N6 | China | N | 2018/04/25 | Influenza A virus (A/chicken/Jiangxi/4.25_NCNP6Q3-OC/2018) | c |
| MW098257 | 1773 | Avian | 4 (HA) | H5N6 | China | N | 2018/04/25 | Influenza A virus (A/chicken/Jiangxi/4.25_NCDZT70Q3-OC/2018) | c |
| MW098267 | 1773 | Avian | 4 (HA) | H5N6 | China | N | 2018/04/26 | Influenza A virus (A/chicken/Hunan/04.26_YYGK71R3-OC/2018) | c |
| MW098273 | 1769 | Avian | 4 (HA) | H5N6 | China | N | 2018/08/27 | Influenza A virus (A/chicken/Hunan/8.27_YYGK70W3-OC/2018) | c |
| MW098274 | 1769 | Avian | 4 (HA) | H5N6 | China | N | 2018/10/27 | Influenza A virus (A/chicken/Hunan/10.27_YYGK83B4-OC/2018) | c |
| MW098276 | 1773 | Avian | 4 (HA) | H5N6 | China | N | 2018/10/27 | Influenza A virus (A/chicken/Hunan/10.27_YYGK55B4-OC/2018) | c |
| MW098277 | 1769 | Avian | 4 (HA) | H5N6 | China | N | 2018/12/24 | Influenza A virus (A/chicken/Hunan/12.24_YYGK63E4-O/2018) | c |
| MW098278 | 1773 | Avian | 4 (HA) | H5N6 | China | N | 2018/12/24 | Influenza A virus (A/chicken/Hunan/12.24_YYGK35E4-O/2018) | c |
| MW098281 | 1769 | Avian | 4 (HA) | H5 | China | N | 2018/03/23 | Influenza A virus (A/duck/Jiangxi/3.23_NCNP99M3-OC/2018) | c |
| MW098299 | 1773 | Avian | 4 (HA) | H5 | China | N | 2018/05/28 | Influenza A virus (A/duck/Jiangxi/5.28_NCNP15N3-OC/2018) | c |
| MW098303 | 1772 | Avian | 4 (HA) | H5 | China | N | 2018/06/21 | Influenza A virus (A/duck/Jiangxi/6.21_NCDZT2S3-OC/2018) | c |
| MW098310 | 1772 | Avian | 4 (HA) | H5 | China | N | 2018/07/25 | Influenza A virus (A/chicken/Jiangxi/7.25_NCNP64U3-OC/2018) | c |
| MW098313 | 1771 | Avian | 4 (HA) | H5 | China | N | 2018/08/28 | Influenza A virus (A/duck/Jiangxi/8.28_NCNP19X3-OC/2018) | c |
| MW098322 | 1769 | Avian | 4 (HA) | H5 | China | N | 2018/09/22 | Influenza A virus (A/duck/Jiangxi/9.22_NCDZT93Z3-OC/2018) | c |
| MW098334 | 1769 | Avian | 4 (HA) | H5 | China | N | 2018/11/27 | Influenza A virus (A/duck/Jiangxi/11.27_NCNP93C4-OC/2018) | c |
| MW098337 | 1769 | Avian | 4 (HA) | H5 | China | N | 2018/11/27 | Influenza A virus (A/chicken/Jiangxi/11.27_NCDZT28C4-OC/2018) | c |
| MW098339 | 1772 | Avian | 4 (HA) | H5 | China | N | 2018/12/17 | Influenza A virus (A/chicken/Jiangxi/12.17_NCDZT57D4-O/2018) | c |
| MW098360 | 1769 | Avian | 4 (HA) | H5 | China | N | 2018/04/26 | Influenza A virus (A/duck/Hunan/04.26_YYGK46R3-OC/2018) | c |
| MW098362 | 1769 | Avian | 4 (HA) | H5 | China | N | 2018/05/29 | Influenza A virus (A/chicken/Hunan/5.29_YYGK1P3-OC/2018) | c |
| MW098364 | 1771 | Avian | 4 (HA) | H5 | China | N | 2018/05/29 | Influenza A virus (A/chicken/Hunan/5.29_YYGK23P3-OC/2018) | c |
| MW098370 | 1769 | Avian | 4 (HA) | H5 | China | N | 2018/05/29 | Influenza A virus (A/chicken/Hunan/5.29_YYGK48P3-OC/2018) | c |
| MW098374 | 1773 | Avian | 4 (HA) | H5 | China | N | 2018/05/29 | Influenza A virus (A/duck/Hunan/5.29_YYGK100P3-OC/2018) | c |
| MW098426 | 1773 | Avian | 4 (HA) | H5N6 | China | N | 2019/01/26 | Influenza A virus (A/chicken/Shandong/1.26_TAWL020-C/2019) | c |
| MW098427 | 1773 | Avian | 4 (HA) | H5N6 | China | N | 2019/01/26 | Influenza A virus (A/chicken/Shandong/1.26_TAWL023-C/2019) | c |
| MW098430 | 1771 | Environment | 4 (HA) | H5N6 | China | N | 2019/01/26 | Influenza A virus (A/environment/Shandong/1.26_TAWL002-E/2019) | c |
| MW098431 | 1773 | Avian | 4 (HA) | H5N6 | China | N | 2018/12/27 | Influenza A virus (A/chicken/Shanxi/12.27_TGRL005-C/2018) | c |
| MW098433 | 1770 | Avian | 4 (HA) | H5N6 | China | N | 2019/02/22 | Influenza A virus (A/chicken/Anhui/2.22_YHZGS004-O/2019) | c |
| MW098435 | 1769 | Avian | 4 (HA) | H5N6 | China | N | 2019/02/22 | Influenza A virus (A/chicken/Anhui/2.22_YHZGS008-O/2019) | c |
| MW098436 | 1769 | Avian | 4 (HA) | H5N6 | China | N | 2019/02/22 | Influenza A virus (A/chicken/Anhui/2.22_YHZGS013-O/2019) | c |
| MW098437 | 1772 | Avian | 4 (HA) | H5N6 | China | N | 2019/02/22 | Influenza A virus (A/chicken/Anhui/2.22_YHZGS015-C/2019) | c |
| MW098439 | 1773 | Avian | 4 (HA) | H5N6 | China | N | 2019/02/22 | Influenza A virus (A/chicken/Anhui/2.22_YHZGS022-C/2019) | c |
| MW098445 | 1769 | Environment | 4 (HA) | H5N6 | China | N | 2019/02/27 | Influenza A virus (A/environment/Henan/2.27_XXHM003-E/2019) | c |
| MW098446 | 1769 | Environment | 4 (HA) | H5N6 | China | N | 2019/02/27 | Influenza A virus (A/environment/Henan/2.27_XXHM004-E/2019) | c |
| MW098447 | 1769 | Environment | 4 (HA) | H5N6 | China | N | 2019/02/27 | Influenza A virus (A/environment/Henan/2.27_XXHM005-E/2019) | c |
| MW098448 | 1769 | Avian | 4 (HA) | H5N6 | China | N | 2019/02/22 | Influenza A virus (A/chicken/Shanxi/2.22_TGRL001-O/2019) | c |
| MW098449 | 1769 | Environment | 4 (HA) | H5N6 | China | N | 2019/02/22 | Influenza A virus (A/environment/Shanxi/2.22_TGRL003-E/2019) | c |
| MW098450 | 1772 | Environment | 4 (HA) | H5N6 | China | N | 2019/02/22 | Influenza A virus (A/environment/Shanxi/2.22_TGRL004-E/2019) | c |
| MW098451 | 1769 | Environment | 4 (HA) | H5N6 | China | N | 2019/02/22 | Influenza A virus (A/environment/Shanxi/2.22_TGRL005-E/2019) | c |
| MW098452 | 1769 | Avian | 4 (HA) | H5N6 | China | N | 2019/02/25 | Influenza A virus (A/chicken/Shandong/2.25_TAWL001-O/2019) | c |
| MW098453 | 1769 | Avian | 4 (HA) | H5N6 | China | N | 2019/02/25 | Influenza A virus (A/chicken/Shandong/2.25_TAWL003-O/2019) | c |
| MW098454 | 1745 | Environment | 4 (HA) | H5 | China | N | 2019/01/26 | Influenza A virus (A/environment/Shandong/1.26_TAWL005-E/2019) | nc |
| MW109328 | 1701 | Environment | 4 (HA) | H5N6 | China | N | 2017/10/26 | Influenza A virus (A/environment/Fujian/10.26_FZHX0002-E/2017) | c |
| MW109336 | 1766 | Environment | 4 (HA) | H5N6 | China | N | 2016/12/12 | Influenza A virus (A/environment/Henan/12.12_HNSQ001-E/2016) | c |
| MW109416 | 1704 | Environment | 4 (HA) | H5N8 | China | N | 2017/01/06 | Influenza A virus (A/environment/Inner_mongolia/1.06_EEDSWSQ001-E/2017) | c |
| MW109424 | 1704 | Environment | 4 (HA) | H5N8 | China | N | 2016/10/12 | Influenza A virus (A/environment/Inner_mongolia/10.12_EEDSWSQ001-E/2016) | c |
| MW109832 | 1704 | Environment | 4 (HA) | H5N8 | China | N | 2017/01/02 | Influenza A virus (A/environment/Shanxi/1.02_YC001-E/2017) | c |
| MW109840 | 1704 | Environment | 4 (HA) | H5N8 | China | N | 2017/01/01 | Influenza A virus (A/environment/XIZang/1.10_LS001-E/2017) | c |
| MW109848 | 1704 | Environment | 4 (HA) | H5N8 | China | N | 2016/10/12 | Influenza A virus (A/environment/Xinjiang/10.12_WLMQXL001-E/2016) | c |
| MW109856 | 1704 | Environment | 4 (HA) | H5N8 | China | N | 2016/12/18 | Influenza A virus (A/environment/Xinjiang/12.18_WLMQXL001-E/2016) | c |
| MW269590 | 1697 | Avian | 4 (HA) | H5N8 | China | N | 2020/10/16 | Influenza A virus (A/common teal/Shanghai/JDS20103116/2020) | p |
| MW275973 | 1698 | Avian | 4 (HA) | H5N8 | China | N | 2020/11/16 | Influenza A virus (A/tundra swan/Shanghai/CM20111601/2020) | p |
| MW314795 | 1701 | Avian | 4 (HA) | H5N6 | China | N | 2019/11 | Influenza A virus (A/chicken/China/1101/2019) | c |
| MW334838 | 1776 | Avian | 4 (HA) | H5N2 | Taiwan | N | 2017/01/11 | Influenza A virus (A/heron/Yilan/17DB0001/2017) | c |
| MW334846 | 1776 | Avian | 4 (HA) | H5N2 | Taiwan | N | 2015/12/14 | Influenza A virus (A/thrush/Taipei City/15120021/2015) | c |

| | | | | | | | | | |
|---|---|---|---|---|---|---|---|---|---|
| MW689547 | 1729 | Avian | 4 (HA) | H5N6 | Mongolia | N | 2020/04 | Influenza A virus (A/Swan goose/Mongolia/02/2020) | c |
| MW926584 | 1761 | Avian | 4 (HA) | H5N6 | Viet Nam | T | 2014/10/06 | Influenza A virus (A/Moscovy duck/Viet Nam/1914/2014) | c |
| MW934713 | 1773 | Avian | 4 (HA) | H5N6 | Viet Nam | T | 2018/12/05 | Influenza A virus (A/Muscovy duck/Vietnam/HN5135/2018) | c |
| MW935434 | 1773 | Avian | 4 (HA) | H5N6 | Viet Nam | T | 2018/12/05 | Influenza A virus (A/Muscovy duck/Vietnam/HN5137/2018) | c |
| MW935531 | 1776 | Avian | 4 (HA) | H5N6 | Viet Nam | T | 2018/12/05 | Influenza A virus (A/duck/Vietnam/HN5149/2018) | c |
| MW935546 | 1773 | Avian | 4 (HA) | H5N6 | Viet Nam | T | 2018/12/05 | Influenza A virus (A/Muscovy duck/Vietnam/HN5132/2018) | c |
| MW935607 | 1776 | Avian | 4 (HA) | H5N6 | Viet Nam | T | 2018/12/05 | Influenza A virus (A/duck/Vietnam/HN5142/2018) | c |
| MW935842 | 1776 | Avian | 4 (HA) | H5N6 | Viet Nam | T | 2018/12/05 | Influenza A virus (A/duck/Vietnam/HN5143/2018) | c |
| MW935851 | 1773 | Avian | 4 (HA) | H5N6 | Viet Nam | T | 2018/12/05 | Influenza A virus (A/Muscovy duck/Vietnam/HN5138/2018) | c |
| MW935867 | 1776 | Avian | 4 (HA) | H5N6 | Viet Nam | T | 2018/12/05 | Influenza A virus (A/duck/Vietnam/HN5147/2018) | c |
| MW935877 | 1776 | Avian | 4 (HA) | H5N6 | Viet Nam | T | 2018/12/05 | Influenza A virus (A/duck/Vietnam/HN5145/2018) | c |
| MW935923 | 1773 | Avian | 4 (HA) | H5N6 | Viet Nam | T | 2018/10/29 | Influenza A virus (A/Muscovy duck/Vietnam/HN5049/2018) | c |
| MW935969 | 1773 | Avian | 4 (HA) | H5N6 | Viet Nam | T | 2018/10/29 | Influenza A virus (A/Muscovy duck/Vietnam/HN5047/2018) | c |
| MW935980 | 1776 | Avian | 4 (HA) | H5N6 | Viet Nam | T | 2018/12/05 | Influenza A virus (A/duck/Vietnam/HN5141/2018) | c |
| MW936050 | 1773 | Avian | 4 (HA) | H5N6 | Viet Nam | T | 2018/10/29 | Influenza A virus (A/Muscovy duck/Vietnam/HN5048/2018) | c |
| MW936065 | 1773 | Avian | 4 (HA) | H5N6 | Viet Nam | T | 2018/08/10 | Influenza A virus (A/Muscovy duck/Vietnam/HN4857/2018) | c |
| MW936079 | 1773 | Avian | 4 (HA) | H5N6 | Viet Nam | T | 2018/08/10 | Influenza A virus (A/Muscovy duck/Vietnam/HN4858/2018) | c |
| MW936093 | 1773 | Avian | 4 (HA) | H5N6 | Viet Nam | T | 2018/10/29 | Influenza A virus (A/Muscovy duck/Vietnam/HN5046/2018) | c |
| MW936120 | 1773 | Avian | 4 (HA) | H5N6 | Viet Nam | T | 2018/08/10 | Influenza A virus (A/Muscovy duck/Vietnam/HN4856/2018) | c |
| MW936123 | 1773 | Avian | 4 (HA) | H5N6 | Viet Nam | T | 2018/12/05 | Influenza A virus (A/Muscovy duck/Vietnam/HN5139/2018) | c |
| MW936140 | 1773 | Avian | 4 (HA) | H5N6 | Viet Nam | T | 2018/11/05 | Influenza A virus (A/duck/Vietnam/HN5080/2018) | c |
| MW936233 | 1773 | Avian | 4 (HA) | H5N6 | Viet Nam | T | 2018/06/09 | Influenza A virus (A/Muscovy duck/Vietnam/HN4711/2018) | c |
| MW936246 | 1773 | Avian | 4 (HA) | H5N6 | Viet Nam | T | 2018/06/09 | Influenza A virus (A/duck/Vietnam/HN4721/2018) | c |
| MW936273 | 1773 | Avian | 4 (HA) | H5N6 | Viet Nam | T | 2018/06/09 | Influenza A virus (A/Muscovy duck/Vietnam/HN4719/2018) | c |
| MW936283 | 1773 | Avian | 4 (HA) | H5N6 | Viet Nam | T | 2018/06/09 | Influenza A virus (A/Muscovy duck/Vietnam/HN4714/2018) | c |
| MW936304 | 1773 | Avian | 4 (HA) | H5N6 | Viet Nam | T | 2018/06/09 | Influenza A virus (A/Muscovy duck/Vietnam/HN4716/2018) | c |
| MW936310 | 1773 | Avian | 4 (HA) | H5N6 | Viet Nam | T | 2018/06/09 | Influenza A virus (A/Muscovy duck/Vietnam/HN4720/2018) | c |
| MW936332 | 1773 | Avian | 4 (HA) | H5N6 | Viet Nam | T | 2018/06/09 | Influenza A virus (A/Muscovy duck/Vietnam/HN4715/2018) | c |
| MW936340 | 1773 | Avian | 4 (HA) | H5N6 | Viet Nam | T | 2018/06/09 | Influenza A virus (A/Muscovy duck/Vietnam/HN4718/2018) | c |
| MW936357 | 1773 | Avian | 4 (HA) | H5N6 | Viet Nam | T | 2018/06/09 | Influenza A virus (A/Muscovy duck/Vietnam/HN4713/2018) | c |
| MW960368 | 1730 | Avian | 4 (HA) | H5N8 | China | N | 2021/01 | Influenza A virus (A/Mute Swan/China/Shandong1/2021) | c |
| MZ166239 | 1704 | Avian | 4 (HA) | H5N8 | France | N | 2020/12/15 | Influenza A virus (A/Mule duck/France/20437/2020) | c |
| MZ166240 | 1704 | Avian | 4 (HA) | H5N8 | France | N | 2021/01/28 | Influenza A virus (A/Mule duck/France/21061/2021) | c |
| MZ166252 | 1704 | Avian | 4 (HA) | H5N8 | France | N | 2020/12/06 | Influenza A virus (A/Mule duck/France/20323/2020) | c |
| MZ166260 | 1704 | Avian | 4 (HA) | H5N8 | France | N | 2020/12/10 | Influenza A virus (A/Mule duck/France/20335/2020) | c |
| MZ166268 | 1704 | Avian | 4 (HA) | H5N8 | France | N | 2020/12/13 | Influenza A virus (A/Mule duck/France/20338/2020) | c |
| MZ166276 | 1704 | Avian | 4 (HA) | H5N8 | France | N | 2020/12/13 | Influenza A virus (A/Mule duck/France/20339/2020) | c |
| MZ166284 | 1704 | Avian | 4 (HA) | H5N8 | France | N | 2020/12/21 | Influenza A virus (A/Mule duck/France/20349/2020) | c |
| MZ166292 | 1704 | Avian | 4 (HA) | H5N8 | France | N | 2020/12/26 | Influenza A virus (A/Mule duck/France/20352/2020) | c |
| MZ166300 | 1704 | Avian | 4 (HA) | H5N8 | France | N | 2020/12/27 | Influenza A virus (A/Mule duck/France/20353/2020) | c |
| MZ166308 | 1704 | Avian | 4 (HA) | H5N8 | France | N | 2021/01/30 | Influenza A virus (A/Mule duck/France/21064/2021) | c |
| MZ166316 | 1704 | Avian | 4 (HA) | H5N8 | France | N | 2021/02/20 | Influenza A virus (A/Mule duck/France/21084/2021) | c |
| MZ191540 | 1704 | Avian | 4 (HA) | H5N6 | Viet Nam | T | 2015/08/21 | Influenza A virus (A/Moscovy duck/Viet Nam/4915 HA/2015) | c |
| MZ235334 | 1704 | Avian | 4 (HA) | H5N8 | Japan | N | 2021/02/03 | Influenza A virus (A/white-tailed eagle/Hokkaido/20210127001/2021) | c |
| MZ620718 | 1704 | Avian | 4 (HA) | H5N8 | Japan | N | 2020/12/22 | Influenza A virus (A/Mandarin duck/Kagoshima/d57/2020) | c |
| OK583834 | 1701 | Avian | 4 (HA) | H5N6 | China | N | 2018/10/26 | Influenza A virus (A/duck/Jiangsu/JS80/2018) | c |
| OK583835 | 1701 | Avian | 4 (HA) | H5N6 | China | N | 2018/10/26 | Influenza A virus (A/goose/Jiangsu/JS03/2018) | c |
| OK583836 | 1701 | Avian | 4 (HA) | H5N6 | China | N | 2019/01/26 | Influenza A virus (A/goose/Jiangsu/JS06/2019) | c |
| OK583837 | 1701 | Avian | 4 (HA) | H5N6 | China | N | 2018/03 | Influenza A virus (A/chicken/Shandong/SD25/2018) | c |
| OK583838 | 1701 | Avian | 4 (HA) | H5N6 | China | N | 2018/01 | Influenza A virus (A/chicken/Shandong/SD13/2018) | c |
| OK583839 | 1701 | Avian | 4 (HA) | H5N6 | China | N | 2018/03 | Influenza A virus (A/chicken/Shandong/SD78/2018) | c |
| OL354495 | 1777 | Avian | 4 (HA) | H5N8 | Egypt | N | 2018/02/05 | Influenza A virus (A/chicken/Egypt/N15177A/2018) | c |
| OL354772 | 1777 | Avian | 4 (HA) | H5N8 | Egypt | N | 2018/02/05 | Influenza A virus (A/chicken/Egypt/N15177D/2018) | c |
| OL354818 | 1777 | Avian | 4 (HA) | H5N8 | Egypt | N | 2018/03/20 | Influenza A virus (A/chicken/Egypt/F15366B/2018) | c |
| OL354826 | 1777 | Avian | 4 (HA) | H5N8 | Egypt | N | 2018/02/05 | Influenza A virus (A/chicken/Egypt/N15175A/2018) | c |
| OL354834 | 1777 | Avian | 4 (HA) | H5N8 | Egypt | N | 2018/02/05 | Influenza A virus (A/chicken/Egypt/N15174B/2018) | c |
| OL354842 | 1777 | Avian | 4 (HA) | H5N8 | Egypt | N | 2018/02/05 | Influenza A virus (A/chicken/Egypt/N15175B/2018) | c |
| OL354865 | 1777 | Avian | 4 (HA) | H5N8 | Egypt | N | 2018/02/05 | Influenza A virus (A/chicken/Egypt/N15175D/2018) | c |
| OL354888 | 1777 | Avian | 4 (HA) | H5N8 | Egypt | N | 2018/01/14 | Influenza A virus (A/duck/Egypt/F15089/2018) | c |
| OL354899 | 1744 | Avian | 4 (HA) | H5N8 | Egypt | N | 2021/02/10 | Influenza A virus (A/chicken/Egypt/A19673/2021) | c |
| OL354917 | 1777 | Avian | 4 (HA) | H5N8 | Egypt | N | 2018/01/14 | Influenza A virus (A/duck/Egypt/F15092/2018) | c |
| OL354925 | 1777 | Avian | 4 (HA) | H5N8 | Egypt | N | 2018/11/24 | Influenza A virus (A/duck/Egypt/A16368/2018) | c |
| OL354942 | 1777 | Avian | 4 (HA) | H5N8 | Egypt | N | 2018/01/15 | Influenza A virus (A/chicken/Egypt/A15044/2018) | c |
| OL354965 | 1777 | Avian | 4 (HA) | H5N8 | Egypt | N | 2018/11/24 | Influenza A virus (A/duck/Egypt/A16372/2018) | c |
| OL354974 | 1777 | Avian | 4 (HA) | H5N8 | Egypt | N | 2018/01/15 | Influenza A virus (A/chicken/Egypt/A15037/2018) | c |
| OL354982 | 1777 | Avian | 4 (HA) | H5N8 | Egypt | N | 2021/01/26 | Influenza A virus (A/Duck/Egypt/A19643/2021) | c |
| OL366043 | 1744 | Avian | 4 (HA) | H5N8 | Egypt | N | 2021/02/10 | Influenza A virus (A/chicken/Egypt/A19671/2021) | c |
| OL467327 | 1704 | Avian | 4 (HA) | H5N8 | Egypt | N | 2018 | Influenza A virus (A/Duck/Egypt/BA3C/2018) | p |
| OL467340 | 1748 | Avian | 4 (HA) | H5N8 | Egypt | N | 2018 | Influenza A virus (A/goose/Egypt/BA71C/2018) | c |
| OL519551 | 1704 | Human | 4 (HA) | H5N6 | China | N | 2021/09/19 | Influenza A virus (A/Guangdong/lgf/2021) | c |
| OL638145 | 1742 | Avian | 4 (HA) | H5N1 | Czech Republ | N | 2021/09/27 | Influenza A virus (A/goose/Czech Republic/18520-2/2021) | c |
| OM060672 | 1751 | Avian | 4 (HA) | H5N8 | South Korea | N | 2014/11/15 | Influenza A virus (A/Duck/South Korea/D14-19/2014) | c |
| OM333435 | 1704 | Avian | 4 (HA) | H5N8 | Egypt | N | 2017/02/14 | Influenza A virus (A/chicken/Egypt/N13731C/2017) | c |
| OM373230 | 1704 | Avian | 4 (HA) | H5N8 | China | N | 2020/12/31 | Influenza A virus (A/eurasian coot/Shandong/W5611/2020) | c |
| OM373238 | 1704 | Avian | 4 (HA) | H5N8 | China | N | 2020/12/31 | Influenza A virus (A/eurasian coot/Shandong/W6137/2020) | c |
| OM373246 | 1704 | Avian | 4 (HA) | H5N8 | China | N | 2020/12/31 | Influenza A virus (A/eurasian coot/Shandong/W6143/2020) | c |
| OM373254 | 1704 | Avian | 4 (HA) | H5N8 | China | N | 2020/12/31 | Influenza A virus (A/eurasian coot/Shandong/W6150/2020) | c |
| OM373310 | 1704 | Avian | 4 (HA) | H5N8 | China | N | 2020/11/13 | Influenza A virus (A/wild duck/Shandong/W3580/2020) | c |
| OM403993 | 1695 | Reassortant | 4 (HA) | H5N8 | | | 1934 | Influenza A virus (A/reassortant/IDCDC-RG71A(Astrakhan/3212/2020 X Puerto Rico/ | c |
| ON024679 | 1588 | Avian | 4 (HA) | H5N8 | Egypt | N | 2020/01 | Influenza A virus (A/chicken/Egypt/AF12/2020) | p |
| ON024680 | 1588 | Avian | 4 (HA) | H5N8 | Egypt | N | 2020/02 | Influenza A virus (A/chicken/Egypt/AF14/2020) | p |
| ON024681 | 1588 | Avian | 4 (HA) | H5N8 | Egypt | N | 2020/02 | Influenza A virus (A/duck/Egypt/AI2/2020) | p |
| ON024682 | 1588 | Avian | 4 (HA) | H5N8 | Egypt | N | 2019/12 | Influenza A virus (A/chicken/Egypt/AI9/2019) | p |

| | | | | | | | | | |
|---|---|---|---|---|---|---|---|---|---|
| ON024683 | 1588 | Avian | 4 (HA) | H5N8 | Egypt | N | 2020/01 | Influenza A virus (A/duck/Egypt/FAOS1/2020) | p |
| ON024684 | 1588 | Avian | 4 (HA) | H5N8 | Egypt | N | 2020/02 | Influenza A virus (A/duck/Egypt/FAOS13/2020) | p |
| ON024685 | 1588 | Avian | 4 (HA) | H5N8 | Egypt | N | 2020/03 | Influenza A virus (A/chicken/Egypt/FAOS18/2020) | p |
| ON024686 | 1588 | Avian | 4 (HA) | H5N8 | Egypt | N | 2020/03 | Influenza A virus (A/chicken/Egypt/FAOS20/2020) | p |
| ON024687 | 1588 | Avian | 4 (HA) | H5N8 | Egypt | N | 2020/03 | Influenza A virus (A/turkey/Egypt/FAOS24/2020) | p |
| ON024688 | 1588 | Avian | 4 (HA) | H5N8 | Egypt | N | 2020/01 | Influenza A virus (A/turkey/Egypt/FAOS5/2020) | p |
| ON024689 | 1588 | Avian | 4 (HA) | H5N8 | Egypt | N | 2020/01 | Influenza A virus (A/duck/Egypt/FAOSG3/2020) | p |
| ON024690 | 1588 | Avian | 4 (HA) | H5N8 | Egypt | N | 2020/01 | Influenza A virus (A/turkey/Egypt/FAOSG5/2020) | p |
| ON024691 | 1588 | Avian | 4 (HA) | H5N8 | Egypt | N | 2020/11 | Influenza A virus (A/turkey/Egypt/FAOSG6/2020) | p |
| ON024692 | 1588 | Avian | 4 (HA) | H5N8 | Egypt | N | 2020/02 | Influenza A virus (A/chicken/Egypt/FAOSG11/2020) | p |
| ON024693 | 1588 | Avian | 4 (HA) | H5N8 | Egypt | N | 2020/02 | Influenza A virus (A/duck/Egypt/FAOSG14/2020) | p |
| ON024694 | 1588 | Avian | 4 (HA) | H5N8 | Egypt | N | 2020/01 | Influenza A virus (A/chicken/Egypt/VG37/2020) | p |
| ON024695 | 1588 | Avian | 4 (HA) | H5N8 | Egypt | N | 2019/10 | Influenza A virus (A/chicken/Egypt/AL7/2019) | p |
| ON024696 | 1588 | Avian | 4 (HA) | H5N8 | Egypt | N | 2019/02 | Influenza A virus (A/chicken/Egypt/AL2/2019) | p |
| ON024697 | 1588 | Avian | 4 (HA) | H5N8 | Egypt | N | 2019/10 | Influenza A virus (A/chicken/Egypt/AL6/2019) | p |
| ON024698 | 1588 | Avian | 4 (HA) | H5N8 | Egypt | N | 2019/12 | Influenza A virus (A/chicken/Egypt/F565/2019) | p |
| ON024699 | 1588 | Avian | 4 (HA) | H5N8 | Egypt | N | 2019/06 | Influenza A virus (A/duck/Egypt/FAO-S6/2019) | p |
| ON024700 | 1588 | Avian | 4 (HA) | H5N8 | Egypt | N | 2019/05 | Influenza A virus (A/turkey/Egypt/FAO-S10/2019) | p |
| ON024701 | 1588 | Avian | 4 (HA) | H5N8 | Egypt | N | 2019/06 | Influenza A virus (A/duck/Egypt/FAO-S17/2019) | p |
| ON024702 | 1588 | Avian | 4 (HA) | H5N8 | Egypt | N | 2019/06 | Influenza A virus (A/turkey/Egypt/FAO-S18/2019) | p |
| ON024703 | 1588 | Avian | 4 (HA) | H5N8 | Egypt | N | 2019/08 | Influenza A virus (A/duck/Egypt/FAO-S24/2019) | p |
| ON024704 | 1588 | Avian | 4 (HA) | H5N8 | Egypt | N | 2019/05 | Influenza A virus (A/duck/Egypt/FAO-S32/2019) | p |
| ON024705 | 1588 | Avian | 4 (HA) | H5N8 | Egypt | N | 2019/05 | Influenza A virus (A/duck/Egypt/FAO-S33/2019) | p |
| ON024706 | 1588 | Avian | 4 (HA) | H5N8 | Egypt | N | 2019/07 | Influenza A virus (A/duck/Egypt/FAO-S37/2019) | p |
| ON024707 | 1588 | Avian | 4 (HA) | H5N8 | Egypt | N | 2019/09 | Influenza A virus (A/turkey/Egypt/FAO-S42/2019) | p |
| ON024708 | 1588 | Avian | 4 (HA) | H5N8 | Egypt | N | 2019/11 | Influenza A virus (A/duck/Egypt/FAO-S76/2019) | p |
| ON024709 | 1588 | Avian | 4 (HA) | H5N8 | Egypt | N | 2019/09 | Influenza A virus (A/chicken/Egypt/FAO-SG21/2019) | p |
| ON024710 | 1588 | Avian | 4 (HA) | H5N8 | Egypt | N | 2019/07 | Influenza A virus (A/chicken/Egypt/FAO-SG10/2019) | p |
| ON024711 | 1588 | Avian | 4 (HA) | H5N8 | Egypt | N | 2019/07 | Influenza A virus (A/duck/Egypt/FAOSG11/2019) | p |
| ON024712 | 1588 | Avian | 4 (HA) | H5N8 | Egypt | N | 2019/07 | Influenza A virus (A/turkey/Egypt/FAOSG22/2019) | p |
| ON024713 | 1588 | Avian | 4 (HA) | H5N8 | Egypt | N | 2019/03 | Influenza A virus (A/duck/Egypt/SD36/2019) | p |
| ON024714 | 1588 | Avian | 4 (HA) | H5N8 | Egypt | N | 2019/02 | Influenza A virus (A/duck/Egypt/SM6G/2019) | p |
| ON024715 | 1588 | Avian | 4 (HA) | H5N8 | Egypt | N | 2019/02 | Influenza A virus (A/duck/Egypt/SMG7/2019) | p |
| ON024716 | 1588 | Avian | 4 (HA) | H5N8 | Egypt | N | 2019/01 | Influenza A virus (A/chicken/Egypt/V1345/2019) | p |
| ON024717 | 1588 | Avian | 4 (HA) | H5N8 | Egypt | N | 2021/04 | Influenza A virus (A/turkey/Egypt/A2/2021) | p |
| ON024718 | 1588 | Avian | 4 (HA) | H5N8 | Egypt | N | 2021/04 | Influenza A virus (A/chicken/Egypt/A4/2021) | p |
| ON024719 | 1588 | Avian | 4 (HA) | H5N8 | Egypt | N | 2021/02 | Influenza A virus (A/duck/Egypt/F91/2021) | p |
| ON024720 | 1588 | Avian | 4 (HA) | H5N8 | Egypt | N | 2021/03 | Influenza A virus (A/chicken/Egypt/F304/2021) | p |
| ON024721 | 1588 | Avian | 4 (HA) | H5N8 | Egypt | N | 2021/04 | Influenza A virus (A/turkey/Egypt/FAOS18/2021) | p |
| ON024723 | 1588 | Avian | 4 (HA) | H5N8 | Egypt | N | 2021/01 | Influenza A virus (A/turkey/Egypt/FAOSL3/2021) | p |
| ON024724 | 1588 | Avian | 4 (HA) | H5N8 | Egypt | N | 2021/01 | Influenza A virus (A/duck/Egypt/FAOSL4/2021) | p |
| ON024725 | 1588 | Avian | 4 (HA) | H5N8 | Egypt | N | 2021/04 | Influenza A virus (A/duck/Egypt/FAOSI11/2021) | p |
| ON260842 | 1692 | Reassortant | 4 (HA) | H5N6 | | | 1934 | Influenza A virus (A/reassortant/IDCDC-RG65A(Guangdong/18SF020/2018 X Puerto I | c |
| ON716427 | 1704 | Avian | 4 (HA) | H5N8 | Cambodia | T | 2021/09/16 | Influenza A virus (A/Duck/Cambodia/f1PPOreu241D3_C/2021) | c |
| ON716428 | 1704 | Avian | 4 (HA) | H5N8 | Cambodia | T | 2021/09/16 | Influenza A virus (A/Duck/Cambodia/f1PPChba241D6/2021) | c |
| ON716429 | 1704 | Avian | 4 (HA) | H5N8 | Cambodia | T | 2021/10/15 | Influenza A virus (A/Duck/Cambodia/f4K241D3_C/2021) | c |
| ON716430 | 1704 | Avian | 4 (HA) | H5N8 | Cambodia | T | 2021/10/15 | Influenza A virus (A/Duck/Cambodia/f4K241D4/2021) | c |
| ON716431 | 1704 | Avian | 4 (HA) | H5N8 | Cambodia | T | 2021/11/16 | Influenza A virus (A/Duck/Cambodia/f6T241D4/2021) | c |
| ON716432 | 1701 | Avian | 4 (HA) | H5N6 | Cambodia | T | 2019/01/29 | Influenza A virus (A/chicken/Cambodia/c9T241C17T/2019) | c |
| ON716433 | 1701 | Avian | 4 (HA) | H5N6 | Cambodia | T | 2018/12/13 | Influenza A virus (A/Duck/Cambodia/c18MKAP189/2018) | c |
| ON716434 | 1701 | Avian | 4 (HA) | H5N6 | Cambodia | T | 2018/12/13 | Influenza A virus (A/Duck/Cambodia/c18MKAP211/2018) | c |
| ON716435 | 1701 | Avian | 4 (HA) | H5N6 | Cambodia | T | 2018/12/13 | Influenza A virus (A/Duck/Cambodia/c18MKAP214/2018) | c |
| ON716436 | 1701 | Avian | 4 (HA) | H5N6 | Cambodia | T | 2020/09/12 | Influenza A virus (A/Duck/Cambodia/e10T241C18/2020) | c |
| ON716437 | 1704 | Avian | 4 (HA) | H5N6 | Cambodia | T | 2020/06/13 | Influenza A virus (A/Duck/Cambodia/e5PPOreu241D3/2020) | c |
| ON716438 | 1704 | Avian | 4 (HA) | H5N6 | Cambodia | T | 2020/06/13 | Influenza A virus (A/Duck/Cambodia/e5PPOreu241D4/2020) | c |
| ON716439 | 1704 | Avian | 4 (HA) | H5N6 | Cambodia | T | 2020/06/13 | Influenza A virus (A/Duck/Cambodia/e5PPOreu241D8/2020) | c |
| ON716440 | 1704 | Avian | 4 (HA) | H5N6 | Cambodia | T | 2020/08/21 | Influenza A virus (A/Duck/Cambodia/e8T241D10/2020) | c |
| ON716441 | 1704 | Avian | 4 (HA) | H5N6 | Cambodia | T | 2020/08/21 | Influenza A virus (A/Duck/Cambodia/e8T241D11/2020) | c |
| ON716442 | 1704 | Avian | 4 (HA) | H5N6 | Cambodia | T | 2020/08/21 | Influenza A virus (A/Duck/Cambodia/e8T241D15/2020) | c |
| ON716443 | 1704 | Avian | 4 (HA) | H5N6 | Cambodia | T | 2020/08/21 | Influenza A virus (A/Duck/Cambodia/e8T241D16/2020) | c |
| ON716444 | 1704 | Avian | 4 (HA) | H5N6 | Cambodia | T | 2020/08/21 | Influenza A virus (A/Duck/Cambodia/e8T241D5/2020) | c |
| ON716445 | 1704 | Avian | 4 (HA) | H5N6 | Cambodia | T | 2020/08/21 | Influenza A virus (A/Duck/Cambodia/e8T241D6/2020) | c |
| ON716446 | 1704 | Avian | 4 (HA) | H5N6 | Cambodia | T | 2020/08/21 | Influenza A virus (A/Duck/Cambodia/e8T241D9/2020) | c |
| OP209769 | 1704 | Human | 4 (HA) | H5N6 | China | N | 2022/02/26 | Influenza A virus (A/Yangzhou/125/2022) | c |
| OP221343 | 1751 | Avian | 4 (HA) | H5N1 | USA | N | 2022/03/10 | Influenza A virus (A/American pelican/Kansas/W22-200/2022) | c |
| OP377338 | 1751 | Avian | 4 (HA) | H5N1 | USA | N | 2022/06/30 | Influenza A virus (A/black vulture/Virginia/W22-499C/2022) | c |
| OP377388 | 1751 | Avian | 4 (HA) | H5N1 | USA | N | 2022/06/30 | Influenza A virus (A/black vulture/Virginia/W22-499A/2022) | c |
| OP377412 | 1751 | Avian | 4 (HA) | H5N1 | USA | N | 2022/03/17 | Influenza A virus (A/royal tern/Florida/W22-245B/2022) | c |
| OP377453 | 1751 | Avian | 4 (HA) | H5N1 | USA | N | 2022/03/07 | Influenza A virus (A/black vulture/Florida/W22-168/2022) | c |
| OP377486 | 1751 | Avian | 4 (HA) | H5N1 | USA | N | 2022/02/25 | Influenza A virus (A/black vulture/Florida/W22-161/2022) | c |
| OP377494 | 1751 | Avian | 4 (HA) | H5N1 | USA | N | 2022/06/30 | Influenza A virus (A/black vulture/Virginia/W22-499B/2022) | c |
| OP377510 | 1751 | Avian | 4 (HA) | H5N1 | USA | N | 2022/02/24 | Influenza A virus (A/great horned owl/Flordia/W22-163A/2022) | c |
| OP377542 | 1751 | Avian | 4 (HA) | H5N1 | USA | N | 2022/02/27 | Influenza A virus (A/great blue heron/Florida/W22-160/2022) | c |
| OP377558 | 1751 | Avian | 4 (HA) | H5N1 | USA | N | 2022/05/10 | Influenza A virus (A/black vulture/North Carolina/W22-367C/2022) | c |
| OP377574 | 1751 | Avian | 4 (HA) | H5N1 | USA | N | 2022/03/04 | Influenza A virus (A/black vulture/Florida/W22-167/2022) | c |
| OP377606 | 1751 | Avian | 4 (HA) | H5N1 | USA | N | 2022/02/20 | Influenza A virus (A/brown pelican/North Carolina/W22-164/2022) | c |
| OP377614 | 1751 | Avian | 4 (HA) | H5N1 | USA | N | 2022/03/12 | Influenza A virus (A/royal tern/Florida/W22-245A/2022) | c |
| OP377638 | 1751 | Avian | 4 (HA) | H5N1 | USA | N | 2022/02/23 | Influenza A virus (A/great horned owl/Flordia/W22-163C/2022) | c |
| OP413016 | 1777 | Avian | 4 (HA) | H5N8 | Egypt | N | 2021/03/03 | Influenza A virus (A/chicken/Egypt/Menoufia/2021) | c |
| OP413044 | 1701 | Avian | 4 (HA) | H5N1 | Egypt | N | 2021/10/07 | Influenza A virus (A/migratory wild bird/Egypt/25/2021) | nc |
| OP413045 | 1701 | Avian | 4 (HA) | H5N1 | Egypt | N | 2021/11/22 | Influenza A virus (A/migratory wild bird/Egypt/26/2021) | nc |
| OP413046 | 1701 | Avian | 4 (HA) | H5N1 | Egypt | N | 2022/03/25 | Influenza A virus (A/migratory wild bird/Egypt/30/2022) | nc |
| OP413047 | 1701 | Avian | 4 (HA) | H5N1 | Egypt | N | 2022/03/25 | Influenza A virus (A/migratory wild bird/Egypt/31/2022) | nc |

| | | | | | | | | | |
|---|---|---|---|---|---|---|---|---|---|
| OP413048 | 1701 | Avian | 4 (HA) | H5N1 | Egypt | N | 2022/03/25 | Influenza A virus (A/migratory wild bird/Egypt/32/2022) | nc |
| OP470716 | 1751 | Avian | 4 (HA) | H5N1 | USA | N | 2022/02/12 | Influenza A virus (A/lesser scaup/Georgia/W22-145C/2022) | c |
| OP470733 | 1751 | Avian | 4 (HA) | H5N1 | USA | N | 2022/02/12 | Influenza A virus (A/bald eagle/Florida/W22-134-CL/2022) | c |
| OP470749 | 1751 | Avian | 4 (HA) | H5N1 | USA | N | 2022/02/12 | Influenza A virus (A/lesser scaup/Georgia/W22-145B/2022) | c |
| OP470757 | 1751 | Avian | 4 (HA) | H5N1 | USA | N | 2022/02/12 | Influenza A virus (A/lesser scaup/Georgia/W22-145D/2022) | c |
| OP470765 | 1751 | Avian | 4 (HA) | H5N1 | USA | N | 2022/02/12 | Influenza A virus (A/lesser scaup/Georgia/W22-143/2022) | c |
| OP470773 | 1751 | Avian | 4 (HA) | H5N1 | USA | N | 2022/02/12 | Influenza A virus (A/lesser scaup/Georgia/W22-145A/2022) | c |
| OP470781 | 1751 | Avian | 4 (HA) | H5N1 | USA | N | 2022/02/12 | Influenza A virus (A/lesser scaup/Georgia/W22-145E/2022) | c |
| OP470789 | 1751 | Avian | 4 (HA) | H5N1 | USA | N | 2022/02/08 | Influenza A virus (A/snow goose/Kentucky/W22-092/2022) | c |
| OP470797 | 1751 | Avian | 4 (HA) | H5N1 | USA | N | 2022/02/12 | Influenza A virus (A/bald eagle/Florida/W22-142/2022) | c |
| OP470804 | 1751 | Avian | 4 (HA) | H5N1 | USA | N | 2022/02/17 | Influenza A virus (A/bald eagle/FL/W22-114/2022) | c |
| OP470812 | 1751 | Avian | 4 (HA) | H5N1 | USA | N | 2022/02/12 | Influenza A virus (A/lesser scaup/Florida/W22-129A/2022) | c |
| OP484862 | 1777 | Avian | 4 (HA) | H5N5 | Egypt | N | 2016/12/21 | Influenza A virus (A/purple heron/Egypt/MB 933C/2016) | c |
| OP597554 | 1735 | Avian | 4 (HA) | H5N5 | Russia | N | 2021/04/06 | Influenza A virus (A/dalmatian pelican/Astrakhan/417-1/2021) | c |
| OP597562 | 1735 | Avian | 4 (HA) | H5N5 | Russia | N | 2021/04/06 | Influenza A virus (A/dalmatian pelican/Astrakhan/417-2/2021) | c |
| OP597570 | 1735 | Avian | 4 (HA) | H5N5 | Russia | N | 2021/04/01 | Influenza A virus (A/pelican/Dagestan/397-1/2021) | c |
| OP597578 | 1735 | Avian | 4 (HA) | H5N5 | Russia | N | 2021/04/01 | Influenza A virus (A/gull/Dagestan/397-2/2021) | c |
| OP597586 | 1735 | Avian | 4 (HA) | H5N5 | Russia | N | 2020/12/29 | Influenza A virus (A/swan/Rostov/2299-2/2020) | c |
| OP597594 | 1735 | Avian | 4 (HA) | H5N5 | Russia | N | 2021/11/08 | Influenza A virus (A/shelduck/Kalmykia/1814-1/2021) | c |
| OP597602 | 1735 | Avian | 4 (HA) | H5N5 | Russia | N | 2021/09/28 | Influenza A virus (A/waterfowl/Russia/1526-4/2021) | c |
| OP748410 | 1704 | Avian | 4 (HA) | H5N8 | China | N | 2020/12 | Influenza A virus (A/chicken/China/JM01/2020) | c |
| OQ195215 | 1701 | Environment | 4 (HA) | H5 | China | N | 2020/06/22 | Influenza A virus (A/Environment/China/H5N6/2020) | c |
| OQ195223 | 1701 | Avian | 4 (HA) | H5 | China | N | 2020/08/19 | Influenza A virus (A/duck/China/H5N6/2020) | c |
| OQ195231 | 1701 | Avian | 4 (HA) | H5 | China | N | 2020/06/22 | Influenza A virus (A/Duck/China/H5N6/2020) | c |

### GISAID

| Isolate_Id | Isolate_Name | Subtype | Location | Isolate_Submitter |
|---|---|---|---|---|
| EPI_ISL_1005680 | A/common teal/Chany Lake/213/2020 | A / H5N2 | Europe / Russian Federation / Novosibirsk | Takehiko Saito (National Institute of Animal Health) |
| EPI_ISL_1009679 | A/duck/Korea/H007/2020 | A / H5N8 | Asia / Korea, Republic of / CN | Yu-Na Lee (Animal and Plant Quarantine Agency (S-2158) / Avian Influenza Research & Diagnostic Division) |
| EPI_ISL_1009680 | A/White peacock/Korea/H533/2020 | A / H5N8 | Asia / Korea, Republic of / GG | Yu-Na Lee (Animal and Plant Quarantine Agency (S-2158) / Avian Influenza Research & Diagnostic Division) |
| EPI_ISL_1009681 | A/chicken/Korea/H541/2020 | A / H5N8 | Asia / Korea, Republic of / CN | Yu-Na Lee (Animal and Plant Quarantine Agency (S-2158) / Avian Influenza Research & Diagnostic Division) |
| EPI_ISL_1009682 | A/chicken/Korea/H544/2020 | A / H5N8 | Asia / Korea, Republic of / GG | Yu-Na Lee (Animal and Plant Quarantine Agency (S-2158) / Avian Influenza Research & Diagnostic Division) |
| EPI_ISL_1009683 | A/duck/Korea/H542/2020 | A / H5N8 | Asia / Korea, Republic of / JN | Yu-Na Lee (Animal and Plant Quarantine Agency (S-2158) / Avian Influenza Research & Diagnostic Division) |
| EPI_ISL_1009684 | A/duck/Korea/H548/2020 | A / H5N8 | Asia / Korea, Republic of / JB | Yu-Na Lee (Animal and Plant Quarantine Agency (S-2158) / Avian Influenza Research & Diagnostic Division) |
| EPI_ISL_1009685 | A/duck/Korea/H549/2020 | A / H5N8 | Asia / Korea, Republic of / JB | Yu-Na Lee (Animal and Plant Quarantine Agency (S-2158) / Avian Influenza Research & Diagnostic Division) |
| EPI_ISL_1009686 | A/chicken/Korea/H550/2020 | A / H5N8 | Asia / Korea, Republic of / GG | Yu-Na Lee (Animal and Plant Quarantine Agency (S-2158) / Avian Influenza Research & Diagnostic Division) |
| EPI_ISL_1009687 | A/quail/Korea/H551/2020 | A / H5N8 | Asia / Korea, Republic of / GB | Yu-Na Lee (Animal and Plant Quarantine Agency (S-2158) / Avian Influenza Research & Diagnostic Division) |
| EPI_ISL_1009688 | A/chicken/Korea/H001/2021 | A / H5N8 | Asia / Korea, Republic of / JN | Yu-Na Lee (Animal and Plant Quarantine Agency (S-2158) / Avian Influenza Research & Diagnostic Division) |
| EPI_ISL_1009689 | A/chicken/Korea/H002/2021 | A / H5N8 | Asia / Korea, Republic of / CN | Yu-Na Lee (Animal and Plant Quarantine Agency (S-2158) / Avian Influenza Research & Diagnostic Division) |
| EPI_ISL_1009690 | A/chicken/Korea/H008/2021 | A / H5N8 | Asia / Korea, Republic of / CN | Yu-Na Lee (Animal and Plant Quarantine Agency (S-2158) / Avian Influenza Research & Diagnostic Division) |
| EPI_ISL_1009691 | A/duck/Korea/H009/2021 | A / H5N8 | Asia / Korea, Republic of / JB | Yu-Na Lee (Animal and Plant Quarantine Agency (S-2158) / Avian Influenza Research & Diagnostic Division) |
| EPI_ISL_1009692 | A/duck/Korea/H010/2021 | A / H5N8 | Asia / Korea, Republic of / CB | Yu-Na Lee (Animal and Plant Quarantine Agency (S-2158) / Avian Influenza Research & Diagnostic Division) |
| EPI_ISL_1009693 | A/chicken/Korea/H022/2021 | A / H5N8 | Asia / Korea, Republic of / GG | Yu-Na Lee (Animal and Plant Quarantine Agency (S-2158) / Avian Influenza Research & Diagnostic Division) |
| EPI_ISL_1009694 | A/duck/Korea/H025/2021 | A / H5N8 | Asia / Korea, Republic of / GN | Yu-Na Lee (Animal and Plant Quarantine Agency (S-2158) / Avian Influenza Research & Diagnostic Division) |
| EPI_ISL_1009695 | A/spot-billed duck/Korea/WA612/2020 | A / H5N8 | Asia / Korea, Republic of / CN | Yu-Na Lee (Animal and Plant Quarantine Agency (S-2158) / Avian Influenza Research & Diagnostic Division) |
| EPI_ISL_1009696 | A/mandarin duck/Korea/WB80/2020 | A / H5N8 | Asia / Korea, Republic of / GG | Yu-Na Lee (Animal and Plant Quarantine Agency (S-2158) / Avian Influenza Research & Diagnostic Division) |
| EPI_ISL_1009697 | A/wild bird/Korea/H357/2020 | A / H5N8 | Asia / Korea, Republic of / JB | Yu-Na Lee (Animal and Plant Quarantine Agency (S-2158) / Avian Influenza Research & Diagnostic Division) |
| EPI_ISL_1009698 | A/mallard/Korea/WA820/2020 | A / H5N8 | Asia / Korea, Republic of / GG | Yu-Na Lee (Animal and Plant Quarantine Agency (S-2158) / Avian Influenza Research & Diagnostic Division) |
| EPI_ISL_1009699 | A/mandarin duck/Korea/WA831/2020 | A / H5N8 | Asia / Korea, Republic of / JB | Yu-Na Lee (Animal and Plant Quarantine Agency (S-2158) / Avian Influenza Research & Diagnostic Division) |
| EPI_ISL_1009700 | A/spot-billed duck/Korea/WA854/2020 | A / H5N8 | Asia / Korea, Republic of / GG | Yu-Na Lee (Animal and Plant Quarantine Agency (S-2158) / Avian Influenza Research & Diagnostic Division) |
| EPI_ISL_1009701 | A/mandarin duck/Korea/WA857/2020 | A / H5N8 | Asia / Korea, Republic of / JB | Yu-Na Lee (Animal and Plant Quarantine Agency (S-2158) / Avian Influenza Research & Diagnostic Division) |
| EPI_ISL_1009702 | A/mandarin duck/Korea/WA877/2020 | A / H5N8 | Asia / Korea, Republic of / JN | Yu-Na Lee (Animal and Plant Quarantine Agency (S-2158) / Avian Influenza Research & Diagnostic Division) |
| EPI_ISL_1009703 | A/spot-billed duck/Korea/WA889/2020 | A / H5N8 | Asia / Korea, Republic of / GG | Yu-Na Lee (Animal and Plant Quarantine Agency (S-2158) / Avian Influenza Research & Diagnostic Division) |
| EPI_ISL_1009704 | A/mandarin duck/Korea/WA899/2020 | A / H5N8 | Asia / Korea, Republic of / CN | Yu-Na Lee (Animal and Plant Quarantine Agency (S-2158) / Avian Influenza Research & Diagnostic Division) |
| EPI_ISL_1009705 | A/mandarin duck/Korea/WA913/2020 | A / H5N8 | Asia / Korea, Republic of / CN | Yu-Na Lee (Animal and Plant Quarantine Agency (S-2158) / Avian Influenza Research & Diagnostic Division) |
| EPI_ISL_1009706 | A/wild bird/Korea/H379/2020 | A / H5N8 | Asia / Korea, Republic of / GB | Yu-Na Lee (Animal and Plant Quarantine Agency (S-2158) / Avian Influenza Research & Diagnostic Division) |
| EPI_ISL_1009707 | A/wild bird/Korea/H467/2020 | A / H5N8 | Asia / Korea, Republic of / CB | Yu-Na Lee (Animal and Plant Quarantine Agency (S-2158) / Avian Influenza Research & Diagnostic Division) |
| EPI_ISL_1009708 | A/spot-billed duck/Korea/WA1000/2020 | A / H5N8 | Asia / Korea, Republic of / GG | Yu-Na Lee (Animal and Plant Quarantine Agency (S-2158) / Avian Influenza Research & Diagnostic Division) |
| EPI_ISL_1009709 | A/wild bird/Korea/H496-3/2020 | A / H5N8 | Asia / Korea, Republic of / JJ | Yu-Na Lee (Animal and Plant Quarantine Agency (S-2158) / Avian Influenza Research & Diagnostic Division) |
| EPI_ISL_1009710 | A/wild duck/Korea/H331/2020 | A / H5N8 | Asia / Korea, Republic of / JJ | Yu-Na Lee (Animal and Plant Quarantine Agency (S-2158) / Avian Influenza Research & Diagnostic Division) |
| EPI_ISL_1009711 | A/duck/Korea/H016/2021 | A / H5N8 | Asia / Korea, Republic of / JN | Yu-Na Lee (Animal and Plant Quarantine Agency (S-2158) / Avian Influenza Research & Diagnostic Division) |
| EPI_ISL_10220427 | A/red fox/Netherlands/22002636-005/2022 | A / H5N1 | Europe / Netherlands / Provincie Gelderland | Rene Heutink (Wageningen Bioveterinary Research) |
| EPI_ISL_10220428 | A/red fox/Netherlands/22002722-004/2022 | A / H5N1 | Europe / Netherlands / Provincie Gelderland | Rene Heutink (Wageningen Bioveterinary Research) |
| EPI_ISL_10220429 | A/polecat/Netherlands/22002792-005/2022 | A / H5N1 | Europe / Netherlands / Provincie Gelderland | Rene Heutink (Wageningen Bioveterinary Research) |
| EPI_ISL_10220432 | A/badger/Netherlands/22003078-004/2022 | A / H5N1 | Europe / Netherlands / Provincie Gelderland | Rene Heutink (Wageningen Bioveterinary Research) |
| EPI_ISL_10255580 | A/wild bird/Netherlands/22002833-002/2022 | A / H5N1 | Europe / Netherlands / Provincie Gelderland | Rene Heutink (Wageningen Bioveterinary Research) |
| EPI_ISL_10255581 | A/red fox/Netherlands/22002992-003/2022 | A / H5N1 | Europe / Netherlands / Provincie Gelderland | Rene Heutink (Wageningen Bioveterinary Research) |
| EPI_ISL_10255587 | A/red fox/Netherlands/22002993-003 /2022 | A / H5N1 | Europe / Netherlands / Provincie Noord-Holland | Rene Heutink (Wageningen Bioveterinary Research) |
| EPI_ISL_10261376 | A/pigeon/Germany-NW/AI00951/2022 | A / H5N1 | Europe / Germany / North Rhine-Westphalia / Regierungsbezirk Arnsberg / Dortmund | Jacqueline King (Friedrich-Loeffler-Institut) |
| EPI_ISL_1033028 | A/chicken/Czech Republic/2395/2021 | A / H5N8 | Europe / Czech Republic / Stredocesky Kraj / Okres Benesov / Jankov | Alexander Nagy (State Veterinary Institute Prague) |
| EPI_ISL_1033029 | A/chicken/Czech Republic/2502-1/2021 | A / H5N8 | Europe / Czech Republic / Pardubicky kraj / Okres Chrudim / Ronov nad Doubravou; GPS 49°53'2 | Alexander Nagy (State Veterinary Institute Prague) |
| EPI_ISL_1033030 | A/chicken/Czech Republic/2502-2/2021 | A / H5N8 | Europe / Czech Republic / Pardubicky kraj / Okres Chrudim / Ronov nad Doubravou; GPS 49°53'2 | Alexander Nagy (State Veterinary Institute Prague) |
| EPI_ISL_1033124 | A/mute swan/Czech Republic/2600/2021 | A / H5N8 | Europe / Czech Republic / Jihocesky Kraj / Okres Strakonice / Strakonice | Alexander Nagy (State Veterinary Institute Prague) |
| EPI_ISL_10347218 | A/Barnacle Goose/Netherlands/6/2021 | A / H5N1 | Europe / Netherlands / Provincie Friesland / Kollumerland | Pascal Lexmond (Erasmus Medical Center / Viroscience) |
| EPI_ISL_10347219 | A/Barnacle Goose/Netherlands/8/2022 | A / H5N1 | Europe / Netherlands / Provincie Friesland / Gemeente Terschelling / Oosterend | Pascal Lexmond (Erasmus Medical Center / Viroscience) |
| EPI_ISL_10347220 | A/Barnacle Goose/Netherlands/7/2021 | A / H5N1 | Europe / Netherlands / Provincie Friesland / Kollumerland | Pascal Lexmond (Erasmus Medical Center / Viroscience) |
| EPI_ISL_10347326 | A/long-eared owl/Germany-NI/AI09037/2021 | A / H5N1 | Europe / Germany / Lower Saxony / Friesland | Jacqueline King (Friedrich-Loeffler-Institut) |
| EPI_ISL_10364607 | A/mute swan/Czech Republic/4099/2021 | A / H5N5 | Europe / Czech Republic / Zlinsky Kraj / Okres Kromeriz / 49°14'23.619'', 17°28'34.391''E | Alexander Nagy (State Veterinary Institute Prague) |
| EPI_ISL_1038924 | A/Astrakhan/3212/2020 | A / H5N8 | Europe / Russian Federation / Astrakhan Oblast | Ivan Susloparov (State Research Center of Virology and Biotechnology (VECTOR) / Emerging Zoonotic Diseases and Influenza) |
| EPI_ISL_1039231 | A/chicken/Astrakhan/321-01/2020 | A / H5N8 | Europe / Russian Federation / Astrakhan Oblast | Natalia Goncharova (State Research Center of Virology and Biotechnology (VECTOR) / Emerging Zoonotic Diseases and Influenza) |
| EPI_ISL_1039234 | A/chicken/Astrakhan/321-05/2020 | A / H5N8 | Europe / Russian Federation / Astrakhan Oblast | Natalia Goncharova (State Research Center of Virology and Biotechnology (VECTOR) / Emerging Zoonotic Diseases and Influenza) |
| EPI_ISL_1039236 | A/chicken/Astrakhan/321-06/2020 | A / H5N8 | Europe / Russian Federation / Astrakhan Oblast | Natalia Goncharova (State Research Center of Virology and Biotechnology (VECTOR) / Emerging Zoonotic Diseases and Influenza) |
| EPI_ISL_1039238 | A/chicken/Astrakhan/321-09/2020 | A / H5N8 | Europe / Russian Federation / Astrakhan Oblast | Natalia Goncharova (State Research Center of Virology and Biotechnology (VECTOR) / Emerging Zoonotic Diseases and Influenza) |
| EPI_ISL_1039239 | A/chicken/Astrakhan/321-10/2020 | A / H5N8 | Europe / Russian Federation / Astrakhan Oblast | Natalia Goncharova (State Research Center of Virology and Biotechnology (VECTOR) / Emerging Zoonotic Diseases and Influenza) |
| EPI_ISL_10406912 | rg A/chicken/Egypt/Q16684C/2019 | A / H5N8 | Africa / Egypt / Qalyubia | Yassmin Moatasim (National Research Center (Hospital) / Water Pollution) |
| EPI_ISL_1041108 | A/duck/Chiba/D2A-8C/2021 | A / H5N8 | Asia / Japan / Chiba | Takehiko Saito (National Institute of Animal Health) |
| EPI_ISL_1041120 | A/duck/Chiba/D2B-7C/2021 | A / H5N8 | Asia / Japan / Chiba | Takehiko Saito (National Institute of Animal Health) |
| EPI_ISL_1041121 | A/duck/Chiba/D2B-9C/2021 | A / H5N8 | Asia / Japan / Chiba | Takehiko Saito (National Institute of Animal Health) |
| EPI_ISL_1041124 | A/chicken/Chiba/I1C/2021 | A / H5N8 | Asia / Japan / Chiba | Takehiko Saito (National Institute of Animal Health) |
| EPI_ISL_1041125 | A/chicken/Chiba/I1T/2021 | A / H5N8 | Asia / Japan / Chiba | Takehiko Saito (National Institute of Animal Health) |
| EPI_ISL_1041126 | A/chicken/Chiba/I2T/2021 | A / H5N8 | Asia / Japan / Chiba | Takehiko Saito (National Institute of Animal Health) |
| EPI_ISL_1041127 | A/chicken/Chiba/I3T/2021 | A / H5N8 | Asia / Japan / Chiba | Takehiko Saito (National Institute of Animal Health) |
| EPI_ISL_1041128 | A/chicken/Chiba/J1C/2021 | A / H5N8 | Asia / Japan / Chiba | Takehiko Saito (National Institute of Animal Health) |
| EPI_ISL_1041129 | A/chicken/Chiba/J1T/2021 | A / H5N8 | Asia / Japan / Chiba | Takehiko Saito (National Institute of Animal Health) |

| | | | | |
|---|---|---|---|---|
| EPI_ISL_1041130 | A/chicken/Chiba/J6C/2021 | A / H5N8 | Asia / Japan / Chiba | Takehiko Saito (National Institute of Animal Health) |
| EPI_ISL_1041131 | A/chicken/Chiba/J6T/2021 | A / H5N8 | Asia / Japan / Chiba | Takehiko Saito (National Institute of Animal Health) |
| EPI_ISL_1041132 | A/chicken/Tokushima/B1T/2020 | A / H5N8 | Asia / Japan / Tokushima | Takehiko Saito (National Institute of Animal Health) |
| EPI_ISL_1041133 | A/chicken/Tokushima/B2T/2020 | A / H5N8 | Asia / Japan / Tokushima | Takehiko Saito (National Institute of Animal Health) |
| EPI_ISL_1041134 | A/chicken/Tokushima/B3T/2020 | A / H5N8 | Asia / Japan / Tokushima | Takehiko Saito (National Institute of Animal Health) |
| EPI_ISL_1041135 | A/chicken/Tokushima/B4T/2020 | A / H5N8 | Asia / Japan / Tokushima | Takehiko Saito (National Institute of Animal Health) |
| EPI_ISL_1041136 | A/duck/Chiba/D1A-10T/2021 | A / H5N8 | Asia / Japan / Chiba | Takehiko Saito (National Institute of Animal Health) |
| EPI_ISL_1041137 | A/duck/Chiba/D1A-1T/2021 | A / H5N8 | Asia / Japan / Chiba | Takehiko Saito (National Institute of Animal Health) |
| EPI_ISL_1041138 | A/duck/Chiba/D1A-2T/2021 | A / H5N8 | Asia / Japan / Chiba | Takehiko Saito (National Institute of Animal Health) |
| EPI_ISL_1041139 | A/duck/Chiba/D1A-3T/2021 | A / H5N8 | Asia / Japan / Chiba | Takehiko Saito (National Institute of Animal Health) |
| EPI_ISL_1041140 | A/duck/Chiba/D1A-4T/2021 | A / H5N8 | Asia / Japan / Chiba | Takehiko Saito (National Institute of Animal Health) |
| EPI_ISL_1041141 | A/duck/Chiba/D1A-5T/2021 | A / H5N8 | Asia / Japan / Chiba | Takehiko Saito (National Institute of Animal Health) |
| EPI_ISL_1041142 | A/duck/Chiba/D1A-6T/2021 | A / H5N8 | Asia / Japan / Chiba | Takehiko Saito (National Institute of Animal Health) |
| EPI_ISL_1041143 | A/duck/Chiba/D1A-7T/2021 | A / H5N8 | Asia / Japan / Chiba | Takehiko Saito (National Institute of Animal Health) |
| EPI_ISL_1041144 | A/duck/Chiba/D1A-8T/2021 | A / H5N8 | Asia / Japan / Chiba | Takehiko Saito (National Institute of Animal Health) |
| EPI_ISL_1041145 | A/duck/Chiba/D1A-9T/2021 | A / H5N8 | Asia / Japan / Chiba | Takehiko Saito (National Institute of Animal Health) |
| EPI_ISL_1041146 | A/duck/Chiba/D1B-10T/2021 | A / H5N8 | Asia / Japan / Chiba | Takehiko Saito (National Institute of Animal Health) |
| EPI_ISL_1041147 | A/duck/Chiba/D1B-1T/2021 | A / H5N8 | Asia / Japan / Chiba | Takehiko Saito (National Institute of Animal Health) |
| EPI_ISL_1041148 | A/duck/Chiba/D1B-2T/2021 | A / H5N8 | Asia / Japan / Chiba | Takehiko Saito (National Institute of Animal Health) |
| EPI_ISL_1041149 | A/duck/Chiba/D1B-3T/2021 | A / H5N8 | Asia / Japan / Chiba | Takehiko Saito (National Institute of Animal Health) |
| EPI_ISL_1041150 | A/duck/Chiba/D1B-4T/2021 | A / H5N8 | Asia / Japan / Chiba | Takehiko Saito (National Institute of Animal Health) |
| EPI_ISL_1041151 | A/duck/Chiba/D1B-5T/2021 | A / H5N8 | Asia / Japan / Chiba | Takehiko Saito (National Institute of Animal Health) |
| EPI_ISL_1041152 | A/duck/Chiba/D1B-6T/2021 | A / H5N8 | Asia / Japan / Chiba | Takehiko Saito (National Institute of Animal Health) |
| EPI_ISL_1041153 | A/duck/Chiba/D1B-7T/2021 | A / H5N8 | Asia / Japan / Chiba | Takehiko Saito (National Institute of Animal Health) |
| EPI_ISL_1041154 | A/duck/Chiba/D1B-8T/2021 | A / H5N8 | Asia / Japan / Chiba | Takehiko Saito (National Institute of Animal Health) |
| EPI_ISL_1041155 | A/duck/Chiba/D1B-9T/2021 | A / H5N8 | Asia / Japan / Chiba | Takehiko Saito (National Institute of Animal Health) |
| EPI_ISL_1041158 | A/chicken/Chiba/K11C/2021 | A / H5N8 | Asia / Japan / Chiba | Takehiko Saito (National Institute of Animal Health) |
| EPI_ISL_1041159 | A/chicken/Chiba/K11T/2021 | A / H5N8 | Asia / Japan / Chiba | Takehiko Saito (National Institute of Animal Health) |
| EPI_ISL_1041160 | A/chicken/Chiba/K5C/2021 | A / H5N8 | Asia / Japan / Chiba | Takehiko Saito (National Institute of Animal Health) |
| EPI_ISL_1041161 | A/chicken/Chiba/K5T/2021 | A / H5N8 | Asia / Japan / Chiba | Takehiko Saito (National Institute of Animal Health) |
| EPI_ISL_1041165 | A/duck/Chiba/D2A-10T/2021 | A / H5N8 | Asia / Japan / Chiba | Takehiko Saito (National Institute of Animal Health) |
| EPI_ISL_1041166 | A/duck/Chiba/D2A-1T/2021 | A / H5N8 | Asia / Japan / Chiba | Takehiko Saito (National Institute of Animal Health) |
| EPI_ISL_1041167 | A/duck/Chiba/D2A-2T/2021 | A / H5N8 | Asia / Japan / Chiba | Takehiko Saito (National Institute of Animal Health) |
| EPI_ISL_1041168 | A/duck/Chiba/D2A-3T/2021 | A / H5N8 | Asia / Japan / Chiba | Takehiko Saito (National Institute of Animal Health) |
| EPI_ISL_1041169 | A/duck/Chiba/D2A-5T/2021 | A / H5N8 | Asia / Japan / Chiba | Takehiko Saito (National Institute of Animal Health) |
| EPI_ISL_1041170 | A/duck/Chiba/D2A-7T/2021 | A / H5N8 | Asia / Japan / Chiba | Takehiko Saito (National Institute of Animal Health) |
| EPI_ISL_1041171 | A/duck/Chiba/D2A-9T/2021 | A / H5N8 | Asia / Japan / Chiba | Takehiko Saito (National Institute of Animal Health) |
| EPI_ISL_1041172 | A/eastern buzzard/Toyama/160213C/2021 | A / H5N8 | Asia / Japan / Toyama | Takehiko Saito (National Institute of Animal Health) |
| EPI_ISL_1041173 | A/eastern buzzard/Toyama/160213T/2021 | A / H5N8 | Asia / Japan / Toyama | Takehiko Saito (National Institute of Animal Health) |
| EPI_ISL_1041176 | A/owl/Tochigi/090204C/2021 | A / H5N8 | Asia / Japan / Tochigi | Takehiko Saito (National Institute of Animal Health) |
| EPI_ISL_1041177 | A/peregrine falcon/Tochigi/090205C/2021 | A / H5N8 | Asia / Japan / Tochigi | Takehiko Saito (National Institute of Animal Health) |
| EPI_ISL_1041178 | A/peregrine falcon/Tochigi/090205T/2021 | A / H5N8 | Asia / Japan / Tochigi | Takehiko Saito (National Institute of Animal Health) |
| EPI_ISL_1041180 | A/whooper swan/Niigata/150212T/2021 | A / H5N8 | Asia / Japan / Niigata | Takehiko Saito (National Institute of Animal Health) |
| EPI_ISL_10434072 | A/swan/Germany-BW/AI00996/2022 | A / H5N2 | Europe / Germany / Baden-Wuerttemberg / Regierungsbezirk Karlsruhe / Karlsruhe | Jacqueline King (Friedrich-Loeffler-Institut) |
| EPI_ISL_10434073 | A/swan/Germany-BW/AI00997/2022 | A / H5N2 | Europe / Germany / Baden-Wuerttemberg / Regierungsbezirk Karlsruhe / Karlsruhe | Jacqueline King (Friedrich-Loeffler-Institut) |
| EPI_ISL_10454874 | A/barnacle goose/Netherlands/22002507-002/2022 | A / H5N1 | Europe / Netherlands / Provincie Gelderland | Rene Heutink (Wageningen Bioveterinary Research) |
| EPI_ISL_10454886 | A/barnacle goose/Netherlands/22002511-001/2022 | A / H5N1 | Europe / Netherlands / Provincie Utrecht | Rene Heutink (Wageningen Bioveterinary Research) |
| EPI_ISL_10454887 | A/red fox/Netherlands/22003495-003/2022 | A / H5N1 | Europe / Netherlands / Provincie Utrecht | Rene Heutink (Wageningen Bioveterinary Research) |
| EPI_ISL_1046889 | A/chicken/Netherlands/21023815-001005/2021 | A / H5N8 | Europe / Netherlands / North Brabant / Gemeente Sint-Oedenrode | Rene Heutink (Wageningen Bioveterinary Research) |
| EPI_ISL_1048238 | A/common kestrel/Netherlands/20020264-002/2020 | A / H5N8 | Europe / Netherlands / Provincie Noord-Holland / Gemeente Blaricum | Rene Heutink (Wageningen Bioveterinary Research) |
| EPI_ISL_1048239 | A/common buzzard/Netherlands/21021023-002/2021 | A / H5N3 | Europe / Netherlands / Provincie Groningen / Gemeente Delfzijl | Rene Heutink (Wageningen Bioveterinary Research) |
| EPI_ISL_1048240 | A/common buzzard/Netherlands/21021396-002/2021 | A / H5N1 | Europe / Netherlands / Provincie Gelderland / Gemeente Zaltbommel | Rene Heutink (Wageningen Bioveterinary Research) |
| EPI_ISL_1048241 | A/barnacle goose/Netherlands/21021591-001/2021 | A / H5N1 | Europe / Netherlands / Provincie Noord-Holland / Gemeente Naarden | Rene Heutink (Wageningen Bioveterinary Research) |
| EPI_ISL_10497305 | A/red knot/Germany-SH/AI01010/2022 | A / H5N1 | Europe / Germany / Schleswig-Holstein / Nordfriesland | Jacqueline King (Friedrich-Loeffler-Institut) |
| EPI_ISL_10497306 | A/buzzard/Germany-BB/AI01212/2022 | A / H5N1 | Europe / Germany / Brandenburg / Ostprignitz-Ruppin | Jacqueline King (Friedrich-Loeffler-Institut) |
| EPI_ISL_1057106 | A/mute swan/Czech Republic/2669-1/2021 | A / H5N8 | Europe / Czech Republic / Jihocesky Kraj / Okres Tabor / Sobeslav | Alexander Nagy (State Veterinary Institute Prague) |
| EPI_ISL_10576444 | A/Anser_anser/Belgium/1809_0002/2022 | A / H5N1 | Europe / Belgium / Province de Liege / Ougrée | Steven Van Borm (Sciensano, Department of Animal Infectious Diseases / Animal Infectious Diseases) |
| EPI_ISL_1058019 | A/mute swan/Czech Republic/2669-2/2021 | A / H5N8 | Europe / Czech Republic / Jihocesky Kraj / Okres Tabor / Sobeslav | Alexander Nagy (State Veterinary Institute Prague) |
| EPI_ISL_1058020 | A/chicken/Czech Republic/29392021 | A / H5N8 | Europe / Czech Republic / Jihocesky Kraj / Okres Strakonice / Katovice | Alexander Nagy (State Veterinary Institute Prague) |
| EPI_ISL_1058021 | A/chicken/Czech Republic/3099-1/2021 | A / H5N8 | Europe / Czech Republic / Plzensky Kraj / Okres Plzen-Mesto / Bzi | Alexander Nagy (State Veterinary Institute Prague) |
| EPI_ISL_1058022 | A/chicken/Czech Republic/3099-2/2021 | A / H5N8 | Europe / Czech Republic / Plzensky Kraj / Okres Plzen-Mesto / Bzi | Alexander Nagy (State Veterinary Institute Prague) |
| EPI_ISL_1063533 | A/Mandarin duck/Kagoshima/KU-d57/2020 | A / H5N8 | Asia / Japan / Kagoshima | Ahmed Magdy Ahmed Khalil (Faculty of Veterinary Medicine - Zagazig University / Zoonoses) |
| EPI_ISL_1063993 | A/common kestrel/Denmark/16023-01/2021-01-01(H5N3) | A / H5N3 | Europe / Denmark / Region Syddanmark / Kolding Kommune | Charlotte Kristiane Hjulsager (Statens Serum Institute / Microbiological Diagnostic and Virology) |
| EPI_ISL_10724257 | A/sanderling/Netherlands/22002272-001/2021 | A / H5N1 | Europe / Netherlands / Provincie Friesland | Rene Heutink (Wageningen Bioveterinary Research) |
| EPI_ISL_10724259 | A/seagull/Netherlands/22002274-002/2022 | A / H5N1 | Europe / Netherlands / Provincie Gelderland | Rene Heutink (Wageningen Bioveterinary Research) |
| EPI_ISL_1080480 | A/mute swan/Czech Republic/3160-1/2021 | A / H5N8 | Europe / Czech Republic / Liberecky Kraj / Okres Liberec / Prisovice, pond Velky pisecak, GPS: 50 | Alexander Nagy (State Veterinary Institute Prague) |
| EPI_ISL_1080579 | A/common coot/Czech Republic/3160-2/2021 | A / H5N8 | Europe / Czech Republic / Liberecky Kraj / Okres Liberec / Prisovice, pond Velky pisecak, GPS: 50 | Alexander Nagy (State Veterinary Institute Prague) |
| EPI_ISL_1081353 | A/whooper swan/Tochigi/090203C/2021 | A / H5N8 | Asia / Japan / Tochigi | Takehiko Saito (National Institute of Animal Health) |
| EPI_ISL_1081369 | A/Chongqing/00013/2021 | A / H5N6 | Asia / China / Chongqinq Municipality | Lei Yang (WHO Chinese National Influenza Center / Virology Institute, Chinese CDC) |
| EPI_ISL_1081370 | A/Anhui/2021-00011/2020 | A / H5N6 | Asia / China / Anhui Province | Lei Yang (WHO Chinese National Influenza Center / Virology Institute, Chinese CDC) |
| EPI_ISL_10915976 | A/red fox/Netherlands/22004172-003/2022 | A / H5N1 | Europe / Netherlands / Provincie Gelderland | Rene Heutink (Wageningen Bioveterinary Research) |
| EPI_ISL_1093422 | A/Turkey/Sweden/SVA210214SZ0002/KN035667-IP5/2021 | A / H5N8 | Europe / Sweden / Skane Lan / Simrishamns Kommun | Siamak Zohari (National Veterinary Institute) |

| | | | | |
|---|---|---|---|---|
| EPI_ISL_1093423 | A/Chicken/Sweden/SVA210217SZ0001/KN001190-IP6/2021 | A / H5N8 | Europe / Sweden / Vastra Gotalands Lan / Ale Kommun | Siamak Zohari (National Veterinary Institute) |
| EPI_ISL_1093424 | A/Common goldeneye/SVA210202SZ0464/KN000277/2021 | A / H5N5 | Europe / Sweden / Kalmar Lan / Vasterviks Kommun | Siamak Zohari (National Veterinary Institute) |
| EPI_ISL_1093425 | A/Taiga bean goose/SVA210217SZ0303/KN000463/2021 | A / H5N8 | Europe / Sweden / Skane Lan / Ystads Kommun | Siamak Zohari (National Veterinary Institute) |
| EPI_ISL_1093426 | A/black-headed gull/Sweden/SVA210216SZ0398/KN000437/2( | A / H5N8 | Europe / Sweden / Skane Lan / Malmo Kommun | Siamak Zohari (National Veterinary Institute) |
| EPI_ISL_10943657 | A/northern goshawk/Sweden/SVA220210SZ0305/FP000317/O | A / H5N1 | Europe / Sweden / Vastra Gotalands Lan / Goteborgs Kommun | Siamak Zohari (National Veterinary Institute) |
| EPI_ISL_1095363 | A/Greylag goose/Sweden/SVA210209SZ0422/KN000346/2021 | A / H5N8 | Europe / Sweden / Blekinge Lan / Solvesborgs Kommun | Siamak Zohari (National Veterinary Institute) |
| EPI_ISL_1095365 | A/Canada goose/Sweden/SVA210209SZ0420/KN000345/2021 | A / H5N8 | Europe / Sweden / Blekinge Lan / Solvesborgs Kommun | Siamak Zohari (National Veterinary Institute) |
| EPI_ISL_1095450 | A/barnacle goose/Sweden/SVA210202SZ0453/KN000276/202 | A / H5N8 | Europe / Sweden / Skane Lan / Lunds Kommun | Siamak Zohari (National Veterinary Institute) |
| EPI_ISL_1095451 | A/barnacle goose/Sweden/SVA210126SZ0462/KN000202/202 | A / H5N8 | Europe / Sweden / Skane Lan / Malmo Kommun | Siamak Zohari (National Veterinary Institute) |
| EPI_ISL_1095453 | A/Turkey/Sweden/SVA201118SZ0002/KN305633-IP1/2020 | A / H5N8 | Europe / Sweden / Skane Lan / Tomelilla Kommun | Siamak Zohari (National Veterinary Institute) |
| EPI_ISL_1095454 | A/Common buzzard/Sweden/SVA210212SZ0284/KN000390/2( | A / H5N8 | Europe / Sweden / Skane Lan / Svedala Kommun | Siamak Zohari (National Veterinary Institute) |
| EPI_ISL_1098830 | A/crane/Kagoshima/KU-93/2021 | A / H5N8 | Asia / Japan / Kagoshima | Ahmed Magdy Ahmed Khalil (Faculty of Veterinary Medicine - Zagazig University / Zoonoses) |
| EPI_ISL_10992813 | A/Mallard/Netherlands/1/2022 | A / H5N1 | Europe / Netherlands / Oud Alblas | Mark Pronk (Erasmus Medical Center / Department of Virology) |
| EPI_ISL_10993074 | A/Mallard/Netherlands/2/2022 | A / H5N1 | Europe / Netherlands / Oud Alblas | Mark Pronk (Erasmus Medical Center / Department of Virology) |
| EPI_ISL_10993075 | A/Mallard/Netherlands/4/2021 | A / H5N1 | Europe / Netherlands / Zwaagdijk, Eendenkooi Zwaagdijk | Mark Pronk (Erasmus Medical Center / Department of Virology) |
| EPI_ISL_10993076 | A/Mallard/Netherlands/5/2021 | A / H5N1 | Europe / Netherlands / Zwaagdijk, Eendenkooi Zwaagdijk | Mark Pronk (Erasmus Medical Center / Department of Virology) |
| EPI_ISL_10993105 | A/Mallard/Netherlands/6/2021 | A / H5N1 | Europe / Netherlands / Zwaagdijk, Eendenkooi Zwaagdijk | Mark Pronk (Erasmus Medical Center / Department of Virology) |
| EPI_ISL_10993134 | A/Mallard/Netherlands/7/2021 | A / H5N1 | Europe / Netherlands / Zwaagdijk, Eendenkooi Zwaagdijk | Mark Pronk (Erasmus Medical Center / Department of Virology) |
| EPI_ISL_10993166 | A/Mallard/Netherlands/8/2021 | A / H5N1 | Europe / Netherlands / Zwaagdijk, Eendenkooi Zwaagdijk | Mark Pronk (Erasmus Medical Center / Department of Virology) |
| EPI_ISL_10993196 | A/Sanderling/Netherlands/1/2022 | A / H5N1 | Europe / Netherlands / Lauwersmeer | Mark Pronk (Erasmus Medical Center / Department of Virology) |
| EPI_ISL_11007527 | A/laying_hen/Moldova/68-1_22VIR638-1/2022 | A / H5N1 | Europe / Moldova, Republic of | Giacomo Barbierato (Istituto Zooprofilattico Sperimentale delle Venezie / Ricerca e innovazione) |
| EPI_ISL_11007528 | A/rooster/Slovenia/2039_22VIR777-1/2021 | A / H5N1 | Europe / Slovenia | Giacomo Barbierato (Istituto Zooprofilattico Sperimentale delle Venezie / Ricerca e innovazione) |
| EPI_ISL_11007529 | A/swan/Slovenia/2041_22VIR777-2/2021 | A / H5N1 | Europe / Slovenia | Giacomo Barbierato (Istituto Zooprofilattico Sperimentale delle Venezie / Ricerca e innovazione) |
| EPI_ISL_11007530 | A/swan/Slovenia/2049_22VIR777-3/2021 | A / H5N1 | Europe / Slovenia | Giacomo Barbierato (Istituto Zooprofilattico Sperimentale delle Venezie / Ricerca e innovazione) |
| EPI_ISL_11007531 | A/swan/Slovenia/2060_22VIR777-4/2021 | A / H5N1 | Europe / Slovenia | Giacomo Barbierato (Istituto Zooprofilattico Sperimentale delle Venezie / Ricerca e innovazione) |
| EPI_ISL_11007532 | A/swan/Slovenia/2072_22VIR777-5/2021 | A / H5N1 | Europe / Slovenia | Giacomo Barbierato (Istituto Zooprofilattico Sperimentale delle Venezie / Ricerca e innovazione) |
| EPI_ISL_11007533 | A/swan/Slovenia/2073_22VIR777-6/2021 | A / H5N1 | Europe / Slovenia | Giacomo Barbierato (Istituto Zooprofilattico Sperimentale delle Venezie / Ricerca e innovazione) |
| EPI_ISL_11007534 | A/seagull/Slovenia/2075_22VIR777-7/2021 | A / H5N1 | Europe / Slovenia | Giacomo Barbierato (Istituto Zooprofilattico Sperimentale delle Venezie / Ricerca e innovazione) |
| EPI_ISL_11007535 | A/swan/Slovenia/13_22VIR777-8/2022 | A / H5N1 | Europe / Slovenia | Giacomo Barbierato (Istituto Zooprofilattico Sperimentale delle Venezie / Ricerca e innovazione) |
| EPI_ISL_11007537 | A/hen/Bulgaria/854-1_22VIR778-10/2021 | A / H5N1 | Europe / Bulgaria | Giacomo Barbierato (Istituto Zooprofilattico Sperimentale delle Venezie / Ricerca e innovazione) |
| EPI_ISL_11007538 | A/hen/Bulgaria/722-1_22VIR778-1/2021 | A / H5N1 | Europe / Bulgaria | Giacomo Barbierato (Istituto Zooprofilattico Sperimentale delle Venezie / Ricerca e innovazione) |
| EPI_ISL_11007539 | A/turkey/Bulgaria/755-1_22VIR778-4/2021 | A / H5N1 | Europe / Bulgaria | Giacomo Barbierato (Istituto Zooprofilattico Sperimentale delle Venezie / Ricerca e innovazione) |
| EPI_ISL_11007540 | A/duck/Bulgaria/756-4_22VIR778-6/2021 | A / H5N1 | Europe / Bulgaria | Giacomo Barbierato (Istituto Zooprofilattico Sperimentale delle Venezie / Ricerca e innovazione) |
| EPI_ISL_11007541 | A/hen/Bulgaria/757-6_22VIR778-7/2021 | A / H5N1 | Europe / Bulgaria | Giacomo Barbierato (Istituto Zooprofilattico Sperimentale delle Venezie / Ricerca e innovazione) |
| EPI_ISL_11007542 | A/chicken/Niger/22VIR1409-13/2022 | A / H5N1 | Africa / Niger | Giacomo Barbierato (Istituto Zooprofilattico Sperimentale delle Venezie / Ricerca e innovazione) |
| EPI_ISL_11007543 | A/chicken/Niger/22VIR1409-23/2022 | A / H5N1 | Africa / Niger | Giacomo Barbierato (Istituto Zooprofilattico Sperimentale delle Venezie / Ricerca e innovazione) |
| EPI_ISL_11007544 | A/quail/Niger/22VIR1409-28/2022 | A / H5N1 | Africa / Niger | Giacomo Barbierato (Istituto Zooprofilattico Sperimentale delle Venezie / Ricerca e innovazione) |
| EPI_ISL_11007545 | A/quail/Niger/22VIR1409-30/2022 | A / H5N1 | Africa / Niger | Giacomo Barbierato (Istituto Zooprofilattico Sperimentale delle Venezie / Ricerca e innovazione) |
| EPI_ISL_11007546 | A/chicken/Niger/22VIR1409-5/2022 | A / H5N1 | Africa / Niger | Giacomo Barbierato (Istituto Zooprofilattico Sperimentale delle Venezie / Ricerca e innovazione) |
| EPI_ISL_11007547 | A/chicken/Niger/22VIR1409-9/2022 | A / H5N1 | Africa / Niger | Giacomo Barbierato (Istituto Zooprofilattico Sperimentale delle Venezie / Ricerca e innovazione) |
| EPI_ISL_11007548 | A/turkey/Italy/21VIR9520/2021 | A / H5N1 | Europe / Italy | Giacomo Barbierato (Istituto Zooprofilattico Sperimentale delle Venezie / Ricerca e innovazione) |
| EPI_ISL_11007721 | A/laying_hen/Moldova/68-2_22VIR638-2/2022 | A / H5N1 | Europe / Moldova, Republic of | Giacomo Barbierato (Istituto Zooprofilattico Sperimentale delle Venezie / Ricerca e innovazione) |
| EPI_ISL_11007722 | A/partridge/Bulgaria/745_22VIR778-3/2021 | A / H5N1 | Europe / Bulgaria | Giacomo Barbierato (Istituto Zooprofilattico Sperimentale delle Venezie / Ricerca e innovazione) |
| EPI_ISL_11019784 | A/red fox/Netherlands/22000097-005/2022 | A / H5N1 | Europe / Netherlands / Provincie Noord-Holland | Rene Heutink (Wageningen Bioveterinary Research) |
| EPI_ISL_11019882 | A/red fox/Netherlands/22002636-002/2022 | A / H5N1 | Europe / Netherlands / Provincie Gelderland | Rene Heutink (Wageningen Bioveterinary Research) |
| EPI_ISL_11112340 | A/Barnacle Goose/Netherlands/10/2022 | A / H5N1 | Europe / Netherlands / Schellinkhout, Zuiderdijk | Mark Pronk (Erasmus Medical Center / Department of Virology) |
| EPI_ISL_11112342 | A/Barnacle Goose/Netherlands/11/2022 | A / H5N1 | Europe / Netherlands / Hippolytushoef, Normerven | Mark Pronk (Erasmus Medical Center / Department of Virology) |
| EPI_ISL_11112344 | A/Barnacle Goose/Netherlands/12/2022 | A / H5N1 | Europe / Netherlands / Stroe, Waddenkust | Mark Pronk (Erasmus Medical Center / Department of Virology) |
| EPI_ISL_11112346 | A/Barnacle Goose/Netherlands/13/2022 | A / H5N1 | Europe / Netherlands / IJsselmeer, Enkhuizen, De Kreupel | Mark Pronk (Erasmus Medical Center / Department of Virology) |
| EPI_ISL_11112348 | A/Barnacle Goose/Netherlands/14/2022 | A / H5N1 | Europe / Netherlands / IJsselmeer, Enkhuizen, De Kreupel | Mark Pronk (Erasmus Medical Center / Department of Virology) |
| EPI_ISL_11112350 | A/Barnacle Goose/Netherlands/15/2022 | A / H5N1 | Europe / Netherlands / Noord Buren, Waddenkust | Mark Pronk (Erasmus Medical Center / Department of Virology) |
| EPI_ISL_11112352 | A/Barnacle Goose/Netherlands/9/2022 | A / H5N1 | Europe / Netherlands / Schellinkhout, Zuiderdijk | Mark Pronk (Erasmus Medical Center / Department of Virology) |
| EPI_ISL_11112354 | A/Common Gull/Netherlands/1/2022 | A / H5N1 | Europe / Netherlands / Hippolytushoef, Waddenzeekust | Mark Pronk (Erasmus Medical Center / Department of Virology) |
| EPI_ISL_11112356 | A/Eurasian Curlew/Netherlands/1/2022 | A / H5N1 | Europe / Netherlands / Hippolytushoef, Normerven | Mark Pronk (Erasmus Medical Center / Department of Virology) |
| EPI_ISL_11112358 | A/European Herring Gull/Netherlands/2/2022 | A / H5N1 | Europe / Netherlands / Den Oever, Voorhaven | Mark Pronk (Erasmus Medical Center / Department of Virology) |
| EPI_ISL_11112360 | A/Great black-backed Gull/Netherlands/2/2022 | A / H5N1 | Europe / Netherlands / IJsselmeer, Enkhuizen, De Kreupel | Mark Pronk (Erasmus Medical Center / Department of Virology) |
| EPI_ISL_11112362 | A/Greylag Goose/Netherlands/1/2022 | A / H5N1 | Europe / Netherlands / Tersluis, Zuiderdijk | Mark Pronk (Erasmus Medical Center / Department of Virology) |
| EPI_ISL_11112484 | A/Caspian Gull/Netherlands/2/2022 | A / H5N1 | Europe / Netherlands / IJsselmeer, Enkhuizen, De Kreupel | Mark Pronk (Erasmus Medical Center / Department of Virology) |
| EPI_ISL_1114725 | A/mallard/Kagoshima/KU-d89/2021 | A / H5N8 | Asia / Japan / Kagoshima | Ahmed Magdy Ahmed Khalil (Faculty of Veterinary Medicine - Zagazig University / Zoonoses) |
| EPI_ISL_1114735 | A/chicken/Tyumen/302-01/2020 | A / H5N8 | Europe / Russian Federation / Tyumen Oblast / Tyumen | Natalia Goncharova (State Research Center of Virology and Biotechnology (VECTOR) / Emerging Zoonotic Diseases and Influenza) |
| EPI_ISL_1114736 | A/chicken/Tyumen/302-02/2020 | A / H5N8 | Europe / Russian Federation / Tyumen Oblast / Tyumen | Natalia Goncharova (State Research Center of Virology and Biotechnology (VECTOR) / Emerging Zoonotic Diseases and Influenza) |
| EPI_ISL_1114737 | A/chicken/Kostroma/304-01/2020 | A / H5N8 | Europe / Russian Federation / Kostroma Oblast / Kostroma | Natalia Goncharova (State Research Center of Virology and Biotechnology (VECTOR) / Emerging Zoonotic Diseases and Influenza) |
| EPI_ISL_1114738 | A/chicken/Kostroma/304-03/2020 | A / H5N8 | Europe / Russian Federation / Kostroma Oblast / Kostroma | Natalia Goncharova (State Research Center of Virology and Biotechnology (VECTOR) / Emerging Zoonotic Diseases and Influenza) |
| EPI_ISL_1114740 | A/chicken/Kostroma/304-04/2020 | A / H5N8 | Europe / Russian Federation / Kostroma Oblast / Kostroma | Natalia Goncharova (State Research Center of Virology and Biotechnology (VECTOR) / Emerging Zoonotic Diseases and Influenza) |
| EPI_ISL_1114741 | A/chicken/Kostroma/304-06/2020 | A / H5N8 | Europe / Russian Federation / Kostroma Oblast / Kostroma | Natalia Goncharova (State Research Center of Virology and Biotechnology (VECTOR) / Emerging Zoonotic Diseases and Influenza) |
| EPI_ISL_1114742 | A/chicken/Kostroma/304-08/2020 | A / H5N8 | Europe / Russian Federation / Kostroma Oblast / Kostroma | Natalia Goncharova (State Research Center of Virology and Biotechnology (VECTOR) / Emerging Zoonotic Diseases and Influenza) |
| EPI_ISL_1114745 | A/chicken/Kostroma/304-10/2020 | A / H5N8 | Europe / Russian Federation / Kostroma Oblast / Kostroma | Natalia Goncharova (State Research Center of Virology and Biotechnology (VECTOR) / Emerging Zoonotic Diseases and Influenza) |
| EPI_ISL_1114746 | A/chicken/Rostov-on-Don/308-02/2020 | A / H5N8 | Europe / Russian Federation / Rostov Oblast / Rostov-on-Don | Natalia Goncharova (State Research Center of Virology and Biotechnology (VECTOR) / Emerging Zoonotic Diseases and Influenza) |
| EPI_ISL_1114747 | A/chicken/Rostov-on-Don/308-03/2020 | A / H5N8 | Europe / Russian Federation / Rostov Oblast / Rostov-on-Don | Natalia Goncharova (State Research Center of Virology and Biotechnology (VECTOR) / Emerging Zoonotic Diseases and Influenza) |
| EPI_ISL_1114748 | A/chicken/Rostov-on-Don/308-04/2020 | A / H5N8 | Europe / Russian Federation / Rostov Oblast / Rostov-on-Don | Natalia Goncharova (State Research Center of Virology and Biotechnology (VECTOR) / Emerging Zoonotic Diseases and Influenza) |
| EPI_ISL_1114749 | A/turkey/Stavropol/320-01/2020 | A / H5N8 | Europe / Russian Federation / Stavropol Krai / Stavropol | Natalia Goncharova (State Research Center of Virology and Biotechnology (VECTOR) / Emerging Zoonotic Diseases and Influenza) |
| EPI_ISL_1114750 | A/turkey/Stavropol/320-02/2020 | A / H5N8 | Europe / Russian Federation / Stavropol Krai / Stavropol | Natalia Goncharova (State Research Center of Virology and Biotechnology (VECTOR) / Emerging Zoonotic Diseases and Influenza) |

| | | | | |
|---|---|---|---|---|
| EPI_ISL_1114751 | A/turkey/Stavropol/320-03/2020 | A / H5N8 | Europe / Russian Federation / Stavropol Krai / Stavropol | Natalia Goncharova (State Research Center of Virology and Biotechnology (VECTOR) / Emerging Zoonotic Diseases and Influenza) |
| EPI_ISL_1114752 | A/mute swan/North Ossetia-Alania/325-01/2020 | A / H5N8 | Europe / Russian Federation / Republic of North Ossetia?Alania | Natalia Goncharova (State Research Center of Virology and Biotechnology (VECTOR) / Emerging Zoonotic Diseases and Influenza) |
| EPI_ISL_1114753 | A/mute swan/North Ossetia-Alania/325-02/2020 | A / H5N8 | Europe / Russian Federation / Republic of North Ossetia?Alania | Natalia Goncharova (State Research Center of Virology and Biotechnology (VECTOR) / Emerging Zoonotic Diseases and Influenza) |
| EPI_ISL_1114754 | A/mute swan/North Ossetia-Alania/325-03/2020 | A / H5N8 | Europe / Russian Federation / Republic of North Ossetia?Alania | Natalia Goncharova (State Research Center of Virology and Biotechnology (VECTOR) / Emerging Zoonotic Diseases and Influenza) |
| EPI_ISL_1114755 | A/turkey/Rostov-on-Don/332-08/2021 | A / H5N8 | Europe / Russian Federation / Rostov Oblast / Rostov-on-Don | Natalia Goncharova (State Research Center of Virology and Biotechnology (VECTOR) / Emerging Zoonotic Diseases and Influenza) |
| EPI_ISL_1114757 | A/turkey/Rostov-on-Don/332-09/2021 | A / H5N8 | Europe / Russian Federation / Rostov Oblast / Rostov-on-Don | Natalia Goncharova (State Research Center of Virology and Biotechnology (VECTOR) / Emerging Zoonotic Diseases and Influenza) |
| EPI_ISL_1114758 | A/turkey/Rostov-on-Don/332-10/2021 | A / H5N8 | Europe / Russian Federation / Rostov Oblast / Rostov-on-Don | Natalia Goncharova (State Research Center of Virology and Biotechnology (VECTOR) / Emerging Zoonotic Diseases and Influenza) |
| EPI_ISL_1114759 | A/turkey/Rostov-on-Don/332-12/2021 | A / H5N8 | Europe / Russian Federation / Rostov Oblast / Rostov-on-Don | Natalia Goncharova (State Research Center of Virology and Biotechnology (VECTOR) / Emerging Zoonotic Diseases and Influenza) |
| EPI_ISL_1114760 | A/chicken/Krasnodar/334-01/2021 | A / H5N8 | Europe / Russian Federation / Krasnodar Krai / Krasnodar | Natalia Goncharova (State Research Center of Virology and Biotechnology (VECTOR) / Emerging Zoonotic Diseases and Influenza) |
| EPI_ISL_1114761 | A/chicken/Krasnodar/334-02/2021 | A / H5N8 | Europe / Russian Federation / Krasnodar Krai / Krasnodar | Natalia Goncharova (State Research Center of Virology and Biotechnology (VECTOR) / Emerging Zoonotic Diseases and Influenza) |
| EPI_ISL_1114763 | A/chicken/Krasnodar/334-03/2021 | A / H5N8 | Europe / Russian Federation / Krasnodar Krai / Krasnodar | Natalia Goncharova (State Research Center of Virology and Biotechnology (VECTOR) / Emerging Zoonotic Diseases and Influenza) |
| EPI_ISL_11208193 | A/duck/Guangxi/S30428/2021(H5N6) | A / H5N6 | Asia / China / Guangxi | Pengfei Cui (Harbin Veterinary Research Institute (CAAS) / Ministry of Agriculture) |
| EPI_ISL_11208194 | A/duck/Guangxi/S31116/2021(H5N6) | A / H5N6 | Asia / China / Guangxi | Pengfei Cui (Harbin Veterinary Research Institute (CAAS) / Ministry of Agriculture) |
| EPI_ISL_11208195 | A/duck/Guizhou/S4702/2021(H5N6) | A / H5N6 | Asia / China / Guizhou | Pengfei Cui (Harbin Veterinary Research Institute (CAAS) / Ministry of Agriculture) |
| EPI_ISL_11208196 | A/duck/Hunan/S40199/2021(H5N6) | A / H5N6 | Asia / China / Hunan | Pengfei Cui (Harbin Veterinary Research Institute (CAAS) / Ministry of Agriculture) |
| EPI_ISL_11208197 | A/duck/Hunan/S40268/2021(H5N6) | A / H5N6 | Asia / China / Hunan | Pengfei Cui (Harbin Veterinary Research Institute (CAAS) / Ministry of Agriculture) |
| EPI_ISL_11208198 | A/duck/Yunnan/S4318/2021(H5N6) | A / H5N6 | Asia / China / Yunnan | Pengfei Cui (Harbin Veterinary Research Institute (CAAS) / Ministry of Agriculture) |
| EPI_ISL_11208199 | A/duck/Zhejiang/S4854/2021(H5N6) | A / H5N6 | Asia / China / Zhejiang | Pengfei Cui (Harbin Veterinary Research Institute (CAAS) / Ministry of Agriculture) |
| EPI_ISL_11208200 | A/goose/Guangdong/S4751/2021(H5N6) | A / H5N6 | Asia / China / Guangdong | Pengfei Cui (Harbin Veterinary Research Institute (CAAS) / Ministry of Agriculture) |
| EPI_ISL_1121143 | A/chicken/Ukraine/1/2016 | A / H5N8 | Europe / Ukraine / Kherson Oblast / Kalanchatskiy District, Novooleksandrivka | Anna Andriivna Kovalenko (Institute of Veterinary Medicine / Research Training Center for Animal Disease Diagnostics) |
| EPI_ISL_1121144 | A/chicken/Ukraine/3/2016 | A / H5N8 | Europe / Ukraine / Kherson Oblast / Kalanchatskiy District, Novooleksandrivka | Anna Andriivna Kovalenko (Institute of Veterinary Medicine / Research Training Center for Animal Disease Diagnostics) |
| EPI_ISL_1121145 | A/duck/Ukraine/4/2016 | A / H5N8 | Europe / Ukraine / Kherson Oblast / Kalanchatskiy District, Novooleksandrivka | Anna Andriivna Kovalenko (Institute of Veterinary Medicine / Research Training Center for Animal Disease Diagnostics) |
| EPI_ISL_11211731 | A/duck/Bangladesh/19D1818/2021 | A / H5N1 | Asia / Bangladesh | Mohammad Enayet Hossain (International Centre for Diarrhoeal Disease Research (ICDDR,B) / Virology) |
| EPI_ISL_1123263 | A/chicken/Scotland/043405/2020 | A / H5N8 | Europe / United Kingdom / Orkney | Alex Byrne (Animal and Plant Health Agency (APHA) / Virology Department) |
| EPI_ISL_1123350 | A/chicken/England/043683/2020 | A / H5N8 | Europe / United Kingdom / Dorset / England | Alex Byrne (Animal and Plant Health Agency (APHA) / Virology Department) |
| EPI_ISL_1123351 | A/duck/England/043628/2020 | A / H5N8 | Europe / United Kingdom / Norfolk | Alex Byrne (Animal and Plant Health Agency (APHA) / Virology Department) |
| EPI_ISL_1123352 | A/chicken/England/045984/2020 | A / H5N8 | Europe / United Kingdom / Norfolk | Alex Byrne (Animal and Plant Health Agency (APHA) / Virology Department) |
| EPI_ISL_1123353 | A/duck/England/046311/2020 | A / H5N8 | Europe / United Kingdom / Norfolk | Alex Byrne (Animal and Plant Health Agency (APHA) / Virology Department) |
| EPI_ISL_1123354 | A/chicken/England/046491/2020 | A / H5N8 | Europe / United Kingdom / Devon | Alex Byrne (Animal and Plant Health Agency (APHA) / Virology Department) |
| EPI_ISL_1123355 | A/pheasant/Wales/000252/2021 | A / H5N8 | Europe / United Kingdom / Isle of Anglesey | Alex Byrne (Animal and Plant Health Agency (APHA) / Virology Department) |
| EPI_ISL_1123357 | A/brent_goose/England/233339/2020 | A / H5N8 | Europe / United Kingdom / Lincolnshire | Alex Byrne (Animal and Plant Health Agency (APHA) / Virology Department) |
| EPI_ISL_1123358 | A/brent_goose/England/095684/2020 | A / H5N5 | Europe / United Kingdom / Northumberland | Alex Byrne (Animal and Plant Health Agency (APHA) / Virology Department) |
| EPI_ISL_1123359 | A/mute swan/England/263814/2020 | A / H5N8 | Europe / United Kingdom / Dorset | Alex Byrne (Animal and Plant Health Agency (APHA) / Virology Department) |
| EPI_ISL_1123360 | A/mute_swan/England/234135/2020 | A / H5N8 | Europe / United Kingdom / Norfolk | Alex Byrne (Animal and Plant Health Agency (APHA) / Virology Department) |
| EPI_ISL_1123361 | A/pheasant/Scotland/000348/2021 | A / H5N1 | Europe / United Kingdom / Fife | Alex Byrne (Animal and Plant Health Agency (APHA) / Virology Department) |
| EPI_ISL_11259261 | A/herring_gull/Ireland/033533_22VIR1325-10/2021 | A / H5N1 | Europe / Ireland | Giacomo Barbierato (Istituto Zooprofilattico Sperimentale delle Venezie / Ricerca e innovazione) |
| EPI_ISL_11259262 | A/crow/Ireland/035624_22VIR1325-11/2021 | A / H5N1 | Europe / Ireland | Giacomo Barbierato (Istituto Zooprofilattico Sperimentale delle Venezie / Ricerca e innovazione) |
| EPI_ISL_11259263 | A/buzzard/Ireland/000656_22VIR1325-12/2022 | A / H5N1 | Europe / Ireland | Giacomo Barbierato (Istituto Zooprofilattico Sperimentale delle Venezie / Ricerca e innovazione) |
| EPI_ISL_11259264 | A/mute_swan/Ireland/033169_22VIR1325-14/2021 | A / H5N1 | Europe / Ireland | Giacomo Barbierato (Istituto Zooprofilattico Sperimentale delle Venezie / Ricerca e innovazione) |
| EPI_ISL_11259265 | A/peregrine_falcon/Ireland/000191_22VIR1325-15/2022 | A / H5N1 | Europe / Ireland | Giacomo Barbierato (Istituto Zooprofilattico Sperimentale delle Venezie / Ricerca e innovazione) |
| EPI_ISL_11259266 | A/mute_swan/Ireland/033945_22VIR1325-16/2021 | A / H5N1 | Europe / Ireland | Giacomo Barbierato (Istituto Zooprofilattico Sperimentale delle Venezie / Ricerca e innovazione) |
| EPI_ISL_11259267 | A/turkey/Ireland/035425_22VIR1325-17/2021 | A / H5N1 | Europe / Ireland | Giacomo Barbierato (Istituto Zooprofilattico Sperimentale delle Venezie / Ricerca e innovazione) |
| EPI_ISL_11259268 | A/duck/Ireland/036646_22VIR1325-18/2021 | A / H5N1 | Europe / Ireland | Giacomo Barbierato (Istituto Zooprofilattico Sperimentale delle Venezie / Ricerca e innovazione) |
| EPI_ISL_11259269 | A/turkey/Ireland/033674_22VIR1325-19/2021 | A / H5N1 | Europe / Ireland | Giacomo Barbierato (Istituto Zooprofilattico Sperimentale delle Venezie / Ricerca e innovazione) |
| EPI_ISL_11259270 | A/mute_swan/Ireland/032363_22VIR1325-1/2021 | A / H5N1 | Europe / Ireland | Giacomo Barbierato (Istituto Zooprofilattico Sperimentale delle Venezie / Ricerca e innovazione) |
| EPI_ISL_11259271 | A/broiler/Ireland/033734_22VIR1325-20/2021 | A / H5N1 | Europe / Ireland | Giacomo Barbierato (Istituto Zooprofilattico Sperimentale delle Venezie / Ricerca e innovazione) |
| EPI_ISL_11259272 | A/layer/Ireland/034424_22VIR1325-21/2021 | A / H5N1 | Europe / Ireland | Giacomo Barbierato (Istituto Zooprofilattico Sperimentale delle Venezie / Ricerca e innovazione) |
| EPI_ISL_11259273 | A/duck/Ireland/036105_22VIR1325-22/2021 | A / H5N1 | Europe / Ireland | Giacomo Barbierato (Istituto Zooprofilattico Sperimentale delle Venezie / Ricerca e innovazione) |
| EPI_ISL_11259274 | A/white-tailed_eagle/Ireland/032034_22VIR1325-23/2021 | A / H5N1 | Europe / Ireland | Giacomo Barbierato (Istituto Zooprofilattico Sperimentale delle Venezie / Ricerca e innovazione) |
| EPI_ISL_11259275 | A/whooper_swan/Ireland/032444_22VIR1325-2/2021 | A / H5N1 | Europe / Ireland | Giacomo Barbierato (Istituto Zooprofilattico Sperimentale delle Venezie / Ricerca e innovazione) |
| EPI_ISL_11259276 | A/peregrine_falcon/Ireland/032476_22VIR1325-3/2021 | A / H5N1 | Europe / Ireland | Giacomo Barbierato (Istituto Zooprofilattico Sperimentale delle Venezie / Ricerca e innovazione) |
| EPI_ISL_11259277 | A/whooper_swan/Ireland/032960_22VIR1325-4/2021 | A / H5N1 | Europe / Ireland | Giacomo Barbierato (Istituto Zooprofilattico Sperimentale delle Venezie / Ricerca e innovazione) |
| EPI_ISL_11259278 | A/magpie/Ireland/032958_22VIR1325-5/2021 | A / H5N1 | Europe / Ireland | Giacomo Barbierato (Istituto Zooprofilattico Sperimentale delle Venezie / Ricerca e innovazione) |
| EPI_ISL_11259279 | A/greylag_goose/Ireland/032969_22VIR1325-6/2021 | A / H5N1 | Europe / Ireland | Giacomo Barbierato (Istituto Zooprofilattico Sperimentale delle Venezie / Ricerca e innovazione) |
| EPI_ISL_11259280 | A/greylag_goose/Ireland/033062_22VIR1325-7/2021 | A / H5N1 | Europe / Ireland | Giacomo Barbierato (Istituto Zooprofilattico Sperimentale delle Venezie / Ricerca e innovazione) |
| EPI_ISL_11259281 | A/white-fronted_goose/Ireland/033181_22VIR1325-8/2021 | A / H5N1 | Europe / Ireland | Giacomo Barbierato (Istituto Zooprofilattico Sperimentale delle Venezie / Ricerca e innovazione) |
| EPI_ISL_11259282 | A/brent_goose/Ireland/033257_22VIR1325-9/2021 | A / H5N1 | Europe / Ireland | Giacomo Barbierato (Istituto Zooprofilattico Sperimentale delle Venezie / Ricerca e innovazione) |
| EPI_ISL_11259283 | A/fox/Ireland/3866_22VIR2064-1/2022 | A / H5N1 | Europe / Ireland | Giacomo Barbierato (Istituto Zooprofilattico Sperimentale delle Venezie / Ricerca e innovazione) |
| EPI_ISL_11259284 | A/goose/Ireland/3869_22VIR2064-2/2022 | A / H5N1 | Europe / Ireland | Giacomo Barbierato (Istituto Zooprofilattico Sperimentale delle Venezie / Ricerca e innovazione) |
| EPI_ISL_11259285 | A/stork/Spain/442-8_22VIR2142-12/2022 | A / H5N1 | Europe / Spain | Giacomo Barbierato (Istituto Zooprofilattico Sperimentale delle Venezie / Ricerca e innovazione) |
| EPI_ISL_11259286 | A/goose/Spain/512-2_22VIR2142-13/2022 | A / H5N1 | Europe / Spain | Giacomo Barbierato (Istituto Zooprofilattico Sperimentale delle Venezie / Ricerca e innovazione) |
| EPI_ISL_11259287 | A/stork/Spain/538-2_22VIR2142-14/2022 | A / H5N1 | Europe / Spain | Giacomo Barbierato (Istituto Zooprofilattico Sperimentale delle Venezie / Ricerca e innovazione) |
| EPI_ISL_11259288 | A/common_crane/Spain/597-2_22VIR2142-15/2022 | A / H5N1 | Europe / Spain | Giacomo Barbierato (Istituto Zooprofilattico Sperimentale delle Venezie / Ricerca e innovazione) |
| EPI_ISL_11259289 | A/gray_heron/Spain/602-1_22VIR2142-17/2022 | A / H5N1 | Europe / Spain | Giacomo Barbierato (Istituto Zooprofilattico Sperimentale delle Venezie / Ricerca e innovazione) |
| EPI_ISL_11259290 | A/stork/Spain/729-1_22VIR2142-18/2022 | A / H5N1 | Europe / Spain | Giacomo Barbierato (Istituto Zooprofilattico Sperimentale delle Venezie / Ricerca e innovazione) |
| EPI_ISL_11259291 | A/turkey/Spain/140-38_22VIR2142-19/2022 | A / H5N1 | Europe / Spain | Giacomo Barbierato (Istituto Zooprofilattico Sperimentale delle Venezie / Ricerca e innovazione) |
| EPI_ISL_11259292 | A/swan/Spain/4087-1_22VIR2142-1/2021 | A / H5N1 | Europe / Spain | Giacomo Barbierato (Istituto Zooprofilattico Sperimentale delle Venezie / Ricerca e innovazione) |
| EPI_ISL_11259293 | A/chicken/Spain/340-37_22VIR2142-20/2022 | A / H5N1 | Europe / Spain | Giacomo Barbierato (Istituto Zooprofilattico Sperimentale delle Venezie / Ricerca e innovazione) |
| EPI_ISL_11259294 | A/chicken/Spain/452-1_22VIR2142-21/2022 | A / H5N1 | Europe / Spain | Giacomo Barbierato (Istituto Zooprofilattico Sperimentale delle Venezie / Ricerca e innovazione) |
| EPI_ISL_11259295 | A/chicken/Spain/452-17_22VIR2142-22/2022 | A / H5N1 | Europe / Spain | Giacomo Barbierato (Istituto Zooprofilattico Sperimentale delle Venezie / Ricerca e innovazione) |
| EPI_ISL_11259296 | A/turkey/Spain/455-83_22VIR2142-23/2022 | A / H5N1 | Europe / Spain | Giacomo Barbierato (Istituto Zooprofilattico Sperimentale delle Venezie / Ricerca e innovazione) |
| EPI_ISL_11259297 | A/turkey/Spain/455-96_22VIR2142-24/2022 | A / H5N1 | Europe / Spain | Giacomo Barbierato (Istituto Zooprofilattico Sperimentale delle Venezie / Ricerca e innovazione) |

| | | | | |
|---|---|---|---|---|
| EPI_ISL_11259298 | A/turkey/Spain/489-21_22VIR2142-25/2022 | A / H5N1 | Europe / Spain | Giacomo Barbierato (Istituto Zooprofilattico Sperimentale delle Venezie / Ricerca e innovazione) |
| EPI_ISL_11259299 | A/turkey/Spain/490-22_22VIR2142-26/2022 | A / H5N1 | Europe / Spain | Giacomo Barbierato (Istituto Zooprofilattico Sperimentale delle Venezie / Ricerca e innovazione) |
| EPI_ISL_11259300 | A/turkey/Spain/490-24_22VIR2142-27/2022 | A / H5N1 | Europe / Spain | Giacomo Barbierato (Istituto Zooprofilattico Sperimentale delle Venezie / Ricerca e innovazione) |
| EPI_ISL_11259301 | A/turkey/Spain/540-26_22VIR2142-28/2022 | A / H5N1 | Europe / Spain | Giacomo Barbierato (Istituto Zooprofilattico Sperimentale delle Venezie / Ricerca e innovazione) |
| EPI_ISL_11259302 | A/chicken/Spain/562-1_22VIR2142-29/2022 | A / H5N1 | Europe / Spain | Giacomo Barbierato (Istituto Zooprofilattico Sperimentale delle Venezie / Ricerca e innovazione) |
| EPI_ISL_11259303 | A/goose/Spain/65-3_22VIR2142-2/2022 | A / H5N1 | Europe / Spain | Giacomo Barbierato (Istituto Zooprofilattico Sperimentale delle Venezie / Ricerca e innovazione) |
| EPI_ISL_11259304 | A/chicken/Spain/564-4_22VIR2142-30/2022 | A / H5N1 | Europe / Spain | Giacomo Barbierato (Istituto Zooprofilattico Sperimentale delle Venezie / Ricerca e innovazione) |
| EPI_ISL_11259305 | A/chicken/Spain/564-11_22VIR2142-31/2022 | A / H5N1 | Europe / Spain | Giacomo Barbierato (Istituto Zooprofilattico Sperimentale delle Venezie / Ricerca e innovazione) |
| EPI_ISL_11259306 | A/turkey/Spain/586-4_22VIR2142-32/2022 | A / H5N1 | Europe / Spain | Giacomo Barbierato (Istituto Zooprofilattico Sperimentale delle Venezie / Ricerca e innovazione) |
| EPI_ISL_11259307 | A/chicken/Spain/587-1_22VIR2142-33/2022 | A / H5N1 | Europe / Spain | Giacomo Barbierato (Istituto Zooprofilattico Sperimentale delle Venezie / Ricerca e innovazione) |
| EPI_ISL_11259308 | A/chicken/Spain/622-8_22VIR2142-34/2022 | A / H5N1 | Europe / Spain | Giacomo Barbierato (Istituto Zooprofilattico Sperimentale delle Venezie / Ricerca e innovazione) |
| EPI_ISL_11259309 | A/chicken/Spain/644-8_22VIR2142-35/2022 | A / H5N1 | Europe / Spain | Giacomo Barbierato (Istituto Zooprofilattico Sperimentale delle Venezie / Ricerca e innovazione) |
| EPI_ISL_11259310 | A/turkey/Spain/645-1_22VIR2142-36/2022 | A / H5N1 | Europe / Spain | Giacomo Barbierato (Istituto Zooprofilattico Sperimentale delle Venezie / Ricerca e innovazione) |
| EPI_ISL_11259311 | A/turkey/Spain/646-7_22VIR2142-37/2022 | A / H5N1 | Europe / Spain | Giacomo Barbierato (Istituto Zooprofilattico Sperimentale delle Venezie / Ricerca e innovazione) |
| EPI_ISL_11259312 | A/chicken/Spain/649-6_22VIR2142-38/2022 | A / H5N1 | Europe / Spain | Giacomo Barbierato (Istituto Zooprofilattico Sperimentale delle Venezie / Ricerca e innovazione) |
| EPI_ISL_11259313 | A/gray_heron/Spain/88-2_22VIR2142-3/2022 | A / H5N1 | Europe / Spain | Giacomo Barbierato (Istituto Zooprofilattico Sperimentale delle Venezie / Ricerca e innovazione) |
| EPI_ISL_11259314 | A/duck/Spain/570-2_22VIR2142-40/2022 | A / H5N1 | Europe / Spain | Giacomo Barbierato (Istituto Zooprofilattico Sperimentale delle Venezie / Ricerca e innovazione) |
| EPI_ISL_11259315 | A/goose/Spain/88-3_22VIR2142-4/2022 | A / H5N1 | Europe / Spain | Giacomo Barbierato (Istituto Zooprofilattico Sperimentale delle Venezie / Ricerca e innovazione) |
| EPI_ISL_11259316 | A/goose/Spain/141-9_22VIR2142-6/2022 | A / H5N1 | Europe / Spain | Giacomo Barbierato (Istituto Zooprofilattico Sperimentale delle Venezie / Ricerca e innovazione) |
| EPI_ISL_11259317 | A/stork/Spain/234-2_22VIR2142-7/2022 | A / H5N1 | Europe / Spain | Giacomo Barbierato (Istituto Zooprofilattico Sperimentale delle Venezie / Ricerca e innovazione) |
| EPI_ISL_11259318 | A/goose/Spain/239-1_22VIR2142-8/2022 | A / H5N1 | Europe / Spain | Giacomo Barbierato (Istituto Zooprofilattico Sperimentale delle Venezie / Ricerca e innovazione) |
| EPI_ISL_11259319 | A/goose/Spain/294-2_22VIR2142-9/2022 | A / H5N1 | Europe / Spain | Giacomo Barbierato (Istituto Zooprofilattico Sperimentale delle Venezie / Ricerca e innovazione) |
| EPI_ISL_11260218 | A/mute_swan/Ireland/037311_22VIR1325-13/2021 | A / H5N1 | Europe / Ireland | Giacomo Barbierato (Istituto Zooprofilattico Sperimentale delle Venezie / Ricerca e innovazione) |
| EPI_ISL_11268632 | A/mute swan/Croatia/26/2022 | A / H5N1 | Europe / Croatia / Vukovarsko-Srijemska Zupanija / Ivankovo (45°15'31.29 18°42'28.27) | Vladimir Savi? (Croatian Veterinary Institute / Poultry Centre) |
| EPI_ISL_11325034 | A/turkey/Egypt/Cairo/AH/2019 | A / H5N8 | Africa / Egypt / Cairo | El-Sayed M. Abdelwhab (Friedrich-Loeffler-Institut) |
| EPI_ISL_11325035 | A/turkey/Egypt/Alex-AH/2019 | A / H5N8 | Africa / Egypt / Alexandria | El-Sayed M. Abdelwhab (Friedrich-Loeffler-Institut) |
| EPI_ISL_11325036 | A/turkey/Egypt/Alex-AH1/2019 | A / H5N8 | Africa / Egypt / Alexandria | El-Sayed M. Abdelwhab (Friedrich-Loeffler-Institut) |
| EPI_ISL_11325037 | A/duck/Egypt/Elbehera-AH2/2019 | A / H5N8 | Africa / Egypt / Elbehera | El-Sayed M. Abdelwhab (Friedrich-Loeffler-Institut) |
| EPI_ISL_11325038 | A/duck/Egypt/Behera-HB2-AH/2020 | A / H5N8 | Africa / Egypt / Elbehera | El-Sayed M. Abdelwhab (Friedrich-Loeffler-Institut) |
| EPI_ISL_11325039 | A/duck/Egypt/Behera-AH1/2019 | A / H5N8 | Africa / Egypt / Elbehera | El-Sayed M. Abdelwhab (Friedrich-Loeffler-Institut) |
| EPI_ISL_11325040 | A/chicken/Egypt/sohag-AH/2020 | A / H5N8 | Africa / Egypt / Sohag | El-Sayed M. Abdelwhab (Friedrich-Loeffler-Institut) |
| EPI_ISL_11325041 | A/chicken/Egypt/qalyubia-layer-AH/2020 | A / H5N8 | Africa / Egypt / Qalyubia | El-Sayed M. Abdelwhab (Friedrich-Loeffler-Institut) |
| EPI_ISL_11325042 | A/chicken/Egypt/kafrelsheik-AH/2019 | A / H5N8 | Africa / Egypt / Kafr El-Sheikh | El-Sayed M. Abdelwhab (Friedrich-Loeffler-Institut) |
| EPI_ISL_11325043 | A/chicken/Egypt/Giza-HG4L-AH/2020 | A / H5N8 | Africa / Egypt / Giza | El-Sayed M. Abdelwhab (Friedrich-Loeffler-Institut) |
| EPI_ISL_11325044 | A/chicken/Egypt/Giza-AH/2020 | A / H5N8 | Africa / Egypt / Giza | El-Sayed M. Abdelwhab (Friedrich-Loeffler-Institut) |
| EPI_ISL_11325045 | A/chicken/Egypt/Elmonufia-backyard-AH/2019 | A / H5N8 | Africa / Egypt / El-Monoufiya | El-Sayed M. Abdelwhab (Friedrich-Loeffler-Institut) |
| EPI_ISL_11325046 | A/chicken/Egypt/Elmonufia-AH/2020 | A / H5N8 | Africa / Egypt / El-Monoufiya | El-Sayed M. Abdelwhab (Friedrich-Loeffler-Institut) |
| EPI_ISL_11325047 | A/chicken/Egypt/Cairo-HC11B-AH/2020 | A / H5N8 | Africa / Egypt / Cairo | El-Sayed M. Abdelwhab (Friedrich-Loeffler-Institut) |
| EPI_ISL_11325048 | A/chicken/Egypt/behera-AH/2021 | A / H5N8 | Africa / Egypt / Elbehera | El-Sayed M. Abdelwhab (Friedrich-Loeffler-Institut) |
| EPI_ISL_11325049 | A/chicken/Egypt/Assiut-AH/2019 | A / H5N8 | Africa / Egypt / Assiut | El-Sayed M. Abdelwhab (Friedrich-Loeffler-Institut) |
| EPI_ISL_11325050 | A/chicken/Egypt/Alex-Breeder-AH/2021 | A / H5N8 | Africa / Egypt / Alexandria | El-Sayed M. Abdelwhab (Friedrich-Loeffler-Institut) |
| EPI_ISL_11325051 | A/chicken/Egypt/Alex-AH/2020 | A / H5N8 | Africa / Egypt / Alexandria | El-Sayed M. Abdelwhab (Friedrich-Loeffler-Institut) |
| EPI_ISL_11325052 | A/chicken/Egypt/Alex-AH2/2020 | A / H5N8 | Africa / Egypt / Alexandria | El-Sayed M. Abdelwhab (Friedrich-Loeffler-Institut) |
| EPI_ISL_11325053 | A/chicken/Egypt/Alex-AH1/2020 | A / H5N8 | Africa / Egypt / Alexandria | El-Sayed M. Abdelwhab (Friedrich-Loeffler-Institut) |
| EPI_ISL_11327088 | A/chicken/Czech Republic/61-1/2022 | A / H5N1 | Europe / Czech Republic / South Moravian Region / Okres Znoimo / Bojanovice, GPS: 48.969002 | Alexander Nagy (State Veterinary Institute Prague) |
| EPI_ISL_11327089 | A/chicken/Czech Republic/61-2/2022 | A / H5N1 | Europe / Czech Republic / South Moravian Region / Okres Znoimo / Bojanovice, GPS: 48.969002 | Alexander Nagy (State Veterinary Institute Prague) |
| EPI_ISL_11327090 | A/chicken/Czech Republic/63/2022 | A / H5N1 | Europe / Czech Republic / Ustecky Kraj / Okres Louny / Blazim GPS: 50.4061775N, 13.6296733E | Alexander Nagy (State Veterinary Institute Prague) |
| EPI_ISL_11330431 | A/white-tailed eagle/Hokkaido/22-RU-WTE-2/2022 | A / H5N1 | Asia / Japan / Hokkaido / Tomamae town | Norikazu Isoda (Graduate School of Veterinary Medicine, Hokkaido University / Laboratory of Microbiology) |
| EPI_ISL_1139014 | A/barnacle goose/Netherlands/20016511-002/2020 | A / H5N8 | Europe / Netherlands / Provincie Groningen / Lauwersoog | Rene Heutink (Wageningen Bioveterinary Research) |
| EPI_ISL_1139015 | A/wild bird/Netherlands/20016515-002/2020 | A / H5N8 | Europe / Netherlands / Provincie Friesland / Cornwerd | Rene Heutink (Wageningen Bioveterinary Research) |
| EPI_ISL_1139016 | A/mute swan/Netherlands/20016516-001/2020 | A / H5N8 | Europe / Netherlands / Provincie Friesland / Aldeboarn | Rene Heutink (Wageningen Bioveterinary Research) |
| EPI_ISL_1139017 | A/mute swan/Netherlands/20016634-001/2020 | A / H5N8 | Europe / Netherlands / South Holland / Berkenwoude | Rene Heutink (Wageningen Bioveterinary Research) |
| EPI_ISL_1139018 | A/cormorant/Netherlands/20016582-005/2020 | A / H5N8 | Europe / Netherlands / Provincie Noord-Holland / Edam-Volendam | Rene Heutink (Wageningen Bioveterinary Research) |
| EPI_ISL_1139019 | A/mute swan/Netherlands/20016618-001/2020 | A / H5N8 | Europe / Netherlands / Provincie Utrecht / Kockengen | Rene Heutink (Wageningen Bioveterinary Research) |
| EPI_ISL_1139020 | A/greylag goose/Netherlands/20016756-001/2020 | A / H5N8 | Europe / Netherlands / Provincie Friesland / Vegelinsoord | Rene Heutink (Wageningen Bioveterinary Research) |
| EPI_ISL_1139021 | A/eurasian wigeon/Netherlands/20016758-001/2020 | A / H5N8 | Europe / Netherlands / Provincie Friesland / Terherne | Rene Heutink (Wageningen Bioveterinary Research) |
| EPI_ISL_1139022 | A/barnacle goose/Netherlands/20016888-001/2020 | A / H5N8 | Europe / Netherlands / Provincie Friesland / Buren | Rene Heutink (Wageningen Bioveterinary Research) |
| EPI_ISL_1139023 | A/short-eared owl/Netherlands/20016896-017/2020 | A / H5N8 | Europe / Netherlands / Provincie Friesland / Holwerd | Rene Heutink (Wageningen Bioveterinary Research) |
| EPI_ISL_1139024 | A/mute swan/Netherlands/20016973-001/2020 | A / H5N8 | Europe / Netherlands / South Holland / Gemeente Rotterdam | Rene Heutink (Wageningen Bioveterinary Research) |
| EPI_ISL_1139025 | A/barnacle goose/Netherlands/20016974-002/2020 | A / H5N8 | Europe / Netherlands / Provincie Friesland / Gemeente Leeuwarden | Rene Heutink (Wageningen Bioveterinary Research) |
| EPI_ISL_1139026 | A/greylag goose/Netherlands/20016975-004/2020 | A / H5N8 | Europe / Netherlands / South Holland / Alphen a/d Rijn | Rene Heutink (Wageningen Bioveterinary Research) |
| EPI_ISL_1139027 | A/egyptian goose/Netherlands/20017027-002/2020 | A / H5N8 | Europe / Netherlands / South Holland / Gemeente Leiderdorp | Rene Heutink (Wageningen Bioveterinary Research) |
| EPI_ISL_1139028 | A/barnacle goose/Netherlands/20017051-001/2020 | A / H5N8 | Europe / Netherlands / Provincie Noord-Holland / Gemeente Landsmeer | Rene Heutink (Wageningen Bioveterinary Research) |
| EPI_ISL_1139029 | A/barnacle goose/Netherlands/20017051-006/2020 | A / H5N8 | Europe / Netherlands / Provincie Noord-Holland / Gemeente Landsmeer | Rene Heutink (Wageningen Bioveterinary Research) |
| EPI_ISL_1139030 | A/barnacle goose/Netherlands/20017052-001/2020 | A / H5N8 | Europe / Netherlands / Provincie Friesland / Gemeente Schiermonnikoog | Rene Heutink (Wageningen Bioveterinary Research) |
| EPI_ISL_1139031 | A/mute swan/Netherlands/20017131-006/2020 | A / H5N8 | Europe / Netherlands / South Holland / Nieuwerkerk a/d Ijssel | Rene Heutink (Wageningen Bioveterinary Research) |
| EPI_ISL_1139032 | A/barnacle goose/Netherlands/20017159-006/2020 | A / H5N8 | Europe / Netherlands / Provincie Friesland / Rijs | Rene Heutink (Wageningen Bioveterinary Research) |
| EPI_ISL_1139033 | A/greylag goose/Netherlands/20017399-006/2020 | A / H5N8 | Europe / Netherlands / South Holland / Hazerswoude-Dorp | Rene Heutink (Wageningen Bioveterinary Research) |
| EPI_ISL_1139034 | A/barnacle goose/Netherlands/20017405-002/2020 | A / H5N8 | Europe / Netherlands / Provincie Noord-Holland / Badhoevedorp | Rene Heutink (Wageningen Bioveterinary Research) |
| EPI_ISL_1139035 | A/greylag goose/Netherlands/20016877-001/2020 | A / H5N8 | Europe / Netherlands / Provincie Noord-Holland / Gemeente Heerhugowaard | Rene Heutink (Wageningen Bioveterinary Research) |
| EPI_ISL_1139036 | A/brant goose/Netherlands/20016948-002/2020 | A / H5N8 | Europe / Netherlands / Provincie Noord-Holland / Gemeente Amstelveen | Rene Heutink (Wageningen Bioveterinary Research) |
| EPI_ISL_1139037 | A/barnacle goose/Netherlands/20016951-001/2020 | A / H5N8 | Europe / Netherlands / South Holland / Pijnacker | Rene Heutink (Wageningen Bioveterinary Research) |

| | | | | |
|---|---|---|---|---|
| EPI_ISL_1139038 | A/wild goose/Netherlands/20016959-001/2020 | A / H5N8 | Europe / Netherlands / Provincie Limburg / Horst | Rene Heutink (Wageningen Bioveterinary Research) |
| EPI_ISL_1139039 | A/mute swan/Netherlands/20016960-001/2020 | A / H5N8 | Europe / Netherlands / South Holland / Zevenhoven | Rene Heutink (Wageningen Bioveterinary Research) |
| EPI_ISL_1139040 | A/mute swan/Netherlands/20017061-001/2020 | A / H5N8 | Europe / Netherlands / Provincie Drenthe / Wapserveen | Rene Heutink (Wageningen Bioveterinary Research) |
| EPI_ISL_1139041 | A/greylag goose/Netherlands/20017064-002/2020 | A / H5N8 | Europe / Netherlands / Provinvie Utrecht | Rene Heutink (Wageningen Bioveterinary Research) |
| EPI_ISL_1139042 | A/mute swan/Netherlands/20017153-002/2020 | A / H5N8 | Europe / Netherlands / South Holland / Rijpwetering | Rene Heutink (Wageningen Bioveterinary Research) |
| EPI_ISL_1139043 | A/mute swan/Netherlands/20017547-002/2020 | A / H5N8 | Europe / Netherlands / South Holland / Gemeente Zoeterwoude | Rene Heutink (Wageningen Bioveterinary Research) |
| EPI_ISL_1139044 | A/eurasian oystercatcher/Netherlands/20017557-003/2020 | A / H5N8 | Europe / Netherlands / Provincie Friesland / West Terschelling | Rene Heutink (Wageningen Bioveterinary Research) |
| EPI_ISL_1139045 | A/greylag goose/Netherlands/20017476-001/2020 | A / H5N8 | Europe / Netherlands / Provincie Noord-Holland / West-Graftdijk | Rene Heutink (Wageningen Bioveterinary Research) |
| EPI_ISL_1139046 | A/greater canada goose/Netherlands/20017479-002/2020 | A / H5N8 | Europe / Netherlands / South Holland / Krimpen a/d Ijssel | Rene Heutink (Wageningen Bioveterinary Research) |
| EPI_ISL_1139047 | A/northern lapwing/Netherlands/20017480-001/2020 | A / H5N8 | Europe / Netherlands / South Holland / Gemeente Spijkenisse | Rene Heutink (Wageningen Bioveterinary Research) |
| EPI_ISL_1139048 | A/wild goose/Netherlands/20017495-002/2020 | A / H5N8 | Europe / Netherlands / Provincie Groningen / Gemeente Winschoten | Rene Heutink (Wageningen Bioveterinary Research) |
| EPI_ISL_1139049 | A/gadwall/Netherlands/20017254-001/2020 | A / H5N8 | Europe / Netherlands / South Holland / Gemeente Delft | Rene Heutink (Wageningen Bioveterinary Research) |
| EPI_ISL_1139050 | A/northern lapwing/Netherlands/20017255-001/2020 | A / H5N8 | Europe / Netherlands / South Holland / Gemeente Delft | Rene Heutink (Wageningen Bioveterinary Research) |
| EPI_ISL_1139051 | A/greylag goose/Netherlands/20017256-002/2020 | A / H5N8 | Europe / Netherlands / Provincie Noord-Holland / Gemeente Zaanstad | Rene Heutink (Wageningen Bioveterinary Research) |
| EPI_ISL_1139076 | A/common kestrel/Netherlands/20017381-001/2020 | A / H5N8 | Europe / Netherlands / South Holland / Roelofarendsveen | Rene Heutink (Wageningen Bioveterinary Research) |
| EPI_ISL_1139078 | A/pink-footed goose/Netherlands/20017382-001/2020 | A / H5N8 | Europe / Netherlands / Provincie Limburg / Leudal | Rene Heutink (Wageningen Bioveterinary Research) |
| EPI_ISL_1139079 | A/barnacle goose/Netherlands/20016888-002/2020 | A / H5N8 | Europe / Netherlands / Provincie Friesland / Buren | Rene Heutink (Wageningen Bioveterinary Research) |
| EPI_ISL_1139080 | A/barnacle goose/Netherlands/20016888-003/2020 | A / H5N8 | Europe / Netherlands / Provincie Friesland / Buren | Rene Heutink (Wageningen Bioveterinary Research) |
| EPI_ISL_1139081 | A/greylag goose/Netherlands/20016896-001/2020 | A / H5N8 | Europe / Netherlands / Provincie Friesland / Holwerd | Rene Heutink (Wageningen Bioveterinary Research) |
| EPI_ISL_1139082 | A/barnacle goose/Netherlands/20016896-011/2020 | A / H5N8 | Europe / Netherlands / Provincie Friesland / Holwerd | Rene Heutink (Wageningen Bioveterinary Research) |
| EPI_ISL_1139083 | A/barnacle goose/Netherlands/20016896-012/2020 | A / H5N8 | Europe / Netherlands / Provincie Friesland / Holwerd | Rene Heutink (Wageningen Bioveterinary Research) |
| EPI_ISL_1139084 | A/eurasian wigeon/Netherlands/20016896-025/2020 | A / H5N8 | Europe / Netherlands / Provincie Friesland / Holwerd | Rene Heutink (Wageningen Bioveterinary Research) |
| EPI_ISL_1139085 | A/barnacle goose/Netherlands/20016935-003/2020 | A / H5N8 | Europe / Netherlands / Provincie Friesland / Holwerd | Rene Heutink (Wageningen Bioveterinary Research) |
| EPI_ISL_1139086 | A/greater canada goose/Netherlands/20017403-003/2020 | A / H5N8 | Europe / Netherlands / North Brabant / Rijswijk (NB) | Rene Heutink (Wageningen Bioveterinary Research) |
| EPI_ISL_1139087 | A/barnacle goose/Netherlands/20017557-001/2020 | A / H5N8 | Europe / Netherlands / Provincie Friesland / West Terschelling | Rene Heutink (Wageningen Bioveterinary Research) |
| EPI_ISL_1139088 | A/barnacle goose/Netherlands/20017557-002/2020 | A / H5N8 | Europe / Netherlands / Provincie Friesland / West Terschelling | Rene Heutink (Wageningen Bioveterinary Research) |
| EPI_ISL_1139089 | A/wild duck/Netherlands/20017794-001/2020 | A / H5N8 | Europe / Netherlands / Provincie Friesland / Grou | Rene Heutink (Wageningen Bioveterinary Research) |
| EPI_ISL_1139090 | A/common buzzard/Netherlands/20017824-001/2020 | A / H5N8 | Europe / Netherlands / North Brabant / Oudgastel | Rene Heutink (Wageningen Bioveterinary Research) |
| EPI_ISL_1139091 | A/muscovy duck/Netherlands/20017611-002/2020 | A / H5N8 | Europe / Netherlands / Provincie Friesland / Grou | Rene Heutink (Wageningen Bioveterinary Research) |
| EPI_ISL_1139092 | A/barnacle goose/Netherlands/20017713-002/2020 | A / H5N8 | Europe / Netherlands / Provincie Groningen / Den Andel | Rene Heutink (Wageningen Bioveterinary Research) |
| EPI_ISL_1139093 | A/gadwall/Netherlands/20017716-001/2020 | A / H5N8 | Europe / Netherlands / Provincie Noord-Holland / Zwaanshoek | Rene Heutink (Wageningen Bioveterinary Research) |
| EPI_ISL_1139094 | A/mute swan/Netherlands/20017717-002/2020 | A / H5N8 | Europe / Netherlands / Provincie Groningen / Noordlaren | Rene Heutink (Wageningen Bioveterinary Research) |
| EPI_ISL_1139095 | A/great egret/Netherlands/20017754-002/2020 | A / H5N8 | Europe / Netherlands / Provincie Utrecht / Haarzuilens | Rene Heutink (Wageningen Bioveterinary Research) |
| EPI_ISL_1139096 | A/wild goose/Netherlands/20017755-002/2020 | A / H5N8 | Europe / Netherlands / Provincie Utrecht / Haarzuilens | Rene Heutink (Wageningen Bioveterinary Research) |
| EPI_ISL_1139097 | A/wild goose/Netherlands/20017761-002/2020 | A / H5N8 | Europe / Netherlands / Provincie Utrecht / Waverveen | Rene Heutink (Wageningen Bioveterinary Research) |
| EPI_ISL_1139098 | A/eurasian wigeon/Netherlands/20017908-002/2020 | A / H5N8 | Europe / Netherlands / Provincie Groningen / Den Andel | Rene Heutink (Wageningen Bioveterinary Research) |
| EPI_ISL_1139099 | A/barnacle goose/Netherlands/20017984-004/2020 | A / H5N8 | Europe / Netherlands / Provincie Zeeland / Zierikzee | Rene Heutink (Wageningen Bioveterinary Research) |
| EPI_ISL_1139100 | A/barnacle goose/Netherlands/20017604-001/2020 | A / H5N8 | Europe / Netherlands / South Holland / Gemeente Oud-Beijerland | Rene Heutink (Wageningen Bioveterinary Research) |
| EPI_ISL_1139101 | A/swan/Netherlands/20017605-002/2020 | A / H5N1 | Europe / Netherlands / Provincie Noord-Holland / Groet | Rene Heutink (Wageningen Bioveterinary Research) |
| EPI_ISL_1139102 | A/swan/Netherlands/20017772-002/2020 | A / H5N1 | Europe / Netherlands / Provincie Noord-Holland / Schoorl | Rene Heutink (Wageningen Bioveterinary Research) |
| EPI_ISL_1139103 | A/peregrine falcon/Netherlands/20017773-002/2020 | A / H5N8 | Europe / Netherlands / Provincie Friesland / Oost-Vlieland | Rene Heutink (Wageningen Bioveterinary Research) |
| EPI_ISL_1139104 | A/wild goose/Netherlands/20017816-001/2020 | A / H5N8 | Europe / Netherlands / Provincie Friesland / Grou | Rene Heutink (Wageningen Bioveterinary Research) |
| EPI_ISL_1139105 | A/greylag goose/Netherlands/20016523-001/2020 | A / H5N8 | Europe / Netherlands / Provincie Noord-Holland / Velserbroek | Rene Heutink (Wageningen Bioveterinary Research) |
| EPI_ISL_1139106 | A/eurasian curlew/Netherlands/20016896-019/2020 | A / H5N8 | Europe / Netherlands / Provincie Friesland / Holwerd | Rene Heutink (Wageningen Bioveterinary Research) |
| EPI_ISL_1139107 | A/muscovy duck/Netherlands/20018067-001/2020 | A / H5N8 | Europe / Netherlands / South Holland / Gemeente Nieuwkoop | Rene Heutink (Wageningen Bioveterinary Research) |
| EPI_ISL_1139108 | A/mute swan/Netherlands/20018754-004/2020 | A / H5N8 | Europe / Netherlands / Provincie Friesland / Schraard | Rene Heutink (Wageningen Bioveterinary Research) |
| EPI_ISL_1139109 | A/mute swan/Netherlands/20018754-006/2020 | A / H5N8 | Europe / Netherlands / Provincie Friesland / Schraard | Rene Heutink (Wageningen Bioveterinary Research) |
| EPI_ISL_1139110 | A/swan/Netherlands/20018830-004/2020 | A / H5N8 | Europe / Netherlands / Provincie Utrecht / Mijdrecht | Rene Heutink (Wageningen Bioveterinary Research) |
| EPI_ISL_1139112 | A/greylag goose/Netherlands/20017386-001/2020 | A / H5N8 | Europe / Netherlands / Provincie Friesland / Gemeente Harlingen | Rene Heutink (Wageningen Bioveterinary Research) |
| EPI_ISL_1139113 | A/wild goose/Netherlands/20017819-001/2020 | A / H5N8 | Europe / Netherlands / Provincie Utrecht / Vinkeveen | Rene Heutink (Wageningen Bioveterinary Research) |
| EPI_ISL_1139159 | A/greylag goose/Netherlands/20017058-002/2020 | A / H5N8 | Europe / Netherlands / Provincie Utrecht / Kockengen | Rene Heutink (Wageningen Bioveterinary Research) |
| EPI_ISL_1139160 | A/pink-footed goose/Netherlands/20018068-001/2020 | A / H5N8 | Europe / Netherlands / Provincie Noord-Holland / Ankeveen | Rene Heutink (Wageningen Bioveterinary Research) |
| EPI_ISL_1139161 | A/greylag goose/Netherlands/20018070-002/2020 | A / H5N8 | Europe / Netherlands / South Holland / Harddinxveld | Rene Heutink (Wageningen Bioveterinary Research) |
| EPI_ISL_1139162 | A/black swan/Netherlands/20018185-001/2020 | A / H5N8 | Europe / Netherlands / South Holland / Gemeente Zoetermeer | Rene Heutink (Wageningen Bioveterinary Research) |
| EPI_ISL_1139163 | A/common buzzard/Netherlands/20018339-002/2020 | A / H5N8 | Europe / Netherlands / Provincie Friesland / Grou | Rene Heutink (Wageningen Bioveterinary Research) |
| EPI_ISL_1139164 | A/northern goshawk/Netherlands/20018560-002/2020 | A / H5N8 | Europe / Netherlands / South Holland / Leidschendam-Voorburg | Rene Heutink (Wageningen Bioveterinary Research) |
| EPI_ISL_1139165 | A/common snipe/Netherlands/20018931-003/2020 | A / H5N8 | Europe / Netherlands / Provincie Utrecht / Ameide | Rene Heutink (Wageningen Bioveterinary Research) |
| EPI_ISL_1139166 | A/wild goose/Netherlands/20018735-002/2020 | A / H5N8 | Europe / Netherlands / Provincie Overijssel / Gemeente Almelo | Rene Heutink (Wageningen Bioveterinary Research) |
| EPI_ISL_1139167 | A/barnacle goose/Netherlands/20018737-002/2020 | A / H5N8 | Europe / Netherlands / South Holland / Gemeente Papendrecht | Rene Heutink (Wageningen Bioveterinary Research) |
| EPI_ISL_1139168 | A/mute swan/Netherlands/20018738-001/2020 | A / H5N8 | Europe / Netherlands / South Holland / Rockanje | Rene Heutink (Wageningen Bioveterinary Research) |
| EPI_ISL_1139169 | A/peregrine falcon/Netherlands/20018821-002/2020 | A / H5N8 | Europe / Netherlands / Provincie Gelderland / Garderen | Rene Heutink (Wageningen Bioveterinary Research) |
| EPI_ISL_1139170 | A/wild goose/Netherlands/20018822-002/2020 | A / H5N8 | Europe / Netherlands / Provincie Utrecht / Gemeente Oudewater | Rene Heutink (Wageningen Bioveterinary Research) |
| EPI_ISL_1139171 | A/mute swan/Netherlands/20018824-002/2020 | A / H5N8 | Europe / Netherlands / Provincie Utrecht / Harmelen | Rene Heutink (Wageningen Bioveterinary Research) |
| EPI_ISL_1139172 | A/mute swan/Netherlands/20018923-001/2020 | A / H5N8 | Europe / Netherlands / South Holland / Stellendam | Rene Heutink (Wageningen Bioveterinary Research) |
| EPI_ISL_1139173 | A/mute swan/Netherlands/20019137-005/2020 | A / H5N8 | Europe / Netherlands / South Holland / Alphen a/d Rijn | Rene Heutink (Wageningen Bioveterinary Research) |
| EPI_ISL_11406398 | A/chicken/England/002070/2022 | A / H5N1 | Europe / United Kingdom / England | Alex Byrne (Animal and Plant Health Agency (APHA) / Virology Department) |
| EPI_ISL_11406399 | A/turkey/England/004737/2022 | A / H5N1 | Europe / United Kingdom / England | Alex Byrne (Animal and Plant Health Agency (APHA) / Virology Department) |
| EPI_ISL_11406401 | A/chicken/England/000187/2022 | A / H5N1 | Europe / United Kingdom / England | Alex Byrne (Animal and Plant Health Agency (APHA) / Virology Department) |
| EPI_ISL_11406402 | A/domestic_duck/England/007588/2022 | A / H5N1 | Europe / United Kingdom / England | Alex Byrne (Animal and Plant Health Agency (APHA) / Virology Department) |
| EPI_ISL_11406403 | A/black-headed_gull/England/306270/2022 | A / H5N1 | Europe / United Kingdom / England | Alex Byrne (Animal and Plant Health Agency (APHA) / Virology Department) |
| EPI_ISL_11449674 | A/Gallus_gallus/Belgium/3194_0001/2022 | A / H5N1 | Europe / Belgium / Provincie Oost-Vlaanderen / Sint-Gillis-Waas | Steven Van Borm (Sciensano, Department of Animal Infectious Diseases / Animal Infectious Diseases) |
| EPI_ISL_11504587 | A/chicken/Vietnam/HU14-LB11/2021 | A / H5N8 | Asia / Vietnam / Tinh Lang Son | Norikazu Isoda (Graduate School of Veterinary Medicine, Hokkaido University / Laboratory of Microbiology) |

| | | | | |
|---|---|---|---|---|
| EPI_ISL_11560233 | A/Brent goose/Netherlands/1/2022 | A / H5N1 | Europe / Netherlands / Stroe, Waddenkust | Mark Pronk (Erasmus Medical Center / Department of Virology) |
| EPI_ISL_11560235 | A/Barnacle Goose/Netherlands/16/2022 | A / H5N1 | Europe / Netherlands / Ameland | Mark Pronk (Erasmus Medical Center / Department of Virology) |
| EPI_ISL_11560236 | A/Barnacle Goose/Netherlands/17/2022 | A / H5N1 | Europe / Netherlands / Ameland | Mark Pronk (Erasmus Medical Center / Department of Virology) |
| EPI_ISL_11560323 | A/Barnacle Goose/Netherlands/18/2022 | A / H5N1 | Europe / Netherlands / Enkhuizen, Ijsselmeer, De Kreupel | Mark Pronk (Erasmus Medical Center / Department of Virology) |
| EPI_ISL_11560324 | A/Graylag goose/Netherlands/2/20222 | A / H5N1 | Europe / Netherlands / Woerden, Breeveld | Mark Pronk (Erasmus Medical Center / Department of Virology) |
| EPI_ISL_11560325 | A/Great black-backed Gull/Netherlands/3/2022 | A / H5N1 | Europe / Netherlands / Wieringerwerf, Zuiderdijkweg | Mark Pronk (Erasmus Medical Center / Department of Virology) |
| EPI_ISL_11561589 | A/chicken/England/011981/2022 | A / H5N1 | Europe / United Kingdom / England | Alex Byrne (Animal and Plant Health Agency (APHA) / Virology Department) |
| EPI_ISL_11561590 | A/chicken/England/012967/2022 | A / H5N1 | Europe / United Kingdom / England | Alex Byrne (Animal and Plant Health Agency (APHA) / Virology Department) |
| EPI_ISL_11561591 | A/domestic_duck/England/012973/2022 | A / H5N1 | Europe / United Kingdom / England | Alex Byrne (Animal and Plant Health Agency (APHA) / Virology Department) |
| EPI_ISL_11561592 | A/turkey/England/016515/2022 | A / H5N1 | Europe / United Kingdom / England | Alex Byrne (Animal and Plant Health Agency (APHA) / Virology Department) |
| EPI_ISL_11561593 | A/black-headed_gull/England/388256/2022 | A / H5N1 | Europe / United Kingdom / England | Alex Byrne (Animal and Plant Health Agency (APHA) / Virology Department) |
| EPI_ISL_11580255 | A/Greylag goose/Sweden/SVA220308SZ0382/FB000559/M-20[illegible] | A / H5N1 | Europe / Sweden / Skane Lan / Trelleborgs Kommun | Siamak Zohari (National Veterinary Institute) |
| EPI_ISL_11628212 | A/chicken/Virginia/22-004415-001/2022 | A / H5N1 | North America / United States / Virginia / Fauquier | Mary Lea Killian (National Veterinary Services Laboratories - USDA / Diagnostic Virology Laboratory) |
| EPI_ISL_11628213 | A/chicken/Kentucky/22-004416-001/2022 | A / H5N1 | North America / United States / Kentucky / Fulton | Mary Lea Killian (National Veterinary Services Laboratories - USDA / Diagnostic Virology Laboratory) |
| EPI_ISL_11628214 | A/chicken/Kentucky/22-004416-002/2022 | A / H5N1 | North America / United States / Kentucky / Fulton | Mary Lea Killian (National Veterinary Services Laboratories - USDA / Diagnostic Virology Laboratory) |
| EPI_ISL_11628215 | A/chicken/Kentucky/22-004416-003/2022 | A / H5N1 | North America / United States / Kentucky / Fulton | Mary Lea Killian (National Veterinary Services Laboratories - USDA / Diagnostic Virology Laboratory) |
| EPI_ISL_11628216 | A/turkey/Kentucky/22-004546-001/2022 | A / H5N1 | North America / United States / Kentucky / Webster | Mary Lea Killian (National Veterinary Services Laboratories - USDA / Diagnostic Virology Laboratory) |
| EPI_ISL_11628217 | A/turkey/Kentucky/22-004546-002/2022 | A / H5N1 | North America / United States / Kentucky / Webster | Mary Lea Killian (National Veterinary Services Laboratories - USDA / Diagnostic Virology Laboratory) |
| EPI_ISL_11628218 | A/turkey/Indiana/22-004688-002/2022 | A / H5N1 | North America / United States / Indiana / Dubois | Mary Lea Killian (National Veterinary Services Laboratories - USDA / Diagnostic Virology Laboratory) |
| EPI_ISL_11628219 | A/turkey/Indiana/22-004688-003/2022 | A / H5N1 | North America / United States / Indiana / Dubois | Mary Lea Killian (National Veterinary Services Laboratories - USDA / Diagnostic Virology Laboratory) |
| EPI_ISL_11628220 | A/turkey/Indiana/22-004688-006/2022 | A / H5N1 | North America / United States / Indiana / Dubois | Mary Lea Killian (National Veterinary Services Laboratories - USDA / Diagnostic Virology Laboratory) |
| EPI_ISL_11628221 | A/chicken/Maine/22-005158-001/2022 | A / H5N1 | North America / United States / Maine / Knox | Mary Lea Killian (National Veterinary Services Laboratories - USDA / Diagnostic Virology Laboratory) |
| EPI_ISL_11628222 | A/chicken/Delaware/22-005260-001/2022 | A / H5N1 | North America / United States / Deleware | Mary Lea Killian (National Veterinary Services Laboratories - USDA / Diagnostic Virology Laboratory) |
| EPI_ISL_11628223 | A/chicken/Delaware/22-005260-002/2022 | A / H5N1 | North America / United States / Deleware | Mary Lea Killian (National Veterinary Services Laboratories - USDA / Diagnostic Virology Laboratory) |
| EPI_ISL_11628224 | A/chicken/Delaware/22-005260-003/2022 | A / H5N1 | North America / United States / Deleware | Mary Lea Killian (National Veterinary Services Laboratories - USDA / Diagnostic Virology Laboratory) |
| EPI_ISL_11628225 | A/turkey/Indiana/22-005289-001/2022 | A / H5N1 | North America / United States / Indiana / Greene | Mary Lea Killian (National Veterinary Services Laboratories - USDA / Diagnostic Virology Laboratory) |
| EPI_ISL_11628226 | A/turkey/Indiana/22-005289-002/2022 | A / H5N1 | North America / United States / Indiana / Greene | Mary Lea Killian (National Veterinary Services Laboratories - USDA / Diagnostic Virology Laboratory) |
| EPI_ISL_11628227 | A/turkey/Indiana/22-005328-001/2022 | A / H5N1 | North America / United States / Indiana / Greene | Mary Lea Killian (National Veterinary Services Laboratories - USDA / Diagnostic Virology Laboratory) |
| EPI_ISL_11628228 | A/turkey/Indiana/22-005328-002/2022 | A / H5N1 | North America / United States / Indiana / Greene | Mary Lea Killian (National Veterinary Services Laboratories - USDA / Diagnostic Virology Laboratory) |
| EPI_ISL_11628229 | A/turkey/Michigan/22-005440-001/2022 | A / H5N1 | North America / United States / Michigan / Kalamazoo | Mary Lea Killian (National Veterinary Services Laboratories - USDA / Diagnostic Virology Laboratory) |
| EPI_ISL_11628230 | A/chicken/Michigan/22-005440-002/2022 | A / H5N1 | North America / United States / Michigan / Kalamazoo | Mary Lea Killian (National Veterinary Services Laboratories - USDA / Diagnostic Virology Laboratory) |
| EPI_ISL_11628231 | A/guinea fowl/Michigan/22-005440-003/2022 | A / H5N1 | North America / United States / Michigan / Kalamazoo | Mary Lea Killian (National Veterinary Services Laboratories - USDA / Diagnostic Virology Laboratory) |
| EPI_ISL_11628232 | A/chicken/Maine/22-005443-001/2022 | A / H5N1 | North America / United States / Maine / Knox | Mary Lea Killian (National Veterinary Services Laboratories - USDA / Diagnostic Virology Laboratory) |
| EPI_ISL_11628233 | A/pheasant/New York/22-005647-001/2022 | A / H5N1 | North America / United States / New York / Suffolk | Mary Lea Killian (National Veterinary Services Laboratories - USDA / Diagnostic Virology Laboratory) |
| EPI_ISL_11628234 | A/pheasant/New York/22-005647-002/2022 | A / H5N1 | North America / United States / New York / Suffolk | Mary Lea Killian (National Veterinary Services Laboratories - USDA / Diagnostic Virology Laboratory) |
| EPI_ISL_11628235 | A/turkey/Indiana/22-005696-001/2022 | A / H5N1 | North America / United States / Indiana / Dubois | Mary Lea Killian (National Veterinary Services Laboratories - USDA / Diagnostic Virology Laboratory) |
| EPI_ISL_11628236 | A/turkey/Indiana/22-005696-002/2022 | A / H5N1 | North America / United States / Indiana / Dubois | Mary Lea Killian (National Veterinary Services Laboratories - USDA / Diagnostic Virology Laboratory) |
| EPI_ISL_11628237 | A/turkey/Indiana/22-005696-003/2022 | A / H5N1 | North America / United States / Indiana / Dubois | Mary Lea Killian (National Veterinary Services Laboratories - USDA / Diagnostic Virology Laboratory) |
| EPI_ISL_11628238 | A/silkie chicken/Iowa/22-006114-001/2022 | A / H5N1 | North America / United States / Iowa / Pottawattamie | Mary Lea Killian (National Veterinary Services Laboratories - USDA / Diagnostic Virology Laboratory) |
| EPI_ISL_11628239 | A/chicken/Connecticut/22-006118-001/2022 | A / H5N1 | North America / United States / Conneticut | Mary Lea Killian (National Veterinary Services Laboratories - USDA / Diagnostic Virology Laboratory) |
| EPI_ISL_11628240 | A/turkey/Indiana/22-006259-001/2022 | A / H5N1 | North America / United States / Indiana / Dubois | Mary Lea Killian (National Veterinary Services Laboratories - USDA / Diagnostic Virology Laboratory) |
| EPI_ISL_11628241 | A/turkey/Indiana/22-006259-002/2022 | A / H5N1 | North America / United States / Indiana / Dubois | Mary Lea Killian (National Veterinary Services Laboratories - USDA / Diagnostic Virology Laboratory) |
| EPI_ISL_11628242 | A/turkey/Indiana/22-006261-001/2022 | A / H5N1 | North America / United States / Indiana / Dubois | Mary Lea Killian (National Veterinary Services Laboratories - USDA / Diagnostic Virology Laboratory) |
| EPI_ISL_11628243 | A/turkey/Indiana/22-006261-002/2022 | A / H5N1 | North America / United States / Indiana / Dubois | Mary Lea Killian (National Veterinary Services Laboratories - USDA / Diagnostic Virology Laboratory) |
| EPI_ISL_11628244 | A/chicken/Missouri/22-006569-001/2022 | A / H5N1 | North America / United States / Missouri / Stoddard | Mary Lea Killian (National Veterinary Services Laboratories - USDA / Diagnostic Virology Laboratory) |
| EPI_ISL_11628245 | A/chicken/Missouri/22-006569-002/2022 | A / H5N1 | North America / United States / Missouri / Stoddard | Mary Lea Killian (National Veterinary Services Laboratories - USDA / Diagnostic Virology Laboratory) |
| EPI_ISL_11628246 | A/chicken/Maryland/22-006578-001/2022 | A / H5N1 | North America / United States / Maryland / Cecil | Mary Lea Killian (National Veterinary Services Laboratories - USDA / Diagnostic Virology Laboratory) |
| EPI_ISL_11628247 | A/chicken/Maryland/22-006578-002/2022 | A / H5N1 | North America / United States / Maryland / Cecil | Mary Lea Killian (National Veterinary Services Laboratories - USDA / Diagnostic Virology Laboratory) |
| EPI_ISL_11628248 | A/chicken/Missouri/22-006639-001/2022 | A / H5N1 | North America / United States / Missouri / Bates | Mary Lea Killian (National Veterinary Services Laboratories - USDA / Diagnostic Virology Laboratory) |
| EPI_ISL_11628249 | A/turkey/South Dakota/22-006792-001/2022 | A / H5N1 | North America / United States / South Dakota / Charles | Mary Lea Killian (National Veterinary Services Laboratories - USDA / Diagnostic Virology Laboratory) |
| EPI_ISL_11628250 | A/turkey/South Dakota/22-006792-002/2022 | A / H5N1 | North America / United States / South Dakota / Charles Mix | Mary Lea Killian (National Veterinary Services Laboratories - USDA / Diagnostic Virology Laboratory) |
| EPI_ISL_11628251 | A/turkey/Iowa/22-006795-001/2022 | A / H5N1 | North America / United States / Iowa / Buena Vista | Mary Lea Killian (National Veterinary Services Laboratories - USDA / Diagnostic Virology Laboratory) |
| EPI_ISL_11628252 | A/turkey/Iowa/22-006795-002/2022 | A / H5N1 | North America / United States / Iowa / Buena Vista | Mary Lea Killian (National Veterinary Services Laboratories - USDA / Diagnostic Virology Laboratory) |
| EPI_ISL_11633613 | A/Cygnus columbianus/Hubei/117/2021 | A / H5N1 | Asia / China / Hubei Province | Ma Liping (Wuhan Institute of Virology, CAS ) |
| EPI_ISL_11633614 | A/Cygnus columbianus/Hubei/121/2021 | A / H5N1 | Asia / China / Hubei Province | Ma Liping (Wuhan Institute of Virology, CAS ) |
| EPI_ISL_11633616 | A/Cygnus columbianus/Hubei/123/2021 | A / H5N1 | Asia / China / Hubei Province | Ma Liping (Wuhan Institute of Virology, CAS ) |
| EPI_ISL_11633707 | A/Cygnus columbianus/Hubei/127/2021 | A / H5N1 | Asia / China / Hubei Province | Ma Liping (Wuhan Institute of Virology, CAS ) |
| EPI_ISL_11633708 | A/Cygnus columbianus/Hubei/126/2021 | A / H5N1 | Asia / China / Hubei Province | Ma Liping (Wuhan Institute of Virology, CAS ) |
| EPI_ISL_11633709 | A/Cygnus columbianus/Hubei/128/2021 | A / H5N1 | Asia / China / Hubei Province | Ma Liping (Wuhan Institute of Virology, CAS ) |
| EPI_ISL_11725869 | A/domestic duck/Germany-BB/AI06239/2021 | A / H5N1 | Europe / Germany / Brandenburg / Spree-Neiße | Jacqueline King (Friedrich-Loeffler-Institut) |
| EPI_ISL_11725962 | A/chicken/Germany-BB/AI06219/2021 | A / H5N1 | Europe / Germany / Brandenburg / Spree-Neiße | Jacqueline King (Friedrich-Loeffler-Institut) |
| EPI_ISL_11725988 | A/chicken/Germany-BB/AI06242/2021 | A / H5N1 | Europe / Germany / Brandenburg / Spree-Neiße | Jacqueline King (Friedrich-Loeffler-Institut) |
| EPI_ISL_11725991 | A/barnacle goose/Germany-MV/AI06175/2021 | A / H5N1 | Europe / Germany / Mecklenburg-Vorpommern / Vorpommern-Rügen | Jacqueline King (Friedrich-Loeffler-Institut) |
| EPI_ISL_11725994 | A/buzzard/Germany-SH/AI07099/2021 | A / H5N1 | Europe / Germany / Schleswig-Holstein / Dithmarschen | Jacqueline King (Friedrich-Loeffler-Institut) |
| EPI_ISL_11798566 | A/barnacle/goose_Denmark/19027-1.02/2021-04-18 | A / H5N1 | Europe / Denmark / Region Midtjylland / Viborg Kommune | Charlotte Kristiane Hjulsager (Statens Serum Institute / Microbiological Diagnostic and Virology) |
| EPI_ISL_11798568 | A/barnacle_goose/Denmark/24273-1.02/2021-10-26 | A / H5N1 | Europe / Denmark / Region Sjaland / Guldborgsund | Charlotte Kristiane Hjulsager (Statens Serum Institute / Microbiological Diagnostic and Virology) |
| EPI_ISL_11798569 | A/barnacle_goose/Denmark/24342-1.02/2021-10-30 | A / H5N1 | Europe / Denmark / Region Midtjylland / Arhus Kommune | Charlotte Kristiane Hjulsager (Statens Serum Institute / Microbiological Diagnostic and Virology) |
| EPI_ISL_11798570 | A/chicken/Denmark/24357-11/2021-11-02 | A / H5N1 | Europe / Denmark / Region Sjaland / Slagelse Kommune | Charlotte Kristiane Hjulsager (Statens Serum Institute / Microbiological Diagnostic and Virology) |
| EPI_ISL_11798571 | A/common_buzzard/Denmark/24271-1.02/2021-10-25 | A / H5N1 | Europe / Denmark / Region Sjaland / Slagelse Kommune | Charlotte Kristiane Hjulsager (Statens Serum Institute / Microbiological Diagnostic and Virology) |
| EPI_ISL_11798572 | A/Eurasian_wigeon/Denmark/24066-9/2021-10-13 | A / H5N1 | Europe / Denmark / Region Syddanmark / Tonder Kommune | Charlotte Kristiane Hjulsager (Statens Serum Institute / Microbiological Diagnostic and Virology) |
| EPI_ISL_11798573 | A/Eurasian_teal/Denmark/24115-2/2021-10-16 | A / H5N1 | Europe / Denmark / Region Syddanmark / Fano Kommune | Charlotte Kristiane Hjulsager (Statens Serum Institute / Microbiological Diagnostic and Virology) |
| EPI_ISL_11798574 | A/Eurasian_wigeon/Denmark/24279-1/2021-10-24 | A / H5N1 | Europe / Denmark / Region Syddanmark / Fano Kommune | Charlotte Kristiane Hjulsager (Statens Serum Institute / Microbiological Diagnostic and Virology) |

| | | | | |
|---|---|---|---|---|
| EPI_ISL_11798575 | A/European_herring_gull/Denmark/19968-1.02/2021-05-14 | A / H5N1 | Europe / Denmark / Region Midtjylland / Hostebro Kommune | Charlotte Kristiane Hjulsager (Statens Serum Institute / Microbiological Diagnostic and Virology) |
| EPI_ISL_11798577 | A/gray_heron/Denmark/24326-1.02/2021-10-28 | A / H5N1 | Europe / Denmark / Region Midtjylland / Hostebro Kommune | Charlotte Kristiane Hjulsager (Statens Serum Institute / Microbiological Diagnostic and Virology) |
| EPI_ISL_11798578 | A/greylag_goose/Denmark/24309-1.01/2021-10-27 | A / H5N1 | Europe / Denmark / Region Syddanmark / Faaborg-Midtfyn | Charlotte Kristiane Hjulsager (Statens Serum Institute / Microbiological Diagnostic and Virology) |
| EPI_ISL_11798579 | A/greylag_goose/Denmark/24343-1.02/2021-11-01 | A / H5N1 | Europe / Denmark / Region Nordjylland / Mariagerfjord | Charlotte Kristiane Hjulsager (Statens Serum Institute / Microbiological Diagnostic and Virology) |
| EPI_ISL_11798580 | A/turkey/Denmark/24325-25/2021-10-30 | A / H5N1 | Europe / Denmark / Region Sjaland / Slagelse Kommune | Charlotte Kristiane Hjulsager (Statens Serum Institute / Microbiological Diagnostic and Virology) |
| EPI_ISL_1180234 | A/mute swan/Czech Republic/1656-1/2021 | A / H5N8 | Europe / Czech Republic / Jihocesky Kraj / Okres Ceske Budejovice / Haklovy Dvory, Starohaklovs | Alexander Nagy (State Veterinary Institute Prague) |
| EPI_ISL_1184508 | A/chicken/Miyazaki/L10T/2021 | A / H5N8 | Asia / Japan / Miyazaki | Takehiko Saito (National Institute of Animal Health) |
| EPI_ISL_1184509 | A/chicken/Miyazaki/L1T/2021 | A / H5N8 | Asia / Japan / Miyazaki | Takehiko Saito (National Institute of Animal Health) |
| EPI_ISL_1184510 | A/chicken/Miyazaki/L4T/2021 | A / H5N8 | Asia / Japan / Miyazaki | Takehiko Saito (National Institute of Animal Health) |
| EPI_ISL_1184511 | A/chicken/Miyazaki/L9T/2021 | A / H5N8 | Asia / Japan / Miyazaki | Takehiko Saito (National Institute of Animal Health) |
| EPI_ISL_1184517 | A/eastern buzzard/Toyama/160208C/2021 | A / H5N8 | Asia / Japan / Toyama | Takehiko Saito (National Institute of Animal Health) |
| EPI_ISL_1184518 | A/eastern buzzard/Toyama/160208T/2021 | A / H5N8 | Asia / Japan / Toyama | Takehiko Saito (National Institute of Animal Health) |
| EPI_ISL_1185026 | A/chicken/Astrakhan/2171-1/2020 | A / H5N8 | Europe / Russian Federation / Astrakhan Oblast | Nikolay Zinyakov (Federal Centre for Animal Health (ARRIAH) / OIE Regional Reference Laboratory) |
| EPI_ISL_11880371 | A/Gallus_gallus/Belgium/4190_0002/2022 | A / H5N1 | Europe / Belgium / Provincie West-Vlaanderen / Meulebeke | Steven Van Borm (Sciensano, Department of Animal Infectious Diseases / Animal Infectious Diseases) |
| EPI_ISL_11897507 | A/chicken/Maryland/22-006578-001-original/2022 | A / H5N1 | North America / United States / Maryland / Cecil | Mary Lea Killian (National Veterinary Services Laboratories - USDA / Diagnostic Virology Laboratory) |
| EPI_ISL_11897508 | A/chicken/Maryland/22-006578-002-original/2022 | A / H5N1 | North America / United States / Maryland / Cecil | Mary Lea Killian (National Veterinary Services Laboratories - USDA / Diagnostic Virology Laboratory) |
| EPI_ISL_11897667 | A/chicken/Missouri/22-006639-001-original/2022 | A / H5N1 | North America / United States / Missouri / Bates | Mary Lea Killian (National Veterinary Services Laboratories - USDA / Diagnostic Virology Laboratory) |
| EPI_ISL_11897668 | A/turkey/South Dakota/22-006792-001-original/2022 | A / H5N1 | North America / United States / South Dakota / Charles Mix | Mary Lea Killian (National Veterinary Services Laboratories - USDA / Diagnostic Virology Laboratory) |
| EPI_ISL_11897669 | A/turkey/South Dakota/22-006792-002-original/2022 | A / H5N1 | North America / United States / South Dakota / Charles Mix | Mary Lea Killian (National Veterinary Services Laboratories - USDA / Diagnostic Virology Laboratory) |
| EPI_ISL_11897670 | A/turkey/Iowa/22-006795-001-original/2022 | A / H5N1 | North America / United States / Iowa / Buena Vista | Mary Lea Killian (National Veterinary Services Laboratories - USDA / Diagnostic Virology Laboratory) |
| EPI_ISL_11897671 | A/turkey/Iowa/22-006795-002-original/2022 | A / H5N1 | North America / United States / Iowa / Buena Vista | Mary Lea Killian (National Veterinary Services Laboratories - USDA / Diagnostic Virology Laboratory) |
| EPI_ISL_11897672 | A/turkey/Missouri/22-006944-002-original/2022 | A / H5N1 | North America / United States / Missouri / Jasper | Mary Lea Killian (National Veterinary Services Laboratories - USDA / Diagnostic Virology Laboratory) |
| EPI_ISL_11897673 | A/chicken/Delaware/22-006945-001-original/2022 | A / H5N1 | North America / United States / Delaware / New Castle | Mary Lea Killian (National Veterinary Services Laboratories - USDA / Diagnostic Virology Laboratory) |
| EPI_ISL_11897674 | A/chicken/Delaware/22-006945-002-original/2022 | A / H5N1 | North America / United States / Delaware / New Castle | Mary Lea Killian (National Veterinary Services Laboratories - USDA / Diagnostic Virology Laboratory) |
| EPI_ISL_11897675 | A/chicken/Maryland/22-006948-001-original/2022 | A / H5N1 | North America / United States / Maryland / Queen Annes | Mary Lea Killian (National Veterinary Services Laboratories - USDA / Diagnostic Virology Laboratory) |
| EPI_ISL_11897676 | A/chicken/Maryland/22-007086-001-original/2022 | A / H5N1 | North America / United States / Maryland / Queen Annes | Mary Lea Killian (National Veterinary Services Laboratories - USDA / Diagnostic Virology Laboratory) |
| EPI_ISL_11897677 | A/turkey/Missouri/22-007087-001-original/2022 | A / H5N1 | North America / United States / Missouri / Lawrence | Mary Lea Killian (National Veterinary Services Laboratories - USDA / Diagnostic Virology Laboratory) |
| EPI_ISL_11897678 | A/turkey/Missouri/22-007087-002-original/2022 | A / H5N1 | North America / United States / Missouri / Lawrence | Mary Lea Killian (National Veterinary Services Laboratories - USDA / Diagnostic Virology Laboratory) |
| EPI_ISL_11897679 | A/chicken/Maryland/22-007273-001-original/2022 | A / H5N1 | North America / United States / Maryland / Cecil | Mary Lea Killian (National Veterinary Services Laboratories - USDA / Diagnostic Virology Laboratory) |
| EPI_ISL_11897680 | A/chicken/Maryland/22-007273-002-original/2022 | A / H5N1 | North America / United States / Maryland / Cecil | Mary Lea Killian (National Veterinary Services Laboratories - USDA / Diagnostic Virology Laboratory) |
| EPI_ISL_11897681 | A/chicken/Iowa/22-007376-001-original/2022 | A / H5N1 | North America / United States / Iowa / Taylor | Mary Lea Killian (National Veterinary Services Laboratories - USDA / Diagnostic Virology Laboratory) |
| EPI_ISL_11897682 | A/chicken/Iowa/22-007376-002-original/2022 | A / H5N1 | North America / United States / Iowa / Taylor | Mary Lea Killian (National Veterinary Services Laboratories - USDA / Diagnostic Virology Laboratory) |
| EPI_ISL_11897683 | A/guinea fowl/Illinois/22-007382-001-original/2022 | A / H5N1 | North America / United States / Illinois / Maclean | Mary Lea Killian (National Veterinary Services Laboratories - USDA / Diagnostic Virology Laboratory) |
| EPI_ISL_11897684 | A/poultry/Kansas/22-007391-001-original/2022 | A / H5N1 | North America / United States / Kansas / Franklin | Mary Lea Killian (National Veterinary Services Laboratories - USDA / Diagnostic Virology Laboratory) |
| EPI_ISL_11897685 | A/poultry/Kansas/22-007391-002-original/2022 | A / H5N1 | North America / United States / Kansas / Franklin | Mary Lea Killian (National Veterinary Services Laboratories - USDA / Diagnostic Virology Laboratory) |
| EPI_ISL_11897686 | A/turkey/Maine/22-007410-001-original/2022 | A / H5N1 | North America / United States / Maine / Lincoln | Mary Lea Killian (National Veterinary Services Laboratories - USDA / Diagnostic Virology Laboratory) |
| EPI_ISL_11897687 | A/chicken/Maine/22-007410-002-original/2022 | A / H5N1 | North America / United States / Maine / Lincoln | Mary Lea Killian (National Veterinary Services Laboratories - USDA / Diagnostic Virology Laboratory) |
| EPI_ISL_11897688 | A/turkey/South Dakota/22-007534-001-original/2022 | A / H5N1 | North America / United States / South Dakota / Charles Mix | Mary Lea Killian (National Veterinary Services Laboratories - USDA / Diagnostic Virology Laboratory) |
| EPI_ISL_11897689 | A/turkey/South Dakota/22-007534-002-original/2022 | A / H5N1 | North America / United States / South Dakota / Charles Mix | Mary Lea Killian (National Veterinary Services Laboratories - USDA / Diagnostic Virology Laboratory) |
| EPI_ISL_11897690 | A/chicken/Wisconsin/22-007545-001-original/2022 | A / H5N1 | North America / United States / Wisconsin / Jefferson | Mary Lea Killian (National Veterinary Services Laboratories - USDA / Diagnostic Virology Laboratory) |
| EPI_ISL_11897691 | A/chicken/Wisconsin/22-007545-002-original/2022 | A / H5N1 | North America / United States / Wisconsin / Jefferson | Mary Lea Killian (National Veterinary Services Laboratories - USDA / Diagnostic Virology Laboratory) |
| EPI_ISL_11897692 | A/chicken/Maine/22-007582-001-original/2022 | A / H5N1 | North America / United States / Maine / York | Mary Lea Killian (National Veterinary Services Laboratories - USDA / Diagnostic Virology Laboratory) |
| EPI_ISL_11897693 | A/chicken/Missouri/22-007677-001-original/2022 | A / H5N1 | North America / United States / Missouri / New London | Mary Lea Killian (National Veterinary Services Laboratories - USDA / Diagnostic Virology Laboratory) |
| EPI_ISL_11897694 | A/chicken/Missouri/22-007677-002-original/2022 | A / H5N1 | North America / United States / Missouri / New London | Mary Lea Killian (National Veterinary Services Laboratories - USDA / Diagnostic Virology Laboratory) |
| EPI_ISL_11897695 | A/chicken/Nebraska/22-007805-001-original/2022 | A / H5N1 | North America / United States / Nebraska / Merrick | Mary Lea Killian (National Veterinary Services Laboratories - USDA / Diagnostic Virology Laboratory) |
| EPI_ISL_11897696 | A/waterfowl/Nebraska/22-007805-002-original/2022 | A / H5N1 | North America / United States / Nebraska / Merrick | Mary Lea Killian (National Veterinary Services Laboratories - USDA / Diagnostic Virology Laboratory) |
| EPI_ISL_11897697 | A/turkey/New Hampshire/22-007886-001-original/2022 | A / H5N1 | North America / United States / New Hampshire / Rockingham | Mary Lea Killian (National Veterinary Services Laboratories - USDA / Diagnostic Virology Laboratory) |
| EPI_ISL_11897698 | A/chicken/Delaware/22-008054-002-original/2022 | A / H5N1 | North America / United States / Delaware / Kent | Mary Lea Killian (National Veterinary Services Laboratories - USDA / Diagnostic Virology Laboratory) |
| EPI_ISL_11897699 | A/chicken/Delaware/22-008054-003-original/2022 | A / H5N1 | North America / United States / Delaware / Kent | Mary Lea Killian (National Veterinary Services Laboratories - USDA / Diagnostic Virology Laboratory) |
| EPI_ISL_11897700 | A/guinea fowl/Maine/22-008064-001-original/2022 | A / H5N1 | North America / United States / Maine / Lincoln | Mary Lea Killian (National Veterinary Services Laboratories - USDA / Diagnostic Virology Laboratory) |
| EPI_ISL_11897701 | A/chicken/Maine/22-008064-002-original/2022 | A / H5N1 | North America / United States / Maine / Lincoln | Mary Lea Killian (National Veterinary Services Laboratories - USDA / Diagnostic Virology Laboratory) |
| EPI_ISL_11897702 | A/duck/Kansas/22-008114-001-original/2022 | A / H5N1 | North America / United States / Kansas / Sedgewick | Mary Lea Killian (National Veterinary Services Laboratories - USDA / Diagnostic Virology Laboratory) |
| EPI_ISL_11897703 | A/chicken/Kansas/22-008114-002-original/2022 | A / H5N1 | North America / United States / Kansas / Sedgewick | Mary Lea Killian (National Veterinary Services Laboratories - USDA / Diagnostic Virology Laboratory) |
| EPI_ISL_11897704 | A/chicken/Iowa/22-008176-003-original/2022 | A / H5N1 | North America / United States / Iowa / Buena Vista | Mary Lea Killian (National Veterinary Services Laboratories - USDA / Diagnostic Virology Laboratory) |
| EPI_ISL_1191587 | A/chicken/Czech Republic/1566-2/2021 | A / H5N8 | Europe / Czech Republic / Jihocesky Kraj / Okres Tabor / Dlouha Lhota/Chynov; GPS 49.3561703 | Alexander Nagy (State Veterinary Institute Prague) |
| EPI_ISL_11922807 | A/chicken/Poland/H182_22VIR2515-1/2022 | A / H5N2 | Europe / Poland | Giacomo Barbierato (Istituto Zooprofilattico Sperimentale delle Venezie / Ricerca e innovazione) |
| EPI_ISL_11922808 | A/duck/Poland/H188_22VIR2515-2/2022 | A / H5N1 | Europe / Poland | Giacomo Barbierato (Istituto Zooprofilattico Sperimentale delle Venezie / Ricerca e innovazione) |
| EPI_ISL_11922809 | A/chicken/Poland/H157_22VIR2515-3/2022 | A / H5N1 | Europe / Poland | Giacomo Barbierato (Istituto Zooprofilattico Sperimentale delle Venezie / Ricerca e innovazione) |
| EPI_ISL_11922810 | A/duck/Poland/H126_22VIR2515-4/2022 | A / H5N1 | Europe / Poland | Giacomo Barbierato (Istituto Zooprofilattico Sperimentale delle Venezie / Ricerca e innovazione) |
| EPI_ISL_11922811 | A/goose/Poland/H124_22VIR2515-5/2022 | A / H5N1 | Europe / Poland | Giacomo Barbierato (Istituto Zooprofilattico Sperimentale delle Venezie / Ricerca e innovazione) |
| EPI_ISL_11922812 | A/chicken/Poland/H071_22VIR2515-6/2022 | A / H5N1 | Europe / Poland | Giacomo Barbierato (Istituto Zooprofilattico Sperimentale delle Venezie / Ricerca e innovazione) |
| EPI_ISL_11922813 | A/swan/Poland/MB078_22VIR2515-7/2022 | A / H5N1 | Europe / Poland | Giacomo Barbierato (Istituto Zooprofilattico Sperimentale delle Venezie / Ricerca e innovazione) |
| EPI_ISL_11922814 | A/swan/Poland/MB083_22VIR2515-8/2022 | A / H5N1 | Europe / Poland | Giacomo Barbierato (Istituto Zooprofilattico Sperimentale delle Venezie / Ricerca e innovazione) |
| EPI_ISL_11922815 | A/swan/Romania/16905_22VIR2749-1/2021 | A / H5N1 | Europe / Romania | Giacomo Barbierato (Istituto Zooprofilattico Sperimentale delle Venezie / Ricerca e innovazione) |
| EPI_ISL_11922816 | A/swan/Romania/10324_22VIR2749-2/2022 | A / H5N1 | Europe / Romania | Giacomo Barbierato (Istituto Zooprofilattico Sperimentale delle Venezie / Ricerca e innovazione) |
| EPI_ISL_11922817 | A/swan/Romania/10394_22VIR2749-3/2022 | A / H5N1 | Europe / Romania | Giacomo Barbierato (Istituto Zooprofilattico Sperimentale delle Venezie / Ricerca e innovazione) |
| EPI_ISL_11922818 | A/swan/Romania/10455_22VIR2749-4/2022 | A / H5N1 | Europe / Romania | Giacomo Barbierato (Istituto Zooprofilattico Sperimentale delle Venezie / Ricerca e innovazione) |
| EPI_ISL_11922819 | A/laying_hen/Romania/10470_22VIR2749-5/2022 | A / H5N1 | Europe / Romania | Giacomo Barbierato (Istituto Zooprofilattico Sperimentale delle Venezie / Ricerca e innovazione) |
| EPI_ISL_11922820 | A/swan/Romania/10656_22VIR2749-6/2022 | A / H5N1 | Europe / Romania | Giacomo Barbierato (Istituto Zooprofilattico Sperimentale delle Venezie / Ricerca e innovazione) |
| EPI_ISL_11922821 | A/swan/Romania/10678_22VIR2749-7/2022 | A / H5N1 | Europe / Romania | Giacomo Barbierato (Istituto Zooprofilattico Sperimentale delle Venezie / Ricerca e innovazione) |
| EPI_ISL_11922822 | A/swan/Romania/10986_22VIR2749-8/2022 | A / H5N1 | Europe / Romania | Giacomo Barbierato (Istituto Zooprofilattico Sperimentale delle Venezie / Ricerca e innovazione) |
| EPI_ISL_11971464 | A/turkey/South Dakota/22-008239-001-original/2022 | A / H5N1 | North America / United States / South Dakota / Kingsbury | Mary Lea Killian (National Veterinary Services Laboratories - USDA / Diagnostic Virology Laboratory) |

| | | | | |
|---|---|---|---|---|
| EPI_ISL_11971465 | A/turkey/South Dakota/22-008239-002-original/2022 | A / H5N1 | North America / United States / South Dakota / Kingsbury | Mary Lea Killian (National Veterinary Services Laboratories - USDA / Diagnostic Virology Laboratory) |
| EPI_ISL_11971466 | A/turkey/South Dakota/22-008242-001-original/2022 | A / H5N1 | North America / United States / South Dakota / Hanson | Mary Lea Killian (National Veterinary Services Laboratories - USDA / Diagnostic Virology Laboratory) |
| EPI_ISL_11971467 | A/turkey/South Dakota/22-008242-002-original/2022 | A / H5N1 | North America / United States / South Dakota / Hanson | Mary Lea Killian (National Veterinary Services Laboratories - USDA / Diagnostic Virology Laboratory) |
| EPI_ISL_11971468 | A/chicken/Kansas/22-008259-001-original/2022 | A / H5N1 | North America / United States / Kansas / Dickinson | Mary Lea Killian (National Veterinary Services Laboratories - USDA / Diagnostic Virology Laboratory) |
| EPI_ISL_11971469 | A/chicken/Kansas/22-008259-002-original/2022 | A / H5N1 | North America / United States / Kansas / Dickinson | Mary Lea Killian (National Veterinary Services Laboratories - USDA / Diagnostic Virology Laboratory) |
| EPI_ISL_11971470 | A/chicken/Maine/22-008366-001-original/2022 | A / H5N1 | North America / United States / Maine / Knox | Mary Lea Killian (National Veterinary Services Laboratories - USDA / Diagnostic Virology Laboratory) |
| EPI_ISL_11971471 | A/chicken/Maine/22-008367-001-original/2022 | A / H5N1 | North America / United States / Maine / York | Mary Lea Killian (National Veterinary Services Laboratories - USDA / Diagnostic Virology Laboratory) |
| EPI_ISL_11971472 | A/turkey/Maine/22-008367-002-original/2022 | A / H5N1 | North America / United States / Maine / York | Mary Lea Killian (National Veterinary Services Laboratories - USDA / Diagnostic Virology Laboratory) |
| EPI_ISL_11971475 | A/chicken/Iowa/22-008373-001-original/2022 | A / H5N1 | North America / United States / Iowa / Warren | Mary Lea Killian (National Veterinary Services Laboratories - USDA / Diagnostic Virology Laboratory) |
| EPI_ISL_11971476 | A/chicken/Iowa/22-008373-002-original/2022 | A / H5N1 | North America / United States / Iowa / Warren | Mary Lea Killian (National Veterinary Services Laboratories - USDA / Diagnostic Virology Laboratory) |
| EPI_ISL_11971477 | A/chicken/Nebraska/22-008467-001-original/2022 | A / H5N1 | North America / United States / Nebraska / Butler | Mary Lea Killian (National Veterinary Services Laboratories - USDA / Diagnostic Virology Laboratory) |
| EPI_ISL_11971478 | A/turkey/South Dakota/22-008479-001-original/2022 | A / H5N1 | North America / United States / South Dakota / Charles Mix | Mary Lea Killian (National Veterinary Services Laboratories - USDA / Diagnostic Virology Laboratory) |
| EPI_ISL_11971479 | A/turkey/South Dakota/22-008479-002-original/2022 | A / H5N1 | North America / United States / South Dakota / Charles Mix | Mary Lea Killian (National Veterinary Services Laboratories - USDA / Diagnostic Virology Laboratory) |
| EPI_ISL_11971480 | A/turkey/South Dakota/22-008483-001-original/2022 | A / H5N1 | North America / United States / South Dakota / Beadle | Mary Lea Killian (National Veterinary Services Laboratories - USDA / Diagnostic Virology Laboratory) |
| EPI_ISL_11971481 | A/turkey/South Dakota/22-008485-001-original/2022 | A / H5N1 | North America / United States / South Dakota / Hanson | Mary Lea Killian (National Veterinary Services Laboratories - USDA / Diagnostic Virology Laboratory) |
| EPI_ISL_11971482 | A/turkey/South Dakota/22-008485-002-original/2022 | A / H5N1 | North America / United States / South Dakota / Hanson | Mary Lea Killian (National Veterinary Services Laboratories - USDA / Diagnostic Virology Laboratory) |
| EPI_ISL_11971483 | A/chicken/Maine/22-008540-001-original/2022 | A / H5N1 | North America / United States / Maine / Cumberland | Mary Lea Killian (National Veterinary Services Laboratories - USDA / Diagnostic Virology Laboratory) |
| EPI_ISL_11971484 | A/turkey/South Dakota/22-008702-001-original/2022 | A / H5N1 | North America / United States / South Dakota / Edmonds | Mary Lea Killian (National Veterinary Services Laboratories - USDA / Diagnostic Virology Laboratory) |
| EPI_ISL_11971485 | A/turkey/South Dakota/22-008702-002-original/2022 | A / H5N1 | North America / United States / South Dakota / Edmonds | Mary Lea Killian (National Veterinary Services Laboratories - USDA / Diagnostic Virology Laboratory) |
| EPI_ISL_11971486 | A/chicken/South Dakota/22-008704-001-original/2022 | A / H5N1 | North America / United States / South Dakota / Kingsbury | Mary Lea Killian (National Veterinary Services Laboratories - USDA / Diagnostic Virology Laboratory) |
| EPI_ISL_11971487 | A/chicken/South Dakota/22-008704-002-original/2022 | A / H5N1 | North America / United States / South Dakota / Kingsbury | Mary Lea Killian (National Veterinary Services Laboratories - USDA / Diagnostic Virology Laboratory) |
| EPI_ISL_11971488 | A/turkey/South Dakota/22-008705-001-original/2022 | A / H5N1 | North America / United States / South Dakota / Beadle | Mary Lea Killian (National Veterinary Services Laboratories - USDA / Diagnostic Virology Laboratory) |
| EPI_ISL_11971489 | A/turkey/South Dakota/22-008705-002-original/2022 | A / H5N1 | North America / United States / South Dakota / Beadle | Mary Lea Killian (National Veterinary Services Laboratories - USDA / Diagnostic Virology Laboratory) |
| EPI_ISL_11971490 | A/pheasant/New York/22-008760-008-original/2022 | A / H5N1 | North America / United States / New York / Suffolk | Mary Lea Killian (National Veterinary Services Laboratories - USDA / Diagnostic Virology Laboratory) |
| EPI_ISL_11971491 | A/mallard/New York/22-008760-007-original/2022 | A / H5N1 | North America / United States / New York / Suffolk | Mary Lea Killian (National Veterinary Services Laboratories - USDA / Diagnostic Virology Laboratory) |
| EPI_ISL_11971492 | A/turkey/Iowa/22-008862-002-original/2022 | A / H5N1 | North America / United States / Iowa / Buena Vista | Mary Lea Killian (National Veterinary Services Laboratories - USDA / Diagnostic Virology Laboratory) |
| EPI_ISL_11971493 | A/turkey/South Dakota/22-008866-001-original/2022 | A / H5N1 | North America / United States / South Dakota / Jerauld | Mary Lea Killian (National Veterinary Services Laboratories - USDA / Diagnostic Virology Laboratory) |
| EPI_ISL_11971494 | A/turkey/South Dakota/22-008866-002-original/2022 | A / H5N1 | North America / United States / South Dakota / Jerauld | Mary Lea Killian (National Veterinary Services Laboratories - USDA / Diagnostic Virology Laboratory) |
| EPI_ISL_11971495 | A/domestic duck/Michigan/22-008890-001-original/2022 | A / H5N1 | North America / United States / Michigan / Macomb | Mary Lea Killian (National Veterinary Services Laboratories - USDA / Diagnostic Virology Laboratory) |
| EPI_ISL_11971496 | A/domestic goose/Michigan/22-008890-004-original/2022 | A / H5N1 | North America / United States / Michigan / Macomb | Mary Lea Killian (National Veterinary Services Laboratories - USDA / Diagnostic Virology Laboratory) |
| EPI_ISL_11971497 | A/chicken/Michigan/22-008890-006-original/2022 | A / H5N1 | North America / United States / Michigan / Macomb | Mary Lea Killian (National Veterinary Services Laboratories - USDA / Diagnostic Virology Laboratory) |
| EPI_ISL_11971498 | A/turkey/Michigan/22-008890-007-original/2022 | A / H5N1 | North America / United States / Michigan / Macomb | Mary Lea Killian (National Veterinary Services Laboratories - USDA / Diagnostic Virology Laboratory) |
| EPI_ISL_11971499 | A/chicken/Nebraska/22-009006-001-original/2022 | A / H5N1 | North America / United States / Nebraska / Butler | Mary Lea Killian (National Veterinary Services Laboratories - USDA / Diagnostic Virology Laboratory) |
| EPI_ISL_11971500 | A/turkey/South Dakota/22-009023-001-original/2022 | A / H5N1 | North America / United States / South Dakota / Hutchison | Mary Lea Killian (National Veterinary Services Laboratories - USDA / Diagnostic Virology Laboratory) |
| EPI_ISL_11971501 | A/turkey/South Dakota/22-009023-002-original/2022 | A / H5N1 | North America / United States / South Dakota / Hutchison | Mary Lea Killian (National Veterinary Services Laboratories - USDA / Diagnostic Virology Laboratory) |
| EPI_ISL_11971502 | A/pheasant/New York/22-009066-001-original/2022 | A / H5N1 | North America / United States / New York / Suffolk | Mary Lea Killian (National Veterinary Services Laboratories - USDA / Diagnostic Virology Laboratory) |
| EPI_ISL_12028893 | A/turkey/Italy/21VIR11507/2021 | A / H5N1 | Europe / Italy | Giacomo Barbierato (Istituto Zooprofilattico Sperimentale delle Venezie / Ricerca e innovazione) |
| EPI_ISL_12045319 | A/chicken/Botswana/2163-B/2021 | A / H5N1 | Africa / Botswana | Alex Byrne (Animal and Plant Health Agency (APHA) / Virology Department) |
| EPI_ISL_12045320 | A/chicken/Botswana/2248-A/2021 | A / H5N1 | Africa / Botswana | Alex Byrne (Animal and Plant Health Agency (APHA) / Virology Department) |
| EPI_ISL_12045918 | A/Dove/Botswana/2097/2021 | A / H5N1 | Africa / Botswana | Alex Byrne (Animal and Plant Health Agency (APHA) / Virology Department) |
| EPI_ISL_12045919 | A/Dove/Botswana/2334/2021 | A / H5N1 | Africa / Botswana | Alex Byrne (Animal and Plant Health Agency (APHA) / Virology Department) |
| EPI_ISL_12045920 | A/fish_eagle/Botswana/1338/2021 | A / H5N1 | Africa / Botswana | Alex Byrne (Animal and Plant Health Agency (APHA) / Virology Department) |
| EPI_ISL_1205485 | A/red knot/Germany-SH/AI03419/2020 | A / H5N3 | Europe / Germany / Schleswig-Holstein / Nordfriesland | Jacqueline King (Friedrich-Loeffler-Institut) |
| EPI_ISL_1205487 | A/red knot/Germany-SH/AI03421/2020 | A / H5N3 | Europe / Germany / Schleswig-Holstein / Nordfriesland | Jacqueline King (Friedrich-Loeffler-Institut) |
| EPI_ISL_1205489 | A/red knot/Germany-SH/AI03424/2020 | A / H5N3 | Europe / Germany / Schleswig-Holstein / Nordfriesland | Jacqueline King (Friedrich-Loeffler-Institut) |
| EPI_ISL_1206285 | A/red knot/Germany-NI/AI00394/2020 | A / H5N3 | Europe / Germany / Lower Saxony / Aurich | Jacqueline King (Friedrich-Loeffler-Institut) |
| EPI_ISL_12066188 | A/Fox/Netherlands/EMC1/2022 | A / H5N1 | Europe / Netherlands / Huissen | Oanh Vuong-Zoet (Erasmus Medical Center / Department of Virology) |
| EPI_ISL_12069288 | A/Fox/Netherlands/EMC2/2022 | A / H5N1 | Europe / Netherlands / Oirschot | Oanh Vuong-Zoet (Erasmus Medical Center / Department of Virology) |
| EPI_ISL_12069289 | A/Fox/Netherlands/EMC3/2022 | A / H5N1 | Europe / Netherlands / Almere | Oanh Vuong-Zoet (Erasmus Medical Center / Department of Virology) |
| EPI_ISL_12069978 | A/greylag goose/Netherlands/22006190-002/2022 | A / H5N1 | Europe / Netherlands / Provincie Gelderland | Rene Heutink (Wageningen Bioveterinary Research) |
| EPI_ISL_12069979 | A/black-backed gull/Netherlands/22006192-001/2022 | A / H5N1 | Europe / Netherlands / Provincie Gelderland | Rene Heutink (Wageningen Bioveterinary Research) |
| EPI_ISL_12140437 | A/duck/Czech Republic/913/2022 | A / H5N1 | Europe / Czech Republic / Jihocesky Kraj / Okres Strakonice / Kliovice; ZipCode:38601 | Alexander Nagy (State Veterinary Institute Prague) |
| EPI_ISL_12150657 | A/goose/Czech Republic/913/2022 | A / H5N1 | Europe / Czech Republic / Jihocesky Kraj / Okres Strakonice / Kliovice; ZipCode:38601 | Alexander Nagy (State Veterinary Institute Prague) |
| EPI_ISL_12150658 | A/chicken/Czech Republic/913/2022 | A / H5N1 | Europe / Czech Republic / Jihocesky Kraj / Okres Strakonice / Kliovice; ZipCode:38601 | Alexander Nagy (State Veterinary Institute Prague) |
| EPI_ISL_12150661 | A/mute swan/Czech Republic/2755/2022 | A / H5N1 | Europe / Czech Republic / Zlinsky Kraj / Okres Zlin / Spytihnev Zip Code: 76364 | Alexander Nagy (State Veterinary Institute Prague) |
| EPI_ISL_12150664 | A/goose/Czech Republic/25322-179/2021 | A / H5N1 | Europe / Czech Republic / Jihocesky Kraj / Okres Ceske Budejovice / Nove Hrady, Zip Code: 3740 | Alexander Nagy (State Veterinary Institute Prague) |
| EPI_ISL_12150666 | A/goose/Czech Republic/25322-205/2021 | A / H5N1 | Europe / Czech Republic / Jihocesky Kraj / Okres Ceske Budejovice / Nove Hrady, Zip Code: 3740 | Alexander Nagy (State Veterinary Institute Prague) |
| EPI_ISL_12174842 | A/Ezo red fox/Hokkaido/1/2022 | A / H5N1 | Asia / Japan / Hokkaido / Sapporo city | Norikazu Isoda (Graduate School of Veterinary Medicine, Hokkaido University / Laboratory of Microbiology) |
| EPI_ISL_12174843 | A/crow/Hokkaido/0103B065/2022 | A / H5N1 | Asia / Japan / Hokkaido / Sapporo city | Norikazu Isoda (Graduate School of Veterinary Medicine, Hokkaido University / Laboratory of Microbiology) |
| EPI_ISL_12176837 | A/pelecanus_crispus/Albania/D383-22_22VIR3125-2/2022 | A / H5N1 | Europe / Albania | Giacomo Barbierato (Istituto Zooprofilattico Sperimentale delle Venezie / Ricerca e innovazione) |
| EPI_ISL_12176838 | A/chicken/Kosovo/278_22VIR3124-1/2021 | A / H5N8 | Europe / Kosovo | Giacomo Barbierato (Istituto Zooprofilattico Sperimentale delle Venezie / Ricerca e innovazione) |
| EPI_ISL_12176839 | A/chicken/Kosovo/279_22VIR3124-2/2021 | A / H5N8 | Europe / Kosovo | Giacomo Barbierato (Istituto Zooprofilattico Sperimentale delle Venezie / Ricerca e innovazione) |
| EPI_ISL_12176840 | A/chicken/Kosovo/280_22VIR3124-3/2021 | A / H5N8 | Europe / Kosovo | Giacomo Barbierato (Istituto Zooprofilattico Sperimentale delle Venezie / Ricerca e innovazione) |
| EPI_ISL_12176841 | A/chicken/Kosovo/283_22VIR3124-4/2021 | A / H5N8 | Europe / Kosovo | Giacomo Barbierato (Istituto Zooprofilattico Sperimentale delle Venezie / Ricerca e innovazione) |
| EPI_ISL_12176843 | A/chicken/Kosovo/284_22VIR3124-5/2021 | A / H5N8 | Europe / Kosovo | Giacomo Barbierato (Istituto Zooprofilattico Sperimentale delle Venezie / Ricerca e innovazione) |
| EPI_ISL_12176844 | A/common_quail/Kosovo/288_22VIR3124-6/2021 | A / H5N8 | Europe / Kosovo | Giacomo Barbierato (Istituto Zooprofilattico Sperimentale delle Venezie / Ricerca e innovazione) |
| EPI_ISL_12176845 | A/chicken/Kosovo/290_22VIR3124-7/2021 | A / H5N8 | Europe / Kosovo | Giacomo Barbierato (Istituto Zooprofilattico Sperimentale delle Venezie / Ricerca e innovazione) |
| EPI_ISL_12176846 | A/chicken/Kosovo/299_22VIR3124-8/2021 | A / H5N8 | Europe / Kosovo | Giacomo Barbierato (Istituto Zooprofilattico Sperimentale delle Venezie / Ricerca e innovazione) |
| EPI_ISL_12176847 | A/chicken/Kosovo/302_22VIR3124-9/2021 | A / H5N8 | Europe / Kosovo | Giacomo Barbierato (Istituto Zooprofilattico Sperimentale delle Venezie / Ricerca e innovazione) |
| EPI_ISL_12176848 | A/chicken/Kosovo/303_22VIR3124-10/2021 | A / H5N8 | Europe / Kosovo | Giacomo Barbierato (Istituto Zooprofilattico Sperimentale delle Venezie / Ricerca e innovazione) |
| EPI_ISL_12176849 | A/common_quail/Kosovo/304-2_22VIR3124-12/2021 | A / H5N8 | Europe / Kosovo | Giacomo Barbierato (Istituto Zooprofilattico Sperimentale delle Venezie / Ricerca e innovazione) |
| EPI_ISL_12176850 | A/chicken/Kosovo/22-2_22VIR3124-13/2022 | A / H5N8 | Europe / Kosovo | Giacomo Barbierato (Istituto Zooprofilattico Sperimentale delle Venezie / Ricerca e innovazione) |

| | | | | |
|---|---|---|---|---|
| EPI_ISL_12176851 | A/chicken/Kosovo/22-8_22VIR3124-14/2022 | A / H5N8 | Europe / Kosovo | Giacomo Barbierato (Istituto Zooprofilattico Sperimentale delle Venezie / Ricerca e innovazione) |
| EPI_ISL_12176852 | A/chicken/Kosovo/22-9_22VIR3124-15/2022 | A / H5N8 | Europe / Kosovo | Giacomo Barbierato (Istituto Zooprofilattico Sperimentale delle Venezie / Ricerca e innovazione) |
| EPI_ISL_12176853 | A/chicken/Kosovo/126_22VIR3124-18/2021 | A / H5N8 | Europe / Kosovo | Giacomo Barbierato (Istituto Zooprofilattico Sperimentale delle Venezie / Ricerca e innovazione) |
| EPI_ISL_12176854 | A/chicken/Kosovo/22-50_22VIR3124-19/2022 | A / H5N8 | Europe / Kosovo | Giacomo Barbierato (Istituto Zooprofilattico Sperimentale delle Venezie / Ricerca e innovazione) |
| EPI_ISL_12176855 | A/chicken/Kosovo/22-59_22VIR3124-20/2022 | A / H5N8 | Europe / Kosovo | Giacomo Barbierato (Istituto Zooprofilattico Sperimentale delle Venezie / Ricerca e innovazione) |
| EPI_ISL_12176931 | A/turkey/Kosovo/13-2_22VIR3124-31/2022 | A / H5N8 | Europe / Kosovo | Giacomo Barbierato (Istituto Zooprofilattico Sperimentale delle Venezie / Ricerca e innovazione) |
| EPI_ISL_12176932 | A/chicken/Albania/D381-22_22VIR3125-1/2022 | A / H5N8 | Europe / Albania | Giacomo Barbierato (Istituto Zooprofilattico Sperimentale delle Venezie / Ricerca e innovazione) |
| EPI_ISL_12177629 | A/black-backed gull/Netherlands/22006711-001/2022 | A / H5N1 | Europe / Netherlands / Provincie Gelderland | Rene Heutink (Wageningen Bioveterinary Research) |
| EPI_ISL_1219038 | A/Chicken/Sweden/SVA210226SZ0168/KN048380-IP9/2021 | A / H5N5 | Europe / Sweden / Skane Lan / Lunds Kommun | Siamak Zohari (National Veterinary Institute) |
| EPI_ISL_1220094 | A/tundra_bean_goose/Poland/MB132/2020 | A / H5N8 | Europe / Poland / West Pomeranian Voivodeship | Edyta ?wi?to? (National Veterinary Research Institut Poland, PIWet-PIB) |
| EPI_ISL_12210335 | A/Falco_peregrinus/Belgium/4055_0002/2022 | A / H5N1 | Europe / Belgium / Provincie Antwerpen / Essen | Steven Van Borm (Sciensano, Department of Animal Infectious Diseases / Animal Infectious Diseases) |
| EPI_ISL_12215416 | A/greylag goose/Netherlands/22005844-002/2022 | A / H5N1 | Europe / Netherlands / South Holland | Rene Heutink (Wageningen Bioveterinary Research) |
| EPI_ISL_12223688 | A/grey heron/Czech Republic/25338-1/2021 | A / H5N1 | Europe / Czech Republic / Liberecky Kraj / Okres Ceska Lipa / Zahradky, GPS: 50.6372833N, 14.5( | Alexander Nagy (State Veterinary Institute Prague) |
| EPI_ISL_12223734 | A/grey heron/Czech Republic/25338-2/2021 | A / H5N1 | Europe / Czech Republic / Liberecky Kraj / Okres Ceska Lipa / Zahradky, GPS: 50.6372833N, 14.5 | Alexander Nagy (State Veterinary Institute Prague) |
| EPI_ISL_1224942 | A/peacock/Netherlands/21022591-002/2021 | A / H5N8 | Europe / Netherlands / Provincie Noord-Holland / Gemeente Zaanstad | Rene Heutink (Wageningen Bioveterinary Research) |
| EPI_ISL_1224946 | A/barnacle goose/Netherlands/21022611-001/2021 | A / H5N1 | Europe / Netherlands / Provincie Friesland / Paesens | Rene Heutink (Wageningen Bioveterinary Research) |
| EPI_ISL_1224949 | A/mute swan/Netherlands/21022898-002/2021 | A / H5N8 | Europe / Netherlands / South Holland / Gemeente Hellevoetsluis | Rene Heutink (Wageningen Bioveterinary Research) |
| EPI_ISL_1224989 | A/common buzzard/Netherlands/21022834-002/2021 | A / H5N1 | Europe / Netherlands / Provincie Friesland / Gemeente Weststellingwerf | Rene Heutink (Wageningen Bioveterinary Research) |
| EPI_ISL_1225077 | A/european herring gull/Netherlands/21023937-002/2021 | A / H5N4 | Europe / Netherlands / South Holland / Midden-Delfland | Rene Heutink (Wageningen Bioveterinary Research) |
| EPI_ISL_1225079 | A/eurasian curlew/Netherlands/21024069-002/2021 | A / H5N4 | Europe / Netherlands / Provincie Friesland / Gemeente Harlingen | Rene Heutink (Wageningen Bioveterinary Research) |
| EPI_ISL_12324302 | A/duck/Czech Republic/3306-1/2022 | A / H5N1 | Europe / Czech Republic / Stredocesky Kraj / Okres Kolin / Masojey, GPS: 50.0259761N, 14.7778 | Alexander Nagy (State Veterinary Institute Prague) |
| EPI_ISL_12325210 | A/chicken/Czech Republic/3306-2/2022 | A / H5N1 | Europe / Czech Republic / Stredocesky Kraj / Okres Kolin / Masojey, GPS: 50.0259761N, 14.7778 | Alexander Nagy (State Veterinary Institute Prague) |
| EPI_ISL_12325995 | A/goose/Czech Republic/25322-229/2021 | A / H5N1 | Europe / Czech Republic / Jihocesky Kraj / Okres Ceske Budejovice / Nove Hrady, Zip Code: 3740 | Alexander Nagy (State Veterinary Institute Prague) |
| EPI_ISL_1235689 | A/Chicken/Sweden/SVA210301SZ0001/KN049780-IP10/2021 | A / H5N8 | Europe / Sweden / Hallands Lan / Kungsbacka Kommun | Siamak Zohari (National Veterinary Institute) |
| EPI_ISL_1236901 | A/goose/Sweden/SVA210301SZ0005/KN049802-IP11/2021 | A / H5N8 | Europe / Sweden / Skane Lan / Landskrona Kommun | Siamak Zohari (National Veterinary Institute) |
| EPI_ISL_1238195 | A/Chicken/Sweden/SVA210302SZ0564/KN052345-IP12/2021 | A / H5N5 | Europe / Sweden / Skane Lan / Trelleborgs Kommun | Siamak Zohari (National Veterinary Institute) |
| EPI_ISL_1238902 | A/common pheasant/Sweden/SVA210224SZ0005/KN000542-II | A / H5N8 | Europe / Sweden / Skane Lan / Trelleborgs Kommun | Siamak Zohari (National Veterinary Institute) |
| EPI_ISL_1239109 | A/northern goshawk/Sweden/SVA210224SZ0431/KN000626/2 | A / H5N8 | Europe / Sweden / Skane Lan / Kavlinge Kommun | Siamak Zohari (National Veterinary Institute) |
| EPI_ISL_1239110 | A/northern goshawk/Sweden/SVA210224SZ0479/KN000627/2 | A / H5N8 | Europe / Sweden / Blekinge Lan / Karlshamns Kommun | Siamak Zohari (National Veterinary Institute) |
| EPI_ISL_1240677 | A/common buzzard /Sweden/SVA210224SZ0485/KN000628/2( | A / H5N8 | Europe / Sweden / Skane Lan / Kristianstads Kommun | Siamak Zohari (National Veterinary Institute) |
| EPI_ISL_1240717 | A/northern goshawk/Sweden/SVA210225SZ0302/KN000664/2 | A / H5N8 | Europe / Sweden / Kalmar Lan / Kalmar Kommun | Siamak Zohari (National Veterinary Institute) |
| EPI_ISL_1240718 | A/barnacle_goose/Sweden/SVA210225SZ0307/KN000666/202 | A / H5N8 | Europe / Sweden / Skane Lan / Malmo Kommun | Siamak Zohari (National Veterinary Institute) |
| EPI_ISL_1240997 | A/eastern buzzard/Tochigi/090311C/2021 | A / H5N8 | Asia / Japan / Tochigi | Takehiko Saito (National Institute of Animal Health) |
| EPI_ISL_1240998 | A/mallard/Novosibirsk region/3509k/2020 | A / H5N8 | Europe / Russian Federation / Novosibirsk | Takehiko Saito (National Institute of Animal Health) |
| EPI_ISL_1241004 | A/owl/Tochigi/090204T/2021 | A / H5N8 | Asia / Japan / Tochigi | Takehiko Saito (National Institute of Animal Health) |
| EPI_ISL_12436968 | A/duck/Czech Republic/5361/2021 | A / H5N8 | Europe / Czech Republic / Kralovehradecky Kraj / Okres Hradec Kralove / Zabedov | Alexander Nagy (State Veterinary Institute Prague) |
| EPI_ISL_12437699 | A/duck/Czech Republic/5467/2021 | A / H5N8 | Europe / Czech Republic / Kralovehradecky Kraj / Okres Hradec Kralove / Chudonice, GPS: N 50°1 | Alexander Nagy (State Veterinary Institute Prague) |
| EPI_ISL_12437700 | A/duck/Czech Republic/6017/2021 | A / H5N8 | Europe / Czech Republic / Stredocesky Kraj / Okres Nymburk / Slibovice | Alexander Nagy (State Veterinary Institute Prague) |
| EPI_ISL_12471655 | A/pelican/Greece/41-TR-313_22VIR3126-1/2022 | A / H5N1 | Europe / Greece | Giacomo Barbierato (Istituto Zooprofilattico Sperimentale delle Venezie / Ricerca e innovazione) |
| EPI_ISL_12471656 | A/pelican/Greece/69_CL_22VIR3126-10/2022 | A / H5N1 | Europe / Greece | Giacomo Barbierato (Istituto Zooprofilattico Sperimentale delle Venezie / Ricerca e innovazione) |
| EPI_ISL_12471657 | A/pelican/Greece/72_CL_22VIR3126-11/2022 | A / H5N1 | Europe / Greece | Giacomo Barbierato (Istituto Zooprofilattico Sperimentale delle Venezie / Ricerca e innovazione) |
| EPI_ISL_12471658 | A/pelican/Greece/41_AL1_22VIR3126-2/2022 | A / H5N1 | Europe / Greece | Giacomo Barbierato (Istituto Zooprofilattico Sperimentale delle Venezie / Ricerca e innovazione) |
| EPI_ISL_12471659 | A/pelican/Greece/41_AL2_22VIR3126-3/2022 | A / H5N1 | Europe / Greece | Giacomo Barbierato (Istituto Zooprofilattico Sperimentale delle Venezie / Ricerca e innovazione) |
| EPI_ISL_12471660 | A/pelican/Greece/64_KI_22VIR3126-6/2022 | A / H5N1 | Europe / Greece | Giacomo Barbierato (Istituto Zooprofilattico Sperimentale delle Venezie / Ricerca e innovazione) |
| EPI_ISL_12471661 | A/pelican/Greece/64_SP_22VIR3126-7/2022 | A / H5N1 | Europe / Greece | Giacomo Barbierato (Istituto Zooprofilattico Sperimentale delle Venezie / Ricerca e innovazione) |
| EPI_ISL_12471662 | A/pelican/Greece/64_LI_22VIR3126-8/2022 | A / H5N1 | Europe / Greece | Giacomo Barbierato (Istituto Zooprofilattico Sperimentale delle Venezie / Ricerca e innovazione) |
| EPI_ISL_12471663 | A/pelican/Greece/64_TR_22VIR3126-9/2022 | A / H5N1 | Europe / Greece | Giacomo Barbierato (Istituto Zooprofilattico Sperimentale delle Venezie / Ricerca e innovazione) |
| EPI_ISL_12474793 | A/barnacle goose/Netherlands/22007405-004/2022 | A / H5N1 | Europe / Netherlands / Provincie Gelderland | Rene Heutink (Wageningen Bioveterinary Research) |
| EPI_ISL_12474794 | A/greylag goose /Netherlands/22006859-001/2022 | A / H5N1 | Europe / Netherlands / South Holland | Rene Heutink (Wageningen Bioveterinary Research) |
| EPI_ISL_12512879 | A/Black-headed gull/Netherlands/2/2022 | A / H5N1 | Europe / Netherlands / Provincie Noord-Holland / Normerven | Sanne Thewessen (Erasmus Medical Center / Viroscience) |
| EPI_ISL_12514425 | A/Black-headed gull/Netherlands/3/2022 | A / H5N1 | Europe / Netherlands / De Kreupel | Sanne Thewessen (Erasmus Medical Center / Viroscience) |
| EPI_ISL_12514442 | A/Black-headed gull/Netherlands/4/2022 | A / H5N1 | Europe / Netherlands / De Kreupel | Sanne Thewessen (Erasmus Medical Center / Viroscience) |
| EPI_ISL_12514483 | A/Caspian Gull/Netherlands/3/2022 | A / H5N1 | Europe / Netherlands / De Kreupel | Sanne Thewessen (Erasmus Medical Center / Viroscience) |
| EPI_ISL_12514532 | A/Common raven/Netherlands/1/2022 | A / H5N1 | Europe / Netherlands / Putten | Sanne Thewessen (Erasmus Medical Center / Viroscience) |
| EPI_ISL_12514573 | A/Song Thrush/Netherlands/1/2022 | A / H5N1 | Europe / Netherlands / Wageningen | Sanne Thewessen (Erasmus Medical Center / Viroscience) |
| EPI_ISL_12514622 | A/Common raven/Netherlands/2/2022 | A / H5N1 | Europe / Netherlands / Putten | Sanne Thewessen (Erasmus Medical Center / Viroscience) |
| EPI_ISL_12514681 | A/European Herring Gull/Netherlands/3/2022 | A / H5N1 | Europe / Netherlands / De Kreupel | Sanne Thewessen (Erasmus Medical Center / Viroscience) |
| EPI_ISL_1254 | A/Goose/Guangdong/1/96 | A / H5N1 | Asia / China / Guangdong Province | |
| EPI_ISL_1255051 | A/tufted_duck/Poland/MB061/2021(H5N5) | A / H5N5 | Europe / Poland / West Pomeranian Voivodeship | Edyta ?wi?to? (National Veterinary Research Institut Poland, PIWet-PIB) |
| EPI_ISL_12567390 | A/chicken/Maryland/22-008243-001-original/2022 | A / H5N1 | North America / United States / Maryland / Cecil | Mary Lea Killian (National Veterinary Services Laboratories - USDA / Diagnostic Virology Laboratory) |
| EPI_ISL_12567391 | A/chicken/Iowa/22-009180-001-original/2022 | A / H5N1 | North America / United States / Iowa / Franklin | Mary Lea Killian (National Veterinary Services Laboratories - USDA / Diagnostic Virology Laboratory) |
| EPI_ISL_12567392 | A/chicken/Iowa/22-009180-002-original/2022 | A / H5N1 | North America / United States / Iowa / Franklin | Mary Lea Killian (National Veterinary Services Laboratories - USDA / Diagnostic Virology Laboratory) |
| EPI_ISL_12567393 | A/chicken/Minnesota/22-009181-001-original/2022 | A / H5N1 | North America / United States / Minnesota / Mower | Mary Lea Killian (National Veterinary Services Laboratories - USDA / Diagnostic Virology Laboratory) |
| EPI_ISL_12567529 | A/Embden goose/Minnesota/22-009181-003-original/2022 | A / H5N1 | North America / United States / Minnesota / Mower | Mary Lea Killian (National Veterinary Services Laboratories - USDA / Diagnostic Virology Laboratory) |
| EPI_ISL_12567530 | A/turkey/Minnesota/22-009182-001-original/2022 | A / H5N1 | North America / United States / Minnesota / Meeker | Mary Lea Killian (National Veterinary Services Laboratories - USDA / Diagnostic Virology Laboratory) |
| EPI_ISL_12567531 | A/turkey/Minnesota/22-009182-003-original/2022 | A / H5N1 | North America / United States / Minnesota / Meeker | Mary Lea Killian (National Veterinary Services Laboratories - USDA / Diagnostic Virology Laboratory) |
| EPI_ISL_12567532 | A/domestic duck/Nebraska/22-009190-001-original/2022 | A / H5N1 | North America / United States / Nebraska / Holt | Mary Lea Killian (National Veterinary Services Laboratories - USDA / Diagnostic Virology Laboratory) |
| EPI_ISL_12567533 | A/chicken/Nebraska/22-009190-002-original/2022 | A / H5N1 | North America / United States / Nebraska / Holt | Mary Lea Killian (National Veterinary Services Laboratories - USDA / Diagnostic Virology Laboratory) |
| EPI_ISL_12572652 | A/chicken/Anhui/S1740/2022(H5N1) | A / H5N1 | Asia / China / Anhui | Pengfei Cui (Harbin Veterinary Research Institute (CAAS) / Ministry of Agriculture) |
| EPI_ISL_12572653 | A/chicken/Jiangxi/S40653/2021(H5N1) | A / H5N1 | Asia / China / Jiangxi | Pengfei Cui (Harbin Veterinary Research Institute (CAAS) / Ministry of Agriculture) |
| EPI_ISL_12572654 | A/duck/Guangdong/S4518/2021(H5N1) | A / H5N1 | Asia / China / Guangdong | Pengfei Cui (Harbin Veterinary Research Institute (CAAS) / Ministry of Agriculture) |
| EPI_ISL_12572655 | A/duck/Guangdong/S4525/2021(H5N1) | A / H5N1 | Asia / China / Guangdong | Pengfei Cui (Harbin Veterinary Research Institute (CAAS) / Ministry of Agriculture) |

| | | | | |
|---|---|---|---|---|
| EPI_ISL_12572656 | A/duck/Guizhou/S1321/2022(H5N1) | A / H5N1 | Asia / China / Guizhou | Pengfei Cui (Harbin Veterinary Research Institute (CAAS) / Ministry of Agriculture) |
| EPI_ISL_12572657 | A/duck/Hubei/S4465/2021(H5N1) | A / H5N1 | Asia / China / Hubei | Pengfei Cui (Harbin Veterinary Research Institute (CAAS) / Ministry of Agriculture) |
| EPI_ISL_12572658 | A/duck/Hubei/SE128/2022(H5N1) | A / H5N1 | Asia / China / Hubei | Pengfei Cui (Harbin Veterinary Research Institute (CAAS) / Ministry of Agriculture) |
| EPI_ISL_12572659 | A/duck/Hubei/SE220/2022(H5N1) | A / H5N1 | Asia / China / Hubei | Pengfei Cui (Harbin Veterinary Research Institute (CAAS) / Ministry of Agriculture) |
| EPI_ISL_12572660 | A/duck/Jiangxi/S40833/2021(H5N1) | A / H5N1 | Asia / China / Jiangxi | Pengfei Cui (Harbin Veterinary Research Institute (CAAS) / Ministry of Agriculture) |
| EPI_ISL_12572661 | A/goose/Guizhou/S1541/2022(H5N1) | A / H5N1 | Asia / China / Guizhou | Pengfei Cui (Harbin Veterinary Research Institute (CAAS) / Ministry of Agriculture) |
| EPI_ISL_12572662 | A/goose/Hunan/SE284/2022(H5N1) | A / H5N1 | Asia / China / Hunan | Pengfei Cui (Harbin Veterinary Research Institute (CAAS) / Ministry of Agriculture) |
| EPI_ISL_12572663 | A/pigeon/Jiangxi/S40784/2021(H5N1) | A / H5N1 | Asia / China / Jiangxi | Pengfei Cui (Harbin Veterinary Research Institute (CAAS) / Ministry of Agriculture) |
| EPI_ISL_12572664 | A/wild duck/Hebei/SD012/2021(H5N1) | A / H5N1 | Asia / China / Hebei | Pengfei Cui (Harbin Veterinary Research Institute (CAAS) / Ministry of Agriculture) |
| EPI_ISL_1259397 | A/curlew/France/21P003648/2021 | A / H5N3 | Europe / France / Basse-Normandie / Departement de la Manche | Francois-Xavier Briand (ANSES Agence Nationale De Securite Sanitaire De L'alimentation / Laboratoire de Ploufragan-Plouzané) |
| EPI_ISL_1259399 | A/red knot/France/21P003249/2021 | A / H5N3 | Europe / France / Basse-Normandie / Departement de la Manche | Francois-Xavier Briand (ANSES Agence Nationale De Securite Sanitaire De L'alimentation / Laboratoire de Ploufragan-Plouzané) |
| EPI_ISL_12690318 | A/chicken/Missouri/22-009192-002-original/2022 | A / H5N1 | North America / United States / Missouri / Gentry | Mary Lea Killian (National Veterinary Services Laboratories - USDA / Diagnostic Virology Laboratory) |
| EPI_ISL_12690319 | A/duck/Missouri/22-009192-003-original/2022 | A / H5N1 | North America / United States / Missouri / Gentry | Mary Lea Killian (National Veterinary Services Laboratories - USDA / Diagnostic Virology Laboratory) |
| EPI_ISL_12690320 | A/chicken/South Dakota/22-009194-002-original/2022 | A / H5N1 | North America / United States / South Dakota / Bon Homme | Mary Lea Killian (National Veterinary Services Laboratories - USDA / Diagnostic Virology Laboratory) |
| EPI_ISL_12690321 | A/guinea fowl/South Dakota/22-009194-003-original/2022 | A / H5N1 | North America / United States / South Dakota / Bon Homme | Mary Lea Killian (National Veterinary Services Laboratories - USDA / Diagnostic Virology Laboratory) |
| EPI_ISL_12690322 | A/turkey/Minnesota/22-009195-002-original/2022 | A / H5N1 | North America / United States / Minnesota / Stearns | Mary Lea Killian (National Veterinary Services Laboratories - USDA / Diagnostic Virology Laboratory) |
| EPI_ISL_12690323 | A/turkey/Minnesota/22-009195-003-original/2022 | A / H5N1 | North America / United States / Minnesota / Stearns | Mary Lea Killian (National Veterinary Services Laboratories - USDA / Diagnostic Virology Laboratory) |
| EPI_ISL_12690324 | A/turkey/South Dakota/22-009196-001-original/2022 | A / H5N1 | North America / United States / South Dakota / Hutchison | Mary Lea Killian (National Veterinary Services Laboratories - USDA / Diagnostic Virology Laboratory) |
| EPI_ISL_12690325 | A/turkey/South Dakota/22-009196-002-original/2022 | A / H5N1 | North America / United States / South Dakota / Hutchison | Mary Lea Killian (National Veterinary Services Laboratories - USDA / Diagnostic Virology Laboratory) |
| EPI_ISL_12690568 | A/turkey/South Dakota/22-009198-001-original/2022 | A / H5N1 | North America / United States / South Dakota / Clark | Mary Lea Killian (National Veterinary Services Laboratories - USDA / Diagnostic Virology Laboratory) |
| EPI_ISL_12690569 | A/turkey/Iowa/22-009227-001-original/2022 | A / H5N1 | North America / United States / Iowa / Buena Vista | Mary Lea Killian (National Veterinary Services Laboratories - USDA / Diagnostic Virology Laboratory) |
| EPI_ISL_12690570 | A/turkey/Iowa/22-009286-001-original/2022 | A / H5N1 | North America / United States / Iowa / Hamilton | Mary Lea Killian (National Veterinary Services Laboratories - USDA / Diagnostic Virology Laboratory) |
| EPI_ISL_12690571 | A/turkey/Iowa/22-009286-002-original/2022 | A / H5N1 | North America / United States / Iowa / Hamilton | Mary Lea Killian (National Veterinary Services Laboratories - USDA / Diagnostic Virology Laboratory) |
| EPI_ISL_12690589 | A/chicken/Iowa/22-009287-001-original/2022 | A / H5N1 | North America / United States / Iowa / Guthrie | Mary Lea Killian (National Veterinary Services Laboratories - USDA / Diagnostic Virology Laboratory) |
| EPI_ISL_12690590 | A/chicken/Iowa/22-009287-002-original/2022 | A / H5N1 | North America / United States / Iowa / Guthrie | Mary Lea Killian (National Veterinary Services Laboratories - USDA / Diagnostic Virology Laboratory) |
| EPI_ISL_12690591 | A/turkey/Minnesota/22-009313-001-original/2022 | A / H5N1 | North America / United States / Minnesota / Kandiyohi | Mary Lea Killian (National Veterinary Services Laboratories - USDA / Diagnostic Virology Laboratory) |
| EPI_ISL_12690903 | A/turkey/Minnesota/22-009314-001-original/2022 | A / H5N1 | North America / United States / Minnesota / Lac Qui Parle | Mary Lea Killian (National Veterinary Services Laboratories - USDA / Diagnostic Virology Laboratory) |
| EPI_ISL_12690904 | A/turkey/Minnesota/22-009314-002-original/2022 | A / H5N1 | North America / United States / Minnesota / Lac Qui Parle | Mary Lea Killian (National Veterinary Services Laboratories - USDA / Diagnostic Virology Laboratory) |
| EPI_ISL_12690905 | A/chicken/New York/22-009324-002-original/2022 | A / H5N1 | North America / United States / New York / Monroe | Mary Lea Killian (National Veterinary Services Laboratories - USDA / Diagnostic Virology Laboratory) |
| EPI_ISL_12690906 | A/domestic duck/New York/22-009324-004-original/2022 | A / H5N1 | North America / United States / New York / Monroe | Mary Lea Killian (National Veterinary Services Laboratories - USDA / Diagnostic Virology Laboratory) |
| EPI_ISL_12690907 | A/American Buff goose/New York/22-009324-006-original/202 | A / H5N1 | North America / United States / New York / Monroe | Mary Lea Killian (National Veterinary Services Laboratories - USDA / Diagnostic Virology Laboratory) |
| EPI_ISL_12690908 | A/quail/New York/22-009324-009-original/2022 | A / H5N1 | North America / United States / New York / Monroe | Mary Lea Killian (National Veterinary Services Laboratories - USDA / Diagnostic Virology Laboratory) |
| EPI_ISL_12690909 | A/chicken/Wyoming/22-009326-002-original/2022 | A / H5N1 | North America / United States / Wyoming / Johnson | Mary Lea Killian (National Veterinary Services Laboratories - USDA / Diagnostic Virology Laboratory) |
| EPI_ISL_12690910 | A/turkey/South Dakota/22-009327-001-original/2022 | A / H5N1 | North America / United States / South Dakota / Spink | Mary Lea Killian (National Veterinary Services Laboratories - USDA / Diagnostic Virology Laboratory) |
| EPI_ISL_12690911 | A/turkey/South Dakota/22-009327-002-original/2022 | A / H5N1 | North America / United States / South Dakota / Spink | Mary Lea Killian (National Veterinary Services Laboratories - USDA / Diagnostic Virology Laboratory) |
| EPI_ISL_12690912 | A/turkey/Minnesota/22-009313-002-original/2022 | A / H5N1 | North America / United States / Minnesota / Kandiyohi | Mary Lea Killian (National Veterinary Services Laboratories - USDA / Diagnostic Virology Laboratory) |
| EPI_ISL_12691028 | A/turkey/South Dakota/22-009329-001-original/2022 | A / H5N1 | North America / United States / South Dakota / Bon Homme | Mary Lea Killian (National Veterinary Services Laboratories - USDA / Diagnostic Virology Laboratory) |
| EPI_ISL_12691029 | A/turkey/South Dakota/22-009329-002-original/2022 | A / H5N1 | North America / United States / South Dakota / Bon Homme | Mary Lea Killian (National Veterinary Services Laboratories - USDA / Diagnostic Virology Laboratory) |
| EPI_ISL_12691030 | A/turkey/South Dakota/22-009330-001-original/2022 | A / H5N1 | North America / United States / South Dakota / Edmunds | Mary Lea Killian (National Veterinary Services Laboratories - USDA / Diagnostic Virology Laboratory) |
| EPI_ISL_12691031 | A/turkey/South Dakota/22-009330-002-original/2022 | A / H5N1 | North America / United States / South Dakota / Edmunds | Mary Lea Killian (National Veterinary Services Laboratories - USDA / Diagnostic Virology Laboratory) |
| EPI_ISL_12691032 | A/turkey/North Carolina/22-009332-001-original/2022 | A / H5N1 | North America / United States / North Carolina / Johnston | Mary Lea Killian (National Veterinary Services Laboratories - USDA / Diagnostic Virology Laboratory) |
| EPI_ISL_12691033 | A/turkey/North Carolina/22-009332-002-original/2022 | A / H5N1 | North America / United States / North Carolina / Johnston | Mary Lea Killian (National Veterinary Services Laboratories - USDA / Diagnostic Virology Laboratory) |
| EPI_ISL_12691034 | A/domestic goose/North Dakota/22-009337-001-original/2022 | A / H5N1 | North America / United States / North Dakota / Kidder | Mary Lea Killian (National Veterinary Services Laboratories - USDA / Diagnostic Virology Laboratory) |
| EPI_ISL_12691035 | A/chicken/North Dakota/22-009337-002-original/2022 | A / H5N1 | North America / United States / North Dakota / Kidder | Mary Lea Killian (National Veterinary Services Laboratories - USDA / Diagnostic Virology Laboratory) |
| EPI_ISL_12693855 | A/chicken/Iowa/22-009819-001-original/2022 | A / H5N1 | North America / United States / Iowa / Osceola | Mary Lea Killian (National Veterinary Services Laboratories - USDA / Diagnostic Virology Laboratory) |
| EPI_ISL_1273409 | A/chicken/Chiba/F3T/2021 | A / H5N8 | Asia / Japan / Chiba | Takehiko Saito (National Institute of Animal Health) |
| EPI_ISL_1273410 | A/chicken/Fukuoka/C1/2020 | A / H5N8 | Asia / Japan / Fukuoka | Takehiko Saito (National Institute of Animal Health) |
| EPI_ISL_1273411 | A/chicken/Fukuoka/C2/2020 | A / H5N8 | Asia / Japan / Fukuoka | Takehiko Saito (National Institute of Animal Health) |
| EPI_ISL_1273412 | A/chicken/Hyogo/3T/2020 | A / H5N8 | Asia / Japan / Hyogo | Takehiko Saito (National Institute of Animal Health) |
| EPI_ISL_1273413 | A/chicken/Hyogo/4T/2020 | A / H5N8 | Asia / Japan / Hyogo | Takehiko Saito (National Institute of Animal Health) |
| EPI_ISL_1273414 | A/chicken/Ibaraki/1T/2021 | A / H5N8 | Asia / Japan / Ibaraki | Takehiko Saito (National Institute of Animal Health) |
| EPI_ISL_1273415 | A/chicken/Ibaraki/2T/2021 | A / H5N8 | Asia / Japan / Ibaraki | Takehiko Saito (National Institute of Animal Health) |
| EPI_ISL_1273416 | A/chicken/Kagawa/10T/2020 | A / H5N8 | Asia / Japan / Kagawa | Takehiko Saito (National Institute of Animal Health) |
| EPI_ISL_1273417 | A/chicken/Kagawa/2T/2020 | A / H5N8 | Asia / Japan / Kagawa | Takehiko Saito (National Institute of Animal Health) |
| EPI_ISL_1273418 | A/chicken/Kagawa/6T/2020 | A / H5N8 | Asia / Japan / Kagawa | Takehiko Saito (National Institute of Animal Health) |
| EPI_ISL_1273419 | A/chicken/Kagawa/7T/2020 | A / H5N8 | Asia / Japan / Kagawa | Takehiko Saito (National Institute of Animal Health) |
| EPI_ISL_1273420 | A/chicken/Kagawa/B7T/2020 | A / H5N8 | Asia / Japan / Kagawa | Takehiko Saito (National Institute of Animal Health) |
| EPI_ISL_1273421 | A/chicken/Kagawa/B9T/2020 | A / H5N8 | Asia / Japan / Kagawa | Takehiko Saito (National Institute of Animal Health) |
| EPI_ISL_1273422 | A/chicken/Kagawa/C1T/2020 | A / H5N8 | Asia / Japan / Kagawa | Takehiko Saito (National Institute of Animal Health) |
| EPI_ISL_1273423 | A/chicken/Kagawa/C2T/2020 | A / H5N8 | Asia / Japan / Kagawa | Takehiko Saito (National Institute of Animal Health) |
| EPI_ISL_1273424 | A/chicken/Kagawa/D1T/2020 | A / H5N8 | Asia / Japan / Kagawa | Takehiko Saito (National Institute of Animal Health) |
| EPI_ISL_1273425 | A/chicken/Kagawa/D2T/2020 | A / H5N8 | Asia / Japan / Kagawa | Takehiko Saito (National Institute of Animal Health) |
| EPI_ISL_1273426 | A/chicken/Kagawa/D3T/2020 | A / H5N8 | Asia / Japan / Kagawa | Takehiko Saito (National Institute of Animal Health) |
| EPI_ISL_1273427 | A/chicken/Kagawa/E4T/2020 | A / H5N8 | Asia / Japan / Kagawa | Takehiko Saito (National Institute of Animal Health) |
| EPI_ISL_1273428 | A/chicken/Kagawa/E5T/2020 | A / H5N8 | Asia / Japan / Kagawa | Takehiko Saito (National Institute of Animal Health) |
| EPI_ISL_1273429 | A/chicken/Kagawa/F3T/2020 | A / H5N8 | Asia / Japan / Kagawa | Takehiko Saito (National Institute of Animal Health) |
| EPI_ISL_1273430 | A/chicken/Kagawa/F4T/2020 | A / H5N8 | Asia / Japan / Kagawa | Takehiko Saito (National Institute of Animal Health) |
| EPI_ISL_1273431 | A/chicken/Kagawa/F5T/2020 | A / H5N8 | Asia / Japan / Kagawa | Takehiko Saito (National Institute of Animal Health) |
| EPI_ISL_1273432 | A/chicken/Kagawa/G10T/2020 | A / H5N8 | Asia / Japan / Kagawa | Takehiko Saito (National Institute of Animal Health) |
| EPI_ISL_1273433 | A/chicken/Kagawa/G12T/2020 | A / H5N8 | Asia / Japan / Kagawa | Takehiko Saito (National Institute of Animal Health) |
| EPI_ISL_1273434 | A/chicken/Kagawa/H1T/2020 | A / H5N8 | Asia / Japan / Kagawa | Takehiko Saito (National Institute of Animal Health) |
| EPI_ISL_1273435 | A/chicken/Kagawa/H4T/2020 | A / H5N8 | Asia / Japan / Kagawa | Takehiko Saito (National Institute of Animal Health) |

| | | | | |
|---|---|---|---|---|
| EPI_ISL_1273436 | A/chicken/Kagawa/H8T/2020 | A / H5N8 | Asia / Japan / Kagawa | Takehiko Saito (National Institute of Animal Health) |
| EPI_ISL_1273437 | A/chicken/Tochigi/2T/2021 | A / H5N8 | Asia / Japan / Tochigi | Takehiko Saito (National Institute of Animal Health) |
| EPI_ISL_1273438 | A/chicken/Tochigi/3T/2021 | A / H5N8 | Asia / Japan / Tochigi | Takehiko Saito (National Institute of Animal Health) |
| EPI_ISL_1273439 | A/chicken/Tochigi/4T/2021 | A / H5N8 | Asia / Japan / Tochigi | Takehiko Saito (National Institute of Animal Health) |
| EPI_ISL_1273440 | A/chicken/Tochigi/5T/2021 | A / H5N8 | Asia / Japan / Tochigi | Takehiko Saito (National Institute of Animal Health) |
| EPI_ISL_1273441 | A/eastern buzzard/Tochigi/090311T/2021 | A / H5N8 | Asia / Japan / Tochigi | Takehiko Saito (National Institute of Animal Health) |
| EPI_ISL_12749687 | A/turkey/Israel/537/2021 | A / H5N1 | Asia / Israel / Northern District / Nahalal | Irina Shkoda (Kimron Veterinary Institute) |
| EPI_ISL_12754531 | A/laying_hen/Romania/11343_22VIR4106-1/2022 | A / H5N1 | Europe / Romania / Judetul Giurgiu / Branistea | Bianca Zecchin (Istituto Zooprofilattico Sperimentale Delle Venezie) |
| EPI_ISL_12754532 | A/pelican/Romania/11335_22VIR4106-2/2022 | A / H5N1 | Europe / Romania / Judetul Tulcea / Sinoe Lake | Bianca Zecchin (Istituto Zooprofilattico Sperimentale Delle Venezie) |
| EPI_ISL_12754533 | A/pelican/Romania/11334_22VIR4106-3/2022 | A / H5N1 | Europe / Romania / Judetul Tulcea / Sinoe Lake | Bianca Zecchin (Istituto Zooprofilattico Sperimentale Delle Venezie) |
| EPI_ISL_12754534 | A/laying_hen/Romania/11562_22VIR4106-4/2022 | A / H5N1 | Europe / Romania / Judetul Giurgiu / Branistea | Bianca Zecchin (Istituto Zooprofilattico Sperimentale Delle Venezie) |
| EPI_ISL_12754535 | A/sea_eagle/Norway/2022-07-196_22VIR3866-1/2022 | A / H5N5 | Europe / Norway / Tromso | Bianca Zecchin (Istituto Zooprofilattico Sperimentale Delle Venezie) |
| EPI_ISL_12754536 | A/sea_eagle/Norway/2022-07-198_22VIR3866-2/2022 | A / H5N5 | Europe / Norway / Hitra | Bianca Zecchin (Istituto Zooprofilattico Sperimentale Delle Venezie) |
| EPI_ISL_1279262 | A/mute swan/Croatia/14/2021 | A / H5N8 | Europe / Croatia / Vukovarsko-Srijemska Zupanija / River Bosut, Podgra?e | Vladimir Savi? (Croatian Veterinary Institute / Poultry Centre) |
| EPI_ISL_1279936 | A/mute swan/Croatia/19/2021 | A / H5N8 | Europe / Croatia / Vukovarsko-Srijemska Zupanija / River Bosut, Podgra?e | Vladimir Savi? (Croatian Veterinary Institute / Poultry Centre) |
| EPI_ISL_12799972 | A/Colorado/18/2022 | A / H5N1 | North America / United States / Colorado | Juliana DaSilva (Centers for Disease Control and Prevention / WHO Collaborating Center for Surveillance, Epidemiology and Control of I |
| EPI_ISL_12866390 | A/chicken/Egypt/Army/1201/2022 | A / H5N8 | Africa / Egypt / Giza | Mohamed Gomaa Seadawy (Egyptian Armed Forces (EAF) / Biological Prevention Department) |
| EPI_ISL_1290863 | A/duck/Laos/XBY118/2015 | A / H5N6 | Asia / Lao, People's Democratic Republic | Yunho Jang (Centers for Disease Control and Prevention / WHO Collaborating Center for Surveillance, Epidemiology and Control of Influ |
| EPI_ISL_1290892 | A/duck/Laos/2310/2019 | A / H5N6 | Asia / Lao, People's Democratic Republic | Yunho Jang (Centers for Disease Control and Prevention / WHO Collaborating Center for Surveillance, Epidemiology and Control of Influ |
| EPI_ISL_12917757 | A/turkey/Israel/564/2021 | A / H5N1 | Asia / Israel / Northern District / Ma'ayan Tzvi | Irina Shkoda (Kimron Veterinary Institute) |
| EPI_ISL_1295638 | A/mute_swan/Norway/FU48/2021 | A / H5N8 | Europe / Norway | Alex Byrne (Animal and Plant Health Agency (APHA) / Virology Department) |
| EPI_ISL_1295639 | A/turkey/Norway/FU496/2020 | A / H5N8 | Europe / Norway | Alex Byrne (Animal and Plant Health Agency (APHA) / Virology Department) |
| EPI_ISL_1296295 | A/Chicken/Sweden/SVA210308SZ0214/KN057720-IP13/2021 | A / H5N8 | Europe / Sweden / Ostergotlands Lan / Mjolby Kommun | Siamak Zohari (National Veterinary Institute) |
| EPI_ISL_12968592 | A/Fancy chicken/NL/FAV-0035/2021 | A / H5N1 | North America / Canada / Newfoundland and Labrador | Tamiko Hisanaga (Canadian Food Inspection Agency) |
| EPI_ISL_12968593 | A/domestic goose/NL/FAV-0035-17/2021 | A / H5N1 | North America / Canada / Manitoba | Tamiko Hisanaga (Canadian Food Inspection Agency) |
| EPI_ISL_12968817 | A/emu/NL/FAV-0035-12/2021 | A / H5N1 | North America / Canada / Newfoundland and Labrador | Tamiko Hisanaga (Canadian Food Inspection Agency) |
| EPI_ISL_12968823 | A/chicken/NL/FAV-0033/2021 | A / H5N1 | North America / Canada / Newfoundland and Labrador | Tamiko Hisanaga (Canadian Food Inspection Agency) |
| EPI_ISL_12980521 | A/Chicken/Sweden/SVA210201SZ0080/FB000693-IP4/2021 | A / H5N5 | Europe / Sweden / Kalmar Lan | Siamak Zohari (National Veterinary Institute) |
| EPI_ISL_129982 | A/duck/Eastern China/1111/2011 | A / H5N2 | Asia / China | |
| EPI_ISL_129983 | A/goose/Eastern China/1112/2011 | A / H5N2 | Asia / China | |
| EPI_ISL_13009618 | A/Sebastopol goose/Massachusetts/22-009371-001-original/2( | A / H5N1 | North America / United States / Massachusetts / Berkshire County | Mary Lea Killian (National Veterinary Services Laboratories - USDA / Diagnostic Virology Laboratory) |
| EPI_ISL_13009620 | A/chicken/Maine/22-009412-001-original/2022 | A / H5N1 | North America / United States / Maine / Knox County | Mary Lea Killian (National Veterinary Services Laboratories - USDA / Diagnostic Virology Laboratory) |
| EPI_ISL_13009634 | A/guinea fowl/Maine/22-009412-002-original/2022 | A / H5N1 | North America / United States / Maine / Knox County | Mary Lea Killian (National Veterinary Services Laboratories - USDA / Diagnostic Virology Laboratory) |
| EPI_ISL_13009636 | A/chicken/Ohio/22-009419-001-original/2022 | A / H5N1 | North America / United States / Ohio / Franklin County | Mary Lea Killian (National Veterinary Services Laboratories - USDA / Diagnostic Virology Laboratory) |
| EPI_ISL_13009652 | A/turkey/South Dakota/22-009534-001-original/2022 | A / H5N1 | North America / United States / South Dakota / Brule County | Mary Lea Killian (National Veterinary Services Laboratories - USDA / Diagnostic Virology Laboratory) |
| EPI_ISL_13009658 | A/turkey/Iowa/22-009550-001-original/2022 | A / H5N1 | North America / United States / Iowa / Buena Vista County | Mary Lea Killian (National Veterinary Services Laboratories - USDA / Diagnostic Virology Laboratory) |
| EPI_ISL_13009673 | A/turkey/North Carolina/22-009583-001-original/2022 | A / H5N1 | North America / United States / North Carolina / Johnston County | Mary Lea Killian (National Veterinary Services Laboratories - USDA / Diagnostic Virology Laboratory) |
| EPI_ISL_13009677 | A/turkey/North Carolina/22-009583-002-original/2022 | A / H5N1 | North America / United States / North Carolina / Johnston County | Mary Lea Killian (National Veterinary Services Laboratories - USDA / Diagnostic Virology Laboratory) |
| EPI_ISL_13009689 | A/chicken/Wyoming/22-009599-001-original/2022 | A / H5N1 | North America / United States / Wyoming / Park County | Mary Lea Killian (National Veterinary Services Laboratories - USDA / Diagnostic Virology Laboratory) |
| EPI_ISL_13009694 | A/chicken/Wyoming/22-009599-002-original/2022 | A / H5N1 | North America / United States / Wyoming / Park County | Mary Lea Killian (National Veterinary Services Laboratories - USDA / Diagnostic Virology Laboratory) |
| EPI_ISL_13026099 | A/Greylag goose/Luxembourg/21217773/2021 | A / H5N1 | Europe / Luxembourg / District de Grevenmacher | Chantal J. Snoeck (Luxembourg Institute of Health / Department of Infection and Immunity) |
| EPI_ISL_13026104 | A/Canada goose/Luxembourg/21217776/2021 | A / H5N1 | Europe / Luxembourg / District de Grevenmacher / Wintrange | Chantal J. Snoeck (Luxembourg Institute of Health / Department of Infection and Immunity) |
| EPI_ISL_13026105 | A/Canada goose/Luxembourg/21237199/2021 | A / H5N1 | Europe / Luxembourg / District de Grevenmacher / Remerschen | Chantal J. Snoeck (Luxembourg Institute of Health / Department of Infection and Immunity) |
| EPI_ISL_13026106 | A/Canada goose/Luxembourg/21239614/2021 | A / H5N1 | Europe / Luxembourg / District de Grevenmacher / Remerschen | Chantal J. Snoeck (Luxembourg Institute of Health / Department of Infection and Immunity) |
| EPI_ISL_13026236 | A/Grey heron/Luxembourg/21243177/2021 | A / H5N1 | Europe / Luxembourg / District de Grevenmacher / Remerschen | Chantal J. Snoeck (Luxembourg Institute of Health / Department of Infection and Immunity) |
| EPI_ISL_13026237 | A/Great cormorant/Luxembourg/21243185/2021 | A / H5N1 | Europe / Luxembourg / District de Grevenmacher / Remerschen | Chantal J. Snoeck (Luxembourg Institute of Health / Department of Infection and Immunity) |
| EPI_ISL_13026245 | A/Canada goose/Luxembourg/22012198/2022 | A / H5N1 | Europe / Luxembourg / District de Grevenmacher / Wintrange | Chantal J. Snoeck (Luxembourg Institute of Health / Department of Infection and Immunity) |
| EPI_ISL_13026246 | A/Barnacle goose/Luxembourg/22033922/2022 | A / H5N1 | Europe / Luxembourg / District de Diekirch / Alscheid | Chantal J. Snoeck (Luxembourg Institute of Health / Department of Infection and Immunity) |
| EPI_ISL_1303550 | A/Chichen/Sweden/SVA210313SZ0001/KN066654-IP15/2021 | A / H5N8 | Europe / Sweden / Ostergotlands Lan / Linkopings Kommun | Siamak Zohari (National Veterinary Institute) |
| EPI_ISL_1303776 | A/Chicken/Sweden/SVA210313SZ0003/KN066666-IP16/2021 | A / H5N8 | Europe / Sweden / Skane Lan / Sjobo Kommun | Siamak Zohari (National Veterinary Institute) |
| EPI_ISL_130380 | A/duck/Jiangsu/m234/2012 | A / H5N2 | Asia / China | |
| EPI_ISL_13048381 | A/avian/Burkina_Faso/21VIR11911-1/2021 | A / H5N1 | Africa / Burkina Faso / Bonyollo | Bianca Zecchin (Istituto Zooprofilattico Sperimentale Delle Venezie) |
| EPI_ISL_13048382 | A/avian/Burkina_Faso/21VIR11911-3/2021 | A / H5N1 | Africa / Burkina Faso / Gomboussougou | Bianca Zecchin (Istituto Zooprofilattico Sperimentale Delle Venezie) |
| EPI_ISL_13048383 | A/chicken/Burkina_Faso/21VIR11911-5/2021 | A / H5N1 | Africa / Burkina Faso / Koubri | Bianca Zecchin (Istituto Zooprofilattico Sperimentale Delle Venezie) |
| EPI_ISL_13052412 | A/turkey/Maine/22-009605-001-original/2022 | A / H5N1 | North America / United States / Maine / Lincoln County | Mary Lea Killian (National Veterinary Services Laboratories - USDA / Diagnostic Virology Laboratory) |
| EPI_ISL_13052413 | A/chicken/Maine/22-009605-002-original/2022 | A / H5N1 | North America / United States / Maine / Lincoln County | Mary Lea Killian (National Veterinary Services Laboratories - USDA / Diagnostic Virology Laboratory) |
| EPI_ISL_13052414 | A/chicken/North Dakota/22-009677-001-original/2022 | A / H5N1 | North America / United States / North Dakota / Dickey County | Mary Lea Killian (National Veterinary Services Laboratories - USDA / Diagnostic Virology Laboratory) |
| EPI_ISL_13052415 | A/turkey/South Dakota/22-009688-001-original/2022 | A / H5N1 | North America / United States / South Dakota / McPherson County | Mary Lea Killian (National Veterinary Services Laboratories - USDA / Diagnostic Virology Laboratory) |
| EPI_ISL_13052416 | A/chicken/Illinois/22-009720-001-original/2022 | A / H5N1 | North America / United States / Illinois / Carroll County | Mary Lea Killian (National Veterinary Services Laboratories - USDA / Diagnostic Virology Laboratory) |
| EPI_ISL_13052417 | A/goose/Illinois/22-009720-003-original/2022 | A / H5N1 | North America / United States / Illinois / Carroll County | Mary Lea Killian (National Veterinary Services Laboratories - USDA / Diagnostic Virology Laboratory) |
| EPI_ISL_13052418 | A/chicken/Minnesota/22-009735-001-original/2022 | A / H5N1 | North America / United States / Minnesota / Stearns County | Mary Lea Killian (National Veterinary Services Laboratories - USDA / Diagnostic Virology Laboratory) |
| EPI_ISL_13052419 | A/turkey/Minnesota/22-009820-001-original/2022 | A / H5N1 | North America / United States / Minnesota / Morrison County | Mary Lea Killian (National Veterinary Services Laboratories - USDA / Diagnostic Virology Laboratory) |
| EPI_ISL_13052420 | A/turkey/Iowa/22-009825-005-original/2022 | A / H5N1 | North America / United States / Iowa / Cherokee County | Mary Lea Killian (National Veterinary Services Laboratories - USDA / Diagnostic Virology Laboratory) |
| EPI_ISL_13052422 | A/turkey/South Dakota/22-009839-001-original/2022 | A / H5N1 | North America / United States / South Dakota / Edmunds County | Mary Lea Killian (National Veterinary Services Laboratories - USDA / Diagnostic Virology Laboratory) |
| EPI_ISL_13052423 | A/turkey/North Dakota/22-009840-001-original/2022 | A / H5N1 | North America / United States / North Dakota / Dickey County | Mary Lea Killian (National Veterinary Services Laboratories - USDA / Diagnostic Virology Laboratory) |
| EPI_ISL_13052424 | A/turkey/South Dakota/22-009841-001-original/2022 | A / H5N1 | North America / United States / South Dakota / Charles Mix County | Mary Lea Killian (National Veterinary Services Laboratories - USDA / Diagnostic Virology Laboratory) |
| EPI_ISL_13052425 | A/turkey/Missouri/22-009845-004-original/2022 | A / H5N1 | North America / United States / Missouri / Jasper County | Mary Lea Killian (National Veterinary Services Laboratories - USDA / Diagnostic Virology Laboratory) |
| EPI_ISL_13052426 | A/chicken/Wyoming/22-009849-001-original/2022 | A / H5N1 | North America / United States / Wyoming / Fremont County | Mary Lea Killian (National Veterinary Services Laboratories - USDA / Diagnostic Virology Laboratory) |
| EPI_ISL_13052427 | A/silkie chicken/Wisconsin/22-009934-002-original/2022 | A / H5N1 | North America / United States / Wisconsin / Rock County | Mary Lea Killian (National Veterinary Services Laboratories - USDA / Diagnostic Virology Laboratory) |
| EPI_ISL_13052429 | A/turkey/Minnesota/22-009976-003-original/2022 | A / H5N1 | North America / United States / Minnesota / Stearns County | Mary Lea Killian (National Veterinary Services Laboratories - USDA / Diagnostic Virology Laboratory) |
| EPI_ISL_13052430 | A/turkey/North Dakota/22-009978-001-original/2022 | A / H5N1 | North America / United States / North Dakota / LaMoure County | Mary Lea Killian (National Veterinary Services Laboratories - USDA / Diagnostic Virology Laboratory) |
| EPI_ISL_13052431 | A/turkey/North Dakota/22-009978-002-original/2022 | A / H5N1 | North America / United States / North Dakota / LaMoure County | Mary Lea Killian (National Veterinary Services Laboratories - USDA / Diagnostic Virology Laboratory) |

| | | | | |
|---|---|---|---|---|
| EPI_ISL_13052432 | A/turkey/South Dakota/22-009979-001-original/2022 | A / H5N1 | North America / United States / South Dakota / Spink County | Mary Lea Killian (National Veterinary Services Laboratories - USDA / Diagnostic Virology Laboratory) |
| EPI_ISL_13052433 | A/turkey/South Dakota/22-009979-002-original/2022 | A / H5N1 | North America / United States / South Dakota / Spink County | Mary Lea Killian (National Veterinary Services Laboratories - USDA / Diagnostic Virology Laboratory) |
| EPI_ISL_13052717 | A/fox/Wisconsin/22-013774-002-original/2022 | A / H5N1 | North America / United States / Wisconsin / Jefferson County | Mary Lea Killian (National Veterinary Services Laboratories - USDA / Diagnostic Virology Laboratory) |
| EPI_ISL_13052718 | A/fox/Wisconsin/22-013774-009-original/2022 | A / H5N1 | North America / United States / Wisconsin / Waushara County | Mary Lea Killian (National Veterinary Services Laboratories - USDA / Diagnostic Virology Laboratory) |
| EPI_ISL_13052719 | A/fox/Wisconsin/22-013774-014-original/2022 | A / H5N1 | North America / United States / Wisconsin / Rock County | Mary Lea Killian (National Veterinary Services Laboratories - USDA / Diagnostic Virology Laboratory) |
| EPI_ISL_13052720 | A/fox/Minnesota/22-014182-001-original/2022 | A / H5N1 | North America / United States / Minnesota / Anoka County | Mary Lea Killian (National Veterinary Services Laboratories - USDA / Diagnostic Virology Laboratory) |
| EPI_ISL_13052721 | A/fox/Iowa/22-014421-001-original/2022 | A / H5N1 | North America / United States / Iowa / Hancock County | Mary Lea Killian (National Veterinary Services Laboratories - USDA / Diagnostic Virology Laboratory) |
| EPI_ISL_13052722 | A/fox/Minnesota/22-014660-001-original/2022 | A / H5N1 | North America / United States / Minnesota / Dakota County | Mary Lea Killian (National Veterinary Services Laboratories - USDA / Diagnostic Virology Laboratory) |
| EPI_ISL_13052723 | A/fox/Minnesota/22-014660-002-original/2022 | A / H5N1 | North America / United States / Minnesota / Dakota County | Mary Lea Killian (National Veterinary Services Laboratories - USDA / Diagnostic Virology Laboratory) |
| EPI_ISL_13052724 | A/fox/Minnesota/22-014661-001-original/2022 | A / H5N1 | North America / United States / Minnesota / Washington County | Mary Lea Killian (National Veterinary Services Laboratories - USDA / Diagnostic Virology Laboratory) |
| EPI_ISL_13052725 | A/fox/Wisconsin/22-014746-010-original/2022 | A / H5N1 | North America / United States / Wisconsin / Adam County | Mary Lea Killian (National Veterinary Services Laboratories - USDA / Diagnostic Virology Laboratory) |
| EPI_ISL_13052726 | A/fox/Wisconsin/22-014746-016-original/2022 | A / H5N1 | North America / United States / Wisconsin / Adam County | Mary Lea Killian (National Veterinary Services Laboratories - USDA / Diagnostic Virology Laboratory) |
| EPI_ISL_13052727 | A/fox/Wisconsin/22-014746-023-original/2022 | A / H5N1 | North America / United States / Wisconsin / Grant County | Mary Lea Killian (National Veterinary Services Laboratories - USDA / Diagnostic Virology Laboratory) |
| EPI_ISL_13052728 | A/fox/Wisconsin/22-014746-029-original/2022 | A / H5N1 | North America / United States / Wisconsin / Grant County | Mary Lea Killian (National Veterinary Services Laboratories - USDA / Diagnostic Virology Laboratory) |
| EPI_ISL_1305775 | A/Chicken/Sweden/SVA210314SZ0001/KN066672-IP17/2021 | A / H5N8 | Europe / Sweden / Ostergotlands Lan / Mjolby Kommun | Siamak Zohari (National Veterinary Institute) |
| EPI_ISL_1305776 | A/Chicken/Sweden/SVA210314SZ0002/KN066679-IP18/2021 | A / H5N8 | Europe / Sweden / Ostergotlands Lan / Mjolby Kommun | Siamak Zohari (National Veterinary Institute) |
| EPI_ISL_1305777 | A/Common peacock/Sweden/SVA210311SZ0002/KN001568-IP | A / H5N5 | Europe / Sweden / Skane Lan / Skurups Kommun | Siamak Zohari (National Veterinary Institute) |
| EPI_ISL_1305778 | A/Goose/Sweden/SVA210311SZ0003/KN0001570-IP14/2021 | A / H5N5 | Europe / Sweden / Skane Lan / Skurups Kommun | Siamak Zohari (National Veterinary Institute) |
| EPI_ISL_1305779 | A/Duck/Sweden/SVA210311SZ0004/KN0001570-IP14/2021 | A / H5N5 | Europe / Sweden / Skane Lan / Skurups Kommun | Siamak Zohari (National Veterinary Institute) |
| EPI_ISL_1305780 | A/Canada_goose/Sweden/SVA210302SZ0455/KN000714/SKsin | A / H5N8 | Europe / Sweden / Skane Lan / Simrishamns Kommun | Siamak Zohari (National Veterinary Institute) |
| EPI_ISL_1305781 | A/Mute Swan/Sweden/SVA210302Z0465/KN000716/SOTR/20 | A / H5N5 | Europe / Sweden / Sodermanlands Lan / Trosa Kommun | Siamak Zohari (National Veterinary Institute) |
| EPI_ISL_13068727 | A/common crane/Israel/670/2021 | A / H5N1 | Asia / Israel / Northern District / Agamon Hula | Irina Shkoda (Kimron Veterinary Institute) |
| EPI_ISL_1307695 | A/Mute Swan/Sweden/SVA210304SZ0311/KN000739/VG/202 | A / H5N8 | Europe / Sweden / Vastra Gotalands Lan / Ockero Kommun | Siamak Zohari (National Veterinary Institute) |
| EPI_ISL_1307703 | A/Canade goose/Sweden/SVA210304SZ0320/KN000779/SK/20 | A / H5N8 | Europe / Sweden / Skane Lan / Klippans Kommun | Siamak Zohari (National Veterinary Institute) |
| EPI_ISL_1307709 | A/Canada goose/Sweden/SVA210305SZ0255/KN000797/SK/20 | A / H5N8 | Europe / Sweden / Skane Lan / Tomelilla Kommun | Siamak Zohari (National Veterinary Institute) |
| EPI_ISL_1307716 | A/Mute Swan/Sweden/SVA210303SZ0380/KN000800/St/2021 | A / H5N8 | Europe / Sweden / Stockholms Lan / Sodertalje Kommun | Siamak Zohari (National Veterinary Institute) |
| EPI_ISL_1307720 | A/northern goshawk/Sweden/SVA210303SZ0381/KN000801/J | A / H5N8 | Europe / Sweden / Jonkopings Lan / Jonkopings Kommun | Siamak Zohari (National Veterinary Institute) |
| EPI_ISL_1308010 | A/Mute Swan/Sweden/SVA210303SZ0392/KN000806/Kal/202 | A / H5N5 | Europe / Sweden / Kalmar Lan / Monsteras Kommun | Siamak Zohari (National Veterinary Institute) |
| EPI_ISL_13114176 | A/Broiler chicken/BC/FAV-0228-OS/2022 | A / H5N1 | North America / Canada / British Columbia | Yohannes Berhane (Canadian Food Inspection Agency / Avian Unit) |
| EPI_ISL_13116131 | A/Chicken/BC/FAV-0346-OS/2022 | A / H5N1 | North America / Canada / British Columbia | Yohannes Berhane (Canadian Food Inspection Agency / Avian Unit) |
| EPI_ISL_13117168 | A/Chicken/BC/FAV-0348-OS/2022 | A / H5N1 | North America / Canada / British Columbia | Yohannes Berhane (Canadian Food Inspection Agency / Avian Unit) |
| EPI_ISL_13117288 | A/Chicken/BC/FAV-0402-OS/2022 | A / H5N1 | North America / Canada / British Columbia | Yohannes Berhane (Canadian Food Inspection Agency / Avian Unit) |
| EPI_ISL_13117289 | A/Chicken/BC/FAV-0369-OS/2022 | A / H5N1 | North America / Canada / British Columbia | Yohannes Berhane (Canadian Food Inspection Agency / Avian Unit) |
| EPI_ISL_13117290 | A/Chicken/BC/FAV-0460-OS/2022 | A / H5N1 | North America / Canada / British Columbia | Yohannes Berhane (Canadian Food Inspection Agency / Avian Unit) |
| EPI_ISL_13117299 | A/Chicken/BC/FAV-0488-OS/2022 | A / H5N1 | North America / Canada / British Columbia | Yohannes Berhane (Canadian Food Inspection Agency / Avian Unit) |
| EPI_ISL_13175606 | A/Turkey/Sweden/SVA210315SZ0260/FB067516-IP19/2021 | A / H5N8 | Europe / Sweden / Skane Lan / Tomelilla Kommun | Siamak Zohari (National Veterinary Institute) |
| EPI_ISL_13175608 | A/Chicken/Sweden/SVA210223SZ0647/FB046588-IP7/2021 | A / H5N8 | Europe / Sweden / Ostergotlands Lan / Linkopings Kommun | Siamak Zohari (National Veterinary Institute) |
| EPI_ISL_13175667 | A/Common pheasant/Sweden/SVA210224SZ0005/KN000542-I | A / H5N8 | Europe / Sweden / Skane Lan / Trelleborgs Kommun | Siamak Zohari (National Veterinary Institute) |
| EPI_ISL_13201050 | A/Lesser Black-backed Gull/Netherlands/1/2022 | A / H5N1 | Europe / Netherlands / Provincie Noord-Holland / De Kreupel | Sanne Thewessen (Erasmus Medical Center / Viroscience) |
| EPI_ISL_13201074 | A/European polecat/Netherlands/1/2022 | A / H5N1 | Europe / Netherlands / Zuid-Holland | Sanne Thewessen (Erasmus Medical Center / Viroscience) |
| EPI_ISL_13243350 | A/duck/Bangladesh/19D1874/2022 | A / H5N1 | Asia / Bangladesh | Mohammad Enayet Hossain (International Centre for Diarrhoeal Disease Research (ICDDR,B) / Virology) |
| EPI_ISL_13243698 | A/duck/Bangladesh/19D1819/2021 | A / H5N1 | Asia / Bangladesh | Mohammad Enayet Hossain (International Centre for Diarrhoeal Disease Research (ICDDR,B) / Virology) |
| EPI_ISL_13245602 | A/white-tailed eagle/Iceland/2022AI02104/2021 | A / H5N1 | Europe / Iceland / Westfjords / Kerlingarfjörður | Jacqueline King (Friedrich-Loeffler-Institut) |
| EPI_ISL_13246267 | A/chicken/Iceland/2022AI02564/2022 | A / H5N1 | Europe / Iceland / South | Jacqueline King (Friedrich-Loeffler-Institut) |
| EPI_ISL_13246657 | A/chicken/Iceland/2022AI02565/2022 | A / H5N1 | Europe / Iceland / South | Jacqueline King (Friedrich-Loeffler-Institut) |
| EPI_ISL_13285237 | A/turkey/South Dakota/22-009980-001-original/2022 | A / H5N1 | North America / United States / South Dakota / Lake County | Mary Lea Killian (National Veterinary Services Laboratories - USDA / Diagnostic Virology Laboratory) |
| EPI_ISL_13285238 | A/turkey/South Dakota/22-009980-002-original/2022 | A / H5N1 | North America / United States / South Dakota / Lake County | Mary Lea Killian (National Veterinary Services Laboratories - USDA / Diagnostic Virology Laboratory) |
| EPI_ISL_13285239 | A/turkey/North Carolina/22-009981-001-original/2022 | A / H5N1 | North America / United States / North Carolina / Johnston County | Mary Lea Killian (National Veterinary Services Laboratories - USDA / Diagnostic Virology Laboratory) |
| EPI_ISL_13285240 | A/turkey/North Carolina/22-009981-004-original/2022 | A / H5N1 | North America / United States / North Carolina / Johnston County | Mary Lea Killian (National Veterinary Services Laboratories - USDA / Diagnostic Virology Laboratory) |
| EPI_ISL_13285241 | A/turkey/North Carolina/22-009982-001-original/2022 | A / H5N1 | North America / United States / North Carolina / Johnston County | Mary Lea Killian (National Veterinary Services Laboratories - USDA / Diagnostic Virology Laboratory) |
| EPI_ISL_13285242 | A/turkey/North Carolina/22-009982-002-original/2022 | A / H5N1 | North America / United States / North Carolina / Johnston County | Mary Lea Killian (National Veterinary Services Laboratories - USDA / Diagnostic Virology Laboratory) |
| EPI_ISL_13285243 | A/turkey/North Carolina/22-009983-007-original/2022 | A / H5N1 | North America / United States / North Carolina / Wayne County | Mary Lea Killian (National Veterinary Services Laboratories - USDA / Diagnostic Virology Laboratory) |
| EPI_ISL_13285244 | A/turkey/North Carolina/22-009983-008-original/2022 | A / H5N1 | North America / United States / North Carolina / Wayne County | Mary Lea Killian (National Veterinary Services Laboratories - USDA / Diagnostic Virology Laboratory) |
| EPI_ISL_13285245 | A/pheasant/Texas/22-009984-001-original/2022 | A / H5N1 | North America / United States / Texas / Erath County | Mary Lea Killian (National Veterinary Services Laboratories - USDA / Diagnostic Virology Laboratory) |
| EPI_ISL_13285246 | A/chicken/Iowa/22-009985-001-original/2022 | A / H5N1 | North America / United States / Iowa / Humboldt County | Mary Lea Killian (National Veterinary Services Laboratories - USDA / Diagnostic Virology Laboratory) |
| EPI_ISL_13285536 | A/turkey/Iowa/22-009986-001-original/2022 | A / H5N1 | North America / United States / Iowa / Sac County | Mary Lea Killian (National Veterinary Services Laboratories - USDA / Diagnostic Virology Laboratory) |
| EPI_ISL_13285537 | A/turkey/Iowa/22-009986-002-original/2022 | A / H5N1 | North America / United States / Iowa / Sac County | Mary Lea Killian (National Veterinary Services Laboratories - USDA / Diagnostic Virology Laboratory) |
| EPI_ISL_13285538 | A/turkey/Minnesota/22-009996-001-original/2022 | A / H5N1 | North America / United States / Minnesota / Le Sueur County | Mary Lea Killian (National Veterinary Services Laboratories - USDA / Diagnostic Virology Laboratory) |
| EPI_ISL_13285539 | A/turkey/Minnesota/22-009997-001-original/2022 | A / H5N1 | North America / United States / Minnesota / Dodge County | Mary Lea Killian (National Veterinary Services Laboratories - USDA / Diagnostic Virology Laboratory) |
| EPI_ISL_13285540 | A/turkey/Minnesota/22-009997-002-original/2022 | A / H5N1 | North America / United States / Minnesota / Dodge County | Mary Lea Killian (National Veterinary Services Laboratories - USDA / Diagnostic Virology Laboratory) |
| EPI_ISL_13285541 | A/turkey/Minnesota/22-009998-001-original/2022 | A / H5N1 | North America / United States / Minnesota / Kandiyohi County | Mary Lea Killian (National Veterinary Services Laboratories - USDA / Diagnostic Virology Laboratory) |
| EPI_ISL_13285542 | A/turkey/Minnesota/22-009999-001-original/2022 | A / H5N1 | North America / United States / Minnesota / Becker County | Mary Lea Killian (National Veterinary Services Laboratories - USDA / Diagnostic Virology Laboratory) |
| EPI_ISL_13285543 | A/turkey/Minnesota/22-010000-001-original/2022 | A / H5N1 | North America / United States / Minnesota / Kandiyohi County | Mary Lea Killian (National Veterinary Services Laboratories - USDA / Diagnostic Virology Laboratory) |
| EPI_ISL_13285548 | A/turkey/Minnesota/22-009999-002-original/2022 | A / H5N1 | North America / United States / Minnesota / Becker County | Mary Lea Killian (National Veterinary Services Laboratories - USDA / Diagnostic Virology Laboratory) |
| EPI_ISL_13285549 | A/pheasant/Texas/22-010008-001-original/2022 | A / H5N1 | North America / United States / Texas / Erath County | Mary Lea Killian (National Veterinary Services Laboratories - USDA / Diagnostic Virology Laboratory) |
| EPI_ISL_13285550 | A/pheasant/Texas/22-010008-002-original/2022 | A / H5N1 | North America / United States / Texas / Erath County | Mary Lea Killian (National Veterinary Services Laboratories - USDA / Diagnostic Virology Laboratory) |
| EPI_ISL_13285551 | A/turkey/South Dakota/22-010013-001-original/2022 | A / H5N1 | North America / United States / South Dakota / McPherson County | Mary Lea Killian (National Veterinary Services Laboratories - USDA / Diagnostic Virology Laboratory) |
| EPI_ISL_13285552 | A/turkey/South Dakota/22-010013-002-original/2022 | A / H5N1 | North America / United States / South Dakota / McPherson County | Mary Lea Killian (National Veterinary Services Laboratories - USDA / Diagnostic Virology Laboratory) |
| EPI_ISL_13285553 | A/turkey/North Dakota/22-010015-001-original/2022 | A / H5N1 | North America / United States / North Dakota / LaMoure County | Mary Lea Killian (National Veterinary Services Laboratories - USDA / Diagnostic Virology Laboratory) |
| EPI_ISL_13285554 | A/turkey/North Dakota/22-010015-002-original/2022 | A / H5N1 | North America / United States / North Dakota / LaMoure County | Mary Lea Killian (National Veterinary Services Laboratories - USDA / Diagnostic Virology Laboratory) |
| EPI_ISL_13285555 | A/turkey/Minnesota/22-010085-001-original/2022 | A / H5N1 | North America / United States / Minnesota / Morrison County | Mary Lea Killian (National Veterinary Services Laboratories - USDA / Diagnostic Virology Laboratory) |
| EPI_ISL_13285556 | A/turkey/Minnesota/22-010085-002-original/2022 | A / H5N1 | North America / United States / Minnesota / Morrison County | Mary Lea Killian (National Veterinary Services Laboratories - USDA / Diagnostic Virology Laboratory) |

| | | | | |
|---|---|---|---|---|
| EPI_ISL_13285557 | A/turkey/Minnesota/22-010092-002-original/2022 | A / H5N1 | North America / United States / Minnesota / Kandiyohi County | Mary Lea Killian (National Veterinary Services Laboratories - USDA / Diagnostic Virology Laboratory) |
| EPI_ISL_13285558 | A/turkey/Minnesota/22-010092-003-original/2022 | A / H5N1 | North America / United States / Minnesota / Kandiyohi County | Mary Lea Killian (National Veterinary Services Laboratories - USDA / Diagnostic Virology Laboratory) |
| EPI_ISL_13285559 | A/turkey/Iowa/22-010094-001-original/2022 | A / H5N1 | North America / United States / Iowa / Hamilton County | Mary Lea Killian (National Veterinary Services Laboratories - USDA / Diagnostic Virology Laboratory) |
| EPI_ISL_13285560 | A/turkey/Iowa/22-010094-002-original/2022 | A / H5N1 | North America / United States / Iowa / Hamilton County | Mary Lea Killian (National Veterinary Services Laboratories - USDA / Diagnostic Virology Laboratory) |
| EPI_ISL_13286627 | A/turkey/Minnesota/22-009996-002-original/2022 | A / H5N1 | North America / United States / Minnesota / Le Sueur County | Mary Lea Killian (National Veterinary Services Laboratories - USDA / Diagnostic Virology Laboratory) |
| EPI_ISL_13300319 | A/brant goose/Netherlands/22009881-002/2022 | A / H5N1 | Europe / Netherlands / Provincie Friesland | Rene Heutink (Wageningen Bioveterinary Research) |
| EPI_ISL_13300324 | A/white stork/Netherlands/22009973-002/2022 | A / H5N1 | Europe / Netherlands / Provincie Gelderland | Rene Heutink (Wageningen Bioveterinary Research) |
| EPI_ISL_13338081 | A/Northern gannet/Sweden/SVA220525SZ0402/FB001671/O-2 | A / H5N1 | Europe / Sweden / Vastra Gotalands Lan / Molndals Kommun | Siamak Zohari (National Veterinary Institute) |
| EPI_ISL_13370915 | A/domestic_duck/England/032919/2022 | A / H5N1 | Europe / United Kingdom / England | Alex Byrne (Animal and Plant Health Agency (APHA) / Virology Department) |
| EPI_ISL_13370918 | A/domestic_duck/England/041295/2022 | A / H5N1 | Europe / United Kingdom / England | Alex Byrne (Animal and Plant Health Agency (APHA) / Virology Department) |
| EPI_ISL_13370924 | A/chicken/England/053826/2022 | A / H5N1 | Europe / United Kingdom / England | Alex Byrne (Animal and Plant Health Agency (APHA) / Virology Department) |
| EPI_ISL_13370925 | A/chicken/England/063896/2022 | A / H5N1 | Europe / United Kingdom / England | Alex Byrne (Animal and Plant Health Agency (APHA) / Virology Department) |
| EPI_ISL_13418526 | A/Little egret/Israel/172/2022 | A / H5N1 | Asia / Israel / Northern District / Kinneret Beach | Irina Shkoda (Kimron Veterinary Institute) |
| EPI_ISL_13429290 | A/Caspian Gull/Netherlands/4/2022 | A / H5N1 | Europe / Netherlands / De Kreupel | Sanne Thewessen (Erasmus Medical Center / Viroscience) |
| EPI_ISL_13429291 | A/Sandwich Tern/Netherlands/8/2022 | A / H5N1 | Europe / Netherlands / Stellendam, Scheelhoekeiland | Sanne Thewessen (Erasmus Medical Center / Viroscience) |
| EPI_ISL_13429292 | A/Sandwich Tern/Netherlands/5/2022 | A / H5N1 | Europe / Netherlands / Stellendam, Scheelhoekeiland | Sanne Thewessen (Erasmus Medical Center / Viroscience) |
| EPI_ISL_13429293 | A/Black-headed gull/Netherlands/5/2022 | A / H5N1 | Europe / Netherlands / De Kreupel | Sanne Thewessen (Erasmus Medical Center / Viroscience) |
| EPI_ISL_13429294 | A/Black-headed gull/Netherlands/6/2022 | A / H5N1 | Europe / Netherlands / De Kreupel | Sanne Thewessen (Erasmus Medical Center / Viroscience) |
| EPI_ISL_13429295 | A/Sandwich Tern/Netherlands/1/2022 | A / H5N1 | Europe / Netherlands / Den Oever, Waddenhaven | Sanne Thewessen (Erasmus Medical Center / Viroscience) |
| EPI_ISL_13429296 | A/Sandwich Tern/Netherlands/2/2022 | A / H5N1 | Europe / Netherlands / Haringvliet, Slijkplaat | Sanne Thewessen (Erasmus Medical Center / Viroscience) |
| EPI_ISL_13429297 | A/Sandwich Tern/Netherlands/3/2022 | A / H5N1 | Europe / Netherlands / Haringvliet, Slijkplaat | Sanne Thewessen (Erasmus Medical Center / Viroscience) |
| EPI_ISL_13429298 | A/Common Tern/Netherlands/1/2022 | A / H5N1 | Europe / Netherlands / Haringvliet, Bliek | Sanne Thewessen (Erasmus Medical Center / Viroscience) |
| EPI_ISL_13429299 | A/Sandwich Tern/Netherlands/6/2022 | A / H5N1 | Europe / Netherlands / Stellendam, Scheelhoekeiland | Sanne Thewessen (Erasmus Medical Center / Viroscience) |
| EPI_ISL_13429300 | A/Sandwich Tern/Netherlands/7/2022 | A / H5N1 | Europe / Netherlands / Stellendam, Scheelhoekeiland | Sanne Thewessen (Erasmus Medical Center / Viroscience) |
| EPI_ISL_13431305 | A/turkey/Minnesota/22-010122-001-original/2022 | A / H5N1 | North America / United States / Minnesota / Stearns County | Mary Lea Killian (National Veterinary Services Laboratories - USDA / Diagnostic Virology Laboratory) |
| EPI_ISL_13431306 | A/turkey/Minnesota/22-010122-002-original/2022 | A / H5N1 | North America / United States / Minnesota / Stearns County | Mary Lea Killian (National Veterinary Services Laboratories - USDA / Diagnostic Virology Laboratory) |
| EPI_ISL_13431307 | A/turkey/Minnesota/22-010123-001-original/2022 | A / H5N1 | North America / United States / Minnesota / Waseca County | Mary Lea Killian (National Veterinary Services Laboratories - USDA / Diagnostic Virology Laboratory) |
| EPI_ISL_13431308 | A/turkey/Minnesota/22-010123-004-original/2022 | A / H5N1 | North America / United States / Minnesota / Waseca County | Mary Lea Killian (National Veterinary Services Laboratories - USDA / Diagnostic Virology Laboratory) |
| EPI_ISL_13431309 | A/turkey/Minnesota/22-010124-001-original/2022 | A / H5N1 | North America / United States / Minnesota / Meeker County | Mary Lea Killian (National Veterinary Services Laboratories - USDA / Diagnostic Virology Laboratory) |
| EPI_ISL_13431310 | A/turkey/Minnesota/22-010124-002-original/2022 | A / H5N1 | North America / United States / Minnesota / Meeker County | Mary Lea Killian (National Veterinary Services Laboratories - USDA / Diagnostic Virology Laboratory) |
| EPI_ISL_13431311 | A/turkey/Minnesota/22-010125-001-original/2022 | A / H5N1 | North America / United States / Minnesota / Morrison County | Mary Lea Killian (National Veterinary Services Laboratories - USDA / Diagnostic Virology Laboratory) |
| EPI_ISL_13431312 | A/turkey/Minnesota/22-010125-002-original/2022 | A / H5N1 | North America / United States / Minnesota / Morrison County | Mary Lea Killian (National Veterinary Services Laboratories - USDA / Diagnostic Virology Laboratory) |
| EPI_ISL_13431313 | A/turkey/Minnesota/22-010126-001-original/2022 | A / H5N1 | North America / United States / Minnesota / Big Stone County | Mary Lea Killian (National Veterinary Services Laboratories - USDA / Diagnostic Virology Laboratory) |
| EPI_ISL_13431314 | A/turkey/Minnesota/22-010126-002-original/2022 | A / H5N1 | North America / United States / Minnesota / Big Stone County | Mary Lea Killian (National Veterinary Services Laboratories - USDA / Diagnostic Virology Laboratory) |
| EPI_ISL_13431688 | A/turkey/Minnesota/22-010127-001-original/2022 | A / H5N1 | North America / United States / Minnesota / Morrison County | Mary Lea Killian (National Veterinary Services Laboratories - USDA / Diagnostic Virology Laboratory) |
| EPI_ISL_13431689 | A/turkey/Minnesota/22-010127-002-original/2022 | A / H5N1 | North America / United States / Minnesota / Morrison County | Mary Lea Killian (National Veterinary Services Laboratories - USDA / Diagnostic Virology Laboratory) |
| EPI_ISL_13431690 | A/turkey/Iowa/22-010132-001-original/2022 | A / H5N1 | North America / United States / Iowa / Hardin County | Mary Lea Killian (National Veterinary Services Laboratories - USDA / Diagnostic Virology Laboratory) |
| EPI_ISL_13431691 | A/turkey/Iowa/22-010132-002-original/2022 | A / H5N1 | North America / United States / Iowa / Hardin County | Mary Lea Killian (National Veterinary Services Laboratories - USDA / Diagnostic Virology Laboratory) |
| EPI_ISL_13431692 | A/turkey/South Dakota/22-010135-001-original/2022 | A / H5N1 | North America / United States / South Dakota / Faulk County | Mary Lea Killian (National Veterinary Services Laboratories - USDA / Diagnostic Virology Laboratory) |
| EPI_ISL_13431693 | A/turkey/South Dakota/22-010135-002-original/2022 | A / H5N1 | North America / United States / South Dakota / Faulk County | Mary Lea Killian (National Veterinary Services Laboratories - USDA / Diagnostic Virology Laboratory) |
| EPI_ISL_13431694 | A/turkey/South Dakota/22-010137-001-original/2022 | A / H5N1 | North America / United States / South Dakota / Clark County | Mary Lea Killian (National Veterinary Services Laboratories - USDA / Diagnostic Virology Laboratory) |
| EPI_ISL_13431695 | A/turkey/South Dakota/22-010137-002-original/2022 | A / H5N1 | North America / United States / South Dakota / Clark County | Mary Lea Killian (National Veterinary Services Laboratories - USDA / Diagnostic Virology Laboratory) |
| EPI_ISL_13431696 | A/turkey/South Dakota/22-010138-001-original/2022 | A / H5N1 | North America / United States / South Dakota / Spink County | Mary Lea Killian (National Veterinary Services Laboratories - USDA / Diagnostic Virology Laboratory) |
| EPI_ISL_13431697 | A/turkey/South Dakota/22-010138-002-original/2022 | A / H5N1 | North America / United States / South Dakota / Spink County | Mary Lea Killian (National Veterinary Services Laboratories - USDA / Diagnostic Virology Laboratory) |
| EPI_ISL_13432603 | A/turkey/South Dakota/22-010139-001-original/2022 | A / H5N1 | North America / United States / South Dakota / Beadle County | Mary Lea Killian (National Veterinary Services Laboratories - USDA / Diagnostic Virology Laboratory) |
| EPI_ISL_13432604 | A/turkey/South Dakota/22-010139-002-original/2022 | A / H5N1 | North America / United States / South Dakota / Beadle County | Mary Lea Killian (National Veterinary Services Laboratories - USDA / Diagnostic Virology Laboratory) |
| EPI_ISL_13432605 | A/chicken/North Dakota/22-010141-001-original/2022 | A / H5N1 | North America / United States / North Dakota / Cass County | Mary Lea Killian (National Veterinary Services Laboratories - USDA / Diagnostic Virology Laboratory) |
| EPI_ISL_13432606 | A/turkey/Missouri/22-010142-001-original/2022 | A / H5N1 | North America / United States / Missouri / Lawrence County | Mary Lea Killian (National Veterinary Services Laboratories - USDA / Diagnostic Virology Laboratory) |
| EPI_ISL_13432607 | A/turkey/North Carolina/22-010144-001-original/2022 | A / H5N1 | North America / United States / North Carolina / Wayne County | Mary Lea Killian (National Veterinary Services Laboratories - USDA / Diagnostic Virology Laboratory) |
| EPI_ISL_13432608 | A/turkey/North Carolina/22-010144-002-original/2022 | A / H5N1 | North America / United States / North Carolina / Wayne County | Mary Lea Killian (National Veterinary Services Laboratories - USDA / Diagnostic Virology Laboratory) |
| EPI_ISL_13432609 | A/turkey/North Carolina/22-010145-001-original/2022 | A / H5N1 | North America / United States / North Carolina / Wayne County | Mary Lea Killian (National Veterinary Services Laboratories - USDA / Diagnostic Virology Laboratory) |
| EPI_ISL_13432610 | A/turkey/North Carolina/22-010145-002-original/2022 | A / H5N1 | North America / United States / North Carolina / Wayne County | Mary Lea Killian (National Veterinary Services Laboratories - USDA / Diagnostic Virology Laboratory) |
| EPI_ISL_13432611 | A/turkey/North Carolina/22-010146-001-original/2022 | A / H5N1 | North America / United States / North Carolina / Wayne County | Mary Lea Killian (National Veterinary Services Laboratories - USDA / Diagnostic Virology Laboratory) |
| EPI_ISL_13432612 | A/turkey/North Carolina/22-010146-002-original/2022 | A / H5N1 | North America / United States / North Carolina / Wayne County | Mary Lea Killian (National Veterinary Services Laboratories - USDA / Diagnostic Virology Laboratory) |
| EPI_ISL_13432752 | A/chicken/Nebraska/22-010155-001-original/2022 | A / H5N1 | North America / United States / Nebraska / Scotts Bluff County | Mary Lea Killian (National Veterinary Services Laboratories - USDA / Diagnostic Virology Laboratory) |
| EPI_ISL_13432764 | A/chicken/Wyoming/22-010157-001-original/2022 | A / H5N1 | North America / United States / Wyoming / Sheridan County | Mary Lea Killian (National Veterinary Services Laboratories - USDA / Diagnostic Virology Laboratory) |
| EPI_ISL_13432776 | A/chicken/Wyoming/22-010157-002-original/2022 | A / H5N1 | North America / United States / Wyoming / Sheridan County | Mary Lea Killian (National Veterinary Services Laboratories - USDA / Diagnostic Virology Laboratory) |
| EPI_ISL_13432788 | A/guinea fowl/Maine/22-010181-001-original/2022 | A / H5N1 | North America / United States / Maine / Waldo County | Mary Lea Killian (National Veterinary Services Laboratories - USDA / Diagnostic Virology Laboratory) |
| EPI_ISL_13432800 | A/goose/Maine/22-010181-002-original/2022 | A / H5N1 | North America / United States / Maine / Waldo County | Mary Lea Killian (National Veterinary Services Laboratories - USDA / Diagnostic Virology Laboratory) |
| EPI_ISL_13432814 | A/chicken/New York/22-010200-002-original/2022 | A / H5N1 | North America / United States / New York / Orleans County | Mary Lea Killian (National Veterinary Services Laboratories - USDA / Diagnostic Virology Laboratory) |
| EPI_ISL_13432826 | A/guinea fowl/New York/22-010200-003-original/2022 | A / H5N1 | North America / United States / New York / Orleans County | Mary Lea Killian (National Veterinary Services Laboratories - USDA / Diagnostic Virology Laboratory) |
| EPI_ISL_13432838 | A/turkey/Missouri/22-010296-001-original/2022 | A / H5N1 | North America / United States / Missouri / Dade County | Mary Lea Killian (National Veterinary Services Laboratories - USDA / Diagnostic Virology Laboratory) |
| EPI_ISL_13432849 | A/chicken/North Carolina/22-010298-001-original/2022 | A / H5N1 | North America / United States / North Carolina / Wayne County | Mary Lea Killian (National Veterinary Services Laboratories - USDA / Diagnostic Virology Laboratory) |
| EPI_ISL_13432861 | A/chicken/North Carolina/22-010298-002-original/2022 | A / H5N1 | North America / United States / North Carolina / Wayne County | Mary Lea Killian (National Veterinary Services Laboratories - USDA / Diagnostic Virology Laboratory) |
| EPI_ISL_13433340 | A/chicken/North Carolina/22-010299-002-original/2022 | A / H5N1 | North America / United States / North Carolina / Wayne County | Mary Lea Killian (National Veterinary Services Laboratories - USDA / Diagnostic Virology Laboratory) |
| EPI_ISL_13433352 | A/turkey/Minnesota/22-010312-001-original/2022 | A / H5N1 | North America / United States / Minnesota / Waseca County | Mary Lea Killian (National Veterinary Services Laboratories - USDA / Diagnostic Virology Laboratory) |
| EPI_ISL_13433365 | A/turkey/Minnesota/22-010312-004-original/2022 | A / H5N1 | North America / United States / Minnesota / Waseca County | Mary Lea Killian (National Veterinary Services Laboratories - USDA / Diagnostic Virology Laboratory) |
| EPI_ISL_13433376 | A/chicken/New York/22-010321-001-original/2022 | A / H5N1 | North America / United States / New York / Fulton County | Mary Lea Killian (National Veterinary Services Laboratories - USDA / Diagnostic Virology Laboratory) |
| EPI_ISL_13433388 | A/chicken/New York/22-010321-002-original/2022 | A / H5N1 | North America / United States / New York / Fulton County | Mary Lea Killian (National Veterinary Services Laboratories - USDA / Diagnostic Virology Laboratory) |
| EPI_ISL_13433400 | A/domestic duck/New York/22-010321-003-original/2022 | A / H5N1 | North America / United States / New York / Fulton County | Mary Lea Killian (National Veterinary Services Laboratories - USDA / Diagnostic Virology Laboratory) |
| EPI_ISL_13433412 | A/chicken/Wisconsin/22-010325-001-original/2022 | A / H5N1 | North America / United States / Wisconsin / Racine County | Mary Lea Killian (National Veterinary Services Laboratories - USDA / Diagnostic Virology Laboratory) |
| EPI_ISL_13433423 | A/turkey/Wisconsin/22-010325-003-original/2022 | A / H5N1 | North America / United States / Wisconsin / Racine County | Mary Lea Killian (National Veterinary Services Laboratories - USDA / Diagnostic Virology Laboratory) |

| | | | | |
|---|---|---|---|---|
| EPI_ISL_13433435 | A/turkey/South Dakota/22-010346-001-original/2022 | A / H5N1 | North America / United States / South Dakota / McPherson County | Mary Lea Killian (National Veterinary Services Laboratories - USDA / Diagnostic Virology Laboratory) |
| EPI_ISL_13433447 | A/turkey/South Dakota/22-010346-002-original/2022 | A / H5N1 | North America / United States / South Dakota / McPherson County | Mary Lea Killian (National Veterinary Services Laboratories - USDA / Diagnostic Virology Laboratory) |
| EPI_ISL_13434393 | A/chicken/Wyoming/22-010378-001-original/2022 | A / H5N1 | North America / United States / Wyoming / Park County | Mary Lea Killian (National Veterinary Services Laboratories - USDA / Diagnostic Virology Laboratory) |
| EPI_ISL_13434407 | A/chicken/Montana/22-010445-001-original/2022 | A / H5N1 | North America / United States / Montana / Judith Basin County | Mary Lea Killian (National Veterinary Services Laboratories - USDA / Diagnostic Virology Laboratory) |
| EPI_ISL_13434418 | A/chicken/Montana/22-010445-003-original/2022 | A / H5N1 | North America / United States / Montana / Judith Basin County | Mary Lea Killian (National Veterinary Services Laboratories - USDA / Diagnostic Virology Laboratory) |
| EPI_ISL_13434430 | A/turkey/North Carolina/22-010448-001-original/2022 | A / H5N1 | North America / United States / North Carolina / Wayne County | Mary Lea Killian (National Veterinary Services Laboratories - USDA / Diagnostic Virology Laboratory) |
| EPI_ISL_13434442 | A/turkey/North Carolina/22-010448-003-original/2022 | A / H5N1 | North America / United States / North Carolina / Wayne County | Mary Lea Killian (National Veterinary Services Laboratories - USDA / Diagnostic Virology Laboratory) |
| EPI_ISL_13434454 | A/Cascade duck/Montana/22-010454-001-original/2022 | A / H5N1 | North America / United States / Montana / Cascade County | Mary Lea Killian (National Veterinary Services Laboratories - USDA / Diagnostic Virology Laboratory) |
| EPI_ISL_13434466 | A/Cascade duck/Montana/22-010454-003-original/2022 | A / H5N1 | North America / United States / Montana / Cascade County | Mary Lea Killian (National Veterinary Services Laboratories - USDA / Diagnostic Virology Laboratory) |
| EPI_ISL_13434477 | A/turkey/Minnesota/22-010463-002-original/2022 | A / H5N1 | North America / United States / Minnesota / Kandiyohi County | Mary Lea Killian (National Veterinary Services Laboratories - USDA / Diagnostic Virology Laboratory) |
| EPI_ISL_13434489 | A/turkey/Minnesota/22-010464-001-original/2022 | A / H5N1 | North America / United States / Minnesota / Renville County | Mary Lea Killian (National Veterinary Services Laboratories - USDA / Diagnostic Virology Laboratory) |
| EPI_ISL_13434551 | A/turkey/Minnesota/22-010464-002-original/2022 | A / H5N1 | North America / United States / Minnesota / Renville County | Mary Lea Killian (National Veterinary Services Laboratories - USDA / Diagnostic Virology Laboratory) |
| EPI_ISL_13453559 | A/domestic_duck/England/073865/2022 | A / H5N1 | Europe / United Kingdom / England | Alex Byrne (Animal and Plant Health Agency (APHA) / Virology Department) |
| EPI_ISL_13453560 | A/chicken/Scotland/078873/2022 | A / H5N1 | Europe / United Kingdom / Scotland | Alex Byrne (Animal and Plant Health Agency (APHA) / Virology Department) |
| EPI_ISL_13453565 | A/buzzard/Scotland/043118/2022 | A / H5N1 | Europe / United Kingdom / Scotland | Alex Byrne (Animal and Plant Health Agency (APHA) / Virology Department) |
| EPI_ISL_13453567 | A/turkey/England/081395/2022 | A / H5N1 | Europe / United Kingdom / England | Alex Byrne (Animal and Plant Health Agency (APHA) / Virology Department) |
| EPI_ISL_13453568 | A/turkey/England/081490/2022 | A / H5N1 | Europe / United Kingdom / England | Alex Byrne (Animal and Plant Health Agency (APHA) / Virology Department) |
| EPI_ISL_13465456 | A/Great white pelican/Israel/619/2021 | A / H5N1 | Asia / Israel / Central District / Mishmar HaSharon | Irina Shkoda (Kimron Veterinary Institute) |
| EPI_ISL_13477015 | A/chicken/Israel/88/2022 | A / H5N1 | Asia / Israel / Haifa District / Binyamina | Irina Shkoda (Kimron Veterinary Institute) |
| EPI_ISL_13486807 | A/Common_buzzard/Scotland/020817/2022 | A / H5N1 | Europe / United Kingdom / Scotland | Alex Byrne (Animal and Plant Health Agency (APHA) / Virology Department) |
| EPI_ISL_13486869 | A/Greylag_goose/Scotland/024915/2022 | A / H5N1 | Europe / United Kingdom / Scotland | Alex Byrne (Animal and Plant Health Agency (APHA) / Virology Department) |
| EPI_ISL_13514370 | A/Thalasseus sandvicensis/Belgium/7473_0009/2022 | A / H5N1 | Europe / Belgium / Provincie West-Vlaanderen / Oostende | Steven Van Borm (Sciensano, Department of Animal Infectious Diseases / Animal Infectious Diseases) |
| EPI_ISL_13514375 | A/Gallus gallus/Belgium/8092_0002/2022 | A / H5N1 | Europe / Belgium / Provincie Vlaams-Brabant / Zoutleeuw | Steven Van Borm (Sciensano, Department of Animal Infectious Diseases / Animal Infectious Diseases) |
| EPI_ISL_13519451 | A/gull/France/22P015977/2022 | A / H5N1 | Europe / France / Picardie / Departement de la Somme | Francois-Xavier Briand (ANSES Agence Nationale De Securite Sanitaire De L'alimentation / Laboratoire de Ploufragan-Plouzané) |
| EPI_ISL_13609610 | A/Great white pelican/Israel/123/2022 | A / H5N1 | Asia / Israel / Haifa District / Ma'ayan Tzvi | Irina Shkoda (Kimron Veterinary Institute) |
| EPI_ISL_13613219 | A/Greylag Goose/Netherlands/4/2022 | A / H5N1 | Europe / Netherlands / Obdam | Sanne Thewessen (Erasmus Medical Center / Viroscience) |
| EPI_ISL_13613220 | A/Greylag Goose/Netherlands/5/2022 | A / H5N1 | Europe / Netherlands / Obdam | Sanne Thewessen (Erasmus Medical Center / Viroscience) |
| EPI_ISL_13613221 | A/Greylag Goose/Netherlands/6/2022 | A / H5N1 | Europe / Netherlands / Obdam | Sanne Thewessen (Erasmus Medical Center / Viroscience) |
| EPI_ISL_13613222 | A/Greylag Goose/Netherlands/7/2022 | A / H5N1 | Europe / Netherlands / Obdam | Sanne Thewessen (Erasmus Medical Center / Viroscience) |
| EPI_ISL_13613223 | A/Greylag Goose/Netherlands/8/2022 | A / H5N1 | Europe / Netherlands / Obdam | Sanne Thewessen (Erasmus Medical Center / Viroscience) |
| EPI_ISL_13613224 | A/Common Tern/Netherlands/2/2022 | A / H5N1 | Europe / Netherlands / De Kreupel, Enkhuizen | Sanne Thewessen (Erasmus Medical Center / Viroscience) |
| EPI_ISL_13613225 | A/Black-headed gull/Netherlands/7/2022 | A / H5N1 | Europe / Netherlands / De Kreupel, Enkhuizen | Sanne Thewessen (Erasmus Medical Center / Viroscience) |
| EPI_ISL_13613226 | A/Black-headed gull/Netherlands/8/2022 | A / H5N1 | Europe / Netherlands / De Kreupel, Enkhuizen | Sanne Thewessen (Erasmus Medical Center / Viroscience) |
| EPI_ISL_13613227 | A/Black-headed gull/Netherlands/9/2022 | A / H5N1 | Europe / Netherlands / De Kreupel, Enkhuizen | Sanne Thewessen (Erasmus Medical Center / Viroscience) |
| EPI_ISL_13613228 | A/Sandwich Tern/Netherlands/9/2022 | A / H5N1 | Europe / Netherlands / Den Oever, Waddenhaven | Sanne Thewessen (Erasmus Medical Center / Viroscience) |
| EPI_ISL_13613229 | A/Sandwich Tern/Netherlands/10/2022 | A / H5N1 | Europe / Netherlands / Den Oever, Waddenhaven | Sanne Thewessen (Erasmus Medical Center / Viroscience) |
| EPI_ISL_13613230 | A/Common Tern/Netherlands/3/2022 | A / H5N1 | Europe / Netherlands / Wieringerwerf, Noorderdijkweg | Sanne Thewessen (Erasmus Medical Center / Viroscience) |
| EPI_ISL_13618384 | A/duck/Israel/589/2021 | A / H5N1 | Asia / Israel / Northern District / Kfar Baruch | Irina Shkoda (Kimron Veterinary Institute) |
| EPI_ISL_13632084 | A/chicken/Kazakhstan/23/2020 | A / H5N8 | Asia / Kazakhstan / Almaty Province | Kairat Tabynov (Kazakh National Agrarian University / International Center for Vaccinology) |
| EPI_ISL_13658359 | A/common kestel/Israel/49-2/2022 | A / H5N1 | Asia / Israel / Northern District / Agamon Hula | Irina Shkoda (Kimron Veterinary Institute) |
| EPI_ISL_13692620 | A/turkey/Stavropol/211-1V/2022 | A / H5N1 | Europe / Russian Federation / Stavropol Krai | Natalia Goncharova (State Research Center of Virology and Biotechnology (VECTOR) / Emerging Zoonotic Diseases and Influenza) |
| EPI_ISL_13692621 | A/turkey/Stavropol/211-4V/2022 | A / H5N1 | Europe / Russian Federation / Stavropol Krai | Natalia Goncharova (State Research Center of Virology and Biotechnology (VECTOR) / Emerging Zoonotic Diseases and Influenza) |
| EPI_ISL_13692622 | A/turkey/Stavropol/211-7V/2022 | A / H5N1 | Europe / Russian Federation / Stavropol Krai | Natalia Goncharova (State Research Center of Virology and Biotechnology (VECTOR) / Emerging Zoonotic Diseases and Influenza) |
| EPI_ISL_13692623 | A/turkey/Stavropol/211-12V/2022 | A / H5N1 | Europe / Russian Federation / Stavropol Krai | Natalia Goncharova (State Research Center of Virology and Biotechnology (VECTOR) / Emerging Zoonotic Diseases and Influenza) |
| EPI_ISL_13692625 | A/turkey/Stavropol/211-15V/2022 | A / H5N1 | Europe / Russian Federation / Stavropol Krai | Natalia Goncharova (State Research Center of Virology and Biotechnology (VECTOR) / Emerging Zoonotic Diseases and Influenza) |
| EPI_ISL_13692626 | A/turkey/Stavropol/211-18V/2022 | A / H5N1 | Europe / Russian Federation / Stavropol Krai | Natalia Goncharova (State Research Center of Virology and Biotechnology (VECTOR) / Emerging Zoonotic Diseases and Influenza) |
| EPI_ISL_13692628 | A/turkey/Stavropol/211-5V/2022 | A / H5N1 | Europe / Russian Federation / Stavropol Krai | Natalia Goncharova (State Research Center of Virology and Biotechnology (VECTOR) / Emerging Zoonotic Diseases and Influenza) |
| EPI_ISL_13692629 | A/turkey/Stavropol/211-9V/2022 | A / H5N1 | Europe / Russian Federation / Stavropol Krai | Natalia Goncharova (State Research Center of Virology and Biotechnology (VECTOR) / Emerging Zoonotic Diseases and Influenza) |
| EPI_ISL_13692630 | A/crow/Khabarovsk/216-11V/2022 | A / H5N1 | Europe / Russian Federation / Khabarovsk Krai | Natalia Goncharova (State Research Center of Virology and Biotechnology (VECTOR) / Emerging Zoonotic Diseases and Influenza) |
| EPI_ISL_13692632 | A/crow/Khabarovsk/216-12V/2022 | A / H5N1 | Europe / Russian Federation / Khabarovsk Krai | Natalia Goncharova (State Research Center of Virology and Biotechnology (VECTOR) / Emerging Zoonotic Diseases and Influenza) |
| EPI_ISL_13692634 | A/crow/Khabarovsk/216-13V/2022 | A / H5N1 | Europe / Russian Federation / Khabarovsk Krai | Natalia Goncharova (State Research Center of Virology and Biotechnology (VECTOR) / Emerging Zoonotic Diseases and Influenza) |
| EPI_ISL_13692637 | A/dalmatian pelican/Astrakhan/213-1V/2022 | A / H5N1 | Europe / Russian Federation / Astrakhan Oblast | Natalia Goncharova (State Research Center of Virology and Biotechnology (VECTOR) / Emerging Zoonotic Diseases and Influenza) |
| EPI_ISL_13692638 | A/dalmatian pelican/Astrakhan/213-2V/2022 | A / H5N1 | Europe / Russian Federation / Astrakhan Oblast | Natalia Goncharova (State Research Center of Virology and Biotechnology (VECTOR) / Emerging Zoonotic Diseases and Influenza) |
| EPI_ISL_137254 | A/duck/Hebei/2/2011 | A / H5N2 | Asia / China / Hebei Province | Di Liu (Institute of Microbiology, Chinese Academy of Sciences) |
| EPI_ISL_137255 | A/duck/Hebei/3/2011 | A / H5N2 | Asia / China / Hebei Province | Di Liu (Institute of Microbiology, Chinese Academy of Sciences) |
| EPI_ISL_13778462 | A/Lesser Black-backed Gull/Netherlands/2/2022 | A / H5N1 | Europe / Netherlands / Enkhuizen, Ijsselmeer, De Kreupel | Sanne Thewessen (Erasmus Medical Center / Viroscience) |
| EPI_ISL_13778464 | A/European Herring Gull/Netherlands/4/2022 | A / H5N1 | Europe / Netherlands / Den Oever, Waddenhaven | Sanne Thewessen (Erasmus Medical Center / Viroscience) |
| EPI_ISL_13778466 | A/Oystercatcher/Netherlands/2/2022 | A / H5N1 | Europe / Netherlands / Den Oever, Waddenhaven | Sanne Thewessen (Erasmus Medical Center / Viroscience) |
| EPI_ISL_13778467 | A/Sandwich Tern/Netherlands/11/2022 | A / H5N1 | Europe / Netherlands / Den Oever, Waddenhaven | Sanne Thewessen (Erasmus Medical Center / Viroscience) |
| EPI_ISL_13778468 | A/European Herring Gull/Netherlands/5/2022 | A / H5N1 | Europe / Netherlands / Walcheren, 't Vroon Westkapelle | Sanne Thewessen (Erasmus Medical Center / Viroscience) |
| EPI_ISL_13778469 | A/European Herring Gull/Netherlands/6/2022 | A / H5N1 | Europe / Netherlands / Walcheren, 't Vroon Westkapelle | Sanne Thewessen (Erasmus Medical Center / Viroscience) |
| EPI_ISL_13778474 | A/European Herring Gull/Netherlands/7/2022 | A / H5N1 | Europe / Netherlands / Walcheren, 't Vroon Westkapelle | Sanne Thewessen (Erasmus Medical Center / Viroscience) |
| EPI_ISL_13778475 | A/Sandwich Tern/Netherlands/12/2022 | A / H5N1 | Europe / Netherlands / Walcheren, 't Vroon Westkapelle | Sanne Thewessen (Erasmus Medical Center / Viroscience) |
| EPI_ISL_13778476 | A/Sandwich Tern/Netherlands/13/2022 | A / H5N1 | Europe / Netherlands / Walcheren, 't Vroon Westkapelle | Sanne Thewessen (Erasmus Medical Center / Viroscience) |
| EPI_ISL_13778477 | A/Common Tern/Netherlands/8/2022 | A / H5N1 | Europe / Netherlands / Haringvliet, Ventjagersplaten, Den Bommel | Sanne Thewessen (Erasmus Medical Center / Viroscience) |
| EPI_ISL_13778478 | A/Sandwich Tern/Netherlands/14/2022 | A / H5N1 | Europe / Netherlands / Walcheren, 't Vroon Westkapelle | Sanne Thewessen (Erasmus Medical Center / Viroscience) |
| EPI_ISL_13778479 | A/Common Tern/Netherlands/9/2022 | A / H5N1 | Europe / Netherlands / Walcheren, 't Vroon Westkapelle | Sanne Thewessen (Erasmus Medical Center / Viroscience) |
| EPI_ISL_13778522 | A/Sandwich Tern/Netherlands/15/2022 | A / H5N1 | Europe / Netherlands / Walcheren, 't Vroon Westkapelle | Sanne Thewessen (Erasmus Medical Center / Viroscience) |
| EPI_ISL_13778523 | A/Common Tern/Netherlands/4/2022 | A / H5N1 | Europe / Netherlands / Wieringerwerf, Noorderdijkweg | Sanne Thewessen (Erasmus Medical Center / Viroscience) |
| EPI_ISL_13778524 | A/Common Tern/Netherlands/5/2022 | A / H5N1 | Europe / Netherlands / Wieringerwerf, Noorderdijkweg | Sanne Thewessen (Erasmus Medical Center / Viroscience) |
| EPI_ISL_13778525 | A/Common Tern/Netherlands/6/2022 | A / H5N1 | Europe / Netherlands / Wieringerwerf, Noorderdijkweg | Sanne Thewessen (Erasmus Medical Center / Viroscience) |
| EPI_ISL_13778526 | A/Common Tern/Netherlands/7/2022 | A / H5N1 | Europe / Netherlands / Wieringerwerf, Noorderdijkweg | Sanne Thewessen (Erasmus Medical Center / Viroscience) |

| | | | | |
|---|---|---|---|---|
| EPI_ISL_1379443 | A/chicken/Vietnam/Raho4-Cd-20-421/2020 | A / H5N6 | Asia / Vietnam | Yunho Jang (Centers for Disease Control and Prevention / WHO Collaborating Center for Surveillance, Epidemiology and Control of Infl |
| EPI_ISL_1385792 | A/Chicken/Sweden/SVA210323SZ001/KN001982-IP21/2021 | A / H5N5 | Europe / Sweden / Stockholms Lan / Ekero Kommun | Siamak Zohari (National Veterinary Institute) |
| EPI_ISL_1386128 | A/Chiken/Sweden/SVA210321SZ0001/KN073551-IP20/2021 | A / H5N8 | Europe / Sweden / Skane Lan / Hassleholms Kommun | Siamak Zohari (National Veterinary Institute) |
| EPI_ISL_1386249 | A/western marsh harrier/Sweden/SVA210316SZ0486/KN0010 | A / H5N8 | Europe / Sweden / Ostergotlands Lan / Linkopings Kommun | Siamak Zohari (National Veterinary Institute) |
| EPI_ISL_13876272 | A/chicken/Ryazan/224-1V/2022 | A / H5N1 | Europe / Russian Federation / Ryazan Oblast | Natalia Goncharova (State Research Center of Virology and Biotechnology (VECTOR) / Emerging Zoonotic Diseases and Influenza) |
| EPI_ISL_1389398 | A/northern goshawk/Sweden/SVA210316SZ0489/KN001044/N | A / H5N8 | Europe / Sweden / Skane Lan / Kavlinge Kommun | Siamak Zohari (National Veterinary Institute) |
| EPI_ISL_13902858 | A/Eurasian Curlew/Netherlands/2/2022 | A / H5N1 | Europe / Netherlands / Den Oever, Waddenhaven | Sanne Thewessen (Erasmus Medical Center / Viroscience) |
| EPI_ISL_1390546 | A/northern goshawk/Sweden/SVA210316SZ0489/KN001045/N | A / H5N8 | Europe / Sweden / Skane Lan / Lomma Kommun | Siamak Zohari (National Veterinary Institute) |
| EPI_ISL_1391019 | A/common golden eye/Sweden/SVA210316SZ0489/KN001046 | A / H5N5 | Europe / Sweden / Skane Lan / Lomma Kommun | Siamak Zohari (National Veterinary Institute) |
| EPI_ISL_1391276 | A/northern goshawk/Sweden/SVA210318SZ0322/KN001092/N | A / H5N5 | Europe / Sweden / Skane Lan / Simrishamns Kommun | Siamak Zohari (National Veterinary Institute) |
| EPI_ISL_139383 | A/goose/Shandong/k1204/2009 | A / H5N5 | Asia / China | |
| EPI_ISL_139384 | A/goose/Guangdong/k0103/2010 | A / H5N5 | Asia / China | |
| EPI_ISL_139385 | A/duck/Jiangsu/k1203/2010 | A / H5N8 | Asia / China | |
| EPI_ISL_139386 | A/quail/Jiangsu/k0104/2010 | A / H5N5 | Asia / China | |
| EPI_ISL_13955171 | A/chicken/Czech Republic/2968/2022 | A / H5N1 | Europe / Czech Republic / Plzensky Kraj / Okres Plzen-Jih / Lišice, Dolní Lukavice, ZipCode: 629 71 | Alexander Nagy (State Veterinary Institute Prague) |
| EPI_ISL_13955204 | A/chicken/Czech Republic/8028-1/2022 | A / H5N1 | Europe / Czech Republic / Plzensky Kraj / Okres Klatovy / Plánice | Alexander Nagy (State Veterinary Institute Prague) |
| EPI_ISL_13957819 | A/chicken/Mali/T1-177_22VIR6104-1/2022 | A / H5N1 | Africa / Mali / Kati | Bianca Zecchin (Istituto Zooprofilattico Sperimentale Delle Venezie) |
| EPI_ISL_13957821 | A/chicken/Mali/T2-178_22VIR6104-3/2022 | A / H5N1 | Africa / Mali / Kati | Bianca Zecchin (Istituto Zooprofilattico Sperimentale Delle Venezie) |
| EPI_ISL_13957822 | A/chicken/Mali/S3-179_22VIR6104-5/2022 | A / H5N1 | Africa / Mali / Kati | Bianca Zecchin (Istituto Zooprofilattico Sperimentale Delle Venezie) |
| EPI_ISL_13957823 | A/chicken/Mali/T4_180_22VIR6104-7/2022 | A / H5N1 | Africa / Mali / Kati | Bianca Zecchin (Istituto Zooprofilattico Sperimentale Delle Venezie) |
| EPI_ISL_13967740 | A/chicken/Czech Republic/8028-2/2022 | A / H5N1 | Europe / Czech Republic / Plzensky Kraj / Okres Klatovy / Plánice | Alexander Nagy (State Veterinary Institute Prague) |
| EPI_ISL_13969422 | A/Goose/Scotland/036879/2022 | A / H5N1 | Europe / United Kingdom / Scotland | Alex Byrne (Animal and Plant Health Agency (APHA) / Virology Department) |
| EPI_ISL_13969423 | A/Gull/Scotland/060376/2022 | A / H5N1 | Europe / United Kingdom / Scotland | Alex Byrne (Animal and Plant Health Agency (APHA) / Virology Department) |
| EPI_ISL_13969424 | A/Gannet/Scotland/084490/2022 | A / H5N1 | Europe / United Kingdom / Scotland | Alex Byrne (Animal and Plant Health Agency (APHA) / Virology Department) |
| EPI_ISL_13969426 | A/Gannet/Scotland/090910/2022 | A / H5N1 | Europe / United Kingdom / Scotland | Alex Byrne (Animal and Plant Health Agency (APHA) / Virology Department) |
| EPI_ISL_13969428 | A/Greylag_goose/England/247696/2022 | A / H5N1 | Europe / United Kingdom / England | Alex Byrne (Animal and Plant Health Agency (APHA) / Virology Department) |
| EPI_ISL_13969430 | A/Mute_swan/Wales/058560/2022 | A / H5N1 | Europe / United Kingdom / Wales | Alex Byrne (Animal and Plant Health Agency (APHA) / Virology Department) |
| EPI_ISL_13990713 | A/Anser_anser/Spain/1277-10_22VIR6312-10/2022 | A / H5N1 | Europe / Spain | Giacomo Barbierato (Istituto Zooprofilattico Sperimentale delle Venezie / Ricerca e innovazione) |
| EPI_ISL_13990714 | A/wild_bird/Spain/1302-1_22VIR6312-11/2022 | A / H5N1 | Europe / Spain | Giacomo Barbierato (Istituto Zooprofilattico Sperimentale delle Venezie / Ricerca e innovazione) |
| EPI_ISL_13990715 | A/Cygnus_olor/Spain/1675-1_22VIR6312-14/2022 | A / H5N1 | Europe / Spain | Giacomo Barbierato (Istituto Zooprofilattico Sperimentale delle Venezie / Ricerca e innovazione) |
| EPI_ISL_13990716 | A/Rhea/Spain/1950-6_22VIR6312-17/2022 | A / H5N1 | Europe / Spain | Giacomo Barbierato (Istituto Zooprofilattico Sperimentale delle Venezie / Ricerca e innovazione) |
| EPI_ISL_13990717 | A/Gypaetus_barbatus/Spain/1878-9_22VIR6312-18/2022 | A / H5N1 | Europe / Spain | Giacomo Barbierato (Istituto Zooprofilattico Sperimentale delle Venezie / Ricerca e innovazione) |
| EPI_ISL_13990718 | A/Gypaetus_barbatus/Spain/1956-25_22VIR6312-19/2022 | A / H5N1 | Europe / Spain | Giacomo Barbierato (Istituto Zooprofilattico Sperimentale delle Venezie / Ricerca e innovazione) |
| EPI_ISL_13990719 | A/Falco_peregrinus/Spain/2010-11_22VIR6312-20/2022 | A / H5N1 | Europe / Spain | Giacomo Barbierato (Istituto Zooprofilattico Sperimentale delle Venezie / Ricerca e innovazione) |
| EPI_ISL_13990720 | A/Anser_anser/Spain/2010-3_22VIR6312-21/2022 | A / H5N1 | Europe / Spain | Giacomo Barbierato (Istituto Zooprofilattico Sperimentale delle Venezie / Ricerca e innovazione) |
| EPI_ISL_13990721 | A/turkey/Spain/711-5_22VIR6312-23/2022 | A / H5N1 | Europe / Spain | Giacomo Barbierato (Istituto Zooprofilattico Sperimentale delle Venezie / Ricerca e innovazione) |
| EPI_ISL_13990722 | A/turkey/Spain/712-7_22VIR6312-24/2022 | A / H5N1 | Europe / Spain | Giacomo Barbierato (Istituto Zooprofilattico Sperimentale delle Venezie / Ricerca e innovazione) |
| EPI_ISL_13990723 | A/turkey/Spain/801-6_22VIR6312-25/2022 | A / H5N1 | Europe / Spain | Giacomo Barbierato (Istituto Zooprofilattico Sperimentale delle Venezie / Ricerca e innovazione) |
| EPI_ISL_13990724 | A/turkey/Spain/802-7_22VIR6312-26/2022 | A / H5N1 | Europe / Spain | Giacomo Barbierato (Istituto Zooprofilattico Sperimentale delle Venezie / Ricerca e innovazione) |
| EPI_ISL_13990725 | A/turkey/Spain/803-5_22VIR6312-27/2022 | A / H5N1 | Europe / Spain | Giacomo Barbierato (Istituto Zooprofilattico Sperimentale delle Venezie / Ricerca e innovazione) |
| EPI_ISL_13990726 | A/turkey/Spain/805-8_22VIR6312-28/2022 | A / H5N1 | Europe / Spain | Giacomo Barbierato (Istituto Zooprofilattico Sperimentale delle Venezie / Ricerca e innovazione) |
| EPI_ISL_13990727 | A/turkey/Spain/830-5_22VIR6312-29/2022 | A / H5N1 | Europe / Spain | Giacomo Barbierato (Istituto Zooprofilattico Sperimentale delle Venezie / Ricerca e innovazione) |
| EPI_ISL_13990728 | A/Anser_anser/Spain/750-4_22VIR6312-2/2022 | A / H5N1 | Europe / Spain | Giacomo Barbierato (Istituto Zooprofilattico Sperimentale delle Venezie / Ricerca e innovazione) |
| EPI_ISL_13990729 | A/turkey/Spain/859-8_22VIR6312-30/2022 | A / H5N1 | Europe / Spain | Giacomo Barbierato (Istituto Zooprofilattico Sperimentale delle Venezie / Ricerca e innovazione) |
| EPI_ISL_13990730 | A/chicken/Spain/897-7_22VIR6312-31/2022 | A / H5N1 | Europe / Spain | Giacomo Barbierato (Istituto Zooprofilattico Sperimentale delle Venezie / Ricerca e innovazione) |
| EPI_ISL_13990731 | A/chicken/Spain/899-7_22VIR6312-32/2022 | A / H5N1 | Europe / Spain | Giacomo Barbierato (Istituto Zooprofilattico Sperimentale delle Venezie / Ricerca e innovazione) |
| EPI_ISL_13990732 | A/chicken/Spain/924-3_22VIR6312-33/2022 | A / H5N1 | Europe / Spain | Giacomo Barbierato (Istituto Zooprofilattico Sperimentale delle Venezie / Ricerca e innovazione) |
| EPI_ISL_13990733 | A/chicken/Spain/942-8_22VIR6312-34/2022 | A / H5N1 | Europe / Spain | Giacomo Barbierato (Istituto Zooprofilattico Sperimentale delle Venezie / Ricerca e innovazione) |
| EPI_ISL_13990734 | A/chicken/Spain/1096-8_22VIR6312-35/2022 | A / H5N1 | Europe / Spain | Giacomo Barbierato (Istituto Zooprofilattico Sperimentale delle Venezie / Ricerca e innovazione) |
| EPI_ISL_13990735 | A/turkey/Spain/1097-7_22VIR6312-36/2022 | A / H5N1 | Europe / Spain | Giacomo Barbierato (Istituto Zooprofilattico Sperimentale delle Venezie / Ricerca e innovazione) |
| EPI_ISL_13990736 | A/chicken/Spain/340-37_22VIR6312-37/2022 | A / H5N1 | Europe / Spain | Giacomo Barbierato (Istituto Zooprofilattico Sperimentale delle Venezie / Ricerca e innovazione) |
| EPI_ISL_13990737 | A/Anser_anser/Spain/638-6_22VIR6312-39/2022 | A / H5N1 | Europe / Spain | Giacomo Barbierato (Istituto Zooprofilattico Sperimentale delle Venezie / Ricerca e innovazione) |
| EPI_ISL_13990738 | A/chicken/Spain/806-7_22VIR6312-41/2022 | A / H5N1 | Europe / Spain | Giacomo Barbierato (Istituto Zooprofilattico Sperimentale delle Venezie / Ricerca e innovazione) |
| EPI_ISL_13990739 | A/Ciconia_ciconia/Spain/853-14_22VIR6312-42/2022 | A / H5N1 | Europe / Spain | Giacomo Barbierato (Istituto Zooprofilattico Sperimentale delle Venezie / Ricerca e innovazione) |
| EPI_ISL_13990740 | A/Falco_peregrinus/Spain/1932-1_22VIR6312-45/2022 | A / H5N1 | Europe / Spain | Giacomo Barbierato (Istituto Zooprofilattico Sperimentale delle Venezie / Ricerca e innovazione) |
| EPI_ISL_13990741 | A/Cygnus_olor/Spain/1950-10_22VIR6312-46/2022 | A / H5N1 | Europe / Spain | Giacomo Barbierato (Istituto Zooprofilattico Sperimentale delle Venezie / Ricerca e innovazione) |
| EPI_ISL_13990743 | A/Ardea_cinerea/Spain/863-1_22VIR6312-5/2022 | A / H5N1 | Europe / Spain | Giacomo Barbierato (Istituto Zooprofilattico Sperimentale delle Venezie / Ricerca e innovazione) |
| EPI_ISL_13990744 | A/Anser_anser/Spain/863-2_22VIR6312-6/2022 | A / H5N1 | Europe / Spain | Giacomo Barbierato (Istituto Zooprofilattico Sperimentale delle Venezie / Ricerca e innovazione) |
| EPI_ISL_13990745 | A/Ciconia_ciconia/Spain/971-6_22VIR6312-7/2022 | A / H5N1 | Europe / Spain | Giacomo Barbierato (Istituto Zooprofilattico Sperimentale delle Venezie / Ricerca e innovazione) |
| EPI_ISL_13990746 | A/Anser_anser/Spain/1035-5_22VIR6312-8/2022 | A / H5N1 | Europe / Spain | Giacomo Barbierato (Istituto Zooprofilattico Sperimentale delle Venezie / Ricerca e innovazione) |
| EPI_ISL_13990747 | A/Anser_anser/Spain/1124-65_22VIR6312-9/2022 | A / H5N1 | Europe / Spain | Giacomo Barbierato (Istituto Zooprofilattico Sperimentale delle Venezie / Ricerca e innovazione) |
| EPI_ISL_1399231 | A/chicken/Czech Republic/3531-1/2021 | A / H5N8 | Europe / Czech Republic / Vysocina Kraj / Okres Pelhrimov / Horni Cerekev; GPS: N 49°19.49288' | Alexander Nagy (State Veterinary Institute Prague) |
| EPI_ISL_1399232 | A/chicken/Czech Republic/3893/2021 | A / H5N8 | Europe / Czech Republic / Jihocesky Kraj / Okres Strakonice / Rojice | Alexander Nagy (State Veterinary Institute Prague) |
| EPI_ISL_1399233 | A/chicken/Czech Republic/4092-1/2021 | A / H5N8 | Europe / Czech Republic / Plzensky Kraj / Okres Klatovy / Hnevnice; GPS: N 49°44.08495', E 13°6. | Alexander Nagy (State Veterinary Institute Prague) |
| EPI_ISL_1399234 | A/chicken/Czech Republic/4092-2/2021 | A / H5N8 | Europe / Czech Republic / Plzensky Kraj / Okres Klatovy / Hnevnice; GPS: N 49°44.08495', E 13°6. | Alexander Nagy (State Veterinary Institute Prague) |
| EPI_ISL_1399235 | A/chicken/Czech Republic/4527-1/2021 | A / H5N8 | Europe / Czech Republic / Stredocesky Kraj / Okres Mlada Boleslav / Brezno | Alexander Nagy (State Veterinary Institute Prague) |
| EPI_ISL_1399236 | A/chicken/Czech Republic/4756/2021 | A / H5N8 | Europe / Czech Republic / Stredocesky Kraj / Okres Pribram / Pocaply, Breznice | Alexander Nagy (State Veterinary Institute Prague) |
| EPI_ISL_1399237 | A/mute swan/Czech Republic/3549/2021 | A / H5N8 | Europe / Czech Republic / Stredocesky Kraj / Okres Nymburk / hydroelectric power Nymburk; GF | Alexander Nagy (State Veterinary Institute Prague) |
| EPI_ISL_1399238 | A/mute swan/Czech Republic/3777/2021 | A / H5N8 | Europe / Czech Republic / Hlavni mesto Praha / Okres Praha / Kvetnice, Mlynsky rybnik; GPS: N 5 | Alexander Nagy (State Veterinary Institute Prague) |
| EPI_ISL_1399239 | A/mute swan/Czech Republic/4100/2021 | A / H5N8 | Europe / Czech Republic / Olomoucky Kraj / Okres Jesenik / Javornik; GPS: 50°24'35.8''N, 17°2'44 | Alexander Nagy (State Veterinary Institute Prague) |
| EPI_ISL_1399240 | A/mute swan/Czech Republic/4270/2021 | A / H5N8 | Europe / Czech Republic / Stredocesky Kraj / Okres Mlada Boleslav / rybnik Ohrada, Bosen | Alexander Nagy (State Veterinary Institute Prague) |
| EPI_ISL_1399241 | A/chicken/Czech Republic/4526/2021 | A / H5N8 | Europe / Czech Republic / Morovskoslezsky Kraj / Okres Opava / Vyhon, Velke Hostice; GPS: N 4 | Alexander Nagy (State Veterinary Institute Prague) |

| | | | | |
|---|---|---|---|---|
| EPI_ISL_1399242 | A/mute swan/Czech Republic/4606/2021 | A / H5N8 | Europe / Czech Republic / Zlinsky Kraj / Okres Uherske Hradiste / Ostrozska jezera, Ostrozska No | Alexander Nagy (State Veterinary Institute Prague) |
| EPI_ISL_1399243 | A/mute swan/Czech Republic/4607-1/2021 | A / H5N8 | Europe / Czech Republic / Olomoucky Kraj / Okres Olomouc / Chomoutov | Alexander Nagy (State Veterinary Institute Prague) |
| EPI_ISL_1399244 | A/mute swan/Czech Republic/4607-2/2021 | A / H5N8 | Europe / Czech Republic / Olomoucky Kraj / Okres Olomouc / Chomoutov | Alexander Nagy (State Veterinary Institute Prague) |
| EPI_ISL_1399245 | A/mute swan/Czech Republic/4799/2021 | A / H5N8 | Europe / Czech Republic / Morovskoslezsky Kraj / Okres Opava / Katerinky u Opavy, Stribrne jeze | Alexander Nagy (State Veterinary Institute Prague) |
| EPI_ISL_14028104 | A/Ciconia_ciconia/Spain/485-2_22VIR6312-1/2022 | A / H5N1 | Europe / Spain | Giacomo Barbierato (Istituto Zooprofilattico Sperimentale delle Venezie / Ricerca e innovazione) |
| EPI_ISL_14036187 | A/chicken/Egypt/HS.1/2022 | A / H5N8 | Africa / Egypt / Monufia / 35 days chicken | Hazem Salah Khalil (Animal Health Research Institute (Egypt) / Poultry Diseases) |
| EPI_ISL_14036311 | A/Duck/Egypt/HS.2/2022 | A / H5N8 | Africa / Egypt / Monufia | Hazem Salah Khalil (Animal Health Research Institute (Egypt) / Poultry Diseases) |
| EPI_ISL_14036428 | A/Quail/Egypt/HS.3/2022 | A / H5N8 | Africa / Egypt / Monufia | Hazem Salah Khalil (Animal Health Research Institute (Egypt) / Poultry Diseases) |
| EPI_ISL_14037832 | A/chicken/Egypt/HS.4/2022 | A / H5N8 | Africa / Egypt | Hazem Salah Khalil (Animal Health Research Institute (Egypt) / Poultry Diseases) |
| EPI_ISL_14037833 | A/Duck/Egypt/HS.5/2022 | A / H5N8 | Africa / Egypt | Hazem Salah Khalil (Animal Health Research Institute (Egypt) / Poultry Diseases) |
| EPI_ISL_14037834 | A/Quail/Egypt/HS.6/2022 | A / H5N8 | Africa / Egypt | Hazem Salah Khalil (Animal Health Research Institute (Egypt) / Poultry Diseases) |
| EPI_ISL_14064695 | A/crow/Hokkaido/0101Q045/2022 | A / H5N1 | Asia / Japan / Hokkaido / Nemuro | Norikazu Isoda (Graduate School of Veterinary Medicine, Hokkaido University / Laboratory of Microbiology) |
| EPI_ISL_14064697 | A/crow/Hokkaido/0101Q044/2022 (H5N1) | A / H5N1 | Asia / Japan / Hokkaido / Nemuro | Norikazu Isoda (Graduate School of Veterinary Medicine, Hokkaido University / Laboratory of Microbiology) |
| EPI_ISL_14064698 | A/crow/Hokkaido/0101Q054/2022 | A / H5N1 | Asia / Japan / Hokkaido / Nemuro | Norikazu Isoda (Graduate School of Veterinary Medicine, Hokkaido University / Laboratory of Microbiology) |
| EPI_ISL_14064699 | A/crow/Hokkaido/0102F046/2022 | A / H5N1 | Asia / Japan / Hokkaido / Erimo | Norikazu Isoda (Graduate School of Veterinary Medicine, Hokkaido University / Laboratory of Microbiology) |
| EPI_ISL_14064768 | A/crow/Hokkaido/0102F048/2022 | A / H5N1 | Asia / Japan / Hokkaido / Erimo | Norikazu Isoda (Graduate School of Veterinary Medicine, Hokkaido University / Laboratory of Microbiology) |
| EPI_ISL_14064803 | A/crow/Hokkaido/0102F043/2022 | A / H5N1 | Asia / Japan / Hokkaido / Erimo | Norikazu Isoda (Graduate School of Veterinary Medicine, Hokkaido University / Laboratory of Microbiology) |
| EPI_ISL_14064890 | A/crow/Hokkaido/0103B073/2022 (H5N1) | A / H5N1 | Asia / Japan / Hokkaido / Sapporo | Norikazu Isoda (Graduate School of Veterinary Medicine, Hokkaido University / Laboratory of Microbiology) |
| EPI_ISL_14064983 | A/chicken/Scotland/093091/2022 | A / H5N1 | Europe / United Kingdom / Scotland | Alex Byrne (Animal and Plant Health Agency (APHA) / Virology Department) |
| EPI_ISL_14064984 | A/chicken/England/093459/2022 | A / H5N1 | Europe / United Kingdom / England | Alex Byrne (Animal and Plant Health Agency (APHA) / Virology Department) |
| EPI_ISL_14064985 | A/Domestic_goose/England/093469/2022 | A / H5N1 | Europe / United Kingdom / England | Alex Byrne (Animal and Plant Health Agency (APHA) / Virology Department) |
| EPI_ISL_14064986 | A/crow/Hokkaido/0102L010/2022 | A / H5N1 | Asia / Japan / Hokkaido / Rishirifuji | Norikazu Isoda (Graduate School of Veterinary Medicine, Hokkaido University / Laboratory of Microbiology) |
| EPI_ISL_14064988 | A/crow/Hokkaido/0102M086/2022 | A / H5N1 | Asia / Japan / Hokkaido / Syari | Norikazu Isoda (Graduate School of Veterinary Medicine, Hokkaido University / Laboratory of Microbiology) |
| EPI_ISL_14065167 | A/white-tailed eagle/Hokkaido/20220210001/2022 | A / H5N1 | Asia / Japan / Hokkaido / Rausu | Norikazu Isoda (Graduate School of Veterinary Medicine, Hokkaido University / Laboratory of Microbiology) |
| EPI_ISL_14070455 | A/tanuki/Hokkaido/1/2022 | A / H5N1 | Asia / Japan / Hokkaido / Sapporo | Norikazu Isoda (Graduate School of Veterinary Medicine, Hokkaido University / Laboratory of Microbiology) |
| EPI_ISL_14098915 | A/harbor seal/Maine/22-020455-001-original/2022 | A / H5N1 | North America / United States / Maine / Cumberland County | Mary Lea Killian (National Veterinary Services Laboratories - USDA / Diagnostic Virology Laboratory) |
| EPI_ISL_14098916 | A/harbor seal/Maine/22-020455-002-original/2022 | A / H5N1 | North America / United States / Maine / Cumberland County | Mary Lea Killian (National Veterinary Services Laboratories - USDA / Diagnostic Virology Laboratory) |
| EPI_ISL_14098917 | A/harbor seal/Maine/22-020455-003-original/2022 | A / H5N1 | North America / United States / Maine / York County | Mary Lea Killian (National Veterinary Services Laboratories - USDA / Diagnostic Virology Laboratory) |
| EPI_ISL_14098918 | A/harbor seal/Maine/22-020455-004-original/2022 | A / H5N1 | North America / United States / Maine / Sagadahoc County | Mary Lea Killian (National Veterinary Services Laboratories - USDA / Diagnostic Virology Laboratory) |
| EPI_ISL_14098919 | A/harbor seal/Maine/22-020455-005-original/2022 | A / H5N1 | North America / United States / Maine / Lincoln County | Mary Lea Killian (National Veterinary Services Laboratories - USDA / Diagnostic Virology Laboratory) |
| EPI_ISL_14098920 | A/harbor seal/Maine/22-020983-001-original/2022 | A / H5N1 | North America / United States / Maine / Cumberland County | Mary Lea Killian (National Veterinary Services Laboratories - USDA / Diagnostic Virology Laboratory) |
| EPI_ISL_14098921 | A/harbor seal/Maine/22-020983-002-original/2022 | A / H5N1 | North America / United States / Maine / Cumberland County | Mary Lea Killian (National Veterinary Services Laboratories - USDA / Diagnostic Virology Laboratory) |
| EPI_ISL_14098922 | A/grey seal/Maine/22-020983-003-original/2022 | A / H5N1 | North America / United States / Maine / Sagadahoc County | Mary Lea Killian (National Veterinary Services Laboratories - USDA / Diagnostic Virology Laboratory) |
| EPI_ISL_14098923 | A/harbor seal/Maine/22-020983-006-original/2022 | A / H5N1 | North America / United States / Maine / Cumberland County | Mary Lea Killian (National Veterinary Services Laboratories - USDA / Diagnostic Virology Laboratory) |
| EPI_ISL_14098924 | A/harbor seal/Maine/22-020983-007-original/2022 | A / H5N1 | North America / United States / Maine / Penobscot County / Plymouth County | Mary Lea Killian (National Veterinary Services Laboratories - USDA / Diagnostic Virology Laboratory) |
| EPI_ISL_14163711 | A/Common Tern/Netherlands/10/2022 | A / H5N1 | Europe / Netherlands / Noordburen, Waddenkust | Sanne Thewessen (Erasmus Medical Center / Viroscience) |
| EPI_ISL_14163712 | A/European Herring Gull/Netherlands/8/2022 | A / H5N1 | Europe / Netherlands / Noordburen, Waddenkust | Sanne Thewessen (Erasmus Medical Center / Viroscience) |
| EPI_ISL_14163713 | A/Caspian Gull/Netherlands/5/2022 | A / H5N1 | Europe / Netherlands / Enkhuizen, Ijsselmeer, De Kreupel | Sanne Thewessen (Erasmus Medical Center / Viroscience) |
| EPI_ISL_14163714 | A/European Herring Gull/Netherlands/9/2022 | A / H5N1 | Europe / Netherlands / Ijmuiden, Forteiland | Sanne Thewessen (Erasmus Medical Center / Viroscience) |
| EPI_ISL_14163715 | A/Greylag Goose/Netherlands/9/2022 | A / H5N1 | Europe / Netherlands / Inlaag, Neeltje Jans | Sanne Thewessen (Erasmus Medical Center / Viroscience) |
| EPI_ISL_14171729 | A/Eurasian Spoonbill/Netherlands/1A/2022 | A / H5N1 | Europe / Netherlands / Schiermonnikoog | Sanne Thewessen (Erasmus Medical Center / Viroscience) |
| EPI_ISL_14171741 | A/Eurasian Spoonbill/Netherlands/1B/2022 | A / H5N1 | Europe / Netherlands / Schiermonnikoog | Sanne Thewessen (Erasmus Medical Center / Viroscience) |
| EPI_ISL_14174187 | A/chicken/China/007/2019 | A / H5N6 | Asia / China | |
| EPI_ISL_14174306 | A/goose/China/0701/2018 | A / H5N6 | Asia / China | |
| EPI_ISL_14174436 | A/mink/China/181/2019 | A / H5N6 | Asia / China | |
| EPI_ISL_14174437 | A/mink/China/183/2019 | A / H5N6 | Asia / China | |
| EPI_ISL_14174476 | A/mink/China/191/2019 | A / H5N6 | Asia / China | |
| EPI_ISL_14174522 | A/mink/China/456/2018 | A / H5N6 | Asia / China | |
| EPI_ISL_14174523 | A/swine/China/RZ/2018 | A / H5N6 | Asia / China | |
| EPI_ISL_1420637 | A/wigeon/Latvia/23903/2021 | A / H5N8 | Europe / Latvia / Jurmala | Juris Kibilds (Institute of Food Safety, Animal Health and Environment BIOR / Laboratory of Microbiology and Pathology) |
| EPI_ISL_14224170 | A/chicken-Cobb/Egypt/55/2019 | A / H5N8 | Africa / Egypt | |
| EPI_ISL_14224174 | A/chicken/Egypt/Dakahlia segment 4 HA gene/2021 | A / H5N8 | Africa / Egypt | |
| EPI_ISL_14233919 | A/Eurasian Spoonbill/Netherlands/3/2022 | A / H5N1 | Europe / Netherlands / Schiermonnikoog, Oosterkwelder | Sanne Thewessen (Erasmus Medical Center / Viroscience) |
| EPI_ISL_14233920 | A/Common Tern/Netherlands/17/2022 | A / H5N1 | Europe / Netherlands / Enkhuizen, Ijsselmeer, De Kreupel | Sanne Thewessen (Erasmus Medical Center / Viroscience) |
| EPI_ISL_14233921 | A/Common Tern/Netherlands/18/2022 | A / H5N1 | Europe / Netherlands / Enkhuizen, Ijsselmeer, De Kreupel | Sanne Thewessen (Erasmus Medical Center / Viroscience) |
| EPI_ISL_14233922 | A/European Herring Gull/Netherlands/11/2022 | A / H5N1 | Europe / Netherlands / Den Oever, Het Schor | Sanne Thewessen (Erasmus Medical Center / Viroscience) |
| EPI_ISL_14233923 | A/Caspian Gull/Netherlands/7/2022 | A / H5N1 | Europe / Netherlands / Enkhuizen, Ijsselmeer, De Kreupel | Sanne Thewessen (Erasmus Medical Center / Viroscience) |
| EPI_ISL_14233924 | A/Common Tern/Netherlands/19/2022 | A / H5N1 | Europe / Netherlands / Enkhuizen, Ijsselmeer, De Kreupel | Sanne Thewessen (Erasmus Medical Center / Viroscience) |
| EPI_ISL_14233925 | A/Common Tern/Netherlands/20/2022 | A / H5N1 | Europe / Netherlands / Enkhuizen, Ijsselmeer, De Kreupel | Sanne Thewessen (Erasmus Medical Center / Viroscience) |
| EPI_ISL_14233926 | A/European Herring Gull/Netherlands/10/2022 | A / H5N1 | Europe / Netherlands / Den Oever, Waddenhaven | Sanne Thewessen (Erasmus Medical Center / Viroscience) |
| EPI_ISL_14233927 | A/Common Tern/Netherlands/11/2022 | A / H5N1 | Europe / Netherlands / Wieringerwerf, Noorderdijkweg | Sanne Thewessen (Erasmus Medical Center / Viroscience) |
| EPI_ISL_14233928 | A/Common Tern/Netherlands/13/2022 | A / H5N1 | Europe / Netherlands / Wieringerwerf, Noorderdijkweg | Sanne Thewessen (Erasmus Medical Center / Viroscience) |
| EPI_ISL_14233929 | A/Common Tern/Netherlands/12/2022 | A / H5N1 | Europe / Netherlands / Wieringerwerf, Noorderdijkweg | Sanne Thewessen (Erasmus Medical Center / Viroscience) |
| EPI_ISL_14233930 | A/Common Tern/Netherlands/14/2022 | A / H5N1 | Europe / Netherlands / Wieringerwerf, Noorderdijkweg | Sanne Thewessen (Erasmus Medical Center / Viroscience) |
| EPI_ISL_14233931 | A/Common Tern/Netherlands/15/2022 | A / H5N1 | Europe / Netherlands / Wieringerwerf, Noorderdijkweg | Sanne Thewessen (Erasmus Medical Center / Viroscience) |
| EPI_ISL_14233932 | A/Common Tern/Netherlands/16/2022 | A / H5N1 | Europe / Netherlands / Wieringerwerf, Noorderdijkweg | Sanne Thewessen (Erasmus Medical Center / Viroscience) |
| EPI_ISL_14233944 | A/Caspian Gull/Netherlands/6/2022 | A / H5N1 | Europe / Netherlands / Enkhuizen, Ijsselmeer, De Kreupel | Sanne Thewessen (Erasmus Medical Center / Viroscience) |
| EPI_ISL_14388346 | A/Tyto_alba/Belgium/334_0012/2021 | A / H5N1 | Europe / Belgium / Provincie West-Vlaanderen / Waarschoot | Steven Van Borm (Sciensano, Department of Animal Infectious Diseases / Animal Infectious Diseases) |
| EPI_ISL_14388434 | A/Buteo_buteo/belgium/334_0013/2021 | A / H5N1 | Europe / Belgium / Provincie West-Vlaanderen / Desselgem | Steven Van Borm (Sciensano, Department of Animal Infectious Diseases / Animal Infectious Diseases) |
| EPI_ISL_14389133 | A/Larus_argentatus/Belgium/595_0008/2022 | A / H5N1 | Europe / Belgium / Provincie Antwerpen / Kapellen | Steven Van Borm (Sciensano, Department of Animal Infectious Diseases / Animal Infectious Diseases) |
| EPI_ISL_14389148 | A/Tachybaptus_ruficollis/Belgium/1234_0008/2022 | A / H5N1 | Europe / Belgium / Provincie Vlaams-Brabant / Oud-Heverlee | Steven Van Borm (Sciensano, Department of Animal Infectious Diseases / Animal Infectious Diseases) |
| EPI_ISL_14389524 | A/Anser_anser_domesticus/Belgium/1668_0016/2022 | A / H5N1 | Europe / Belgium / Provincie Vlaams-Brabant / Oud-Heverlee | Steven Van Borm (Sciensano, Department of Animal Infectious Diseases / Animal Infectious Diseases) |

| | | | |
|---|---|---|---|
| EPI_ISL_14390100 | A/Larus_canus/Belgium/1668_0019/2022 | A / H5N1 | Europe / Belgium / Provincie Antwerpen / Berchem | Steven Van Borm (Sciensano, Department of Animal Infectious Diseases / Animal Infectious Diseases) |
| EPI_ISL_14390385 | A/Phalacrocorax_carbo/Belgium/1734_0002/2022 | A / H5N1 | Europe / Belgium / Provincie Oost-Vlaanderen / Evergem | Steven Van Borm (Sciensano, Department of Animal Infectious Diseases / Animal Infectious Diseases) |
| EPI_ISL_14391865 | A/Buteo_buteo/Belgium/2606_0006/2022 | A / H5N1 | Europe / Belgium / Provincie Antwerpen / Kontich | Steven Van Borm (Sciensano, Department of Animal Infectious Diseases / Animal Infectious Diseases) |
| EPI_ISL_14392127 | A/Branta_leucopsis/Belgium/2606_0009/2022 | A / H5N1 | Europe / Belgium / Provincie West-Vlaanderen / Blankenberge | Steven Van Borm (Sciensano, Department of Animal Infectious Diseases / Animal Infectious Diseases) |
| EPI_ISL_14393097 | A/Fox/Netherlands/EMC4/2022 | A / H5N1 | Europe / Netherlands / Den Haag | Sanne Thewessen (Erasmus Medical Center / Viroscience) |
| EPI_ISL_14393115 | A/Fox/Netherlands/EMC5/2022 | A / H5N1 | Europe / Netherlands / Oosterbeek | Sanne Thewessen (Erasmus Medical Center / Viroscience) |
| EPI_ISL_14393126 | A/Fox/Netherlands/EMC6/2022 | A / H5N1 | Europe / Netherlands / Leusden | Sanne Thewessen (Erasmus Medical Center / Viroscience) |
| EPI_ISL_14393465 | A/Branta_canadensis/Belgium/4821_0001/2022 | A / H5N1 | Europe / Belgium / Province de Liege / Visé | Steven Van Borm (Sciensano, Department of Animal Infectious Diseases / Animal Infectious Diseases) |
| EPI_ISL_14393671 | A/Branta_canadensis/Belgium/5177_0003/2022 | A / H5N1 | Europe / Belgium / Province de Liege / Visé | Steven Van Borm (Sciensano, Department of Animal Infectious Diseases / Animal Infectious Diseases) |
| EPI_ISL_14469689 | A/Common Tern/Netherlands/21/2022 | A / H5N1 | Europe / Netherlands / Enkhuizen, Ijsselmeer, De Kreupel | Sanne Thewessen (Erasmus Medical Center / Viroscience) |
| EPI_ISL_14469690 | A/Common Tern/Netherlands/22/2022 | A / H5N1 | Europe / Netherlands / Enkhuizen, Ijsselmeer, De Kreupel | Sanne Thewessen (Erasmus Medical Center / Viroscience) |
| EPI_ISL_14469691 | A/Common Tern/Netherlands/23/2022 | A / H5N1 | Europe / Netherlands / Enkhuizen, Ijsselmeer, De Kreupel | Sanne Thewessen (Erasmus Medical Center / Viroscience) |
| EPI_ISL_14469692 | A/Common Tern/Netherlands/24/2022 | A / H5N1 | Europe / Netherlands / Enkhuizen, Ijsselmeer, De Kreupel | Sanne Thewessen (Erasmus Medical Center / Viroscience) |
| EPI_ISL_14469693 | A/Common Tern/Netherlands/25/2022 | A / H5N1 | Europe / Netherlands / Marker Wadden | Sanne Thewessen (Erasmus Medical Center / Viroscience) |
| EPI_ISL_14493899 | A/Vulpes_vulpes/Belgium/8660_0016/2022 | A / H5N1 | Europe / Belgium / Provincie Vlaams-Brabant / Kortenberg | Steven Van Borm (Sciensano, Department of Animal Infectious Diseases / Animal Infectious Diseases) |
| EPI_ISL_14494351 | A/Larus_argentatus/Belgium/9013_0001/2022 | A / H5N1 | Europe / Belgium / Provincie Oost-Vlaanderen / Gent | Steven Van Borm (Sciensano, Department of Animal Infectious Diseases / Animal Infectious Diseases) |
| EPI_ISL_14494796 | A/Vulpes_vulpes/Belgium/9031_0008/2022 | A / H5N1 | Europe / Belgium / Provincie West-Vlaanderen / Knokke-Heist | Steven Van Borm (Sciensano, Department of Animal Infectious Diseases / Animal Infectious Diseases) |
| EPI_ISL_14494933 | A/Gallus_gallus/Belgium/9548_0001/2022 | A / H5N1 | Europe / Belgium / Provincie West-Vlaanderen / Roeselare | Steven Van Borm (Sciensano, Department of Animal Infectious Diseases / Animal Infectious Diseases) |
| EPI_ISL_14497321 | A/Eurasian Spoonbill/Netherlands/4/2022 | A / H5N1 | Europe / Netherlands / Werkeiland, Neeltje Jans | Sanne Thewessen (Erasmus Medical Center / Viroscience) |
| EPI_ISL_14497837 | A/domestic_duck/England/100990/2022 | A / H5N1 | Europe / United Kingdom / England | Alex Byrne (Animal and Plant Health Agency (APHA) / Virology Department) |
| EPI_ISL_14497855 | A/domestic_duck/England/104859/2022 | A / H5N1 | Europe / United Kingdom / England | Alex Byrne (Animal and Plant Health Agency (APHA) / Virology Department) |
| EPI_ISL_14497870 | A/domestic_duck/England/105414/2022 | A / H5N1 | Europe / United Kingdom / England | Alex Byrne (Animal and Plant Health Agency (APHA) / Virology Department) |
| EPI_ISL_14497885 | A/domestic_goose/England/105863/2022 | A / H5N1 | Europe / United Kingdom / England | Alex Byrne (Animal and Plant Health Agency (APHA) / Virology Department) |
| EPI_ISL_14533193 | A/fox/New_York/074441/2022 | A / H5N1 | North America / United States / New York / Cayuga County | Brittany D Cronk (Cornell University / Population Medicine and Diagnostics) |
| EPI_ISL_14551957 | A/fox/New_York/088592/2022 | A / H5N1 | North America / United States / New York / Saint Lawrence County | Brittany D Cronk (Cornell University / Population Medicine and Diagnostics) |
| EPI_ISL_14552806 | A/fox/New_York/095525/2022 | A / H5N1 | North America / United States / New York / Saint Lawrence County | Brittany D Cronk (Cornell University / Population Medicine and Diagnostics) |
| EPI_ISL_14552975 | A/fox/New_York/099451/2022 | A / H5N1 | North America / United States / New York / Erie County | Brittany D Cronk (Cornell University / Population Medicine and Diagnostics) |
| EPI_ISL_14553810 | A/fox/New_York/099488/2022 | A / H5N1 | North America / United States / New York / Cayuga County | Brittany D Cronk (Cornell University / Population Medicine and Diagnostics) |
| EPI_ISL_14553811 | A/fox/New_York/103994/2022 | A / H5N1 | North America / United States / New York / Niagara County | Brittany D Cronk (Cornell University / Population Medicine and Diagnostics) |
| EPI_ISL_14553813 | A/fox/New_York/107242/2022 | A / H5N1 | North America / United States / New York / Allegany County | Brittany D Cronk (Cornell University / Population Medicine and Diagnostics) |
| EPI_ISL_14553997 | A/fox/New_York/115912/2022 | A / H5N1 | North America / United States / New York / Allegany County | Brittany D Cronk (Cornell University / Population Medicine and Diagnostics) |
| EPI_ISL_14604371 | A/crow/Hokkaido/0101Q061/2022 | A / H5N1 | Asia / Japan / Hokkaido / Rausu | Norikazu Isoda (Graduate School of Veterinary Medicine, Hokkaido University / Laboratory of Microbiology) |
| EPI_ISL_14615026 | A/crow/Hokkaido/0102L015/2022 | A / H5N1 | Asia / Japan / Hokkaido / Rebun | Norikazu Isoda (Graduate School of Veterinary Medicine, Hokkaido University / Laboratory of Microbiology) |
| EPI_ISL_14615027 | A/crow/Hokkaido/0103L018/2022 | A / H5N1 | Asia / Japan / Hokkaido / Hamatonbetsu | Norikazu Isoda (Graduate School of Veterinary Medicine, Hokkaido University / Laboratory of Microbiology) |
| EPI_ISL_14615036 | A/crow/Hokkaido/0101Q056/2022 | A / H5N1 | Asia / Japan / Hokkaido / Shibetsu | Norikazu Isoda (Graduate School of Veterinary Medicine, Hokkaido University / Laboratory of Microbiology) |
| EPI_ISL_14702899 | A/nothern_gannet/France/22P019331/2022 | A / H5N1 | Europe / France / Brittany / Departement des Cotes-d'Armor | Francois-Xavier Briand (ANSES Agence Nationale De Securite Sanitaire De L'alimentation / Laboratoire de Ploufragan-Plouzané) |
| EPI_ISL_14722960 | A/vulture/France/22P018210/2022 | A / H5N1 | Europe / France / Rhone-Alpes / Departement de la Drome | Francois-Xavier Briand (ANSES Agence Nationale De Securite Sanitaire De L'alimentation / Laboratoire de Ploufragan-Plouzané) |
| EPI_ISL_14749595 | A/duck/China/DK09-HA.seq/2016 | A / H5N6 | Asia / China | |
| EPI_ISL_14749603 | A/goose/China/GS38-HA.seq/2016 | A / H5N6 | Asia / China | |
| EPI_ISL_14760542 | A/turkey/Italy/21VIR8728-1/2021 | A / H5N1 | Europe / Italy | Giacomo Barbierato (Istituto Zooprofilattico Sperimentale delle Venezie / Ricerca e innovazione) |
| EPI_ISL_14760543 | A/turkey/Italy/21VIR8816-1/2021 | A / H5N1 | Europe / Italy | Giacomo Barbierato (Istituto Zooprofilattico Sperimentale delle Venezie / Ricerca e innovazione) |
| EPI_ISL_14760544 | A/turkey/Italy/21VIR8925-6/2021 | A / H5N1 | Europe / Italy | Giacomo Barbierato (Istituto Zooprofilattico Sperimentale delle Venezie / Ricerca e innovazione) |
| EPI_ISL_14760545 | A/turkey/Italy/21VIR8926-1/2021 | A / H5N1 | Europe / Italy | Giacomo Barbierato (Istituto Zooprofilattico Sperimentale delle Venezie / Ricerca e innovazione) |
| EPI_ISL_14760546 | A/turkey/Italy/21VIR8926-2/2021 | A / H5N1 | Europe / Italy | Giacomo Barbierato (Istituto Zooprofilattico Sperimentale delle Venezie / Ricerca e innovazione) |
| EPI_ISL_14760547 | A/turkey/Italy/21VIR9073-3/2021 | A / H5N1 | Europe / Italy | Giacomo Barbierato (Istituto Zooprofilattico Sperimentale delle Venezie / Ricerca e innovazione) |
| EPI_ISL_14760548 | A/chicken/Italy/21VIR9074-10/2021 | A / H5N1 | Europe / Italy | Giacomo Barbierato (Istituto Zooprofilattico Sperimentale delle Venezie / Ricerca e innovazione) |
| EPI_ISL_14760549 | A/turkey/Italy/21VIR9144-2/2021 | A / H5N1 | Europe / Italy | Giacomo Barbierato (Istituto Zooprofilattico Sperimentale delle Venezie / Ricerca e innovazione) |
| EPI_ISL_14760550 | A/chicken/Italy/21VIR9133-21/2021 | A / H5N1 | Europe / Italy | Giacomo Barbierato (Istituto Zooprofilattico Sperimentale delle Venezie / Ricerca e innovazione) |
| EPI_ISL_14760551 | A/turkey/Italy/21VIR9209-1/2021 | A / H5N1 | Europe / Italy | Giacomo Barbierato (Istituto Zooprofilattico Sperimentale delle Venezie / Ricerca e innovazione) |
| EPI_ISL_14760552 | A/chicken/Italy/21VIR9212-1/2021 | A / H5N1 | Europe / Italy | Giacomo Barbierato (Istituto Zooprofilattico Sperimentale delle Venezie / Ricerca e innovazione) |
| EPI_ISL_14760553 | A/turkey/Italy/21VIR9211-1/2021 | A / H5N1 | Europe / Italy | Giacomo Barbierato (Istituto Zooprofilattico Sperimentale delle Venezie / Ricerca e innovazione) |
| EPI_ISL_14760554 | A/turkey/Italy/21VIR9217-1/2021 | A / H5N1 | Europe / Italy | Giacomo Barbierato (Istituto Zooprofilattico Sperimentale delle Venezie / Ricerca e innovazione) |
| EPI_ISL_14760555 | A/turkey/Italy/21VIR9213-1/2021 | A / H5N1 | Europe / Italy | Giacomo Barbierato (Istituto Zooprofilattico Sperimentale delle Venezie / Ricerca e innovazione) |
| EPI_ISL_14760556 | A/turkey/Italy/21VIR9215-1/2021 | A / H5N1 | Europe / Italy | Giacomo Barbierato (Istituto Zooprofilattico Sperimentale delle Venezie / Ricerca e innovazione) |
| EPI_ISL_14760557 | A/chicken/Italy/21VIR9219-6/2021 | A / H5N1 | Europe / Italy | Giacomo Barbierato (Istituto Zooprofilattico Sperimentale delle Venezie / Ricerca e innovazione) |
| EPI_ISL_14760558 | A/chicken/Italy/21VIR9371-1/2021 | A / H5N1 | Europe / Italy | Giacomo Barbierato (Istituto Zooprofilattico Sperimentale delle Venezie / Ricerca e innovazione) |
| EPI_ISL_14760559 | A/avian/Italy/21VIR9425-2/2021 | A / H5N1 | Europe / Italy | Giacomo Barbierato (Istituto Zooprofilattico Sperimentale delle Venezie / Ricerca e innovazione) |
| EPI_ISL_14760560 | A/turkey/Italy/21VIR9372-1/2021 | A / H5N1 | Europe / Italy | Giacomo Barbierato (Istituto Zooprofilattico Sperimentale delle Venezie / Ricerca e innovazione) |
| EPI_ISL_14760561 | A/quail/Italy/21VIR9474-1/2021 | A / H5N1 | Europe / Italy | Giacomo Barbierato (Istituto Zooprofilattico Sperimentale delle Venezie / Ricerca e innovazione) |
| EPI_ISL_14760562 | A/duck/Italy/21VIR9373-2/2021 | A / H5N1 | Europe / Italy | Giacomo Barbierato (Istituto Zooprofilattico Sperimentale delle Venezie / Ricerca e innovazione) |
| EPI_ISL_14760563 | A/turkey/Italy/21VIR9426-3/2021 | A / H5N1 | Europe / Italy | Giacomo Barbierato (Istituto Zooprofilattico Sperimentale delle Venezie / Ricerca e innovazione) |
| EPI_ISL_14760564 | A/turkey/Italy/21VIR9476-1/2021 | A / H5N1 | Europe / Italy | Giacomo Barbierato (Istituto Zooprofilattico Sperimentale delle Venezie / Ricerca e innovazione) |
| EPI_ISL_14760565 | A/turkey/Italy/21VIR9475-1/2021 | A / H5N1 | Europe / Italy | Giacomo Barbierato (Istituto Zooprofilattico Sperimentale delle Venezie / Ricerca e innovazione) |
| EPI_ISL_14760566 | A/chicken/Italy/21VIR9509-1/2021 | A / H5N1 | Europe / Italy | Giacomo Barbierato (Istituto Zooprofilattico Sperimentale delle Venezie / Ricerca e innovazione) |
| EPI_ISL_14760567 | A/chicken/Italy/21VIR9507-7/2021 | A / H5N1 | Europe / Italy | Giacomo Barbierato (Istituto Zooprofilattico Sperimentale delle Venezie / Ricerca e innovazione) |
| EPI_ISL_14760568 | A/chicken/Italy/21VIR9508-3/2021 | A / H5N1 | Europe / Italy | Giacomo Barbierato (Istituto Zooprofilattico Sperimentale delle Venezie / Ricerca e innovazione) |
| EPI_ISL_14760569 | A/turkey/Italy/21VIR9607-2/2021 | A / H5N1 | Europe / Italy | Giacomo Barbierato (Istituto Zooprofilattico Sperimentale delle Venezie / Ricerca e innovazione) |
| EPI_ISL_14760570 | A/turkey/Italy/21VIR9606-2/2021 | A / H5N1 | Europe / Italy | Giacomo Barbierato (Istituto Zooprofilattico Sperimentale delle Venezie / Ricerca e innovazione) |
| EPI_ISL_14760571 | A/turkey/Italy/21VIR9604-3/2021 | A / H5N1 | Europe / Italy | Giacomo Barbierato (Istituto Zooprofilattico Sperimentale delle Venezie / Ricerca e innovazione) |
| EPI_ISL_14760572 | A/turkey/Italy/21VIR9609-2/2021 | A / H5N1 | Europe / Italy | Giacomo Barbierato (Istituto Zooprofilattico Sperimentale delle Venezie / Ricerca e innovazione) |
| EPI_ISL_14760573 | A/turkey/Italy/21VIR9605-4/2021 | A / H5N1 | Europe / Italy | Giacomo Barbierato (Istituto Zooprofilattico Sperimentale delle Venezie / Ricerca e innovazione) |
| EPI_ISL_14760574 | A/turkey/Italy/21VIR9608-2/2021 | A / H5N1 | Europe / Italy | Giacomo Barbierato (Istituto Zooprofilattico Sperimentale delle Venezie / Ricerca e innovazione) |

| | | | | |
|---|---|---|---|---|
| EPI_ISL_14760575 | A/quail/Italy/21VIR9515-1/2021 | A / H5N1 | Europe / Italy | Giacomo Barbierato (Istituto Zooprofilattico Sperimentale delle Venezie / Ricerca e innovazione) |
| EPI_ISL_14760576 | A/turkey/Italy/21VIR9518-1/2021 | A / H5N1 | Europe / Italy | Giacomo Barbierato (Istituto Zooprofilattico Sperimentale delle Venezie / Ricerca e innovazione) |
| EPI_ISL_14760577 | A/turkey/Italy/21VIR9519-1/2021 | A / H5N1 | Europe / Italy | Giacomo Barbierato (Istituto Zooprofilattico Sperimentale delle Venezie / Ricerca e innovazione) |
| EPI_ISL_14760578 | A/turkey/Italy/21VIR9520-1/2021 | A / H5N1 | Europe / Italy | Giacomo Barbierato (Istituto Zooprofilattico Sperimentale delle Venezie / Ricerca e innovazione) |
| EPI_ISL_14760579 | A/turkey/Italy/21VIR9643-2/2021 | A / H5N1 | Europe / Italy | Giacomo Barbierato (Istituto Zooprofilattico Sperimentale delle Venezie / Ricerca e innovazione) |
| EPI_ISL_14760580 | A/turkey/Italy/21VIR9644-2/2021 | A / H5N1 | Europe / Italy | Giacomo Barbierato (Istituto Zooprofilattico Sperimentale delle Venezie / Ricerca e innovazione) |
| EPI_ISL_14760581 | A/turkey/Italy/21VIR9645-2/2021 | A / H5N1 | Europe / Italy | Giacomo Barbierato (Istituto Zooprofilattico Sperimentale delle Venezie / Ricerca e innovazione) |
| EPI_ISL_14760582 | A/chicken/Italy/21VIR9580-23/2021 | A / H5N1 | Europe / Italy | Giacomo Barbierato (Istituto Zooprofilattico Sperimentale delle Venezie / Ricerca e innovazione) |
| EPI_ISL_14760583 | A/turkey/Italy/21VIR9579-10/2021 | A / H5N1 | Europe / Italy | Giacomo Barbierato (Istituto Zooprofilattico Sperimentale delle Venezie / Ricerca e innovazione) |
| EPI_ISL_14760584 | A/turkey/Italy/21VIR9648-2/2021 | A / H5N1 | Europe / Italy | Giacomo Barbierato (Istituto Zooprofilattico Sperimentale delle Venezie / Ricerca e innovazione) |
| EPI_ISL_14760585 | A/turkey/Italy/21VIR9651-2/2021 | A / H5N1 | Europe / Italy | Giacomo Barbierato (Istituto Zooprofilattico Sperimentale delle Venezie / Ricerca e innovazione) |
| EPI_ISL_14760586 | A/turkey/Italy/21VIR9650-2/2021 | A / H5N1 | Europe / Italy | Giacomo Barbierato (Istituto Zooprofilattico Sperimentale delle Venezie / Ricerca e innovazione) |
| EPI_ISL_14760587 | A/turkey/Italy/21VIR9646-2/2021 | A / H5N1 | Europe / Italy | Giacomo Barbierato (Istituto Zooprofilattico Sperimentale delle Venezie / Ricerca e innovazione) |
| EPI_ISL_14760588 | A/turkey/Italy/21VIR9647-2/2021 | A / H5N1 | Europe / Italy | Giacomo Barbierato (Istituto Zooprofilattico Sperimentale delle Venezie / Ricerca e innovazione) |
| EPI_ISL_14760589 | A/turkey/Italy/21VIR9694-2/2021 | A / H5N1 | Europe / Italy | Giacomo Barbierato (Istituto Zooprofilattico Sperimentale delle Venezie / Ricerca e innovazione) |
| EPI_ISL_14760590 | A/turkey/Italy/21VIR9692-16/2021 | A / H5N1 | Europe / Italy | Giacomo Barbierato (Istituto Zooprofilattico Sperimentale delle Venezie / Ricerca e innovazione) |
| EPI_ISL_14760591 | A/turkey/Italy/21VIR9695-4/2021 | A / H5N1 | Europe / Italy | Giacomo Barbierato (Istituto Zooprofilattico Sperimentale delle Venezie / Ricerca e innovazione) |
| EPI_ISL_14760592 | A/turkey/Italy/21VIR9693-3/2021 | A / H5N1 | Europe / Italy | Giacomo Barbierato (Istituto Zooprofilattico Sperimentale delle Venezie / Ricerca e innovazione) |
| EPI_ISL_14760593 | A/turkey/Italy/21VIR9696-2/2021 | A / H5N1 | Europe / Italy | Giacomo Barbierato (Istituto Zooprofilattico Sperimentale delle Venezie / Ricerca e innovazione) |
| EPI_ISL_14760594 | A/turkey/Italy/21VIR9697-3_/2022 | A / H5N1 | Europe / Italy | Giacomo Barbierato (Istituto Zooprofilattico Sperimentale delle Venezie / Ricerca e innovazione) |
| EPI_ISL_14760595 | A/turkey/Italy/21VIR9769-4/2021 | A / H5N1 | Europe / Italy | Giacomo Barbierato (Istituto Zooprofilattico Sperimentale delle Venezie / Ricerca e innovazione) |
| EPI_ISL_14760596 | A/turkey/Italy/21VIR9766-3/2021 | A / H5N1 | Europe / Italy | Giacomo Barbierato (Istituto Zooprofilattico Sperimentale delle Venezie / Ricerca e innovazione) |
| EPI_ISL_14760597 | A/chicken/Italy/21VIR9868-8/2021 | A / H5N1 | Europe / Italy | Giacomo Barbierato (Istituto Zooprofilattico Sperimentale delle Venezie / Ricerca e innovazione) |
| EPI_ISL_14760598 | A/chicken/Italy/21VIR9902-2/2021 | A / H5N1 | Europe / Italy | Giacomo Barbierato (Istituto Zooprofilattico Sperimentale delle Venezie / Ricerca e innovazione) |
| EPI_ISL_14760599 | A/turkey/Italy/21VIR9870-3/2021 | A / H5N1 | Europe / Italy | Giacomo Barbierato (Istituto Zooprofilattico Sperimentale delle Venezie / Ricerca e innovazione) |
| EPI_ISL_14760600 | A/turkey/Italy/21VIR9869-4/2021 | A / H5N1 | Europe / Italy | Giacomo Barbierato (Istituto Zooprofilattico Sperimentale delle Venezie / Ricerca e innovazione) |
| EPI_ISL_14760601 | A/turkey/Italy/21VIR9903-2/2021 | A / H5N1 | Europe / Italy | Giacomo Barbierato (Istituto Zooprofilattico Sperimentale delle Venezie / Ricerca e innovazione) |
| EPI_ISL_14760602 | A/chicken/Italy/21VIR9830-27/2021 | A / H5N1 | Europe / Italy | Giacomo Barbierato (Istituto Zooprofilattico Sperimentale delle Venezie / Ricerca e innovazione) |
| EPI_ISL_14760603 | A/chicken/Italy/21VIR9904-16/2021 | A / H5N1 | Europe / Italy | Giacomo Barbierato (Istituto Zooprofilattico Sperimentale delle Venezie / Ricerca e innovazione) |
| EPI_ISL_14760604 | A/chicken/Italy/21VIR9904-17/2021 | A / H5N1 | Europe / Italy | Giacomo Barbierato (Istituto Zooprofilattico Sperimentale delle Venezie / Ricerca e innovazione) |
| EPI_ISL_14760605 | A/chicken/Italy/21VIR9904-18/2021 | A / H5N1 | Europe / Italy | Giacomo Barbierato (Istituto Zooprofilattico Sperimentale delle Venezie / Ricerca e innovazione) |
| EPI_ISL_14760606 | A/turkey/Italy/21VIR9906-2/2021 | A / H5N1 | Europe / Italy | Giacomo Barbierato (Istituto Zooprofilattico Sperimentale delle Venezie / Ricerca e innovazione) |
| EPI_ISL_14760607 | A/turkey/Italy/21VIR9905-2/2021 | A / H5N1 | Europe / Italy | Giacomo Barbierato (Istituto Zooprofilattico Sperimentale delle Venezie / Ricerca e innovazione) |
| EPI_ISL_14760608 | A/duck/Italy/21VIR9836-15/2021 | A / H5N1 | Europe / Italy | Giacomo Barbierato (Istituto Zooprofilattico Sperimentale delle Venezie / Ricerca e innovazione) |
| EPI_ISL_14760609 | A/turkey/Italy/21VIR9839-1/2021 | A / H5N1 | Europe / Italy | Giacomo Barbierato (Istituto Zooprofilattico Sperimentale delle Venezie / Ricerca e innovazione) |
| EPI_ISL_14760610 | A/chicken/Italy/21VIR9837-3/2021 | A / H5N1 | Europe / Italy | Giacomo Barbierato (Istituto Zooprofilattico Sperimentale delle Venezie / Ricerca e innovazione) |
| EPI_ISL_14760611 | A/goose/Italy/IZSLT_21VIR10273/2021 | A / H5N1 | Europe / Italy | Giacomo Barbierato (Istituto Zooprofilattico Sperimentale delle Venezie / Ricerca e innovazione) |
| EPI_ISL_14760612 | A/chicken/Italy/21VIR9858-3/2021 | A / H5N1 | Europe / Italy | Giacomo Barbierato (Istituto Zooprofilattico Sperimentale delle Venezie / Ricerca e innovazione) |
| EPI_ISL_14760613 | A/turkey/Italy/21VIR9889-1/2021 | A / H5N1 | Europe / Italy | Giacomo Barbierato (Istituto Zooprofilattico Sperimentale delle Venezie / Ricerca e innovazione) |
| EPI_ISL_14760614 | A/turkey/Italy/21VIR9891-21/2021 | A / H5N1 | Europe / Italy | Giacomo Barbierato (Istituto Zooprofilattico Sperimentale delle Venezie / Ricerca e innovazione) |
| EPI_ISL_14760615 | A/turkey/Italy/21VIR9996/2021 | A / H5N1 | Europe / Italy | Giacomo Barbierato (Istituto Zooprofilattico Sperimentale delle Venezie / Ricerca e innovazione) |
| EPI_ISL_14760616 | A/chicken/Italy/21VIR10000/2021 | A / H5N1 | Europe / Italy | Giacomo Barbierato (Istituto Zooprofilattico Sperimentale delle Venezie / Ricerca e innovazione) |
| EPI_ISL_14760617 | A/turkey/Italy/21VIR9999/2021 | A / H5N1 | Europe / Italy | Giacomo Barbierato (Istituto Zooprofilattico Sperimentale delle Venezie / Ricerca e innovazione) |
| EPI_ISL_14760618 | A/turkey/Italy/21VIR9997/2021 | A / H5N1 | Europe / Italy | Giacomo Barbierato (Istituto Zooprofilattico Sperimentale delle Venezie / Ricerca e innovazione) |
| EPI_ISL_14760619 | A/turkey/Italy/21VIR10001/2021 | A / H5N1 | Europe / Italy | Giacomo Barbierato (Istituto Zooprofilattico Sperimentale delle Venezie / Ricerca e innovazione) |
| EPI_ISL_14760620 | A/chicken/Italy/21VIR9998/2021 | A / H5N1 | Europe / Italy | Giacomo Barbierato (Istituto Zooprofilattico Sperimentale delle Venezie / Ricerca e innovazione) |
| EPI_ISL_14760621 | A/turkey/Italy/21VIR9894-3/2021 | A / H5N1 | Europe / Italy | Giacomo Barbierato (Istituto Zooprofilattico Sperimentale delle Venezie / Ricerca e innovazione) |
| EPI_ISL_14760622 | A/chicken/Italy/21VIR9944-2/2021 | A / H5N1 | Europe / Italy | Giacomo Barbierato (Istituto Zooprofilattico Sperimentale delle Venezie / Ricerca e innovazione) |
| EPI_ISL_14760623 | A/turkey/Italy/21VIR9949-9/2021 | A / H5N1 | Europe / Italy | Giacomo Barbierato (Istituto Zooprofilattico Sperimentale delle Venezie / Ricerca e innovazione) |
| EPI_ISL_14760624 | A/chicken/Italy/21VIR9940-5/2021 | A / H5N1 | Europe / Italy | Giacomo Barbierato (Istituto Zooprofilattico Sperimentale delle Venezie / Ricerca e innovazione) |
| EPI_ISL_14760625 | A/chicken/Italy/21VIR10096/2021 | A / H5N1 | Europe / Italy | Giacomo Barbierato (Istituto Zooprofilattico Sperimentale delle Venezie / Ricerca e innovazione) |
| EPI_ISL_14760626 | A/chicken/Italy/21VIR10093/2021 | A / H5N1 | Europe / Italy | Giacomo Barbierato (Istituto Zooprofilattico Sperimentale delle Venezie / Ricerca e innovazione) |
| EPI_ISL_14760627 | A/turkey/Italy/21VIR10097/2021 | A / H5N1 | Europe / Italy | Giacomo Barbierato (Istituto Zooprofilattico Sperimentale delle Venezie / Ricerca e innovazione) |
| EPI_ISL_14760628 | A/turkey/Italy/21VIR10095/2021 | A / H5N1 | Europe / Italy | Giacomo Barbierato (Istituto Zooprofilattico Sperimentale delle Venezie / Ricerca e innovazione) |
| EPI_ISL_14760629 | A/chicken/Italy/21VIR10094/2021 | A / H5N1 | Europe / Italy | Giacomo Barbierato (Istituto Zooprofilattico Sperimentale delle Venezie / Ricerca e innovazione) |
| EPI_ISL_14760630 | A/turkey/Italy/21VIR10092/2021 | A / H5N1 | Europe / Italy | Giacomo Barbierato (Istituto Zooprofilattico Sperimentale delle Venezie / Ricerca e innovazione) |
| EPI_ISL_14760631 | A/turkey/Italy/21VIR9901-3/2021 | A / H5N1 | Europe / Italy | Giacomo Barbierato (Istituto Zooprofilattico Sperimentale delle Venezie / Ricerca e innovazione) |
| EPI_ISL_14760632 | A/Guinea_fowl/Italy/21VIR10216/2021 | A / H5N1 | Europe / Italy | Giacomo Barbierato (Istituto Zooprofilattico Sperimentale delle Venezie / Ricerca e innovazione) |
| EPI_ISL_14760633 | A/Guinea_fowl/Italy/21VIR10231/2021 | A / H5N1 | Europe / Italy | Giacomo Barbierato (Istituto Zooprofilattico Sperimentale delle Venezie / Ricerca e innovazione) |
| EPI_ISL_14760634 | A/turkey/Italy/21VIR10103/2021 | A / H5N1 | Europe / Italy | Giacomo Barbierato (Istituto Zooprofilattico Sperimentale delle Venezie / Ricerca e innovazione) |
| EPI_ISL_14760635 | A/chicken/Italy/21VIR10252/2021 | A / H5N1 | Europe / Italy | Giacomo Barbierato (Istituto Zooprofilattico Sperimentale delle Venezie / Ricerca e innovazione) |
| EPI_ISL_14760636 | A/chicken/Italy/21VIR10146/2021 | A / H5N1 | Europe / Italy | Giacomo Barbierato (Istituto Zooprofilattico Sperimentale delle Venezie / Ricerca e innovazione) |
| EPI_ISL_14760637 | A/pheasant/Italy/21VIR10359/2021 | A / H5N1 | Europe / Italy | Giacomo Barbierato (Istituto Zooprofilattico Sperimentale delle Venezie / Ricerca e innovazione) |
| EPI_ISL_14760638 | A/guinea_fowl/Italy/21VIR10351/2021 | A / H5N1 | Europe / Italy | Giacomo Barbierato (Istituto Zooprofilattico Sperimentale delle Venezie / Ricerca e innovazione) |
| EPI_ISL_14760639 | A/quail/Italy/21VIR10356/2021 | A / H5N1 | Europe / Italy | Giacomo Barbierato (Istituto Zooprofilattico Sperimentale delle Venezie / Ricerca e innovazione) |
| EPI_ISL_14760640 | A/chicken/Italy/21VIR10358/2021 | A / H5N1 | Europe / Italy | Giacomo Barbierato (Istituto Zooprofilattico Sperimentale delle Venezie / Ricerca e innovazione) |
| EPI_ISL_14760641 | A/chicken/Italy/21VIR10357/2021 | A / H5N1 | Europe / Italy | Giacomo Barbierato (Istituto Zooprofilattico Sperimentale delle Venezie / Ricerca e innovazione) |
| EPI_ISL_14760642 | A/chicken/Italy/21VIR10262/2021 | A / H5N1 | Europe / Italy | Giacomo Barbierato (Istituto Zooprofilattico Sperimentale delle Venezie / Ricerca e innovazione) |
| EPI_ISL_14760643 | A/chicken/Italy/21VIR10263/2021 | A / H5N1 | Europe / Italy | Giacomo Barbierato (Istituto Zooprofilattico Sperimentale delle Venezie / Ricerca e innovazione) |
| EPI_ISL_14760644 | A/chicken/Italy/21VIR10271/2021 | A / H5N1 | Europe / Italy | Giacomo Barbierato (Istituto Zooprofilattico Sperimentale delle Venezie / Ricerca e innovazione) |
| EPI_ISL_14760645 | A/chicken/Italy/21VIR10264/2021 | A / H5N1 | Europe / Italy | Giacomo Barbierato (Istituto Zooprofilattico Sperimentale delle Venezie / Ricerca e innovazione) |
| EPI_ISL_14760646 | A/chicken/Italy/21VIR10265/2021 | A / H5N1 | Europe / Italy | Giacomo Barbierato (Istituto Zooprofilattico Sperimentale delle Venezie / Ricerca e innovazione) |

| | | | | |
|---|---|---|---|---|
| EPI_ISL_14760647 | A/turkey/Italy/21VIR10338/2021 | A / H5N1 | Europe / Italy | Giacomo Barbierato (Istituto Zooprofilattico Sperimentale delle Venezie / Ricerca e innovazione) |
| EPI_ISL_14760648 | A/chicken/Italy/21VIR10382/2021 | A / H5N1 | Europe / Italy | Giacomo Barbierato (Istituto Zooprofilattico Sperimentale delle Venezie / Ricerca e innovazione) |
| EPI_ISL_14760649 | A/turkey/Italy/21VIR10396/2021 | A / H5N1 | Europe / Italy | Giacomo Barbierato (Istituto Zooprofilattico Sperimentale delle Venezie / Ricerca e innovazione) |
| EPI_ISL_14760650 | A/turkey/Italy/21VIR10395/2021 | A / H5N1 | Europe / Italy | Giacomo Barbierato (Istituto Zooprofilattico Sperimentale delle Venezie / Ricerca e innovazione) |
| EPI_ISL_14760651 | A/chicken/Italy/21VIR10394/2021 | A / H5N1 | Europe / Italy | Giacomo Barbierato (Istituto Zooprofilattico Sperimentale delle Venezie / Ricerca e innovazione) |
| EPI_ISL_14760652 | A/turkey/Italy/21VIR10397/2021 | A / H5N1 | Europe / Italy | Giacomo Barbierato (Istituto Zooprofilattico Sperimentale delle Venezie / Ricerca e innovazione) |
| EPI_ISL_14760653 | A/turkey/Italy/21VIR10398/2021 | A / H5N1 | Europe / Italy | Giacomo Barbierato (Istituto Zooprofilattico Sperimentale delle Venezie / Ricerca e innovazione) |
| EPI_ISL_14760654 | A/chicken/Italy/21VIR10470/2021 | A / H5N1 | Europe / Italy | Giacomo Barbierato (Istituto Zooprofilattico Sperimentale delle Venezie / Ricerca e innovazione) |
| EPI_ISL_14760655 | A/chicken/Italy/21VIR10387/2021 | A / H5N1 | Europe / Italy | Giacomo Barbierato (Istituto Zooprofilattico Sperimentale delle Venezie / Ricerca e innovazione) |
| EPI_ISL_14760656 | A/goose/Italy/Italy/21VIR10458/2021 | A / H5N1 | Europe / Italy | Giacomo Barbierato (Istituto Zooprofilattico Sperimentale delle Venezie / Ricerca e innovazione) |
| EPI_ISL_14760658 | A/chicken/Italy/21VIR10385/2021 | A / H5N1 | Europe / Italy | Giacomo Barbierato (Istituto Zooprofilattico Sperimentale delle Venezie / Ricerca e innovazione) |
| EPI_ISL_14760659 | A/chicken/Italy/21VIR10469/2021 | A / H5N1 | Europe / Italy | Giacomo Barbierato (Istituto Zooprofilattico Sperimentale delle Venezie / Ricerca e innovazione) |
| EPI_ISL_14760660 | A/turkey/Italy/21VIR10471/2021 | A / H5N1 | Europe / Italy | Giacomo Barbierato (Istituto Zooprofilattico Sperimentale delle Venezie / Ricerca e innovazione) |
| EPI_ISL_14760661 | A/chicken/Italy/21VIR10386/2021 | A / H5N1 | Europe / Italy | Giacomo Barbierato (Istituto Zooprofilattico Sperimentale delle Venezie / Ricerca e innovazione) |
| EPI_ISL_14760662 | A/duck/Italy/21VIR10523-1/2021 | A / H5N1 | Europe / Italy | Giacomo Barbierato (Istituto Zooprofilattico Sperimentale delle Venezie / Ricerca e innovazione) |
| EPI_ISL_14760663 | A/turkey/Italy/21VIR10524-1/2021 | A / H5N1 | Europe / Italy | Giacomo Barbierato (Istituto Zooprofilattico Sperimentale delle Venezie / Ricerca e innovazione) |
| EPI_ISL_14760664 | A/turkey/Italy/21VIR10459-1/2021 | A / H5N1 | Europe / Italy | Giacomo Barbierato (Istituto Zooprofilattico Sperimentale delle Venezie / Ricerca e innovazione) |
| EPI_ISL_14760665 | A/chicken/Italy/21VIR10463-1/2021 | A / H5N1 | Europe / Italy | Giacomo Barbierato (Istituto Zooprofilattico Sperimentale delle Venezie / Ricerca e innovazione) |
| EPI_ISL_14760666 | A/chicken/Italy/21VIR10522-13/2021 | A / H5N1 | Europe / Italy | Giacomo Barbierato (Istituto Zooprofilattico Sperimentale delle Venezie / Ricerca e innovazione) |
| EPI_ISL_14760667 | A/turkey/Italy/21VIR10622-1/2021 | A / H5N1 | Europe / Italy | Giacomo Barbierato (Istituto Zooprofilattico Sperimentale delle Venezie / Ricerca e innovazione) |
| EPI_ISL_14760668 | A/turkey/Italy/21VIR10617-2/2021 | A / H5N1 | Europe / Italy | Giacomo Barbierato (Istituto Zooprofilattico Sperimentale delle Venezie / Ricerca e innovazione) |
| EPI_ISL_14760669 | A/turkey/Italy/21VIR10621-1/2021 | A / H5N1 | Europe / Italy | Giacomo Barbierato (Istituto Zooprofilattico Sperimentale delle Venezie / Ricerca e innovazione) |
| EPI_ISL_14760670 | A/chicken/Italy/21VIR10616-1/2021 | A / H5N1 | Europe / Italy | Giacomo Barbierato (Istituto Zooprofilattico Sperimentale delle Venezie / Ricerca e innovazione) |
| EPI_ISL_14760671 | A/chicken/Italy/21VIR10618-19/2021 | A / H5N1 | Europe / Italy | Giacomo Barbierato (Istituto Zooprofilattico Sperimentale delle Venezie / Ricerca e innovazione) |
| EPI_ISL_14760672 | A/chicken/Italy/21VIR10619-1/2021 | A / H5N1 | Europe / Italy | Giacomo Barbierato (Istituto Zooprofilattico Sperimentale delle Venezie / Ricerca e innovazione) |
| EPI_ISL_14760673 | A/chicken/Italy/21VIR10620-1/2021 | A / H5N1 | Europe / Italy | Giacomo Barbierato (Istituto Zooprofilattico Sperimentale delle Venezie / Ricerca e innovazione) |
| EPI_ISL_14760674 | A/chicken/Italy/21VIR10724-13/2021 | A / H5N1 | Europe / Italy | Giacomo Barbierato (Istituto Zooprofilattico Sperimentale delle Venezie / Ricerca e innovazione) |
| EPI_ISL_14760675 | A/chicken/Italy/21VIR10723-1/2021 | A / H5N1 | Europe / Italy | Giacomo Barbierato (Istituto Zooprofilattico Sperimentale delle Venezie / Ricerca e innovazione) |
| EPI_ISL_14760676 | A/chicken/Italy/21VIR10634-1/2021 | A / H5N1 | Europe / Italy | Giacomo Barbierato (Istituto Zooprofilattico Sperimentale delle Venezie / Ricerca e innovazione) |
| EPI_ISL_14760677 | A/chicken/Italy/21VIR10635-1/2021 | A / H5N1 | Europe / Italy | Giacomo Barbierato (Istituto Zooprofilattico Sperimentale delle Venezie / Ricerca e innovazione) |
| EPI_ISL_14760678 | A/chicken/Italy/21VIR10637-2/2021 | A / H5N1 | Europe / Italy | Giacomo Barbierato (Istituto Zooprofilattico Sperimentale delle Venezie / Ricerca e innovazione) |
| EPI_ISL_14760679 | A/turkey/Italy/21VIR10636-17/2021 | A / H5N1 | Europe / Italy | Giacomo Barbierato (Istituto Zooprofilattico Sperimentale delle Venezie / Ricerca e innovazione) |
| EPI_ISL_14760680 | A/turkey/Italy/21VIR10645-15/2021 | A / H5N1 | Europe / Italy | Giacomo Barbierato (Istituto Zooprofilattico Sperimentale delle Venezie / Ricerca e innovazione) |
| EPI_ISL_14760681 | A/chicken/Italy/21VIR10853/2021 | A / H5N1 | Europe / Italy | Giacomo Barbierato (Istituto Zooprofilattico Sperimentale delle Venezie / Ricerca e innovazione) |
| EPI_ISL_14760682 | A/turkey/Italy/21VIR10699-1/2021 | A / H5N1 | Europe / Italy | Giacomo Barbierato (Istituto Zooprofilattico Sperimentale delle Venezie / Ricerca e innovazione) |
| EPI_ISL_14760683 | A/turkey/Italy/21VIR10856/2021 | A / H5N1 | Europe / Italy | Giacomo Barbierato (Istituto Zooprofilattico Sperimentale delle Venezie / Ricerca e innovazione) |
| EPI_ISL_14760684 | A/turkey/Italy/21VIR10852/2021 | A / H5N1 | Europe / Italy | Giacomo Barbierato (Istituto Zooprofilattico Sperimentale delle Venezie / Ricerca e innovazione) |
| EPI_ISL_14760685 | A/turkey/Italy/21VIR10857/2021 | A / H5N1 | Europe / Italy | Giacomo Barbierato (Istituto Zooprofilattico Sperimentale delle Venezie / Ricerca e innovazione) |
| EPI_ISL_14760686 | A/turkey/Italy/21VIR10854/2021 | A / H5N1 | Europe / Italy | Giacomo Barbierato (Istituto Zooprofilattico Sperimentale delle Venezie / Ricerca e innovazione) |
| EPI_ISL_14760687 | A/turkey/Italy/21VIR10855/2021 | A / H5N1 | Europe / Italy | Giacomo Barbierato (Istituto Zooprofilattico Sperimentale delle Venezie / Ricerca e innovazione) |
| EPI_ISL_14760688 | A/chicken/Italy/21VIR10696-7/2021 | A / H5N1 | Europe / Italy | Giacomo Barbierato (Istituto Zooprofilattico Sperimentale delle Venezie / Ricerca e innovazione) |
| EPI_ISL_14760689 | A/turkey/Italy/21VIR10697-1/2021 | A / H5N1 | Europe / Italy | Giacomo Barbierato (Istituto Zooprofilattico Sperimentale delle Venezie / Ricerca e innovazione) |
| EPI_ISL_14760690 | A/turkey/Italy/21VIR10737-1/2021 | A / H5N1 | Europe / Italy | Giacomo Barbierato (Istituto Zooprofilattico Sperimentale delle Venezie / Ricerca e innovazione) |
| EPI_ISL_14760691 | A/chicken/Italy/21VIR10814-1/2021 | A / H5N1 | Europe / Italy | Giacomo Barbierato (Istituto Zooprofilattico Sperimentale delle Venezie / Ricerca e innovazione) |
| EPI_ISL_14760692 | A/chicken/Italy/21VIR10808-1/2021 | A / H5N1 | Europe / Italy | Giacomo Barbierato (Istituto Zooprofilattico Sperimentale delle Venezie / Ricerca e innovazione) |
| EPI_ISL_14760693 | A/duck/Italy/21VIR10809-1/2021 | A / H5N1 | Europe / Italy | Giacomo Barbierato (Istituto Zooprofilattico Sperimentale delle Venezie / Ricerca e innovazione) |
| EPI_ISL_14760694 | A/turkey/Italy/21VIR10810-1/2021 | A / H5N1 | Europe / Italy | Giacomo Barbierato (Istituto Zooprofilattico Sperimentale delle Venezie / Ricerca e innovazione) |
| EPI_ISL_14760695 | A/chicken/Italy/21VIR10811-28/2021 | A / H5N1 | Europe / Italy | Giacomo Barbierato (Istituto Zooprofilattico Sperimentale delle Venezie / Ricerca e innovazione) |
| EPI_ISL_14760696 | A/chicken/Italy/21VIR10812-11/2021 | A / H5N1 | Europe / Italy | Giacomo Barbierato (Istituto Zooprofilattico Sperimentale delle Venezie / Ricerca e innovazione) |
| EPI_ISL_14760697 | A/turkey/Italy/21VIR10929/2021 | A / H5N1 | Europe / Italy | Giacomo Barbierato (Istituto Zooprofilattico Sperimentale delle Venezie / Ricerca e innovazione) |
| EPI_ISL_14760698 | A/chicken/Italy/21VIR10928/2021 | A / H5N1 | Europe / Italy | Giacomo Barbierato (Istituto Zooprofilattico Sperimentale delle Venezie / Ricerca e innovazione) |
| EPI_ISL_14760699 | A/turkey/Italy/21VIR10930/2021 | A / H5N1 | Europe / Italy | Giacomo Barbierato (Istituto Zooprofilattico Sperimentale delle Venezie / Ricerca e innovazione) |
| EPI_ISL_14760700 | A/turkey/Italy/21VIR10757-1/2021 | A / H5N1 | Europe / Italy | Giacomo Barbierato (Istituto Zooprofilattico Sperimentale delle Venezie / Ricerca e innovazione) |
| EPI_ISL_14760701 | A/chicken/Italy/21VIR10843-1/2021 | A / H5N1 | Europe / Italy | Giacomo Barbierato (Istituto Zooprofilattico Sperimentale delle Venezie / Ricerca e innovazione) |
| EPI_ISL_14760702 | A/turkey/Italy/21VIR10847-1/2021 | A / H5N1 | Europe / Italy | Giacomo Barbierato (Istituto Zooprofilattico Sperimentale delle Venezie / Ricerca e innovazione) |
| EPI_ISL_14760703 | A/turkey/Italy/21VIR10970-3/2021 | A / H5N1 | Europe / Italy | Giacomo Barbierato (Istituto Zooprofilattico Sperimentale delle Venezie / Ricerca e innovazione) |
| EPI_ISL_14760704 | A/chicken/Italy/21VIR10971-3/2021 | A / H5N1 | Europe / Italy | Giacomo Barbierato (Istituto Zooprofilattico Sperimentale delle Venezie / Ricerca e innovazione) |
| EPI_ISL_14760705 | A/chicken/Italy/21VIR10973-4/2021 | A / H5N1 | Europe / Italy | Giacomo Barbierato (Istituto Zooprofilattico Sperimentale delle Venezie / Ricerca e innovazione) |
| EPI_ISL_14760706 | A/chicken/Italy/21VIR10972-3/2021 | A / H5N1 | Europe / Italy | Giacomo Barbierato (Istituto Zooprofilattico Sperimentale delle Venezie / Ricerca e innovazione) |
| EPI_ISL_14760707 | A/chicken/Italy/21VIR10969-3/2021 | A / H5N1 | Europe / Italy | Giacomo Barbierato (Istituto Zooprofilattico Sperimentale delle Venezie / Ricerca e innovazione) |
| EPI_ISL_14760708 | A/turkey/Italy/21VIR10988-1/2021 | A / H5N1 | Europe / Italy | Giacomo Barbierato (Istituto Zooprofilattico Sperimentale delle Venezie / Ricerca e innovazione) |
| EPI_ISL_14760709 | A/chicken/Italy/21VIR10989/2021 | A / H5N1 | Europe / Italy | Giacomo Barbierato (Istituto Zooprofilattico Sperimentale delle Venezie / Ricerca e innovazione) |
| EPI_ISL_14760710 | A/chicken/Italy/21VIR11072-1/2021 | A / H5N1 | Europe / Italy | Giacomo Barbierato (Istituto Zooprofilattico Sperimentale delle Venezie / Ricerca e innovazione) |
| EPI_ISL_14760711 | A/chicken/Italy/21VIR11074-1/2021 | A / H5N1 | Europe / Italy | Giacomo Barbierato (Istituto Zooprofilattico Sperimentale delle Venezie / Ricerca e innovazione) |
| EPI_ISL_14760712 | A/chicken/Italy/21VIR11071-2/2021 | A / H5N1 | Europe / Italy | Giacomo Barbierato (Istituto Zooprofilattico Sperimentale delle Venezie / Ricerca e innovazione) |
| EPI_ISL_14760713 | A/avian/Italy/21VIR10913-3/2021 | A / H5N1 | Europe / Italy | Giacomo Barbierato (Istituto Zooprofilattico Sperimentale delle Venezie / Ricerca e innovazione) |
| EPI_ISL_14760714 | A/chicken/Italy/21VIR10987-1/2021 | A / H5N1 | Europe / Italy | Giacomo Barbierato (Istituto Zooprofilattico Sperimentale delle Venezie / Ricerca e innovazione) |
| EPI_ISL_14760715 | A/chicken/Italy/21VIR11076-1/2021 | A / H5N1 | Europe / Italy | Giacomo Barbierato (Istituto Zooprofilattico Sperimentale delle Venezie / Ricerca e innovazione) |
| EPI_ISL_14760716 | A/chicken/Italy/21VIR11079-11/2021 | A / H5N1 | Europe / Italy | Giacomo Barbierato (Istituto Zooprofilattico Sperimentale delle Venezie / Ricerca e innovazione) |
| EPI_ISL_14760717 | A/chicken/Italy/21VIR11078-10/2021 | A / H5N1 | Europe / Italy | Giacomo Barbierato (Istituto Zooprofilattico Sperimentale delle Venezie / Ricerca e innovazione) |
| EPI_ISL_14760718 | A/chicken/Italy/21VIR11073-1/2021 | A / H5N1 | Europe / Italy | Giacomo Barbierato (Istituto Zooprofilattico Sperimentale delle Venezie / Ricerca e innovazione) |
| EPI_ISL_14760719 | A/chicken/Italy/21VIR11075-2/2021 | A / H5N1 | Europe / Italy | Giacomo Barbierato (Istituto Zooprofilattico Sperimentale delle Venezie / Ricerca e innovazione) |

| | | | | |
|---|---|---|---|---|
| EPI_ISL_14760720 | A/turkey/Italy/21VIR11077-1/2021 | A / H5N1 | Europe / Italy | Giacomo Barbierato (Istituto Zooprofilattico Sperimentale delle Venezie / Ricerca e innovazione) |
| EPI_ISL_14760721 | A/chicken/Italy/21VIR10995-1/2021 | A / H5N1 | Europe / Italy | Giacomo Barbierato (Istituto Zooprofilattico Sperimentale delle Venezie / Ricerca e innovazione) |
| EPI_ISL_14760722 | A/chicken/Italy/21VIR10999-1/2021 | A / H5N1 | Europe / Italy | Giacomo Barbierato (Istituto Zooprofilattico Sperimentale delle Venezie / Ricerca e innovazione) |
| EPI_ISL_14760723 | A/chicken/Italy/21VIR10997-1/2021 | A / H5N1 | Europe / Italy | Giacomo Barbierato (Istituto Zooprofilattico Sperimentale delle Venezie / Ricerca e innovazione) |
| EPI_ISL_14760724 | A/turkey/Italy/21VIR11006-1/2021 | A / H5N1 | Europe / Italy | Giacomo Barbierato (Istituto Zooprofilattico Sperimentale delle Venezie / Ricerca e innovazione) |
| EPI_ISL_14760725 | A/turkey/Italy/21VIR11007-1/2021 | A / H5N1 | Europe / Italy | Giacomo Barbierato (Istituto Zooprofilattico Sperimentale delle Venezie / Ricerca e innovazione) |
| EPI_ISL_14760726 | A/turkey/Italy/21VIR11166/2021 | A / H5N1 | Europe / Italy | Giacomo Barbierato (Istituto Zooprofilattico Sperimentale delle Venezie / Ricerca e innovazione) |
| EPI_ISL_14760727 | A/chicken/Italy/21VIR11055-1/2021 | A / H5N1 | Europe / Italy | Giacomo Barbierato (Istituto Zooprofilattico Sperimentale delle Venezie / Ricerca e innovazione) |
| EPI_ISL_14760728 | A/pheasant/Italy/21VIR11162/2021 | A / H5N1 | Europe / Italy | Giacomo Barbierato (Istituto Zooprofilattico Sperimentale delle Venezie / Ricerca e innovazione) |
| EPI_ISL_14760729 | A/turkey/Italy/21VIR11164/2021 | A / H5N1 | Europe / Italy | Giacomo Barbierato (Istituto Zooprofilattico Sperimentale delle Venezie / Ricerca e innovazione) |
| EPI_ISL_14760730 | A/pheasant/Italy/21VIR11165/2021 | A / H5N1 | Europe / Italy | Giacomo Barbierato (Istituto Zooprofilattico Sperimentale delle Venezie / Ricerca e innovazione) |
| EPI_ISL_14760731 | A/laying_hen/Italy/21VIR11167/2021 | A / H5N1 | Europe / Italy | Giacomo Barbierato (Istituto Zooprofilattico Sperimentale delle Venezie / Ricerca e innovazione) |
| EPI_ISL_14760732 | A/laying_hen/Italy/21VIR11168/2021 | A / H5N1 | Europe / Italy | Giacomo Barbierato (Istituto Zooprofilattico Sperimentale delle Venezie / Ricerca e innovazione) |
| EPI_ISL_14760733 | A/laying_hen/Italy/21VIR11169/2021 | A / H5N1 | Europe / Italy | Giacomo Barbierato (Istituto Zooprofilattico Sperimentale delle Venezie / Ricerca e innovazione) |
| EPI_ISL_14760734 | A/chicken/Italy/21VIR11234/2021 | A / H5N1 | Europe / Italy | Giacomo Barbierato (Istituto Zooprofilattico Sperimentale delle Venezie / Ricerca e innovazione) |
| EPI_ISL_14760735 | A/laying_hen/Italy/21VIR11300/2021 | A / H5N1 | Europe / Italy | Giacomo Barbierato (Istituto Zooprofilattico Sperimentale delle Venezie / Ricerca e innovazione) |
| EPI_ISL_14760736 | A/chicken/Italy/21VIR11054-1/2021 | A / H5N1 | Europe / Italy | Giacomo Barbierato (Istituto Zooprofilattico Sperimentale delle Venezie / Ricerca e innovazione) |
| EPI_ISL_14760737 | A/chicken/Italy/21VIR11056-1/2021 | A / H5N1 | Europe / Italy | Giacomo Barbierato (Istituto Zooprofilattico Sperimentale delle Venezie / Ricerca e innovazione) |
| EPI_ISL_14760738 | A/chicken/Italy/21VIR11056-6/2021 | A / H5N1 | Europe / Italy | Giacomo Barbierato (Istituto Zooprofilattico Sperimentale delle Venezie / Ricerca e innovazione) |
| EPI_ISL_14760739 | A/turkey/Italy/21VIR11058/2021 | A / H5N1 | Europe / Italy | Giacomo Barbierato (Istituto Zooprofilattico Sperimentale delle Venezie / Ricerca e innovazione) |
| EPI_ISL_14760740 | A/turkey/Italy/21VIR11163/2021 | A / H5N1 | Europe / Italy | Giacomo Barbierato (Istituto Zooprofilattico Sperimentale delle Venezie / Ricerca e innovazione) |
| EPI_ISL_14760741 | A/chicken/Italy/21VIR11235/2021 | A / H5N1 | Europe / Italy | Giacomo Barbierato (Istituto Zooprofilattico Sperimentale delle Venezie / Ricerca e innovazione) |
| EPI_ISL_14760742 | A/avian/Italy/21VIR11236/2021 | A / H5N1 | Europe / Italy | Giacomo Barbierato (Istituto Zooprofilattico Sperimentale delle Venezie / Ricerca e innovazione) |
| EPI_ISL_14760743 | A/turkey/Italy/21VIR11237/2021 | A / H5N1 | Europe / Italy | Giacomo Barbierato (Istituto Zooprofilattico Sperimentale delle Venezie / Ricerca e innovazione) |
| EPI_ISL_14760744 | A/chicken/Italy/21VIR11238/2021 | A / H5N1 | Europe / Italy | Giacomo Barbierato (Istituto Zooprofilattico Sperimentale delle Venezie / Ricerca e innovazione) |
| EPI_ISL_14760745 | A/guinea_fowl/Italy/21VIR11298/2021 | A / H5N1 | Europe / Italy | Giacomo Barbierato (Istituto Zooprofilattico Sperimentale delle Venezie / Ricerca e innovazione) |
| EPI_ISL_14760746 | A/turkey/Italy/21VIR11302/2021 | A / H5N1 | Europe / Italy | Giacomo Barbierato (Istituto Zooprofilattico Sperimentale delle Venezie / Ricerca e innovazione) |
| EPI_ISL_14760747 | A/turkey/Italy/21VIR11251-1/2021 | A / H5N1 | Europe / Italy | Giacomo Barbierato (Istituto Zooprofilattico Sperimentale delle Venezie / Ricerca e innovazione) |
| EPI_ISL_14760748 | A/turkey/Italy/21VIR11252-1/2021 | A / H5N1 | Europe / Italy | Giacomo Barbierato (Istituto Zooprofilattico Sperimentale delle Venezie / Ricerca e innovazione) |
| EPI_ISL_14760749 | A/turkey/Italy/21VIR11254-1/2021 | A / H5N1 | Europe / Italy | Giacomo Barbierato (Istituto Zooprofilattico Sperimentale delle Venezie / Ricerca e innovazione) |
| EPI_ISL_14760750 | A/laying_hen/Italy/21VIR11253-1/2021 | A / H5N1 | Europe / Italy | Giacomo Barbierato (Istituto Zooprofilattico Sperimentale delle Venezie / Ricerca e innovazione) |
| EPI_ISL_14760751 | A/laying_hen/Italy/21VIR11299/2021 | A / H5N1 | Europe / Italy | Giacomo Barbierato (Istituto Zooprofilattico Sperimentale delle Venezie / Ricerca e innovazione) |
| EPI_ISL_14760752 | A/avian/Italy/21VIR11301/2021 | A / H5N1 | Europe / Italy | Giacomo Barbierato (Istituto Zooprofilattico Sperimentale delle Venezie / Ricerca e innovazione) |
| EPI_ISL_14760753 | A/turkey/Italy/21VIR11384-1/2021 | A / H5N1 | Europe / Italy | Giacomo Barbierato (Istituto Zooprofilattico Sperimentale delle Venezie / Ricerca e innovazione) |
| EPI_ISL_14760754 | A/turkey/Italy/21VIR11387-1/2021 | A / H5N1 | Europe / Italy | Giacomo Barbierato (Istituto Zooprofilattico Sperimentale delle Venezie / Ricerca e innovazione) |
| EPI_ISL_14760755 | A/laying_hen/Italy/21VIR11390-2/2021 | A / H5N1 | Europe / Italy | Giacomo Barbierato (Istituto Zooprofilattico Sperimentale delle Venezie / Ricerca e innovazione) |
| EPI_ISL_14760756 | A/turkey/Italy/21VIR11391-3/2021 | A / H5N1 | Europe / Italy | Giacomo Barbierato (Istituto Zooprofilattico Sperimentale delle Venezie / Ricerca e innovazione) |
| EPI_ISL_14760757 | A/turkey/Italy/21VIR11392-2/2021 | A / H5N1 | Europe / Italy | Giacomo Barbierato (Istituto Zooprofilattico Sperimentale delle Venezie / Ricerca e innovazione) |
| EPI_ISL_14760758 | A/turkey/Italy/21VIR11393-3/2021 | A / H5N1 | Europe / Italy | Giacomo Barbierato (Istituto Zooprofilattico Sperimentale delle Venezie / Ricerca e innovazione) |
| EPI_ISL_14760762 | A/guinea_fowl/Italy/21VIR11383-1/2021 | A / H5N1 | Europe / Italy | Giacomo Barbierato (Istituto Zooprofilattico Sperimentale delle Venezie / Ricerca e innovazione) |
| EPI_ISL_14760774 | A/chicken/Italy/21VIR11385-1/2021 | A / H5N1 | Europe / Italy | Giacomo Barbierato (Istituto Zooprofilattico Sperimentale delle Venezie / Ricerca e innovazione) |
| EPI_ISL_14760784 | A/turkey/Italy/21VIR11386-1/2021 | A / H5N1 | Europe / Italy | Giacomo Barbierato (Istituto Zooprofilattico Sperimentale delle Venezie / Ricerca e innovazione) |
| EPI_ISL_14760793 | A/laying_hen/Italy/21VIR11394-1/2021 | A / H5N1 | Europe / Italy | Giacomo Barbierato (Istituto Zooprofilattico Sperimentale delle Venezie / Ricerca e innovazione) |
| EPI_ISL_14760803 | A/laying_hen/Italy/21VIR11395-1/2021 | A / H5N1 | Europe / Italy | Giacomo Barbierato (Istituto Zooprofilattico Sperimentale delle Venezie / Ricerca e innovazione) |
| EPI_ISL_14760814 | A/turkey/Italy/21VIR11396-1/2021 | A / H5N1 | Europe / Italy | Giacomo Barbierato (Istituto Zooprofilattico Sperimentale delle Venezie / Ricerca e innovazione) |
| EPI_ISL_14760826 | A/laying_hen/Italy/21VIR11486/2021 | A / H5N1 | Europe / Italy | Giacomo Barbierato (Istituto Zooprofilattico Sperimentale delle Venezie / Ricerca e innovazione) |
| EPI_ISL_14760837 | A/chicken/Italy/21VIR11485/2021 | A / H5N1 | Europe / Italy | Giacomo Barbierato (Istituto Zooprofilattico Sperimentale delle Venezie / Ricerca e innovazione) |
| EPI_ISL_14760849 | A/turkey/Italy/21VIR11509/2021 | A / H5N1 | Europe / Italy | Giacomo Barbierato (Istituto Zooprofilattico Sperimentale delle Venezie / Ricerca e innovazione) |
| EPI_ISL_14760859 | A/turkey/Italy/21VIR11510/2021 | A / H5N1 | Europe / Italy | Giacomo Barbierato (Istituto Zooprofilattico Sperimentale delle Venezie / Ricerca e innovazione) |
| EPI_ISL_14760870 | A/turkey/Italy/21VIR11440-2/2021 | A / H5N1 | Europe / Italy | Giacomo Barbierato (Istituto Zooprofilattico Sperimentale delle Venezie / Ricerca e innovazione) |
| EPI_ISL_14760880 | A/chicken/Italy/21VIR11508/2021 | A / H5N1 | Europe / Italy | Giacomo Barbierato (Istituto Zooprofilattico Sperimentale delle Venezie / Ricerca e innovazione) |
| EPI_ISL_14760891 | A/turkey/Italy/21VIR11511/2021 | A / H5N1 | Europe / Italy | Giacomo Barbierato (Istituto Zooprofilattico Sperimentale delle Venezie / Ricerca e innovazione) |
| EPI_ISL_14760902 | A/turkey/Italy/21VIR11506/2022 | A / H5N1 | Europe / Italy | Giacomo Barbierato (Istituto Zooprofilattico Sperimentale delle Venezie / Ricerca e innovazione) |
| EPI_ISL_14760913 | A/turkey/Italy/21VIR11499-5/2021 | A / H5N1 | Europe / Italy | Giacomo Barbierato (Istituto Zooprofilattico Sperimentale delle Venezie / Ricerca e innovazione) |
| EPI_ISL_14760923 | A/turkey/Italy/21VIR11503-1/2021 | A / H5N1 | Europe / Italy | Giacomo Barbierato (Istituto Zooprofilattico Sperimentale delle Venezie / Ricerca e innovazione) |
| EPI_ISL_14760936 | A/laying_hen/Italy/21VIR11505-2/2021 | A / H5N1 | Europe / Italy | Giacomo Barbierato (Istituto Zooprofilattico Sperimentale delle Venezie / Ricerca e innovazione) |
| EPI_ISL_14760946 | A/laying_hen/Italy/21VIR11498-1/2021 | A / H5N1 | Europe / Italy | Giacomo Barbierato (Istituto Zooprofilattico Sperimentale delle Venezie / Ricerca e innovazione) |
| EPI_ISL_14760956 | A/turkey/Italy/21VIR11492-4/2021 | A / H5N1 | Europe / Italy | Giacomo Barbierato (Istituto Zooprofilattico Sperimentale delle Venezie / Ricerca e innovazione) |
| EPI_ISL_14760967 | A/turkey/Italy/21VIR11495-5/2021 | A / H5N1 | Europe / Italy | Giacomo Barbierato (Istituto Zooprofilattico Sperimentale delle Venezie / Ricerca e innovazione) |
| EPI_ISL_14760977 | A/turkey/Italy/21VIR11500-3/2021 | A / H5N1 | Europe / Italy | Giacomo Barbierato (Istituto Zooprofilattico Sperimentale delle Venezie / Ricerca e innovazione) |
| EPI_ISL_14760988 | A/duck/Italy/21VIR11501/2021 | A / H5N1 | Europe / Italy | Giacomo Barbierato (Istituto Zooprofilattico Sperimentale delle Venezie / Ricerca e innovazione) |
| EPI_ISL_14760999 | A/turkey/Italy/21VIR11504/2021 | A / H5N1 | Europe / Italy | Giacomo Barbierato (Istituto Zooprofilattico Sperimentale delle Venezie / Ricerca e innovazione) |
| EPI_ISL_14761009 | A/chicken/Italy/21VIR11571/2021 | A / H5N1 | Europe / Italy | Giacomo Barbierato (Istituto Zooprofilattico Sperimentale delle Venezie / Ricerca e innovazione) |
| EPI_ISL_14761020 | A/chicken/Italy/21VIR11572/2021 | A / H5N1 | Europe / Italy | Giacomo Barbierato (Istituto Zooprofilattico Sperimentale delle Venezie / Ricerca e innovazione) |
| EPI_ISL_14761031 | A/turkey/Italy/21VIR11596-3/2021 | A / H5N1 | Europe / Italy | Giacomo Barbierato (Istituto Zooprofilattico Sperimentale delle Venezie / Ricerca e innovazione) |
| EPI_ISL_14761041 | A/broiler/Italy/21VIR11583-1/2021 | A / H5N1 | Europe / Italy | Giacomo Barbierato (Istituto Zooprofilattico Sperimentale delle Venezie / Ricerca e innovazione) |
| EPI_ISL_14761052 | A/laying_hen/Italy/21VIR11595-5/2021 | A / H5N1 | Europe / Italy | Giacomo Barbierato (Istituto Zooprofilattico Sperimentale delle Venezie / Ricerca e innovazione) |
| EPI_ISL_14761065 | A/laying_hen/Italy/21VIR11597-1/2021 | A / H5N1 | Europe / Italy | Giacomo Barbierato (Istituto Zooprofilattico Sperimentale delle Venezie / Ricerca e innovazione) |
| EPI_ISL_14761075 | A/laying_hen/Italy/21VIR11592-1/2021 | A / H5N1 | Europe / Italy | Giacomo Barbierato (Istituto Zooprofilattico Sperimentale delle Venezie / Ricerca e innovazione) |
| EPI_ISL_14761086 | A/turkey/Italy/21VIR11586-1/2021 | A / H5N1 | Europe / Italy | Giacomo Barbierato (Istituto Zooprofilattico Sperimentale delle Venezie / Ricerca e innovazione) |
| EPI_ISL_14761097 | A/turkey/Italy/21VIR11587-1/2021 | A / H5N1 | Europe / Italy | Giacomo Barbierato (Istituto Zooprofilattico Sperimentale delle Venezie / Ricerca e innovazione) |
| EPI_ISL_14761107 | A/turkey/Italy/21VIR11587-2/2021 | A / H5N1 | Europe / Italy | Giacomo Barbierato (Istituto Zooprofilattico Sperimentale delle Venezie / Ricerca e innovazione) |

| | | | | |
|---|---|---|---|---|
| EPI_ISL_14761119 | A/turkey/Italy/21VIR11691/2021 | A / H5N1 | Europe / Italy | Giacomo Barbierato (Istituto Zooprofilattico Sperimentale delle Venezie / Ricerca e innovazione) |
| EPI_ISL_14761129 | A/turkey/Italy/21VIR11692/2021 | A / H5N1 | Europe / Italy | Giacomo Barbierato (Istituto Zooprofilattico Sperimentale delle Venezie / Ricerca e innovazione) |
| EPI_ISL_14761141 | A/turkey/Italy/21VIR11690/2021 | A / H5N1 | Europe / Italy | Giacomo Barbierato (Istituto Zooprofilattico Sperimentale delle Venezie / Ricerca e innovazione) |
| EPI_ISL_14761152 | A/turkey/Italy/21VIR11661-7/2021 | A / H5N1 | Europe / Italy | Giacomo Barbierato (Istituto Zooprofilattico Sperimentale delle Venezie / Ricerca e innovazione) |
| EPI_ISL_14761165 | A/broiler/Italy/21VIR11800/2021 | A / H5N1 | Europe / Italy | Giacomo Barbierato (Istituto Zooprofilattico Sperimentale delle Venezie / Ricerca e innovazione) |
| EPI_ISL_14761176 | A/laying_hen/Italy/21VIR11801-7/2021 | A / H5N1 | Europe / Italy | Giacomo Barbierato (Istituto Zooprofilattico Sperimentale delle Venezie / Ricerca e innovazione) |
| EPI_ISL_14761188 | A/broiler/Italy/21VIR11802-1/2021 | A / H5N1 | Europe / Italy | Giacomo Barbierato (Istituto Zooprofilattico Sperimentale delle Venezie / Ricerca e innovazione) |
| EPI_ISL_14761199 | A/turkey/Italy/21VIR11772-7/2021 | A / H5N1 | Europe / Italy | Giacomo Barbierato (Istituto Zooprofilattico Sperimentale delle Venezie / Ricerca e innovazione) |
| EPI_ISL_14761208 | A/avian/Italy/21VIR11773-4/2021 | A / H5N1 | Europe / Italy | Giacomo Barbierato (Istituto Zooprofilattico Sperimentale delle Venezie / Ricerca e innovazione) |
| EPI_ISL_14761220 | A/chicken/Italy/21VIR11774-2/2021 | A / H5N1 | Europe / Italy | Giacomo Barbierato (Istituto Zooprofilattico Sperimentale delle Venezie / Ricerca e innovazione) |
| EPI_ISL_14761232 | A/turkey/Italy/21VIR11935/2021 | A / H5N1 | Europe / Italy | Giacomo Barbierato (Istituto Zooprofilattico Sperimentale delle Venezie / Ricerca e innovazione) |
| EPI_ISL_14761243 | A/broiler/Italy/21VIR11847-1/2021 | A / H5N1 | Europe / Italy | Giacomo Barbierato (Istituto Zooprofilattico Sperimentale delle Venezie / Ricerca e innovazione) |
| EPI_ISL_14761249 | A/broiler/Italy/21VIR11886-1/2021 | A / H5N1 | Europe / Italy | Giacomo Barbierato (Istituto Zooprofilattico Sperimentale delle Venezie / Ricerca e innovazione) |
| EPI_ISL_14761250 | A/heron/Italy/22VIR1562/2022 | A / H5N1 | Europe / Italy | Giacomo Barbierato (Istituto Zooprofilattico Sperimentale delle Venezie / Ricerca e innovazione) |
| EPI_ISL_14761251 | A/broiler/Italy/22VIR54-1/2022 | A / H5N1 | Europe / Italy | Giacomo Barbierato (Istituto Zooprofilattico Sperimentale delle Venezie / Ricerca e innovazione) |
| EPI_ISL_14761261 | A/laying_hen/Italy/22VIR48-1/2022 | A / H5N1 | Europe / Italy | Giacomo Barbierato (Istituto Zooprofilattico Sperimentale delle Venezie / Ricerca e innovazione) |
| EPI_ISL_14761272 | A/turkey/Italy/IZSLT_22VIR366-3/2022 | A / H5N1 | Europe / Italy | Giacomo Barbierato (Istituto Zooprofilattico Sperimentale delle Venezie / Ricerca e innovazione) |
| EPI_ISL_14761281 | A/turkey/Italy/22VIR171-2/2022 | A / H5N1 | Europe / Italy | Giacomo Barbierato (Istituto Zooprofilattico Sperimentale delle Venezie / Ricerca e innovazione) |
| EPI_ISL_14761291 | A/guinea_fowl/Italy/22VIR205-4/2022 | A / H5N1 | Europe / Italy | Giacomo Barbierato (Istituto Zooprofilattico Sperimentale delle Venezie / Ricerca e innovazione) |
| EPI_ISL_14761302 | A/laying_hen/Italy/22VIR203-3/2022 | A / H5N1 | Europe / Italy | Giacomo Barbierato (Istituto Zooprofilattico Sperimentale delle Venezie / Ricerca e innovazione) |
| EPI_ISL_14761312 | A/chicken/Italy/22VIR234-1/2022 | A / H5N1 | Europe / Italy | Giacomo Barbierato (Istituto Zooprofilattico Sperimentale delle Venezie / Ricerca e innovazione) |
| EPI_ISL_14761318 | A/laying_hen/Italy/22VIR204-1/2022 | A / H5N1 | Europe / Italy | Giacomo Barbierato (Istituto Zooprofilattico Sperimentale delle Venezie / Ricerca e innovazione) |
| EPI_ISL_14761327 | A/broiler/Italy/22VIR278-3/2022 | A / H5N1 | Europe / Italy | Giacomo Barbierato (Istituto Zooprofilattico Sperimentale delle Venezie / Ricerca e innovazione) |
| EPI_ISL_14761338 | A/hawk/Italy/22VIR428-5/2022 | A / H5N1 | Europe / Italy | Giacomo Barbierato (Istituto Zooprofilattico Sperimentale delle Venezie / Ricerca e innovazione) |
| EPI_ISL_14761349 | A/hawk/Italy/22VIR428-7/2022 | A / H5N1 | Europe / Italy | Giacomo Barbierato (Istituto Zooprofilattico Sperimentale delle Venezie / Ricerca e innovazione) |
| EPI_ISL_14761361 | A/buzzard/Italy/22VIR767-1/2022 | A / H5N1 | Europe / Italy | Giacomo Barbierato (Istituto Zooprofilattico Sperimentale delle Venezie / Ricerca e innovazione) |
| EPI_ISL_14761373 | A/buzzard/Italy/22VIR767-2/2022 | A / H5N1 | Europe / Italy | Giacomo Barbierato (Istituto Zooprofilattico Sperimentale delle Venezie / Ricerca e innovazione) |
| EPI_ISL_14761383 | A/swan/Italy/22VIR1560/2022 | A / H5N1 | Europe / Italy | Giacomo Barbierato (Istituto Zooprofilattico Sperimentale delle Venezie / Ricerca e innovazione) |
| EPI_ISL_14761395 | A/herring_gull/Italy/22VIR1710-3/2022 | A / H5N1 | Europe / Italy | Giacomo Barbierato (Istituto Zooprofilattico Sperimentale delle Venezie / Ricerca e innovazione) |
| EPI_ISL_14761407 | A/duck/Italy/22VIR1294-1/2022 | A / H5N1 | Europe / Italy | Giacomo Barbierato (Istituto Zooprofilattico Sperimentale delle Venezie / Ricerca e innovazione) |
| EPI_ISL_14761417 | A/duck/Italy/22VIR1295-1/2022 | A / H5N1 | Europe / Italy | Giacomo Barbierato (Istituto Zooprofilattico Sperimentale delle Venezie / Ricerca e innovazione) |
| EPI_ISL_14761428 | A/goose/Italy/22VIR1520/2022 | A / H5N1 | Europe / Italy | Giacomo Barbierato (Istituto Zooprofilattico Sperimentale delle Venezie / Ricerca e innovazione) |
| EPI_ISL_14761439 | A/laying_hen/Italy/22VIR1521-2/2022 | A / H5N1 | Europe / Italy | Giacomo Barbierato (Istituto Zooprofilattico Sperimentale delle Venezie / Ricerca e innovazione) |
| EPI_ISL_14761451 | A/chicken/Italy/22VIR1694-5/2022 | A / H5N1 | Europe / Italy | Giacomo Barbierato (Istituto Zooprofilattico Sperimentale delle Venezie / Ricerca e innovazione) |
| EPI_ISL_14761462 | A/pheasant/Italy/22VIR1953-1/2022 | A / H5N1 | Europe / Italy | Giacomo Barbierato (Istituto Zooprofilattico Sperimentale delle Venezie / Ricerca e innovazione) |
| EPI_ISL_14761473 | A/peacock/Italy/22VIR1953-2/2022 | A / H5N1 | Europe / Italy | Giacomo Barbierato (Istituto Zooprofilattico Sperimentale delle Venezie / Ricerca e innovazione) |
| EPI_ISL_14761485 | A/chicken/Italy/22VIR1953-3/2022 | A / H5N1 | Europe / Italy | Giacomo Barbierato (Istituto Zooprofilattico Sperimentale delle Venezie / Ricerca e innovazione) |
| EPI_ISL_14761496 | A/chicken/Italy/22VIR1953-4/2022 | A / H5N1 | Europe / Italy | Giacomo Barbierato (Istituto Zooprofilattico Sperimentale delle Venezie / Ricerca e innovazione) |
| EPI_ISL_14761507 | A/broiler/Italy/22VIR1892-3/2022 | A / H5N1 | Europe / Italy | Giacomo Barbierato (Istituto Zooprofilattico Sperimentale delle Venezie / Ricerca e innovazione) |
| EPI_ISL_14761519 | A/chicken/Italy/IZSLT22VIR2562-1/2022 | A / H5N1 | Europe / Italy | Giacomo Barbierato (Istituto Zooprofilattico Sperimentale delle Venezie / Ricerca e innovazione) |
| EPI_ISL_14769966 | A/chicken/Lesotho/341.10/2021 | A / H5N1 | Africa / Lesotho | |
| EPI_ISL_14769967 | A/chicken/Lesotho/352.3/2021 | A / H5N1 | Africa / Lesotho | |
| EPI_ISL_14770020 | A/chicken/Egypt/Q16684C/2019 | A / H5N8 | Africa / Egypt | |
| EPI_ISL_14770033 | A/chicken/Egypt/N13731D/2017 | A / H5N8 | Africa / Egypt | |
| EPI_ISL_14770035 | A/duck/Egypt/N13736A/2017 | A / H5N8 | Africa / Egypt | |
| EPI_ISL_14770036 | A/chicken/Egypt/H13794D/2017 | A / H5N8 | Africa / Egypt | |
| EPI_ISL_14770037 | A/chicken/Egypt/Q13845A/2017 | A / H5N8 | Africa / Egypt | |
| EPI_ISL_14770039 | A/chicken/Egypt/A13773E/2017 | A / H5N8 | Africa / Egypt | |
| EPI_ISL_14770042 | A/duck/Egypt/N14205C/2017 | A / H5N8 | Africa / Egypt | |
| EPI_ISL_14770043 | A/chicken/Egypt/Q13936C/2017 | A / H5N8 | Africa / Egypt | |
| EPI_ISL_14770046 | A/chicken/Egypt/N13720E/2017 | A / H5N8 | Africa / Egypt | |
| EPI_ISL_14770047 | A/chicken/Egypt/N15172D/2018 | A / H5N8 | Africa / Egypt | |
| EPI_ISL_14770049 | A/duck/Egypt/N14200A/2017 | A / H5N8 | Africa / Egypt | |
| EPI_ISL_14770051 | A/chicken/Egypt/F14110D/2017 | A / H5N8 | Africa / Egypt | |
| EPI_ISL_14770153 | A/chicken/Egypt/F13829B/2017 | A / H5N8 | Africa / Egypt | |
| EPI_ISL_14770166 | A/swan/Shandong/W3762/2020 | A / H5N8 | Asia / China | |
| EPI_ISL_14775174 | A/chicken/Viet Nam/QN_21_3105/2021 | A / H5N8 | Asia / Vietnam | |
| EPI_ISL_14775175 | A/Duck/Viet Nam/QN_21_5289/2021 | A / H5N8 | Asia / Vietnam | |
| EPI_ISL_14775176 | A/chicken/Viet Nam/QN-20-985/2020 | A / H5N6 | Asia / Vietnam | |
| EPI_ISL_14775177 | A/chicken/Viet Nam/QN-20-1144/2020 | A / H5N6 | Asia / Vietnam | |
| EPI_ISL_14775178 | A/chicken/Viet Nam/QN-20-3597/2020 | A / H5N6 | Asia / Vietnam | |
| EPI_ISL_14775179 | A/chicken/Viet Nam/QN-21-279/2021 | A / H5N6 | Asia / Vietnam | |
| EPI_ISL_14775180 | A/chicken/Viet Nam/QN-21-109/2021 | A / H5N6 | Asia / Vietnam | |
| EPI_ISL_14775181 | A/chicken/Viet Nam/QN-21-187/2021 | A / H5N6 | Asia / Vietnam | |
| EPI_ISL_14775182 | A/chicken/Viet Nam/QN-21-237/2021 | A / H5N6 | Asia / Vietnam | |
| EPI_ISL_14775183 | A/chicken/Viet Nam/QN-21-255/2021 | A / H5N6 | Asia / Vietnam | |
| EPI_ISL_14775184 | A/chicken/Viet Nam/QN-21-438/2021 | A / H5N6 | Asia / Vietnam | |
| EPI_ISL_14775185 | A/chicken/Viet Nam/QN-21-956/2021 | A / H5N6 | Asia / Vietnam | |
| EPI_ISL_14775186 | A/chicken/Viet Nam/QN-21-1044/2021 | A / H5N6 | Asia / Vietnam | |
| EPI_ISL_14775274 | A/chicken/Viet Nam/20-161/2020 | A / H5N6 | Asia / Vietnam | |
| EPI_ISL_14775517 | A/goose/Hebei/HG12/2021 | A / H5N8 | Asia / China / Hebei Province | |
| EPI_ISL_14775522 | A/Duck/Ismailia/F6/2021 | A / H5N8 | Africa / Egypt / Ismailia | |

| | | | | |
|---|---|---|---|---|
| EPI_ISL_14775523 | A/Duck/Ismailia/F16/2021 | A / H5N8 | Africa / Egypt / Ismailia | |
| EPI_ISL_14775524 | A/chicken/Asyut/F2/2021 | A / H5N8 | Africa / Egypt / Asyut | |
| EPI_ISL_14778363 | A/Wild bird/China/Cixi02/2020 | A / H5N8 | Asia / China | |
| EPI_ISL_14810369 | A/porpoise /Sweden/SVA220712SZ0367/FB002184/O-2022 | A / H5N1 | Europe / Sweden / Vastra Gotalands Lan / Tanums Kommun | Siamak Zohari (National Veterinary Institute) |
| EPI_ISL_14811896 | A/northern gannet/Sweden/SVA220623SZ0232/FB001982/O-2 | A / H5N1 | Europe / Sweden / Vastra Gotalands Lan / Sotenas Kommun | Siamak Zohari (National Veterinary Institute) |
| EPI_ISL_14811929 | A/northern gannet/Sweden/SVA220623SZ0232/FB001984/N-2 | A / H5N1 | Europe / Sweden / Hallands Lan / Falkenbergs Kommun | Siamak Zohari (National Veterinary Institute) |
| EPI_ISL_148126 | A/wild duck/Shandong/628/2011 | A / H5N1 | Asia / China | |
| EPI_ISL_14819895 | A/Mallard/Ningxia/BY246/2020 | A / H5N8 | Asia / China / Ningxia Hui Autonomous Region | Zeyu Yang (Chinese Academy of Forestry / Research Institute of Forest Ecology, Environment and Protection) |
| EPI_ISL_14819898 | A/Wild Duck/Ningxia/Y95/2021 | A / H5N8 | Asia / China / Ningxia Hui Autonomous Region | Zeyu Yang (Chinese Academy of Forestry / Research Institute of Forest Ecology, Environment and Protection) |
| EPI_ISL_14819899 | A/Wild Duck/Ningxia/Y99/2021 | A / H5N8 | Asia / China / Ningxia Hui Autonomous Region | Zeyu Yang (Chinese Academy of Forestry / Research Institute of Forest Ecology, Environment and Protection) |
| EPI_ISL_14819900 | A/Spot-billed Duck/Ningxia/Y16/2021 | A / H5N8 | Asia / China / Ningxia Hui Autonomous Region | Zeyu Yang (Chinese Academy of Forestry / Research Institute of Forest Ecology, Environment and Protection) |
| EPI_ISL_14822356 | A/Spot-billed Duck/Ningxia/Y26/2021 | A / H5N8 | Asia / China / Ningxia Hui Autonomous Region | Zeyu Yang (Chinese Academy of Forestry / Research Institute of Forest Ecology, Environment and Protection) |
| EPI_ISL_14822357 | A/Wild Duck/Ningxia/Y54/2021 | A / H5N8 | Asia / China / Ningxia Hui Autonomous Region | Zeyu Yang (Chinese Academy of Forestry / Research Institute of Forest Ecology, Environment and Protection) |
| EPI_ISL_14822426 | A/Bar-headed Goose/Tibet/P2320/2021 | A / H5N8 | Asia / China / Tibet Autonomous Region | Zeyu Yang (Chinese Academy of Forestry / Research Institute of Forest Ecology, Environment and Protection) |
| EPI_ISL_14822451 | A/Bar-headed Goose/Tibet/P1908/2021 | A / H5N8 | Asia / China / Tibet Autonomous Region | Zeyu Yang (Chinese Academy of Forestry / Research Institute of Forest Ecology, Environment and Protection) |
| EPI_ISL_14822452 | A/Bar-headed Goose/Tibet/P1910/2021 | A / H5N8 | Asia / China / Tibet Autonomous Region | Zeyu Yang (Chinese Academy of Forestry / Research Institute of Forest Ecology, Environment and Protection) |
| EPI_ISL_14822547 | A/Bar-headed Goose/Tibet/P1987/2021 | A / H5N8 | Asia / China / Tibet Autonomous Region | Zeyu Yang (Chinese Academy of Forestry / Research Institute of Forest Ecology, Environment and Protection) |
| EPI_ISL_14822549 | A/bar_headed_goose/Tibet/T1640/2021 | A / H5N8 | Asia / China / Tibet Autonomous Region | Zeyu Yang (Chinese Academy of Forestry / Research Institute of Forest Ecology, Environment and Protection) |
| EPI_ISL_14822756 | A/bar_headed_goose/Tibet/T1707/2021 | A / H5N8 | Asia / China / Tibet Autonomous Region | Zeyu Yang (Chinese Academy of Forestry / Research Institute of Forest Ecology, Environment and Protection) |
| EPI_ISL_14822757 | A/brown-headed_gull/Tibet/N38/2021 | A / H5N8 | Asia / China / Tibet Autonomous Region | Zeyu Yang (Chinese Academy of Forestry / Research Institute of Forest Ecology, Environment and Protection) |
| EPI_ISL_14823405 | A/Wild geese/Hubei/H358/2021 | A / H5N8 | Asia / China / Hubei Province | Zeyu Yang (Chinese Academy of Forestry / Research Institute of Forest Ecology, Environment and Protection) |
| EPI_ISL_14823421 | A/Wild geese/Hubei/H418/2021 | A / H5N8 | Asia / China / Hubei Province | Zeyu Yang (Chinese Academy of Forestry / Research Institute of Forest Ecology, Environment and Protection) |
| EPI_ISL_14823422 | A/Wild geese/Hubei/H549/2021 | A / H5N8 | Asia / China / Hubei Province | Zeyu Yang (Chinese Academy of Forestry / Research Institute of Forest Ecology, Environment and Protection) |
| EPI_ISL_14823424 | A/Whooper swan/Sanmenxia/Y1/2020 | A / H5N8 | Asia / China / Henan Province | Zeyu Yang (Chinese Academy of Forestry / Research Institute of Forest Ecology, Environment and Protection) |
| EPI_ISL_14823425 | A/Whooper swan/Sanmenxia/G2/2020 | A / H5N8 | Asia / China / Henan Province | Zeyu Yang (Chinese Academy of Forestry / Research Institute of Forest Ecology, Environment and Protection) |
| EPI_ISL_14823426 | A/Whooper swan/Sanmenxia/Y6/2020 | A / H5N8 | Asia / China / Henan Province | Zeyu Yang (Chinese Academy of Forestry / Research Institute of Forest Ecology, Environment and Protection) |
| EPI_ISL_14823549 | A/Whooper swan/Sanmenxia/Y8/2020 | A / H5N8 | Asia / China / Henan Province | Zeyu Yang (Chinese Academy of Forestry / Research Institute of Forest Ecology, Environment and Protection) |
| EPI_ISL_14823646 | A/Whooper swan/Sanmenxia/Y10/2020 | A / H5N8 | Asia / China / Henan Province | Zeyu Yang (Chinese Academy of Forestry / Research Institute of Forest Ecology, Environment and Protection) |
| EPI_ISL_14823672 | A/Whooper swan/Sanmenxia/Y11/2020 | A / H5N8 | Asia / China / Henan Province | Zeyu Yang (Chinese Academy of Forestry / Research Institute of Forest Ecology, Environment and Protection) |
| EPI_ISL_14823811 | A/Whooper swan/Sanmenxia/Y12/2020 | A / H5N8 | Asia / China / Henan Province | Zeyu Yang (Chinese Academy of Forestry / Research Institute of Forest Ecology, Environment and Protection) |
| EPI_ISL_14823812 | A/Whooper swan/Sanmenxia/G13/2020 | A / H5N8 | Asia / China / Henan Province | Zeyu Yang (Chinese Academy of Forestry / Research Institute of Forest Ecology, Environment and Protection) |
| EPI_ISL_14823813 | A/Whooper swan/Sanmenxia/Y14/2020 | A / H5N8 | Asia / China / Henan Province | Zeyu Yang (Chinese Academy of Forestry / Research Institute of Forest Ecology, Environment and Protection) |
| EPI_ISL_14823863 | A/Whooper swan/Sanmenxia/Y51/2020 | A / H5N8 | Asia / China / Henan Province | Zeyu Yang (Chinese Academy of Forestry / Research Institute of Forest Ecology, Environment and Protection) |
| EPI_ISL_14824026 | A/Whooper swan/Sanmenxia/G15/2020 | A / H5N8 | Asia / China / Henan Province | Zeyu Yang (Chinese Academy of Forestry / Research Institute of Forest Ecology, Environment and Protection) |
| EPI_ISL_14824091 | A/Whooper swan/Sanmenxia/Y15/2020 | A / H5N8 | Asia / China / Henan Province | Zeyu Yang (Chinese Academy of Forestry / Research Institute of Forest Ecology, Environment and Protection) |
| EPI_ISL_14824095 | A/Whooper swan/Sanmenxia/Y16/2020 | A / H5N8 | Asia / China / Henan Province | Zeyu Yang (Chinese Academy of Forestry / Research Institute of Forest Ecology, Environment and Protection) |
| EPI_ISL_14824122 | A/Whooper swan/Sanmenxia/Y52/2020 | A / H5N8 | Asia / China / Henan Province | Zeyu Yang (Chinese Academy of Forestry / Research Institute of Forest Ecology, Environment and Protection) |
| EPI_ISL_14835331 | A/ostrich/Egypt/Egypt-Ostrich-2021-HA/2021 | A / H5N8 | Africa / Egypt | |
| EPI_ISL_14835960 | A/Whooper swan/Sanmenxia/H1/2020 | A / H5N8 | Asia / China / Henan Province | Zeyu Yang (Chinese Academy of Forestry / Research Institute of Forest Ecology, Environment and Protection) |
| EPI_ISL_14835963 | A/Whooper swan/Sanmenxia/H615/2020 | A / H5N8 | Asia / China / Henan Province | Zeyu Yang (Chinese Academy of Forestry / Research Institute of Forest Ecology, Environment and Protection) |
| EPI_ISL_14835964 | A/Whooper swan/Sanmenxia/H735/2020 | A / H5N8 | Asia / China / Henan Province | Zeyu Yang (Chinese Academy of Forestry / Research Institute of Forest Ecology, Environment and Protection) |
| EPI_ISL_14835965 | A/Whooper swan/Sanmenxia/H810/2020 | A / H5N8 | Asia / China / Henan Province | Zeyu Yang (Chinese Academy of Forestry / Research Institute of Forest Ecology, Environment and Protection) |
| EPI_ISL_14835968 | A/Whooper swan/Sanmenxia/Y24/2020 | A / H5N8 | Asia / China / Henan Province | Zeyu Yang (Chinese Academy of Forestry / Research Institute of Forest Ecology, Environment and Protection) |
| EPI_ISL_14835974 | A/Whooper swan/Sanmenxia/Y18/2020 | A / H5N8 | Asia / China / Henan Province | Zeyu Yang (Chinese Academy of Forestry / Research Institute of Forest Ecology, Environment and Protection) |
| EPI_ISL_14835976 | A/Whooper swan/Sanmenxia/Y56/2020 | A / H5N8 | Asia / China / Henan Province | Zeyu Yang (Chinese Academy of Forestry / Research Institute of Forest Ecology, Environment and Protection) |
| EPI_ISL_14836024 | A/Whooper swan/Sanmenxia/Y19/2020 | A / H5N8 | Asia / China / Henan Province | Zeyu Yang (Chinese Academy of Forestry / Research Institute of Forest Ecology, Environment and Protection) |
| EPI_ISL_14836025 | A/Whooper swan/Sanmenxia/Y20/2020 | A / H5N8 | Asia / China / Henan Province | Zeyu Yang (Chinese Academy of Forestry / Research Institute of Forest Ecology, Environment and Protection) |
| EPI_ISL_14836026 | A/Whooper swan/Sanmenxia/Y21/2020 | A / H5N8 | Asia / China / Henan Province | Zeyu Yang (Chinese Academy of Forestry / Research Institute of Forest Ecology, Environment and Protection) |
| EPI_ISL_14836027 | A/Whooper swan/Sanmenxia/Y23-1/2020 | A / H5N8 | Asia / China / Henan Province | Zeyu Yang (Chinese Academy of Forestry / Research Institute of Forest Ecology, Environment and Protection) |
| EPI_ISL_14836077 | A/Whooper swan/Sanmenxia/Y23-2/2020 | A / H5N8 | Asia / China / Henan Province | Zeyu Yang (Chinese Academy of Forestry / Research Institute of Forest Ecology, Environment and Protection) |
| EPI_ISL_14836078 | A/Whooper swan/Sanmenxia/G27/2020 | A / H5N8 | Asia / China / Henan Province | Zeyu Yang (Chinese Academy of Forestry / Research Institute of Forest Ecology, Environment and Protection) |
| EPI_ISL_14836079 | A/Whooper swan/Sanmenxia/G28/2020 | A / H5N8 | Asia / China / Henan Province | Zeyu Yang (Chinese Academy of Forestry / Research Institute of Forest Ecology, Environment and Protection) |
| EPI_ISL_14836080 | A/Whooper swan/Sanmenxia/G29/2020 | A / H5N8 | Asia / China / Henan Province | Zeyu Yang (Chinese Academy of Forestry / Research Institute of Forest Ecology, Environment and Protection) |
| EPI_ISL_14836081 | A/Whooper swan/Sanmenxia/Y26/2020 | A / H5N8 | Asia / China / Henan Province | Zeyu Yang (Chinese Academy of Forestry / Research Institute of Forest Ecology, Environment and Protection) |
| EPI_ISL_14836082 | A/Whooper swan/Sanmenxia/Y27/2020 | A / H5N8 | Asia / China / Henan Province | Zeyu Yang (Chinese Academy of Forestry / Research Institute of Forest Ecology, Environment and Protection) |
| EPI_ISL_14836083 | A/Whooper swan/Sanmenxia/Y31/2020 | A / H5N8 | Asia / China / Henan Province | Zeyu Yang (Chinese Academy of Forestry / Research Institute of Forest Ecology, Environment and Protection) |
| EPI_ISL_14836086 | A/Whooper swan/Sanmenxia/Y36/2020 | A / H5N8 | Asia / China / Henan Province | Zeyu Yang (Chinese Academy of Forestry / Research Institute of Forest Ecology, Environment and Protection) |
| EPI_ISL_14836088 | A/Whooper swan/Sanmenxia/Y48A/2020 | A / H5N8 | Asia / China / Henan Province | Zeyu Yang (Chinese Academy of Forestry / Research Institute of Forest Ecology, Environment and Protection) |
| EPI_ISL_14836098 | A/Whooper swan/Sanmenxia/Y54/2020 | A / H5N8 | Asia / China / Henan Province | Zeyu Yang (Chinese Academy of Forestry / Research Institute of Forest Ecology, Environment and Protection) |
| EPI_ISL_14836101 | A/Whooper swan/Sanmenxia/Y57/2020 | A / H5N8 | Asia / China / Henan Province | Zeyu Yang (Chinese Academy of Forestry / Research Institute of Forest Ecology, Environment and Protection) |
| EPI_ISL_14836104 | A/Whooper swan/Sanmenxia/G48B/2020 | A / H5N8 | Asia / China / Henan Province | Zeyu Yang (Chinese Academy of Forestry / Research Institute of Forest Ecology, Environment and Protection) |
| EPI_ISL_14836369 | A/Whooper swan/Sanmenxia/Y48B/2020 | A / H5N8 | Asia / China / Henan Province | Zeyu Yang (Chinese Academy of Forestry / Research Institute of Forest Ecology, Environment and Protection) |
| EPI_ISL_14836371 | A/Whooper swan/Sanmenxia/Y49/2020 | A / H5N8 | Asia / China / Henan Province | Zeyu Yang (Chinese Academy of Forestry / Research Institute of Forest Ecology, Environment and Protection) |
| EPI_ISL_14836374 | A/Whooper swan/Sanmenxia/G25/2020 | A / H5N8 | Asia / China / Henan Province | Zeyu Yang (Chinese Academy of Forestry / Research Institute of Forest Ecology, Environment and Protection) |
| EPI_ISL_14836376 | A/Whooper swan/Sanmenxia/Y25/2020 | A / H5N8 | Asia / China / Henan Province | Zeyu Yang (Chinese Academy of Forestry / Research Institute of Forest Ecology, Environment and Protection) |
| EPI_ISL_14836426 | A/Whooper swan/Sanmenxia/B560/2020 | A / H5N8 | Asia / China / Henan Province | Zeyu Yang (Chinese Academy of Forestry / Research Institute of Forest Ecology, Environment and Protection) |
| EPI_ISL_14836593 | A/Whooper swan/Sanmenxia/B312/2020 | A / H5N8 | Asia / China / Henan Province | Zeyu Yang (Chinese Academy of Forestry / Research Institute of Forest Ecology, Environment and Protection) |
| EPI_ISL_14838179 | A/Whooper swan/Sanmenxia/B497/2020 | A / H5N8 | Asia / China / Henan Province | Zeyu Yang (Chinese Academy of Forestry / Research Institute of Forest Ecology, Environment and Protection) |
| EPI_ISL_14839943 | A/Whooper swan/Sanmenxia/B1109/2021 | A / H5N8 | Asia / China / Henan Province | Zeyu Yang (Chinese Academy of Forestry / Research Institute of Forest Ecology, Environment and Protection) |
| EPI_ISL_14840034 | A/Whooper swan/Sanmenxia/B1770/2020 | A / H5N8 | Asia / China / Henan Province | Zeyu Yang (Chinese Academy of Forestry / Research Institute of Forest Ecology, Environment and Protection) |
| EPI_ISL_14840053 | A/Whooper swan/Sanmenxia/B2465/2021 | A / H5N8 | Asia / China / Henan Province | Zeyu Yang (Chinese Academy of Forestry / Research Institute of Forest Ecology, Environment and Protection) |
| EPI_ISL_14841913 | A/mallard/22P019377/France/2022 | A / H5N1 | Europe / France / Picardie / Departement de la Somme | Francois-Xavier Briand (ANSES Agence Nationale De Securite Sanitaire De L'alimentation / Laboratoire de Ploufragan-Plouzané) |

| | | | | |
|---|---|---|---|---|
| EPI_ISL_14845579 | A/whooper_swan/Rongcheng/M82/2021 | A / H5 | Asia / China / Shandong Province | Zeyu Yang (Chinese Academy of Forestry / Research Institute of Forest Ecology, Environment and Protection) |
| EPI_ISL_14857053 | A/chicken/Kursk/230-2V/2022 | A / H5N1 | Europe / Russian Federation / Kursk Oblast | Natalia Goncharova (State Research Center of Virology and Biotechnology (VECTOR) / Emerging Zoonotic Diseases and Influenza) |
| EPI_ISL_14857054 | A/chicken/Kursk/230-4V/2022 | A / H5N1 | Europe / Russian Federation / Kursk Oblast | Natalia Goncharova (State Research Center of Virology and Biotechnology (VECTOR) / Emerging Zoonotic Diseases and Influenza) |
| EPI_ISL_14857055 | A/chicken/Kursk/230-5V/2022 | A / H5N1 | Europe / Russian Federation / Kursk Oblast | Natalia Goncharova (State Research Center of Virology and Biotechnology (VECTOR) / Emerging Zoonotic Diseases and Influenza) |
| EPI_ISL_14857056 | A/chicken/Kursk/230-7V/2022 | A / H5N1 | Europe / Russian Federation / Kursk Oblast | Natalia Goncharova (State Research Center of Virology and Biotechnology (VECTOR) / Emerging Zoonotic Diseases and Influenza) |
| EPI_ISL_14857057 | A/chicken/Kursk/230-8V/2022 | A / H5N1 | Europe / Russian Federation / Kursk Oblast | Natalia Goncharova (State Research Center of Virology and Biotechnology (VECTOR) / Emerging Zoonotic Diseases and Influenza) |
| EPI_ISL_14857058 | A/chicken/Kursk/230-10V/2022 | A / H5N1 | Europe / Russian Federation / Kursk Oblast | Natalia Goncharova (State Research Center of Virology and Biotechnology (VECTOR) / Emerging Zoonotic Diseases and Influenza) |
| EPI_ISL_14857059 | A/chicken/Kursk/230-15V/2022 | A / H5N1 | Europe / Russian Federation / Kursk Oblast | Natalia Goncharova (State Research Center of Virology and Biotechnology (VECTOR) / Emerging Zoonotic Diseases and Influenza) |
| EPI_ISL_14857060 | A/chicken/Kursk/234-19V/2022 | A / H5N1 | Europe / Russian Federation / Kursk Oblast | Natalia Goncharova (State Research Center of Virology and Biotechnology (VECTOR) / Emerging Zoonotic Diseases and Influenza) |
| EPI_ISL_14857061 | A/quail/Kursk/234-20V/2022 | A / H5N1 | Europe / Russian Federation / Kursk Oblast | Natalia Goncharova (State Research Center of Virology and Biotechnology (VECTOR) / Emerging Zoonotic Diseases and Influenza) |
| EPI_ISL_14857062 | A/chicken/Magadan/235-57V/2022 | A / H5N1 | Europe / Russian Federation / Magadan Oblast | Natalia Goncharova (State Research Center of Virology and Biotechnology (VECTOR) / Emerging Zoonotic Diseases and Influenza) |
| EPI_ISL_14857063 | A/chicken/Magadan/235-58V/2022 | A / H5N1 | Europe / Russian Federation / Magadan Oblast | Natalia Goncharova (State Research Center of Virology and Biotechnology (VECTOR) / Emerging Zoonotic Diseases and Influenza) |
| EPI_ISL_14857064 | A/chicken/Magadan/235-59V/2022 | A / H5N1 | Europe / Russian Federation / Magadan Oblast | Natalia Goncharova (State Research Center of Virology and Biotechnology (VECTOR) / Emerging Zoonotic Diseases and Influenza) |
| EPI_ISL_14857065 | A/chicken/Magadan/235-60V/2022 | A / H5N1 | Europe / Russian Federation / Magadan Oblast | Natalia Goncharova (State Research Center of Virology and Biotechnology (VECTOR) / Emerging Zoonotic Diseases and Influenza) |
| EPI_ISL_14864637 | A/goose/England/317610/2022 | A / H5N1 | Europe / United Kingdom / England | Alex Byrne (Animal and Plant Health Agency (APHA) / Virology Department) |
| EPI_ISL_14864646 | A/Canada_goose/England/320660/2022 | A / H5N1 | Europe / United Kingdom / England | Alex Byrne (Animal and Plant Health Agency (APHA) / Virology Department) |
| EPI_ISL_14864650 | A/turkey/England/111923/2022 | A / H5N1 | Europe / United Kingdom / England | Alex Byrne (Animal and Plant Health Agency (APHA) / Virology Department) |
| EPI_ISL_14867042 | A/duck/Kagoshima/NIES229/2020 | A / H5N8 | Asia / Japan / Kagoshima | |
| EPI_ISL_14871335 | A/poultry/Benin/21-A-08-035-O/2021 | A / H5N1 | Africa / Benin | |
| EPI_ISL_14871336 | A/poultry/Benin/21-A-08-009-O/2021 | A / H5N1 | Africa / Benin | |
| EPI_ISL_14871337 | A/poultry/Benin/21-A-08-033-O/2021 | A / H5N1 | Africa / Benin | |
| EPI_ISL_14871338 | A/poultry/Benin/21-A-08-034-O/2021 | A / H5N1 | Africa / Benin | |
| EPI_ISL_14871339 | A/poultry/Benin/21-A-09-031-O/2021 | A / H5N1 | Africa / Benin | |
| EPI_ISL_14886367 | A/duck/Bangladesh/51602/2021 | A / H5N1 | Asia / Bangladesh | |
| EPI_ISL_14886675 | A/duck/Bangladesh/51601/2021 | A / H5N1 | Asia / Bangladesh | |
| EPI_ISL_14886825 | A/duck/Bangladesh/51600/2021 | A / H5N1 | Asia / Bangladesh | |
| EPI_ISL_14901700 | A/duck/Egypt/Ismailia/2021 | A / H5N8 | Africa / Egypt | |
| EPI_ISL_14917968 | A/common_murre/Poland/MB151/2022 | A / H5N1 | Europe / Poland / Pomeranian Voivodeship | Edyta ?wi?to? (National Veterinary Research Institut Poland, PIWet-PIB) |
| EPI_ISL_14917979 | A/herring_gull/Poland/MB138/2022 | A / H5N1 | Europe / Poland / Pomeranian Voivodeship | Edyta ?wi?to? (National Veterinary Research Institut Poland, PIWet-PIB) |
| EPI_ISL_14917999 | A/black-headed_gull/Poland/MB139/2022 | A / H5N1 | Europe / Poland / Pomeranian Voivodeship | Edyta ?wi?to? (National Veterinary Research Institut Poland, PIWet-PIB) |
| EPI_ISL_14933724 | A/duck/France/22P020165/2022 | A / H5N1 | Europe / France / Rhone-Alpes / Departement de l'Ain | Francois-Xavier Briand (ANSES Agence Nationale De Securite Sanitaire De L'alimentation / Laboratoire de Ploufragan-Plouzané) |
| EPI_ISL_14936999 | A/bald eagle/Kansas/W22-197/2022 | A / H5N1 | North America / United States | |
| EPI_ISL_14937000 | A/snow goose/Kansas/W22-199D/2022 | A / H5N1 | North America / United States | |
| EPI_ISL_14937001 | A/snow goose/Kansas/W22-199F/2022 | A / H5N1 | North America / United States | |
| EPI_ISL_14937002 | A/bald eagle/Georgia/W22-194B/2022 | A / H5N1 | North America / United States | |
| EPI_ISL_14937069 | A/bald eagle/Florida/W22-191/2022 | A / H5N1 | North America / United States | |
| EPI_ISL_14937070 | A/bald eagle/Florida/W22-195/2022 | A / H5N1 | North America / United States | |
| EPI_ISL_14937071 | A/bald eagle/Georgia/W22-202/2022 | A / H5N1 | North America / United States | |
| EPI_ISL_14937099 | A/snow goose/Kansas/W22-199A/2022 | A / H5N1 | North America / United States | |
| EPI_ISL_14937100 | A/snow goose/Kansas/W22-199B/2022 | A / H5N1 | North America / United States | |
| EPI_ISL_14937101 | A/bald eagle/South Carolina/W22-205/2022 | A / H5N1 | North America / United States | |
| EPI_ISL_14937102 | A/snow goose/Kansas/W22-199E/2022 | A / H5N1 | North America / United States | |
| EPI_ISL_14937103 | A/bald eagle/Georgia/W22-194A/2022 | A / H5N1 | North America / United States | |
| EPI_ISL_14937104 | A/snow goose/Kansas/W22-199C/2022 | A / H5N1 | North America / United States | |
| EPI_ISL_14937105 | A/bald eagle/Florida/W22-189/2022 | A / H5N1 | North America / United States | |
| EPI_ISL_14937233 | A/Turkey/Egypt/Giza/2021 | A / H5N8 | Africa / Egypt | |
| EPI_ISL_15004053 | A/Pekin duck/Indiana/22-010611-001-original/2022 | A / H5N1 | North America / United States / Indiana / Elkhart County | Mary Lea Killian (National Veterinary Services Laboratories - USDA / Diagnostic Virology Laboratory) |
| EPI_ISL_15004054 | A/Pekin duck/Indiana/22-010624-001-original/2022 | A / H5N1 | North America / United States / Indiana / Elkhart County | Mary Lea Killian (National Veterinary Services Laboratories - USDA / Diagnostic Virology Laboratory) |
| EPI_ISL_15004055 | A/turkey/South Dakota/22-010639-001-original/2022 | A / H5N1 | North America / United States / South Dakota / Clark County | Mary Lea Killian (National Veterinary Services Laboratories - USDA / Diagnostic Virology Laboratory) |
| EPI_ISL_15004056 | A/turkey/South Dakota/22-010639-002-original/2022 | A / H5N1 | North America / United States / South Dakota / Clark County | Mary Lea Killian (National Veterinary Services Laboratories - USDA / Diagnostic Virology Laboratory) |
| EPI_ISL_15004057 | A/turkey/Minnesota/22-010652-001-original/2022 | A / H5N1 | North America / United States / Minnesota / Otter Tail County | Mary Lea Killian (National Veterinary Services Laboratories - USDA / Diagnostic Virology Laboratory) |
| EPI_ISL_15004058 | A/turkey/Minnesota/22-010652-002-original/2022 | A / H5N1 | North America / United States / Minnesota / Otter Tail County | Mary Lea Killian (National Veterinary Services Laboratories - USDA / Diagnostic Virology Laboratory) |
| EPI_ISL_15004059 | A/turkey/Minnesota/22-010654-001-original/2022 | A / H5N1 | North America / United States / Minnesota / Kandiyohi County | Mary Lea Killian (National Veterinary Services Laboratories - USDA / Diagnostic Virology Laboratory) |
| EPI_ISL_15004060 | A/chicken/North Dakota/22-010657-001-original/2022 | A / H5N1 | North America / United States / North Dakota / Stutsman County | Mary Lea Killian (National Veterinary Services Laboratories - USDA / Diagnostic Virology Laboratory) |
| EPI_ISL_15004061 | A/goose/North Dakota/22-010657-002-original/2022 | A / H5N1 | North America / United States / North Dakota / Stutsman County | Mary Lea Killian (National Veterinary Services Laboratories - USDA / Diagnostic Virology Laboratory) |
| EPI_ISL_15004062 | A/chicken/Colorado/22-010668-001-original/2022 | A / H5N1 | North America / United States / Colorado / Pitkin County | Mary Lea Killian (National Veterinary Services Laboratories - USDA / Diagnostic Virology Laboratory) |
| EPI_ISL_15004063 | A/turkey/South Dakota/22-010765-001-original/2022 | A / H5N1 | North America / United States / South Dakota / Edmunds County | Mary Lea Killian (National Veterinary Services Laboratories - USDA / Diagnostic Virology Laboratory) |
| EPI_ISL_15004064 | A/turkey/Minnesota/22-010770-002-original/2022 | A / H5N1 | North America / United States / Minnesota / Yankton County | Mary Lea Killian (National Veterinary Services Laboratories - USDA / Diagnostic Virology Laboratory) |
| EPI_ISL_15004065 | A/turkey/Minnesota/22-010770-003-original/2022 | A / H5N1 | North America / United States / Minnesota / Yankton County | Mary Lea Killian (National Veterinary Services Laboratories - USDA / Diagnostic Virology Laboratory) |
| EPI_ISL_15004066 | A/turkey/Minnesota/22-010771-001-original/2022 | A / H5N1 | North America / United States / Minnesota / Le Sueur County | Mary Lea Killian (National Veterinary Services Laboratories - USDA / Diagnostic Virology Laboratory) |
| EPI_ISL_15004067 | A/turkey/Minnesota/22-010771-003-original/2022 | A / H5N1 | North America / United States / Minnesota / Le Sueur County | Mary Lea Killian (National Veterinary Services Laboratories - USDA / Diagnostic Virology Laboratory) |
| EPI_ISL_15004068 | A/turkey/Minnesota/22-010772-004-original/2022 | A / H5N1 | North America / United States / Minnesota / Stearns County | Mary Lea Killian (National Veterinary Services Laboratories - USDA / Diagnostic Virology Laboratory) |
| EPI_ISL_15004069 | A/turkey/Minnesota/22-010773-001-original/2022 | A / H5N1 | North America / United States / Minnesota / Swift County | Mary Lea Killian (National Veterinary Services Laboratories - USDA / Diagnostic Virology Laboratory) |
| EPI_ISL_15004070 | A/goose/Michigan/22-010845-001-original/2022 | A / H5N1 | North America / United States / Michigan / Menominee County | Mary Lea Killian (National Veterinary Services Laboratories - USDA / Diagnostic Virology Laboratory) |
| EPI_ISL_15004071 | A/goose/Michigan/22-010845-002-original/2022 | A / H5N1 | North America / United States / Michigan / Menominee County | Mary Lea Killian (National Veterinary Services Laboratories - USDA / Diagnostic Virology Laboratory) |
| EPI_ISL_15004072 | A/duck/Michigan/22-010845-003-original/2022 | A / H5N1 | North America / United States / Michigan / Menominee County | Mary Lea Killian (National Veterinary Services Laboratories - USDA / Diagnostic Virology Laboratory) |
| EPI_ISL_15004372 | A/Amazon parrot/Michigan/22-010848-001-original/2022 | A / H5N1 | North America / United States / Michigan / Washtenaw County | Mary Lea Killian (National Veterinary Services Laboratories - USDA / Diagnostic Virology Laboratory) |
| EPI_ISL_15004373 | A/Catalina macaw/Michigan/22-010848-002-original/2022 | A / H5N1 | North America / United States / Michigan / Washtenaw County | Mary Lea Killian (National Veterinary Services Laboratories - USDA / Diagnostic Virology Laboratory) |
| EPI_ISL_15004374 | A/chicken/Nebraska/22-010905-001-original/2022 | A / H5N1 | North America / United States / Nebraska / Dixon County | Mary Lea Killian (National Veterinary Services Laboratories - USDA / Diagnostic Virology Laboratory) |
| EPI_ISL_15004375 | A/chicken/Montana/22-010923-005-original/2022 | A / H5N1 | North America / United States / Montana / Toole County | Mary Lea Killian (National Veterinary Services Laboratories - USDA / Diagnostic Virology Laboratory) |
| EPI_ISL_15004376 | A/chicken/Minnesota/22-010928-001-original/2022 | A / H5N1 | North America / United States / Minnesota / Benton County | Mary Lea Killian (National Veterinary Services Laboratories - USDA / Diagnostic Virology Laboratory) |
| EPI_ISL_15004377 | A/turkey/Minnesota/22-010928-002-original/2022 | A / H5N1 | North America / United States / Minnesota / Benton County | Mary Lea Killian (National Veterinary Services Laboratories - USDA / Diagnostic Virology Laboratory) |

| | | | | |
|---|---|---|---|---|
| EPI_ISL_15004378 | A/guinea fowl/Minnesota/22-010928-003-original/2022 | A / H5N1 | North America / United States / Minnesota / Benton County | Mary Lea Killian (National Veterinary Services Laboratories - USDA / Diagnostic Virology Laboratory) |
| EPI_ISL_15004379 | A/domestic duck/Minnesota/22-010928-004-original/2022 | A / H5N1 | North America / United States / Minnesota / Benton County | Mary Lea Killian (National Veterinary Services Laboratories - USDA / Diagnostic Virology Laboratory) |
| EPI_ISL_15004380 | A/chicken/North Carolina/22-010929-001-original/2022 | A / H5N1 | North America / United States / North Carolina / Wayne County | Mary Lea Killian (National Veterinary Services Laboratories - USDA / Diagnostic Virology Laboratory) |
| EPI_ISL_15004381 | A/chicken/North Carolina/22-010929-002-original/2022 | A / H5N1 | North America / United States / North Carolina / Wayne County | Mary Lea Killian (National Veterinary Services Laboratories - USDA / Diagnostic Virology Laboratory) |
| EPI_ISL_15004383 | A/turkey/Wisconsin/22-010935-002-original/2022 | A / H5N1 | North America / United States / Wisconsin / Barron County | Mary Lea Killian (National Veterinary Services Laboratories - USDA / Diagnostic Virology Laboratory) |
| EPI_ISL_15004384 | A/chicken/North Dakota/22-010977-001-original/2022 | A / H5N1 | North America / United States / North Dakota / Sheridan County | Mary Lea Killian (National Veterinary Services Laboratories - USDA / Diagnostic Virology Laboratory) |
| EPI_ISL_15004385 | A/chicken/North Dakota/22-010977-002-original/2022 | A / H5N1 | North America / United States / North Dakota / Sheridan County | Mary Lea Killian (National Veterinary Services Laboratories - USDA / Diagnostic Virology Laboratory) |
| EPI_ISL_15004386 | A/turkey/Minnesota/22-010991-001-original/2022 | A / H5N1 | North America / United States / Minnesota / Morrison County | Mary Lea Killian (National Veterinary Services Laboratories - USDA / Diagnostic Virology Laboratory) |
| EPI_ISL_15004387 | A/turkey/Minnesota/22-010991-002-original/2022 | A / H5N1 | North America / United States / Minnesota / Morrison County | Mary Lea Killian (National Veterinary Services Laboratories - USDA / Diagnostic Virology Laboratory) |
| EPI_ISL_15004388 | A/turkey/Minnesota/22-010992-001-original/2022 | A / H5N1 | North America / United States / Minnesota / Yellow Medicine County | Mary Lea Killian (National Veterinary Services Laboratories - USDA / Diagnostic Virology Laboratory) |
| EPI_ISL_15004389 | A/turkey/Minnesota/22-010992-002-original/2022 | A / H5N1 | North America / United States / Minnesota / Yellow Medicine County | Mary Lea Killian (National Veterinary Services Laboratories - USDA / Diagnostic Virology Laboratory) |
| EPI_ISL_15004390 | A/chicken/Minnesota/22-010993-001-original/2022 | A / H5N1 | North America / United States / Minnesota / Morrison County | Mary Lea Killian (National Veterinary Services Laboratories - USDA / Diagnostic Virology Laboratory) |
| EPI_ISL_15004391 | A/chicken/Minnesota/22-010993-004-original/2022 | A / H5N1 | North America / United States / Minnesota / Morrison County | Mary Lea Killian (National Veterinary Services Laboratories - USDA / Diagnostic Virology Laboratory) |
| EPI_ISL_15004392 | A/chicken/Minnesota/22-010994-002-original/2022 | A / H5N1 | North America / United States / Minnesota / Morrison County | Mary Lea Killian (National Veterinary Services Laboratories - USDA / Diagnostic Virology Laboratory) |
| EPI_ISL_15004393 | A/turkey/Minnesota/22-010995-001-original/2022 | A / H5N1 | North America / United States / Minnesota / Kandiyohi County | Mary Lea Killian (National Veterinary Services Laboratories - USDA / Diagnostic Virology Laboratory) |
| EPI_ISL_15004394 | A/turkey/Minnesota/22-010995-002-original/2022 | A / H5N1 | North America / United States / Minnesota / Kandiyohi County | Mary Lea Killian (National Veterinary Services Laboratories - USDA / Diagnostic Virology Laboratory) |
| EPI_ISL_15004395 | A/pheasant/South Dakota/22-011098-001-original/2022 | A / H5N1 | North America / United States / South Dakota / Deuel County | Mary Lea Killian (National Veterinary Services Laboratories - USDA / Diagnostic Virology Laboratory) |
| EPI_ISL_15004396 | A/chicken/North Dakota/22-011109-001-original/2022 | A / H5N1 | North America / United States / North Dakota / Barnes County | Mary Lea Killian (National Veterinary Services Laboratories - USDA / Diagnostic Virology Laboratory) |
| EPI_ISL_15004397 | A/turkey/Kansas/22-011119-004-original/2022 | A / H5N1 | North America / United States / Kansas / McPherson County | Mary Lea Killian (National Veterinary Services Laboratories - USDA / Diagnostic Virology Laboratory) |
| EPI_ISL_15004398 | A/chicken/Michigan/22-011121-001-original/2022 | A / H5N1 | North America / United States / Michigan / Menominee County | Mary Lea Killian (National Veterinary Services Laboratories - USDA / Diagnostic Virology Laboratory) |
| EPI_ISL_15004399 | A/domestic duck/Michigan/22-011121-002-original/2022 | A / H5N1 | North America / United States / Michigan / Menominee County | Mary Lea Killian (National Veterinary Services Laboratories - USDA / Diagnostic Virology Laboratory) |
| EPI_ISL_15004400 | A/chicken/Michigan/22-011121-003-original/2022 | A / H5N1 | North America / United States / Michigan / Menominee County | Mary Lea Killian (National Veterinary Services Laboratories - USDA / Diagnostic Virology Laboratory) |
| EPI_ISL_15004401 | A/domestic goose/Michigan/22-011121-005-original/2022 | A / H5N1 | North America / United States / Michigan / Menominee County | Mary Lea Killian (National Veterinary Services Laboratories - USDA / Diagnostic Virology Laboratory) |
| EPI_ISL_15004402 | A/chicken/Montana/22-010923-004-original/2022 | A / H5N1 | North America / United States / Montana / Toole County | Mary Lea Killian (National Veterinary Services Laboratories - USDA / Diagnostic Virology Laboratory) |
| EPI_ISL_15004403 | A/turkey/Minnesota/22-011140-001-original/2022 | A / H5N1 | North America / United States / Minnesota / Blue Earth County | Mary Lea Killian (National Veterinary Services Laboratories - USDA / Diagnostic Virology Laboratory) |
| EPI_ISL_15004404 | A/turkey/Minnesota/22-011141-001-original/2022 | A / H5N1 | North America / United States / Minnesota / Otter Tail County | Mary Lea Killian (National Veterinary Services Laboratories - USDA / Diagnostic Virology Laboratory) |
| EPI_ISL_15004405 | A/turkey/Minnesota/22-011141-002-original/2022 | A / H5N1 | North America / United States / Minnesota / Otter Tail County | Mary Lea Killian (National Veterinary Services Laboratories - USDA / Diagnostic Virology Laboratory) |
| EPI_ISL_15004406 | A/turkey/Minnesota/22-011142-001-original/2022 | A / H5N1 | North America / United States / Minnesota / Waseca County | Mary Lea Killian (National Veterinary Services Laboratories - USDA / Diagnostic Virology Laboratory) |
| EPI_ISL_15004407 | A/turkey/Minnesota/22-011142-002-original/2022 | A / H5N1 | North America / United States / Minnesota / Waseca County | Mary Lea Killian (National Veterinary Services Laboratories - USDA / Diagnostic Virology Laboratory) |
| EPI_ISL_15004408 | A/chicken/Wisconsin/22-011213-001-original/2022 | A / H5N1 | North America / United States / Wisconsin / Sheboygan County | Mary Lea Killian (National Veterinary Services Laboratories - USDA / Diagnostic Virology Laboratory) |
| EPI_ISL_15004409 | A/chicken/Wisconsin/22-011213-002-original/2022 | A / H5N1 | North America / United States / Wisconsin / Sheboygan County | Mary Lea Killian (National Veterinary Services Laboratories - USDA / Diagnostic Virology Laboratory) |
| EPI_ISL_15004410 | A/chicken/North Dakota/22-011286-001-original/2022 | A / H5N1 | North America / United States / North Dakota / LaMoure County | Mary Lea Killian (National Veterinary Services Laboratories - USDA / Diagnostic Virology Laboratory) |
| EPI_ISL_15004411 | A/chicken/Wisconsin/22-011334-002-original/2022 | A / H5N1 | North America / United States / Wisconsin / Columbia County | Mary Lea Killian (National Veterinary Services Laboratories - USDA / Diagnostic Virology Laboratory) |
| EPI_ISL_15004412 | A/chicken/Wisconsin/22-011334-003-original/2022 | A / H5N1 | North America / United States / Wisconsin / Columbia County | Mary Lea Killian (National Veterinary Services Laboratories - USDA / Diagnostic Virology Laboratory) |
| EPI_ISL_15004413 | A/chicken/Minnesota/22-010994-001-original/2022 | A / H5N1 | North America / United States / Minnesota / Morrison County | Mary Lea Killian (National Veterinary Services Laboratories - USDA / Diagnostic Virology Laboratory) |
| EPI_ISL_15022831 | A/swan/Lithuania/1220PG1_22VIR7255-1/2022 | A / H5N1 | Europe / Lithuania | Giacomo Barbierato (Istituto Zooprofilattico Sperimentale delle Venezie / Ricerca e innovazione) |
| EPI_ISL_15023114 | A/stork/Lithuania/4371PG1_22VIR7255-3/2022 | A / H5N1 | Europe / Lithuania | Giacomo Barbierato (Istituto Zooprofilattico Sperimentale delle Venezie / Ricerca e innovazione) |
| EPI_ISL_15038822 | A/chicken/England/114595/2022 | A / H5N1 | Europe / United Kingdom / England | Alex Byrne (Animal and Plant Health Agency (APHA) / Virology Department) |
| EPI_ISL_15063427 | A/chicken/Ehime/TU10-2-13/2022 | A / H5N1 | Asia / Japan / Ehime | |
| EPI_ISL_15063428 | A/environment/Akita/TU1-49/2021 | A / H5N8 | Asia / Japan / Akita | |
| EPI_ISL_15063618 | A/snow goose/North Dakota/N22-06/2022 | A / H5N1 | North America / United States | |
| EPI_ISL_15063648 | A/snow goose/North Dakota/N22-04/2022 | A / H5N1 | North America / United States | |
| EPI_ISL_15063668 | A/snow goose/North Dakota/N22-05/2022 | A / H5N1 | North America / United States | |
| EPI_ISL_15063669 | A/ring-billed gull/Florida/W22-169/2022 | A / H5N1 | North America / United States | |
| EPI_ISL_15063839 | A/bald eagle/Virginia/W22-306/2022 | A / H5N1 | North America / United States | |
| EPI_ISL_15063841 | A/muscovy duck/Florida/W22-306/2022 | A / H5N1 | North America / United States | |
| EPI_ISL_15063842 | A/bald eagle/North Carolina/W22-186/2022 | A / H5N1 | North America / United States | |
| EPI_ISL_15063844 | A/gull/Florida/W22-162/2022 | A / H5N1 | North America / United States | |
| EPI_ISL_15063845 | A/snow goose/Kansas/W22-177B/2022 | A / H5N1 | North America / United States | |
| EPI_ISL_15063846 | A/bald eagle/Florida/W22-134-OP/2022 | A / H5N1 | North America / United States | |
| EPI_ISL_15063847 | A/bald eagle/North Carolina/W22-229/2022 | A / H5N1 | North America / United States | |
| EPI_ISL_15069397 | A/bottlenose dolphin/Florida/UFTt2203/2022 | A / H5N1 | North America / United States / Florida / Dixie County | Thomas Paul Fabrizio (St. Jude Children's Research Hospital / Infectious Diseases) |
| EPI_ISL_15069398 | A/European Herring Gull/Netherlands/12/2022 | A / H5N1 | Europe / Netherlands / Neeltje Jans | Sanne Thewessen (Erasmus Medical Center / Viroscience) |
| EPI_ISL_15069399 | A/Black-headed gull/Netherlands/10/2022 | A / H5N1 | Europe / Netherlands / Neeltje Jans | Sanne Thewessen (Erasmus Medical Center / Viroscience) |
| EPI_ISL_15069400 | A/Mute Swan/Netherlands/1/2022 | A / H5N1 | Europe / Netherlands / Weversinlaag, Schouwen | Sanne Thewessen (Erasmus Medical Center / Viroscience) |
| EPI_ISL_15069401 | A/Common Tern/Netherlands/26/2022 | A / H5N1 | Europe / Netherlands / Weversinlaag | Sanne Thewessen (Erasmus Medical Center / Viroscience) |
| EPI_ISL_15072537 | A/chicken/Hokkaido/A-1/2022 | A / H5 | Asia / Japan / Hokkaido / Abashiri | Norikazu Isoda (Graduate School of Veterinary Medicine, Hokkaido University / Laboratory of Microbiology) |
| EPI_ISL_15072538 | A/chicken/Hokkaido/I-1/2022 | A / H5N1 | Asia / Japan / Hokkaido / Shiraoi | Norikazu Isoda (Graduate School of Veterinary Medicine, Hokkaido University / Laboratory of Microbiology) |
| EPI_ISL_15072539 | A/emu/Hokkaido/A-2/2022 | A / H5N1 | Asia / Japan / Hokkaido / Abashiri | Norikazu Isoda (Graduate School of Veterinary Medicine, Hokkaido University / Laboratory of Microbiology) |
| EPI_ISL_15072540 | A/crow/Hokkaido/HU-1/2022 | A / H5N1 | Asia / Japan / Hokkaido / Sapporo | Norikazu Isoda (Graduate School of Veterinary Medicine, Hokkaido University / Laboratory of Microbiology) |
| EPI_ISL_15077371 | A/chicken/Idaho/22-011347-004-original/2022 | A / H5N1 | North America / United States / Idaho / Gooding County | Mary Lea Killian (National Veterinary Services Laboratories - USDA / Diagnostic Virology Laboratory) |
| EPI_ISL_15077373 | A/chicken/Pennsylvania/22-012092-006-original/2022 | A / H5N1 | North America / United States / Pennsylvania / Lancaster County | Mary Lea Killian (National Veterinary Services Laboratories - USDA / Diagnostic Virology Laboratory) |
| EPI_ISL_15077374 | A/black vulture/Florida/22-010358-001-original/2022 | A / H5N1 | North America / United States / Florida / Citrus County | Mary Lea Killian (National Veterinary Services Laboratories - USDA / Diagnostic Virology Laboratory) |
| EPI_ISL_15077375 | A/red-shouldered hawk/Minnesota/22-012000-004-original/2( | A / H5N1 | North America / United States / Minnesota | Mary Lea Killian (National Veterinary Services Laboratories - USDA / Diagnostic Virology Laboratory) |
| EPI_ISL_15077376 | A/black vulture/Florida/22-012331-001-original/2022 | A / H5N1 | North America / United States / Florida / Brevard County | Mary Lea Killian (National Veterinary Services Laboratories - USDA / Diagnostic Virology Laboratory) |
| EPI_ISL_15077377 | A/black vulture/Florida/22-012333-001-original/2022 | A / H5N1 | North America / United States / Florida / Brevard County | Mary Lea Killian (National Veterinary Services Laboratories - USDA / Diagnostic Virology Laboratory) |
| EPI_ISL_15078237 | A/Canada goose/Wyoming/22-011671-001-original/2022 | A / H5N1 | North America / United States / Wyoming / Fremont County | Mary Lea Killian (National Veterinary Services Laboratories - USDA / Diagnostic Virology Laboratory) |
| EPI_ISL_15078238 | A/turkey/Iowa/22-012098-001-original/2022 | A / H5N1 | North America / United States / Iowa / Bremer County | Mary Lea Killian (National Veterinary Services Laboratories - USDA / Diagnostic Virology Laboratory) |
| EPI_ISL_15078239 | A/black vulture/Maryland/22-012407-001-original/2022 | A / H5N1 | North America / United States / Maryland / Cecil County | Mary Lea Killian (National Veterinary Services Laboratories - USDA / Diagnostic Virology Laboratory) |
| EPI_ISL_15078240 | A/Cooper's hawk/Minnesota/22-012931-001-original/2022 | A / H5N1 | North America / United States / Minnesota / Ramsey County | Mary Lea Killian (National Veterinary Services Laboratories - USDA / Diagnostic Virology Laboratory) |
| EPI_ISL_15078241 | A/bald eagle/Wyoming/22-013015-001-original/2022 | A / H5N1 | North America / United States / Wyoming / Park County | Mary Lea Killian (National Veterinary Services Laboratories - USDA / Diagnostic Virology Laboratory) |
| EPI_ISL_15078244 | A/chicken/Michigan/22-013961-001-original/2022 | A / H5N1 | North America / United States / Michigan / Oakland County / Oakland Count | Mary Lea Killian (National Veterinary Services Laboratories - USDA / Diagnostic Virology Laboratory) |

| | | | | |
|---|---|---|---|---|
| EPI_ISL_15078245 | A/fox/Michigan/22-014536-004-original/2022 | A / H5N1 | North America / United States / Michigan / Lapeer County | Mary Lea Killian (National Veterinary Services Laboratories - USDA / Diagnostic Virology Laboratory) |
| EPI_ISL_15078246 | A/fox/Wisconsin/22-014746-008-original/2022 | A / H5N1 | North America / United States / Wisconsin / Adam County | Mary Lea Killian (National Veterinary Services Laboratories - USDA / Diagnostic Virology Laboratory) |
| EPI_ISL_15078247 | A/fox/Wisconsin/22-014746-030-original/2022 | A / H5N1 | North America / United States / Wisconsin / Grant County | Mary Lea Killian (National Veterinary Services Laboratories - USDA / Diagnostic Virology Laboratory) |
| EPI_ISL_15078248 | A/bobcat/Wisconsin/22-016051-001-original/2022 | A / H5N1 | North America / United States / Wisconsin | Mary Lea Killian (National Veterinary Services Laboratories - USDA / Diagnostic Virology Laboratory) |
| EPI_ISL_15078249 | A/fox/Minnesota/22-016487-001-original/2022 | A / H5N1 | North America / United States / Minnesota / Itasca County | Mary Lea Killian (National Veterinary Services Laboratories - USDA / Diagnostic Virology Laboratory) |
| EPI_ISL_15078250 | A/Virginia opossum/Iowa/22-016780-001-original/2022 | A / H5N1 | North America / United States / Iowa / Buchanan County | Mary Lea Killian (National Veterinary Services Laboratories - USDA / Diagnostic Virology Laboratory) |
| EPI_ISL_15078251 | A/red fox/North Dakota/22-017354-001-original/2022 | A / H5N1 | North America / United States / North Dakota / Burleigh County | Mary Lea Killian (National Veterinary Services Laboratories - USDA / Diagnostic Virology Laboratory) |
| EPI_ISL_15078252 | A/raccoon/Washington/22-018406-002-original/2022 | A / H5N1 | North America / United States / Washington / Franklin County | Mary Lea Killian (National Veterinary Services Laboratories - USDA / Diagnostic Virology Laboratory) |
| EPI_ISL_15078253 | A/red fox/Michigan/22-018712-001-original/2022 | A / H5N1 | North America / United States / Michigan / Muskegon County | Mary Lea Killian (National Veterinary Services Laboratories - USDA / Diagnostic Virology Laboratory) |
| EPI_ISL_15078254 | A/skunk/Washington/22-019274-001-original/2022 | A / H5N1 | North America / United States / Washington / Whitman County | Mary Lea Killian (National Veterinary Services Laboratories - USDA / Diagnostic Virology Laboratory) |
| EPI_ISL_15078255 | A/dolphin/Florida/22-025319-002-original/2022 | A / H5N1 | North America / United States / Florida / Dixie County | Mary Lea Killian (National Veterinary Services Laboratories - USDA / Diagnostic Virology Laboratory) |
| EPI_ISL_15078261 | A/chicken/Pennsylvania/22-012092-010-original/2022 | A / H5N1 | North America / United States / Pennsylvania / Lancaster County | Mary Lea Killian (National Veterinary Services Laboratories - USDA / Diagnostic Virology Laboratory) |
| EPI_ISL_15081424 | A/wigeon/Sakhalin/37M/2021 | A / H5N1 | Europe / Russian Federation | Ivan Sobolev (Research Institute of Experimental and Clinical Medicine) |
| EPI_ISL_15088295 | A/Common Buzzard/Netherlands/1/2022 | A / H5N1 | Europe / Netherlands / Hippolytushoef, Mulders Eendenkooi | Sanne Thewessen (Erasmus Medical Center / Viroscience) |
| EPI_ISL_15088296 | A/European Herring Gull/Netherlands/13/2022 | A / H5N1 | Europe / Netherlands / Den Oever, Waddenhaven | Sanne Thewessen (Erasmus Medical Center / Viroscience) |
| EPI_ISL_15088297 | A/European Herring Gull/Netherlands/14/2022 | A / H5N1 | Europe / Netherlands / Hippolytushoef, normerven | Sanne Thewessen (Erasmus Medical Center / Viroscience) |
| EPI_ISL_15088298 | A/Northern Gannet/Netherlands/1/2022 | A / H5N1 | Europe / Netherlands / Hippolytushoef, normerven | Sanne Thewessen (Erasmus Medical Center / Viroscience) |
| EPI_ISL_15088299 | A/Greylag Goose/Netherlands/10/2022 | A / H5N1 | Europe / Netherlands / Wieringerwerf, Noorderdijkweg | Sanne Thewessen (Erasmus Medical Center / Viroscience) |
| EPI_ISL_15088300 | A/Greylag Goose/Netherlands/11/2022 | A / H5N1 | Europe / Netherlands / Wieringerwerf, Noorderdijkweg | Sanne Thewessen (Erasmus Medical Center / Viroscience) |
| EPI_ISL_15088301 | A/Greylag Goose/Netherlands/12/2022 | A / H5N1 | Europe / Netherlands / Wieringerwerf, Noorderdijkweg | Sanne Thewessen (Erasmus Medical Center / Viroscience) |
| EPI_ISL_15088302 | A/Greylag Goose/Netherlands/13/2022 | A / H5N1 | Europe / Netherlands / Noord Buren, Waddenkust | Sanne Thewessen (Erasmus Medical Center / Viroscience) |
| EPI_ISL_15088303 | A/Lesser Black-backed Gull/Netherlands/3/2022 | A / H5N1 | Europe / Netherlands / IJmuiden, Forteiland | Sanne Thewessen (Erasmus Medical Center / Viroscience) |
| EPI_ISL_15088304 | A/European Herring Gull/Netherlands/15/2022 | A / H5N1 | Europe / Netherlands / IJmuiden, Forteiland | Sanne Thewessen (Erasmus Medical Center / Viroscience) |
| EPI_ISL_15088305 | A/Common Teal/Netherlands/1/2022 | A / H5N1 | Europe / Netherlands / Wieringerwerf, Noorderdijkweg | Sanne Thewessen (Erasmus Medical Center / Viroscience) |
| EPI_ISL_15088306 | A/Mallard/Netherlands/3/2022 | A / H5N1 | Europe / Netherlands / Hippolytushoef, Mulders Eendenkooi | Sanne Thewessen (Erasmus Medical Center / Viroscience) |
| EPI_ISL_15088307 | A/Mallard/Netherlands/4/2022 | A / H5N1 | Europe / Netherlands / Hippolytushoef, Mulders Eendenkooi | Sanne Thewessen (Erasmus Medical Center / Viroscience) |
| EPI_ISL_15088308 | A/Mallard/Netherlands/5/2022 | A / H5N1 | Europe / Netherlands / Hippolytushoef, Mulders Eendenkooi | Sanne Thewessen (Erasmus Medical Center / Viroscience) |
| EPI_ISL_15088309 | A/Mallard/Netherlands/6/2022 | A / H5N1 | Europe / Netherlands / Hippolytushoef, Mulders Eendenkooi | Sanne Thewessen (Erasmus Medical Center / Viroscience) |
| EPI_ISL_15115556 | A/turkey/England/112259/2022 | A / H5N1 | Europe / United Kingdom / East Riding of Yorkshire | Alex Byrne (Animal and Plant Health Agency (APHA) / Virology Department) |
| EPI_ISL_152085 | A/duck/Guangdong/wy11/2008 | A / H5N5 | Asia / China | |
| EPI_ISL_152086 | A/duck/Guangdong/wy19/2008 | A / H5N5 | Asia / China | |
| EPI_ISL_152087 | A/duck/Guangdong/wy24/2008 | A / H5N5 | Asia / China | |
| EPI_ISL_15214832 | A/bald eagle/Florida/W22-153B/2022 | A / H5N1 | North America / United States | |
| EPI_ISL_15214835 | A/common tern/Maine/W22-480A/2022 | A / H5N1 | North America / United States | |
| EPI_ISL_15214837 | A/snow goose/Kansas/W22-260/2022 | A / H5N1 | North America / United States | |
| EPI_ISL_15214838 | A/common eiders/Maine/W22-481B/2022 | A / H5N1 | North America / United States | |
| EPI_ISL_15214839 | A/bald eagle/Florida/W22-153A/2022 | A / H5N1 | North America / United States | |
| EPI_ISL_15214841 | A/common tern/Maine/W22-480B/2022 | A / H5N1 | North America / United States | |
| EPI_ISL_15214843 | A/Ross's goose/North Dakota/N22-08/2022 | A / H5N1 | North America / United States | |
| EPI_ISL_15214845 | A/lesser snow goose/North Dakota/ND-10/2022 | A / H5N1 | North America / United States | |
| EPI_ISL_15214846 | A/snow goose/Kansas/W22-174B/2022 | A / H5N1 | North America / United States | |
| EPI_ISL_15215193 | A/common eiders/Maine/W22-481A/2022 | A / H5N1 | North America / United States | |
| EPI_ISL_15215277 | A/bald eagle/North Carolina/W22-140/2022 | A / H5N1 | North America / United States | |
| EPI_ISL_15215278 | A/hooded merganser/Florida/W22-154/2022 | A / H5N1 | North America / United States | |
| EPI_ISL_15215280 | A/bald eagle/Kansas/W22-185/2022 | A / H5N1 | North America / United States | |
| EPI_ISL_15234355 | A/red-backed-hawk/Spain/2313-1_22VIR8632-10/2022 | A / H5N1 | Europe / Spain | Giacomo Barbierato (Istituto Zooprofilattico Sperimentale delle Venezie / Ricerca e innovazione) |
| EPI_ISL_15234356 | A/Anser_anser/Spain/2636-3_22VIR8632-11/2022 | A / H5N1 | Europe / Spain | Giacomo Barbierato (Istituto Zooprofilattico Sperimentale delle Venezie / Ricerca e innovazione) |
| EPI_ISL_15234357 | A/Anser_anser/Spain/2753-3_22VIR8632-12/2022 | A / H5N1 | Europe / Spain | Giacomo Barbierato (Istituto Zooprofilattico Sperimentale delle Venezie / Ricerca e innovazione) |
| EPI_ISL_15234358 | A/Anser_anser/Spain/2825-1_22VIR8632-13/2022 | A / H5N1 | Europe / Spain | Giacomo Barbierato (Istituto Zooprofilattico Sperimentale delle Venezie / Ricerca e innovazione) |
| EPI_ISL_15234359 | A/Anser_anser/Spain/810-6_22VIR8632-16/2022 | A / H5N1 | Europe / Spain | Giacomo Barbierato (Istituto Zooprofilattico Sperimentale delle Venezie / Ricerca e innovazione) |
| EPI_ISL_15234360 | A/turkey/Spain/2755-5_22VIR8632-1/2022 | A / H5N1 | Europe / Spain | Giacomo Barbierato (Istituto Zooprofilattico Sperimentale delle Venezie / Ricerca e innovazione) |
| EPI_ISL_15234361 | A/turkey/Spain/2996-43_22VIR8632-21/2022 | A / H5N1 | Europe / Spain | Giacomo Barbierato (Istituto Zooprofilattico Sperimentale delle Venezie / Ricerca e innovazione) |
| EPI_ISL_15234362 | A/turkey/Spain/2996-42_22VIR8632-22/2022 | A / H5N1 | Europe / Spain | Giacomo Barbierato (Istituto Zooprofilattico Sperimentale delle Venezie / Ricerca e innovazione) |
| EPI_ISL_15234363 | A/turkey/Spain/2755-6_22VIR8632-2/2022 | A / H5N1 | Europe / Spain | Giacomo Barbierato (Istituto Zooprofilattico Sperimentale delle Venezie / Ricerca e innovazione) |
| EPI_ISL_15234364 | A/chicken/Spain/2854-5_22VIR8632-3/2022 | A / H5N1 | Europe / Spain | Giacomo Barbierato (Istituto Zooprofilattico Sperimentale delle Venezie / Ricerca e innovazione) |
| EPI_ISL_15234365 | A/chicken/Spain/2854-7_22VIR8632-4/2022 | A / H5N1 | Europe / Spain | Giacomo Barbierato (Istituto Zooprofilattico Sperimentale delle Venezie / Ricerca e innovazione) |
| EPI_ISL_15234366 | A/duck/Spain/2095-2_22VIR8632-5/2022 | A / H5N1 | Europe / Spain | Giacomo Barbierato (Istituto Zooprofilattico Sperimentale delle Venezie / Ricerca e innovazione) |
| EPI_ISL_15234656 | A/bearded_vulture/Spain/2116-3_22VIR8632-8/2022 | A / H5N1 | Europe / Spain | Giacomo Barbierato (Istituto Zooprofilattico Sperimentale delle Venezie / Ricerca e innovazione) |
| EPI_ISL_15267012 | A/Lesser Black-backed Gull/Netherlands/4/2022 | A / H5N1 | Europe / Netherlands / Den Oever, Waddenhaven | Sanne Thewessen (Erasmus Medical Center / Viroscience) |
| EPI_ISL_15267013 | A/Eurasian Spoonbill/Netherlands/5/2022 | A / H5N1 | Europe / Netherlands / Texel, de Schorren | Sanne Thewessen (Erasmus Medical Center / Viroscience) |
| EPI_ISL_15267014 | A/Eurasian Spoonbill/Netherlands/6/2022 | A / H5N1 | Europe / Netherlands / Texel, de Schorren | Sanne Thewessen (Erasmus Medical Center / Viroscience) |
| EPI_ISL_15267016 | A/Eurasian Spoonbill/Netherlands/7/2022 | A / H5N1 | Europe / Netherlands / Texel, de Schorren | Sanne Thewessen (Erasmus Medical Center / Viroscience) |
| EPI_ISL_15267017 | A/Eurasian Spoonbill/Netherlands/8/2022 | A / H5N1 | Europe / Netherlands / Wieringerwerf, Noorderdijkweg | Sanne Thewessen (Erasmus Medical Center / Viroscience) |
| EPI_ISL_15267018 | A/Greylag Goose/Netherlands/14/2022 | A / H5N1 | Europe / Netherlands / Wieringerwerf, Noorderdijkweg | Sanne Thewessen (Erasmus Medical Center / Viroscience) |
| EPI_ISL_15267019 | A/Greylag Goose/Netherlands/15/2022 | A / H5N1 | Europe / Netherlands / Enkhuizen, Ijsselmeer, De Kreupel | Sanne Thewessen (Erasmus Medical Center / Viroscience) |
| EPI_ISL_15267020 | A/Greylag Goose/Netherlands/16/2022 | A / H5N1 | Europe / Netherlands / Enkhuizen, Ijsselmeer, De Kreupel | Sanne Thewessen (Erasmus Medical Center / Viroscience) |
| EPI_ISL_15267021 | A/Common Tern/Netherlands/27/2022 | A / H5N1 | Europe / Netherlands / Enkhuizen, Ijsselmeer, De Kreupel | Sanne Thewessen (Erasmus Medical Center / Viroscience) |
| EPI_ISL_15267022 | A/European Herring Gull/Netherlands/16/2022 | A / H5N1 | Europe / Netherlands / Hippolytushoef, Mulders Eendenkooi | Sanne Thewessen (Erasmus Medical Center / Viroscience) |
| EPI_ISL_15267023 | A/Northern Gannet/Netherlands/2/2022 | A / H5N1 | Europe / Netherlands / Vlieland, Noordzee strand | Sanne Thewessen (Erasmus Medical Center / Viroscience) |
| EPI_ISL_15267024 | A/Northern Gannet/Netherlands/3/2022 | A / H5N1 | Europe / Netherlands / Vlieland, Noordzee strand | Sanne Thewessen (Erasmus Medical Center / Viroscience) |
| EPI_ISL_15267025 | A/Northern Gannet/Netherlands/4/2022 | A / H5N1 | Europe / Netherlands / Vlieland, Noordzee strand | Sanne Thewessen (Erasmus Medical Center / Viroscience) |
| EPI_ISL_15267026 | A/Northern Gannet/Netherlands/5/2022 | A / H5N1 | Europe / Netherlands / Vlieland, Noordzee strand | Sanne Thewessen (Erasmus Medical Center / Viroscience) |

| | | | | |
|---|---|---|---|---|
| EPI_ISL_15267027 | A/Northern Gannet/Netherlands/6/2022 | A / H5N1 | Europe / Netherlands / Vlieland, Noordzee strand | Sanne Thewessen (Erasmus Medical Center / Viroscience) |
| EPI_ISL_15350905 | A/chicken/Ghana/AVL-763_21VIR7050-39/2021 | A / H5N1 | Africa / Ghana | Giacomo Barbierato (Istituto Zooprofilattico Sperimentale delle Venezie / Ricerca e innovazione) |
| EPI_ISL_153620 | A/chicken/China/AH/2012 | A / H5N1 | Asia / China | |
| EPI_ISL_15364788 | A/European Herring Gull/Netherlands/17/2022 | A / H5N1 | Europe / Netherlands / Vlieland, Noordzee strand | Sanne Thewessen (Erasmus Medical Center / Viroscience) |
| EPI_ISL_15364789 | A/European Herring Gull/Netherlands/18/2022 | A / H5N1 | Europe / Netherlands / Vlieland, eerste Kroonspolder | Sanne Thewessen (Erasmus Medical Center / Viroscience) |
| EPI_ISL_15364790 | A/Greylag Goose/Netherlands/17/2022 | A / H5N1 | Europe / Netherlands / Hippolytushoef, Normerven | Sanne Thewessen (Erasmus Medical Center / Viroscience) |
| EPI_ISL_15364792 | A/Mallard/Netherlands/11/2022 | A / H5N1 | Europe / Netherlands / Hippolytushoef, Mulders Eendenkooi | Sanne Thewessen (Erasmus Medical Center / Viroscience) |
| EPI_ISL_15364793 | A/Mallard/Netherlands/12/2022 | A / H5N1 | Europe / Netherlands / Lithse eendenkooi | Sanne Thewessen (Erasmus Medical Center / Viroscience) |
| EPI_ISL_15364794 | A/Mallard/Netherlands/13/2022 | A / H5N1 | Europe / Netherlands / Lithse eendenkooi | Sanne Thewessen (Erasmus Medical Center / Viroscience) |
| EPI_ISL_15364795 | A/Mallard/Netherlands/8/2022 | A / H5N1 | Europe / Netherlands / Hippolytushoef, Mulders Eendenkooi | Sanne Thewessen (Erasmus Medical Center / Viroscience) |
| EPI_ISL_15364796 | A/Mallard/Netherlands/9/2022 | A / H5N1 | Europe / Netherlands / Hippolytushoef, Mulders Eendenkooi | Sanne Thewessen (Erasmus Medical Center / Viroscience) |
| EPI_ISL_15364797 | A/Mute Swan/Netherlands/2/2022 | A / H5N1 | Europe / Netherlands / Kollum | Sanne Thewessen (Erasmus Medical Center / Viroscience) |
| EPI_ISL_15430045 | A/herring_gull/Poland/MB82B/2016 | A / H5N8 | Europe / Poland / West Pomeranian Voivodeship | Edyta ?wi?to? (National Veterinary Research Institut Poland, PIWet-PIB) |
| EPI_ISL_15430209 | A/common crane/Yunnan-Huize/11/2021(H5N8) | A / H5N8 | Asia / China / Yunnan Province / Huize Black-necked Crane National Nature Reserve, Yunnan Pro | Jinglin Wang (Yunnan Animal Science and Veterinary Institute / Yunnan Tropical and Subtropical Animal Viruses Disease Laborator) |
| EPI_ISL_15433349 | A/common crane /Yunnan-Huize/22/2021(H5N8) | A / H5N8 | Asia / China / Yunnan Province / Huize Black-necked Crane National Nature Reserve, Yunnan Pro | Jinglin Wang (Yunnan Animal Science and Veterinary Institute / Yunnan Tropical and Subtropical Animal Viruses Disease Laborator) |
| EPI_ISL_15435500 | A/common crane/Yunnan-Huize/24/2021(H5N8) | A / H5N8 | Asia / China / Yunnan Province / Huize Black-necked Crane National Nature Reserve, Yunnan Pro | Jinglin Wang (Yunnan Animal Science and Veterinary Institute / Yunnan Tropical and Subtropical Animal Viruses Disease Laborator) |
| EPI_ISL_15435511 | A/common crane/Yunnan-Huize/27/2021(H5N8) | A / H5N8 | Asia / China / Yunnan Province / Huize Black-necked Crane National Nature Reserve, Yunnan Pro | Jinglin Wang (Yunnan Animal Science and Veterinary Institute / Yunnan Tropical and Subtropical Animal Viruses Disease Laborator) |
| EPI_ISL_15535272 | A/domestic_goose/Poland/H397-N/2022 | A / H5N1 | Europe / Poland / Lodz Voivodeship | Edyta ?wi?to? (National Veterinary Research Institut Poland, PIWet-PIB) |
| EPI_ISL_15542438 | A/CastillaLaMancha/223739/2022 | A / H5N1 | Europe / Spain / Castille-La Mancha | Francisco Pozo (Instituto de Salud Carlos III) |
| EPI_ISL_15576616 | A/white-fronted goose/Miyagi/0410D001/2022 | A / H5N1 | Asia / Japan / Miyagi / Kurihara city | Norikazu Isoda (Graduate School of Veterinary Medicine, Hokkaido University / Laboratory of Microbiology) |
| EPI_ISL_15576617 | A/Eurasian wigeon/Hokkaido/Q71/2022 | A / H5N1 | Asia / Japan / Hokkaido / Noduke Peninsula, Bekkai town | Norikazu Isoda (Graduate School of Veterinary Medicine, Hokkaido University / Laboratory of Microbiology) |
| EPI_ISL_15579535 | A/European Herring Gull/Netherlands/19/2022 | A / H5N1 | Europe / Netherlands / Hippolytushoef, Normerven | Sanne Thewessen (Erasmus Medical Center / Viroscience) |
| EPI_ISL_15579536 | A/Greylag Goose/Netherlands/18/2022 | A / H5N1 | Europe / Netherlands / Stroe, Waddenkust | Sanne Thewessen (Erasmus Medical Center / Viroscience) |
| EPI_ISL_15579537 | A/Barnacle Goose/Netherlands/19/2022 | A / H5N1 | Europe / Netherlands / Enkhuizen, Ijsselmeer, De Kreupel | Sanne Thewessen (Erasmus Medical Center / Viroscience) |
| EPI_ISL_15579538 | A/Eurasian Curlew/Netherlands/4/2022 | A / H5N1 | Europe / Netherlands / Hippolytushoef, Normerven | Sanne Thewessen (Erasmus Medical Center / Viroscience) |
| EPI_ISL_15579539 | A/European Herring Gull/Netherlands/20/2022 | A / H5N1 | Europe / Netherlands / Noord Buren, Waddenkust | Sanne Thewessen (Erasmus Medical Center / Viroscience) |
| EPI_ISL_15579540 | A/Mallard/Netherlands/15/2022 | A / H5N1 | Europe / Netherlands / Eendenkooi Wieringen | Sanne Thewessen (Erasmus Medical Center / Viroscience) |
| EPI_ISL_15579543 | A/Mallard/Netherlands/18/2022 | A / H5N1 | Europe / Netherlands / Eendenkooi Wieringen | Sanne Thewessen (Erasmus Medical Center / Viroscience) |
| EPI_ISL_15579544 | A/Great Black-backed Gull/4/2022 | A / H5N1 | Europe / Netherlands / Kwade hoek, Stellendam | Sanne Thewessen (Erasmus Medical Center / Viroscience) |
| EPI_ISL_15579545 | A/Sanderling/Netherlands/2/2022 | A / H5N1 | Europe / Netherlands / Kwade hoek, Stellendam | Sanne Thewessen (Erasmus Medical Center / Viroscience) |
| EPI_ISL_15581789 | A/Mallard/Netherlands/19/2022 | A / H5N1 | Europe / Netherlands / Eendenkooi Wieringen | Sanne Thewessen (Erasmus Medical Center / Viroscience) |
| EPI_ISL_15581790 | A/Mallard/Netherlands/20/2022 | A / H5N1 | Europe / Netherlands / Oud-Albas | Sanne Thewessen (Erasmus Medical Center / Viroscience) |
| EPI_ISL_15581791 | A/Mallard/Netherlands/21/2022 | A / H5N1 | Europe / Netherlands / Oud-Albas | Sanne Thewessen (Erasmus Medical Center / Viroscience) |
| EPI_ISL_15581792 | A/Mallard/Netherlands/22/2022 | A / H5N1 | Europe / Netherlands / Oud-Albas | Sanne Thewessen (Erasmus Medical Center / Viroscience) |
| EPI_ISL_15585858 | A/chicken/England/119017/2022 | A / H5N1 | Europe / United Kingdom / England | Alex Byrne (Animal and Plant Health Agency (APHA) / Virology Department) |
| EPI_ISL_15585864 | A/chicken/England/123975/2022 | A / H5N1 | Europe / United Kingdom / England | Alex Byrne (Animal and Plant Health Agency (APHA) / Virology Department) |
| EPI_ISL_15585890 | A/chicken/Wales/116603/2022 | A / H5N1 | Europe / United Kingdom / Wales | Alex Byrne (Animal and Plant Health Agency (APHA) / Virology Department) |
| EPI_ISL_15586122 | A/domestic_duck/England/117887/2022 | A / H5N1 | Europe / United Kingdom / England | Alex Byrne (Animal and Plant Health Agency (APHA) / Virology Department) |
| EPI_ISL_15586133 | A/mute_swan/England/117298/2022 | A / H5N1 | Europe / United Kingdom / England | Alex Byrne (Animal and Plant Health Agency (APHA) / Virology Department) |
| EPI_ISL_15602898 | A/turkey/Montana/22-028879-001-original/2022 | A / H5N4 | North America / United States / Montana / Teton County | Mary Lea Killian (National Veterinary Services Laboratories - USDA / Diagnostic Virology Laboratory) |
| EPI_ISL_15613488 | A/mallard/Miyazaki/4501C607-c1/2021 | A / H5N8 | Asia / Japan / Miyazaki | |
| EPI_ISL_15613489 | A/mallard/Miyazaki/4501C607-c3/2021 | A / H5N8 | Asia / Japan / Miyazaki | |
| EPI_ISL_15613490 | A/mallard/Miyazaki/4501C607-c5/2021 | A / H5N8 | Asia / Japan / Miyazaki | |
| EPI_ISL_15613491 | A/mallard/Miyazaki/4501C607-c6/2021 | A / H5N8 | Asia / Japan / Miyazaki | |
| EPI_ISL_15613492 | A/mallard/Miyazaki/4501C607-c7/2021 | A / H5N8 | Asia / Japan / Miyazaki | |
| EPI_ISL_15613493 | A/mallard/Miyazaki/4501C607-c8/2021 | A / H5N8 | Asia / Japan / Miyazaki | |
| EPI_ISL_15613494 | A/teal/Miyazaki/211109-32/2021 | A / H5N1 | Asia / Japan / Miyazaki | |
| EPI_ISL_15613502 | A/whooper swan/Iwate/0302I010/2022 | A / H5 | Asia / Japan / Iwate | |
| EPI_ISL_15613505 | A/white-tailed eagle/Hokkaido/20220322001/2022 | A / H5 | Asia / Japan / Hokkaido | |
| EPI_ISL_15613506 | A/white-tailed eagle/Hokkaido/20220324001/2022 | A / H5 | Asia / Japan / Hokkaido | |
| EPI_ISL_15613515 | A/white-tailed eagle/Hokkaido/20220426001/2022 | A / H5 | Asia / Japan / Hokkaido | |
| EPI_ISL_15613516 | A/black kite/Hokkaido/0104P081/2022 | A / H5 | Asia / Japan / Hokkaido | |
| EPI_ISL_15613574 | A/Guangdong/1/2021 | A / H5N6 | Asia / China / Guangdong Province | |
| EPI_ISL_15614803 | A/ibis/Egypt/RLQP-229S/2022 | A / H5N1 | Africa / Egypt | |
| EPI_ISL_15614805 | A/Lesser scaup/MD/-LC-EESC-024/2022 | A / H5N1 | North America / United States | |
| EPI_ISL_15614807 | A/red-tailed hawk/Kansas/W22-198/2022 | A / H5N1 | North America / United States | |
| EPI_ISL_15647834 | A/duck/Korea/H493/2022 | A / H5N1 | Asia / Korea, Republic of | Se-hee An (Animal and Plant Quarantine Agency (APQA) / Avian Influenza Research and Diagsnotic Division) |
| EPI_ISL_15647835 | A/common teal/Korea/WA537/2022 | A / H5N1 | Asia / Korea, Republic of | Se-hee An (Animal and Plant Quarantine Agency (APQA) / Avian Influenza Research and Diagsnotic Division) |
| EPI_ISL_15647836 | A/Mandarin duck/Korea/WA496 | A / H5N1 | Asia / Korea, Republic of | Se-hee An (Animal and Plant Quarantine Agency (APQA) / Avian Influenza Research and Diagsnotic Division) |
| EPI_ISL_15647837 | A/duck/Korea/H125/2022 | A / H5N1 | Asia / Korea, Republic of | Se-hee An (Animal and Plant Quarantine Agency (APQA) / Avian Influenza Research and Diagsnotic Division) |
| EPI_ISL_15676637 | A/chicken/Croatia/55/2022 | A / H5N1 | Europe / Croatia / Osjecko-Baranjska Zupanija / Beli Manstir | Vladimir Savi? (Croatian Veterinary Institute / Poultry Centre) |
| EPI_ISL_15676638 | A/mute swan/Croatia/108/2022 | A / H5N1 | Europe / Croatia / Zagrebacka Zupanija / Strmec Samoborski, Kipiš?e lake | Vladimir Savi? (Croatian Veterinary Institute / Poultry Centre) |
| EPI_ISL_156815 | A/mallard duck/Shanghai/SH-9/2013 | A / H5N8 | Asia / China / Shanghai Municipality | Fan Sheng Tao (Institute of Laboratory Animal Sciences, Chinese Academy) |
| EPI_ISL_15697406 | A/Pavo/Belgium/10906_0006/2022 | A / H5N1 | Europe / Belgium / Provincie Limburg / Bocholt | Steven Van Borm (Sciensano, Department of Animal Infectious Diseases / Animal Infectious Diseases) |
| EPI_ISL_15697407 | A/Gallus_gallus/Belgium/10968_0005/2022 | A / H5N1 | Europe / Belgium / Provincie West-Vlaanderen / Sint Laureins | Steven Van Borm (Sciensano, Department of Animal Infectious Diseases / Animal Infectious Diseases) |
| EPI_ISL_15698209 | A/Gallus_gallus/Belgium/11248_0001/2022 | A / H5N1 | Europe / Belgium / Provincie West-Vlaanderen / Sint Laureins | Steven Van Borm (Sciensano, Department of Animal Infectious Diseases / Animal Infectious Diseases) |
| EPI_ISL_15698370 | A/Ardea_cinerea/Belgium/11304_0003/2022 | A / H5N1 | Europe / Belgium / Provincie Oost-Vlaanderen / Zelzate | Steven Van Borm (Sciensano, Department of Animal Infectious Diseases / Animal Infectious Diseases) |
| EPI_ISL_15699041 | A/Cygnus_olor/Belgium/11304_0009/2022 | A / H5N1 | Europe / Belgium / Provincie West-Vlaanderen / Brugge | Steven Van Borm (Sciensano, Department of Animal Infectious Diseases / Animal Infectious Diseases) |
| EPI_ISL_15699790 | A/Gallus_gallus/Belgium/11387_0002/2022 | A / H5N1 | Europe / Belgium / Provincie Oost-Vlaanderen / Sint Laureins | Steven Van Borm (Sciensano, Department of Animal Infectious Diseases / Animal Infectious Diseases) |
| EPI_ISL_15699799 | A/Gallus_gallus/Belgium/11445_0002/2022 | A / H5N1 | Europe / Belgium / Provincie West-Vlaanderen / Wingene | Steven Van Borm (Sciensano, Department of Animal Infectious Diseases / Animal Infectious Diseases) |
| EPI_ISL_15699929 | A/Gallus_gallus/Belgium/11943_0003/2022 | A / H5N1 | Europe / Belgium / Provincie Limburg / Tongeren | Steven Van Borm (Sciensano, Department of Animal Infectious Diseases / Animal Infectious Diseases) |
| EPI_ISL_15732741 | A/large-billed crow/Hokkaido/B003/2022 | A / H5N2 | Asia / Japan / Hokkaido / Sapporo | Norikazu Isoda (Graduate School of Veterinary Medicine, Hokkaido University / Laboratory of Microbiology) |

| | | | | |
|---|---|---|---|---|
| EPI_ISL_15732766 | A/Eurasian wigeon/Hokkaido/M184/2022 | A / H5N1 | Asia / Japan / Hokkaido / the Komuke Lake | Norikazu Isoda (Graduate School of Veterinary Medicine, Hokkaido University / Laboratory of Microbiology) |
| EPI_ISL_1575129 | A/Common Buzzard/Netherlands/4/2020 | A / H5N8 | Europe / Netherlands / Rotterdam | Theo Bestebroer (Erasmus Medical Center / Department of Virology) |
| EPI_ISL_1575130 | A/Mute Swan/Netherlands/1/2020 | A / H5N8 | Europe / Netherlands / Rotterdam | Theo Bestebroer (Erasmus Medical Center / Department of Virology) |
| EPI_ISL_1575131 | A/Herring Gull/Netherlands/1/2020 | A / H5N8 | Europe / Netherlands / Rotterdam | Theo Bestebroer (Erasmus Medical Center / Department of Virology) |
| EPI_ISL_1575132 | A/Herring Gull/Netherlands/2/2020 | A / H5N8 | Europe / Netherlands / Rotterdam | Theo Bestebroer (Erasmus Medical Center / Department of Virology) |
| EPI_ISL_1575133 | A/Herring Gull/Netherlands/3/2020 | A / H5N8 | Europe / Netherlands / Rotterdam | Theo Bestebroer (Erasmus Medical Center / Department of Virology) |
| EPI_ISL_1575134 | A/Mute Swan/Netherlands/3/2020 | A / H5N8 | Europe / Netherlands / Rotterdam | Theo Bestebroer (Erasmus Medical Center / Department of Virology) |
| EPI_ISL_1575135 | A/Greylag Goose/Netherlands/1/2020 | A / H5N8 | Europe / Netherlands / Rotterdam | Theo Bestebroer (Erasmus Medical Center / Department of Virology) |
| EPI_ISL_1575136 | A/Mute Swan/Netherlands/5/2020 | A / H5N8 | Europe / Netherlands / Rotterdam | Theo Bestebroer (Erasmus Medical Center / Department of Virology) |
| EPI_ISL_15759949 | A/Common Teal/Netherlands/3/2022 | A / H5N1 | Europe / Netherlands / Eendenkooi, de Sompen | Sanne Thewessen (Erasmus Medical Center / Viroscience) |
| EPI_ISL_15759950 | A/Common Buzzard/2/2022 | A / H5N1 | Europe / Netherlands / Woerden | Sanne Thewessen (Erasmus Medical Center / Viroscience) |
| EPI_ISL_15759951 | A/Mallard/Netherlands/24/2022 | A / H5N1 | Europe / Netherlands / Oud-Albas | Sanne Thewessen (Erasmus Medical Center / Viroscience) |
| EPI_ISL_15759952 | A/Mallard/Netherlands/25/2022 | A / H5N1 | Europe / Netherlands / Oud-Albas | Sanne Thewessen (Erasmus Medical Center / Viroscience) |
| EPI_ISL_15759953 | A/Mallard/Netherlands/26/2022 | A / H5N1 | Europe / Netherlands / Oud-Alblas | Sanne Thewessen (Erasmus Medical Center / Viroscience) |
| EPI_ISL_157609 | A/breeder duck/Korea/Gochang1/2014 | A / H5N8 | Asia / Korea, Republic of | |
| EPI_ISL_157610 | A/broiler duck/Korea/Buan2/2014 | A / H5N8 | Asia / Korea, Republic of | |
| EPI_ISL_157611 | A/baikal teal/Korea/Donglim3/2014 | A / H5N8 | Asia / Korea, Republic of | |
| EPI_ISL_15812579 | A/Anser_anser_domesticus/Belgium/10918_0002/2022 | A / H5N1 | Europe / Belgium / Provincie West-Vlaanderen / Beernem | Steven Van Borm (Sciensano, Department of Animal Infectious Diseases / Animal Infectious Diseases) |
| EPI_ISL_15812678 | A/Anas_platyrhynchos_domesticus/Belgium/11599_0005/202 | A / H5N1 | Europe / Belgium / Provincie West-Vlaanderen / Diksmuide | Steven Van Borm (Sciensano, Department of Animal Infectious Diseases / Animal Infectious Diseases) |
| EPI_ISL_15813174 | A/Gallus_gallus/Belgium/12311_0001/2022 | A / H5N1 | Europe / Belgium / Province de Liege / Aywaille | Steven Van Borm (Sciensano, Department of Animal Infectious Diseases / Animal Infectious Diseases) |
| EPI_ISL_15814043 | A/Gallus_gallus/Belgium/12395_0002/2022 | A / H5N1 | Europe / Belgium / Provincie Antwerpen / Hoogstraten | Steven Van Borm (Sciensano, Department of Animal Infectious Diseases / Animal Infectious Diseases) |
| EPI_ISL_15843373 | A/Bald Eagle/BC/OTH-33-36/2022 (H5N1) | A / H5N1 | North America / Canada / British Columbia | Yohannes Berhane (Canadian Food Inspection Agency / Avian Unit) |
| EPI_ISL_15844573 | A/Broiler Chicken/BC/FAV-0228/2022 (H5N1) | A / H5N1 | North America / Canada / British Columbia | Yohannes Berhane (Canadian Food Inspection Agency / Avian Unit) |
| EPI_ISL_15857836 | A/teal/Italy/22VIR10396-33/2022 | A / H5N1 | Europe / Italy | Giacomo Barbierato (Istituto Zooprofilattico Sperimentale delle Venezie / Ricerca e innovazione) |
| EPI_ISL_15857837 | A/teal/Italy/22VIR10396-5/2022 | A / H5N1 | Europe / Italy | Giacomo Barbierato (Istituto Zooprofilattico Sperimentale delle Venezie / Ricerca e innovazione) |
| EPI_ISL_15857838 | A/teal/Italy/22VIR10576-13/2022 | A / H5N1 | Europe / Italy | Giacomo Barbierato (Istituto Zooprofilattico Sperimentale delle Venezie / Ricerca e innovazione) |
| EPI_ISL_15857839 | A/chicken/Italy/22VIR11246-6/2022 | A / H5N1 | Europe / Italy | Giacomo Barbierato (Istituto Zooprofilattico Sperimentale delle Venezie / Ricerca e innovazione) |
| EPI_ISL_15857840 | A/turkey/Italy/22VIR11262-1/2022 | A / H5N1 | Europe / Italy | Giacomo Barbierato (Istituto Zooprofilattico Sperimentale delle Venezie / Ricerca e innovazione) |
| EPI_ISL_15857841 | A/swan/Italy/22VIR10875-2/2022 | A / H5N1 | Europe / Italy | Giacomo Barbierato (Istituto Zooprofilattico Sperimentale delle Venezie / Ricerca e innovazione) |
| EPI_ISL_15857842 | A/swan/Italy/22VIR10995-7/2022 | A / H5N1 | Europe / Italy | Giacomo Barbierato (Istituto Zooprofilattico Sperimentale delle Venezie / Ricerca e innovazione) |
| EPI_ISL_15857843 | A/rural_laying_hen/Italy/22VIR10996-6/2022 | A / H5N1 | Europe / Italy | Giacomo Barbierato (Istituto Zooprofilattico Sperimentale delle Venezie / Ricerca e innovazione) |
| EPI_ISL_15857844 | A/turkey/Italy/22VIR11056-4/2022 | A / H5N1 | Europe / Italy | Giacomo Barbierato (Istituto Zooprofilattico Sperimentale delle Venezie / Ricerca e innovazione) |
| EPI_ISL_15857845 | A/turkey/Italy/22VIR11144-1/2022 | A / H5N1 | Europe / Italy | Giacomo Barbierato (Istituto Zooprofilattico Sperimentale delle Venezie / Ricerca e innovazione) |
| EPI_ISL_15857846 | A/turkey/Italy/22VIR10627-4/2022 | A / H5N1 | Europe / Italy | Giacomo Barbierato (Istituto Zooprofilattico Sperimentale delle Venezie / Ricerca e innovazione) |
| EPI_ISL_15857847 | A/rural_chicken/Italy/22VIR10651-4/2022 | A / H5N1 | Europe / Italy | Giacomo Barbierato (Istituto Zooprofilattico Sperimentale delle Venezie / Ricerca e innovazione) |
| EPI_ISL_15857848 | A/turkey/Italy/22VIR10733-4/2022 | A / H5N1 | Europe / Italy | Giacomo Barbierato (Istituto Zooprofilattico Sperimentale delle Venezie / Ricerca e innovazione) |
| EPI_ISL_15857849 | A/swan/Italy/22VIR10737-10/2022 | A / H5N1 | Europe / Italy | Giacomo Barbierato (Istituto Zooprofilattico Sperimentale delle Venezie / Ricerca e innovazione) |
| EPI_ISL_15857864 | A/rural_goose/Italy/22VIR10789-2/2022 | A / H5N1 | Europe / Italy | Giacomo Barbierato (Istituto Zooprofilattico Sperimentale delle Venezie / Ricerca e innovazione) |
| EPI_ISL_15857953 | A/laying_hen/Italy/22VIR10800-2/2022 | A / H5N1 | Europe / Italy | Giacomo Barbierato (Istituto Zooprofilattico Sperimentale delle Venezie / Ricerca e innovazione) |
| EPI_ISL_15857954 | A/laying_hen/Italy/22VIR10267-7/2022 | A / H5N1 | Europe / Italy | Giacomo Barbierato (Istituto Zooprofilattico Sperimentale delle Venezie / Ricerca e innovazione) |
| EPI_ISL_15857955 | A/rural_duck/Italy/22VIR10534-3/2022 | A / H5N1 | Europe / Italy | Giacomo Barbierato (Istituto Zooprofilattico Sperimentale delle Venezie / Ricerca e innovazione) |
| EPI_ISL_15857956 | A/teal/Italy/22VIR9917-19/2022 | A / H5N1 | Europe / Italy | Giacomo Barbierato (Istituto Zooprofilattico Sperimentale delle Venezie / Ricerca e innovazione) |
| EPI_ISL_15857957 | A/teal/Italy/22VIR9917-52/2022 | A / H5N1 | Europe / Italy | Giacomo Barbierato (Istituto Zooprofilattico Sperimentale delle Venezie / Ricerca e innovazione) |
| EPI_ISL_15857958 | A/swan/Italy/22VIR10097-3/2022 | A / H5N1 | Europe / Italy | Giacomo Barbierato (Istituto Zooprofilattico Sperimentale delle Venezie / Ricerca e innovazione) |
| EPI_ISL_15857959 | A/swan/Italy/22VIR10097-4/2022 | A / H5N1 | Europe / Italy | Giacomo Barbierato (Istituto Zooprofilattico Sperimentale delle Venezie / Ricerca e innovazione) |
| EPI_ISL_15857961 | A/Goose/Italy/22VIR9673-3/2022 | A / H5N1 | Europe / Italy | Giacomo Barbierato (Istituto Zooprofilattico Sperimentale delle Venezie / Ricerca e innovazione) |
| EPI_ISL_15857990 | A/Chicken/Italy/22VIR9832-3/2022 | A / H5N1 | Europe / Italy | Giacomo Barbierato (Istituto Zooprofilattico Sperimentale delle Venezie / Ricerca e innovazione) |
| EPI_ISL_15857991 | A/Hawaiian_goose/Italy/22VIR9126-7/2022 | A / H5N1 | Europe / Italy | Giacomo Barbierato (Istituto Zooprofilattico Sperimentale delle Venezie / Ricerca e innovazione) |
| EPI_ISL_15857992 | A/chicken/France/22P023203/2022 | A / H5N1 | Europe / France / Brittany / Departement des Cotes-d'Armor | Francois-Xavier Briand (ANSES Agence Nationale De Securite Sanitaire De L'alimentation / Laboratoire de Ploufragan-Plouzané) |
| EPI_ISL_15878539 | A/Mink/Spain/3691-8_22VIR10586-10/2022 | A / H5N1 | Europe / Spain | Giacomo Barbierato (Istituto Zooprofilattico Sperimentale delle Venezie / Ricerca e innovazione) |
| EPI_ISL_15878540 | A/Mink/Spain/3691-10_22VIR10586-11/2022 | A / H5N1 | Europe / Spain | Giacomo Barbierato (Istituto Zooprofilattico Sperimentale delle Venezie / Ricerca e innovazione) |
| EPI_ISL_15878541 | A/Mink/Spain/3691-2_22VIR10586-8/2022 | A / H5N1 | Europe / Spain | Giacomo Barbierato (Istituto Zooprofilattico Sperimentale delle Venezie / Ricerca e innovazione) |
| EPI_ISL_15878542 | A/Mink/Spain/3691-3_22VIR10586-9/2022 | A / H5N1 | Europe / Spain | Giacomo Barbierato (Istituto Zooprofilattico Sperimentale delle Venezie / Ricerca e innovazione) |
| EPI_ISL_15878543 | A/laying_hen/Spain/3135-17_22VIR10586-1/2022 | A / H5N1 | Europe / Spain | Giacomo Barbierato (Istituto Zooprofilattico Sperimentale delle Venezie / Ricerca e innovazione) |
| EPI_ISL_15878544 | A/laying_hen/Spain/3135-41_22VIR10586-2/2022 | A / H5N1 | Europe / Spain | Giacomo Barbierato (Istituto Zooprofilattico Sperimentale delle Venezie / Ricerca e innovazione) |
| EPI_ISL_15878545 | A/laying_hen/Spain/3232-35_22VIR10586-3/2022 | A / H5N1 | Europe / Spain | Giacomo Barbierato (Istituto Zooprofilattico Sperimentale delle Venezie / Ricerca e innovazione) |
| EPI_ISL_15878546 | A/laying_hen/Spain/3232-37_22VIR10586-4/2022 | A / H5N1 | Europe / Spain | Giacomo Barbierato (Istituto Zooprofilattico Sperimentale delle Venezie / Ricerca e innovazione) |
| EPI_ISL_15878548 | A/Northern_gannet/Spain/3372-12_22VIR10586-6/2022 | A / H5N1 | Europe / Spain | Giacomo Barbierato (Istituto Zooprofilattico Sperimentale delle Venezie / Ricerca e innovazione) |
| EPI_ISL_15878549 | A/White_stork/Spain/3438-3_22VIR10586-7/2022 | A / H5N1 | Europe / Spain | Giacomo Barbierato (Istituto Zooprofilattico Sperimentale delle Venezie / Ricerca e innovazione) |
| EPI_ISL_15895378 | A/peafowl/ Croatia/123/2022 | A / H5N1 | Europe / Croatia / Zagrebacka Zupanija / Gornji Stupnik | Vladimir Savi? (Croatian Veterinary Institute / Poultry Centre) |
| EPI_ISL_15895379 | A/mute swan/Croatia/125/2022 | A / H5N1 | Europe / Croatia / Vukovarsko-Srijemska Zupanija / Vukovar, Vuka river | Vladimir Savi? (Croatian Veterinary Institute / Poultry Centre) |
| EPI_ISL_15895380 | A/mute swan/Croatia/126/2022 | A / H5N1 | Europe / Croatia / Vukovarsko-Srijemska Zupanija / Vukovar, Vuka river | Vladimir Savi? (Croatian Veterinary Institute / Poultry Centre) |
| EPI_ISL_15923322 | A/peregrine falcon/Kanagawa/1409C001T1/2022 | A / H5N1 | Asia / Japan / Kanagawa | Takehiko Saito (National Institute of Animal Health) |
| EPI_ISL_15925875 | A/Eurasian Wigeon/Netherlands/1/2022 | A / H5N1 | Europe / Netherlands / Eendenkooi, Wieringen | Oanh Vuong-Zoet (Erasmus Medical Center / Department of Virology) |
| EPI_ISL_15925876 | A/Eurasian Wigeon/Netherlands/2/2022 | A / H5N1 | Europe / Netherlands / Eendenkooi, Wieringen | Oanh Vuong-Zoet (Erasmus Medical Center / Department of Virology) |
| EPI_ISL_15925878 | A/Common Teal/Netherlands/5/2022 | A / H5N1 | Europe / Netherlands / Eendenkooi, Wieringen | Oanh Vuong-Zoet (Erasmus Medical Center / Department of Virology) |
| EPI_ISL_15925879 | A/Common Teal/Netherlands/6/2022 | A / H5N1 | Europe / Netherlands / Eendenkooi, Wieringen | Oanh Vuong-Zoet (Erasmus Medical Center / Department of Virology) |
| EPI_ISL_15925880 | A/Gadwall/Netherlands/1/2022 | A / H5N1 | Europe / Netherlands / Eendenkooi, Wieringen | Oanh Vuong-Zoet (Erasmus Medical Center / Department of Virology) |
| EPI_ISL_15925881 | A/Gadwall/Netherlands/2/2022 | A / H5N1 | Europe / Netherlands / Eendenkooi, Wieringen | Oanh Vuong-Zoet (Erasmus Medical Center / Department of Virology) |
| EPI_ISL_15925882 | A/Eurasian Wigeon/Netherlands/3/2022 | A / H5N1 | Europe / Netherlands / Krommenier, Woudpolder | Oanh Vuong-Zoet (Erasmus Medical Center / Department of Virology) |
| EPI_ISL_15925883 | A/Eurasian Wigeon/Netherlands/4/2022 | A / H5N1 | Europe / Netherlands / Oud-Alblas | Oanh Vuong-Zoet (Erasmus Medical Center / Department of Virology) |
| EPI_ISL_15925884 | A/Great Cormorant/Netherlands/1/2022 | A / H5N1 | Europe / Netherlands / Enkhuizen, Ijsselmeer, De Kreupel | Oanh Vuong-Zoet (Erasmus Medical Center / Department of Virology) |

| | | | | |
|---|---|---|---|---|
| EPI_ISL_15925885 | A/Black-headed Gull/Netherlands/11/2022 | A / H5N1 | Europe / Netherlands / Enkhuizen, Ijsselmeer, De Kreupel | Oanh Vuong-Zoet (Erasmus Medical Center / Department of Virology) |
| EPI_ISL_15942297 | A/domestic_duck/England/152473/2022 | A / H5N1 | Europe / United Kingdom | Daniel Maskell (Animal and Plant Health Agency (APHA) / International Reference Laboratory) |
| EPI_ISL_15943002 | A/Spot-billed_duck/Korea/K22-730-1/2022 | A / H5N1 | Asia / Korea, Republic of / Chungcheongnam-do | Dong-Hun Lee (Konkuk University / College of Veterinary Medicine) |
| EPI_ISL_15943015 | A/Wild_bird/Korea/K22-742/2022 | A / H5N1 | Asia / Korea, Republic of / Chungcheongnam-do | Dong-Hun Lee (Konkuk University / College of Veterinary Medicine) |
| EPI_ISL_15944663 | A/Spot-billed_duck/Korea/K22-856-2/2022 | A / H5N1 | Asia / Korea, Republic of | Dong-Hun Lee (Konkuk University / College of Veterinary Medicine) |
| EPI_ISL_15944665 | A/Spot-billed_duck/Korea/K22-862-1/2022 | A / H5N1 | Asia / Korea, Republic of | Dong-Hun Lee (Konkuk University / College of Veterinary Medicine) |
| EPI_ISL_15944667 | A/Spot-billed_duck/Korea/K22-920/2022 | A / H5N1 | Asia / Korea, Republic of | Dong-Hun Lee (Konkuk University / College of Veterinary Medicine) |
| EPI_ISL_159719 | A/Chicken/Kumamoto/1-7/2014 | A / H5N8 | Asia / Japan / Kumamoto | Michiyo Yoshizawa Harada (National Agriculture and Food Research Organization / National Institute of Animal Health) |
| EPI_ISL_16003253 | A/Red Fox/MB/FAV-370-01/2022 | A / H5N1 | North America / Canada / Manitoba | Yohannes Berhane (Canadian Food Inspection Agency / Avian Unit) |
| EPI_ISL_16003265 | A/Red Fox/MB/FAV-414-004/2022 | A / H5N1 | North America / Canada / Manitoba | Yohannes Berhane (Canadian Food Inspection Agency / Avian Unit) |
| EPI_ISL_16003266 | A/Red Fox/MB/FAV-414-08/2022 | A / H5N1 | North America / Canada / Manitoba | Yohannes Berhane (Canadian Food Inspection Agency / Avian Unit) |
| EPI_ISL_16003267 | A/Red Fox/MB/FAV-0414-12/2022 | A / H5N1 | North America / Canada / Manitoba | Yohannes Berhane (Canadian Food Inspection Agency / Avian Unit) |
| EPI_ISL_16003268 | A/Skunk/MB/FAV-470/2022 | A / H5N1 | North America / Canada / Manitoba | Yohannes Berhane (Canadian Food Inspection Agency / Avian Unit) |
| EPI_ISL_16013752 | A/Pelican/Venezuela/Pel3/2022 | A / H5N1 | South America / Venezuela, Bolivarian Republic of / Estado Anzoategui | Flor Helene Pujol (Instituto Venezolano de investigaciones Científicas / Microbiology) |
| EPI_ISL_16013753 | A/Pelican/Venezuela/Pel4/2022 | A / H5N1 | South America / Venezuela, Bolivarian Republic of / Estado Anzoategui | Flor Helene Pujol (Instituto Venezolano de investigaciones Científicas / Microbiology) |
| EPI_ISL_16056568 | A/Eurasian Wigeon/Netherlands/5/2022 | A / H5N1 | Europe / Netherlands / Krommeniedijk-Woudpolder, Noord-Holland | Oanh Vuong-Zoet (Erasmus Medical Center / Department of Virology) |
| EPI_ISL_16056569 | A/Gadwall/Netherlands/3/2022 | A / H5N1 | Europe / Netherlands / Eendenkooi Wieringen | Oanh Vuong-Zoet (Erasmus Medical Center / Department of Virology) |
| EPI_ISL_16056570 | A/Mallard Duck/Netherlands/27/2022 | A / H5N1 | Europe / Netherlands / Oud Alblas | Oanh Vuong-Zoet (Erasmus Medical Center / Department of Virology) |
| EPI_ISL_16056571 | A/Mallard Duck/Netherlands/28/2022 | A / H5N1 | Europe / Netherlands / Oud Alblas | Oanh Vuong-Zoet (Erasmus Medical Center / Department of Virology) |
| EPI_ISL_16056572 | A/Mallard Duck/Netherlands/29/2022 | A / H5N1 | Europe / Netherlands / Oud Alblas | Oanh Vuong-Zoet (Erasmus Medical Center / Department of Virology) |
| EPI_ISL_16101521 | A/chicken/Idaho/22-011336-001-original/2022 | A / H5N1 | North America / United States / Idaho / Caribou County | Mary Lea Killian (National Veterinary Services Laboratories - USDA / Diagnostic Virology Laboratory) |
| EPI_ISL_16101522 | A/turkey/Minnesota/22-011337-001-original/2022 | A / H5N1 | North America / United States / Minnesota / Meeker County | Mary Lea Killian (National Veterinary Services Laboratories - USDA / Diagnostic Virology Laboratory) |
| EPI_ISL_16101523 | A/turkey/Minnesota/22-011337-002-original/2022 | A / H5N1 | North America / United States / Minnesota / Meeker County | Mary Lea Killian (National Veterinary Services Laboratories - USDA / Diagnostic Virology Laboratory) |
| EPI_ISL_16101524 | A/turkey/Minnesota/22-011338-001-original/2022 | A / H5N1 | North America / United States / Minnesota / Blue Earth County | Mary Lea Killian (National Veterinary Services Laboratories - USDA / Diagnostic Virology Laboratory) |
| EPI_ISL_16101525 | A/turkey/Minnesota/22-011338-002-original/2022 | A / H5N1 | North America / United States / Minnesota / Blue Earth County | Mary Lea Killian (National Veterinary Services Laboratories - USDA / Diagnostic Virology Laboratory) |
| EPI_ISL_16101526 | A/domestic duck/Indiana/22-011343-001-original/2022 | A / H5N1 | North America / United States / Indiana / Elkhart County | Mary Lea Killian (National Veterinary Services Laboratories - USDA / Diagnostic Virology Laboratory) |
| EPI_ISL_16101527 | A/chicken/Idaho/22-011347-002-original/2022 | A / H5N1 | North America / United States / Idaho / Gooding County | Mary Lea Killian (National Veterinary Services Laboratories - USDA / Diagnostic Virology Laboratory) |
| EPI_ISL_16101528 | A/chicken/Utah/22-011468-001-original/2022 | A / H5N1 | North America / United States / Utah / Utah County | Mary Lea Killian (National Veterinary Services Laboratories - USDA / Diagnostic Virology Laboratory) |
| EPI_ISL_16101529 | A/chicken/Utah/22-011468-002-original/2022 | A / H5N1 | North America / United States / Utah / Utah County | Mary Lea Killian (National Veterinary Services Laboratories - USDA / Diagnostic Virology Laboratory) |
| EPI_ISL_16102154 | A/turkey/Wisconsin/22-011508-001-original/2022 | A / H5N1 | North America / United States / Wisconsin / Polk County | Mary Lea Killian (National Veterinary Services Laboratories - USDA / Diagnostic Virology Laboratory) |
| EPI_ISL_16102155 | A/turkey/Wisconsin/22-011508-002-original/2022 | A / H5N1 | North America / United States / Wisconsin / Polk County | Mary Lea Killian (National Veterinary Services Laboratories - USDA / Diagnostic Virology Laboratory) |
| EPI_ISL_16102156 | A/domestic duck/Michigan/22-011521-001-original/2022 | A / H5N1 | North America / United States / Michigan / Macomb County | Mary Lea Killian (National Veterinary Services Laboratories - USDA / Diagnostic Virology Laboratory) |
| EPI_ISL_16102157 | A/chicken/Michigan/22-011521-003-original/2022 | A / H5N1 | North America / United States / Michigan / Macomb County | Mary Lea Killian (National Veterinary Services Laboratories - USDA / Diagnostic Virology Laboratory) |
| EPI_ISL_16102158 | A/chicken/Michigan/22-011521-004-original/2022 | A / H5N1 | North America / United States / Michigan / Macomb County | Mary Lea Killian (National Veterinary Services Laboratories - USDA / Diagnostic Virology Laboratory) |
| EPI_ISL_16102159 | A/quail/Michigan/22-011521-005-original/2022 | A / H5N1 | North America / United States / Michigan / Macomb County | Mary Lea Killian (National Veterinary Services Laboratories - USDA / Diagnostic Virology Laboratory) |
| EPI_ISL_16102160 | A/chicken/Michigan/22-011526-001-original/2022 | A / H5N1 | North America / United States / Michigan / Livingston County | Mary Lea Killian (National Veterinary Services Laboratories - USDA / Diagnostic Virology Laboratory) |
| EPI_ISL_16102161 | A/domestic duck/Michigan/22-011526-002-original/2022 | A / H5N1 | North America / United States / Michigan / Livingston County | Mary Lea Killian (National Veterinary Services Laboratories - USDA / Diagnostic Virology Laboratory) |
| EPI_ISL_16102162 | A/chicken/Michigan/22-011526-003-original/2022 | A / H5N1 | North America / United States / Michigan / Livingston County | Mary Lea Killian (National Veterinary Services Laboratories - USDA / Diagnostic Virology Laboratory) |
| EPI_ISL_16102163 | A/chicken/Pennsylvania/22-011531-001-original/2022 | A / H5N1 | North America / United States / Pennsylvania / Lancaster County | Mary Lea Killian (National Veterinary Services Laboratories - USDA / Diagnostic Virology Laboratory) |
| EPI_ISL_16102164 | A/chicken/Pennsylvania/22-011531-002-original/2022 | A / H5N1 | North America / United States / Pennsylvania / Lancaster County | Mary Lea Killian (National Veterinary Services Laboratories - USDA / Diagnostic Virology Laboratory) |
| EPI_ISL_16102165 | A/chicken/Michigan/22-011627-001-original/2022 | A / H5N1 | North America / United States / Michigan / Menominee County | Mary Lea Killian (National Veterinary Services Laboratories - USDA / Diagnostic Virology Laboratory) |
| EPI_ISL_16102166 | A/chicken/Pennsylvania/22-011767-001-original/2022 | A / H5N1 | North America / United States / Pennsylvania / Lancaster County | Mary Lea Killian (National Veterinary Services Laboratories - USDA / Diagnostic Virology Laboratory) |
| EPI_ISL_16102167 | A/guinea fowl/Idaho/22-011775-001-original/2022 | A / H5N1 | North America / United States / Idaho / Gooding County | Mary Lea Killian (National Veterinary Services Laboratories - USDA / Diagnostic Virology Laboratory) |
| EPI_ISL_16102168 | A/chicken/Idaho/22-011775-002-original/2022 | A / H5N1 | North America / United States / Idaho / Gooding County | Mary Lea Killian (National Veterinary Services Laboratories - USDA / Diagnostic Virology Laboratory) |
| EPI_ISL_16102169 | A/chicken/North Dakota/22-011778-001-original/2022 | A / H5N1 | North America / United States / North Dakota / Stutsman County | Mary Lea Killian (National Veterinary Services Laboratories - USDA / Diagnostic Virology Laboratory) |
| EPI_ISL_16102170 | A/turkey/Minnesota/22-011787-001-original/2022 | A / H5N1 | North America / United States / Minnesota / Otter Tail County | Mary Lea Killian (National Veterinary Services Laboratories - USDA / Diagnostic Virology Laboratory) |
| EPI_ISL_16102171 | A/turkey/Minnesota/22-011787-002-original/2022 | A / H5N1 | North America / United States / Minnesota / Otter Tail County | Mary Lea Killian (National Veterinary Services Laboratories - USDA / Diagnostic Virology Laboratory) |
| EPI_ISL_16102172 | A/turkey/Minnesota/22-011789-001-original/2022 | A / H5N1 | North America / United States / Minnesota / Stearns County | Mary Lea Killian (National Veterinary Services Laboratories - USDA / Diagnostic Virology Laboratory) |
| EPI_ISL_16102173 | A/turkey/Minnesota/22-011789-002-original/2022 | A / H5N1 | North America / United States / Minnesota / Stearns County | Mary Lea Killian (National Veterinary Services Laboratories - USDA / Diagnostic Virology Laboratory) |
| EPI_ISL_16102174 | A/turkey/Minnesota/22-011790-002-original/2022 | A / H5N1 | North America / United States / Minnesota / Morrison County | Mary Lea Killian (National Veterinary Services Laboratories - USDA / Diagnostic Virology Laboratory) |
| EPI_ISL_16102227 | A/turkey/Minnesota/22-011791-003-original/2022 | A / H5N1 | North America / United States / Minnesota / Kandiyohi County | Mary Lea Killian (National Veterinary Services Laboratories - USDA / Diagnostic Virology Laboratory) |
| EPI_ISL_16102228 | A/turkey/Minnesota/22-011791-004-original/2022 | A / H5N1 | North America / United States / Minnesota / Kandiyohi County | Mary Lea Killian (National Veterinary Services Laboratories - USDA / Diagnostic Virology Laboratory) |
| EPI_ISL_16102229 | A/turkey/Minnesota/22-011792-001-original/2022 | A / H5N1 | North America / United States / Minnesota / Morrison County | Mary Lea Killian (National Veterinary Services Laboratories - USDA / Diagnostic Virology Laboratory) |
| EPI_ISL_16102230 | A/turkey/Minnesota/22-011792-002-original/2022 | A / H5N1 | North America / United States / Minnesota / Morrison County | Mary Lea Killian (National Veterinary Services Laboratories - USDA / Diagnostic Virology Laboratory) |
| EPI_ISL_16102261 | A/turkey/Minnesota/22-011793-001-original/2022 | A / H5N1 | North America / United States / Minnesota / Meeker County | Mary Lea Killian (National Veterinary Services Laboratories - USDA / Diagnostic Virology Laboratory) |
| EPI_ISL_16102262 | A/turkey/Minnesota/22-011793-002-original/2022 | A / H5N1 | North America / United States / Minnesota / Meeker County | Mary Lea Killian (National Veterinary Services Laboratories - USDA / Diagnostic Virology Laboratory) |
| EPI_ISL_16102263 | A/turkey/Minnesota/22-011794-001-original/2022 | A / H5N1 | North America / United States / Minnesota / Swift County | Mary Lea Killian (National Veterinary Services Laboratories - USDA / Diagnostic Virology Laboratory) |
| EPI_ISL_16102264 | A/turkey/Minnesota/22-011794-002-original/2022 | A / H5N1 | North America / United States / Minnesota / Swift County | Mary Lea Killian (National Veterinary Services Laboratories - USDA / Diagnostic Virology Laboratory) |
| EPI_ISL_16102265 | A/turkey/Minnesota/22-011795-001-original/2022 | A / H5N1 | North America / United States / Minnesota / Kandiyohi County | Mary Lea Killian (National Veterinary Services Laboratories - USDA / Diagnostic Virology Laboratory) |
| EPI_ISL_16102266 | A/turkey/Minnesota/22-011796-001-original/2022 | A / H5N1 | North America / United States / Minnesota / Morrison County | Mary Lea Killian (National Veterinary Services Laboratories - USDA / Diagnostic Virology Laboratory) |
| EPI_ISL_16102267 | A/turkey/Minnesota/22-011796-002-original/2022 | A / H5N1 | North America / United States / Minnesota / Morrison County | Mary Lea Killian (National Veterinary Services Laboratories - USDA / Diagnostic Virology Laboratory) |
| EPI_ISL_16102275 | A/chicken/Colorado/22-011826-001-original/2022 | A / H5N1 | North America / United States / Colorado / Montrose County | Mary Lea Killian (National Veterinary Services Laboratories - USDA / Diagnostic Virology Laboratory) |
| EPI_ISL_16102276 | A/chicken/Colorado/22-011826-002-original/2022 | A / H5N1 | North America / United States / Colorado / Montrose County | Mary Lea Killian (National Veterinary Services Laboratories - USDA / Diagnostic Virology Laboratory) |
| EPI_ISL_16102277 | A/Indian Runner duck/Colorado/22-011827-001-original/2022 | A / H5N1 | North America / United States / Colorado / La Plata County | Mary Lea Killian (National Veterinary Services Laboratories - USDA / Diagnostic Virology Laboratory) |
| EPI_ISL_16102278 | A/chicken/Colorado/22-011827-002-original/2022 | A / H5N1 | North America / United States / Colorado / La Plata County | Mary Lea Killian (National Veterinary Services Laboratories - USDA / Diagnostic Virology Laboratory) |
| EPI_ISL_16102279 | A/chicken/Pennsylvania/22-011920-001-original/2022 | A / H5N1 | North America / United States / Pennsylvania / Lancaster County | Mary Lea Killian (National Veterinary Services Laboratories - USDA / Diagnostic Virology Laboratory) |
| EPI_ISL_16102280 | A/chicken/Pennsylvania/22-011921-002-original/2022 | A / H5N1 | North America / United States / Pennsylvania / Lancaster County | Mary Lea Killian (National Veterinary Services Laboratories - USDA / Diagnostic Virology Laboratory) |
| EPI_ISL_16102281 | A/duck/Indiana/22-011946-002-original/2022 | A / H5N1 | North America / United States / Indiana / Elkhart County | Mary Lea Killian (National Veterinary Services Laboratories - USDA / Diagnostic Virology Laboratory) |
| EPI_ISL_16102282 | A/duck/Indiana/22-011946-004-original/2022 | A / H5N1 | North America / United States / Indiana / Elkhart County | Mary Lea Killian (National Veterinary Services Laboratories - USDA / Diagnostic Virology Laboratory) |
| EPI_ISL_16102283 | A/turkey/Minnesota/22-012034-001-original/2022 | A / H5N1 | North America / United States / Minnesota / Todd County | Mary Lea Killian (National Veterinary Services Laboratories - USDA / Diagnostic Virology Laboratory) |
| EPI_ISL_16102926 | A/turkey/Minnesota/22-012036-001-original/2022 | A / H5N1 | North America / United States / Minnesota / Stearns County | Mary Lea Killian (National Veterinary Services Laboratories - USDA / Diagnostic Virology Laboratory) |
| EPI_ISL_16102927 | A/turkey/Minnesota/22-012036-002-original/2022 | A / H5N1 | North America / United States / Minnesota / Stearns County | Mary Lea Killian (National Veterinary Services Laboratories - USDA / Diagnostic Virology Laboratory) |

| | | | | |
|---|---|---|---|---|
| EPI_ISL_16102928 | A/chicken/Montana/22-012040-001-original/2022 | A / H5N1 | North America / United States / Montana / Glacier County | Mary Lea Killian (National Veterinary Services Laboratories - USDA / Diagnostic Virology Laboratory) |
| EPI_ISL_16102929 | A/mallard/Montana/22-012040-005-original/2022 | A / H5N1 | North America / United States / Montana / Glacier County | Mary Lea Killian (National Veterinary Services Laboratories - USDA / Diagnostic Virology Laboratory) |
| EPI_ISL_16102930 | A/chicken/Idaho/22-012045-001-original/2022 | A / H5N1 | North America / United States / Idaho / Madison County | Mary Lea Killian (National Veterinary Services Laboratories - USDA / Diagnostic Virology Laboratory) |
| EPI_ISL_16102931 | A/chicken/Idaho/22-012045-002-original/2022 | A / H5N1 | North America / United States / Idaho / Madison County | Mary Lea Killian (National Veterinary Services Laboratories - USDA / Diagnostic Virology Laboratory) |
| EPI_ISL_16102932 | A/duck/Idaho/22-012046-001-original/2022 | A / H5N1 | North America / United States / Idaho / Madison County | Mary Lea Killian (National Veterinary Services Laboratories - USDA / Diagnostic Virology Laboratory) |
| EPI_ISL_16102933 | A/goose/Idaho/22-012047-001-original/2022 | A / H5N1 | North America / United States / Idaho / Madison County | Mary Lea Killian (National Veterinary Services Laboratories - USDA / Diagnostic Virology Laboratory) |
| EPI_ISL_16112044 | A/Arctic_tern/Denmark/05803-1.01/2022 | A / H5N1 | Europe / Denmark / Region Nordjylland / Thisted Kommune | Charlotte Kristiane Hjulsager (Statens Serum Institute / Microbiological Diagnostic and Virology) |
| EPI_ISL_16113089 | A/barnacle_goose/Denmark/03332-1.02/2022 | A / H5N1 | Europe / Denmark / Region Midtjylland / Ringkobing-Skjern | Charlotte Kristiane Hjulsager (Statens Serum Institute / Microbiological Diagnostic and Virology) |
| EPI_ISL_16113090 | A/peregrine_falcon/Denmark/06880-1.02/2022 | A / H5N1 | Europe / Denmark / Region Sjaland / Vordingborg Kommune | Charlotte Kristiane Hjulsager (Statens Serum Institute / Microbiological Diagnostic and Virology) |
| EPI_ISL_16113225 | A/sandwich_tern/Denmark/05712-1.02/2022 | A / H5N1 | Europe / Denmark / Region Sjaland / Slagelse Kommune | Charlotte Kristiane Hjulsager (Statens Serum Institute / Microbiological Diagnostic and Virology) |
| EPI_ISL_16113229 | A/sandwich_tern/Denmark/05802-1.01_2022 | A / H5N1 | Europe / Denmark / Region Nordjylland / Thisted Kommune | Charlotte Kristiane Hjulsager (Statens Serum Institute / Microbiological Diagnostic and Virology) |
| EPI_ISL_16113230 | A/sandwich_tern/Denmark/05806-1.01/2022 | A / H5N1 | Europe / Denmark / Region Nordjylland / Thisted Kommune | Charlotte Kristiane Hjulsager (Statens Serum Institute / Microbiological Diagnostic and Virology) |
| EPI_ISL_16113231 | A/tawny_owl/Denmark/05372-1.01/2022 | A / H5N1 | Europe / Denmark / Region Syddanmark / Odense Kommune | Charlotte Kristiane Hjulsager (Statens Serum Institute / Microbiological Diagnostic and Virology) |
| EPI_ISL_16113242 | A/tawny_owl/Denmark/05373-1.01/2022 | A / H5N1 | Europe / Denmark / Region Syddanmark / Odense Kommune | Charlotte Kristiane Hjulsager (Statens Serum Institute / Microbiological Diagnostic and Virology) |
| EPI_ISL_16113366 | A/white-tailed_eagle/Denmark/05716-1.04/2022 | A / H5N1 | Europe / Denmark / Region Midtjylland / Horsens Kommune | Charlotte Kristiane Hjulsager (Statens Serum Institute / Microbiological Diagnostic and Virology) |
| EPI_ISL_16115054 | A/common_buzzard/Denmark/08287-1.01/2022 | A / H5N1 | Europe / Denmark / Region Syddanmark / Sonderborg Kommune | Charlotte Kristiane Hjulsager (Statens Serum Institute / Microbiological Diagnostic and Virology) |
| EPI_ISL_16157545 | A/chicken/Ecuador/NIC-INSPI-AGROCALIDAD-FBC-2/2022 | A / H5N1 | South America / Ecuador / Provincia de Cotopaxi | Alfredo Bruno Caicedo (Instituto Nacional de Investigacion en Salud Publica (INSPI) / Centro Nacional de Influenza (NIC)) |
| EPI_ISL_16161673 | A/chicken/Ecuador/NIC-INSPI-AGROCALIDAD-FBT-3/2022 | A / H5N1 | South America / Ecuador / Provincia de Cotopaxi | Alfredo Bruno Caicedo (Instituto Nacional de Investigacion en Salud Publica (INSPI) / Centro Nacional de Influenza (NIC)) |
| EPI_ISL_16161675 | A/chicken/Ecuador/NIC-INSPI-AGROCALIDAD-FBC-4/2022 | A / H5N1 | South America / Ecuador / Provincia de Cotopaxi | Alfredo Bruno Caicedo (Instituto Nacional de Investigacion en Salud Publica (INSPI) / Centro Nacional de Influenza (NIC)) |
| EPI_ISL_16171247 | A/chicken/Pennsylvania/22-012091-001-original/2022 | A / H5N1 | North America / United States / Pennsylvania / Lancaster County | Mary Lea Killian (National Veterinary Services Laboratories - USDA / Diagnostic Virology Laboratory) |
| EPI_ISL_16171248 | A/chicken/Pennsylvania/22-012092-003-original/2022 | A / H5N1 | North America / United States / Pennsylvania / Lancaster County | Mary Lea Killian (National Veterinary Services Laboratories - USDA / Diagnostic Virology Laboratory) |
| EPI_ISL_16171249 | A/chicken/Pennsylvania/22-012092-005-original/2022 | A / H5N1 | North America / United States / Pennsylvania / Lancaster County | Mary Lea Killian (National Veterinary Services Laboratories - USDA / Diagnostic Virology Laboratory) |
| EPI_ISL_16171250 | A/turkey/North Dakota/22-012099-001-original/2022 | A / H5N1 | North America / United States / North Dakota / Richland County | Mary Lea Killian (National Veterinary Services Laboratories - USDA / Diagnostic Virology Laboratory) |
| EPI_ISL_16171251 | A/turkey/North Dakota/22-012099-002-original/2022 | A / H5N1 | North America / United States / North Dakota / Richland County | Mary Lea Killian (National Veterinary Services Laboratories - USDA / Diagnostic Virology Laboratory) |
| EPI_ISL_16171252 | A/chicken/North Dakota/22-012100-001-original/2022 | A / H5N1 | North America / United States / North Dakota / Renville County | Mary Lea Killian (National Veterinary Services Laboratories - USDA / Diagnostic Virology Laboratory) |
| EPI_ISL_16171254 | A/turkey/Minnesota/22-012123-001-original/2022 | A / H5N1 | North America / United States / Minnesota / Todd County | Mary Lea Killian (National Veterinary Services Laboratories - USDA / Diagnostic Virology Laboratory) |
| EPI_ISL_16171255 | A/turkey/Minnesota/22-012125-001-original/2022 | A / H5N1 | North America / United States / Minnesota / Morrison County | Mary Lea Killian (National Veterinary Services Laboratories - USDA / Diagnostic Virology Laboratory) |
| EPI_ISL_16171256 | A/turkey/Minnesota/22-012125-002-original/2022 | A / H5N1 | North America / United States / Minnesota / Morrison County | Mary Lea Killian (National Veterinary Services Laboratories - USDA / Diagnostic Virology Laboratory) |
| EPI_ISL_16171322 | A/turkey/Minnesota/22-012126-002-original/2022 | A / H5N1 | North America / United States / Minnesota / Todd County | Mary Lea Killian (National Veterinary Services Laboratories - USDA / Diagnostic Virology Laboratory) |
| EPI_ISL_16171323 | A/turkey/Minnesota/22-012126-003-original/2022 | A / H5N1 | North America / United States / Minnesota / Todd County | Mary Lea Killian (National Veterinary Services Laboratories - USDA / Diagnostic Virology Laboratory) |
| EPI_ISL_16171324 | A/chicken/Pennsylvania/22-012248-001-original/2022 | A / H5N1 | North America / United States / Pennsylvania / Lancaster County | Mary Lea Killian (National Veterinary Services Laboratories - USDA / Diagnostic Virology Laboratory) |
| EPI_ISL_16171325 | A/chicken/Pennsylvania/22-012248-003-original/2022 | A / H5N1 | North America / United States / Pennsylvania / Lancaster County | Mary Lea Killian (National Veterinary Services Laboratories - USDA / Diagnostic Virology Laboratory) |
| EPI_ISL_16171326 | A/turkey/Minnesota/22-012285-001-original/2022 | A / H5N1 | North America / United States / Minnesota / Swift County | Mary Lea Killian (National Veterinary Services Laboratories - USDA / Diagnostic Virology Laboratory) |
| EPI_ISL_16171327 | A/turkey/Minnesota/22-012285-002-original/2022 | A / H5N1 | North America / United States / Minnesota / Swift County | Mary Lea Killian (National Veterinary Services Laboratories - USDA / Diagnostic Virology Laboratory) |
| EPI_ISL_16171328 | A/turkey/Minnesota/22-012286-001-original/2022 | A / H5N1 | North America / United States / Minnesota / Otter Tail County | Mary Lea Killian (National Veterinary Services Laboratories - USDA / Diagnostic Virology Laboratory) |
| EPI_ISL_16171329 | A/turkey/Minnesota/22-012286-002-original/2022 | A / H5N1 | North America / United States / Minnesota / Otter Tail County | Mary Lea Killian (National Veterinary Services Laboratories - USDA / Diagnostic Virology Laboratory) |
| EPI_ISL_16171330 | A/turkey/Minnesota/22-012287-001-original/2022 | A / H5N1 | North America / United States / Minnesota / Yellow Medicine County | Mary Lea Killian (National Veterinary Services Laboratories - USDA / Diagnostic Virology Laboratory) |
| EPI_ISL_16171331 | A/chicken/Colorado/22-012308-001-original/2022 | A / H5N1 | North America / United States / Colorado / Montrose County | Mary Lea Killian (National Veterinary Services Laboratories - USDA / Diagnostic Virology Laboratory) |
| EPI_ISL_16171332 | A/chicken/Colorado/22-012308-002-original/2022 | A / H5N1 | North America / United States / Colorado / Montrose County | Mary Lea Killian (National Veterinary Services Laboratories - USDA / Diagnostic Virology Laboratory) |
| EPI_ISL_16171399 | A/chicken/Minnesota/22-012373-001-original/2022 | A / H5N1 | North America / United States / Minnesota / Rice County | Mary Lea Killian (National Veterinary Services Laboratories - USDA / Diagnostic Virology Laboratory) |
| EPI_ISL_16171400 | A/goose/Minnesota/22-012373-002-original/2022 | A / H5N1 | North America / United States / Minnesota / Rice County | Mary Lea Killian (National Veterinary Services Laboratories - USDA / Diagnostic Virology Laboratory) |
| EPI_ISL_16171401 | A/turkey/Minnesota/22-012374-001-original/2022 | A / H5N1 | North America / United States / Minnesota / Stearns County | Mary Lea Killian (National Veterinary Services Laboratories - USDA / Diagnostic Virology Laboratory) |
| EPI_ISL_16171402 | A/chicken/Utah/22-012375-001-original/2022 | A / H5N1 | North America / United States / Utah / Cache County | Mary Lea Killian (National Veterinary Services Laboratories - USDA / Diagnostic Virology Laboratory) |
| EPI_ISL_16171403 | A/chicken/Wisconsin/22-012376-001-original/2022 | A / H5N1 | North America / United States / Wisconsin / Polk County | Mary Lea Killian (National Veterinary Services Laboratories - USDA / Diagnostic Virology Laboratory) |
| EPI_ISL_16171404 | A/duck/Wisconsin/22-012376-002-original/2022 | A / H5N1 | North America / United States / Wisconsin / Polk County | Mary Lea Killian (National Veterinary Services Laboratories - USDA / Diagnostic Virology Laboratory) |
| EPI_ISL_16171405 | A/chicken/Wisconsin/22-012376-003-original/2022 | A / H5N1 | North America / United States / Wisconsin / Polk County | Mary Lea Killian (National Veterinary Services Laboratories - USDA / Diagnostic Virology Laboratory) |
| EPI_ISL_16171406 | A/domestic duck/Michigan/22-012454-001-original/2022 | A / H5N1 | North America / United States / Michigan / Menominee County | Mary Lea Killian (National Veterinary Services Laboratories - USDA / Diagnostic Virology Laboratory) |
| EPI_ISL_16171407 | A/domestic goose/Michigan/22-012454-002-original/2022 | A / H5N1 | North America / United States / Michigan / Menominee County | Mary Lea Killian (National Veterinary Services Laboratories - USDA / Diagnostic Virology Laboratory) |
| EPI_ISL_16171408 | A/chicken/Iowa/22-012510-002-original/2022 | A / H5N1 | North America / United States / Iowa / Kossuth County | Mary Lea Killian (National Veterinary Services Laboratories - USDA / Diagnostic Virology Laboratory) |
| EPI_ISL_16171409 | A/chicken/North Dakota/22-012538-001-original/2022 | A / H5N1 | North America / United States / North Dakota / Richland County | Mary Lea Killian (National Veterinary Services Laboratories - USDA / Diagnostic Virology Laboratory) |
| EPI_ISL_16171411 | A/chicken/North Dakota/22-012538-002-original/2022 | A / H5N1 | North America / United States / North Dakota / Richland County | Mary Lea Killian (National Veterinary Services Laboratories - USDA / Diagnostic Virology Laboratory) |
| EPI_ISL_16171412 | A/chicken/Montana/22-012540-001-original/2022 | A / H5N1 | North America / United States / Montana / Missoula County | Mary Lea Killian (National Veterinary Services Laboratories - USDA / Diagnostic Virology Laboratory) |
| EPI_ISL_16171413 | A/chicken/Montana/22-012540-002-original/2022 | A / H5N1 | North America / United States / Montana / Missoula County | Mary Lea Killian (National Veterinary Services Laboratories - USDA / Diagnostic Virology Laboratory) |
| EPI_ISL_16171415 | A/chicken/Pennsylvania/22-012578-004-original/2022 | A / H5N1 | North America / United States / Pennsylvania / Lancaster County | Mary Lea Killian (National Veterinary Services Laboratories - USDA / Diagnostic Virology Laboratory) |
| EPI_ISL_16171416 | A/chicken/Kansas/22-012741-003-original/2022 | A / H5N1 | North America / United States / Kansas / Republic County | Mary Lea Killian (National Veterinary Services Laboratories - USDA / Diagnostic Virology Laboratory) |
| EPI_ISL_16171417 | A/chicken/Pennsylvania/22-012742-001-original/2022 | A / H5N1 | North America / United States / Pennsylvania / Lancaster County | Mary Lea Killian (National Veterinary Services Laboratories - USDA / Diagnostic Virology Laboratory) |
| EPI_ISL_16171418 | A/chicken/Pennsylvania/22-012742-002-original/2022 | A / H5N1 | North America / United States / Pennsylvania / Lancaster County | Mary Lea Killian (National Veterinary Services Laboratories - USDA / Diagnostic Virology Laboratory) |
| EPI_ISL_16171419 | A/chicken/Nebraska/22-012756-001-original/2022 | A / H5N1 | North America / United States / Nebraska / Knox County | Mary Lea Killian (National Veterinary Services Laboratories - USDA / Diagnostic Virology Laboratory) |
| EPI_ISL_16171449 | A/chicken/Michigan/22-012855-001-original/2022 | A / H5N1 | North America / United States / Michigan / Saginaw County | Mary Lea Killian (National Veterinary Services Laboratories - USDA / Diagnostic Virology Laboratory) |
| EPI_ISL_16171450 | A/guinea fowl/Michigan/22-012855-002-original/2022 | A / H5N1 | North America / United States / Michigan / Saginaw County | Mary Lea Killian (National Veterinary Services Laboratories - USDA / Diagnostic Virology Laboratory) |
| EPI_ISL_16171451 | A/turkey/Michigan/22-012855-003-original/2022 | A / H5N1 | North America / United States / Michigan / Saginaw County | Mary Lea Killian (National Veterinary Services Laboratories - USDA / Diagnostic Virology Laboratory) |
| EPI_ISL_16171452 | A/domestic goose/Michigan/22-012855-004-original/2022 | A / H5N1 | North America / United States / Michigan / Saginaw County | Mary Lea Killian (National Veterinary Services Laboratories - USDA / Diagnostic Virology Laboratory) |
| EPI_ISL_16171453 | A/domestic duck/Michigan/22-012855-005-original/2022 | A / H5N1 | North America / United States / Michigan / Saginaw County | Mary Lea Killian (National Veterinary Services Laboratories - USDA / Diagnostic Virology Laboratory) |
| EPI_ISL_16171454 | A/chicken/Minnesota/22-012932-001-original/2022 | A / H5N1 | North America / United States / Minnesota / Carver County | Mary Lea Killian (National Veterinary Services Laboratories - USDA / Diagnostic Virology Laboratory) |
| EPI_ISL_16171455 | A/chicken/Minnesota/22-012932-002-original/2022 | A / H5N1 | North America / United States / Minnesota / Carver County | Mary Lea Killian (National Veterinary Services Laboratories - USDA / Diagnostic Virology Laboratory) |
| EPI_ISL_16171456 | A/chicken/Indiana/22-012933-001-original/2022 | A / H5N1 | North America / United States / Indiana / Johnson County | Mary Lea Killian (National Veterinary Services Laboratories - USDA / Diagnostic Virology Laboratory) |
| EPI_ISL_16171457 | A/chicken/Minnesota/22-012966-001-original/2022 | A / H5N1 | North America / United States / Minnesota / Swift County | Mary Lea Killian (National Veterinary Services Laboratories - USDA / Diagnostic Virology Laboratory) |
| EPI_ISL_16171458 | A/turkey/Wisconsin/22-012973-001-original/2022 | A / H5N1 | North America / United States / Wisconsin / Barron County | Mary Lea Killian (National Veterinary Services Laboratories - USDA / Diagnostic Virology Laboratory) |
| EPI_ISL_16171459 | A/turkey/Wisconsin/22-012974-001-original/2022 | A / H5N1 | North America / United States / Wisconsin / Barron County | Mary Lea Killian (National Veterinary Services Laboratories - USDA / Diagnostic Virology Laboratory) |
| EPI_ISL_16171460 | A/turkey/Wisconsin/22-012974-002-original/2022 | A / H5N1 | North America / United States / Wisconsin / Barron County | Mary Lea Killian (National Veterinary Services Laboratories - USDA / Diagnostic Virology Laboratory) |
| EPI_ISL_16171461 | A/chicken/Michigan/22-012977-001-original/2022 | A / H5N1 | North America / United States / Michigan / Wexford County | Mary Lea Killian (National Veterinary Services Laboratories - USDA / Diagnostic Virology Laboratory) |
| EPI_ISL_16171462 | A/chicken/Montana/22-013053-001-original/2022 | A / H5N1 | North America / United States / Montana / Gallatin County | Mary Lea Killian (National Veterinary Services Laboratories - USDA / Diagnostic Virology Laboratory) |

| | | | | |
|---|---|---|---|---|
| EPI_ISL_16171463 | A/chicken/Montana/22-013053-002-original/2022 | A / H5N1 | North America / United States / Montana / Gallatin County | Mary Lea Killian (National Veterinary Services Laboratories - USDA / Diagnostic Virology Laboratory) |
| EPI_ISL_16171464 | A/duck/Montana/22-013055-005-original/2022 | A / H5N1 | North America / United States / Montana / Pondera County | Mary Lea Killian (National Veterinary Services Laboratories - USDA / Diagnostic Virology Laboratory) |
| EPI_ISL_16171465 | A/duck/Montana/22-013055-006-original/2022 | A / H5N1 | North America / United States / Montana / Pondera County | Mary Lea Killian (National Veterinary Services Laboratories - USDA / Diagnostic Virology Laboratory) |
| EPI_ISL_16171466 | A/chicken/Montana/22-013056-002-original/2022 | A / H5N1 | North America / United States / Montana / Fergus County | Mary Lea Killian (National Veterinary Services Laboratories - USDA / Diagnostic Virology Laboratory) |
| EPI_ISL_16171467 | A/chicken/Colorado/22-013057-001-original/2022 | A / H5N1 | North America / United States / Colorado / Weld County | Mary Lea Killian (National Veterinary Services Laboratories - USDA / Diagnostic Virology Laboratory) |
| EPI_ISL_16171468 | A/guinea fowl/Wisconsin/22-013061-003-original/2022 | A / H5N1 | North America / United States / Wisconsin / Oconto County | Mary Lea Killian (National Veterinary Services Laboratories - USDA / Diagnostic Virology Laboratory) |
| EPI_ISL_16171469 | A/peacock/Wisconsin/22-013061-004-original/2022 | A / H5N1 | North America / United States / Wisconsin / Oconto County | Mary Lea Killian (National Veterinary Services Laboratories - USDA / Diagnostic Virology Laboratory) |
| EPI_ISL_16171470 | A/Lady Amherst's pheasant/Wisconsin/22-013061-005-origina | A / H5N1 | North America / United States / Wisconsin / Oconto County | Mary Lea Killian (National Veterinary Services Laboratories - USDA / Diagnostic Virology Laboratory) |
| EPI_ISL_16171471 | A/turkey/Minnesota/22-013084-001-original/2022 | A / H5N1 | North America / United States / Minnesota / Stearns County | Mary Lea Killian (National Veterinary Services Laboratories - USDA / Diagnostic Virology Laboratory) |
| EPI_ISL_16171472 | A/turkey/Minnesota/22-013084-002-original/2022 | A / H5N1 | North America / United States / Minnesota / Stearns County | Mary Lea Killian (National Veterinary Services Laboratories - USDA / Diagnostic Virology Laboratory) |
| EPI_ISL_16171473 | A/chicken/Nebraska/22-013105-001-original/2022 | A / H5N1 | North America / United States / Nebraska / Washington County | Mary Lea Killian (National Veterinary Services Laboratories - USDA / Diagnostic Virology Laboratory) |
| EPI_ISL_16171474 | A/chicken/Alaska/22-013107-001-original/2022 | A / H5N1 | North America / United States / Alaska / Matanuska-Susitna Borough | Mary Lea Killian (National Veterinary Services Laboratories - USDA / Diagnostic Virology Laboratory) |
| EPI_ISL_16171475 | A/chicken/Alaska/22-013107-002-original/2022 | A / H5N1 | North America / United States / Alaska / Matanuska-Susitna Borough | Mary Lea Killian (National Veterinary Services Laboratories - USDA / Diagnostic Virology Laboratory) |
| EPI_ISL_16171476 | A/turkey/Wisconsin/22-013183-002-original/2022 | A / H5N1 | North America / United States / Wisconsin / Barron County | Mary Lea Killian (National Veterinary Services Laboratories - USDA / Diagnostic Virology Laboratory) |
| EPI_ISL_16171477 | A/guinea fowl/Wisconsin/22-013183-003-original/2022 | A / H5N1 | North America / United States / Wisconsin / Barron County | Mary Lea Killian (National Veterinary Services Laboratories - USDA / Diagnostic Virology Laboratory) |
| EPI_ISL_16171478 | A/domestic duck/Wisconsin/22-013183-004-original/2022 | A / H5N1 | North America / United States / Wisconsin / Barron County | Mary Lea Killian (National Veterinary Services Laboratories - USDA / Diagnostic Virology Laboratory) |
| EPI_ISL_16171479 | A/peacock/Wisconsin/22-013183-005-original/2022 | A / H5N1 | North America / United States / Wisconsin / Barron County | Mary Lea Killian (National Veterinary Services Laboratories - USDA / Diagnostic Virology Laboratory) |
| EPI_ISL_16171480 | A/Pekin duck/Pennsylvania/22-013189-003-original/2022 | A / H5N1 | North America / United States / Pennsylvania / Lancaster County | Mary Lea Killian (National Veterinary Services Laboratories - USDA / Diagnostic Virology Laboratory) |
| EPI_ISL_16171481 | A/Pekin duck/Pennsylvania/22-013189-004-original/2022 | A / H5N1 | North America / United States / Pennsylvania / Lancaster County | Mary Lea Killian (National Veterinary Services Laboratories - USDA / Diagnostic Virology Laboratory) |
| EPI_ISL_16171482 | A/chicken/Oklahoma/22-013220-001-original/2022 | A / H5N1 | North America / United States / Oklahoma / Sequoyah County | Mary Lea Killian (National Veterinary Services Laboratories - USDA / Diagnostic Virology Laboratory) |
| EPI_ISL_16171483 | A/chicken/Oklahoma/22-013220-002-original/2022 | A / H5N1 | North America / United States / Oklahoma / Sequoyah County | Mary Lea Killian (National Veterinary Services Laboratories - USDA / Diagnostic Virology Laboratory) |
| EPI_ISL_16171485 | A/goose/Minnesota/22-013221-002-original/2022 | A / H5N1 | North America / United States / Minnesota / Morrison County | Mary Lea Killian (National Veterinary Services Laboratories - USDA / Diagnostic Virology Laboratory) |
| EPI_ISL_16171486 | A/duck/Minnesota/22-013221-003-original/2022 | A / H5N1 | North America / United States / Minnesota / Morrison County | Mary Lea Killian (National Veterinary Services Laboratories - USDA / Diagnostic Virology Laboratory) |
| EPI_ISL_16171487 | A/duck/Minnesota/22-013221-004-original/2022 | A / H5N1 | North America / United States / Minnesota / Morrison County | Mary Lea Killian (National Veterinary Services Laboratories - USDA / Diagnostic Virology Laboratory) |
| EPI_ISL_16171488 | A/chicken/Minnesota/22-013222-001-original/2022 | A / H5N1 | North America / United States / Minnesota / Chisago County | Mary Lea Killian (National Veterinary Services Laboratories - USDA / Diagnostic Virology Laboratory) |
| EPI_ISL_16171490 | A/goose/Minnesota/22-013222-004-original/2022 | A / H5N1 | North America / United States / Minnesota / Chisago County | Mary Lea Killian (National Veterinary Services Laboratories - USDA / Diagnostic Virology Laboratory) |
| EPI_ISL_16171491 | A/goose/Minnesota/22-013222-005-original/2022 | A / H5N1 | North America / United States / Minnesota / Chisago County | Mary Lea Killian (National Veterinary Services Laboratories - USDA / Diagnostic Virology Laboratory) |
| EPI_ISL_16171492 | A/chicken/Minnesota/22-013223-001-original/2022 | A / H5N1 | North America / United States / Minnesota / Chisago County | Mary Lea Killian (National Veterinary Services Laboratories - USDA / Diagnostic Virology Laboratory) |
| EPI_ISL_16171493 | A/chicken/Minnesota/22-013223-003-original/2022 | A / H5N1 | North America / United States / Minnesota / Chisago County | Mary Lea Killian (National Veterinary Services Laboratories - USDA / Diagnostic Virology Laboratory) |
| EPI_ISL_16171494 | A/chicken/Michigan/22-013224-001-original/2022 | A / H5N1 | North America / United States / Michigan / Branch County | Mary Lea Killian (National Veterinary Services Laboratories - USDA / Diagnostic Virology Laboratory) |
| EPI_ISL_16171495 | A/duck/Pennsylvania/22-013368-002-original/2022 | A / H5N1 | North America / United States / Pennsylvania / Berks County | Mary Lea Killian (National Veterinary Services Laboratories - USDA / Diagnostic Virology Laboratory) |
| EPI_ISL_16171496 | A/duck/Wisconsin/22-013373-001-original/2022 | A / H5N1 | North America / United States / Wisconsin / Sauk County | Mary Lea Killian (National Veterinary Services Laboratories - USDA / Diagnostic Virology Laboratory) |
| EPI_ISL_16171497 | A/chicken/Wisconsin/22-013373-002-original/2022 | A / H5N1 | North America / United States / Wisconsin / Sauk County | Mary Lea Killian (National Veterinary Services Laboratories - USDA / Diagnostic Virology Laboratory) |
| EPI_ISL_16171498 | A/chicken/Illinois/22-013392-001-original/2022 | A / H5N1 | North America / United States / Illinois / Kane County | Mary Lea Killian (National Veterinary Services Laboratories - USDA / Diagnostic Virology Laboratory) |
| EPI_ISL_16171499 | A/chicken/Illinois/22-013392-002-original/2022 | A / H5N1 | North America / United States / Illinois / Kane County | Mary Lea Killian (National Veterinary Services Laboratories - USDA / Diagnostic Virology Laboratory) |
| EPI_ISL_16171500 | A/chicken/Minnesota/22-013412-001-original/2022 | A / H5N1 | North America / United States / Minnesota / Chisago County | Mary Lea Killian (National Veterinary Services Laboratories - USDA / Diagnostic Virology Laboratory) |
| EPI_ISL_16171501 | A/chicken/Minnesota/22-013412-002-original/2022 | A / H5N1 | North America / United States / Minnesota / Chisago County | Mary Lea Killian (National Veterinary Services Laboratories - USDA / Diagnostic Virology Laboratory) |
| EPI_ISL_16171502 | A/turkey/Minnesota/22-013413-002-original/2022 | A / H5N1 | North America / United States / Minnesota / Lyon County | Mary Lea Killian (National Veterinary Services Laboratories - USDA / Diagnostic Virology Laboratory) |
| EPI_ISL_16171503 | A/turkey/Minnesota/22-013413-003-original/2022 | A / H5N1 | North America / United States / Minnesota / Lyon County | Mary Lea Killian (National Veterinary Services Laboratories - USDA / Diagnostic Virology Laboratory) |
| EPI_ISL_16171504 | A/chicken/Wisconsin/22-013445-001-original/2022 | A / H5N1 | North America / United States / Wisconsin / Fond du Lac County | Mary Lea Killian (National Veterinary Services Laboratories - USDA / Diagnostic Virology Laboratory) |
| EPI_ISL_16171505 | A/chicken/Wisconsin/22-013446-001-original/2022 | A / H5N1 | North America / United States / Wisconsin / Polk County | Mary Lea Killian (National Veterinary Services Laboratories - USDA / Diagnostic Virology Laboratory) |
| EPI_ISL_16171506 | A/turkey/Wisconsin/22-013447-001-original/2022 | A / H5N1 | North America / United States / Wisconsin / Barron County | Mary Lea Killian (National Veterinary Services Laboratories - USDA / Diagnostic Virology Laboratory) |
| EPI_ISL_16171507 | A/turkey/Wisconsin/22-013447-002-original/2022 | A / H5N1 | North America / United States / Wisconsin / Barron County | Mary Lea Killian (National Veterinary Services Laboratories - USDA / Diagnostic Virology Laboratory) |
| EPI_ISL_16171508 | A/turkey/Wisconsin/22-013448-001-original/2022 | A / H5N1 | North America / United States / Wisconsin / Barron County | Mary Lea Killian (National Veterinary Services Laboratories - USDA / Diagnostic Virology Laboratory) |
| EPI_ISL_16171509 | A/turkey/Wisconsin/22-013448-002-original/2022 | A / H5N1 | North America / United States / Wisconsin / Barron County | Mary Lea Killian (National Veterinary Services Laboratories - USDA / Diagnostic Virology Laboratory) |
| EPI_ISL_16171510 | A/turkey/South Dakota/22-013454-001-original/2022 | A / H5N1 | North America / United States / South Dakota / Beadle County | Mary Lea Killian (National Veterinary Services Laboratories - USDA / Diagnostic Virology Laboratory) |
| EPI_ISL_16171511 | A/turkey/South Dakota/22-013454-002-original/2022 | A / H5N1 | North America / United States / South Dakota / Beadle County | Mary Lea Killian (National Veterinary Services Laboratories - USDA / Diagnostic Virology Laboratory) |
| EPI_ISL_16171512 | A/turkey/South Dakota/22-013547-001-original/2022 | A / H5N1 | North America / United States / South Dakota / Hamlin County | Mary Lea Killian (National Veterinary Services Laboratories - USDA / Diagnostic Virology Laboratory) |
| EPI_ISL_16171513 | A/chicken/South Dakota/22-013554-001-original/2022 | A / H5N1 | North America / United States / South Dakota / Day County | Mary Lea Killian (National Veterinary Services Laboratories - USDA / Diagnostic Virology Laboratory) |
| EPI_ISL_16171514 | A/duck/Minnesota/22-013605-001-original/2022 | A / H5N1 | North America / United States / Minnesota / Carver County | Mary Lea Killian (National Veterinary Services Laboratories - USDA / Diagnostic Virology Laboratory) |
| EPI_ISL_16171516 | A/duck/Minnesota/22-013605-003-original/2022 | A / H5N1 | North America / United States / Minnesota / Carver County | Mary Lea Killian (National Veterinary Services Laboratories - USDA / Diagnostic Virology Laboratory) |
| EPI_ISL_16171517 | A/turkey/Utah/22-013612-001-original/2022 | A / H5N1 | North America / United States / Utah / Cache County | Mary Lea Killian (National Veterinary Services Laboratories - USDA / Diagnostic Virology Laboratory) |
| EPI_ISL_16171518 | A/chicken/Minnesota/22-013752-001-original/2022 | A / H5N1 | North America / United States / Minnesota / Anoka County | Mary Lea Killian (National Veterinary Services Laboratories - USDA / Diagnostic Virology Laboratory) |
| EPI_ISL_16171519 | A/chicken/Washington/22-013763-002-original/2022 | A / H5N1 | North America / United States / Washington / Pacific County | Mary Lea Killian (National Veterinary Services Laboratories - USDA / Diagnostic Virology Laboratory) |
| EPI_ISL_16171520 | A/chicken/Washington/22-013763-003-original/2022 | A / H5N1 | North America / United States / Washington / Pacific County | Mary Lea Killian (National Veterinary Services Laboratories - USDA / Diagnostic Virology Laboratory) |
| EPI_ISL_16171521 | A/chicken/Wisconsin/22-013776-001-original/2022 | A / H5N1 | North America / United States / Wisconsin / Pierce County | Mary Lea Killian (National Veterinary Services Laboratories - USDA / Diagnostic Virology Laboratory) |
| EPI_ISL_16171522 | A/goose/Oregon/22-013779-001-original/2022 | A / H5N1 | North America / United States / Oregon / Linn County | Mary Lea Killian (National Veterinary Services Laboratories - USDA / Diagnostic Virology Laboratory) |
| EPI_ISL_16171523 | A/chicken/Montana/22-013942-001-original/2022 | A / H5N1 | North America / United States / Montana / Fergus County | Mary Lea Killian (National Veterinary Services Laboratories - USDA / Diagnostic Virology Laboratory) |
| EPI_ISL_16171524 | A/chicken/Montana/22-013942-002-original/2022 | A / H5N1 | North America / United States / Montana / Fergus County | Mary Lea Killian (National Veterinary Services Laboratories - USDA / Diagnostic Virology Laboratory) |
| EPI_ISL_16196676 | A/duck/Cyprus/750-2_22VIR12133-10/2022 | A / H5N1 | Europe / Cyprus | Ambra Pastori (Istituto Zooprofilattico Sperimentale Delle Venezie) |
| EPI_ISL_16196756 | A/goose/Cyprus/750-3_22VIR12133-11/2022 | A / H5N1 | Europe / Cyprus | Ambra Pastori (Istituto Zooprofilattico Sperimentale Delle Venezie) |
| EPI_ISL_16196757 | A/duck/Cyprus/752-2_22VIR12133-13/2022 | A / H5N1 | Europe / Cyprus | Ambra Pastori (Istituto Zooprofilattico Sperimentale Delle Venezie) |
| EPI_ISL_16196758 | A/duck/Cyprus/745-1_22VIR12133-1/2022 | A / H5N1 | Europe / Cyprus | Ambra Pastori (Istituto Zooprofilattico Sperimentale Delle Venezie) |
| EPI_ISL_16196759 | A/goose/Cyprus/746-1_22VIR12133-3/2022 | A / H5N1 | Europe / Cyprus | Ambra Pastori (Istituto Zooprofilattico Sperimentale Delle Venezie) |
| EPI_ISL_16196760 | A/duck/Cyprus/749-1_22VIR12133-5/2022 | A / H5N1 | Europe / Cyprus | Ambra Pastori (Istituto Zooprofilattico Sperimentale Delle Venezie) |
| EPI_ISL_16196761 | A/goose/Cyprus/749-3_22VIR12133-7/2022 | A / H5N1 | Europe / Cyprus | Ambra Pastori (Istituto Zooprofilattico Sperimentale Delle Venezie) |
| EPI_ISL_161972 | A/duck/Jiangxi/95/2014 | A / H5N6 | Asia / China / Jiangxi Province | Guang Liu (Beijing Genomics Institute (BGI)) |
| EPI_ISL_161973 | A/environment/Shenzhen/25-24/2013 | A / H5N6 | Asia / China / Guangdong Province | Guang Liu (Beijing Genomics Institute (BGI)) |
| EPI_ISL_16199040 | A/chicken/Austria/2514-19_22VIR11382-1/2022 | A / H5N1 | Europe / Austria | Ambra Pastori (Istituto Zooprofilattico Sperimentale Delle Venezie) |
| EPI_ISL_16199509 | A/chicken/Austria/2514-20_22VIR11382-2/2022 | A / H5N1 | Europe / Austria | Ambra Pastori (Istituto Zooprofilattico Sperimentale Delle Venezie) |
| EPI_ISL_16206845 | A/goose/Luxembourg/22253283/2022 | A / H5N1 | Europe / Luxembourg / District de Diekirch | Chantal J. Snoeck (Luxembourg Institute of Health / Department of Infection and Immunity) |
| EPI_ISL_162300 | A/duck/Zhejiang/W24/2013 | A / H5N8 | Asia / China | |

| | | | | |
|---|---|---|---|---|
| EPI_ISL_162301 | A/duck/Zhejiang/6D18/2013 | A / H5N8 | Asia / China | |
| EPI_ISL_16240161 | A/domestic goose/Washington/22-014023-001-original/2022 | A / H5N1 | North America / United States / Washington / Spokane County | Mary Lea Killian (National Veterinary Services Laboratories - USDA / Diagnostic Virology Laboratory) |
| EPI_ISL_16240162 | A/turkey/Wisconsin/22-014125-001-original/2022 | A / H5N1 | North America / United States / Wisconsin / Barron County | Mary Lea Killian (National Veterinary Services Laboratories - USDA / Diagnostic Virology Laboratory) |
| EPI_ISL_16240163 | A/turkey/Wisconsin/22-014126-001-original/2022 | A / H5N1 | North America / United States / Wisconsin / Barron County | Mary Lea Killian (National Veterinary Services Laboratories - USDA / Diagnostic Virology Laboratory) |
| EPI_ISL_16240164 | A/turkey/Wisconsin/22-014126-002-original/2022 | A / H5N1 | North America / United States / Wisconsin / Barron County | Mary Lea Killian (National Veterinary Services Laboratories - USDA / Diagnostic Virology Laboratory) |
| EPI_ISL_16240165 | A/chicken/Nebraska/22-014140-001-original/2022 | A / H5N1 | North America / United States / Nebraska / Knox County | Mary Lea Killian (National Veterinary Services Laboratories - USDA / Diagnostic Virology Laboratory) |
| EPI_ISL_16240166 | A/chicken/Nebraska/22-014140-002-original/2022 | A / H5N1 | North America / United States / Nebraska / Knox County | Mary Lea Killian (National Veterinary Services Laboratories - USDA / Diagnostic Virology Laboratory) |
| EPI_ISL_16240167 | A/chicken/Nebraska/22-014141-002-original/2022 | A / H5N1 | North America / United States / Nebraska / Knox County | Mary Lea Killian (National Veterinary Services Laboratories - USDA / Diagnostic Virology Laboratory) |
| EPI_ISL_16240168 | A/chicken/Nebraska/22-014141-003-original/2022 | A / H5N1 | North America / United States / Nebraska / Knox County | Mary Lea Killian (National Veterinary Services Laboratories - USDA / Diagnostic Virology Laboratory) |
| EPI_ISL_16240169 | A/chicken/Idaho/22-014175-001-original/2022 | A / H5N1 | North America / United States / Idaho / Canyon County | Mary Lea Killian (National Veterinary Services Laboratories - USDA / Diagnostic Virology Laboratory) |
| EPI_ISL_16240220 | A/chicken/Idaho/22-014175-003-original/2022 | A / H5N1 | North America / United States / Idaho / Canyon County | Mary Lea Killian (National Veterinary Services Laboratories - USDA / Diagnostic Virology Laboratory) |
| EPI_ISL_16240221 | A/turkey/Michigan/22-014263-001-original/2022 | A / H5N1 | North America / United States / Michigan / Muskegon County | Mary Lea Killian (National Veterinary Services Laboratories - USDA / Diagnostic Virology Laboratory) |
| EPI_ISL_16240223 | A/chicken/Idaho/22-014275-001-original/2022 | A / H5N1 | North America / United States / Idaho / Ada County | Mary Lea Killian (National Veterinary Services Laboratories - USDA / Diagnostic Virology Laboratory) |
| EPI_ISL_16240224 | A/chicken/Idaho/22-014275-002-original/2022 | A / H5N1 | North America / United States / Idaho / Ada County | Mary Lea Killian (National Veterinary Services Laboratories - USDA / Diagnostic Virology Laboratory) |
| EPI_ISL_16240225 | A/duck/Idaho/22-014275-003-original/2022 | A / H5N1 | North America / United States / Idaho / Ada County | Mary Lea Killian (National Veterinary Services Laboratories - USDA / Diagnostic Virology Laboratory) |
| EPI_ISL_16240227 | A/chicken/Minnesota/22-014294-002-original/2022 | A / H5N1 | North America / United States / Minnesota / Crow Wing County | Mary Lea Killian (National Veterinary Services Laboratories - USDA / Diagnostic Virology Laboratory) |
| EPI_ISL_16240228 | A/turkey/Minnesota/22-014294-003-original/2022 | A / H5N1 | North America / United States / Minnesota / Crow Wing County | Mary Lea Killian (National Veterinary Services Laboratories - USDA / Diagnostic Virology Laboratory) |
| EPI_ISL_16240229 | A/turkey/Minnesota/22-014294-006-original/2022 | A / H5N1 | North America / United States / Minnesota / Crow Wing County | Mary Lea Killian (National Veterinary Services Laboratories - USDA / Diagnostic Virology Laboratory) |
| EPI_ISL_16240239 | A/goose/Minnesota/22-014294-007-original/2022 | A / H5N1 | North America / United States / Minnesota / Crow Wing County | Mary Lea Killian (National Veterinary Services Laboratories - USDA / Diagnostic Virology Laboratory) |
| EPI_ISL_16240351 | A/chicken/Minnesota/22-014295-001-original/2022 | A / H5N1 | North America / United States / Minnesota / Todd County | Mary Lea Killian (National Veterinary Services Laboratories - USDA / Diagnostic Virology Laboratory) |
| EPI_ISL_16240352 | A/chicken/Minnesota/22-014295-002-original/2022 | A / H5N1 | North America / United States / Minnesota / Todd County | Mary Lea Killian (National Veterinary Services Laboratories - USDA / Diagnostic Virology Laboratory) |
| EPI_ISL_16240353 | A/duck/Minnesota/22-014295-003-original/2022 | A / H5N1 | North America / United States / Minnesota / Todd County | Mary Lea Killian (National Veterinary Services Laboratories - USDA / Diagnostic Virology Laboratory) |
| EPI_ISL_16240361 | A/turkey/Washington/22-014316-001-original/2022 | A / H5N1 | North America / United States / Washington / Pierce County | Mary Lea Killian (National Veterinary Services Laboratories - USDA / Diagnostic Virology Laboratory) |
| EPI_ISL_16240372 | A/duck/Washington/22-014327-001-original/2022 | A / H5N1 | North America / United States / Washington / Pierce County | Mary Lea Killian (National Veterinary Services Laboratories - USDA / Diagnostic Virology Laboratory) |
| EPI_ISL_16240382 | A/duck/Washington/22-014327-002-original/2022 | A / H5N1 | North America / United States / Washington / Pierce County | Mary Lea Killian (National Veterinary Services Laboratories - USDA / Diagnostic Virology Laboratory) |
| EPI_ISL_16240392 | A/chicken/Washington/22-014327-003-original/2022 | A / H5N1 | North America / United States / Washington / Pierce County | Mary Lea Killian (National Veterinary Services Laboratories - USDA / Diagnostic Virology Laboratory) |
| EPI_ISL_16240404 | A/duck/Pennsylvania/22-014337-007-original/2022 | A / H5N1 | North America / United States / Pennsylvania / Berks County | Mary Lea Killian (National Veterinary Services Laboratories - USDA / Diagnostic Virology Laboratory) |
| EPI_ISL_16240414 | A/chicken/Pennsylvania/22-014338-001-original/2022 | A / H5N1 | North America / United States / Pennsylvania / Lancaster County | Mary Lea Killian (National Veterinary Services Laboratories - USDA / Diagnostic Virology Laboratory) |
| EPI_ISL_16240423 | A/chicken/Pennsylvania/22-014338-002-original/2022 | A / H5N1 | North America / United States / Pennsylvania / Lancaster County | Mary Lea Killian (National Veterinary Services Laboratories - USDA / Diagnostic Virology Laboratory) |
| EPI_ISL_16240434 | A/duck/Pennsylvania/22-014339-001-original/2022 | A / H5N1 | North America / United States / Pennsylvania / Berks County | Mary Lea Killian (National Veterinary Services Laboratories - USDA / Diagnostic Virology Laboratory) |
| EPI_ISL_16240445 | A/duck/Pennsylvania/22-014339-002-original/2022 | A / H5N1 | North America / United States / Pennsylvania / Berks County | Mary Lea Killian (National Veterinary Services Laboratories - USDA / Diagnostic Virology Laboratory) |
| EPI_ISL_16240456 | A/chicken/Idaho/22-014425-002-original/2022 | A / H5N1 | North America / United States / Idaho / Ada County | Mary Lea Killian (National Veterinary Services Laboratories - USDA / Diagnostic Virology Laboratory) |
| EPI_ISL_16240466 | A/chicken/Washington/22-014449-001-original/2022 | A / H5N1 | North America / United States / Washington / Clallam County | Mary Lea Killian (National Veterinary Services Laboratories - USDA / Diagnostic Virology Laboratory) |
| EPI_ISL_16240476 | A/chicken/Washington/22-014449-002-original/2022 | A / H5N1 | North America / United States / Washington / Clallam County | Mary Lea Killian (National Veterinary Services Laboratories - USDA / Diagnostic Virology Laboratory) |
| EPI_ISL_16240486 | A/goose/Washington/22-014450-001-original/2022 | A / H5N1 | North America / United States / Washington / Clallam county | Mary Lea Killian (National Veterinary Services Laboratories - USDA / Diagnostic Virology Laboratory) |
| EPI_ISL_16240497 | A/duck/Utah/22-014481-001-original/2022 | A / H5N1 | North America / United States / Utah / Cache County | Mary Lea Killian (National Veterinary Services Laboratories - USDA / Diagnostic Virology Laboratory) |
| EPI_ISL_16240508 | A/guinea fowl/Utah/22-014481-002-original/2022 | A / H5N1 | North America / United States / Utah / Cache County | Mary Lea Killian (National Veterinary Services Laboratories - USDA / Diagnostic Virology Laboratory) |
| EPI_ISL_16240519 | A/chicken/Minnesota/22-014503-001-original/2022 | A / H5N1 | North America / United States / Minnesota / Chisago County | Mary Lea Killian (National Veterinary Services Laboratories - USDA / Diagnostic Virology Laboratory) |
| EPI_ISL_16240529 | A/chicken/Minnesota/22-014503-002-original/2022 | A / H5N1 | North America / United States / Minnesota / Chisago County | Mary Lea Killian (National Veterinary Services Laboratories - USDA / Diagnostic Virology Laboratory) |
| EPI_ISL_16240634 | A/fox/Michigan/22-014536-001-original/2022 | A / H5N1 | North America / United States / Michigan / Macomb County | Mary Lea Killian (National Veterinary Services Laboratories - USDA / Diagnostic Virology Laboratory) |
| EPI_ISL_16240635 | A/fox/Michigan/22-014536-002-original/2022 | A / H5N1 | North America / United States / Michigan / Macomb County | Mary Lea Killian (National Veterinary Services Laboratories - USDA / Diagnostic Virology Laboratory) |
| EPI_ISL_16240636 | A/fox/Michigan/22-014536-007-original/2022 | A / H5N1 | North America / United States / Michigan / Saint Clair County | Mary Lea Killian (National Veterinary Services Laboratories - USDA / Diagnostic Virology Laboratory) |
| EPI_ISL_16240637 | A/fox/Michigan/22-014536-008-original/2022 | A / H5N1 | North America / United States / Michigan / Saint Clair County | Mary Lea Killian (National Veterinary Services Laboratories - USDA / Diagnostic Virology Laboratory) |
| EPI_ISL_16240638 | A/chicken/Minnesota/22-014663-001-original/2022 | A / H5N1 | North America / United States / Minnesota / Chisago County | Mary Lea Killian (National Veterinary Services Laboratories - USDA / Diagnostic Virology Laboratory) |
| EPI_ISL_16240639 | A/chicken/Minnesota/22-014663-002-original/2022 | A / H5N1 | North America / United States / Minnesota / Chisago County | Mary Lea Killian (National Veterinary Services Laboratories - USDA / Diagnostic Virology Laboratory) |
| EPI_ISL_16240640 | A/duck/Minnesota/22-014663-003-original/2022 | A / H5N1 | North America / United States / Minnesota / Chisago County | Mary Lea Killian (National Veterinary Services Laboratories - USDA / Diagnostic Virology Laboratory) |
| EPI_ISL_16240641 | A/duck/Minnesota/22-014663-004-original/2022 | A / H5N1 | North America / United States / Minnesota / Chisago County | Mary Lea Killian (National Veterinary Services Laboratories - USDA / Diagnostic Virology Laboratory) |
| EPI_ISL_16240642 | A/turkey/Idaho/22-014701-001-original/2022 | A / H5N1 | North America / United States / Idaho / Ada County | Mary Lea Killian (National Veterinary Services Laboratories - USDA / Diagnostic Virology Laboratory) |
| EPI_ISL_16240643 | A/chicken/Idaho/22-014701-002-original/2022 | A / H5N1 | North America / United States / Idaho / Ada County | Mary Lea Killian (National Veterinary Services Laboratories - USDA / Diagnostic Virology Laboratory) |
| EPI_ISL_16240644 | A/chicken/Minnesota/22-014734-001-original/2022 | A / H5N1 | North America / United States / Minnesota / Grant County | Mary Lea Killian (National Veterinary Services Laboratories - USDA / Diagnostic Virology Laboratory) |
| EPI_ISL_16240880 | A/chicken/Minnesota/22-014734-002-original/2022 | A / H5N1 | North America / United States / Minnesota / Grant County | Mary Lea Killian (National Veterinary Services Laboratories - USDA / Diagnostic Virology Laboratory) |
| EPI_ISL_16240881 | A/chicken/Washington/22-014739-001-original/2022 | A / H5N1 | North America / United States / Washington / Okanogan County | Mary Lea Killian (National Veterinary Services Laboratories - USDA / Diagnostic Virology Laboratory) |
| EPI_ISL_16240882 | A/chicken/Washington/22-014744-001-original/2022 | A / H5N1 | North America / United States / Washington / Whatcom County | Mary Lea Killian (National Veterinary Services Laboratories - USDA / Diagnostic Virology Laboratory) |
| EPI_ISL_16240883 | A/goose/Washington/22-014744-003-original/2022 | A / H5N1 | North America / United States / Washington / Whatcom County | Mary Lea Killian (National Veterinary Services Laboratories - USDA / Diagnostic Virology Laboratory) |
| EPI_ISL_16240884 | A/goose/Washington/22-014744-004-original/2022 | A / H5N1 | North America / United States / Washington / Whatcom County | Mary Lea Killian (National Veterinary Services Laboratories - USDA / Diagnostic Virology Laboratory) |
| EPI_ISL_16240885 | A/chicken/Wisconsin/22-014859-001-original/2022 | A / H5N1 | North America / United States / Wisconsin / Marinette County | Mary Lea Killian (National Veterinary Services Laboratories - USDA / Diagnostic Virology Laboratory) |
| EPI_ISL_16240886 | A/chicken/Wyoming/22-014916-001-original/2022 | A / H5N1 | North America / United States / Wyoming / Sheridan County | Mary Lea Killian (National Veterinary Services Laboratories - USDA / Diagnostic Virology Laboratory) |
| EPI_ISL_16240887 | A/chicken/Wyoming/22-014917-001-original/2022 | A / H5N1 | North America / United States / Wyoming / Sheridan County | Mary Lea Killian (National Veterinary Services Laboratories - USDA / Diagnostic Virology Laboratory) |
| EPI_ISL_16240888 | A/chicken/Wyoming/22-014918-001-original/2022 | A / H5N1 | North America / United States / Wyoming / Lincoln County | Mary Lea Killian (National Veterinary Services Laboratories - USDA / Diagnostic Virology Laboratory) |
| EPI_ISL_16240889 | A/chicken/Wyoming/22-014920-001-original/2022 | A / H5N1 | North America / United States / Wyoming / Lincoln County | Mary Lea Killian (National Veterinary Services Laboratories - USDA / Diagnostic Virology Laboratory) |
| EPI_ISL_16240985 | A/guinea fowl/Minnesota/22-014734-003-original/2022 | A / H5N1 | North America / United States / Minnesota / Grant County | Mary Lea Killian (National Veterinary Services Laboratories - USDA / Diagnostic Virology Laboratory) |
| EPI_ISL_16243202 | A/chicken/Idaho/22-014942-001-original/2022 | A / H5N1 | North America / United States / Idaho / Canyon County | Mary Lea Killian (National Veterinary Services Laboratories - USDA / Diagnostic Virology Laboratory) |
| EPI_ISL_16243210 | A/goose/Idaho/22-014944-001-original/2022 | A / H5N1 | North America / United States / Idaho / Ada County | Mary Lea Killian (National Veterinary Services Laboratories - USDA / Diagnostic Virology Laboratory) |
| EPI_ISL_16243221 | A/turkey/Wisconsin/22-014948-001-original/2022 | A / H5N1 | North America / United States / Wisconsin / Barron County | Mary Lea Killian (National Veterinary Services Laboratories - USDA / Diagnostic Virology Laboratory) |
| EPI_ISL_16243231 | A/chicken/Pennsylvania/22-015036-001-original/2022 | A / H5N1 | North America / United States / Pennsylvania / Berks County | Mary Lea Killian (National Veterinary Services Laboratories - USDA / Diagnostic Virology Laboratory) |
| EPI_ISL_16243241 | A/chicken/Pennsylvania/22-015036-002-original/2022 | A / H5N1 | North America / United States / Pennsylvania / Berks County | Mary Lea Killian (National Veterinary Services Laboratories - USDA / Diagnostic Virology Laboratory) |
| EPI_ISL_16243253 | A/fox/New York/22-015063-001-original/2022 | A / H5N1 | North America / United States / New York / Saint Lawrence County | Mary Lea Killian (National Veterinary Services Laboratories - USDA / Diagnostic Virology Laboratory) |
| EPI_ISL_16243264 | A/fox/New York/22-015064-001-original/2022 | A / H5N1 | North America / United States / New York / Erie County | Mary Lea Killian (National Veterinary Services Laboratories - USDA / Diagnostic Virology Laboratory) |
| EPI_ISL_16243273 | A/fox/New York/22-015065-001-original/2022 | A / H5N1 | North America / United States / New York / Erie County | Mary Lea Killian (National Veterinary Services Laboratories - USDA / Diagnostic Virology Laboratory) |
| EPI_ISL_16243294 | A/chicken/Wisconsin/22-015068-003-original/2022 | A / H5N1 | North America / United States / Wisconsin / Dunn County | Mary Lea Killian (National Veterinary Services Laboratories - USDA / Diagnostic Virology Laboratory) |
| EPI_ISL_16243306 | A/chicken/Wisconsin/22-015068-005-original/2022 | A / H5N1 | North America / United States / Wisconsin / Dunn County | Mary Lea Killian (National Veterinary Services Laboratories - USDA / Diagnostic Virology Laboratory) |
| EPI_ISL_16243316 | A/chicken/Idaho/22-015073-001-original/2022 | A / H5N1 | North America / United States / Idaho / Ada County | Mary Lea Killian (National Veterinary Services Laboratories - USDA / Diagnostic Virology Laboratory) |

| | | | | |
|---|---|---|---|---|
| EPI_ISL_16243326 | A/chicken/North Dakota/22-015199-001-original/2022 | A / H5N1 | North America / United States / North Dakota / Burke County | Mary Lea Killian (National Veterinary Services Laboratories - USDA / Diagnostic Virology Laboratory) |
| EPI_ISL_16243588 | A/chicken/North Dakota/22-015199-002-original/2022 | A / H5N1 | North America / United States / North Dakota / Burke County | Mary Lea Killian (National Veterinary Services Laboratories - USDA / Diagnostic Virology Laboratory) |
| EPI_ISL_16243598 | A/chicken/Pennsylvania/22-015201-001-original/2022 | A / H5N1 | North America / United States / Pennsylvania / Berks County | Mary Lea Killian (National Veterinary Services Laboratories - USDA / Diagnostic Virology Laboratory) |
| EPI_ISL_16243610 | A/chicken/Pennsylvania/22-015201-002-original/202 | A / H5N1 | North America / United States / Pennsylvania / Berks County | Mary Lea Killian (National Veterinary Services Laboratories - USDA / Diagnostic Virology Laboratory) |
| EPI_ISL_16243619 | A/chicken/Washington/22-015210-001-original/2022 | A / H5N1 | North America / United States / Washington / Thurston County | Mary Lea Killian (National Veterinary Services Laboratories - USDA / Diagnostic Virology Laboratory) |
| EPI_ISL_16243630 | A/chicken/Oregon/22-015228-001-original/2022 | A / H5N1 | North America / United States / Oregon / Lane County | Mary Lea Killian (National Veterinary Services Laboratories - USDA / Diagnostic Virology Laboratory) |
| EPI_ISL_16243641 | A/chicken/Illinois/22-015233-005-original/2022 | A / H5N1 | North America / United States / Illinois / Boone County | Mary Lea Killian (National Veterinary Services Laboratories - USDA / Diagnostic Virology Laboratory) |
| EPI_ISL_16243651 | A/chicken/Illinois/22-015233-006-original/2022 | A / H5N1 | North America / United States / Illinois / Boone County | Mary Lea Killian (National Veterinary Services Laboratories - USDA / Diagnostic Virology Laboratory) |
| EPI_ISL_16243660 | A/chicken/Utah/22-015245-001-original/2022 | A / H5N1 | North America / United States / Utah / Salt Lake County | Mary Lea Killian (National Veterinary Services Laboratories - USDA / Diagnostic Virology Laboratory) |
| EPI_ISL_16243667 | A/goose/Utah/22-015245-002-original/2022 | A / H5N1 | North America / United States / Utah / Salt Lake County | Mary Lea Killian (National Veterinary Services Laboratories - USDA / Diagnostic Virology Laboratory) |
| EPI_ISL_16243872 | A/duck/New Jersey/22-015259-001-original/2022 | A / H5N1 | North America / United States / New Jersey / Monmouth County | Mary Lea Killian (National Veterinary Services Laboratories - USDA / Diagnostic Virology Laboratory) |
| EPI_ISL_16243883 | A/duck/New Jersey/22-015259-002-original/2022 | A / H5N1 | North America / United States / New Jersey / Monmouth County | Mary Lea Killian (National Veterinary Services Laboratories - USDA / Diagnostic Virology Laboratory) |
| EPI_ISL_16243894 | A/chicken/Idaho/22-015271-002-original/2022 | A / H5N1 | North America / United States / Idaho / Ada County | Mary Lea Killian (National Veterinary Services Laboratories - USDA / Diagnostic Virology Laboratory) |
| EPI_ISL_16243915 | A/duck/Idaho/22-015275-001-original/2022 | A / H5N1 | North America / United States / Idaho / Canyon County | Mary Lea Killian (National Veterinary Services Laboratories - USDA / Diagnostic Virology Laboratory) |
| EPI_ISL_16243926 | A/duck/Idaho/22-015275-002-original/2022 | A / H5N1 | North America / United States / Idaho / Canyon County | Mary Lea Killian (National Veterinary Services Laboratories - USDA / Diagnostic Virology Laboratory) |
| EPI_ISL_16243948 | A/Pekin duck/Pennsylvania/22-015337-001-original/2022 | A / H5N1 | North America / United States / Pennsylvania / Berks County | Mary Lea Killian (National Veterinary Services Laboratories - USDA / Diagnostic Virology Laboratory) |
| EPI_ISL_16243958 | A/Pekin duck/Pennsylvania/22-015337-003-original/2022 | A / H5N1 | North America / United States / Pennsylvania / Berks County | Mary Lea Killian (National Veterinary Services Laboratories - USDA / Diagnostic Virology Laboratory) |
| EPI_ISL_16243970 | A/fox/Iowa/22-015357-002-original/2022 | A / H5N1 | North America / United States / Iowa / Cedar County | Mary Lea Killian (National Veterinary Services Laboratories - USDA / Diagnostic Virology Laboratory) |
| EPI_ISL_16243981 | A/chicken/Minnesota/22-015398-001-original/2022 | A / H5N1 | North America / United States / Minnesota / Clay County | Mary Lea Killian (National Veterinary Services Laboratories - USDA / Diagnostic Virology Laboratory) |
| EPI_ISL_16243992 | A/chicken/Minnesota/22-015398-002-original/2022 | A / H5N1 | North America / United States / Minnesota / Clay County | Mary Lea Killian (National Veterinary Services Laboratories - USDA / Diagnostic Virology Laboratory) |
| EPI_ISL_16244002 | A/turkey/Minnesota/22-015400-001-original/2022 | A / H5N1 | North America / United States / Minnesota / Kandiyohi County | Mary Lea Killian (National Veterinary Services Laboratories - USDA / Diagnostic Virology Laboratory) |
| EPI_ISL_16244003 | A/turkey/Minnesota/22-015400-002-original/2022 | A / H5N1 | North America / United States / Minnesota / Kandiyohi County | Mary Lea Killian (National Veterinary Services Laboratories - USDA / Diagnostic Virology Laboratory) |
| EPI_ISL_16244004 | A/chicken/Idaho/22-015407-001-original/2022 | A / H5N1 | North America / United States / Idaho / Canyon County | Mary Lea Killian (National Veterinary Services Laboratories - USDA / Diagnostic Virology Laboratory) |
| EPI_ISL_16244005 | A/chicken/Idaho/22-015408-001-original/2022 | A / H5N1 | North America / United States / Idaho / Ada County | Mary Lea Killian (National Veterinary Services Laboratories - USDA / Diagnostic Virology Laboratory) |
| EPI_ISL_16244006 | A/chicken/Idaho/22-015409-001-original/2022 | A / H5N1 | North America / United States / Idaho / Canyon County | Mary Lea Killian (National Veterinary Services Laboratories - USDA / Diagnostic Virology Laboratory) |
| EPI_ISL_16244007 | A/chicken/Idaho/22-015410-001-original/2022 | A / H5N1 | North America / United States / Idaho / Canyon County | Mary Lea Killian (National Veterinary Services Laboratories - USDA / Diagnostic Virology Laboratory) |
| EPI_ISL_16244008 | A/duck/Idaho/22-015410-002-original/2022 | A / H5N1 | North America / United States / Idaho / Canyon County | Mary Lea Killian (National Veterinary Services Laboratories - USDA / Diagnostic Virology Laboratory) |
| EPI_ISL_16244009 | A/Pekin duck/Pennsylvania/22-015427-002-original/2022 | A / H5N1 | North America / United States / Pennsylvania / Berks County | Mary Lea Killian (National Veterinary Services Laboratories - USDA / Diagnostic Virology Laboratory) |
| EPI_ISL_16244010 | A/chicken/Indiana/22-015386-002-original/2022 | A / H5N1 | North America / United States / Indiana / Allen County | Mary Lea Killian (National Veterinary Services Laboratories - USDA / Diagnostic Virology Laboratory) |
| EPI_ISL_16244011 | A/duck/Colorado/22-015599-001-original/2022 | A / H5N1 | North America / United States / Colorado / Jefferson County | Mary Lea Killian (National Veterinary Services Laboratories - USDA / Diagnostic Virology Laboratory) |
| EPI_ISL_16244012 | A/chicken/Idaho/22-015654-001-original/2022 | A / H5N1 | North America / United States / Idaho / Ada County | Mary Lea Killian (National Veterinary Services Laboratories - USDA / Diagnostic Virology Laboratory) |
| EPI_ISL_16244013 | A/chicken/Idaho/22-015655-001-original/2022 | A / H5N1 | North America / United States / Idaho / Canyon County | Mary Lea Killian (National Veterinary Services Laboratories - USDA / Diagnostic Virology Laboratory) |
| EPI_ISL_16244014 | A/chicken/Idaho/22-015656-001-original/2022 | A / H5N1 | North America / United States / Idaho / Canyon County | Mary Lea Killian (National Veterinary Services Laboratories - USDA / Diagnostic Virology Laboratory) |
| EPI_ISL_16244015 | A/chicken/Idaho/22-015657-001-original/2022 | A / H5N1 | North America / United States / Idaho / Ada County | Mary Lea Killian (National Veterinary Services Laboratories - USDA / Diagnostic Virology Laboratory) |
| EPI_ISL_16244016 | A/chicken/Idaho/22-015657-002-original/2022 | A / H5N1 | North America / United States / Idaho / Ada County | Mary Lea Killian (National Veterinary Services Laboratories - USDA / Diagnostic Virology Laboratory) |
| EPI_ISL_16244017 | A/chicken/South Dakota/22-015737-001-original/2022 | A / H5N1 | North America / United States / South Dakota / Codington County | Mary Lea Killian (National Veterinary Services Laboratories - USDA / Diagnostic Virology Laboratory) |
| EPI_ISL_16244018 | A/chicken/Idaho/22-015744-001-original/2022 | A / H5N1 | North America / United States / Idaho / Ada County | Mary Lea Killian (National Veterinary Services Laboratories - USDA / Diagnostic Virology Laboratory) |
| EPI_ISL_16244019 | A/chicken/Minnesota/22-015759-001-original/2022 | A / H5N1 | North America / United States / Minnesota / Polk County | Mary Lea Killian (National Veterinary Services Laboratories - USDA / Diagnostic Virology Laboratory) |
| EPI_ISL_16244020 | A/chicken/Minnesota/22-015759-002-original/2022 | A / H5N1 | North America / United States / Minnesota / Polk County | Mary Lea Killian (National Veterinary Services Laboratories - USDA / Diagnostic Virology Laboratory) |
| EPI_ISL_16244021 | A/turkey/Minnesota/22-015759-003-original/2022 | A / H5N1 | North America / United States / Minnesota / Polk County | Mary Lea Killian (National Veterinary Services Laboratories - USDA / Diagnostic Virology Laboratory) |
| EPI_ISL_16244022 | A/chicken/Wyoming/22-015810-001-original/2022 | A / H5N1 | North America / United States / Wyoming / Campbell County | Mary Lea Killian (National Veterinary Services Laboratories - USDA / Diagnostic Virology Laboratory) |
| EPI_ISL_16244023 | A/chicken/Wyoming/22-015810-002-original/2022 | A / H5N1 | North America / United States / Wyoming / Campbell County | Mary Lea Killian (National Veterinary Services Laboratories - USDA / Diagnostic Virology Laboratory) |
| EPI_ISL_16244024 | A/turkey/Minnesota/22-015885-001-original/2022 | A / H5N1 | North America / United States / Minnesota / Dakota County | Mary Lea Killian (National Veterinary Services Laboratories - USDA / Diagnostic Virology Laboratory) |
| EPI_ISL_16244025 | A/turkey/Minnesota/22-015885-002-original/2022 | A / H5N1 | North America / United States / Minnesota / Dakota County | Mary Lea Killian (National Veterinary Services Laboratories - USDA / Diagnostic Virology Laboratory) |
| EPI_ISL_16244026 | A/chicken/Wisconsin/22-015940-001-original/2022 | A / H5N1 | North America / United States / Wisconsin / Bayfield County | Mary Lea Killian (National Veterinary Services Laboratories - USDA / Diagnostic Virology Laboratory) |
| EPI_ISL_16244028 | A/Pekin duck/Pennsylvania/22-015943-001-original/2022 | A / H5N1 | North America / United States / Pennsylvania / Berks County | Mary Lea Killian (National Veterinary Services Laboratories - USDA / Diagnostic Virology Laboratory) |
| EPI_ISL_16244029 | A/duck/Washington/22-016021-001-original/2022 | A / H5N1 | North America / United States / Washington / King County | Mary Lea Killian (National Veterinary Services Laboratories - USDA / Diagnostic Virology Laboratory) |
| EPI_ISL_16244030 | A/duck/Washington/22-016021-002-original/2022 | A / H5N1 | North America / United States / Washington / King County | Mary Lea Killian (National Veterinary Services Laboratories - USDA / Diagnostic Virology Laboratory) |
| EPI_ISL_162467 | A/duck/Guangdong/GD01/2014 | A / H5N6 | Asia / China | |
| EPI_ISL_16249274 | A/Peru/PIU-001/2022 | A / H5N1 | South America / Peru / Departamento de Piura | Carlos Patricio Padilla Rojas (Instituto Nacional de Salud Peru / Laboratorio de Biotecnolgia y Biologia Molecular) |
| EPI_ISL_16249681 | A/Peru/LAM-002/2022 | A / H5N1 | South America / Peru / Departamento de Lambayeque | Carlos Patricio Padilla Rojas (Instituto Nacional de Salud Peru / Laboratorio de Biotecnolgia y Biologia Molecular) |
| EPI_ISL_16249730 | A/Peru/LIM-003/2022 | A / H5N1 | South America / Peru / Departamento de Lima | Carlos Patricio Padilla Rojas (Instituto Nacional de Salud Peru / Laboratorio de Biotecnolgia y Biologia Molecular) |
| EPI_ISL_16253341 | A/Mallard Duck/Netherlands/31/2022 | A / H5N1 | Europe / Netherlands / Eendenkooi, Wieringen (Slijkerman) | Sanne Thewessen (Erasmus Medical Center / Viroscience) |
| EPI_ISL_16253342 | A/Eurasian Wigeon/Netherlands/6/2022 | A / H5N2 | Europe / Netherlands / Eendenkooi Stolwijk | Sanne Thewessen (Erasmus Medical Center / Viroscience) |
| EPI_ISL_16271814 | A/goose/Alabama/22-038970-002-original/2022 | A / H5N1 | North America / United States / Alabama / Lawrence County | Mary Lea Killian (National Veterinary Services Laboratories - USDA / Diagnostic Virology Laboratory) |
| EPI_ISL_16271815 | A/chicken/Alabama/22-038970-003-original/2022 | A / H5N1 | North America / United States / Alabama / Lawrence County | Mary Lea Killian (National Veterinary Services Laboratories - USDA / Diagnostic Virology Laboratory) |
| EPI_ISL_16271816 | A/duck/Alabama/22-038970-004-original/2022 | A / H5N1 | North America / United States / Alabama / Lawrence County | Mary Lea Killian (National Veterinary Services Laboratories - USDA / Diagnostic Virology Laboratory) |
| EPI_ISL_16271817 | A/chicken/Missouri/22-037522-001-original/2022 | A / H5N1 | North America / United States / Missouri / Webster County | Mary Lea Killian (National Veterinary Services Laboratories - USDA / Diagnostic Virology Laboratory) |
| EPI_ISL_16271818 | A/turkey/South Dakota/22-037527-001-original/2022 | A / H5N1 | North America / United States / South Dakota / Beadle County | Mary Lea Killian (National Veterinary Services Laboratories - USDA / Diagnostic Virology Laboratory) |
| EPI_ISL_16271819 | A/turkey/South Dakota/22-037357-001-original/2022 | A / H5N1 | North America / United States / South Dakota / Edmunds County | Mary Lea Killian (National Veterinary Services Laboratories - USDA / Diagnostic Virology Laboratory) |
| EPI_ISL_16271820 | A/domestic goose/Kentucky/22-037359-001-original/2022 | A / H5N1 | North America / United States / Kentucky / Jefferson County | Mary Lea Killian (National Veterinary Services Laboratories - USDA / Diagnostic Virology Laboratory) |
| EPI_ISL_16271821 | A/domestic goose/Maine/22-037360-001-original/2022 | A / H5N1 | North America / United States / Maine / Washington County | Mary Lea Killian (National Veterinary Services Laboratories - USDA / Diagnostic Virology Laboratory) |
| EPI_ISL_16271822 | A/chicken/Maine/22-037360-002-original/2022 | A / H5N1 | North America / United States / Maine / Washington County | Mary Lea Killian (National Veterinary Services Laboratories - USDA / Diagnostic Virology Laboratory) |
| EPI_ISL_16271823 | A/chicken/Nebraska/22-038152-001-original/2022 | A / H5N1 | North America / United States / Nebraska / Dixon County | Mary Lea Killian (National Veterinary Services Laboratories - USDA / Diagnostic Virology Laboratory) |
| EPI_ISL_16271824 | A/chicken/Nebraska/22-038149-001-original/2022 | A / H5N1 | North America / United States / Nebraska / Dixon County | Mary Lea Killian (National Veterinary Services Laboratories - USDA / Diagnostic Virology Laboratory) |
| EPI_ISL_16271825 | A/chicken/Iowa/22-035154-002-original/2022 | A / H5N1 | North America / United States / Iowa / Wright County | Mary Lea Killian (National Veterinary Services Laboratories - USDA / Diagnostic Virology Laboratory) |
| EPI_ISL_16271826 | A/turkey/Missouri/22-038012-001-original/2022 | A / H5N1 | North America / United States / Missouri / Webster County | Mary Lea Killian (National Veterinary Services Laboratories - USDA / Diagnostic Virology Laboratory) |
| EPI_ISL_16271827 | A/rhea/Arizona/22-037270-002-original/2022 | A / H5N1 | North America / United States / Arizona / Maricopa County | Mary Lea Killian (National Veterinary Services Laboratories - USDA / Diagnostic Virology Laboratory) |
| EPI_ISL_16271828 | A/domestic duck/North Carolina/22-037942-001-original/2022 | A / H5N1 | North America / United States / North Carolina / Durham County | Mary Lea Killian (National Veterinary Services Laboratories - USDA / Diagnostic Virology Laboratory) |
| EPI_ISL_16271829 | A/chicken/Alaska/22-037940-001-original/2022 | A / H5N1 | North America / United States / Alaska / Matanuska-Susitna Borough | Mary Lea Killian (National Veterinary Services Laboratories - USDA / Diagnostic Virology Laboratory) |
| EPI_ISL_16271830 | A/chicken/South Dakota/22-037931-001-original/2022 | A / H5N1 | North America / United States / South Dakota / Turner County | Mary Lea Killian (National Veterinary Services Laboratories - USDA / Diagnostic Virology Laboratory) |
| EPI_ISL_16271831 | A/turkey/South Dakota/22-037933-001-original/2022 | A / H5N1 | North America / United States / South Dakota / Beadle County | Mary Lea Killian (National Veterinary Services Laboratories - USDA / Diagnostic Virology Laboratory) |

| | | | | |
|---|---|---|---|---|
| EPI_ISL_16271832 | A/domestic goose/Kentucky/22-037359-002-original/2022 | A / H5N1 | North America / United States / Kentucky / Jefferson County | Mary Lea Killian (National Veterinary Services Laboratories - USDA / Diagnostic Virology Laboratory) |
| EPI_ISL_16271833 | A/turkey/South Dakota/22-037934-001-original/2022 | A / H5N1 | North America / United States / South Dakota / Beadle County | Mary Lea Killian (National Veterinary Services Laboratories - USDA / Diagnostic Virology Laboratory) |
| EPI_ISL_16271834 | A/turkey/South Dakota/22-037930-001-original/2022 | A / H5N1 | North America / United States / South Dakota / Spink County | Mary Lea Killian (National Veterinary Services Laboratories - USDA / Diagnostic Virology Laboratory) |
| EPI_ISL_16271835 | A/striped skunk/Idaho/22-037165-002-original/2022 | A / H5N1 | North America / United States / Idaho / Jefferson County | Mary Lea Killian (National Veterinary Services Laboratories - USDA / Diagnostic Virology Laboratory) |
| EPI_ISL_16271836 | A/chicken/Tennessee/22-037736-001-original/2022 | A / H5N1 | North America / United States / Tennessee / Davidson County | Mary Lea Killian (National Veterinary Services Laboratories - USDA / Diagnostic Virology Laboratory) |
| EPI_ISL_16271837 | A/chicken/Florida/22-037745-001-original/2022 | A / H5N1 | North America / United States / Florida / Broward County | Mary Lea Killian (National Veterinary Services Laboratories - USDA / Diagnostic Virology Laboratory) |
| EPI_ISL_16271838 | A/turkey/Minnesota/22-037734-001-original/2022 | A / H5N1 | North America / United States / Minnesota / Todd County | Mary Lea Killian (National Veterinary Services Laboratories - USDA / Diagnostic Virology Laboratory) |
| EPI_ISL_16271839 | A/chicken/South Dakota/22-037701-001-original/2022 | A / H5N1 | North America / United States / South Dakota / Brule County | Mary Lea Killian (National Veterinary Services Laboratories - USDA / Diagnostic Virology Laboratory) |
| EPI_ISL_16271840 | A/chicken/Oregon/22-037712-001-original/2022 | A / H5N1 | North America / United States / Oregon / Columbia County | Mary Lea Killian (National Veterinary Services Laboratories - USDA / Diagnostic Virology Laboratory) |
| EPI_ISL_16271841 | A/turkey/South Dakota/22-037700-001-original/2022 | A / H5N1 | North America / United States / South Dakota / Faulk County | Mary Lea Killian (National Veterinary Services Laboratories - USDA / Diagnostic Virology Laboratory) |
| EPI_ISL_16271842 | A/chicken/Ohio/22-035584-001-original/2022 | A / H5N1 | North America / United States / Ohio / Butler County | Mary Lea Killian (National Veterinary Services Laboratories - USDA / Diagnostic Virology Laboratory) |
| EPI_ISL_16271843 | A/Chinese ringneck pheasant/Wisconsin/22-035394-001-origir | A / H5N1 | North America / United States / Wisconsin / Jefferson County | Mary Lea Killian (National Veterinary Services Laboratories - USDA / Diagnostic Virology Laboratory) |
| EPI_ISL_16271844 | A/Chinese ringneck pheasant/Wisconsin/22-035394-002-origir | A / H5N1 | North America / United States / Wisconsin / Jefferson County | Mary Lea Killian (National Veterinary Services Laboratories - USDA / Diagnostic Virology Laboratory) |
| EPI_ISL_16271845 | A/chicken/New York/22-035475-001-original/2022 | A / H5N1 | North America / United States / New York / Sullivan County | Mary Lea Killian (National Veterinary Services Laboratories - USDA / Diagnostic Virology Laboratory) |
| EPI_ISL_16271846 | A/chicken/New York/22-035476-001-original/2022 | A / H5N1 | North America / United States / New York / Sullivan County | Mary Lea Killian (National Veterinary Services Laboratories - USDA / Diagnostic Virology Laboratory) |
| EPI_ISL_16271847 | A/chicken/Wyoming/22-035578-001-original/2022 | A / H5N1 | North America / United States / Wyoming / Converse County | Mary Lea Killian (National Veterinary Services Laboratories - USDA / Diagnostic Virology Laboratory) |
| EPI_ISL_16271848 | A/black bear/Alaska/22-036396-001-original/2022 | A / H5N1 | North America / United States / Alaska / Juneau City and Borough | Mary Lea Killian (National Veterinary Services Laboratories - USDA / Diagnostic Virology Laboratory) |
| EPI_ISL_16271849 | A/chicken/North Carolina/22-037124-002-original/2022 | A / H5N1 | North America / United States / North Carolina / Union County | Mary Lea Killian (National Veterinary Services Laboratories - USDA / Diagnostic Virology Laboratory) |
| EPI_ISL_16271850 | A/chicken/Florida/22-036965-001-original/2022 | A / H5N1 | North America / United States / Florida / Broward County | Mary Lea Killian (National Veterinary Services Laboratories - USDA / Diagnostic Virology Laboratory) |
| EPI_ISL_16271851 | A/chicken/Florida/22-036767-001-original/2022 | A / H5N1 | North America / United States / Florida / Hillsborough County | Mary Lea Killian (National Veterinary Services Laboratories - USDA / Diagnostic Virology Laboratory) |
| EPI_ISL_16271852 | A/turkey/Florida/22-036768-001-original/2022 | A / H5N1 | North America / United States / Florida / Seminole County | Mary Lea Killian (National Veterinary Services Laboratories - USDA / Diagnostic Virology Laboratory) |
| EPI_ISL_16271853 | A/pheasant/Michigan/22-036108-001-original/2022 | A / H5N1 | North America / United States / Michigan / Lapeer County | Mary Lea Killian (National Veterinary Services Laboratories - USDA / Diagnostic Virology Laboratory) |
| EPI_ISL_16271854 | A/chicken/North Dakota/22-036126-001-original/2022 | A / H5N1 | North America / United States / North Dakota / Mountrail County | Mary Lea Killian (National Veterinary Services Laboratories - USDA / Diagnostic Virology Laboratory) |
| EPI_ISL_16271855 | A/domestic goose/Tennessee/22-036130-001-original/2022 | A / H5N1 | North America / United States / Tennessee / Lincoln County | Mary Lea Killian (National Veterinary Services Laboratories - USDA / Diagnostic Virology Laboratory) |
| EPI_ISL_16271856 | A/Muscovy duck/New York/22-036304-003-original/2022 | A / H5N1 | North America / United States / New York / Queens County | Mary Lea Killian (National Veterinary Services Laboratories - USDA / Diagnostic Virology Laboratory) |
| EPI_ISL_16271857 | A/chicken/New York/22-036304-007-original/2022 | A / H5N1 | North America / United States / New York / Queens County | Mary Lea Killian (National Veterinary Services Laboratories - USDA / Diagnostic Virology Laboratory) |
| EPI_ISL_16271858 | A/chicken/Tennessee/22-036544-002-original/2022 | A / H5N1 | North America / United States / Tennessee / Bledsoe County | Mary Lea Killian (National Veterinary Services Laboratories - USDA / Diagnostic Virology Laboratory) |
| EPI_ISL_16271859 | A/turkey/Minnesota/22-036547-001-original/2022 | A / H5N1 | North America / United States / Minnesota / Otter Tail County | Mary Lea Killian (National Veterinary Services Laboratories - USDA / Diagnostic Virology Laboratory) |
| EPI_ISL_16271860 | A/chicken/Florida/22-036555-001-original/2022 | A / H5N1 | North America / United States / Florida / Duval County | Mary Lea Killian (National Veterinary Services Laboratories - USDA / Diagnostic Virology Laboratory) |
| EPI_ISL_16271861 | A/Pekin duck/California/22-036722-001-original/2022 | A / H5N1 | North America / United States / California / Fresno County | Mary Lea Killian (National Veterinary Services Laboratories - USDA / Diagnostic Virology Laboratory) |
| EPI_ISL_16271862 | A/chicken/Wyoming/22-035579-001-original/2022 | A / H5N1 | North America / United States / Wyoming / Converse County | Mary Lea Killian (National Veterinary Services Laboratories - USDA / Diagnostic Virology Laboratory) |
| EPI_ISL_16271863 | A/chicken/Wyoming/22-035580-001-original/2022 | A / H5N1 | North America / United States / Wyoming / Converse County | Mary Lea Killian (National Veterinary Services Laboratories - USDA / Diagnostic Virology Laboratory) |
| EPI_ISL_16271864 | A/chicken/Wyoming/22-035581-001-original/2022 | A / H5N1 | North America / United States / Wyoming / Converse County | Mary Lea Killian (National Veterinary Services Laboratories - USDA / Diagnostic Virology Laboratory) |
| EPI_ISL_16271865 | A/domestic goose/Kentucky/22-037028-001-original/2022 | A / H5N1 | North America / United States / Kentucky / Jefferson County | Mary Lea Killian (National Veterinary Services Laboratories - USDA / Diagnostic Virology Laboratory) |
| EPI_ISL_16297060 | A/chicken/Oregon/22-030743-001-original/2022 | A / H5N1 | North America / United States / Oregon / Douglas County | Mary Lea Killian (National Veterinary Services Laboratories - USDA / Diagnostic Virology Laboratory) |
| EPI_ISL_16297061 | A/chicken/North Dakota/22-030784-001-original/2022 | A / H5N1 | North America / United States / North Dakota / Nelson County | Mary Lea Killian (National Veterinary Services Laboratories - USDA / Diagnostic Virology Laboratory) |
| EPI_ISL_16297062 | A/pheasant/South Dakota/22-034797-001-original/2022 | A / H5N1 | North America / United States / South Dakota / Gregory County | Mary Lea Killian (National Veterinary Services Laboratories - USDA / Diagnostic Virology Laboratory) |
| EPI_ISL_16297063 | A/chicken/Montana/22-034813-003-original/2022 | A / H5N1 | North America / United States / Montana / Granite County | Mary Lea Killian (National Veterinary Services Laboratories - USDA / Diagnostic Virology Laboratory) |
| EPI_ISL_16297064 | A/chicken/Oregon/22-034833-001-original/2022 | A / H5N1 | North America / United States / Oregon / Deschutes County | Mary Lea Killian (National Veterinary Services Laboratories - USDA / Diagnostic Virology Laboratory) |
| EPI_ISL_16297065 | A/turkey/Minnesota/22-034849-001-original/2022 | A / H5N1 | North America / United States / Minnesota / Stearns County | Mary Lea Killian (National Veterinary Services Laboratories - USDA / Diagnostic Virology Laboratory) |
| EPI_ISL_16297066 | A/skunk/Minnesota/22-034925-001-original/2022 | A / H5N1 | North America / United States / Minnesota / Itasca County | Mary Lea Killian (National Veterinary Services Laboratories - USDA / Diagnostic Virology Laboratory) |
| EPI_ISL_16297067 | A/Muscovy duck/California/22-034335-001-original/2022 | A / H5N1 | North America / United States / California / Fresno County | Mary Lea Killian (National Veterinary Services Laboratories - USDA / Diagnostic Virology Laboratory) |
| EPI_ISL_16297068 | A/chicken/Pennsylvania/22-034995-001-original/2022 | A / H5N1 | North America / United States / Pennsylvania / Allegheny County | Mary Lea Killian (National Veterinary Services Laboratories - USDA / Diagnostic Virology Laboratory) |
| EPI_ISL_16297069 | A/chicken/New Jersey/22-035003-009-original/2022 | A / H5N1 | North America / United States / New Jersey / Bergen County | Mary Lea Killian (National Veterinary Services Laboratories - USDA / Diagnostic Virology Laboratory) |
| EPI_ISL_16297070 | A/chicken/Arizona/22-035006-002-original/2022 | A / H5N1 | North America / United States / Arizona / Yavapai County | Mary Lea Killian (National Veterinary Services Laboratories - USDA / Diagnostic Virology Laboratory) |
| EPI_ISL_16297071 | A/chicken/North Dakota/22-035070-001-original/2022 | A / H5N1 | North America / United States / North Dakota / Bottineau County | Mary Lea Killian (National Veterinary Services Laboratories - USDA / Diagnostic Virology Laboratory) |
| EPI_ISL_16297072 | A/turkey/North Dakota/22-035070-002-original/2022 | A / H5N1 | North America / United States / North Dakota / Bottineau County | Mary Lea Killian (National Veterinary Services Laboratories - USDA / Diagnostic Virology Laboratory) |
| EPI_ISL_16297073 | A/chicken/California/22-035144-001-original/2022 | A / H5N1 | North America / United States / California / San Diego County | Mary Lea Killian (National Veterinary Services Laboratories - USDA / Diagnostic Virology Laboratory) |
| EPI_ISL_16297074 | A/domestic goose/California/22-035144-003-original/2022 | A / H5N1 | North America / United States / California / San Diego County | Mary Lea Killian (National Veterinary Services Laboratories - USDA / Diagnostic Virology Laboratory) |
| EPI_ISL_16297075 | A/chicken/South Carolina/22-035153-001-original/2022 | A / H5N1 | North America / United States / South Carolina / Beaufort County | Mary Lea Killian (National Veterinary Services Laboratories - USDA / Diagnostic Virology Laboratory) |
| EPI_ISL_16297076 | A/guinea fowl/South Carolina/22-035153-002-original/2022 | A / H5N1 | North America / United States / South Carolina / Beaufort County | Mary Lea Killian (National Veterinary Services Laboratories - USDA / Diagnostic Virology Laboratory) |
| EPI_ISL_16297077 | A/turkey/Pennsylvania/22-035178-001-original/2022 | A / H5N1 | North America / United States / Pennsylvania / Lehigh County | Mary Lea Killian (National Veterinary Services Laboratories - USDA / Diagnostic Virology Laboratory) |
| EPI_ISL_16297078 | A/chicken/Oregon/22-035197-001-original/2022 | A / H5N1 | North America / United States / Oregon / Yamhill County | Mary Lea Killian (National Veterinary Services Laboratories - USDA / Diagnostic Virology Laboratory) |
| EPI_ISL_16297079 | A/chicken/Massachussetts/22-035225-001-original/2022 | A / H5N1 | North America / United States / Massachussetts | Mary Lea Killian (National Veterinary Services Laboratories - USDA / Diagnostic Virology Laboratory) |
| EPI_ISL_16297080 | A/chicken/Massachussetts/22-035234-001-original/2022 | A / H5N1 | North America / United States / Massachussetts | Mary Lea Killian (National Veterinary Services Laboratories - USDA / Diagnostic Virology Laboratory) |
| EPI_ISL_16297081 | A/chicken/Arizona/22-035241-003-original/2022 | A / H5N1 | North America / United States / Arizona / Yavapai County | Mary Lea Killian (National Veterinary Services Laboratories - USDA / Diagnostic Virology Laboratory) |
| EPI_ISL_16297082 | A/chicken/Iowa/22-035700-001-original/2022 | A / H5N1 | North America / United States / Iowa / Louisa County | Mary Lea Killian (National Veterinary Services Laboratories - USDA / Diagnostic Virology Laboratory) |
| EPI_ISL_16297083 | A/domestic duck/Iowa/22-035700-002-original/2022 | A / H5N1 | North America / United States / Iowa / Louisa County | Mary Lea Killian (National Veterinary Services Laboratories - USDA / Diagnostic Virology Laboratory) |
| EPI_ISL_16297084 | A/turkey/Pennsylvania/22-035743-001-original/2022 | A / H5N1 | North America / United States / Pennsylvania / Lehigh County | Mary Lea Killian (National Veterinary Services Laboratories - USDA / Diagnostic Virology Laboratory) |
| EPI_ISL_16297085 | A/chicken/Wisconsin/22-034812-002-original/2022 | A / H5N1 | North America / United States / Wisconsin / Waukesha County | Mary Lea Killian (National Veterinary Services Laboratories - USDA / Diagnostic Virology Laboratory) |
| EPI_ISL_16297086 | A/wild turkey/New York/22-034864-002-original/202 | A / H5N1 | North America / United States / New York / Dutchess County | Mary Lea Killian (National Veterinary Services Laboratories - USDA / Diagnostic Virology Laboratory) |
| EPI_ISL_16297117 | A/mallard/Texas/22-035744-001-original/2022 | A / H5N1 | North America / United States / Texas / Tarrant County | Mary Lea Killian (National Veterinary Services Laboratories - USDA / Diagnostic Virology Laboratory) |
| EPI_ISL_16297118 | A/chicken/Washington/22-035749-001-original/2022 | A / H5N1 | North America / United States / Washington / Snohomish County | Mary Lea Killian (National Veterinary Services Laboratories - USDA / Diagnostic Virology Laboratory) |
| EPI_ISL_16297119 | A/turkey/Pennsylvania/22-036105-007-original/2022 | A / H5N1 | North America / United States / Pennsylvania / Lehigh County | Mary Lea Killian (National Veterinary Services Laboratories - USDA / Diagnostic Virology Laboratory) |
| EPI_ISL_16297120 | A/turkey/Pennsylvania/22-036115-004-original/2022 | A / H5N1 | North America / United States / Pennsylvania / Lehigh County | Mary Lea Killian (National Veterinary Services Laboratories - USDA / Diagnostic Virology Laboratory) |
| EPI_ISL_16297121 | A/chicken/Nebraska/22-037884-001-original/2022 | A / H5N1 | North America / United States / Nebraska / Dixon County | Mary Lea Killian (National Veterinary Services Laboratories - USDA / Diagnostic Virology Laboratory) |
| EPI_ISL_16297122 | A/skunk/Wisconsin/22-037029-001-original/2022 | A / H5N1 | North America / United States / Wisconsin / Douglas County | Mary Lea Killian (National Veterinary Services Laboratories - USDA / Diagnostic Virology Laboratory) |
| EPI_ISL_16297123 | A/chicken/Iowa/22-035569-001-original/2022 | A / H5N1 | North America / United States / Iowa / Wright County | Mary Lea Killian (National Veterinary Services Laboratories - USDA / Diagnostic Virology Laboratory) |
| EPI_ISL_16297124 | A/chicken/Iowa/22-035567-003-original/2022 | A / H5N1 | North America / United States / Iowa / Wright County | Mary Lea Killian (National Veterinary Services Laboratories - USDA / Diagnostic Virology Laboratory) |
| EPI_ISL_16297125 | A/turkey/Pennsylvania/22-035424-001-original/2022 | A / H5N1 | North America / United States / Pennsylvania / Lehigh County | Mary Lea Killian (National Veterinary Services Laboratories - USDA / Diagnostic Virology Laboratory) |
| EPI_ISL_16297126 | A/chicken/Iowa/22-034682-001-original/2022 | A / H5N1 | North America / United States / Iowa / Wright County | Mary Lea Killian (National Veterinary Services Laboratories - USDA / Diagnostic Virology Laboratory) |
| EPI_ISL_16297127 | A/turkey/Minnesota/22-034666-001-original/2022 | A / H5N1 | North America / United States / Minnesota / Swift County | Mary Lea Killian (National Veterinary Services Laboratories - USDA / Diagnostic Virology Laboratory) |

| | | | | |
|---|---|---|---|---|
| EPI_ISL_16297128 | A/chicken/Colorado/22-034082-001-original/2022 | A / H5N1 | North America / United States / Colorado / La Plata County | Mary Lea Killian (National Veterinary Services Laboratories - USDA / Diagnostic Virology Laboratory) |
| EPI_ISL_16297129 | A/turkey/Minnesota/22-034260-001-original/2022 | A / H5N1 | North America / United States / Minnesota / Le Sueur County | Mary Lea Killian (National Veterinary Services Laboratories - USDA / Diagnostic Virology Laboratory) |
| EPI_ISL_16297130 | A/chicken/Florida/22-034076-001-original/2022 | A / H5N1 | North America / United States / Florida / Pasco County | Mary Lea Killian (National Veterinary Services Laboratories - USDA / Diagnostic Virology Laboratory) |
| EPI_ISL_16297131 | A/chicken/Florida/22-033918-001-original/2022 | A / H5N1 | North America / United States / Florida / Indian River County | Mary Lea Killian (National Veterinary Services Laboratories - USDA / Diagnostic Virology Laboratory) |
| EPI_ISL_16297132 | A/domestic duck/Florida/22-033918-002-original/2022 | A / H5N1 | North America / United States / Florida / Indian River County | Mary Lea Killian (National Veterinary Services Laboratories - USDA / Diagnostic Virology Laboratory) |
| EPI_ISL_16297133 | A/chicken/Florida/22-033923-001-original/2022 | A / H5N1 | North America / United States / Florida / Lake County | Mary Lea Killian (National Veterinary Services Laboratories - USDA / Diagnostic Virology Laboratory) |
| EPI_ISL_16297134 | A/chicken/Florida/22-033924-001-original/2022 | A / H5N1 | North America / United States / Florida / Indian River County | Mary Lea Killian (National Veterinary Services Laboratories - USDA / Diagnostic Virology Laboratory) |
| EPI_ISL_16297135 | A/turkey/Minnesota/22-033929-001-original/2022 | A / H5N1 | North America / United States / Minnesota / Swift County | Mary Lea Killian (National Veterinary Services Laboratories - USDA / Diagnostic Virology Laboratory) |
| EPI_ISL_16297136 | A/turkey/Utah/22-033909-001-original/2022 | A / H5N1 | North America / United States / Utah / Sanpete County | Mary Lea Killian (National Veterinary Services Laboratories - USDA / Diagnostic Virology Laboratory) |
| EPI_ISL_16297137 | A/chicken/South Dakota/22-033852-001-original/2022 | A / H5N1 | North America / United States / South Dakota / Grant County | Mary Lea Killian (National Veterinary Services Laboratories - USDA / Diagnostic Virology Laboratory) |
| EPI_ISL_16297138 | A/African goose/New Jersey/22-033679-001-original/2022 | A / H5N1 | North America / United States / New Jersey / Ocean County | Mary Lea Killian (National Veterinary Services Laboratories - USDA / Diagnostic Virology Laboratory) |
| EPI_ISL_16297139 | A/chicken/Virginia/22-033680-002-original/2022 | A / H5N1 | North America / United States / Virginia / Southampton County | Mary Lea Killian (National Veterinary Services Laboratories - USDA / Diagnostic Virology Laboratory) |
| EPI_ISL_16297140 | A/chicken/Oklahoma/22-033558-001-original/2022 | A / H5N1 | North America / United States / Oklahoma / Cleveland County | Mary Lea Killian (National Veterinary Services Laboratories - USDA / Diagnostic Virology Laboratory) |
| EPI_ISL_16297141 | A/peafowl/Florida/22-033621-001-original/2022 | A / H5N1 | North America / United States / Florida / Indian River County | Mary Lea Killian (National Veterinary Services Laboratories - USDA / Diagnostic Virology Laboratory) |
| EPI_ISL_16297142 | A/chicken/North Carolina/22-033524-001-original/2022 | A / H5N1 | North America / United States / North Carolina / Wake County | Mary Lea Killian (National Veterinary Services Laboratories - USDA / Diagnostic Virology Laboratory) |
| EPI_ISL_16297326 | A/chicken/North Carolina/22-033522-001-original/2022 | A / H5N1 | North America / United States / North Carolina / Wake County | Mary Lea Killian (National Veterinary Services Laboratories - USDA / Diagnostic Virology Laboratory) |
| EPI_ISL_16297327 | A/chicken/Florida/22-035222-003-original/2022 | A / H5N1 | North America / United States / Florida / Broward County | Mary Lea Killian (National Veterinary Services Laboratories - USDA / Diagnostic Virology Laboratory) |
| EPI_ISL_16297328 | A/chicken/Colorado/22-033394-001-original/2022 | A / H5N1 | North America / United States / Colorado / Jefferson County | Mary Lea Killian (National Veterinary Services Laboratories - USDA / Diagnostic Virology Laboratory) |
| EPI_ISL_16297329 | A/chicken/Idaho/22-033409-001-original/2022 | A / H5N1 | North America / United States / Idaho / Valley County | Mary Lea Killian (National Veterinary Services Laboratories - USDA / Diagnostic Virology Laboratory) |
| EPI_ISL_16297330 | A/chicken/Oklahoma/22-033387-001-original/2022 | A / H5N1 | North America / United States / Oklahoma / Creek County | Mary Lea Killian (National Veterinary Services Laboratories - USDA / Diagnostic Virology Laboratory) |
| EPI_ISL_16297331 | A/chicken/Florida/22-033389-001-original/2022 | A / H5N1 | North America / United States / Florida / Okeechobee County | Mary Lea Killian (National Veterinary Services Laboratories - USDA / Diagnostic Virology Laboratory) |
| EPI_ISL_16297332 | A/goose/Louisiana/22-033391-001-original/2022 | A / H5N1 | North America / United States / Louisiana / Ouachita Parish | Mary Lea Killian (National Veterinary Services Laboratories - USDA / Diagnostic Virology Laboratory) |
| EPI_ISL_16297333 | A/chicken/Florida/22-033392-001-original/2022 | A / H5N1 | North America / United States / Florida / Indian River County | Mary Lea Killian (National Veterinary Services Laboratories - USDA / Diagnostic Virology Laboratory) |
| EPI_ISL_16297334 | A/domestic duck/South Dakota/22-033350-004-original/2022 | A / H5N1 | North America / United States / South Dakota / Spink County | Mary Lea Killian (National Veterinary Services Laboratories - USDA / Diagnostic Virology Laboratory) |
| EPI_ISL_16297335 | A/turkey/South Dakota/22-033350-005-original/2022 | A / H5N1 | North America / United States / South Dakota / Spink County | Mary Lea Killian (National Veterinary Services Laboratories - USDA / Diagnostic Virology Laboratory) |
| EPI_ISL_16297336 | A/chicken/Iowa/22-033362-001-original/2022 | A / H5N1 | North America / United States / Iowa / Dallas County | Mary Lea Killian (National Veterinary Services Laboratories - USDA / Diagnostic Virology Laboratory) |
| EPI_ISL_16297337 | A/chicken/South Dakota/22-033350-001-original/2022 | A / H5N1 | North America / United States / South Dakota / Spink County | Mary Lea Killian (National Veterinary Services Laboratories - USDA / Diagnostic Virology Laboratory) |
| EPI_ISL_16297338 | A/chicken/Virginia/22-033345-001-original/2022 | A / H5N1 | North America / United States / Virginia / Virginia County | Mary Lea Killian (National Veterinary Services Laboratories - USDA / Diagnostic Virology Laboratory) |
| EPI_ISL_16297339 | A/chicken/Rhode Island/22-033346-001-original/2022 | A / H5N1 | North America / United States / Rhode Island / Newport County | Mary Lea Killian (National Veterinary Services Laboratories - USDA / Diagnostic Virology Laboratory) |
| EPI_ISL_16297340 | A/chicken/Montana/22-033348-001-original/2022 | A / H5N1 | North America / United States / Montana / Hill County | Mary Lea Killian (National Veterinary Services Laboratories - USDA / Diagnostic Virology Laboratory) |
| EPI_ISL_16297341 | A/chicken/guinea fowl/Kansas/22-033226-001-original/2022 | A / H5N1 | North America / United States / Kansas / Shawnee County | Mary Lea Killian (National Veterinary Services Laboratories - USDA / Diagnostic Virology Laboratory) |
| EPI_ISL_16297342 | A/chicken/Texas/22-033221-001-original/2022 | A / H5N1 | North America / United States / Texas / Rockwall County | Mary Lea Killian (National Veterinary Services Laboratories - USDA / Diagnostic Virology Laboratory) |
| EPI_ISL_16297343 | A/domestic duck/Florida/22-032962-001-original/2022 | A / H5N1 | North America / United States / Florida / Saint Lucie County | Mary Lea Killian (National Veterinary Services Laboratories - USDA / Diagnostic Virology Laboratory) |
| EPI_ISL_16297344 | A/chicken/Florida/22-032963-001-original/2022 | A / H5N1 | North America / United States / Florida / Osceola County | Mary Lea Killian (National Veterinary Services Laboratories - USDA / Diagnostic Virology Laboratory) |
| EPI_ISL_16297345 | A/chicken/Florida/22-032965-001-original/2022 | A / H5N1 | North America / United States / Florida / Indian River County | Mary Lea Killian (National Veterinary Services Laboratories - USDA / Diagnostic Virology Laboratory) |
| EPI_ISL_16297346 | A/domestic goose/South Dakota/22-033171-001-original/2022 | A / H5N1 | North America / United States / South Dakota / Roberts County | Mary Lea Killian (National Veterinary Services Laboratories - USDA / Diagnostic Virology Laboratory) |
| EPI_ISL_16297347 | A/chicken/North Dakota/22-032950-001-original/2022 | A / H5N1 | North America / United States / North Dakota / Traill County | Mary Lea Killian (National Veterinary Services Laboratories - USDA / Diagnostic Virology Laboratory) |
| EPI_ISL_16297348 | A/guinea fowl/North Dakota/22-032950-002-original/2022 | A / H5N1 | North America / United States / North Dakota / Traill County | Mary Lea Killian (National Veterinary Services Laboratories - USDA / Diagnostic Virology Laboratory) |
| EPI_ISL_16297349 | A/turkey/North Dakota/22-032950-003-original/2022 | A / H5N1 | North America / United States / North Dakota / Traill County | Mary Lea Killian (National Veterinary Services Laboratories - USDA / Diagnostic Virology Laboratory) |
| EPI_ISL_16297350 | A/turkey/Utah/22-032951-001-original/2022 | A / H5N1 | North America / United States / Utah / Sanpete County | Mary Lea Killian (National Veterinary Services Laboratories - USDA / Diagnostic Virology Laboratory) |
| EPI_ISL_16297351 | A/pheasant/Nebraska/22-032910-001-original/2022 | A / H5N1 | North America / United States / Nebraska / York County | Mary Lea Killian (National Veterinary Services Laboratories - USDA / Diagnostic Virology Laboratory) |
| EPI_ISL_16297352 | A/turkey/Pennsylvania/22-032809-001-original/2022 | A / H5N1 | North America / United States / Pennsylvania / Adams County | Mary Lea Killian (National Veterinary Services Laboratories - USDA / Diagnostic Virology Laboratory) |
| EPI_ISL_16297353 | A/chicken/Nevada/22-032799-001-original/2022 | A / H5N1 | North America / United States / Nevada / Nye County | Mary Lea Killian (National Veterinary Services Laboratories - USDA / Diagnostic Virology Laboratory) |
| EPI_ISL_16297354 | A/turkey/Alaska/22-032677-004-original/2022 | A / H5N1 | North America / United States / Alaska / Matanuska-Susitna Borough | Mary Lea Killian (National Veterinary Services Laboratories - USDA / Diagnostic Virology Laboratory) |
| EPI_ISL_16297355 | A/chicken/Colorado/22-032675-001-original/2022 | A / H5N1 | North America / United States / Colorado / La Plata County | Mary Lea Killian (National Veterinary Services Laboratories - USDA / Diagnostic Virology Laboratory) |
| EPI_ISL_16297356 | A/turkey/Utah/22-032657-001-original/2022 | A / H5N1 | North America / United States / Utah / Sanpete County | Mary Lea Killian (National Veterinary Services Laboratories - USDA / Diagnostic Virology Laboratory) |
| EPI_ISL_16297357 | A/chicken/Florida/22-032661-001-original/2022 | A / H5N1 | North America / United States / Florida / Boward County | Mary Lea Killian (National Veterinary Services Laboratories - USDA / Diagnostic Virology Laboratory) |
| EPI_ISL_16297358 | A/chicken/Colorado/22-032670-001-original/2022 | A / H5N1 | North America / United States / Colorado / Larimer County | Mary Lea Killian (National Veterinary Services Laboratories - USDA / Diagnostic Virology Laboratory) |
| EPI_ISL_16297359 | A/guinea fowl/Oklahoma/22-032674-001-original/2022 | A / H5N1 | North America / United States / Oklahoma / Tulsa County | Mary Lea Killian (National Veterinary Services Laboratories - USDA / Diagnostic Virology Laboratory) |
| EPI_ISL_16297360 | A/chicken/Arkansas/22-032566-001-original/2022 | A / H5N1 | North America / United States / Arkansas / Pope County | Mary Lea Killian (National Veterinary Services Laboratories - USDA / Diagnostic Virology Laboratory) |
| EPI_ISL_16297361 | A/goose/Oklahoma/22-032413-001-original/2022 | A / H5N1 | North America / United States / Oklahoma / Payne County | Mary Lea Killian (National Veterinary Services Laboratories - USDA / Diagnostic Virology Laboratory) |
| EPI_ISL_16297362 | A/swan/Oklahoma/22-032414-001-original/2022 | A / H5N1 | North America / United States / Oklahoma / Oklahoma County | Mary Lea Killian (National Veterinary Services Laboratories - USDA / Diagnostic Virology Laboratory) |
| EPI_ISL_16297363 | A/domestic duck/New Jersey/22-032412-001-original/2022 | A / H5N1 | North America / United States / New Jersey / Warren County | Mary Lea Killian (National Veterinary Services Laboratories - USDA / Diagnostic Virology Laboratory) |
| EPI_ISL_16297364 | A/turkey/Utah/22-032405-001-original/2022 | A / H5N1 | North America / United States / Utah / Sanpete County | Mary Lea Killian (National Veterinary Services Laboratories - USDA / Diagnostic Virology Laboratory) |
| EPI_ISL_16297365 | A/chicken/Colorado/22-032233-001-original/2022 | A / H5N1 | North America / United States / Colorado / Weld County | Mary Lea Killian (National Veterinary Services Laboratories - USDA / Diagnostic Virology Laboratory) |
| EPI_ISL_16297366 | A/rhea/Pennsylvania/22-032402-001-original/2022 | A / H5N1 | North America / United States / Pennsylvania / Dauphin County | Mary Lea Killian (National Veterinary Services Laboratories - USDA / Diagnostic Virology Laboratory) |
| EPI_ISL_16297367 | A/cape teal/Texas/22-032200-001-original/2022 | A / H5N1 | North America / United States / Texas / El Paso County | Mary Lea Killian (National Veterinary Services Laboratories - USDA / Diagnostic Virology Laboratory) |
| EPI_ISL_16297368 | A/chicken/Wyoming/22-032216-001-original/2022 | A / H5N1 | North America / United States / Wyoming / Fremont County | Mary Lea Killian (National Veterinary Services Laboratories - USDA / Diagnostic Virology Laboratory) |
| EPI_ISL_16297369 | A/chicken/South Dakota/22-032160-001-original/2022 | A / H5N1 | North America / United States / South Dakota / Hamlin County | Mary Lea Killian (National Veterinary Services Laboratories - USDA / Diagnostic Virology Laboratory) |
| EPI_ISL_16297370 | A/turkey/California/22-032159-001-original/2022 | A / H5N1 | North America / United States / California / Stanislaus County | Mary Lea Killian (National Veterinary Services Laboratories - USDA / Diagnostic Virology Laboratory) |
| EPI_ISL_16297371 | A/chicken/Kentucky/22-031959-001-original/2022 | A / H5N1 | North America / United States / Kentucky / Logan County | Mary Lea Killian (National Veterinary Services Laboratories - USDA / Diagnostic Virology Laboratory) |
| EPI_ISL_16297372 | A/Egyptian goose/Florida/22-031886-001-original/2022 | A / H5N1 | North America / United States / Florida / Martin County | Mary Lea Killian (National Veterinary Services Laboratories - USDA / Diagnostic Virology Laboratory) |
| EPI_ISL_16297373 | A/chicken/Florida/22-031886-002-original/2022 | A / H5N1 | North America / United States / Florida / Martin County | Mary Lea Killian (National Veterinary Services Laboratories - USDA / Diagnostic Virology Laboratory) |
| EPI_ISL_16297374 | A/swan/Oklahoma/22-031912-001-original/2022 | A / H5N1 | North America / United States / Oklahoma / Oklahoma County | Mary Lea Killian (National Veterinary Services Laboratories - USDA / Diagnostic Virology Laboratory) |
| EPI_ISL_16297375 | A/turkey/Michigan/22-031751-001-original/2022 | A / H5N1 | North America / United States / Michigan / Lapeer County | Mary Lea Killian (National Veterinary Services Laboratories - USDA / Diagnostic Virology Laboratory) |
| EPI_ISL_16297376 | A/chicken/Florida/22-031859-001-original/2022 | A / H5N1 | North America / United States / Florida / Broward County | Mary Lea Killian (National Veterinary Services Laboratories - USDA / Diagnostic Virology Laboratory) |
| EPI_ISL_16297377 | A/turkey/Utah/22-031727-001-original/2022 | A / H5N1 | North America / United States / Utah / Sanpete County | Mary Lea Killian (National Veterinary Services Laboratories - USDA / Diagnostic Virology Laboratory) |
| EPI_ISL_16297378 | A/chicken/Arkansas/22-031676-001-original/2022 | A / H5N1 | North America / United States / Arkansas / Madison County | Mary Lea Killian (National Veterinary Services Laboratories - USDA / Diagnostic Virology Laboratory) |
| EPI_ISL_16297379 | A/chicken/Arkansas/22-031667-003-original/2022 | A / H5N1 | North America / United States / Arkansas / Madison County | Mary Lea Killian (National Veterinary Services Laboratories - USDA / Diagnostic Virology Laboratory) |
| EPI_ISL_16297380 | A/turkey/Utah/22-033401-002-original/2022 | A / H5N1 | North America / United States / Utah / Sanpete County | Mary Lea Killian (National Veterinary Services Laboratories - USDA / Diagnostic Virology Laboratory) |
| EPI_ISL_16297381 | A/chicken/Michigan/22-032218-001-original/2022 | A / H5N1 | North America / United States / Michigan / Genesee County | Mary Lea Killian (National Veterinary Services Laboratories - USDA / Diagnostic Virology Laboratory) |
| EPI_ISL_16297382 | A/turkey/Utah/22-031721-004-original/2022 | A / H5N1 | North America / United States / Utah / Sanpete County | Mary Lea Killian (National Veterinary Services Laboratories - USDA / Diagnostic Virology Laboratory) |

| | | | | |
|---|---|---|---|---|
| EPI_ISL_163493 | A/Sichuan/26221/2014 | A / H5N6 | Asia / China / Sichuan Province | Lei Yang (WHO Chinese National Influenza Center / Virology Institute, Chinese CDC) |
| EPI_ISL_16388458 | A/whooper swan/Akita/0511B002/2022 | A / H5N1 | Asia / Japan / Akita / Odate city | Norikazu Isoda (Graduate School of Veterinary Medicine, Hokkaido University / Laboratory of Microbiology) |
| EPI_ISL_16395206 | A/cat/France/22P026544/2022 | A / H5N1 | Europe / France / Region Pays de la Loire | Francois-Xavier Briand (ANSES Agence Nationale De Securite Sanitaire De L'alimentation / Laboratoire de Ploufragan-Plouzané) |
| EPI_ISL_16466440 | A/Changsha/1/2022 | A / H5N6 | Asia / China / Hunan Province / Changsha city | Zheng Huang (Changsha Disease Prevention and Control Center) |
| EPI_ISL_16524187 | A/Common gull/Netherlands/1/2023 | A / H5N1 | Europe / Netherlands / Hippolytushoef, Mulders Eendenkooi | Sanne Thewessen (Erasmus Medical Center / Viroscience) |
| EPI_ISL_16524203 | A/Black-headed gull/Netherlands/1/2023 | A / H5N1 | Europe / Netherlands / Groningen, Pioenpark | Sanne Thewessen (Erasmus Medical Center / Viroscience) |
| EPI_ISL_16524213 | A/European herring gull/Netherlands/1/2023 | A / H5N1 | Europe / Netherlands / Den Oever, Waddenhaven | Sanne Thewessen (Erasmus Medical Center / Viroscience) |
| EPI_ISL_16524230 | A/Black-headed gull/Netherlands/2/2023 | A / H5N1 | Europe / Netherlands / Den Oever, Waddenhaven | Sanne Thewessen (Erasmus Medical Center / Viroscience) |
| EPI_ISL_16524243 | A/European herring gull/Netherlands/2/2023 | A / H5N1 | Europe / Netherlands / Den Oever, Waddenhaven | Sanne Thewessen (Erasmus Medical Center / Viroscience) |
| EPI_ISL_16524245 | A/European herring gull/Netherlands/3/2023 | A / H5N1 | Europe / Netherlands / Den Oever, Waddenhaven | Sanne Thewessen (Erasmus Medical Center / Viroscience) |
| EPI_ISL_16524246 | A/Caspian gull/Netherlands/1/2023 | A / H5N1 | Europe / Netherlands / Den Oever, Waddenhaven | Sanne Thewessen (Erasmus Medical Center / Viroscience) |
| EPI_ISL_16524247 | A/Peregrine falcon/Netherlands/1/2023 | A / H5N1 | Europe / Netherlands / Wieingerwerf, Sluitgatweg | Sanne Thewessen (Erasmus Medical Center / Viroscience) |
| EPI_ISL_16524248 | A/European herring gull/Netherlands/4/2023 | A / H5N1 | Europe / Netherlands / Den Oever, Waddenhaven | Sanne Thewessen (Erasmus Medical Center / Viroscience) |
| EPI_ISL_16547552 | A/Mustela_furo/Belgium/14312-0001/2022 | A / H5N1 | Europe / Belgium / Provincie Limburg / Riemst | Steven Van Borm (Sciensano, Department of Animal Infectious Diseases / Animal Infectious Diseases) |
| EPI_ISL_16547666 | A/Mustela_furo/Belgium/14312-0002/2022 | A / H5N1 | Europe / Belgium / Provincie Limburg / Riemst | Steven Van Borm (Sciensano, Department of Animal Infectious Diseases / Animal Infectious Diseases) |
| EPI_ISL_16547851 | A/Gallus_gallus/Belgium/14313-001/2022 | A / H5N1 | Europe / Belgium / Provincie Limburg / Riemst | Steven Van Borm (Sciensano, Department of Animal Infectious Diseases / Animal Infectious Diseases) |
| EPI_ISL_16548347 | A/Pelecanus_conspicillatus/Belgium/12905-0001/2022 | A / H5N1 | Europe / Belgium / Province de Hainaut / Cambron-Casteau | Steven Van Borm (Sciensano, Department of Animal Infectious Diseases / Animal Infectious Diseases) |
| EPI_ISL_16548356 | A/Phasianus_colchicus/Belgium/13027-0001/2022 | A / H5N1 | Europe / Belgium / Province de Hainaut / Soignies | Steven Van Borm (Sciensano, Department of Animal Infectious Diseases / Animal Infectious Diseases) |
| EPI_ISL_16548429 | A/Gallus_gallus/Belgium/13078-0001/2022 | A / H5N1 | Europe / Belgium / Provincie Antwerpen / Turnhout | Steven Van Borm (Sciensano, Department of Animal Infectious Diseases / Animal Infectious Diseases) |
| EPI_ISL_16548600 | A/Gallus_gallus/Belgium/12485-0001/2022 | A / H5N1 | Europe / Belgium / Provincie West-Vlaanderen / Eke | Steven Van Borm (Sciensano, Department of Animal Infectious Diseases / Animal Infectious Diseases) |
| EPI_ISL_16548638 | A/Phasianus_colchicus/Belgium/11704-0003/2022 | A / H5N1 | Europe / Belgium / Province de Liege / Clavier | Steven Van Borm (Sciensano, Department of Animal Infectious Diseases / Animal Infectious Diseases) |
| EPI_ISL_16549157 | A/Gallus_gallus/Belgium/13357-0001/2022 | A / H5N1 | Europe / Belgium / Provincie Vlaams-Brabant / Lennik | Steven Van Borm (Sciensano, Department of Animal Infectious Diseases / Animal Infectious Diseases) |
| EPI_ISL_16549245 | A/Otus_scops/Belgium/13463-0002/2022 | A / H5N1 | Europe / Belgium / Province de Liege / Lontzen | Steven Van Borm (Sciensano, Department of Animal Infectious Diseases / Animal Infectious Diseases) |
| EPI_ISL_16549349 | A/Phasianus_colchicus/Belgium/10993-0003/2022 | A / H5N1 | Europe / Belgium / Province de Liege / Sainte-Fontaine | Steven Van Borm (Sciensano, Department of Animal Infectious Diseases / Animal Infectious Diseases) |
| EPI_ISL_16549379 | A/Phasianus_colchicus/Belgium/12352-0002/2022 | A / H5N1 | Europe / Belgium / Province de Liege / Clavier | Steven Van Borm (Sciensano, Department of Animal Infectious Diseases / Animal Infectious Diseases) |
| EPI_ISL_16549380 | A/Vulpes_vulpes/Belgium/14775-0001/2022 | A / H5N1 | Europe / Belgium / Provincie Vlaams-Brabant / Vilvoorde | Steven Van Borm (Sciensano, Department of Animal Infectious Diseases / Animal Infectious Diseases) |
| EPI_ISL_16555202 | A/turkey/Minnesota/22-010654-002-original/2022 | A / H5N1 | North America / United States / Minnesota / Kandiyohi County | Mary Lea Killian (National Veterinary Services Laboratories - USDA / Diagnostic Virology Laboratory) |
| EPI_ISL_16555203 | A/neotropic cormorant/Arizona/245467/2022 | A / H5N1 | North America / United States / Arizona / Maricopa County | Mary Lea Killian (National Veterinary Services Laboratories - USDA / Diagnostic Virology Laboratory) |
| EPI_ISL_16555204 | A/eared grebe/North Dakota/245625/2022 | A / H5N1 | North America / United States / North Dakota / Stutsman County | Mary Lea Killian (National Veterinary Services Laboratories - USDA / Diagnostic Virology Laboratory) |
| EPI_ISL_16555205 | A/Canada goose/California/246038/2022 | A / H5N1 | North America / United States / California / Colusa County | Mary Lea Killian (National Veterinary Services Laboratories - USDA / Diagnostic Virology Laboratory) |
| EPI_ISL_1665247 | A/chicken/Italy/21VIR1151-2/2021 | A / H5N8 | Europe / Italy / Pordenone | Bianca Zecchin (Istituto Zooprofilattico Sperimentale Delle Venezie) |
| EPI_ISL_1665248 | A/chicken/Italy/21VIR1293-9/2021 | A / H5N8 | Europe / Italy / Padova | Bianca Zecchin (Istituto Zooprofilattico Sperimentale Delle Venezie) |
| EPI_ISL_1665249 | A/chicken/Romania/10101_21VIR2044-1/2021 | A / H5N5 | Europe / Romania / Judetul Timis | Bianca Zecchin (Istituto Zooprofilattico Sperimentale Delle Venezie) |
| EPI_ISL_1665250 | A/chicken/Slovakia/Pah10_21VIR1086-5/2021 | A / H5N5 | Europe / Slovakia / Dobrohost | Bianca Zecchin (Istituto Zooprofilattico Sperimentale Delle Venezie) |
| EPI_ISL_1665251 | A/common_teal/Italy/20VIR7439-191/2020 | A / H5N8 | Europe / Italy / Rovigo | Bianca Zecchin (Istituto Zooprofilattico Sperimentale Delle Venezie) |
| EPI_ISL_1665252 | A/duck/Italy/21VIR1293-15/2021 | A / H5N8 | Europe / Italy / Padova | Bianca Zecchin (Istituto Zooprofilattico Sperimentale Delle Venezie) |
| EPI_ISL_1665254 | A/duck/Romania/10206_21VIR849-4/2021 | A / H5N8 | Europe / Romania / Judetul Ilfov | Bianca Zecchin (Istituto Zooprofilattico Sperimentale Delle Venezie) |
| EPI_ISL_1665255 | A/guinea_fowl/Italy/21VIR1293-20/2021 | A / H5N8 | Europe / Italy / Padova | Bianca Zecchin (Istituto Zooprofilattico Sperimentale Delle Venezie) |
| EPI_ISL_1665256 | A/muscovy_duck/Slovakia/Pah1_21VIR1086-1/2021 | A / H5N8 | Europe / Slovakia / Kosice | Bianca Zecchin (Istituto Zooprofilattico Sperimentale Delle Venezie) |
| EPI_ISL_1665257 | A/mute_swan/Austria/21011165_21VIR1085-2/2021 | A / H5N8 | Europe / Austria / Tulln | Bianca Zecchin (Istituto Zooprofilattico Sperimentale Delle Venezie) |
| EPI_ISL_1665258 | A/mute_swan/Austria/21013162_21VIR1085-5/2021 | A / H5N5 | Europe / Austria / Leibnitz | Bianca Zecchin (Istituto Zooprofilattico Sperimentale Delle Venezie) |
| EPI_ISL_1665259 | A/mute_swan/Austria/21014124_21VIR1085-8/2021 | A / H5N8 | Europe / Austria / Vienna | Bianca Zecchin (Istituto Zooprofilattico Sperimentale Delle Venezie) |
| EPI_ISL_1665260 | A/mute_swan/Norway/FU5_21VIR850-3/2021 | A / H5N8 | Europe / Norway | Bianca Zecchin (Istituto Zooprofilattico Sperimentale Delle Venezie) |
| EPI_ISL_1665261 | A/mute_swan/Slovakia/Pah15_21VIR1086-3/2021 | A / H5N5 | Europe / Slovakia / Kalinkovo | Bianca Zecchin (Istituto Zooprofilattico Sperimentale Delle Venezie) |
| EPI_ISL_1665262 | A/mute_swan/Slovakia/Pah6_21VIR1086-2/2021 | A / H5N5 | Europe / Slovakia / Petrzalka | Bianca Zecchin (Istituto Zooprofilattico Sperimentale Delle Venezie) |
| EPI_ISL_1665263 | A/mute_swan/Slovenia/1639-20_21VIR959-1/2020 | A / H5N8 | Europe / Slovenia | Bianca Zecchin (Istituto Zooprofilattico Sperimentale Delle Venezie) |
| EPI_ISL_1665264 | A/mute_swan/Slovenia/1756-20_21VIR959-3/2020 | A / H5N8 | Europe / Slovenia | Bianca Zecchin (Istituto Zooprofilattico Sperimentale Delle Venezie) |
| EPI_ISL_1665265 | A/mute_swan/Slovenia/1799-20_21VIR959-2/2020 | A / H5N8 | Europe / Slovenia | Bianca Zecchin (Istituto Zooprofilattico Sperimentale Delle Venezie) |
| EPI_ISL_1665266 | A/mute_swan/Slovenia/1820-20_21VIR959-4/2020 | A / H5N8 | Europe / Slovenia | Bianca Zecchin (Istituto Zooprofilattico Sperimentale Delle Venezie) |
| EPI_ISL_1665267 | A/mute_swan/Slovenia/1914-20_21VIR959-5/2020 | A / H5N5 | Europe / Slovenia | Bianca Zecchin (Istituto Zooprofilattico Sperimentale Delle Venezie) |
| EPI_ISL_1665268 | A/whooper_swan/Romania/10123_21VIR849-1/2021 | A / H5N5 | Europe / Romania / Judetul Constanta | Bianca Zecchin (Istituto Zooprofilattico Sperimentale Delle Venezie) |
| EPI_ISL_1665269 | A/whooper_swan/Romania/10311_21VIR849-2/2021 | A / H5N5 | Europe / Romania / Judetul Constanta | Bianca Zecchin (Istituto Zooprofilattico Sperimentale Delle Venezie) |
| EPI_ISL_1665272 | A/common_buzzard/Italy/21VIR431-7/2020 | A / H5N8 | Europe / Italy / Lecce | Bianca Zecchin (Istituto Zooprofilattico Sperimentale Delle Venezie) |
| EPI_ISL_1665273 | A/geese/Romania/10206_21VIR849-3/2021 | A / H5N8 | Europe / Romania / Judetul Ilfov | Bianca Zecchin (Istituto Zooprofilattico Sperimentale Delle Venezie) |
| EPI_ISL_1665274 | A/house_sparrow/Romania/10327_21VIR849-5/2021 | A / H5N8 | Europe / Romania / Judetul Ilfov | Bianca Zecchin (Istituto Zooprofilattico Sperimentale Delle Venezie) |
| EPI_ISL_166584 | A/duck/Beijing/FS01/2013 | A / H5N8 | Asia / China / Beijing Municipality / Fangshan District | Di Liu (Institute of Microbiology, Chinese Academy of Sciences) |
| EPI_ISL_166630 | A/mallard/Korea/W452/2014 | A / H5N8 | Asia / Korea, Republic of | |
| EPI_ISL_166693 | A/duck/Beijing/FS01/2014 | A / H5N8 | Asia / China / Beijing Municipality / Fangshan District | Di Liu (Institute of Microbiology, Chinese Academy of Sciences) |
| EPI_ISL_166694 | A/duck/Beijing/CT01/2014 | A / H5N8 | Asia / China / Beijing Municipality / Shunyi District | Di Liu (Institute of Microbiology, Chinese Academy of Sciences) |
| EPI_ISL_167140 | A/turkey/Germany-MV/R2472/2014 | A / H5N8 | Europe / Germany / Mecklenburg-Vorpommern | Elke Starick (Friedrich-Loeffler-Institut) |
| EPI_ISL_167178 | A/duck/Shandong/Q1/2013 | A / H5N8 | Asia / China | |
| EPI_ISL_167767 | A/environment/Zhenjiang/C13/2013 | A / H5N6 | Asia / China | |
| EPI_ISL_167904 | A/duck/England/36254/14 | A / H5N8 | Europe / United Kingdom / East Riding of Yorkshire | Amanda Seekings (Animal and Plant Health Agency (APHA) / Virology Department) |
| EPI_ISL_167905 | A/Chicken/Netherlands/14015526/2014 | A / H5N8 | Europe / Netherlands / Provincie Utrecht / Gemeente Oudewater | Guus Koch (Wageningen Bioveterinary Research) |
| EPI_ISL_168024 | A/duck/Chiba/26-372-48/2014 | A / H5N8 | Asia / Japan / Chiba | Takehiko Saito (National Institute of Animal Health) |
| EPI_ISL_168025 | A/duck/Chiba/26-372-61/2014 | A / H5N8 | Asia / Japan / Chiba | Takehiko Saito (National Institute of Animal Health) |
| EPI_ISL_168075 | A/chicken/Netherlands/14015531/2014 | A / H5N8 | Europe / Netherlands / Provincie Utrecht / Gemeente Oudewater | Guus Koch (Wageningen Bioveterinary Research) |
| EPI_ISL_168384 | A/chicken/Laos/LPQ001/2014 | A / H5N6 | Asia / Lao, People's Democratic Republic | |
| EPI_ISL_168385 | A/duck/Laos/XBY004/2014 | A / H5N6 | Asia / Lao, People's Democratic Republic | |
| EPI_ISL_168663 | A/duck/Laos/LPQ002/2014 | A / H5N6 | Asia / Lao, People's Democratic Republic | |
| EPI_ISL_168664 | A/chicken/Laos/XBY003/2014 | A / H5N6 | Asia / Lao, People's Democratic Republic | |
| EPI_ISL_168696 | A/duck/England/36038/14 | A / H5N8 | Europe / United Kingdom / East Riding of Yorkshire | Amanda Seekings (Animal and Plant Health Agency (APHA) / Virology Department) |

| | | | | |
|---|---|---|---|---|
| EPI_ISL_168697 | A/duck/England/36226/14 | A / H5N8 | Europe / United Kingdom / East Riding of Yorkshire | Amanda Seekings (Animal and Plant Health Agency (APHA) / Virology Department) |
| EPI_ISL_168746 | A/eurasian wigeon/Netherlands/emc-1/2014 | A / H5N8 | Europe / Netherlands / Provincie Utrecht / Gemeente Woerden / Between Kamerik and Kockeng | Josanne Verhagen (Erasmus Medical Center / Department of Virology) |
| EPI_ISL_168747 | A/eurasian wigeon/Netherlands/emc-2/2014 | A / H5N8 | Europe / Netherlands / Provincie Utrecht / Gemeente Woerden / Between Kamerik and Kockeng | Josanne Verhagen (Erasmus Medical Center / Department of Virology) |
| EPI_ISL_169273 | A/turkey/Germany/AR2485-86-L00899/2014 | A / H5N8 | Europe / Germany / Mecklenburg-Vorpommern | Anne Pohlmann (Friedrich-Loeffler-Institut) |
| EPI_ISL_169282 | A/chicken/Netherlands/emc-3/2014 | A / H5N8 | Europe / Netherlands / South Holland / Gemeente Nieuwkoop | Josanne Verhagen (Erasmus Medical Center / Department of Virology) |
| EPI_ISL_169350 | A/turkey/Italy/14VIR7898-10/2014 | A / H5N8 | Europe / Italy / Veneto / Province of Rovigo | Alice Fusaro (Istituto Zooprofilattico Sperimentale Delle Venezie) |
| EPI_ISL_169390 | A/crane/Kagoshima/KU1/2014 | A / H5N8 | Asia / Japan / Kagoshima | Makoto Ozawa (Kagoshima University ) |
| EPI_ISL_169422 | A/chicken/Miyazaki/3/2014 | A / H5N8 | Asia / Japan / Miyazaki | Takehiko Saito (National Institute of Animal Health) |
| EPI_ISL_169423 | A/chicken/Miyazaki/4/2014 | A / H5N8 | Asia / Japan / Miyazaki | Takehiko Saito (National Institute of Animal Health) |
| EPI_ISL_169424 | A/chicken/Miyazaki/7/2014 | A / H5N8 | Asia / Japan / Miyazaki | Takehiko Saito (National Institute of Animal Health) |
| EPI_ISL_169427 | A/wigeon/Sakha/1/2014 | A / H5N8 | Europe / Russian Federation / Sakha (Yakutia) Republic | Ivan Susloparov (State Research Center of Virology and Biotechnology (VECTOR) / Emerging Zoonotic Diseases and Influenza) |
| EPI_ISL_169429 | A/environment/Kagoshima/KU-ngr-H/2014 | A / H5N8 | Asia / Japan / Kagoshima | Makoto Ozawa (Kagoshima University ) |
| EPI_ISL_169632 | A/domestic duck/Germany-NI/R3468/2014 | A / H5N8 | Europe / Germany / Lower Saxony | Elke Starick (Friedrich-Loeffler-Institut) |
| EPI_ISL_1697181 | A/chicken/Czech Republic/6151-1/2021 | A / H5N8 | Europe / Czech Republic / Kralovehradecky Kraj / Okres Hradec Kralove / Kosicky | Alexander Nagy (State Veterinary Institute Prague) |
| EPI_ISL_1697182 | A/chicken/Czech Republic/5903/2021 | A / H5N8 | Europe / Czech Republic / Kralovehradecky Kraj / Okres Jicin / Vysoke Veseli | Alexander Nagy (State Veterinary Institute Prague) |
| EPI_ISL_1697183 | A/chicken/Czech Republic/6684/2021 | A / H5N8 | Europe / Czech Republic / Ustecky Kraj / Okres Most / Polerady | Alexander Nagy (State Veterinary Institute Prague) |
| EPI_ISL_1697184 | A/chicken/Czech Republic/6654/2021 | A / H5N8 | Europe / Czech Republic / Plzensky Kraj / Okres Tacov / Bor; GPS: 49.7115939N, 12.7751636E | Alexander Nagy (State Veterinary Institute Prague) |
| EPI_ISL_1697185 | A/duck/Czech Republic/6653-4/2021 | A / H5N8 | Europe / Czech Republic / Kralovehradecky Kraj / Okres Hradec Kralove / Stary Bydzov | Alexander Nagy (State Veterinary Institute Prague) |
| EPI_ISL_1697186 | A/duck/Czech Republic/6653-6/2021 | A / H5N8 | Europe / Czech Republic / Kralovehradecky Kraj / Okres Hradec Kralove / Stary Bydzov | Alexander Nagy (State Veterinary Institute Prague) |
| EPI_ISL_1697187 | A/chicken/Czech Republic/6542-1/2021 | A / H5N8 | Europe / Czech Republic / Hlavni mesto Praha / Okres Praha / Trebotov; GPS: 49.6993983N, 14.2 | Alexander Nagy (State Veterinary Institute Prague) |
| EPI_ISL_1697188 | A/chicken/Czech Republic/6532-1/2021 | A / H5N8 | Europe / Czech Republic / Morovskoslezsky Kraj / Okres Bruntal / Zator | Alexander Nagy (State Veterinary Institute Prague) |
| EPI_ISL_1697189 | A/chicken/Czech Republic/6532-2/2021 | A / H5N8 | Europe / Czech Republic / Morovskoslezsky Kraj / Okres Bruntal / Zator | Alexander Nagy (State Veterinary Institute Prague) |
| EPI_ISL_1697190 | A/peacock/Czech Republic/6529-2/2021 | A / H5N8 | Europe / Czech Republic / Ustecky Kraj / Okres Teplice / Bilina, Razice; GPS: 50.5215139N, 13.825 | Alexander Nagy (State Veterinary Institute Prague) |
| EPI_ISL_1697191 | A/chicken/Czech Republic/6527/2021 | A / H5N8 | Europe / Czech Republic / Kralovehradecky Kraj / Okres Hradec Kralove / Dobrenice | Alexander Nagy (State Veterinary Institute Prague) |
| EPI_ISL_1697192 | A/duck/Czech Republic/5792-13/2021 | A / H5N8 | Europe / Czech Republic / Kralovehradecky Kraj / Okres Hradec Kralove / Lukova | Alexander Nagy (State Veterinary Institute Prague) |
| EPI_ISL_1697193 | A/duck/Czech Republic/5792-19/2021 | A / H5N8 | Europe / Czech Republic / Kralovehradecky Kraj / Okres Hradec Kralove / Lukova | Alexander Nagy (State Veterinary Institute Prague) |
| EPI_ISL_1697194 | A/chicken/Czech Republic/4980/2021 | A / H5N8 | Europe / Czech Republic / Zlinsky Kraj / Okres Uherske Hradiste / Osvetimany | Alexander Nagy (State Veterinary Institute Prague) |
| EPI_ISL_1697195 | A/duck/Czech Republic/5466/2021 | A / H5N8 | Europe / Czech Republic / Pardubicky kraj / Okres Pardubice / Vapno; GPS: 50.1074822N, 15.527 | Alexander Nagy (State Veterinary Institute Prague) |
| EPI_ISL_1697196 | A/duck/Czech Republic/5448/2021 | A / H5N8 | Europe / Czech Republic / Stredocesky Kraj / Okres Nymburk / Vinice u Mestce Kralove | Alexander Nagy (State Veterinary Institute Prague) |
| EPI_ISL_1697197 | A/australian brushturkey/Czech Republic/5904/2021 | A / H5N8 | Europe / Czech Republic / Plzensky Kraj | Alexander Nagy (State Veterinary Institute Prague) |
| EPI_ISL_1697198 | A/duck/Czech Republic/5360-1/2021 | A / H5N8 | Europe / Czech Republic / Kralovehradecky Kraj / Okres Hradec Kralove / Dobrenice | Alexander Nagy (State Veterinary Institute Prague) |
| EPI_ISL_1697199 | A/duck/Czech Republic/5360-2/2021 | A / H5N8 | Europe / Czech Republic / Kralovehradecky Kraj / Okres Hradec Kralove / Dobrenice | Alexander Nagy (State Veterinary Institute Prague) |
| EPI_ISL_1697200 | A/chicken/Czech Republic/6542-2/2021 | A / H5N8 | Europe / Czech Republic / Hlavni mesto Praha / Okres Praha / Trebotov; GPS: 49.6993983N, 14.2 | Alexander Nagy (State Veterinary Institute Prague) |
| EPI_ISL_169851 | A/stork/Germany-MV/R24/2015 | A / H5N8 | Europe / Germany / Mecklenburg-Vorpommern | Elke Starick (Friedrich-Loeffler-Institut) |
| EPI_ISL_170155 | A/Environment/Xuzhou/yj010/2014 | A / H5N6 | Asia / China / jiangsu | Lunbiao Cui (Jiangsu Provincial Center for Disease Control & Prevention) |
| EPI_ISL_170184 | A/chicken/Germany-MV/R153/2015 | A / H5N8 | Europe / Germany / Mecklenburg-Vorpommern | Elke Starick (Friedrich-Loeffler-Institut) |
| EPI_ISL_170230 | A/Environment/Zhenjiang/zj35/2014 | A / H5N8 | Asia / China / jiangsu | Lunbiao Cui (Jiangsu Provincial Center for Disease Control & Prevention) |
| EPI_ISL_170234 | A/Chicken/Changzhou/cz93/2014 | A / H5N8 | Asia / China / jiangsu | Lunbiao Cui (Jiangsu Provincial Center for Disease Control & Prevention) |
| EPI_ISL_170236 | A/Environment/Huaian/ha118/2014 | A / H5N8 | Asia / China / jiangsu | Lunbiao Cui (Jiangsu Provincial Center for Disease Control & Prevention) |
| EPI_ISL_171655 | A/waterfowl/Korea/S005/2014 | A / H5N8 | Asia / Korea, Republic of | |
| EPI_ISL_171656 | A/broiler duck/Korea/H29/2014 | A / H5N8 | Asia / Korea, Republic of | |
| EPI_ISL_171657 | A/broiler duck/Korea/H31/2014 | A / H5N8 | Asia / Korea, Republic of | |
| EPI_ISL_171674 | A/Baikal teal/Korea/H41/2014 | A / H5N8 | Asia / Korea, Republic of | |
| EPI_ISL_171675 | A/broiler duck/Korea/H47/2014 | A / H5N8 | Asia / Korea, Republic of | |
| EPI_ISL_171676 | A/broiler duck/Korea/H48/2014 | A / H5N8 | Asia / Korea, Republic of | |
| EPI_ISL_171677 | A/broiler duck/Korea/H49/2014 | A / H5N8 | Asia / Korea, Republic of | |
| EPI_ISL_171678 | A/bean goose/Korea/H40/2014 | A / H5N8 | Asia / Korea, Republic of | |
| EPI_ISL_171695 | A/Baikal teal/Korea/H52/2014 | A / H5N8 | Asia / Korea, Republic of | |
| EPI_ISL_171696 | A/bean goose/Korea/H53/2014 | A / H5N8 | Asia / Korea, Republic of | |
| EPI_ISL_171697 | A/Baikal teal/Korea/H62/2014 | A / H5N8 | Asia / Korea, Republic of | |
| EPI_ISL_171698 | A/broiler duck/Korea/H65/2014 | A / H5N8 | Asia / Korea, Republic of | |
| EPI_ISL_171699 | A/Baikal teal/Korea/H66/2014 | A / H5N8 | Asia / Korea, Republic of | |
| EPI_ISL_171700 | A/Baikal teal/Korea/H68/2014 | A / H5N8 | Asia / Korea, Republic of | |
| EPI_ISL_171701 | A/Baikal teal/Korea/H80/2014 | A / H5N8 | Asia / Korea, Republic of | |
| EPI_ISL_171702 | A/Coot/Korea/H81/2014 | A / H5N8 | Asia / Korea, Republic of | |
| EPI_ISL_171703 | A/Baikal teal/Korea/H84/2014 | A / H5N8 | Asia / Korea, Republic of | |
| EPI_ISL_171704 | A/Baikal teal/Korea/H96/2014 | A / H5N8 | Asia / Korea, Republic of | |
| EPI_ISL_171705 | A/breeder chicken/Korea/H122/2014 | A / H5N8 | Asia / Korea, Republic of | |
| EPI_ISL_171706 | A/breeder duck/Korea/H128/2014 | A / H5N8 | Asia / Korea, Republic of | |
| EPI_ISL_171707 | A/broiler duck/Korea/H133/2014 | A / H5N8 | Asia / Korea, Republic of | |
| EPI_ISL_171708 | A/broiler duck/Korea/H145/2014 | A / H5N8 | Asia / Korea, Republic of | |
| EPI_ISL_171709 | A/breeder duck/Korea/H158/2014 | A / H5N8 | Asia / Korea, Republic of | |
| EPI_ISL_171710 | A/breeder duck/Korea/H200/2014 | A / H5N8 | Asia / Korea, Republic of | |
| EPI_ISL_171711 | A/mallard/Korea/H207/2014 | A / H5N8 | Asia / Korea, Republic of | |
| EPI_ISL_171712 | A/white-fronted goose/Korea/H231/2014 | A / H5N8 | Asia / Korea, Republic of | |
| EPI_ISL_171713 | A/breeder duck/Korea/H249/2014 | A / H5N8 | Asia / Korea, Republic of | |
| EPI_ISL_171714 | A/breeder chicken/Korea/H250/2014 | A / H5N8 | Asia / Korea, Republic of | |
| EPI_ISL_171715 | A/Korean native chicken/Korea/H257/2014 | A / H5N8 | Asia / Korea, Republic of | |
| EPI_ISL_171716 | A/mallard/Korea/H297/2014 | A / H5N8 | Asia / Korea, Republic of | |
| EPI_ISL_171717 | A/bean goose/Korea/H328/2014 | A / H5N8 | Asia / Korea, Republic of | |
| EPI_ISL_171718 | A/tundra swan/Korea/H411/2014 | A / H5N8 | Asia / Korea, Republic of | |
| EPI_ISL_171719 | A/common teal/Korea/H455-30/2014 | A / H5N8 | Asia / Korea, Republic of | |

| | | | | |
|---|---|---|---|---|
| EPI_ISL_171720 | A/spot-billed duck/Korea/H455-42/2014 | A / H5N8 | Asia / Korea, Republic of | |
| EPI_ISL_1719909 | A/swan/Lithuania/1258PG1_21VIR2606-2/2021 | A / H5N8 | Europe / Lithuania / Vilnius | Bianca Zecchin (Istituto Zooprofilattico Sperimentale Delle Venezie) |
| EPI_ISL_1719910 | A/swan/Lithuania/1298PG1_21VIR2606-3/2021 | A / H5N8 | Europe / Lithuania / Vilnius | Bianca Zecchin (Istituto Zooprofilattico Sperimentale Delle Venezie) |
| EPI_ISL_1719911 | A/swan/Lithuania/1842PG1_21VIR2606-5/2021 | A / H5N8 | Europe / Lithuania / Vilnius | Bianca Zecchin (Istituto Zooprofilattico Sperimentale Delle Venezie) |
| EPI_ISL_1719912 | A/swan/Lithuania/2306PG1_21VIR2606-7/2021 | A / H5N8 | Europe / Lithuania / Vilnius | Bianca Zecchin (Istituto Zooprofilattico Sperimentale Delle Venezie) |
| EPI_ISL_1719913 | A/chicken/Bulgaria/39_21VIR1454-3/2021 | A / H5N8 | Europe / Bulgaria / Slavyanovo | Bianca Zecchin (Istituto Zooprofilattico Sperimentale Delle Venezie) |
| EPI_ISL_1719914 | A/pekin_duck/Bulgaria/48-3_21VIR1454-7/2021 | A / H5N8 | Europe / Bulgaria / Slavyanovo | Bianca Zecchin (Istituto Zooprofilattico Sperimentale Delle Venezie) |
| EPI_ISL_1719915 | A/chicken/Bulgaria/50-1_21VIR1454-9/2021 | A / H5N8 | Europe / Bulgaria / Slavyanovo | Bianca Zecchin (Istituto Zooprofilattico Sperimentale Delle Venezie) |
| EPI_ISL_172815 | A/ibis/Germany-MV/R44/2015 | A / H5N8 | Europe / Germany / Mecklenburg-Vorpommern | Elke Starick (Friedrich-Loeffler-Institut) |
| EPI_ISL_173478 | A/duck/Jiangxi/13469/2014 | A / H5N6 | Asia / China | |
| EPI_ISL_173479 | A/duck/Jiangxi/13475/2014 | A / H5N6 | Asia / China | |
| EPI_ISL_173480 | A/duck/Jiangxi/13484/2014 | A / H5N6 | Asia / China | |
| EPI_ISL_173482 | A/black chicken/Jiangxi/10129/2014 | A / H5N6 | Asia / China | |
| EPI_ISL_173483 | A/environment/Jiangxi/10170/2014 | A / H5N6 | Asia / China | |
| EPI_ISL_173531 | A/goose/Shantou/1763/2014 | A / H5N6 | Asia / China | |
| EPI_ISL_173532 | A/goose/Shantou/1791/2014 | A / H5N6 | Asia / China | |
| EPI_ISL_173533 | A/goose/Shantou/1806/2014 | A / H5N6 | Asia / China | |
| EPI_ISL_173568 | A/green-winged teal/Korea/KU-12/2015 | A / H5 | Asia / Korea, Republic of / Chungcheongnam-do | Dong-Hun Lee (Konkuk University / College of Veterinary Medicine) |
| EPI_ISL_173664 | A/chicken/Washington/61-9/2014 | A / H5N2 | North America / United States | |
| EPI_ISL_173665 | A/turkey/Washington/61-22/2014 | A / H5N2 | North America / United States | |
| EPI_ISL_173666 | A/chicken/Oregon/41613-2/2014 | A / H5N8 | North America / United States | |
| EPI_ISL_173667 | A/domestic duck/Washington/61-16/2014 | A / H5N2 | North America / United States | |
| EPI_ISL_173668 | A/guinea fowl/Oregon/41613-1/2014 | A / H5N8 | North America / United States | |
| EPI_ISL_173669 | A/duck/Zhejiang/6D2/2013 | A / H5N6 | Asia / China | |
| EPI_ISL_173670 | A/chicken/Zhejiang/6C2/2013 | A / H5N6 | Asia / China | |
| EPI_ISL_173877 | A/Northern pintail/Washington/40964/2014 | A / H5N2 | North America / United States | |
| EPI_ISL_173878 | A/gyrfalcon/Washington/41088-6/2014 | A / H5N8 | North America / United States | |
| EPI_ISL_174349 | A/chicken/Netherlands/14015766/2014 | A / H5N8 | Europe / Netherlands / South Holland / Gemeente Nieuwkoop / Ter Aar | Guus Koch (Wageningen Bioveterinary Research) |
| EPI_ISL_174350 | A/Chicken/Netherlands/14015824/2014 | A / H5N8 | Europe / Netherlands / Provincie Overijssel / Gemeente Kampen / Kamperveen | Guus Koch (Wageningen Bioveterinary Research) |
| EPI_ISL_174351 | A/duck/Netherlands/14015989/2014 | A / H5N8 | Europe / Netherlands / Provincie Overijssel / Gemeente Kampen / Kamperveen | Guus Koch (Wageningen Bioveterinary Research) |
| EPI_ISL_174352 | A/chicken/Netherlands/14016437/2014 | A / H5N8 | Europe / Netherlands / South Holland / Gemeente Zoeterwoude / 2381 NZ | Guus Koch (Wageningen Bioveterinary Research) |
| EPI_ISL_174353 | A/breeder chicken/Korea/H0503/2014 | A / H5N8 | Asia / Korea, Republic of / Jeollabuk-do | Eun-Kyoung Lee (Animal and Plant Quarantine Agency (APQA) / Avian Influenza Research and Diagsnotic Division) |
| EPI_ISL_174354 | A/breeder chicken/Korea/H0818/2014 | A / H5N8 | Asia / Korea, Republic of / Chungcheongnam-do | Eun-Kyoung Lee (Animal and Plant Quarantine Agency (APQA) / Avian Influenza Research and Diagsnotic Division) |
| EPI_ISL_174355 | A/broiler duck/Korea/H0651/2014 | A / H5N8 | Asia / Korea, Republic of / Jeollanam-do | Eun-Kyoung Lee (Animal and Plant Quarantine Agency (APQA) / Avian Influenza Research and Diagsnotic Division) |
| EPI_ISL_174356 | A/chicken/Korea/H0881/2014 | A / H5N8 | Asia / Korea, Republic of | Eun-Kyoung Lee (Animal and Plant Quarantine Agency (APQA) / Avian Influenza Research and Diagsnotic Division) |
| EPI_ISL_174357 | A/broiler duck/Korea/H0959/2014 | A / H5N8 | Asia / Korea, Republic of / Jeollanam-do | Eun-Kyoung Lee (Animal and Plant Quarantine Agency (APQA) / Avian Influenza Research and Diagsnotic Division) |
| EPI_ISL_174358 | A/breeder chicken/Korea/H1068/2014 | A / H5N8 | Asia / Korea, Republic of / Chungcheongnam-do | Eun-Kyoung Lee (Animal and Plant Quarantine Agency (APQA) / Avian Influenza Research and Diagsnotic Division) |
| EPI_ISL_174359 | A/Korean native chicken/Korea/H1139/2014 | A / H5N8 | Asia / Korea, Republic of / Chungcheongnam-do | Eun-Kyoung Lee (Animal and Plant Quarantine Agency (APQA) / Avian Influenza Research and Diagsnotic Division) |
| EPI_ISL_174360 | A/chicken/Korea/H1236/2014 | A / H5N8 | Asia / Korea, Republic of / Chungcheongnam-do | Eun-Kyoung Lee (Animal and Plant Quarantine Agency (APQA) / Avian Influenza Research and Diagsnotic Division) |
| EPI_ISL_174361 | A/chicken/Korea/H1268/2014 | A / H5N8 | Asia / Korea, Republic of / Gyeongsangnam-do | Eun-Kyoung Lee (Animal and Plant Quarantine Agency (APQA) / Avian Influenza Research and Diagsnotic Division) |
| EPI_ISL_174362 | A/chicken/Korea/H1292/2014 | A / H5N8 | Asia / Korea, Republic of / Gyeonggi-do | Eun-Kyoung Lee (Animal and Plant Quarantine Agency (APQA) / Avian Influenza Research and Diagsnotic Division) |
| EPI_ISL_174363 | A/goose/Korea/H1296/2014 | A / H5N8 | Asia / Korea, Republic of / Chungcheongbuk-do | Eun-Kyoung Lee (Animal and Plant Quarantine Agency (APQA) / Avian Influenza Research and Diagsnotic Division) |
| EPI_ISL_174364 | A/Korean native chicken/Korea/H1299/2014 | A / H5N8 | Asia / Korea, Republic of / Ulsan | Eun-Kyoung Lee (Animal and Plant Quarantine Agency (APQA) / Avian Influenza Research and Diagsnotic Division) |
| EPI_ISL_174365 | A/breeder duck/Korea/H1343/2014 | A / H5N8 | Asia / Korea, Republic of / Jeollanam-do | Eun-Kyoung Lee (Animal and Plant Quarantine Agency (APQA) / Avian Influenza Research and Diagsnotic Division) |
| EPI_ISL_174366 | A/chicken/Korea/H1350/2014 | A / H5N8 | Asia / Korea, Republic of / Chungcheongnam-do | Eun-Kyoung Lee (Animal and Plant Quarantine Agency (APQA) / Avian Influenza Research and Diagsnotic Division) |
| EPI_ISL_174367 | A/gadwall/Korea/H1351/2014 | A / H5N8 | Asia / Korea, Republic of / Jeju-teukbyeoljachido | Eun-Kyoung Lee (Animal and Plant Quarantine Agency (APQA) / Avian Influenza Research and Diagsnotic Division) |
| EPI_ISL_174368 | A/broiler duck/Korea/H1413/2014 | A / H5N8 | Asia / Korea, Republic of / Jeollanam-do | Eun-Kyoung Lee (Animal and Plant Quarantine Agency (APQA) / Avian Influenza Research and Diagsnotic Division) |
| EPI_ISL_174369 | A/broiler duck/Korea/H1414/2014 | A / H5N8 | Asia / Korea, Republic of / Jeollanam-do | Eun-Kyoung Lee (Animal and Plant Quarantine Agency (APQA) / Avian Influenza Research and Diagsnotic Division) |
| EPI_ISL_174370 | A/goose/Korea/H1545/2014 | A / H5N8 | Asia / Korea, Republic of / Gangwon-do | Eun-Kyoung Lee (Animal and Plant Quarantine Agency (APQA) / Avian Influenza Research and Diagsnotic Division) |
| EPI_ISL_174371 | A/Korean native chicken/Korea/H1554/2014 | A / H5N8 | Asia / Korea, Republic of / Daegu | Eun-Kyoung Lee (Animal and Plant Quarantine Agency (APQA) / Avian Influenza Research and Diagsnotic Division) |
| EPI_ISL_174372 | A/broiler duck/Korea/H1556/2014 | A / H5N8 | Asia / Korea, Republic of / Jeollanam-do | Eun-Kyoung Lee (Animal and Plant Quarantine Agency (APQA) / Avian Influenza Research and Diagsnotic Division) |
| EPI_ISL_174373 | A/broiler duck/Korea/H1582/2014 | A / H5N8 | Asia / Korea, Republic of / Gyeonggi-do | Eun-Kyoung Lee (Animal and Plant Quarantine Agency (APQA) / Avian Influenza Research and Diagsnotic Division) |
| EPI_ISL_174374 | A/breeder duck/Korea/H1596/2014 | A / H5N8 | Asia / Korea, Republic of / Jeollanam-do | Eun-Kyoung Lee (Animal and Plant Quarantine Agency (APQA) / Avian Influenza Research and Diagsnotic Division) |
| EPI_ISL_174375 | A/broiler duck/Korea/H1683/2014 | A / H5N8 | Asia / Korea, Republic of / Jeollanam-do | Eun-Kyoung Lee (Animal and Plant Quarantine Agency (APQA) / Avian Influenza Research and Diagsnotic Division) |
| EPI_ISL_174376 | A/broiler duck/Korea/H1685/2014 | A / H5N8 | Asia / Korea, Republic of / Jeollanam-do | Eun-Kyoung Lee (Animal and Plant Quarantine Agency (APQA) / Avian Influenza Research and Diagsnotic Division) |
| EPI_ISL_174377 | A/Korean native chicken/Korea/H1687/2014 | A / H5N8 | Asia / Korea, Republic of / Jeollanam-do | Eun-Kyoung Lee (Animal and Plant Quarantine Agency (APQA) / Avian Influenza Research and Diagsnotic Division) |
| EPI_ISL_174378 | A/goose/Korea/H1689/2014 | A / H5N8 | Asia / Korea, Republic of / Jeollanam-do | Eun-Kyoung Lee (Animal and Plant Quarantine Agency (APQA) / Avian Influenza Research and Diagsnotic Division) |
| EPI_ISL_174379 | A/goose/Korea/H1698/2014 | A / H5N8 | Asia / Korea, Republic of / Jeollanam-do | Eun-Kyoung Lee (Animal and Plant Quarantine Agency (APQA) / Avian Influenza Research and Diagsnotic Division) |
| EPI_ISL_174380 | A/broiler duck/Korea/H1731/2014 | A / H5N8 | Asia / Korea, Republic of / Jeollanam-do | Eun-Kyoung Lee (Animal and Plant Quarantine Agency (APQA) / Avian Influenza Research and Diagsnotic Division) |
| EPI_ISL_174381 | A/broiler duck/Korea/H1733/2014 | A / H5N8 | Asia / Korea, Republic of / Jeollanam-do | Eun-Kyoung Lee (Animal and Plant Quarantine Agency (APQA) / Avian Influenza Research and Diagsnotic Division) |
| EPI_ISL_174382 | A/broiler duck/Korea/H1734/2014 | A / H5N8 | Asia / Korea, Republic of / Jeollanam-do | Eun-Kyoung Lee (Animal and Plant Quarantine Agency (APQA) / Avian Influenza Research and Diagsnotic Division) |
| EPI_ISL_174383 | A/broiler duck/Korea/H1739/2014 | A / H5N8 | Asia / Korea, Republic of / Jeollanam-do | Eun-Kyoung Lee (Animal and Plant Quarantine Agency (APQA) / Avian Influenza Research and Diagsnotic Division) |
| EPI_ISL_174384 | A/broiler duck/Korea/H1745/2014 | A / H5N8 | Asia / Korea, Republic of / Jeollanam-do | Eun-Kyoung Lee (Animal and Plant Quarantine Agency (APQA) / Avian Influenza Research and Diagsnotic Division) |
| EPI_ISL_174385 | A/Korean native chicken/Korea/H1747/2014 | A / H5N8 | Asia / Korea, Republic of / Jeollanam-do | Eun-Kyoung Lee (Animal and Plant Quarantine Agency (APQA) / Avian Influenza Research and Diagsnotic Division) |
| EPI_ISL_174386 | A/breeder duck/Korea/H1752/2014 | A / H5N8 | Asia / Korea, Republic of / Jeollanam-do | Eun-Kyoung Lee (Animal and Plant Quarantine Agency (APQA) / Avian Influenza Research and Diagsnotic Division) |
| EPI_ISL_174387 | A/broiler duck/Korea/H1755/2014 | A / H5N8 | Asia / Korea, Republic of / Jeollanam-do | Eun-Kyoung Lee (Animal and Plant Quarantine Agency (APQA) / Avian Influenza Research and Diagsnotic Division) |
| EPI_ISL_174388 | A/broiler duck/Korea/H1763/2014 | A / H5N8 | Asia / Korea, Republic of / Jeollanam-do | Eun-Kyoung Lee (Animal and Plant Quarantine Agency (APQA) / Avian Influenza Research and Diagsnotic Division) |
| EPI_ISL_174389 | A/broiler duck/Korea/H1803/2014 | A / H5N8 | Asia / Korea, Republic of / Jeollanam-do | Eun-Kyoung Lee (Animal and Plant Quarantine Agency (APQA) / Avian Influenza Research and Diagsnotic Division) |
| EPI_ISL_174390 | A/broiler duck/Korea/H1839/2014 | A / H5N8 | Asia / Korea, Republic of / Jeollanam-do | Eun-Kyoung Lee (Animal and Plant Quarantine Agency (APQA) / Avian Influenza Research and Diagsnotic Division) |
| EPI_ISL_174391 | A/broiler duck/Korea/H1840/2014 | A / H5N8 | Asia / Korea, Republic of / Jeollabuk-do | Eun-Kyoung Lee (Animal and Plant Quarantine Agency (APQA) / Avian Influenza Research and Diagsnotic Division) |
| EPI_ISL_174392 | A/Korean native chicken/Korea/H1847/2014 | A / H5N8 | Asia / Korea, Republic of / Jeollanam-do | Eun-Kyoung Lee (Animal and Plant Quarantine Agency (APQA) / Avian Influenza Research and Diagsnotic Division) |
| EPI_ISL_174393 | A/broiler duck/Korea/H1864/2014 | A / H5N8 | Asia / Korea, Republic of / Jeollabuk-do | Eun-Kyoung Lee (Animal and Plant Quarantine Agency (APQA) / Avian Influenza Research and Diagsnotic Division) |

| | | | | |
|---|---|---|---|---|
| EPI_ISL_174394 | A/Korean native chicken/Korea/H1903/2014 | A / H5N8 | Asia / Korea, Republic of / Gyeongsangbuk-do | Eun-Kyoung Lee (Animal and Plant Quarantine Agency (APQA) / Avian Influenza Research and Diagsnotic Division) |
| EPI_ISL_174395 | A/spot-billed duck/Korea/H1981/2014 | A / H5N8 | Asia / Korea, Republic of / Chungcheongbuk-do | Eun-Kyoung Lee (Animal and Plant Quarantine Agency (APQA) / Avian Influenza Research and Diagsnotic Division) |
| EPI_ISL_174396 | A/mallard/Korea/H1991/2014 | A / H5N8 | Asia / Korea, Republic of / Gyeonggi-do | Eun-Kyoung Lee (Animal and Plant Quarantine Agency (APQA) / Avian Influenza Research and Diagsnotic Division) |
| EPI_ISL_174397 | A/mallard/Korea/H2003/2014 | A / H5N8 | Asia / Korea, Republic of / Gyeonggi-do | Eun-Kyoung Lee (Animal and Plant Quarantine Agency (APQA) / Avian Influenza Research and Diagsnotic Division) |
| EPI_ISL_174398 | A/mallard/Korea/H1924-6/2014 | A / H5N8 | Asia / Korea, Republic of / Jeollanam-do | Eun-Kyoung Lee (Animal and Plant Quarantine Agency (APQA) / Avian Influenza Research and Diagsnotic Division) |
| EPI_ISL_174399 | A/breeder duck/Korea/H0345/2014 | A / H5N8 | Asia / Korea, Republic of / Chungcheongbuk-do | Eun-Kyoung Lee (Animal and Plant Quarantine Agency (APQA) / Avian Influenza Research and Diagsnotic Division) |
| EPI_ISL_174400 | A/breeder duck/Korea/H0566/2014 | A / H5N8 | Asia / Korea, Republic of / Chungcheongbuk-do | Eun-Kyoung Lee (Animal and Plant Quarantine Agency (APQA) / Avian Influenza Research and Diagsnotic Division) |
| EPI_ISL_174401 | A/Common Teal/Korea/H844/2014 | A / H5N8 | Asia / Korea, Republic of / Chungcheongnam-do | Eun-Kyoung Lee (Animal and Plant Quarantine Agency (APQA) / Avian Influenza Research and Diagsnotic Division) |
| EPI_ISL_174409 | A/American green-winged teal/Washington/195750/2014 | A / H5N1 | North America / United States | |
| EPI_ISL_174411 | A/turkey/BC/FAV10/2014 | A / H5N2 | North America / Canada | |
| EPI_ISL_174516 | A/crane/Kagoshima/KU13/2014(H5N8) | A / H5N8 | Asia / Japan / Kagoshima | Makoto Ozawa (Kagoshima University ) |
| EPI_ISL_174517 | A/crane/Kagoshima/KU21/2014(H5N8) | A / H5N8 | Asia / Japan / Kagoshima | Makoto Ozawa (Kagoshima University ) |
| EPI_ISL_174518 | A/crane/Kagoshima/KU41/2014(H5N8) | A / H5N8 | Asia / Japan / Kagoshima | Makoto Ozawa (Kagoshima University ) |
| EPI_ISL_174519 | A/crane/Kagoshima/KU53/2015(H5N8) | A / H5N8 | Asia / Japan / Kagoshima | Makoto Ozawa (Kagoshima University ) |
| EPI_ISL_174520 | A/mallard duck/Kagoshima/KU70/2015(H5N8) | A / H5N8 | Asia / Japan / Kagoshima | Makoto Ozawa (Kagoshima University ) |
| EPI_ISL_174521 | A/mallard duck/Kagoshima/KU116/2015(H5N8) | A / H5N8 | Asia / Japan / Kagoshima | Makoto Ozawa (Kagoshima University ) |
| EPI_ISL_175335 | A/Guangzhou/39715/2014 | A / H5N6 | Asia / China | |
| EPI_ISL_175336 | A/duck/Guangzhou/41227/2014 | A / H5N6 | Asia / China | |
| EPI_ISL_175339 | A/chicken/BC/FAV8/2014 | A / H5N2 | North America / Canada | |
| EPI_ISL_175340 | A/chicken/BC/FAV9/2014 | A / H5N2 | North America / Canada | |
| EPI_ISL_175534 | A/MuteSwan/Sweden/SVA150311KU0277/SZ502/2015 | A / H5N8 | Europe / Sweden / Stockholms Lan / Stockholms Kommun / Strömkajen, Stockholm | Siamak Zohari (National Veterinary Institute) |
| EPI_ISL_175535 | A/MuteSwan/Sweden/SVA150313KU0141/SZ543/2015 | A / H5N8 | Europe / Sweden / Stockholms Lan / Stockholms Kommun / Ladugårdslandsviken, Stockholm | Siamak Zohari (National Veterinary Institute) |
| EPI_ISL_1760446 | A/whooper swan/Shaanxi/SXY26/2020 | A / H5N8 | Asia / China / Shaanxi Province | Hongliang Chai (Northeast Forestry University / College of Wildlife Resources) |
| EPI_ISL_1760447 | A/whooper swan/Shaanxi/SXY66/2020 | A / H5N8 | Asia / China / Shaanxi Province | Hongliang Chai (Northeast Forestry University / College of Wildlife Resources) |
| EPI_ISL_1760448 | A/common teal/Shaanxi/SXY1-1/2020 | A / H5N8 | Asia / China / Shaanxi Province | Hongliang Chai (Northeast Forestry University / College of Wildlife Resources) |
| EPI_ISL_1760450 | A/whooper swan/Shaanxi/SXY2-1/2020 | A / H5N8 | Asia / China / Shaanxi Province | Hongliang Chai (Northeast Forestry University / College of Wildlife Resources) |
| EPI_ISL_1760451 | A/whooper swan/Shanxi/SX16/2020 | A / H5N8 | Asia / China / Shanxi Province | Hongliang Chai (Northeast Forestry University / College of Wildlife Resources) |
| EPI_ISL_1760452 | A/whooper swan/Shanxi/SX31/2020 | A / H5N8 | Asia / China / Shanxi Province | Hongliang Chai (Northeast Forestry University / College of Wildlife Resources) |
| EPI_ISL_1760453 | A/whooper swan/Shanxi/SX56/2020 | A / H5N8 | Asia / China / Shanxi Province | Hongliang Chai (Northeast Forestry University / College of Wildlife Resources) |
| EPI_ISL_1760454 | A/whooper swan/Shanxi/SX106/2020 | A / H5N8 | Asia / China / Shanxi Province | Hongliang Chai (Northeast Forestry University / College of Wildlife Resources) |
| EPI_ISL_1760455 | A/whooper swan/Shanxi/SX116/2020 | A / H5N2 | Asia / China / Shanxi Province | Hongliang Chai (Northeast Forestry University / College of Wildlife Resources) |
| EPI_ISL_176840 | A/feline/Guangdong/1/2014(H5N6) | A / H5N6 | Asia / China / Guangdong Province | Yuhai Bi (Institute of Microbiology, Chinese Academy of Sciences) |
| EPI_ISL_176841 | A/feline/Guangdong/2/2014(H5N6) | A / H5N6 | Asia / China / Guangdong Province | Yuhai Bi (Institute of Microbiology, Chinese Academy of Sciences) |
| EPI_ISL_177455 | A/turkey/BC/FAV14/2014 | A / H5N2 | North America / Canada / British Columbia | Tamiko Hisanaga (Canadian Food Inspection Agency) |
| EPI_ISL_177584 | A/domestic duck/Hungary/7341/2015 | A / H5N8 | Europe / Hungary / Bekes megye / Füzesgyarmat | Adam Dan (Danam.Vet.Molbiol) |
| EPI_ISL_177647 | A/turkey/California/K1500169-1.2/2015 | A / H5N8 | North America / United States | |
| EPI_ISL_177648 | A/pheasant/Washington/3147-2/2015 | A / H5N2 | North America / United States | |
| EPI_ISL_177649 | A/eurasian wigeon/Netherlands/1/2015 | A / H5N8 | Europe / Netherlands / Provincie Noord-Holland / Gemeente Purmerend / Ilpendam | Josanne Verhagen (Erasmus Medical Center / Department of Virology) |
| EPI_ISL_177699 | A/duck/Vietnam/1152/2014 | A / H5N6 | Asia / Vietnam | |
| EPI_ISL_177700 | A/duck/Vietnam/1151/2014 | A / H5N6 | Asia / Vietnam | |
| EPI_ISL_177701 | A/duck/Vietnam/1507/2014 | A / H5N6 | Asia / Vietnam | |
| EPI_ISL_177702 | A/duck/Vietnam/1511/2014 | A / H5N6 | Asia / Vietnam | |
| EPI_ISL_177703 | A/duck/Vietnam/1434/2014 | A / H5N6 | Asia / Vietnam | |
| EPI_ISL_177704 | A/duck/Vietnam/1144/2014 | A / H5N6 | Asia / Vietnam | |
| EPI_ISL_177725 | A/baikal teal/Korea/1437/2014 | A / H5N8 | Asia / Korea, Republic of | |
| EPI_ISL_177726 | A/baikal teal/Korea/1441/2014 | A / H5N8 | Asia / Korea, Republic of | |
| EPI_ISL_177727 | A/baikal teal/Korea/1445/2014 | A / H5N8 | Asia / Korea, Republic of | |
| EPI_ISL_177728 | A/baikal teal/Korea/1446/2014 | A / H5N8 | Asia / Korea, Republic of | |
| EPI_ISL_177729 | A/baikal teal/Korea/1447/2014 | A / H5N8 | Asia / Korea, Republic of | |
| EPI_ISL_177730 | A/baikal teal/Korea/1448/2014 | A / H5N8 | Asia / Korea, Republic of | |
| EPI_ISL_177731 | A/baikal teal/Korea/1449/2014 | A / H5N8 | Asia / Korea, Republic of | |
| EPI_ISL_177732 | A/baikal teal/Korea/1452/2014 | A / H5N8 | Asia / Korea, Republic of | |
| EPI_ISL_177733 | A/baikal teal/Korea/1454/2014 | A / H5N8 | Asia / Korea, Republic of | |
| EPI_ISL_177734 | A/baikal teal/Korea/1456/2014 | A / H5N8 | Asia / Korea, Republic of | |
| EPI_ISL_177735 | A/baikal teal/Korea/1457/2014 | A / H5N8 | Asia / Korea, Republic of | |
| EPI_ISL_177736 | A/baikal teal/Korea/1458/2014 | A / H5N8 | Asia / Korea, Republic of | |
| EPI_ISL_177737 | A/baikal teal/Korea/2399/2014 | A / H5N8 | Asia / Korea, Republic of | |
| EPI_ISL_177738 | A/baikal teal/Korea/2402/2014 | A / H5N8 | Asia / Korea, Republic of | |
| EPI_ISL_177739 | A/baikal teal/Korea/2403/2014 | A / H5N8 | Asia / Korea, Republic of | |
| EPI_ISL_177740 | A/baikal teal/Korea/2406/2014 | A / H5N8 | Asia / Korea, Republic of | |
| EPI_ISL_177741 | A/baikal teal/Korea/2414/2014 | A / H5N8 | Asia / Korea, Republic of | |
| EPI_ISL_177742 | A/baikal teal/Korea/2416/2014 | A / H5N8 | Asia / Korea, Republic of | |
| EPI_ISL_177743 | A/baikal teal/Korea/2417/2014 | A / H5N8 | Asia / Korea, Republic of | |
| EPI_ISL_177761 | A/cat/Sichuan/SC18/2014 | A / H5N6 | Asia / China | |
| EPI_ISL_177762 | A/swan goose/Jilin/JL01/2014 | A / H5N6 | Asia / China | |
| EPI_ISL_177869 | A/chicken/TongHai/302/2014 | A / H5N1 | Asia / China | |
| EPI_ISL_177989 | A/Spotted dove/Vietnam/WBT191/2014 | A / H5N6 | Asia / Vietnam | |
| EPI_ISL_177990 | A/Black-crowned night heron/Vietnam/WBT198/2014 | A / H5N6 | Asia / Vietnam | |
| EPI_ISL_177991 | A/Little egret/Vietnam/WBT210/2014 | A / H5N6 | Asia / Vietnam | |
| EPI_ISL_177992 | A/Common moorhen/Vietnam/WBT226/2014 | A / H5N6 | Asia / Vietnam | |
| EPI_ISL_177993 | A/Chinese pond heron/Vietnam/WBT231/2014 | A / H5N6 | Asia / Vietnam | |
| EPI_ISL_178079 | A/poultry/BC/FAV15/2014 | A / H5N2 | North America / Canada / British Columbia | Tamiko Hisanaga (Canadian Food Inspection Agency) |

| | | | | |
|---|---|---|---|---|
| EPI_ISL_178080 | A/poultry/BC/FAV17/2014 | A / H5N2 | North America / Canada / British Columbia | Tamiko Hisanaga (Canadian Food Inspection Agency) |
| EPI_ISL_178081 | A/poultry/BC/FAV19/2014 | A / H5N2 | North America / Canada / British Columbia | Tamiko Hisanaga (Canadian Food Inspection Agency) |
| EPI_ISL_178082 | A/chicken/BC/FAV20/2014 | A / H5N2 | North America / Canada / British Columbia | Tamiko Hisanaga (Canadian Food Inspection Agency) |
| EPI_ISL_178083 | A/chicken/BC/FAV21/2014 | A / H5N2 | North America / Canada / British Columbia | Tamiko Hisanaga (Canadian Food Inspection Agency) |
| EPI_ISL_178084 | A/chicken/BC/FAV22/2014 | A / H5N2 | North America / Canada / British Columbia | Tamiko Hisanaga (Canadian Food Inspection Agency) |
| EPI_ISL_178085 | A/chicken/BC/FAV23/2014 | A / H5N2 | North America / Canada / British Columbia | Tamiko Hisanaga (Canadian Food Inspection Agency) |
| EPI_ISL_178086 | A/chicken/BC/FAV24/2014 | A / H5N2 | North America / Canada / British Columbia | Tamiko Hisanaga (Canadian Food Inspection Agency) |
| EPI_ISL_178087 | A/chicken/BC/FAV25/2014 | A / H5N2 | North America / Canada / British Columbia | Tamiko Hisanaga (Canadian Food Inspection Agency) |
| EPI_ISL_178249 | A/American wigeon/BC/050-31/2015 | A / H5N8 | North America / Canada / British Columbia | Tamiko Hisanaga (Canadian Food Inspection Agency) |
| EPI_ISL_178250 | A/Environment/Yunnan/DQ5/2015 | A / H5N6 | Asia / China / Yunnan | Lei Yang (WHO Chinese National Influenza Center / Virology Institute, Chinese CDC) |
| EPI_ISL_178251 | A/Environment/Yunnan/DQ8/2015 | A / H5N6 | Asia / China / Yunnan | Lei Yang (WHO Chinese National Influenza Center / Virology Institute, Chinese CDC) |
| EPI_ISL_178252 | A/Environment/Yunnan/DQ9/2015 | A / H5N6 | Asia / China / Yunnan | Lei Yang (WHO Chinese National Influenza Center / Virology Institute, Chinese CDC) |
| EPI_ISL_178253 | A/Environment/Yunnan/DQ16/2015 | A / H5N6 | Asia / China / Yunnan | Lei Yang (WHO Chinese National Influenza Center / Virology Institute, Chinese CDC) |
| EPI_ISL_178255 | A/Environment/Yunnan/DQ53/2015 | A / H5N6 | Asia / China / Yunnan | Lei Yang (WHO Chinese National Influenza Center / Virology Institute, Chinese CDC) |
| EPI_ISL_178261 | A/Yunnan/14563/2015 | A / H5N6 | Asia / China / Yunnan | Lei Yang (WHO Chinese National Influenza Center / Virology Institute, Chinese CDC) |
| EPI_ISL_178262 | A/Yunnan/14564/2015 | A / H5N6 | Asia / China / Yunnan | Lei Yang (WHO Chinese National Influenza Center / Virology Institute, Chinese CDC) |
| EPI_ISL_178263 | A/chicken/Iowa/04-20/2015 | A / H5N2 | North America / United States | |
| EPI_ISL_179023 | A/goose/Taiwan/a015/2015 | A / H5N8 | Asia / Taiwan | Yu-Pin Liu (Animal Health Research Institute) |
| EPI_ISL_179024 | A/duck/Taiwan/a043/2015 | A / H5N2 | Asia / Taiwan | Yu-Pin Liu (Animal Health Research Institute) |
| EPI_ISL_179025 | A/chicken/Taiwan/a174/2015 | A / H5N3 | Asia / Taiwan | Yu-Pin Liu (Animal Health Research Institute) |
| EPI_ISL_179026 | A/duck/Taiwan/a068/2015 | A / H5N8 | Asia / Taiwan | Yu-Pin Liu (Animal Health Research Institute) |
| EPI_ISL_179175 | A/muscovy duck/Quang Ninh/4c111/2013 | A / H5N6 | Asia / Vietnam | |
| EPI_ISL_179176 | A/muscovy duck/Quang Ninh/5c112/2013 | A / H5N6 | Asia / Vietnam | |
| EPI_ISL_179374 | A/duck/Nha Trang/75c131/2014 | A / H5N6 | Asia / Vietnam | |
| EPI_ISL_179395 | A/chicken/AnNing/1/2014 | A / H5N1 | Asia / China | |
| EPI_ISL_179396 | A/chicken/AnNing/4/2014 | A / H5N1 | Asia / China | |
| EPI_ISL_179397 | A/chicken/AnNing/6/2014 | A / H5N1 | Asia / China | |
| EPI_ISL_179398 | A/chicken/DaLi/302/2014 | A / H5N1 | Asia / China | |
| EPI_ISL_179399 | A/chicken/DaLi/502/2014 | A / H5N1 | Asia / China | |
| EPI_ISL_179400 | A/chicken/Tonghai/3/2014 | A / H5N1 | Asia / China | |
| EPI_ISL_179401 | A/chicken/Tonghai/5/2014 | A / H5N1 | Asia / China | |
| EPI_ISL_179402 | A/chicken/Tonghai/802/2014 | A / H5N1 | Asia / China | |
| EPI_ISL_179543 | A/turkey/Minnesota/9892-2/2015 | A / H5N2 | North America / United States | |
| EPI_ISL_179544 | A/chicken/Washington/3490-18/2015 | A / H5N2 | North America / United States | |
| EPI_ISL_179545 | A/chicken/Oregon/A01819044/2015 | A / H5N2 | North America / United States | |
| EPI_ISL_179546 | A/turkey/Minnesota/7172-1/2015 | A / H5N2 | North America / United States | |
| EPI_ISL_179547 | A/turkey/Missouri/7458-1/2015 | A / H5N2 | North America / United States | |
| EPI_ISL_179548 | A/turkey/Arkansas/7791-1/2015 | A / H5N2 | North America / United States | |
| EPI_ISL_179549 | A/chicken/Kansas/8395-3/2015 | A / H5N2 | North America / United States | |
| EPI_ISL_179550 | A/turkey/Minnesota/9845-4/2015 | A / H5N2 | North America / United States | |
| EPI_ISL_179634 | A/duck/Jiangxi/NCDZT1123/2014 | A / H5N6 | Asia / China | |
| EPI_ISL_179635 | A/duck/Jiangxi/NCDZT1126/2014 | A / H5N6 | Asia / China | |
| EPI_ISL_179644 | A/Anas_crecca/Hubei/Chenhu1623-5/2014 | A / H5N6 | Asia / China | |
| EPI_ISL_179645 | A/chicken/Sichuan/NCJPL1/2014 | A / H5N6 | Asia / China | |
| EPI_ISL_179646 | A/duck/Sichuan/NCJPL7/2014 | A / H5N6 | Asia / China | |
| EPI_ISL_179647 | A/duck/Sichuan/NCXJ15/2014 | A / H5N6 | Asia / China | |
| EPI_ISL_179648 | A/duck/Sichuan/NCXJ24/2014 | A / H5N6 | Asia / China | |
| EPI_ISL_179649 | A/duck/Sichuan/NCXJ16/2014 | A / H5N6 | Asia / China | |
| EPI_ISL_179650 | A/environment/Sichuan/NCLL1/2014 | A / H5N6 | Asia / China | |
| EPI_ISL_179651 | A/duck/Sichuan/NCXN10/2014 | A / H5N1 | Asia / China | |
| EPI_ISL_179652 | A/duck/Sichuan/NCXN11/2014 | A / H5N1 | Asia / China | |
| EPI_ISL_179653 | A/pigeon/Sichuan/NCXN29/2014 | A / H5N1 | Asia / China | |
| EPI_ISL_180495 | A/Environment/Jiangxi/20983/2013 | A / H5N1 | Asia / China / jiangxi | Lei Yang (WHO Chinese National Influenza Center / Virology Institute, Chinese CDC) |
| EPI_ISL_180499 | A/Environment/Jiangxi/23094/2013 | A / H5N6 | Asia / China / jiangxi | Lei Yang (WHO Chinese National Influenza Center / Virology Institute, Chinese CDC) |
| EPI_ISL_180649 | A/goose/Shandong/k1201/2009 | A / H5N1 | Asia / China | |
| EPI_ISL_180655 | A/goose/Yangzhou/ZG62/2011 | A / H5N5 | Asia / China | |
| EPI_ISL_180656 | A/goose/Yangzhou/ZG60/2009 | A / H5N5 | Asia / China | |
| EPI_ISL_181080 | A/common teal/Korea/KU-12/2015 | A / H5N8 | Asia / Korea, Republic of / Chungcheongnam-do / Galsangyo | Jung Hoon Kwon (Konkuk University / College of Veterinary Medicine) |
| EPI_ISL_181081 | A/mallard/Korea/KU3-2/2015 | A / H5N8 | Asia / Korea, Republic of / Gyeongsangnam-do / Junam reservoir | Jung Hoon Kwon (Konkuk University / College of Veterinary Medicine) |
| EPI_ISL_181082 | A/mallard/Korea/N15-99/2015 | A / H5N8 | Asia / Korea, Republic of / Seoul / Jungnangcheon | Jung Hoon Kwon (Konkuk University / College of Veterinary Medicine) |
| EPI_ISL_181083 | A/mandarin duck/Korea/K14-363-1/2014 | A / H5N8 | Asia / Korea, Republic of / Chungcheongnam-do / pungsecheon | Jung Hoon Kwon (Konkuk University / College of Veterinary Medicine) |
| EPI_ISL_181084 | A/mandarin duck/Korea/K14-366-1/2014 | A / H5N8 | Asia / Korea, Republic of / Chungcheongnam-do / pungsecheon | Jung Hoon Kwon (Konkuk University / College of Veterinary Medicine) |
| EPI_ISL_181085 | A/mandarin duck/Korea/K14-367-1/2014 | A / H5N8 | Asia / Korea, Republic of / Chungcheongnam-do / pungsecheon | Jung Hoon Kwon (Konkuk University / College of Veterinary Medicine) |
| EPI_ISL_181086 | A/greater white-fronted goose/Korea/K14-367-4/2014 | A / H5N8 | Asia / Korea, Republic of / Chungcheongnam-do / pungsecheon | Jung Hoon Kwon (Konkuk University / College of Veterinary Medicine) |
| EPI_ISL_181087 | A/greater white-fronted goose/Korea/K14-369-3/2014 | A / H5N8 | Asia / Korea, Republic of / Chungcheongnam-do / pungsecheon | Jung Hoon Kwon (Konkuk University / College of Veterinary Medicine) |
| EPI_ISL_181088 | A/greater white-fronted goose/Korea/K14-371-4/2014 | A / H5N8 | Asia / Korea, Republic of / Chungcheongnam-do / pungsecheon | Jung Hoon Kwon (Konkuk University / College of Veterinary Medicine) |
| EPI_ISL_181089 | A/greater white-fronted goose/Korea/K14-372-2/2014 | A / H5N8 | Asia / Korea, Republic of / Chungcheongnam-do / pungsecheon | Jung Hoon Kwon (Konkuk University / College of Veterinary Medicine) |
| EPI_ISL_181090 | A/greater white-fronted goose/Korea/K14-374-1/2014 | A / H5N8 | Asia / Korea, Republic of / Chungcheongnam-do / pungsecheon | Jung Hoon Kwon (Konkuk University / College of Veterinary Medicine) |
| EPI_ISL_181093 | A/eurasian wigeon/Netherlands/1/2014 | A / H5N8 | Europe / Netherlands | |
| EPI_ISL_181094 | A/eurasian wigeon/Netherlands/2/2014 | A / H5N8 | Europe / Netherlands | |
| EPI_ISL_181127 | A/duck/Vietnam/LBM751/2014 | A / H5N6 | Asia / Vietnam | |
| EPI_ISL_181128 | A/duck/Vietnam/LBM752/2014 | A / H5N6 | Asia / Vietnam | |

| | | | | |
|---|---|---|---|---|
| EPI_ISL_181129 | A/muscovy duck/Vietnam/LBM754/2014 | A / H5N6 | Asia / Vietnam | |
| EPI_ISL_181130 | A/muscovy duck/Vietnam/LBM755/2014 | A / H5N6 | Asia / Vietnam | |
| EPI_ISL_181131 | A/muscovy duck/Vietnam/LBM756/2014 | A / H5N6 | Asia / Vietnam | |
| EPI_ISL_181132 | A/muscovy duck/Vietnam/LBM757/2014 | A / H5N6 | Asia / Vietnam | |
| EPI_ISL_181133 | A/duck/Vietnam/LBM758/2014 | A / H5N6 | Asia / Vietnam | |
| EPI_ISL_181134 | A/duck/Vietnam/LBM759/2014 | A / H5N6 | Asia / Vietnam | |
| EPI_ISL_181135 | A/duck/Vietnam/LBM760/2014 | A / H5N6 | Asia / Vietnam | |
| EPI_ISL_181665 | A/chicken/Shenzhen/715/2013 | A / H5N6 | Asia / China | |
| EPI_ISL_181666 | A/chicken/Shenzhen/1061/2013 | A / H5N6 | Asia / China | |
| EPI_ISL_181667 | A/chicken/Shenzhen/2269/2013 | A / H5N6 | Asia / China | |
| EPI_ISL_181668 | A/chicken/Dongguan/2690/2013 | A / H5N6 | Asia / China | |
| EPI_ISL_181669 | A/chicken/Dongguan/3363/2013 | A / H5N6 | Asia / China | |
| EPI_ISL_182059 | A/chicken/Shenzhen/433/2013 | A / H5N6 | Asia / China | |
| EPI_ISL_182060 | A/chicken/Shenzhen/1395/2013 | A / H5N6 | Asia / China | |
| EPI_ISL_182061 | A/chicken/Shenzhen/1845/2013 | A / H5N6 | Asia / China | |
| EPI_ISL_182062 | A/chicken/Shenzhen/2396/2013 | A / H5N6 | Asia / China | |
| EPI_ISL_182063 | A/chicken/Shenzhen/2464/2013 | A / H5N6 | Asia / China | |
| EPI_ISL_182064 | A/duck/Dongguan/2685/2013 | A / H5N6 | Asia / China | |
| EPI_ISL_182065 | A/silkie chicken/Dongguan/2809/2013 | A / H5N6 | Asia / China | |
| EPI_ISL_182066 | A/duck/Dongguan/3069/2013 | A / H5N6 | Asia / China | |
| EPI_ISL_182067 | A/chicken/Dongguan/4259/2013 | A / H5N6 | Asia / China | |
| EPI_ISL_182068 | A/duck/Jiangxi/10160/2014 | A / H5N6 | Asia / China | |
| EPI_ISL_182069 | A/environment/Jiangxi/10164/2014 | A / H5N6 | Asia / China | |
| EPI_ISL_182070 | A/environment/Jiangxi/10171/2014 | A / H5N6 | Asia / China | |
| EPI_ISL_182086 | A/chicken/Shenzhen/552/2013 | A / H5N6 | Asia / China | |
| EPI_ISL_1822592 | A/whooper swan/Shanxi/SX166/2020 | A / H5N8 | Asia / China / Shanxi Province | Hongliang Chai (Northeast Forestry University / College of Wildlife Resources) |
| EPI_ISL_1822593 | A/whooper swan/Shanxi/SX206/2020 | A / H5N8 | Asia / China / Shanxi Province | Hongliang Chai (Northeast Forestry University / College of Wildlife Resources) |
| EPI_ISL_1822594 | A/whooper swan/Shanxi/SX216/2020 | A / H5N8 | Asia / China / Shanxi Province | Hongliang Chai (Northeast Forestry University / College of Wildlife Resources) |
| EPI_ISL_1822595 | A/whooper swan/Shanxi/SX231/2020 | A / H5N8 | Asia / China / Shanxi Province | Hongliang Chai (Northeast Forestry University / College of Wildlife Resources) |
| EPI_ISL_1822596 | A/whooper swan/Shanxi/SX251/2020 | A / H5N8 | Asia / China / Shanxi Province | Hongliang Chai (Northeast Forestry University / College of Wildlife Resources) |
| EPI_ISL_1822597 | A/whooper swan/Shanxi/SX276/2020 | A / H5N8 | Asia / China / Shanxi Province | Hongliang Chai (Northeast Forestry University / College of Wildlife Resources) |
| EPI_ISL_1822598 | A/whooper swan/Shanxi/SX291/2020 | A / H5N8 | Asia / China / Shanxi Province | Hongliang Chai (Northeast Forestry University / College of Wildlife Resources) |
| EPI_ISL_1822599 | A/whooper swan/Shanxi/SX346/2020 | A / H5N8 | Asia / China / Shanxi Province | Hongliang Chai (Northeast Forestry University / College of Wildlife Resources) |
| EPI_ISL_1829205 | A/whooper swan/Henan/SMQ5/2020 | A / H5N8 | Asia / China / Henan Province | Hongliang Chai (Northeast Forestry University / College of Wildlife Resources) |
| EPI_ISL_1834185 | A/whooper swan/Henan/SMQ6/2020 | A / H5N8 | Asia / China / Henan Province | Hongliang Chai (Northeast Forestry University / College of Wildlife Resources) |
| EPI_ISL_190456 | A/turkey/Iowa/11762-1/2015 | A / H5N2 | North America / United States | |
| EPI_ISL_190457 | A/turkey/Iowa/13541-1/2015 | A / H5N2 | North America / United States | |
| EPI_ISL_190458 | A/chicken/Iowa/13542-2/2015 | A / H5N2 | North America / United States | |
| EPI_ISL_190459 | A/turkey/Iowa/14319-1/2015 | A / H5N2 | North America / United States | |
| EPI_ISL_190460 | A/turkey/Iowa/14318-1/2015 | A / H5N2 | North America / United States | |
| EPI_ISL_190461 | A/chicken/Iowa/14322-6/2015 | A / H5N2 | North America / United States | |
| EPI_ISL_190462 | A/chicken/Iowa/14399-4/2015 | A / H5N2 | North America / United States | |
| EPI_ISL_190464 | A/chicken/Iowa/14589-1/2015 | A / H5N2 | North America / United States | |
| EPI_ISL_191765 | A/scarlet_ibis/Germany/AR44-L01279/2015 | A / H5N8 | Europe / Germany / Mecklenburg-Vorpommern / zoo | Anne Pohlmann (Friedrich-Loeffler-Institut) |
| EPI_ISL_1937883 | A/whooper swan/Henan/SMQ7/2020 | A / H5N8 | Asia / China / Henan Province | Hongliang Chai (Northeast Forestry University / College of Wildlife Resources) |
| EPI_ISL_1937884 | A/whooper swan/Henan/SM1/2020 | A / H5N8 | Asia / China / Henan Province | Hongliang Chai (Northeast Forestry University / College of Wildlife Resources) |
| EPI_ISL_1937889 | A/whooper swan/Henan/SM16/2020 | A / H5N8 | Asia / China / Henan Province | Hongliang Chai (Northeast Forestry University / College of Wildlife Resources) |
| EPI_ISL_1937890 | A/whooper swan/Henan/SM31/2020 | A / H5N8 | Asia / China / Henan Province | Hongliang Chai (Northeast Forestry University / College of Wildlife Resources) |
| EPI_ISL_1937891 | A/whooper swan/Henan/SM61/2020 | A / H5N8 | Asia / China / Henan Province | Hongliang Chai (Northeast Forestry University / College of Wildlife Resources) |
| EPI_ISL_1937892 | A/whooper swan/Henan/SM76/2020 | A / H5N8 | Asia / China / Henan Province | Hongliang Chai (Northeast Forestry University / College of Wildlife Resources) |
| EPI_ISL_1938307 | A/whooper swan/Henan/SM86/2020 | A / H5N8 | Asia / China / Henan Province | Hongliang Chai (Northeast Forestry University / College of Wildlife Resources) |
| EPI_ISL_1938309 | A/whooper swan/Henan/SM111/2020 | A / H5N8 | Asia / China / Henan Province | Hongliang Chai (Northeast Forestry University / College of Wildlife Resources) |
| EPI_ISL_1938478 | A/whooper swan/Henan/SMQ9/2020 | A / H5N8 | Asia / China / Henan Province | Hongliang Chai (Northeast Forestry University / College of Wildlife Resources) |
| EPI_ISL_1939080 | A/whooper swan/Henan/SMQ10/2020 | A / H5N8 | Asia / China / Henan Province | Hongliang Chai (Northeast Forestry University / College of Wildlife Resources) |
| EPI_ISL_1939617 | A/eurasian eagle-owl/Henan/SMQ11/2020 | A / H5N8 | Asia / China / Henan Province | Hongliang Chai (Northeast Forestry University / College of Wildlife Resources) |
| EPI_ISL_1941351 | A/duck/Czech Republic/7682-9/2021 | A / H5N8 | Europe / Czech Republic / Kralovehradecky Kraj / Okres Hradec Kralove / Nove Mesto | Alexander Nagy (State Veterinary Institute Prague) |
| EPI_ISL_1941365 | A/duck/Czech Republic/7682-5/2021 | A / H5N8 | Europe / Czech Republic / Kralovehradecky Kraj / Okres Hradec Kralove / Nove Mesto | Alexander Nagy (State Veterinary Institute Prague) |
| EPI_ISL_1941375 | A/duck/Czech Republic/7682-2/2021 | A / H5N8 | Europe / Czech Republic / Kralovehradecky Kraj / Okres Hradec Kralove / Nove Mesto | Alexander Nagy (State Veterinary Institute Prague) |
| EPI_ISL_1941417 | A/duck/Czech Republic/7681-8/2021 | A / H5N8 | Europe / Czech Republic / Kralovehradecky Kraj / Okres Hradec Kralove / Klamos | Alexander Nagy (State Veterinary Institute Prague) |
| EPI_ISL_1941444 | A/duck/Czech Republic/7681-7/2021 | A / H5N8 | Europe / Czech Republic / Kralovehradecky Kraj / Okres Hradec Kralove / Klamos | Alexander Nagy (State Veterinary Institute Prague) |
| EPI_ISL_1941480 | A/duck/Czech Republic/7681-5/2021 | A / H5N8 | Europe / Czech Republic / Kralovehradecky Kraj / Okres Hradec Kralove / Klamos | Alexander Nagy (State Veterinary Institute Prague) |
| EPI_ISL_1941542 | A/duck/Czech Republic/7681-3/2021 | A / H5N8 | Europe / Czech Republic / Kralovehradecky Kraj / Okres Hradec Kralove / Klamos | Alexander Nagy (State Veterinary Institute Prague) |
| EPI_ISL_1941578 | A/duck/Czech Republic/7681-10/2021 | A / H5N8 | Europe / Czech Republic / Kralovehradecky Kraj / Okres Hradec Kralove / Klamos | Alexander Nagy (State Veterinary Institute Prague) |
| EPI_ISL_1941579 | A/duck/Czech Republic/6653-5/2021 | A / H5N8 | Europe / Czech Republic / Kralovehradecky Kraj / Okres Hradec Kralove / Stary Bydzov | Alexander Nagy (State Veterinary Institute Prague) |
| EPI_ISL_1941580 | A/duck/Czech Republic/6653-15/2021 | A / H5N8 | Europe / Czech Republic / Kralovehradecky Kraj / Okres Hradec Kralove / Stary Bydzov | Alexander Nagy (State Veterinary Institute Prague) |
| EPI_ISL_1941581 | A/goose/Czech Republic/5363-10/2021 | A / H5N8 | Europe / Czech Republic / Plzensky Kraj / Okres Plzen-Jih / Zbuch, pond; GPS: 49.6897778N, 13.2 | Alexander Nagy (State Veterinary Institute Prague) |
| EPI_ISL_1941582 | A/chicken/Czech Republic/7100/2021 | A / H5N8 | Europe / Czech Republic / Jihocesky Kraj / Okres Strakonice / Vodnany | Alexander Nagy (State Veterinary Institute Prague) |
| EPI_ISL_195294 | A/Yunnan/0127/2015 | A / H5N6 | Asia / China | |
| EPI_ISL_195308 | A/chicken/Yunnan/19/2015 | A / H5N6 | Asia / China | |
| EPI_ISL_195309 | A/chicken/Yunnan/22/2015 | A / H5N6 | Asia / China | |
| EPI_ISL_195310 | A/chicken/Yunnan/25A/2015 | A / H5N6 | Asia / China | |
| EPI_ISL_195769 | A/Environment/Guangdong/GZ55/2013(H5N8) | A / H5N8 | Asia / China / Guangdong Province | Xianqiao Zeng (Guangdong Provincial Center for Disease Control and Prevention) |

| | | | | |
|---|---|---|---|---|
| EPI_ISL_196100 | A/swine/Guangdong/1/2014 | A / H5N6 | Asia / China | |
| EPI_ISL_196101 | A/swine/Guangdong/2/2014 | A / H5N6 | Asia / China | |
| EPI_ISL_196571 | A/duck/Jiangsu/WX156/2013 | A / H5N6 | Asia / China / Jiangsu Provice | Maijuan Ma (Beijing Institute of Microbiology and Epidemiology / State Key Laboratory of Pathogen and Biosecurity) |
| EPI_ISL_196572 | A/environment/Jiangsu/WX175/2013 | A / H5N6 | Asia / China / Jiangsu Provice | Maijuan Ma (Beijing Institute of Microbiology and Epidemiology / State Key Laboratory of Pathogen and Biosecurity) |
| EPI_ISL_196573 | A/chicken/Jiangsu/WXS7393/2014 | A / H5N6 | Asia / China / Jiangsu Provice | Maijuan Ma (Beijing Institute of Microbiology and Epidemiology / State Key Laboratory of Pathogen and Biosecurity) |
| EPI_ISL_196591 | A/chicken/Jiangsu/WXBING2/2014 | A / H5N6 | Asia / China / Jiangsu Provice | Maijuan Ma (Beijing Institute of Microbiology and Epidemiology / State Key Laboratory of Pathogen and Biosecurity) |
| EPI_ISL_196592 | A/goose/Jiangsu/WX202/2014 | A / H5N8 | Asia / China / Jiangsu Provice | Maijuan Ma (Beijing Institute of Microbiology and Epidemiology / State Key Laboratory of Pathogen and Biosecurity) |
| EPI_ISL_198663 | A/turkey/Germany/AR3382-L00937/2014 | A / H5N8 | Europe / Germany / Lower Saxony | Anne Pohlmann (Friedrich-Loeffler-Institut) |
| EPI_ISL_198665 | A/turkey/Germany/AR3390-L00939/2014 | A / H5N8 | Europe / Germany / Lower Saxony | Anne Pohlmann (Friedrich-Loeffler-Institut) |
| EPI_ISL_198754 | A/duck/Hunan/12.17 YYFQH012-O/2014(H5N6) | A / H5N6 | Asia / China | Yuhai Bi (Institute of Microbiology, Chinese Academy of Sciences) |
| EPI_ISL_198757 | A/goose/Jilin/04.04 SY003-O/2015(H5N6) | A / H5N6 | Asia / China | Yuhai Bi (Institute of Microbiology, Chinese Academy of Sciences) |
| EPI_ISL_198758 | A/chicken/Jiangxi/04.01 NCJD131-1-O/2015(H5N6) | A / H5N6 | Asia / China | Yuhai Bi (Institute of Microbiology, Chinese Academy of Sciences) |
| EPI_ISL_198761 | A/chicken/Hunan/12.07 YYGK114-P/2013(H5N6) | A / H5N6 | Asia / China | Yuhai Bi (Institute of Microbiology, Chinese Academy of Sciences) |
| EPI_ISL_198762 | A/chicken/Hunan/12.28 YYGK034-P/2013(H5N6) | A / H5N6 | Asia / China | Yuhai Bi (Institute of Microbiology, Chinese Academy of Sciences) |
| EPI_ISL_198765 | A/duck/Hunan/12.07 YYGK108-P/2013(H5N6) | A / H5N6 | Asia / China | Yuhai Bi (Institute of Microbiology, Chinese Academy of Sciences) |
| EPI_ISL_198766 | A/duck/Hunan/12.07 YYGK110-P/2013(H5N6) | A / H5N6 | Asia / China | Yuhai Bi (Institute of Microbiology, Chinese Academy of Sciences) |
| EPI_ISL_198767 | A/duck/Hunan/12.07 YYGK111-P/2013(H5N6) | A / H5N6 | Asia / China | Yuhai Bi (Institute of Microbiology, Chinese Academy of Sciences) |
| EPI_ISL_198768 | A/duck/Hunan/12.07 YYGK112-P/2013(H5N6) | A / H5N6 | Asia / China | Yuhai Bi (Institute of Microbiology, Chinese Academy of Sciences) |
| EPI_ISL_198769 | A/duck/Hunan/12.07 YYGK113-P/2013(H5N6) | A / H5N6 | Asia / China | Yuhai Bi (Institute of Microbiology, Chinese Academy of Sciences) |
| EPI_ISL_198770 | A/duck/Hunan/12.07 YYGK118-P/2013(H5N6) | A / H5N6 | Asia / China | Yuhai Bi (Institute of Microbiology, Chinese Academy of Sciences) |
| EPI_ISL_198771 | A/duck/Hunan/12.07 YYGK032-P/2013(H5N6) | A / H5N6 | Asia / China | Yuhai Bi (Institute of Microbiology, Chinese Academy of Sciences) |
| EPI_ISL_198772 | A/duck/Hunan/12.07 YYGK0043-P/2013(H5N6) | A / H5N6 | Asia / China | Yuhai Bi (Institute of Microbiology, Chinese Academy of Sciences) |
| EPI_ISL_198773 | A/duck/Hunan/12.07 YYGK68-P/2013(H5N6) | A / H5N6 | Asia / China | Yuhai Bi (Institute of Microbiology, Chinese Academy of Sciences) |
| EPI_ISL_198777 | A/duck/Hunan/12.07 YGK098-P/2013(H5N6) | A / H5N6 | Asia / China | Yuhai Bi (Institute of Microbiology, Chinese Academy of Sciences) |
| EPI_ISL_198781 | A/chicken/Hunan/02.26 YYGKJ281-P/2014(H5N6) | A / H5N6 | Asia / China | Yuhai Bi (Institute of Microbiology, Chinese Academy of Sciences) |
| EPI_ISL_198782 | A/chicken/Hunan/02.26 YYGKJ295-P/2014(H5N6) | A / H5N6 | Asia / China | Yuhai Bi (Institute of Microbiology, Chinese Academy of Sciences) |
| EPI_ISL_198783 | A/duck/Hunan/01.16 YYFQH185-P/2014(H5N6) | A / H5N6 | Asia / China | Yuhai Bi (Institute of Microbiology, Chinese Academy of Sciences) |
| EPI_ISL_198784 | A/duck/Hunan/01.16 YYGK0204-P/2014(H5N6) | A / H5N6 | Asia / China | Yuhai Bi (Institute of Microbiology, Chinese Academy of Sciences) |
| EPI_ISL_198785 | A/duck/Hunan/01.16 YYGK218-P/2014(H5N6) | A / H5N6 | Asia / China | Yuhai Bi (Institute of Microbiology, Chinese Academy of Sciences) |
| EPI_ISL_198789 | A/duck/Hunan/02.26 YYGK249-P/2014(H5N6) | A / H5N6 | Asia / China | Yuhai Bi (Institute of Microbiology, Chinese Academy of Sciences) |
| EPI_ISL_198791 | A/duck/Hunan/02.26 YYGK250-P/2014(H5N6) | A / H5N6 | Asia / China | Yuhai Bi (Institute of Microbiology, Chinese Academy of Sciences) |
| EPI_ISL_198792 | A/duck/Hunan/02.26 YYGK251-P/2014(H5N6) | A / H5N6 | Asia / China | Yuhai Bi (Institute of Microbiology, Chinese Academy of Sciences) |
| EPI_ISL_198793 | A/duck/Hunan/02.26 YYGK254-P/2014(H5N6) | A / H5N6 | Asia / China | Yuhai Bi (Institute of Microbiology, Chinese Academy of Sciences) |
| EPI_ISL_198794 | A/duck/Hunan/02.26 YYGK0255-P/2014(H5N6) | A / H5N6 | Asia / China | Yuhai Bi (Institute of Microbiology, Chinese Academy of Sciences) |
| EPI_ISL_198795 | A/duck/Hunan/02.26 YYGK263-P/2014(H5N6) | A / H5N6 | Asia / China | Yuhai Bi (Institute of Microbiology, Chinese Academy of Sciences) |
| EPI_ISL_198796 | A/duck/Hunan/02.26 YYGK265-P/2014(H5N6) | A / H5N6 | Asia / China | Yuhai Bi (Institute of Microbiology, Chinese Academy of Sciences) |
| EPI_ISL_198797 | A/duck/Hunan/02.26 YYGK266-P/2014(H5N6) | A / H5N6 | Asia / China | Yuhai Bi (Institute of Microbiology, Chinese Academy of Sciences) |
| EPI_ISL_198798 | A/environment/Jiangxi/02.10 NCDZT023/2015(H5N6) | A / H5N6 | Asia / China | Yuhai Bi (Institute of Microbiology, Chinese Academy of Sciences) |
| EPI_ISL_198799 | A/Pigeon/Jiangxi/02.10 NCDZT0017-P/2015(H5N6) | A / H5N6 | Asia / China | Yuhai Bi (Institute of Microbiology, Chinese Academy of Sciences) |
| EPI_ISL_198800 | A/environment/Jiangxi/02.10 NCDZT009/2015(H5N6) | A / H5N6 | Asia / China | Yuhai Bi (Institute of Microbiology, Chinese Academy of Sciences) |
| EPI_ISL_198801 | A/duck/Hunan/02.26 YYGK267-P/2014(H5N6) | A / H5N6 | Asia / China | Yuhai Bi (Institute of Microbiology, Chinese Academy of Sciences) |
| EPI_ISL_198802 | A/duck/Hunan/02.26 YYGK271-P/2014(H5N6) | A / H5N6 | Asia / China | Yuhai Bi (Institute of Microbiology, Chinese Academy of Sciences) |
| EPI_ISL_198803 | A/duck/Hunan/02.26 YYGK275/2014(H5N6) | A / H5N6 | Asia / China | Yuhai Bi (Institute of Microbiology, Chinese Academy of Sciences) |
| EPI_ISL_198804 | A/duck/Hunan/02.26 YYGK276-P/2014(H5N6) | A / H5N6 | Asia / China | Yuhai Bi (Institute of Microbiology, Chinese Academy of Sciences) |
| EPI_ISL_198805 | A/duck/Hunan/02.16 YYGK281-P/2014(H5N6) | A / H5N6 | Asia / China | Yuhai Bi (Institute of Microbiology, Chinese Academy of Sciences) |
| EPI_ISL_198806 | A/duck/Hunan/02.26 YYGK283-P/2014(H5N6) | A / H5N6 | Asia / China | Yuhai Bi (Institute of Microbiology, Chinese Academy of Sciences) |
| EPI_ISL_198807 | A/duck/Hunan/02.26 YYGK286-P/2014(H5N6) | A / H5N6 | Asia / China | Yuhai Bi (Institute of Microbiology, Chinese Academy of Sciences) |
| EPI_ISL_198808 | A/duck/Hunan/02.26 YYGK289-P/2014(H5N6) | A / H5N6 | Asia / China | Yuhai Bi (Institute of Microbiology, Chinese Academy of Sciences) |
| EPI_ISL_198809 | A/duck/Hunan/02.26 YYFQH290-P/2014(H5N6) | A / H5N6 | Asia / China | Yuhai Bi (Institute of Microbiology, Chinese Academy of Sciences) |
| EPI_ISL_198810 | A/duck/Hunan/02.26 YYFQHY291-P/2014(H5N6) | A / H5N6 | Asia / China | Yuhai Bi (Institute of Microbiology, Chinese Academy of Sciences) |
| EPI_ISL_198811 | A/duck/Hunan/02.26 YYFQH293-P/2014(H5N6) | A / H5N6 | Asia / China | Yuhai Bi (Institute of Microbiology, Chinese Academy of Sciences) |
| EPI_ISL_198812 | A/duck/Hunan/02.26 YYFQH294-P/2014(H5N6) | A / H5N6 | Asia / China | Yuhai Bi (Institute of Microbiology, Chinese Academy of Sciences) |
| EPI_ISL_198813 | A/duck/Hunan/02.26 YYGQH295-P/2014(H5N6) | A / H5N6 | Asia / China | Yuhai Bi (Institute of Microbiology, Chinese Academy of Sciences) |
| EPI_ISL_198814 | A/duck/Hunan/02.26 YYFQH297-P/2014(H5N6) | A / H5N6 | Asia / China | Yuhai Bi (Institute of Microbiology, Chinese Academy of Sciences) |
| EPI_ISL_198815 | A/duck/Hunan/02.26 YYGKY298-P/2014(H5N6) | A / H5N6 | Asia / China | Yuhai Bi (Institute of Microbiology, Chinese Academy of Sciences) |
| EPI_ISL_198816 | A/duck/Hunan/01.16 YYFQH299-P/2014(H5N6) | A / H5N6 | Asia / China | Yuhai Bi (Institute of Microbiology, Chinese Academy of Sciences) |
| EPI_ISL_198817 | A/duck/Hunan/01.16 YYFQH300-P/2014(H5N6) | A / H5N6 | Asia / China | Yuhai Bi (Institute of Microbiology, Chinese Academy of Sciences) |
| EPI_ISL_198818 | A/duck/Hunan/01.16 YYFQH302-P/2014(H5N6) | A / H5N6 | Asia / China | Yuhai Bi (Institute of Microbiology, Chinese Academy of Sciences) |
| EPI_ISL_198819 | A/duck/Hunan/01.16 YYFQH303-P/2014(H5N6) | A / H5N6 | Asia / China | Yuhai Bi (Institute of Microbiology, Chinese Academy of Sciences) |
| EPI_ISL_198820 | A/duck/Hunan/01.16 YYFQH304-P/2014(H5N6) | A / H5N6 | Asia / China | Yuhai Bi (Institute of Microbiology, Chinese Academy of Sciences) |
| EPI_ISL_198821 | A/duck/Hunan/01.16 YYFQH305-P/2014(H5N6) | A / H5N6 | Asia / China | Yuhai Bi (Institute of Microbiology, Chinese Academy of Sciences) |
| EPI_ISL_198822 | A/duck/Hunan/01.16 YYFQH306-P/2014(H5N6) | A / H5N6 | Asia / China | Yuhai Bi (Institute of Microbiology, Chinese Academy of Sciences) |
| EPI_ISL_198823 | A/duck/Hunan/01.16 YYFQH307-P/2014(H5N6) | A / H5N6 | Asia / China | Yuhai Bi (Institute of Microbiology, Chinese Academy of Sciences) |
| EPI_ISL_198824 | A/duck/Hunan/02.26 YYFQH308-P/2014(H5N6) | A / H5N6 | Asia / China | Yuhai Bi (Institute of Microbiology, Chinese Academy of Sciences) |
| EPI_ISL_198825 | A/duck/Hunan/01.16 YYFQH309-P/2014(H5N6) | A / H5N6 | Asia / China | Yuhai Bi (Institute of Microbiology, Chinese Academy of Sciences) |
| EPI_ISL_198826 | A/duck/Hunan/01.16 YYFQH310-P/2014(H5N6) | A / H5N6 | Asia / China | Yuhai Bi (Institute of Microbiology, Chinese Academy of Sciences) |
| EPI_ISL_198827 | A/duck/Hunan/02.26 YYFQH311-P/2014(H5N6) | A / H5N6 | Asia / China | Yuhai Bi (Institute of Microbiology, Chinese Academy of Sciences) |
| EPI_ISL_198828 | A/duck/Hunan/01.16 YYFQH315-P/2014(H5N6) | A / H5N6 | Asia / China | Yuhai Bi (Institute of Microbiology, Chinese Academy of Sciences) |
| EPI_ISL_198829 | A/duck/Hunan/01.16 YYFQH317-P/2014(H5N6) | A / H5N6 | Asia / China | Yuhai Bi (Institute of Microbiology, Chinese Academy of Sciences) |
| EPI_ISL_198830 | A/duck/Hunan/01.16 YYFQH318-P/2014(H5N6) | A / H5N6 | Asia / China | Yuhai Bi (Institute of Microbiology, Chinese Academy of Sciences) |
| EPI_ISL_198831 | A/duck/Hunan/01.16 YYFQH319-P/2014(H5N6) | A / H5N6 | Asia / China | Yuhai Bi (Institute of Microbiology, Chinese Academy of Sciences) |
| EPI_ISL_198832 | A/duck/Hunan/02.26 YYFQH320-P/2014(H5N6) | A / H5N6 | Asia / China | Yuhai Bi (Institute of Microbiology, Chinese Academy of Sciences) |

| | | | | |
|---|---|---|---|---|
| EPI_ISL_198833 | A/duck/Hunan/12.17 YYGK014/2014(H5N6) | A / H5N6 | Asia / China | Yuhai Bi (Institute of Microbiology, Chinese Academy of Sciences) |
| EPI_ISL_198835 | A/environment/Jiangxi/02.10 NCDZT0018/2015(H5N6) | A / H5N6 | Asia / China | Yuhai Bi (Institute of Microbiology, Chinese Academy of Sciences) |
| EPI_ISL_198836 | A/environment/Jiangxi/02.10 NCDZT018/2015(H5N6) | A / H5N6 | Asia / China | Yuhai Bi (Institute of Microbiology, Chinese Academy of Sciences) |
| EPI_ISL_198838 | A/environment/JiangXi/02.10 NCNP0019/2015(H5N6) | A / H5N6 | Asia / China | Yuhai Bi (Institute of Microbiology, Chinese Academy of Sciences) |
| EPI_ISL_198849 | A/pigeon/Jilin/04.25 CCHL018-O/2015(H5N6) | A / H5N6 | Asia / China | Yuhai Bi (Institute of Microbiology, Chinese Academy of Sciences) |
| EPI_ISL_198856 | A/chicken/Jiangxi/04.01 NCDZT0040-O/2015(H5N6) | A / H5N6 | Asia / China | Yuhai Bi (Institute of Microbiology, Chinese Academy of Sciences) |
| EPI_ISL_198857 | A/duck/Hunan/12.17 YYGK040/2014(H5N6) | A / H5N6 | Asia / China | Yuhai Bi (Institute of Microbiology, Chinese Academy of Sciences) |
| EPI_ISL_198860 | A/duck/Hunan/12.17 YYFQH052-P/2014(H5N6) | A / H5N6 | Asia / China | Yuhai Bi (Institute of Microbiology, Chinese Academy of Sciences) |
| EPI_ISL_198861 | A/chicken/Hunan/04.14 YYKG0533-Z-O/2015(H5N6) | A / H5N6 | Asia / China | Yuhai Bi (Institute of Microbiology, Chinese Academy of Sciences) |
| EPI_ISL_198862 | A/duck/Hunan/12.17 YYGK056/2014(H5N6) | A / H5N6 | Asia / China | Yuhai Bi (Institute of Microbiology, Chinese Academy of Sciences) |
| EPI_ISL_198863 | A/duck/Hunan/12.17 YYGK0057/2014(H5N6) | A / H5N6 | Asia / China | Yuhai Bi (Institute of Microbiology, Chinese Academy of Sciences) |
| EPI_ISL_198866 | A/duck/Sichuan/04.27 NBXJ717/2014(H5N6) | A / H5N6 | Asia / China | Yuhai Bi (Institute of Microbiology, Chinese Academy of Sciences) |
| EPI_ISL_198867 | A/duck/Sichuan/04.27 NBXJ718/2014(H5N6) | A / H5N6 | Asia / China | Yuhai Bi (Institute of Microbiology, Chinese Academy of Sciences) |
| EPI_ISL_198868 | A/duck/Sichuan/04.27 NBXJ71/2014(H5N6) | A / H5N6 | Asia / China | Yuhai Bi (Institute of Microbiology, Chinese Academy of Sciences) |
| EPI_ISL_198869 | A/duck/Sichuan/04.27 NBXJ720/2014(H5N6) | A / H5N6 | Asia / China | Yuhai Bi (Institute of Microbiology, Chinese Academy of Sciences) |
| EPI_ISL_198870 | A/duck/Sichuan/04.27 NBXJ731/2014(H5N6) | A / H5N6 | Asia / China | Yuhai Bi (Institute of Microbiology, Chinese Academy of Sciences) |
| EPI_ISL_198871 | A/duck/Sichuan/04.27 NCXJ073/2014(H5N6) | A / H5N6 | Asia / China | Yuhai Bi (Institute of Microbiology, Chinese Academy of Sciences) |
| EPI_ISL_198872 | A/duck/Sichuan/04.27 NBXJ79/2014(H5N6) | A / H5N6 | Asia / China | Yuhai Bi (Institute of Microbiology, Chinese Academy of Sciences) |
| EPI_ISL_198873 | A/duck/Hunan/12.17 YYGK079-1/2014(H5N6) | A / H5N6 | Asia / China | Yuhai Bi (Institute of Microbiology, Chinese Academy of Sciences) |
| EPI_ISL_198874 | A/duck/Sichuan/04.27 NBXJ713/2014(H5N6) | A / H5N6 | Asia / China | Yuhai Bi (Institute of Microbiology, Chinese Academy of Sciences) |
| EPI_ISL_198875 | A/environment/Jilin/04.11 CCHSL012/2015(H5N6) | A / H5N6 | Asia / China | Yuhai Bi (Institute of Microbiology, Chinese Academy of Sciences) |
| EPI_ISL_198876 | A/duck/Hunan/12.17 YYGK082/2014(H5N6) | A / H5N6 | Asia / China | Yuhai Bi (Institute of Microbiology, Chinese Academy of Sciences) |
| EPI_ISL_198878 | A/environment/Jilin/04.25 NA004/2015(H5N6) | A / H5N6 | Asia / China | Yuhai Bi (Institute of Microbiology, Chinese Academy of Sciences) |
| EPI_ISL_198880 | A/chicken/Yunnan/03.16 DQXYL0034-O/2015(H5N6) | A / H5N6 | Asia / China | Yuhai Bi (Institute of Microbiology, Chinese Academy of Sciences) |
| EPI_ISL_198881 | A/chicken/Yunnan/03.16 DQJT0048-O/2015(H5N6) | A / H5N6 | Asia / China | Yuhai Bi (Institute of Microbiology, Chinese Academy of Sciences) |
| EPI_ISL_198882 | A/duck/Yunnan/03.16 DQXYL0039-O/2015(H5N6) | A / H5N6 | Asia / China | Yuhai Bi (Institute of Microbiology, Chinese Academy of Sciences) |
| EPI_ISL_198883 | A/duck/Yunnan/03.15 DQWGH005-O/2015(H5N6) | A / H5N6 | Asia / China | Yuhai Bi (Institute of Microbiology, Chinese Academy of Sciences) |
| EPI_ISL_198884 | A/chicken/Jiangxi/12.05 NCDZT0012/2014(H5N6) | A / H5N6 | Asia / China | Yuhai Bi (Institute of Microbiology, Chinese Academy of Sciences) |
| EPI_ISL_198885 | A/chicken/Jiangxi/12.05 NCDZT0020/2014(H5N6) | A / H5N6 | Asia / China | Yuhai Bi (Institute of Microbiology, Chinese Academy of Sciences) |
| EPI_ISL_198886 | A/chicken/Jiangxi/12.05 NCDZT0021/2014(H5N6) | A / H5N6 | Asia / China | Yuhai Bi (Institute of Microbiology, Chinese Academy of Sciences) |
| EPI_ISL_198887 | A/chicken/Jiangxi/12.05 NCDZT0031/2014(H5N6) | A / H5N6 | Asia / China | Yuhai Bi (Institute of Microbiology, Chinese Academy of Sciences) |
| EPI_ISL_198888 | A/chicken/Jiangxi/12.05 NCDZT0032/2014(H5N6) | A / H5N6 | Asia / China | Yuhai Bi (Institute of Microbiology, Chinese Academy of Sciences) |
| EPI_ISL_198889 | A/chicken/Jiangxi/12.05 NCDZT0040/2014(H5N6) | A / H5N6 | Asia / China | Yuhai Bi (Institute of Microbiology, Chinese Academy of Sciences) |
| EPI_ISL_198890 | A/chicken/Jiangxi/12.05 NCDZT007/2014(H5N6) | A / H5N6 | Asia / China | Yuhai Bi (Institute of Microbiology, Chinese Academy of Sciences) |
| EPI_ISL_198891 | A/chicken/Jiangxi/12.05 NCDZT009/2014(H5N6) | A / H5N6 | Asia / China | Yuhai Bi (Institute of Microbiology, Chinese Academy of Sciences) |
| EPI_ISL_198893 | A/pigeon/Sichuan/04.27 NBXJ743/2014(H5N6) | A / H5N6 | Asia / China | Yuhai Bi (Institute of Microbiology, Chinese Academy of Sciences) |
| EPI_ISL_198894 | A/environment/Jiangxi/02.10 NCDZT16/2015(H5N6) | A / H5N6 | Asia / China | Yuhai Bi (Institute of Microbiology, Chinese Academy of Sciences) |
| EPI_ISL_198895 | A/environment/Yunnan/03.15 DQWGH012/2015(H5N6) | A / H5N6 | Asia / China | Yuhai Bi (Institute of Microbiology, Chinese Academy of Sciences) |
| EPI_ISL_198896 | A/environment/Yunnan/03.15 DQWGH010/2015(H5N6) | A / H5N6 | Asia / China | Yuhai Bi (Institute of Microbiology, Chinese Academy of Sciences) |
| EPI_ISL_198897 | A/environment/Yunnan/03.15 DQWGH011-2/2015(H5N6) | A / H5N6 | Asia / China | Yuhai Bi (Institute of Microbiology, Chinese Academy of Sciences) |
| EPI_ISL_198898 | A/environment/Yunnan/03.15 DQWGH014/2015(H5N6) | A / H5N6 | Asia / China | Yuhai Bi (Institute of Microbiology, Chinese Academy of Sciences) |
| EPI_ISL_198899 | A/environment/Yunnan/03.17 DQJT039/2015(H5N6) | A / H5N6 | Asia / China | Yuhai Bi (Institute of Microbiology, Chinese Academy of Sciences) |
| EPI_ISL_198900 | A/environment/Yunnan/03.15 DQWGH009/2015(H5N6) | A / H5N6 | Asia / China | Yuhai Bi (Institute of Microbiology, Chinese Academy of Sciences) |
| EPI_ISL_198901 | A/environment/Yunnan/03.15 DQWGH020/2015(H5N6) | A / H5N6 | Asia / China | Yuhai Bi (Institute of Microbiology, Chinese Academy of Sciences) |
| EPI_ISL_198902 | A/environment/Yunnan/03.15 DQWGH017/2015(H5N6) | A / H5N6 | Asia / China | Yuhai Bi (Institute of Microbiology, Chinese Academy of Sciences) |
| EPI_ISL_198903 | A/environment/Yunnan/03.15 DQWGH004/2015(H5N6) | A / H5N6 | Asia / China | Yuhai Bi (Institute of Microbiology, Chinese Academy of Sciences) |
| EPI_ISL_198904 | A/environment/Yunnan/03.15 DQWGH008-1/2015(H5N6) | A / H5N6 | Asia / China | Yuhai Bi (Institute of Microbiology, Chinese Academy of Sciences) |
| EPI_ISL_198906 | A/duck/Hunan/12.17 YYFQH0018-Z-P/2014(H5N6) | A / H5N6 | Asia / China | Yuhai Bi (Institute of Microbiology, Chinese Academy of Sciences) |
| EPI_ISL_198908 | A/duck/Hunan/12.17 YYGK0063-O/2014(H5N6) | A / H5N6 | Asia / China | Yuhai Bi (Institute of Microbiology, Chinese Academy of Sciences) |
| EPI_ISL_198909 | A/duck/Hunan/12.17 YYGK0056/2014(H5N6) | A / H5N6 | Asia / China | Yuhai Bi (Institute of Microbiology, Chinese Academy of Sciences) |
| EPI_ISL_198911 | A/duck/Hunan/12.17 YYGK0066-P/2014(H5N6) | A / H5N6 | Asia / China | Yuhai Bi (Institute of Microbiology, Chinese Academy of Sciences) |
| EPI_ISL_198916 | A/chicken/Jiangxi/05.06 NCDZT1174-O/2015(H5N6) | A / H5N6 | Asia / China | Yuhai Bi (Institute of Microbiology, Chinese Academy of Sciences) |
| EPI_ISL_198917 | A/chicken/Jiangxi/04.01 NCJD0119-O/2015(H5N6) | A / H5N6 | Asia / China | Yuhai Bi (Institute of Microbiology, Chinese Academy of Sciences) |
| EPI_ISL_198919 | A/chicken/Jiangxi/04.01 NCJD0010-O/2015(H5N6) | A / H5N6 | Asia / China | Yuhai Bi (Institute of Microbiology, Chinese Academy of Sciences) |
| EPI_ISL_198920 | A/chicken/Jiangxi/04.01 NCJD0137-O/2015(H5N6) | A / H5N6 | Asia / China | Yuhai Bi (Institute of Microbiology, Chinese Academy of Sciences) |
| EPI_ISL_198921 | A/chicken/Jiangxi/04.01 NCJD0146-O/2015(H5N6) | A / H5N6 | Asia / China | Yuhai Bi (Institute of Microbiology, Chinese Academy of Sciences) |
| EPI_ISL_198923 | A/chicken/Jiangxi/04.01 NCDZT0173-P/2015(H5N6) | A / H5N6 | Asia / China | Yuhai Bi (Institute of Microbiology, Chinese Academy of Sciences) |
| EPI_ISL_198924 | A/chicken/Jiangxi/04.01 NCDZT0175-P/2015(H5N6) | A / H5N6 | Asia / China | Yuhai Bi (Institute of Microbiology, Chinese Academy of Sciences) |
| EPI_ISL_198926 | A/chicken/Jiangxi/04.01 NCDZT0192-P/2015(H5N6) | A / H5N6 | Asia / China | Yuhai Bi (Institute of Microbiology, Chinese Academy of Sciences) |
| EPI_ISL_198927 | A/chicken/Jiangxi/04.01 NCDZT0203-P/2015(H5N6) | A / H5N6 | Asia / China | Yuhai Bi (Institute of Microbiology, Chinese Academy of Sciences) |
| EPI_ISL_198929 | A/chicken/Jiangxi/04.01 NCDZT0206-P/2015(H5N6) | A / H5N6 | Asia / China | Yuhai Bi (Institute of Microbiology, Chinese Academy of Sciences) |
| EPI_ISL_198930 | A/chicken/Jiangxi/04.01 NCDZT0238-P/2015(H5N6) | A / H5N6 | Asia / China | Yuhai Bi (Institute of Microbiology, Chinese Academy of Sciences) |
| EPI_ISL_198932 | A/chicken/Jiangxi/04.01 NCDZT0279-P/2015(H5N6) | A / H5N6 | Asia / China | Yuhai Bi (Institute of Microbiology, Chinese Academy of Sciences) |
| EPI_ISL_198933 | A/chicken/Jiangxi/04.01 NCDZT0285-P/2015(H5N6) | A / H5N6 | Asia / China | Yuhai Bi (Institute of Microbiology, Chinese Academy of Sciences) |
| EPI_ISL_198934 | A/chicken/Jiangxi/04.01 NCDZT0291-P/2015(H5N6) | A / H5N6 | Asia / China | Yuhai Bi (Institute of Microbiology, Chinese Academy of Sciences) |
| EPI_ISL_198935 | A/chicken/Jiangxi/04.01 NCDZT0292-P/2015(H5N6) | A / H5N6 | Asia / China | Yuhai Bi (Institute of Microbiology, Chinese Academy of Sciences) |
| EPI_ISL_198937 | A/duck/Jiangxi/04.01 NCDZT0312-O/2015(H5N6) | A / H5N6 | Asia / China | Yuhai Bi (Institute of Microbiology, Chinese Academy of Sciences) |
| EPI_ISL_198939 | A/duck/Jiangxi/04.01 NCDZT0334-P/2015(H5N6) | A / H5N6 | Asia / China | Yuhai Bi (Institute of Microbiology, Chinese Academy of Sciences) |
| EPI_ISL_198949 | A/environment/Yunnan/03.15 DQWGH001/2015(H5N6) | A / H5N6 | Asia / China | Yuhai Bi (Institute of Microbiology, Chinese Academy of Sciences) |
| EPI_ISL_198952 | A/environment/Yunnan/03.17 DQJT040/2015(H5N6) | A / H5N6 | Asia / China | Yuhai Bi (Institute of Microbiology, Chinese Academy of Sciences) |
| EPI_ISL_198953 | A/environment/Jiangxi/05.07 NCJD0010D/2015(H5N6) | A / H5N6 | Asia / China | Yuhai Bi (Institute of Microbiology, Chinese Academy of Sciences) |
| EPI_ISL_198954 | A/environment/Jiangxi/05.07 NCJD0012D/2015(H5N6) | A / H5N6 | Asia / China | Yuhai Bi (Institute of Microbiology, Chinese Academy of Sciences) |

| | | | | |
|---|---|---|---|---|
| EPI_ISL_198956 | A/environment/Jiangxi/05.07 NCJD0017D/2015(H5N6) | A / H5N6 | Asia / China | Yuhai Bi (Institute of Microbiology, Chinese Academy of Sciences) |
| EPI_ISL_198958 | A/environment/Jiangxi/05.07 NCJD0022D/2015(H5N6) | A / H5N6 | Asia / China | Yuhai Bi (Institute of Microbiology, Chinese Academy of Sciences) |
| EPI_ISL_198959 | A/environment/Jiangxi/05.07 NCJD0023D/2015(H5N6) | A / H5N6 | Asia / China | Yuhai Bi (Institute of Microbiology, Chinese Academy of Sciences) |
| EPI_ISL_198960 | A/environment/Jiangxi/05.07 NCJD0027D/2015(H5N6) | A / H5N6 | Asia / China | Yuhai Bi (Institute of Microbiology, Chinese Academy of Sciences) |
| EPI_ISL_198962 | A/Pigeon/Jiangxi/04.01 NCDZT0347-O/2015(H5N6) | A / H5N6 | Asia / China | Yuhai Bi (Institute of Microbiology, Chinese Academy of Sciences) |
| EPI_ISL_198964 | A/Pigeon/Jiangxi/04.01 NCDZT0348-O/2015(H5N6) | A / H5N6 | Asia / China | Yuhai Bi (Institute of Microbiology, Chinese Academy of Sciences) |
| EPI_ISL_198967 | A/Pigeon/Jiangxi/04.01 NCDZT0352-O/2015(H5N6) | A / H5N6 | Asia / China | Yuhai Bi (Institute of Microbiology, Chinese Academy of Sciences) |
| EPI_ISL_198968 | A/Pigeon/Jiangxi/04.01 NCDZT0353-O/2015(H5N6) | A / H5N6 | Asia / China | Yuhai Bi (Institute of Microbiology, Chinese Academy of Sciences) |
| EPI_ISL_198971 | A/chicken/Jiangxi/05.06 NCDZT0039B-P/2015(H5N6) | A / H5N6 | Asia / China | Yuhai Bi (Institute of Microbiology, Chinese Academy of Sciences) |
| EPI_ISL_198972 | A/chicken/Jiangxi/05.06 NCDZT0003C-P/2015(H5N6) | A / H5N6 | Asia / China | Yuhai Bi (Institute of Microbiology, Chinese Academy of Sciences) |
| EPI_ISL_198976 | A/chicken/Jiangxi/04.01 NCDZT0042-O/2015(H5N6) | A / H5N6 | Asia / China | Yuhai Bi (Institute of Microbiology, Chinese Academy of Sciences) |
| EPI_ISL_198977 | A/chicken/Jiangxi/05.06 NCDZT0043B-P/2015(H5N6) | A / H5N6 | Asia / China | Yuhai Bi (Institute of Microbiology, Chinese Academy of Sciences) |
| EPI_ISL_198981 | A/environment/Jiangxi/05.07 NCJD0052C/2015(H5N6) | A / H5N6 | Asia / China | Yuhai Bi (Institute of Microbiology, Chinese Academy of Sciences) |
| EPI_ISL_198984 | A/duck/Jiangxi/05.07 NCJD0056A-P/2015(H5N6) | A / H5N6 | Asia / China | Yuhai Bi (Institute of Microbiology, Chinese Academy of Sciences) |
| EPI_ISL_198985 | A/chicken/Jiangxi/05.06 NCDZT0058B-P/2015(H5N6) | A / H5N6 | Asia / China | Yuhai Bi (Institute of Microbiology, Chinese Academy of Sciences) |
| EPI_ISL_198987 | A/duck/Jiangxi/05.07 NCJD0061A-P/2015(H5N6) | A / H5N6 | Asia / China | Yuhai Bi (Institute of Microbiology, Chinese Academy of Sciences) |
| EPI_ISL_198990 | A/chicken/Jiangxi/04.01 NCNP0065-Z-O/2015(H5N6) | A / H5N6 | Asia / China | Yuhai Bi (Institute of Microbiology, Chinese Academy of Sciences) |
| EPI_ISL_198991 | A/duck/Jiangxi/05.07 NCJD0067A-P/2015(H5N6) | A / H5N6 | Asia / China | Yuhai Bi (Institute of Microbiology, Chinese Academy of Sciences) |
| EPI_ISL_198992 | A/environment/Jiangxi/05.07 NCJD0070C/2015(H5N6) | A / H5N6 | Asia / China | Yuhai Bi (Institute of Microbiology, Chinese Academy of Sciences) |
| EPI_ISL_198996 | A/environment/Jiangxi/05.07 NCJD0078C/2015(H5N6) | A / H5N6 | Asia / China | Yuhai Bi (Institute of Microbiology, Chinese Academy of Sciences) |
| EPI_ISL_199006 | A/environment/Jiangxi/05.07 NCJD0095C/2015(H5N6) | A / H5N6 | Asia / China | Yuhai Bi (Institute of Microbiology, Chinese Academy of Sciences) |
| EPI_ISL_199007 | A/environment/Jiangxi/05.07 NCJD0098C/2015(H5N6) | A / H5N6 | Asia / China | Yuhai Bi (Institute of Microbiology, Chinese Academy of Sciences) |
| EPI_ISL_199009 | A/environment/Jiangxi/05.07 NCJD0093C/2015(H5N6) | A / H5N6 | Asia / China | Yuhai Bi (Institute of Microbiology, Chinese Academy of Sciences) |
| EPI_ISL_199010 | A/chicken/Jiangxi/12.05 NCNP006/2014(H5N6) | A / H5N6 | Asia / China | Yuhai Bi (Institute of Microbiology, Chinese Academy of Sciences) |
| EPI_ISL_199011 | A/chicken/Hubei/03.06 WHWTZ0115-O/2015(H5N6) | A / H5N6 | Asia / China | Yuhai Bi (Institute of Microbiology, Chinese Academy of Sciences) |
| EPI_ISL_199015 | A/duck/Hubei/03.06 WHWTZ0129-P/2015(H5N6) | A / H5N6 | Asia / China | Yuhai Bi (Institute of Microbiology, Chinese Academy of Sciences) |
| EPI_ISL_199017 | A/chicken/Hubei/03.06 WHWTZ0148-O/2015(H5N6) | A / H5N6 | Asia / China | Yuhai Bi (Institute of Microbiology, Chinese Academy of Sciences) |
| EPI_ISL_199026 | A/duck/Hubei/03.06 WHWTZ0123-O/2015(H5N6) | A / H5N6 | Asia / China | Yuhai Bi (Institute of Microbiology, Chinese Academy of Sciences) |
| EPI_ISL_199028 | A/duck/Hubei/03.06 WHWTZ0124-O/2015(H5N6) | A / H5N6 | Asia / China | Yuhai Bi (Institute of Microbiology, Chinese Academy of Sciences) |
| EPI_ISL_199029 | A/duck/HuBei/03.06 WHWTZ0158-P/2015(H5N6) | A / H5N6 | Asia / China | Yuhai Bi (Institute of Microbiology, Chinese Academy of Sciences) |
| EPI_ISL_199030 | A/duck/Hubei/03.06 WHWTZ0125-O/2015(H5N6) | A / H5N6 | Asia / China | Yuhai Bi (Institute of Microbiology, Chinese Academy of Sciences) |
| EPI_ISL_199031 | A/duck/Hubei/03.06 WHWTZ0130-O/2015(H5N6) | A / H5N6 | Asia / China | Yuhai Bi (Institute of Microbiology, Chinese Academy of Sciences) |
| EPI_ISL_199032 | A/duck/Hubei/03.06 WHWTZ0148-O/2015(H5N6) | A / H5N6 | Asia / China | Yuhai Bi (Institute of Microbiology, Chinese Academy of Sciences) |
| EPI_ISL_199034 | A/chicken/Yunnan/03.16 DQJT0068-O/2015(H5N6) | A / H5N6 | Asia / China | Yuhai Bi (Institute of Microbiology, Chinese Academy of Sciences) |
| EPI_ISL_199035 | A/chicken/Yunnan/03.15 DQXYL0073-O/2015(H5N6) | A / H5N6 | Asia / China | Yuhai Bi (Institute of Microbiology, Chinese Academy of Sciences) |
| EPI_ISL_199036 | A/chicken/Yunnan/03.16 DQJT0066-O/2015(H5N6) | A / H5N6 | Asia / China | Yuhai Bi (Institute of Microbiology, Chinese Academy of Sciences) |
| EPI_ISL_199037 | A/duck/Yunnan/03.16 DQXYL0013-O/2015 | A / H5N6 | Asia / China | Yuhai Bi (Institute of Microbiology, Chinese Academy of Sciences) |
| EPI_ISL_199038 | A/duck/Hunan/04.22 LDDX037-P/2015(H5N6) | A / H5N6 | Asia / China | Yuhai Bi (Institute of Microbiology, Chinese Academy of Sciences) |
| EPI_ISL_199041 | A/chicken/Yunnan/03.16 DQXYL023-O/2015(H5N6) | A / H5N6 | Asia / China | Yuhai Bi (Institute of Microbiology, Chinese Academy of Sciences) |
| EPI_ISL_199042 | A/duck/Yunnan/03.16 DQXYL003-O/2015(H5N6) | A / H5N6 | Asia / China | Yuhai Bi (Institute of Microbiology, Chinese Academy of Sciences) |
| EPI_ISL_199043 | A/duck/Yunnan/03.16 DQXYL017-O/2015(H5N6) | A / H5N6 | Asia / China | Yuhai Bi (Institute of Microbiology, Chinese Academy of Sciences) |
| EPI_ISL_199044 | A/duck/Yunnan/03.16 DQXYL022-O/2015(H5N6) | A / H5N6 | Asia / China | Yuhai Bi (Institute of Microbiology, Chinese Academy of Sciences) |
| EPI_ISL_199045 | A/duck/Hunan/02.07 YYFQH010-P/2015(H5N6) | A / H5N6 | Asia / China | Yuhai Bi (Institute of Microbiology, Chinese Academy of Sciences) |
| EPI_ISL_199046 | A/duck/Hunan/01.21 YYFQH010-O/2015(H5N6) | A / H5N6 | Asia / China | Yuhai Bi (Institute of Microbiology, Chinese Academy of Sciences) |
| EPI_ISL_199047 | A/duck/Hunan/01.21 YYFQH011-O/2015(H5N6) | A / H5N6 | Asia / China | Yuhai Bi (Institute of Microbiology, Chinese Academy of Sciences) |
| EPI_ISL_199049 | A/duck/Hunan/02.07 YYFQHY12Y-O/2015(H5N6) | A / H5N6 | Asia / China | Yuhai Bi (Institute of Microbiology, Chinese Academy of Sciences) |
| EPI_ISL_199050 | A/duck/Hunan/01.21 YYFQH012-O/2015(H5N6) | A / H5N6 | Asia / China | Yuhai Bi (Institute of Microbiology, Chinese Academy of Sciences) |
| EPI_ISL_199051 | A/duck/Hunan/03.09 YYFQH0047-O/2015(H5N6) | A / H5N6 | Asia / China | Yuhai Bi (Institute of Microbiology, Chinese Academy of Sciences) |
| EPI_ISL_199052 | A/duck/Hunan/02.07 YYFQH015-P/2015(H5N6) | A / H5N6 | Asia / China | Yuhai Bi (Institute of Microbiology, Chinese Academy of Sciences) |
| EPI_ISL_199053 | A/duck/Hunan/01.21 YYFQH015-O/2015(H5N6) | A / H5N6 | Asia / China | Yuhai Bi (Institute of Microbiology, Chinese Academy of Sciences) |
| EPI_ISL_199054 | A/duck/Hunan/01.21 YYFQH016-O/2015(H5N6) | A / H5N6 | Asia / China | Yuhai Bi (Institute of Microbiology, Chinese Academy of Sciences) |
| EPI_ISL_199055 | A/duck/Hunan/01.21 YYFQH018-O/2015(H5N6) | A / H5N6 | Asia / China | Yuhai Bi (Institute of Microbiology, Chinese Academy of Sciences) |
| EPI_ISL_199056 | A/duck/Hunan/01.21 YYFQH019-P/2015(H5N6) | A / H5N6 | Asia / China | Yuhai Bi (Institute of Microbiology, Chinese Academy of Sciences) |
| EPI_ISL_199057 | A/duck/Hunan/02.07 YYFQHY19Y-O/2015(H5N6) | A / H5N6 | Asia / China | Yuhai Bi (Institute of Microbiology, Chinese Academy of Sciences) |
| EPI_ISL_199058 | A/duck/Hunan/02.07 YYFQHY20Y-O/2015(H5N6) | A / H5N6 | Asia / China | Yuhai Bi (Institute of Microbiology, Chinese Academy of Sciences) |
| EPI_ISL_199059 | A/duck/Hunan/01.21 YYFQH022-O/2015(H5N6) | A / H5N6 | Asia / China | Yuhai Bi (Institute of Microbiology, Chinese Academy of Sciences) |
| EPI_ISL_199062 | A/Phalacrocorax/Hubei/01.09 V5/2015(H5N6) | A / H5N6 | Asia / China | Yuhai Bi (Institute of Microbiology, Chinese Academy of Sciences) |
| EPI_ISL_199063 | A/Phalacrocorax/Hubei/01.09 V7/2015(H5N6) | A / H5N6 | Asia / China | Yuhai Bi (Institute of Microbiology, Chinese Academy of Sciences) |
| EPI_ISL_199064 | A/duck/Hunan/01.21 YYFQH023-O/2015(H5N6) | A / H5N6 | Asia / China | Yuhai Bi (Institute of Microbiology, Chinese Academy of Sciences) |
| EPI_ISL_199065 | A/duck/Hunan/01.21 YYFQH024-P/2015(H5N6) | A / H5N6 | Asia / China | Yuhai Bi (Institute of Microbiology, Chinese Academy of Sciences) |
| EPI_ISL_199066 | A/duck/Hunan/01.21 YYFQH025-P/2015(H5N6) | A / H5N6 | Asia / China | Yuhai Bi (Institute of Microbiology, Chinese Academy of Sciences) |
| EPI_ISL_199067 | A/duck/Hunan/02.07 YYFQH026-O/2015(H5N6) | A / H5N6 | Asia / China | Yuhai Bi (Institute of Microbiology, Chinese Academy of Sciences) |
| EPI_ISL_199068 | A/duck/Hunan/01.21 YYFQH027-O/2015(H5N6) | A / H5N6 | Asia / China | Yuhai Bi (Institute of Microbiology, Chinese Academy of Sciences) |
| EPI_ISL_199069 | A/duck/Hunan/02.07 YYFQH029-O/2015(H5N6) | A / H5N6 | Asia / China | Yuhai Bi (Institute of Microbiology, Chinese Academy of Sciences) |
| EPI_ISL_199070 | A/duck/Hunan/01.21 YYFQH030-O/2015(H5N6) | A / H5N6 | Asia / China | Yuhai Bi (Institute of Microbiology, Chinese Academy of Sciences) |
| EPI_ISL_199071 | A/duck/Hunan/01.21 YYFQH032-P/2015(H5N6) | A / H5N6 | Asia / China | Yuhai Bi (Institute of Microbiology, Chinese Academy of Sciences) |
| EPI_ISL_199072 | A/duck/Hunan/02.07 YYFQH032-O/2015(H5N6) | A / H5N6 | Asia / China | Yuhai Bi (Institute of Microbiology, Chinese Academy of Sciences) |
| EPI_ISL_199073 | A/duck/Hunan/02.07 YYFQH003-P/2015(H5N6) | A / H5N6 | Asia / China | Yuhai Bi (Institute of Microbiology, Chinese Academy of Sciences) |
| EPI_ISL_199074 | A/duck/Hunan/01.21 YYFQH003-O/2015(H5N6) | A / H5N6 | Asia / China | Yuhai Bi (Institute of Microbiology, Chinese Academy of Sciences) |
| EPI_ISL_199075 | A/duck/Hunan/01.21 YYFQH005-P/2015(H5N6) | A / H5N6 | Asia / China | Yuhai Bi (Institute of Microbiology, Chinese Academy of Sciences) |
| EPI_ISL_199076 | A/duck/Hunan/01.21 YYFQH005-O/2015(H5N6) | A / H5N6 | Asia / China | Yuhai Bi (Institute of Microbiology, Chinese Academy of Sciences) |
| EPI_ISL_199077 | A/duck/Hunan/02.07 YYFQH006-P/2015(H5N6) | A / H5N6 | Asia / China | Yuhai Bi (Institute of Microbiology, Chinese Academy of Sciences) |

| | | | | |
|---|---|---|---|---|
| EPI_ISL_199079 | A/duck/Hunan/01.21 YYFQH006-O/2015(H5N6) | A / H5N6 | Asia / China | Yuhai Bi (Institute of Microbiology, Chinese Academy of Sciences) |
| EPI_ISL_199080 | A/duck/Hunan/01.21 YYFQH007-O/2015(H5N6) | A / H5N6 | Asia / China | Yuhai Bi (Institute of Microbiology, Chinese Academy of Sciences) |
| EPI_ISL_199081 | A/chicken/Hunan/04.14 YYXS0835-O/2015(H5N6) | A / H5N6 | Asia / China | Yuhai Bi (Institute of Microbiology, Chinese Academy of Sciences) |
| EPI_ISL_199083 | A/duck/Hunan/01.21 YYFQH009-O/2015(H5N6) | A / H5N6 | Asia / China | Yuhai Bi (Institute of Microbiology, Chinese Academy of Sciences) |
| EPI_ISL_199084 | A/environment/Yunnan/03.16 DQXYL0053-Z-O/2015(H5N6) | A / H5N6 | Asia / China | Yuhai Bi (Institute of Microbiology, Chinese Academy of Sciences) |
| EPI_ISL_199085 | A/chicken/Yunnan/03.15 DQJT0054-Z-P/2015(H5N6) | A / H5N6 | Asia / China | Yuhai Bi (Institute of Microbiology, Chinese Academy of Sciences) |
| EPI_ISL_199086 | A/chicken/Yunnan/03.16 DQXYL0063-Z-O/2015(H5N6) | A / H5N6 | Asia / China | Yuhai Bi (Institute of Microbiology, Chinese Academy of Sciences) |
| EPI_ISL_199087 | A/environment/Yunnan/03.17 DQJT0015/2015(H5N6) | A / H5N6 | Asia / China | Yuhai Bi (Institute of Microbiology, Chinese Academy of Sciences) |
| EPI_ISL_199093 | A/duck/Yunnan/03.15 DQXYL007-Z-O/2015(H5N6) | A / H5N6 | Asia / China | Yuhai Bi (Institute of Microbiology, Chinese Academy of Sciences) |
| EPI_ISL_199094 | A/environment/Yunnan/03.15 DQWGH008-Z/2015(H5N6) | A / H5N6 | Asia / China | Yuhai Bi (Institute of Microbiology, Chinese Academy of Sciences) |
| EPI_ISL_199095 | A/environment/Yunnan/03.16 DQXYL0027/2015(H5N6) | A / H5N6 | Asia / China | Yuhai Bi (Institute of Microbiology, Chinese Academy of Sciences) |
| EPI_ISL_199099 | A/environment/Yunnan/03.16 DQXYL0032/2015(H5N6) | A / H5N6 | Asia / China | Yuhai Bi (Institute of Microbiology, Chinese Academy of Sciences) |
| EPI_ISL_199100 | A/chicken/Yunnan/03.17 DQJT0041-O/2015(H5N6) | A / H5N6 | Asia / China | Yuhai Bi (Institute of Microbiology, Chinese Academy of Sciences) |
| EPI_ISL_199101 | A/chicken/Yunnan/03.16 DQXYL0044-O/2015(N6) | A / H5N6 | Asia / China | Yuhai Bi (Institute of Microbiology, Chinese Academy of Sciences) |
| EPI_ISL_199102 | A/chicken/Yunnan/03.16 DQXYL0050-O/2015(H5N6) | A / H5N6 | Asia / China | Yuhai Bi (Institute of Microbiology, Chinese Academy of Sciences) |
| EPI_ISL_199103 | A/environment/Yunnan/03.17 DQJT0051-P/2015(H5N6) | A / H5N6 | Asia / China | Yuhai Bi (Institute of Microbiology, Chinese Academy of Sciences) |
| EPI_ISL_199104 | A/chicken/Yunnan/03.16 DQXYL0052-O/2015(H5N6) | A / H5N6 | Asia / China | Yuhai Bi (Institute of Microbiology, Chinese Academy of Sciences) |
| EPI_ISL_199106 | A/chicken/Yunnan/03.16 DQXYL057-O/2015(H5N6) | A / H5N6 | Asia / China | Yuhai Bi (Institute of Microbiology, Chinese Academy of Sciences) |
| EPI_ISL_199109 | A/chicken/Yunnan/03.16 DQJT069-O/2015(H5N6) | A / H5N6 | Asia / China | Yuhai Bi (Institute of Microbiology, Chinese Academy of Sciences) |
| EPI_ISL_199110 | A/Anser fabalis/Hunan/03.09 YYDTH0007/2015(H5N6) | A / H5N6 | Asia / China | Yuhai Bi (Institute of Microbiology, Chinese Academy of Sciences) |
| EPI_ISL_199112 | A/environment/Hunan/04.14 YYGK391/2015(H5N6) | A / H5N6 | Asia / China | Yuhai Bi (Institute of Microbiology, Chinese Academy of Sciences) |
| EPI_ISL_199113 | A/environment/Hunan/04.14 YYGK0393-1/2015(H5N6) | A / H5N6 | Asia / China | Yuhai Bi (Institute of Microbiology, Chinese Academy of Sciences) |
| EPI_ISL_199119 | A/duck/Hunan/04.14 YYGK460-O/2015(H5N6) | A / H5N6 | Asia / China | Yuhai Bi (Institute of Microbiology, Chinese Academy of Sciences) |
| EPI_ISL_199120 | A/duck/Hunan/04.14 YYGK464-O/2015(H5N6) | A / H5N6 | Asia / China | Yuhai Bi (Institute of Microbiology, Chinese Academy of Sciences) |
| EPI_ISL_199122 | A/duck/Hunan/04.14 YYGK468-O/2015(H5N6) | A / H5N6 | Asia / China | Yuhai Bi (Institute of Microbiology, Chinese Academy of Sciences) |
| EPI_ISL_199130 | A/chicken/Hunan/04.14 YYGK607-P/2015(H5N6) | A / H5N6 | Asia / China | Yuhai Bi (Institute of Microbiology, Chinese Academy of Sciences) |
| EPI_ISL_199137 | A/duck/Hunan/04.14 YYXS877-P/2015(H5N6) | A / H5N6 | Asia / China | Yuhai Bi (Institute of Microbiology, Chinese Academy of Sciences) |
| EPI_ISL_199138 | A/duck/Hunan/04.14 YYGK0878-P/2015(H5N6) | A / H5N6 | Asia / China | Yuhai Bi (Institute of Microbiology, Chinese Academy of Sciences) |
| EPI_ISL_199146 | A/chicken/Miyazaki/2-4/2014 | A / H5N8 | Asia / Japan / Miyazaki / Miyazaki city | Takehiko Saito (National Institute of Animal Health) |
| EPI_ISL_199147 | A/chicken/Yamaguchi/6/2014 | A / H5N8 | Asia / Japan / Yamaguchi | Takehiko Saito (National Institute of Animal Health) |
| EPI_ISL_199148 | A/chicken/Okayama/1-2/2015 | A / H5N8 | Asia / Japan / Okayama | Takehiko Saito (National Institute of Animal Health) |
| EPI_ISL_199149 | A/chicken/Saga/1-1/2015 | A / H5N8 | Asia / Japan / Saga | Takehiko Saito (National Institute of Animal Health) |
| EPI_ISL_199151 | A/duck/Shandong/11.14 DL-DKd11/2013(H5N8) | A / H5N8 | Asia / China | Yuhai Bi (Institute of Microbiology, Chinese Academy of Sciences) |
| EPI_ISL_199153 | A/goose/Shandong/11.02 CQ-GS/2014(H5N6) | A / H5N6 | Asia / China | Yuhai Bi (Institute of Microbiology, Chinese Academy of Sciences) |
| EPI_ISL_199155 | A/goose/Shandong/12.05 YG-GS/2014(H5N6) | A / H5N6 | Asia / China | Yuhai Bi (Institute of Microbiology, Chinese Academy of Sciences) |
| EPI_ISL_199156 | A/goose/Shandong/12.08 YG-GS/2014(H5N8) | A / H5N8 | Asia / China | Yuhai Bi (Institute of Microbiology, Chinese Academy of Sciences) |
| EPI_ISL_199158 | A/goose/Shandong/05.23 DT-GS/2014(H5N8) | A / H5N8 | Asia / China | Yuhai Bi (Institute of Microbiology, Chinese Academy of Sciences) |
| EPI_ISL_199159 | A/goose/Shandong/GD-GS/2014(H5N8) | A / H5N8 | Asia / China | Yuhai Bi (Institute of Microbiology, Chinese Academy of Sciences) |
| EPI_ISL_199160 | A/goose/Shandong/01.01 JY-GS/2015(H5N2) | A / H5N2 | Asia / China | Yuhai Bi (Institute of Microbiology, Chinese Academy of Sciences) |
| EPI_ISL_199161 | A/goose/Shandong/01.07 JQ-GS/2015(H5N2) | A / H5N2 | Asia / China | Yuhai Bi (Institute of Microbiology, Chinese Academy of Sciences) |
| EPI_ISL_199163 | A/goose/Shandong/02.16 YG-GS/2015(H5N6) | A / H5N6 | Asia / China | Yuhai Bi (Institute of Microbiology, Chinese Academy of Sciences) |
| EPI_ISL_199174 | A/duck/Guangdong/03.26 DGCPLB021-O/2015(H5N6) | A / H5N6 | Asia / China | Yuhai Bi (Institute of Microbiology, Chinese Academy of Sciences) |
| EPI_ISL_199175 | A/duck/Guangdong/03.26 DGCP066-O/2015(H5N6) | A / H5N6 | Asia / China | Yuhai Bi (Institute of Microbiology, Chinese Academy of Sciences) |
| EPI_ISL_199178 | A/duck/Guangdong/03.26 DGCP074-O/2015(H5N6) | A / H5N6 | Asia / China | Yuhai Bi (Institute of Microbiology, Chinese Academy of Sciences) |
| EPI_ISL_199179 | A/duck/Guangdong/03.26 DGCP079-O/2015(H5N6) | A / H5N6 | Asia / China | Yuhai Bi (Institute of Microbiology, Chinese Academy of Sciences) |
| EPI_ISL_199180 | A/duck/Guangdong/03.26 DGCP080-O/2015(H5N6) | A / H5N6 | Asia / China | Yuhai Bi (Institute of Microbiology, Chinese Academy of Sciences) |
| EPI_ISL_199181 | A/duck/Guangdong/03.26 DGCP076-O/2015(H5N6) | A / H5N6 | Asia / China | Yuhai Bi (Institute of Microbiology, Chinese Academy of Sciences) |
| EPI_ISL_199183 | A/duck/Guangdong/03.26 DGCPLB020-P/2015(H5N6) | A / H5N6 | Asia / China | Yuhai Bi (Institute of Microbiology, Chinese Academy of Sciences) |
| EPI_ISL_199185 | A/duck/Guangdong/03.26 DGCP709-P/2015(H5N6) | A / H5N6 | Asia / China | Yuhai Bi (Institute of Microbiology, Chinese Academy of Sciences) |
| EPI_ISL_199186 | A/duck/Guangdong/03.26 DGCPLB022-P/2015(H5N6) | A / H5N6 | Asia / China | Yuhai Bi (Institute of Microbiology, Chinese Academy of Sciences) |
| EPI_ISL_199187 | A/duck/Guangdong/03.26 DGCPLB023-P/2015(H5N6) | A / H5N6 | Asia / China | Yuhai Bi (Institute of Microbiology, Chinese Academy of Sciences) |
| EPI_ISL_199188 | A/duck/Guangdong/03.26 DGCPLB019-P/2015(H5N6) | A / H5N6 | Asia / China | Yuhai Bi (Institute of Microbiology, Chinese Academy of Sciences) |
| EPI_ISL_199189 | A/duck/Guangdong/03.26 DGCP078-P/2015(H5N6) | A / H5N6 | Asia / China | Yuhai Bi (Institute of Microbiology, Chinese Academy of Sciences) |
| EPI_ISL_199190 | A/duck/Guangdong/03.26 DGCPLB009-O/2015(H5N6) | A / H5N6 | Asia / China | Yuhai Bi (Institute of Microbiology, Chinese Academy of Sciences) |
| EPI_ISL_199192 | A/duck/Guangdong/03.26 DGCPLB007-P/2015(H5N6) | A / H5N6 | Asia / China | Yuhai Bi (Institute of Microbiology, Chinese Academy of Sciences) |
| EPI_ISL_199195 | A/duck/Guangdong/03.26 DGCP080-P/2015(H5N6) | A / H5N6 | Asia / China | Yuhai Bi (Institute of Microbiology, Chinese Academy of Sciences) |
| EPI_ISL_199196 | A/goose/Shandong/04.22 GT/2015(H5N2) | A / H5N2 | Asia / China | Yuhai Bi (Institute of Microbiology, Chinese Academy of Sciences) |
| EPI_ISL_199198 | A/duck/Guangdong/03.26 DGCP081-P/2015(H5N6) | A / H5N6 | Asia / China | Yuhai Bi (Institute of Microbiology, Chinese Academy of Sciences) |
| EPI_ISL_199199 | A/duck/Guangdong/03.26 DGCP078-O/2015(H5N6) | A / H5N6 | Asia / China | Yuhai Bi (Institute of Microbiology, Chinese Academy of Sciences) |
| EPI_ISL_199200 | A/duck/Guangdong/03.26 DGCP084-P/2015(H5N6) | A / H5N6 | Asia / China | Yuhai Bi (Institute of Microbiology, Chinese Academy of Sciences) |
| EPI_ISL_199201 | A/duck/Guangdong/03.26 DGCP061-P/2015(H5N6) | A / H5N6 | Asia / China | Yuhai Bi (Institute of Microbiology, Chinese Academy of Sciences) |
| EPI_ISL_199207 | A/goose/Guangdong/04.22 DGCP117-O/2015(H5N6) | A / H5N6 | Asia / China | Yuhai Bi (Institute of Microbiology, Chinese Academy of Sciences) |
| EPI_ISL_199208 | A/duck/Guangdong/04.23 DGQTXC229-O/2015(H5N6) | A / H5N6 | Asia / China | Yuhai Bi (Institute of Microbiology, Chinese Academy of Sciences) |
| EPI_ISL_199209 | A/duck/Guangdong/04.22 DGCP068-O/2015(H5N6) | A / H5N6 | Asia / China | Yuhai Bi (Institute of Microbiology, Chinese Academy of Sciences) |
| EPI_ISL_199211 | A/duck/Guangdong/04.23 DGQTSJ150-O/2015(H5N6) | A / H5N6 | Asia / China | Yuhai Bi (Institute of Microbiology, Chinese Academy of Sciences) |
| EPI_ISL_199214 | A/duck/Guangdong/04.23 DGQTXC202-O/2015(H5N6) | A / H5N6 | Asia / China | Yuhai Bi (Institute of Microbiology, Chinese Academy of Sciences) |
| EPI_ISL_199215 | A/duck/Guangdong/04.22 DGCP069-O/2015(H5N6) | A / H5N6 | Asia / China | Yuhai Bi (Institute of Microbiology, Chinese Academy of Sciences) |
| EPI_ISL_199222 | A/duck/Guangdong/04.23 DGQTXC203-O/2015(H5N6) | A / H5N6 | Asia / China | Yuhai Bi (Institute of Microbiology, Chinese Academy of Sciences) |
| EPI_ISL_199230 | A/duck/Guangdong/04.22 DGCPLB018-O/2015(H5N6) | A / H5N6 | Asia / China | Yuhai Bi (Institute of Microbiology, Chinese Academy of Sciences) |
| EPI_ISL_199234 | A/duck/Guangdong/04.23 DGQTXC229-P/2015(H5N6) | A / H5N6 | Asia / China | Yuhai Bi (Institute of Microbiology, Chinese Academy of Sciences) |
| EPI_ISL_199235 | A/duck/Guangdong/04.23 DGQTSJ145-O/2015(H5N6) | A / H5N6 | Asia / China | Yuhai Bi (Institute of Microbiology, Chinese Academy of Sciences) |
| EPI_ISL_199236 | A/duck/Guangdong/04.23 DGQTXC231-O/2015(H5N6) | A / H5N6 | Asia / China | Yuhai Bi (Institute of Microbiology, Chinese Academy of Sciences) |

| | | | | |
|---|---|---|---|---|
| EPI_ISL_199241 | A/duck/Guangdong/04.23 DGQTXC226-O/2015(H5N6) | A / H5N6 | Asia / China | Yuhai Bi (Institute of Microbiology, Chinese Academy of Sciences) |
| EPI_ISL_199242 | A/duck/Guangdong/04.22 DGCP061-O/2015(H5N6) | A / H5N6 | Asia / China | Yuhai Bi (Institute of Microbiology, Chinese Academy of Sciences) |
| EPI_ISL_199243 | A/environment/Guangdong/04.22 DGCP027/2015(H5N6) | A / H5N6 | Asia / China | Yuhai Bi (Institute of Microbiology, Chinese Academy of Sciences) |
| EPI_ISL_199244 | A/goose/Guangdong/04.22 DGCP120-P/2015(H5N6) | A / H5N6 | Asia / China | Yuhai Bi (Institute of Microbiology, Chinese Academy of Sciences) |
| EPI_ISL_199250 | A/duck/Guangdong/04.22 DGCP079-O/2015(H5N6) | A / H5N6 | Asia / China | Yuhai Bi (Institute of Microbiology, Chinese Academy of Sciences) |
| EPI_ISL_199252 | A/duck/Guangdong/04.22 DGCPLB020-O/2015(H5N6) | A / H5N6 | Asia / China | Yuhai Bi (Institute of Microbiology, Chinese Academy of Sciences) |
| EPI_ISL_199255 | A/duck/Guangdong/04.22 DGCP071-O/2015(H5N6) | A / H5N6 | Asia / China | Yuhai Bi (Institute of Microbiology, Chinese Academy of Sciences) |
| EPI_ISL_199259 | A/duck/Guangdong/04.22 DGCP062-O/2015(H5N6) | A / H5N6 | Asia / China | Yuhai Bi (Institute of Microbiology, Chinese Academy of Sciences) |
| EPI_ISL_199260 | A/goose/Guangdong/04.22 DGCPLB045-O/2015(H5N6) | A / H5N6 | Asia / China | Yuhai Bi (Institute of Microbiology, Chinese Academy of Sciences) |
| EPI_ISL_199262 | A/duck/Guangdong/04.23 DGQTXC228-O/2015(H5N6) | A / H5N6 | Asia / China | Yuhai Bi (Institute of Microbiology, Chinese Academy of Sciences) |
| EPI_ISL_199268 | A/goose/Guangdong/04.22 DGCP118-P/2015(H5N6) | A / H5N6 | Asia / China | Yuhai Bi (Institute of Microbiology, Chinese Academy of Sciences) |
| EPI_ISL_199269 | A/goose/Guangdong/04.23 DGQTXC216-P/2015(H5N6) | A / H5N6 | Asia / China | Yuhai Bi (Institute of Microbiology, Chinese Academy of Sciences) |
| EPI_ISL_199271 | A/duck/Guangdong/04.23 DGQTXC202-P/2015(H5N6) | A / H5N6 | Asia / China | Yuhai Bi (Institute of Microbiology, Chinese Academy of Sciences) |
| EPI_ISL_199280 | A/duck/Guangdong/04.22 DGCP067-O/2015(H5N6) | A / H5N6 | Asia / China | Yuhai Bi (Institute of Microbiology, Chinese Academy of Sciences) |
| EPI_ISL_199281 | A/duck/Guangdong/04.22 DGCP080-O/2015(H5N6) | A / H5N6 | Asia / China | Yuhai Bi (Institute of Microbiology, Chinese Academy of Sciences) |
| EPI_ISL_199289 | A/duck/Guangdong/04.23 DGQTXC213-P/2015(H5N6) | A / H5N6 | Asia / China | Yuhai Bi (Institute of Microbiology, Chinese Academy of Sciences) |
| EPI_ISL_199290 | A/duck/Guangdong/04.23 DGQTXC236-P/2015(H5N6) | A / H5N6 | Asia / China | Yuhai Bi (Institute of Microbiology, Chinese Academy of Sciences) |
| EPI_ISL_199291 | A/duck/Guangdong/04.23 DGQTXC213-O/2015(H5N6) | A / H5N6 | Asia / China | Yuhai Bi (Institute of Microbiology, Chinese Academy of Sciences) |
| EPI_ISL_199292 | A/duck/Guangdong/04.23 DGQTXC232-P/2015(H5N6) | A / H5N6 | Asia / China | Yuhai Bi (Institute of Microbiology, Chinese Academy of Sciences) |
| EPI_ISL_199293 | A/duck/Guangdong/04.23 DGQTSJ131-O/2015(H5N6) | A / H5N6 | Asia / China | Yuhai Bi (Institute of Microbiology, Chinese Academy of Sciences) |
| EPI_ISL_199294 | A/goose/Guangdong/04.23 DGQTXC236-O/2015(H5N6) | A / H5N6 | Asia / China | Yuhai Bi (Institute of Microbiology, Chinese Academy of Sciences) |
| EPI_ISL_199295 | A/duck/Guangdong/04.23 DGQTXC228-P/2015(H5N6) | A / H5N6 | Asia / China | Yuhai Bi (Institute of Microbiology, Chinese Academy of Sciences) |
| EPI_ISL_199296 | A/pigeon/Guangdong/04.23 DGQTXC197-O/2015(H5N6) | A / H5N6 | Asia / China | Yuhai Bi (Institute of Microbiology, Chinese Academy of Sciences) |
| EPI_ISL_199297 | A/duck/Guangdong/04.23 DGQTXC235-O/2015(H5N6) | A / H5N6 | Asia / China | Yuhai Bi (Institute of Microbiology, Chinese Academy of Sciences) |
| EPI_ISL_199299 | A/duck/Guangdong/04.22 DGCP081-O/2015(H5N6) | A / H5N6 | Asia / China | Yuhai Bi (Institute of Microbiology, Chinese Academy of Sciences) |
| EPI_ISL_199301 | A/goose/Guangdong/04.23 DGQTXC237-O1/2015(H5N6) | A / H5N6 | Asia / China | Yuhai Bi (Institute of Microbiology, Chinese Academy of Sciences) |
| EPI_ISL_199302 | A/duck/Guangdong/04.23 DGQTSJ130-O/2015(H5N6) | A / H5N6 | Asia / China | Yuhai Bi (Institute of Microbiology, Chinese Academy of Sciences) |
| EPI_ISL_199303 | A/duck/Guangdong/04.22 DGCP084-O/2015(H5N6) | A / H5N6 | Asia / China | Yuhai Bi (Institute of Microbiology, Chinese Academy of Sciences) |
| EPI_ISL_199304 | A/duck/Guangdong/04.22 DGCPLB041-P/2015(H5N6) | A / H5N6 | Asia / China | Yuhai Bi (Institute of Microbiology, Chinese Academy of Sciences) |
| EPI_ISL_199305 | A/duck/Guangdong/04.22 DGCPLB044-P/2015(H5N6) | A / H5N6 | Asia / China | Yuhai Bi (Institute of Microbiology, Chinese Academy of Sciences) |
| EPI_ISL_199314 | A/environment/Jiangxi/01.14 NCJD005/2015(H5N6) | A / H5N6 | Asia / China | Yuhai Bi (Institute of Microbiology, Chinese Academy of Sciences) |
| EPI_ISL_199328 | A/duck/Jiangxi/02.05 YGYXG015-P/2015(H6N6) | A / H5N6 | Asia / China | Yuhai Bi (Institute of Microbiology, Chinese Academy of Sciences) |
| EPI_ISL_199337 | A/duck/Jiangxi/01.14 NCJD054-P/2015(H5N6) | A / H5N6 | Asia / China | Yuhai Bi (Institute of Microbiology, Chinese Academy of Sciences) |
| EPI_ISL_199341 | A/environment/Jiangxi/01.14 NCJD027/2015(H5N6) | A / H5N6 | Asia / China | Yuhai Bi (Institute of Microbiology, Chinese Academy of Sciences) |
| EPI_ISL_199342 | A/chicken/Jiangxi/01.14 NCJD181-P/2015(H5N6) | A / H5N6 | Asia / China | Yuhai Bi (Institute of Microbiology, Chinese Academy of Sciences) |
| EPI_ISL_199347 | A/duck/Guangdong/04.23 DGQTXC230-O/2015(H5N6) | A / H5N6 | Asia / China | Yuhai Bi (Institute of Microbiology, Chinese Academy of Sciences) |
| EPI_ISL_199380 | A/duck/Guangdong/04.23 DGQTXC235-P/2015(H5N6) | A / H5N6 | Asia / China | Yuhai Bi (Institute of Microbiology, Chinese Academy of Sciences) |
| EPI_ISL_199389 | A/environment/Guangdong/03.27 DGQTSJ046/2015(H5N6) | A / H5N6 | Asia / China | Yuhai Bi (Institute of Microbiology, Chinese Academy of Sciences) |
| EPI_ISL_199392 | A/environment/Guangdong/03.27 DGQTXC056/2015(H5N6) | A / H5N6 | Asia / China | Yuhai Bi (Institute of Microbiology, Chinese Academy of Sciences) |
| EPI_ISL_199397 | A/chicken/Jiangxi/01.14 NCJD125-P/2015(H5N6) | A / H5N6 | Asia / China | Yuhai Bi (Institute of Microbiology, Chinese Academy of Sciences) |
| EPI_ISL_199398 | A/duck/Jiangxi/02.06 NCJD039-P/2015(H5N6) | A / H5N6 | Asia / China | Yuhai Bi (Institute of Microbiology, Chinese Academy of Sciences) |
| EPI_ISL_199399 | A/duck/Jiangxi/02.06 NCJD036-P/2015(H5N6) | A / H5N6 | Asia / China | Yuhai Bi (Institute of Microbiology, Chinese Academy of Sciences) |
| EPI_ISL_199400 | A/chicken/Jiangxi/02.05 YGYXG035-P/2015(H5N6) | A / H5N6 | Asia / China | Yuhai Bi (Institute of Microbiology, Chinese Academy of Sciences) |
| EPI_ISL_199401 | A/chicken/Jiangxi/02.05 YGYXG023-P/2015(H5N6) | A / H5N6 | Asia / China | Yuhai Bi (Institute of Microbiology, Chinese Academy of Sciences) |
| EPI_ISL_199402 | A/duck/Jiangxi/02.05 YGYXG038-P/2015(H5N6) | A / H5N6 | Asia / China | Yuhai Bi (Institute of Microbiology, Chinese Academy of Sciences) |
| EPI_ISL_199403 | A/duck/Guangdong/03.26 DGCP062-P/2015(H5N6) | A / H5N6 | Asia / China | Yuhai Bi (Institute of Microbiology, Chinese Academy of Sciences) |
| EPI_ISL_199404 | A/duck/Guangdong/03.26 DGCPLB007-O/2015(H5N6) | A / H5N6 | Asia / China | Yuhai Bi (Institute of Microbiology, Chinese Academy of Sciences) |
| EPI_ISL_199405 | A/duck/Guangdong/03.26 DGCPLB022-O/2015(H5N6) | A / H5N6 | Asia / China | Yuhai Bi (Institute of Microbiology, Chinese Academy of Sciences) |
| EPI_ISL_199406 | A/duck/Guangdong/03.26 DGCP076-P/2015(H5N6) | A / H5N6 | Asia / China | Yuhai Bi (Institute of Microbiology, Chinese Academy of Sciences) |
| EPI_ISL_199407 | A/duck/Guangdong/03.26 DGCP077-O/2015(H5N6) | A / H5N6 | Asia / China | Yuhai Bi (Institute of Microbiology, Chinese Academy of Sciences) |
| EPI_ISL_199409 | A/duck/Guangdong/03.26 DGCP082-O/2015(H5N6) | A / H5N6 | Asia / China | Yuhai Bi (Institute of Microbiology, Chinese Academy of Sciences) |
| EPI_ISL_199410 | A/goose/Shandong/03.16 JNRe-5/2014(H5N6) | A / H5N6 | Asia / China | Yuhai Bi (Institute of Microbiology, Chinese Academy of Sciences) |
| EPI_ISL_199412 | A/duck/Shandong/11.14 RD/2013(H5N8) | A / H5N8 | Asia / China | Yuhai Bi (Institute of Microbiology, Chinese Academy of Sciences) |
| EPI_ISL_199414 | A/goose/Shandong/02.26 GD/2015(H5N6) | A / H5N6 | Asia / China | Yuhai Bi (Institute of Microbiology, Chinese Academy of Sciences) |
| EPI_ISL_199415 | A/duck/Guangdong/03.26 DGCPLB023-O/2015(H5N6) | A / H5N6 | Asia / China | Yuhai Bi (Institute of Microbiology, Chinese Academy of Sciences) |
| EPI_ISL_199416 | A/duck/Guangdong/03.26 DGCPLB024-O/2015(H5N6) | A / H5N6 | Asia / China | Yuhai Bi (Institute of Microbiology, Chinese Academy of Sciences) |
| EPI_ISL_199417 | A/duck/Guangdong/03.26 DGCP062-O/2015(H5N6) | A / H5N6 | Asia / China | Yuhai Bi (Institute of Microbiology, Chinese Academy of Sciences) |
| EPI_ISL_199419 | A/duck/Guangxi/04.10 JX046/2015(H5N6) | A / H5N6 | Asia / China | Yuhai Bi (Institute of Microbiology, Chinese Academy of Sciences) |
| EPI_ISL_199444 | A/chicken/Guangxi/04.10 NM140/2015(H5N6) | A / H5N6 | Asia / China | Yuhai Bi (Institute of Microbiology, Chinese Academy of Sciences) |
| EPI_ISL_199445 | A/environment/Guangdong/04.15 SZBAXQshui2/2015(H5N6) | A / H5N6 | Asia / China | Yuhai Bi (Institute of Microbiology, Chinese Academy of Sciences) |
| EPI_ISL_199446 | A/chicken/Guangdong/04.15 SZBAXQ019/2015(H5N6) | A / H5N6 | Asia / China | Yuhai Bi (Institute of Microbiology, Chinese Academy of Sciences) |
| EPI_ISL_199449 | A/duck/Guangdong/04.15 SZBAXQ011/2015(H5N6) | A / H5N6 | Asia / China | Yuhai Bi (Institute of Microbiology, Chinese Academy of Sciences) |
| EPI_ISL_199450 | A/duck/Guangdong/04.15 SZBAXQ017/2015(H5N6) | A / H5N6 | Asia / China | Yuhai Bi (Institute of Microbiology, Chinese Academy of Sciences) |
| EPI_ISL_199451 | A/duck/Guangdong/04.15 SZBAXQ020/2015(H5N6) | A / H5N6 | Asia / China | Yuhai Bi (Institute of Microbiology, Chinese Academy of Sciences) |
| EPI_ISL_199452 | A/duck/Guangdong/04.15 SZBAXQ021/2015(H5N6) | A / H5N6 | Asia / China | Yuhai Bi (Institute of Microbiology, Chinese Academy of Sciences) |
| EPI_ISL_199453 | A/duck/Guangdong/04.15 SZBAXQ003/2015(H5N6) | A / H5N6 | Asia / China | Yuhai Bi (Institute of Microbiology, Chinese Academy of Sciences) |
| EPI_ISL_199456 | A/pigeon/Guangdong/04.15 SZBAXQ004/2015(H5N6) | A / H5N6 | Asia / China | Yuhai Bi (Institute of Microbiology, Chinese Academy of Sciences) |
| EPI_ISL_199458 | A/duck/Guangdong/04.16 SZLGDBL013/2015(H5N6) | A / H5N6 | Asia / China | Yuhai Bi (Institute of Microbiology, Chinese Academy of Sciences) |
| EPI_ISL_199459 | A/duck/Guangdong/04.16 SZLGDBL002/2015(H5N6) | A / H5N6 | Asia / China | Yuhai Bi (Institute of Microbiology, Chinese Academy of Sciences) |
| EPI_ISL_199460 | A/duck/Guangdong/04.16 SZLGDBL003/2015(H5N6) | A / H5N6 | Asia / China | Yuhai Bi (Institute of Microbiology, Chinese Academy of Sciences) |
| EPI_ISL_199461 | A/duck/Guangdong/04.16 SZLGDBL005/2015(H5N6) | A / H5N6 | Asia / China | Yuhai Bi (Institute of Microbiology, Chinese Academy of Sciences) |
| EPI_ISL_199462 | A/duck/Guangdong/04.16 SZLGDBL009/2015(H5N6) | A / H5N6 | Asia / China | Yuhai Bi (Institute of Microbiology, Chinese Academy of Sciences) |

| | | | | |
|---|---|---|---|---|
| EPI_ISL_199463 | A/duck/Guangdong/04.16 SZLGWL002/2015(H5N6) | A / H5N6 | Asia / China | Yuhai Bi (Institute of Microbiology, Chinese Academy of Sciences) |
| EPI_ISL_199465 | A/pigeon/Guangdong/04.16 SZLGWL006/2015(H5N6) | A / H5N6 | Asia / China | Yuhai Bi (Institute of Microbiology, Chinese Academy of Sciences) |
| EPI_ISL_199466 | A/environment/Guangdong/04.16 SZLGWLShui1/2015(H5N6) | A / H5N6 | Asia / China | Yuhai Bi (Institute of Microbiology, Chinese Academy of Sciences) |
| EPI_ISL_199468 | A/environment/Guangdong/04.16 SZLGWLShui5/2015(H5N6) | A / H5N6 | Asia / China | Yuhai Bi (Institute of Microbiology, Chinese Academy of Sciences) |
| EPI_ISL_199536 | A/chicken/Montana/15-010559-1/2015 | A / H5N2 | North America / United States / Montana / Judith Basin County | |
| EPI_ISL_199546 | A/turkey/North Dakota/15-011420-13/2015 | A / H5N2 | North America / United States / North Dakota / Dickey County | |
| EPI_ISL_199547 | A/chicken/Nebraska/15-017897-1/2015 | A / H5N2 | North America / United States / Nebraska / Dixon County | |
| EPI_ISL_199548 | A/turkey/Wisconsin/15-012012-2/2015 | A / H5N2 | North America / United States / Wisconsin / Barron County | |
| EPI_ISL_199549 | A/turkey/North Dakota/15-013049-1/2015 | A / H5N2 | North America / United States / North Dakota / LaMoure County / commercial; 12 week old | |
| EPI_ISL_199550 | A/chicken/Minnesota/15-013533-1/2015 | A / H5N2 | North America / United States / Minnesota / Stearns County | |
| EPI_ISL_199551 | A/turkey/South Dakota/15-010371/2015 | A / H5N2 | North America / United States / South Dakota / Beadle County | |
| EPI_ISL_199552 | A/chicken/Nebraska/15-017990-5/2015 | A / H5N2 | North America / United States / Nebraska / Dixon County | |
| EPI_ISL_199553 | A/chicken/Wisconsin/15-011595-1/2015 | A / H5N2 | North America / United States / Wisconsin / Jefferson County | |
| EPI_ISL_199554 | A/chicken/Wisconsin/15-012160-1/2015 | A / H5N2 | North America / United States / Wisconsin / Juneau County | |
| EPI_ISL_200819 | A/chicken/Yunnan/02.09 DQXY008L/2015(H5N6) | A / H5N6 | Asia / China | Yuhai Bi (Institute of Microbiology, Chinese Academy of Sciences) |
| EPI_ISL_200820 | A/goose/Yunnan/07.14 DQWCK119/2015(H5N6) | A / H5N6 | Asia / China | Yuhai Bi (Institute of Microbiology, Chinese Academy of Sciences) |
| EPI_ISL_200821 | A/goose/Yunnan/07.14 DQWCK122/2015(H5N6) | A / H5N6 | Asia / China | Yuhai Bi (Institute of Microbiology, Chinese Academy of Sciences) |
| EPI_ISL_200822 | A/duck/Yunnan/07.15 DQNPH124/2015(H5N6) | A / H5N6 | Asia / China | Yuhai Bi (Institute of Microbiology, Chinese Academy of Sciences) |
| EPI_ISL_200823 | A/duck/Yunnan/07.15 DQNPH125/2015(H5N6) | A / H5N6 | Asia / China | Yuhai Bi (Institute of Microbiology, Chinese Academy of Sciences) |
| EPI_ISL_200824 | A/duck/Yunnan/07.15 DQNPH127/2015(H5N6) | A / H5N6 | Asia / China | Yuhai Bi (Institute of Microbiology, Chinese Academy of Sciences) |
| EPI_ISL_200825 | A/duck/Yunnan/07.15 DQNPH129/2015(H5N6) | A / H5N6 | Asia / China | Yuhai Bi (Institute of Microbiology, Chinese Academy of Sciences) |
| EPI_ISL_200826 | A/environment/Yunnan/07.11 DQENM015/2015(H5N6) | A / H5N6 | Asia / China | Yuhai Bi (Institute of Microbiology, Chinese Academy of Sciences) |
| EPI_ISL_200827 | A/environment/Yunnan/07.11 DQENM006/2015(H5N6) | A / H5N6 | Asia / China | Yuhai Bi (Institute of Microbiology, Chinese Academy of Sciences) |
| EPI_ISL_200828 | A/environment/Yunnan/07.11 DQENM004/2015(H5N6) | A / H5N6 | Asia / China | Yuhai Bi (Institute of Microbiology, Chinese Academy of Sciences) |
| EPI_ISL_200829 | A/chicken/Yunnan/07.11 DQENM005/2015(H5N6) | A / H5N6 | Asia / China | Yuhai Bi (Institute of Microbiology, Chinese Academy of Sciences) |
| EPI_ISL_200831 | A/environment/Yunnan/07.11 DQXY011/2015(H5N6) | A / H5N6 | Asia / China | Yuhai Bi (Institute of Microbiology, Chinese Academy of Sciences) |
| EPI_ISL_200832 | A/environment/Yunnan/07.11 DQXY20/2015(H5N6) | A / H5N6 | Asia / China | Yuhai Bi (Institute of Microbiology, Chinese Academy of Sciences) |
| EPI_ISL_200833 | A/environment/Yunnan/02.09 DQWGH013-2E/2015(H5N6) | A / H5N6 | Asia / China | Yuhai Bi (Institute of Microbiology, Chinese Academy of Sciences) |
| EPI_ISL_200834 | A/environment/Yunnan/02.09 DQXY005-1E/2015(H5N6) | A / H5N6 | Asia / China | Yuhai Bi (Institute of Microbiology, Chinese Academy of Sciences) |
| EPI_ISL_200835 | A/environment/Yunnan/02.09 DQXY009-1E/2015(H5N6) | A / H5N6 | Asia / China | Yuhai Bi (Institute of Microbiology, Chinese Academy of Sciences) |
| EPI_ISL_200836 | A/environment/Yunnan/02.10 DQXY053-1E/2015(H5N6) | A / H5N6 | Asia / China | Yuhai Bi (Institute of Microbiology, Chinese Academy of Sciences) |
| EPI_ISL_200837 | A/Yunnan/DQ001/2015(H5N6) | A / H5N6 | Asia / China / Yunnan Province | Yuhai Bi (Institute of Microbiology, Chinese Academy of Sciences) |
| EPI_ISL_200838 | A/Yunnan/DQ002/2015(H5N6) | A / H5N6 | Asia / China / Yunnan Province | Yuhai Bi (Institute of Microbiology, Chinese Academy of Sciences) |
| EPI_ISL_201355 | A/duck/Fujian/05.07 FZHMDK14-O/2015(H5N6) | A / H5N6 | Asia / China | Yuhai Bi (Institute of Microbiology, Chinese Academy of Sciences) |
| EPI_ISL_201378 | A/duck/Jiangxi/09.04 JX-14-11/2014(H5N6) | A / H5N6 | Asia / China / Jiangxi | Yuhai Bi (Institute of Microbiology, Chinese Academy of Sciences) |
| EPI_ISL_201798 | A/goose/Guangdong/s13124/2013 | A / H5N8 | Asia / China | |
| EPI_ISL_201799 | A/duck/Guangdong/s14044/2014 | A / H5N8 | Asia / China | |
| EPI_ISL_202553 | A/chicken/Guizhou/4/2013 | A / H5N1 | Asia / China | Huihui Kong (Harbin Veterinary Research Institute (CAAS) / Ministry of Agriculture) |
| EPI_ISL_202554 | A/chicken/Sichuan/J1/2014 | A / H5N6 | Asia / China | Huihui Kong (Harbin Veterinary Research Institute (CAAS) / Ministry of Agriculture) |
| EPI_ISL_202555 | A/chicken/Yunnan/3/2014 | A / H5N1 | Asia / China | Huihui Kong (Harbin Veterinary Research Institute (CAAS) / Ministry of Agriculture) |
| EPI_ISL_202556 | A/duck/Anhui/S1300/2014 | A / H5N6 | Asia / China | Huihui Kong (Harbin Veterinary Research Institute (CAAS) / Ministry of Agriculture) |
| EPI_ISL_202558 | A/duck/Liaoning/S1001/2014 | A / H5N8 | Asia / China | Huihui Kong (Harbin Veterinary Research Institute (CAAS) / Ministry of Agriculture) |
| EPI_ISL_202559 | A/wild bird/Liaoning/S1660/2014 | A / H5N2 | Asia / China | Huihui Kong (Harbin Veterinary Research Institute (CAAS) / Ministry of Agriculture) |
| EPI_ISL_202763 | A/environment/GZ/75/2014 | A / H5N6 | Asia / China / Guangdong | Jay Zhou (The University of Hong Kong / Department of Microbiology) |
| EPI_ISL_202764 | A/environment/GZ/76/2014 | A / H5N2 | Asia / China / Guangdong | Jay Zhou (The University of Hong Kong / Department of Microbiology) |
| EPI_ISL_202765 | A/environment/GZ/77/2014 | A / H5N6 | Asia / China / Guangdong | Jay Zhou (The University of Hong Kong / Department of Microbiology) |
| EPI_ISL_202766 | A/environment/GZ/NIOSH-187/2014 | A / H5N6 | Asia / China / Guangdong Province / Guangdong | Jay Zhou (The University of Hong Kong / Department of Microbiology) |
| EPI_ISL_203646 | A/Environment/Guangdong/79900/2014 | A / H5N6 | Asia / China / Guangdong | Lei Yang (WHO Chinese National Influenza Center / Virology Institute, Chinese CDC) |
| EPI_ISL_203647 | A/Environment/Guangdong/79478/2014 | A / H5N6 | Asia / China / Guangdong | Lei Yang (WHO Chinese National Influenza Center / Virology Institute, Chinese CDC) |
| EPI_ISL_203648 | A/Environment/Guangdong/79929/2014 | A / H5N6 | Asia / China / Guangdong | Lei Yang (WHO Chinese National Influenza Center / Virology Institute, Chinese CDC) |
| EPI_ISL_203649 | A/Environment/Chongqing/28975/2014 | A / H5N6 | Asia / China / Chongqing | Lei Yang (WHO Chinese National Influenza Center / Virology Institute, Chinese CDC) |
| EPI_ISL_203650 | A/Environment/Chongqing/72076/2014 | A / H5N6 | Asia / China / Chongqing | Lei Yang (WHO Chinese National Influenza Center / Virology Institute, Chinese CDC) |
| EPI_ISL_203651 | A/Environment/Chongqing/28957/2014 | A / H5N6 | Asia / China / Chongqing | Lei Yang (WHO Chinese National Influenza Center / Virology Institute, Chinese CDC) |
| EPI_ISL_203652 | A/Environment/Hunan/18327/2014 | A / H5N6 | Asia / China / Hunan | Lei Yang (WHO Chinese National Influenza Center / Virology Institute, Chinese CDC) |
| EPI_ISL_203653 | A/Environment/Zhejiang/21574/2014 | A / H5N6 | Asia / China / Zhejiang | Lei Yang (WHO Chinese National Influenza Center / Virology Institute, Chinese CDC) |
| EPI_ISL_203654 | A/Environment/Jiangxi/10733/2014 | A / H5N6 | Asia / China / Jiangxi | Lei Yang (WHO Chinese National Influenza Center / Virology Institute, Chinese CDC) |
| EPI_ISL_203655 | A/Environment/Chongqing/28970/2014 | A / H5N6 | Asia / China / Chongqing | Lei Yang (WHO Chinese National Influenza Center / Virology Institute, Chinese CDC) |
| EPI_ISL_203731 | A/duck/Zhejiang/77138/2014 | A / H5N2 | Asia / China | |
| EPI_ISL_203732 | A/duck/Zhejiang/727041/2014 | A / H5N2 | Asia / China | |
| EPI_ISL_203733 | A/goose/Zhejiang/77166/2014 | A / H5N2 | Asia / China | |
| EPI_ISL_203734 | A/goose/Zhejiang/77167/2014 | A / H5N2 | Asia / China | |
| EPI_ISL_203735 | A/goose/Zhejiang/77168/2014 | A / H5N2 | Asia / China | |
| EPI_ISL_203736 | A/chicken/Zhejiang/727159/2014 | A / H5N2 | Asia / China | |
| EPI_ISL_203737 | A/chicken/Zhejiang/727079/2014 | A / H5N2 | Asia / China | |
| EPI_ISL_203738 | A/duck/Zhejiang/727158/2014 | A / H5N6 | Asia / China | |
| EPI_ISL_203739 | A/goose/Zhejiang/727092/2014 | A / H5N6 | Asia / China | |
| EPI_ISL_203740 | A/goose/Zhejiang/727110/2014 | A / H5N6 | Asia / China | |
| EPI_ISL_203741 | A/goose/Zhejiang/925036/2014 | A / H5N6 | Asia / China | |
| EPI_ISL_203742 | A/goose/Zhejiang/925105/2014 | A / H5N6 | Asia / China | |
| EPI_ISL_203743 | A/goose/Zhejiang/925106/2014 | A / H5N6 | Asia / China | |
| EPI_ISL_203744 | A/goose/Zhejiang/925108/2014 | A / H5N6 | Asia / China | |
| EPI_ISL_203745 | A/goose/Zhejiang/112080/2014 | A / H5N6 | Asia / China | |

| | | | | |
|---|---|---|---|---|
| EPI_ISL_203746 | A/goose/Zhejiang/1120132/2014 | A / H5N6 | Asia / China | |
| EPI_ISL_203747 | A/chicken/Zhejiang/727022/2014 | A / H5N6 | Asia / China | |
| EPI_ISL_203748 | A/chicken/Zhejiang/727026/2014 | A / H5N6 | Asia / China | |
| EPI_ISL_203749 | A/chicken/Zhejiang/727155/2014 | A / H5N6 | Asia / China | |
| EPI_ISL_203750 | A/duck/Zhejiang/925019/2014 | A / H5N8 | Asia / China | |
| EPI_ISL_203751 | A/duck/Zhejiang/925169/2014 | A / H5N8 | Asia / China | |
| EPI_ISL_203752 | A/goose/Zhejiang/925037/2014 | A / H5N8 | Asia / China | |
| EPI_ISL_203753 | A/goose/Zhejiang/925104/2014 | A / H5N8 | Asia / China | |
| EPI_ISL_205115 | A/duck/Wuhan/WHYF02/2015 | A / H5N6 | Asia / China | |
| EPI_ISL_205116 | A/duck/Wuhan/WHYF03/2015 | A / H5N6 | Asia / China | |
| EPI_ISL_205117 | A/chicken/Wuhan/WHYJ01/2015 | A / H5N6 | Asia / China | |
| EPI_ISL_205118 | A/chicken/Wuhan/WHYJ02/2015 | A / H5N6 | Asia / China | |
| EPI_ISL_205119 | A/duck/Wenzhou/YHQL22/2014 | A / H5N6 | Asia / China | |
| EPI_ISL_205120 | A/duck/Wuhan/JXYFB22/2015 | A / H5N6 | Asia / China | |
| EPI_ISL_205121 | A/duck/Taizhou/TZYG12/2015 | A / H5N6 | Asia / China | |
| EPI_ISL_205137 | A/turtledove/Wuhan/WHBJ12/2014 | A / H5N6 | Asia / China | |
| EPI_ISL_205138 | A/turtledove/Wuhan/WHBJ28/2014 | A / H5N6 | Asia / China | |
| EPI_ISL_205139 | A/turtledove/Wuhan/HKBJ27/2015 | A / H5N6 | Asia / China | |
| EPI_ISL_205140 | A/turtledove/Wuhan/HKBJ43/2015 | A / H5N6 | Asia / China | |
| EPI_ISL_205141 | A/turtledove/Wuhan/WHBJ26/2014 | A / H5N6 | Asia / China | |
| EPI_ISL_205142 | A/turtledove/Wuhan/HKBJ07/2015 | A / H5N6 | Asia / China | |
| EPI_ISL_205313 | A/Shenzhen/1/2015 | A / H5N6 | Asia / China / Guangdong Province / Guangdong | Lei Yang (WHO Chinese National Influenza Center / Virology Institute, Chinese CDC) |
| EPI_ISL_205314 | A/Environment/Shenzhen/1/2015 | A / H5N6 | Asia / China / Guangdong | Lei Yang (WHO Chinese National Influenza Center / Virology Institute, Chinese CDC) |
| EPI_ISL_205315 | A/Environment/Shenzhen/2/2015 | A / H5N6 | Asia / China / Guangdong | Lei Yang (WHO Chinese National Influenza Center / Virology Institute, Chinese CDC) |
| EPI_ISL_205316 | A/Environment/Shenzhen/3/2015 | A / H5N6 | Asia / China / Guangdong | Lei Yang (WHO Chinese National Influenza Center / Virology Institute, Chinese CDC) |
| EPI_ISL_205503 | A/Shenzhen/TH001/2015 (H5N6) | A / H5N6 | Asia / China / Guangdong Province | Yuhai Bi (Institute of Microbiology, Chinese Academy of Sciences) |
| EPI_ISL_205821 | A/peregrine_falcon/Hong_Kong/04955/2015 | A / H5N6 | Asia / Hong Kong (SAR) | Tak Wai Cheng (Agriculture, Fisheries and Conservation Department / Tai Lung Veterinary Laboratory) |
| EPI_ISL_205823 | A/Oriental_magpie_robin/Hong_Kong/06154/2015 | A / H5N6 | Asia / Hong Kong (SAR) | Tak Wai Cheng (Agriculture, Fisheries and Conservation Department / Tai Lung Veterinary Laboratory) |
| EPI_ISL_205824 | A/Oriental_magpie_robin/Hong_Kong/16759/2015 | A / H5N6 | Asia / Hong Kong (SAR) | Tak Wai Cheng (Agriculture, Fisheries and Conservation Department / Tai Lung Veterinary Laboratory) |
| EPI_ISL_205825 | A/great_egret/Hong_Kong/00032/2016 | A / H5N6 | Asia / Hong Kong (SAR) | Tak Wai Cheng (Agriculture, Fisheries and Conservation Department / Tai Lung Veterinary Laboratory) |
| EPI_ISL_205856 | A/chicken/Yunnan/1/2014 | A / H5N1 | Asia / China | |
| EPI_ISL_205959 | A/Duck/Guangdong/SS1A1/2014?H5N6? | A / H5N6 | Asia / China / Guangdong Province | Qi Wenbao (South China Agricultural University / Veterinary Medicine College) |
| EPI_ISL_205961 | A/Goose/ Guangdong /JG125/2013?H5N6? | A / H5N6 | Asia / China / Guangdong Province | Qi Wenbao (South China Agricultural University / Veterinary Medicine College) |
| EPI_ISL_205962 | A/Duck/Guangdong/PY501/2014?H5N6? | A / H5N6 | Asia / China / Guangdong Province | Qi Wenbao (South China Agricultural University / Veterinary Medicine College) |
| EPI_ISL_205963 | A/Duck/Guangdong/PY503/2014?H5N6? | A / H5N6 | Asia / China / Guangdong Province | Qi Wenbao (South China Agricultural University / Veterinary Medicine College) |
| EPI_ISL_205964 | A/Chicken/Guangdong/FG594/2015?H5N6? | A / H5N6 | Asia / China / Guangdong Province | Qi Wenbao (South China Agricultural University / Veterinary Medicine College) |
| EPI_ISL_205965 | A/Enviroment/Guangdong/SS127/2013(H5N6) | A / H5N6 | Asia / China / Guangdong Province | Qi Wenbao (South China Agricultural University / Veterinary Medicine College) |
| EPI_ISL_205966 | A/Goose/ Guangdong /SSLBY/2015?H5N6? | A / H5N6 | Asia / China / Guangdong Province | Qi Wenbao (South China Agricultural University / Veterinary Medicine College) |
| EPI_ISL_205967 | A/Goose/ Guangdong /SSZYP/2015?H5N6? | A / H5N6 | Asia / China / Guangdong Province | Qi Wenbao (South China Agricultural University / Veterinary Medicine College) |
| EPI_ISL_206036 | A/Shenzhen/1/2016 | A / H5N6 | Asia / China / Guangdong | Lei Yang (WHO Chinese National Influenza Center / Virology Institute, Chinese CDC) |
| EPI_ISL_206394 | A/snow goose/Missouri/15-011246-1/2015 | A / H5N2 | North America / United States | |
| EPI_ISL_206395 | A/American wigeon/Utah/AH0007824/2015 | A / H5N8 | North America / United States | |
| EPI_ISL_206396 | A/mallard/Oregon/AH0003952/2015 | A / H5N2 | North America / United States | |
| EPI_ISL_206397 | A/Northern pintail/Oregon/AH0003967/2015 | A / H5N2 | North America / United States | |
| EPI_ISL_206398 | A/wood duck/Oregon/AH0007244/2015 | A / H5N2 | North America / United States | |
| EPI_ISL_206399 | A/wood duck/Oregon/AH0007257/2015 | A / H5N2 | North America / United States | |
| EPI_ISL_206400 | A/wood duck/Oregon/AH0007263/2015 | A / H5N2 | North America / United States | |
| EPI_ISL_206401 | A/Northern shoveler/Oregon/AH0007332/2015 | A / H5N2 | North America / United States | |
| EPI_ISL_206402 | A/Northern shoveler/Oregon/AH0007337/2015 | A / H5N2 | North America / United States | |
| EPI_ISL_206403 | A/Northern shoveler/Oregon/AH0007339/2015 | A / H5N2 | North America / United States | |
| EPI_ISL_206404 | A/mallard/Idaho/AH0008597/2015 | A / H5N8 | North America / United States | |
| EPI_ISL_206405 | A/mallard/Oregon/AH0008887/2015 | A / H5N2 | North America / United States | |
| EPI_ISL_206406 | A/Northern pintail/Oregon/AH0003871/2015 | A / H5N2 | North America / United States | |
| EPI_ISL_206407 | A/mallard/Idaho/AH0007412/2015 | A / H5N2 | North America / United States | |
| EPI_ISL_206408 | A/mallard/Idaho/AH0007413/2015 | A / H5N2 | North America / United States | |
| EPI_ISL_206409 | A/American green-winged teal/Idaho/AH0011899/2015 | A / H5N2 | North America / United States | |
| EPI_ISL_206410 | A/American green-winged teal/Oregon/AH0012403/2015 | A / H5N2 | North America / United States | |
| EPI_ISL_206411 | A/Canada goose/Oregon/AH0012452/2015 | A / H5N8 | North America / United States | |
| EPI_ISL_206412 | A/American wigeon/Oregon/AH0012525/2015 | A / H5N8 | North America / United States | |
| EPI_ISL_206413 | A/mallard/Nevada/AH0006855/2015 | A / H5N8 | North America / United States | |
| EPI_ISL_206415 | A/chicken/California/15-004912/2015 | A / H5N8 | North America / United States | |
| EPI_ISL_206429 | A/mallard/Idaho/AH0005954/2014 | A / H5N8 | North America / United States | |
| EPI_ISL_206430 | A/mallard/Idaho/AH0005955/2014 | A / H5N8 | North America / United States | |
| EPI_ISL_206431 | A/mallard/Oregon/AH0003821/2014 | A / H5N2 | North America / United States | |
| EPI_ISL_206432 | A/bald eagle/Idaho/15-002892-2/2015 | A / H5N8 | North America / United States | |
| EPI_ISL_206433 | A/Northern pintail/Washington/195365/2014 | A / H5N2 | North America / United States | |
| EPI_ISL_206434 | A/mallard/Oregon/195547/2014 | A / H5N2 | North America / United States | |
| EPI_ISL_206435 | A/American wigeon/Washington/195968/2014 | A / H5N8 | North America / United States | |
| EPI_ISL_206436 | A/mallard/Washington/196262/2015 | A / H5N2 | North America / United States | |
| EPI_ISL_206437 | A/Northern pintail/Washington/196271/2014 | A / H5N8 | North America / United States | |
| EPI_ISL_206438 | A/mallard/Washington/196865/2015 | A / H5N2 | North America / United States | |
| EPI_ISL_206439 | A/American wigeon/Washington/196336/2015 | A / H5N1 | North America / United States | |

| | | | | |
|---|---|---|---|---|
| EPI_ISL_206440 | A/American wigeon/Washington/196340/2015 | A / H5N1 | North America / United States | |
| EPI_ISL_206441 | A/mallard/Oregon/195536/2014 | A / H5N8 | North America / United States | |
| EPI_ISL_206442 | A/American wigeon/Washington/195198/2014 | A / H5N8 | North America / United States | |
| EPI_ISL_206443 | A/American wigeon/Washington/195205/2014 | A / H5N8 | North America / United States | |
| EPI_ISL_206444 | A/mallard/Washington/195246/2014 | A / H5N2 | North America / United States | |
| EPI_ISL_206445 | A/peregrine falcon/Washington/196426/2014 | A / H5N8 | North America / United States | |
| EPI_ISL_206446 | A/mallard/Washington/195810/2014 | A / H5N2 | North America / United States | |
| EPI_ISL_206447 | A/Canada goose/Washington/197619/2014 | A / H5N2 | North America / United States | |
| EPI_ISL_206448 | A/Cooper's hawk/Minnesota/198225/2015 | A / H5N2 | North America / United States | |
| EPI_ISL_206449 | A/snowy owl/Wisconsin/198399/2015 | A / H5N2 | North America / United States | |
| EPI_ISL_206450 | A/Canada goose/Kansas/197850/2015 | A / H5N2 | North America / United States | |
| EPI_ISL_206456 | A/goose/Taiwan/TNC1/2015 | A / H5N8 | Asia / Taiwan | |
| EPI_ISL_206457 | A/goose/Taiwan/TNC2/2015 | A / H5N8 | Asia / Taiwan | |
| EPI_ISL_206458 | A/goose/Taiwan/TNC3/2015 | A / H5N8 | Asia / Taiwan | |
| EPI_ISL_206459 | A/goose/Taiwan/TNC4/2015 | A / H5N8 | Asia / Taiwan | |
| EPI_ISL_206460 | A/goose/Taiwan/TNC5/2015 | A / H5N8 | Asia / Taiwan | |
| EPI_ISL_206461 | A/goose/Taiwan/TNC6/2015 | A / H5N8 | Asia / Taiwan | |
| EPI_ISL_206462 | A/goose/Taiwan/TNC7/2015 | A / H5N8 | Asia / Taiwan | |
| EPI_ISL_206463 | A/goose/Taiwan/TNC8/2015 | A / H5N8 | Asia / Taiwan | |
| EPI_ISL_206464 | A/goose/Taiwan/TNC9/2015 | A / H5N8 | Asia / Taiwan | |
| EPI_ISL_206465 | A/goose/Taiwan/TNC10/2015 | A / H5N8 | Asia / Taiwan | |
| EPI_ISL_206466 | A/goose/Taiwan/TNC11/2015 | A / H5N8 | Asia / Taiwan | |
| EPI_ISL_206467 | A/goose/Taiwan/TNC12/2015 | A / H5N8 | Asia / Taiwan | |
| EPI_ISL_206468 | A/goose/Taiwan/TNC13/2015 | A / H5N8 | Asia / Taiwan | |
| EPI_ISL_206469 | A/goose/Taiwan/TNC14/2015 | A / H5N8 | Asia / Taiwan | |
| EPI_ISL_206470 | A/goose/Taiwan/TNO1/2015 | A / H5N8 | Asia / Taiwan | |
| EPI_ISL_206471 | A/goose/Taiwan/TNO2/2015 | A / H5N8 | Asia / Taiwan | |
| EPI_ISL_206472 | A/goose/Taiwan/TNO3/2015 | A / H5N8 | Asia / Taiwan | |
| EPI_ISL_206473 | A/goose/Taiwan/TNO4/2015 | A / H5N8 | Asia / Taiwan | |
| EPI_ISL_206475 | A/goose/Taiwan/TNO6/2015 | A / H5N8 | Asia / Taiwan | |
| EPI_ISL_206476 | A/goose/Taiwan/TNO7/2015 | A / H5N8 | Asia / Taiwan | |
| EPI_ISL_206477 | A/goose/Taiwan/TNO8/2015 | A / H5N8 | Asia / Taiwan | |
| EPI_ISL_206478 | A/goose/Taiwan/TNO9/2015 | A / H5N8 | Asia / Taiwan | |
| EPI_ISL_206479 | A/goose/Taiwan/TNO10/2015 | A / H5N8 | Asia / Taiwan | |
| EPI_ISL_206480 | A/goose/Taiwan/TNO11/2015 | A / H5N8 | Asia / Taiwan | |
| EPI_ISL_206481 | A/goose/Taiwan/TNO12/2015 | A / H5N8 | Asia / Taiwan | |
| EPI_ISL_206482 | A/goose/Taiwan/TNO13/2015 | A / H5N8 | Asia / Taiwan | |
| EPI_ISL_206483 | A/goose/Taiwan/TNO14/2015 | A / H5N8 | Asia / Taiwan | |
| EPI_ISL_206484 | A/goose/Taiwan/TNO15/2015 | A / H5N8 | Asia / Taiwan | |
| EPI_ISL_206485 | A/goose/Taiwan/TNO16/2015 | A / H5N8 | Asia / Taiwan | |
| EPI_ISL_206486 | A/goose/Taiwan/TNO17/2015 | A / H5N8 | Asia / Taiwan | |
| EPI_ISL_206487 | A/goose/Taiwan/TNO18/2015 | A / H5N8 | Asia / Taiwan | |
| EPI_ISL_206488 | A/goose/Taiwan/TNO19/2015 | A / H5N8 | Asia / Taiwan | |
| EPI_ISL_206489 | A/goose/Taiwan/TNO20/2015 | A / H5N8 | Asia / Taiwan | |
| EPI_ISL_206490 | A/goose/Jiangsu/QD5/2014 | A / H5N8 | Asia / China | |
| EPI_ISL_206491 | A/goose/Shandong/WFSG1/2014 | A / H5N8 | Asia / China | |
| EPI_ISL_206492 | A/goose/Yangzhou/0420/2014 | A / H5N8 | Asia / China | |
| EPI_ISL_206568 | A/ Guangdong /SZ872/2015?H5N6? | A / H5N6 | Asia / China / Guangdong Province | Qi Wenbao (South China Agricultural University / Veterinary Medicine College) |
| EPI_ISL_206569 | A/ Guangdong /ZQ874/2015?H5N6? | A / H5N6 | Asia / China / Guangdong Province | Qi Wenbao (South China Agricultural University / Veterinary Medicine College) |
| EPI_ISL_206675 | A/environment/Yunnan/YN19/2015 | A / H5N6 | Asia / China | |
| EPI_ISL_206676 | A/environment/Yunnan/YN22/2015 | A / H5N6 | Asia / China | |
| EPI_ISL_206677 | A/environment/Yunnan/YN25/2015 | A / H5N6 | Asia / China | |
| EPI_ISL_206773 | A/gull/Germany-NI/R45/2015 | A / H5N8 | Europe / Germany / Lower Saxony | Elke Starick (Friedrich-Loeffler-Institut) |
| EPI_ISL_207048 | A/Shenzhen/TH003/2016(H5N6) | A / H5N6 | Asia / China / Guangdong Province | Yuhai Bi (Institute of Microbiology, Chinese Academy of Sciences) |
| EPI_ISL_207051 | A/Shenzhen/TH002/2016(H5N6) | A / H5N6 | Asia / China / Guangdong Province | Yuhai Bi (Institute of Microbiology, Chinese Academy of Sciences) |
| EPI_ISL_2081527 | A/red_fox/England/AVP-M1-21-01/2020 | A / H5N8 | Europe / United Kingdom / Norfolk | Alex Byrne (Animal and Plant Health Agency (APHA) / Virology Department) |
| EPI_ISL_2081528 | A/seal/England/AVP-031141/2020 | A / H5N8 | Europe / United Kingdom / Norfolk | Alex Byrne (Animal and Plant Health Agency (APHA) / Virology Department) |
| EPI_ISL_208822 | A/duck/Eastern China/L0321/2010 | A / H5N2 | Asia / China | |
| EPI_ISL_208823 | A/duck/Eastern China/L0230/2010 | A / H5N2 | Asia / China | |
| EPI_ISL_208824 | A/duck/Eastern China/L0405/2010 | A / H5N8 | Asia / China | |
| EPI_ISL_208825 | A/duck/Eastern China/L0423/2011 | A / H5N8 | Asia / China | |
| EPI_ISL_208826 | A/duck/Eastern China/L0611/2011 | A / H5N8 | Asia / China | |
| EPI_ISL_208827 | A/duck/Eastern China/L0722/2012 | A / H5N8 | Asia / China | |
| EPI_ISL_208828 | A/duck/Eastern China/L1021/2012 | A / H5N8 | Asia / China | |
| EPI_ISL_208829 | A/duck/Eastern China/L1120/2012 | A / H5N8 | Asia / China | |
| EPI_ISL_208830 | A/goose/Eastern China/L1214/2012 | A / H5N8 | Asia / China | |
| EPI_ISL_208831 | A/goose/Eastern China/L1204/2012 | A / H5N8 | Asia / China | |
| EPI_ISL_208832 | A/duck/Eastern China/S1210/2013 | A / H5N8 | Asia / China | |
| EPI_ISL_208833 | A/goose/Eastern China/S0513/2013 | A / H5N6 | Asia / China | |
| EPI_ISL_208834 | A/duck/Eastern China/S0131/2014 | A / H5N2 | Asia / China | |
| EPI_ISL_208835 | A/duck/Eastern China/S0215/2014 | A / H5N8 | Asia / China | |
| EPI_ISL_208836 | A/duck/Eastern China/S0711/2014 | A / H5N6 | Asia / China | |

| | | | | |
|---|---|---|---|---|
| EPI_ISL_208837 | A/duck/Eastern China/S0808/2014 | A / H5N2 | Asia / China | |
| EPI_ISL_208838 | A/duck/Eastern China/S0908/2014 | A / H5N6 | Asia / China | |
| EPI_ISL_208839 | A/duck/Eastern China/S0322/2014 | A / H5N6 | Asia / China | |
| EPI_ISL_208840 | A/goose/Eastern China/S0408/2014 | A / H5N8 | Asia / China | |
| EPI_ISL_208841 | A/duck/Eastern China/S1109/2014 | A / H5N8 | Asia / China | |
| EPI_ISL_2111625 | A/mute_swan/Poland/MB131/2021 | A / H5N8 | Europe / Poland / Pomeranian Voivodeship / Puck Bay (W?adys?awowo) | Kamila Dziadek (National Veterinary Research Institute / Department of Poultry Diseases) |
| EPI_ISL_2113167 | A/mute_swan/Poland/MB189/2021 | A / H5N8 | Europe / Poland / Pomeranian Voivodeship / Gdynia | Kamila Dziadek (National Veterinary Research Institute / Department of Poultry Diseases) |
| EPI_ISL_2113493 | A/mute_swan/Poland/MB268/2021 | A / H5N8 | Europe / Poland / Pomeranian Voivodeship / Puck Bay (Jastarnia) | Kamila Dziadek (National Veterinary Research Institute / Department of Poultry Diseases) |
| EPI_ISL_2114013 | A/mute_swan/Poland/MB272/2021 | A / H5N8 | Europe / Poland / Pomeranian Voivodeship / Stegna | Kamila Dziadek (National Veterinary Research Institute / Department of Poultry Diseases) |
| EPI_ISL_2131573 | A/Chicken/Sweden/SVA210420SZ0002/KN099429-IP22/2021 | A / H5N8 | Europe / Sweden / Skane Lan / Eslovs Kommun | Siamak Zohari (National Veterinary Institute) |
| EPI_ISL_213782 | A/ostrich/Korea/H829/2014 | A / H5N8 | Asia / Korea, Republic of | |
| EPI_ISL_217025 | A/Changsha/1/2014 | A / H5N6 | Asia / China | |
| EPI_ISL_2172517 | A/common buzzard/Netherlands/21021278-002/2021 | A / H5N8 | Europe / Netherlands / Provincie Friesland / Ferwert | Rene Heutink (Wageningen Bioveterinary Research) |
| EPI_ISL_2172518 | A/common kestrel/Netherlands/21021301-039/2021 | A / H5N8 | Europe / Netherlands / South Holland / Strijen | Rene Heutink (Wageningen Bioveterinary Research) |
| EPI_ISL_2172522 | A/common buzzard/Netherlands/21021497-001/2021 | A / H5N8 | Europe / Netherlands / Provincie Noord-Holland / Rijssenhout | Rene Heutink (Wageningen Bioveterinary Research) |
| EPI_ISL_2172523 | A/sanderling/Netherlands/21021794-002/2021 | A / H5N8 | Europe / Netherlands / South Holland / Hoogvliet | Rene Heutink (Wageningen Bioveterinary Research) |
| EPI_ISL_2172525 | A/barnacle goose/Netherlands/21022039-002/2021 | A / H5N8 | Europe / Netherlands / South Holland / Oostvoorne | Rene Heutink (Wageningen Bioveterinary Research) |
| EPI_ISL_2172526 | A/common buzzard/Netherlands/21023939-001/2021 | A / H5N8 | Europe / Netherlands / Provincie Gelderland / Rha (Bronckhorst) | Rene Heutink (Wageningen Bioveterinary Research) |
| EPI_ISL_2172528 | A/barnacle goose/Netherlands/21024066-001/2021 | A / H5N1 | Europe / Netherlands / Provincie Friesland / Nes | Rene Heutink (Wageningen Bioveterinary Research) |
| EPI_ISL_2172529 | A/wild goose/Netherlands/21024076-001/2021 | A / H5N8 | Europe / Netherlands / Provincie Gelderland / Apeldoorn | Rene Heutink (Wageningen Bioveterinary Research) |
| EPI_ISL_2172530 | A/common buzzard/Netherlands/21024357-002/2021 | A / H5N3 | Europe / Netherlands / Provincie Friesland / Oostbierum | Rene Heutink (Wageningen Bioveterinary Research) |
| EPI_ISL_2172531 | A/barnacle goose/Netherlands/21024358-001/2021 | A / H5N8 | Europe / Netherlands / Provincie Noord-Holland / Hoorn | Rene Heutink (Wageningen Bioveterinary Research) |
| EPI_ISL_2172532 | A/gadwall/Netherlands/21024401-002/2021 | A / H5N8 | Europe / Netherlands / South Holland / Abbenbroek | Rene Heutink (Wageningen Bioveterinary Research) |
| EPI_ISL_2172533 | A/common buzzard/Netherlands/21024712-002/2021 | A / H5N8 | Europe / Netherlands / Provincie Groningen / Leek | Rene Heutink (Wageningen Bioveterinary Research) |
| EPI_ISL_2172608 | A/barnacle goose/Netherlands/21024897-001/2021 | A / H5N8 | Europe / Netherlands / South Holland / Rijswijk | Rene Heutink (Wageningen Bioveterinary Research) |
| EPI_ISL_2173364 | A/peregrine falcon/Netherlands/21025108-001/2021 | A / H5N4 | Europe / Netherlands / South Holland / Brielle | Rene Heutink (Wageningen Bioveterinary Research) |
| EPI_ISL_2174084 | A/common murre/Netherlands/21025491-002/2021 | A / H5N1 | Europe / Netherlands / Provincie Friesland / Schiermonnikoog | Rene Heutink (Wageningen Bioveterinary Research) |
| EPI_ISL_2174728 | A/barnacle goose/Netherlands/21025769-002/2021 | A / H5N1 | Europe / Netherlands / Provincie Groningen / Termunten | Rene Heutink (Wageningen Bioveterinary Research) |
| EPI_ISL_2175628 | A/common buzzard/Netherlands/21021187-001/2021 | A / H5N8 | Europe / Netherlands / Provincie Groningen / Niehoeve | Rene Heutink (Wageningen Bioveterinary Research) |
| EPI_ISL_2176321 | A/barnacle goose/Netherlands/21023498-002/2021 | A / H5N8 | Europe / Netherlands / Provincie Noord-Holland / Oosterend | Rene Heutink (Wageningen Bioveterinary Research) |
| EPI_ISL_2176841 | A/barnacle goose/Netherlands/21023501-002/2021 | A / H5N8 | Europe / Netherlands / Provincie Noord-Holland / Den Oever | Rene Heutink (Wageningen Bioveterinary Research) |
| EPI_ISL_2177398 | A/peacock/Netherlands/21026542-001/2021 | A / H5N8 | Europe / Netherlands / Provincie Noord-Holland / Hoorn | Rene Heutink (Wageningen Bioveterinary Research) |
| EPI_ISL_217939 | A/Waterfowl/Hubei/Chenhu1306/2014_H5N6 | A / H5N6 | Asia / China / Hubei Province | Jianjun Chen (Wuhan Institute of Virology / Chinese Academy of Sciences) |
| EPI_ISL_217940 | A/Waterfowl/Hubei/Chenhu1347/2014_H5N6 | A / H5N6 | Asia / China / Hubei Province | Jianjun Chen (Wuhan Institute of Virology / Chinese Academy of Sciences) |
| EPI_ISL_217941 | A/Anas crecca/Hubei/Chenhu1623-5/2014_H5N6 | A / H5N6 | Asia / China / Hubei Province | Jianjun Chen (Wuhan Institute of Virology / Chinese Academy of Sciences) |
| EPI_ISL_218200 | A/Bar-headed_goose/Qinghai/X402/2015_H5N6 | A / H5N6 | Asia / China / Qinghai Province | Jianjun Chen (Wuhan Institute of Virology / Chinese Academy of Sciences) |
| EPI_ISL_218201 | A/Bar-headed_goose/Qinghai/X417/2015_H5N6 | A / H5N6 | Asia / China / Qinghai Province | Jianjun Chen (Wuhan Institute of Virology / Chinese Academy of Sciences) |
| EPI_ISL_218202 | A/Bar-headed_goose/Qinghai/X418-1/2015_H5N6 | A / H5N6 | Asia / China / Qinghai Province | Jianjun Chen (Wuhan Institute of Virology / Chinese Academy of Sciences) |
| EPI_ISL_218203 | A/Bar-headed_goose/Qinghai/X418-2/2015_H5N6 | A / H5N6 | Asia / China / Qinghai Province | Jianjun Chen (Wuhan Institute of Virology / Chinese Academy of Sciences) |
| EPI_ISL_218204 | A/Bar-headed_goose/Qinghai/X424/2015_H5N6 | A / H5N6 | Asia / China / Qinghai Province | Jianjun Chen (Wuhan Institute of Virology / Chinese Academy of Sciences) |
| EPI_ISL_218205 | A/Bar-headed_goose/Qinghai/X428/2015_H5N6 | A / H5N6 | Asia / China / Qinghai Province | Jianjun Chen (Wuhan Institute of Virology / Chinese Academy of Sciences) |
| EPI_ISL_218206 | A/Bar-headed_goose/Qinghai/X430/2015_H5N6 | A / H5N6 | Asia / China / Qinghai Province | Jianjun Chen (Wuhan Institute of Virology / Chinese Academy of Sciences) |
| EPI_ISL_218207 | A/Bar-headed_goose/Qinghai/X437/2015_H5N6 | A / H5N6 | Asia / China / Qinghai Province | Jianjun Chen (Wuhan Institute of Virology / Chinese Academy of Sciences) |
| EPI_ISL_2193998 | A/barnacle goose/Netherlands/21027016-002/2021 | A / H5N1 | Europe / Netherlands / Provincie Groningen / Pieterburen | Rene Heutink (Wageningen Bioveterinary Research) |
| EPI_ISL_2194014 | A/white-tailed eagle/Netherlands/21027616-001/2021 | A / H5N1 | Europe / Netherlands / Provincie Groningen / Noordlaren | Rene Heutink (Wageningen Bioveterinary Research) |
| EPI_ISL_2194036 | A/barnacle goose/Netherlands/21028196-002/2021 | A / H5N1 | Europe / Netherlands / Provincie Groningen / Onnen | Rene Heutink (Wageningen Bioveterinary Research) |
| EPI_ISL_2194218 | A/red fox/Netherlands/21028774-002/2021 | A / H5N1 | Europe / Netherlands / Provincie Groningen / Bellingwolde | Rene Heutink (Wageningen Bioveterinary Research) |
| EPI_ISL_2194219 | A/red fox/Netherlands/21028774-004/2021 | A / H5N1 | Europe / Netherlands / Provincie Groningen / Bellingwolde | Rene Heutink (Wageningen Bioveterinary Research) |
| EPI_ISL_2194543 | A/chicken/Czech Republic/6151-2/2021 | A / H5N8 | Europe / Czech Republic / Kralovehradecky Kraj / Okres Hradec Kralove / Kosicky | Alexander Nagy (State Veterinary Institute Prague) |
| EPI_ISL_219760 | A/Environment/Hunan/28147/2014 | A / H5N1 | Asia / China / Hunan | Lei Yang (WHO Chinese National Influenza Center / Virology Institute, Chinese CDC) |
| EPI_ISL_219770 | A/Environment/Hunan/18459/2014 | A / H5N1 | Asia / China / Hunan | Lei Yang (WHO Chinese National Influenza Center / Virology Institute, Chinese CDC) |
| EPI_ISL_219771 | A/Environment/Hunan/25952/2014 | A / H5N1 | Asia / China / Hunan | Lei Yang (WHO Chinese National Influenza Center / Virology Institute, Chinese CDC) |
| EPI_ISL_219776 | A/Environment/Hunan/18478/2014 | A / H5N1 | Asia / China / Hunan | Lei Yang (WHO Chinese National Influenza Center / Virology Institute, Chinese CDC) |
| EPI_ISL_219777 | A/Environment/Chongqing/45355/2015 | A / H5N2 | Asia / China / Chongqing | Lei Yang (WHO Chinese National Influenza Center / Virology Institute, Chinese CDC) |
| EPI_ISL_219778 | A/Environment/Hubei/45047/2015 | A / H5N2 | Asia / China / Hubei | Lei Yang (WHO Chinese National Influenza Center / Virology Institute, Chinese CDC) |
| EPI_ISL_219779 | A/Environment/Hubei/45046/2015 | A / H5N2 | Asia / China / Hubei | Lei Yang (WHO Chinese National Influenza Center / Virology Institute, Chinese CDC) |
| EPI_ISL_219780 | A/Environment/Qinghai/42934/2015 | A / H5N2 | Asia / China / Qinghai | Lei Yang (WHO Chinese National Influenza Center / Virology Institute, Chinese CDC) |
| EPI_ISL_219781 | A/Environment/Qinghai/39266/2015 | A / H5N2 | Asia / China / Qinghai | Lei Yang (WHO Chinese National Influenza Center / Virology Institute, Chinese CDC) |
| EPI_ISL_219782 | A/Environment/Chongqing/47485/2015 | A / H5N6 | Asia / China / Chongqing | Lei Yang (WHO Chinese National Influenza Center / Virology Institute, Chinese CDC) |
| EPI_ISL_219783 | A/Environment/Guangxi/46690/2015 | A / H5N6 | Asia / China / Guangxi | Lei Yang (WHO Chinese National Influenza Center / Virology Institute, Chinese CDC) |
| EPI_ISL_219784 | A/Environment/Chongqing/45440/2014 | A / H5N6 | Asia / China / Chongqing | Lei Yang (WHO Chinese National Influenza Center / Virology Institute, Chinese CDC) |
| EPI_ISL_219785 | A/Environment/Chongqing/45439/2014 | A / H5N6 | Asia / China / Chongqing | Lei Yang (WHO Chinese National Influenza Center / Virology Institute, Chinese CDC) |
| EPI_ISL_219786 | A/Environment/Chongqing/45378/2015 | A / H5N6 | Asia / China / Chongqing | Lei Yang (WHO Chinese National Influenza Center / Virology Institute, Chinese CDC) |
| EPI_ISL_219787 | A/Environment/Chongqing/45373/2015 | A / H5N6 | Asia / China / Chongqing | Lei Yang (WHO Chinese National Influenza Center / Virology Institute, Chinese CDC) |
| EPI_ISL_219788 | A/Environment/Chongqing/45349/2015 | A / H5N6 | Asia / China / Chongqing | Lei Yang (WHO Chinese National Influenza Center / Virology Institute, Chinese CDC) |
| EPI_ISL_219789 | A/Environment/Chongqing/45379/2015 | A / H5N6 | Asia / China / Chongqing | Lei Yang (WHO Chinese National Influenza Center / Virology Institute, Chinese CDC) |
| EPI_ISL_219790 | A/Environment/Chongqing/45208/2015 | A / H5N6 | Asia / China / Chongqing | Lei Yang (WHO Chinese National Influenza Center / Virology Institute, Chinese CDC) |
| EPI_ISL_219791 | A/Environment/Chongqing/45221/2015 | A / H5N6 | Asia / China / Chongqing | Lei Yang (WHO Chinese National Influenza Center / Virology Institute, Chinese CDC) |
| EPI_ISL_219792 | A/Environment/Chongqing/45185/2015 | A / H5N6 | Asia / China / Chongqing | Lei Yang (WHO Chinese National Influenza Center / Virology Institute, Chinese CDC) |
| EPI_ISL_219793 | A/Environment/Chongqing/45155/2015 | A / H5N6 | Asia / China / Chongqing | Lei Yang (WHO Chinese National Influenza Center / Virology Institute, Chinese CDC) |
| EPI_ISL_219794 | A/Environment/Chongqing/45114/2015 | A / H5N6 | Asia / China / Chongqing | Lei Yang (WHO Chinese National Influenza Center / Virology Institute, Chinese CDC) |
| EPI_ISL_219795 | A/Environment/Chongqing/45134/2015 | A / H5N6 | Asia / China / Chongqing | Lei Yang (WHO Chinese National Influenza Center / Virology Institute, Chinese CDC) |

| | | | | |
|---|---|---|---|---|
| EPI_ISL_219796 | A/Environment/Hubei/45035/2015 | A / H5N6 | Asia / China / Hubei | Lei Yang (WHO Chinese National Influenza Center / Virology Institute, Chinese CDC) |
| EPI_ISL_219797 | A/Environment/Hubei/45034/2015 | A / H5N6 | Asia / China / Hubei | Lei Yang (WHO Chinese National Influenza Center / Virology Institute, Chinese CDC) |
| EPI_ISL_219798 | A/Environment/Guangxi/44389/2015 | A / H5N6 | Asia / China / Guangxi | Lei Yang (WHO Chinese National Influenza Center / Virology Institute, Chinese CDC) |
| EPI_ISL_219799 | A/Environment/Chongqing/99173/2014 | A / H5N6 | Asia / China / Chongqing | Lei Yang (WHO Chinese National Influenza Center / Virology Institute, Chinese CDC) |
| EPI_ISL_219800 | A/Environment/Guangxi/42586/2015 | A / H5N6 | Asia / China / Guangxi | Lei Yang (WHO Chinese National Influenza Center / Virology Institute, Chinese CDC) |
| EPI_ISL_219801 | A/Environment/Anhui/72105/2014 | A / H5N6 | Asia / China / Anhui | Lei Yang (WHO Chinese National Influenza Center / Virology Institute, Chinese CDC) |
| EPI_ISL_219802 | A/Environment/Chongqing/77193/2014 | A / H5N6 | Asia / China / Chongqing | Lei Yang (WHO Chinese National Influenza Center / Virology Institute, Chinese CDC) |
| EPI_ISL_219803 | A/Environment/Guangdong/77245/2014 | A / H5N6 | Asia / China / Guangdong | Lei Yang (WHO Chinese National Influenza Center / Virology Institute, Chinese CDC) |
| EPI_ISL_219804 | A/Environment/Hunan/18487/2014 | A / H5N6 | Asia / China / Hunan | Lei Yang (WHO Chinese National Influenza Center / Virology Institute, Chinese CDC) |
| EPI_ISL_219805 | A/Environment/Jiangxi/10649/2014 | A / H5N6 | Asia / China / Jiangxi | Lei Yang (WHO Chinese National Influenza Center / Virology Institute, Chinese CDC) |
| EPI_ISL_219806 | A/Environment/Jiangxi/10717/2014 | A / H5N6 | Asia / China / Jiangxi | Lei Yang (WHO Chinese National Influenza Center / Virology Institute, Chinese CDC) |
| EPI_ISL_219807 | A/Environment/Guangdong/40929/2015 | A / H5N6 | Asia / China / Guangdong | Lei Yang (WHO Chinese National Influenza Center / Virology Institute, Chinese CDC) |
| EPI_ISL_219808 | A/Environment/Guangdong/40113/2015 | A / H5N6 | Asia / China / Guangdong | Lei Yang (WHO Chinese National Influenza Center / Virology Institute, Chinese CDC) |
| EPI_ISL_219809 | A/Environment/Chongqing/72096/2014 | A / H5N6 | Asia / China / Chongqing | Lei Yang (WHO Chinese National Influenza Center / Virology Institute, Chinese CDC) |
| EPI_ISL_219810 | A/Environment/Yunnan/39607/2015 | A / H5N6 | Asia / China / Yunnan | Lei Yang (WHO Chinese National Influenza Center / Virology Institute, Chinese CDC) |
| EPI_ISL_219811 | A/Environment/Hubei/38005/2014 | A / H5N6 | Asia / China / Hubei | Lei Yang (WHO Chinese National Influenza Center / Virology Institute, Chinese CDC) |
| EPI_ISL_219812 | A/Environment/Hubei/37983/2014 | A / H5N6 | Asia / China / Hubei | Lei Yang (WHO Chinese National Influenza Center / Virology Institute, Chinese CDC) |
| EPI_ISL_219813 | A/Environment/Guangdong/33311/2015 | A / H5N6 | Asia / China / Guangdong | Lei Yang (WHO Chinese National Influenza Center / Virology Institute, Chinese CDC) |
| EPI_ISL_219814 | A/Environment/Guangdong/21131/2015 | A / H5N6 | Asia / China / Guangdong | Lei Yang (WHO Chinese National Influenza Center / Virology Institute, Chinese CDC) |
| EPI_ISL_219815 | A/Environment/Guangdong/21109/2015 | A / H5N6 | Asia / China / Guangdong | Lei Yang (WHO Chinese National Influenza Center / Virology Institute, Chinese CDC) |
| EPI_ISL_219816 | A/Environment/Hunan/07767/2015 | A / H5N6 | Asia / China / Hunan | Lei Yang (WHO Chinese National Influenza Center / Virology Institute, Chinese CDC) |
| EPI_ISL_219817 | A/Environment/Hunan/07681/2015 | A / H5N6 | Asia / China / Hunan | Lei Yang (WHO Chinese National Influenza Center / Virology Institute, Chinese CDC) |
| EPI_ISL_219818 | A/Environment/Xinjiang/07002/2014 | A / H5N6 | Asia / China / Xinjiang | Lei Yang (WHO Chinese National Influenza Center / Virology Institute, Chinese CDC) |
| EPI_ISL_219819 | A/Environment/Chongqing/99178/2014 | A / H5N6 | Asia / China / Chongqing | Lei Yang (WHO Chinese National Influenza Center / Virology Institute, Chinese CDC) |
| EPI_ISL_219820 | A/Environment/Xinjiang/00320/2014 | A / H5N6 | Asia / China / Xinjiang | Lei Yang (WHO Chinese National Influenza Center / Virology Institute, Chinese CDC) |
| EPI_ISL_219821 | A/Environment/Xinjiang/00318/2014 | A / H5N6 | Asia / China / Xinjiang | Lei Yang (WHO Chinese National Influenza Center / Virology Institute, Chinese CDC) |
| EPI_ISL_219822 | A/Environment/Xinjiang/00300/2014 | A / H5N6 | Asia / China / Xinjiang | Lei Yang (WHO Chinese National Influenza Center / Virology Institute, Chinese CDC) |
| EPI_ISL_219823 | A/Environment/Guangdong/00299/2014 | A / H5N6 | Asia / China / Guangdong | Lei Yang (WHO Chinese National Influenza Center / Virology Institute, Chinese CDC) |
| EPI_ISL_219824 | A/Environment/Xinjiang/00324/2014 | A / H5N6 | Asia / China / Xinjiang | Lei Yang (WHO Chinese National Influenza Center / Virology Institute, Chinese CDC) |
| EPI_ISL_219825 | A/Environment/Hunan/98885/2014 | A / H5N6 | Asia / China / Hunan | Lei Yang (WHO Chinese National Influenza Center / Virology Institute, Chinese CDC) |
| EPI_ISL_219826 | A/Environment/Chongqing/99181/2014 | A / H5N6 | Asia / China / Chongqing | Lei Yang (WHO Chinese National Influenza Center / Virology Institute, Chinese CDC) |
| EPI_ISL_219827 | A/Environment/Chongqing/99166/2014 | A / H5N6 | Asia / China / Chongqing | Lei Yang (WHO Chinese National Influenza Center / Virology Institute, Chinese CDC) |
| EPI_ISL_219828 | A/Guangdong/99710/2014 | A / H5N6 | Asia / China / Guangdong | Lei Yang (WHO Chinese National Influenza Center / Virology Institute, Chinese CDC) |
| EPI_ISL_219829 | A/Environment/Jiangsu/98335/2014 | A / H5N8 | Asia / China / Jiangsu | Lei Yang (WHO Chinese National Influenza Center / Virology Institute, Chinese CDC) |
| EPI_ISL_219844 | A/Environment/Hunan/72935/2014 | A / H5N2 | Asia / China / Hunan | Lei Yang (WHO Chinese National Influenza Center / Virology Institute, Chinese CDC) |
| EPI_ISL_219872 | A/goose/Taiwan/01038/2015 | A / H5N3 | Asia / Taiwan | |
| EPI_ISL_219873 | A/chicken/Taiwan/01174/2015 | A / H5N3 | Asia / Taiwan | |
| EPI_ISL_219874 | A/duck/Taiwan/A3400/2015 | A / H5N8 | Asia / Taiwan | |
| EPI_ISL_219875 | A/goose/Taiwan/01019/2015 | A / H5N8 | Asia / Taiwan | |
| EPI_ISL_219876 | A/goose/Taiwan/01026/2015 | A / H5N8 | Asia / Taiwan | |
| EPI_ISL_219877 | A/goose/Taiwan/01039/2015 | A / H5N8 | Asia / Taiwan | |
| EPI_ISL_219878 | A/duck/Taiwan/01006/2015 | A / H5N2 | Asia / Taiwan | |
| EPI_ISL_219879 | A/goose/Taiwan/01022/2015 | A / H5N2 | Asia / Taiwan | |
| EPI_ISL_219880 | A/goose/Taiwan/01023/2015 | A / H5N2 | Asia / Taiwan | |
| EPI_ISL_219881 | A/goose/Taiwan/01031/2015 | A / H5N2 | Asia / Taiwan | |
| EPI_ISL_219882 | A/goose/Taiwan/01040/2015 | A / H5N2 | Asia / Taiwan | |
| EPI_ISL_221389 | A/environment/Guangdong/GZ693/2015 | A / H5N6 | Asia / China | |
| EPI_ISL_221390 | A/environment/Guangdong/ZS558/2015 | A / H5N6 | Asia / China | |
| EPI_ISL_221391 | A/environment/Guangdong/GZ670/2015 | A / H5N6 | Asia / China | |
| EPI_ISL_221705 | A/duck/Guangdong/01.01 SZSGXJK001-Y /2016 | A / H5N6 | Asia / China / Guangdong Province | Yuhai Bi (Institute of Microbiology, Chinese Academy of Sciences) |
| EPI_ISL_221706 | A/duck/Guangdong/01.01 SZSGXJK001-G/2016 | A / H5N6 | Asia / China / Guangdong Province | Yuhai Bi (Institute of Microbiology, Chinese Academy of Sciences) |
| EPI_ISL_221707 | A/duck/Guangdong/01.01 SZSGXJK002-Y/2016 | A / H5N6 | Asia / China / Guangdong Province | Yuhai Bi (Institute of Microbiology, Chinese Academy of Sciences) |
| EPI_ISL_221708 | A/duck/Guangdong/01.01 SZSGXJK002-G/2016 | A / H5N6 | Asia / China / Guangdong Province | Yuhai Bi (Institute of Microbiology, Chinese Academy of Sciences) |
| EPI_ISL_221709 | A/duck/Guangdong/01.01 SZSGXJK003-G/2016 | A / H5N6 | Asia / China / Guangdong Province | Yuhai Bi (Institute of Microbiology, Chinese Academy of Sciences) |
| EPI_ISL_221710 | A/duck/Guangdong/01.01 SZSGXJK004-Y/2016 | A / H5N6 | Asia / China / Guangdong Province | Yuhai Bi (Institute of Microbiology, Chinese Academy of Sciences) |
| EPI_ISL_221711 | A/duck/Guangdong/01.01 SZSGXJK003-W/2016 | A / H5N6 | Asia / China / Guangdong Province | Yuhai Bi (Institute of Microbiology, Chinese Academy of Sciences) |
| EPI_ISL_221712 | A/duck/Guangdong/01.01 SZSGXJK005-Y/2016 | A / H5N6 | Asia / China / Guangdong Province | Yuhai Bi (Institute of Microbiology, Chinese Academy of Sciences) |
| EPI_ISL_221713 | A/duck/Guangdong/01.01 SZSGXJK005-G /2016 | A / H5N6 | Asia / China / Guangdong Province | Yuhai Bi (Institute of Microbiology, Chinese Academy of Sciences) |
| EPI_ISL_221714 | A/duck/Guangdong/01.01 SZSGXJK006-Y/2016 | A / H5N6 | Asia / China / Guangdong Province | Yuhai Bi (Institute of Microbiology, Chinese Academy of Sciences) |
| EPI_ISL_221715 | A/duck/Guangdong/01.01 SZSGXJK006-G/2016 | A / H5N6 | Asia / China / Guangdong Province | Yuhai Bi (Institute of Microbiology, Chinese Academy of Sciences) |
| EPI_ISL_221716 | A/duck/Guangdong/01.01 SZSGXJK007-G/2016 | A / H5N6 | Asia / China / Guangdong Province | Yuhai Bi (Institute of Microbiology, Chinese Academy of Sciences) |
| EPI_ISL_221719 | A/environment/Guangdong/01.01 SZSGXJK006-E/2016 | A / H5N6 | Asia / China / Guangdong Province | Yuhai Bi (Institute of Microbiology, Chinese Academy of Sciences) |
| EPI_ISL_221720 | A/feline/Guangdong/1/2015 | A / H5N6 | Asia / China | |
| EPI_ISL_221721 | A/feline/Guangdong/2/2015 | A / H5N6 | Asia / China | |
| EPI_ISL_221723 | A/tiger/Yunnan/tig1404/2014 | A / H5N1 | Asia / China | |
| EPI_ISL_221885 | A/chicken/Guangdong/10.16 SZLGKQHY-072/2015 | A / H5N6 | Asia / China / Guangdong Province | Yuhai Bi (Institute of Microbiology, Chinese Academy of Sciences) |
| EPI_ISL_222132 | A/duck/Eastern China/JY/2014 | A / H5N8 | Asia / China | |
| EPI_ISL_222134 | A/goose/Eastern China/CZ/2013 | A / H5N8 | Asia / China | |
| EPI_ISL_2227275 | A/turkey/Netherlands/21028936-001005/2021 | A / H5N8 | Europe / Netherlands / Provincie Limburg / Weert | Rene Heutink (Wageningen Bioveterinary Research) |
| EPI_ISL_2227276 | A/barnacle goose/Netherlands/21027357-002/2021 | A / H5N1 | Europe / Netherlands / Provincie Groningen / Termunten | Rene Heutink (Wageningen Bioveterinary Research) |
| EPI_ISL_2227277 | A/barnacle goose/Netherlands/21028534-002/2021 | A / H5N1 | Europe / Netherlands / Provincie Groningen / Waterhuizen | Rene Heutink (Wageningen Bioveterinary Research) |
| EPI_ISL_2227278 | A/goose/Netherlands/21028502-002/21028502/2021 | A / H5N1 | Europe / Netherlands / Provincie Groningen / Houwerzijl | Rene Heutink (Wageningen Bioveterinary Research) |

| | | | | |
|---|---|---|---|---|
| EPI_ISL_2227279 | A/western marsh harrier/Netherlands/21028606-002/2021 | A / H5N1 | Europe / Netherlands / Provincie Friesland / Ooststellingwerf | Rene Heutink (Wageningen Bioveterinary Research) |
| EPI_ISL_223364 | A/chicken/Wuxi/SC7393/2015 | A / H5N6 | Asia / China / Jiangsu Provice | Maijuan Ma (Beijing Institute of Microbiology and Epidemiology / State Key Laboratory of Pathogen and Biosecurity) |
| EPI_ISL_2234807 | A/knot_wader/Ireland/472_21VIR2956-1/2021 | A / H5N3 | Europe / Ireland | Bianca Zecchin (Istituto Zooprofilattico Sperimentale Delle Venezie) |
| EPI_ISL_2234808 | A/knot_wader/Ireland/473_21VIR2956-2/2021 | A / H5N3 | Europe / Ireland | Bianca Zecchin (Istituto Zooprofilattico Sperimentale Delle Venezie) |
| EPI_ISL_2234809 | A/yellow-legged_gull/Switzerland/15-0039_21VIR3035/2021 | A / H5N4 | Europe / Switzerland | Bianca Zecchin (Istituto Zooprofilattico Sperimentale Delle Venezie) |
| EPI_ISL_2234810 | A/seagull/Italy/21VIR2479/2021 | A / H5N8 | Europe / Italy | Bianca Zecchin (Istituto Zooprofilattico Sperimentale Delle Venezie) |
| EPI_ISL_2234811 | A/anser_anser/Spain/102-8_21VIR1230-4/2021 | A / H5N8 | Europe / Spain | Bianca Zecchin (Istituto Zooprofilattico Sperimentale Delle Venezie) |
| EPI_ISL_2234812 | A/anser_anser/Spain/297-1_21VIR1230-5/2021 | A / H5N8 | Europe / Spain | Bianca Zecchin (Istituto Zooprofilattico Sperimentale Delle Venezie) |
| EPI_ISL_2234813 | A/ciconia_ciconia/Spain/102-1_21VIR1230-2/2021 | A / H5N8 | Europe / Spain | Bianca Zecchin (Istituto Zooprofilattico Sperimentale Delle Venezie) |
| EPI_ISL_2234814 | A/peregrine_falcon/Spain/3365-1_21VIR1230-1/2020 | A / H5N8 | Europe / Spain | Bianca Zecchin (Istituto Zooprofilattico Sperimentale Delle Venezie) |
| EPI_ISL_2234815 | A/duck/Romania/10202_21VIR2593-31/2021 | A / H5N8 | Europe / Romania | Bianca Zecchin (Istituto Zooprofilattico Sperimentale Delle Venezie) |
| EPI_ISL_2234816 | A/duck/Romania/10205-t4_21VIR2593-3/2021 | A / H5N8 | Europe / Romania | Bianca Zecchin (Istituto Zooprofilattico Sperimentale Delle Venezie) |
| EPI_ISL_2234817 | A/goose/Romania/10205-t3_21VIR2593-7/2021 | A / H5N8 | Europe / Romania | Bianca Zecchin (Istituto Zooprofilattico Sperimentale Delle Venezie) |
| EPI_ISL_2234818 | A/goose/Romania/10205-t5_21VIR2593-8/2021 | A / H5N8 | Europe / Romania | Bianca Zecchin (Istituto Zooprofilattico Sperimentale Delle Venezie) |
| EPI_ISL_2234819 | A/goose/Romania/10330_21VIR2593-35/2021 | A / H5N8 | Europe / Romania | Bianca Zecchin (Istituto Zooprofilattico Sperimentale Delle Venezie) |
| EPI_ISL_2234820 | A/hawk/Romania/10363_21VIR2593-11/2021 | A / H5N8 | Europe / Romania | Bianca Zecchin (Istituto Zooprofilattico Sperimentale Delle Venezie) |
| EPI_ISL_2234821 | A/whooper_swan/Romania/10122_21VIR2593-23/2021 | A / H5N5 | Europe / Romania | Bianca Zecchin (Istituto Zooprofilattico Sperimentale Delle Venezie) |
| EPI_ISL_2234822 | A/whooper_swan/Romania/10171_21VIR2593-15/2021 | A / H5N5 | Europe / Romania | Bianca Zecchin (Istituto Zooprofilattico Sperimentale Delle Venezie) |
| EPI_ISL_2234823 | A/whooper_swan/Romania/10213_21VIR2593-27/2021 | A / H5N5 | Europe / Romania | Bianca Zecchin (Istituto Zooprofilattico Sperimentale Delle Venezie) |
| EPI_ISL_2234824 | A/whooper_swan/Romania/10362_21VIR2593-20/2021 | A / H5N5 | Europe / Romania | Bianca Zecchin (Istituto Zooprofilattico Sperimentale Delle Venezie) |
| EPI_ISL_2234831 | A/peregrine_falcon/Northern_Ireland/21VIR1662-3/2021 | A / H5N3 | Europe / United Kingdom | Bianca Zecchin (Istituto Zooprofilattico Sperimentale Delle Venezie) |
| EPI_ISL_224580 | A/great crested grebe/Uvs-Nuur Lake/341/2016 | A / H5N8 | Europe / Russian Federation / Tyva Republic | Ivan Sobolev (Research Institute of Experimental and Clinical Medicine) |
| EPI_ISL_224704 | A/Bar-headed Goose/Qinghai/BTY1-B/2016 | A / H5N8 | Asia / China / Qinghai Province | Jianjun Chen (Wuhan Institute of Virology / Chinese Academy of Sciences) |
| EPI_ISL_224705 | A/Bar-headed Goose/Qinghai/BTY1-LV/2016 | A / H5N8 | Asia / China / Qinghai Province | Jianjun Chen (Wuhan Institute of Virology / Chinese Academy of Sciences) |
| EPI_ISL_224707 | A/Bar-headed Goose/Qinghai/BTY2-B/2016 | A / H5N8 | Asia / China / Qinghai Province | Jianjun Chen (Wuhan Institute of Virology / Chinese Academy of Sciences) |
| EPI_ISL_224708 | A/Bar-headed Goose/Qinghai/BTY2-LU/2016 | A / H5N8 | Asia / China / Qinghai Province | Jianjun Chen (Wuhan Institute of Virology / Chinese Academy of Sciences) |
| EPI_ISL_224709 | A/Bar-headed Goose/Qinghai/BTY3-B/2016 | A / H5N8 | Asia / China / Qinghai Province | Jianjun Chen (Wuhan Institute of Virology / Chinese Academy of Sciences) |
| EPI_ISL_224710 | A/Bar-headed Goose/Qinghai/BTY3-LU/2016 | A / H5N8 | Asia / China / Qinghai Province | Jianjun Chen (Wuhan Institute of Virology / Chinese Academy of Sciences) |
| EPI_ISL_224711 | A/Bar-headed Goose/Qinghai/BTY4-B/2016 | A / H5N8 | Asia / China / Qinghai Province | Jianjun Chen (Wuhan Institute of Virology / Chinese Academy of Sciences) |
| EPI_ISL_224712 | A/Bar-headed Goose/Qinghai/BTY4-LU/2016 | A / H5N8 | Asia / China / Qinghai Province | Jianjun Chen (Wuhan Institute of Virology / Chinese Academy of Sciences) |
| EPI_ISL_224713 | A/Bar-headed Goose/Qinghai/BTY5-B/2016 | A / H5N8 | Asia / China / Qinghai Province | Jianjun Chen (Wuhan Institute of Virology / Chinese Academy of Sciences) |
| EPI_ISL_224714 | A/Bar-headed Goose/Qinghai/BTY6-B/2016 | A / H5N8 | Asia / China / Qinghai Province | Jianjun Chen (Wuhan Institute of Virology / Chinese Academy of Sciences) |
| EPI_ISL_224715 | A/Bar-headed Goose/Qinghai/BTY6-LU/2016 | A / H5N8 | Asia / China / Qinghai Province | Jianjun Chen (Wuhan Institute of Virology / Chinese Academy of Sciences) |
| EPI_ISL_224716 | A/Bar-headed Goose/Qinghai/BTY7-B/2016 | A / H5N8 | Asia / China / Qinghai Province | Jianjun Chen (Wuhan Institute of Virology / Chinese Academy of Sciences) |
| EPI_ISL_224717 | A/Bar-headed Goose/Qinghai/BTY7-LU1/2016 | A / H5N8 | Asia / China / Qinghai Province | Jianjun Chen (Wuhan Institute of Virology / Chinese Academy of Sciences) |
| EPI_ISL_224718 | A/Bar-headed Goose/Qinghai/BTY7-LU2/2016 | A / H5N8 | Asia / China / Qinghai Province | Jianjun Chen (Wuhan Institute of Virology / Chinese Academy of Sciences) |
| EPI_ISL_224719 | A/Bar-headed Goose/Qinghai/BTY8-B/2016 | A / H5N8 | Asia / China / Qinghai Province | Jianjun Chen (Wuhan Institute of Virology / Chinese Academy of Sciences) |
| EPI_ISL_224720 | A/Bar-headed Goose/Qinghai/BTY8-LU/2016 | A / H5N8 | Asia / China / Qinghai Province | Jianjun Chen (Wuhan Institute of Virology / Chinese Academy of Sciences) |
| EPI_ISL_224721 | A/Bar-headed Goose/Qinghai/BTY9-B/2016 | A / H5N8 | Asia / China / Qinghai Province | Jianjun Chen (Wuhan Institute of Virology / Chinese Academy of Sciences) |
| EPI_ISL_224722 | A/Bar-headed Goose/Qinghai/BTY9-LU/2016 | A / H5N8 | Asia / China / Qinghai Province | Jianjun Chen (Wuhan Institute of Virology / Chinese Academy of Sciences) |
| EPI_ISL_224723 | A/Bar-headed Goose/Qinghai/BTY10-B/2016 | A / H5N8 | Asia / China / Qinghai Province | Jianjun Chen (Wuhan Institute of Virology / Chinese Academy of Sciences) |
| EPI_ISL_224724 | A/Bar-headed Goose/Qinghai/BTY10-LU/2016 | A / H5N8 | Asia / China / Qinghai Province | Jianjun Chen (Wuhan Institute of Virology / Chinese Academy of Sciences) |
| EPI_ISL_224725 | A/Bar-headed Goose/Qinghai/BTY11-B/2016 | A / H5N8 | Asia / China / Qinghai Province | Jianjun Chen (Wuhan Institute of Virology / Chinese Academy of Sciences) |
| EPI_ISL_224726 | A/Bar-headed Goose/Qinghai/BTY11-LU/2016 | A / H5N8 | Asia / China / Qinghai Province | Jianjun Chen (Wuhan Institute of Virology / Chinese Academy of Sciences) |
| EPI_ISL_224727 | A/Bar-headed Goose/Qinghai/BTY12-B/2016 | A / H5N8 | Asia / China / Qinghai Province | Jianjun Chen (Wuhan Institute of Virology / Chinese Academy of Sciences) |
| EPI_ISL_224728 | A/Bar-headed Goose/Qinghai/BTY12-LU/2016 | A / H5N8 | Asia / China / Qinghai Province | Jianjun Chen (Wuhan Institute of Virology / Chinese Academy of Sciences) |
| EPI_ISL_224729 | A/Bar-headed Goose/Qinghai/BTY13-B/2016 | A / H5N8 | Asia / China / Qinghai Province | Jianjun Chen (Wuhan Institute of Virology / Chinese Academy of Sciences) |
| EPI_ISL_224730 | A/Bar-headed Goose/Qinghai/BTY13-LU/2016 | A / H5N8 | Asia / China / Qinghai Province | Jianjun Chen (Wuhan Institute of Virology / Chinese Academy of Sciences) |
| EPI_ISL_224731 | A/Bar-headed Goose/Qinghai/BTY14-B/2016 | A / H5N8 | Asia / China / Qinghai Province | Jianjun Chen (Wuhan Institute of Virology / Chinese Academy of Sciences) |
| EPI_ISL_224732 | A/Bar-headed Goose/Qinghai/BTY14-LU/2016 | A / H5N8 | Asia / China / Qinghai Province | Jianjun Chen (Wuhan Institute of Virology / Chinese Academy of Sciences) |
| EPI_ISL_224733 | A/Bar-headed Goose/Qinghai/BTY15-B/2016 | A / H5N8 | Asia / China / Qinghai Province | Jianjun Chen (Wuhan Institute of Virology / Chinese Academy of Sciences) |
| EPI_ISL_224734 | A/Bar-headed Goose/Qinghai/BTY15-LU/2016 | A / H5N8 | Asia / China / Qinghai Province | Jianjun Chen (Wuhan Institute of Virology / Chinese Academy of Sciences) |
| EPI_ISL_224735 | A/Bar-headed Goose/Qinghai/BTY16-B/2016 | A / H5N8 | Asia / China / Qinghai Province | Jianjun Chen (Wuhan Institute of Virology / Chinese Academy of Sciences) |
| EPI_ISL_224736 | A/Bar-headed Goose/Qinghai/BTY16-LU/2016 | A / H5N8 | Asia / China / Qinghai Province | Jianjun Chen (Wuhan Institute of Virology / Chinese Academy of Sciences) |
| EPI_ISL_224737 | A/Bar-headed Goose/Qinghai/BTY17-B/2016 | A / H5N8 | Asia / China / Qinghai Province | Jianjun Chen (Wuhan Institute of Virology / Chinese Academy of Sciences) |
| EPI_ISL_224738 | A/Bar-headed Goose/Qinghai/BTY17-LU/2016 | A / H5N8 | Asia / China / Qinghai Province | Jianjun Chen (Wuhan Institute of Virology / Chinese Academy of Sciences) |
| EPI_ISL_224739 | A/Bar-headed Goose/Qinghai/BTY18-B/2016 | A / H5N8 | Asia / China / Qinghai Province | Jianjun Chen (Wuhan Institute of Virology / Chinese Academy of Sciences) |
| EPI_ISL_224740 | A/Bar-headed Goose/Qinghai/BTY18-LU/2016 | A / H5N8 | Asia / China / Qinghai Province | Jianjun Chen (Wuhan Institute of Virology / Chinese Academy of Sciences) |
| EPI_ISL_224742 | A/Brown-headed Gull/Qinghai/ZTO1-B/2016 | A / H5N8 | Asia / China / Qinghai Province | Jianjun Chen (Wuhan Institute of Virology / Chinese Academy of Sciences) |
| EPI_ISL_224743 | A/Brown-headed Gull/Qinghai/ZTO1-LU/2016 | A / H5N8 | Asia / China / Qinghai Province | Jianjun Chen (Wuhan Institute of Virology / Chinese Academy of Sciences) |
| EPI_ISL_224744 | A/Brown-headed Gull/Qinghai/ZTO3-B/2016 | A / H5N8 | Asia / China / Qinghai Province | Jianjun Chen (Wuhan Institute of Virology / Chinese Academy of Sciences) |
| EPI_ISL_224745 | A/Brown-headed Gull/Qinghai/ZTO3-LU/2016 | A / H5N8 | Asia / China / Qinghai Province | Jianjun Chen (Wuhan Institute of Virology / Chinese Academy of Sciences) |
| EPI_ISL_224746 | A/Brown-headed Gull/Qinghai/ZTO4-B/2016 | A / H5N8 | Asia / China / Qinghai Province | Jianjun Chen (Wuhan Institute of Virology / Chinese Academy of Sciences) |
| EPI_ISL_224747 | A/Brown-headed Gull/Qinghai/ZTO5-B/2016 | A / H5N8 | Asia / China / Qinghai Province | Jianjun Chen (Wuhan Institute of Virology / Chinese Academy of Sciences) |
| EPI_ISL_224748 | A/Brown-headed Gull/Qinghai/ZTO5-K/2016 | A / H5N8 | Asia / China / Qinghai Province | Jianjun Chen (Wuhan Institute of Virology / Chinese Academy of Sciences) |
| EPI_ISL_224749 | A/Brown-headed Gull/Qinghai/ZTO6-B/2016 | A / H5N8 | Asia / China / Qinghai Province | Jianjun Chen (Wuhan Institute of Virology / Chinese Academy of Sciences) |
| EPI_ISL_224750 | A/Brown-headed Gull/Qinghai/ZTO6-SP/2016 | A / H5N8 | Asia / China / Qinghai Province | Jianjun Chen (Wuhan Institute of Virology / Chinese Academy of Sciences) |
| EPI_ISL_224751 | A/Brown-headed Gull/Qinghai/ZTO6-MU/2016 | A / H5N8 | Asia / China / Qinghai Province | Jianjun Chen (Wuhan Institute of Virology / Chinese Academy of Sciences) |
| EPI_ISL_224752 | A/Great Black-headed Gull/Qinghai/YO1-B/2016 | A / H5N8 | Asia / China / Qinghai Province | Jianjun Chen (Wuhan Institute of Virology / Chinese Academy of Sciences) |
| EPI_ISL_224879 | A/duck/Zhejiang/6DK19/2013 | A / H5N2 | Asia / China | |
| EPI_ISL_224984 | A/duck/Guangzhou/018/2014 | A / H5N6 | Asia / China | |
| EPI_ISL_224985 | A/duck/Guangzhou/021/2014 | A / H5N6 | Asia / China | |

| | | | | |
|---|---|---|---|---|
| EPI_ISL_2254497 | A/western jackdaw/SVA210317SZ0321/KN001057/E-2021 | A / H5N8 | Europe / Sweden / Ostergotlands Lan / Mjolby Kommun | Siamak Zohari (National Veterinary Institute) |
| EPI_ISL_225570 | A/turkey/South Dakota/15-012511-2/2015 | A / H5N2 | North America / United States | |
| EPI_ISL_225571 | A/turkey/Minnesota/15-012582-1/2015 | A / H5N2 | North America / United States | |
| EPI_ISL_225572 | A/chicken/Iowa/15-013388-1/2015 | A / H5N2 | North America / United States | |
| EPI_ISL_2276068 | A/chicken/Senegal/21VIR1084-3/2021 | A / H5N1 | Africa / Senegal / Region de Thies | Bianca Zecchin (Istituto Zooprofilattico Sperimentale Delle Venezie) |
| EPI_ISL_2276069 | A/chicken/Senegal/21VIR1084-4/2021 | A / H5N1 | Africa / Senegal / Region de Thies | Bianca Zecchin (Istituto Zooprofilattico Sperimentale Delle Venezie) |
| EPI_ISL_2276070 | A/chicken/Senegal/21VIR1084-5/2021 | A / H5N1 | Africa / Senegal / Region de Thies | Bianca Zecchin (Istituto Zooprofilattico Sperimentale Delle Venezie) |
| EPI_ISL_2276071 | A/great-white_pelican/Senegal/21-67_21VIR1084-8/2021 | A / H5N1 | Africa / Senegal / Djoudj bird sanctuary | Bianca Zecchin (Istituto Zooprofilattico Sperimentale Delle Venezie) |
| EPI_ISL_229213 | A/duck/Vietnam/LBM808/2015 | A / H5N6 | Asia / Vietnam | |
| EPI_ISL_2301187 | A/common buzzard/Sweden/SVA210323SZ0426/KN001120/O- | A / H5N8 | Europe / Sweden / Vastra Gotalands Lan / Lidkopings Kommun | Siamak Zohari (National Veterinary Institute) |
| EPI_ISL_230158 | A/muscovy duck/LBM803/2015 | A / H5N6 | Asia / Vietnam | |
| EPI_ISL_230754 | A/muscovy duck/LBM806/2015 | A / H5N6 | Asia / Vietnam | |
| EPI_ISL_230755 | A/muscovy duck/Vietnam/LBM809/2015 | A / H5N6 | Asia / Vietnam | |
| EPI_ISL_230756 | A/muscovy duck/Vietnam/LBM810/2015 | A / H5N6 | Asia / Vietnam | |
| EPI_ISL_230757 | A/muscovy duck/Vietnam/LBM817/2015 | A / H5N6 | Asia / Vietnam | |
| EPI_ISL_230758 | A/muscovy duck/Vietnam/LBM818/2015 | A / H5N6 | Asia / Vietnam | |
| EPI_ISL_230759 | A/muscovy duck/Vietnam/LBM823/2015 | A / H5N6 | Asia / Vietnam | |
| EPI_ISL_230760 | A/duck/Vietnam/LBM812/2015 | A / H5N6 | Asia / Vietnam | |
| EPI_ISL_230820 | A/great crested grebe/Tyva/34/2016 | A / H5N8 | Europe / Russian Federation / Tyva Republic | Artem Fadeev (WHO National Influenza Centre Russian Federation) |
| EPI_ISL_231525 | A/duck/Vietnam/LBM813/2015 | A / H5N6 | Asia / Vietnam | |
| EPI_ISL_231526 | A/duck/Vietnam/LBM816/2015 | A / H5N6 | Asia / Vietnam | |
| EPI_ISL_231527 | A/duck/Vietnam/LBM824/2015 | A / H5N6 | Asia / Vietnam | |
| EPI_ISL_231684 | A/wild duck/Tyva/35/2016 | A / H5N8 | Europe / Russian Federation / Tyva Republic | Artem Fadeev (WHO National Influenza Centre Russian Federation) |
| EPI_ISL_231685 | A/black-headed gull/Tyva/41/2016 | A / H5N8 | Europe / Russian Federation / Tyva Republic | Artem Fadeev (WHO National Influenza Centre Russian Federation) |
| EPI_ISL_231747 | A/duck/Hubei/SZY250/2016 | A / H5N2 | Asia / China | |
| EPI_ISL_234057 | A/grey heron /Uvs-Nuur Lake/20/2016 | A / H5N8 | Europe / Russian Federation / Tyva Republic | Ivan Sobolev (Research Institute of Experimental and Clinical Medicine) |
| EPI_ISL_234058 | A/common tern /Uvs-Nuur Lake/26/2016 | A / H5N8 | Europe / Russian Federation / Tyva Republic | Ivan Sobolev (Research Institute of Experimental and Clinical Medicine) |
| EPI_ISL_234282 | A/chicken/Hubei/XG18/2015 | A / H5N6 | Asia / China | |
| EPI_ISL_234283 | A/duck/Hubei/XG18/2015 | A / H5N6 | Asia / China | |
| EPI_ISL_234284 | A/duck/Hubei/WH18/2015 | A / H5N6 | Asia / China | |
| EPI_ISL_234329 | A/breeder duck/Korea/H2086/2015 | A / H5N8 | Asia / Korea, Republic of / Jeollanam-do | Eun-Kyoung Lee (Animal and Plant Quarantine Agency (APQA) / Avian Influenza Research and Diagsnotic Division) |
| EPI_ISL_234330 | A/mallard/Korea/H2102/2015 | A / H5N8 | Asia / Korea, Republic of / Chungcheongnam-do | Eun-Kyoung Lee (Animal and Plant Quarantine Agency (APQA) / Avian Influenza Research and Diagsnotic Division) |
| EPI_ISL_234331 | A/broiler duck/Korea/H2116/2015 | A / H5N8 | Asia / Korea, Republic of / Jeollabuk-do | Eun-Kyoung Lee (Animal and Plant Quarantine Agency (APQA) / Avian Influenza Research and Diagsnotic Division) |
| EPI_ISL_234332 | A/broiler duck/Korea/H2194/2015 | A / H5N8 | Asia / Korea, Republic of / Chungcheongnam-do | Eun-Kyoung Lee (Animal and Plant Quarantine Agency (APQA) / Avian Influenza Research and Diagsnotic Division) |
| EPI_ISL_234333 | A/broiler duck/Korea/H2266/2015 | A / H5N8 | Asia / Korea, Republic of / Chungcheongnam-do | Eun-Kyoung Lee (Animal and Plant Quarantine Agency (APQA) / Avian Influenza Research and Diagsnotic Division) |
| EPI_ISL_234334 | A/broiler duck/Korea/H2278/2015 | A / H5N8 | Asia / Korea, Republic of / Jeollanam-do | Eun-Kyoung Lee (Animal and Plant Quarantine Agency (APQA) / Avian Influenza Research and Diagsnotic Division) |
| EPI_ISL_234335 | A/wild bird/Korea/H2290/2015 | A / H5N8 | Asia / Korea, Republic of / Chungcheongnam-do | Eun-Kyoung Lee (Animal and Plant Quarantine Agency (APQA) / Avian Influenza Research and Diagsnotic Division) |
| EPI_ISL_234336 | A/wild bird/Korea/H2291/2015 | A / H5N8 | Asia / Korea, Republic of / Jeju-teukbyeoljachido | Eun-Kyoung Lee (Animal and Plant Quarantine Agency (APQA) / Avian Influenza Research and Diagsnotic Division) |
| EPI_ISL_234337 | A/mallard/Korea/H2304/2015 | A / H5N8 | Asia / Korea, Republic of / Chungcheongnam-do | Eun-Kyoung Lee (Animal and Plant Quarantine Agency (APQA) / Avian Influenza Research and Diagsnotic Division) |
| EPI_ISL_234338 | A/mallard/Korea/H2321/2015 | A / H5N8 | Asia / Korea, Republic of / Seoul | Eun-Kyoung Lee (Animal and Plant Quarantine Agency (APQA) / Avian Influenza Research and Diagsnotic Division) |
| EPI_ISL_234339 | A/broiler duck/Korea/H2385/2015 | A / H5N8 | Asia / Korea, Republic of / Jeollanam-do | Eun-Kyoung Lee (Animal and Plant Quarantine Agency (APQA) / Avian Influenza Research and Diagsnotic Division) |
| EPI_ISL_234340 | A/broiler duck/Korea/H2393/2015 | A / H5N8 | Asia / Korea, Republic of / Chungcheongbuk-do | Eun-Kyoung Lee (Animal and Plant Quarantine Agency (APQA) / Avian Influenza Research and Diagsnotic Division) |
| EPI_ISL_234341 | A/broiler duck/Korea/H2400/2015 | A / H5N8 | Asia / Korea, Republic of / Chungcheongbuk-do | Eun-Kyoung Lee (Animal and Plant Quarantine Agency (APQA) / Avian Influenza Research and Diagsnotic Division) |
| EPI_ISL_234342 | A/broiler duck/Korea/H2405/2015 | A / H5N8 | Asia / Korea, Republic of / Chungcheongbuk-do | Eun-Kyoung Lee (Animal and Plant Quarantine Agency (APQA) / Avian Influenza Research and Diagsnotic Division) |
| EPI_ISL_234343 | A/breeder chicken/Korea/H2496/2015 | A / H5N8 | Asia / Korea, Republic of / Chungcheongbuk-do | Eun-Kyoung Lee (Animal and Plant Quarantine Agency (APQA) / Avian Influenza Research and Diagsnotic Division) |
| EPI_ISL_234344 | A/broiler duck/Korea/H2517/2015 | A / H5N8 | Asia / Korea, Republic of / Chungcheongbuk-do | Eun-Kyoung Lee (Animal and Plant Quarantine Agency (APQA) / Avian Influenza Research and Diagsnotic Division) |
| EPI_ISL_234345 | A/broiler duck/Korea/H2531/2015 | A / H5N8 | Asia / Korea, Republic of / Jeollabuk-do | Eun-Kyoung Lee (Animal and Plant Quarantine Agency (APQA) / Avian Influenza Research and Diagsnotic Division) |
| EPI_ISL_234346 | A/chicken/Korea/H2553/2015 | A / H5N8 | Asia / Korea, Republic of / Jeollabuk-do | Eun-Kyoung Lee (Animal and Plant Quarantine Agency (APQA) / Avian Influenza Research and Diagsnotic Division) |
| EPI_ISL_234347 | A/chicken/Korea/H2565/2015 | A / H5N8 | Asia / Korea, Republic of / Jeollabuk-do | Eun-Kyoung Lee (Animal and Plant Quarantine Agency (APQA) / Avian Influenza Research and Diagsnotic Division) |
| EPI_ISL_234348 | A/chicken/Korea/H2590/2015 | A / H5N8 | Asia / Korea, Republic of / Gyeonggi-do | Eun-Kyoung Lee (Animal and Plant Quarantine Agency (APQA) / Avian Influenza Research and Diagsnotic Division) |
| EPI_ISL_234349 | A/korean native chicken/Korea/H2598/2015 | A / H5N8 | Asia / Korea, Republic of / Gyeonggi-do | Eun-Kyoung Lee (Animal and Plant Quarantine Agency (APQA) / Avian Influenza Research and Diagsnotic Division) |
| EPI_ISL_234350 | A/broiler duck/Korea/H2606/2015 | A / H5N8 | Asia / Korea, Republic of / Jeollabuk-do | Eun-Kyoung Lee (Animal and Plant Quarantine Agency (APQA) / Avian Influenza Research and Diagsnotic Division) |
| EPI_ISL_234351 | A/broiler duck/Korea/H2618/2015 | A / H5N8 | Asia / Korea, Republic of / Jeollanam-do | Eun-Kyoung Lee (Animal and Plant Quarantine Agency (APQA) / Avian Influenza Research and Diagsnotic Division) |
| EPI_ISL_234352 | A/duck/Korea/H2627/2015 | A / H5N8 | Asia / Korea, Republic of / Gyeonggi-do | Eun-Kyoung Lee (Animal and Plant Quarantine Agency (APQA) / Avian Influenza Research and Diagsnotic Division) |
| EPI_ISL_234353 | A/duck/Korea/H2628/2015 | A / H5N8 | Asia / Korea, Republic of | Eun-Kyoung Lee (Animal and Plant Quarantine Agency (APQA) / Avian Influenza Research and Diagsnotic Division) |
| EPI_ISL_234354 | A/broildr duck/Korea/H2637/2015 | A / H5N8 | Asia / Korea, Republic of / Jeollanam-do | Eun-Kyoung Lee (Animal and Plant Quarantine Agency (APQA) / Avian Influenza Research and Diagsnotic Division) |
| EPI_ISL_234355 | A/broiler duck/Korea/H2641/2015 | A / H5N8 | Asia / Korea, Republic of / Jeollanam-do | Eun-Kyoung Lee (Animal and Plant Quarantine Agency (APQA) / Avian Influenza Research and Diagsnotic Division) |
| EPI_ISL_234356 | A/broiler duck/Korea/H2649/2015 | A / H5N8 | Asia / Korea, Republic of / Jeollanam-do | Eun-Kyoung Lee (Animal and Plant Quarantine Agency (APQA) / Avian Influenza Research and Diagsnotic Division) |
| EPI_ISL_234357 | A/breeder duck/Korea/H2675/2015 | A / H5N8 | Asia / Korea, Republic of / Jeollanam-do | Eun-Kyoung Lee (Animal and Plant Quarantine Agency (APQA) / Avian Influenza Research and Diagsnotic Division) |
| EPI_ISL_234358 | A/broiler duck/Korea/H2826/2015 | A / H5N8 | Asia / Korea, Republic of / Jeollanam-do | Eun-Kyoung Lee (Animal and Plant Quarantine Agency (APQA) / Avian Influenza Research and Diagsnotic Division) |
| EPI_ISL_234359 | A/broiler duck/Korea/H2827/2015 | A / H5N8 | Asia / Korea, Republic of / Jeollanam-do | Eun-Kyoung Lee (Animal and Plant Quarantine Agency (APQA) / Avian Influenza Research and Diagsnotic Division) |
| EPI_ISL_234360 | A/broiler duck/Korea/LBM334/2015 | A / H5N8 | Asia / Korea, Republic of / Jeollanam-do | Eun-Kyoung Lee (Animal and Plant Quarantine Agency (APQA) / Avian Influenza Research and Diagsnotic Division) |
| EPI_ISL_234361 | A/mallard/Korea/H2891/2015 | A / H5N8 | Asia / Korea, Republic of / Jeollanam-do | Eun-Kyoung Lee (Animal and Plant Quarantine Agency (APQA) / Avian Influenza Research and Diagsnotic Division) |
| EPI_ISL_234362 | A/broiler duck/Korea/H2910/2015 | A / H5N8 | Asia / Korea, Republic of / Jeollanam-do | Eun-Kyoung Lee (Animal and Plant Quarantine Agency (APQA) / Avian Influenza Research and Diagsnotic Division) |
| EPI_ISL_234363 | A/broiler duck/Korea/H3078/2015 | A / H5N8 | Asia / Korea, Republic of / Jeollanam-do | Eun-Kyoung Lee (Animal and Plant Quarantine Agency (APQA) / Avian Influenza Research and Diagsnotic Division) |
| EPI_ISL_234364 | A/broiler duck/Korea/H3092/2015 | A / H5N8 | Asia / Korea, Republic of / Jeollanam-do | Eun-Kyoung Lee (Animal and Plant Quarantine Agency (APQA) / Avian Influenza Research and Diagsnotic Division) |
| EPI_ISL_234365 | A/broiler duck/Korea/H3093/2015 | A / H5N8 | Asia / Korea, Republic of / Jeollanam-do | Eun-Kyoung Lee (Animal and Plant Quarantine Agency (APQA) / Avian Influenza Research and Diagsnotic Division) |
| EPI_ISL_234366 | A/broiler duck/Korea/H3098/2015 | A / H5N8 | Asia / Korea, Republic of / Jeollanam-do | Eun-Kyoung Lee (Animal and Plant Quarantine Agency (APQA) / Avian Influenza Research and Diagsnotic Division) |
| EPI_ISL_234367 | A/broiler duck/Korea/H3104/2015 | A / H5N8 | Asia / Korea, Republic of / Jeollanam-do | Eun-Kyoung Lee (Animal and Plant Quarantine Agency (APQA) / Avian Influenza Research and Diagsnotic Division) |
| EPI_ISL_234368 | A/broiler duck/Korea/H3298/2015 | A / H5N8 | Asia / Korea, Republic of / Jeollanam-do | Eun-Kyoung Lee (Animal and Plant Quarantine Agency (APQA) / Avian Influenza Research and Diagsnotic Division) |
| EPI_ISL_234369 | A/broiler duck/Korea/H3301/2015 | A / H5N8 | Asia / Korea, Republic of / Jeollanam-do | Eun-Kyoung Lee (Animal and Plant Quarantine Agency (APQA) / Avian Influenza Research and Diagsnotic Division) |
| EPI_ISL_234370 | A/broiler duck/Korea/H3302/2015 | A / H5N8 | Asia / Korea, Republic of / Jeollanam-do | Eun-Kyoung Lee (Animal and Plant Quarantine Agency (APQA) / Avian Influenza Research and Diagsnotic Division) |

| | | | | |
|---|---|---|---|---|
| EPI_ISL_234371 | A/breeder duck/Korea/16AQ17/2016 | A / H5N8 | Asia / Korea, Republic of / Gyeonggi-do | Eun-Kyoung Lee (Animal and Plant Quarantine Agency (APQA) / Avian Influenza Research and Diagsnotic Division) |
| EPI_ISL_234372 | A/duck/Korea/16A02/2016 | A / H5N8 | Asia / Korea, Republic of / Gyeonggi-do | Eun-Kyoung Lee (Animal and Plant Quarantine Agency (APQA) / Avian Influenza Research and Diagsnotic Division) |
| EPI_ISL_234376 | A/migratory waterfowl/Hubei/Chenhu1306/2014 | A / H5N6 | Asia / China | |
| EPI_ISL_234377 | A/migratory waterfowl/Hubei/Chenhu1347/2014 | A / H5N6 | Asia / China | |
| EPI_ISL_234670 | A/Copsychus saularis/Guangdong/SW8/2014 | A / H5N6 | Asia / China | |
| EPI_ISL_234671 | A/Gallinula chloropus/Guangdong/GZ174/2014 | A / H5N6 | Asia / China | |
| EPI_ISL_234679 | A/Syrrhaptes paradoxus/Guangdong/ZH283/2015 | A / H5N6 | Asia / China | |
| EPI_ISL_235199 | A/duck/Zhejiang/6DK19-MA/2013 | A / H5N2 | Asia / China / Zhejiang Province | |
| EPI_ISL_235740 | A/mallard/Alaska/AH0088535/2016 | A / H5N2 | North America / United States | |
| EPI_ISL_235771 | A/mandarin duck/Gifu/2112D001/2014 | A / H5N8 | Asia / Japan / Gifu | |
| EPI_ISL_235772 | A/tundra swan/Tottori/C6nk/2014 | A / H5N8 | Asia / Japan / Tottori | |
| EPI_ISL_236802 | A/chicken/BC/FAV2/2015 | A / H5N1 | North America / Canada | |
| EPI_ISL_237551 | A/goose/Shandong/LQ01694/2013 | A / H5N8 | Asia / China / Shandong Province | Yuhai Bi (Institute of Microbiology, Chinese Academy of Sciences) |
| EPI_ISL_237552 | A/chicken/Shandong/KD02699/2013 | A / H5N8 | Asia / China / Shandong Province | Yuhai Bi (Institute of Microbiology, Chinese Academy of Sciences) |
| EPI_ISL_237553 | A/duck/India/10CA01/2016 | A / H5N8 | Asia / India / National Capital Territory of Delhi | Shanmuga sundaram Nagarajan (ICAR-National Institute of High Security Animal Diseases / Indian Council of Agricultural Research) |
| EPI_ISL_237554 | A/painted stork/India/10CA03/2016 | A / H5N8 | Asia / India / Madhya Pradesh / Gwalior | Shanmuga sundaram Nagarajan (ICAR-National Institute of High Security Animal Diseases / Indian Council of Agricultural Research) |
| EPI_ISL_237730 | A/mute_swan/Hungary/51049/2016 | A / H5N8 | Europe / Hungary / Csongrad megye | Adam Dan (Danam.Vet.Molbiol) |
| EPI_ISL_237731 | A/domestic_turkey/Hungary/53433/2016 | A / H5N8 | Europe / Hungary / Bekes megye | Adam Dan (Danam.Vet.Molbiol) |
| EPI_ISL_237732 | A/tufted_duck/Germany-SH/R8446/2016 | A / H5N8 | Europe / Germany / Schleswig-Holstein / Lake Plön | Anne Pohlmann (Friedrich-Loeffler-Institut) |
| EPI_ISL_237733 | A/tufted_duck/Germany/AR8444-L01986/2016 | A / H5N8 | Europe / Germany / Schleswig-Holstein / Lake Plön | Anne Pohlmann (Friedrich-Loeffler-Institut) |
| EPI_ISL_237792 | A/wild duck/Germany-BW/R8455/2016 | A / H5N8 | Europe / Germany / Baden-Wuerttemberg | Elke Starick (Friedrich-Loeffler-Institut) |
| EPI_ISL_237921 | A/wild duck/Poland/82A/2016 | A / H5N8 | Europe / Poland / West Pomeranian Voivodeship | Edyta ?wi?to? (National Veterinary Research Institut Poland, PIWet-PIB) |
| EPI_ISL_237922 | A/tufted duck/Denmark/17740-1/2016 | A / H5N8 | Europe / Denmark / Region Hovedstaden / Kobenhavns Kommune / Stadsgraven, Christiania | Charlotte Kristiane Hjulsager (Statens Serum Institute / Microbiological Diagnostic and Virology) |
| EPI_ISL_237944 | A/tufted_duck/Germany/AR8444-L01987/2016 | A / H5N8 | Europe / Germany / Schleswig-Holstein / Lake Plön | Anne Pohlmann (Friedrich-Loeffler-Institut) |
| EPI_ISL_237945 | A/tufted_duck/Germany/AR8459-L01988/2016 | A / H5N8 | Europe / Germany / Baden-Wuerttemberg / Lake Constance | Anne Pohlmann (Friedrich-Loeffler-Institut) |
| EPI_ISL_237947 | A/duck/Japan/AnimalQuarantine-HE72/2015 | A / H5N6 | Asia / Japan | Takehiko Saito (National Institute of Animal Health) |
| EPI_ISL_237952 | A/chicken/Japan/AnimalQuarantine-HE144/2016 | A / H5N6 | Asia / Japan | Takehiko Saito (National Institute of Animal Health) |
| EPI_ISL_237958 | A/tufted duck/Germany-SH/R8444/2016 | A / H5N8 | Europe / Germany / Schleswig-Holstein | Elke Starick (Friedrich-Loeffler-Institut) |
| EPI_ISL_237964 | A/Mulard_duck/Hungary/54494/2016 | A / H5N8 | Europe / Hungary / Bacs-Kiskun / Kiskunmajsa | Adam Dan (Danam.Vet.Molbiol) |
| EPI_ISL_237965 | A/goose/Hungary/55128/2016 | A / H5N8 | Europe / Hungary / Bacs-Kiskun / Bugac | Adam Dan (Danam.Vet.Molbiol) |
| EPI_ISL_237966 | A/duck/Hungary/55191/2016 | A / H5N8 | Europe / Hungary / Bacs-Kiskun / Kisszállás | Adam Dan (Danam.Vet.Molbiol) |
| EPI_ISL_237987 | A/gyrfalcon/Washington/41088/6/2014 | A / H5N8 | North America / United States | Takehiko Saito (National Institute of Animal Health) |
| EPI_ISL_238037 | A/turkey/Germany-SH/R8595/2016 | A / H5N8 | Europe / Germany / Schleswig-Holstein | Elke Starick (Friedrich-Loeffler-Institut) |
| EPI_ISL_238038 | A/chicken/Germany-MV/R8790/2016 | A / H5N8 | Europe / Germany / Mecklenburg-Vorpommern | Elke Starick (Friedrich-Loeffler-Institut) |
| EPI_ISL_238039 | A/chicken/Germany-SH/R8758/2016 | A / H5N8 | Europe / Germany / Schleswig-Holstein | Elke Starick (Friedrich-Loeffler-Institut) |
| EPI_ISL_238040 | A/tufted duck/Germany-SN/R8795/2016 | A / H5N8 | Europe / Germany / Saxony | Elke Starick (Friedrich-Loeffler-Institut) |
| EPI_ISL_238148 | A/Mandarin_duck/Korea/K16-187-3/2016 | A / H5N6 | Asia / Korea, Republic of / Chungcheongbuk-do / Gokgyo-cheon (36°45'12.3"N 127°07'12.7"E) | Jung Hoon Kwon (Konkuk University / College of Veterinary Medicine) |
| EPI_ISL_238196 | A/mute swan/Croatia/70/2016 | A / H5N8 | Europe / Croatia / Vukovarsko-Srijemska Zupanija / Cerna, river Bid | Vladimir Savi? (Croatian Veterinary Institute / Poultry Centre) |
| EPI_ISL_238197 | A/mute swan/Croatia/78/2016 | A / H5N8 | Europe / Croatia / Osjecko-Baranjska Zupanija / Bijelo Brdo | Vladimir Savi? (Croatian Veterinary Institute / Poultry Centre) |
| EPI_ISL_238605 | A/environment/Kagoshima/KU-ngr-I/2016(H5N6) | A / H5N6 | Asia / Japan / Kagoshima | Makoto Ozawa (Kagoshima University ) |
| EPI_ISL_238893 | A/Chicken/Sweden/SVA161122KU0453/SZ0209316/2016 | A / H5N8 | Europe / Sweden / Skane Lan / Helsingborgs Kommun / City of Mörarp, Lat. 56.053495 Long. 12 | Siamak Zohari (National Veterinary Institute) |
| EPI_ISL_238894 | A/Chicken/Sweden/SVA161122KU0453/SZ0209317/2016 | A / H5N8 | Europe / Sweden / Skane Lan / Helsingborgs Kommun / City of Mörarp, Lat. 56.053495 Long. 12. | Siamak Zohari (National Veterinary Institute) |
| EPI_ISL_238895 | A/Chicken/Sweden/SVA161122KU0453/SZ0209318/2016 | A / H5N8 | Europe / Sweden / Skane Lan / Helsingborgs Kommun / City of Mörarp, Lat. 56.053495 Long. 12. | Siamak Zohari (National Veterinary Institute) |
| EPI_ISL_238896 | A/Chicken/Sweden/SVA161122KU0453/SZ0209321/2016 | A / H5N8 | Europe / Sweden / Skane Lan / Helsingborgs Kommun / City of Mörarp, Lat. 56.053495 Long. 12. | Siamak Zohari (National Veterinary Institute) |
| EPI_ISL_238897 | A/Common Goldeneye/Sweden/SVA161117KU0322/SZ000216 | A / H5N8 | Europe / Sweden / Skane Lan / Vellinge Kommun / Vellinge, Höllviken | Siamak Zohari (National Veterinary Institute) |
| EPI_ISL_239069 | A/mute swan/Croatia/85/2016 | A / H5N8 | Europe / Croatia / Osjecko-Baranjska Zupanija / Bijelo Brdo | Vladimir Savi? (Croatian Veterinary Institute / Poultry Centre) |
| EPI_ISL_239260 | A/chicken/Korea/H23/2016(H5N6) | A / H5N6 | Asia / Korea, Republic of / Gyeonggi-do | Youn-Jeong Lee (Animal and Plant Quarantine Agency (APQA) / Avian Influenza Research and Diagsnotic Division) |
| EPI_ISL_239261 | A/chicken/Korea/HN1/2016(H5N6) | A / H5N6 | Asia / Korea, Republic of / Jeollanam-do | Youn-Jeong Lee (Animal and Plant Quarantine Agency (APQA) / Avian Influenza Research and Diagsnotic Division) |
| EPI_ISL_239262 | A/duck/Korea/ES2/2016(H5N6) | A / H5N6 | Asia / Korea, Republic of / Chungcheongbuk-do | Youn-Jeong Lee (Animal and Plant Quarantine Agency (APQA) / Avian Influenza Research and Diagsnotic Division) |
| EPI_ISL_239263 | A/duck/Korea/H15/2016(H5N6) | A / H5N6 | Asia / Korea, Republic of / Chungcheongbuk-do | Youn-Jeong Lee (Animal and Plant Quarantine Agency (APQA) / Avian Influenza Research and Diagsnotic Division) |
| EPI_ISL_239266 | A/Eurasian eagle owl/Korea/960/2016(H5N6) | A / H5N6 | Asia / Korea, Republic of / Gangwon-do | Youn-Jeong Lee (Animal and Plant Quarantine Agency (APQA) / Avian Influenza Research and Diagsnotic Division) |
| EPI_ISL_239267 | A/mandarin duck/Korea/T102-1/2016(H5N6) | A / H5N6 | Asia / Korea, Republic of / Chungcheongnam-do | Youn-Jeong Lee (Animal and Plant Quarantine Agency (APQA) / Avian Influenza Research and Diagsnotic Division) |
| EPI_ISL_239269 | A/mandarin duck/Korea/WB246/2016(H5N6) | A / H5N6 | Asia / Korea, Republic of / Chungcheongnam-do | Youn-Jeong Lee (Animal and Plant Quarantine Agency (APQA) / Avian Influenza Research and Diagsnotic Division) |
| EPI_ISL_239271 | A/spot_billed_duck/Korea/WB141/2016(H5N6) | A / H5N6 | Asia / Korea, Republic of / Jeollabuk-do | Youn-Jeong Lee (Animal and Plant Quarantine Agency (APQA) / Avian Influenza Research and Diagsnotic Division) |
| EPI_ISL_239351 | A/duck/Hyogo/1/2016 | A / H5N6 | Asia / Japan / Hyogo | Ikuyo Takayama (National Institute of Infectious Diseases (NIID)) |
| EPI_ISL_239419 | A/chicken/Hungary/59048/2016 | A / H5N8 | Europe / Hungary / Jasz-Nagykun-Szolnok | Adam Dan (Danam.Vet.Molbiol) |
| EPI_ISL_239420 | A/duck/Hungary/60441/2016 | A / H5N8 | Europe / Hungary / Bacs-Kiskun | Adam Dan (Danam.Vet.Molbiol) |
| EPI_ISL_239421 | A/muscovy duck/Aomori/1-2T/2016 | A / H5N6 | Asia / Japan | Takehiko Saito (National Institute of Animal Health) |
| EPI_ISL_239422 | A/muscovy duck/Aomori/2-4T/2016 | A / H5N6 | Asia / Japan | Takehiko Saito (National Institute of Animal Health) |
| EPI_ISL_239423 | A/chicken/Niigata/2-7T/2016 | A / H5N6 | Asia / Japan | Takehiko Saito (National Institute of Animal Health) |
| EPI_ISL_239424 | A/muscovy duck/Aomori/1-10C/2016 | A / H5N6 | Asia / Japan | Takehiko Saito (National Institute of Animal Health) |
| EPI_ISL_239425 | A/chicken/Niigata/2-6T/2016 | A / H5N6 | Asia / Japan | Takehiko Saito (National Institute of Animal Health) |
| EPI_ISL_239426 | A/chicken/Niigata/2-8T/2016 | A / H5N6 | Asia / Japan | Takehiko Saito (National Institute of Animal Health) |
| EPI_ISL_239427 | A/chicken/Niigata/2-5C/2016 | A / H5N6 | Asia / Japan | Takehiko Saito (National Institute of Animal Health) |
| EPI_ISL_239428 | A/muscovy duck/Aomori/2-2T/2016 | A / H5N6 | Asia / Japan | Takehiko Saito (National Institute of Animal Health) |
| EPI_ISL_239429 | A/muscovy duck/Aomori/1-7T/2016 | A / H5N6 | Asia / Japan | Takehiko Saito (National Institute of Animal Health) |
| EPI_ISL_239430 | A/muscovy duck/Aomori/2-1C/2016 | A / H5N6 | Asia / Japan | Takehiko Saito (National Institute of Animal Health) |
| EPI_ISL_239431 | A/muscovy duck/Aomori/2-3T/2016 | A / H5N6 | Asia / Japan | Takehiko Saito (National Institute of Animal Health) |
| EPI_ISL_239433 | A/muscovy duck/Aomori/1-3T/2016 | A / H5N6 | Asia / Japan | Takehiko Saito (National Institute of Animal Health) |
| EPI_ISL_239434 | A/tufted_duck/Switzerland/V246-L02001/2016 | A / H5N8 | Europe / Switzerland / Neuenburgersee | Anne Pohlmann (Friedrich-Loeffler-Institut) |
| EPI_ISL_239435 | A/tufted_duck/Switzerland/V254-L02002/2016 | A / H5N8 | Europe / Switzerland / Lake Constance | Anne Pohlmann (Friedrich-Loeffler-Institut) |
| EPI_ISL_239436 | A/tufted_duck/Switzerland/V237-L02003/2016 | A / H5N8 | Europe / Switzerland / Bielersee | Anne Pohlmann (Friedrich-Loeffler-Institut) |

| | | | | |
|---|---|---|---|---|
| EPI_ISL_239437 | A/caspian_gull/Switzerland/V247-L02004/2016 | A / H5N8 | Europe / Switzerland / Neuenburgersee | Anne Pohlmann (Friedrich-Loeffler-Institut) |
| EPI_ISL_239573 | A/crane/Kagoshima/KU-4/2016(H5N6) | A / H5N6 | Asia / Japan / Kagoshima | Makoto Ozawa (Kagoshima University ) |
| EPI_ISL_239801 | A/turkey/England/052131/2016 | A / H5N8 | Europe / United Kingdom / Lincolnshire / Louth | James Seekings (Animal and Plant Health Agency (APHA) / Virology Department) |
| EPI_ISL_239802 | A/Common-coot/Egypt/CA285/2016 | A / H5N8 | Africa / Egypt / Damietta | Mahmoud Mohamed Naguib (Friedrich-Loeffler-Institut) |
| EPI_ISL_239997 | A/decoy_duck/France/161104e/2016 | A / H5N8 | Europe / France / Region Nord-Pas-De Calais / Departement du Pas-de-Calais | Francois-Xavier Briand (ANSES Agence Nationale De Securite Sanitaire De L'alimentation / Laboratoire de Ploufragan-Plouzané) |
| EPI_ISL_239999 | A/black headed gull/Ibaraki/235T/2016 | A / H5N6 | Asia / Japan | Takehiko Saito (National Institute of Animal Health) |
| EPI_ISL_240000 | A/black headed gull/Ibaraki/253T/2016 | A / H5N6 | Asia / Japan | Takehiko Saito (National Institute of Animal Health) |
| EPI_ISL_240001 | A/mute swan/Ibaraki/239C/2016 | A / H5N6 | Asia / Japan | Takehiko Saito (National Institute of Animal Health) |
| EPI_ISL_240002 | A/mute swan/Ibaraki/236T/2016 | A / H5N6 | Asia / Japan | Takehiko Saito (National Institute of Animal Health) |
| EPI_ISL_240003 | A/mute swan/Ibaraki/250T/2016 | A / H5N6 | Asia / Japan | Takehiko Saito (National Institute of Animal Health) |
| EPI_ISL_240004 | A/mute swan/Ibaraki/245C/2016 | A / H5N6 | Asia / Japan | Takehiko Saito (National Institute of Animal Health) |
| EPI_ISL_240005 | A/mute swan/Ibaraki/232C/2016 | A / H5N6 | Asia / Japan | Takehiko Saito (National Institute of Animal Health) |
| EPI_ISL_240006 | A/mute swan/Ibaraki/247C/2016 | A / H5N6 | Asia / Japan | Takehiko Saito (National Institute of Animal Health) |
| EPI_ISL_240007 | A/mute swan/Ibaraki/240C/2016 | A / H5N6 | Asia / Japan | Takehiko Saito (National Institute of Animal Health) |
| EPI_ISL_240008 | A/mute swan/Ibaraki/249T/2016 | A / H5N6 | Asia / Japan | Takehiko Saito (National Institute of Animal Health) |
| EPI_ISL_240009 | A/mute swan/Ibaraki/242T/2016 | A / H5N6 | Asia / Japan | Takehiko Saito (National Institute of Animal Health) |
| EPI_ISL_240010 | A/black headed gull/Ibaraki/235C/2016 | A / H5N6 | Asia / Japan | Takehiko Saito (National Institute of Animal Health) |
| EPI_ISL_240011 | A/mute swan/Ibaraki/254C/2016 | A / H5N6 | Asia / Japan | Takehiko Saito (National Institute of Animal Health) |
| EPI_ISL_240012 | A/duck/France/161108h/2016 | A / H5N8 | Europe / France / Midi-Pyrenees / Departement du Tarn | Francois-Xavier Briand (ANSES Agence Nationale De Securite Sanitaire De L'alimentation / Laboratoire de Ploufragan-Plouzané) |
| EPI_ISL_240101 | A/mute swan/Croatia/102/2016 | A / H5N5 | Europe / Croatia / Sisacko-Moslavacka Zupanija / Crkveni Bok | Vladimir Savi? (Croatian Veterinary Institute / Poultry Centre) |
| EPI_ISL_240102 | A/domestic goose/Poland/33/2016 | A / H5N8 | Europe / Poland / Lubusz Voivodeship | Edyta ?wi?to? (National Veterinary Research Institut Poland, PIWet-PIB) |
| EPI_ISL_240103 | A/domestic goose/Poland/72/2016 | A / H5N8 | Europe / Poland / Lubusz Voivodeship | Edyta ?wi?to? (National Veterinary Research Institut Poland, PIWet-PIB) |
| EPI_ISL_240104 | A/chicken/Poland/79A/2016 | A / H5N8 | Europe / Poland / Subcarpathian Voivodeship | Edyta ?wi?to? (National Veterinary Research Institut Poland, PIWet-PIB) |
| EPI_ISL_240105 | A/chicken/Poland/85A/2016 | A / H5N8 | Europe / Poland / Lesser Poland Voivodeship | Edyta ?wi?to? (National Veterinary Research Institut Poland, PIWet-PIB) |
| EPI_ISL_240106 | A/mute swan/Poland/108/2016 | A / H5N8 | Europe / Poland / Masovian Voivodeship | Edyta ?wi?to? (National Veterinary Research Institut Poland, PIWet-PIB) |
| EPI_ISL_240107 | A/turkey/Poland/78/2016 | A / H5N8 | Europe / Poland / Lubusz Voivodeship | Edyta ?wi?to? (National Veterinary Research Institut Poland, PIWet-PIB) |
| EPI_ISL_240108 | A/turkey/Poland/83/2016 | A / H5N8 | Europe / Poland / Lubusz Voivodeship | Edyta ?wi?to? (National Veterinary Research Institut Poland, PIWet-PIB) |
| EPI_ISL_240109 | A/chicken/Kalmykia/2661/2016 | A / H5N8 | Europe / Russian Federation / Republic of Kalmykia | Ivan Susloparov (State Research Center of Virology and Biotechnology (VECTOR) / Emerging Zoonotic Diseases and Influenza) |
| EPI_ISL_240110 | A/chicken/Astrakhan/3131/2016 | A / H5N8 | Europe / Russian Federation / Astrakhan Oblast | Ivan Susloparov (State Research Center of Virology and Biotechnology (VECTOR) / Emerging Zoonotic Diseases and Influenza) |
| EPI_ISL_240111 | A/herring gull/Poland/84/2016 | A / H5N8 | Europe / Poland / West Pomeranian Voivodeship / Swinoujscie | Edyta ?wi?to? (National Veterinary Research Institut Poland, PIWet-PIB) |
| EPI_ISL_2402936 | A/chicken/Czech Republic/10251-1/2021 | A / H5N8 | Europe / Czech Republic / Stredocesky Kraj / Okres Melnik / Chorusice | Alexander Nagy (State Veterinary Institute Prague) |
| EPI_ISL_2402937 | A/chicken/Czech Republic/10251-2/2021 | A / H5N8 | Europe / Czech Republic / Stredocesky Kraj / Okres Melnik / Chorusice | Alexander Nagy (State Veterinary Institute Prague) |
| EPI_ISL_2402938 | A/chicken/Czech Republic/10405/2021 | A / H5N8 | Europe / Czech Republic / Stredocesky Kraj / Okres Mlada Boleslav | Alexander Nagy (State Veterinary Institute Prague) |
| EPI_ISL_240527 | A/chicken/Niigata/1-1T/2016 | A / H5N6 | Asia / Japan | Takehiko Saito (National Institute of Animal Health) |
| EPI_ISL_240528 | A/chicken/Niigata/1-2C/2016 | A / H5N6 | Asia / Japan | Takehiko Saito (National Institute of Animal Health) |
| EPI_ISL_240529 | A/muscovy duck/Aomori/1-3T/2016s | A / H5N6 | Asia / Japan | Takehiko Saito (National Institute of Animal Health) |
| EPI_ISL_240530 | A/chicken/Niigata/1-3C/2016 | A / H5N6 | Asia / Japan | Takehiko Saito (National Institute of Animal Health) |
| EPI_ISL_240531 | A/chicken/Niigata/1-4C/2016 | A / H5N6 | Asia / Japan | Takehiko Saito (National Institute of Animal Health) |
| EPI_ISL_240532 | A/muscovy duck/Aomori/1-1T/2016 | A / H5N6 | Asia / Japan | Takehiko Saito (National Institute of Animal Health) |
| EPI_ISL_240540 | A/chicken/Croatia/103/2016 | A / H5N8 | Europe / Croatia / Viroviticko-Podravska Zupanija / Križnica | Vladimir Savi? (Croatian Veterinary Institute / Poultry Centre) |
| EPI_ISL_240599 | A/chicken/Hokkaido/1-3-7T/2016 | A / H5N6 | Asia / Japan | Takehiko Saito (National Institute of Animal Health) |
| EPI_ISL_240601 | A/mute swan/Kyoto/8T/2016 | A / H5N6 | Asia / Japan | Takehiko Saito (National Institute of Animal Health) |
| EPI_ISL_240604 | A/mute swan/Kyoto/6T/2016 | A / H5N6 | Asia / Japan | Takehiko Saito (National Institute of Animal Health) |
| EPI_ISL_240605 | A/mute swan/Kyoto/5T/2016 | A / H5N6 | Asia / Japan | Takehiko Saito (National Institute of Animal Health) |
| EPI_ISL_240606 | A/chicken/Miyazaki/1-4C/2016 | A / H5N6 | Asia / Japan | Takehiko Saito (National Institute of Animal Health) |
| EPI_ISL_240607 | A/mute swan/Kyoto/4T/2016 | A / H5N6 | Asia / Japan | Takehiko Saito (National Institute of Animal Health) |
| EPI_ISL_240608 | A/black headed gull/Ibaraki/258T/2016 | A / H5N6 | Asia / Japan | Takehiko Saito (National Institute of Animal Health) |
| EPI_ISL_240609 | A/mute swan/Kyoto/3T/2016 | A / H5N6 | Asia / Japan | Takehiko Saito (National Institute of Animal Health) |
| EPI_ISL_240611 | A/mute swan/Ibaraki/269T/2016 | A / H5N6 | Asia / Japan | Takehiko Saito (National Institute of Animal Health) |
| EPI_ISL_240612 | A/chicken/Hokkaido/1-1C2C/2016 | A / H5N6 | Asia / Japan | Takehiko Saito (National Institute of Animal Health) |
| EPI_ISL_240613 | A/black headed gull/Ibaraki/265T/2016 | A / H5N6 | Asia / Japan | Takehiko Saito (National Institute of Animal Health) |
| EPI_ISL_240615 | A/mute swan/Ibaraki/255C/2016 | A / H5N6 | Asia / Japan | Takehiko Saito (National Institute of Animal Health) |
| EPI_ISL_240617 | A/mute swan/Kyoto/2T/2016 | A / H5N6 | Asia / Japan | Takehiko Saito (National Institute of Animal Health) |
| EPI_ISL_240618 | A/chicken/Hokkaido/1-3-7C/2016 | A / H5N6 | Asia / Japan | Takehiko Saito (National Institute of Animal Health) |
| EPI_ISL_240619 | A/black swan/Ibaraki/272C/2016 | A / H5N6 | Asia / Japan | Takehiko Saito (National Institute of Animal Health) |
| EPI_ISL_240621 | A/chicken/Miyazaki/1-7C/2016 | A / H5N6 | Asia / Japan | Takehiko Saito (National Institute of Animal Health) |
| EPI_ISL_240622 | A/chicken/Miyazaki/1-6T/2016 | A / H5N6 | Asia / Japan | Takehiko Saito (National Institute of Animal Health) |
| EPI_ISL_240624 | A/chicken/Miyazaki/1-6C/2016 | A / H5N6 | Asia / Japan | Takehiko Saito (National Institute of Animal Health) |
| EPI_ISL_240625 | A/chicken/Miyazaki/1-7T/2016 | A / H5N6 | Asia / Japan | Takehiko Saito (National Institute of Animal Health) |
| EPI_ISL_240626 | A/black swan/Ibaraki/259T/2016 | A / H5N6 | Asia / Japan | Takehiko Saito (National Institute of Animal Health) |
| EPI_ISL_240627 | A/mute swan/Ibaraki/211T/2016 | A / H5N6 | Asia / Japan | Takehiko Saito (National Institute of Animal Health) |
| EPI_ISL_240628 | A/mute swan/Kyoto/1T/2016 | A / H5N6 | Asia / Japan | Takehiko Saito (National Institute of Animal Health) |
| EPI_ISL_240670 | A/mute swan/Croatia/9/2017 | A / H5N8 | Europe / Croatia / Grad Zagreb | Vladimir Savi? (Croatian Veterinary Institute / Poultry Centre) |
| EPI_ISL_240671 | A/mute swan/Croatia/104/2016 | A / H5N8 | Europe / Croatia / Sisacko-Moslavacka Zupanija / Mužilov?ica | Vladimir Savi? (Croatian Veterinary Institute / Poultry Centre) |
| EPI_ISL_240677 | A/domestic duck/Siberia/103/2016 | A / H5N8 | Europe / Russian Federation / Novosibirsk Oblast | Ivan Sobolev (Research Institute of Experimental and Clinical Medicine) |
| EPI_ISL_240678 | A/domestic duck/Siberia/50K/2016 | A / H5N8 | Europe / Russian Federation / Novosibirsk Oblast | Ivan Sobolev (Research Institute of Experimental and Clinical Medicine) |
| EPI_ISL_240702 | A/Eurasian wigeon/Netherlands/1/2016 | A / H5N8 | Europe / Netherlands / Provincie Noord-Holland / Edam-Volendam | Maria Johanna Poen (Erasmus Medical Center / Department of Virology) |
| EPI_ISL_240703 | A/Hunan/55555/2016 | A / H5N6 | Asia / China / Hunan Province | Lei Yang (WHO Chinese National Influenza Center / Virology Institute, Chinese CDC) |
| EPI_ISL_240704 | A/Guangxi/55726/2016 | A / H5N6 | Asia / China / Guangxi Zhuang Autonomous Region | Lei Yang (WHO Chinese National Influenza Center / Virology Institute, Chinese CDC) |
| EPI_ISL_240892 | A/turkey/Germany-NI/R10523/2016 | A / H5N8 | Europe / Germany / Lower Saxony | Elke Starick (Friedrich-Loeffler-Institut) |
| EPI_ISL_240893 | A/swan/Germany-SN/R10645/2016 | A / H5N5 | Europe / Germany / Saxony | Elke Starick (Friedrich-Loeffler-Institut) |
| EPI_ISL_241249 | A/domestic duck/Germany-MV/R9869/2016 | A / H5N8 | Europe / Germany / Mecklenburg-Vorpommern | Elke Starick (Friedrich-Loeffler-Institut) |

| | | | | |
|---|---|---|---|---|
| EPI_ISL_241745 | A/chicken/Kumamoto/1-7T/2016 | A / H5N6 | Asia / Japan | Takehiko Saito (National Institute of Animal Health) |
| EPI_ISL_241746 | A/black headed gull/Ibaraki/282T/2016 | A / H5N6 | Asia / Japan | Takehiko Saito (National Institute of Animal Health) |
| EPI_ISL_241747 | A/black-headed-gull/Ibaraki/267T/2016 | A / H5N6 | Asia / Japan | Takehiko Saito (National Institute of Animal Health) |
| EPI_ISL_241748 | A/chicken/Kumamoto/1-1T/2016 | A / H5N6 | Asia / Japan | Takehiko Saito (National Institute of Animal Health) |
| EPI_ISL_241749 | A/mute swan/Ibaraki/263T/2016 | A / H5N6 | Asia / Japan | Takehiko Saito (National Institute of Animal Health) |
| EPI_ISL_241750 | A/mute swan/Ibaraki/262C/2016 | A / H5N6 | Asia / Japan | Takehiko Saito (National Institute of Animal Health) |
| EPI_ISL_241751 | A/chicken/Kumamoto/1-4C/2016 | A / H5N6 | Asia / Japan | Takehiko Saito (National Institute of Animal Health) |
| EPI_ISL_241752 | A/whooper swan/Ibaraki/301T/2016 | A / H5N6 | Asia / Japan | Takehiko Saito (National Institute of Animal Health) |
| EPI_ISL_241753 | A/black swan/Ibaraki/290C/2016 | A / H5N6 | Asia / Japan | Takehiko Saito (National Institute of Animal Health) |
| EPI_ISL_241754 | A/great crested grebe/Ibaraki/286T/2016 | A / H5N6 | Asia / Japan | Takehiko Saito (National Institute of Animal Health) |
| EPI_ISL_241755 | A/great crested grebe/Ibaraki/287T/2016 | A / H5N6 | Asia / Japan | Takehiko Saito (National Institute of Animal Health) |
| EPI_ISL_241756 | A/mute swan/Ibaraki/252T/2016 | A / H5N6 | Asia / Japan | Takehiko Saito (National Institute of Animal Health) |
| EPI_ISL_241757 | A/black swan/Ibaraki/256C/2016 | A / H5N6 | Asia / Japan | Takehiko Saito (National Institute of Animal Health) |
| EPI_ISL_241758 | A/chicken/Miyazaki/1-5C/2016 | A / H5N6 | Asia / Japan | Takehiko Saito (National Institute of Animal Health) |
| EPI_ISL_241759 | A/black headed gull/Ibaraki/291T/2016 | A / H5N6 | Asia / Japan | Takehiko Saito (National Institute of Animal Health) |
| EPI_ISL_241760 | A/chicken/Kumamoto/1-2C/2016 | A / H5N6 | Asia / Japan | Takehiko Saito (National Institute of Animal Health) |
| EPI_ISL_241761 | A/chicken/Kumamoto/1-3T/2016 | A / H5N6 | Asia / Japan | Takehiko Saito (National Institute of Animal Health) |
| EPI_ISL_241762 | A/black swan/Ibaraki/302C/2016 | A / H5N6 | Asia / Japan | Takehiko Saito (National Institute of Animal Health) |
| EPI_ISL_241763 | A/chicken/Kumamoto/1-6T/2016 | A / H5N6 | Asia / Japan | Takehiko Saito (National Institute of Animal Health) |
| EPI_ISL_241764 | A/chicken/Kumamoto/1-7C/2016 | A / H5N6 | Asia / Japan | Takehiko Saito (National Institute of Animal Health) |
| EPI_ISL_241765 | A/chicken/Kumamoto/1-4T/2016 | A / H5N6 | Asia / Japan | Takehiko Saito (National Institute of Animal Health) |
| EPI_ISL_241766 | A/chicken/Kumamoto/1-5C/2016 | A / H5N6 | Asia / Japan | Takehiko Saito (National Institute of Animal Health) |
| EPI_ISL_241767 | A/black headed gull/Ibaraki/284T/2016 | A / H5N6 | Asia / Japan | Takehiko Saito (National Institute of Animal Health) |
| EPI_ISL_241768 | A/mute swan/Ibaraki/270T/2016 | A / H5N6 | Asia / Japan | Takehiko Saito (National Institute of Animal Health) |
| EPI_ISL_241769 | A/mute swan/Ibaraki/248C/2016 | A / H5N6 | Asia / Japan | Takehiko Saito (National Institute of Animal Health) |
| EPI_ISL_241770 | A/chicken/Kumamoto/1-3C/2016 | A / H5N6 | Asia / Japan | Takehiko Saito (National Institute of Animal Health) |
| EPI_ISL_241771 | A/black swan/Ibaraki/271C/2016 | A / H5N6 | Asia / Japan | Takehiko Saito (National Institute of Animal Health) |
| EPI_ISL_241772 | A/black swan/Ibaraki/310C/2017 | A / H5N6 | Asia / Japan | Takehiko Saito (National Institute of Animal Health) |
| EPI_ISL_241773 | A/pochard/Ibaraki/268T/2016 | A / H5N6 | Asia / Japan | Takehiko Saito (National Institute of Animal Health) |
| EPI_ISL_241774 | A/chicken/Kumamoto/1-1C/2016 | A / H5N6 | Asia / Japan | Takehiko Saito (National Institute of Animal Health) |
| EPI_ISL_241775 | A/chicken/Kumamoto/1-2T/2016 | A / H5N6 | Asia / Japan | Takehiko Saito (National Institute of Animal Health) |
| EPI_ISL_241776 | A/black headed gull/Ibaraki/289T/2016 | A / H5N6 | Asia / Japan | Takehiko Saito (National Institute of Animal Health) |
| EPI_ISL_241777 | A/great crested grebe/Ibaraki/298C/2016 | A / H5N6 | Asia / Japan | Takehiko Saito (National Institute of Animal Health) |
| EPI_ISL_241778 | A/muteswan/Ibaraki/277T/2016 | A / H5N6 | Asia / Japan | Takehiko Saito (National Institute of Animal Health) |
| EPI_ISL_241779 | A/chicken/Kumamoto/1-5T/2016 | A / H5N6 | Asia / Japan | Takehiko Saito (National Institute of Animal Health) |
| EPI_ISL_241952 | A/breeder duck/Croatia/21/2017 | A / H5N8 | Europe / Croatia / Zagrebacka Zupanija / Sop Bukevski | Vladimir Savi? (Croatian Veterinary Institute / Poultry Centre) |
| EPI_ISL_241953 | A/mute swan/Croatia/15/2017 | A / H5N8 | Europe / Croatia / Koprivnicko-Krizevacka Zupanija / Lake Šoderica, Legrad | Vladimir Savi? (Croatian Veterinary Institute / Poultry Centre) |
| EPI_ISL_242401 | A/chicken/Gifu/1-1T/2017 | A / H5N6 | Asia / Japan | Takehiko Saito (National Institute of Animal Health) |
| EPI_ISL_242403 | A/chicken/Gifu/1-1C/2017 | A / H5N6 | Asia / Japan | Takehiko Saito (National Institute of Animal Health) |
| EPI_ISL_242404 | A/chicken/Gifu/1-10C/2017 | A / H5N6 | Asia / Japan | Takehiko Saito (National Institute of Animal Health) |
| EPI_ISL_242405 | A/whooper swan/Ibaraki/331C/2017 | A / H5N6 | Asia / Japan | Takehiko Saito (National Institute of Animal Health) |
| EPI_ISL_242406 | A/black swan/Ibaraki/341C/2017 | A / H5N6 | Asia / Japan | Takehiko Saito (National Institute of Animal Health) |
| EPI_ISL_242407 | A/chicken/Gifu/1-2T/2017 | A / H5N6 | Asia / Japan | Takehiko Saito (National Institute of Animal Health) |
| EPI_ISL_242408 | A/chicken/Gifu/1-10T/2017 | A / H5N6 | Asia / Japan | Takehiko Saito (National Institute of Animal Health) |
| EPI_ISL_242409 | A/chicken/Gifu/1-6C/2017 | A / H5N6 | Asia / Japan | Takehiko Saito (National Institute of Animal Health) |
| EPI_ISL_242411 | A/chicken/Gifu/1-6T/2017 | A / H5N6 | Asia / Japan | Takehiko Saito (National Institute of Animal Health) |
| EPI_ISL_242412 | A/chicken/Gifu/1-9C/2017 | A / H5N6 | Asia / Japan | Takehiko Saito (National Institute of Animal Health) |
| EPI_ISL_242413 | A/chicken/Gifu/1-8C/2017 | A / H5N6 | Asia / Japan | Takehiko Saito (National Institute of Animal Health) |
| EPI_ISL_242414 | A/chicken/Gifu/1-2C/2017 | A / H5N6 | Asia / Japan | Takehiko Saito (National Institute of Animal Health) |
| EPI_ISL_242415 | A/black swan/Ibaraki/341T/2017 | A / H5N6 | Asia / Japan | Takehiko Saito (National Institute of Animal Health) |
| EPI_ISL_242666 | A/chicken/Kumamoto/1-6C/2016 | A / H5N6 | Asia / Japan | Takehiko Saito (National Institute of Animal Health) |
| EPI_ISL_242817 | A/chicken/Czech Republic/206-17_2/2017(H5N8) | A / H5N8 | Europe / Czech Republic / South Moravian Region / Okres Breclav / Brod nad Dyjí | Alexander Nagy (State Veterinary Institute Prague) |
| EPI_ISL_242902 | A/wigeon/Italy/17VIR57-3/2017 | A / H5N8 | Europe / Italy / Friuli-Venezia Giulia / Province of Gorizia | Alice Fusaro (Istituto Zooprofilattico Sperimentale Delle Venezie) |
| EPI_ISL_242903 | A/Northern Pintail/Tottori/b37/2016 | A / H5N6 | Asia / Japan / Tottori | |
| EPI_ISL_243038 | A/domestic mallard/Korea/LBM176/2014 | A / H5N8 | Asia / Korea, Republic of | Eun-Kyoung Lee (Animal and Plant Quarantine Agency (APQA) / Avian Influenza Research and Diagsnotic Division) |
| EPI_ISL_243039 | A/broiler duck/Korea/LBM281/2014 | A / H5N8 | Asia / Korea, Republic of | Eun-Kyoung Lee (Animal and Plant Quarantine Agency (APQA) / Avian Influenza Research and Diagsnotic Division) |
| EPI_ISL_243040 | A/Korean native chicken/Korea/LBM282/2014 | A / H5N8 | Asia / Korea, Republic of | Eun-Kyoung Lee (Animal and Plant Quarantine Agency (APQA) / Avian Influenza Research and Diagsnotic Division) |
| EPI_ISL_243041 | A/Korean native chicken/Korea/LBM287/2014 | A / H5N8 | Asia / Korea, Republic of | Eun-Kyoung Lee (Animal and Plant Quarantine Agency (APQA) / Avian Influenza Research and Diagsnotic Division) |
| EPI_ISL_243042 | A/Korean native chicken/Korea/LBM288/2014 | A / H5N8 | Asia / Korea, Republic of | Eun-Kyoung Lee (Animal and Plant Quarantine Agency (APQA) / Avian Influenza Research and Diagsnotic Division) |
| EPI_ISL_243043 | A/Korean native chicken/Korea/LBM367/2014 | A / H5N8 | Asia / Korea, Republic of | Eun-Kyoung Lee (Animal and Plant Quarantine Agency (APQA) / Avian Influenza Research and Diagsnotic Division) |
| EPI_ISL_243048 | A/turkey/Czech Republic/38-17_5/2017 (H5N8) | A / H5N8 | Europe / Czech Republic / South Moravian Region / Okres Brno-Venkov / Ivan?ice-N?m?ice | Alexander Nagy (State Veterinary Institute Prague) |
| EPI_ISL_243049 | A/turkey/Germany-SH/R425/2017 | A / H5N5 | Europe / Germany / Schleswig-Holstein | Elke Starick (Friedrich-Loeffler-Institut) |
| EPI_ISL_243058 | A/black swan/Akita/1/2016 | A / H5N6 | Asia / Japan / Akita | |
| EPI_ISL_243059 | A/black swan/Akita/2/2016 | A / H5N6 | Asia / Japan / Akita | |
| EPI_ISL_243060 | A/teal/Tottori/1/2016 | A / H5N6 | Asia / Japan / Tottori | |
| EPI_ISL_243061 | A/teal/Tottori/2/2016 | A / H5N6 | Asia / Japan / Tottori | |
| EPI_ISL_243085 | A/wigeon/Italy/16VIR9616-3/2016 | A / H5N5 | Europe / Italy / Friuli-Venezia Giulia / Province of Gorizia | Alice Fusaro (Istituto Zooprofilattico Sperimentale Delle Venezie) |
| EPI_ISL_243344 | A/wild pigeon/Jilin/CC01/2014 | A / H5N6 | Asia / China | |
| EPI_ISL_243345 | A/environment/Jilin/CC02/2014 | A / H5N6 | Asia / China | |
| EPI_ISL_243672 | A/black swan/Ibaraki/365C/2017 | A / H5N6 | Asia / Japan | Takehiko Saito (National Institute of Animal Health) |
| EPI_ISL_243673 | A/chicken/Miyazaki/2-2C/2017 | A / H5N6 | Asia / Japan | Takehiko Saito (National Institute of Animal Health) |
| EPI_ISL_243674 | A/black swan/Ibaraki/356C/2017 | A / H5N6 | Asia / Japan | Takehiko Saito (National Institute of Animal Health) |

| | | | | |
|---|---|---|---|---|
| EPI_ISL_243675 | A/black swan/Ibaraki/348C/2017 | A / H5N6 | Asia / Japan | Takehiko Saito (National Institute of Animal Health) |
| EPI_ISL_243676 | A/black swan/Ibaraki/357T/2017 | A / H5N6 | Asia / Japan | Takehiko Saito (National Institute of Animal Health) |
| EPI_ISL_243677 | A/black swan/Ibaraki/345T/2017 | A / H5N6 | Asia / Japan | Takehiko Saito (National Institute of Animal Health) |
| EPI_ISL_243678 | A/black swan/Ibaraki/350C/2017 | A / H5N6 | Asia / Japan | Takehiko Saito (National Institute of Animal Health) |
| EPI_ISL_243679 | A/black swan/Ibaraki/345C/2017 | A / H5N6 | Asia / Japan | Takehiko Saito (National Institute of Animal Health) |
| EPI_ISL_243680 | A/chicken/Miyazaki/2-1T/2017 | A / H5N6 | Asia / Japan | Takehiko Saito (National Institute of Animal Health) |
| EPI_ISL_243681 | A/chicken/Miyazaki/2-4C/2017 | A / H5N6 | Asia / Japan | Takehiko Saito (National Institute of Animal Health) |
| EPI_ISL_243682 | A/black swan/Ibaraki/365T/2017 | A / H5N6 | Asia / Japan | Takehiko Saito (National Institute of Animal Health) |
| EPI_ISL_243683 | A/chicken/Miyazaki/2-1C/2017 | A / H5N6 | Asia / Japan | Takehiko Saito (National Institute of Animal Health) |
| EPI_ISL_243684 | A/chicken/Miyazaki/2-5T/2017 | A / H5N6 | Asia / Japan | Takehiko Saito (National Institute of Animal Health) |
| EPI_ISL_243685 | A/black swan/Ibaraki/357C/2017 | A / H5N6 | Asia / Japan | Takehiko Saito (National Institute of Animal Health) |
| EPI_ISL_243686 | A/whooper swan/Ibaraki/351C/2017 | A / H5N6 | Asia / Japan | Takehiko Saito (National Institute of Animal Health) |
| EPI_ISL_243687 | A/chicken/Miyazaki/2-3C/2017 | A / H5N6 | Asia / Japan | Takehiko Saito (National Institute of Animal Health) |
| EPI_ISL_243688 | A/black swan/Ibaraki/356T/2017 | A / H5N6 | Asia / Japan | Takehiko Saito (National Institute of Animal Health) |
| EPI_ISL_243689 | A/black swan/Ibaraki/350T/2017 | A / H5N6 | Asia / Japan | Takehiko Saito (National Institute of Animal Health) |
| EPI_ISL_243693 | A/mute swan/Croatia/30/2017 | A / H5N8 | Europe / Croatia / Osjecko-Baranjska Zupanija / Bilje | Vladimir Savi? (Croatian Veterinary Institute / Poultry Centre) |
| EPI_ISL_243695 | A/greylag goose/Croatia/33/2017 | A / H5N8 | Europe / Croatia / Bjelovarsko-Bilogorska Zupanija / Prekobrdo | Vladimir Savi? (Croatian Veterinary Institute / Poultry Centre) |
| EPI_ISL_243698 | A/mute swan/Croatia/42/2017 | A / H5N5 | Europe / Croatia / Osjecko-Baranjska Zupanija / road between villages Kozjak and Tikves | Vladimir Savi? (Croatian Veterinary Institute / Poultry Centre) |
| EPI_ISL_244486 | A/chicken/Vietnam/NCVD14-A324/2014 | A / H5N6 | Asia / Vietnam | |
| EPI_ISL_244487 | A/chicken/Vietnam/NCVD-15A22/2015 | A / H5N6 | Asia / Vietnam | |
| EPI_ISL_244488 | A/Pheasant/Vietnam/NCVD14-A367/2014 | A / H5N6 | Asia / Vietnam | |
| EPI_ISL_244489 | A/duck/Vietnam/NCVD14-A392/2014 | A / H5N6 | Asia / Vietnam | |
| EPI_ISL_244510 | A/duck/Vietnam/NCVD14-A415/2014 | A / H5N6 | Asia / Vietnam | |
| EPI_ISL_244511 | A/duck/Vietnam/NCVD14-A418/2014 | A / H5N6 | Asia / Vietnam | |
| EPI_ISL_244512 | A/duck/Vietnam/NCVD14-A503/2014 | A / H5N6 | Asia / Vietnam | |
| EPI_ISL_244513 | A/chicken/Vietnam/NCVD-15A17/2015 | A / H5N6 | Asia / Vietnam | |
| EPI_ISL_244514 | A/goose/Vietnam/NCVD-15A27/2015 | A / H5N6 | Asia / Vietnam | |
| EPI_ISL_244515 | A/chicken/Vietnam/NCVD-15A51/2015 | A / H5N6 | Asia / Vietnam | |
| EPI_ISL_244516 | A/chicken/Vietnam/NCVD-15A55/2015 | A / H5N6 | Asia / Vietnam | |
| EPI_ISL_244517 | A/duck/Vietnam/NCVD-15A57/2015 | A / H5N6 | Asia / Vietnam | |
| EPI_ISL_244518 | A/chicken/Vietnam/NCVD-15A59/2015 | A / H5N6 | Asia / Vietnam | |
| EPI_ISL_244519 | A/environment/Korea/W541/2016 | A / H5N6 | Asia / Korea, Republic of | |
| EPI_ISL_244520 | A/environment/Korea/W542/2016 | A / H5N6 | Asia / Korea, Republic of | |
| EPI_ISL_244521 | A/environment/Korea/W543/2016 | A / H5N6 | Asia / Korea, Republic of | |
| EPI_ISL_244522 | A/environment/Korea/W544/2016 | A / H5N6 | Asia / Korea, Republic of | |
| EPI_ISL_244523 | A/whooper swan/Korea/Gangjin 48/2016 | A / H5N6 | Asia / Korea, Republic of | |
| EPI_ISL_244524 | A/duck/Hubei/ZYSYF1/2015 | A / H5N6 | Asia / China | |
| EPI_ISL_244525 | A/duck/Hubei/ZYSYF24/2015 | A / H5N6 | Asia / China | |
| EPI_ISL_244526 | A/duck/Hubei/ZYSYG18/2015 | A / H5N6 | Asia / China | |
| EPI_ISL_244527 | A/duck/Hubei/ZYSYG5/2015 | A / H5N6 | Asia / China | |
| EPI_ISL_244528 | A/chicken/Hubei/ZYSJF11/2016 | A / H5N6 | Asia / China | |
| EPI_ISL_244529 | A/chicken/Hubei/ZYSJF16/2016 | A / H5N6 | Asia / China | |
| EPI_ISL_244530 | A/chicken/Hubei/ZYSJF22/2016 | A / H5N6 | Asia / China | |
| EPI_ISL_244531 | A/chicken/Hubei/ZYSJF38/2016 | A / H5N6 | Asia / China | |
| EPI_ISL_244532 | A/chicken/Taishun/TS12/2016 | A / H5N6 | Asia / China | |
| EPI_ISL_244533 | A/chicken/Taishun/TS90/2016 | A / H5N6 | Asia / China | |
| EPI_ISL_244534 | A/chicken/Taishun/TS2/2016 | A / H5N6 | Asia / China | |
| EPI_ISL_244535 | A/chicken/Ganzhou/GZ27/2015 | A / H5N6 | Asia / China | |
| EPI_ISL_244536 | A/chicken/Ganzhou/GZ21/2015 | A / H5N6 | Asia / China | |
| EPI_ISL_244537 | A/chicken/Ganzhou/GZ50/2015 | A / H5N6 | Asia / China | |
| EPI_ISL_244538 | A/whooper swan/Korea/Gangjin 49_2/2016 | A / H5N6 | Asia / Korea, Republic of | |
| EPI_ISL_244539 | A/whooper swan/Korea/Gangjin 49_1/2016 | A / H5N6 | Asia / Korea, Republic of | |
| EPI_ISL_247371 | A/mute swan/Ibaraki/219T/2016 | A / H5N6 | Asia / Japan / Ibaraki | Takehiko Saito (National Institute of Animal Health) |
| EPI_ISL_247372 | A/mute swan/Ibaraki/220C/2016 | A / H5N6 | Asia / Japan / Ibaraki | Takehiko Saito (National Institute of Animal Health) |
| EPI_ISL_247373 | A/black headed gull/Ibaraki/194C/2016 | A / H5N6 | Asia / Japan / Ibaraki | Takehiko Saito (National Institute of Animal Health) |
| EPI_ISL_247374 | A/mute swan/Ibaraki/214C/2016 | A / H5N6 | Asia / Japan / Ibaraki | Takehiko Saito (National Institute of Animal Health) |
| EPI_ISL_247375 | A/mute swan/Ibaraki/222C/2016 | A / H5N6 | Asia / Japan / Ibaraki | Takehiko Saito (National Institute of Animal Health) |
| EPI_ISL_247376 | A/mute swan/Ibaraki/208T/2016 | A / H5N6 | Asia / Japan / Ibaraki | Takehiko Saito (National Institute of Animal Health) |
| EPI_ISL_247377 | A/mute swan/Ibaraki/223C/2016 | A / H5N6 | Asia / Japan / Ibaraki | Takehiko Saito (National Institute of Animal Health) |
| EPI_ISL_247378 | A/chicken/Saga/1-1T/2017 | A / H5N6 | Asia / Japan / Saga | Takehiko Saito (National Institute of Animal Health) |
| EPI_ISL_247379 | A/chicken/Saga/1-5T/2017 | A / H5N6 | Asia / Japan / Saga | Takehiko Saito (National Institute of Animal Health) |
| EPI_ISL_247380 | A/chicken/Saga/1-4C/2017 | A / H5N6 | Asia / Japan / Saga | Takehiko Saito (National Institute of Animal Health) |
| EPI_ISL_247381 | A/mute swan/Ibaraki/217C/2016 | A / H5N6 | Asia / Japan / Ibaraki | Takehiko Saito (National Institute of Animal Health) |
| EPI_ISL_247382 | A/chicken/Saga/1-1C/2017 | A / H5N6 | Asia / Japan / Saga | Takehiko Saito (National Institute of Animal Health) |
| EPI_ISL_247383 | A/mute swan/Ibaraki/202C/2016 | A / H5N6 | Asia / Japan / Ibaraki | Takehiko Saito (National Institute of Animal Health) |
| EPI_ISL_247384 | A/chicken/Saga/1-3C/2017 | A / H5N6 | Asia / Japan / Saga | Takehiko Saito (National Institute of Animal Health) |
| EPI_ISL_247385 | A/chicken/Saga/1-2C/2017 | A / H5N6 | Asia / Japan / Saga | Takehiko Saito (National Institute of Animal Health) |
| EPI_ISL_247386 | A/mute swan/Ibaraki/214T/2016 | A / H5N6 | Asia / Japan / Ibaraki | Takehiko Saito (National Institute of Animal Health) |
| EPI_ISL_247387 | A/mute swan/Ibaraki/202T/2016 | A / H5N6 | Asia / Japan / Ibaraki | Takehiko Saito (National Institute of Animal Health) |
| EPI_ISL_247388 | A/mute swan/Ibaraki/222T/2016 | A / H5N6 | Asia / Japan / Ibaraki | Takehiko Saito (National Institute of Animal Health) |
| EPI_ISL_247389 | A/mute swan/Ibaraki/223T/2016 | A / H5N6 | Asia / Japan / Ibaraki | Takehiko Saito (National Institute of Animal Health) |
| EPI_ISL_247390 | A/mute swan/Ibaraki/218T/2016 | A / H5N6 | Asia / Japan / Ibaraki | Takehiko Saito (National Institute of Animal Health) |

| | | | | |
|---|---|---|---|---|
| EPI_ISL_247391 | A/mute swan/Ibaraki/217T/2016 | A / H5N6 | Asia / Japan / Ibaraki | Takehiko Saito (National Institute of Animal Health) |
| EPI_ISL_247392 | A/chicken/Saga/1-6T/2017 | A / H5N6 | Asia / Japan / Saga | Takehiko Saito (National Institute of Animal Health) |
| EPI_ISL_247393 | A/chicken/Saga/1-5C/2017 | A / H5N6 | Asia / Japan / Saga | Takehiko Saito (National Institute of Animal Health) |
| EPI_ISL_247394 | A/whooper swan/Ibaraki/188C/2016 | A / H5N6 | Asia / Japan / Ibaraki | Takehiko Saito (National Institute of Animal Health) |
| EPI_ISL_247395 | A/chicken/Saga/1-6C/2017 | A / H5N6 | Asia / Japan / Saga | Takehiko Saito (National Institute of Animal Health) |
| EPI_ISL_247396 | A/chicken/Saga/1-2T/2017 | A / H5N6 | Asia / Japan / Saga | Takehiko Saito (National Institute of Animal Health) |
| EPI_ISL_247397 | A/chicken/Saga/1-3T/2017 | A / H5N6 | Asia / Japan / Saga | Takehiko Saito (National Institute of Animal Health) |
| EPI_ISL_247398 | A/chicken/Saga/1-4T/2017 | A / H5N6 | Asia / Japan / Saga | Takehiko Saito (National Institute of Animal Health) |
| EPI_ISL_247399 | A/chicken/Saga/1-7C/2017 | A / H5N6 | Asia / Japan / Saga | Takehiko Saito (National Institute of Animal Health) |
| EPI_ISL_247400 | A/mute swan/Ibaraki/210T/2016 | A / H5N6 | Asia / Japan / Ibaraki | Takehiko Saito (National Institute of Animal Health) |
| EPI_ISL_247401 | A/mute swan/Ibaraki/221C/2016 | A / H5N6 | Asia / Japan / Ibaraki | Takehiko Saito (National Institute of Animal Health) |
| EPI_ISL_247402 | A/mute swan/Ibaraki/219C/2016 | A / H5N6 | Asia / Japan / Ibaraki | Takehiko Saito (National Institute of Animal Health) |
| EPI_ISL_247403 | A/chicken/Saga/1-7T/2017 | A / H5N6 | Asia / Japan / Saga | Takehiko Saito (National Institute of Animal Health) |
| EPI_ISL_247404 | A/mute swan/Ibaraki/210C/2016 | A / H5N6 | Asia / Japan / Ibaraki | Takehiko Saito (National Institute of Animal Health) |
| EPI_ISL_247405 | A/mute swan/Ibaraki/220T/2016 | A / H5N6 | Asia / Japan / Ibaraki | Takehiko Saito (National Institute of Animal Health) |
| EPI_ISL_247406 | A/black headed gull/Ibaraki/194T/2016 | A / H5N6 | Asia / Japan / Ibaraki | Takehiko Saito (National Institute of Animal Health) |
| EPI_ISL_247407 | A/mute swan/Ibaraki/218C/2016 | A / H5N6 | Asia / Japan / Ibaraki | Takehiko Saito (National Institute of Animal Health) |
| EPI_ISL_247429 | A/domestic duck/Germany-BB/R681ff/2017 | A / H5N8 | Europe / Germany / Brandenburg | Elke Starick (Friedrich-Loeffler-Institut) |
| EPI_ISL_247715 | A/long-eared owl/Voronezh/15/2017 | A / H5N8 | Europe / Russian Federation / Voronezh Oblast | Ivan Susloparov (State Research Center of Virology and Biotechnology (VECTOR) / Emerging Zoonotic Diseases and Influenza) |
| EPI_ISL_247716 | A/Ural owl/Voronezh/14/2017 | A / H5N8 | Europe / Russian Federation / Voronezh Oblast | Ivan Susloparov (State Research Center of Virology and Biotechnology (VECTOR) / Emerging Zoonotic Diseases and Influenza) |
| EPI_ISL_247717 | A/long-eared owl/Voronezh/16/2017 | A / H5N8 | Europe / Russian Federation / Voronezh Oblast | Ivan Susloparov (State Research Center of Virology and Biotechnology (VECTOR) / Emerging Zoonotic Diseases and Influenza) |
| EPI_ISL_247718 | A/chicken/Voronezh/18/2017 | A / H5N8 | Europe / Russian Federation / Voronezh Oblast | Ivan Susloparov (State Research Center of Virology and Biotechnology (VECTOR) / Emerging Zoonotic Diseases and Influenza) |
| EPI_ISL_247719 | A/chicken/Voronezh/19/2017 | A / H5N8 | Europe / Russian Federation / Voronezh Oblast | Ivan Susloparov (State Research Center of Virology and Biotechnology (VECTOR) / Emerging Zoonotic Diseases and Influenza) |
| EPI_ISL_247720 | A/chicken/Voronezh/20/2017 | A / H5N8 | Europe / Russian Federation / Voronezh Oblast | Ivan Susloparov (State Research Center of Virology and Biotechnology (VECTOR) / Emerging Zoonotic Diseases and Influenza) |
| EPI_ISL_247721 | A/turkey/Rostov-on-Don/11/2017 | A / H5N8 | Europe / Russian Federation / Rostov Oblast | Ivan Susloparov (State Research Center of Virology and Biotechnology (VECTOR) / Emerging Zoonotic Diseases and Influenza) |
| EPI_ISL_247722 | A/goose/Krasnodar/3144/2017 | A / H5N8 | Europe / Russian Federation / Krasnodar Krai | Ivan Susloparov (State Research Center of Virology and Biotechnology (VECTOR) / Emerging Zoonotic Diseases and Influenza) |
| EPI_ISL_247723 | A/mute swan/Krasnodar/25/2017 | A / H5N8 | Europe / Russian Federation / Krasnodar Krai | Ivan Susloparov (State Research Center of Virology and Biotechnology (VECTOR) / Emerging Zoonotic Diseases and Influenza) |
| EPI_ISL_247724 | A/wild duck/Tatarstan/3059/2016 | A / H5N8 | Europe / Russian Federation / Republic of Tatarstan | Ivan Susloparov (State Research Center of Virology and Biotechnology (VECTOR) / Emerging Zoonotic Diseases and Influenza) |
| EPI_ISL_247725 | A/chicken/Kalmykia/2643/2016 | A / H5N8 | Europe / Russian Federation / Republic of Kalmykia | Ivan Susloparov (State Research Center of Virology and Biotechnology (VECTOR) / Emerging Zoonotic Diseases and Influenza) |
| EPI_ISL_248634 | A/duck/Taiwan/1702004/2017 | A / H5N6 | Asia / Taiwan | Yu-Pin Liu (Animal Health Research Institute) |
| EPI_ISL_248657 | A/mute swan/Czech Republic/653-17/2017 (H5N8) | A / H5N8 | Europe / Czech Republic / Hlavni mesto Praha / Okres Praha / Praha 1, St?elecký ostrov | Alexander Nagy (State Veterinary Institute Prague) |
| EPI_ISL_248663 | A/mallard/Czech Republic/722-17_2/2017 (H5N8) | A / H5N8 | Europe / Czech Republic / Jihocesky Kraj / Okres Tabor / Praha 1, St?elecký ostrov | Alexander Nagy (State Veterinary Institute Prague) |
| EPI_ISL_248664 | A/mute swan/Croatia/61/2017 | A / H5N8 | Europe / Croatia / Karlovacka Zupanija / Karlovac | Vladimir Savi? (Croatian Veterinary Institute / Poultry Centre) |
| EPI_ISL_248666 | A/bronze turkey/Czech Republic/1414-17/2017 (H5N8) | A / H5N8 | Europe / Czech Republic / Ustecky Kraj / Okres Chomutov / Bílenice, GPS: 50°25'18.168''N, 13°30 | Alexander Nagy (State Veterinary Institute Prague) |
| EPI_ISL_248879 | A/chicken/Croatia/70/2017 | A / H5N8 | Europe / Croatia / Vukovarsko-Srijemska Zupanija / Mlaka Antinska | Vladimir Savi? (Croatian Veterinary Institute / Poultry Centre) |
| EPI_ISL_248934 | A/mute swan/Czech Republic/722-17_1/2017 (H5N8) | A / H5N8 | Europe / Czech Republic / Jihocesky Kraj / Okres Tabor / Lake Jordán, Van?urova Street | Alexander Nagy (State Veterinary Institute Prague) |
| EPI_ISL_249324 | A/chicken/Czech Republic/585-17_1/2017 (H5N8) | A / H5N8 | Europe / Czech Republic / Jihocesky Kraj / Okres Strakonice / Vod?any | Alexander Nagy (State Veterinary Institute Prague) |
| EPI_ISL_249681 | A/turkey/Germany-BB/R234ff/2017 | A / H5N8 | Europe / Germany / Brandenburg / LK Ostprignitz-Ruppin | Elke Starick (Friedrich-Loeffler-Institut) |
| EPI_ISL_249691 | A/chicken/Germany-MV/R10048/2016 | A / H5N8 | Europe / Germany / Mecklenburg-Vorpommern / Neukloster | Elke Starick (Friedrich-Loeffler-Institut) |
| EPI_ISL_249692 | A/chicken/Germany-NI/R11406/2016 | A / H5N8 | Europe / Germany / Lower Saxony / LK Northeim | Elke Starick (Friedrich-Loeffler-Institut) |
| EPI_ISL_250231 | A/domestic_duck/Siberia/49_feather/2016 | A / H5N8 | Europe / Russian Federation / Novosibirsk Oblast / Chany lake | Artem Fadeev (WHO National Influenza Centre Russian Federation) |
| EPI_ISL_250885 | A/mute swan/Czech Republic/54-17_2/2017 (H5N8) | A / H5N8 | Europe / Czech Republic / South Moravian Region / Okres Blansko / N?m?ice | Alexander Nagy (State Veterinary Institute Prague) |
| EPI_ISL_250886 | A/Indian Runner Duck/Czech Republic/749-17/2017 (H5N8) | A / H5N8 | Europe / Czech Republic / Kralovehradecky Kraj / Okres Nachod / Náchod-Babí GPS: 52°25'56.20 | Alexander Nagy (State Veterinary Institute Prague) |
| EPI_ISL_250887 | A/chicken/Czech Republic/55-17_1/2017 (H5N8) | A / H5N8 | Europe / Czech Republic / South Moravian Region / Okres Znoimo / Moravský Krumlov-Rakšice | Alexander Nagy (State Veterinary Institute Prague) |
| EPI_ISL_250902 | A/mute swan/Czech Republic/879-17/2017 (H5N8) | A / H5N8 | Europe / Czech Republic / Kralovehradecky Kraj / Okres Hradec Kralove / Most u Soutoku, GPS: | Alexander Nagy (State Veterinary Institute Prague) |
| EPI_ISL_250917 | A/duck/Czech Republic/1467-17/2017 (H5N8) | A / H5N8 | Europe / Czech Republic / Jihocesky Kraj / Okres Prachatice / Vlachovo B?ezí | Alexander Nagy (State Veterinary Institute Prague) |
| EPI_ISL_250918 | A/chicken/Czech Republic/1863-17_1/2017 (H5N8) | A / H5N8 | Europe / Czech Republic / Stredocesky Kraj / Okres Benesov / Dolní Kralovice | Alexander Nagy (State Veterinary Institute Prague) |
| EPI_ISL_250919 | A/chicken/Czech Republic/1896-17_1/2017 (H5N8) | A / H5N8 | Europe / Czech Republic / Jihocesky Kraj / Okres Jindrichuv Hradec / Nová Ves nad Lužnicí, GPS: | Alexander Nagy (State Veterinary Institute Prague) |
| EPI_ISL_250920 | A/mute swan/Czech Republic/2031-17/2017 (H5N5) | A / H5N5 | Europe / Czech Republic / Liberecky Kraj / Okres Liberec / ZOO Liberec, GPS: 50°46'49.640''N, 15 | Alexander Nagy (State Veterinary Institute Prague) |
| EPI_ISL_252881 | A/chicken/Czech Republic/2764-17_1/2017 (H5N8) | A / H5N8 | Europe / Czech Republic / Karlovarsky Kraj / Okres Karlovy Vary / Boži?any, GPS: 50°16'18.814''N | Alexander Nagy (State Veterinary Institute Prague) |
| EPI_ISL_252882 | A/chicken/Czech Republic/2764-17_2/2017 (H5N8) | A / H5N8 | Europe / Czech Republic / Karlovarsky Kraj / Okres Karlovy Vary / Boži?any, GPS: 50°16'18.814''N | Alexander Nagy (State Veterinary Institute Prague) |
| EPI_ISL_253029 | A/chicken/Zhejiang/194/2016 | A / H5N2 | Asia / China | |
| EPI_ISL_253030 | A/chicken/Zhejiang/209/2016 | A / H5N2 | Asia / China | |
| EPI_ISL_253031 | A/chicken/Zhejiang/217/2016 | A / H5N2 | Asia / China | |
| EPI_ISL_253032 | A/chicken/Zhejiang/232/2016 | A / H5N2 | Asia / China | |
| EPI_ISL_253036 | A/turkey/England/003778/2017 | A / H5N8 | Europe / United Kingdom / Lincolnshire | James Seekings (Animal and Plant Health Agency (APHA) / Virology Department) |
| EPI_ISL_253037 | A/chicken/Wales/000023/2016 | A / H5N8 | Europe / United Kingdom / County of Carmarthenshire | James Seekings (Animal and Plant Health Agency (APHA) / Virology Department) |
| EPI_ISL_253038 | A/pochard_duck/England/SA12_157809/2016 | A / H5N8 | Europe / United Kingdom / Southport, Merseyside | James Seekings (Animal and Plant Health Agency (APHA) / Virology Department) |
| EPI_ISL_253946 | A/chicken/Japan/AQ-HE144/2015 | A / H5N6 | Asia / Japan | |
| EPI_ISL_253949 | A/duck/Japan/AQ-HE72/2015 | A / H5N6 | Asia / Japan | |
| EPI_ISL_254725 | A/chicken/Korea/H903/2017 | A / H5N8 | Asia / Korea, Republic of / Jeollabuk-do | Youn-Jeong Lee (Animal and Plant Quarantine Agency (APQA) / Avian Influenza Research and Diagsnotic Division) |
| EPI_ISL_254745 | A/mallard duck/Korea/WA137/2017 | A / H5N8 | Asia / Korea, Republic of / Chungcheongnam-do | Youn-Jeong Lee (Animal and Plant Quarantine Agency (APQA) / Avian Influenza Research and Diagsnotic Division) |
| EPI_ISL_254814 | A/mallard/Hungary/57857/2016 | A / H5N8 | Europe / Hungary / Hajdu-Bihar | Adam Dan (Danam.Vet.Molbiol) |
| EPI_ISL_255012 | A/American wigeon/California/UCD58P/2015 | A / H5N8 | North America / United States / California / Solano County / Grizzly Island Wildlife Refuge | Andrew Ramey (U.S. Geological Survey Alaska Science Center) |
| EPI_ISL_255174 | A/Goose/Hungary/65817/2016 | A / H5N8 | Europe / Hungary / Hajdu-Bihar | Adam Dan (Danam.Vet.Molbiol) |
| EPI_ISL_255181 | A/swan/Italy/17VIR537-2/2017 | A / H5N8 | Europe / Italy / Udine | Bianca Zecchin (Istituto Zooprofilattico Sperimentale Delle Venezie) |
| EPI_ISL_255182 | A/turkey/Italy/17VIR538-1/2017 | A / H5N8 | Europe / Italy / Venezia | Bianca Zecchin (Istituto Zooprofilattico Sperimentale Delle Venezie) |
| EPI_ISL_255183 | A/turkey/Italy/17VIR576-11/2017 | A / H5N8 | Europe / Italy / Padova | Bianca Zecchin (Istituto Zooprofilattico Sperimentale Delle Venezie) |
| EPI_ISL_255184 | A/chicken/Italy/17VIR653-12/2017 | A / H5N8 | Europe / Italy / Rovigo | Bianca Zecchin (Istituto Zooprofilattico Sperimentale Delle Venezie) |
| EPI_ISL_255185 | A/turkey/Italy/17VIR973-2/2017 | A / H5N8 | Europe / Italy / Parma | Bianca Zecchin (Istituto Zooprofilattico Sperimentale Delle Venezie) |
| EPI_ISL_255186 | A/turkey/Italy/17VIR1338-3/2017 | A / H5N8 | Europe / Italy / Mantova | Bianca Zecchin (Istituto Zooprofilattico Sperimentale Delle Venezie) |
| EPI_ISL_255187 | A/turkey/Italy/17VIR1452-22/2017 | A / H5N8 | Europe / Italy / Verona | Bianca Zecchin (Istituto Zooprofilattico Sperimentale Delle Venezie) |

| | | | |
|---|---|---|---|
| EPI_ISL_255189 | A/gadwall/Italy/17VIR133-2/2017 | A / H5N5 Europe / Italy / Gorizia | Bianca Zecchin (Istituto Zooprofilattico Sperimentale Delle Venezie) |
| EPI_ISL_255190 | A/Mute swan/Hungary/119/2017 | A / H5N8 Europe / Hungary / Somogy megye / Somogy | Adam Dan (Danam.Vet.Molbiol) |
| EPI_ISL_255191 | A/Harris Hawk/Hungary/120/2017 | A / H5N8 Europe / Hungary / Budapest fovaros / Budapest | Adam Dan (Danam.Vet.Molbiol) |
| EPI_ISL_255192 | A/Goose/Hungary/982/2017 | A / H5N8 Europe / Hungary / Jasz-Nagykun-Szolnok | Adam Dan (Danam.Vet.Molbiol) |
| EPI_ISL_255193 | A/Duck/Hungary/984/2017 | A / H5N8 Europe / Hungary / Bekes megye | Adam Dan (Danam.Vet.Molbiol) |
| EPI_ISL_255194 | A/Goose/Hungary/1030/2017 | A / H5N8 Europe / Hungary / Csongrad megye | Adam Dan (Danam.Vet.Molbiol) |
| EPI_ISL_255195 | A/Duck/Hungary/1588/2017 | A / H5N8 Europe / Hungary / Bacs-Kiskun | Adam Dan (Danam.Vet.Molbiol) |
| EPI_ISL_255196 | A/Greylag_goose/Hungary/1941/2017 | A / H5N8 Europe / Hungary / Fejer megye | Adam Dan (Danam.Vet.Molbiol) |
| EPI_ISL_255197 | A/Mute swan/Hungary/1955/2017 | A / H5N8 Europe / Hungary / Veszprem megye | Adam Dan (Danam.Vet.Molbiol) |
| EPI_ISL_255198 | A/Turkey/Hungary/2030/2017 | A / H5N8 Europe / Hungary / Veszprem megye | Adam Dan (Danam.Vet.Molbiol) |
| EPI_ISL_255199 | A/Mute swan/Hungary/2193/2017 | A / H5N8 Europe / Hungary / Borsod-Abauj-Zemplen | Adam Dan (Danam.Vet.Molbiol) |
| EPI_ISL_255200 | A/Mute swan/Hungary/2508/2017 | A / H5N8 Europe / Hungary / Somogy megye / Somogy | Adam Dan (Danam.Vet.Molbiol) |
| EPI_ISL_255201 | A/Mute swan/Hungary/2825/2017 | A / H5N8 Europe / Hungary / Tolna megye / Tolna | Adam Dan (Danam.Vet.Molbiol) |
| EPI_ISL_255202 | A/Mute swan/Hungary/3139/2017 | A / H5N8 Europe / Hungary / Zala | Adam Dan (Danam.Vet.Molbiol) |
| EPI_ISL_255203 | A/Mute swan/Hungary/3513/2017 | A / H5N8 Europe / Hungary / Vas | Adam Dan (Danam.Vet.Molbiol) |
| EPI_ISL_255204 | A/Mute swan/Hungary/3542/2017 | A / H5N8 Europe / Hungary / Budapest fovaros | Adam Dan (Danam.Vet.Molbiol) |
| EPI_ISL_255205 | A/Chicken/Hungary/2496/2017 | A / H5N8 Europe / Hungary / Jasz-Nagykun-Szolnok | Adam Dan (Danam.Vet.Molbiol) |
| EPI_ISL_255206 | A/Peregrine_falcon/Hungary/4882/2017 | A / H5N8 Europe / Hungary / Gyor-Moson-Sopron | Adam Dan (Danam.Vet.Molbiol) |
| EPI_ISL_255207 | A/Mute swan/Hungary/5316/2017 | A / H5N8 Europe / Hungary / Tolna megye / Tolna | Adam Dan (Danam.Vet.Molbiol) |
| EPI_ISL_255208 | A/Common_buzzard/Hungary/7061/2017 | A / H5N8 Europe / Hungary / Tolna megye / Tolna | Adam Dan (Danam.Vet.Molbiol) |
| EPI_ISL_255209 | A/Common_tern/Hungary/8187/2017 | A / H5N8 Europe / Hungary / Tolna megye | Adam Dan (Danam.Vet.Molbiol) |
| EPI_ISL_255210 | A/Harris_hawk/Hungary/2762a/2017 | A / H5N8 Europe / Hungary / Pest megye / Pest | Adam Dan (Danam.Vet.Molbiol) |
| EPI_ISL_255211 | A/Harris_hawk/Hungary/2762b/2017 | A / H5N8 Europe / Hungary / Pest megye / Pest | Adam Dan (Danam.Vet.Molbiol) |
| EPI_ISL_255212 | A/Greylag_goose/Hungary/320/2017 | A / H5N8 Europe / Hungary / Somogy megye / Somogy | Adam Dan (Danam.Vet.Molbiol) |
| EPI_ISL_255213 | A/GuineaFowl/Hungary/596/2017 | A / H5N8 Europe / Hungary / Hajdu-Bihar | Adam Dan (Danam.Vet.Molbiol) |
| EPI_ISL_255214 | A/Chicken/Hungary/1751/2017 | A / H5N8 Europe / Hungary / Gyor-Moson-Sopron | Adam Dan (Danam.Vet.Molbiol) |
| EPI_ISL_255215 | A/Mute swan/Hungary/3137/2017 | A / H5N8 Europe / Hungary / Zala | Adam Dan (Danam.Vet.Molbiol) |
| EPI_ISL_255216 | A/Mallard/Hungary/5821/2017 | A / H5N8 Europe / Hungary / Zala | Adam Dan (Danam.Vet.Molbiol) |
| EPI_ISL_255218 | A/Mallard/Hungary/1574b/2017 | A / H5N8 Europe / Hungary / Somogy megye / Somogy | Adam Dan (Danam.Vet.Molbiol) |
| EPI_ISL_255219 | A/White_fronted_goose/Hungary/801/2017 | A / H5N8 Europe / Hungary / Hajdu-Bihar | Adam Dan (Danam.Vet.Molbiol) |
| EPI_ISL_255220 | A/Mallard/Hungary/1574a/2017 | A / H5N8 Europe / Hungary / Somogy megye / Somogy | Adam Dan (Danam.Vet.Molbiol) |
| EPI_ISL_255388 | A/goose/Hunan/161/2014 | A / H5N6 Asia / China | Jie Cui (Wuhan Institute of Virology, CAS. ) |
| EPI_ISL_255389 | A/chicken/Anhui/AH325/2015 | A / H5N6 Asia / China | Jie Cui (Wuhan Institute of Virology, CAS. ) |
| EPI_ISL_255398 | A/Goose/Hungary/59712/2016 | A / H5N8 Europe / Hungary / Csongrad megye / Csongrad | Adam Dan (Danam.Vet.Molbiol) |
| EPI_ISL_255412 | A/chicken/Hunan/HN158/2015 | A / H5N6 Asia / China | Jie Cui (Wuhan Institute of Virology, CAS. ) |
| EPI_ISL_255413 | A/chicken/Hunan/HN244/2015 | A / H5N6 Asia / China | Jie Cui (Wuhan Institute of Virology, CAS. ) |
| EPI_ISL_255414 | A/chicken/Hunan/HN245/2015 | A / H5N6 Asia / China | Jie Cui (Wuhan Institute of Virology, CAS. ) |
| EPI_ISL_255415 | A/chicken/Hunan/HN247/2015 | A / H5N6 Asia / China | Jie Cui (Wuhan Institute of Virology, CAS. ) |
| EPI_ISL_255416 | A/chicken/Hunan/HN248/2015 | A / H5N6 Asia / China | Jie Cui (Wuhan Institute of Virology, CAS. ) |
| EPI_ISL_255417 | A/chicken/Hunan/HN249/2015 | A / H5N6 Asia / China | Jie Cui (Wuhan Institute of Virology, CAS. ) |
| EPI_ISL_255418 | A/chicken/Hunan/HN250/2015 | A / H5N6 Asia / China | Jie Cui (Wuhan Institute of Virology, CAS. ) |
| EPI_ISL_255419 | A/chicken/Hunan/HN251/2015 | A / H5N6 Asia / China | Jie Cui (Wuhan Institute of Virology, CAS. ) |
| EPI_ISL_255420 | A/chicken/Hunan/HN252/2015 | A / H5N6 Asia / China | Jie Cui (Wuhan Institute of Virology, CAS. ) |
| EPI_ISL_255421 | A/chicken/Hunan/HN254/2015 | A / H5N6 Asia / China | Jie Cui (Wuhan Institute of Virology, CAS. ) |
| EPI_ISL_255422 | A/chicken/Hunan/HN263/2015 | A / H5N6 Asia / China | Jie Cui (Wuhan Institute of Virology, CAS. ) |
| EPI_ISL_255423 | A/chicken/Hunan/HN255/2015 | A / H5N6 Asia / China | Jie Cui (Wuhan Institute of Virology, CAS. ) |
| EPI_ISL_255424 | A/chicken/Hunan/HN264/2015 | A / H5N6 Asia / China | Jie Cui (Wuhan Institute of Virology, CAS. ) |
| EPI_ISL_255425 | A/chicken/Hunan/HN265/2015 | A / H5N6 Asia / China | Jie Cui (Wuhan Institute of Virology, CAS. ) |
| EPI_ISL_255426 | A/chicken/Hunan/HN266/2015 | A / H5N6 Asia / China | Jie Cui (Wuhan Institute of Virology, CAS. ) |
| EPI_ISL_255427 | A/chicken/Hunan/HN301/2015 | A / H5N6 Asia / China | Jie Cui (Wuhan Institute of Virology, CAS. ) |
| EPI_ISL_255428 | A/chicken/Hunan/HN302/2015 | A / H5N6 Asia / China | Jie Cui (Wuhan Institute of Virology, CAS. ) |
| EPI_ISL_255429 | A/chicken/Hunan/HN303/2015 | A / H5N6 Asia / China | Jie Cui (Wuhan Institute of Virology, CAS. ) |
| EPI_ISL_255430 | A/chicken/Hunan/HN304/2015 | A / H5N6 Asia / China | Jie Cui (Wuhan Institute of Virology, CAS. ) |
| EPI_ISL_255433 | A/chicken/Hunan/HN305/2015 | A / H5N6 Asia / China | Jie Cui (Wuhan Institute of Virology, CAS. ) |
| EPI_ISL_255466 | A/chicken/Hunan/HN306/2015 | A / H5N6 Asia / China | Jie Cui (Wuhan Institute of Virology, CAS. ) |
| EPI_ISL_255467 | A/chicken/Hunan/HN307/2015 | A / H5N6 Asia / China | Jie Cui (Wuhan Institute of Virology, CAS. ) |
| EPI_ISL_255468 | A/chicken/Hunan/HN308/2015 | A / H5N6 Asia / China | Jie Cui (Wuhan Institute of Virology, CAS. ) |
| EPI_ISL_255469 | A/chicken/Hunan/HN40/2015 | A / H5N6 Asia / China | Jie Cui (Wuhan Institute of Virology, CAS. ) |
| EPI_ISL_255470 | A/chicken/Hunan/HN58/2015 | A / H5N6 Asia / China | Jie Cui (Wuhan Institute of Virology, CAS. ) |
| EPI_ISL_255489 | A/chicken/Hunan/HN75/2015 | A / H5N6 Asia / China | Jie Cui (Wuhan Institute of Virology, CAS. ) |
| EPI_ISL_255491 | A/duck/Anhui/AH328/2015 | A / H5N6 Asia / China | Jie Cui (Wuhan Institute of Virology, CAS. ) |
| EPI_ISL_255492 | A/duck/Hunan/193/2014 | A / H5N6 Asia / China | Jie Cui (Wuhan Institute of Virology, CAS. ) |
| EPI_ISL_255493 | A/duck/Hunan/233/2014 | A / H5N6 Asia / China | Jie Cui (Wuhan Institute of Virology, CAS. ) |
| EPI_ISL_255509 | A/duck/Hunan/HN05/2014 | A / H5N6 Asia / China | Jie Cui (Wuhan Institute of Virology, CAS. ) |
| EPI_ISL_255529 | A/duck/Hunan/HN13/2015 | A / H5N6 Asia / China | Jie Cui (Wuhan Institute of Virology, CAS. ) |
| EPI_ISL_255530 | A/duck/Hunan/HN16/2015 | A / H5N6 Asia / China | Jie Cui (Wuhan Institute of Virology, CAS. ) |
| EPI_ISL_255534 | A/duck/Hunan/HN162/2015 | A / H5N6 Asia / China | Jie Cui (Wuhan Institute of Virology, CAS. ) |
| EPI_ISL_2555532 | A/Cygnus columbianus/Hubei/116/2020 | A / H5N8 Asia / China / Jiangxi Province / Longgan Lake National Nature Reserve | Ma Liping (Wuhan Institute of Virology, CAS ) |
| EPI_ISL_2555534 | A/Cygnus columbianus/Hubei/56/2020 | A / H5N8 Asia / China / Jiangxi Province | Ma Liping (Wuhan Institute of Virology, CAS ) |
| EPI_ISL_2555536 | A/Chlidonias hybrida/Hubei/55/2020 | A / H5N8 Asia / China / Jiangxi Province | Ma Liping (Wuhan Institute of Virology, CAS ) |
| EPI_ISL_2555538 | A/Cygnus columbianus/Hubei/53/2020 | A / H5N8 Asia / China / Jiangxi Province | Ma Liping (Wuhan Institute of Virology, CAS ) |
| EPI_ISL_2555541 | A/Cygnus columbianus/Hubei/52/2020 | A / H5N8 Asia / China / Jiangxi Province | Ma Liping (Wuhan Institute of Virology, CAS ) |

| | | | | |
|---|---|---|---|---|
| EPI_ISL_2555542 | A/Cygnus columbianus/Hubei/51/2020 | A / H5N8 | Asia / China / Jiangxi Province | Ma Liping (Wuhan Institute of Virology, CAS ) |
| EPI_ISL_2555543 | A/Cygnus columbianus/Hubei/50/2020 | A / H5N8 | Asia / China / Jiangxi Province | Ma Liping (Wuhan Institute of Virology, CAS ) |
| EPI_ISL_255559 | A/duck/Hunan/HN163/2015 | A / H5N6 | Asia / China | Jie Cui (Wuhan Institute of Virology, CAS. ) |
| EPI_ISL_255561 | A/duck/Hunan/HN167/2015 | A / H5N6 | Asia / China | Jie Cui (Wuhan Institute of Virology, CAS. ) |
| EPI_ISL_255562 | A/duck/Hunan/HN171/2015 | A / H5N6 | Asia / China | Jie Cui (Wuhan Institute of Virology, CAS. ) |
| EPI_ISL_255563 | A/duck/Hunan/HN172/2015 | A / H5N6 | Asia / China | Jie Cui (Wuhan Institute of Virology, CAS. ) |
| EPI_ISL_255564 | A/duck/Hunan/HN174/2015 | A / H5N6 | Asia / China | Jie Cui (Wuhan Institute of Virology, CAS. ) |
| EPI_ISL_255587 | A/duck/Hunan/HN176/2015 | A / H5N6 | Asia / China | Jie Cui (Wuhan Institute of Virology, CAS. ) |
| EPI_ISL_255592 | A/duck/Hunan/HN177/2015 | A / H5N6 | Asia / China | Jie Cui (Wuhan Institute of Virology, CAS. ) |
| EPI_ISL_255597 | A/duck/Hunan/HN178/2015 | A / H5N6 | Asia / China | Jie Cui (Wuhan Institute of Virology, CAS. ) |
| EPI_ISL_255599 | A/duck/Hunan/HN182/2015 | A / H5N6 | Asia / China | Jie Cui (Wuhan Institute of Virology, CAS. ) |
| EPI_ISL_255600 | A/duck/Hunan/HN183/2015 | A / H5N6 | Asia / China | Jie Cui (Wuhan Institute of Virology, CAS. ) |
| EPI_ISL_255603 | A/duck/Hunan/HN186/2015 | A / H5N6 | Asia / China | Jie Cui (Wuhan Institute of Virology, CAS. ) |
| EPI_ISL_255605 | A/duck/Hunan/HN190/2015 | A / H5N6 | Asia / China | Jie Cui (Wuhan Institute of Virology, CAS. ) |
| EPI_ISL_255607 | A/duck/Hunan/HN191/2015 | A / H5N6 | Asia / China | Jie Cui (Wuhan Institute of Virology, CAS. ) |
| EPI_ISL_255609 | A/duck/Hunan/HN192/2015 | A / H5N6 | Asia / China | Jie Cui (Wuhan Institute of Virology, CAS. ) |
| EPI_ISL_255610 | A/duck/Hunan/HN194/2015 | A / H5N6 | Asia / China | Jie Cui (Wuhan Institute of Virology, CAS. ) |
| EPI_ISL_2556113 | A/Cygnus columbianus/Hubei/49/2020 | A / H5N8 | Asia / China / Jiangxi Province | Ma Liping (Wuhan Institute of Virology, CAS ) |
| EPI_ISL_255612 | A/duck/Hunan/HN195/2015 | A / H5N6 | Asia / China | Jie Cui (Wuhan Institute of Virology, CAS. ) |
| EPI_ISL_255613 | A/duck/Hunan/HN197/2015 | A / H5N6 | Asia / China | Jie Cui (Wuhan Institute of Virology, CAS. ) |
| EPI_ISL_255614 | A/duck/Hunan/HN201/2015 | A / H5N6 | Asia / China | Jie Cui (Wuhan Institute of Virology, CAS. ) |
| EPI_ISL_255615 | A/duck/Hunan/HN204/2015 | A / H5N6 | Asia / China | Jie Cui (Wuhan Institute of Virology, CAS. ) |
| EPI_ISL_255616 | A/duck/Hunan/HN205/2015 | A / H5N6 | Asia / China | Jie Cui (Wuhan Institute of Virology, CAS. ) |
| EPI_ISL_255617 | A/duck/Hunan/HN206/2015 | A / H5N6 | Asia / China | Jie Cui (Wuhan Institute of Virology, CAS. ) |
| EPI_ISL_255618 | A/duck/Hunan/HN207/2015 | A / H5N6 | Asia / China | Jie Cui (Wuhan Institute of Virology, CAS. ) |
| EPI_ISL_255619 | A/duck/Hunan/HN208/2015 | A / H5N6 | Asia / China | Jie Cui (Wuhan Institute of Virology, CAS. ) |
| EPI_ISL_255620 | A/duck/Hunan/HN209/2015 | A / H5N6 | Asia / China | Jie Cui (Wuhan Institute of Virology, CAS. ) |
| EPI_ISL_255621 | A/duck/Hunan/HN210/2015 | A / H5N6 | Asia / China | Jie Cui (Wuhan Institute of Virology, CAS. ) |
| EPI_ISL_255624 | A/duck/Hunan/HN212/2015 | A / H5N6 | Asia / China | Jie Cui (Wuhan Institute of Virology, CAS. ) |
| EPI_ISL_255627 | A/duck/France/161587/2016 | A / H5N8 | Europe / France / Aquitaine / Departement des Landes | Francois-Xavier Briand (ANSES Agence Nationale De Securite Sanitaire De L'alimentation / Laboratoire de Ploufragan-Plouzané) |
| EPI_ISL_255689 | A/duck/Hunan/HN214/2015 | A / H5N6 | Asia / China | Jie Cui (Wuhan Institute of Virology, CAS. ) |
| EPI_ISL_255691 | A/duck/Hunan/HN215/2015 | A / H5N6 | Asia / China | Jie Cui (Wuhan Institute of Virology, CAS. ) |
| EPI_ISL_255692 | A/duck/Hunan/HN216/2015 | A / H5N6 | Asia / China | Jie Cui (Wuhan Institute of Virology, CAS. ) |
| EPI_ISL_255694 | A/duck/Hunan/HN217/2015 | A / H5N6 | Asia / China | Jie Cui (Wuhan Institute of Virology, CAS. ) |
| EPI_ISL_255695 | A/duck/Hunan/HN220/2015 | A / H5N6 | Asia / China | Jie Cui (Wuhan Institute of Virology, CAS. ) |
| EPI_ISL_255698 | A/duck/Hunan/HN221/2015 | A / H5N6 | Asia / China | Jie Cui (Wuhan Institute of Virology, CAS. ) |
| EPI_ISL_255699 | A/duck/Hunan/HN224/2015 | A / H5N6 | Asia / China | Jie Cui (Wuhan Institute of Virology, CAS. ) |
| EPI_ISL_255701 | A/duck/Hunan/HN227/2015 | A / H5N6 | Asia / China | Jie Cui (Wuhan Institute of Virology, CAS. ) |
| EPI_ISL_255702 | A/duck/Hunan/HN230/2015 | A / H5N6 | Asia / China | Jie Cui (Wuhan Institute of Virology, CAS. ) |
| EPI_ISL_255738 | A/duck/Hunan/HN236/2015 | A / H5N6 | Asia / China | Jie Cui (Wuhan Institute of Virology, CAS. ) |
| EPI_ISL_255741 | A/duck/Hunan/HN238/2015 | A / H5N6 | Asia / China | Jie Cui (Wuhan Institute of Virology, CAS. ) |
| EPI_ISL_255742 | A/duck/Hunan/HN239/2015 | A / H5N6 | Asia / China | Jie Cui (Wuhan Institute of Virology, CAS. ) |
| EPI_ISL_255743 | A/duck/Hunan/HN240/2015 | A / H5N6 | Asia / China | Jie Cui (Wuhan Institute of Virology, CAS. ) |
| EPI_ISL_255744 | A/duck/Hunan/HN241/2015 | A / H5N6 | Asia / China | Jie Cui (Wuhan Institute of Virology, CAS. ) |
| EPI_ISL_255746 | A/duck/Hunan/HN242/2015 | A / H5N6 | Asia / China | Jie Cui (Wuhan Institute of Virology, CAS. ) |
| EPI_ISL_255747 | A/duck/Hunan/HN271/2015 | A / H5N6 | Asia / China | Jie Cui (Wuhan Institute of Virology, CAS. ) |
| EPI_ISL_255749 | A/duck/Hunan/HN277/2015 | A / H5N6 | Asia / China | Jie Cui (Wuhan Institute of Virology, CAS. ) |
| EPI_ISL_255751 | A/duck/Hunan/HN278/2015 | A / H5N6 | Asia / China | Jie Cui (Wuhan Institute of Virology, CAS. ) |
| EPI_ISL_255752 | A/duck/Hunan/HN285/2015 | A / H5N6 | Asia / China | Jie Cui (Wuhan Institute of Virology, CAS. ) |
| EPI_ISL_255754 | A/duck/Hunan/HN288/2015 | A / H5N6 | Asia / China | Jie Cui (Wuhan Institute of Virology, CAS. ) |
| EPI_ISL_255755 | A/duck/Hunan/HN29/2015 | A / H5N6 | Asia / China | Jie Cui (Wuhan Institute of Virology, CAS. ) |
| EPI_ISL_255756 | A/duck/Hunan/HN333/2015 | A / H5N6 | Asia / China | Jie Cui (Wuhan Institute of Virology, CAS. ) |
| EPI_ISL_255757 | A/duck/Hunan/HN334/2015 | A / H5N6 | Asia / China | Jie Cui (Wuhan Institute of Virology, CAS. ) |
| EPI_ISL_255758 | A/duck/Hunan/HN335/2015 | A / H5N6 | Asia / China | Jie Cui (Wuhan Institute of Virology, CAS. ) |
| EPI_ISL_255759 | A/duck/Hunan/HN336/2015 | A / H5N6 | Asia / China | Jie Cui (Wuhan Institute of Virology, CAS. ) |
| EPI_ISL_255760 | A/duck/Hunan/HN339/2015 | A / H5N6 | Asia / China | Jie Cui (Wuhan Institute of Virology, CAS. ) |
| EPI_ISL_255761 | A/duck/Hunan/HN341/2015 | A / H5N6 | Asia / China | Jie Cui (Wuhan Institute of Virology, CAS. ) |
| EPI_ISL_255762 | A/duck/Hunan/HN342/2015 | A / H5N6 | Asia / China | Jie Cui (Wuhan Institute of Virology, CAS. ) |
| EPI_ISL_255763 | A/duck/Hunan/HN343/2015 | A / H5N6 | Asia / China | Jie Cui (Wuhan Institute of Virology, CAS. ) |
| EPI_ISL_255764 | A/duck/Hunan/HN42/2015 | A / H5N6 | Asia / China | Jie Cui (Wuhan Institute of Virology, CAS. ) |
| EPI_ISL_255765 | A/duck/Hunan/HN45/2015 | A / H5N6 | Asia / China | Jie Cui (Wuhan Institute of Virology, CAS. ) |
| EPI_ISL_255766 | A/duck/Hunan/HN53/2015 | A / H5N6 | Asia / China | Jie Cui (Wuhan Institute of Virology, CAS. ) |
| EPI_ISL_255768 | A/duck/Hunan/HN54/2015 | A / H5N6 | Asia / China | Jie Cui (Wuhan Institute of Virology, CAS. ) |
| EPI_ISL_255769 | A/duck/Hunan/HN59/2015 | A / H5N6 | Asia / China | Jie Cui (Wuhan Institute of Virology, CAS. ) |
| EPI_ISL_255770 | A/duck/Hunan/HN60/2015 | A / H5N6 | Asia / China | Jie Cui (Wuhan Institute of Virology, CAS. ) |
| EPI_ISL_255771 | A/duck/Hunan/HN61/2015 | A / H5N6 | Asia / China | Jie Cui (Wuhan Institute of Virology, CAS. ) |
| EPI_ISL_255772 | A/duck/Hunan/HN64/2015 | A / H5N6 | Asia / China | Jie Cui (Wuhan Institute of Virology, CAS. ) |
| EPI_ISL_255773 | A/duck/Hunan/HN66/2015 | A / H5N6 | Asia / China | Jie Cui (Wuhan Institute of Virology, CAS. ) |
| EPI_ISL_255774 | A/duck/Hunan/HN67/2015 | A / H5N6 | Asia / China | Jie Cui (Wuhan Institute of Virology, CAS. ) |
| EPI_ISL_255775 | A/duck/Hunan/HN71/2015 | A / H5N6 | Asia / China | Jie Cui (Wuhan Institute of Virology, CAS. ) |
| EPI_ISL_255816 | A/duck/Hunan/HN72/2015 | A / H5N6 | Asia / China | Jie Cui (Wuhan Institute of Virology, CAS. ) |
| EPI_ISL_255817 | A/duck/Hunan/HN74/2015 | A / H5N6 | Asia / China | Jie Cui (Wuhan Institute of Virology, CAS. ) |

| | | | | |
|---|---|---|---|---|
| EPI_ISL_255818 | A/duck/Hunan/HN76/2015 | A / H5N6 | Asia / China | Jie Cui (Wuhan Institute of Virology, CAS. ) |
| EPI_ISL_255819 | A/duck/Hunan/HN78/2015 | A / H5N6 | Asia / China | Jie Cui (Wuhan Institute of Virology, CAS. ) |
| EPI_ISL_255820 | A/duck/Hunan/HN80/2015 | A / H5N6 | Asia / China | Jie Cui (Wuhan Institute of Virology, CAS. ) |
| EPI_ISL_255821 | A/environment/Hunan/HN261/2015 | A / H5N6 | Asia / China | Jie Cui (Wuhan Institute of Virology, CAS. ) |
| EPI_ISL_255822 | A/goose/Anhui/AH350/2015 | A / H5N6 | Asia / China | Jie Cui (Wuhan Institute of Virology, CAS. ) |
| EPI_ISL_255823 | A/goose/Hubei/113/2014 | A / H5N6 | Asia / China | Jie Cui (Wuhan Institute of Virology, CAS. ) |
| EPI_ISL_255824 | A/goose/HuBei/117/2014 | A / H5N6 | Asia / China | Jie Cui (Wuhan Institute of Virology, CAS. ) |
| EPI_ISL_255825 | A/goose/Hunan/107/2014 | A / H5N6 | Asia / China | Jie Cui (Wuhan Institute of Virology, CAS. ) |
| EPI_ISL_255826 | A/goose/Hunan/109/2014 | A / H5N6 | Asia / China | Jie Cui (Wuhan Institute of Virology, CAS. ) |
| EPI_ISL_255827 | A/goose/Hunan/110/2014 | A / H5N6 | Asia / China | Jie Cui (Wuhan Institute of Virology, CAS. ) |
| EPI_ISL_255828 | A/goose/Hunan/111/2014 | A / H5N6 | Asia / China | Jie Cui (Wuhan Institute of Virology, CAS. ) |
| EPI_ISL_255829 | A/goose/Hunan/115/2014 | A / H5N6 | Asia / China | Jie Cui (Wuhan Institute of Virology, CAS. ) |
| EPI_ISL_255830 | A/goose/Hunan/116/2014 | A / H5N6 | Asia / China | Jie Cui (Wuhan Institute of Virology, CAS. ) |
| EPI_ISL_255831 | A/goose/Hunan/155/2014 | A / H5N6 | Asia / China | Jie Cui (Wuhan Institute of Virology, CAS. ) |
| EPI_ISL_255832 | A/goose/Hunan/159/2014 | A / H5N6 | Asia / China | Jie Cui (Wuhan Institute of Virology, CAS. ) |
| EPI_ISL_255833 | A/goose/Hunan/160/2014 | A / H5N6 | Asia / China | Jie Cui (Wuhan Institute of Virology, CAS. ) |
| EPI_ISL_255834 | A/goose/Hunan/81/2014 | A / H5N6 | Asia / China | Jie Cui (Wuhan Institute of Virology, CAS. ) |
| EPI_ISL_255835 | A/goose/Hunan/82/2014 | A / H5N6 | Asia / China | Jie Cui (Wuhan Institute of Virology, CAS. ) |
| EPI_ISL_255836 | A/goose/Hunan/HN11/2015 | A / H5N6 | Asia / China | Jie Cui (Wuhan Institute of Virology, CAS. ) |
| EPI_ISL_255837 | A/goose/Hunan/HN22/2015 | A / H5N6 | Asia / China | Jie Cui (Wuhan Institute of Virology, CAS. ) |
| EPI_ISL_255838 | A/goose/Hunan/HN310/2015 | A / H5N6 | Asia / China | Jie Cui (Wuhan Institute of Virology, CAS. ) |
| EPI_ISL_255839 | A/goose/Hunan/HN311/2015 | A / H5N6 | Asia / China | Jie Cui (Wuhan Institute of Virology, CAS. ) |
| EPI_ISL_255840 | A/goose/Hunan/HN314/2015 | A / H5N6 | Asia / China | Jie Cui (Wuhan Institute of Virology, CAS. ) |
| EPI_ISL_255841 | A/goose/Hunan/HN315/2015 | A / H5N6 | Asia / China | Jie Cui (Wuhan Institute of Virology, CAS. ) |
| EPI_ISL_255842 | A/goose/Hunan/HN316/2015 | A / H5N6 | Asia / China | Jie Cui (Wuhan Institute of Virology, CAS. ) |
| EPI_ISL_255843 | A/goose/Hunan/HN317/2015 | A / H5N6 | Asia / China | Jie Cui (Wuhan Institute of Virology, CAS. ) |
| EPI_ISL_255844 | A/goose/Hunan/HN318/2015 | A / H5N6 | Asia / China | Jie Cui (Wuhan Institute of Virology, CAS. ) |
| EPI_ISL_255845 | A/goose/Hunan/HN319/2015 | A / H5N6 | Asia / China | Jie Cui (Wuhan Institute of Virology, CAS. ) |
| EPI_ISL_255846 | A/goose/Hunan/HN325/2015 | A / H5N6 | Asia / China | Jie Cui (Wuhan Institute of Virology, CAS. ) |
| EPI_ISL_255847 | A/goose/Hunan/HN57/2015 | A / H5N6 | Asia / China | Jie Cui (Wuhan Institute of Virology, CAS. ) |
| EPI_ISL_255848 | A/goose/Hunan/HN88/2015 | A / H5N6 | Asia / China | Jie Cui (Wuhan Institute of Virology, CAS. ) |
| EPI_ISL_255849 | A/goose/Hunan/HN90/2015 | A / H5N6 | Asia / China | Jie Cui (Wuhan Institute of Virology, CAS. ) |
| EPI_ISL_255850 | A/goose/SiChuan/15/2015 | A / H5N6 | Asia / China | Jie Cui (Wuhan Institute of Virology, CAS. ) |
| EPI_ISL_255851 | A/muscovy/duck/Hunan/101/2014 | A / H5N6 | Asia / China | Jie Cui (Wuhan Institute of Virology, CAS. ) |
| EPI_ISL_255852 | A/muscovy/duck/Hunan/102/2014 | A / H5N6 | Asia / China | Jie Cui (Wuhan Institute of Virology, CAS. ) |
| EPI_ISL_255853 | A/muscovy/duck/Hunan/103/2014 | A / H5N6 | Asia / China | Jie Cui (Wuhan Institute of Virology, CAS. ) |
| EPI_ISL_255854 | A/muscovy/duck/Hunan/104/2014 | A / H5N6 | Asia / China | Jie Cui (Wuhan Institute of Virology, CAS. ) |
| EPI_ISL_255857 | A/muscovy/duck/Hunan/125/2014 | A / H5N6 | Asia / China | Jie Cui (Wuhan Institute of Virology, CAS. ) |
| EPI_ISL_255858 | A/muscovy/duck/Hunan/147/2014 | A / H5N6 | Asia / China | Jie Cui (Wuhan Institute of Virology, CAS. ) |
| EPI_ISL_255859 | A/muscovy/duck/Hunan/148/2014 | A / H5N6 | Asia / China | Jie Cui (Wuhan Institute of Virology, CAS. ) |
| EPI_ISL_255860 | A/muscovy/duck/Hunan/167/2014 | A / H5N6 | Asia / China | Jie Cui (Wuhan Institute of Virology, CAS. ) |
| EPI_ISL_255861 | A/muscovy/duck/Hunan/183/2014 | A / H5N6 | Asia / China | Jie Cui (Wuhan Institute of Virology, CAS. ) |
| EPI_ISL_255862 | A/muscovy/duck/Hunan/202/2014 | A / H5N6 | Asia / China | Jie Cui (Wuhan Institute of Virology, CAS. ) |
| EPI_ISL_255863 | A/muscovy/duck/Hunan/216/2014 | A / H5N6 | Asia / China | Jie Cui (Wuhan Institute of Virology, CAS. ) |
| EPI_ISL_255864 | A/muscovy/duck/Hunan/229/2014 | A / H5N6 | Asia / China | Jie Cui (Wuhan Institute of Virology, CAS. ) |
| EPI_ISL_255865 | A/muscovy/duck/Hunan/232/2014 | A / H5N6 | Asia / China | Jie Cui (Wuhan Institute of Virology, CAS. ) |
| EPI_ISL_255866 | A/muscovy/duck/Hunan/85/2014 | A / H5N6 | Asia / China | Jie Cui (Wuhan Institute of Virology, CAS. ) |
| EPI_ISL_255867 | A/muscovy/duck/Hunan/88/2014 | A / H5N6 | Asia / China | Jie Cui (Wuhan Institute of Virology, CAS. ) |
| EPI_ISL_255868 | A/muscovy/duck/Hunan/89/2014 | A / H5N6 | Asia / China | Jie Cui (Wuhan Institute of Virology, CAS. ) |
| EPI_ISL_255869 | A/muscovy/duck/Hunan/90/2014 | A / H5N6 | Asia / China | Jie Cui (Wuhan Institute of Virology, CAS. ) |
| EPI_ISL_255870 | A/muscovy/duck/Hunan/93/2014 | A / H5N6 | Asia / China | Jie Cui (Wuhan Institute of Virology, CAS. ) |
| EPI_ISL_255871 | A/muscovy/duck/Hunan/94/2014 | A / H5N6 | Asia / China | Jie Cui (Wuhan Institute of Virology, CAS. ) |
| EPI_ISL_255872 | A/muscovy/duck/Hunan/HN346/2015 | A / H5N6 | Asia / China | Jie Cui (Wuhan Institute of Virology, CAS. ) |
| EPI_ISL_255873 | A/muscovy/duck/Hunan/HN347/2015 | A / H5N6 | Asia / China | Jie Cui (Wuhan Institute of Virology, CAS. ) |
| EPI_ISL_255874 | A/muscovy/duck/Hunan/HN348/2015 | A / H5N6 | Asia / China | Jie Cui (Wuhan Institute of Virology, CAS. ) |
| EPI_ISL_255875 | A/muscovy/duck/Hunan/HN349/2015 | A / H5N6 | Asia / China | Jie Cui (Wuhan Institute of Virology, CAS. ) |
| EPI_ISL_255876 | A/muscovy/duck/Hunan/HN350/2015 | A / H5N6 | Asia / China | Jie Cui (Wuhan Institute of Virology, CAS. ) |
| EPI_ISL_255877 | A/muscovy/duck/Hunan/HN351/2015 | A / H5N6 | Asia / China | Jie Cui (Wuhan Institute of Virology, CAS. ) |
| EPI_ISL_255878 | A/muscovy/duck/Hunan/HN352/2015 | A / H5N6 | Asia / China | Jie Cui (Wuhan Institute of Virology, CAS. ) |
| EPI_ISL_255879 | A/muscovy/duck/Hunan/HN353/2015 | A / H5N6 | Asia / China | Jie Cui (Wuhan Institute of Virology, CAS. ) |
| EPI_ISL_255880 | A/muscovy/duck/Hunan/HN354/2015 | A / H5N6 | Asia / China | Jie Cui (Wuhan Institute of Virology, CAS. ) |
| EPI_ISL_255881 | A/muscovy/duck/Hunan/HN355/2015 | A / H5N6 | Asia / China | Jie Cui (Wuhan Institute of Virology, CAS. ) |
| EPI_ISL_255882 | A/muscovy/duck/Hunan/HN356/2015 | A / H5N6 | Asia / China | Jie Cui (Wuhan Institute of Virology, CAS. ) |
| EPI_ISL_255883 | A/muscovy/duck/Hunan/HN358/2015 | A / H5N6 | Asia / China | Jie Cui (Wuhan Institute of Virology, CAS. ) |
| EPI_ISL_255884 | A/muscovy/duck/Hunan/HN359/2015 | A / H5N6 | Asia / China | Jie Cui (Wuhan Institute of Virology, CAS. ) |
| EPI_ISL_255885 | A/muscovy/duck/Hunan/HN360/2015 | A / H5N6 | Asia / China | Jie Cui (Wuhan Institute of Virology, CAS. ) |
| EPI_ISL_255886 | A/muscovy/duck/Hunan/HN364/2015 | A / H5N6 | Asia / China | Jie Cui (Wuhan Institute of Virology, CAS. ) |
| EPI_ISL_255887 | A/muscovy/duck/Hunan/HN365/2015 | A / H5N6 | Asia / China | Jie Cui (Wuhan Institute of Virology, CAS. ) |
| EPI_ISL_255888 | A/muscovy/duck/Hunan/HN366/2015 | A / H5N6 | Asia / China | Jie Cui (Wuhan Institute of Virology, CAS. ) |
| EPI_ISL_255889 | A/muscovy/duck/Hunan/HN368/2015 | A / H5N6 | Asia / China | Jie Cui (Wuhan Institute of Virology, CAS. ) |
| EPI_ISL_255891 | A/Tufted Duck/Netherlands/1/2016 | A / H5N8 | Europe / Netherlands / Provincie Noord-Holland / Edam-Volendam | Maria Johanna Poen (Erasmus Medical Center / Department of Virology) |
| EPI_ISL_255892 | A/Great Black-backed Gull/Netherlands/2/2016 | A / H5N8 | Europe / Netherlands / Provincie Noord-Holland | Maria Johanna Poen (Erasmus Medical Center / Department of Virology) |

| | | | | |
|---|---|---|---|---|
| EPI_ISL_255910 | A/Mew Gull/Netherlands/1/2016 | A / H5N8 | Europe / Netherlands / Provincie Noord-Holland | Maria Johanna Poen (Erasmus Medical Center / Department of Virology) |
| EPI_ISL_255911 | A/Eurasian Wigeon/Netherlands/2/2016 | A / H5N8 | Europe / Netherlands / Provincie Noord-Holland | Maria Johanna Poen (Erasmus Medical Center / Department of Virology) |
| EPI_ISL_255912 | A/Eurasian Wigeon/Netherlands/4/2016 | A / H5N8 | Europe / Netherlands / South Holland | Maria Johanna Poen (Erasmus Medical Center / Department of Virology) |
| EPI_ISL_255913 | A/Mallard/Netherlands/2/2017 | A / H5N8 | Europe / Netherlands / South Holland | Maria Johanna Poen (Erasmus Medical Center / Department of Virology) |
| EPI_ISL_255914 | A/Eurasian Wigeon/Netherlands/9/2016 | A / H5N8 | Europe / Netherlands / Provincie Noord-Holland | Maria Johanna Poen (Erasmus Medical Center / Department of Virology) |
| EPI_ISL_255915 | A/chicken/Poland/114/2016 | A / H5N8 | Europe / Poland / Lesser Poland Voivodeship | Edyta ?wi?to? (National Veterinary Research Institut Poland, PIWet-PIB) |
| EPI_ISL_255916 | A/chicken/Poland/002/2017 | A / H5N8 | Europe / Poland / Opole Voivodeship | Edyta ?wi?to? (National Veterinary Research Institut Poland, PIWet-PIB) |
| EPI_ISL_255917 | A/mute swan/Poland/64/2017 | A / H5N5 | Europe / Poland / Lower Silesian Voivodeship | Edyta ?wi?to? (National Veterinary Research Institut Poland, PIWet-PIB) |
| EPI_ISL_255933 | A/Cormorant/Hungary/6102/2017 | A / H5N8 | Europe / Hungary / Komarom-Esztergom | Adam Dan (Danam.Vet.Molbiol) |
| EPI_ISL_255934 | A/Mute swan/Hungary/6092/2017 | A / H5N8 | Europe / Hungary / Komarom-Esztergom | Adam Dan (Danam.Vet.Molbiol) |
| EPI_ISL_255935 | A/Mute swan/Hungary/6276/2017 | A / H5N8 | Europe / Hungary / Zala | Adam Dan (Danam.Vet.Molbiol) |
| EPI_ISL_255936 | A/Pheasant/Hungary/6553/2017 | A / H5N8 | Europe / Hungary / Jasz-Nagykun-Szolnok | Adam Dan (Danam.Vet.Molbiol) |
| EPI_ISL_255937 | A/Pheasant/Hungary/7685/2017 | A / H5N8 | Europe / Hungary / Jasz-Nagykun-Szolnok | Adam Dan (Danam.Vet.Molbiol) |
| EPI_ISL_255938 | A/Rook/Hungary/4975/2017 | A / H5N8 | Europe / Hungary / Szabolcs-Szatmar-Bereg | Adam Dan (Danam.Vet.Molbiol) |
| EPI_ISL_256213 | A/Hubei/29578/2016 | A / H5N6 | Asia / China / Hubei Province | Lei Yang (WHO Chinese National Influenza Center / Virology Institute, Chinese CDC) |
| EPI_ISL_256290 | A/environment/Changsha/156/2014 | A / H5N1 | Asia / China | |
| EPI_ISL_256298 | A/gadwall/Kurgan/2442/2016 | A / H5N8 | Europe / Russian Federation / Kurgan Oblast | Ivan Susloparov (State Research Center of Virology and Biotechnology (VECTOR) / Emerging Zoonotic Diseases and Influenza) |
| EPI_ISL_256299 | A/chicken/Sergiyev Posad/38/2017 | A / H5N8 | Europe / Russian Federation / Moscow Oblast | Ivan Susloparov (State Research Center of Virology and Biotechnology (VECTOR) / Emerging Zoonotic Diseases and Influenza) |
| EPI_ISL_256300 | A/chicken/Sergiyev Posad/39/2017 | A / H5N8 | Europe / Russian Federation / Moscow Oblast | Ivan Susloparov (State Research Center of Virology and Biotechnology (VECTOR) / Emerging Zoonotic Diseases and Influenza) |
| EPI_ISL_256301 | A/environment/Kamchatka/18/2016 | A / H5N5 | Europe / Russian Federation / Kamchatka Krai | Ivan Susloparov (State Research Center of Virology and Biotechnology (VECTOR) / Emerging Zoonotic Diseases and Influenza) |
| EPI_ISL_256302 | A/mute swan/Czech Republic/499-17/2017 (H5N8) | A / H5N8 | Europe / Czech Republic / Olomoucky Kraj / Okres Olomouc / Olomouc - Nové Sady | Alexander Nagy (State Veterinary Institute Prague) |
| EPI_ISL_256304 | A/mute swan/Czech Republic/581-17/2017 (H5N8) | A / H5N8 | Europe / Czech Republic / Olomoucky Kraj / Okres Olomouc / Olomouc city - U D?tského domov[illegible] | Alexander Nagy (State Veterinary Institute Prague) |
| EPI_ISL_256305 | A/shelduck/Italy/17VIR1572-24/2017 | A / H5N8 | Europe / Italy / Rovigo | Bianca Zecchin (Istituto Zooprofilattico Sperimentale Delle Venezie) |
| EPI_ISL_256306 | A/turkey/Italy/17VIR1574-1/2017 | A / H5N8 | Europe / Italy / Mantova | Bianca Zecchin (Istituto Zooprofilattico Sperimentale Delle Venezie) |
| EPI_ISL_256307 | A/chicken/Italy/17VIR1684-2/2017 | A / H5N8 | Europe / Italy / Venezia | Bianca Zecchin (Istituto Zooprofilattico Sperimentale Delle Venezie) |
| EPI_ISL_256308 | A/chicken/Italy/17VIR1751-3/2017 | A / H5N8 | Europe / Italy / Venezia | Bianca Zecchin (Istituto Zooprofilattico Sperimentale Delle Venezie) |
| EPI_ISL_256309 | A/goose/Czech Republic/821-17_2/2017 (H5N8) | A / H5N8 | Europe / Czech Republic / Jihocesky Kraj / Okres Tabor / Sedle?ko u Sob?slav? | Alexander Nagy (State Veterinary Institute Prague) |
| EPI_ISL_256396 | A/chicken/Taiwan/u7/2016 | A / H5N2 | Asia / Taiwan | Yu-Pin Liu (Animal Health Research Institute) |
| EPI_ISL_256398 | A/chicken/Taiwan/x37/2016 | A / H5N8 | Asia / Taiwan | Yu-Pin Liu (Animal Health Research Institute) |
| EPI_ISL_256453 | A/Duck/Hungary/54738/2016 | A / H5N8 | Europe / Hungary / Bacs-Kiskun | Adam Dan (Danam.Vet.Molbiol) |
| EPI_ISL_256457 | A/Duck/Hungary/55764/2016 | A / H5N8 | Europe / Hungary / Bacs-Kiskun | Adam Dan (Danam.Vet.Molbiol) |
| EPI_ISL_256460 | A/Turkey/Hungary/53136/2016 | A / H5N8 | Europe / Hungary / Bekes megye / Bekes | Adam Dan (Danam.Vet.Molbiol) |
| EPI_ISL_256462 | A/Mute_swan/Hungary/5879/2017 | A / H5N5 | Europe / Hungary / Budapest fovaros | Adam Dan (Danam.Vet.Molbiol) |
| EPI_ISL_256495 | A/owl/Hokkaido/X6/2016 | A / H5N6 | Asia / Japan / Hokkaido | Masatoshi Okamatsu (Hokkaido University / Graduate School of Veterinary Medicine) |
| EPI_ISL_256496 | A/peregrine falcon/Hokkaido/X7/2016 | A / H5N6 | Asia / Japan / Hokkaido | Masatoshi Okamatsu (Hokkaido University / Graduate School of Veterinary Medicine) |
| EPI_ISL_256497 | A/pintail/Hokkaido/X8/2016 | A / H5N6 | Asia / Japan / Hokkaido | Masatoshi Okamatsu (Hokkaido University / Graduate School of Veterinary Medicine) |
| EPI_ISL_256498 | A/whooper swan/Hokkaido/X12/2017 | A / H5N6 | Asia / Japan / Hokkaido | Masatoshi Okamatsu (Hokkaido University / Graduate School of Veterinary Medicine) |
| EPI_ISL_256499 | A/whooper swan/Hokkaido/X13/2017 | A / H5N6 | Asia / Japan / Hokkaido | Masatoshi Okamatsu (Hokkaido University / Graduate School of Veterinary Medicine) |
| EPI_ISL_256500 | A/chicken/Hokkaido/002/2017 | A / H5N6 | Asia / Japan / Hokkaido | Masatoshi Okamatsu (Hokkaido University / Graduate School of Veterinary Medicine) |
| EPI_ISL_256501 | A/mute swan/Aomori/4/2016 | A / H5N6 | Asia / Japan / Aomori | Masatoshi Okamatsu (Hokkaido University / Graduate School of Veterinary Medicine) |
| EPI_ISL_256505 | A/whooper swan/Aomori/9/2016 | A / H5N6 | Asia / Japan | Masatoshi Okamatsu (Hokkaido University / Graduate School of Veterinary Medicine) |
| EPI_ISL_256506 | A/grey-faced buzzard/Aomori/10/2016 | A / H5N6 | Asia / Japan | Masatoshi Okamatsu (Hokkaido University / Graduate School of Veterinary Medicine) |
| EPI_ISL_256507 | A/whooper swan/Iwate/1/2016 | A / H5N6 | Asia / Japan | Masatoshi Okamatsu (Hokkaido University / Graduate School of Veterinary Medicine) |
| EPI_ISL_256508 | A/whooper swan/Iwate/5/2016 | A / H5N6 | Asia / Japan | Masatoshi Okamatsu (Hokkaido University / Graduate School of Veterinary Medicine) |
| EPI_ISL_256509 | A/whooper swan/Iwate/8/2016 | A / H5N6 | Asia / Japan | Masatoshi Okamatsu (Hokkaido University / Graduate School of Veterinary Medicine) |
| EPI_ISL_256510 | A/tundra swan/Iwate/9/2016 | A / H5N6 | Asia / Japan | Masatoshi Okamatsu (Hokkaido University / Graduate School of Veterinary Medicine) |
| EPI_ISL_256511 | A/whooper swan/Iwate/11/2016 | A / H5N6 | Asia / Japan | Masatoshi Okamatsu (Hokkaido University / Graduate School of Veterinary Medicine) |
| EPI_ISL_256512 | A/coot/Iwate/13/2016 | A / H5N6 | Asia / Japan | Masatoshi Okamatsu (Hokkaido University / Graduate School of Veterinary Medicine) |
| EPI_ISL_256513 | A/whooper swan/Iwate/17/2016 | A / H5N6 | Asia / Japan | Masatoshi Okamatsu (Hokkaido University / Graduate School of Veterinary Medicine) |
| EPI_ISL_256514 | A/white-fronted goose/Miyagi/1/2016 | A / H5N6 | Asia / Japan | Masatoshi Okamatsu (Hokkaido University / Graduate School of Veterinary Medicine) |
| EPI_ISL_256515 | A/white-fronted goose/Miyagi/2/2016 | A / H5N6 | Asia / Japan | Masatoshi Okamatsu (Hokkaido University / Graduate School of Veterinary Medicine) |
| EPI_ISL_256516 | A/whooper swan/Fukushima/1/2016 | A / H5N6 | Asia / Japan | Masatoshi Okamatsu (Hokkaido University / Graduate School of Veterinary Medicine) |
| EPI_ISL_256517 | A/whooper swan/Fukushima/3/2016 | A / H5N6 | Asia / Japan | Masatoshi Okamatsu (Hokkaido University / Graduate School of Veterinary Medicine) |
| EPI_ISL_256518 | A/tundra swan/Niigata/1/2016 | A / H5N6 | Asia / Japan | Masatoshi Okamatsu (Hokkaido University / Graduate School of Veterinary Medicine) |
| EPI_ISL_256519 | A/tundra swan/Niigata/4/2016 | A / H5N6 | Asia / Japan | Masatoshi Okamatsu (Hokkaido University / Graduate School of Veterinary Medicine) |
| EPI_ISL_256520 | A/tundra swan/Niigata/8/2016 | A / H5N6 | Asia / Japan | Masatoshi Okamatsu (Hokkaido University / Graduate School of Veterinary Medicine) |
| EPI_ISL_256521 | A/peregrine falcon/Niigata/12/2017 | A / H5N6 | Asia / Japan | Masatoshi Okamatsu (Hokkaido University / Graduate School of Veterinary Medicine) |
| EPI_ISL_256522 | A/whooper swan/Niigata/13/2017 | A / H5N6 | Asia / Japan | Masatoshi Okamatsu (Hokkaido University / Graduate School of Veterinary Medicine) |
| EPI_ISL_256523 | A/mute swan/Ibaraki/7/2016 | A / H5N6 | Asia / Japan | Masatoshi Okamatsu (Hokkaido University / Graduate School of Veterinary Medicine) |
| EPI_ISL_256524 | A/whooper swan/Tochigi/1/2017 | A / H5N6 | Asia / Japan | Masatoshi Okamatsu (Hokkaido University / Graduate School of Veterinary Medicine) |
| EPI_ISL_256817 | A/turkey/Germany-NI/R9807/2016 | A / H5N8 | Europe / Germany / Lower Saxony | Elke Starick (Friedrich-Loeffler-Institut) |
| EPI_ISL_256818 | A/domestic duck/Germany-MV/R9764/2016 | A / H5N8 | Europe / Germany / Mecklenburg-Vorpommern | Elke Starick (Friedrich-Loeffler-Institut) |
| EPI_ISL_2574053 | A/Towny owel/Sweden/SVAU210406SZ0042/KN001362/M-20[illegible] | A / H5N8 | Europe / Sweden / Skane Lan / Sjobo Kommun | Siamak Zohari (National Veterinary Institute) |
| EPI_ISL_257416 | A/black_swan/Ibaraki/28-446T/2017 | A / H5N6 | Asia / Japan | Takehiko Saito (National Institute of Animal Health) |
| EPI_ISL_257417 | A/black_swan/Ibaraki/28-446C/2017 | A / H5N6 | Asia / Japan | Takehiko Saito (National Institute of Animal Health) |
| EPI_ISL_257438 | A/chicken/Miyagi/1-5C/2017 | A / H5N6 | Asia / Japan / Miyagi | Takehiko Saito (National Institute of Animal Health) |
| EPI_ISL_257439 | A/chicken/Chiba/1-4C/2017 | A / H5N6 | Asia / Japan / Chiba | Takehiko Saito (National Institute of Animal Health) |
| EPI_ISL_257440 | A/chicken/Chiba/1-5T/2017 | A / H5N6 | Asia / Japan / Chiba | Takehiko Saito (National Institute of Animal Health) |
| EPI_ISL_257441 | A/chicken/Chiba/1-3C/2017 | A / H5N6 | Asia / Japan / Chiba | Takehiko Saito (National Institute of Animal Health) |
| EPI_ISL_257442 | A/chicken/Chiba/1-3T/2017 | A / H5N6 | Asia / Japan / Chiba | Takehiko Saito (National Institute of Animal Health) |
| EPI_ISL_257443 | A/chicken/Chiba/1-5C/2017 | A / H5N6 | Asia / Japan / Chiba | Takehiko Saito (National Institute of Animal Health) |
| EPI_ISL_257444 | A/chicken/Chiba/1-2T/2017 | A / H5N6 | Asia / Japan / Chiba | Takehiko Saito (National Institute of Animal Health) |

| | | | | |
|---|---|---|---|---|
| EPI_ISL_257445 | A/chicken/Chiba/1-1T/2017 | A / H5N6 | Asia / Japan / Chiba | Takehiko Saito (National Institute of Animal Health) |
| EPI_ISL_257446 | A/chicken/Chiba/1-4T/2017 | A / H5N6 | Asia / Japan / Chiba | Takehiko Saito (National Institute of Animal Health) |
| EPI_ISL_257447 | A/chicken/Chiba/1-2C/2017 | A / H5N6 | Asia / Japan / Chiba | Takehiko Saito (National Institute of Animal Health) |
| EPI_ISL_257448 | A/chicken/Miyagi/1-5T/2017 | A / H5N6 | Asia / Japan / Miyagi | Takehiko Saito (National Institute of Animal Health) |
| EPI_ISL_257449 | A/chicken/Chiba/1-1C/2017 | A / H5N6 | Asia / Japan / Chiba | Takehiko Saito (National Institute of Animal Health) |
| EPI_ISL_257657 | A/Pavo cristatus/Jiangxi/JA1/2016 | A / H5N6 | Asia / China | |
| EPI_ISL_257699 | A/Tufted Duck/Switzerland/V237/2016 | A / H5N8 | Europe / Switzerland / Kanton Bern / Amt Biel / Lake Biel | Ronald Dijkman (Faculty of Veterinary Medicine at the University of Bern / Institute of Virology and Immunology IVI) |
| EPI_ISL_257700 | A/mute swan/Czech Republic/1813-17/2017 (H5N8) | A / H5N8 | Europe / Czech Republic / Karlovarsky Kraj / Okres Cheb / GPS: 50°4'56.294''N, 12°22'7.530E | Alexander Nagy (State Veterinary Institute Prague) |
| EPI_ISL_257701 | A/chicken/Czech Republic/2643-17_1/2017 (H5N8) | A / H5N8 | Europe / Czech Republic / Jihocesky Kraj / Okres Jindrichuv Hradec / Da?ice | Alexander Nagy (State Veterinary Institute Prague) |
| EPI_ISL_257702 | A/chicken/Czech Republic/2644-17_1/2017 (H5N8) | A / H5N8 | Europe / Czech Republic / Plzensky Kraj / Bohy (Kralovice), GPS: 49°56'15.55''N, 13°35'17.58E | Alexander Nagy (State Veterinary Institute Prague) |
| EPI_ISL_257703 | A/mallard/Czech Republic/2705-17/2017 (H5N8) | A / H5N8 | Europe / Czech Republic / Kralovehradecky Kraj / Havlí?k?v Brod, GPS: 49°36'13.705''N, 15°35'0.. | Alexander Nagy (State Veterinary Institute Prague) |
| EPI_ISL_257731 | A/turkey/Germany-BB/R377ff/2017 | A / H5N8 | Europe / Germany / Brandenburg | Elke Starick (Friedrich-Loeffler-Institut) |
| EPI_ISL_258415 | A/grey heron/Germany-TH/R1125/2017 | A / H5N8 | Europe / Germany / Thuringia | Elke Starick (Friedrich-Loeffler-Institut) |
| EPI_ISL_258526 | A/white stork/Germany-TH/R1149/2017 | A / H5N8 | Europe / Germany / Thuringia | Elke Starick (Friedrich-Loeffler-Institut) |
| EPI_ISL_258527 | A/tawny owl/Germany-SN/R1186/2017 | A / H5N8 | Europe / Germany / Saxony | Elke Starick (Friedrich-Loeffler-Institut) |
| EPI_ISL_258661 | A/mute swan/Germany-TH/R1126/2017 | A / H5N8 | Europe / Germany / Thuringia | Elke Starick (Friedrich-Loeffler-Institut) |
| EPI_ISL_259074 | A/common buzzard/Germany-SN/R1117/2017 | A / H5N5 | Europe / Germany / Saxony | Elke Starick (Friedrich-Loeffler-Institut) |
| EPI_ISL_259525 | A/egret/Germany-SH/R1459/2017 | A / H5N5 | Europe / Germany / Schleswig-Holstein | Elke Starick (Friedrich-Loeffler-Institut) |
| EPI_ISL_259923 | A/duck/Guangdong/673/2014(H5N6) | A / H5N6 | Asia / China / Guangdong Province | Qi Wenbao (South China Agricultural University / Veterinary Medicine College) |
| EPI_ISL_259924 | A/goose/Guangdong/674/2014(H5N6) | A / H5N6 | Asia / China / Guangdong Province | Qi Wenbao (South China Agricultural University / Veterinary Medicine College) |
| EPI_ISL_259927 | A/goose/Guangdong/SH7/2013(H5N1) | A / H5N1 | Asia / China / Guangdong Province | Qi Wenbao (South China Agricultural University / Veterinary Medicine College) |
| EPI_ISL_260058 | A/grey heron/Germany-SN/R572/2017 | A / H5N5 | Europe / Germany / Saxony | Elke Starick (Friedrich-Loeffler-Institut) |
| EPI_ISL_260059 | A/cormorant/Germany-SH/R896/2017 | A / H5N5 | Europe / Germany / Schleswig-Holstein | Elke Starick (Friedrich-Loeffler-Institut) |
| EPI_ISL_260773 | A/whooper swan/Iwate/7/2016 | A / H5N6 | Asia / Japan | Masatoshi Okamatsu (Hokkaido University / Graduate School of Veterinary Medicine) |
| EPI_ISL_261332 | A/chicken/Croatia/104/2017 | A / H5N5 | Europe / Croatia / Krapinsko-Zagorska Zupanija / Špi?kovina | Vladimir Savi? (Croatian Veterinary Institute / Poultry Centre) |
| EPI_ISL_261634 | A/chicken/Republic of Macedonia/AR1167-L02131/2017 | A / H5N8 | Europe / Macedonia, the former Yogoslav Republic of / N 41°12'10, E20°42'01 | Anne Pohlmann (Friedrich-Loeffler-Institut) |
| EPI_ISL_261687 | A/black swan/Germany-BW/R1364/2017 | A / H5N8 | Europe / Germany / Baden-Wuerttemberg | Elke Starick (Friedrich-Loeffler-Institut) |
| EPI_ISL_262055 | A/eurasian wigeon/Germany-NI/AR249-L02143/2017 | A / H5N8 | Europe / Germany / Lower Saxony / Wilhelmshaven | Anne Pohlmann (Friedrich-Loeffler-Institut) |
| EPI_ISL_262056 | A/greylag goose/Germany-NI/AR11353-L02142/2016 | A / H5N5 | Europe / Germany / Lower Saxony / Wangerland | Anne Pohlmann (Friedrich-Loeffler-Institut) |
| EPI_ISL_262057 | A/mute swan/Germany-NI/AR1529-L02145/2017 | A / H5N8 | Europe / Germany / Lower Saxony / Ammerland | Anne Pohlmann (Friedrich-Loeffler-Institut) |
| EPI_ISL_262058 | A/greylag goose/Germany-NI/AR1395-L02144/2017 | A / H5N8 | Europe / Germany / Lower Saxony / Peine | Anne Pohlmann (Friedrich-Loeffler-Institut) |
| EPI_ISL_262059 | A/greylag goose/Germany-NI/AR703-L02138/2017 | A / H5N8 | Europe / Germany / Lower Saxony / Hemmingen | Anne Pohlmann (Friedrich-Loeffler-Institut) |
| EPI_ISL_2652003 | A/White-Tailed Eagle/Sweden/SVA210528SZ0223/KN002027/ | A / H5N1 | Europe / Sweden / Stockholms Lan / Ekero Kommun | Siamak Zohari (National Veterinary Institute) |
| EPI_ISL_266417 | A/common teal/Korea/W547/2016 | A / H5N8 | Asia / Korea, Republic of | |
| EPI_ISL_266418 | A/common teal/Korea/W549/2016 | A / H5N8 | Asia / Korea, Republic of | |
| EPI_ISL_266419 | A/common teal/Korea/W548/2016 | A / H5N8 | Asia / Korea, Republic of | |
| EPI_ISL_266420 | A/common teal/Korea/W550/2016 | A / H5N8 | Asia / Korea, Republic of | |
| EPI_ISL_266421 | A/common teal/Korea/W555/2017 | A / H5N8 | Asia / Korea, Republic of | |
| EPI_ISL_266424 | A/common teal/Korea/W558/2017 | A / H5N6 | Asia / Korea, Republic of | |
| EPI_ISL_266425 | A/common teal/Korea/W559/2017 | A / H5N6 | Asia / Korea, Republic of | |
| EPI_ISL_266514 | A/environment/Zhongshan/ZS01/2016 | A / H5N6 | Asia / China | |
| EPI_ISL_266536 | A/chicken/Belgium/807/2017 | A / H5N8 | Europe / Belgium | |
| EPI_ISL_266537 | A/peacock/Belgium/1017/2017 | A / H5N8 | Europe / Belgium | |
| EPI_ISL_266538 | A/Cygnus olor/Belgium/1567/2017 | A / H5N8 | Europe / Belgium | |
| EPI_ISL_266596 | A/swine/Guangdong/G3/2015 | A / H5N6 | Asia / China / Guangdong Province | Feng Qin (South China Agricultural University / Veterinary Medicine College) |
| EPI_ISL_266600 | A/enviroment/Guangdong/F3/2015 | A / H5N6 | Asia / China / Guangdong Province | Feng Qin (South China Agricultural University / Veterinary Medicine College) |
| EPI_ISL_266819 | A/enviroment/Guangdong/F4/2016 | A / H5N6 | Asia / China / Guangdong Province | Feng Qin (South China Agricultural University / Veterinary Medicine College) |
| EPI_ISL_266820 | A/Bean goose/Hubei/CH-i122/2017_H5N8 | A / H5N8 | Asia / China / Hubei Province | Jianjun Chen (Wuhan Institute of Virology / Chinese Academy of Sciences) |
| EPI_ISL_266821 | A/Bean goose/Hubei/CH-i119/2017 | A / H5N8 | Asia / China / Hubei Province | Jianjun Chen (Wuhan Institute of Virology / Chinese Academy of Sciences) |
| EPI_ISL_266822 | A/Herring Gull/Hubei/CH-i149/2017_H5N8 | A / H5N8 | Asia / China / Hubei Province | Jianjun Chen (Wuhan Institute of Virology / Chinese Academy of Sciences) |
| EPI_ISL_266823 | A/Bean goose/Hubei/CH-i336/2017 | A / H5N6 | Asia / China / Hubei Province | Jianjun Chen (Wuhan Institute of Virology / Chinese Academy of Sciences) |
| EPI_ISL_266824 | A/Bean goose/Hubei/CH-i320/2017_H5N8 | A / H5N8 | Asia / China / Hubei Province | Jianjun Chen (Wuhan Institute of Virology / Chinese Academy of Sciences) |
| EPI_ISL_266825 | A/Bean goose/Hubei/CH-I225/2017 | A / H5N6 | Asia / China / Hubei Province | Jianjun Chen (Wuhan Institute of Virology / Chinese Academy of Sciences) |
| EPI_ISL_266939 | A/tundra swan/Tottori/3111S001/2016 | A / H5N6 | Asia / Japan / Tottori | |
| EPI_ISL_266940 | A/tundra swan/Tottori/3111S002/2016 | A / H5N6 | Asia / Japan / Tottori | |
| EPI_ISL_266941 | A/snowy owl/Akita/0051D007/2016 | A / H5N6 | Asia / Japan / Akita | |
| EPI_ISL_266942 | A/snowy owl/Akita/0051D008/2016 | A / H5N6 | Asia / Japan / Akita | |
| EPI_ISL_266943 | A/snowy owl/Akita/0051D010-4/2016 | A / H5N6 | Asia / Japan / Akita | |
| EPI_ISL_267135 | A/green-winged teal/Egypt/871/2016 | A / H5N8 | Africa / Egypt | |
| EPI_ISL_267136 | A/green-winged teal/Egypt/877/2016 | A / H5N8 | Africa / Egypt | |
| EPI_ISL_268021 | A/duck/Egypt/SS19/2017 | A / H5N8 | Africa / Egypt / Sharqia | Abdel-Satar Mohamed Arafa (Animal Health Research Institute) |
| EPI_ISL_2681044 | A/chicken/Romania/12448_21VIR3734-3/2021 | A / H5N8 | Europe / Romania / Mures, Ungheni | Bianca Zecchin (Istituto Zooprofilattico Sperimentale Delle Venezie) |
| EPI_ISL_2681045 | A/white_stork/Poland/MB391/2021 | A / H5N1 | Europe / Poland / Swietokrzyskie Voivodeship | Edyta ?wi?to? (National Veterinary Research Institut Poland, PIWet-PIB) |
| EPI_ISL_268513 | A/duck/Egypt/F446/2017 | A / H5N8 | Africa / Egypt / Cairo | Abdel-Satar Mohamed Arafa (Animal Health Research Institute) |
| EPI_ISL_2685842 | A/tundra swan/Hubei/BQ2/2020 | A / H5N8 | Asia / China / Hubei Province | Hongliang Chai (Northeast Forestry University / College of Wildlife Resources) |
| EPI_ISL_2685846 | A/tundra swan/Hubei/BQ3/2020 | A / H5N8 | Asia / China / Hubei Province | Hongliang Chai (Northeast Forestry University / College of Wildlife Resources) |
| EPI_ISL_2685847 | A/tundra swan/Hubei/BQ4/2020 | A / H5N8 | Asia / China / Hubei Province | Hongliang Chai (Northeast Forestry University / College of Wildlife Resources) |
| EPI_ISL_2685848 | A/tundra swan/Hubei/BQ6/2020 | A / H5N8 | Asia / China / Hubei Province | Hongliang Chai (Northeast Forestry University / College of Wildlife Resources) |
| EPI_ISL_2685900 | A/tundra swan/Hubei/BQ7/2020 | A / H5N8 | Asia / China / Hubei Province | Hongliang Chai (Northeast Forestry University / College of Wildlife Resources) |
| EPI_ISL_2685947 | A/tundra swan/Hubei/BQ8/2020 | A / H5N8 | Asia / China / Hubei Province | Hongliang Chai (Northeast Forestry University / College of Wildlife Resources) |
| EPI_ISL_268618 | A/Bk_swan/NL-Den Oever/16013973-002/2016 | A / H5N8 | Europe / Netherlands / Provincie Noord-Holland / Den Oever | Saskia Bergervoet (Wageningen Bioveterinary Research) |
| EPI_ISL_268619 | A/Bl_H_gull/NL-Slootdorp/16014102-002/2016 | A / H5N8 | Europe / Netherlands / Provincie Noord-Holland / Slootdorp | Saskia Bergervoet (Wageningen Bioveterinary Research) |

| | | | | |
|---|---|---|---|---|
| EPI_ISL_268620 | A/Buzzard/NL-Durgerdam/16015100-004/2016 | A / H5N8 | Europe / Netherlands / Provincie Noord-Holland / Durgerdam | Saskia Bergervoet (Wageningen Bioveterinary Research) |
| EPI_ISL_268621 | A/C_Gull/NL-Slootdorp/16014102-003/2016 | A / H5N8 | Europe / Netherlands / Provincie Noord-Holland / Slootdorp | Saskia Bergervoet (Wageningen Bioveterinary Research) |
| EPI_ISL_268622 | A/Ch/NL-Abbega/X16015736/2016 | A / H5N8 | Europe / Netherlands / Provincie Friesland / Abbega | Saskia Bergervoet (Wageningen Bioveterinary Research) |
| EPI_ISL_268623 | A/Ch/NL-Boven Leeuwen/16016151-006-010/2016 | A / H5N8 | Europe / Netherlands / Provincie Gelderland / Boven Leeuwen | Saskia Bergervoet (Wageningen Bioveterinary Research) |
| EPI_ISL_268624 | A/Ch/NL-Den Oever/16014231-001/2016 | A / H5N8 | Europe / Netherlands / Provincie Noord-Holland / Den Oever | Saskia Bergervoet (Wageningen Bioveterinary Research) |
| EPI_ISL_268625 | A/Ch/NL-Hiaure/16016112-001-005/2016 | A / H5N8 | Europe / Netherlands / Provincie Friesland / Hiaure | Saskia Bergervoet (Wageningen Bioveterinary Research) |
| EPI_ISL_268626 | A/Ch/NL-Rhenen/16016141-006/2016 | A / H5N8 | Europe / Netherlands / Provincie Utrecht / Gemeente Rhenen | Saskia Bergervoet (Wageningen Bioveterinary Research) |
| EPI_ISL_268627 | A/Ch/NL-Zoeterwoude/16016484-021-025/2016 | A / H5N8 | Europe / Netherlands / South Holland / Gemeente Zoeterwoude | Saskia Bergervoet (Wageningen Bioveterinary Research) |
| EPI_ISL_268628 | A/Crow/NL-Oostwoud/16015372-004/2016 | A / H5N8 | Europe / Netherlands / Provincie Noord-Holland / Oostwoud | Saskia Bergervoet (Wageningen Bioveterinary Research) |
| EPI_ISL_268629 | A/Dk/NL-Biddinghuizen/16014829-011-015/2016 | A / H5N8 | Europe / Netherlands / Provincie Flevoland / Biddinghuizen | Saskia Bergervoet (Wageningen Bioveterinary Research) |
| EPI_ISL_268630 | A/Dk/NL-Biddinghuizen/16015083-016-020/2016 | A / H5N8 | Europe / Netherlands / Provincie Flevoland / Biddinghuizen | Saskia Bergervoet (Wageningen Bioveterinary Research) |
| EPI_ISL_268631 | A/Dk/NL-Biddinghuizen/16015145-021-025/2016 | A / H5N8 | Europe / Netherlands / Provincie Flevoland / Biddinghuizen | Saskia Bergervoet (Wageningen Bioveterinary Research) |
| EPI_ISL_268632 | A/Dk/NL-Kamperveen/16016104-001-005/2016 | A / H5N8 | Europe / Netherlands / Provincie Overijssel / Kamperveen | Saskia Bergervoet (Wageningen Bioveterinary Research) |
| EPI_ISL_268633 | A/Dk/NL-Rotterdam/16014008-001-005/2016 | A / H5N8 | Europe / Netherlands / South Holland / Gemeente Rotterdam | Saskia Bergervoet (Wageningen Bioveterinary Research) |
| EPI_ISL_268634 | A/Dk/NL-Stolwijk/16016291-016-020/2016 | A / H5N8 | Europe / Netherlands / South Holland / Stolwijk | Saskia Bergervoet (Wageningen Bioveterinary Research) |
| EPI_ISL_268635 | A/Eur_Wig/NL-Akkrum/16015817-003/2016 | A / H5N8 | Europe / Netherlands / Provincie Friesland / Akkrum | Saskia Bergervoet (Wageningen Bioveterinary Research) |
| EPI_ISL_268636 | A/Eur_Wig/NL-De Waal (Texel)/16014891-003/2016 | A / H5N8 | Europe / Netherlands / Provincie Noord-Holland / De Waal (Texel) | Saskia Bergervoet (Wageningen Bioveterinary Research) |
| EPI_ISL_268637 | A/Eur_Wig/NL-De Waal (Texel)/16014891-004/2016 | A / H5N8 | Europe / Netherlands / Provincie Noord-Holland / De Waal (Texel) | Saskia Bergervoet (Wageningen Bioveterinary Research) |
| EPI_ISL_268638 | A/Eur_Wig/NL-Drieborg (Dollard)/16015513-001/2016 | A / H5N8 | Europe / Netherlands / Provincie Groningen / Drieborg (Dollard) | Saskia Bergervoet (Wageningen Bioveterinary Research) |
| EPI_ISL_268639 | A/Eur_Wig/NL-Enumatil-Groningen/16015704-001/2016 | A / H5N8 | Europe / Netherlands / Provincie Groningen / Enumatil-Groningen | Saskia Bergervoet (Wageningen Bioveterinary Research) |
| EPI_ISL_268640 | A/Eur_Wig/NL-Ferwert/16015273-002/2016 | A / H5N8 | Europe / Netherlands / Provincie Friesland / Ferwert | Saskia Bergervoet (Wageningen Bioveterinary Research) |
| EPI_ISL_268641 | A/Eur_Wig/NL-Gouda/16015824-001/2016 | A / H5N8 | Europe / Netherlands / South Holland / Gemeente Gouda | Saskia Bergervoet (Wageningen Bioveterinary Research) |
| EPI_ISL_268642 | A/Eur_Wig/NL-Greonterp/16015653-001/2016 | A / H5N8 | Europe / Netherlands / Provincie Friesland / Greonterp | Saskia Bergervoet (Wageningen Bioveterinary Research) |
| EPI_ISL_268643 | A/Eur_Wig/NL-Groningen/16015376-003/2016 | A / H5N8 | Europe / Netherlands / Provincie Groningen / Gemeente Groningen | Saskia Bergervoet (Wageningen Bioveterinary Research) |
| EPI_ISL_268644 | A/Eur_Wig/NL-Leeuwarden/16015699-002/2016 | A / H5N8 | Europe / Netherlands / Provincie Friesland / Gemeente Leeuwarden | Saskia Bergervoet (Wageningen Bioveterinary Research) |
| EPI_ISL_268645 | A/Eur_Wig/NL-Leidschendam/16015697-007/2016 | A / H5N8 | Europe / Netherlands / South Holland / Leidschendam | Saskia Bergervoet (Wageningen Bioveterinary Research) |
| EPI_ISL_268646 | A/Eur_Wig/NL-Reeuwijk/16015903-003/2016 | A / H5N8 | Europe / Netherlands / South Holland / Gemeente Reeuwijk | Saskia Bergervoet (Wageningen Bioveterinary Research) |
| EPI_ISL_268647 | A/Eur_Wig/NL-Terschelling/16015692-010/2016 | A / H5N8 | Europe / Netherlands / Provincie Friesland / Gemeente Terschelling | Saskia Bergervoet (Wageningen Bioveterinary Research) |
| EPI_ISL_268648 | A/Eur_Wig/NL-Vianen/16015917-006/2016 | A / H5N8 | Europe / Netherlands / Provincie Utrecht / Gemeente Vianen | Saskia Bergervoet (Wageningen Bioveterinary Research) |
| EPI_ISL_268649 | A/Eur_Wig/NL-Walterswald/16015923-003/2016 | A / H5N8 | Europe / Netherlands / Provincie Friesland / Walterswald | Saskia Bergervoet (Wageningen Bioveterinary Research) |
| EPI_ISL_268650 | A/Eur_Wig/NL-West Graftdijk/16015746-003/2016 | A / H5N8 | Europe / Netherlands / Provincie Noord-Holland / West Graftdijk | Saskia Bergervoet (Wageningen Bioveterinary Research) |
| EPI_ISL_268651 | A/Eur_Wig/NL-Wormer/16016143-002/2016 | A / H5N8 | Europe / Netherlands / Provincie Noord-Holland / Wormer | Saskia Bergervoet (Wageningen Bioveterinary Research) |
| EPI_ISL_268652 | A/Eur_Wig/NL-Zoeterwoude/16015702-010/2016 | A / H5N8 | Europe / Netherlands / South Holland / Gemeente Zoeterwoude | Saskia Bergervoet (Wageningen Bioveterinary Research) |
| EPI_ISL_268653 | A/Eur_Wig/NL-Zwolle/16015820-002/2016 | A / H5N8 | Europe / Netherlands / Provincie Overijssel / Gemeente Zwolle | Saskia Bergervoet (Wageningen Bioveterinary Research) |
| EPI_ISL_268654 | A/G_c_grebe/NL-Monnickendam/16013865-009-010/2016 | A / H5N8 | Europe / Netherlands / Provincie Noord-Holland / Monnickendam | Saskia Bergervoet (Wageningen Bioveterinary Research) |
| EPI_ISL_268655 | A/Go/NL-Roggebotsluis/16014462-010/2016 | A / H5N8 | Europe / Netherlands / Provincie Flevoland / Roggebotsluis | Saskia Bergervoet (Wageningen Bioveterinary Research) |
| EPI_ISL_268656 | A/Gr_bk_bd_gull/NL-Slootdorp/16014102-005/2016 | A / H5N8 | Europe / Netherlands / Provincie Noord-Holland / Slootdorp | Saskia Bergervoet (Wageningen Bioveterinary Research) |
| EPI_ISL_268657 | A/Grey_Go/NL-Groot-Ammers/16015901-012/2016 | A / H5N8 | Europe / Netherlands / South Holland / Groot-Ammers | Saskia Bergervoet (Wageningen Bioveterinary Research) |
| EPI_ISL_268658 | A/Gull/NL-Marker Wadden/16014466-020/2016 | A / H5N8 | Europe / Netherlands / Marker Wadden | Saskia Bergervoet (Wageningen Bioveterinary Research) |
| EPI_ISL_268659 | A/Gull1/NL-Marker Wadden/16014466-011/2016 | A / H5N8 | Europe / Netherlands / Marker Wadden | Saskia Bergervoet (Wageningen Bioveterinary Research) |
| EPI_ISL_268660 | A/Gull10/NL-Marker Wadden/16014466-014/2016 | A / H5N8 | Europe / Netherlands / Marker Wadden | Saskia Bergervoet (Wageningen Bioveterinary Research) |
| EPI_ISL_268661 | A/L-bl-ba-gull/NL-Sovon/16014324-014/2016 | A / H5N8 | Europe / Netherlands | Saskia Bergervoet (Wageningen Bioveterinary Research) |
| EPI_ISL_268662 | A/M_Swan/NL-Roggebotsluis/16014462-019/2016 | A / H5N8 | Europe / Netherlands / Provincie Flevoland / Roggebotsluis | Saskia Bergervoet (Wageningen Bioveterinary Research) |
| EPI_ISL_268663 | A/Magpie/NL-Volendam/16014331-002/2016 | A / H5N8 | Europe / Netherlands / Provincie Noord-Holland / Edam-Volendam | Saskia Bergervoet (Wageningen Bioveterinary Research) |
| EPI_ISL_268664 | A/Mal/NL-IJsselmuiden/16015448-002/2016 | A / H5N8 | Europe / Netherlands / Provincie Overijssel / IJsselmuiden | Saskia Bergervoet (Wageningen Bioveterinary Research) |
| EPI_ISL_268665 | A/Mal/NL-Mastenbroek/16015378-002/2016 | A / H5N8 | Europe / Netherlands / Provincie Overijssel / Mastenbroek | Saskia Bergervoet (Wageningen Bioveterinary Research) |
| EPI_ISL_268666 | A/P_falcon/NL-Vrouwenpolder (Zeeland)/16015510-001/2016 | A / H5N8 | Europe / Netherlands / Provincie Zeeland / Vrouwenpolder | Saskia Bergervoet (Wageningen Bioveterinary Research) |
| EPI_ISL_268667 | A/Sea_eagle/NL-Assen/16015398-002/2016 | A / H5N8 | Europe / Netherlands / Provincie Drenthe / Gemeente Assen | Saskia Bergervoet (Wageningen Bioveterinary Research) |
| EPI_ISL_268668 | A/T_Dk/NL-Almeerder Zand/16014341-003/2016 | A / H5N8 | Europe / Netherlands / Provincie Flevoland / Almeerder Zand | Saskia Bergervoet (Wageningen Bioveterinary Research) |
| EPI_ISL_268669 | A/T_Dk/NL-Monnickendam/16013865-006-008/2016 | A / H5N8 | Europe / Netherlands / Provincie Noord-Holland / Monnickendam | Saskia Bergervoet (Wageningen Bioveterinary Research) |
| EPI_ISL_268670 | A/T_Dk/NL-Roggebotsluis/16014462-015/2016 | A / H5N8 | Europe / Netherlands / Provincie Flevoland / Roggebotsluis | Saskia Bergervoet (Wageningen Bioveterinary Research) |
| EPI_ISL_268671 | A/T_Dk/NL-Rotterdam/16014155-001/2016 | A / H5N8 | Europe / Netherlands / South Holland / Gemeente Rotterdam | Saskia Bergervoet (Wageningen Bioveterinary Research) |
| EPI_ISL_268672 | A/T_Dk/NL-Werkendam/16014159-002/2016 | A / H5N8 | Europe / Netherlands / North Brabant / Gemeente Werkendam | Saskia Bergervoet (Wageningen Bioveterinary Research) |
| EPI_ISL_268673 | A/T_Dk/NL-Werkendam/16014159-003/2016 | A / H5N8 | Europe / Netherlands / North Brabant / Gemeente Werkendam | Saskia Bergervoet (Wageningen Bioveterinary Research) |
| EPI_ISL_268674 | A/T_Dk/NL-Zeewolde/16013976-001/2016 | A / H5N8 | Europe / Netherlands / Provincie Flevoland / Gemeente Zeewolde | Saskia Bergervoet (Wageningen Bioveterinary Research) |
| EPI_ISL_268675 | A/T_Dk/NL-Zeewolde/16013976-001-003/2016 | A / H5N8 | Europe / Netherlands / Provincie Flevoland / Gemeente Zeewolde | Saskia Bergervoet (Wageningen Bioveterinary Research) |
| EPI_ISL_268676 | A/T_Dk/NL-Zeewolde/16013976-004/2016 | A / H5N8 | Europe / Netherlands / Provincie Flevoland / Gemeente Zeewolde | Saskia Bergervoet (Wageningen Bioveterinary Research) |
| EPI_ISL_268677 | A/T_Dk/NL-Zeewolde/16013976-004-006/2016 | A / H5N8 | Europe / Netherlands / Provincie Flevoland / Gemeente Zeewolde | Saskia Bergervoet (Wageningen Bioveterinary Research) |
| EPI_ISL_268678 | A/T_Dk/NL-Zeewolde/16013976-005/2016 | A / H5N8 | Europe / Netherlands / Provincie Flevoland / Gemeente Zeewolde | Saskia Bergervoet (Wageningen Bioveterinary Research) |
| EPI_ISL_268679 | A/T_Dk/NL-Zeewolde/16013976-006/2016 | A / H5N8 | Europe / Netherlands / Provincie Flevoland / Gemeente Zeewolde | Saskia Bergervoet (Wageningen Bioveterinary Research) |
| EPI_ISL_268680 | A/T_Dk/NL-Zuidoost Beemster/16014148-002/2016 | A / H5N8 | Europe / Netherlands / Provincie Noord-Holland / Zuidoost Beemster | Saskia Bergervoet (Wageningen Bioveterinary Research) |
| EPI_ISL_268681 | A/T_Dk/NL-Zuidoost Beemster/16014148-009/2016 | A / H5N8 | Europe / Netherlands / Provincie Noord-Holland / Zuidoost Beemster | Saskia Bergervoet (Wageningen Bioveterinary Research) |
| EPI_ISL_268682 | A/Teal/NL-Ferwert/16015273-013/2016 | A / H5N8 | Europe / Netherlands / Provincie Friesland / Ferwert | Saskia Bergervoet (Wageningen Bioveterinary Research) |
| EPI_ISL_268799 | A/Back-headed_Gull/Netherlands/8/2016 | A / H5N8 | Europe / Netherlands | Maria Johanna Poen (Erasmus Medical Center / Department of Virology) |
| EPI_ISL_268800 | A/Black-headed_Gull/Netherlands/17/2016 | A / H5N8 | Europe / Netherlands | Maria Johanna Poen (Erasmus Medical Center / Department of Virology) |
| EPI_ISL_268866 | A/Back-headed_Gull/Netherlands/9/2016 | A / H5N8 | Europe / Netherlands | Maria Johanna Poen (Erasmus Medical Center / Department of Virology) |
| EPI_ISL_268916 | A/Caspian _Gull/Netherlands/1/2016 | A / H5N8 | Europe / Netherlands | Maria Johanna Poen (Erasmus Medical Center / Department of Virology) |
| EPI_ISL_268927 | A/Common_Buzzard/Netherlands/1/2016 | A / H5N8 | Europe / Netherlands | Maria Johanna Poen (Erasmus Medical Center / Department of Virology) |
| EPI_ISL_268928 | A/mute swan/Czech Republic/54-17_1/2017 (H5N8) | A / H5N8 | Europe / Czech Republic / South Moravian Region / Mesto Brno / ??m?ice u Ivan?ic | Alexander Nagy (State Veterinary Institute Prague) |
| EPI_ISL_268929 | A/Common_Eider/Netherlands/2/2016 | A / H5N8 | Europe / Netherlands | Maria Johanna Poen (Erasmus Medical Center / Department of Virology) |
| EPI_ISL_268930 | A/goose/Czech Republic/136-17_1/2017 (H5N8) | A / H5N8 | Europe / Czech Republic / South Moravian Region / Mesto Brno / Ivan?ice-N?m?ice | Alexander Nagy (State Veterinary Institute Prague) |
| EPI_ISL_268931 | A/mallard/Czech Republic/136-17_2/2017 (H5N8) | A / H5N8 | Europe / Czech Republic / South Moravian Region / Mesto Brno / Ivan?ice-N?m?ice | Alexander Nagy (State Veterinary Institute Prague) |

| | | | | |
|---|---|---|---|---|
| EPI_ISL_268932 | A/goose/Czech Republic/197-17/2017 (H5N8) | A / H5N8 | Europe / Czech Republic / Hlavni mesto Praha / Okres Praha / Lázn? Touše? | Alexander Nagy (State Veterinary Institute Prague) |
| EPI_ISL_268933 | A/chicken/Czech Republic/508-17_1/2017 (H5N8) | A / H5N8 | Europe / Czech Republic / Jihocesky Kraj / Okres Tabor / Chýnov | Alexander Nagy (State Veterinary Institute Prague) |
| EPI_ISL_268934 | A/mallard/Czech Republic/508-17_4/2017 (H5N8) | A / H5N8 | Europe / Czech Republic / Jihocesky Kraj / Okres Tabor / Chýnov | Alexander Nagy (State Veterinary Institute Prague) |
| EPI_ISL_268935 | A/mute swan/Czech Republic/572-17_3/2017 (H5N8) | A / H5N8 | Europe / Czech Republic / Hlavni mesto Praha / Okres Praha / Prague 1, Alšovo náb?eží | Alexander Nagy (State Veterinary Institute Prague) |
| EPI_ISL_268936 | A/mute swan/Czech Republic/964-17/2017 (H5N8) | A / H5N8 | Europe / Czech Republic / Stredocesky Kraj / Okres Kolin | Alexander Nagy (State Veterinary Institute Prague) |
| EPI_ISL_268937 | A/Eurasian_Wigeon/Netherlands/10/2016 | A / H5N8 | Europe / Netherlands | Maria Johanna Poen (Erasmus Medical Center / Department of Virology) |
| EPI_ISL_268938 | A/mute swan/Czech Republic/967-17/2017 (H5N8) | A / H5N8 | Europe / Czech Republic / Kralovehradecky Kraj / Okres Hradec Kralove / Tylovo náb?eží | Alexander Nagy (State Veterinary Institute Prague) |
| EPI_ISL_268939 | A/mute swan/Czech Republic/987-17_2/2017 (H5N8) | A / H5N8 | Europe / Czech Republic / Hlavni mesto Praha / Okres Praha / Prague 1, p?ístavišt? Four seasons | Alexander Nagy (State Veterinary Institute Prague) |
| EPI_ISL_268940 | A/mute swan/Czech Republic/1058-17/2017 (H5N8) | A / H5N8 | Europe / Czech Republic / Jihocesky Kraj / Okres Ceske Budejovice / ?eské Bud?jovice, 48°59'24.( | Alexander Nagy (State Veterinary Institute Prague) |
| EPI_ISL_268941 | A/mute swan/Czech Republic/1060-17/2017 (H5N8) | A / H5N8 | Europe / Czech Republic / Jihocesky Kraj / Okres Ceske Budejovice / ?eské Bud?jovice | Alexander Nagy (State Veterinary Institute Prague) |
| EPI_ISL_268942 | A/mute swan/Czech Republic/1155-17/2017 (H5N8) | A / H5N8 | Europe / Czech Republic / Jihocesky Kraj / Okres Ceske Budejovice / ?eské Bud?jovice | Alexander Nagy (State Veterinary Institute Prague) |
| EPI_ISL_268943 | A/mute swan/Czech Republic/1156-17/2017 (H5N8) | A / H5N8 | Europe / Czech Republic / Jihocesky Kraj / Okres Pisek / Písek | Alexander Nagy (State Veterinary Institute Prague) |
| EPI_ISL_268944 | A/mute swan/Czech Republic/1170-17_2/2017 (H5N8) | A / H5N8 | Europe / Czech Republic / Olomoucky Kraj / Okres Jesenik / Javorník-ves | Alexander Nagy (State Veterinary Institute Prague) |
| EPI_ISL_268945 | A/mute swan/Czech Republic/1171-17/2017 (H5N8) | A / H5N8 | Europe / Czech Republic / Morovskoslezsky Kraj / Okres Opava / Jilešovice | Alexander Nagy (State Veterinary Institute Prague) |
| EPI_ISL_268946 | A/chicken/Czech Republic/1208-17_1/2017 (H5N8) | A / H5N8 | Europe / Czech Republic / Liberecky Kraj / Okres Semily / Turnov | Alexander Nagy (State Veterinary Institute Prague) |
| EPI_ISL_268947 | A/mallard/Czech Republic/1219-17_1/2017 (H5N8) | A / H5N8 | Europe / Czech Republic / Jihocesky Kraj / Okres Strakonice / Blatná | Alexander Nagy (State Veterinary Institute Prague) |
| EPI_ISL_268948 | A/mallard/Czech Republic/1226-17/2017 (H5N8) | A / H5N8 | Europe / Czech Republic / Zlinsky Kraj / Okres Zlin / Otrokovice | Alexander Nagy (State Veterinary Institute Prague) |
| EPI_ISL_268949 | A/mute swan/Czech Republic/1227-17/2017 (H5N8) | A / H5N8 | Europe / Czech Republic / Zlinsky Kraj / Okres Uherske Hradiste / Uherské Hradišt? | Alexander Nagy (State Veterinary Institute Prague) |
| EPI_ISL_268950 | A/mute swan/Czech Republic/1296-17_1/2017 (H5N8) | A / H5N8 | Europe / Czech Republic / Olomoucky Kraj / Okres Prerov / Tova?ov Lakes | Alexander Nagy (State Veterinary Institute Prague) |
| EPI_ISL_268951 | A/mute swan/Czech Republic/1330-17_1/2017 (H5N8) | A / H5N8 | Europe / Czech Republic / Jihocesky Kraj / Okres Tabor / Roudná | Alexander Nagy (State Veterinary Institute Prague) |
| EPI_ISL_268952 | A/mute swan/Czech Republic/1331-17_1/2017 (H5N8) | A / H5N8 | Europe / Czech Republic / Jihocesky Kraj / Okres Tabor / Tu?apy | Alexander Nagy (State Veterinary Institute Prague) |
| EPI_ISL_268953 | A/mute swan/Czech Republic/1337-17/2017 (H5N8) | A / H5N8 | Europe / Czech Republic / Jihocesky Kraj / Okres Tabor / Veselí nad Lužnicí | Alexander Nagy (State Veterinary Institute Prague) |
| EPI_ISL_268954 | A/mute swan/Czech Republic/1339-17/2017 (H5N8) | A / H5N8 | Europe / Czech Republic / Morovskoslezsky Kraj / Okres Opava / Opava | Alexander Nagy (State Veterinary Institute Prague) |
| EPI_ISL_268955 | A/chicken/Czech Republic/1344-17/2017 (H5N8) | A / H5N8 | Europe / Czech Republic / Morovskoslezsky Kraj / Okres Ostrava-Mesto / Poruba | Alexander Nagy (State Veterinary Institute Prague) |
| EPI_ISL_268956 | A/mute swan/Czech Republic/1461-17/2017 (H5N8) | A / H5N8 | Europe / Czech Republic / Jihocesky Kraj / Okres Strakonice / Katovice | Alexander Nagy (State Veterinary Institute Prague) |
| EPI_ISL_268957 | A/chicken/Czech Republic/1465-17/2017 (H5N8) | A / H5N8 | Europe / Czech Republic / Jihocesky Kraj / Okres Ceske Budejovice / Ledenice | Alexander Nagy (State Veterinary Institute Prague) |
| EPI_ISL_268958 | A/mute swan/Czech Republic/1519-17/2017 (H5N8) | A / H5N8 | Europe / Czech Republic / Kralovehradecky Kraj / Okres Hradec Kralove / písník Kosi?ka | Alexander Nagy (State Veterinary Institute Prague) |
| EPI_ISL_268959 | A/mute swan/Czech Republic/1576-17_C/2017 (H5N8) | A / H5N8 | Europe / Czech Republic / Karlovarsky Kraj / Okres Cheb / Skalka pond | Alexander Nagy (State Veterinary Institute Prague) |
| EPI_ISL_268960 | A/mallard/Czech Republic/1577-17/2017 (H5N8) | A / H5N8 | Europe / Czech Republic / Jihocesky Kraj / Okres Jindrichuv Hradec / Zbuzany, near the bridge ac | Alexander Nagy (State Veterinary Institute Prague) |
| EPI_ISL_268961 | A/mute swan/Czech Republic/1640-17/2017 (H5N8) | A / H5N8 | Europe / Czech Republic / Ustecky Kraj / Okres Chomutov / Kada?, the Oh?e river, near the railw | Alexander Nagy (State Veterinary Institute Prague) |
| EPI_ISL_268962 | A/mallard/Czech Republic/1672-17/2017 (H5N8) | A / H5N8 | Europe / Czech Republic / Jihocesky Kraj / Okres Jindrichuv Hradec / Lásenice, the Nežárka river | Alexander Nagy (State Veterinary Institute Prague) |
| EPI_ISL_268963 | A/chicken/Czech Republic/1675-17_2/2017 (H5N8) | A / H5N8 | Europe / Czech Republic / Hlavni mesto Praha / Okres Praha / Kostelec nad Labem | Alexander Nagy (State Veterinary Institute Prague) |
| EPI_ISL_268964 | A/grey heron/Czech Republic/1680-17_2/2017 (H5N8) | A / H5N8 | Europe / Czech Republic / Jihocesky Kraj / Okres Jindrichuv Hradec / T?ebo?, Zlatá stoka | Alexander Nagy (State Veterinary Institute Prague) |
| EPI_ISL_268965 | A/Indian Runner Duck/Czech Republic/1683-17_1/2017 (H5N8 | A / H5N8 | Europe / Czech Republic / Liberecky Kraj / Okres Liberec / Liberec | Alexander Nagy (State Veterinary Institute Prague) |
| EPI_ISL_268966 | A/chicken/Czech Republic/1687-17_2/2017 (H5N8) | A / H5N8 | Europe / Czech Republic / Morovskoslezsky Kraj / Okres Ostrava-Mesto / Orlová-Olmovec | Alexander Nagy (State Veterinary Institute Prague) |
| EPI_ISL_268967 | A/chicken/Czech Republic/1688-17_1/2017 (H5N8) | A / H5N8 | Europe / Czech Republic / Stredocesky Kraj / Okres Beroun / Hlásná T?ebá? | Alexander Nagy (State Veterinary Institute Prague) |
| EPI_ISL_268968 | A/chicken/Czech Republic/1689-17/2017 (H5N8) | A / H5N8 | Europe / Czech Republic / Pardubicky kraj / Okres Usti nad Orlici / Koldín-Hradišt? | Alexander Nagy (State Veterinary Institute Prague) |
| EPI_ISL_268969 | A/mallard/Czech Republic/1690-17_2/2017 (H5N8) | A / H5N8 | Europe / Czech Republic / Jihocesky Kraj / Okres Strakonice / Blatná | Alexander Nagy (State Veterinary Institute Prague) |
| EPI_ISL_268970 | A/mute swan/Czech Republic/1691-17/2017 (H5N8) | A / H5N8 | Europe / Czech Republic / Pardubicky kraj / Okres Pardubice / Pardubice, lake Bajkal | Alexander Nagy (State Veterinary Institute Prague) |
| EPI_ISL_268971 | A/bronze turkey/Czech Republic/1755-17_1/2017 (H5N8) | A / H5N8 | Europe / Czech Republic / Hlavni mesto Praha / Okres Praha / Touše? | Alexander Nagy (State Veterinary Institute Prague) |
| EPI_ISL_268972 | A/turkey/Czech Republic/1767-17_2/2017 (H5N8) | A / H5N8 | Europe / Czech Republic / Jihocesky Kraj / Okres Jindrichuv Hradec / Horní Lhota, Stráž nad Nežá | Alexander Nagy (State Veterinary Institute Prague) |
| EPI_ISL_268973 | A/mute swan/Czech Republic/1848-17_1/2017 (H5N8) | A / H5N8 | Europe / Czech Republic / Karlovarsky Kraj / Okres Karlovy Vary / Karlovy Vary, the Oh?e river | Alexander Nagy (State Veterinary Institute Prague) |
| EPI_ISL_268974 | A/mute swan/Czech Republic/1848-17_2/2017 (H5N8) | A / H5N8 | Europe / Czech Republic / Karlovarsky Kraj / Okres Karlovy Vary / Karlovy Vary, the Oh?e river | Alexander Nagy (State Veterinary Institute Prague) |
| EPI_ISL_268975 | A/chicken/Czech Republic/1953-17/2017 (H5N8) | A / H5N8 | Europe / Czech Republic / Liberecky Kraj / Okres Ceska Lipa / Velký Grunov | Alexander Nagy (State Veterinary Institute Prague) |
| EPI_ISL_268976 | A/goose/Czech Republic/1954-17/2017 (H5N8) | A / H5N8 | Europe / Czech Republic / Pardubicky kraj / Okres Pardubice / Chvaletice | Alexander Nagy (State Veterinary Institute Prague) |
| EPI_ISL_268977 | A/goose/Czech Republic/1998-17_1/2017 (H5N8) | A / H5N8 | Europe / Czech Republic / Kralovehradecky Kraj / Okres Hradec Kralove / Ž?ár nad Orlicí | Alexander Nagy (State Veterinary Institute Prague) |
| EPI_ISL_268978 | A/mute swan/Czech Republic/2008-17_1/2017 (H5N8) | A / H5N8 | Europe / Czech Republic / Morovskoslezsky Kraj / Okres Karvina / Závada nad Olší | Alexander Nagy (State Veterinary Institute Prague) |
| EPI_ISL_268979 | A/quail/Czech Republic/2063-17_1/2017 (H5N8) | A / H5N8 | Europe / Czech Republic / Jihocesky Kraj / Okres Strakonice / Volyn? | Alexander Nagy (State Veterinary Institute Prague) |
| EPI_ISL_268980 | A/chicken/Czech Republic/2216-17_1/2017 (H5N8) | A / H5N8 | Europe / Czech Republic / Jihocesky Kraj / Okres Strakonice / Volyn? | Alexander Nagy (State Veterinary Institute Prague) |
| EPI_ISL_268981 | A/spot-billed pelican/Czech Republic/2270-17/2017 (H5N5) | A / H5N5 | Europe / Czech Republic / Liberecky Kraj / Okres Liberec / ZOO Liberec | Alexander Nagy (State Veterinary Institute Prague) |
| EPI_ISL_268982 | A/chicken/Czech Republic/2514-17/2017 (H5N8) | A / H5N8 | Europe / Czech Republic / Morovskoslezsky Kraj / Okres Karvina / Doubrava | Alexander Nagy (State Veterinary Institute Prague) |
| EPI_ISL_268983 | A/mallard/Czech Republic/2641-17/2017 (H5N8) | A / H5N8 | Europe / Czech Republic / Vysocina Kraj / Okres Pelhrimov / Želiv | Alexander Nagy (State Veterinary Institute Prague) |
| EPI_ISL_268984 | A/chicken/Czech Republic/2677-17_2/2017 (H5N8) | A / H5N8 | Europe / Czech Republic / Kralovehradecky Kraj / Okres Hradec Kralove / Bernartice | Alexander Nagy (State Veterinary Institute Prague) |
| EPI_ISL_268985 | A/mallard/Czech Republic/2678-17_1/2017 (H5N8) | A / H5N8 | Europe / Czech Republic / Vysocina Kraj / Okres Havlickuv Brod / Havli?k?v Brod | Alexander Nagy (State Veterinary Institute Prague) |
| EPI_ISL_268986 | A/mallard/Czech Republic/2820-17_1/2017 (H5N8) | A / H5N8 | Europe / Czech Republic / Karlovarsky Kraj / Okres Cheb / Poustka | Alexander Nagy (State Veterinary Institute Prague) |
| EPI_ISL_268987 | A/mallard/Czech Republic/2820-17_2/2017 (H5N8) | A / H5N8 | Europe / Czech Republic / Karlovarsky Kraj / Okres Cheb / Poustka | Alexander Nagy (State Veterinary Institute Prague) |
| EPI_ISL_268988 | A/mallard/Czech Republic/2821-17_1/2017 (H5N8) | A / H5N8 | Europe / Czech Republic / Karlovarsky Kraj / Okres Cheb / Klest | Alexander Nagy (State Veterinary Institute Prague) |
| EPI_ISL_268989 | A/chicken/Czech Republic/2821-17_2/2017 (H5N8) | A / H5N8 | Europe / Czech Republic / Karlovarsky Kraj / Okres Cheb / Klest | Alexander Nagy (State Veterinary Institute Prague) |
| EPI_ISL_268990 | A/mallard/Czech Republic/2822-17_1/2017 (H5N8) | A / H5N8 | Europe / Czech Republic / Karlovarsky Kraj / Okres Cheb / Klest | Alexander Nagy (State Veterinary Institute Prague) |
| EPI_ISL_268991 | A/chicken/Czech Republic/2822-17_2/2017 (H5N8) | A / H5N8 | Europe / Czech Republic / Karlovarsky Kraj / Okres Cheb / Klest | Alexander Nagy (State Veterinary Institute Prague) |
| EPI_ISL_268992 | A/chicken/Czech Republic/3507-17/2017 (H5N8) | A / H5N8 | Europe / Czech Republic / Morovskoslezsky Kraj / Okres Frydek-Mistek / Dobrá | Alexander Nagy (State Veterinary Institute Prague) |
| EPI_ISL_268993 | A/chicken/Czech Republic/4231-17_2/2017 (H5N8) | A / H5N7 | Europe / Czech Republic / Karlovarsky Kraj / Okres Karlovy Vary / Otro?ín | Alexander Nagy (State Veterinary Institute Prague) |
| EPI_ISL_269591 | A/Eurasian_Wigeon/Netherlands/6/2016 | A / H5N8 | Europe / Netherlands | Maria Johanna Poen (Erasmus Medical Center / Department of Virology) |
| EPI_ISL_269592 | A/Eurasian_Wigeon/Netherlands/8/2016 | A / H5N8 | Europe / Netherlands | Maria Johanna Poen (Erasmus Medical Center / Department of Virology) |
| EPI_ISL_269593 | A/Eurasian_Wigeon/Netherlands/11/2016 | A / H5N8 | Europe / Netherlands | Maria Johanna Poen (Erasmus Medical Center / Department of Virology) |
| EPI_ISL_269594 | A/Eurasian_Wigeon/Netherlands/22/2016 | A / H5N8 | Europe / Netherlands | Maria Johanna Poen (Erasmus Medical Center / Department of Virology) |
| EPI_ISL_269595 | A/Eurasian_Wigeon/Netherlands/12/2016 | A / H5N8 | Europe / Netherlands | Maria Johanna Poen (Erasmus Medical Center / Department of Virology) |
| EPI_ISL_269596 | A/Eurasian_Wigeon/Netherlands/13/2016 | A / H5N8 | Europe / Netherlands | Maria Johanna Poen (Erasmus Medical Center / Department of Virology) |
| EPI_ISL_269597 | A/Great_Black-backed_Gull/Netherlands/1/2016 | A / H5N8 | Europe / Netherlands | Maria Johanna Poen (Erasmus Medical Center / Department of Virology) |
| EPI_ISL_269598 | A/Great_Black-backed_Gull/Netherlands/3/2016 | A / H5N8 | Europe / Netherlands | Maria Johanna Poen (Erasmus Medical Center / Department of Virology) |
| EPI_ISL_269599 | A/Great_Black-backed_Gull/Netherlands/4/2016 | A / H5N8 | Europe / Netherlands | Maria Johanna Poen (Erasmus Medical Center / Department of Virology) |
| EPI_ISL_269600 | A/Great_Crested_Grebe/Netherlands/2/2016 | A / H5N8 | Europe / Netherlands | Maria Johanna Poen (Erasmus Medical Center / Department of Virology) |

| | | | | |
|---|---|---|---|---|
| EPI_ISL_269601 | A/Eurasian_Herring_Gull/Netherlands/2/2016 | A / H5N8 | Europe / Netherlands | Maria Johanna Poen (Erasmus Medical Center / Department of Virology) |
| EPI_ISL_269602 | A/Lesser_Black-backed_Gull/Netherlands/1/2016 | A / H5N8 | Europe / Netherlands | Maria Johanna Poen (Erasmus Medical Center / Department of Virology) |
| EPI_ISL_269603 | A/Mallard/Netherlands/51/2016 | A / H5N8 | Europe / Netherlands | Maria Johanna Poen (Erasmus Medical Center / Department of Virology) |
| EPI_ISL_269604 | A/Mallard/Netherlands/1/2017 | A / H5N8 | Europe / Netherlands | Maria Johanna Poen (Erasmus Medical Center / Department of Virology) |
| EPI_ISL_269692 | A/Mallard/Netherlands/3/2017 | A / H5N8 | Europe / Netherlands | Maria Johanna Poen (Erasmus Medical Center / Department of Virology) |
| EPI_ISL_269693 | A/Common_Pochard/Netherlands/1/2016 | A / H5N8 | Europe / Netherlands | Maria Johanna Poen (Erasmus Medical Center / Department of Virology) |
| EPI_ISL_269694 | A/Eurasian_Wigeon/Netherlands/1/2016 | A / H5N8 | Europe / Netherlands | Maria Johanna Poen (Erasmus Medical Center / Department of Virology) |
| EPI_ISL_269695 | A/Eurasian_Wigeon/Netherlands/21/2016 | A / H5N8 | Europe / Netherlands | Maria Johanna Poen (Erasmus Medical Center / Department of Virology) |
| EPI_ISL_269696 | A/Eurasian_Wigeon/Netherlands/23/2016 | A / H5N8 | Europe / Netherlands / Provincie Noord-Holland | Maria Johanna Poen (Erasmus Medical Center / Department of Virology) |
| EPI_ISL_269703 | A/Eurasian_Wigeon/Netherlands/25/2016 | A / H5N8 | Europe / Netherlands | Maria Johanna Poen (Erasmus Medical Center / Department of Virology) |
| EPI_ISL_270356 | A/common teal/Shanghai/PD1108-3/2013 | A / H5N8 | Asia / China | |
| EPI_ISL_270357 | A/common teal/Shanghai/PD1108-6/2013 | A / H5N8 | Asia / China | |
| EPI_ISL_270358 | A/common teal/Shanghai/PD1108-13/2013 | A / H5N8 | Asia / China | |
| EPI_ISL_270359 | A/common teal/Shanghai/PD1108-20/2013 | A / H5N8 | Asia / China | |
| EPI_ISL_270360 | A/common teal/Shanghai/PD1108-24/2013 | A / H5N8 | Asia / China | |
| EPI_ISL_270361 | A/common teal/Shanghai/PD1108-25/2013 | A / H5N8 | Asia / China | |
| EPI_ISL_270362 | A/common teal/Shanghai/PD1108-27/2013 | A / H5N8 | Asia / China | |
| EPI_ISL_270363 | A/common teal/Shanghai/PD1118-1/2013 | A / H5N8 | Asia / China | |
| EPI_ISL_270364 | A/common teal/Shanghai/PD1121-10/2013 | A / H5N8 | Asia / China | |
| EPI_ISL_270365 | A/common teal/Shanghai/PD1121-15/2013 | A / H5N8 | Asia / China | |
| EPI_ISL_270366 | A/common teal/Shanghai/PD1121-19/2013 | A / H5N8 | Asia / China | |
| EPI_ISL_270367 | A/common teal/Shanghai/PD1202-9/2013 | A / H5N8 | Asia / China | |
| EPI_ISL_270368 | A/common teal/Shanghai/PD1121-18/2013 | A / H5N8 | Asia / China | |
| EPI_ISL_270369 | A/mallard/Shanghai/SH-9/2013 | A / H5N8 | Asia / China | |
| EPI_ISL_270370 | A/falcated duck/Shanghai/PD1121-26/2013 | A / H5N8 | Asia / China | |
| EPI_ISL_270371 | A/falcated duck/Shanghai/PD1121-27/2013 | A / H5N8 | Asia / China | |
| EPI_ISL_270372 | A/spot-billed duck/Shanghai/PD1202-3/2013 | A / H5N8 | Asia / China | |
| EPI_ISL_2713170 | A/tundra swan/Hubei/BQ9/2020 | A / H5N8 | Asia / China / Hubei Province | Hongliang Chai (Northeast Forestry University / College of Wildlife Resources) |
| EPI_ISL_2713314 | A/whiskered tern/Hubei/BQ10/2020 | A / H5N8 | Asia / China / Hubei Province | Hongliang Chai (Northeast Forestry University / College of Wildlife Resources) |
| EPI_ISL_2713372 | A/bean goose/Hubei/BQ11/2020 | A / H5N8 | Asia / China / Hubei Province | Hongliang Chai (Northeast Forestry University / College of Wildlife Resources) |
| EPI_ISL_271704 | A/Goose/Hungary/17261/2017 | A / H5N8 | Europe / Hungary / Bacs-Kiskun | Adam Dan (Danam.Vet.Molbiol) |
| EPI_ISL_271705 | A/Goose/Hungary/15729/2017 | A / H5N8 | Europe / Hungary / Bacs-Kiskun | Adam Dan (Danam.Vet.Molbiol) |
| EPI_ISL_271706 | A/Goose/Hungary/17051/2017 | A / H5N8 | Europe / Hungary / Bacs-Kiskun | Adam Dan (Danam.Vet.Molbiol) |
| EPI_ISL_271707 | A/Goose/Hungary/17580/2017 | A / H5N8 | Europe / Hungary / Bacs-Kiskun | Adam Dan (Danam.Vet.Molbiol) |
| EPI_ISL_271708 | A/Goose/Hungary/17985/2017 | A / H5N8 | Europe / Hungary / Bacs-Kiskun | Adam Dan (Danam.Vet.Molbiol) |
| EPI_ISL_271709 | A/Goose/Hungary/59763/2016 | A / H5N8 | Europe / Hungary / Bacs-Kiskun | Adam Dan (Danam.Vet.Molbiol) |
| EPI_ISL_271710 | A/Mulard_duck/Hungary/60369/2016 | A / H5N8 | Europe / Hungary / Bacs-Kiskun | Adam Dan (Danam.Vet.Molbiol) |
| EPI_ISL_271711 | A/Mulard_duck/Hungary/62902/2016 | A / H5N8 | Europe / Hungary / Csongrad megye / Csongrad | Adam Dan (Danam.Vet.Molbiol) |
| EPI_ISL_271712 | A/Goose/Hungary/63743/2016 | A / H5N8 | Europe / Hungary / Bacs-Kiskun | Adam Dan (Danam.Vet.Molbiol) |
| EPI_ISL_271713 | A/Goose/Hungary/64909/2016 | A / H5N8 | Europe / Hungary / Bacs-Kiskun | Adam Dan (Danam.Vet.Molbiol) |
| EPI_ISL_271714 | A/Mulard_duck/Hungary/59163/2016 | A / H5N8 | Europe / Hungary / Bacs-Kiskun | Adam Dan (Danam.Vet.Molbiol) |
| EPI_ISL_272740 | A/goose/Guangdong/QY01/2016 | A / H5N6 | Asia / China | |
| EPI_ISL_272859 | A/grey heron/W779/2017 | A / H5N8 | Asia / Korea, Republic of | |
| EPI_ISL_273359 | A/duck/Vietnam/LBM835/2015 | A / H5N6 | Asia / Vietnam | |
| EPI_ISL_273360 | A/duck/Vietnam/LBM806/2015 | A / H5N6 | Asia / Vietnam | |
| EPI_ISL_273846 | A/chicken/Italy/17VIR3078/2017 | A / H5N8 | Europe / Italy / Emilia-Romagna / Province of Bologna / Mordano | Bianca Zecchin (Istituto Zooprofilattico Sperimentale Delle Venezie) |
| EPI_ISL_273847 | A/turkey/Italy/17VIR5878-3/2017 | A / H5N8 | Europe / Italy / Veneto / Province of Verona / Albaredo d'Adige | Bianca Zecchin (Istituto Zooprofilattico Sperimentale Delle Venezie) |
| EPI_ISL_274858 | A/mute swan/Kaliningrad/132/2017 | A / H5N8 | Europe / Russian Federation / Kaliningrad Oblast | Ivan Susloparov (State Research Center of Virology and Biotechnology (VECTOR) / Emerging Zoonotic Diseases and Influenza) |
| EPI_ISL_275283 | A/chicken/Shchyolkovo/47/2017 | A / H5N8 | Europe / Russian Federation / Moscow Oblast | Ivan Susloparov (State Research Center of Virology and Biotechnology (VECTOR) / Emerging Zoonotic Diseases and Influenza) |
| EPI_ISL_275287 | A/chicken/Rostov-on-Don/44/2017 | A / H5N8 | Europe / Russian Federation / Rostov Oblast | Ivan Susloparov (State Research Center of Virology and Biotechnology (VECTOR) / Emerging Zoonotic Diseases and Influenza) |
| EPI_ISL_275288 | A/chicken/Tatarstan/88/2017 | A / H5N8 | Europe / Russian Federation / Republic of Tatarstan | Ivan Susloparov (State Research Center of Virology and Biotechnology (VECTOR) / Emerging Zoonotic Diseases and Influenza) |
| EPI_ISL_277035 | A/chicken/Wuxi/6074/2015 | A / H5N2 | Asia / China / Jiangsu | Maijuan Ma (Beijing Institute of Microbiology and Epidemiology / State Key Laboratory of Pathogen and Biosecurity) |
| EPI_ISL_277036 | A/duck/Wuxi/6462/2015 | A / H5N2 | Asia / China / Jiangsu | Maijuan Ma (Beijing Institute of Microbiology and Epidemiology / State Key Laboratory of Pathogen and Biosecurity) |
| EPI_ISL_277037 | A/duck/Wuxi/6466/2015 | A / H5N2 | Asia / China / Jiangsu | Maijuan Ma (Beijing Institute of Microbiology and Epidemiology / State Key Laboratory of Pathogen and Biosecurity) |
| EPI_ISL_277040 | A/chicken/Wuxi/7157/2015 | A / H5N6 | Asia / China / Jiangsu | Maijuan Ma (Beijing Institute of Microbiology and Epidemiology / State Key Laboratory of Pathogen and Biosecurity) |
| EPI_ISL_277041 | A/duck/Wuxi/7249/2015 | A / H5N6 | Asia / China / Jiangsu | Maijuan Ma (Beijing Institute of Microbiology and Epidemiology / State Key Laboratory of Pathogen and Biosecurity) |
| EPI_ISL_277042 | A/chicken/Wuxi/7765/2016 | A / H5N6 | Asia / China / Jiangsu | Maijuan Ma (Beijing Institute of Microbiology and Epidemiology / State Key Laboratory of Pathogen and Biosecurity) |
| EPI_ISL_278025 | A/Duck/Zhanjiang/15577/2015 | A / H5N6 | Asia / China / Guangdong Province / Zhanjiang | Jia Weixin (South China Agricultural University / Veterinary Medicine College) |
| EPI_ISL_278026 | A/Chicken/Huizhou/16274/2016(H5N6) | A / H5N6 | Asia / China / Guangdong Province / Huizhou | Jia Weixin (South China Agricultural University / Veterinary Medicine College) |
| EPI_ISL_278027 | A/Goose/Qingyuan/16875/2016 | A / H5N6 | Asia / China / Guangdong Province / Qingyuan | Jia Weixin (South China Agricultural University / Veterinary Medicine College) |
| EPI_ISL_278031 | A/Quail/Zhanjiang/16887/2016 | A / H5N6 | Asia / China / Guangdong Province / Zhanjiang | Jia Weixin (South China Agricultural University / Veterinary Medicine College) |
| EPI_ISL_278040 | A/peregrine falcon/HK/4955/2015 | A / H5N6 | Asia / Hong Kong (SAR) | |
| EPI_ISL_278041 | A/oriental magpie robin/HK/6154/2015 | A / H5N6 | Asia / Hong Kong (SAR) | |
| EPI_ISL_278200 | A/Brahma chicken/Belgium/6153/2017 | A / H5N8 | Europe / Belgium | |
| EPI_ISL_278201 | A/chicken/Belgium/5990/2017 | A / H5N8 | Europe / Belgium | |
| EPI_ISL_278204 | A/cat/Zhejiang/C1/2016 | A / H5N6 | Asia / China | |
| EPI_ISL_278205 | A/cat/Zhejiang/C2/2016 | A / H5N6 | Asia / China | |
| EPI_ISL_279010 | A/crane/Kagoshima/KU-6/2016 | A / H5N6 | Asia / Japan / Kagoshima | Makoto Ozawa (Kagoshima University ) |
| EPI_ISL_279011 | A/crane/Kagoshima/KU-8/2016 | A / H5N6 | Asia / Japan / Kagoshima | Makoto Ozawa (Kagoshima University ) |
| EPI_ISL_279012 | A/crane/Kagoshima/KU-9/2016 | A / H5N6 | Asia / Japan / Kagoshima | Makoto Ozawa (Kagoshima University ) |
| EPI_ISL_279013 | A/duck/Kagoshima/KU-d35/2016 | A / H5N6 | Asia / Japan / Kagoshima | Makoto Ozawa (Kagoshima University ) |
| EPI_ISL_279014 | A/duck/Kagoshima/KU-d42/2016 | A / H5N6 | Asia / Japan / Kagoshima | Makoto Ozawa (Kagoshima University ) |

| | | | |
|---|---|---|---|
| EPI_ISL_279015 | A/duck/Kagoshima/KU-d66/2016 | A / H5N6 Asia / Japan / Kagoshima | Makoto Ozawa (Kagoshima University ) |
| EPI_ISL_279016 | A/duck/Kagoshima/KU-d79/2016 | A / H5N6 Asia / Japan / Kagoshima | Makoto Ozawa (Kagoshima University ) |
| EPI_ISL_279017 | A/crane/Kagoshima/KU-10/2016 | A / H5N6 Asia / Japan / Kagoshima | Makoto Ozawa (Kagoshima University ) |
| EPI_ISL_279018 | A/crane/Kagoshima/KU-12/2016 | A / H5N6 Asia / Japan / Kagoshima | Makoto Ozawa (Kagoshima University ) |
| EPI_ISL_279019 | A/crane/Kagoshima/KU-15/2016 | A / H5N6 Asia / Japan / Kagoshima | Makoto Ozawa (Kagoshima University ) |
| EPI_ISL_279020 | A/crane/Kagoshima/KU-16/2016 | A / H5N6 Asia / Japan / Kagoshima | Makoto Ozawa (Kagoshima University ) |
| EPI_ISL_279021 | A/northern pintail/Kagoshima/KU-23/2016 | A / H5N6 Asia / Japan / Kagoshima | Makoto Ozawa (Kagoshima University ) |
| EPI_ISL_279022 | A/crane/Kagoshima/KU-14/2016 | A / H5N6 Asia / Japan / Kagoshima | Makoto Ozawa (Kagoshima University ) |
| EPI_ISL_279023 | A/eurasian wigeon/Kagoshima/KU-20/2016 | A / H5N6 Asia / Japan / Kagoshima | Makoto Ozawa (Kagoshima University ) |
| EPI_ISL_279024 | A/eurasian wigeon/Kagoshima/KU-21/2016 | A / H5N6 Asia / Japan / Kagoshima | Makoto Ozawa (Kagoshima University ) |
| EPI_ISL_279025 | A/crane/Kagoshima/KU-25/2016 | A / H5N6 Asia / Japan / Kagoshima | Makoto Ozawa (Kagoshima University ) |
| EPI_ISL_279026 | A/crane/Kagoshima/KU-26/2016 | A / H5N6 Asia / Japan / Kagoshima | Makoto Ozawa (Kagoshima University ) |
| EPI_ISL_279027 | A/crane/Kagoshima/KU-27/2016 | A / H5N6 Asia / Japan / Kagoshima | Makoto Ozawa (Kagoshima University ) |
| EPI_ISL_279028 | A/crane/Kagoshima/KU-28/2016 | A / H5N6 Asia / Japan / Kagoshima | Makoto Ozawa (Kagoshima University ) |
| EPI_ISL_279029 | A/eurasian wigeon/Kagoshima/KU-32/2016 | A / H5N6 Asia / Japan / Kagoshima | Makoto Ozawa (Kagoshima University ) |
| EPI_ISL_279030 | A/crane/Kagoshima/KU-31/2016 | A / H5N6 Asia / Japan / Kagoshima | Makoto Ozawa (Kagoshima University ) |
| EPI_ISL_279031 | A/crane/Kagoshima/KU-34/2016 | A / H5N6 Asia / Japan / Kagoshima | Makoto Ozawa (Kagoshima University ) |
| EPI_ISL_279032 | A/crane/Kagoshima/KU-33/2016 | A / H5N6 Asia / Japan / Kagoshima | Makoto Ozawa (Kagoshima University ) |
| EPI_ISL_279033 | A/crane/Kagoshima/KU-37/2016 | A / H5N6 Asia / Japan / Kagoshima | Makoto Ozawa (Kagoshima University ) |
| EPI_ISL_279034 | A/crane/Kagoshima/KU-43/2016 | A / H5N6 Asia / Japan / Kagoshima | Makoto Ozawa (Kagoshima University ) |
| EPI_ISL_279035 | A/crane/Kagoshima/KU-44/2016 | A / H5N6 Asia / Japan / Kagoshima | Makoto Ozawa (Kagoshima University ) |
| EPI_ISL_279036 | A/crane/Kagoshima/KU-45/2016 | A / H5N6 Asia / Japan / Kagoshima | Makoto Ozawa (Kagoshima University ) |
| EPI_ISL_279037 | A/crane/Kagoshima/KU-46/2016 | A / H5N6 Asia / Japan / Kagoshima | Makoto Ozawa (Kagoshima University ) |
| EPI_ISL_279038 | A/crane/Kagoshima/KU-48/2016 | A / H5N6 Asia / Japan / Kagoshima | Makoto Ozawa (Kagoshima University ) |
| EPI_ISL_279039 | A/crane/Kagoshima/KU-52/2016 | A / H5N6 Asia / Japan / Kagoshima | Makoto Ozawa (Kagoshima University ) |
| EPI_ISL_279040 | A/crane/Kagoshima/KU-53/2016 | A / H5N6 Asia / Japan / Kagoshima | Makoto Ozawa (Kagoshima University ) |
| EPI_ISL_279178 | A/duck/Zhejiang/1026109/2015 | A / H5N2 Asia / China | |
| EPI_ISL_279179 | A/chicken/Zhejiang/81643/2015 | A / H5N2 Asia / China | |
| EPI_ISL_279180 | A/chicken/Zhejiang/7450/2015 | A / H5N2 Asia / China | |
| EPI_ISL_279181 | A/chicken/Zhejiang/514135/2015 | A / H5N2 Asia / China | |
| EPI_ISL_279424 | A/wildfowl/Shandong/SD01/2015 | A / H5N6 Asia / China | |
| EPI_ISL_279790 | A/wildfowl/Shandong/SD04/2016 | A / H5N8 Asia / China | |
| EPI_ISL_279791 | A/wildfowl/Shandong/SD02/2015 | A / H5N2 Asia / China | |
| EPI_ISL_280485 | A/common teal/Shanghai/JDS36/2016 | A / H5N6 Asia / China | |
| EPI_ISL_280486 | A/common teal/Shanghai/JDS66/2016 | A / H5N6 Asia / China | |
| EPI_ISL_280487 | A/common teal/Shanghai/JDS76/2016 | A / H5N6 Asia / China | |
| EPI_ISL_280488 | A/common teal/Shanghai/JDS81/2016 | A / H5N6 Asia / China | |
| EPI_ISL_280489 | A/Eurasian wigeon/Shanghai/JDS101/2016 | A / H5N6 Asia / China | |
| EPI_ISL_280490 | A/mandarin duck/Shanghai/PD4/2016 | A / H5N6 Asia / China | |
| EPI_ISL_280491 | A/shoveller duck/Shanghai/JDS90/2016 | A / H5N6 Asia / China | |
| EPI_ISL_280674 | A/mute_swan/Switzerland/V0244.2-L02307/2017 | A / H5N8 Europe / Switzerland / Yverdon-les-Bains | Anne Pohlmann (Friedrich-Loeffler-Institut) |
| EPI_ISL_2815336 | A/common teal/Ningxia/105/2020 | A / H5N8 Asia / China / Ningxia Hui Autonomous Region | Hongliang Chai (Northeast Forestry University / College of Wildlife Resources) |
| EPI_ISL_2815374 | A/mallard/Ningxia/175/2020 | A / H5N8 Asia / China / Ningxia Hui Autonomous Region | Hongliang Chai (Northeast Forestry University / College of Wildlife Resources) |
| EPI_ISL_2820250 | A/mallard/Ningxia/176/2020 | A / H5N8 Asia / China / Ningxia Hui Autonomous Region | Hongliang Chai (Northeast Forestry University / College of Wildlife Resources) |
| EPI_ISL_2820259 | A/common teal/Ningxia/181/2020 | A / H5N8 Asia / China / Ningxia Hui Autonomous Region | Hongliang Chai (Northeast Forestry University / College of Wildlife Resources) |
| EPI_ISL_2820261 | A/common teal/Ningxia/189/2020 | A / H5N8 Asia / China / Ningxia Hui Autonomous Region | Hongliang Chai (Northeast Forestry University / College of Wildlife Resources) |
| EPI_ISL_2820262 | A/common teal/Ningxia/237/2020 | A / H5N8 Asia / China / Ningxia Hui Autonomous Region | Hongliang Chai (Northeast Forestry University / College of Wildlife Resources) |
| EPI_ISL_2820288 | A/mallard/Ningxia/239/2020 | A / H5N8 Asia / China / Ningxia Hui Autonomous Region | Hongliang Chai (Northeast Forestry University / College of Wildlife Resources) |
| EPI_ISL_2820465 | A/mallard/Ningxia/241/2020 | A / H5N8 Asia / China / Ningxia Hui Autonomous Region | Hongliang Chai (Northeast Forestry University / College of Wildlife Resources) |
| EPI_ISL_2820479 | A/common pochard/Ningxia/243/2020 | A / H5N8 Asia / China / Ningxia Hui Autonomous Region | Hongliang Chai (Northeast Forestry University / College of Wildlife Resources) |
| EPI_ISL_2820480 | A/common teal/Ningxia/245/2020 | A / H5N8 Asia / China / Ningxia Hui Autonomous Region | Hongliang Chai (Northeast Forestry University / College of Wildlife Resources) |
| EPI_ISL_2820501 | A/mallard/Ningxia/247/2020 | A / H5N8 Asia / China / Ningxia Hui Autonomous Region | Hongliang Chai (Northeast Forestry University / College of Wildlife Resources) |
| EPI_ISL_2820503 | A/mallard/Ningxia/249/2020 | A / H5N8 Asia / China / Ningxia Hui Autonomous Region | Hongliang Chai (Northeast Forestry University / College of Wildlife Resources) |
| EPI_ISL_2820504 | A/common teal/Ningxia/253/2020 | A / H5N8 Asia / China / Ningxia Hui Autonomous Region | Hongliang Chai (Northeast Forestry University / College of Wildlife Resources) |
| EPI_ISL_282133 | A/turkey/Czech Republic/38-17_1/2017 (H5N8) | A / H5N8 Europe / Czech Republic / South Moravian Region / Okres Brno-Venkov / Ivan?ice-N?m?ice | Alexander Nagy (State Veterinary Institute Prague) |
| EPI_ISL_282141 | A/swan/Italy/17VIR7064-1/2017 | A / H5N8 Europe / Italy / Lombardy / Province of Bergamo / Mozzanica | Bianca Zecchin (Istituto Zooprofilattico Sperimentale Delle Venezie) |
| EPI_ISL_282143 | A/goose/Italy/17VIR6358-3/2017 | A / H5N8 Europe / Italy / Lombardy / Province of Pavia / Chignolo Po | Bianca Zecchin (Istituto Zooprofilattico Sperimentale Delle Venezie) |
| EPI_ISL_282394 | A/goose/Yangzhou/YZ587/2016 | A / H5N6 Asia / China | |
| EPI_ISL_282395 | A/chicken/Xuzhou/XZ6/2016 | A / H5N6 Asia / China | |
| EPI_ISL_282396 | A/duck/Anhui/S4/2016 | A / H5N6 Asia / China | |
| EPI_ISL_282397 | A/chicken/Guangdong/GD1602/2016 | A / H5N6 Asia / China | |
| EPI_ISL_282408 | A/poultry/China/XY918.4/2016 | A / H5N6 Asia / China | |
| EPI_ISL_283129 | A/chicken/Czech Republic/988-17/2017 (H5N8) | A / H5N8 Europe / Czech Republic / Olomoucky Kraj / Okres Prerov / Lov?šice | Alexander Nagy (State Veterinary Institute Prague) |
| EPI_ISL_283702 | A/Duck/Yunnan/YN-2/2016 (H5N6) | A / H5N6 Asia / China / Yunnan Province | Qi Wenbao (South China Agricultural University / Veterinary Medicine College) |
| EPI_ISL_283703 | A/Chicken/Yunnan/YN-3/2016 (H5N6) | A / H5N6 Asia / China / Yunnan Province | Qi Wenbao (South China Agricultural University / Veterinary Medicine College) |
| EPI_ISL_283705 | A/Chicken/Yunnan/YN-7/2016 (H5N6) | A / H5N6 Asia / China / Yunnan Province | Qi Wenbao (South China Agricultural University / Veterinary Medicine College) |
| EPI_ISL_283965 | A/Chicken/Yunnan/YN-8/2016 (H5N6) | A / H5N6 Asia / China / Yunnan Province | Qi Wenbao (South China Agricultural University / Veterinary Medicine College) |
| EPI_ISL_283966 | A/Duck/Yunnan/YN-9/2016 (H5N6) | A / H5N6 Asia / China / Yunnan Province | Qi Wenbao (South China Agricultural University / Veterinary Medicine College) |
| EPI_ISL_283967 | A/Chicken/Yunnan/YN-10/2015 (H5N6) | A / H5N6 Asia / China / Yunnan Province | Qi Wenbao (South China Agricultural University / Veterinary Medicine College) |
| EPI_ISL_283968 | A/Chicken/Yunnan/YN-11/2016 (H5N6) | A / H5N6 Asia / China / Yunnan Province | Qi Wenbao (South China Agricultural University / Veterinary Medicine College) |
| EPI_ISL_283969 | A/Chicken/Yunnan/YN-12/2016 (H5N6) | A / H5N6 Asia / China / Yunnan Province | Qi Wenbao (South China Agricultural University / Veterinary Medicine College) |
| EPI_ISL_283970 | A/Ostrich/Guangxi/GX-1/2017 (H5N6) | A / H5N6 Asia / China / Guangxi Zhuang Autonomous Region | Qi Wenbao (South China Agricultural University / Veterinary Medicine College) |

| | | | | |
|---|---|---|---|---|
| EPI_ISL_283971 | A/Quail/Guangxi/GX-2/2017 (H5N6) | A / H5N6 | Asia / China / Guangxi Zhuang Autonomous Region | Qi Wenbao (South China Agricultural University / Veterinary Medicine College) |
| EPI_ISL_284004 | A/Chicken/South_Africa/S2017/08_0336_P3/2017 | A / H5N8 | Africa / South Africa / Province of the Western Cape | Florette Kathleen Treurnicht (National Institute for Communicable Diseases (NICD) / Center for Respiratory Diseases and Meningitis) |
| EPI_ISL_284196 | A/Pigeon/South_Africa/S2017/08_0324_24/2017 | A / H5N8 | Africa / South Africa / Province of the Western Cape | Florette Kathleen Treurnicht (National Institute for Communicable Diseases (NICD) / Center for Respiratory Diseases and Meningitis) |
| EPI_ISL_284197 | A/Ostrich/South_Africa/S2017/08_0046_P2/2017 | A / H5N8 | Africa / South Africa / Province of the Western Cape | Florette Kathleen Treurnicht (National Institute for Communicable Diseases (NICD) / Center for Respiratory Diseases and Meningitis) |
| EPI_ISL_284198 | A/Ostrich/South_Africa/S2017/08_0361_P17/2017 | A / H5N8 | Africa / South Africa / Province of the Western Cape | Florette Kathleen Treurnicht (National Institute for Communicable Diseases (NICD) / Center for Respiratory Diseases and Meningitis) |
| EPI_ISL_284650 | A/Anhui/33162/2016 | A / H5N6 | Asia / China / Anhui Province | Lei Yang (WHO Chinese National Influenza Center / Virology Institute, Chinese CDC) |
| EPI_ISL_284651 | A/Anhui/33163/2016 | A / H5N6 | Asia / China / Anhui Province | Lei Yang (WHO Chinese National Influenza Center / Virology Institute, Chinese CDC) |
| EPI_ISL_284683 | A/Ostrich/South_Africa/S2017/08_0268_P2/2017 | A / H5N8 | Africa / South Africa / Province of the Western Cape | Florette Kathleen Treurnicht (National Institute for Communicable Diseases (NICD) / Center for Respiratory Diseases and Meningitis) |
| EPI_ISL_284684 | A/Ostrich/South_Africa/S2017/08_0268_P9/2017 | A / H5N8 | Africa / South Africa / Province of the Western Cape | Florette Kathleen Treurnicht (National Institute for Communicable Diseases (NICD) / Center for Respiratory Diseases and Meningitis) |
| EPI_ISL_284692 | A/Geese/South_Africa/S2017/09_0055_P2/2017 | A / H5N8 | Africa / South Africa / Province of the Western Cape | Florette Kathleen Treurnicht (National Institute for Communicable Diseases (NICD) / Center for Respiratory Diseases and Meningitis) |
| EPI_ISL_285480 | A/Ostrich/South_Africa/S2017/08_0362_P10/2017 | A / H5N8 | Africa / South Africa / Province of the Western Cape | Florette Kathleen Treurnicht (National Institute for Communicable Diseases (NICD) / Center for Respiratory Diseases and Meningitis) |
| EPI_ISL_285488 | A/Ostrich/South_Africa/S2017/08_0047_P7/2017 | A / H5N8 | Africa / South Africa / Province of the Western Cape | Florette Kathleen Treurnicht (National Institute for Communicable Diseases (NICD) / Center for Respiratory Diseases and Meningitis) |
| EPI_ISL_285509 | A/Ostrich/South_Africa/S2017/08_0362_P11/2017 | A / H5N8 | Africa / South Africa / Province of the Western Cape | Florette Kathleen Treurnicht (National Institute for Communicable Diseases (NICD) / Center for Respiratory Diseases and Meningitis) |
| EPI_ISL_285512 | A/Chicken/South_Africa/S2017/08_0336_P1/2017 | A / H5N8 | Africa / South Africa / Province of the Western Cape | Florette Kathleen Treurnicht (National Institute for Communicable Diseases (NICD) / Center for Respiratory Diseases and Meningitis) |
| EPI_ISL_285601 | A/Geese/South_Africa/S2017/09_0065_P1/2017 | A / H5N8 | Africa / South Africa / Province of the Western Cape | Florette Kathleen Treurnicht (National Institute for Communicable Diseases (NICD) / Center for Respiratory Diseases and Meningitis) |
| EPI_ISL_285602 | A/Geese/South_Africa/S2017/09_0065_P2/2017 | A / H5N8 | Africa / South Africa / Province of the Western Cape | Florette Kathleen Treurnicht (National Institute for Communicable Diseases (NICD) / Center for Respiratory Diseases and Meningitis) |
| EPI_ISL_285604 | A/Ostrich/South_Africa/S2017/08_0046_P3/2017 | A / H5N8 | Africa / South Africa / Province of the Western Cape | Florette Kathleen Treurnicht (National Institute for Communicable Diseases (NICD) / Center for Respiratory Diseases and Meningitis) |
| EPI_ISL_285606 | A/Duck/South_Africa/S2017/08_0340_P1/2017 | A / H5N8 | Africa / South Africa / Province of the Western Cape | Florette Kathleen Treurnicht (National Institute for Communicable Diseases (NICD) / Center for Respiratory Diseases and Meningitis) |
| EPI_ISL_285607 | A/Duck/South_Africa/S2017/08_0340_P2/2017 | A / H5N8 | Africa / South Africa / Province of the Western Cape | Florette Kathleen Treurnicht (National Institute for Communicable Diseases (NICD) / Center for Respiratory Diseases and Meningitis) |
| EPI_ISL_285608 | A/Ostrich/South_Africa/S2017/08_0362_P8_33/2017 | A / H5N8 | Africa / South Africa / Province of the Western Cape | Florette Kathleen Treurnicht (National Institute for Communicable Diseases (NICD) / Center for Respiratory Diseases and Meningitis) |
| EPI_ISL_285609 | A/Geese/South_Africa/S2017/08_0558_P2/2017 | A / H5N8 | Africa / South Africa / Province of the Western Cape | Florette Kathleen Treurnicht (National Institute for Communicable Diseases (NICD) / Center for Respiratory Diseases and Meningitis) |
| EPI_ISL_285610 | A/Guineafowl/South_Africa/S2017/08_0274_P1/2017 | A / H5N8 | Africa / South Africa / Province of the Western Cape | Florette Kathleen Treurnicht (National Institute for Communicable Diseases (NICD) / Center for Respiratory Diseases and Meningitis) |
| EPI_ISL_285611 | A/Wildbirds_Guineafowl_Makou_Egyptian Geese/South_Africa | A / H5N8 | Africa / South Africa / Province of the Western Cape | Florette Kathleen Treurnicht (National Institute for Communicable Diseases (NICD) / Center for Respiratory Diseases and Meningitis) |
| EPI_ISL_285616 | A/Ostrich/South_Africa/S2017/08_0161_P8/2017 | A / H5N8 | Africa / South Africa / Province of the Western Cape | Florette Kathleen Treurnicht (National Institute for Communicable Diseases (NICD) / Center for Respiratory Diseases and Meningitis) |
| EPI_ISL_285617 | A/Guineafowl/South_Africa/S2017/08_0243_P1/2017 | A / H5N8 | Africa / South Africa / Province of the Western Cape | Florette Kathleen Treurnicht (National Institute for Communicable Diseases (NICD) / Center for Respiratory Diseases and Meningitis) |
| EPI_ISL_285618 | A/Ostrich/South_Africa/S2017/08_0362_P7/2017 | A / H5N8 | Africa / South Africa / Province of the Western Cape | Florette Kathleen Treurnicht (National Institute for Communicable Diseases (NICD) / Center for Respiratory Diseases and Meningitis) |
| EPI_ISL_285619 | A/chicken/Egypt/Kafr-Elshiekh-18/2017 | A / H5N8 | Africa / Egypt / Kafr-Elshiekh | El-Sayed M. Abdelwhab (Friedrich-Loeffler-Institut) |
| EPI_ISL_285620 | A/chicken/Egypt/Buheira-12/2017 | A / H5N8 | Africa / Egypt / Buheira | El-Sayed M. Abdelwhab (Friedrich-Loeffler-Institut) |
| EPI_ISL_285621 | A/chicken/Egypt/Gharbiya-15/2017 | A / H5N8 | Africa / Egypt / Gharbiya | El-Sayed M. Abdelwhab (Friedrich-Loeffler-Institut) |
| EPI_ISL_285623 | A/Chicken/South_Africa/S2017/08_0416_38/2017 | A / H5N8 | Africa / South Africa / Province of the Western Cape | Florette Kathleen Treurnicht (National Institute for Communicable Diseases (NICD) / Center for Respiratory Diseases and Meningitis) |
| EPI_ISL_285624 | A/Chicken/South_Africa/S2017/09_0050_56/2017 | A / H5N8 | Africa / South Africa / Province of the Western Cape | Florette Kathleen Treurnicht (National Institute for Communicable Diseases (NICD) / Center for Respiratory Diseases and Meningitis) |
| EPI_ISL_285625 | A/Duck/South_Africa/S2017/08_0481_P2/2017 | A / H5N8 | Africa / South Africa / Province of the Western Cape | Florette Kathleen Treurnicht (National Institute for Communicable Diseases (NICD) / Center for Respiratory Diseases and Meningitis) |
| EPI_ISL_285627 | A/Swan/South_Africa/S2017/08_0517_P2/2017 | A / H5N8 | Africa / South Africa / Province of the Western Cape | Florette Kathleen Treurnicht (National Institute for Communicable Diseases (NICD) / Center for Respiratory Diseases and Meningitis) |
| EPI_ISL_285647 | A/Geese/South_Africa/S2017/08_0520_46/2017 | A / H5N8 | Africa / South Africa / Province of the Western Cape | Florette Kathleen Treurnicht (National Institute for Communicable Diseases (NICD) / Center for Respiratory Diseases and Meningitis) |
| EPI_ISL_285648 | A/Geese/South_Africa/S2017/09_0055_P1/2017 | A / H5N8 | Africa / South Africa / Province of the Western Cape | Florette Kathleen Treurnicht (National Institute for Communicable Diseases (NICD) / Center for Respiratory Diseases and Meningitis) |
| EPI_ISL_285649 | A/Chicken/South_Africa/S2017/09_0184_63/2017 | A / H5N8 | Africa / South Africa / Province of the Western Cape | Florette Kathleen Treurnicht (National Institute for Communicable Diseases (NICD) / Center for Respiratory Diseases and Meningitis) |
| EPI_ISL_285650 | A/Ostrich/South_Africa/S2017/08_0046_AF/2017 | A / H5N8 | Africa / South Africa / Province of the Western Cape | Florette Kathleen Treurnicht (National Institute for Communicable Diseases (NICD) / Center for Respiratory Diseases and Meningitis) |
| EPI_ISL_285651 | A/Guineafowl/South_Africa/S2017/08_0190_9/2017 | A / H5N8 | Africa / South Africa / Province of the Western Cape | Florette Kathleen Treurnicht (National Institute for Communicable Diseases (NICD) / Center for Respiratory Diseases and Meningitis) |
| EPI_ISL_285652 | A/Guineafowl/South_Africa/S2017/08_0243_P2/2017 | A / H5N8 | Africa / South Africa / Province of the Western Cape | Florette Kathleen Treurnicht (National Institute for Communicable Diseases (NICD) / Center for Respiratory Diseases and Meningitis) |
| EPI_ISL_285653 | A/Pigeon/South_Africa/S2017/08_0323_P1/2017 | A / H5N8 | Africa / South Africa / Province of the Western Cape | Florette Kathleen Treurnicht (National Institute for Communicable Diseases (NICD) / Center for Respiratory Diseases and Meningitis) |
| EPI_ISL_285655 | A/Geese/South_Africa/S2017/08_0558_P1/2017 | A / H5N8 | Africa / South Africa / Province of the Western Cape | Florette Kathleen Treurnicht (National Institute for Communicable Diseases (NICD) / Center for Respiratory Diseases and Meningitis) |
| EPI_ISL_285657 | A/Chicken/South_Africa/S2017/08_0561_P2/2017 | A / H5N8 | Africa / South Africa / Province of the Western Cape | Florette Kathleen Treurnicht (National Institute for Communicable Diseases (NICD) / Center for Respiratory Diseases and Meningitis) |
| EPI_ISL_285658 | A/Chicken/South_Africa/S2017/08_0561_P1/2017 | A / H5N8 | Africa / South Africa / Province of the Western Cape | Florette Kathleen Treurnicht (National Institute for Communicable Diseases (NICD) / Center for Respiratory Diseases and Meningitis) |
| EPI_ISL_285915 | A/Chicken/South_Africa/S2017/08_0581_P1/2017 | A / H5N8 | Africa / South Africa / Province of the Western Cape | Florette Kathleen Treurnicht (National Institute for Communicable Diseases (NICD) / Center for Respiratory Diseases and Meningitis) |
| EPI_ISL_285916 | A/Chicken/South_Africa/S2017/08_0581_P2/2017 | A / H5N8 | Africa / South Africa / Province of the Western Cape | Florette Kathleen Treurnicht (National Institute for Communicable Diseases (NICD) / Center for Respiratory Diseases and Meningitis) |
| EPI_ISL_285917 | A/Chicken/South_Africa/S2017/09_0184_62/2017 | A / H5N8 | Africa / South Africa / Province of the Western Cape | Florette Kathleen Treurnicht (National Institute for Communicable Diseases (NICD) / Center for Respiratory Diseases and Meningitis) |
| EPI_ISL_285918 | A/Ostrich/South_Africa/S2017/08_0362_P8_34/2017 | A / H5N8 | Africa / South Africa / Province of the Western Cape | Florette Kathleen Treurnicht (National Institute for Communicable Diseases (NICD) / Center for Respiratory Diseases and Meningitis) |
| EPI_ISL_287564 | A/T_Dk/NL-Werkendam/16014159-001/2016 | A / H5N5 | Europe / Netherlands / North Brabant / Gemeente Werkendam / Werkendam | Saskia Bergervoet (Wageningen Bioveterinary Research) |
| EPI_ISL_287565 | A/M_Swan/NL-Groningen/16015826-001/2016 | A / H5N5 | Europe / Netherlands / Provincie Groningen / Gemeente Groningen / Groningen | Saskia Bergervoet (Wageningen Bioveterinary Research) |
| EPI_ISL_287800 | A/spoonbill/Taiwan/DB645/2017 | A / H5N6 | Asia / Taiwan | Yu-Pin Liu (Animal Health Research Institute) |
| EPI_ISL_287906 | A/Duck/Netherlands/17017236-001-005/2017 | A / H5N6 | Europe / Netherlands / Provincie Flevoland / Gemeente Dronten / Biddinghuizen | Saskia Bergervoet (Wageningen Bioveterinary Research) |
| EPI_ISL_287907 | A/Duck/Netherlands/17017237-001-005/2017 | A / H5N6 | Europe / Netherlands / Provincie Flevoland / Gemeente Dronten / Biddinghuizen | Saskia Bergervoet (Wageningen Bioveterinary Research) |
| EPI_ISL_288362 | A/chicken/Greece/39_2017/2017 | A / H5N6 | Europe / Greece | James Seekings (Animal and Plant Health Agency (APHA) / Virology Department) |
| EPI_ISL_288363 | A/chicken/Greece/39_2017a/2017 | A / H5N6 | Europe / Greece | James Seekings (Animal and Plant Health Agency (APHA) / Virology Department) |
| EPI_ISL_288364 | A/chicken/Greece/39_2017b/2017 | A / H5N6 | Europe / Greece | James Seekings (Animal and Plant Health Agency (APHA) / Virology Department) |
| EPI_ISL_288409 | A/Mute Swan/Netherlands/17017367-012/2017 | A / H5N6 | Europe / Netherlands / Provincie Gelderland / Hulshorst | Saskia Bergervoet (Wageningen Bioveterinary Research) |
| EPI_ISL_288410 | A/Mute Swan/Netherlands/17017377-001/2017 | A / H5N6 | Europe / Netherlands / Provincie Gelderland / Gemeente Elburg | Saskia Bergervoet (Wageningen Bioveterinary Research) |
| EPI_ISL_288411 | A/Go/NL-Utrecht/17006881-001/2017 | A / H5N5 | Europe / Netherlands / Provincie Utrecht / Utrecht | Saskia Bergervoet (Wageningen Bioveterinary Research) |
| EPI_ISL_288412 | A/Tufted Duck/Netherlands/17017367-007/2017 | A / H5N6 | Europe / Netherlands / Provincie Gelderland / Hulshorst | Saskia Bergervoet (Wageningen Bioveterinary Research) |
| EPI_ISL_288436 | A/mallard/Korea/Jeju-H24/2017 | A / H5N6 | Asia / Korea, Republic of / Jeju-teukbyeoljachido | Eun-Kyoung Lee (Animal and Plant Quarantine Agency (APQA) / Avian Influenza Research and Diagsnotic Division) |
| EPI_ISL_288437 | A/duck/Korea/HD1/2017 | A / H5N6 | Asia / Korea, Republic of / Jeollabuk-do | Eun-Kyoung Lee (Animal and Plant Quarantine Agency (APQA) / Avian Influenza Research and Diagsnotic Division) |
| EPI_ISL_288438 | A/chicken/Korea/Gunsan/2017 | A / H5N8 | Asia / Korea, Republic of / Jeollabuk-do | Eun-Kyoung Lee (Animal and Plant Quarantine Agency (APQA) / Avian Influenza Research and Diagsnotic Division) |
| EPI_ISL_288439 | A/chicken/Korea/Gimje2/2017 | A / H5N8 | Asia / Korea, Republic of / Jeollabuk-do | Eun-Kyoung Lee (Animal and Plant Quarantine Agency (APQA) / Avian Influenza Research and Diagsnotic Division) |
| EPI_ISL_289173 | A/mute swan/Shimane/3211A001/2017 | A / H5N6 | Asia / Japan / Shimane | |
| EPI_ISL_289500 | A/Great Black-backed Gull/Netherlands/1/2017 | A / H5N6 | Europe / Netherlands / Provincie Noord-Holland / De Kreupel: 52° 47' 58.3044'' N, 5° 13' 34.550 | Maria Johanna Poen (Erasmus Medical Center / Department of Virology) |
| EPI_ISL_289501 | A/Black-headed Gull/Netherlands/29/2017 | A / H5N6 | Europe / Netherlands / Provincie Noord-Holland / De Kreupel: 52° 47' 58.3044'' N, 5° 13' 34.550 | Maria Johanna Poen (Erasmus Medical Center / Department of Virology) |
| EPI_ISL_290070 | A/duck/Liaoning/LN/2011 | A / H5N5 | Asia / China | |
| EPI_ISL_290071 | A/common porchard/Aichi/2301N025/2017 | A / H5N6 | Asia / Japan / Aichi | |
| EPI_ISL_290072 | A/eastern buzzard/Oita/4401F005/2017 | A / H5N6 | Asia / Japan / Oita | |
| EPI_ISL_290073 | A/mandarin duck/Tochigi/0902C033/2017 | A / H5N6 | Asia / Japan / Tochigi | |
| EPI_ISL_290156 | A/mute swan/Ibaraki/0803_210/2016 | A / H5N6 | Asia / Japan / Ibaraki | |
| EPI_ISL_290157 | A/northern goshawk/Tochigi/0912A004/2016 | A / H5N6 | Asia / Japan / Tochigi | |

| | | | |
|---|---|---|---|
| EPI_ISL_290158 | A/environment/Gifu/21/2017 | A / H5N6 Asia / Japan / Gifu | |
| EPI_ISL_290250 | A/duck/Vietnam/QuangBinh/LBM0818/2016(H5N6) | A / H5N6 Asia / Vietnam | |
| EPI_ISL_290251 | A/duck/Vietnam/QuangBinh/LBM0908/2016(H5N6) | A / H5N6 Asia / Vietnam | |
| EPI_ISL_290252 | A/duck/Vietnam/QuangBinh/LBM0909/2016(H5N6) | A / H5N6 Asia / Vietnam | |
| EPI_ISL_290253 | A/duck/Vietnam/QuangBinh/LBM0911/2016(H5N6) | A / H5N6 Asia / Vietnam | |
| EPI_ISL_290294 | A/Heron/Vietnam/QuangBinh/LBM0910/2016(H5N6) | A / H5N6 Asia / Vietnam | |
| EPI_ISL_290295 | A/mute swan/Hyogo/2801ITM002/2017 | A / H5N6 Asia / Japan / Hyogo | |
| EPI_ISL_290296 | A/mute swan/Hyogo/2801ITM003/2017 | A / H5N6 Asia / Japan / Hyogo | |
| EPI_ISL_290297 | A/mute swan/Hyogo/2801ITM004/2017 | A / H5N6 Asia / Japan / Hyogo | |
| EPI_ISL_290298 | A/mute swan/Hyogo/2801ITM005/2017 | A / H5N6 Asia / Japan / Hyogo | |
| EPI_ISL_290299 | A/mute swan/Hyogo/2801ITM006/2017 | A / H5N6 Asia / Japan / Hyogo | |
| EPI_ISL_290300 | A/mute swan/Hyogo/2801ITM007/2017 | A / H5N6 Asia / Japan / Hyogo | |
| EPI_ISL_290301 | A/mute swan/Hyogo/2801ITM008/2017 | A / H5N6 Asia / Japan / Hyogo | |
| EPI_ISL_290302 | A/mute swan/Hyogo/2801ITM009/2017 | A / H5N6 Asia / Japan / Hyogo | |
| EPI_ISL_290305 | A/mute swan/Hyogo/2801ITM010/2017 | A / H5N6 Asia / Japan / Hyogo | |
| EPI_ISL_290306 | A/mute swan/Hyogo/2801ITM011/2017 | A / H5N6 Asia / Japan / Hyogo | |
| EPI_ISL_290307 | A/mute swan/Hyogo/2801ITM012/2017 | A / H5N6 Asia / Japan / Hyogo | |
| EPI_ISL_290308 | A/mute swan/Hyogo/2801ITM013/2017 | A / H5N6 Asia / Japan / Hyogo | |
| EPI_ISL_290309 | A/mute swan/Hyogo/2801ITM014/2017 | A / H5N6 Asia / Japan / Hyogo | |
| EPI_ISL_290310 | A/peregrine falcon/Niigata/15/2016 | A / H5N6 Asia / Japan / Niigata | |
| EPI_ISL_290311 | A/whooper swan/Niigata/5112008/2016 | A / H5N6 Asia / Japan / Niigata | |
| EPI_ISL_290312 | A/greater scaup/Aichi/2301H050/2017 | A / H5N6 Asia / Japan / Aichi | |
| EPI_ISL_290313 | A/peregrine falcon/Gifu/2102A008/2017 | A / H5N6 Asia / Japan / Gifu | |
| EPI_ISL_291109 | A/common_pochard/Germany-BY/AR09-18-L02421/2017 | A / H5N6 Europe / Germany / Bavaria | Anne Pohlmann (Friedrich-Loeffler-Institut) |
| EPI_ISL_291952 | A/Bar-headed Goose/Qinghai/B44/2016 | A / H5N8 Asia / China / Qinghai Province | Jianjun Chen (Wuhan Institute of Virology / Chinese Academy of Sciences) |
| EPI_ISL_292170 | A/Bar-headed Goose/Qinghai/A22/2016 | A / H5N8 Asia / China / Qinghai Province | Jianjun Chen (Wuhan Institute of Virology / Chinese Academy of Sciences) |
| EPI_ISL_292172 | A/Bar-headed Goose/Qinghai/B7/2016 | A / H5N8 Asia / China / Qinghai Province | Jianjun Chen (Wuhan Institute of Virology / Chinese Academy of Sciences) |
| EPI_ISL_292173 | A/Bar-headed Goose/Qinghai/B54/2016 | A / H5N8 Asia / China / Qinghai Province | Jianjun Chen (Wuhan Institute of Virology / Chinese Academy of Sciences) |
| EPI_ISL_292174 | A/Bar-headed Goose/Qinghai/XX431/2016 | A / H5N8 Asia / China / Qinghai Province | Jianjun Chen (Wuhan Institute of Virology / Chinese Academy of Sciences) |
| EPI_ISL_292175 | A/Bar-headed Goose/Qinghai/XX446/2016 | A / H5N8 Asia / China / Qinghai Province | Jianjun Chen (Wuhan Institute of Virology / Chinese Academy of Sciences) |
| EPI_ISL_292176 | A/Bar-headed Goose/Qinghai/A17/2016 | A / H5N8 Asia / China / Qinghai Province | Jianjun Chen (Wuhan Institute of Virology / Chinese Academy of Sciences) |
| EPI_ISL_292177 | A/Bar-headed Goose/Qinghai/A20/2016 | A / H5N8 Asia / China / Qinghai Province | Jianjun Chen (Wuhan Institute of Virology / Chinese Academy of Sciences) |
| EPI_ISL_292178 | A/Bar-headed Goose/Qinghai/A12/2016 | A / H5N8 Asia / China / Qinghai Province | Jianjun Chen (Wuhan Institute of Virology / Chinese Academy of Sciences) |
| EPI_ISL_292179 | A/Bar-headed Goose/Qinghai/XX13/2016 | A / H5N8 Asia / China / Qinghai Province | Jianjun Chen (Wuhan Institute of Virology / Chinese Academy of Sciences) |
| EPI_ISL_292180 | A/Bar-headed Goose/Qinghai/XX22/2016 | A / H5N8 Asia / China / Qinghai Province | Jianjun Chen (Wuhan Institute of Virology / Chinese Academy of Sciences) |
| EPI_ISL_292181 | A/Bar-headed Goose/Qinghai/A19/2016 | A / H5N8 Asia / China / Qinghai Province | Jianjun Chen (Wuhan Institute of Virology / Chinese Academy of Sciences) |
| EPI_ISL_292182 | A/Bar-headed Goose/Qinghai/A16/2016 | A / H5N8 Asia / China / Qinghai Province | Jianjun Chen (Wuhan Institute of Virology / Chinese Academy of Sciences) |
| EPI_ISL_292183 | A/Bar-headed Goose/Qinghai/A13/2016 | A / H5N8 Asia / China / Qinghai Province | Jianjun Chen (Wuhan Institute of Virology / Chinese Academy of Sciences) |
| EPI_ISL_292184 | A/Bar-headed Goose/Qinghai/B51/2016 | A / H5N8 Asia / China / Qinghai Province | Jianjun Chen (Wuhan Institute of Virology / Chinese Academy of Sciences) |
| EPI_ISL_292185 | A/Great Cormorant/Qinghai/Y01/2016 | A / H5N8 Asia / China / Qinghai Province | Jianjun Chen (Wuhan Institute of Virology / Chinese Academy of Sciences) |
| EPI_ISL_292186 | A/Bar-headed Goose/Qinghai/B34/2016 | A / H5N8 Asia / China / Qinghai Province | Jianjun Chen (Wuhan Institute of Virology / Chinese Academy of Sciences) |
| EPI_ISL_292187 | A/Bar-headed Goose/Qinghai/XX782/2016 | A / H5N8 Asia / China / Qinghai Province | Jianjun Chen (Wuhan Institute of Virology / Chinese Academy of Sciences) |
| EPI_ISL_292188 | A/Great Cormorant/Qinghai/B82/2016 | A / H5N8 Asia / China / Qinghai Province | Jianjun Chen (Wuhan Institute of Virology / Chinese Academy of Sciences) |
| EPI_ISL_292189 | A/Bar-headed Goose/Qinghai/A21/2016 | A / H5N8 Asia / China / Qinghai Province | Jianjun Chen (Wuhan Institute of Virology / Chinese Academy of Sciences) |
| EPI_ISL_292190 | A/Bar-headed Goose/Qinghai/XX111/2016 | A / H5N8 Asia / China / Qinghai Province | Jianjun Chen (Wuhan Institute of Virology / Chinese Academy of Sciences) |
| EPI_ISL_292191 | A/Bar-headed Goose/Qinghai/B12/2016 | A / H5N8 Asia / China / Qinghai Province | Jianjun Chen (Wuhan Institute of Virology / Chinese Academy of Sciences) |
| EPI_ISL_292192 | A/Bar-headed Goose/Qinghai/A23/2016 | A / H5N8 Asia / China / Qinghai Province | Jianjun Chen (Wuhan Institute of Virology / Chinese Academy of Sciences) |
| EPI_ISL_292193 | A/Bar-headed Goose/Qinghai/XX76/2016 | A / H5N8 Asia / China / Qinghai Province | Jianjun Chen (Wuhan Institute of Virology / Chinese Academy of Sciences) |
| EPI_ISL_292194 | A/Bar-headed Goose/Qinghai/a88/2016 | A / H5N8 Asia / China / Qinghai Province | Jianjun Chen (Wuhan Institute of Virology / Chinese Academy of Sciences) |
| EPI_ISL_292196 | A/Bar-headed Goose/Qinghai/a32/2016 | A / H5N8 Asia / China / Qinghai Province | Jianjun Chen (Wuhan Institute of Virology / Chinese Academy of Sciences) |
| EPI_ISL_292198 | A/Bar-headed Goose/Qinghai/a115/2016 | A / H5N8 Asia / China / Qinghai Province | Jianjun Chen (Wuhan Institute of Virology / Chinese Academy of Sciences) |
| EPI_ISL_292223 | A/mute_swan/England/AVP_18_001986/2017 | A / H5N6 Europe / United Kingdom / Dorset / Abbotsbury | James Seekings (Animal and Plant Health Agency (APHA) / Virology Department) |
| EPI_ISL_292224 | A/pochard_duck/England/AVP_18_003254/2018 | A / H5N6 Europe / United Kingdom / Dorset / Abbotsbury | James Seekings (Animal and Plant Health Agency (APHA) / Virology Department) |
| EPI_ISL_292225 | A/canada_goose/England/AV58_18OPpoolEP1/2018 | A / H5N6 Europe / United Kingdom / Dorset / Abbotsbury | James Seekings (Animal and Plant Health Agency (APHA) / Virology Department) |
| EPI_ISL_292229 | A/Bar-headed Goose/Qinghai/a113/2016 | A / H5N8 Asia / China / Qinghai Province | Jianjun Chen (Wuhan Institute of Virology / Chinese Academy of Sciences) |
| EPI_ISL_292230 | A/Bar-headed Goose/Qinghai/a93/2016 | A / H5N8 Asia / China / Qinghai Province | Jianjun Chen (Wuhan Institute of Virology / Chinese Academy of Sciences) |
| EPI_ISL_292231 | A/Bar-headed Goose/Qinghai/a61/2016 | A / H5N8 Asia / China / Qinghai Province | Jianjun Chen (Wuhan Institute of Virology / Chinese Academy of Sciences) |
| EPI_ISL_292232 | A/Bar-headed Goose/Qinghai/HDT001/2016 | A / H5N8 Asia / China / Qinghai Province | Jianjun Chen (Wuhan Institute of Virology / Chinese Academy of Sciences) |
| EPI_ISL_292233 | A/Bar-headed Goose/Qinghai/a91/2016 | A / H5N8 Asia / China / Qinghai Province | Jianjun Chen (Wuhan Institute of Virology / Chinese Academy of Sciences) |
| EPI_ISL_292234 | A/Bar-headed Goose/Qinghai/a92/2016 | A / H5N8 Asia / China / Qinghai Province | Jianjun Chen (Wuhan Institute of Virology / Chinese Academy of Sciences) |
| EPI_ISL_292235 | A/Bar-headed Goose/Qinghai/a43/2016 | A / H5N8 Asia / China / Qinghai Province | Jianjun Chen (Wuhan Institute of Virology / Chinese Academy of Sciences) |
| EPI_ISL_292236 | A/Bar-headed Goose/Qinghai/a24/2016 | A / H5N8 Asia / China / Qinghai Province | Jianjun Chen (Wuhan Institute of Virology / Chinese Academy of Sciences) |
| EPI_ISL_292237 | A/Bar-headed Goose/Qinghai/a15/2016 | A / H5N8 Asia / China / Qinghai Province | Jianjun Chen (Wuhan Institute of Virology / Chinese Academy of Sciences) |
| EPI_ISL_292238 | A/Bar-headed Goose/Qinghai/p18/2016 | A / H5N8 Asia / China / Qinghai Province | Jianjun Chen (Wuhan Institute of Virology / Chinese Academy of Sciences) |
| EPI_ISL_292239 | A/Bar-headed Goose/Qinghai/p9/2016 | A / H5N8 Asia / China / Qinghai Province | Jianjun Chen (Wuhan Institute of Virology / Chinese Academy of Sciences) |
| EPI_ISL_292240 | A/Bar-headed Goose/Qinghai/p23/2016 | A / H5N8 Asia / China / Qinghai Province | Jianjun Chen (Wuhan Institute of Virology / Chinese Academy of Sciences) |
| EPI_ISL_292326 | A/Great Cormorant/Qinghai/a51/2016 | A / H5N8 Asia / China / Qinghai Province | Jianjun Chen (Wuhan Institute of Virology / Chinese Academy of Sciences) |
| EPI_ISL_292327 | A/Bar-headed Goose/Qinghai/p2/2016 | A / H5N8 Asia / China / Qinghai Province | Jianjun Chen (Wuhan Institute of Virology / Chinese Academy of Sciences) |
| EPI_ISL_292328 | A/Bar-headed Goose/Qinghai/a45/2016 | A / H5N8 Asia / China / Qinghai Province | Jianjun Chen (Wuhan Institute of Virology / Chinese Academy of Sciences) |
| EPI_ISL_292329 | A/Bar-headed Goose/Qinghai/a27/2016 | A / H5N8 Asia / China / Qinghai Province | Jianjun Chen (Wuhan Institute of Virology / Chinese Academy of Sciences) |
| EPI_ISL_292330 | A/Bar-headed Goose/Qinghai/a26/2016 | A / H5N8 Asia / China / Qinghai Province | Jianjun Chen (Wuhan Institute of Virology / Chinese Academy of Sciences) |
| EPI_ISL_292331 | A/Bar-headed Goose/Qinghai/a114/2016 | A / H5N8 Asia / China / Qinghai Province | Jianjun Chen (Wuhan Institute of Virology / Chinese Academy of Sciences) |

| | | | | |
|---|---|---|---|---|
| EPI_ISL_292332 | A/water/Qinghai/i34-39/2016 | A / H5N8 | Asia / China / Qinghai Province | Jianjun Chen (Wuhan Institute of Virology / Chinese Academy of Sciences) |
| EPI_ISL_292333 | A/Water/Qinghai/i40-44/2016 | A / H5N8 | Asia / China / Qinghai Province | Jianjun Chen (Wuhan Institute of Virology / Chinese Academy of Sciences) |
| EPI_ISL_292334 | A/Bar-headed Goose/Qinghai/B655/2017 | A / H5N8 | Asia / China / Qinghai Province | Jianjun Chen (Wuhan Institute of Virology / Chinese Academy of Sciences) |
| EPI_ISL_292335 | A/water/Qinghai/XXII2871/2017 | A / H5N8 | Asia / China / Qinghai Province | Jianjun Chen (Wuhan Institute of Virology / Chinese Academy of Sciences) |
| EPI_ISL_292340 | A/Bar-headed Goose/Qinghai/a230/2017 | A / H5N8 | Asia / China / Qinghai Province | Jianjun Chen (Wuhan Institute of Virology / Chinese Academy of Sciences) |
| EPI_ISL_292341 | A/Bar-headed Goose/Qinghai/a218/2017 | A / H5N8 | Asia / China / Qinghai Province | Jianjun Chen (Wuhan Institute of Virology / Chinese Academy of Sciences) |
| EPI_ISL_292345 | A/duck/Korea/H35/2017 | A / H5N6 | Asia / Korea, Republic of / Jeollanam-do | Yu-Na Lee (Animal and Plant Quarantine Agency (S-2158) / Avian Influenza Research & Diagnostic Division) |
| EPI_ISL_292349 | A/duck/Korea/H56/2017 | A / H5N6 | Asia / Korea, Republic of | Yu-Na Lee (Animal and Plant Quarantine Agency (S-2158) / Avian Influenza Research & Diagnostic Division) |
| EPI_ISL_292413 | A/duck/Korea/H70/2017 | A / H5N6 | Asia / Korea, Republic of | Yu-Na Lee (Animal and Plant Quarantine Agency (S-2158) / Avian Influenza Research & Diagnostic Division) |
| EPI_ISL_292414 | A/duck/Korea/H80/2017 | A / H5N6 | Asia / Korea, Republic of | Yu-Na Lee (Animal and Plant Quarantine Agency (S-2158) / Avian Influenza Research & Diagnostic Division) |
| EPI_ISL_292415 | A/duck/Korea/H81/2017 | A / H5N6 | Asia / Korea, Republic of | Yu-Na Lee (Animal and Plant Quarantine Agency (S-2158) / Avian Influenza Research & Diagnostic Division) |
| EPI_ISL_292416 | A/duck/Korea/H103/2017 | A / H5N6 | Asia / Korea, Republic of | Yu-Na Lee (Animal and Plant Quarantine Agency (S-2158) / Avian Influenza Research & Diagnostic Division) |
| EPI_ISL_292417 | A/duck/Korea/H107/2017 | A / H5N6 | Asia / Korea, Republic of | Yu-Na Lee (Animal and Plant Quarantine Agency (S-2158) / Avian Influenza Research & Diagnostic Division) |
| EPI_ISL_292418 | A/mandarin duck/Korea/H69/2017 | A / H5N6 | Asia / Korea, Republic of | Yu-Na Lee (Animal and Plant Quarantine Agency (S-2158) / Avian Influenza Research & Diagnostic Division) |
| EPI_ISL_292419 | A/mandarin duck/Korea/H71/2017 | A / H5N6 | Asia / Korea, Republic of | Yu-Na Lee (Animal and Plant Quarantine Agency (S-2158) / Avian Influenza Research & Diagnostic Division) |
| EPI_ISL_292420 | A/mandarin duck/Korea/H119-3/2017 | A / H5N6 | Asia / Korea, Republic of | Yu-Na Lee (Animal and Plant Quarantine Agency (S-2158) / Avian Influenza Research & Diagnostic Division) |
| EPI_ISL_292476 | A/Bar-headed Goose/Qinghai/a237/2017 | A / H5N8 | Asia / China / Qinghai Province | Jianjun Chen (Wuhan Institute of Virology / Chinese Academy of Sciences) |
| EPI_ISL_292477 | A/Great Crested Grebe/Qinghai/a737/2017 | A / H5N8 | Asia / China / Qinghai Province | Jianjun Chen (Wuhan Institute of Virology / Chinese Academy of Sciences) |
| EPI_ISL_292478 | A/Bar-headed Goose/Qinghai/a765/2017 | A / H5N8 | Asia / China / Qinghai Province | Jianjun Chen (Wuhan Institute of Virology / Chinese Academy of Sciences) |
| EPI_ISL_292479 | A/Bar-headed Goose/Qinghai/a893/2017 | A / H5N8 | Asia / China / Qinghai Province | Jianjun Chen (Wuhan Institute of Virology / Chinese Academy of Sciences) |
| EPI_ISL_292495 | A/Bar-headed Goose/Qinghai/B11/2016 | A / H5N8 | Asia / China / Qinghai Province | Jianjun Chen (Wuhan Institute of Virology / Chinese Academy of Sciences) |
| EPI_ISL_292496 | A/Bar-headed Goose/Qinghai/A11/2016 | A / H5N8 | Asia / China / Qinghai Province | Jianjun Chen (Wuhan Institute of Virology / Chinese Academy of Sciences) |
| EPI_ISL_292497 | A/Bar-headed Goose/Qinghai/XXI122/2016 | A / H5N8 | Asia / China / Qinghai Province | Jianjun Chen (Wuhan Institute of Virology / Chinese Academy of Sciences) |
| EPI_ISL_292498 | A/Bar-headed Goose/Qinghai/a210/2017 | A / H5N8 | Asia / China / Qinghai Province | Jianjun Chen (Wuhan Institute of Virology / Chinese Academy of Sciences) |
| EPI_ISL_2932600 | A/goose/Kazakhstan/4-190-20-B-H5N8-1/2020 | A / H5N8 | Asia / Kazakhstan / Almaty Province / Karasai district, Umtyl, Merei village | Alexandr Shevtsov (National Center of Biotechnology Republic of Kazakhstan) |
| EPI_ISL_2932608 | A/chicken/Kazakhstan/220-B-2-H5N8-4/2020 | A / H5N8 | Asia / Kazakhstan / Almaty Province / Karasai district, Umtyl, Merei village | Alexandr Shevtsov (National Center of Biotechnology Republic of Kazakhstan) |
| EPI_ISL_2932609 | A/duck/Kazakhstan/12-20-B-Talg-11/2020 | A / H5N8 | Asia / Kazakhstan / North Kazakhstan / Timiryazevsky district, Timiryazevo village | Alexandr Shevtsov (National Center of Biotechnology Republic of Kazakhstan) |
| EPI_ISL_2932612 | A/goose/Kazakhstan/7-20-B-Talg-12/2020 | A / H5N8 | Asia / Kazakhstan / North Kazakhstan / Ayyrtausky district, Arykbalyk, Agymtai batyr village | Alexandr Shevtsov (National Center of Biotechnology Republic of Kazakhstan) |
| EPI_ISL_2932614 | A/swan/Kazakhstan/9-20-B-Talg-39/2020 | A / H5N8 | Asia / Kazakhstan / North Kazakhstan / Mamlyutsky district, Mamlyutsky, Mamlyutka village | Alexandr Shevtsov (National Center of Biotechnology Republic of Kazakhstan) |
| EPI_ISL_2932616 | A/chicken/Kazakhstan/12-20-B-Talg-45/2020 | A / H5N8 | Asia / Kazakhstan / North Kazakhstan / Akzhar district, Talshyk, Kazanskoe village | Alexandr Shevtsov (National Center of Biotechnology Republic of Kazakhstan) |
| EPI_ISL_2932617 | A/crow/Kazakhstan/15-20-B-Talg-4/2020 | A / H5N8 | Asia / Kazakhstan / North Kazakhstan / Petropavlovsk city | Alexandr Shevtsov (National Center of Biotechnology Republic of Kazakhstan) |
| EPI_ISL_2932619 | A/swan/Kazakhstan/1-267-20-B-Talg-52/2020 | A / H5N8 | Asia / Kazakhstan / Akmola / Birzhan sal district, Baymyrza, Shoshkaly lake | Alexandr Shevtsov (National Center of Biotechnology Republic of Kazakhstan) |
| EPI_ISL_2932682 | A/pigeon/Kazakhstan/15-20-B-Talg-5/2020 | A / H5N8 | Asia / Kazakhstan / North Kazakhstan / Petropavlovsk city | Alexandr Shevtsov (National Center of Biotechnology Republic of Kazakhstan) |
| EPI_ISL_2932683 | A/chicken/Kazakhstan/1-20-B-Talg-67/2020 | A / H5N8 | Asia / Kazakhstan / North Kazakhstan / Tayinshinsky district, Roshchinsky, Dimitrovka village | Alexandr Shevtsov (National Center of Biotechnology Republic of Kazakhstan) |
| EPI_ISL_293482 | A/chicken/Miyazaki/21/2016 | A / H5N6 | Asia / Japan / Miyazaki | |
| EPI_ISL_293483 | A/chicken/Kumamoto/45/2016 | A / H5N6 | Asia / Japan / Kumamoto | |
| EPI_ISL_293484 | A/environment/Aomori/4/2016 | A / H5N6 | Asia / Japan / Aomori | |
| EPI_ISL_293485 | A/environment/Aomori/R1/2016 | A / H5N6 | Asia / Japan / Aomori | |
| EPI_ISL_293486 | A/environment/Niigata/5/2016 | A / H5N6 | Asia / Japan / Niigata | |
| EPI_ISL_293487 | A/environment/Aomori/2-1/2016 | A / H5N6 | Asia / Japan / Aomori | |
| EPI_ISL_293488 | A/environment/Hokkaido/1-8/2016 | A / H5N6 | Asia / Japan / Hokkaido | |
| EPI_ISL_293489 | A/environment/Saga/4/2017 | A / H5N6 | Asia / Japan / Saga | |
| EPI_ISL_293498 | A/northern goshawk/Mie/2412C007/2016 | A / H5N6 | Asia / Japan / Mie | |
| EPI_ISL_293499 | A/peregrine falcon/Nagasaki/4212A005/2016 | A / H5N6 | Asia / Japan / Nagasaki | |
| EPI_ISL_293500 | A/tufted duck/Hyogo/2812A009/2016 | A / H5N6 | Asia / Japan / Hyogo | |
| EPI_ISL_293501 | A/wigeon/Aichi/2301H025/2017 | A / H5N6 | Asia / Japan / Aichi | |
| EPI_ISL_293502 | A/common porchard/Aichi/2301N017/2017 | A / H5N6 | Asia / Japan / Aichi | |
| EPI_ISL_293503 | A/common porchard/Yamaguchi/3501B002/2017 | A / H5N6 | Asia / Japan / Yamaguchi | |
| EPI_ISL_293504 | A/mute swan/Hyogo/2801ITM001/2017 | A / H5N6 | Asia / Japan / Hyogo | |
| EPI_ISL_293505 | A/coot/Shiga/2501T010/2017 | A / H5N6 | Asia / Japan / Shiga | |
| EPI_ISL_293989 | A/chicken/Vietnam/HU4-42/2015 | A / H5N6 | Asia / Vietnam | |
| EPI_ISL_294412 | A/black swan/Aichi/2312T001/2016 | A / H5N6 | Asia / Japan / Aichi | |
| EPI_ISL_294413 | A/black swan/Aichi/2312T002/2016 | A / H5N6 | Asia / Japan / Aichi | |
| EPI_ISL_294414 | A/black swan/Aichi/2312T003/2016 | A / H5N6 | Asia / Japan / Aichi | |
| EPI_ISL_294600 | A/Von Schrenck's bittern/Jiangxi/Y9/2014 | A / H5N8 | Asia / China | |
| EPI_ISL_294610 | A/turkey/Minnesota/E3/2015 | A / H5N2 | North America / United States | |
| EPI_ISL_294611 | A/turkey/Minnesota/E4/2015 | A / H5N2 | North America / United States | |
| EPI_ISL_294612 | A/turkey/Minnesota/E6/2015 | A / H5N2 | North America / United States | |
| EPI_ISL_294613 | A/environment/Minnesota/E1/2015 | A / H5N2 | North America / United States | |
| EPI_ISL_294614 | A/environment/Minnesota/E2/2015 | A / H5N2 | North America / United States | |
| EPI_ISL_294615 | A/environment/Minnesota/E5/2015 | A / H5N2 | North America / United States | |
| EPI_ISL_294616 | A/environment/Minnesota/E7/2015 | A / H5N2 | North America / United States | |
| EPI_ISL_294617 | A/environment/Minnesota/E8/2015 | A / H5N2 | North America / United States | |
| EPI_ISL_294618 | A/environment/Minnesota/E9/2015 | A / H5N2 | North America / United States | |
| EPI_ISL_294619 | A/environment/Minnesota/E10/2015 | A / H5N2 | North America / United States | |
| EPI_ISL_294628 | A/turkey/Minnesota/E11/2015 | A / H5N2 | North America / United States | |
| EPI_ISL_294630 | A/turkey/Minnesota/E12/2015 | A / H5N2 | North America / United States | |
| EPI_ISL_294631 | A/turkey/Minnesota/E13/2015 | A / H5N2 | North America / United States | |
| EPI_ISL_294632 | A/turkey/Minnesota/E14/2015 | A / H5N2 | North America / United States | |
| EPI_ISL_294635 | A/turkey/Minnesota/E17/2015 | A / H5N2 | North America / United States | |
| EPI_ISL_294636 | A/turkey/Minnesota/E18/2015 | A / H5N2 | North America / United States | |
| EPI_ISL_294683 | A/environment/Minnesota/E15/2015 | A / H5N2 | North America / United States | |

| | | | | |
|---|---|---|---|---|
| EPI_ISL_294685 | A/environment/Minnesota/E16/2015 | A / H5N2 | North America / United States | |
| EPI_ISL_294742 | A/environment/Minnesota/M1/2015 | A / H5N2 | North America / United States | |
| EPI_ISL_294747 | A/environment/Minnesota/M2/2015 | A / H5N2 | North America / United States | |
| EPI_ISL_294748 | A/environment/Minnesota/M3/2015 | A / H5N2 | North America / United States | |
| EPI_ISL_294749 | A/environment/Minnesota/M4/2015 | A / H5N2 | North America / United States | |
| EPI_ISL_294750 | A/environment/Minnesota/M5/2015 | A / H5N2 | North America / United States | |
| EPI_ISL_294751 | A/environment/Minnesota/M6/2015 | A / H5N2 | North America / United States | |
| EPI_ISL_294752 | A/environment/Minnesota/M7/2015 | A / H5N2 | North America / United States | |
| EPI_ISL_294753 | A/environment/Minnesota/M8/2015 | A / H5N2 | North America / United States | |
| EPI_ISL_294754 | A/turkey/Minnesota/M14/2015 | A / H5N2 | North America / United States | |
| EPI_ISL_294755 | A/turkey/Minnesota/M15/2015 | A / H5N2 | North America / United States | |
| EPI_ISL_294756 | A/turkey/Minnesota/M16/2015 | A / H5N2 | North America / United States | |
| EPI_ISL_294757 | A/turkey/Minnesota/M17/2015 | A / H5N2 | North America / United States | |
| EPI_ISL_294759 | A/environment/Minnesota/M9/2015 | A / H5N2 | North America / United States | |
| EPI_ISL_294760 | A/environment/Minnesota/M10/2015 | A / H5N2 | North America / United States | |
| EPI_ISL_294762 | A/environment/Minnesota/M11/2015 | A / H5N2 | North America / United States | |
| EPI_ISL_294763 | A/environment/Minnesota/M12/2015 | A / H5N2 | North America / United States | |
| EPI_ISL_294764 | A/environment/Minnesota/M13/2015 | A / H5N2 | North America / United States | |
| EPI_ISL_294767 | A/environment/Nebraska/L3/2015 | A / H5N2 | North America / United States | |
| EPI_ISL_294768 | A/environment/Nebraska/L4/2015 | A / H5N2 | North America / United States | |
| EPI_ISL_294769 | A/environment/Nebraska/L5/2015 | A / H5N2 | North America / United States | |
| EPI_ISL_294771 | A/environment/Nebraska/L6/2015 | A / H5N2 | North America / United States | |
| EPI_ISL_294773 | A/environment/Nebraska/L8/2015 | A / H5N2 | North America / United States | |
| EPI_ISL_294774 | A/environment/Nebraska/L9/2015 | A / H5N2 | North America / United States | |
| EPI_ISL_294777 | A/environment/Iowa/L2/2015 | A / H5N2 | North America / United States | |
| EPI_ISL_295027 | A/chicken/Kostroma/1718/2017 | A / H5N2 | Europe / Russian Federation / Kostroma Oblast | Ivan Susloparov (State Research Center of Virology and Biotechnology (VECTOR) / Emerging Zoonotic Diseases and Influenza) |
| EPI_ISL_295143 | A/goose/Guangdong/YJD/2014 | A / H5N6 | Asia / China | |
| EPI_ISL_295144 | A/chicken/Yangzhou/YD1/2014 | A / H5N6 | Asia / China | |
| EPI_ISL_295147 | A/chicken/Anhui/QD1/2014 | A / H5N1 | Asia / China | |
| EPI_ISL_295492 | A/Anser cygnoides/Hubei/FW44/2016 | A / H5N8 | Asia / China | |
| EPI_ISL_295493 | A/Cygnus atratus/Hubei/2Z2-O/2016 | A / H5N8 | Asia / China | |
| EPI_ISL_295494 | A/Cygnus atratus/Hubei/HF-1/2016 | A / H5N8 | Asia / China | |
| EPI_ISL_295496 | A/Commom Teal/Ningxia/474-9/2015 | A / H5N6 | Asia / China | |
| EPI_ISL_295497 | A/Shoveller/Ningxia/475-11/2015 | A / H5N6 | Asia / China | |
| EPI_ISL_295498 | A/Wigeon/Ningxia/476-12/2015 | A / H5N6 | Asia / China | |
| EPI_ISL_295499 | A/Commom Teal/Ningxia/478-15/2015 | A / H5N6 | Asia / China | |
| EPI_ISL_295500 | A/Shoveller/Ningxia/481-21/2015 | A / H5N6 | Asia / China | |
| EPI_ISL_295501 | A/Great Crested Grebe/Ningxia/482-26/2015 | A / H5N6 | Asia / China | |
| EPI_ISL_295502 | A/Shoveller/Ningxia/483-28/2015 | A / H5N6 | Asia / China | |
| EPI_ISL_295503 | A/Gadwall/Ningxia/472-7/2015 | A / H5N6 | Asia / China | |
| EPI_ISL_295504 | A/Falcated Duck/Ningxia/484-29/2015 | A / H5N6 | Asia / China | |
| EPI_ISL_295505 | A/Gadwall/Ningxia/485-31/2015 | A / H5N6 | Asia / China | |
| EPI_ISL_295519 | A/Ruddy Shelduck/Ningxia/486-33/2015 | A / H5N6 | Asia / China | |
| EPI_ISL_295520 | A/Gadwall/Ningxia/487-38/2015 | A / H5N6 | Asia / China | |
| EPI_ISL_295521 | A/Shoveller/Ningxia/488-53/2015 | A / H5N6 | Asia / China | |
| EPI_ISL_295701 | A/whooper swan/Ibaraki/28-309/2016 | A / H5N6 | Asia / Japan / Ibaraki | |
| EPI_ISL_295702 | A/black-headed gull/Ibaraki/0803-194/2016 | A / H5N6 | Asia / Japan / Ibaraki | |
| EPI_ISL_295703 | A/mute swan/Ibaraki/0803-201/2016 | A / H5N6 | Asia / Japan / Ibaraki | |
| EPI_ISL_295704 | A/mute swan/Ibaraki/0803-208/2016 | A / H5N6 | Asia / Japan / Ibaraki | |
| EPI_ISL_295705 | A/mute swan/Ibaraki/0803-214/2016 | A / H5N6 | Asia / Japan / Ibaraki | |
| EPI_ISL_295751 | A/greater scaup/Aichi/2301N021/2017 | A / H5N6 | Asia / Japan / Aichi | |
| EPI_ISL_295752 | A/black-headed gull/Hyogo/2801E009/2017 | A / H5N6 | Asia / Japan / Hyogo | |
| EPI_ISL_295754 | A/cackling goose/Aichi/2312T020/2016 | A / H5N6 | Asia / Japan / Aichi | |
| EPI_ISL_295796 | A/muscovy duck/Vietnam/HU2-26/2014 | A / H5N6 | Asia / Vietnam | |
| EPI_ISL_295818 | A/muscovy duck/Vietnam/HU7-17/2017 | A / H5N6 | Asia / Vietnam | |
| EPI_ISL_295819 | A/muscovy duck/Vietnam/HU7-20/2017 | A / H5N6 | Asia / Vietnam | |
| EPI_ISL_295820 | A/muscovy duck/Vietnam/HU7-23/2017 | A / H5N6 | Asia / Vietnam | |
| EPI_ISL_297234 | A/chicken/Rostov-on-Don/1321/2017 | A / H5N8 | Europe / Russian Federation / Rostov Oblast | Ivan Susloparov (State Research Center of Virology and Biotechnology (VECTOR) / Emerging Zoonotic Diseases and Influenza) |
| EPI_ISL_297235 | A/chicken/Rostov-on-Don/1598/2017 | A / H5N8 | Europe / Russian Federation / Rostov Oblast | Ivan Susloparov (State Research Center of Virology and Biotechnology (VECTOR) / Emerging Zoonotic Diseases and Influenza) |
| EPI_ISL_297324 | A/White-eyed Pochard/Ningxia/473-8/2015 | A / H5N6 | Asia / China | |
| EPI_ISL_297325 | A/White-eyed Pochard/Ningxia/477-14/2015 | A / H5N6 | Asia / China | |
| EPI_ISL_297326 | A/White-eyed Pochard/Ningxia/479-16/2015 | A / H5N6 | Asia / China | |
| EPI_ISL_297327 | A/White-eyed Pochard/Ningxia/480-17/2015 | A / H5N6 | Asia / China | |
| EPI_ISL_297388 | A/chicken/Republic of Macedonia/466/2017 | A / H5N8 | Europe / Macedonia, the former Yogoslav Republic of / Opstina Struga / N 41012’10, E20042’01 | Aleksandar Dodovski (Faculty of Veterinary Medicine Skopje / Department for Avian Diseases) |
| EPI_ISL_297463 | A/chicken/Kostroma/1717/2017 | A / H5N2 | Europe / Russian Federation / Kostroma Oblast | Ivan Susloparov (State Research Center of Virology and Biotechnology (VECTOR) / Emerging Zoonotic Diseases and Influenza) |
| EPI_ISL_297464 | A/chicken/Kostroma/1719/2017 | A / H5N2 | Europe / Russian Federation / Kostroma Oblast | Ivan Susloparov (State Research Center of Virology and Biotechnology (VECTOR) / Emerging Zoonotic Diseases and Influenza) |
| EPI_ISL_297465 | A/chicken/Kostroma/1720/2017 | A / H5N2 | Europe / Russian Federation / Kostroma Oblast | Ivan Susloparov (State Research Center of Virology and Biotechnology (VECTOR) / Emerging Zoonotic Diseases and Influenza) |
| EPI_ISL_297466 | A/chicken/Kostroma/1721/2017 | A / H5N2 | Europe / Russian Federation / Kostroma Oblast | Ivan Susloparov (State Research Center of Virology and Biotechnology (VECTOR) / Emerging Zoonotic Diseases and Influenza) |
| EPI_ISL_297930 | A/chicken/Anhui/MZ33/2016 | A / H5N6 | Asia / China | |
| EPI_ISL_297931 | A/chicken/Anhui/MZ34/2016 | A / H5N6 | Asia / China | |
| EPI_ISL_297932 | A/chicken/Henan/YB0597/2016 | A / H5N6 | Asia / China | |
| EPI_ISL_298640 | A/turkey/Israel/1045/2016 | A / H5N8 | Asia / Israel | |

| | | | | |
|---|---|---|---|---|
| EPI_ISL_298641 | A/chicken/Israel/881/2016 | A / H5N8 | Asia / Israel | |
| EPI_ISL_298642 | A/chicken/Israel/1048/2016 | A / H5N8 | Asia / Israel | |
| EPI_ISL_298647 | A/cormorant/Israel/1035/2016 | A / H5N8 | Asia / Israel | |
| EPI_ISL_298648 | A/great egret/Israel/1088/2016 | A / H5N8 | Asia / Israel | |
| EPI_ISL_298649 | A/turkey/Israel/184/2017 | A / H5N8 | Asia / Israel | |
| EPI_ISL_298650 | A/turkey/Israel/1076/2016 | A / H5N8 | Asia / Israel | |
| EPI_ISL_298651 | A/great egret/Israel/1084/2016 | A / H5N8 | Asia / Israel | |
| EPI_ISL_298652 | A/grey goose/Israel/986/2016 | A / H5N8 | Asia / Israel | |
| EPI_ISL_298653 | A/peregrine falcon/Israel/1086/2016 | A / H5N8 | Asia / Israel | |
| EPI_ISL_299583 | A/heron/Guangdong/C1/2013 | A / H5N6 | Asia / China | |
| EPI_ISL_3001273 | A/common buzzard/Sweden/SVA210510SZ0206/FB001829/C-2 | A / H5N8 | Europe / Sweden / Uppsala Lan / Uppsala Kommun | Siamak Zohari (National Veterinary Institute) |
| EPI_ISL_300547 | A/White-fronted Goose/AN/1-15-12/2016 | A / H5N8 | Europe / Ukraine | Susanne Koethe (Friedrich-Loeffler-Institut) |
| EPI_ISL_300548 | A/Ruddy Shelduck/AN/2-14-12/2016 | A / H5N8 | Europe / Ukraine / Kherson Oblast | Susanne Koethe (Friedrich-Loeffler-Institut) |
| EPI_ISL_300562 | A/Environmental (EM) /AN/2/17 | A / H5N8 | Europe / Ukraine / Kherson Oblast | Susanne Koethe (Friedrich-Loeffler-Institut) |
| EPI_ISL_3005950 | A/barnacle goose/Sweden/SVA210511SZ0567/FB001840/M-2( | A / H5N1 | Europe / Sweden / Skane Lan / Malmo Kommun | Siamak Zohari (National Veterinary Institute) |
| EPI_ISL_300642 | A/turkey/South Dakota/15-011769-1/2015 | A / H5N2 | North America / United States | |
| EPI_ISL_300643 | A/turkey/South Dakota/15-015102-5/2015 | A / H5N2 | North America / United States | |
| EPI_ISL_300644 | A/turkey/South Dakota/15-015103-1/2015 | A / H5N2 | North America / United States | |
| EPI_ISL_300645 | A/turkey/South Dakota/15-017406-1/2015 | A / H5N2 | North America / United States | |
| EPI_ISL_300646 | A/turkey/Wisconsin/15-012886-1/2015 | A / H5N2 | North America / United States | |
| EPI_ISL_300647 | A/turkey/Wisconsin/15-013180-1/2015 | A / H5N2 | North America / United States | |
| EPI_ISL_300648 | A/turkey/Wisconsin/15-013771-1/2015 | A / H5N2 | North America / United States | |
| EPI_ISL_300649 | A/turkey/Wisconsin/15-014296-1/2015 | A / H5N2 | North America / United States | |
| EPI_ISL_300650 | A/turkey/Wisconsin/15-014298-1/2015 | A / H5N2 | North America / United States | |
| EPI_ISL_300651 | A/turkey/Minnesota/15-017647-1/2015 | A / H5N2 | North America / United States | |
| EPI_ISL_300652 | A/turkey/Minnesota/15-017649-1/2015 | A / H5N2 | North America / United States | |
| EPI_ISL_300653 | A/turkey/Minnesota/15-017814-1/2015 | A / H5N2 | North America / United States | |
| EPI_ISL_300654 | A/turkey/Minnesota/15-017989-1/2015 | A / H5N2 | North America / United States | |
| EPI_ISL_300655 | A/turkey/Minnesota/15-018201-1/2015 | A / H5N2 | North America / United States | |
| EPI_ISL_300656 | A/turkey/Missouri/15-007513-1/2015 | A / H5N2 | North America / United States | |
| EPI_ISL_300657 | A/turkey/South Dakota/15-011089-3/2015 | A / H5N2 | North America / United States | |
| EPI_ISL_300658 | A/turkey/South Dakota/15-011415-1/2015 | A / H5N2 | North America / United States | |
| EPI_ISL_300659 | A/turkey/South Dakota/15-011417-1/2015 | A / H5N2 | North America / United States | |
| EPI_ISL_300660 | A/tundra swan/Niigata/1511C003/2016 | A / H5N6 | Asia / Japan / Niigata | |
| EPI_ISL_300661 | A/tundra swan/Niigata/5112004/2016 | A / H5N6 | Asia / Japan / Niigata | |
| EPI_ISL_300662 | A/tundra swan/Niigata/5112006/2016 | A / H5N6 | Asia / Japan / Niigata | |
| EPI_ISL_300663 | A/tundra swan/Niigata/5112007/2016 | A / H5N6 | Asia / Japan / Niigata | |
| EPI_ISL_300664 | A/mute swan/Hyogo/2801ITM015/2017 | A / H5N6 | Asia / Japan / Hyogo | |
| EPI_ISL_300665 | A/turkey/Minnesota/15-017046-1/2015 | A / H5N2 | North America / United States | |
| EPI_ISL_300666 | A/turkey/Minnesota/15-017217-1/2015 | A / H5N2 | North America / United States | |
| EPI_ISL_300667 | A/turkey/Minnesota/15-017218-1/2015 | A / H5N2 | North America / United States | |
| EPI_ISL_300668 | A/turkey/Minnesota/15-017219-1/2015 | A / H5N2 | North America / United States | |
| EPI_ISL_300669 | A/turkey/Minnesota/15-017409-1/2015 | A / H5N2 | North America / United States | |
| EPI_ISL_300670 | A/turkey/Minnesota/15-017410-1/2015 | A / H5N2 | North America / United States | |
| EPI_ISL_300671 | A/turkey/Minnesota/15-017540-1/2015 | A / H5N2 | North America / United States | |
| EPI_ISL_300672 | A/turkey/Minnesota/15-014851-1/2015 | A / H5N2 | North America / United States | |
| EPI_ISL_300673 | A/turkey/Minnesota/15-014972-1/2015 | A / H5N2 | North America / United States | |
| EPI_ISL_300674 | A/turkey/Minnesota/15-016050-1/2015 | A / H5N2 | North America / United States | |
| EPI_ISL_300675 | A/turkey/Minnesota/15-016876-1/2015 | A / H5N2 | North America / United States | |
| EPI_ISL_300676 | A/turkey/Minnesota/15-016877-1/2015 | A / H5N2 | North America / United States | |
| EPI_ISL_300677 | A/turkey/Minnesota/15-017039-1/2015 | A / H5N2 | North America / United States | |
| EPI_ISL_300678 | A/turkey/Minnesota/15-017040-1/2015 | A / H5N2 | North America / United States | |
| EPI_ISL_300679 | A/turkey/Minnesota/15-017044-1/2015 | A / H5N2 | North America / United States | |
| EPI_ISL_300680 | A/turkey/Minnesota/15-014299-1/2015 | A / H5N2 | North America / United States | |
| EPI_ISL_300681 | A/turkey/Minnesota/15-014301-1/2015 | A / H5N2 | North America / United States | |
| EPI_ISL_300682 | A/turkey/Minnesota/15-014310-1/2015 | A / H5N2 | North America / United States | |
| EPI_ISL_300683 | A/turkey/Minnesota/15-014311-1/2015 | A / H5N2 | North America / United States | |
| EPI_ISL_300684 | A/turkey/Minnesota/15-014381-1/2015 | A / H5N2 | North America / United States | |
| EPI_ISL_300685 | A/turkey/Minnesota/15-014850-1/2015 | A / H5N2 | North America / United States | |
| EPI_ISL_300686 | A/turkey/Poland/63/2016 | A / H5N8 | Europe / Poland / Lubusz Voivodeship | Edyta ?wi?to? (National Veterinary Research Institut Poland, PIWet-PIB) |
| EPI_ISL_300687 | A/chicken/Poland/34/2017 | A / H5N8 | Europe / Poland / Lesser Poland Voivodeship | Edyta ?wi?to? (National Veterinary Research Institut Poland, PIWet-PIB) |
| EPI_ISL_300688 | A/turkey/Minnesota/15-013755-2/2015 | A / H5N2 | North America / United States | |
| EPI_ISL_300689 | A/turkey/Minnesota/15-013895-1/2015 | A / H5N2 | North America / United States | |
| EPI_ISL_300690 | A/turkey/Minnesota/15-013896-1/2015 | A / H5N2 | North America / United States | |
| EPI_ISL_300691 | A/turkey/Minnesota/15-014095-1/2015 | A / H5N2 | North America / United States | |
| EPI_ISL_300692 | A/turkey/Minnesota/15-014097-1/2015 | A / H5N2 | North America / United States | |
| EPI_ISL_300693 | A/domestic_duck/Poland/47/2017 | A / H5N8 | Europe / Poland / Greater Poland Voivodeship | Edyta ?wi?to? (National Veterinary Research Institut Poland, PIWet-PIB) |
| EPI_ISL_300694 | A/turkey/Minnesota/15-014098-1/2015 | A / H5N2 | North America / United States | |
| EPI_ISL_300695 | A/turkey/Minnesota/15-014110-1/2015 | A / H5N2 | North America / United States | |
| EPI_ISL_300696 | A/turkey/Minnesota/15-014111-1/2015 | A / H5N2 | North America / United States | |
| EPI_ISL_300697 | A/turkey/Minnesota/15-014293-1/2015 | A / H5N2 | North America / United States | |
| EPI_ISL_300698 | A/turkey/Poland/54/2017 | A / H5N8 | Europe / Poland / Masovian Voivodeship | Edyta ?wi?to? (National Veterinary Research Institut Poland, PIWet-PIB) |

| | | | | |
|---|---|---|---|---|
| EPI_ISL_300699 | A/domestic_goose/Poland/69/2017 | A / H5N8 | Europe / Poland / Greater Poland Voivodeship | Edyta ?wi?to? (National Veterinary Research Institut Poland, PIWet-PIB) |
| EPI_ISL_300700 | A/turkey/Poland/72/2017 | A / H5N8 | Europe / Poland / Warmian-Masurian Voivedeship | Edyta ?wi?to? (National Veterinary Research Institut Poland, PIWet-PIB) |
| EPI_ISL_300701 | A/turkey/Poland/94/2017 | A / H5N8 | Europe / Poland / Greater Poland Voivodeship | Edyta ?wi?to? (National Veterinary Research Institut Poland, PIWet-PIB) |
| EPI_ISL_300702 | A/chicken/Poland/101/2017 | A / H5N8 | Europe / Poland / Lower Silesian Voivodeship | Edyta ?wi?to? (National Veterinary Research Institut Poland, PIWet-PIB) |
| EPI_ISL_300703 | A/domestic_goose/Poland/124/2017 | A / H5N8 | Europe / Poland / Lower Silesian Voivodeship | Edyta ?wi?to? (National Veterinary Research Institut Poland, PIWet-PIB) |
| EPI_ISL_300704 | A/turkey/Poland/192/2017 | A / H5N8 | Europe / Poland / Podlasie Voivodeship | Edyta ?wi?to? (National Veterinary Research Institut Poland, PIWet-PIB) |
| EPI_ISL_300705 | A/turkey/Poland/285/2017 | A / H5N8 | Europe / Poland / Greater Poland Voivodeship | Edyta ?wi?to? (National Veterinary Research Institut Poland, PIWet-PIB) |
| EPI_ISL_300706 | A/turkey/Minnesota/15-013181-1/2015 | A / H5N2 | North America / United States | |
| EPI_ISL_300707 | A/turkey/Minnesota/15-013391-1/2015 | A / H5N2 | North America / United States | |
| EPI_ISL_300708 | A/turkey/Minnesota/15-013392-1/2015 | A / H5N2 | North America / United States | |
| EPI_ISL_300709 | A/turkey/Minnesota/15-013393-1/2015 | A / H5N2 | North America / United States | |
| EPI_ISL_300710 | A/turkey/Minnesota/15-013395-1/2015 | A / H5N2 | North America / United States | |
| EPI_ISL_300711 | A/turkey/Minnesota/15-013398-1/2015 | A / H5N2 | North America / United States | |
| EPI_ISL_300712 | A/chicken/Poland/263/2017 | A / H5N8 | Europe / Poland / Lubusz Voivodeship | Edyta ?wi?to? (National Veterinary Research Institut Poland, PIWet-PIB) |
| EPI_ISL_300713 | A/turkey/Minnesota/15-013399-1/2015 | A / H5N2 | North America / United States | |
| EPI_ISL_300714 | A/turkey/Minnesota/15-013401-1/2015 | A / H5N2 | North America / United States | |
| EPI_ISL_300715 | A/turkey/Minnesota/15-013534-1/2015 | A / H5N2 | North America / United States | |
| EPI_ISL_300716 | A/turkey/Minnesota/15-012659-2/2015 | A / H5N2 | North America / United States | |
| EPI_ISL_300717 | A/turkey/Minnesota/15-012661-1/2015 | A / H5N2 | North America / United States | |
| EPI_ISL_300718 | A/turkey/Minnesota/15-012663-1/2015 | A / H5N2 | North America / United States | |
| EPI_ISL_300719 | A/turkey/Minnesota/15-012666-1/2015 | A / H5N2 | North America / United States | |
| EPI_ISL_300720 | A/turkey/Minnesota/15-012887-1/2015 | A / H5N2 | North America / United States | |
| EPI_ISL_300721 | A/turkey/Minnesota/15-012888-1/2015 | A / H5N2 | North America / United States | |
| EPI_ISL_300722 | A/turkey/Minnesota/15-012889-1/2015 | A / H5N2 | North America / United States | |
| EPI_ISL_300723 | A/turkey/Minnesota/15-013050-1/2015 | A / H5N2 | North America / United States | |
| EPI_ISL_300724 | A/turkey/Minnesota/15-013051-1/2015 | A / H5N2 | North America / United States | |
| EPI_ISL_300725 | A/turkey/Minnesota/15-013052-1/2015 | A / H5N2 | North America / United States | |
| EPI_ISL_300726 | A/turkey/Minnesota/15-012576-1/2015 | A / H5N2 | North America / United States | |
| EPI_ISL_300727 | A/turkey/Minnesota/15-012577-1/2015 | A / H5N2 | North America / United States | |
| EPI_ISL_300728 | A/turkey/Minnesota/15-012578-4/2015 | A / H5N2 | North America / United States | |
| EPI_ISL_300729 | A/turkey/Minnesota/15-012580-4/2015 | A / H5N2 | North America / United States | |
| EPI_ISL_300730 | A/turkey/Minnesota/15-012583-1/2015 | A / H5N2 | North America / United States | |
| EPI_ISL_300731 | A/turkey/Minnesota/15-012584-1/2015 | A / H5N2 | North America / United States | |
| EPI_ISL_300732 | A/turkey/Minnesota/15-012585-1/2015 | A / H5N2 | North America / United States | |
| EPI_ISL_300733 | A/turkey/Minnesota/15-012658-1/2015 | A / H5N2 | North America / United States | |
| EPI_ISL_300734 | A/turkey/Minnesota/15-011834-1/2015 | A / H5N2 | North America / United States | |
| EPI_ISL_300735 | A/turkey/Minnesota/15-011849-1/2015 | A / H5N2 | North America / United States | |
| EPI_ISL_300736 | A/turkey/Minnesota/15-011923-1/2015 | A / H5N2 | North America / United States | |
| EPI_ISL_300737 | A/turkey/Minnesota/15-012289-2/2015 | A / H5N2 | North America / United States | |
| EPI_ISL_300738 | A/turkey/Minnesota/15-012508-2/2015 | A / H5N2 | North America / United States | |
| EPI_ISL_300739 | A/turkey/Minnesota/15-012509-1/2015 | A / H5N2 | North America / United States | |
| EPI_ISL_300740 | A/turkey/Minnesota/15-012510-4/2015 | A / H5N2 | North America / United States | |
| EPI_ISL_300741 | A/turkey/Minnesota/15-012574-1/2015 | A / H5N2 | North America / United States | |
| EPI_ISL_300742 | A/turkey/Minnesota/15-012575-1/2015 | A / H5N2 | North America / United States | |
| EPI_ISL_300743 | A/mallard/Poland/17/2017 | A / H5N8 | Europe / Poland / West Pomeranian Voivodeship | Edyta ?wi?to? (National Veterinary Research Institut Poland, PIWet-PIB) |
| EPI_ISL_300744 | A/swan/Poland/32/2017 | A / H5N8 | Europe / Poland / Lesser Poland Voivodeship | Edyta ?wi?to? (National Veterinary Research Institut Poland, PIWet-PIB) |
| EPI_ISL_300745 | A/wild duck/Poland/57/2017 | A / H5N8 | Europe / Poland / Lubusz Voivodeship | Edyta ?wi?to? (National Veterinary Research Institut Poland, PIWet-PIB) |
| EPI_ISL_300746 | A/swan/Poland/81/2017 | A / H5N5 | Europe / Poland / Lower Silesian Voivodeship | Edyta ?wi?to? (National Veterinary Research Institut Poland, PIWet-PIB) |
| EPI_ISL_300747 | A/mute_swan/Poland/137/2017 | A / H5N8 | Europe / Poland / Pomeranian Voivodeship | Edyta ?wi?to? (National Veterinary Research Institut Poland, PIWet-PIB) |
| EPI_ISL_300749 | A/turkey/Poland/81/2016 | A / H5N8 | Europe / Poland / Lubusz Voivodeship | Edyta ?wi?to? (National Veterinary Research Institut Poland, PIWet-PIB) |
| EPI_ISL_300750 | A/domestic_duck/Poland/88/2016 | A / H5N8 | Europe / Poland / Lubusz Voivodeship | Edyta ?wi?to? (National Veterinary Research Institut Poland, PIWet-PIB) |
| EPI_ISL_300783 | A/turkey/Poland/89/2016 | A / H5N8 | Europe / Poland / Lubusz Voivodeship | Edyta ?wi?to? (National Veterinary Research Institut Poland, PIWet-PIB) |
| EPI_ISL_300808 | A/turkey/Poland/93/2016 | A / H5N8 | Europe / Poland / Lubusz Voivodeship | Edyta ?wi?to? (National Veterinary Research Institut Poland, PIWet-PIB) |
| EPI_ISL_300841 | A/turkey/Poland/100/2016 | A / H5N8 | Europe / Poland / Lubusz Voivodeship | Edyta ?wi?to? (National Veterinary Research Institut Poland, PIWet-PIB) |
| EPI_ISL_300872 | A/chicken/Poland/103/2016 | A / H5N8 | Europe / Poland / Swietokrzyskie Voivodeship | Edyta ?wi?to? (National Veterinary Research Institut Poland, PIWet-PIB) |
| EPI_ISL_300898 | A/turkey/Poland/107/2016 | A / H5N8 | Europe / Poland / Lubusz Voivodeship | Edyta ?wi?to? (National Veterinary Research Institut Poland, PIWet-PIB) |
| EPI_ISL_300920 | A/domestic_duck/Poland/112/2016 | A / H5N8 | Europe / Poland / Lubusz Voivodeship | Edyta ?wi?to? (National Veterinary Research Institut Poland, PIWet-PIB) |
| EPI_ISL_300939 | A/chicken/Poland/116/2016 | A / H5N8 | Europe / Poland / Subcarpathian Voivodeship | Edyta ?wi?to? (National Veterinary Research Institut Poland, PIWet-PIB) |
| EPI_ISL_300978 | A/chicken/Poland/117/2016 | A / H5N8 | Europe / Poland / Swietokrzyskie Voivodeship | Edyta ?wi?to? (National Veterinary Research Institut Poland, PIWet-PIB) |
| EPI_ISL_300979 | A/turkey/Poland/004/2017 | A / H5N8 | Europe / Poland / Lubusz Voivodeship | Edyta ?wi?to? (National Veterinary Research Institut Poland, PIWet-PIB) |
| EPI_ISL_300980 | A/turkey/Poland/005/2017 | A / H5N8 | Europe / Poland / Lubusz Voivodeship | Edyta ?wi?to? (National Veterinary Research Institut Poland, PIWet-PIB) |
| EPI_ISL_300981 | A/turkey/Poland/15/2017 | A / H5N8 | Europe / Poland / Lubusz Voivodeship | Edyta ?wi?to? (National Veterinary Research Institut Poland, PIWet-PIB) |
| EPI_ISL_301037 | A/chicken/Poland/16/2017 | A / H5N8 | Europe / Poland / Opole Voivodeship | Edyta ?wi?to? (National Veterinary Research Institut Poland, PIWet-PIB) |
| EPI_ISL_301038 | A/chicken/Poland/77/2017 | A / H5N8 | Europe / Poland / Lesser Poland Voivodeship | Edyta ?wi?to? (National Veterinary Research Institut Poland, PIWet-PIB) |
| EPI_ISL_301039 | A/domestic_goose/Poland/190/2017 | A / H5N8 | Europe / Poland / Greater Poland Voivodeship | Edyta ?wi?to? (National Veterinary Research Institut Poland, PIWet-PIB) |
| EPI_ISL_301047 | A/chicken/Poland/199/2017 | A / H5N8 | Europe / Poland / Masovian Voivodeship | Edyta ?wi?to? (National Veterinary Research Institut Poland, PIWet-PIB) |
| EPI_ISL_301048 | A/chicken/Poland/208/2017 | A / H5N8 | Europe / Poland / Kuyavian-Pomeranian Voivodeship | Edyta ?wi?to? (National Veterinary Research Institut Poland, PIWet-PIB) |
| EPI_ISL_301049 | A/domestic_goose/Poland/270/2017 | A / H5N8 | Europe / Poland / Greater Poland Voivodeship | Edyta ?wi?to? (National Veterinary Research Institut Poland, PIWet-PIB) |
| EPI_ISL_301050 | A/mute_swan/Poland/12/2017 | A / H5N8 | Europe / Poland / Lower Silesian Voivodeship | Edyta ?wi?to? (National Veterinary Research Institut Poland, PIWet-PIB) |
| EPI_ISL_301051 | A/swan/Poland/23/2017 | A / H5N8 | Europe / Poland / Lower Silesian Voivodeship | Edyta ?wi?to? (National Veterinary Research Institut Poland, PIWet-PIB) |
| EPI_ISL_301052 | A/mute_swan/Poland/30/2017 | A / H5N8 | Europe / Poland / Lower Silesian Voivodeship | Edyta ?wi?to? (National Veterinary Research Institut Poland, PIWet-PIB) |
| EPI_ISL_301053 | A/mallard/Poland/33/2017 | A / H5N8 | Europe / Poland / Masovian Voivodeship | Edyta ?wi?to? (National Veterinary Research Institut Poland, PIWet-PIB) |

| | | | | |
|---|---|---|---|---|
| EPI_ISL_301054 | A/swan/Poland/49/2017 | A / H5N8 | Europe / Poland / Lower Silesian Voivodeship | Edyta ?wi?to? (National Veterinary Research Institut Poland, PIWet-PIB) |
| EPI_ISL_301055 | A/mute_swan/Poland/54/2017 | A / H5N8 | Europe / Poland / Lodz Voivodeship | Edyta ?wi?to? (National Veterinary Research Institut Poland, PIWet-PIB) |
| EPI_ISL_301056 | A/swan/Poland/56/2017 | A / H5N8 | Europe / Poland / Lower Silesian Voivodeship | Edyta ?wi?to? (National Veterinary Research Institut Poland, PIWet-PIB) |
| EPI_ISL_301057 | A/mute_swan/Poland/68/2017 | A / H5N8 | Europe / Poland / Lower Silesian Voivodeship | Edyta ?wi?to? (National Veterinary Research Institut Poland, PIWet-PIB) |
| EPI_ISL_301058 | A/turkey/Minnesota/15-011602-1/2015 | A / H5N2 | North America / United States | |
| EPI_ISL_301059 | A/turkey/Minnesota/15-011603-1/2015 | A / H5N2 | North America / United States | |
| EPI_ISL_301060 | A/turkey/Minnesota/15-011656-1/2015 | A / H5N2 | North America / United States | |
| EPI_ISL_301061 | A/turkey/Minnesota/15-011661-1/2015 | A / H5N2 | North America / United States | |
| EPI_ISL_301062 | A/turkey/Minnesota/15-011666-1/2015 | A / H5N2 | North America / United States | |
| EPI_ISL_301063 | A/mute_swan/Poland/72/2017 | A / H5N8 | Europe / Poland / Lubusz Voivodeship | Edyta ?wi?to? (National Veterinary Research Institut Poland, PIWet-PIB) |
| EPI_ISL_301064 | A/turkey/Minnesota/15-011668-1/2015 | A / H5N2 | North America / United States | |
| EPI_ISL_301065 | A/turkey/Minnesota/15-011669-1/2015 | A / H5N2 | North America / United States | |
| EPI_ISL_301066 | A/turkey/Minnesota/15-011833-1/2015 | A / H5N2 | North America / United States | |
| EPI_ISL_301067 | A/mute_swan/Poland/76/2017 | A / H5N8 | Europe / Poland / Masovian Voivodeship | Edyta ?wi?to? (National Veterinary Research Institut Poland, PIWet-PIB) |
| EPI_ISL_301068 | A/wild_duck/Poland/78/2017 | A / H5N8 | Europe / Poland / Lower Silesian Voivodeship | Edyta ?wi?to? (National Veterinary Research Institut Poland, PIWet-PIB) |
| EPI_ISL_301069 | A/swan/Poland/88/2017 | A / H5N8 | Europe / Poland / Greater Poland Voivodeship | Edyta ?wi?to? (National Veterinary Research Institut Poland, PIWet-PIB) |
| EPI_ISL_301070 | A/swan/Poland/99/2017 | A / H5N8 | Europe / Poland / Greater Poland Voivodeship | Edyta ?wi?to? (National Veterinary Research Institut Poland, PIWet-PIB) |
| EPI_ISL_301071 | A/turkey/Minnesota/15-010856-1/2015 | A / H5N2 | North America / United States | |
| EPI_ISL_301072 | A/swan/Poland/107/2017 | A / H5N8 | Europe / Poland / Kuyavian-Pomeranian Voivodeship | Edyta ?wi?to? (National Veterinary Research Institut Poland, PIWet-PIB) |
| EPI_ISL_301073 | A/turkey/Minnesota/15-010915-1/2015 | A / H5N2 | North America / United States | |
| EPI_ISL_301074 | A/turkey/Minnesota/15-011079-1/2015 | A / H5N2 | North America / United States | |
| EPI_ISL_301075 | A/turkey/Minnesota/15-011200-1/2015 | A / H5N2 | North America / United States | |
| EPI_ISL_301076 | A/turkey/Minnesota/15-011202-1/2015 | A / H5N2 | North America / United States | |
| EPI_ISL_301077 | A/turkey/Minnesota/15-011591-1/2015 | A / H5N2 | North America / United States | |
| EPI_ISL_301078 | A/mute_swan/Poland/109/2017 | A / H5N8 | Europe / Poland / Kuyavian-Pomeranian Voivodeship | Edyta ?wi?to? (National Veterinary Research Institut Poland, PIWet-PIB) |
| EPI_ISL_301079 | A/turkey/Minnesota/15-011593-1/2015 | A / H5N2 | North America / United States | |
| EPI_ISL_301080 | A/turkey/Minnesota/15-011596-1/2015 | A / H5N2 | North America / United States | |
| EPI_ISL_301081 | A/turkey/Iowa/15-017224-1/2015 | A / H5N2 | North America / United States | |
| EPI_ISL_301082 | A/turkey/Iowa/15-017421-1/2015 | A / H5N2 | North America / United States | |
| EPI_ISL_301083 | A/turkey/Iowa/15-017556-1/2015 | A / H5N2 | North America / United States | |
| EPI_ISL_301084 | A/turkey/Iowa/15-017557-1/2015 | A / H5N2 | North America / United States | |
| EPI_ISL_301085 | A/turkey/Iowa/15-017655-1/2015 | A / H5N2 | North America / United States | |
| EPI_ISL_301086 | A/turkey/Iowa/15-017824-1/2015 | A / H5N2 | North America / United States | |
| EPI_ISL_301087 | A/turkey/Minnesota/15-010375-2/2015 | A / H5N2 | North America / United States | |
| EPI_ISL_301088 | A/turkey/Minnesota/15-010560-1/2015 | A / H5N2 | North America / United States | |
| EPI_ISL_301089 | A/turkey/Minnesota/15-010777-1/2015 | A / H5N2 | North America / United States | |
| EPI_ISL_301090 | A/turkey/Iowa/15-014868-1/2015 | A / H5N2 | North America / United States | |
| EPI_ISL_301091 | A/turkey/Iowa/15-014869-1/2015 | A / H5N2 | North America / United States | |
| EPI_ISL_301092 | A/turkey/Iowa/15-014954-1/2015 | A / H5N2 | North America / United States | |
| EPI_ISL_301093 | A/turkey/Iowa/15-015071-3/2015 | A / H5N2 | North America / United States | |
| EPI_ISL_301094 | A/turkey/Iowa/15-015116-1/2015 | A / H5N2 | North America / United States | |
| EPI_ISL_301095 | A/turkey/Iowa/15-016077-1/2015 | A / H5N2 | North America / United States | |
| EPI_ISL_301096 | A/turkey/Iowa/15-016082-1/2015 | A / H5N2 | North America / United States | |
| EPI_ISL_301097 | A/turkey/Iowa/15-016365-1/2015 | A / H5N2 | North America / United States | |
| EPI_ISL_301098 | A/turkey/Iowa/15-016743-1/2015 | A / H5N2 | North America / United States | |
| EPI_ISL_301099 | A/turkey/Iowa/15-014789-1/2015 | A / H5N2 | North America / United States | |
| EPI_ISL_301100 | A/turkey/Iowa/15-014862-1/2015 | A / H5N2 | North America / United States | |
| EPI_ISL_301101 | A/turkey/Iowa/15-014863-1/2015 | A / H5N2 | North America / United States | |
| EPI_ISL_301102 | A/turkey/Iowa/15-014864-1/2015 | A / H5N2 | North America / United States | |
| EPI_ISL_301103 | A/turkey/Iowa/15-013773-1/2015 | A / H5N2 | North America / United States | |
| EPI_ISL_301104 | A/turkey/Iowa/15-013926-1/2015 | A / H5N2 | North America / United States | |
| EPI_ISL_301105 | A/turkey/Iowa/15-014119-1/2015 | A / H5N2 | North America / United States | |
| EPI_ISL_301106 | A/turkey/Iowa/15-014120-1/2015 | A / H5N2 | North America / United States | |
| EPI_ISL_301107 | A/turkey/Iowa/15-014562-2/2015 | A / H5N2 | North America / United States | |
| EPI_ISL_301108 | A/turkey/Iowa/15-014570-1/2015 | A / H5N2 | North America / United States | |
| EPI_ISL_301109 | A/turkey/Iowa/15-014772-1/2015 | A / H5N2 | North America / United States | |
| EPI_ISL_301110 | A/turkey/Iowa/15-013179-4/2015 | A / H5N2 | North America / United States | |
| EPI_ISL_301111 | A/great-horned owl/Idaho/15-001583-11/2015 | A / H5N2 | North America / United States | |
| EPI_ISL_301112 | A/guinea fowl/Iowa/15-018413-1/2015 | A / H5N2 | North America / United States | |
| EPI_ISL_301113 | A/peregrine falcon/Idaho/15-001362-3/2015 | A / H5N2 | North America / United States | |
| EPI_ISL_301114 | A/red-tailed hawk/Washington/15-002551-2/2015 | A / H5N2 | North America / United States | |
| EPI_ISL_301115 | A/ring-necked duck/Kentucky/15-012967-1/2015 | A / H5N2 | North America / United States | |
| EPI_ISL_301116 | A/snow goose/Kentucky/15-012967-2/2015 | A / H5N2 | North America / United States | |
| EPI_ISL_301117 | A/great-horned owl/Idaho/15-003155-1/2015 | A / H5N2 | North America / United States | |
| EPI_ISL_301545 | A/chicken/Nebraska/15-016865-1/2015 | A / H5N2 | North America / United States | |
| EPI_ISL_301546 | A/chicken/Nebraska/15-019197-1/2015 | A / H5N2 | North America / United States | |
| EPI_ISL_301547 | A/chicken/Oregon/15-005256-1/2015 | A / H5N2 | North America / United States | |
| EPI_ISL_301548 | A/chicken/South Dakota/15-015847-3/2015 | A / H5N2 | North America / United States | |
| EPI_ISL_301549 | A/chicken/Wisconsin/15-013062-1/2015 | A / H5N2 | North America / United States | |
| EPI_ISL_301550 | A/chicken/Wisconsin/15-014400-1/2015 | A / H5N2 | North America / United States | |
| EPI_ISL_301551 | A/falcon/Missouri/15-009166-2/2015 | A / H5N2 | North America / United States | |
| EPI_ISL_301552 | A/goose/Washington/15-001678-1/2015 | A / H5N2 | North America / United States | |

| | | | | |
|---|---|---|---|---|
| EPI_ISL_301553 | A/chicken/Iowa/15-019293-1/2015 | A / H5N2 | North America / United States | |
| EPI_ISL_301554 | A/chicken/Minnesota/15-012694-1/2015 | A / H5N2 | North America / United States | |
| EPI_ISL_301555 | A/chicken/Minnesota/15-012882-2/2015 | A / H5N2 | North America / United States | |
| EPI_ISL_301556 | A/chicken/Minnesota/15-014297-1/2015 | A / H5N2 | North America / United States | |
| EPI_ISL_301557 | A/chicken/Minnesota/15-015844-1/2015 | A / H5N2 | North America / United States | |
| EPI_ISL_301558 | A/chicken/Minnesota/15-017817-1/2015 | A / H5N2 | North America / United States | |
| EPI_ISL_301559 | A/chicken/Missouri/15-014275-1/2015 | A / H5N2 | North America / United States | |
| EPI_ISL_301560 | A/chicken/Nebraska/15-015085-1/2015 | A / H5N2 | North America / United States | |
| EPI_ISL_301561 | A/chicken/Nebraska/15-015543-1/2015 | A / H5N2 | North America / United States | |
| EPI_ISL_301562 | A/chicken/Iowa/15-016078-1/2015 | A / H5N2 | North America / United States | |
| EPI_ISL_301563 | A/chicken/Iowa/15-016158-1/2015 | A / H5N2 | North America / United States | |
| EPI_ISL_301564 | A/chicken/Iowa/15-016159-1/2015 | A / H5N2 | North America / United States | |
| EPI_ISL_301565 | A/chicken/Iowa/15-016160-1/2015 | A / H5N2 | North America / United States | |
| EPI_ISL_301566 | A/chicken/Iowa/15-016887-1/2015 | A / H5N2 | North America / United States | |
| EPI_ISL_301567 | A/chicken/Iowa/15-016888-1/2015 | A / H5N2 | North America / United States | |
| EPI_ISL_301568 | A/chicken/Iowa/15-017038-1/2015 | A / H5N2 | North America / United States | |
| EPI_ISL_301569 | A/chicken/Iowa/15-017419-1/2015 | A / H5N2 | North America / United States | |
| EPI_ISL_301570 | A/chicken/Iowa/15-017656-1/2015 | A / H5N2 | North America / United States | |
| EPI_ISL_301571 | A/chicken/Iowa/15-014774-1/2015 | A / H5N2 | North America / United States | |
| EPI_ISL_301572 | A/chicken/Iowa/15-015016-1/2015 | A / H5N2 | North America / United States | |
| EPI_ISL_301573 | A/chicken/Iowa/15-015018-1/2015 | A / H5N2 | North America / United States | |
| EPI_ISL_301574 | A/chicken/Iowa/15-015073-1/2015 | A / H5N2 | North America / United States | |
| EPI_ISL_301575 | A/chicken/Iowa/15-015117-1/2015 | A / H5N2 | North America / United States | |
| EPI_ISL_301576 | A/chicken/Iowa/15-015300-1/2015 | A / H5N2 | North America / United States | |
| EPI_ISL_301577 | A/chicken/Iowa/15-015544-2/2015 | A / H5N2 | North America / United States | |
| EPI_ISL_301578 | A/chicken/Iowa/15-015555-1/2015 | A / H5N2 | North America / United States | |
| EPI_ISL_301579 | A/chicken/Iowa/15-015556-1/2015 | A / H5N2 | North America / United States | |
| EPI_ISL_301580 | A/chicken/Iowa/15-013982-1/2015 | A / H5N2 | North America / United States | |
| EPI_ISL_301581 | A/chicken/Iowa/15-014283-1/2015 | A / H5N2 | North America / United States | |
| EPI_ISL_301582 | A/chicken/Iowa/15-014294-1/2015 | A / H5N2 | North America / United States | |
| EPI_ISL_301583 | A/chicken/Iowa/15-014572-1/2015 | A / H5N2 | North America / United States | |
| EPI_ISL_301584 | A/chicken/Iowa/15-014586-1/2015 | A / H5N2 | North America / United States | |
| EPI_ISL_301585 | A/chicken/Iowa/15-014591-1/2015 | A / H5N2 | North America / United States | |
| EPI_ISL_301586 | A/chicken/Iowa/15-014719-1/2015 | A / H5N2 | North America / United States | |
| EPI_ISL_301587 | A/chicken/Iowa/15-014769-1/2015 | A / H5N2 | North America / United States | |
| EPI_ISL_301588 | A/chicken/Idaho/15-001711-7/2015 | A / H5N2 | North America / United States | |
| EPI_ISL_301589 | A/chicken/Iowa/15-012564-7/2015 | A / H5N2 | North America / United States | |
| EPI_ISL_301590 | A/chicken/Iowa/15-013408-3/2015 | A / H5N2 | North America / United States | |
| EPI_ISL_301591 | A/chicken/Iowa/15-013430-16/2015 | A / H5N2 | North America / United States | |
| EPI_ISL_301592 | A/chicken/Iowa/15-013540-6/2015 | A / H5N2 | North America / United States | |
| EPI_ISL_301593 | A/chicken/Iowa/15-013784-2/2015 | A / H5N2 | North America / United States | |
| EPI_ISL_301594 | A/chicken/Iowa/15-013972-1/2015 | A / H5N2 | North America / United States | |
| EPI_ISL_301595 | A/chicken/Iowa/15-013976-6/2015 | A / H5N2 | North America / United States | |
| EPI_ISL_301596 | A/Canada goose/Wyoming/15-009285-1/2015 | A / H5N2 | North America / United States | |
| EPI_ISL_301611 | A/Canada goose/Michigan/15-018340-2/2015 | A / H5N2 | North America / United States | |
| EPI_ISL_301612 | A/Canada goose/Michigan/15-019873-5/2015 | A / H5N2 | North America / United States | |
| EPI_ISL_301782 | A/white-tailed_eagle/Denmark/3073-1w/2018-02-13 | A / H5N6 | Europe / Denmark / Region Sjaland / Slagelse Kommune | Charlotte Kristiane Hjulsager (Statens Serum Institute / Microbiological Diagnostic and Virology) |
| EPI_ISL_301784 | A/Mallard/Korea/K17-1825/2017 | A / H5N6 | Asia / Korea, Republic of / Gyeonggi-do / Cheongmi-cheon (37°06'56.9"N 127°25'18.3"E) | Jung Hoon Kwon (Konkuk University / College of Veterinary Medicine) |
| EPI_ISL_301785 | A/Mandarin_duck/Korea/K17-1815/2017 | A / H5N6 | Asia / Korea, Republic of / Gyeonggi-do / Cheongmi-cheon (37°06'56.9"N 127°25'18.3"E) | Jung Hoon Kwon (Konkuk University / College of Veterinary Medicine) |
| EPI_ISL_301786 | A/Mandarin_duck/Korea/K17-1817/2017 | A / H5N6 | Asia / Korea, Republic of / Gyeonggi-do / Cheongmi-cheon (37°06'56.9"N 127°25'18.3"E) | Jung Hoon Kwon (Konkuk University / College of Veterinary Medicine) |
| EPI_ISL_301787 | A/Mandarin_duck/Korea/K17-1826/2017 | A / H5N6 | Asia / Korea, Republic of / Gyeonggi-do / Cheongmi-cheon (37°06'56.9"N 127°25'18.3"E) | Jung Hoon Kwon (Konkuk University / College of Veterinary Medicine) |
| EPI_ISL_301803 | A/Mandarin_duck/Korea/K17-1828/2017 | A / H5N6 | Asia / Korea, Republic of / Gyeonggi-do / Cheongmi-cheon (37°06'56.9"N 127°25'18.3"E) | Jung Hoon Kwon (Konkuk University / College of Veterinary Medicine) |
| EPI_ISL_302523 | A/Anas platyrhynchos/Korea/W612/2017 | A / H5N6 | Asia / Korea, Republic of | |
| EPI_ISL_302524 | A/Anas platyrhynchos/Korea/W613/2017 | A / H5N6 | Asia / Korea, Republic of | |
| EPI_ISL_302525 | A/Anas platyrhynchos/Korea/W614/2017 | A / H5N6 | Asia / Korea, Republic of | |
| EPI_ISL_302526 | A/Anas platyrhynchos/Korea/W615/2017 | A / H5N6 | Asia / Korea, Republic of | |
| EPI_ISL_3026043 | A/European herring gull/Sweden/SVA210617SZ0354/FB00225 | A / H5N8 | Europe / Sweden / Gotlands Lan / Gotlands Kommun | Siamak Zohari (National Veterinary Institute) |
| EPI_ISL_3026699 | A/common eider/Sweden/SVA210617SZ0354/FB002252/I-202 | A / H5N8 | Europe / Sweden / Gotlands Lan / Gotlands Kommun | Siamak Zohari (National Veterinary Institute) |
| EPI_ISL_302708 | A/Great_Cormorant/Hubei/chenhu_V11/2015 | A / H5N6 | Asia / China / Hubei Province | Jianjun Chen (Wuhan Institute of Virology / Chinese Academy of Sciences) |
| EPI_ISL_302823 | A/Great black-backed gull/Netherlands/1/2018 | A / H5N6 | Europe / Netherlands / Provincie Noord-Holland / De Kreupel | Maria Johanna Poen (Erasmus Medical Center / Department of Virology) |
| EPI_ISL_302824 | A/Eurasian wigeon/Netherlands/1/2018 | A / H5N6 | Europe / Netherlands / Provincie Noord-Holland / Hippolytushoef | Maria Johanna Poen (Erasmus Medical Center / Department of Virology) |
| EPI_ISL_302825 | A/Chicken/Netherlands/EMC-1/2018 | A / H5N6 | Europe / Netherlands / Provincie Groningen / Gemeente Grootegast | Maria Johanna Poen (Erasmus Medical Center / Department of Virology) |
| EPI_ISL_302826 | A/Chicken/Netherlands/EMC-14/2018 | A / H5N6 | Europe / Netherlands / Provincie Groningen / Gemeente Grootegast | Maria Johanna Poen (Erasmus Medical Center / Department of Virology) |
| EPI_ISL_303520 | A/Mallard/Republic of Georgia/1/2018 | A / H5N6 | Asia / Georgia | Maria Johanna Poen (Erasmus Medical Center / Department of Virology) |
| EPI_ISL_303630 | A/Mandarin_duck/Korea/K17-1862/2017 | A / H5N6 | Asia / Korea, Republic of / Chungcheongnam-do / Gokgyo-cheon (36°45?12.3?N, 127°07?12.7?E | Jung Hoon Kwon (Konkuk University / College of Veterinary Medicine) |
| EPI_ISL_303631 | A/Mandarin_duck/Korea/K17-1866/2017 | A / H5N6 | Asia / Korea, Republic of / Chungcheongnam-do / Gokgyo-cheon (36°45?12.3?N, 127°07?12.7?E | Jung Hoon Kwon (Konkuk University / College of Veterinary Medicine) |
| EPI_ISL_303632 | A/Mandarin_duck/Korea/K17-1869/2017 | A / H5N6 | Asia / Korea, Republic of / Chungcheongnam-do / Gokgyo-cheon (36°45?12.3?N, 127°07?12.7?E | Jung Hoon Kwon (Konkuk University / College of Veterinary Medicine) |
| EPI_ISL_303633 | A/Mandarin_duck/Korea/K17-1873/2017 | A / H5N6 | Asia / Korea, Republic of / Chungcheongnam-do / Gokgyo-cheon (36°45?12.3?N, 127°07?12.7?E | Jung Hoon Kwon (Konkuk University / College of Veterinary Medicine) |
| EPI_ISL_303634 | A/Mandarin_duck/Korea/K17-1879/2017 | A / H5N6 | Asia / Korea, Republic of / Chungcheongnam-do / Gokgyo-cheon (36°45?12.3?N, 127°07?12.7?E | Jung Hoon Kwon (Konkuk University / College of Veterinary Medicine) |
| EPI_ISL_303635 | A/Mandarin_duck/Korea/K17-1881/2017 | A / H5N6 | Asia / Korea, Republic of / Chungcheongnam-do / Gokgyo-cheon (36°45?12.3?N, 127°07?12.7?E | Jung Hoon Kwon (Konkuk University / College of Veterinary Medicine) |
| EPI_ISL_303636 | A/Mandarin_duck/Korea/K17-1885/2017 | A / H5N6 | Asia / Korea, Republic of / Chungcheongnam-do / Gokgyo-cheon (36°45?12.3?N, 127°07?12.7?E | Jung Hoon Kwon (Konkuk University / College of Veterinary Medicine) |
| EPI_ISL_303637 | A/Mandarin_duck/Korea/K17-1887/2017 | A / H5N6 | Asia / Korea, Republic of / Chungcheongnam-do / Gokgyo-cheon (36°45?12.3?N, 127°07?12.7?E | Jung Hoon Kwon (Konkuk University / College of Veterinary Medicine) |

| | | | | |
|---|---|---|---|---|
| EPI_ISL_303638 | A/Mandarin_duck/Korea/K17-1889/2017 | A / H5N6 | Asia / Korea, Republic of / Chungcheongnam-do / Gokgyo-cheon (36°45?12.3?N, 127°07?12.7?E | Jung Hoon Kwon (Konkuk University / College of Veterinary Medicine) |
| EPI_ISL_303639 | A/Mandarin_duck/Korea/K17-1891/2017 | A / H5N6 | Asia / Korea, Republic of / Chungcheongnam-do / Gokgyo-cheon (36°45?12.3?N, 127°07?12.7?E | Jung Hoon Kwon (Konkuk University / College of Veterinary Medicine) |
| EPI_ISL_303640 | A/Mandarin_duck/Korea/K17-1893/2017 | A / H5N6 | Asia / Korea, Republic of / Chungcheongnam-do / Gokgyo-cheon (36°45?12.3?N, 127°07?12.7?E | Jung Hoon Kwon (Konkuk University / College of Veterinary Medicine) |
| EPI_ISL_303641 | A/Mandarin_duck/Korea/K17-1894/2017 | A / H5N6 | Asia / Korea, Republic of / Chungcheongnam-do / Gokgyo-cheon (36°45?12.3?N, 127°07?12.7?E | Jung Hoon Kwon (Konkuk University / College of Veterinary Medicine) |
| EPI_ISL_303642 | A/Mandarin_duck/Korea/K17-1896/2017 | A / H5N6 | Asia / Korea, Republic of / Chungcheongnam-do / Gokgyo-cheon (36°45?12.3?N, 127°07?12.7?E | Jung Hoon Kwon (Konkuk University / College of Veterinary Medicine) |
| EPI_ISL_303643 | A/Mandarin_duck/Korea/K18-3/2018 | A / H5N6 | Asia / Korea, Republic of / Chungcheongnam-do / Gokgyo-cheon (36°45?12.3?N, 127°07?12.7?E | Jung Hoon Kwon (Konkuk University / College of Veterinary Medicine) |
| EPI_ISL_303837 | A/grey-headed gull/Uganda/MUWRP-538/2017 | A / H5N8 | Africa / Uganda | |
| EPI_ISL_304404 | A/Fujian-Sanyuan/21099/2017 | A / H5N6 | Asia / China / Fujian Province | Lei Yang (WHO Chinese National Influenza Center / Virology Institute, Chinese CDC) |
| EPI_ISL_304956 | A/duck/Vietnam/QuangBinh/DH130723/2017 | A / H5N6 | Asia / Vietnam | |
| EPI_ISL_304957 | A/duck/Vietnam/QuangBinh/DH330718/2017 | A / H5N6 | Asia / Vietnam | |
| EPI_ISL_304958 | A/chicken/Vietnam/QuangBinh/BD1113/2017 | A / H5N6 | Asia / Vietnam | |
| EPI_ISL_304959 | A/chicken/Vietnam/QuangBinh/BoTrach1113/2017 | A / H5N6 | Asia / Vietnam | |
| EPI_ISL_304960 | A/duck/Vietnam/QuangBinh/QN530206/2018 | A / H5N6 | Asia / Vietnam | |
| EPI_ISL_305416 | A/Domestic_Duck/Netherlands/EMC-2/2018 | A / H5N6 | Europe / Netherlands / Provincie Overijssel / Gemeente Kampen / Commercial poultry farm | Maria Johanna Poen (Erasmus Medical Center / Department of Virology) |
| EPI_ISL_305417 | A/Domestic_Duck/Netherlands/EMC-6/2018 | A / H5N6 | Europe / Netherlands / Provincie Overijssel / Gemeente Kampen / Commercial poultry farm | Maria Johanna Poen (Erasmus Medical Center / Department of Virology) |
| EPI_ISL_305453 | A/chicken/Germany-SH/AR163-L02542/2018 | A / H5N6 | Europe / Germany / Schleswig-Holstein / Nordfriesland | Anne Pohlmann (Friedrich-Loeffler-Institut) |
| EPI_ISL_305454 | A/chicken/Germany-SH/AR164-L02543/2018 | A / H5N6 | Europe / Germany / Schleswig-Holstein / Nordfriesland | Anne Pohlmann (Friedrich-Loeffler-Institut) |
| EPI_ISL_305455 | A/duck/Germany-SH/AR165-L02544/2018 | A / H5N6 | Europe / Germany / Schleswig-Holstein / Nordfriesland | Anne Pohlmann (Friedrich-Loeffler-Institut) |
| EPI_ISL_306910 | A/Turkey/Riyadh/AI1/2017 | A / H5N8 | Asia / Saudi Arabia / Riyadh | Gavin Smith (The University of Hong Kong / Department of Microbiology) |
| EPI_ISL_306911 | A/Duck/Riyadh/AI2/2017 | A / H5N8 | Asia / Saudi Arabia / Riyadh | Gavin Smith (The University of Hong Kong / Department of Microbiology) |
| EPI_ISL_306912 | A/Chicken/Riyadh/AI6/2017 | A / H5N8 | Asia / Saudi Arabia / Riyadh | Gavin Smith (The University of Hong Kong / Department of Microbiology) |
| EPI_ISL_306913 | A/Chicken/Al-Ahsaa/AI7/2017 | A / H5N8 | Asia / Saudi Arabia / Al-Ahsaa | Gavin Smith (The University of Hong Kong / Department of Microbiology) |
| EPI_ISL_306914 | A/Chicken/Riyadh/AI10/2017 | A / H5N8 | Asia / Saudi Arabia / Riyadh | Gavin Smith (The University of Hong Kong / Department of Microbiology) |
| EPI_ISL_306915 | A/Chicken/Riyadh/A15/2018 | A / H5N8 | Asia / Saudi Arabia / Riyadh | Gavin Smith (The University of Hong Kong / Department of Microbiology) |
| EPI_ISL_306955 | A/Bulbul/Riyadh/AI4/2017 | A / H5N8 | Asia / Saudi Arabia / Riyadh | Gavin Smith (The University of Hong Kong / Department of Microbiology) |
| EPI_ISL_306956 | A/Falcon/Riyadh/AI5/2017 | A / H5N8 | Asia / Saudi Arabia / Riyadh | Gavin Smith (The University of Hong Kong / Department of Microbiology) |
| EPI_ISL_306957 | A/Chicken/Al-Ahsaa/AI8/2017 | A / H5N8 | Asia / Saudi Arabia / Al-Ahsaa | Gavin Smith (The University of Hong Kong / Department of Microbiology) |
| EPI_ISL_306958 | A/Ornamental bird/Al-Qasim/AI9/2017 | A / H5N8 | Asia / Saudi Arabia / Al-Qasim | Gavin Smith (The University of Hong Kong / Department of Microbiology) |
| EPI_ISL_306959 | A/Holland pigeon/Riyadh/AI3/2017 | A / H5N8 | Asia / Saudi Arabia / Riyadh | Gavin Smith (The University of Hong Kong / Department of Microbiology) |
| EPI_ISL_306989 | A/turkey/Germany-SH/AR185-L02549/2018 | A / H5N6 | Europe / Germany / Schleswig-Holstein / Nordfriesland | Anne Pohlmann (Friedrich-Loeffler-Institut) |
| EPI_ISL_308781 | A/jungle crow/Hyogo/2803A002/2018 | A / H5N6 | Asia / Japan / Hyogo | |
| EPI_ISL_308782 | A/jungle crow/Hyogo/2803E011/2018 | A / H5N6 | Asia / Japan / Hyogo | |
| EPI_ISL_308783 | A/jungle crow/Hyogo/2803E022/2018 | A / H5N6 | Asia / Japan / Hyogo | |
| EPI_ISL_308802 | A/duck/Democratic Republic of the Congo/17RS882-29/2017 | A / H5N8 | Africa / Congo, the Democatic Republic of | |
| EPI_ISL_308803 | A/duck/Democratic Republic of the Congo/17RS882-33/2017 | A / H5N8 | Africa / Congo, the Democatic Republic of | |
| EPI_ISL_308804 | A/duck/Democratic Republic of the Congo/17RS882-40/2017 | A / H5N8 | Africa / Congo, the Democatic Republic of | |
| EPI_ISL_308805 | A/duck/Democratic Republic of the Congo/17RS882-5/2017 | A / H5N8 | Africa / Congo, the Democatic Republic of | |
| EPI_ISL_308806 | A/chicken/Cameroon/17RS1661-1/2017 | A / H5N8 | Africa / Cameroon | |
| EPI_ISL_308807 | A/duck/Cameroon/17RS1661-3/2017 | A / H5N8 | Africa / Cameroon | |
| EPI_ISL_308808 | A/Indian peafowl/Cameroon/17RS1661-6/2017 | A / H5N8 | Africa / Cameroon | |
| EPI_ISL_308809 | A/pigeon/Cameroon/17RS1661-4/2017 | A / H5N8 | Africa / Cameroon | |
| EPI_ISL_3090099 | A/Sichuan/06681/2021 | A / H5N6 | Asia / China / Sichuan Province | Lei Yang (WHO Chinese National Influenza Center / Virology Institute, Chinese CDC) |
| EPI_ISL_309194 | A/Anas platyrhynchos/Belgium/1899/2017 | A / H5N8 | Europe / Belgium | |
| EPI_ISL_309195 | A/Cygnus olor/Belgium/2967/2017 | A / H5N8 | Europe / Belgium | |
| EPI_ISL_309196 | A/Buteo buteo/Belgium/3022/2017 | A / H5N8 | Europe / Belgium | |
| EPI_ISL_309198 | A/Guinea fowl/Belgium/6102/2017 | A / H5N8 | Europe / Belgium | |
| EPI_ISL_309199 | A/quail/Vietnam/CVVI-01/2015 | A / H5N6 | Asia / Vietnam | |
| EPI_ISL_309200 | A/quail/Vietnam/CVVI-03/2015 | A / H5N6 | Asia / Vietnam | |
| EPI_ISL_3098710 | A/common eider/Sweden/SVA210617SZ0354/FB002253/I-202 | A / H5N8 | Europe / Sweden / Gotlands Lan / Gotlands Kommun | Siamak Zohari (National Veterinary Institute) |
| EPI_ISL_3098719 | A/peregrine falcon/Sweden/SVA210309SZ0403/FB000851/K-2 | A / H5N4 | Europe / Sweden / Blekinge Lan / Solvesborgs Kommun | Siamak Zohari (National Veterinary Institute) |
| EPI_ISL_3102024 | A/barnacle goose/Sweden/SVA210423SZ0250/FB001648/M-2 | A / H5N8 | Europe / Sweden / Skane Lan / Trelleborgs Kommun | Siamak Zohari (National Veterinary Institute) |
| EPI_ISL_3102029 | A/barnacle goose/Sweden/SVA210423SZ0252/FB001647/M20 | A / H5N8 | Europe / Sweden / Skane Lan / Vellinge Kommun | Siamak Zohari (National Veterinary Institute) |
| EPI_ISL_3102054 | A/European_herring_gull/Bulgaria/222_21VIR4270-1/2021 | A / H5N5 | Europe / Bulgaria / Kiten, Burgas | Bianca Zecchin (Istituto Zooprofilattico Sperimentale Delle Venezie) |
| EPI_ISL_3102055 | A/grey_heron/Bulgaria/223_21VIR4270-2/2021 | A / H5N5 | Europe / Bulgaria / Kiten, Burgas | Bianca Zecchin (Istituto Zooprofilattico Sperimentale Delle Venezie) |
| EPI_ISL_3102056 | A/chicken/Bulgaria/274-5_21VIR4270-4/2021 | A / H5N8 | Europe / Bulgaria / Krivo pole, Haskovo | Bianca Zecchin (Istituto Zooprofilattico Sperimentale Delle Venezie) |
| EPI_ISL_3102057 | A/chicken/Bulgaria/275-4_21VIR4270-6/2021 | A / H5N8 | Europe / Bulgaria / Trilistnik, Plovdiv | Bianca Zecchin (Istituto Zooprofilattico Sperimentale Delle Venezie) |
| EPI_ISL_3102058 | A/chicken/Bulgaria/297-2_21VIR4270-8/2021 | A / H5N8 | Europe / Bulgaria / Trilistnik, Plovdiv | Bianca Zecchin (Istituto Zooprofilattico Sperimentale Delle Venezie) |
| EPI_ISL_3102059 | A/chicken/Bulgaria/298-1_21VIR4270-9/2021 | A / H5N8 | Europe / Bulgaria / Trilistnik, Plovdiv | Bianca Zecchin (Istituto Zooprofilattico Sperimentale Delle Venezie) |
| EPI_ISL_3102060 | A/duck/Poland/H160_21RS1385-1/2021 | A / H5N8 | Europe / Poland / Wielkopolskie | Bianca Zecchin (Istituto Zooprofilattico Sperimentale Delle Venezie) |
| EPI_ISL_3102061 | A/duck/Poland/H542_21RS1385-2/2021 | A / H5N8 | Europe / Poland / Wielkopolskie | Bianca Zecchin (Istituto Zooprofilattico Sperimentale Delle Venezie) |
| EPI_ISL_3102062 | A/chicken/Poland/H1124_21RS1385-3/2021 | A / H5N8 | Europe / Poland / Wielkopolskie | Bianca Zecchin (Istituto Zooprofilattico Sperimentale Delle Venezie) |
| EPI_ISL_3102063 | A/goose/Poland/H1044_21RS1385-4/2021 | A / H5N8 | Europe / Poland / Wielkopolskie | Bianca Zecchin (Istituto Zooprofilattico Sperimentale Delle Venezie) |
| EPI_ISL_3102064 | A/duck/Poland/H1029_21RS1385-5/2021 | A / H5N8 | Europe / Poland / Wielkopolskie | Bianca Zecchin (Istituto Zooprofilattico Sperimentale Delle Venezie) |
| EPI_ISL_3102065 | A/goose/Poland/H1036_21RS1385-6/2021 | A / H5N8 | Europe / Poland / Wielkopolskie | Bianca Zecchin (Istituto Zooprofilattico Sperimentale Delle Venezie) |
| EPI_ISL_3102066 | A/chicken/Poland/H812_21RS1385-7/2021 | A / H5N8 | Europe / Poland / Wielkopolskie | Bianca Zecchin (Istituto Zooprofilattico Sperimentale Delle Venezie) |
| EPI_ISL_3102067 | A/chicken/Poland/H293_21RS1385-8/2021 | A / H5N8 | Europe / Poland / Mazowieckie | Bianca Zecchin (Istituto Zooprofilattico Sperimentale Delle Venezie) |
| EPI_ISL_3102068 | A/chicken/Poland/H984_21RS1385-9/2021 | A / H5N8 | Europe / Poland / Mazowieckie | Bianca Zecchin (Istituto Zooprofilattico Sperimentale Delle Venezie) |
| EPI_ISL_3102069 | A/chicken/Poland/H1161_21RS1385-10/2021 | A / H5N8 | Europe / Poland / Mazowieckie | Bianca Zecchin (Istituto Zooprofilattico Sperimentale Delle Venezie) |
| EPI_ISL_3102070 | A/white_stork/Poland/MB412_21RS1385-11/2021 | A / H5N8 | Europe / Poland / Mazowieckie | Bianca Zecchin (Istituto Zooprofilattico Sperimentale Delle Venezie) |
| EPI_ISL_3102071 | A/chicken/Poland/H712_21RS1385-12/2021 | A / H5N8 | Europe / Poland / Mazowieckie | Bianca Zecchin (Istituto Zooprofilattico Sperimentale Delle Venezie) |
| EPI_ISL_3102072 | A/chicken/Poland/H1084_21RS1385-13/2021 | A / H5N8 | Europe / Poland / Mazowieckie | Bianca Zecchin (Istituto Zooprofilattico Sperimentale Delle Venezie) |
| EPI_ISL_3102073 | A/turkey/Poland/H1184_21RS1385-14/2021 | A / H5N8 | Europe / Poland / Mazowieckie | Bianca Zecchin (Istituto Zooprofilattico Sperimentale Delle Venezie) |

| | | | | |
|---|---|---|---|---|
| EPI_ISL_3102074 | A/turkey/Poland/H1168_21RS1385-15/2021 | A / H5N8 | Europe / Poland / Warmi?sko-mazurskie | Bianca Zecchin (Istituto Zooprofilattico Sperimentale Delle Venezie) |
| EPI_ISL_3102075 | A/chicken/Poland/H928_21RS1385-16/2021 | A / H5N8 | Europe / Poland / Warmi?sko-mazurskie | Bianca Zecchin (Istituto Zooprofilattico Sperimentale Delle Venezie) |
| EPI_ISL_3102076 | A/duck/Poland/H515_21RS1385-17/2021 | A / H5N8 | Europe / Poland / Lodzkie | Bianca Zecchin (Istituto Zooprofilattico Sperimentale Delle Venezie) |
| EPI_ISL_3102077 | A/duck/Poland/H514_21RS1385-18/2021 | A / H5N8 | Europe / Poland / Pomorskie | Bianca Zecchin (Istituto Zooprofilattico Sperimentale Delle Venezie) |
| EPI_ISL_3102078 | A/mute_swan/Poland/MB396_21RS1385-19/2021 | A / H5N8 | Europe / Poland / Zachodniopomorskie | Bianca Zecchin (Istituto Zooprofilattico Sperimentale Delle Venezie) |
| EPI_ISL_3102079 | A/turkey/Poland/H1289_21RS1385-20/2021 | A / H5N8 | Europe / Poland / Mazowieckie | Bianca Zecchin (Istituto Zooprofilattico Sperimentale Delle Venezie) |
| EPI_ISL_3102080 | A/chicken/Austria/21052483_21VIR3291-1/2021 | A / H5N8 | Europe / Austria / Oberneukirchen | Bianca Zecchin (Istituto Zooprofilattico Sperimentale Delle Venezie) |
| EPI_ISL_3102081 | A/mute_swan/Romania/11981-1_21VIR3163-5/2021 | A / H5N5 | Europe / Romania / Tulcea Chilia Veche | Bianca Zecchin (Istituto Zooprofilattico Sperimentale Delle Venezie) |
| EPI_ISL_3102082 | A/mute_swan/Romania/11981-2_21VIR3163-6/2021 | A / H5N8 | Europe / Romania / Tulcea Chilia Veche | Bianca Zecchin (Istituto Zooprofilattico Sperimentale Delle Venezie) |
| EPI_ISL_3102208 | A/barnacle goose/Sweden/SVA210210SZ0372/FB000374/M-2( | A / H5N8 | Europe / Sweden / Skane Lan / Svedala Kommun | Siamak Zohari (National Veterinary Institute) |
| EPI_ISL_312376 | A/Armenian Gull/Republic of Georgia/4/2017 | A / H5N6 | Asia / Georgia / Mtkvari River, Tbilisi | Maria Johanna Poen (Erasmus Medical Center / Department of Virology) |
| EPI_ISL_3127850 | A/whooper swan/Sweden/SVA210331SZ0441/FB0001325/W-2 | A / H5N8 | Europe / Sweden / Dalarnas Lan / Leksands Kommun | Siamak Zohari (National Veterinary Institute) |
| EPI_ISL_3128152 | A/Canada goose/Sweden/SVA210331SZ/FB001326/W-2021 | A / H5N8 | Europe / Sweden / Dalarnas Lan / Leksands Kommun | Siamak Zohari (National Veterinary Institute) |
| EPI_ISL_3128522 | A/Canada goose/Sweden/SVA210407SZ0510/FB001379/O-202 | A / H5N8 | Europe / Sweden / Vastra Gotalands Lan / Hjo Kommun | Siamak Zohari (National Veterinary Institute) |
| EPI_ISL_3128523 | A/dalmatian pelican/Greece/47t_21VIR3735-3/2021 | A / H5N8 | Europe / Greece / Serres | Bianca Zecchin (Istituto Zooprofilattico Sperimentale Delle Venezie) |
| EPI_ISL_3128524 | A/partridge/Kosovo/96_21VIR5162-14/2021 | A / H5N8 | Europe / Kosovo / Suhareke | Bianca Zecchin (Istituto Zooprofilattico Sperimentale Delle Venezie) |
| EPI_ISL_3128525 | A/chicken/Kosovo/97_21VIR5162-15/2021 | A / H5N8 | Europe / Kosovo / Gjilan | Bianca Zecchin (Istituto Zooprofilattico Sperimentale Delle Venezie) |
| EPI_ISL_3128526 | A/chicken/Kosovo/98_21VIR5162-16/2021 | A / H5N8 | Europe / Kosovo / Mitrovice | Bianca Zecchin (Istituto Zooprofilattico Sperimentale Delle Venezie) |
| EPI_ISL_3128527 | A/duck/Kosovo/107_21VIR5162-17/2021 | A / H5N8 | Europe / Kosovo / Kacanik | Bianca Zecchin (Istituto Zooprofilattico Sperimentale Delle Venezie) |
| EPI_ISL_3128528 | A/chicken/Kosovo/113_21VIR5162-19/2021 | A / H5N8 | Europe / Kosovo / Shtime | Bianca Zecchin (Istituto Zooprofilattico Sperimentale Delle Venezie) |
| EPI_ISL_3128529 | A/chicken/Kosovo/82_21VIR5162-1/2021 | A / H5N8 | Europe / Kosovo / Mitrovica | Bianca Zecchin (Istituto Zooprofilattico Sperimentale Delle Venezie) |
| EPI_ISL_3128530 | A/chicken/Kosovo/84_21VIR5162-2/2021 | A / H5N8 | Europe / Kosovo / Ferizaj | Bianca Zecchin (Istituto Zooprofilattico Sperimentale Delle Venezie) |
| EPI_ISL_3128531 | A/chicken/Kosovo/90_21VIR5162-4/2021 | A / H5N8 | Europe / Kosovo / Rahovec | Bianca Zecchin (Istituto Zooprofilattico Sperimentale Delle Venezie) |
| EPI_ISL_3128532 | A/chicken/Kosovo/104_21VIR5162-7/2021 | A / H5N8 | Europe / Kosovo / Malisheve | Bianca Zecchin (Istituto Zooprofilattico Sperimentale Delle Venezie) |
| EPI_ISL_3128533 | A/chicken/Albania/2574_21VIR5387-1/2021 | A / H5N8 | Europe / Albania | Bianca Zecchin (Istituto Zooprofilattico Sperimentale Delle Venezie) |
| EPI_ISL_3128534 | A/chicken/Albania/3816_21VIR5387-2/2021 | A / H5N8 | Europe / Albania | Bianca Zecchin (Istituto Zooprofilattico Sperimentale Delle Venezie) |
| EPI_ISL_313226 | A/white stork/Germany-NI/AR251/2018 | A / H5N6 | Europe / Germany / Lower Saxony / Vechta | Jacqueline King (Friedrich-Loeffler-Institut) |
| EPI_ISL_313227 | A/buzzard/Germany-NRW/AR279/2018 | A / H5N6 | Europe / Germany / North Rhine-Westphalia / Herford | Jacqueline King (Friedrich-Loeffler-Institut) |
| EPI_ISL_3135897 | A/Turkey/Hungary/16603/2021 (H5N1) | A / H5N1 | Europe / Hungary / Hajdu-Bihar | Katalin Szentpáli-Gavallér (National Food Chain Safety Office, Hungary / Virology) |
| EPI_ISL_3135926 | A/eagle/Hungary/8569/2021 (H5N5) | A / H5N5 | Europe / Hungary / Somogy megye | Katalin Szentpáli-Gavallér (National Food Chain Safety Office, Hungary / Virology) |
| EPI_ISL_3135931 | A/swan/Hungary/9638/2021 (H5N8) | A / H5N8 | Europe / Hungary / Csongrad megye | Katalin Szentpáli-Gavallér (National Food Chain Safety Office, Hungary / Virology) |
| EPI_ISL_313805 | A/Northern Goshawk/Tokyo/1301B003T/2018 | A / H5N6 | Asia / Japan | Takehiko Saito (National Institute of Animal Health) |
| EPI_ISL_313806 | A/chicken/Kagawa/1T-1/2018 | A / H5N6 | Asia / Japan | Takehiko Saito (National Institute of Animal Health) |
| EPI_ISL_3138567 | A/common buzzard/Sweden/SVA210415SZ0338/FB001522/O- | A / H5N8 | Europe / Sweden / Vastra Gotalands Lan / Boras Kommun | Siamak Zohari (National Veterinary Institute) |
| EPI_ISL_3138618 | A/whooper swan/Sweden/SVA210420SZ0485/FB0001581/W-2 | A / H5N8 | Europe / Sweden / Dalarnas Lan / Borlange Kommun | Siamak Zohari (National Veterinary Institute) |
| EPI_ISL_3142375 | A/mute_swan/Austria/21051907_21VIR3291-6/2021 | A / H5N8 | Europe / Austria / Oberneukirchen | Bianca Zecchin (Istituto Zooprofilattico Sperimentale Delle Venezie) |
| EPI_ISL_3142391 | A/mute_swan/Austria/21051907_21VIR3291-7/2021 | A / H5N8 | Europe / Austria / Oberneukirchen | Bianca Zecchin (Istituto Zooprofilattico Sperimentale Delle Venezie) |
| EPI_ISL_3142407 | A/turkey/Kosovo/90_21VIR5162-5/2021 | A / H5N8 | Europe / Kosovo / Rahovec | Bianca Zecchin (Istituto Zooprofilattico Sperimentale Delle Venezie) |
| EPI_ISL_3143953 | A/common buzzard/Sweden/SVA210505SZ0374/FB0001789/M | A / H5N8 | Europe / Sweden / Skane Lan / Bastads Kommun | Siamak Zohari (National Veterinary Institute) |
| EPI_ISL_3143954 | A/whooper swan/Sweden/SVA2100329SZ0210/FB0001277/H- | A / H5N5 | Europe / Sweden / Kalmar Lan / Hongsby Kommun | Siamak Zohari (National Veterinary Institute) |
| EPI_ISL_314821 | A/chicken/Kagawa/1C-1/2018 | A / H5N6 | Asia / Japan / Kagawa | Takehiko Saito (National Institute of Animal Health) |
| EPI_ISL_314856 | A/chicken/Guangdong/G1012/2018 | A / H5N6 | Asia / China / Guangdong Province / live bird market | Jiming Chen (China Animal Health and Epidemiology Center (CAHEC)) |
| EPI_ISL_314857 | A/duck/Guangdong/G1378/2018 | A / H5N6 | Asia / China / Guangdong Province / live bird market | Jiming Chen (China Animal Health and Epidemiology Center (CAHEC)) |
| EPI_ISL_315236 | A/Enviroment/Xuzhou/4018/2018 | A / H5N6 | Asia / China / Jiangsu Provice | Xian Qi (Jiangsu Provincial Center for Disease Control & Prevention) |
| EPI_ISL_315237 | A/Chicken/Xuzhou/470/2017 | A / H5N6 | Asia / China / Jiangsu Provice | Xian Qi (Jiangsu Provincial Center for Disease Control & Prevention) |
| EPI_ISL_315238 | A/Enviroment/Xuzhou/461/2017 | A / H5N6 | Asia / China / Jiangsu Provice | Xian Qi (Jiangsu Provincial Center for Disease Control & Prevention) |
| EPI_ISL_316513 | A/chicken/Italy/17VIR2294-5/2017 | A / H5N8 | Europe / Italy / Treviso | Bianca Zecchin (Istituto Zooprofilattico Sperimentale Delle Venezie) |
| EPI_ISL_316514 | A/chicken/Italy/17VIR2658-3/2017 | A / H5N8 | Europe / Italy / Pordenone | Bianca Zecchin (Istituto Zooprofilattico Sperimentale Delle Venezie) |
| EPI_ISL_316515 | A/turkey/Italy/17VIR2722-29/2017 | A / H5N8 | Europe / Italy / Verona | Bianca Zecchin (Istituto Zooprofilattico Sperimentale Delle Venezie) |
| EPI_ISL_316516 | A/chicken/Italy/17VIR2788/2017 | A / H5N8 | Europe / Italy / Torino | Bianca Zecchin (Istituto Zooprofilattico Sperimentale Delle Venezie) |
| EPI_ISL_316517 | A/turkey/Italy/17VIR3192-1/2017 | A / H5N8 | Europe / Italy / Verona | Bianca Zecchin (Istituto Zooprofilattico Sperimentale Delle Venezie) |
| EPI_ISL_316518 | A/turkey/Italy/17VIR4407-1/2017 | A / H5N8 | Europe / Italy / Mantova | Bianca Zecchin (Istituto Zooprofilattico Sperimentale Delle Venezie) |
| EPI_ISL_316519 | A/turkey/Italy/17VIR5773-2/2017 | A / H5N8 | Europe / Italy / Mantova | Bianca Zecchin (Istituto Zooprofilattico Sperimentale Delle Venezie) |
| EPI_ISL_316520 | A/duck/Italy/17VIR5769-5/2017 | A / H5N8 | Europe / Italy / Mantova | Bianca Zecchin (Istituto Zooprofilattico Sperimentale Delle Venezie) |
| EPI_ISL_316521 | A/chicken/Italy/17VIR5794-5/2017 | A / H5N8 | Europe / Italy / Mantova | Bianca Zecchin (Istituto Zooprofilattico Sperimentale Delle Venezie) |
| EPI_ISL_316522 | A/turkey/Italy/17VIR5964-8/2017 | A / H5N8 | Europe / Italy / Verona | Bianca Zecchin (Istituto Zooprofilattico Sperimentale Delle Venezie) |
| EPI_ISL_316523 | A/chicken/Italy/17VIR5986-1/2017 | A / H5N8 | Europe / Italy / Mantova | Bianca Zecchin (Istituto Zooprofilattico Sperimentale Delle Venezie) |
| EPI_ISL_316524 | A/turkey/Italy/17VIR6108-3/2017 | A / H5N8 | Europe / Italy / Verona | Bianca Zecchin (Istituto Zooprofilattico Sperimentale Delle Venezie) |
| EPI_ISL_316525 | A/turkey/Italy/17VIR6111-16/2017 | A / H5N8 | Europe / Italy / Parma | Bianca Zecchin (Istituto Zooprofilattico Sperimentale Delle Venezie) |
| EPI_ISL_316526 | A/turkey/Italy/17VIR6355-1/2017 | A / H5N8 | Europe / Italy / Mantova | Bianca Zecchin (Istituto Zooprofilattico Sperimentale Delle Venezie) |
| EPI_ISL_316527 | A/turkey/Italy/17VIR6356-1/2017 | A / H5N8 | Europe / Italy / Mantova | Bianca Zecchin (Istituto Zooprofilattico Sperimentale Delle Venezie) |
| EPI_ISL_316528 | A/turkey/Italy/17VIR6357-2/2017 | A / H5N8 | Europe / Italy / Mantova | Bianca Zecchin (Istituto Zooprofilattico Sperimentale Delle Venezie) |
| EPI_ISL_316529 | A/turkey/Italy/17VIR6469-2/2017 | A / H5N8 | Europe / Italy / Mantova | Bianca Zecchin (Istituto Zooprofilattico Sperimentale Delle Venezie) |
| EPI_ISL_316530 | A/turkey/Italy/17VIR6882-1/2017 | A / H5N8 | Europe / Italy / Verona | Bianca Zecchin (Istituto Zooprofilattico Sperimentale Delle Venezie) |
| EPI_ISL_316531 | A/pheasant/Italy/17VIR7065-21/2017 | A / H5N8 | Europe / Italy / Lodi | Bianca Zecchin (Istituto Zooprofilattico Sperimentale Delle Venezie) |
| EPI_ISL_316532 | A/turkey/Italy/17VIR6930-4/2017 | A / H5N8 | Europe / Italy / Verona | Bianca Zecchin (Istituto Zooprofilattico Sperimentale Delle Venezie) |
| EPI_ISL_316533 | A/turkey/Italy/17VIR6928-5/2017 | A / H5N8 | Europe / Italy / Verona | Bianca Zecchin (Istituto Zooprofilattico Sperimentale Delle Venezie) |
| EPI_ISL_316534 | A/chicken/Italy/17VIR7062-1/2017 | A / H5N8 | Europe / Italy / Pavia | Bianca Zecchin (Istituto Zooprofilattico Sperimentale Delle Venezie) |
| EPI_ISL_316535 | A/turkey/Italy/17VIR7197-1/2017 | A / H5N8 | Europe / Italy / Verona | Bianca Zecchin (Istituto Zooprofilattico Sperimentale Delle Venezie) |
| EPI_ISL_316536 | A/turkey/Italy/17VIR7337-1/2017 | A / H5N8 | Europe / Italy / Cremona | Bianca Zecchin (Istituto Zooprofilattico Sperimentale Delle Venezie) |
| EPI_ISL_316537 | A/goose/Italy/17VIR7626-5/2017 | A / H5N8 | Europe / Italy / Padova | Bianca Zecchin (Istituto Zooprofilattico Sperimentale Delle Venezie) |
| EPI_ISL_316538 | A/turkey/Italy/17VIR8199-1/2017 | A / H5N8 | Europe / Italy / Vicenza | Bianca Zecchin (Istituto Zooprofilattico Sperimentale Delle Venezie) |

| | | | | |
|---|---|---|---|---|
| EPI_ISL_316539 | A/turkey/Italy/17VIR8275-3/2017 | A / H5N8 | Europe / Italy / Verona | Bianca Zecchin (Istituto Zooprofilattico Sperimentale Delle Venezie) |
| EPI_ISL_316540 | A/duck/Italy/17VIR8200-12/2017 | A / H5N8 | Europe / Italy / Vicenza | Bianca Zecchin (Istituto Zooprofilattico Sperimentale Delle Venezie) |
| EPI_ISL_316541 | A/duck/Italy/17VIR8349/2017 | A / H5N8 | Europe / Italy / Cremona | Bianca Zecchin (Istituto Zooprofilattico Sperimentale Delle Venezie) |
| EPI_ISL_316542 | A/chicken/Italy/17VIR8678/2017 | A / H5N8 | Europe / Italy / Ferrara | Bianca Zecchin (Istituto Zooprofilattico Sperimentale Delle Venezie) |
| EPI_ISL_316543 | A/chicken/Italy/17VIR8783-5/2017 | A / H5N8 | Europe / Italy / Vicenza | Bianca Zecchin (Istituto Zooprofilattico Sperimentale Delle Venezie) |
| EPI_ISL_316544 | A/chicken/Italy/17VIR8821/2017 | A / H5N8 | Europe / Italy / Bergamo | Bianca Zecchin (Istituto Zooprofilattico Sperimentale Delle Venezie) |
| EPI_ISL_316545 | A/chicken/Italy/17VIR8744-9/2017 | A / H5N8 | Europe / Italy / Vicenza | Bianca Zecchin (Istituto Zooprofilattico Sperimentale Delle Venezie) |
| EPI_ISL_316546 | A/chicken/Italy/17VIR8885/2017 | A / H5N8 | Europe / Italy / Bergamo | Bianca Zecchin (Istituto Zooprofilattico Sperimentale Delle Venezie) |
| EPI_ISL_316547 | A/turkey/Italy/17VIR8886-3/2017 | A / H5N8 | Europe / Italy / Padova | Bianca Zecchin (Istituto Zooprofilattico Sperimentale Delle Venezie) |
| EPI_ISL_316548 | A/turkey/Italy/17VIR8819-3/2017 | A / H5N8 | Europe / Italy / Brescia | Bianca Zecchin (Istituto Zooprofilattico Sperimentale Delle Venezie) |
| EPI_ISL_316549 | A/chicken/Italy/17VIR9048-2/2017 | A / H5N8 | Europe / Italy / Bergamo | Bianca Zecchin (Istituto Zooprofilattico Sperimentale Delle Venezie) |
| EPI_ISL_316550 | A/turkey/Italy/17VIR9106-2/2017 | A / H5N8 | Europe / Italy / Brescia | Bianca Zecchin (Istituto Zooprofilattico Sperimentale Delle Venezie) |
| EPI_ISL_316551 | A/chicken/Italy/17VIR9107/2017 | A / H5N8 | Europe / Italy / Sondrio | Bianca Zecchin (Istituto Zooprofilattico Sperimentale Delle Venezie) |
| EPI_ISL_316552 | A/turkey/Italy/17VIR9049-2/2017 | A / H5N8 | Europe / Italy / Mantova | Bianca Zecchin (Istituto Zooprofilattico Sperimentale Delle Venezie) |
| EPI_ISL_316553 | A/chicken/Italy/17VIR9113/2017 | A / H5N8 | Europe / Italy / Padova | Bianca Zecchin (Istituto Zooprofilattico Sperimentale Delle Venezie) |
| EPI_ISL_316554 | A/duck/Italy/17VIR8898/2017 | A / H5N8 | Europe / Italy / Padova | Bianca Zecchin (Istituto Zooprofilattico Sperimentale Delle Venezie) |
| EPI_ISL_316555 | A/turkey/Italy/17VIR9245/2017 | A / H5N8 | Europe / Italy / Brescia | Bianca Zecchin (Istituto Zooprofilattico Sperimentale Delle Venezie) |
| EPI_ISL_316556 | A/turkey/Italy/17VIR9246/2017 | A / H5N8 | Europe / Italy / Brescia | Bianca Zecchin (Istituto Zooprofilattico Sperimentale Delle Venezie) |
| EPI_ISL_316557 | A/chicken/Italy/17VIR9456/2017 | A / H5N8 | Europe / Italy / Milano | Bianca Zecchin (Istituto Zooprofilattico Sperimentale Delle Venezie) |
| EPI_ISL_316558 | A/duck/Italy/17VIR9533/2017 | A / H5N8 | Europe / Italy / Brescia | Bianca Zecchin (Istituto Zooprofilattico Sperimentale Delle Venezie) |
| EPI_ISL_316559 | A/chicken/Italy/17VIR9679/2017 | A / H5N8 | Europe / Italy / Brescia | Bianca Zecchin (Istituto Zooprofilattico Sperimentale Delle Venezie) |
| EPI_ISL_316560 | A/chicken/Italy/17VIR9680/2017 | A / H5N8 | Europe / Italy / Brescia | Bianca Zecchin (Istituto Zooprofilattico Sperimentale Delle Venezie) |
| EPI_ISL_316561 | A/turkey/Italy/17VIR9699/2017 | A / H5N8 | Europe / Italy / Brescia | Bianca Zecchin (Istituto Zooprofilattico Sperimentale Delle Venezie) |
| EPI_ISL_316562 | A/turkey/Italy/17VIR9773/2017 | A / H5N8 | Europe / Italy / Brescia | Bianca Zecchin (Istituto Zooprofilattico Sperimentale Delle Venezie) |
| EPI_ISL_316563 | A/turkey/Italy/17VIR9775-1/2017 | A / H5N8 | Europe / Italy / Bergamo | Bianca Zecchin (Istituto Zooprofilattico Sperimentale Delle Venezie) |
| EPI_ISL_316564 | A/turkey/Italy/17VIR9908/2017 | A / H5N8 | Europe / Italy / Brescia | Bianca Zecchin (Istituto Zooprofilattico Sperimentale Delle Venezie) |
| EPI_ISL_316565 | A/turkey/Italy/17VIR9905-1/2017 | A / H5N8 | Europe / Italy / Brescia | Bianca Zecchin (Istituto Zooprofilattico Sperimentale Delle Venezie) |
| EPI_ISL_316566 | A/duck/Italy/17VIR9906-1/2017 | A / H5N8 | Europe / Italy / Brescia | Bianca Zecchin (Istituto Zooprofilattico Sperimentale Delle Venezie) |
| EPI_ISL_316567 | A/duck/Italy/17VIR9907-2/2017 | A / H5N8 | Europe / Italy / Brescia | Bianca Zecchin (Istituto Zooprofilattico Sperimentale Delle Venezie) |
| EPI_ISL_316568 | A/turkey/Italy/17VIR9942-2/2017 | A / H5N8 | Europe / Italy / Brescia | Bianca Zecchin (Istituto Zooprofilattico Sperimentale Delle Venezie) |
| EPI_ISL_316569 | A/chicken/Italy/17VIR9941-1/2017 | A / H5N8 | Europe / Italy / Brescia | Bianca Zecchin (Istituto Zooprofilattico Sperimentale Delle Venezie) |
| EPI_ISL_316570 | A/chicken/Italy/17VIR9846-4/2017 | A / H5N8 | Europe / Italy / Asti | Bianca Zecchin (Istituto Zooprofilattico Sperimentale Delle Venezie) |
| EPI_ISL_316571 | A/chicken/Italy/17VIR10053/2017 | A / H5N8 | Europe / Italy / Brescia | Bianca Zecchin (Istituto Zooprofilattico Sperimentale Delle Venezie) |
| EPI_ISL_316572 | A/chicken/Italy/17VIR9868-1/2017 | A / H5N8 | Europe / Italy / Roma | Bianca Zecchin (Istituto Zooprofilattico Sperimentale Delle Venezie) |
| EPI_ISL_316573 | A/turkey/Italy/17VIR10133-8/2017 | A / H5N8 | Europe / Italy / Cremona | Bianca Zecchin (Istituto Zooprofilattico Sperimentale Delle Venezie) |
| EPI_ISL_316574 | A/chicken/Italy/17VIR10083/2017 | A / H5N8 | Europe / Italy / Brescia | Bianca Zecchin (Istituto Zooprofilattico Sperimentale Delle Venezie) |
| EPI_ISL_316575 | A/chicken/Italy/17VIR10082/2017 | A / H5N8 | Europe / Italy / Brescia | Bianca Zecchin (Istituto Zooprofilattico Sperimentale Delle Venezie) |
| EPI_ISL_316576 | A/chicken/Italy/17VIR10130-2/2017 | A / H5N8 | Europe / Italy / Brescia | Bianca Zecchin (Istituto Zooprofilattico Sperimentale Delle Venezie) |
| EPI_ISL_316577 | A/chicken/Italy/17VIR10132-1/2017 | A / H5N8 | Europe / Italy / Brescia | Bianca Zecchin (Istituto Zooprofilattico Sperimentale Delle Venezie) |
| EPI_ISL_316578 | A/chicken/Italy/17VIR10131-1/2017 | A / H5N8 | Europe / Italy / Brescia | Bianca Zecchin (Istituto Zooprofilattico Sperimentale Delle Venezie) |
| EPI_ISL_316579 | A/chicken/Italy/17VIR10437-2/2017 | A / H5N8 | Europe / Italy / Brescia | Bianca Zecchin (Istituto Zooprofilattico Sperimentale Delle Venezie) |
| EPI_ISL_316580 | A/chicken/Italy/17VIR10408/2017 | A / H5N8 | Europe / Italy / Treviso | Bianca Zecchin (Istituto Zooprofilattico Sperimentale Delle Venezie) |
| EPI_ISL_316581 | A/duck/Italy/17VIR10767-13/2017 | A / H5N8 | Europe / Italy / Treviso | Bianca Zecchin (Istituto Zooprofilattico Sperimentale Delle Venezie) |
| EPI_ISL_316582 | A/turkey/Italy/17VIR10983-22/2017 | A / H5N8 | Europe / Italy / Ravenna | Bianca Zecchin (Istituto Zooprofilattico Sperimentale Delle Venezie) |
| EPI_ISL_316583 | A/swan/Italy/17VIR3594-3/2017 | A / H5N8 | Europe / Italy / Torino | Bianca Zecchin (Istituto Zooprofilattico Sperimentale Delle Venezie) |
| EPI_ISL_316584 | A/swan/Italy/17VIR8513/2017 | A / H5N8 | Europe / Italy / Como | Bianca Zecchin (Istituto Zooprofilattico Sperimentale Delle Venezie) |
| EPI_ISL_316585 | A/swan/Italy/17VIR9017/2017 | A / H5N8 | Europe / Italy / Padova | Bianca Zecchin (Istituto Zooprofilattico Sperimentale Delle Venezie) |
| EPI_ISL_316586 | A/swan/Italy/17VIR9038-1/2017 | A / H5N8 | Europe / Italy / Verbano-Cusio-Ossola | Bianca Zecchin (Istituto Zooprofilattico Sperimentale Delle Venezie) |
| EPI_ISL_316587 | A/swan/Italy/17VIR9038-2/2017 | A / H5N8 | Europe / Italy / Verbano-Cusio-Ossola | Bianca Zecchin (Istituto Zooprofilattico Sperimentale Delle Venezie) |
| EPI_ISL_316588 | A/swan/Italy/17VIR9836-1/2017 | A / H5N8 | Europe / Italy / Verbano-Cusio-Ossola | Bianca Zecchin (Istituto Zooprofilattico Sperimentale Delle Venezie) |
| EPI_ISL_316589 | A/quail/Italy/17VIR9455-2/2017 | A / H5N8 | Europe / Italy / Bergamo | Bianca Zecchin (Istituto Zooprofilattico Sperimentale Delle Venezie) |
| EPI_ISL_316591 | A/kestrel/Italy/17VIR9840/2017 | A / H5N8 | Europe / Italy / Ferrara | Bianca Zecchin (Istituto Zooprofilattico Sperimentale Delle Venezie) |
| EPI_ISL_316592 | A/mallard/Italy/17VIR6103-1/2017 | A / H5N8 | Europe / Italy / Pavia | Bianca Zecchin (Istituto Zooprofilattico Sperimentale Delle Venezie) |
| EPI_ISL_316593 | A/pigeon/Italy/17VIR9842/2017 | A / H5N8 | Europe / Italy / Ferrara | Bianca Zecchin (Istituto Zooprofilattico Sperimentale Delle Venezie) |
| EPI_ISL_316594 | A/wild_goose/Italy/17VIR9841/2017 | A / H5N8 | Europe / Italy / Ferrara | Bianca Zecchin (Istituto Zooprofilattico Sperimentale Delle Venezie) |
| EPI_ISL_319652 | A/tufted duck/Shimane/3211TY001/2017 | A / H5N6 | Asia / Japan / Shimane | |
| EPI_ISL_320545 | A/White-tailed sea eagle/Germany/AR3089/2017 | A / H5 | Europe / Germany / Mecklenburg-Vorpommern | Timm Harder (Friedrich-Loeffler-Institut) |
| EPI_ISL_320546 | A/White-tailed sea eagle/Germany/AR3093/2017 | A / H5N8 | Europe / Germany / Mecklenburg-Vorpommern | Timm Harder (Friedrich-Loeffler-Institut) |
| EPI_ISL_320574 | A/White-tailed sea eagle/Germany/AR10297/2016 | A / H5N8 | Europe / Germany / Schleswig-Holstein | Timm Harder (Friedrich-Loeffler-Institut) |
| EPI_ISL_320607 | A/chicken/Kursk/284/2018 | A / H5N8 | Europe / Russian Federation / Kursk Oblast | Ivan Susloparov (State Research Center of Virology and Biotechnology (VECTOR) / Emerging Zoonotic Diseases and Influenza) |
| EPI_ISL_320608 | A/chicken/Kursk/526/2018 | A / H5N8 | Europe / Russian Federation / Kursk Oblast | Ivan Susloparov (State Research Center of Virology and Biotechnology (VECTOR) / Emerging Zoonotic Diseases and Influenza) |
| EPI_ISL_320657 | A/chicken/Samara/446/2018 | A / H5N8 | Europe / Russian Federation / Samara Oblast | Ivan Susloparov (State Research Center of Virology and Biotechnology (VECTOR) / Emerging Zoonotic Diseases and Influenza) |
| EPI_ISL_320658 | A/chicken/Samara/447/2018 | A / H5N8 | Europe / Russian Federation / Samara Oblast | Ivan Susloparov (State Research Center of Virology and Biotechnology (VECTOR) / Emerging Zoonotic Diseases and Influenza) |
| EPI_ISL_320659 | A/duck/Samara/452/2018 | A / H5N8 | Europe / Russian Federation / Samara Oblast | Ivan Susloparov (State Research Center of Virology and Biotechnology (VECTOR) / Emerging Zoonotic Diseases and Influenza) |
| EPI_ISL_320660 | A/goose/Samara/455/2018 | A / H5N8 | Europe / Russian Federation / Samara Oblast | Ivan Susloparov (State Research Center of Virology and Biotechnology (VECTOR) / Emerging Zoonotic Diseases and Influenza) |
| EPI_ISL_320661 | A/goose/Samara/459/2018 | A / H5N8 | Europe / Russian Federation / Samara Oblast | Ivan Susloparov (State Research Center of Virology and Biotechnology (VECTOR) / Emerging Zoonotic Diseases and Influenza) |
| EPI_ISL_320662 | A/chicken/Penza/300/2018 | A / H5N8 | Europe / Russian Federation / Penza Oblast | Ivan Susloparov (State Research Center of Virology and Biotechnology (VECTOR) / Emerging Zoonotic Diseases and Influenza) |
| EPI_ISL_320663 | A/chicken/Penza/301/2018 | A / H5N8 | Europe / Russian Federation / Penza Oblast | Ivan Susloparov (State Research Center of Virology and Biotechnology (VECTOR) / Emerging Zoonotic Diseases and Influenza) |
| EPI_ISL_320664 | A/chicken/Orel/533/2018 | A / H5N8 | Europe / Russian Federation / Oryol Oblast | Ivan Susloparov (State Research Center of Virology and Biotechnology (VECTOR) / Emerging Zoonotic Diseases and Influenza) |
| EPI_ISL_320674 | A/chicken/Rostov-on-Don/766/2018 | A / H5N8 | Europe / Russian Federation / Rostov Oblast | Ivan Susloparov (State Research Center of Virology and Biotechnology (VECTOR) / Emerging Zoonotic Diseases and Influenza) |
| EPI_ISL_320675 | A/turkey/Rostov-on-Don/817/2018 | A / H5N8 | Europe / Russian Federation / Rostov Oblast | Ivan Susloparov (State Research Center of Virology and Biotechnology (VECTOR) / Emerging Zoonotic Diseases and Influenza) |
| EPI_ISL_320676 | A/chicken/Penza/605/2018 | A / H5N8 | Europe / Russian Federation / Penza Oblast | Ivan Susloparov (State Research Center of Virology and Biotechnology (VECTOR) / Emerging Zoonotic Diseases and Influenza) |

| | | | | |
|---|---|---|---|---|
| EPI_ISL_320677 | A/chicken/Penza/607/2018 | A / H5N8 | Europe / Russian Federation / Penza Oblast | Ivan Susloparov (State Research Center of Virology and Biotechnology (VECTOR) / Emerging Zoonotic Diseases and Influenza) |
| EPI_ISL_320678 | A/chicken/Cheboksary/805/2018 | A / H5N8 | Europe / Russian Federation / Chuvash Republic-Chuvashia | Ivan Susloparov (State Research Center of Virology and Biotechnology (VECTOR) / Emerging Zoonotic Diseases and Influenza) |
| EPI_ISL_320679 | A/chicken/Cheboksary/806/2018 | A / H5N8 | Europe / Russian Federation / Chuvash Republic-Chuvashia | Ivan Susloparov (State Research Center of Virology and Biotechnology (VECTOR) / Emerging Zoonotic Diseases and Influenza) |
| EPI_ISL_320680 | A/chicken/Kursk/757/2018 | A / H5N8 | Europe / Russian Federation / Kursk Oblast | Ivan Susloparov (State Research Center of Virology and Biotechnology (VECTOR) / Emerging Zoonotic Diseases and Influenza) |
| EPI_ISL_320681 | A/chicken/Kursk/760/2018 | A / H5N8 | Europe / Russian Federation / Kursk Oblast | Ivan Susloparov (State Research Center of Virology and Biotechnology (VECTOR) / Emerging Zoonotic Diseases and Influenza) |
| EPI_ISL_320682 | A/chicken/Kursk/762/2018 | A / H5N8 | Europe / Russian Federation / Kursk Oblast | Ivan Susloparov (State Research Center of Virology and Biotechnology (VECTOR) / Emerging Zoonotic Diseases and Influenza) |
| EPI_ISL_320683 | A/goose/Samara/673/2018 | A / H5N8 | Europe / Russian Federation / Samara Oblast | Ivan Susloparov (State Research Center of Virology and Biotechnology (VECTOR) / Emerging Zoonotic Diseases and Influenza) |
| EPI_ISL_320684 | A/goose/Samara/675/2018 | A / H5N8 | Europe / Russian Federation / Samara Oblast | Ivan Susloparov (State Research Center of Virology and Biotechnology (VECTOR) / Emerging Zoonotic Diseases and Influenza) |
| EPI_ISL_320685 | A/chicken/Samara/679/2018 | A / H5N8 | Europe / Russian Federation / Samara Oblast | Ivan Susloparov (State Research Center of Virology and Biotechnology (VECTOR) / Emerging Zoonotic Diseases and Influenza) |
| EPI_ISL_320953 | A/chicken/Cheboksary/849/2018 | A / H5N8 | Europe / Russian Federation / Chuvash Republic-Chuvashia | Ivan Susloparov (State Research Center of Virology and Biotechnology (VECTOR) / Emerging Zoonotic Diseases and Influenza) |
| EPI_ISL_320954 | A/chicken/Cheboksary/850/2018 | A / H5N8 | Europe / Russian Federation / Chuvash Republic-Chuvashia | Ivan Susloparov (State Research Center of Virology and Biotechnology (VECTOR) / Emerging Zoonotic Diseases and Influenza) |
| EPI_ISL_320955 | A/chicken/Cheboksary/851/2018 | A / H5N8 | Europe / Russian Federation / Chuvash Republic-Chuvashia | Ivan Susloparov (State Research Center of Virology and Biotechnology (VECTOR) / Emerging Zoonotic Diseases and Influenza) |
| EPI_ISL_320956 | A/chicken/Cheboksary/853/2018 | A / H5N8 | Europe / Russian Federation / Chuvash Republic-Chuvashia | Ivan Susloparov (State Research Center of Virology and Biotechnology (VECTOR) / Emerging Zoonotic Diseases and Influenza) |
| EPI_ISL_320957 | A/chicken/Cheboksary/854/2018 | A / H5N8 | Europe / Russian Federation / Chuvash Republic-Chuvashia | Ivan Susloparov (State Research Center of Virology and Biotechnology (VECTOR) / Emerging Zoonotic Diseases and Influenza) |
| EPI_ISL_320958 | A/chicken/Tatarstan/7/2018 | A / H5N8 | Europe / Russian Federation / Republic of Tatarstan | Ivan Susloparov (State Research Center of Virology and Biotechnology (VECTOR) / Emerging Zoonotic Diseases and Influenza) |
| EPI_ISL_322179 | A/domestic duck/Germany-MV/AR613-L02727/2018 | A / H5N6 | Europe / Germany / Mecklenburg-Vorpommern / Nordwestmecklenburg | Jacqueline King (Friedrich-Loeffler-Institut) |
| EPI_ISL_322984 | A/Grey seal/361-10/BalticPL/16 | A / H5N8 | Europe / Poland | Dai-Lun Shin (Stiftung Tierärztliche Hochschule Hannover) |
| EPI_ISL_3290184 | A/Canada Goose/Sweden/SVA210330SZ0426/FB001307/E-202 | A / H5N8 | Europe / Sweden / Ostergotlands Lan / Norrkopings Kommun | Siamak Zohari (National Veterinary Institute) |
| EPI_ISL_3294584 | A/Mute Swan/Sweden/SVA210330SZ0417/FB001305/D-2021 | A / H5N5 | Europe / Sweden / Sodermanlands Lan / Oxelosunds Kummun | Siamak Zohari (National Veterinary Institute) |
| EPI_ISL_3298195 | A/common buzzard/Sweden/SVA210329SZ0208/FB0001278/M | A / H5N8 | Europe / Sweden / Skane Lan / Simrishamns Kommun | Siamak Zohari (National Veterinary Institute) |
| EPI_ISL_3298234 | A/peregrine falcon/Sweden/SVA210325SZ0348/FP001210/H-2 | A / H5N4 | Europe / Sweden / Kalmar Lan / Vasterviks Kommun | Siamak Zohari (National Veterinary Institute) |
| EPI_ISL_3298300 | A/Greylag goose/Sweden/SVA210324SZ0403/FB001169/N-202 | A / H5N8 | Europe / Sweden / Hallands Lan / Varbergs Kommun | Siamak Zohari (National Veterinary Institute) |
| EPI_ISL_331223 | A/Duck/England/1279/2014 | A / H5N8 | Europe / United Kingdom / East Riding of Yorkshire | Alex Byrne (Animal and Plant Health Agency (APHA) / Virology Department) |
| EPI_ISL_3314946 | A/common buzzard/Sweden/SVA10324SZ0359/FB001163/M-2 | A / H5N8 | Europe / Sweden / Skane Lan | Siamak Zohari (National Veterinary Institute) |
| EPI_ISL_3314973 | A/common buzzard/Sweden/SVA210323SZ0426/FB001120/O- | A / H5N8 | Europe / Sweden / Vastra Gotalands Lan / Lidkopings Kommun | Siamak Zohari (National Veterinary Institute) |
| EPI_ISL_3315298 | A/whooper swan/Sweden/SVA210318SZ0304/FB0001089/M-2 | A / H5N8 | Europe / Sweden / Skane Lan / Simrishamns Kommun | Siamak Zohari (National Veterinary Institute) |
| EPI_ISL_3319316 | A/common eider/Sweden/SVA210729SZ0320/FB002613/I-202 | A / H5N8 | Europe / Sweden / Gotlands Lan / Gotlands Kommun | Siamak Zohari (National Veterinary Institute) |
| EPI_ISL_3319859 | A/common eider/Sweden/SVA210729SZ0323/FB002615/I-202 | A / H5N8 | Europe / Sweden / Gotlands Lan / Gotlands Kommun | Siamak Zohari (National Veterinary Institute) |
| EPI_ISL_332400 | A/Jiangsu/1/2018 | A / H5N6 | Asia / China | Xian Qi (Jiangsu Provincial Center for Disease Control & Prevention) |
| EPI_ISL_332430 | A/Chicken/Netherlands/18003041-001-005/2018 | A / H5N6 | Europe / Netherlands / Oldekerk | Saskia Bergervoet (Wageningen Bioveterinary Research) |
| EPI_ISL_332431 | A/Duck/Netherlands/18003885-001-005/2018 | A / H5N6 | Europe / Netherlands / Kamperveen | Saskia Bergervoet (Wageningen Bioveterinary Research) |
| EPI_ISL_332432 | A/Guineafowl/Netherlands/17017775-020-024/2017 | A / H5N6 | Europe / Netherlands / Biddinghuizen | Saskia Bergervoet (Wageningen Bioveterinary Research) |
| EPI_ISL_332433 | A/Bird/Netherlands/17017775-035-039/2017 | A / H5N6 | Europe / Netherlands / Biddinghuizen | Saskia Bergervoet (Wageningen Bioveterinary Research) |
| EPI_ISL_332434 | A/Peacock/Netherlands/17017775-064-068/2017 | A / H5N6 | Europe / Netherlands / Biddinghuizen | Saskia Bergervoet (Wageningen Bioveterinary Research) |
| EPI_ISL_332435 | A/Chicken/Netherlands/18000887-005/2018 | A / H5N6 | Europe / Netherlands / Rhoon | Saskia Bergervoet (Wageningen Bioveterinary Research) |
| EPI_ISL_332436 | A/Peacock/Netherlands/18000887-006/2018 | A / H5N6 | Europe / Netherlands / Rhoon | Saskia Bergervoet (Wageningen Bioveterinary Research) |
| EPI_ISL_332437 | A/Turkey/Netherlands/18000887-007-010/2018 | A / H5N6 | Europe / Netherlands / Rhoon | Saskia Bergervoet (Wageningen Bioveterinary Research) |
| EPI_ISL_332438 | A/Mute swan/Netherlands/17017903-002/2017 | A / H5N6 | Europe / Netherlands / Elburg | Saskia Bergervoet (Wageningen Bioveterinary Research) |
| EPI_ISL_332439 | A/Perigrine falcon/Netherlands/18003274-001/2018 | A / H5N6 | Europe / Netherlands / Westernieland | Saskia Bergervoet (Wageningen Bioveterinary Research) |
| EPI_ISL_332440 | A/Buzzard/Netherlands/18004242-002/2018 | A / H5N6 | Europe / Netherlands / Zandfoort | Saskia Bergervoet (Wageningen Bioveterinary Research) |
| EPI_ISL_332441 | A/Mallard/Netherlands/18012508-017/2018 | A / H5N6 | Europe / Netherlands / Eemmeer | Saskia Bergervoet (Wageningen Bioveterinary Research) |
| EPI_ISL_332804 | A/duck/Bangladesh/19D857/2017 | A / H5N6 | Asia / Bangladesh | Yunho Jang (Centers for Disease Control and Prevention / WHO Collaborating Center for Surveillance, Epidemiology and Control of Influ |
| EPI_ISL_332905 | A/duck/Bangladesh/19D853/2017 | A / H5N6 | Asia / Bangladesh | Yunho Jang (Centers for Disease Control and Prevention / WHO Collaborating Center for Surveillance, Epidemiology and Control of Influ |
| EPI_ISL_332947 | A/duck/Bangladesh/19D851/2017 | A / H5N6 | Asia / Bangladesh | Yunho Jang (Centers for Disease Control and Prevention / WHO Collaborating Center for Surveillance, Epidemiology and Control of Influ |
| EPI_ISL_332948 | A/duck/Bangladesh/19D849/2017 | A / H5N6 | Asia / Bangladesh | Yunho Jang (Centers for Disease Control and Prevention / WHO Collaborating Center for Surveillance, Epidemiology and Control of Influ |
| EPI_ISL_333130 | A/goose/Bangladesh/19D855/2017 | A / H5N6 | Asia / Bangladesh | Yunho Jang (Centers for Disease Control and Prevention / WHO Collaborating Center for Surveillance, Epidemiology and Control of Influ |
| EPI_ISL_333131 | A/duck/Bangladesh/19D852/2017 | A / H5N6 | Asia / Bangladesh | Yunho Jang (Centers for Disease Control and Prevention / WHO Collaborating Center for Surveillance, Epidemiology and Control of Influ |
| EPI_ISL_333132 | A/goose/Bangladesh/19D820/2017 | A / H5N6 | Asia / Bangladesh | Yunho Jang (Centers for Disease Control and Prevention / WHO Collaborating Center for Surveillance, Epidemiology and Control of Influ |
| EPI_ISL_333133 | A/duck/Bangladesh/19D770/2017 | A / H5N6 | Asia / Bangladesh | Yunho Jang (Centers for Disease Control and Prevention / WHO Collaborating Center for Surveillance, Epidemiology and Control of Influ |
| EPI_ISL_333134 | A/goose/Bangladesh/19D764/2017 | A / H5N6 | Asia / Bangladesh | Yunho Jang (Centers for Disease Control and Prevention / WHO Collaborating Center for Surveillance, Epidemiology and Control of Influ |
| EPI_ISL_333629 | A/Environment/Fujiansanyuan/08/2017 | A / H5N6 | Asia / China / Fujian Province | Xie Jianfeng (Fujian Center for Disease Control and Prevention) |
| EPI_ISL_333630 | A/Environment/Fujian/05326/2016 | A / H5N6 | Asia / China / Fujian Province | Xie Jianfeng (Fujian Center for Disease Control and Prevention) |
| EPI_ISL_333631 | A/Environment/Fujian/05324/2016 | A / H5N6 | Asia / China / Fujian Province | Xie Jianfeng (Fujian Center for Disease Control and Prevention) |
| EPI_ISL_333632 | A/Environment/Fujian/09991/2016 | A / H5N6 | Asia / China / Fujian Province | Xie Jianfeng (Fujian Center for Disease Control and Prevention) |
| EPI_ISL_333633 | A/Environment/Fujian/28681/2016 | A / H5N6 | Asia / China / Fujian Province | Xie Jianfeng (Fujian Center for Disease Control and Prevention) |
| EPI_ISL_333634 | A/Environment/Fujian/28686/2016 | A / H5N6 | Asia / China / Fujian Province | Xie Jianfeng (Fujian Center for Disease Control and Prevention) |
| EPI_ISL_333636 | A/Environment/Fujian/52356/2016 | A / H5N6 | Asia / China / Fujian Province | Xie Jianfeng (Fujian Center for Disease Control and Prevention) |
| EPI_ISL_333975 | A/chicken/Vietnam/NCVD-16A26/2016 | A / H5N6 | Asia / Vietnam | Yunho Jang (Centers for Disease Control and Prevention / WHO Collaborating Center for Surveillance, Epidemiology and Control of Influ |
| EPI_ISL_334419 | A/chicken/Kagawa/13T/2018 | A / H5N6 | Asia / Japan | Takehiko Saito (National Institute of Animal Health) |
| EPI_ISL_334420 | A/chicken/Kagawa/11C-2/2018 | A / H5N6 | Asia / Japan | Takehiko Saito (National Institute of Animal Health) |
| EPI_ISL_334421 | A/chicken/Kagawa/12C-1/2018 | A / H5N6 | Asia / Japan | Takehiko Saito (National Institute of Animal Health) |
| EPI_ISL_334422 | A/chicken/Kagawa/12T-2/2018 | A / H5N6 | Asia / Japan | Takehiko Saito (National Institute of Animal Health) |
| EPI_ISL_334423 | A/chicken/Kagawa/14T-2/2018 | A / H5N6 | Asia / Japan | Takehiko Saito (National Institute of Animal Health) |
| EPI_ISL_334424 | A/chicken/Kagawa/14T-1/2018 | A / H5N6 | Asia / Japan | Takehiko Saito (National Institute of Animal Health) |
| EPI_ISL_334425 | A/chicken/Kagawa/12T-1/2018 | A / H5N6 | Asia / Japan | Takehiko Saito (National Institute of Animal Health) |
| EPI_ISL_334426 | A/chicken/Kagawa/15T/2018 | A / H5N6 | Asia / Japan | Takehiko Saito (National Institute of Animal Health) |
| EPI_ISL_334427 | A/chicken/Kagawa/11T-1/2018 | A / H5N6 | Asia / Japan | Takehiko Saito (National Institute of Animal Health) |
| EPI_ISL_334428 | A/chicken/Kagawa/11C-1/2018 | A / H5N6 | Asia / Japan | Takehiko Saito (National Institute of Animal Health) |
| EPI_ISL_334430 | A/chicken/Kagawa/11T-2/2018 | A / H5N6 | Asia / Japan | Takehiko Saito (National Institute of Animal Health) |
| EPI_ISL_334431 | A/chicken/Kagawa/12C-2/2018 | A / H5N6 | Asia / Japan | Takehiko Saito (National Institute of Animal Health) |
| EPI_ISL_335257 | A/chicken/Kagawa/2C-2/2018 | A / H5N6 | Asia / Japan | Takehiko Saito (National Institute of Animal Health) |
| EPI_ISL_335258 | A/chicken/Kagawa/4T-1/2018 | A / H5N6 | Asia / Japan | Takehiko Saito (National Institute of Animal Health) |

| | | | |
|---|---|---|---|
| EPI_ISL_335259 | A/chicken/Kagawa/5C-1/2018 | A / H5N6 Asia / Japan | Takehiko Saito (National Institute of Animal Health) |
| EPI_ISL_335260 | A/chicken/Kagawa/4C-1/2018 | A / H5N6 Asia / Japan | Takehiko Saito (National Institute of Animal Health) |
| EPI_ISL_335261 | A/chicken/Kagawa/5T-1/2018 | A / H5N6 Asia / Japan | Takehiko Saito (National Institute of Animal Health) |
| EPI_ISL_335262 | A/chicken/Kagawa/1T-2/2018 | A / H5N6 Asia / Japan | Takehiko Saito (National Institute of Animal Health) |
| EPI_ISL_335263 | A/chicken/Kagawa/2C-1/2018 | A / H5N6 Asia / Japan | Takehiko Saito (National Institute of Animal Health) |
| EPI_ISL_335264 | A/Jungle crow/Hyogo/2803E023C/2018 | A / H5N6 Asia / Japan | Takehiko Saito (National Institute of Animal Health) |
| EPI_ISL_335265 | A/chicken/Kagawa/2T-1/2018 | A / H5N6 Asia / Japan | Takehiko Saito (National Institute of Animal Health) |
| EPI_ISL_335266 | A/chicken/Kagawa/2T-2/2018 | A / H5N6 Asia / Japan | Takehiko Saito (National Institute of Animal Health) |
| EPI_ISL_335268 | A/chicken/Kagawa/5C-2/2018 | A / H5N6 Asia / Japan | Takehiko Saito (National Institute of Animal Health) |
| EPI_ISL_335269 | A/chicken/Kagawa/3T-2/2018 | A / H5N6 Asia / Japan | Takehiko Saito (National Institute of Animal Health) |
| EPI_ISL_335270 | A/chicken/Kagawa/3C-2/2018 | A / H5N6 Asia / Japan | Takehiko Saito (National Institute of Animal Health) |
| EPI_ISL_335271 | A/chicken/Kagawa/3T-1/2018 | A / H5N6 Asia / Japan | Takehiko Saito (National Institute of Animal Health) |
| EPI_ISL_335272 | A/chicken/Kagawa/3C-1/2018 | A / H5N6 Asia / Japan | Takehiko Saito (National Institute of Animal Health) |
| EPI_ISL_335273 | A/Jungle crow/Hyogo/2803E023T/2018 | A / H5N6 Asia / Japan | Takehiko Saito (National Institute of Animal Health) |
| EPI_ISL_335440 | A/mandarin_duck/Korea/H119-4/2017 | A / H5N6 Asia / Korea, Republic of | Yu-Na Lee (Animal and Plant Quarantine Agency (S-2158) / Avian Influenza Research & Diagnostic Division) |
| EPI_ISL_335441 | A/duck/Korea/H125-1/2018 | A / H5N6 Asia / Korea, Republic of | Yu-Na Lee (Animal and Plant Quarantine Agency (S-2158) / Avian Influenza Research & Diagnostic Division) |
| EPI_ISL_335442 | A/duck/Korea/H125-2/2018 | A / H5N6 Asia / Korea, Republic of | Yu-Na Lee (Animal and Plant Quarantine Agency (S-2158) / Avian Influenza Research & Diagnostic Division) |
| EPI_ISL_335443 | A/duck/Korea/H182/2018 | A / H5N6 Asia / Korea, Republic of | Yu-Na Lee (Animal and Plant Quarantine Agency (S-2158) / Avian Influenza Research & Diagnostic Division) |
| EPI_ISL_335444 | A/duck/Korea/H187/2018 | A / H5N6 Asia / Korea, Republic of | Yu-Na Lee (Animal and Plant Quarantine Agency (S-2158) / Avian Influenza Research & Diagnostic Division) |
| EPI_ISL_335445 | A/duck/Korea/H191/2018 | A / H5N6 Asia / Korea, Republic of | Yu-Na Lee (Animal and Plant Quarantine Agency (S-2158) / Avian Influenza Research & Diagnostic Division) |
| EPI_ISL_335446 | A/duck/Korea/H192/2018 | A / H5N6 Asia / Korea, Republic of | Yu-Na Lee (Animal and Plant Quarantine Agency (S-2158) / Avian Influenza Research & Diagnostic Division) |
| EPI_ISL_335447 | A/duck/Korea/H196/2018 | A / H5N6 Asia / Korea, Republic of | Yu-Na Lee (Animal and Plant Quarantine Agency (S-2158) / Avian Influenza Research & Diagnostic Division) |
| EPI_ISL_335448 | A/chicken/Korea/H203/2018 | A / H5N6 Asia / Korea, Republic of | Yu-Na Lee (Animal and Plant Quarantine Agency (S-2158) / Avian Influenza Research & Diagnostic Division) |
| EPI_ISL_335449 | A/chicken/Korea/H204/2018 | A / H5N6 Asia / Korea, Republic of | Yu-Na Lee (Animal and Plant Quarantine Agency (S-2158) / Avian Influenza Research & Diagnostic Division) |
| EPI_ISL_335450 | A/chicken/Korea/H214/2018 | A / H5N6 Asia / Korea, Republic of | Yu-Na Lee (Animal and Plant Quarantine Agency (S-2158) / Avian Influenza Research & Diagnostic Division) |
| EPI_ISL_335451 | A/chicken/Korea/H219/2018 | A / H5N6 Asia / Korea, Republic of | Yu-Na Lee (Animal and Plant Quarantine Agency (S-2158) / Avian Influenza Research & Diagnostic Division) |
| EPI_ISL_335452 | A/duck/Korea/H229/2018 | A / H5N6 Asia / Korea, Republic of | Yu-Na Lee (Animal and Plant Quarantine Agency (S-2158) / Avian Influenza Research & Diagnostic Division) |
| EPI_ISL_335453 | A/chicken/Korea/H238/2018 | A / H5N6 Asia / Korea, Republic of | Yu-Na Lee (Animal and Plant Quarantine Agency (S-2158) / Avian Influenza Research & Diagnostic Division) |
| EPI_ISL_335454 | A/chicken/Korea/H239/2018 | A / H5N6 Asia / Korea, Republic of | Yu-Na Lee (Animal and Plant Quarantine Agency (S-2158) / Avian Influenza Research & Diagnostic Division) |
| EPI_ISL_335455 | A/chicken/Korea/H241/2018 | A / H5N6 Asia / Korea, Republic of | Yu-Na Lee (Animal and Plant Quarantine Agency (S-2158) / Avian Influenza Research & Diagnostic Division) |
| EPI_ISL_335456 | A/spotbill_duck/Korea/WA159/2018 | A / H5N6 Asia / Korea, Republic of | Yu-Na Lee (Animal and Plant Quarantine Agency (S-2158) / Avian Influenza Research & Diagnostic Division) |
| EPI_ISL_335457 | A/mute swan/Shimane/3211A002/2017 | A / H5N6 Asia / Japan | Takehiko Saito (National Institute of Animal Health) |
| EPI_ISL_335458 | A/Jungle crow/Hyogo/2803E026T/2018 | A / H5N6 Asia / Japan | Takehiko Saito (National Institute of Animal Health) |
| EPI_ISL_335459 | A/Jungle crow/Hyogo/2803E025C/2018 | A / H5N6 Asia / Japan | Takehiko Saito (National Institute of Animal Health) |
| EPI_ISL_335460 | A/Jungle crow/Hyogo/2803E028C/2018 | A / H5N6 Asia / Japan | Takehiko Saito (National Institute of Animal Health) |
| EPI_ISL_335461 | A/Jungle crow/Hyogo/2803E026C/2018 | A / H5N6 Asia / Japan | Takehiko Saito (National Institute of Animal Health) |
| EPI_ISL_335462 | A/Jungle crow/Hyogo/2803E028T/2018 | A / H5N6 Asia / Japan | Takehiko Saito (National Institute of Animal Health) |
| EPI_ISL_335463 | A/Northern Goshawk/Tokyo/1301B003/2018 | A / H5N6 Asia / Japan | Takehiko Saito (National Institute of Animal Health) |
| EPI_ISL_335464 | A/Jungle crow/Hyogo/2803E027C/2018 | A / H5N6 Asia / Japan | Takehiko Saito (National Institute of Animal Health) |
| EPI_ISL_335465 | A/Jungle crow/Hyogo/2803E024T/2018 | A / H5N6 Asia / Japan | Takehiko Saito (National Institute of Animal Health) |
| EPI_ISL_335466 | A/Jungle crow/Hyogo/2803E025T/2018 | A / H5N6 Asia / Japan | Takehiko Saito (National Institute of Animal Health) |
| EPI_ISL_335467 | A/Jungle crow/Hyogo/2803E027T/2018 | A / H5N6 Asia / Japan | Takehiko Saito (National Institute of Animal Health) |
| EPI_ISL_335468 | A/Jungle crow/Hyogo/2803E024C/2018 | A / H5N6 Asia / Japan | Takehiko Saito (National Institute of Animal Health) |
| EPI_ISL_336925 | A/common gull/Saratov/1676/2018 | A / H5N6 Europe / Russian Federation / Saratov Oblast / Engelssky District | Natalia Goncharova (State Research Center of Virology and Biotechnology (VECTOR) / Emerging Zoonotic Diseases and Influenza) |
| EPI_ISL_336929 | A/chicken/Voronezh/1488/2018 | A / H5N8 Europe / Russian Federation / Voronezh Oblast | Alexey Danilenko (State Research Center of Virology and Biotechnology (VECTOR) / Emerging Zoonotic Diseases and Influenza) |
| EPI_ISL_336930 | A/turkey/Rostov-on-Don/1117/2018 | A / H5N8 Europe / Russian Federation / Rostov Oblast | Alexey Danilenko (State Research Center of Virology and Biotechnology (VECTOR) / Emerging Zoonotic Diseases and Influenza) |
| EPI_ISL_336931 | A/chiken/Voronezh/1491/2018 | A / H5N8 Europe / Russian Federation / Voronezh Oblast | Alexey Danilenko (State Research Center of Virology and Biotechnology (VECTOR) / Emerging Zoonotic Diseases and Influenza) |
| EPI_ISL_336932 | A/chicken/Voronezh/1504/2018 | A / H5N8 Europe / Russian Federation / Voronezh Oblast | Alexey Danilenko (State Research Center of Virology and Biotechnology (VECTOR) / Emerging Zoonotic Diseases and Influenza) |
| EPI_ISL_336933 | A/chicken/Voronezh/1513/2018 | A / H5N8 Europe / Russian Federation / Voronezh Oblast | Alexey Danilenko (State Research Center of Virology and Biotechnology (VECTOR) / Emerging Zoonotic Diseases and Influenza) |
| EPI_ISL_337151 | A/chicken/Mari El/870/2018 | A / H5N8 Europe / Russian Federation / Mari El Republic | Natalia Goncharova (State Research Center of Virology and Biotechnology (VECTOR) / Emerging Zoonotic Diseases and Influenza) |
| EPI_ISL_337273 | A/Guangxi/31906/2018 | A / H5N6 Asia / China / Guangxi | Lei Yang (WHO Chinese National Influenza Center / Virology Institute, Chinese CDC) |
| EPI_ISL_337274 | A/Guangdong/18SF020/2018 | A / H5N6 Asia / China / Guangdong | Lei Yang (WHO Chinese National Influenza Center / Virology Institute, Chinese CDC) |
| EPI_ISL_337275 | A/Jiangsu/32888/2018 | A / H5N6 Asia / China / Jiangsu | Lei Yang (WHO Chinese National Influenza Center / Virology Institute, Chinese CDC) |
| EPI_ISL_337276 | A/Guangxi/32797/2018 | A / H5N6 Asia / China / Guangxi | Lei Yang (WHO Chinese National Influenza Center / Virology Institute, Chinese CDC) |
| EPI_ISL_337280 | A/Guangxi/13486/2017 | A / H5N6 Asia / China / Guangxi | Lei Yang (WHO Chinese National Influenza Center / Virology Institute, Chinese CDC) |
| EPI_ISL_337410 | A/Turkey/Central Russia/5/2018 | A / H5N8 Europe / Russian Federation | Takehiko Saito (National Institute of Animal Health) |
| EPI_ISL_338359 | A/chicken/Central Russia/1/2017 | A / H5N2 Europe / Russian Federation | Takehiko Saito (National Institute of Animal Health) |
| EPI_ISL_339105 | A/Duck/France/RG1/2016 (H5N8) | A / H5N8 Europe / France | Thomas Figueroa (Ecole Veterinaire de Toulouse / UMR1225 IHAP Interactions hôtes-agents pathogènes) |
| EPI_ISL_339932 | A/Muscovy duck/Long An/AI470/2018 | A / H5N6 Asia / Vietnam | Takehiko Saito (National Institute of Animal Health) |
| EPI_ISL_340075 | A/chicken/Tatarstan/112/2017 | A / H5N8 Europe / Russian Federation / Republic of Tatarstan | Svetlana Svyatchenko (State Research Center of Virology and Biotechnology (VECTOR) / Emerging Zoonotic Diseases and Influenza) |
| EPI_ISL_340453 | A/Env/Guangdong/172/2017 | A / H5N6 Asia / China / Guangdong Province | Ru Bai (Guangdong Provincial Center for Disease Control and Prevention) |
| EPI_ISL_340786 | A/Env/Guangdong/C17059172/YF/2017-03-06 | A / H5N6 Asia / China | Ru Bai (Guangdong Provincial Center for Disease Control and Prevention) |
| EPI_ISL_340788 | A/Env/Guangdong/zhanjiang/C17277346/2017-12-05 | A / H5N6 Asia / China | Ru Bai (Guangdong Provincial Center for Disease Control and Prevention) |
| EPI_ISL_340789 | A/Duck/Guangdong/PO17281256/MZH/2017-8-21 | A / H5N6 Asia / China | Ru Bai (Guangdong Provincial Center for Disease Control and Prevention) |
| EPI_ISL_340791 | A/Env/Guangdong/C172811415/MZH/2017-10-16 | A / H5N6 Asia / China | Ru Bai (Guangdong Provincial Center for Disease Control and Prevention) |
| EPI_ISL_340792 | A/Duck/Guangdong/PO17281388/MZH/2017-10-24 | A / H5N6 Asia / China | Ru Bai (Guangdong Provincial Center for Disease Control and Prevention) |
| EPI_ISL_340793 | A/Env/Guangdong/C172811414/MZH/2017-10-16 | A / H5N6 Asia / China | Ru Bai (Guangdong Provincial Center for Disease Control and Prevention) |
| EPI_ISL_340794 | A/Env/Guangdong/zhanjiang/C18277209/2018-05-21 | A / H5N6 Asia / China | Ru Bai (Guangdong Provincial Center for Disease Control and Prevention) |
| EPI_ISL_340795 | A/Env/Guangdong/Huizhou/C18280077/2018-02-22 | A / H5N6 Asia / China | Ru Bai (Guangdong Provincial Center for Disease Control and Prevention) |
| EPI_ISL_340796 | A/Guangdong/18SF020/2018-09-30 | A / H5N6 Asia / China | Ru Bai (Guangdong Provincial Center for Disease Control and Prevention) |
| EPI_ISL_340797 | A/Env/Guangdong/Huizhou/C18280031/2018-01-22 | A / H5N6 Asia / China | Ru Bai (Guangdong Provincial Center for Disease Control and Prevention) |
| EPI_ISL_340798 | A/Env/Guangdong/zhongshan/C182870234/2018-04-24 | A / H5N6 Asia / China | Ru Bai (Guangdong Provincial Center for Disease Control and Prevention) |

| | | | | |
|---|---|---|---|---|
| EPI_ISL_340799 | A/Env/Guangdong/Huizhou/C17280804/2017-11-21 | A / H5N6 | Asia / China | Ru Bai (Guangdong Provincial Center for Disease Control and Prevention) |
| EPI_ISL_340800 | A/Env/Guangdong/zhanjiang/C18277050/2018-01-29 | A / H5N6 | Asia / China | Ru Bai (Guangdong Provincial Center for Disease Control and Prevention) |
| EPI_ISL_340805 | A/Env/Guangdong/Shenzhen/C18061184/2018-05-08 | A / H5N6 | Asia / China | Ru Bai (Guangdong Provincial Center for Disease Control and Prevention) |
| EPI_ISL_340806 | A/Env/Guangdong/Qingyuan/C18285099/2018-02-28 | A / H5N6 | Asia / China | Ru Bai (Guangdong Provincial Center for Disease Control and Prevention) |
| EPI_ISL_340807 | A/Env/Guangdong/Foshan/C182750200/2018-03-06 | A / H5N6 | Asia / China | Ru Bai (Guangdong Provincial Center for Disease Control and Prevention) |
| EPI_ISL_340808 | A/Env/Guangdong/Foshan/C182753007/2018-01-02 | A / H5N6 | Asia / China | Ru Bai (Guangdong Provincial Center for Disease Control and Prevention) |
| EPI_ISL_340809 | A/Env/Guangdong/Foshan/C172750608/2017-12-26 | A / H5N6 | Asia / China | Ru Bai (Guangdong Provincial Center for Disease Control and Prevention) |
| EPI_ISL_340810 | A/Env/Guangdong/Foshan/C172750600/2017-12-26 | A / H5N6 | Asia / China | Ru Bai (Guangdong Provincial Center for Disease Control and Prevention) |
| EPI_ISL_340811 | A/Env/Guangdong/Foshan/C172750528/2017-12-05 | A / H5N6 | Asia / China | Ru Bai (Guangdong Provincial Center for Disease Control and Prevention) |
| EPI_ISL_340812 | A/Env/Guangdong/Dongguan/C172863577/2017-12-04 | A / H5N6 | Asia / China | Ru Bai (Guangdong Provincial Center for Disease Control and Prevention) |
| EPI_ISL_340813 | A/Env/Guangdong/C17272382/SHG/2017-11-14 | A / H5N6 | Asia / China | Ru Bai (Guangdong Provincial Center for Disease Control and Prevention) |
| EPI_ISL_340814 | A/Env/Guangdong/C172752325/FSH/2017-11-27 | A / H5N6 | Asia / China | Ru Bai (Guangdong Provincial Center for Disease Control and Prevention) |
| EPI_ISL_340815 | A/Env/Guangdong/C172752326/FSH/2017-11-27 | A / H5N6 | Asia / China | Ru Bai (Guangdong Provincial Center for Disease Control and Prevention) |
| EPI_ISL_340816 | A/Env/Guangdong/zhongshan/C182870028/2018-01-15 | A / H5N6 | Asia / China | Ru Bai (Guangdong Provincial Center for Disease Control and Prevention) |
| EPI_ISL_340817 | A/Env/Guangdong/C172752329/FSH/2017-11-27 | A / H5N6 | Asia / China | Ru Bai (Guangdong Provincial Center for Disease Control and Prevention) |
| EPI_ISL_340818 | A/Env/Guangdong/Foshan/C182750024/2018-01-09 | A / H5N6 | Asia / China | Ru Bai (Guangdong Provincial Center for Disease Control and Prevention) |
| EPI_ISL_340819 | A/Env/Guangdong/Jieyang/C17289387/2017-12-05 | A / H5N6 | Asia / China | Ru Bai (Guangdong Provincial Center for Disease Control and Prevention) |
| EPI_ISL_340820 | A/Env/Guangdong/Jieyang/C17289388/2017-12-05 | A / H5N6 | Asia / China | Ru Bai (Guangdong Provincial Center for Disease Control and Prevention) |
| EPI_ISL_340821 | A/Env/Guangdong/Jieyang/C17289389/2017-12-05 | A / H5N6 | Asia / China | Ru Bai (Guangdong Provincial Center for Disease Control and Prevention) |
| EPI_ISL_340822 | A/Env/Guangdong/Jieyang/C17289390/2017-12-05 | A / H5N6 | Asia / China | Ru Bai (Guangdong Provincial Center for Disease Control and Prevention) |
| EPI_ISL_340823 | A/Env/Guangdong/zhanjiang/C18277136/2018-04-02 | A / H5N6 | Asia / China | Ru Bai (Guangdong Provincial Center for Disease Control and Prevention) |
| EPI_ISL_340824 | A/Env/Guangdong/Foshan/C182750085/2018-01-30 | A / H5N6 | Asia / China | Ru Bai (Guangdong Provincial Center for Disease Control and Prevention) |
| EPI_ISL_340825 | A/Env/Guangdong/Jieyang/C18289059/2018-02-06 | A / H5N6 | Asia / China | Ru Bai (Guangdong Provincial Center for Disease Control and Prevention) |
| EPI_ISL_340826 | A/Env/Guangdong/C17059359/YF/2017-05-31 | A / H5N6 | Asia / China | Ru Bai (Guangdong Provincial Center for Disease Control and Prevention) |
| EPI_ISL_340827 | A/Env/Guangdong/C172790853/ZHQ/2017-10-30 | A / H5N6 | Asia / China | Ru Bai (Guangdong Provincial Center for Disease Control and Prevention) |
| EPI_ISL_340829 | A/Env/Guangdong/C17285757/QY/2017-11-21 | A / H5N6 | Asia / China | Ru Bai (Guangdong Provincial Center for Disease Control and Prevention) |
| EPI_ISL_340830 | A/Env/Guangdong/zhanjiang/C17277335/2017-11-27 | A / H5N6 | Asia / China | Ru Bai (Guangdong Provincial Center for Disease Control and Prevention) |
| EPI_ISL_340831 | A/Env/Guangdong/Shaoguan/C17272470/2018-01-16 | A / H5N6 | Asia / China | Ru Bai (Guangdong Provincial Center for Disease Control and Prevention) |
| EPI_ISL_340832 | A/Env/Guangdong/zhanjiang/C18277135/2018-04-02 | A / H5N6 | Asia / China | Ru Bai (Guangdong Provincial Center for Disease Control and Prevention) |
| EPI_ISL_340833 | A/Env/Guangdong/Foshan/C182750020/2018-01-02 | A / H5N6 | Asia / China | Ru Bai (Guangdong Provincial Center for Disease Control and Prevention) |
| EPI_ISL_340834 | A/Duck/Guangdong/PO1707260017/GZH/2017-08-02 | A / H5N6 | Asia / China | Ru Bai (Guangdong Provincial Center for Disease Control and Prevention) |
| EPI_ISL_340835 | A/Env/Guangdong/Zhaoqing/C172790995/2017-12-12 | A / H5N6 | Asia / China | Ru Bai (Guangdong Provincial Center for Disease Control and Prevention) |
| EPI_ISL_340838 | A/Env/Guangdong/C17059521/YF/2017-11-14 | A / H5N6 | Asia / China | Ru Bai (Guangdong Provincial Center for Disease Control and Prevention) |
| EPI_ISL_340839 | A/Env/Guangdong/EN17275405/FSH/2017-10-18 | A / H5N6 | Asia / China | Ru Bai (Guangdong Provincial Center for Disease Control and Prevention) |
| EPI_ISL_340844 | A/Env/Guangdong/C17285752/QY/2017-11-21 | A / H5N6 | Asia / China | Ru Bai (Guangdong Provincial Center for Disease Control and Prevention) |
| EPI_ISL_340845 | A/Env/Guangdong/C17285753/QY/2017-11-21 | A / H5N6 | Asia / China | Ru Bai (Guangdong Provincial Center for Disease Control and Prevention) |
| EPI_ISL_340846 | A/Env/Guangdong/zhongshan/C172870590/2017-11-28 | A / H5N6 | Asia / China | Ru Bai (Guangdong Provincial Center for Disease Control and Prevention) |
| EPI_ISL_340847 | A/Duck/Guangdong/Yangjiang/PO18284009/2018-01-09 | A / H5N6 | Asia / China | Ru Bai (Guangdong Provincial Center for Disease Control and Prevention) |
| EPI_ISL_340848 | A/Env/Guangdong/C172790591/ZHQ/2017-5-8 | A / H5N6 | Asia / China | Ru Bai (Guangdong Provincial Center for Disease Control and Prevention) |
| EPI_ISL_340849 | A/Env/Guangdong/C172811482/MZH/2017-10-16 | A / H5N6 | Asia / China | Ru Bai (Guangdong Provincial Center for Disease Control and Prevention) |
| EPI_ISL_340852 | A/Goose/Guangdong/PO1707260055/GZH/2017-08-02 | A / H5N6 | Asia / China | Ru Bai (Guangdong Provincial Center for Disease Control and Prevention) |
| EPI_ISL_340853 | A/Env/Guangdong/C17272335/SHG/2017-5-16 | A / H5N6 | Asia / China | Ru Bai (Guangdong Provincial Center for Disease Control and Prevention) |
| EPI_ISL_340854 | A/Env/Guangdong/C17285091/QY/2017-1-17 | A / H5N6 | Asia / China | Ru Bai (Guangdong Provincial Center for Disease Control and Prevention) |
| EPI_ISL_340855 | A/Waterfowl/Guangdong/PO17284158/YJ/2017-5-10 | A / H5N6 | Asia / China | Ru Bai (Guangdong Provincial Center for Disease Control and Prevention) |
| EPI_ISL_340856 | A/Env/Guangdong/EN17284259/YJ/2017-7-12 | A / H5N6 | Asia / China | Ru Bai (Guangdong Provincial Center for Disease Control and Prevention) |
| EPI_ISL_340857 | A/Duck/Guangdong/PO17284260/YJ/2017-7-12 | A / H5N6 | Asia / China | Ru Bai (Guangdong Provincial Center for Disease Control and Prevention) |
| EPI_ISL_340858 | A/Env/Guangdong/C17280709/HZH/2017-5-2 | A / H5N6 | Asia / China | Ru Bai (Guangdong Provincial Center for Disease Control and Prevention) |
| EPI_ISL_341293 | A/Hubei/29578/2016 x PR8 (CNIC-HB29578) | A / H5N6 | Asia / China / Hubei Province | Lei Yang (WHO Chinese National Influenza Center / Virology Institute, Chinese CDC) |
| EPI_ISL_341294 | A/Fujian-Sanyuan/21099/2017 x PR8 (CNIC-21099) | A / H5N6 | Asia / China / Fujian Province | Lei Yang (WHO Chinese National Influenza Center / Virology Institute, Chinese CDC) |
| EPI_ISL_344526 | A/Duck/Egypt/AR517/2017 | A / H5N8 | Africa / Egypt / Fayoum | Timm Harder (Friedrich-Loeffler-Institut) |
| EPI_ISL_344527 | A/Duck/Egypt/AR518/2017 | A / H5N8 | Africa / Egypt / Giza | Timm Harder (Friedrich-Loeffler-Institut) |
| EPI_ISL_344528 | A/Chicken/Egypt/AR520/2017 | A / H5N8 | Africa / Egypt / Qualiobia | Timm Harder (Friedrich-Loeffler-Institut) |
| EPI_ISL_344531 | A/Chicken/Egypt/AR539/2018 | A / H5N8 | Africa / Egypt / Mansoura | Timm Harder (Friedrich-Loeffler-Institut) |
| EPI_ISL_344532 | A/Chicken/Egypt/AR541/2018 | A / H5N8 | Africa / Egypt / Giza | Timm Harder (Friedrich-Loeffler-Institut) |
| EPI_ISL_344533 | A/Duck/Egypt/AR542/2018 | A / H5N8 | Africa / Egypt / Fayoum | Timm Harder (Friedrich-Loeffler-Institut) |
| EPI_ISL_344534 | A/Chicken/Egypt/AR543/2018 | A / H5N8 | Africa / Egypt / Fayoum | Timm Harder (Friedrich-Loeffler-Institut) |
| EPI_ISL_344537 | A/Chicken/Egypt/AR548/2018 | A / H5N8 | Africa / Egypt / Qualiobia | Timm Harder (Friedrich-Loeffler-Institut) |
| EPI_ISL_344538 | A/Chicken/Egypt/AR549/2018 | A / H5N8 | Africa / Egypt / Qualiobia | Timm Harder (Friedrich-Loeffler-Institut) |
| EPI_ISL_344539 | A/Turkey/Egypt/AR550/2018 | A / H5N8 | Africa / Egypt / Fayoum | Timm Harder (Friedrich-Loeffler-Institut) |
| EPI_ISL_344540 | A/Chicken/Egypt/AR552/2018 | A / H5N8 | Africa / Egypt / Beni Suef | Timm Harder (Friedrich-Loeffler-Institut) |
| EPI_ISL_344541 | A/Chicken/Egypt/AR553/2018 | A / H5N8 | Africa / Egypt / Giza | Timm Harder (Friedrich-Loeffler-Institut) |
| EPI_ISL_344542 | A/Chicken/Egypt/AR556/2018 | A / H5N8 | Africa / Egypt / Qualiobia | Timm Harder (Friedrich-Loeffler-Institut) |
| EPI_ISL_344543 | A/Chicken/Egypt/AR557/2018 | A / H5N8 | Africa / Egypt / Qualiobia | Timm Harder (Friedrich-Loeffler-Institut) |
| EPI_ISL_344544 | A/Chicken/Egypt/AR558/2018 | A / H5N8 | Africa / Egypt / Giza | Timm Harder (Friedrich-Loeffler-Institut) |
| EPI_ISL_344545 | A/Chicken/Egypt/AR559/2018 | A / H5N8 | Africa / Egypt / Fayoum | Timm Harder (Friedrich-Loeffler-Institut) |
| EPI_ISL_344546 | A/Duck/Egypt/AR560/2018 | A / H5N8 | Africa / Egypt / Giza | Timm Harder (Friedrich-Loeffler-Institut) |
| EPI_ISL_344547 | A/Chicken/Egypt/AR562/2018 | A / H5N8 | Africa / Egypt / Qualiobia | Timm Harder (Friedrich-Loeffler-Institut) |
| EPI_ISL_344548 | A/Chicken/Egypt/AR591/2018 | A / H5N8 | Africa / Egypt / Giza | Timm Harder (Friedrich-Loeffler-Institut) |
| EPI_ISL_346255 | A/duck/Egypt/VG1099/2018 | A / H5N2 | Africa / Egypt / Dakahlia | Mahmoud Mohamed Naguib (Friedrich-Loeffler-Institut) |
| EPI_ISL_347752 | A/Chicken/Yangzhou/f6/2018 | A / H5N6 | Asia / China / Jiangsu Provice | Xian Qi (Jiangsu Provincial Center for Disease Control & Prevention) |
| EPI_ISL_347754 | A/Chicken/Yangzhou/f9/2018 | A / H5N6 | Asia / China / Jiangsu Provice | Xian Qi (Jiangsu Provincial Center for Disease Control & Prevention) |
| EPI_ISL_347757 | A/Chicken/Yangzhou/f14/2018 | A / H5N6 | Asia / China / Jiangsu Provice | Xian Qi (Jiangsu Provincial Center for Disease Control & Prevention) |

| | | | | |
|---|---|---|---|---|
| EPI_ISL_347793 | A/Enviroment/Changzhou/768/2017 | A / H5N6 | Asia / China / Jiangsu Provice | Xian Qi (Jiangsu Provincial Center for Disease Control & Prevention) |
| EPI_ISL_348279 | A/duck/Nigeria/17RS737-43/2016 | A / H5N8 | Africa / Nigeria / Kano state | Bianca Zecchin (Istituto Zooprofilattico Sperimentale Delle Venezie) |
| EPI_ISL_348282 | A/chicken/Niger/17RS167-19/2017 | A / H5N8 | Africa / Niger / Niamey | Bianca Zecchin (Istituto Zooprofilattico Sperimentale Delle Venezie) |
| EPI_ISL_348283 | A/goose/Niger/17RS167-22/2017 | A / H5N8 | Africa / Niger / Niamey | Bianca Zecchin (Istituto Zooprofilattico Sperimentale Delle Venezie) |
| EPI_ISL_348284 | A/chicken/Niger/17RS167-1/2017 | A / H5N1 | Africa / Niger / Niamey | Bianca Zecchin (Istituto Zooprofilattico Sperimentale Delle Venezie) |
| EPI_ISL_348285 | A/chicken/Niger/17RS167-3/2017 | A / H5N1 | Africa / Niger / Niamey | Bianca Zecchin (Istituto Zooprofilattico Sperimentale Delle Venezie) |
| EPI_ISL_351792 | A/Chicken/Egypt/Al00994/2019 | A / H5N2 | Africa / Egypt / Beheira | Timm Harder (Friedrich-Loeffler-Institut) |
| EPI_ISL_353281 | A/Chicken/Suzhou/j1/2019 | A / H5N6 | Asia / China / Jiangsu Provice | Xian Qi (Jiangsu Provincial Center for Disease Control & Prevention) |
| EPI_ISL_353285 | A/Chicken/Suzhou/j5/2019 | A / H5N6 | Asia / China / Jiangsu Provice | Xian Qi (Jiangsu Provincial Center for Disease Control & Prevention) |
| EPI_ISL_353618 | A/Chicken/Suzhou/j6/2019 | A / H5N6 | Asia / China / Jiangsu Provice | Xian Qi (Jiangsu Provincial Center for Disease Control & Prevention) |
| EPI_ISL_354550 | A/Environment/Jiangxi/50653/2016 | A / H5N6 | Asia / China / Jiangxi | Xiaoxu Zeng (WHO Chinese National Influenza Center / Virology Institute, Chinese CDC) |
| EPI_ISL_354551 | A/Environment/Jiangxi/50652/2016 | A / H5N6 | Asia / China / Jiangxi | Xiaoxu Zeng (WHO Chinese National Influenza Center / Virology Institute, Chinese CDC) |
| EPI_ISL_354552 | A/Environment/Jiangxi/50624/2016 | A / H5N6 | Asia / China / Jiangxi | Xiaoxu Zeng (WHO Chinese National Influenza Center / Virology Institute, Chinese CDC) |
| EPI_ISL_354553 | A/Duck/Jiangxi/50455/2016 | A / H5N6 | Asia / China / Jiangxi | Xiaoxu Zeng (WHO Chinese National Influenza Center / Virology Institute, Chinese CDC) |
| EPI_ISL_354554 | A/Environment/Jiangxi/50452/2016 | A / H5N6 | Asia / China / Jiangxi | Xiaoxu Zeng (WHO Chinese National Influenza Center / Virology Institute, Chinese CDC) |
| EPI_ISL_354555 | A/Environment/Jiangxi/50449/2016 | A / H5N6 | Asia / China / Jiangxi | Xiaoxu Zeng (WHO Chinese National Influenza Center / Virology Institute, Chinese CDC) |
| EPI_ISL_354556 | A/Duck/Jiangxi/50446/2016 | A / H5N6 | Asia / China / Jiangxi | Xiaoxu Zeng (WHO Chinese National Influenza Center / Virology Institute, Chinese CDC) |
| EPI_ISL_354557 | A/Environment/Jiangxi/50445/2016 | A / H5N6 | Asia / China / Jiangxi | Xiaoxu Zeng (WHO Chinese National Influenza Center / Virology Institute, Chinese CDC) |
| EPI_ISL_354559 | A/Environment/Jiangxi/45191/2016 | A / H5N6 | Asia / China / Jiangxi | Xiaoxu Zeng (WHO Chinese National Influenza Center / Virology Institute, Chinese CDC) |
| EPI_ISL_354560 | A/Chicken/Jiangxi/45153/2016 | A / H5N6 | Asia / China / Jiangxi | Xiaoxu Zeng (WHO Chinese National Influenza Center / Virology Institute, Chinese CDC) |
| EPI_ISL_354561 | A/Environment/Jiangxi/45148/2015 | A / H5N6 | Asia / China / Jiangxi | Xiaoxu Zeng (WHO Chinese National Influenza Center / Virology Institute, Chinese CDC) |
| EPI_ISL_354562 | A/Chicken/Jiangxi/45093/2016 | A / H5N6 | Asia / China / Jiangxi | Xiaoxu Zeng (WHO Chinese National Influenza Center / Virology Institute, Chinese CDC) |
| EPI_ISL_354563 | A/Environment/Jiangxi/45084/2016 | A / H5N6 | Asia / China / Jiangxi | Xiaoxu Zeng (WHO Chinese National Influenza Center / Virology Institute, Chinese CDC) |
| EPI_ISL_354564 | A/Environment/Jiangxi/44908/2015 | A / H5N6 | Asia / China / Jiangxi | Xiaoxu Zeng (WHO Chinese National Influenza Center / Virology Institute, Chinese CDC) |
| EPI_ISL_354565 | A/Environment/Jiangxi/46456/2015 | A / H5N6 | Asia / China / Jiangxi | Xiaoxu Zeng (WHO Chinese National Influenza Center / Virology Institute, Chinese CDC) |
| EPI_ISL_354566 | A/Environment/Jiangxi/46390/2015 | A / H5N6 | Asia / China / Jiangxi | Xiaoxu Zeng (WHO Chinese National Influenza Center / Virology Institute, Chinese CDC) |
| EPI_ISL_354567 | A/Environment/Jiangxi/25004/2014 | A / H5N6 | Asia / China / Jiangxi | Xiaoxu Zeng (WHO Chinese National Influenza Center / Virology Institute, Chinese CDC) |
| EPI_ISL_354571 | A/Environment/Jiangxi/24995/2014 | A / H5N6 | Asia / China / Jiangxi | Xiaoxu Zeng (WHO Chinese National Influenza Center / Virology Institute, Chinese CDC) |
| EPI_ISL_362127 | A/Grey seal/361-13/BalticPL/16 | A / H5N8 | Europe / Poland / Pomeranian Voivodeship | Dai-Lun Shin (Stiftung Tierärztliche Hochschule Hannover) |
| EPI_ISL_365299 | A/chicken/South Africa/17090108/2017 | A / H5N8 | Africa / South Africa | |
| EPI_ISL_365300 | A/muscovy duck/Vietnam/LBM801c132/2015 | A / H5N6 | Asia / Vietnam | |
| EPI_ISL_366233 | A/Cygnus olor/England/AS00778/2017 | A / H5N8 | Europe / United Kingdom | |
| EPI_ISL_366235 | A/Cygnus olor/England/AS00868/2017 | A / H5N8 | Europe / United Kingdom | |
| EPI_ISL_366236 | A/Cygnus olor/England/AS00918/2016 | A / H5N8 | Europe / United Kingdom | |
| EPI_ISL_366237 | A/Cygnus olor/England/WVUK/2016 | A / H5N8 | Europe / United Kingdom | |
| EPI_ISL_366238 | A/Cygnus olor/England/WVJX/2016 | A / H5N8 | Europe / United Kingdom | |
| EPI_ISL_366239 | A/Cygnus olor/England/YBOV/2017 | A / H5N8 | Europe / United Kingdom | |
| EPI_ISL_366240 | A/Cygnus olor/England/WVZP/2016 | A / H5N8 | Europe / United Kingdom | |
| EPI_ISL_366241 | A/Cygnus olor/England/WULH/2016 | A / H5N8 | Europe / United Kingdom | |
| EPI_ISL_366244 | A/Cygnus olor/England/UnringedBirdB/2017 | A / H5N8 | Europe / United Kingdom | |
| EPI_ISL_366612 | A/Poultry/Iran/clade 2344/2018 | A / H5N8 | Asia / Iran, Islamic Republic of | |
| EPI_ISL_366635 | A/chicken/Alkharj/910/2018 | A / H5N8 | Asia / Saudi Arabia | |
| EPI_ISL_368312 | A/chicken/Nha Trang/185/2017 | A / H5N6 | Asia / Vietnam | |
| EPI_ISL_368313 | A/muscovy duck/Vietnam/LBM941/2017 | A / H5N6 | Asia / Vietnam | |
| EPI_ISL_368368 | A/China/Original/2018 | A / H5N6 | Asia / China | |
| EPI_ISL_368623 | A/chicken/Egypt/AB1/2018 | A / H5N8 | Africa / Egypt | |
| EPI_ISL_368668 | A/duck/Viet Nam/HU1-2361/2014 | A / H5N6 | Asia / Vietnam | |
| EPI_ISL_369336 | A/chicken/South Africa/Villiers/2017 | A / H5N8 | Africa / South Africa | |
| EPI_ISL_369337 | A/chicken/South Africa/Standerton/2017 | A / H5N8 | Africa / South Africa | |
| EPI_ISL_369338 | A/chicken/South Africa/436893/2017 | A / H5N8 | Africa / South Africa | |
| EPI_ISL_369339 | A/chicken/South Africa/440638A/2017 | A / H5N8 | Africa / South Africa | |
| EPI_ISL_369340 | A/chicken/South Africa/440638B/2017 | A / H5N8 | Africa / South Africa | |
| EPI_ISL_369341 | A/chicken/South Africa/441587/2017 | A / H5N8 | Africa / South Africa | |
| EPI_ISL_369342 | A/chicken/South Africa/MC002/2017 | A / H5N8 | Africa / South Africa | |
| EPI_ISL_369343 | A/chicken/South Africa/441839/2017 | A / H5N8 | Africa / South Africa | |
| EPI_ISL_369344 | A/ostrich/South Africa/17080046/2017 | A / H5N8 | Africa / South Africa | |
| EPI_ISL_369345 | A/chicken/South Africa/443397/2017 | A / H5N8 | Africa / South Africa | |
| EPI_ISL_369346 | A/chicken/South Africa/17080336/2017 | A / H5N8 | Africa / South Africa | |
| EPI_ISL_369347 | A/Pekin duck/South Africa/17080340/2017 | A / H5N8 | Africa / South Africa | |
| EPI_ISL_369348 | A/Pekin duck/South Africa/17080481/2017 | A / H5N8 | Africa / South Africa | |
| EPI_ISL_369349 | A/Swan/South Africa/17080517/2017 | A / H5N8 | Africa / South Africa | |
| EPI_ISL_369350 | A/chicken/South Africa/17080561/2017 | A / H5N8 | Africa / South Africa | |
| EPI_ISL_369351 | A/chicken/South Africa/17080581/2017 | A / H5N8 | Africa / South Africa | |
| EPI_ISL_369352 | A/chicken/South Africa/17090050/2017 | A / H5N8 | Africa / South Africa | |
| EPI_ISL_369353 | A/chicken/South Africa/17090100/2017 | A / H5N8 | Africa / South Africa | |
| EPI_ISL_369354 | A/chicken/South Africa/17090202/2017 | A / H5N8 | Africa / South Africa | |
| EPI_ISL_369355 | A/chicken/South Africa/448475/2017 | A / H5N8 | Africa / South Africa | |
| EPI_ISL_369356 | A/chicken/South Africa/17090325/2017 | A / H5N8 | Africa / South Africa | |
| EPI_ISL_369357 | A/chicken/South Africa/449300/2017 | A / H5N8 | Africa / South Africa | |
| EPI_ISL_369358 | A/chicken/South Africa/17090348/2017 | A / H5N8 | Africa / South Africa | |
| EPI_ISL_369359 | A/chicken/South Africa/449418/2017 | A / H5N8 | Africa / South Africa | |
| EPI_ISL_369360 | A/chicken/South Africa/449443/2017 | A / H5N8 | Africa / South Africa | |

| | | | | |
|---|---|---|---|---|
| EPI_ISL_369361 | A/chicken/South Africa/17090335/2017 | A / H5N8 | Africa / South Africa | |
| EPI_ISL_369362 | A/turkey/South Africa/450199/2017 | A / H5N8 | Africa / South Africa | |
| EPI_ISL_369363 | A/chicken/South Africa/115370/2017 | A / H5N8 | Africa / South Africa | |
| EPI_ISL_369364 | A/chicken/South Africa/450628/2017 | A / H5N8 | Africa / South Africa | |
| EPI_ISL_369365 | A/chicken/South Africa/451457/2017 | A / H5N8 | Africa / South Africa | |
| EPI_ISL_369366 | A/ostrich/South Africa/002/2017 | A / H5N8 | Africa / South Africa | |
| EPI_ISL_370414 | A/Pavo cristatus/China/JS02/2018 | A / H5N6 | Asia / China | |
| EPI_ISL_370415 | A/Pavo cristatus/China/JS01/2018 | A / H5N6 | Asia / China | |
| EPI_ISL_370416 | A/Pavo cristatus/China/K10/2018 | A / H5N6 | Asia / China | |
| EPI_ISL_372404 | A/goose/Guangdong/A-Goose-Guangdong-GS013-2015-HA/20 | A / H5N6 | Asia / China | |
| EPI_ISL_372409 | A/goose/Guangdong/A-Goose-Guangdong-GS014-2015-HA/20 | A / H5N6 | Asia / China | |
| EPI_ISL_372430 | A/goose/Guangdong/A-Goose-Guangdong-GS017-2015(H5N6) | A / H5N6 | Asia / China | |
| EPI_ISL_372438 | A/goose/Guangdong/A-Goose-Guangdong-GS114-2015(H5N6) | A / H5N6 | Asia / China | |
| EPI_ISL_372446 | A/goose/Guangdong/A-Goose-Guangdong-GS116-2015(H5N6) | A / H5N6 | Asia / China | |
| EPI_ISL_372454 | A/goose/Guangdong/A-Goose-Guangdong-GS120-2015(H5N6) | A / H5N6 | Asia / China | |
| EPI_ISL_372462 | A/goose/Guangdong/A-Goose-Guangdong-GS119-2015(H5N6) | A / H5N6 | Asia / China | |
| EPI_ISL_372467 | A/goose/Guangdong/A-Goose-Guangdong-GS144-2015(H5N6) | A / H5N6 | Asia / China | |
| EPI_ISL_372475 | A/goose/Guangdong/A-Goose-Guangdong-GS148-2016(H5N6) | A / H5N6 | Asia / China | |
| EPI_ISL_372486 | A/goose/Guangdong/A-Goose-Guangdong-GS018-2015(H5N6) | A / H5N6 | Asia / China | |
| EPI_ISL_372491 | A/chicken/South Korea/AIV49/2017 | A / H5N6 | Asia / Korea, Republic of | |
| EPI_ISL_372791 | A/goose/China/GS25-HA.seq/2016 | A / H5N6 | Asia / China | |
| EPI_ISL_372799 | A/goose/China/GS24-HA.seq/2016 | A / H5N6 | Asia / China | |
| EPI_ISL_372806 | A/goose/China/GS37-4.seq/2016 | A / H5N6 | Asia / China | |
| EPI_ISL_372815 | A/goose/China/GS42-4.seq/2016 | A / H5N6 | Asia / China | |
| EPI_ISL_372823 | A/goose/China/GS45-4.seq/2016 | A / H5N6 | Asia / China | |
| EPI_ISL_372831 | A/chicken/China/CK46-4.seq/2016 | A / H5N6 | Asia / China | |
| EPI_ISL_372840 | A/goose/China/GS74-4.seq/2016 | A / H5N6 | Asia / China | |
| EPI_ISL_372962 | A/mallard/Aichi/2312T010/2016 | A / H5N6 | Asia / Japan / Aichi | |
| EPI_ISL_372965 | A/mallard/Aichi/2312T011/2016 | A / H5N6 | Asia / Japan / Aichi | |
| EPI_ISL_372968 | A/wigeon/Aichi/2312T012/2016 | A / H5N6 | Asia / Japan / Aichi | |
| EPI_ISL_372971 | A/mallard/Aichi/2312T018/2016 | A / H5N6 | Asia / Japan / Aichi | |
| EPI_ISL_376189 | A/duck/China/waterfowl/2016 | A / H5N2 | Asia / China | |
| EPI_ISL_3770721 | A/GX-hechi/01/2021 | A / H5N6 | Asia / China / Guangxi Zhuang Autonomous Region | Min Chen (Guangxi Center for Disease Prevention and Control) |
| EPI_ISL_3770740 | A/GX-guilin/11151/2021 | A / H5N6 | Asia / China / Guangxi Zhuang Autonomous Region | Min Chen (Guangxi Center for Disease Prevention and Control) |
| EPI_ISL_378234 | A/Domestic goose/Sulaimani/Sul.1/2018 | A / H5N8 | Asia / Iraq | |
| EPI_ISL_378252 | A/duck/Vietnam/LBM805c113/2015 | A / H5N6 | Asia / Vietnam | |
| EPI_ISL_378253 | A/duck/Nha Trang/194/2017 | A / H5N6 | Asia / Vietnam | |
| EPI_ISL_378254 | A/duck/Nha Trang/221/2017 | A / H5N6 | Asia / Vietnam | |
| EPI_ISL_378256 | A/Mulard Duck/Hungary/59163/2016 | A / H5N8 | Europe / Hungary | |
| EPI_ISL_378280 | A/cooper's hawk/Washington/1298/2015 | A / H5N2 | North America / United States / Washington | |
| EPI_ISL_378311 | A/pigeon/South Africa/17080323/2017 | A / H5N8 | Africa / South Africa | |
| EPI_ISL_378312 | A/Domestic goose/South Africa/17090065/2017 | A / H5N8 | Africa / South Africa | |
| EPI_ISL_378362 | A/whooper swan/Shanxi/RC01/2016 | A / H5N8 | Asia / China | |
| EPI_ISL_378363 | A/cackling goose/Aichi/2312T008/2016 | A / H5N6 | Asia / Japan / Aichi | |
| EPI_ISL_378364 | A/cackling goose/Aichi/2312T017/2016 | A / H5N6 | Asia / Japan / Aichi | |
| EPI_ISL_378365 | A/cackling goose/Aichi/2312T019/2016 | A / H5N6 | Asia / Japan / Aichi | |
| EPI_ISL_378634 | A/Whooper Swan/Sanmenxia/01/2016 | A / H5N8 | Asia / China | |
| EPI_ISL_379090 | A/peafowl/Cameroon/17RS1661-6/2017 | A / H5N8 | Africa / Cameroon | |
| EPI_ISL_379227 | A/chicken/Uganda/17RS115-15/2017 | A / H5N8 | Africa / Uganda | |
| EPI_ISL_379228 | A/duck/Uganda/17RS115-9/2017 | A / H5N8 | Africa / Uganda | |
| EPI_ISL_379568 | A/muscovy duck/Vietnam/HU4-160/2015 | A / H5N6 | Asia / Vietnam | |
| EPI_ISL_379682 | A/goose/China/1106/2016 | A / H5N2 | Asia / China | |
| EPI_ISL_379683 | A/duck/China/YD1516/2016 | A / H5N2 | Asia / China | |
| EPI_ISL_379744 | A/chicken/Egypt/FL6/2018 | A / H5N8 | Africa / Egypt | |
| EPI_ISL_379963 | A/duck/Vietnam/HU4-879/2015 | A / H5N6 | Asia / Vietnam | |
| EPI_ISL_380991 | A/Crow/Aghakhan/2017 | A / H5N8 | Asia / Iran, Islamic Republic of | |
| EPI_ISL_381741 | A/CHICKEN/Egypt/CA35/2017 | A / H5N8 | Africa / Egypt | |
| EPI_ISL_381742 | A/Duck/Egypt/CAG26/2017 | A / H5N8 | Africa / Egypt | |
| EPI_ISL_381743 | A/Duck/Egypt/SM57/2017 | A / H5N8 | Africa / Egypt | |
| EPI_ISL_381810 | A/Egyptian goose/South Africa/001/2017 | A / H5N8 | Africa / South Africa | |
| EPI_ISL_381811 | A/Speckled pigeon/South Africa/08-004B/2017 | A / H5N8 | Africa / South Africa | |
| EPI_ISL_381812 | A/Sacred ibis/South Africa/009/2017 | A / H5N8 | Africa / South Africa | |
| EPI_ISL_381813 | A/Guinea fowl/South Africa/17080243/2017 | A / H5N8 | Africa / South Africa | |
| EPI_ISL_381814 | A/Guinea fowl/South Africa/17080274/2017 | A / H5N8 | Africa / South Africa | |
| EPI_ISL_381815 | A/dove/South Africa/17080324/2017 | A / H5N8 | Africa / South Africa | |
| EPI_ISL_386922 | A/Luscinia cyane/Jiangxi/U2/2016 ?H5N6? | A / H5N6 | Asia / China / Jiangxi Province | Zhang Tao (Tsinghua University) |
| EPI_ISL_386923 | A/Streptopelia decaocto/Jiangxi/E1/2015(H5N6) | A / H5N6 | Asia / China / Jiangxi Province | Zhang Tao (Tsinghua University) |
| EPI_ISL_386924 | A/Streptopelia decaocto/Jiangxi/J8/2015 ?H5N6? | A / H5N6 | Asia / China / Jiangxi Province | Zhang Tao (Tsinghua University) |
| EPI_ISL_386925 | A/quail/Jiangxi/B9/2015 ?H5N6? | A / H5N6 | Asia / China / Jiangxi Province | Zhang Tao (Tsinghua University) |
| EPI_ISL_386926 | A/Mallard/Jiangxi/JXH22/2014 ?H5N6? | A / H5N6 | Asia / China / Jiangxi Province | Zhang Tao (Tsinghua University) |
| EPI_ISL_386927 | A/Mallard/Jiangxi/JXH9/2014 ?H5N6? | A / H5N6 | Asia / China / Jiangxi Province | Zhang Tao (Tsinghua University) |
| EPI_ISL_386928 | A/duck/Jiangxi/E7/2014 ?H5N6? | A / H5N6 | Asia / China / Jiangxi Province | Zhang Tao (Tsinghua University) |

| | | | | |
|---|---|---|---|---|
| EPI_ISL_386929 | A/chickenl/Jiangxi/E84/2014 ?H5N6? | A / H5N6 | Asia / China / Jiangxi Province | Zhang Tao (Tsinghua University) |
| EPI_ISL_386930 | A/duck/Jiangxi/F31/2014?H5N6? | A / H5N6 | Asia / China / Jiangxi Province | Zhang Tao (Tsinghua University) |
| EPI_ISL_387133 | A/chicken/Zimbabwe/AI4935/2017 | A / H5N8 | Africa / Zimbabwe | |
| EPI_ISL_387135 | A/duck/Egypt/N13736E/2017 | A / H5N8 | Africa / Egypt | |
| EPI_ISL_387136 | A/teal/Egypt/1198C/2017 | A / H5N8 | Africa / Egypt | |
| EPI_ISL_387137 | A/chicken/Egypt/Q13804A/2017 | A / H5N8 | Africa / Egypt | |
| EPI_ISL_387138 | A/teal/Egypt /823C/2016 | A / H5N8 | Africa / Egypt | |
| EPI_ISL_387139 | A/duck/Egypt/F13666A/2017 | A / H5N8 | Africa / Egypt | |
| EPI_ISL_387140 | A/teal/Egypt/1202C/2017 | A / H5N8 | Africa / Egypt | |
| EPI_ISL_387141 | A/northern shoveler/Egypt/813C/2016 | A / H5N8 | Africa / Egypt | |
| EPI_ISL_387143 | A/goose/Spain/IA17CR02699/2017 | A / H5N8 | Europe / Spain | |
| EPI_ISL_387351 | A/duck/Viet Nam/HN-2431/2015 | A / H5N1 | Asia / Vietnam | |
| EPI_ISL_387352 | A/duck/Viet Nam/HN-2512/2015 | A / H5N6 | Asia / Vietnam | |
| EPI_ISL_387353 | A/duck/Viet Nam/HN-2520/2015 | A / H5N6 | Asia / Vietnam | |
| EPI_ISL_387354 | A/duck/Viet Nam/HN-2522/2015 | A / H5N6 | Asia / Vietnam | |
| EPI_ISL_387362 | A/duck/Viet Nam/QN-2624/2016 | A / H5N6 | Asia / Vietnam | |
| EPI_ISL_388773 | A/little grebe/Egypt/1056OP/2016 | A / H5N8 | Africa / Egypt | |
| EPI_ISL_389022 | A/chicken/Nghe An/01VTC/2018 | A / H5N6 | Asia / Vietnam / Tinh Nghe An | Natalia Goncharova (State Research Center of Virology and Biotechnology (VECTOR) / Emerging Zoonotic Diseases and Influenza) |
| EPI_ISL_389023 | A/chicken/Nghe An/02VTC/2018 | A / H5N6 | Asia / Vietnam / Tinh Nghe An | Natalia Goncharova (State Research Center of Virology and Biotechnology (VECTOR) / Emerging Zoonotic Diseases and Influenza) |
| EPI_ISL_389082 | A/chicken/Nghe An/14VTC/2015 | A / H5N6 | Asia / Vietnam / Tinh Nghe An | Natalia Goncharova (State Research Center of Virology and Biotechnology (VECTOR) / Emerging Zoonotic Diseases and Influenza) |
| EPI_ISL_389145 | A/chicken/Nghe An/27VTC/2018 | A / H5N6 | Asia / Vietnam / Tinh Nghe An | Natalia Goncharova (State Research Center of Virology and Biotechnology (VECTOR) / Emerging Zoonotic Diseases and Influenza) |
| EPI_ISL_389290 | A/chicken/Ha Tinh/73VTC/2017 | A / H5N6 | Asia / Vietnam / Tinh Ha Tinh | Natalia Goncharova (State Research Center of Virology and Biotechnology (VECTOR) / Emerging Zoonotic Diseases and Influenza) |
| EPI_ISL_389657 | A/chicken/Vietnam/HU9-842/2018 | A / H5N6 | Asia / Vietnam | |
| EPI_ISL_389659 | A/chicken/Vietnam/HU9-847/2018 | A / H5N6 | Asia / Vietnam | |
| EPI_ISL_389737 | A/chicken/Egypt/H13795A/2017 | A / H5N8 | Africa / Egypt | |
| EPI_ISL_389738 | A/chicken/Egypt/H13791C/2017 | A / H5N8 | Africa / Egypt | |
| EPI_ISL_389739 | A/chicken/Egypt/N13726/2017 | A / H5N8 | Africa / Egypt | |
| EPI_ISL_389740 | A/chicken/ Egypt/N13732A/2017 | A / H5N8 | Africa / Egypt | |
| EPI_ISL_389743 | A/chicken/Egypt/H13793E/2017 | A / H5N8 | Africa / Egypt | |
| EPI_ISL_389746 | A/duck/Egypt/N13733B/2017 | A / H5N8 | Africa / Egypt | |
| EPI_ISL_389747 | A/chicken/Egypt/N13717E/2017 | A / H5N8 | Africa / Egypt | |
| EPI_ISL_389748 | A/chicken/Egypt/A13775D/2017 | A / H5N8 | Africa / Egypt | |
| EPI_ISL_389750 | A/duck/Egypt/N13734B/2017 | A / H5N8 | Africa / Egypt | |
| EPI_ISL_389751 | A/chicken/Egypt/M14081D/2017 | A / H5N8 | Africa / Egypt | |
| EPI_ISL_389752 | A/chicken/Egypt/M13844B/2017 | A / H5N8 | Africa / Egypt | |
| EPI_ISL_389753 | A/duck/Egypt/H13797E/2017 | A / H5N8 | Africa / Egypt | |
| EPI_ISL_389754 | A/duck/Egypt/N13736B/2017 | A / H5N8 | Africa / Egypt | |
| EPI_ISL_389755 | A/chicken/Egypt/Q13944C/2017 | A / H5N8 | Africa / Egypt | |
| EPI_ISL_389756 | A/chicken/Egypt/Q13941B/2017 | A / H5N8 | Africa / Egypt | |
| EPI_ISL_389757 | A/duck/Egypt/N14200B/2017 | A / H5N8 | Africa / Egypt | |
| EPI_ISL_389759 | A/chicken/Egypt/F13660A/2017 | A / H5N8 | Africa / Egypt | |
| EPI_ISL_389760 | A/chicken/Egypt/F13830A/2017 | A / H5N8 | Africa / Egypt | |
| EPI_ISL_389761 | A/chicken/Egypt/N13720A/2017 | A / H5N8 | Africa / Egypt | |
| EPI_ISL_389762 | A/chicken/Egypt/Q13845B/2017 | A / H5N8 | Africa / Egypt | |
| EPI_ISL_389765 | A/chicken/Egypt/F13664A/2017 | A / H5N8 | Africa / Egypt | |
| EPI_ISL_389766 | A/chicken/Egypt/A13777B/2017 | A / H5N8 | Africa / Egypt | |
| EPI_ISL_389767 | A/chicken/Egypt/H13794E/2017 | A / H5N8 | Africa / Egypt | |
| EPI_ISL_389768 | A/chicken/Egypt/H13800C/2017 | A / H5N8 | Africa / Egypt | |
| EPI_ISL_389769 | A/chicken/Egypt/H13796A/2017 | A / H5N8 | Africa / Egypt | |
| EPI_ISL_389770 | A/chicken/Egypt/F13829A/2017 | A / H5N8 | Africa / Egypt | |
| EPI_ISL_389771 | A/chicken/Egypt/A13776B/2017 | A / H5N8 | Africa / Egypt | |
| EPI_ISL_389772 | A/chicken/Egypt/H13799B/2017 | A / H5N8 | Africa / Egypt | |
| EPI_ISL_389773 | A/chicken/Egypt/Q13936B/2017 | A / H5N8 | Africa / Egypt | |
| EPI_ISL_389774 | A/goose/Egypt/A13779B/2017 | A / H5N8 | Africa / Egypt | |
| EPI_ISL_389776 | A/duck/Egypt/N13735A/2017 | A / H5N8 | Africa / Egypt | |
| EPI_ISL_389791 | A/chicken/Bangladesh/1896-17-HA/2017 | A / H5N6 | Asia / Bangladesh | |
| EPI_ISL_393518 | A/Avian/Iran/38AMA/2019 | A / H5N3 | Asia / Iran, Islamic Republic of | |
| EPI_ISL_3950049 | A/Mallard/Sweden/SVA210827SZ0252/KN002973/D-2021 | A / H5N8 | Europe / Sweden / Sodermanlands Lan / Nykopings Kommun | Siamak Zohari (National Veterinary Institute) |
| EPI_ISL_3950066 | A/Mallard/Sweden/SVA210827SZ0252/KN002974/D-2021 | A / H5N8 | Europe / Sweden / Sodermanlands Lan / Nykopings Kommun | Siamak Zohari (National Veterinary Institute) |
| EPI_ISL_397346 | A/duck/Tottori/E9/2016 | A / H5N6 | Asia / Japan / Tottori | |
| EPI_ISL_397347 | A/duck/Tottori/E10/2016 | A / H5N6 | Asia / Japan / Tottori | |
| EPI_ISL_397348 | A/duck/Tottori/E13/2016 | A / H5N6 | Asia / Japan / Tottori | |
| EPI_ISL_398050 | A/chicken-layer/Egypt/MR18/2018 | A / H5 | Africa / Egypt | |
| EPI_ISL_399051 | A/duck/Egypt/F13663C/2017 | A / H5N8 | Africa / Egypt | |
| EPI_ISL_399055 | A/duck/Egypt/F13667A/2017 | A / H5N8 | Africa / Egypt | |
| EPI_ISL_399989 | A/Duck/Egypt/AI20251/2019 | A / H5N8 | Africa / Egypt | Timm Harder (Friedrich-Loeffler-Institut) |
| EPI_ISL_399990 | A/Duck/Egypt/AI20252/2019 | A / H5N8 | Africa / Egypt | Timm Harder (Friedrich-Loeffler-Institut) |
| EPI_ISL_399991 | A/Duck/Egypt/AI20259/2019 | A / H5 | Africa / Egypt | Timm Harder (Friedrich-Loeffler-Institut) |
| EPI_ISL_399993 | A/Duck/Egypt/AI20263/2019 | A / H5N8 | Africa / Egypt | Timm Harder (Friedrich-Loeffler-Institut) |
| EPI_ISL_399994 | A/Duck/Egypt/AI20266/2019 | A / H5N8 | Africa / Egypt | Timm Harder (Friedrich-Loeffler-Institut) |
| EPI_ISL_399995 | A/Chicken/Egypt/AI20268/2019 | A / H5N8 | Africa / Egypt | Timm Harder (Friedrich-Loeffler-Institut) |
| EPI_ISL_399996 | A/Chicken/Egypt/AI20284/2019 | A / H5N8 | Africa / Egypt | Timm Harder (Friedrich-Loeffler-Institut) |

| | | | | |
|---|---|---|---|---|
| EPI_ISL_400027 | A/Turkey/Egypt/AI20285/2019 | A / H5N8 | Africa / Egypt | Timm Harder (Friedrich-Loeffler-Institut) |
| EPI_ISL_400043 | A/Chicken/Egypt/AI20286/2019 | A / H5N8 | Africa / Egypt | Timm Harder (Friedrich-Loeffler-Institut) |
| EPI_ISL_400485 | A/Green pheasant/Hunan/10/2015 | A / H5N6 | Asia / China / Hunan | Yanbing Li (Harbin Veterinary Research Institute (CAAS) / Ministry of Agriculture) |
| EPI_ISL_400486 | A/Common pheasant/Hunan/11/2015 | A / H5N6 | Asia / China / Hunan | Yanbing Li (Harbin Veterinary Research Institute (CAAS) / Ministry of Agriculture) |
| EPI_ISL_400487 | A/Common pheasant/Hunan/2/2016 | A / H5N6 | Asia / China / Hunan | Yanbing Li (Harbin Veterinary Research Institute (CAAS) / Ministry of Agriculture) |
| EPI_ISL_400488 | A/Silver Pheasant/Hunan/3/2016 | A / H5N6 | Asia / China / Hunan | Yanbing Li (Harbin Veterinary Research Institute (CAAS) / Ministry of Agriculture) |
| EPI_ISL_400489 | A/Whooper Swan/Hunan/4/2016 | A / H5N6 | Asia / China / Hunan | Yanbing Li (Harbin Veterinary Research Institute (CAAS) / Ministry of Agriculture) |
| EPI_ISL_400490 | A/Whooper swan/Shanxi/6/2016 | A / H5N8 | Asia / China / Shanxi | Yanbing Li (Harbin Veterinary Research Institute (CAAS) / Ministry of Agriculture) |
| EPI_ISL_400491 | A/Whooper swan/Shanxi/7/2016 | A / H5N8 | Asia / China / Shanxi | Yanbing Li (Harbin Veterinary Research Institute (CAAS) / Ministry of Agriculture) |
| EPI_ISL_400492 | A/Greylag goose/Hunan/1/2017 | A / H5N6 | Asia / China / Hunan | Yanbing Li (Harbin Veterinary Research Institute (CAAS) / Ministry of Agriculture) |
| EPI_ISL_400493 | A/Black swan/Hubei/2/2017 | A / H5N8 | Asia / China / Hubei | Yanbing Li (Harbin Veterinary Research Institute (CAAS) / Ministry of Agriculture) |
| EPI_ISL_400494 | A/Black swan/Hubei/3/2017 | A / H5N8 | Asia / China / Hubei | Yanbing Li (Harbin Veterinary Research Institute (CAAS) / Ministry of Agriculture) |
| EPI_ISL_400548 | A/Peacock/Hunan/12/2015 | A / H5N6 | Asia / China / Hunan | Yanbing Li (Harbin Veterinary Research Institute (CAAS) / Ministry of Agriculture) |
| EPI_ISL_400549 | A/Peacock/Hunan/13/2015 | A / H5N6 | Asia / China / Hunan | Yanbing Li (Harbin Veterinary Research Institute (CAAS) / Ministry of Agriculture) |
| EPI_ISL_400550 | A/Peacock/Hunan/15/2015 | A / H5N6 | Asia / China / Hunan | Yanbing Li (Harbin Veterinary Research Institute (CAAS) / Ministry of Agriculture) |
| EPI_ISL_400551 | A/Peacock/Hunan/14/2015 | A / H5N2 | Asia / China / Hunan | Yanbing Li (Harbin Veterinary Research Institute (CAAS) / Ministry of Agriculture) |
| EPI_ISL_400552 | A/Peacock/Hunan/1/2016 | A / H5N6 | Asia / China / Hunan | Yanbing Li (Harbin Veterinary Research Institute (CAAS) / Ministry of Agriculture) |
| EPI_ISL_402134 | A/turkey/Poland/23/2019 | A / H5N8 | Europe / Poland / Lublin Voivodeship | Edyta ?wi?to? (National Veterinary Research Institut Poland, PIWet-PIB) |
| EPI_ISL_4031377 | A/Duck/China/B1/2018 | A / H5N6 | Asia / China | |
| EPI_ISL_4031378 | A/Duck/China/B2/2018 | A / H5N6 | Asia / China | |
| EPI_ISL_4031379 | A/Duck/China/B3/2019 | A / H5N6 | Asia / China | |
| EPI_ISL_4031380 | A/Duck/China/B4/2019 | A / H5N6 | Asia / China | |
| EPI_ISL_4031381 | A/Duck/China/B5/2019 | A / H5N6 | Asia / China | |
| EPI_ISL_4031382 | A/Duck/China/B6/2019 | A / H5N6 | Asia / China | |
| EPI_ISL_4031383 | A/Duck/China/B7/2019 | A / H5N6 | Asia / China | |
| EPI_ISL_4031384 | A/Duck/China/B8/2019 | A / H5N6 | Asia / China | |
| EPI_ISL_4031385 | A/Duck/China/B9/2019 | A / H5N6 | Asia / China | |
| EPI_ISL_4031450 | A/Chicken/Viet Nam/AI-1606/2016 | A / H5N6 | Asia / Vietnam | |
| EPI_ISL_4031458 | A/mallard/Denmark/12106-1/2018 | A / H5N6 | Europe / Denmark | |
| EPI_ISL_4031459 | A/mallard/Denmark/12106-2/2018 | A / H5N6 | Europe / Denmark | |
| EPI_ISL_4031461 | A/black-headed gull/Denmark/4663-1/2018 | A / H5N6 | Europe / Denmark | |
| EPI_ISL_4031467 | A/pheasant/Denmark/12106-03/2018 | A / H5N6 | Europe / Denmark | |
| EPI_ISL_4032096 | A/duck/Nantou/15A03445-11-20T/2015 | A / H5N8 | Asia / Taiwan | |
| EPI_ISL_4032097 | A/chicken/Changhua/15010120-1/2015 | A / H5N2 | Asia / Taiwan | |
| EPI_ISL_4032098 | A/chicken/Changhua/15010288/2015 | A / H5N2 | Asia / Taiwan | |
| EPI_ISL_4032099 | A/chicken/Changhua/15010324-2/2015 | A / H5N2 | Asia / Taiwan | |
| EPI_ISL_4032104 | A/chicken/Changhua/15010361-1/2015 | A / H5N2 | Asia / Taiwan | |
| EPI_ISL_4032148 | A/chicken/Changhua/15010362-2/2015 | A / H5N2 | Asia / Taiwan | |
| EPI_ISL_4032190 | A/chicken/Changhua/15010363-2/2015 | A / H5N2 | Asia / Taiwan | |
| EPI_ISL_4032194 | A/chicken/Changhua/15040036-1/2015 | A / H5N8 | Asia / Taiwan | |
| EPI_ISL_4032195 | A/chicken/Changhua/15040043-1/2015 | A / H5N8 | Asia / Taiwan | |
| EPI_ISL_4032196 | A/chicken/Changhua/15040037-2/2015 | A / H5N8 | Asia / Taiwan | |
| EPI_ISL_4032197 | A/chicken/Changhua/15040042/2015 | A / H5N8 | Asia / Taiwan | |
| EPI_ISL_4032198 | A/chicken/Changhua/15050013-1/2015 | A / H5N2 | Asia / Taiwan | |
| EPI_ISL_4032199 | A/chicken/Changhua/15050038-1/2015 | A / H5N8 | Asia / Taiwan | |
| EPI_ISL_4032200 | A/chicken/Changhua/15060027-2/2015 | A / H5N2 | Asia / Taiwan | |
| EPI_ISL_4032201 | A/chicken/Changhua/15060029/2015 | A / H5N8 | Asia / Taiwan | |
| EPI_ISL_4032202 | A/chicken/Changhua/15060037-1/2015 | A / H5N2 | Asia / Taiwan | |
| EPI_ISL_4032203 | A/chicken/Changhua/15120008/2015 | A / H5N2 | Asia / Taiwan | |
| EPI_ISL_4032204 | A/chicken/Changhua/16010060-2/2016 | A / H5N2 | Asia / Taiwan | |
| EPI_ISL_4032205 | A/chicken/Changhua/16020042-2/2016 | A / H5N2 | Asia / Taiwan | |
| EPI_ISL_4032206 | A/chicken/Changhua/17030063-1/2017 | A / H5N2 | Asia / Taiwan | |
| EPI_ISL_4032207 | A/chicken/Changhua/17060006/2017 | A / H5N2 | Asia / Taiwan | |
| EPI_ISL_4032208 | A/chicken/Changhua/18010011-1/2018 | A / H5N2 | Asia / Taiwan | |
| EPI_ISL_4032212 | A/chicken/Changhua/18040008-1/2018 | A / H5N2 | Asia / Taiwan | |
| EPI_ISL_4032213 | A/chicken/Chiayi/15010287-2/2015 | A / H5N2 | Asia / Taiwan | |
| EPI_ISL_4032214 | A/chicken/Chiayi/15010409-2/2015 | A / H5N2 | Asia / Taiwan | |
| EPI_ISL_4032215 | A/chicken/Chiayi/15010454-1/2015 | A / H5N8 | Asia / Taiwan | |
| EPI_ISL_4032216 | A/chicken/Chiayi/17030059-2/2017 | A / H5N2 | Asia / Taiwan | |
| EPI_ISL_4032217 | A/chicken/Chiayi/17040009-2/2017 | A / H5N2 | Asia / Taiwan | |
| EPI_ISL_4032218 | A/chicken/Chiayi/18120003-1/2018 | A / H5N2 | Asia / Taiwan | |
| EPI_ISL_4032219 | A/chicken/Hsinchu/16080004-2/2016 | A / H5N2 | Asia / Taiwan | |
| EPI_ISL_4032220 | A/chicken/Kaohsiung/16010047/2016 | A / H5N8 | Asia / Taiwan | |
| EPI_ISL_4032221 | A/chicken/Kaohsiung/16020017-2/2016 | A / H5N8 | Asia / Taiwan | |
| EPI_ISL_4032222 | A/chicken/Kaohsiung/16040021-1/2016 | A / H5N8 | Asia / Taiwan | |
| EPI_ISL_4032223 | A/chicken/Kaohsiung/16070003/2016 | A / H5N8 | Asia / Taiwan | |
| EPI_ISL_4032224 | A/chicken/Kaohsiung/16120012/2016 | A / H5N8 | Asia / Taiwan | |
| EPI_ISL_4032225 | A/chicken/Kaohsiung/16120034/2016 | A / H5N8 | Asia / Taiwan | |
| EPI_ISL_4032226 | A/chicken/Kaohsiung/16MB0889/2016 | A / H5N8 | Asia / Taiwan | |
| EPI_ISL_4032227 | A/chicken/Kaohsiung/17030023/2017 | A / H5N8 | Asia / Taiwan | |
| EPI_ISL_4032228 | A/chicken/Kaohsiung/17040006-2/2017 | A / H5N2 | Asia / Taiwan | |
| EPI_ISL_4032229 | A/chicken/Kaohsiung/17040007/2017 | A / H5N2 | Asia / Taiwan | |

| | | | |
|---|---|---|---|
| EPI_ISL_4032230 | A/chicken/Kaohsiung/17100002/2017 | A / H5N2 | Asia / Taiwan |
| EPI_ISL_4032235 | A/chicken/Kaohsiung/18080004-1/2018 | A / H5N2 | Asia / Taiwan |
| EPI_ISL_4032236 | A/chicken/Kaohsiung/18080004-2/2018 | A / H5N2 | Asia / Taiwan |
| EPI_ISL_4032237 | A/chicken/Miaoli/15010421-1/2015 | A / H5N8 | Asia / Taiwan |
| EPI_ISL_4032238 | A/chicken/Miaoli/18040011-1/2018 | A / H5N2 | Asia / Taiwan |
| EPI_ISL_4032239 | A/chicken/Nantou/16110023-1/2016 | A / H5N8 | Asia / Taiwan |
| EPI_ISL_4032240 | A/chicken/New Taipei City/16110019-1/2016 | A / H5N8 | Asia / Taiwan |
| EPI_ISL_4032241 | A/chicken/New Taipei City/16110020-2/2016 | A / H5N8 | Asia / Taiwan |
| EPI_ISL_4032242 | A/chicken/New Taipei City/17040002-2/2017 | A / H5N2 | Asia / Taiwan |
| EPI_ISL_4032243 | A/chicken/New Taipei City/17050004-2/2017 | A / H5N8 | Asia / Taiwan |
| EPI_ISL_4032244 | A/chicken/New Taipei City/17090001/2017 | A / H5N8 | Asia / Taiwan |
| EPI_ISL_4032245 | A/chicken/New Taipei City/17090002/2017 | A / H5N8 | Asia / Taiwan |
| EPI_ISL_4032246 | A/chicken/New Taipei City/17090003/2017 | A / H5N8 | Asia / Taiwan |
| EPI_ISL_4032247 | A/chicken/New Taipei City/18010021-3/2018 | A / H5N2 | Asia / Taiwan |
| EPI_ISL_4032248 | A/chicken/Pingtung/15010402/2015 | A / H5N2 | Asia / Taiwan |
| EPI_ISL_4032264 | A/chicken/Pingtung/15050006-1/2015 | A / H5N2 | Asia / Taiwan |
| EPI_ISL_4032266 | A/chicken/Pingtung/16020009-3/2016 | A / H5N2 | Asia / Taiwan |
| EPI_ISL_4032267 | A/chicken/Pingtung/18010017-1/2018 | A / H5N2 | Asia / Taiwan |
| EPI_ISL_4032268 | A/chicken/Pingtung/18010019-2/2018 | A / H5N2 | Asia / Taiwan |
| EPI_ISL_4032269 | A/chicken/Pingtung/18020015-1/2018 | A / H5N2 | Asia / Taiwan |
| EPI_ISL_4032270 | A/chicken/Pingtung/18050006-1/2018 | A / H5N2 | Asia / Taiwan |
| EPI_ISL_4032271 | A/chicken/Pingtung/18080001-1/2018 | A / H5N2 | Asia / Taiwan |
| EPI_ISL_4032272 | A/chicken/Taichung/15010399/2015 | A / H5N2 | Asia / Taiwan |
| EPI_ISL_4032273 | A/chicken/Taichung/16040032-3/2016 | A / H5N2 | Asia / Taiwan |
| EPI_ISL_4032274 | A/chicken/Taichung/16050044-2/2016 | A / H5N2 | Asia / Taiwan |
| EPI_ISL_4032275 | A/chicken/Taichung/16110017-3/2016 | A / H5N8 | Asia / Taiwan |
| EPI_ISL_4032276 | A/chicken/Tainan/15020214/2015 | A / H5N8 | Asia / Taiwan |
| EPI_ISL_4032277 | A/chicken/Tainan/16040035-1/2016 | A / H5N2 | Asia / Taiwan |
| EPI_ISL_4032278 | A/chicken/Tainan/17030058-1/2017 | A / H5N8 | Asia / Taiwan |
| EPI_ISL_4032279 | A/chicken/Tainan/18040014-2/2018 | A / H5N2 | Asia / Taiwan |
| EPI_ISL_4032280 | A/chicken/Taipei City/17090004/2017 | A / H5N8 | Asia / Taiwan |
| EPI_ISL_4032281 | A/chicken/Taipei City/18010007-3/2018 | A / H5N2 | Asia / Taiwan |
| EPI_ISL_4032282 | A/chicken/Taipei City/18010008-3/2018 | A / H5N2 | Asia / Taiwan |
| EPI_ISL_4032283 | A/chicken/Taipei City/18050007-1/2018 | A / H5N2 | Asia / Taiwan |
| EPI_ISL_4032284 | A/chicken/Taipei City/18050010-8/2018 | A / H5N2 | Asia / Taiwan |
| EPI_ISL_4032285 | A/chicken/Taipei/16060022-4/2016 | A / H5N8 | Asia / Taiwan |
| EPI_ISL_4032286 | A/chicken/Taipei/16060023-1/2016 | A / H5N8 | Asia / Taiwan |
| EPI_ISL_4032287 | A/chicken/Taipei/16120021/2016 | A / H5N8 | Asia / Taiwan |
| EPI_ISL_4032288 | A/chicken/Taipei/17050005-1/2017 | A / H5N8 | Asia / Taiwan |
| EPI_ISL_4032289 | A/chicken/Taipei/17050006-2/2017 | A / H5N8 | Asia / Taiwan |
| EPI_ISL_4032290 | A/chicken/Taipei/17050007-2/2017 | A / H5N8 | Asia / Taiwan |
| EPI_ISL_4032291 | A/chicken/Taipei/17050008-3/2017 | A / H5N8 | Asia / Taiwan |
| EPI_ISL_4032292 | A/chicken/Taipei/17050009-3/2017 | A / H5N8 | Asia / Taiwan |
| EPI_ISL_4032293 | A/chicken/Yunlin/15020048-1/2015 | A / H5N2 | Asia / Taiwan |
| EPI_ISL_4032294 | A/chicken/Yunlin/15040002-2/2015 | A / H5N2 | Asia / Taiwan |
| EPI_ISL_4032295 | A/chicken/Yunlin/15040027/2015 | A / H5N2 | Asia / Taiwan |
| EPI_ISL_4032296 | A/chicken/Yunlin/15050012/2015 | A / H5N8 | Asia / Taiwan |
| EPI_ISL_4032297 | A/chicken/Yunlin/15050037-2/2015 | A / H5N2 | Asia / Taiwan |
| EPI_ISL_4032298 | A/chicken/Yunlin/15050039-2/2015 | A / H5N2 | Asia / Taiwan |
| EPI_ISL_4032299 | A/chicken/Yunlin/15060014-1/2015 | A / H5N2 | Asia / Taiwan |
| EPI_ISL_4032300 | A/chicken/Yunlin/15080024/2015 | A / H5N2 | Asia / Taiwan |
| EPI_ISL_4032301 | A/chicken/Yunlin/16010001-1/2016 | A / H5N2 | Asia / Taiwan |
| EPI_ISL_4032302 | A/chicken/Yunlin/16010008-1/2016 | A / H5N2 | Asia / Taiwan |
| EPI_ISL_4032303 | A/chicken/Yunlin/16020038/2016 | A / H5N8 | Asia / Taiwan |
| EPI_ISL_4032304 | A/chicken/Yunlin/16050053/2016 | A / H5N2 | Asia / Taiwan |
| EPI_ISL_4032305 | A/chicken/Yunlin/17010001/2017 | A / H5N8 | Asia / Taiwan |
| EPI_ISL_4032306 | A/chicken/Yunlin/17010014/2017 | A / H5N2 | Asia / Taiwan |
| EPI_ISL_4032307 | A/chicken/Yunlin/17020033-2/2017 | A / H5N2 | Asia / Taiwan |
| EPI_ISL_4032308 | A/chicken/Yunlin/17020094-1/2017 | A / H5N2 | Asia / Taiwan |
| EPI_ISL_4032309 | A/chicken/Yunlin/17020095-2/2017 | A / H5N2 | Asia / Taiwan |
| EPI_ISL_4032310 | A/chicken/Yunlin/17030001-1/2017 | A / H5N2 | Asia / Taiwan |
| EPI_ISL_4032311 | A/chicken/Yunlin/17030007-1/2017 | A / H5N2 | Asia / Taiwan |
| EPI_ISL_4032312 | A/chicken/Yunlin/17030010-1/2017 | A / H5N2 | Asia / Taiwan |
| EPI_ISL_4032313 | A/chicken/Yunlin/17030014-1/2017 | A / H5N2 | Asia / Taiwan |
| EPI_ISL_4032314 | A/chicken/Yunlin/17030020-2/2017 | A / H5N2 | Asia / Taiwan |
| EPI_ISL_4032315 | A/chicken/Yunlin/17030021-1/2017 | A / H5N2 | Asia / Taiwan |
| EPI_ISL_4032316 | A/chicken/Yunlin/17030032-2/2017 | A / H5N2 | Asia / Taiwan |
| EPI_ISL_4032317 | A/chicken/Yunlin/17030065-2/2017 | A / H5N2 | Asia / Taiwan |
| EPI_ISL_4032318 | A/chicken/Yunlin/17040003-1/2017 | A / H5N2 | Asia / Taiwan |
| EPI_ISL_4032319 | A/chicken/Yunlin/17040004-2/2017 | A / H5N2 | Asia / Taiwan |
| EPI_ISL_4032320 | A/chicken/Yunlin/17040005-1/2017 | A / H5N2 | Asia / Taiwan |
| EPI_ISL_4032321 | A/chicken/Yunlin/17040010-1/2017 | A / H5N2 | Asia / Taiwan |

| | | | |
|---|---|---|---|
| EPI_ISL_4032322 | A/chicken/Yunlin/17040011-1/2017 | A / H5N2 | Asia / Taiwan |
| EPI_ISL_4032323 | A/chicken/Yunlin/17040012-1/2017 | A / H5N2 | Asia / Taiwan |
| EPI_ISL_4032324 | A/chicken/Yunlin/17060019/2017 | A / H5N2 | Asia / Taiwan |
| EPI_ISL_4032325 | A/chicken/Yunlin/17070010/2017 | A / H5N2 | Asia / Taiwan |
| EPI_ISL_4032326 | A/chicken/Yunlin/18010001-2/2018 | A / H5N2 | Asia / Taiwan |
| EPI_ISL_4032327 | A/chicken/Yunlin/18010003-3/2018 | A / H5N2 | Asia / Taiwan |
| EPI_ISL_4032328 | A/chicken/Yunlin/18010023-1/2018 | A / H5N2 | Asia / Taiwan |
| EPI_ISL_4032329 | A/chicken/Yunlin/18010024-2/2018 | A / H5N2 | Asia / Taiwan |
| EPI_ISL_4032330 | A/chicken/Yunlin/18020003-2/2018 | A / H5N2 | Asia / Taiwan |
| EPI_ISL_4032331 | A/chicken/Yunlin/18020004-1/2018 | A / H5N2 | Asia / Taiwan |
| EPI_ISL_4032332 | A/chicken/Yunlin/18020005-2/2018 | A / H5N2 | Asia / Taiwan |
| EPI_ISL_4032333 | A/chicken/Yunlin/18020008-2/2018 | A / H5N2 | Asia / Taiwan |
| EPI_ISL_4032334 | A/chicken/Yunlin/18020021-1/2018 | A / H5N2 | Asia / Taiwan |
| EPI_ISL_4032335 | A/chicken/Yunlin/18030002-1/2018 | A / H5N2 | Asia / Taiwan |
| EPI_ISL_4032336 | A/chicken/Yunlin/18040006-3/2018 | A / H5N2 | Asia / Taiwan |
| EPI_ISL_4032337 | A/chicken/Yunlin/18050013-1/2018 | A / H5N2 | Asia / Taiwan |
| EPI_ISL_4032338 | A/chicken/Yunlin/18060001-2/2018 | A / H5N2 | Asia / Taiwan |
| EPI_ISL_4032379 | A/chicken/Yunlin/18070003-3/2018 | A / H5N2 | Asia / Taiwan |
| EPI_ISL_4032419 | A/chicken/Yunlin/18080006-1/2018 | A / H5N2 | Asia / Taiwan |
| EPI_ISL_4032420 | A/duck/Changhua/15A03431-1-20T/2015 | A / H5N2 | Asia / Taiwan |
| EPI_ISL_4032421 | A/duck/Changhua/15A03473-1-20T/2015 | A / H5N2 | Asia / Taiwan |
| EPI_ISL_4032422 | A/duck/Chiayi/15080023/2015 | A / H5N8 | Asia / Taiwan |
| EPI_ISL_4032423 | A/duck/Hsinchu/15020216/2015 | A / H5N2 | Asia / Taiwan |
| EPI_ISL_4032424 | A/duck/Kaohsiung/15010213/2015 | A / H5N2 | Asia / Taiwan |
| EPI_ISL_4032425 | A/duck/Nantou/15A3437/2015 | A / H5N2 | Asia / Taiwan |
| EPI_ISL_4032426 | A/duck/Nantou/17A0243/2017 | A / H5N2 | Asia / Taiwan |
| EPI_ISL_4032427 | A/duck/Pingtung/15010145/2015 | A / H5N2 | Asia / Taiwan |
| EPI_ISL_4032428 | A/duck/Pingtung/15010157-1/2015 | A / H5N3 | Asia / Taiwan |
| EPI_ISL_4032429 | A/duck/Pingtung/15010180/2015 | A / H5N3 | Asia / Taiwan |
| EPI_ISL_4032430 | A/duck/Pingtung/15010444/2015 | A / H5N2 | Asia / Taiwan |
| EPI_ISL_4032431 | A/duck/Pingtung/15010461-1/2015 | A / H5N2 | Asia / Taiwan |
| EPI_ISL_4032432 | A/duck/Pingtung/15120006/2015 | A / H5N8 | Asia / Taiwan |
| EPI_ISL_4032433 | A/duck/Pingtung/15A03399-1-10T/2015 | A / H5N2 | Asia / Taiwan |
| EPI_ISL_4032434 | A/duck/Pingtung/17A00408-1-10T/2017 | A / H5N2 | Asia / Taiwan |
| EPI_ISL_4032435 | A/duck/Pingtung/17A0160/2017 | A / H5N2 | Asia / Taiwan |
| EPI_ISL_4032436 | A/duck/Pingtung/18A00003-11-20T/2018 | A / H5N2 | Asia / Taiwan |
| EPI_ISL_4032437 | A/duck/Pingtung/18A00076-11-20T/2018 | A / H5N2 | Asia / Taiwan |
| EPI_ISL_4032438 | A/duck/Pingtung/18X00077/2018 | A / H5N2 | Asia / Taiwan |
| EPI_ISL_4032439 | A/duck/Taichung/15010400-1/2015 | A / H5N2 | Asia / Taiwan |
| EPI_ISL_4032440 | A/duck/Tainan/15010403/2015 | A / H5N8 | Asia / Taiwan |
| EPI_ISL_4032441 | A/duck/Tainan/16010020-1/2016 | A / H5N2 | Asia / Taiwan |
| EPI_ISL_4032442 | A/duck/Tainan/16010024-1/2016 | A / H5N2 | Asia / Taiwan |
| EPI_ISL_4032443 | A/duck/Taoyuan/16010023-1/2016 | A / H5N2 | Asia / Taiwan |
| EPI_ISL_4032444 | A/duck/Yilan/17A0092/2017 | A / H5N2 | Asia / Taiwan |
| EPI_ISL_4032445 | A/duck/Yunlin/15010115-2/2015 | A / H5N2 | Asia / Taiwan |
| EPI_ISL_4032446 | A/duck/Yunlin/15010244-1/2015 | A / H5N2 | Asia / Taiwan |
| EPI_ISL_4032447 | A/duck/Yunlin/15010277/2015 | A / H5N8 | Asia / Taiwan |
| EPI_ISL_4032448 | A/duck/Yunlin/15010310-4/2015 | A / H5N2 | Asia / Taiwan |
| EPI_ISL_4032449 | A/duck/Yunlin/15010318-2/2015 | A / H5N2 | Asia / Taiwan |
| EPI_ISL_4032468 | A/duck/Yunlin/15010369-1/2015 | A / H5N2 | Asia / Taiwan |
| EPI_ISL_4032522 | A/duck/Yunlin/15010458-1/2015 | A / H5N8 | Asia / Taiwan |
| EPI_ISL_4032531 | A/duck/Yunlin/15010538/2015 | A / H5N8 | Asia / Taiwan |
| EPI_ISL_4032532 | A/duck/Yunlin/15070002-2/2015 | A / H5N2 | Asia / Taiwan |
| EPI_ISL_4032533 | A/duck/Yunlin/15A03403-1-20T/2015 | A / H5N2 | Asia / Taiwan |
| EPI_ISL_4032534 | A/duck/Yunlin/15A04883-1-10T/2015 | A / H5N2 | Asia / Taiwan |
| EPI_ISL_4032535 | A/duck/Yunlin/15A5019/2015 | A / H5N2 | Asia / Taiwan |
| EPI_ISL_4032536 | A/duck/Yunlin/17110001/2017 | A / H5N2 | Asia / Taiwan |
| EPI_ISL_4032537 | A/duck/Yunlin/17A0086/2017 | A / H5N2 | Asia / Taiwan |
| EPI_ISL_4032538 | A/duck/Yunlin/17A0686/2017 | A / H5N2 | Asia / Taiwan |
| EPI_ISL_4032539 | A/duck/Yunlin/18010004-2/2018 | A / H5N2 | Asia / Taiwan |
| EPI_ISL_4032540 | A/duck/Yunlin/18060006-1/2018 | A / H5N2 | Asia / Taiwan |
| EPI_ISL_4032541 | A/duck/Yunlin/18060007-3/2018 | A / H5N2 | Asia / Taiwan |
| EPI_ISL_4032542 | A/duck/Yunlin/18X00007-3/2018 | A / H5N2 | Asia / Taiwan |
| EPI_ISL_4032543 | A/environment/Yunlin/18X00007/2018 | A / H5N2 | Asia / Taiwan |
| EPI_ISL_4032544 | A/environment/Yunlin/18X00070/2018 | A / H5N2 | Asia / Taiwan |
| EPI_ISL_4032545 | A/goose/Changhua/15040008/2015 | A / H5N2 | Asia / Taiwan |
| EPI_ISL_4032546 | A/goose/Chiayi/15010008-3/2015 | A / H5N2 | Asia / Taiwan |
| EPI_ISL_4032547 | A/goose/Chiayi/15010016-1/2015 | A / H5N2 | Asia / Taiwan |
| EPI_ISL_4032548 | A/goose/Chiayi/15020022/2015 | A / H5N8 | Asia / Taiwan |
| EPI_ISL_4032549 | A/goose/Chiayi/16010017-2/2016 | A / H5N2 | Asia / Taiwan |
| EPI_ISL_4032550 | A/goose/Chiayi/16040003-1/2016 | A / H5N2 | Asia / Taiwan |
| EPI_ISL_4032551 | A/goose/Chiayi/16040005/2016 | A / H5N2 | Asia / Taiwan |

| | | | | |
|---|---|---|---|---|
| EPI_ISL_4032552 | A/goose/Chiayi/16040007-1/2016 | A / H5N2 | Asia / Taiwan | |
| EPI_ISL_4032553 | A/goose/Chiayi/17020027-1/2017 | A / H5N2 | Asia / Taiwan | |
| EPI_ISL_4032554 | A/goose/Chiayi/17040014/2017 | A / H5N2 | Asia / Taiwan | |
| EPI_ISL_4032555 | A/goose/Chiayi/18010006/2018 | A / H5N2 | Asia / Taiwan | |
| EPI_ISL_4032556 | A/goose/Chiayi/18020002-2/2018 | A / H5N2 | Asia / Taiwan | |
| EPI_ISL_4032558 | A/goose/Chiayi/18020014-2/2018 | A / H5N2 | Asia / Taiwan | |
| EPI_ISL_4032559 | A/goose/Chiayi/18020014-3/2018 | A / H5N2 | Asia / Taiwan | |
| EPI_ISL_4032560 | A/goose/Pingtung/15010007-4/2015 | A / H5N2 | Asia / Taiwan | |
| EPI_ISL_4032561 | A/goose/Pingtung/15010037-1/2015 | A / H5N2 | Asia / Taiwan | |
| EPI_ISL_4032562 | A/goose/Pingtung/15050014/2015 | A / H5N8 | Asia / Taiwan | |
| EPI_ISL_4032563 | A/goose/Pingtung/16010061/2016 | A / H5N8 | Asia / Taiwan | |
| EPI_ISL_4032564 | A/goose/Tainan/16010031-2/2016 | A / H5N2 | Asia / Taiwan | |
| EPI_ISL_4032565 | A/goose/Tainan/16020041-2/2016 | A / H5N2 | Asia / Taiwan | |
| EPI_ISL_4032566 | A/goose/Tainan/16040018-1/2016 | A / H5N8 | Asia / Taiwan | |
| EPI_ISL_4032567 | A/goose/Tainan/16070002/2016 | A / H5N8 | Asia / Taiwan | |
| EPI_ISL_4032568 | A/goose/Tainan/17030022-1/2017 | A / H5N2 | Asia / Taiwan | |
| EPI_ISL_4032569 | A/goose/Tainan/17030024-1/2017 | A / H5N2 | Asia / Taiwan | |
| EPI_ISL_4032570 | A/goose/Tainan/18010022-1/2018 | A / H5N2 | Asia / Taiwan | |
| EPI_ISL_4032571 | A/goose/Tainan/18050014-1/2018 | A / H5N2 | Asia / Taiwan | |
| EPI_ISL_4032572 | A/goose/Taoyuan/15010041/2015 | A / H5N8 | Asia / Taiwan | |
| EPI_ISL_4032573 | A/goose/Taoyuan/18120002-2/2018 | A / H5N2 | Asia / Taiwan | |
| EPI_ISL_4032574 | A/goose/Yunlin/15010005/2015 | A / H5N2 | Asia / Taiwan | |
| EPI_ISL_4032575 | A/goose/Yunlin/15010009-1/2015 | A / H5N2 | Asia / Taiwan | |
| EPI_ISL_4032576 | A/goose/Yunlin/15010010-2/2015 | A / H5N2 | Asia / Taiwan | |
| EPI_ISL_4032577 | A/goose/Yunlin/15010011-1/2015 | A / H5N2 | Asia / Taiwan | |
| EPI_ISL_4032578 | A/goose/Yunlin/15010029-1/2015 | A / H5N2 | Asia / Taiwan | |
| EPI_ISL_404207 | A/duck/Nghe An/5382VTC/2019 | A / H5N6 | Asia / Vietnam / Tinh Ha Tinh | Natalia Goncharova (State Research Center of Virology and Biotechnology (VECTOR) / Emerging Zoonotic Diseases and Influenza) |
| EPI_ISL_404986 | A/chicken/Ba Ria-Vung Tau/21078VTC/2019 | A / H5N6 | Asia / Vietnam / Tinh Ba Ria-Vung Tau | Natalia Goncharova (State Research Center of Virology and Biotechnology (VECTOR) / Emerging Zoonotic Diseases and Influenza) |
| EPI_ISL_404987 | A/chicken/Dong Nai/25437VTC/2019 | A / H5N6 | Asia / Vietnam / Tinh Dong Nai | Natalia Goncharova (State Research Center of Virology and Biotechnology (VECTOR) / Emerging Zoonotic Diseases and Influenza) |
| EPI_ISL_404988 | A/chicken/Thanh Hoa/13836VTC/2019 | A / H5N6 | Asia / Vietnam / Tinh Thanh Hoa | Natalia Goncharova (State Research Center of Virology and Biotechnology (VECTOR) / Emerging Zoonotic Diseases and Influenza) |
| EPI_ISL_404993 | A/white-fronted goose/Germany-BB/AI00018/2020 | A / H5N8 | Europe / Germany / Brandenburg / Spree/Neiße | Jacqueline King (Friedrich-Loeffler-Institut) |
| EPI_ISL_405278 | A/guinea_fowl/Nigeria/OG-GF11T_19VIR8424-7/2019_H5N8 | A / H5N8 | Africa / Nigeria / Ogun State | Bianca Zecchin (Istituto Zooprofilattico Sperimentale Delle Venezie) |
| EPI_ISL_405391 | A/chicken/Czech Republic/1175-1/2020 | A / H5N8 | Europe / Czech Republic / Vysocina Kraj / Okres Zd'ar nad Szavou / Št?pánov nad Svratkou | Alexander Nagy (State Veterinary Institute Prague) |
| EPI_ISL_4055115 | A/goose/Yunlin/15040040/2015 | A / H5N8 | Asia / Taiwan | |
| EPI_ISL_4055116 | A/goose/Yunlin/16020008-4/2016 | A / H5N2 | Asia / Taiwan | |
| EPI_ISL_4055117 | A/goose/Yunlin/16020011-1/2016 | A / H5N2 | Asia / Taiwan | |
| EPI_ISL_4055118 | A/goose/Yunlin/16030042-2/2016 | A / H5N2 | Asia / Taiwan | |
| EPI_ISL_4055119 | A/goose/Yunlin/16040002-2/2016 | A / H5N2 | Asia / Taiwan | |
| EPI_ISL_4055120 | A/goose/Yunlin/17100005/2017 | A / H5N2 | Asia / Taiwan | |
| EPI_ISL_4055121 | A/goose/Yunlin/18030020-1/2018 | A / H5N2 | Asia / Taiwan | |
| EPI_ISL_4055122 | A/goose/Yunlin/18030020-2/2018 | A / H5N2 | Asia / Taiwan | |
| EPI_ISL_4055123 | A/goose/Yunlin/18030020-3/2018 | A / H5N2 | Asia / Taiwan | |
| EPI_ISL_4055124 | A/goose/Yunlin/18040002-3/2018 | A / H5N2 | Asia / Taiwan | |
| EPI_ISL_4055125 | A/goose/Yunlin/18040018-2/2018 | A / H5N2 | Asia / Taiwan | |
| EPI_ISL_4055126 | A/goose/Yunlin/18050003-2/2018 | A / H5N2 | Asia / Taiwan | |
| EPI_ISL_4055127 | A/goose/Yunlin/18090002-3/2018 | A / H5N2 | Asia / Taiwan | |
| EPI_ISL_4055128 | A/goose/Yunlin/18090006-1/2018 | A / H5N2 | Asia / Taiwan | |
| EPI_ISL_4055129 | A/goose/Yunlin/18110002-3/2018 | A / H5N2 | Asia / Taiwan | |
| EPI_ISL_4055132 | A/turkey/Changhua/16010011-1/2016 | A / H5N2 | Asia / Taiwan | |
| EPI_ISL_4055133 | A/turkey/Chiayi/17020086-2/2017 | A / H5N8 | Asia / Taiwan | |
| EPI_ISL_4055135 | A/turkey/Chiayi/17020092-2/2017 | A / H5N8 | Asia / Taiwan | |
| EPI_ISL_4055136 | A/turkey/Chiayi/18010013-3/2018 | A / H5N2 | Asia / Taiwan | |
| EPI_ISL_4055137 | A/turkey/Yunlin/15030069-1/2015 | A / H5N2 | Asia / Taiwan | |
| EPI_ISL_4055138 | A/turkey/Yunlin/15060049-1/2015 | A / H5N2 | Asia / Taiwan | |
| EPI_ISL_4055139 | A/turkey/Yunlin/18020020-2/2018 | A / H5N2 | Asia / Taiwan | |
| EPI_ISL_4055140 | A/turkey/Yunlin/18070001-2/2018 | A / H5N2 | Asia / Taiwan | |
| EPI_ISL_4055141 | A/turkey/Yunlin/18090003-3/2018 | A / H5N2 | Asia / Taiwan | |
| EPI_ISL_4055142 | A/turkey/Yunlin/18110001-3/2018 | A / H5N2 | Asia / Taiwan | |
| EPI_ISL_4055143 | A/turkey/Yunlin/18120005-2/2018 | A / H5N2 | Asia / Taiwan | |
| EPI_ISL_4055845 | A/duck/Ugii lake/#66/2018 | A / H5N6 | Asia / Mongolia | |
| EPI_ISL_4055846 | A/duck/Khunt lake/#500/2019 | A / H5N6 | Asia / Mongolia | |
| EPI_ISL_405813 | A/hawk/Poland/003/2020 | A / H5N8 | Europe / Poland / Lublin Voivodeship | Edyta ?wi?to? (National Veterinary Research Institut Poland, PIWet-PIB) |
| EPI_ISL_4061481 | A/chicken/Nigeria/VRD21-43_21VIR2288-4/2021 | A / H5N8 | Africa / Nigeria | |
| EPI_ISL_4061482 | A/chicken/Nigeria/VRD21-035B_21VIR2288-1/2021 | A / H5N8 | Africa / Nigeria | |
| EPI_ISL_4061483 | A/chicken/Nigeria/VRD21-100_21VIR2370-423/2021 | A / H5N8 | Africa / Nigeria | |
| EPI_ISL_4061484 | A/chicken/Nigeria/VRD21-102_21VIR2370-424/2021 | A / H5N1 | Africa / Nigeria | |
| EPI_ISL_4061485 | A/chicken/Nigeria/VRD21-109_21VIR2370-425/2021 | A / H5N1 | Africa / Nigeria | |
| EPI_ISL_4061486 | A/chicken/Nigeria/VRD21-37_21VIR2288-2/2021 | A / H5N1 | Africa / Nigeria | |
| EPI_ISL_4061488 | A/chicken/Nigeria/VRD21-53B_21VIR2288-5/2021 | A / H5N8 | Africa / Nigeria | |
| EPI_ISL_4061490 | A/chicken/Nigeria/VRD21-88_21VIR2288-8/2021 | A / H5N8 | Africa / Nigeria | |
| EPI_ISL_4061491 | A/chicken/Nigeria/VRD21-98_21VIR2288-6/2021 | A / H5N1 | Africa / Nigeria | |
| EPI_ISL_4061541 | A/chicken/Netherlands/2113/2017 | A / H5N8 | Europe / Netherlands | |

| | | | |
|---|---|---|---|
| EPI_ISL_4061544 | A/Quail/Viet Nam/4615/2015 | A / H5N6 | Asia / Vietnam |
| EPI_ISL_4061567 | A/chicken/Viet Nam/6115/2015 | A / H5N6 | Asia / Vietnam |
| EPI_ISL_4061568 | A/Duck/Viet Nam/5314/2014 | A / H5N6 | Asia / Vietnam |
| EPI_ISL_4061712 | A/whooper swan/Miyagi/0402B001/2021 | A / H5N8 | Asia / Japan / Miyagi |
| EPI_ISL_4061713 | A/whooper swan/Fukushima/0701B002/2021 | A / H5N8 | Asia / Japan / Fukushima |
| EPI_ISL_4062109 | A/chicken/HeBei/HB1905/2019 | A / H5N6 | Asia / China |
| EPI_ISL_4062111 | A/chicken/HeBei/HB1907/2019 | A / H5N6 | Asia / China |
| EPI_ISL_4062113 | A/chicken/Pakistan/531/2018 | A / H5N8 | Asia / Pakistan |
| EPI_ISL_4062114 | A/quail/Pakistan/998/2018 | A / H5N8 | Asia / Pakistan |
| EPI_ISL_4062115 | A/chicken/Pakistan/1000/2018 | A / H5N8 | Asia / Pakistan |
| EPI_ISL_4062116 | A/environment/Pakistan/927/2018 | A / H5N8 | Asia / Pakistan |
| EPI_ISL_4062127 | A/chicken/HeBei/CK05/2019 | A / H5N6 | Asia / China |
| EPI_ISL_4062128 | A/goose/HeBei/GD07/2019 | A / H5N6 | Asia / China |
| EPI_ISL_4062437 | A/goose/China/21FU001/2020 | A / H5N8 | Asia / China |
| EPI_ISL_4062438 | A/goose/China/21FU002/2020 | A / H5N8 | Asia / China |
| EPI_ISL_4062439 | A/goose/China/21FU003/2020 | A / H5N8 | Asia / China |
| EPI_ISL_4062440 | A/goose/China/21FU004/2020 | A / H5N8 | Asia / China |
| EPI_ISL_4062441 | A/goose/China/21FU005/2020 | A / H5N8 | Asia / China |
| EPI_ISL_4062442 | A/goose/China/21FU006/2020 | A / H5N8 | Asia / China |
| EPI_ISL_4062443 | A/goose/China/21FU007/2020 | A / H5N8 | Asia / China |
| EPI_ISL_4062444 | A/goose/China/21FU008/2020 | A / H5N8 | Asia / China |
| EPI_ISL_4063606 | A/chicken/Northern China/F0130c/2018 | A / H5N6 | Asia / China |
| EPI_ISL_4064431 | A/duck/France/161113/2016 | A / H5N8 | Europe / France |
| EPI_ISL_4064432 | A/duck/France/161116/2016 | A / H5N8 | Europe / France |
| EPI_ISL_4064433 | A/duck/France/161137/2016 | A / H5N8 | Europe / France |
| EPI_ISL_4064434 | A/duck/France/161141/2016 | A / H5N8 | Europe / France |
| EPI_ISL_4064435 | A/duck/France/161142/2016 | A / H5N8 | Europe / France |
| EPI_ISL_4064436 | A/duck/France/161143/2016 | A / H5N8 | Europe / France |
| EPI_ISL_4064437 | A/duck/France/161147/2016 | A / H5N8 | Europe / France |
| EPI_ISL_4064438 | A/duck/France/161173/2016 | A / H5N8 | Europe / France |
| EPI_ISL_4064439 | A/duck/France/161174/2016 | A / H5N8 | Europe / France |
| EPI_ISL_4064440 | A/duck/France/161182/2016 | A / H5N8 | Europe / France |
| EPI_ISL_4064441 | A/duck/France/161207/2016 | A / H5N8 | Europe / France |
| EPI_ISL_4064442 | A/duck/France/161212/2016 | A / H5N8 | Europe / France |
| EPI_ISL_4064443 | A/duck/France/161224/2016 | A / H5N8 | Europe / France |
| EPI_ISL_4064444 | A/duck/France/161225/2016 | A / H5N8 | Europe / France |
| EPI_ISL_4064445 | A/duck/France/161227/2016 | A / H5N8 | Europe / France |
| EPI_ISL_4064446 | A/duck/France/161228/2016 | A / H5N8 | Europe / France |
| EPI_ISL_4064447 | A/duck/France/161229/2016 | A / H5N8 | Europe / France |
| EPI_ISL_4064448 | A/duck/France/161230/2016 | A / H5N8 | Europe / France |
| EPI_ISL_4064449 | A/duck/France/161231/2016 | A / H5N8 | Europe / France |
| EPI_ISL_4064450 | A/duck/France/161233/2016 | A / H5N8 | Europe / France |
| EPI_ISL_4064451 | A/duck/France/161234/2016 | A / H5N8 | Europe / France |
| EPI_ISL_4064452 | A/duck/France/161239/2016 | A / H5N8 | Europe / France |
| EPI_ISL_4064453 | A/duck/France/161240/2016 | A / H5N8 | Europe / France |
| EPI_ISL_4064454 | A/duck/France/161242/2016 | A / H5N8 | Europe / France |
| EPI_ISL_4064455 | A/duck/France/161243/2016 | A / H5N8 | Europe / France |
| EPI_ISL_4064456 | A/duck/France/161256/2016 | A / H5N8 | Europe / France |
| EPI_ISL_4064457 | A/duck/France/161263/2016 | A / H5N8 | Europe / France |
| EPI_ISL_4064458 | A/duck/France/161271/2016 | A / H5N8 | Europe / France |
| EPI_ISL_4064459 | A/duck/France/161275/2016 | A / H5N8 | Europe / France |
| EPI_ISL_4064460 | A/duck/France/161277/2016 | A / H5N8 | Europe / France |
| EPI_ISL_4064461 | A/duck/France/161298/2016 | A / H5N8 | Europe / France |
| EPI_ISL_4064462 | A/duck/France/161383/2016 | A / H5N8 | Europe / France |
| EPI_ISL_4064463 | A/duck/France/161443/2016 | A / H5N8 | Europe / France |
| EPI_ISL_4064464 | A/duck/France/161444/2016 | A / H5N8 | Europe / France |
| EPI_ISL_4064465 | A/duck/France/161445/2016 | A / H5N8 | Europe / France |
| EPI_ISL_4064466 | A/duck/France/161449/2016 | A / H5N8 | Europe / France |
| EPI_ISL_4064467 | A/duck/France/161450/2016 | A / H5N8 | Europe / France |
| EPI_ISL_4064468 | A/duck/France/161455/2016 | A / H5N8 | Europe / France |
| EPI_ISL_4064469 | A/duck/France/161456/2016 | A / H5N8 | Europe / France |
| EPI_ISL_4064470 | A/duck/France/161457/2016 | A / H5N8 | Europe / France |
| EPI_ISL_4064471 | A/duck/France/161473/2016 | A / H5N8 | Europe / France |
| EPI_ISL_4064472 | A/duck/France/161475/2016 | A / H5N8 | Europe / France |
| EPI_ISL_4064473 | A/duck/France/161476/2016 | A / H5N8 | Europe / France |
| EPI_ISL_4064474 | A/duck/France/161477/2016 | A / H5N8 | Europe / France |
| EPI_ISL_4064475 | A/duck/France/161478/2016 | A / H5N8 | Europe / France |
| EPI_ISL_4064476 | A/duck/France/161497/2016 | A / H5N8 | Europe / France |
| EPI_ISL_4064477 | A/duck/France/161498/2016 | A / H5N8 | Europe / France |
| EPI_ISL_4064478 | A/duck/France/161500/2016 | A / H5N8 | Europe / France |
| EPI_ISL_4064479 | A/duck/France/161501/2016 | A / H5N8 | Europe / France |
| EPI_ISL_4064480 | A/duck/France/161577/2016 | A / H5N8 | Europe / France |

| | | | |
|---|---|---|---|
| EPI_ISL_4064482 | A/duck/France/161600/2016 | A / H5N8 | Europe / France |
| EPI_ISL_4064483 | A/duck/France/161601/2016 | A / H5N8 | Europe / France |
| EPI_ISL_4064484 | A/duck/France/170013/2016 | A / H5N8 | Europe / France |
| EPI_ISL_4064485 | A/duck/France/170016/2016 | A / H5N8 | Europe / France |
| EPI_ISL_4064486 | A/duck/France/170017/2016 | A / H5N8 | Europe / France |
| EPI_ISL_4064487 | A/duck/France/170018/2017 | A / H5N8 | Europe / France |
| EPI_ISL_4064488 | A/duck/France/170030/2016 | A / H5N8 | Europe / France |
| EPI_ISL_4064489 | A/duck/France/170031/2017 | A / H5N8 | Europe / France |
| EPI_ISL_4064490 | A/duck/France/170032/2017 | A / H5N8 | Europe / France |
| EPI_ISL_4064491 | A/duck/France/170033/2016 | A / H5N8 | Europe / France |
| EPI_ISL_4064492 | A/duck/France/170034/2017 | A / H5N8 | Europe / France |
| EPI_ISL_4064493 | A/duck/France/170036/2017 | A / H5N8 | Europe / France |
| EPI_ISL_4064494 | A/duck/France/170038/2017 | A / H5N8 | Europe / France |
| EPI_ISL_4064495 | A/chicken/France/170063/2017 | A / H5N8 | Europe / France |
| EPI_ISL_4064496 | A/duck/France/170064/2017 | A / H5N8 | Europe / France |
| EPI_ISL_4064497 | A/duck/France/170067/2017 | A / H5N8 | Europe / France |
| EPI_ISL_4064498 | A/duck/France/170098/2017 | A / H5N8 | Europe / France |
| EPI_ISL_4064499 | A/duck/France/170117/2017 | A / H5N8 | Europe / France |
| EPI_ISL_4064500 | A/duck/France/170118/2017 | A / H5N8 | Europe / France |
| EPI_ISL_4064501 | A/chicken/France/170176/2017 | A / H5N8 | Europe / France |
| EPI_ISL_4064502 | A/duck/France/170177/2017 | A / H5N8 | Europe / France |
| EPI_ISL_4064503 | A/duck/France/170178/2017 | A / H5N8 | Europe / France |
| EPI_ISL_4064504 | A/duck/France/170180/2017 | A / H5N8 | Europe / France |
| EPI_ISL_4064505 | A/duck/France/170245/2017 | A / H5N8 | Europe / France |
| EPI_ISL_4064506 | A/duck/France/170249/2017 | A / H5N8 | Europe / France |
| EPI_ISL_4064507 | A/duck/France/170250/2017 | A / H5N8 | Europe / France |
| EPI_ISL_4064508 | A/turkey/France/170278/2017 | A / H5N8 | Europe / France |
| EPI_ISL_4064509 | A/duck/France/170280/2017 | A / H5N8 | Europe / France |
| EPI_ISL_4064510 | A/duck/France/170284/2017 | A / H5N8 | Europe / France |
| EPI_ISL_4064511 | A/duck/France/170308/2017 | A / H5N8 | Europe / France |
| EPI_ISL_4064512 | A/duck/France/170309/2017 | A / H5N8 | Europe / France |
| EPI_ISL_4064513 | A/duck/France/170325/2017 | A / H5N8 | Europe / France |
| EPI_ISL_4064514 | A/duck/France/170330/2017 | A / H5N8 | Europe / France |
| EPI_ISL_4064515 | A/duck/France/170331/2017 | A / H5N8 | Europe / France |
| EPI_ISL_4064516 | A/duck/France/170338/2017 | A / H5N8 | Europe / France |
| EPI_ISL_4064517 | A/duck/France/170339/2017 | A / H5N8 | Europe / France |
| EPI_ISL_4064518 | A/chicken/France/170340/2017 | A / H5N8 | Europe / France |
| EPI_ISL_4064519 | A/duck/France/170341/2017 | A / H5N8 | Europe / France |
| EPI_ISL_4064520 | A/duck/France/170349/2017 | A / H5N8 | Europe / France |
| EPI_ISL_4064521 | A/duck/France/170368/2017 | A / H5N8 | Europe / France |
| EPI_ISL_4064522 | A/duck/France/170369/2017 | A / H5N8 | Europe / France |
| EPI_ISL_4064523 | A/duck/France/170370/2017 | A / H5N8 | Europe / France |
| EPI_ISL_4064525 | A/chicken/France/170407/2017 | A / H5N8 | Europe / France |
| EPI_ISL_4064526 | A/chicken/France/170408/2017 | A / H5N8 | Europe / France |
| EPI_ISL_4064527 | A/guineafowl/France/170409/2017 | A / H5N8 | Europe / France |
| EPI_ISL_4064535 | A/duck/France/170432/2017 | A / H5N8 | Europe / France |
| EPI_ISL_4064541 | A/chicken/France/170433/2017 | A / H5N8 | Europe / France |
| EPI_ISL_4064548 | A/chicken/France/170436/2017 | A / H5N8 | Europe / France |
| EPI_ISL_4064555 | A/duck/France/170437/2017 | A / H5N8 | Europe / France |
| EPI_ISL_4064569 | A/duck/France/170518/2017 | A / H5N8 | Europe / France |
| EPI_ISL_4064575 | A/duck/France/170612/2017 | A / H5N8 | Europe / France |
| EPI_ISL_4064582 | A/duck/France/170683/2017 | A / H5N8 | Europe / France |
| EPI_ISL_4064589 | A/duck/France/170685/2017 | A / H5N8 | Europe / France |
| EPI_ISL_4064597 | A/chicken/France/170686/2017 | A / H5N8 | Europe / France |
| EPI_ISL_4064603 | A/duck/France/170733/2017 | A / H5N8 | Europe / France |
| EPI_ISL_4064610 | A/duck/France/170734/2017 | A / H5N8 | Europe / France |
| EPI_ISL_4064617 | A/duck/France/170735/2017 | A / H5N8 | Europe / France |
| EPI_ISL_4064619 | A/duck/France/170738/2017 | A / H5N8 | Europe / France |
| EPI_ISL_4064621 | A/duck/France/170772/2017 | A / H5N8 | Europe / France |
| EPI_ISL_4064622 | A/chicken/France/170773/2017 | A / H5N8 | Europe / France |
| EPI_ISL_4064623 | A/chicken/France/170774/2017 | A / H5N8 | Europe / France |
| EPI_ISL_4064624 | A/duck/France/170775/2017 | A / H5N8 | Europe / France |
| EPI_ISL_4064625 | A/duck/France/170806/2017 | A / H5N8 | Europe / France |
| EPI_ISL_4064626 | A/duck/France/170820/2017 | A / H5N8 | Europe / France |
| EPI_ISL_4064627 | A/duck/France/170822/2017 | A / H5N8 | Europe / France |
| EPI_ISL_4064628 | A/duck/France/170974/2017 | A / H5N8 | Europe / France |
| EPI_ISL_4064629 | A/duck/France/170975/2017 | A / H5N8 | Europe / France |
| EPI_ISL_4064630 | A/goose/France/170976/2017 | A / H5N8 | Europe / France |
| EPI_ISL_4064631 | A/avian/France/170977/2017 | A / H5N8 | Europe / France |
| EPI_ISL_4064632 | A/duck/France/171131/2017 | A / H5N8 | Europe / France |
| EPI_ISL_4064633 | A/duck/France/171133/2017 | A / H5N8 | Europe / France |
| EPI_ISL_4064634 | A/duck/France/171134/2017 | A / H5N8 | Europe / France |

| | | | |
|---|---|---|---|
| EPI_ISL_4064635 | A/duck/France/171202/2017 | A / H5N8 | Europe / France |
| EPI_ISL_4064636 | A/duck/France/171253/2017 | A / H5N8 | Europe / France |
| EPI_ISL_4064637 | A/duck/France/171270/2017 | A / H5N8 | Europe / France |
| EPI_ISL_4064638 | A/duck/France/171275/2017 | A / H5N8 | Europe / France |
| EPI_ISL_4064639 | A/goose/France/171278/2017 | A / H5N8 | Europe / France |
| EPI_ISL_4064640 | A/duck/France/171378/2017 | A / H5N8 | Europe / France |
| EPI_ISL_4064641 | A/duck/France/171379/2017 | A / H5N8 | Europe / France |
| EPI_ISL_4064642 | A/duck/France/171408/2017 | A / H5N8 | Europe / France |
| EPI_ISL_4064643 | A/duck/France/171410/2017 | A / H5N8 | Europe / France |
| EPI_ISL_4064645 | A/duck/France/171455/2017 | A / H5N8 | Europe / France |
| EPI_ISL_4064646 | A/duck/France/171457/2017 | A / H5N8 | Europe / France |
| EPI_ISL_4064647 | A/duck/France/171461/2017 | A / H5N8 | Europe / France |
| EPI_ISL_4064648 | A/duck/France/171462/2017 | A / H5N8 | Europe / France |
| EPI_ISL_4064649 | A/duck/France/171463/2017 | A / H5N8 | Europe / France |
| EPI_ISL_4064650 | A/duck/France/171466/2017 | A / H5N8 | Europe / France |
| EPI_ISL_4064651 | A/duck/France/171467/2017 | A / H5N8 | Europe / France |
| EPI_ISL_4064652 | A/duck/France/171468/2017 | A / H5N8 | Europe / France |
| EPI_ISL_4064653 | A/duck/France/171469/2017 | A / H5N8 | Europe / France |
| EPI_ISL_4064654 | A/chicken/France/171471/2017 | A / H5N8 | Europe / France |
| EPI_ISL_4064655 | A/duck/France/171472/2017 | A / H5N8 | Europe / France |
| EPI_ISL_4064656 | A/duck/France/171486/2017 | A / H5N8 | Europe / France |
| EPI_ISL_4064657 | A/duck/France/171514/2017 | A / H5N8 | Europe / France |
| EPI_ISL_4064658 | A/chicken/France/171525/2017 | A / H5N8 | Europe / France |
| EPI_ISL_4064659 | A/duck/France/171533/2017 | A / H5N8 | Europe / France |
| EPI_ISL_4064660 | A/chicken/France/171534/2017 | A / H5N8 | Europe / France |
| EPI_ISL_4064661 | A/duck/France/171535/2017 | A / H5N8 | Europe / France |
| EPI_ISL_4064662 | A/duck/France/171536/2017 | A / H5N8 | Europe / France |
| EPI_ISL_4064663 | A/chicken/France/171537/2017 | A / H5N8 | Europe / France |
| EPI_ISL_4064664 | A/duck/France/171539/2017 | A / H5N8 | Europe / France |
| EPI_ISL_4064665 | A/duck/France/171541/2017 | A / H5N8 | Europe / France |
| EPI_ISL_4064666 | A/duck/France/171542/2017 | A / H5N8 | Europe / France |
| EPI_ISL_4064667 | A/duck/France/171543/2017 | A / H5N8 | Europe / France |
| EPI_ISL_4064668 | A/duck/France/171546/2017 | A / H5N8 | Europe / France |
| EPI_ISL_4064669 | A/duck/France/171550/2017 | A / H5N8 | Europe / France |
| EPI_ISL_4064670 | A/avian/France/171599/2017 | A / H5N8 | Europe / France |
| EPI_ISL_4064671 | A/duck/France/171603/2017 | A / H5N8 | Europe / France |
| EPI_ISL_4064672 | A/avian/France/171606/2017 | A / H5N8 | Europe / France |
| EPI_ISL_4064673 | A/duck/France/171607/2017 | A / H5N8 | Europe / France |
| EPI_ISL_4064674 | A/duck/France/171639/2017 | A / H5N8 | Europe / France |
| EPI_ISL_4064675 | A/duck/France/171641/2017 | A / H5N8 | Europe / France |
| EPI_ISL_4064676 | A/duck/France/171650/2017 | A / H5N8 | Europe / France |
| EPI_ISL_4064677 | A/duck/France/171860/2017 | A / H5N8 | Europe / France |
| EPI_ISL_4064678 | A/duck/France/171874/2017 | A / H5N8 | Europe / France |
| EPI_ISL_4064679 | A/duck/France/171884/2017 | A / H5N8 | Europe / France |
| EPI_ISL_4064680 | A/chicken/France/171915/2017 | A / H5N8 | Europe / France |
| EPI_ISL_4064681 | A/duck/France/171928/2017 | A / H5N8 | Europe / France |
| EPI_ISL_4064682 | A/duck/France/171929/2017 | A / H5N8 | Europe / France |
| EPI_ISL_4064683 | A/duck/France/171932/2017 | A / H5N8 | Europe / France |
| EPI_ISL_4064684 | A/chicken/France/172002/2017 | A / H5N8 | Europe / France |
| EPI_ISL_4064685 | A/chicken/France/172003/2017 | A / H5N8 | Europe / France |
| EPI_ISL_4064686 | A/duck/France/172004/2017 | A / H5N8 | Europe / France |
| EPI_ISL_4064687 | A/duck/France/172012/2017 | A / H5N8 | Europe / France |
| EPI_ISL_4064688 | A/duck/France/172038/2017 | A / H5N8 | Europe / France |
| EPI_ISL_4064689 | A/duck/France/172057/2017 | A / H5N8 | Europe / France |
| EPI_ISL_4064690 | A/chicken/France/172092/2017 | A / H5N8 | Europe / France |
| EPI_ISL_4064691 | A/chicken/France/172093/2017 | A / H5N8 | Europe / France |
| EPI_ISL_4064692 | A/duck/France/172094/2017 | A / H5N8 | Europe / France |
| EPI_ISL_4064693 | A/duck/France/172096/2017 | A / H5N8 | Europe / France |
| EPI_ISL_4064694 | A/chicken/France/172137/2017 | A / H5N8 | Europe / France |
| EPI_ISL_4064695 | A/duck/France/172144/2017 | A / H5N8 | Europe / France |
| EPI_ISL_4064696 | A/duck/France/172182/2017 | A / H5N8 | Europe / France |
| EPI_ISL_4064697 | A/chicken/France/172287/2017 | A / H5N8 | Europe / France |
| EPI_ISL_4064698 | A/duck/France/172289/2017 | A / H5N8 | Europe / France |
| EPI_ISL_4064699 | A/duck/France/172331/2017 | A / H5N8 | Europe / France |
| EPI_ISL_4064700 | A/duck/France/172362/2017 | A / H5N8 | Europe / France |
| EPI_ISL_4064701 | A/chicken/France/172383/2017 | A / H5N8 | Europe / France |
| EPI_ISL_4064702 | A/duck/France/172384/2017 | A / H5N8 | Europe / France |
| EPI_ISL_4064703 | A/duck/France/172390/2017 | A / H5N8 | Europe / France |
| EPI_ISL_4064704 | A/duck/France/172394/2017 | A / H5N8 | Europe / France |
| EPI_ISL_4064705 | A/chicken/France/172610/2017 | A / H5N8 | Europe / France |
| EPI_ISL_4064707 | A/swan/France/171265/2017 | A / H5N8 | Europe / France |
| EPI_ISL_4064708 | A/gull/France/161176/2017 | A / H5N8 | Europe / France |

| | | | |
|---|---|---|---|
| EPI_ISL_4064709 | A/swan/France/171377/2017 | A / H5N8 | Europe / France |
| EPI_ISL_4064710 | A/swan/France/170496/2017 | A / H5N8 | Europe / France |
| EPI_ISL_4064711 | A/duck/France/171779/2017 | A / H5N8 | Europe / France |
| EPI_ISL_4064712 | A/swan/France/171960/2017 | A / H5N8 | Europe / France |
| EPI_ISL_4064713 | A/swan/France/170166/2017 | A / H5N8 | Europe / France |
| EPI_ISL_4064714 | A/goose/France/171953/2017 | A / H5N8 | Europe / France |
| EPI_ISL_4064715 | A/swan/France/171585/2017 | A / H5N8 | Europe / France |
| EPI_ISL_4064716 | A/swan/France/171376/2017 | A / H5N8 | Europe / France |
| EPI_ISL_4064717 | A/swan/France/171267/2017 | A / H5N8 | Europe / France |
| EPI_ISL_4069650 | A/chicken/Egypt/N16732/2019 | A / H5N8 | Africa / Egypt |
| EPI_ISL_4069651 | A/duck/Egypt/N16719/2019 | A / H5N8 | Africa / Egypt |
| EPI_ISL_4069652 | A/chicken/Egypt/Q16711B/2019 | A / H5N8 | Africa / Egypt |
| EPI_ISL_4069678 | A/chicken/Egypt/Q16710C/2019 | A / H5N8 | Africa / Egypt |
| EPI_ISL_4069710 | A/chicken/Egypt/Q16711A/2019 | A / H5N8 | Africa / Egypt |
| EPI_ISL_4069736 | A/duck/Egypt/Q16716A/2019 | A / H5N8 | Africa / Egypt |
| EPI_ISL_4069750 | A/duck/Egypt/N16720/2019 | A / H5N8 | Africa / Egypt |
| EPI_ISL_4069752 | A/chicken/Egypt/N16730/2019 | A / H5N8 | Africa / Egypt |
| EPI_ISL_4069756 | A/chicken/Egypt/Q16711C/2019 | A / H5N8 | Africa / Egypt |
| EPI_ISL_4069758 | A/duck/Egypt/N16717/2019 | A / H5N8 | Africa / Egypt |
| EPI_ISL_4069759 | A/chicken/Egypt/Q16712A/2019 | A / H5N8 | Africa / Egypt |
| EPI_ISL_4069762 | A/duck/Egypt/N16722/2019 | A / H5N8 | Africa / Egypt |
| EPI_ISL_4069765 | A/chicken/Egypt/Q16807E/2019 | A / H5N8 | Africa / Egypt |
| EPI_ISL_4069778 | A/duck/Egypt/A16793/2019 | A / H5N8 | Africa / Egypt |
| EPI_ISL_4070128 | A/duck/Denmark/19062-5Sp1c/2017 | A / H5N8 | Europe / Denmark |
| EPI_ISL_4070172 | A/goose/Denmark/1365-1p1c/2017 | A / H5N8 | Europe / Denmark |
| EPI_ISL_4070378 | A/pigeon/Egypt/A16805/2019 | A / H5N8 | Africa / Egypt |
| EPI_ISL_4070379 | A/chicken/Egypt/Q16710B/2019 | A / H5N8 | Africa / Egypt |
| EPI_ISL_4070380 | A/chicken/Egypt/Q16808B/2019 | A / H5N8 | Africa / Egypt |
| EPI_ISL_4070383 | A/chicken/Egypt/Q16807B/2019 | A / H5N8 | Africa / Egypt |
| EPI_ISL_4070608 | A/chicken/Egypt/Q16710A/2019 | A / H5N8 | Africa / Egypt |
| EPI_ISL_4070728 | A/pigeon/Egypt/A16804/2019 | A / H5N8 | Africa / Egypt |
| EPI_ISL_4070816 | A/chicken/Egypt/Q16807C/2019 | A / H5N8 | Africa / Egypt |
| EPI_ISL_4070847 | A/chicken/Egypt/F17230B/2019 | A / H5N8 | Africa / Egypt |
| EPI_ISL_4070896 | A/chicken/Egypt/F17230D/2019 | A / H5N8 | Africa / Egypt |
| EPI_ISL_4070932 | A/duck/Egypt/N16721/2019 | A / H5N8 | Africa / Egypt |
| EPI_ISL_4070933 | A/chicken/Egypt/F17229A/2019 | A / H5N8 | Africa / Egypt |
| EPI_ISL_4070935 | A/pigeon/Egypt/A16800/2019 | A / H5N8 | Africa / Egypt |
| EPI_ISL_4070936 | A/chicken/Egypt/F17229B/2019 | A / H5N8 | Africa / Egypt |
| EPI_ISL_4070940 | A/chicken/Egypt/F17230C/2019 | A / H5N8 | Africa / Egypt |
| EPI_ISL_4070945 | A/chicken/Egypt/F17230A/2019 | A / H5N8 | Africa / Egypt |
| EPI_ISL_4071019 | A/falcated duck/Kagoshima/KU-d3/2020 | A / H5N8 | Asia / Japan / Kagoshima |
| EPI_ISL_4071447 | A/environment/Bangladesh/42410/2020 | A / H5N6 | Asia / Bangladesh |
| EPI_ISL_4071515 | A/duck/Bangladesh/43127/2020 | A / H5N6 | Asia / Bangladesh |
| EPI_ISL_4071518 | A/environment/Bangladesh/42416/2020 | A / H5N6 | Asia / Bangladesh |
| EPI_ISL_4071595 | A/duck/Bangladesh/43050/2020 | A / H5N6 | Asia / Bangladesh |
| EPI_ISL_4071615 | A/duck/Bangladesh/43123/2020 | A / H5N6 | Asia / Bangladesh |
| EPI_ISL_4071616 | A/Ferruginous duck/Bangladesh/42380/2020 | A / H5N6 | Asia / Bangladesh |
| EPI_ISL_4071631 | A/duck/Bangladesh/43128/2020 | A / H5N6 | Asia / Bangladesh |
| EPI_ISL_4071634 | A/duck/Bangladesh/43082/2020 | A / H5N6 | Asia / Bangladesh |
| EPI_ISL_4071635 | A/duck/Bangladesh/43120/2020 | A / H5N6 | Asia / Bangladesh |
| EPI_ISL_4071638 | A/duck/Bangladesh/43099/2020 | A / H5N6 | Asia / Bangladesh |
| EPI_ISL_4071641 | A/duck/Bangladesh/43129/2020 | A / H5N6 | Asia / Bangladesh |
| EPI_ISL_4071723 | A/duck/Bangladesh/43122/2020 | A / H5N6 | Asia / Bangladesh |
| EPI_ISL_4071724 | A/Common pochard/Bangladesh/42386/2020 | A / H5N6 | Asia / Bangladesh |
| EPI_ISL_4071728 | A/duck/Bangladesh/43119/2020 | A / H5N6 | Asia / Bangladesh |
| EPI_ISL_4071929 | A/chicken/Zhejiang/6107/2016 | A / H5N6 | Asia / China / Zhejiang Province |
| EPI_ISL_4071930 | A/chicken/Zhejiang/13160/2016 | A / H5N6 | Asia / China / Zhejiang Province |
| EPI_ISL_4071931 | A/chicken/Zhejiang/528127/2016 | A / H5N6 | Asia / China / Zhejiang Province |
| EPI_ISL_4071932 | A/Muscovy duck/China/H5N6/2020 | A / H5N6 | Asia / China |
| EPI_ISL_4072009 | A/chicken/Zhejiang/102637/2016 | A / H5N6 | Asia / China / Zhejiang Province |
| EPI_ISL_4072173 | A/chicken/Zhejiang/1130131/2016 | A / H5N6 | Asia / China / Zhejiang Province |
| EPI_ISL_4073121 | A/duck/Bangladesh/44433/2020 | A / H5N6 | Asia / Bangladesh |
| EPI_ISL_4073122 | A/duck/Bangladesh/44442/2020 | A / H5N6 | Asia / Bangladesh |
| EPI_ISL_4073123 | A/duck/Bangladesh/44448/2020 | A / H5N6 | Asia / Bangladesh |
| EPI_ISL_4073127 | A/duck/Bangladesh/44423/2020 | A / H5N6 | Asia / Bangladesh |
| EPI_ISL_4073128 | A/duck/Bangladesh/44432/2020 | A / H5N6 | Asia / Bangladesh |
| EPI_ISL_4073129 | A/duck/Bangladesh/44440/2020 | A / H5N6 | Asia / Bangladesh |
| EPI_ISL_4073130 | A/duck/Bangladesh/44524/2020 | A / H5N6 | Asia / Bangladesh |
| EPI_ISL_4073131 | A/duck/Bangladesh/44453/2020 | A / H5N6 | Asia / Bangladesh |
| EPI_ISL_4073132 | A/duck/Bangladesh/44469/2020 | A / H5N6 | Asia / Bangladesh |
| EPI_ISL_4073133 | A/duck/Bangladesh/44424/2020 | A / H5N6 | Asia / Bangladesh |
| EPI_ISL_4073156 | A/duck/Bangladesh/44417/2020 | A / H5N6 | Asia / Bangladesh |

| | | | | |
|---|---|---|---|---|
| EPI_ISL_4073171 | A/duck/Bangladesh/44502/2020 | A / H5N6 | Asia / Bangladesh | |
| EPI_ISL_4073173 | A/duck/Bangladesh/44484/2020 | A / H5N6 | Asia / Bangladesh | |
| EPI_ISL_4073174 | A/duck/Bangladesh/44418/2020 | A / H5N6 | Asia / Bangladesh | |
| EPI_ISL_4073175 | A/duck/Bangladesh/44447/2020 | A / H5N6 | Asia / Bangladesh | |
| EPI_ISL_4073176 | A/duck/Bangladesh/44430/2020 | A / H5N6 | Asia / Bangladesh | |
| EPI_ISL_4073177 | A/duck/Bangladesh/44456/2020 | A / H5N6 | Asia / Bangladesh | |
| EPI_ISL_4073178 | A/duck/Bangladesh/44434/2020 | A / H5N6 | Asia / Bangladesh | |
| EPI_ISL_4073179 | A/duck/Bangladesh/44471/2020 | A / H5N6 | Asia / Bangladesh | |
| EPI_ISL_4073181 | A/duck/Bangladesh/44477/2020 | A / H5N6 | Asia / Bangladesh | |
| EPI_ISL_4073186 | A/duck/Bangladesh/44523/2020 | A / H5N6 | Asia / Bangladesh | |
| EPI_ISL_4073189 | A/duck/Bangladesh/44508/2020 | A / H5N6 | Asia / Bangladesh | |
| EPI_ISL_4073195 | A/duck/Bangladesh/44504/2020 | A / H5N6 | Asia / Bangladesh | |
| EPI_ISL_408528 | A/spot_billed_duck/Korea/H0027/2014 | A / H5N8 | Asia / Korea, Republic of | Yu-Na Lee (Animal and Plant Quarantine Agency (S-2158) / Avian Influenza Research & Diagnostic Division) |
| EPI_ISL_408530 | A/duck/Korea/H0032/2014 | A / H5N8 | Asia / Korea, Republic of | Yu-Na Lee (Animal and Plant Quarantine Agency (S-2158) / Avian Influenza Research & Diagnostic Division) |
| EPI_ISL_408531 | A/duck/Korea/H0035/2014 | A / H5N8 | Asia / Korea, Republic of | Yu-Na Lee (Animal and Plant Quarantine Agency (S-2158) / Avian Influenza Research & Diagnostic Division) |
| EPI_ISL_408532 | A/duck/Korea/H0075/2014 | A / H5N8 | Asia / Korea, Republic of | Yu-Na Lee (Animal and Plant Quarantine Agency (S-2158) / Avian Influenza Research & Diagnostic Division) |
| EPI_ISL_408533 | A/duck/Korea/H0078/2014 | A / H5N8 | Asia / Korea, Republic of | Yu-Na Lee (Animal and Plant Quarantine Agency (S-2158) / Avian Influenza Research & Diagnostic Division) |
| EPI_ISL_408534 | A/duck/Korea/H0085/2014 | A / H5N8 | Asia / Korea, Republic of | Yu-Na Lee (Animal and Plant Quarantine Agency (S-2158) / Avian Influenza Research & Diagnostic Division) |
| EPI_ISL_408535 | A/duck/Korea/H0089/2014 | A / H5N8 | Asia / Korea, Republic of | Yu-Na Lee (Animal and Plant Quarantine Agency (S-2158) / Avian Influenza Research & Diagnostic Division) |
| EPI_ISL_408536 | A/duck/Korea/H0090/2014 | A / H5N8 | Asia / Korea, Republic of | Yu-Na Lee (Animal and Plant Quarantine Agency (S-2158) / Avian Influenza Research & Diagnostic Division) |
| EPI_ISL_408537 | A/duck/Korea/H0106/2014 | A / H5N8 | Asia / Korea, Republic of | Yu-Na Lee (Animal and Plant Quarantine Agency (S-2158) / Avian Influenza Research & Diagnostic Division) |
| EPI_ISL_408538 | A/duck/Korea/H0108/2014 | A / H5N8 | Asia / Korea, Republic of | Yu-Na Lee (Animal and Plant Quarantine Agency (S-2158) / Avian Influenza Research & Diagnostic Division) |
| EPI_ISL_408539 | A/duck/Korea/H0109/2014 | A / H5N8 | Asia / Korea, Republic of | Yu-Na Lee (Animal and Plant Quarantine Agency (S-2158) / Avian Influenza Research & Diagnostic Division) |
| EPI_ISL_408540 | A/duck/Korea/H0110/2014 | A / H5N8 | Asia / Korea, Republic of | Yu-Na Lee (Animal and Plant Quarantine Agency (S-2158) / Avian Influenza Research & Diagnostic Division) |
| EPI_ISL_408574 | A/duck/Korea/H0111/2014 | A / H5N8 | Asia / Korea, Republic of | Yu-Na Lee (Animal and Plant Quarantine Agency (S-2158) / Avian Influenza Research & Diagnostic Division) |
| EPI_ISL_408575 | A/duck/Korea/H0112/2014 | A / H5N8 | Asia / Korea, Republic of | Yu-Na Lee (Animal and Plant Quarantine Agency (S-2158) / Avian Influenza Research & Diagnostic Division) |
| EPI_ISL_408576 | A/chicken/Korea/H0151/2014 | A / H5N8 | Asia / Korea, Republic of | Yu-Na Lee (Animal and Plant Quarantine Agency (S-2158) / Avian Influenza Research & Diagnostic Division) |
| EPI_ISL_408577 | A/duck/Korea/H0193/2014 | A / H5N8 | Asia / Korea, Republic of | Yu-Na Lee (Animal and Plant Quarantine Agency (S-2158) / Avian Influenza Research & Diagnostic Division) |
| EPI_ISL_408578 | A/duck/Korea/H0196/2014 | A / H5N8 | Asia / Korea, Republic of | Yu-Na Lee (Animal and Plant Quarantine Agency (S-2158) / Avian Influenza Research & Diagnostic Division) |
| EPI_ISL_408579 | A/baikal_teal/Korea/H0203/2014 | A / H5N8 | Asia / Korea, Republic of | Yu-Na Lee (Animal and Plant Quarantine Agency (S-2158) / Avian Influenza Research & Diagnostic Division) |
| EPI_ISL_408580 | A/duck/Korea/H0262/2014 | A / H5N8 | Asia / Korea, Republic of | Yu-Na Lee (Animal and Plant Quarantine Agency (S-2158) / Avian Influenza Research & Diagnostic Division) |
| EPI_ISL_408581 | A/duck/Korea/H0266/2014 | A / H5N8 | Asia / Korea, Republic of | Yu-Na Lee (Animal and Plant Quarantine Agency (S-2158) / Avian Influenza Research & Diagnostic Division) |
| EPI_ISL_408582 | A/duck/Korea/H0268/2014 | A / H5N8 | Asia / Korea, Republic of | Yu-Na Lee (Animal and Plant Quarantine Agency (S-2158) / Avian Influenza Research & Diagnostic Division) |
| EPI_ISL_408583 | A/duck/Korea/H0277/2014 | A / H5N8 | Asia / Korea, Republic of | Yu-Na Lee (Animal and Plant Quarantine Agency (S-2158) / Avian Influenza Research & Diagnostic Division) |
| EPI_ISL_408584 | A/Korean_native_chicken/Korea/H0300/2014 | A / H5N8 | Asia / Korea, Republic of | Yu-Na Lee (Animal and Plant Quarantine Agency (S-2158) / Avian Influenza Research & Diagnostic Division) |
| EPI_ISL_408585 | A/duck/Korea/H0308/2014 | A / H5N8 | Asia / Korea, Republic of | Yu-Na Lee (Animal and Plant Quarantine Agency (S-2158) / Avian Influenza Research & Diagnostic Division) |
| EPI_ISL_408586 | A/duck/Korea/H0309/2014 | A / H5N8 | Asia / Korea, Republic of | Yu-Na Lee (Animal and Plant Quarantine Agency (S-2158) / Avian Influenza Research & Diagnostic Division) |
| EPI_ISL_408587 | A/duck/Korea/H0329/2014 | A / H5N8 | Asia / Korea, Republic of | Yu-Na Lee (Animal and Plant Quarantine Agency (S-2158) / Avian Influenza Research & Diagnostic Division) |
| EPI_ISL_408588 | A/duck/Korea/H0330/2014 | A / H5N8 | Asia / Korea, Republic of | Yu-Na Lee (Animal and Plant Quarantine Agency (S-2158) / Avian Influenza Research & Diagnostic Division) |
| EPI_ISL_408590 | A/duck/Korea/H0351/2014 | A / H5N8 | Asia / Korea, Republic of | Yu-Na Lee (Animal and Plant Quarantine Agency (S-2158) / Avian Influenza Research & Diagnostic Division) |
| EPI_ISL_408591 | A/duck/Korea/H0356/2014 | A / H5N8 | Asia / Korea, Republic of | Yu-Na Lee (Animal and Plant Quarantine Agency (S-2158) / Avian Influenza Research & Diagnostic Division) |
| EPI_ISL_408592 | A/duck/Korea/H0357/2014 | A / H5N8 | Asia / Korea, Republic of | Yu-Na Lee (Animal and Plant Quarantine Agency (S-2158) / Avian Influenza Research & Diagnostic Division) |
| EPI_ISL_408593 | A/duck/Korea/H0361/2014 | A / H5N8 | Asia / Korea, Republic of | Yu-Na Lee (Animal and Plant Quarantine Agency (S-2158) / Avian Influenza Research & Diagnostic Division) |
| EPI_ISL_408594 | A/duck/Korea/H0362/2014 | A / H5N8 | Asia / Korea, Republic of | Yu-Na Lee (Animal and Plant Quarantine Agency (S-2158) / Avian Influenza Research & Diagnostic Division) |
| EPI_ISL_408595 | A/duck/Korea/H0390/2014 | A / H5N8 | Asia / Korea, Republic of | Yu-Na Lee (Animal and Plant Quarantine Agency (S-2158) / Avian Influenza Research & Diagnostic Division) |
| EPI_ISL_408596 | A/duck/Korea/H0397/2014 | A / H5N8 | Asia / Korea, Republic of | Yu-Na Lee (Animal and Plant Quarantine Agency (S-2158) / Avian Influenza Research & Diagnostic Division) |
| EPI_ISL_408597 | A/duck/Korea/H0398/2014 | A / H5N8 | Asia / Korea, Republic of | Yu-Na Lee (Animal and Plant Quarantine Agency (S-2158) / Avian Influenza Research & Diagnostic Division) |
| EPI_ISL_408598 | A/duck/Korea/H0400/2014 | A / H5N8 | Asia / Korea, Republic of | Yu-Na Lee (Animal and Plant Quarantine Agency (S-2158) / Avian Influenza Research & Diagnostic Division) |
| EPI_ISL_408599 | A/duck/Korea/H0414/2014 | A / H5N8 | Asia / Korea, Republic of | Yu-Na Lee (Animal and Plant Quarantine Agency (S-2158) / Avian Influenza Research & Diagnostic Division) |
| EPI_ISL_408600 | A/chicken/Korea/H0421/2014 | A / H5N8 | Asia / Korea, Republic of | Yu-Na Lee (Animal and Plant Quarantine Agency (S-2158) / Avian Influenza Research & Diagnostic Division) |
| EPI_ISL_408601 | A/chicken/Korea/H0422/2014 | A / H5N8 | Asia / Korea, Republic of | Yu-Na Lee (Animal and Plant Quarantine Agency (S-2158) / Avian Influenza Research & Diagnostic Division) |
| EPI_ISL_408602 | A/duck/Korea/H0425/2014 | A / H5N8 | Asia / Korea, Republic of | Yu-Na Lee (Animal and Plant Quarantine Agency (S-2158) / Avian Influenza Research & Diagnostic Division) |
| EPI_ISL_408603 | A/duck/Korea/H0426/2014 | A / H5N8 | Asia / Korea, Republic of | Yu-Na Lee (Animal and Plant Quarantine Agency (S-2158) / Avian Influenza Research & Diagnostic Division) |
| EPI_ISL_408604 | A/duck/Korea/H0428/2014 | A / H5N8 | Asia / Korea, Republic of | Yu-Na Lee (Animal and Plant Quarantine Agency (S-2158) / Avian Influenza Research & Diagnostic Division) |
| EPI_ISL_408605 | A/duck/Korea/H0429/2014 | A / H5N8 | Asia / Korea, Republic of | Yu-Na Lee (Animal and Plant Quarantine Agency (S-2158) / Avian Influenza Research & Diagnostic Division) |
| EPI_ISL_408606 | A/duck/Korea/H0430/2014 | A / H5N8 | Asia / Korea, Republic of | Yu-Na Lee (Animal and Plant Quarantine Agency (S-2158) / Avian Influenza Research & Diagnostic Division) |
| EPI_ISL_408607 | A/duck/Korea/H0431/2014 | A / H5N8 | Asia / Korea, Republic of | Yu-Na Lee (Animal and Plant Quarantine Agency (S-2158) / Avian Influenza Research & Diagnostic Division) |
| EPI_ISL_408608 | A/duck/Korea/H0439/2014 | A / H5N8 | Asia / Korea, Republic of | Yu-Na Lee (Animal and Plant Quarantine Agency (S-2158) / Avian Influenza Research & Diagnostic Division) |
| EPI_ISL_408609 | A/duck/Korea/H0445/2014 | A / H5N8 | Asia / Korea, Republic of | Yu-Na Lee (Animal and Plant Quarantine Agency (S-2158) / Avian Influenza Research & Diagnostic Division) |
| EPI_ISL_408610 | A/environment/Korea/H0469/2014 | A / H5N8 | Asia / Korea, Republic of | Yu-Na Lee (Animal and Plant Quarantine Agency (S-2158) / Avian Influenza Research & Diagnostic Division) |
| EPI_ISL_408611 | A/chicken/Korea/H0470/2014 | A / H5N8 | Asia / Korea, Republic of | Yu-Na Lee (Animal and Plant Quarantine Agency (S-2158) / Avian Influenza Research & Diagnostic Division) |
| EPI_ISL_408612 | A/duck/Korea/H0488/2014 | A / H5N8 | Asia / Korea, Republic of | Yu-Na Lee (Animal and Plant Quarantine Agency (S-2158) / Avian Influenza Research & Diagnostic Division) |
| EPI_ISL_408613 | A/spot_billed_duck/Korea/H0497/2014 | A / H5N8 | Asia / Korea, Republic of | Yu-Na Lee (Animal and Plant Quarantine Agency (S-2158) / Avian Influenza Research & Diagnostic Division) |
| EPI_ISL_408615 | A/duck/Korea/H0512/2014 | A / H5N8 | Asia / Korea, Republic of | Yu-Na Lee (Animal and Plant Quarantine Agency (S-2158) / Avian Influenza Research & Diagnostic Division) |
| EPI_ISL_408616 | A/duck/Korea/H0513/2014 | A / H5N8 | Asia / Korea, Republic of | Yu-Na Lee (Animal and Plant Quarantine Agency (S-2158) / Avian Influenza Research & Diagnostic Division) |
| EPI_ISL_408617 | A/duck/Korea/H0525/2014 | A / H5N8 | Asia / Korea, Republic of | Yu-Na Lee (Animal and Plant Quarantine Agency (S-2158) / Avian Influenza Research & Diagnostic Division) |
| EPI_ISL_408618 | A/chicken/Korea/H0526/2014 | A / H5N8 | Asia / Korea, Republic of | Yu-Na Lee (Animal and Plant Quarantine Agency (S-2158) / Avian Influenza Research & Diagnostic Division) |
| EPI_ISL_408619 | A/duck/Korea/H0527/2014 | A / H5N8 | Asia / Korea, Republic of | Yu-Na Lee (Animal and Plant Quarantine Agency (S-2158) / Avian Influenza Research & Diagnostic Division) |
| EPI_ISL_408620 | A/chicken/Korea/H0533/2014 | A / H5N8 | Asia / Korea, Republic of | Yu-Na Lee (Animal and Plant Quarantine Agency (S-2158) / Avian Influenza Research & Diagnostic Division) |
| EPI_ISL_408621 | A/duck/Korea/H0534/2014 | A / H5N8 | Asia / Korea, Republic of | Yu-Na Lee (Animal and Plant Quarantine Agency (S-2158) / Avian Influenza Research & Diagnostic Division) |
| EPI_ISL_408622 | A/duck/Korea/H0535/2014 | A / H5N8 | Asia / Korea, Republic of | Yu-Na Lee (Animal and Plant Quarantine Agency (S-2158) / Avian Influenza Research & Diagnostic Division) |
| EPI_ISL_408623 | A/duck/Korea/H0537/2014 | A / H5N8 | Asia / Korea, Republic of | Yu-Na Lee (Animal and Plant Quarantine Agency (S-2158) / Avian Influenza Research & Diagnostic Division) |

| | | | |
|---|---|---|---|
| EPI_ISL_408624 | A/duck/Korea/H0539/2014 | A / H5N8 Asia / Korea, Republic of | Yu-Na Lee (Animal and Plant Quarantine Agency (S-2158) / Avian Influenza Research & Diagnostic Division) |
| EPI_ISL_408625 | A/chicken/Korea/H0541/2014 | A / H5N8 Asia / Korea, Republic of | Yu-Na Lee (Animal and Plant Quarantine Agency (S-2158) / Avian Influenza Research & Diagnostic Division) |
| EPI_ISL_408626 | A/chicken/Korea/H0550/2014 | A / H5N8 Asia / Korea, Republic of | Yu-Na Lee (Animal and Plant Quarantine Agency (S-2158) / Avian Influenza Research & Diagnostic Division) |
| EPI_ISL_408627 | A/duck/Korea/H0551/2014 | A / H5N8 Asia / Korea, Republic of | Yu-Na Lee (Animal and Plant Quarantine Agency (S-2158) / Avian Influenza Research & Diagnostic Division) |
| EPI_ISL_408628 | A/mandarin_duck/Korea/H0555/2014 | A / H5N8 Asia / Korea, Republic of | Yu-Na Lee (Animal and Plant Quarantine Agency (S-2158) / Avian Influenza Research & Diagnostic Division) |
| EPI_ISL_408629 | A/duck/Korea/H0556/2014 | A / H5N8 Asia / Korea, Republic of | Yu-Na Lee (Animal and Plant Quarantine Agency (S-2158) / Avian Influenza Research & Diagnostic Division) |
| EPI_ISL_408630 | A/duck/Korea/H0557/2014 | A / H5N8 Asia / Korea, Republic of | Yu-Na Lee (Animal and Plant Quarantine Agency (S-2158) / Avian Influenza Research & Diagnostic Division) |
| EPI_ISL_408631 | A/duck/Korea/H0561/2014 | A / H5N8 Asia / Korea, Republic of | Yu-Na Lee (Animal and Plant Quarantine Agency (S-2158) / Avian Influenza Research & Diagnostic Division) |
| EPI_ISL_408633 | A/environment/Korea/H0576/2014 | A / H5N8 Asia / Korea, Republic of | Yu-Na Lee (Animal and Plant Quarantine Agency (S-2158) / Avian Influenza Research & Diagnostic Division) |
| EPI_ISL_408634 | A/environment/Korea/H0577/2014 | A / H5N8 Asia / Korea, Republic of | Yu-Na Lee (Animal and Plant Quarantine Agency (S-2158) / Avian Influenza Research & Diagnostic Division) |
| EPI_ISL_408635 | A/duck/Korea/H0580/2014 | A / H5N8 Asia / Korea, Republic of | Yu-Na Lee (Animal and Plant Quarantine Agency (S-2158) / Avian Influenza Research & Diagnostic Division) |
| EPI_ISL_408636 | A/duck/Korea/H0582/2014 | A / H5N8 Asia / Korea, Republic of | Yu-Na Lee (Animal and Plant Quarantine Agency (S-2158) / Avian Influenza Research & Diagnostic Division) |
| EPI_ISL_408637 | A/duck/Korea/H0583/2014 | A / H5N8 Asia / Korea, Republic of | Yu-Na Lee (Animal and Plant Quarantine Agency (S-2158) / Avian Influenza Research & Diagnostic Division) |
| EPI_ISL_408638 | A/duck/Korea/H0584/2014 | A / H5N8 Asia / Korea, Republic of | Yu-Na Lee (Animal and Plant Quarantine Agency (S-2158) / Avian Influenza Research & Diagnostic Division) |
| EPI_ISL_408639 | A/duck/Korea/H0585/2014 | A / H5N8 Asia / Korea, Republic of | Yu-Na Lee (Animal and Plant Quarantine Agency (S-2158) / Avian Influenza Research & Diagnostic Division) |
| EPI_ISL_408640 | A/duck/Korea/H0586/2014 | A / H5N8 Asia / Korea, Republic of | Yu-Na Lee (Animal and Plant Quarantine Agency (S-2158) / Avian Influenza Research & Diagnostic Division) |
| EPI_ISL_408641 | A/environment/Korea/H0591/2014 | A / H5N8 Asia / Korea, Republic of | Yu-Na Lee (Animal and Plant Quarantine Agency (S-2158) / Avian Influenza Research & Diagnostic Division) |
| EPI_ISL_408642 | A/duck/Korea/H0602/2014 | A / H5N8 Asia / Korea, Republic of | Yu-Na Lee (Animal and Plant Quarantine Agency (S-2158) / Avian Influenza Research & Diagnostic Division) |
| EPI_ISL_408643 | A/duck/Korea/H0603/2014 | A / H5N8 Asia / Korea, Republic of | Yu-Na Lee (Animal and Plant Quarantine Agency (S-2158) / Avian Influenza Research & Diagnostic Division) |
| EPI_ISL_408644 | A/duck/Korea/H0605/2014 | A / H5N8 Asia / Korea, Republic of | Yu-Na Lee (Animal and Plant Quarantine Agency (S-2158) / Avian Influenza Research & Diagnostic Division) |
| EPI_ISL_408645 | A/duck/Korea/H0606/2014 | A / H5N8 Asia / Korea, Republic of | Yu-Na Lee (Animal and Plant Quarantine Agency (S-2158) / Avian Influenza Research & Diagnostic Division) |
| EPI_ISL_408646 | A/duck/Korea/H0607/2014 | A / H5N8 Asia / Korea, Republic of | Yu-Na Lee (Animal and Plant Quarantine Agency (S-2158) / Avian Influenza Research & Diagnostic Division) |
| EPI_ISL_408647 | A/duck/Korea/H0609/2014 | A / H5N8 Asia / Korea, Republic of | Yu-Na Lee (Animal and Plant Quarantine Agency (S-2158) / Avian Influenza Research & Diagnostic Division) |
| EPI_ISL_408648 | A/duck/Korea/H0610/2014 | A / H5N8 Asia / Korea, Republic of | Yu-Na Lee (Animal and Plant Quarantine Agency (S-2158) / Avian Influenza Research & Diagnostic Division) |
| EPI_ISL_408649 | A/duck/Korea/H0611/2014 | A / H5N8 Asia / Korea, Republic of | Yu-Na Lee (Animal and Plant Quarantine Agency (S-2158) / Avian Influenza Research & Diagnostic Division) |
| EPI_ISL_408650 | A/duck/Korea/H0613/2014 | A / H5N8 Asia / Korea, Republic of | Yu-Na Lee (Animal and Plant Quarantine Agency (S-2158) / Avian Influenza Research & Diagnostic Division) |
| EPI_ISL_408651 | A/duck/Korea/H0614/2014 | A / H5N8 Asia / Korea, Republic of | Yu-Na Lee (Animal and Plant Quarantine Agency (S-2158) / Avian Influenza Research & Diagnostic Division) |
| EPI_ISL_408652 | A/duck/Korea/H0615/2014 | A / H5N8 Asia / Korea, Republic of | Yu-Na Lee (Animal and Plant Quarantine Agency (S-2158) / Avian Influenza Research & Diagnostic Division) |
| EPI_ISL_408653 | A/duck/Korea/H0616/2014 | A / H5N8 Asia / Korea, Republic of | Yu-Na Lee (Animal and Plant Quarantine Agency (S-2158) / Avian Influenza Research & Diagnostic Division) |
| EPI_ISL_408654 | A/duck/Korea/H0617/2014 | A / H5N8 Asia / Korea, Republic of | Yu-Na Lee (Animal and Plant Quarantine Agency (S-2158) / Avian Influenza Research & Diagnostic Division) |
| EPI_ISL_408655 | A/duck/Korea/H0618/2014 | A / H5N8 Asia / Korea, Republic of | Yu-Na Lee (Animal and Plant Quarantine Agency (S-2158) / Avian Influenza Research & Diagnostic Division) |
| EPI_ISL_408656 | A/duck/Korea/H0619/2014 | A / H5N8 Asia / Korea, Republic of | Yu-Na Lee (Animal and Plant Quarantine Agency (S-2158) / Avian Influenza Research & Diagnostic Division) |
| EPI_ISL_408657 | A/duck/Korea/H0620/2014 | A / H5N8 Asia / Korea, Republic of | Yu-Na Lee (Animal and Plant Quarantine Agency (S-2158) / Avian Influenza Research & Diagnostic Division) |
| EPI_ISL_408658 | A/duck/Korea/H0621/2014 | A / H5N8 Asia / Korea, Republic of | Yu-Na Lee (Animal and Plant Quarantine Agency (S-2158) / Avian Influenza Research & Diagnostic Division) |
| EPI_ISL_408659 | A/duck/Korea/H0622/2014 | A / H5N8 Asia / Korea, Republic of | Yu-Na Lee (Animal and Plant Quarantine Agency (S-2158) / Avian Influenza Research & Diagnostic Division) |
| EPI_ISL_408660 | A/duck/Korea/H0623/2014 | A / H5N8 Asia / Korea, Republic of | Yu-Na Lee (Animal and Plant Quarantine Agency (S-2158) / Avian Influenza Research & Diagnostic Division) |
| EPI_ISL_408661 | A/duck/Korea/H0624/2014 | A / H5N8 Asia / Korea, Republic of | Yu-Na Lee (Animal and Plant Quarantine Agency (S-2158) / Avian Influenza Research & Diagnostic Division) |
| EPI_ISL_408662 | A/duck/Korea/H0625/2014 | A / H5N8 Asia / Korea, Republic of | Yu-Na Lee (Animal and Plant Quarantine Agency (S-2158) / Avian Influenza Research & Diagnostic Division) |
| EPI_ISL_408663 | A/duck/Korea/H0626/2014 | A / H5N8 Asia / Korea, Republic of | Yu-Na Lee (Animal and Plant Quarantine Agency (S-2158) / Avian Influenza Research & Diagnostic Division) |
| EPI_ISL_408664 | A/duck/Korea/H0646/2014 | A / H5N8 Asia / Korea, Republic of | Yu-Na Lee (Animal and Plant Quarantine Agency (S-2158) / Avian Influenza Research & Diagnostic Division) |
| EPI_ISL_408673 | A/duck/Korea/H0652/2014 | A / H5N8 Asia / Korea, Republic of | Yu-Na Lee (Animal and Plant Quarantine Agency (S-2158) / Avian Influenza Research & Diagnostic Division) |
| EPI_ISL_408674 | A/duck/Korea/H0653/2014 | A / H5N8 Asia / Korea, Republic of | Yu-Na Lee (Animal and Plant Quarantine Agency (S-2158) / Avian Influenza Research & Diagnostic Division) |
| EPI_ISL_408675 | A/chicken/Korea/H0660/2014 | A / H5N8 Asia / Korea, Republic of | Yu-Na Lee (Animal and Plant Quarantine Agency (S-2158) / Avian Influenza Research & Diagnostic Division) |
| EPI_ISL_408676 | A/duck/Korea/H0663/2014 | A / H5N8 Asia / Korea, Republic of | Yu-Na Lee (Animal and Plant Quarantine Agency (S-2158) / Avian Influenza Research & Diagnostic Division) |
| EPI_ISL_408677 | A/duck/Korea/H0673/2014 | A / H5N8 Asia / Korea, Republic of | Yu-Na Lee (Animal and Plant Quarantine Agency (S-2158) / Avian Influenza Research & Diagnostic Division) |
| EPI_ISL_408678 | A/duck/Korea/H0674/2014 | A / H5N8 Asia / Korea, Republic of | Yu-Na Lee (Animal and Plant Quarantine Agency (S-2158) / Avian Influenza Research & Diagnostic Division) |
| EPI_ISL_408679 | A/duck/Korea/H0691/2014 | A / H5N8 Asia / Korea, Republic of | Yu-Na Lee (Animal and Plant Quarantine Agency (S-2158) / Avian Influenza Research & Diagnostic Division) |
| EPI_ISL_408680 | A/duck/Korea/H0692/2014 | A / H5N8 Asia / Korea, Republic of | Yu-Na Lee (Animal and Plant Quarantine Agency (S-2158) / Avian Influenza Research & Diagnostic Division) |
| EPI_ISL_408681 | A/duck/Korea/H0694/2014 | A / H5N8 Asia / Korea, Republic of | Yu-Na Lee (Animal and Plant Quarantine Agency (S-2158) / Avian Influenza Research & Diagnostic Division) |
| EPI_ISL_408682 | A/environment/Korea/H0701/2014 | A / H5N8 Asia / Korea, Republic of | Yu-Na Lee (Animal and Plant Quarantine Agency (S-2158) / Avian Influenza Research & Diagnostic Division) |
| EPI_ISL_408683 | A/duck/Korea/H0705/2014 | A / H5N8 Asia / Korea, Republic of | Yu-Na Lee (Animal and Plant Quarantine Agency (S-2158) / Avian Influenza Research & Diagnostic Division) |
| EPI_ISL_408684 | A/duck/Korea/H0708/2014 | A / H5N8 Asia / Korea, Republic of | Yu-Na Lee (Animal and Plant Quarantine Agency (S-2158) / Avian Influenza Research & Diagnostic Division) |
| EPI_ISL_408685 | A/chicken/Korea/H0709/2014 | A / H5N8 Asia / Korea, Republic of | Yu-Na Lee (Animal and Plant Quarantine Agency (S-2158) / Avian Influenza Research & Diagnostic Division) |
| EPI_ISL_408686 | A/duck/Korea/H0711/2014 | A / H5N8 Asia / Korea, Republic of | Yu-Na Lee (Animal and Plant Quarantine Agency (S-2158) / Avian Influenza Research & Diagnostic Division) |
| EPI_ISL_408687 | A/chicken/Korea/H0713/2014 | A / H5N8 Asia / Korea, Republic of | Yu-Na Lee (Animal and Plant Quarantine Agency (S-2158) / Avian Influenza Research & Diagnostic Division) |
| EPI_ISL_408688 | A/duck/Korea/H0716/2014 | A / H5N8 Asia / Korea, Republic of | Yu-Na Lee (Animal and Plant Quarantine Agency (S-2158) / Avian Influenza Research & Diagnostic Division) |
| EPI_ISL_408689 | A/duck/Korea/H0717/2014 | A / H5N8 Asia / Korea, Republic of | Yu-Na Lee (Animal and Plant Quarantine Agency (S-2158) / Avian Influenza Research & Diagnostic Division) |
| EPI_ISL_408690 | A/duck/Korea/H0731/2014 | A / H5N8 Asia / Korea, Republic of | Yu-Na Lee (Animal and Plant Quarantine Agency (S-2158) / Avian Influenza Research & Diagnostic Division) |
| EPI_ISL_408691 | A/backyard_poultry/Korea/H0734/2014 | A / H5N8 Asia / Korea, Republic of | Yu-Na Lee (Animal and Plant Quarantine Agency (S-2158) / Avian Influenza Research & Diagnostic Division) |
| EPI_ISL_408692 | A/duck/Korea/H0741/2014 | A / H5N8 Asia / Korea, Republic of | Yu-Na Lee (Animal and Plant Quarantine Agency (S-2158) / Avian Influenza Research & Diagnostic Division) |
| EPI_ISL_408693 | A/environment/Korea/H0745/2014 | A / H5N8 Asia / Korea, Republic of | Yu-Na Lee (Animal and Plant Quarantine Agency (S-2158) / Avian Influenza Research & Diagnostic Division) |
| EPI_ISL_408694 | A/environment/Korea/H0760/2014 | A / H5N8 Asia / Korea, Republic of | Yu-Na Lee (Animal and Plant Quarantine Agency (S-2158) / Avian Influenza Research & Diagnostic Division) |
| EPI_ISL_408695 | A/duck/Korea/H0761/2014 | A / H5N8 Asia / Korea, Republic of | Yu-Na Lee (Animal and Plant Quarantine Agency (S-2158) / Avian Influenza Research & Diagnostic Division) |
| EPI_ISL_408696 | A/duck/Korea/H0763/2014 | A / H5N8 Asia / Korea, Republic of | Yu-Na Lee (Animal and Plant Quarantine Agency (S-2158) / Avian Influenza Research & Diagnostic Division) |
| EPI_ISL_408697 | A/duck/Korea/H0768/2014 | A / H5N8 Asia / Korea, Republic of | Yu-Na Lee (Animal and Plant Quarantine Agency (S-2158) / Avian Influenza Research & Diagnostic Division) |
| EPI_ISL_408698 | A/Eurasian_coot/Korea/H0770/2014 | A / H5N8 Asia / Korea, Republic of | Yu-Na Lee (Animal and Plant Quarantine Agency (S-2158) / Avian Influenza Research & Diagnostic Division) |
| EPI_ISL_408699 | A/chicken/Korea/H0773/2014 | A / H5N8 Asia / Korea, Republic of | Yu-Na Lee (Animal and Plant Quarantine Agency (S-2158) / Avian Influenza Research & Diagnostic Division) |
| EPI_ISL_408700 | A/duck/Korea/H0787/2014 | A / H5N8 Asia / Korea, Republic of | Yu-Na Lee (Animal and Plant Quarantine Agency (S-2158) / Avian Influenza Research & Diagnostic Division) |
| EPI_ISL_408701 | A/backyard_poultry/Korea/H0790/2014 | A / H5N8 Asia / Korea, Republic of | Yu-Na Lee (Animal and Plant Quarantine Agency (S-2158) / Avian Influenza Research & Diagnostic Division) |
| EPI_ISL_408702 | A/chicken/Korea/H0795/2014 | A / H5N8 Asia / Korea, Republic of | Yu-Na Lee (Animal and Plant Quarantine Agency (S-2158) / Avian Influenza Research & Diagnostic Division) |
| EPI_ISL_408703 | A/duck/Korea/H0797/2014 | A / H5N8 Asia / Korea, Republic of | Yu-Na Lee (Animal and Plant Quarantine Agency (S-2158) / Avian Influenza Research & Diagnostic Division) |
| EPI_ISL_408704 | A/chicken/Korea/H0811/2014 | A / H5N8 Asia / Korea, Republic of | Yu-Na Lee (Animal and Plant Quarantine Agency (S-2158) / Avian Influenza Research & Diagnostic Division) |

| | | | | |
|---|---|---|---|---|
| EPI_ISL_408705 | A/duck/Korea/H0813/2014 | A / H5N8 | Asia / Korea, Republic of | Yu-Na Lee (Animal and Plant Quarantine Agency (S-2158) / Avian Influenza Research & Diagnostic Division) |
| EPI_ISL_408707 | A/duck/Korea/H0822/2014 | A / H5N8 | Asia / Korea, Republic of | Yu-Na Lee (Animal and Plant Quarantine Agency (S-2158) / Avian Influenza Research & Diagnostic Division) |
| EPI_ISL_408708 | A/duck/Korea/H0824/2014 | A / H5N8 | Asia / Korea, Republic of | Yu-Na Lee (Animal and Plant Quarantine Agency (S-2158) / Avian Influenza Research & Diagnostic Division) |
| EPI_ISL_408711 | A/environment/Korea/H0832/2014 | A / H5N8 | Asia / Korea, Republic of | Yu-Na Lee (Animal and Plant Quarantine Agency (S-2158) / Avian Influenza Research & Diagnostic Division) |
| EPI_ISL_408712 | A/chicken/Korea/H0835/2014 | A / H5N8 | Asia / Korea, Republic of | Yu-Na Lee (Animal and Plant Quarantine Agency (S-2158) / Avian Influenza Research & Diagnostic Division) |
| EPI_ISL_408713 | A/duck/Korea/H0839/2014 | A / H5N8 | Asia / Korea, Republic of | Yu-Na Lee (Animal and Plant Quarantine Agency (S-2158) / Avian Influenza Research & Diagnostic Division) |
| EPI_ISL_408714 | A/duck/Korea/H0845/2014 | A / H5N8 | Asia / Korea, Republic of | Yu-Na Lee (Animal and Plant Quarantine Agency (S-2158) / Avian Influenza Research & Diagnostic Division) |
| EPI_ISL_408715 | A/environment/Korea/H0853/2014 | A / H5N8 | Asia / Korea, Republic of | Yu-Na Lee (Animal and Plant Quarantine Agency (S-2158) / Avian Influenza Research & Diagnostic Division) |
| EPI_ISL_408716 | A/environment/Korea/H0867_2/2014 | A / H5N8 | Asia / Korea, Republic of | Yu-Na Lee (Animal and Plant Quarantine Agency (S-2158) / Avian Influenza Research & Diagnostic Division) |
| EPI_ISL_408717 | A/duck/Korea/H0879/2014 | A / H5N8 | Asia / Korea, Republic of | Yu-Na Lee (Animal and Plant Quarantine Agency (S-2158) / Avian Influenza Research & Diagnostic Division) |
| EPI_ISL_408719 | A/duck/Korea/H0882/2014 | A / H5N8 | Asia / Korea, Republic of | Yu-Na Lee (Animal and Plant Quarantine Agency (S-2158) / Avian Influenza Research & Diagnostic Division) |
| EPI_ISL_408720 | A/chicken/Korea/H0884/2014 | A / H5N8 | Asia / Korea, Republic of | Yu-Na Lee (Animal and Plant Quarantine Agency (S-2158) / Avian Influenza Research & Diagnostic Division) |
| EPI_ISL_408721 | A/duck/Korea/H0887/2014 | A / H5N8 | Asia / Korea, Republic of | Yu-Na Lee (Animal and Plant Quarantine Agency (S-2158) / Avian Influenza Research & Diagnostic Division) |
| EPI_ISL_408722 | A/duck/Korea/H0888/2014 | A / H5N8 | Asia / Korea, Republic of | Yu-Na Lee (Animal and Plant Quarantine Agency (S-2158) / Avian Influenza Research & Diagnostic Division) |
| EPI_ISL_408723 | A/chicken/Korea/H0891/2014 | A / H5N8 | Asia / Korea, Republic of | Yu-Na Lee (Animal and Plant Quarantine Agency (S-2158) / Avian Influenza Research & Diagnostic Division) |
| EPI_ISL_408724 | A/chicken/Korea/H0893/2014 | A / H5N8 | Asia / Korea, Republic of | Yu-Na Lee (Animal and Plant Quarantine Agency (S-2158) / Avian Influenza Research & Diagnostic Division) |
| EPI_ISL_408725 | A/chicken/Korea/H0898/2014 | A / H5N8 | Asia / Korea, Republic of | Yu-Na Lee (Animal and Plant Quarantine Agency (S-2158) / Avian Influenza Research & Diagnostic Division) |
| EPI_ISL_408726 | A/duck/Korea/H0901/2014 | A / H5N8 | Asia / Korea, Republic of | Yu-Na Lee (Animal and Plant Quarantine Agency (S-2158) / Avian Influenza Research & Diagnostic Division) |
| EPI_ISL_408727 | A/chicken/Korea/H0905/2014 | A / H5N8 | Asia / Korea, Republic of | Yu-Na Lee (Animal and Plant Quarantine Agency (S-2158) / Avian Influenza Research & Diagnostic Division) |
| EPI_ISL_408728 | A/great_egret/Korea/H0912/2014 | A / H5N8 | Asia / Korea, Republic of | Yu-Na Lee (Animal and Plant Quarantine Agency (S-2158) / Avian Influenza Research & Diagnostic Division) |
| EPI_ISL_408729 | A/bean_goose/Korea/H0924/2014 | A / H5N8 | Asia / Korea, Republic of | Yu-Na Lee (Animal and Plant Quarantine Agency (S-2158) / Avian Influenza Research & Diagnostic Division) |
| EPI_ISL_408730 | A/duck/Korea/H0936/2014 | A / H5N8 | Asia / Korea, Republic of | Yu-Na Lee (Animal and Plant Quarantine Agency (S-2158) / Avian Influenza Research & Diagnostic Division) |
| EPI_ISL_408731 | A/chicken/Korea/H0940/2014 | A / H5N8 | Asia / Korea, Republic of | Yu-Na Lee (Animal and Plant Quarantine Agency (S-2158) / Avian Influenza Research & Diagnostic Division) |
| EPI_ISL_408732 | A/chicken/Korea/H0950/2014 | A / H5N8 | Asia / Korea, Republic of | Yu-Na Lee (Animal and Plant Quarantine Agency (S-2158) / Avian Influenza Research & Diagnostic Division) |
| EPI_ISL_408733 | A/chicken/Korea/H0952/2014 | A / H5N8 | Asia / Korea, Republic of | Yu-Na Lee (Animal and Plant Quarantine Agency (S-2158) / Avian Influenza Research & Diagnostic Division) |
| EPI_ISL_408734 | A/chicken/Korea/H0953/2014 | A / H5N8 | Asia / Korea, Republic of | Yu-Na Lee (Animal and Plant Quarantine Agency (S-2158) / Avian Influenza Research & Diagnostic Division) |
| EPI_ISL_408735 | A/duck/Korea/H0958/2014 | A / H5N8 | Asia / Korea, Republic of | Yu-Na Lee (Animal and Plant Quarantine Agency (S-2158) / Avian Influenza Research & Diagnostic Division) |
| EPI_ISL_408737 | A/chicken/Korea/H1008/2014 | A / H5N8 | Asia / Korea, Republic of | Yu-Na Lee (Animal and Plant Quarantine Agency (S-2158) / Avian Influenza Research & Diagnostic Division) |
| EPI_ISL_408738 | A/duck/Korea/H1027/2014 | A / H5N8 | Asia / Korea, Republic of | Yu-Na Lee (Animal and Plant Quarantine Agency (S-2158) / Avian Influenza Research & Diagnostic Division) |
| EPI_ISL_408739 | A/duck/Korea/H1030/2014 | A / H5N8 | Asia / Korea, Republic of | Yu-Na Lee (Animal and Plant Quarantine Agency (S-2158) / Avian Influenza Research & Diagnostic Division) |
| EPI_ISL_408740 | A/duck/Korea/H1031/2014 | A / H5N8 | Asia / Korea, Republic of | Yu-Na Lee (Animal and Plant Quarantine Agency (S-2158) / Avian Influenza Research & Diagnostic Division) |
| EPI_ISL_408741 | A/duck/Korea/H1032/2014 | A / H5N8 | Asia / Korea, Republic of | Yu-Na Lee (Animal and Plant Quarantine Agency (S-2158) / Avian Influenza Research & Diagnostic Division) |
| EPI_ISL_408742 | A/duck/Korea/H1033/2014 | A / H5N8 | Asia / Korea, Republic of | Yu-Na Lee (Animal and Plant Quarantine Agency (S-2158) / Avian Influenza Research & Diagnostic Division) |
| EPI_ISL_408743 | A/duck/Korea/H1037/2014 | A / H5N8 | Asia / Korea, Republic of | Yu-Na Lee (Animal and Plant Quarantine Agency (S-2158) / Avian Influenza Research & Diagnostic Division) |
| EPI_ISL_408744 | A/duck/Korea/H1041/2014 | A / H5N8 | Asia / Korea, Republic of | Yu-Na Lee (Animal and Plant Quarantine Agency (S-2158) / Avian Influenza Research & Diagnostic Division) |
| EPI_ISL_408745 | A/duck/Korea/H1056/2014 | A / H5N8 | Asia / Korea, Republic of | Yu-Na Lee (Animal and Plant Quarantine Agency (S-2158) / Avian Influenza Research & Diagnostic Division) |
| EPI_ISL_408746 | A/duck/Korea/H1062/2014 | A / H5N8 | Asia / Korea, Republic of | Yu-Na Lee (Animal and Plant Quarantine Agency (S-2158) / Avian Influenza Research & Diagnostic Division) |
| EPI_ISL_408747 | A/duck/Korea/H1067/2014 | A / H5N8 | Asia / Korea, Republic of | Yu-Na Lee (Animal and Plant Quarantine Agency (S-2158) / Avian Influenza Research & Diagnostic Division) |
| EPI_ISL_408749 | A/duck/Korea/H1072/2014 | A / H5N8 | Asia / Korea, Republic of | Yu-Na Lee (Animal and Plant Quarantine Agency (S-2158) / Avian Influenza Research & Diagnostic Division) |
| EPI_ISL_408750 | A/chicken/Korea/H1078/2014 | A / H5N8 | Asia / Korea, Republic of | Yu-Na Lee (Animal and Plant Quarantine Agency (S-2158) / Avian Influenza Research & Diagnostic Division) |
| EPI_ISL_408751 | A/duck/Korea/H1111/2014 | A / H5N8 | Asia / Korea, Republic of | Yu-Na Lee (Animal and Plant Quarantine Agency (S-2158) / Avian Influenza Research & Diagnostic Division) |
| EPI_ISL_408752 | A/duck/Korea/H1112/2014 | A / H5N8 | Asia / Korea, Republic of | Yu-Na Lee (Animal and Plant Quarantine Agency (S-2158) / Avian Influenza Research & Diagnostic Division) |
| EPI_ISL_408753 | A/chicken/Korea/H1121/2014 | A / H5N8 | Asia / Korea, Republic of | Yu-Na Lee (Animal and Plant Quarantine Agency (S-2158) / Avian Influenza Research & Diagnostic Division) |
| EPI_ISL_408755 | A/chicken/Korea/H1152/2014 | A / H5N8 | Asia / Korea, Republic of | Yu-Na Lee (Animal and Plant Quarantine Agency (S-2158) / Avian Influenza Research & Diagnostic Division) |
| EPI_ISL_408756 | A/duck/Korea/H1161/2014 | A / H5N8 | Asia / Korea, Republic of | Yu-Na Lee (Animal and Plant Quarantine Agency (S-2158) / Avian Influenza Research & Diagnostic Division) |
| EPI_ISL_408757 | A/duck/Korea/H1173/2014 | A / H5N8 | Asia / Korea, Republic of | Yu-Na Lee (Animal and Plant Quarantine Agency (S-2158) / Avian Influenza Research & Diagnostic Division) |
| EPI_ISL_408758 | A/duck/Korea/H1174/2014 | A / H5N8 | Asia / Korea, Republic of | Yu-Na Lee (Animal and Plant Quarantine Agency (S-2158) / Avian Influenza Research & Diagnostic Division) |
| EPI_ISL_408759 | A/chicken/Korea/H1202/2014 | A / H5N8 | Asia / Korea, Republic of | Yu-Na Lee (Animal and Plant Quarantine Agency (S-2158) / Avian Influenza Research & Diagnostic Division) |
| EPI_ISL_408760 | A/chicken/Korea/H1205/2014 | A / H5N8 | Asia / Korea, Republic of | Yu-Na Lee (Animal and Plant Quarantine Agency (S-2158) / Avian Influenza Research & Diagnostic Division) |
| EPI_ISL_408761 | A/duck/Korea/H1219/2014 | A / H5N8 | Asia / Korea, Republic of | Yu-Na Lee (Animal and Plant Quarantine Agency (S-2158) / Avian Influenza Research & Diagnostic Division) |
| EPI_ISL_408762 | A/environment/Korea/H1220/2014 | A / H5N8 | Asia / Korea, Republic of | Yu-Na Lee (Animal and Plant Quarantine Agency (S-2158) / Avian Influenza Research & Diagnostic Division) |
| EPI_ISL_408763 | A/chicken/Korea/H1230/2014 | A / H5N8 | Asia / Korea, Republic of | Yu-Na Lee (Animal and Plant Quarantine Agency (S-2158) / Avian Influenza Research & Diagnostic Division) |
| EPI_ISL_408764 | A/duck/Korea/H1267/2014 | A / H5N8 | Asia / Korea, Republic of | Yu-Na Lee (Animal and Plant Quarantine Agency (S-2158) / Avian Influenza Research & Diagnostic Division) |
| EPI_ISL_408765 | A/duck/Korea/H1275/2014 | A / H5N8 | Asia / Korea, Republic of | Yu-Na Lee (Animal and Plant Quarantine Agency (S-2158) / Avian Influenza Research & Diagnostic Division) |
| EPI_ISL_408766 | A/chicken/Korea/H1289/2014 | A / H5N8 | Asia / Korea, Republic of | Yu-Na Lee (Animal and Plant Quarantine Agency (S-2158) / Avian Influenza Research & Diagnostic Division) |
| EPI_ISL_408767 | A/chicken/Korea/H1250/2014 | A / H5N8 | Asia / Korea, Republic of | Yu-Na Lee (Animal and Plant Quarantine Agency (S-2158) / Avian Influenza Research & Diagnostic Division) |
| EPI_ISL_408768 | A/environment/Korea/H1247/2014 | A / H5N8 | Asia / Korea, Republic of | Yu-Na Lee (Animal and Plant Quarantine Agency (S-2158) / Avian Influenza Research & Diagnostic Division) |
| EPI_ISL_408770 | A/quail/Korea/H1329/2014 | A / H5N8 | Asia / Korea, Republic of | Yu-Na Lee (Animal and Plant Quarantine Agency (S-2158) / Avian Influenza Research & Diagnostic Division) |
| EPI_ISL_408779 | A/duck/Korea/H1597/2014 | A / H5N8 | Asia / Korea, Republic of | Yu-Na Lee (Animal and Plant Quarantine Agency (S-2158) / Avian Influenza Research & Diagnostic Division) |
| EPI_ISL_408780 | A/duck/Korea/H1613/2014 | A / H5N8 | Asia / Korea, Republic of | Yu-Na Lee (Animal and Plant Quarantine Agency (S-2158) / Avian Influenza Research & Diagnostic Division) |
| EPI_ISL_408786 | A/duck/Korea/H1736/2014 | A / H5N8 | Asia / Korea, Republic of | Yu-Na Lee (Animal and Plant Quarantine Agency (S-2158) / Avian Influenza Research & Diagnostic Division) |
| EPI_ISL_408788 | A/duck/Korea/H1740/2014 | A / H5N8 | Asia / Korea, Republic of | Yu-Na Lee (Animal and Plant Quarantine Agency (S-2158) / Avian Influenza Research & Diagnostic Division) |
| EPI_ISL_408789 | A/duck/Korea/H1742/2014 | A / H5N8 | Asia / Korea, Republic of | Yu-Na Lee (Animal and Plant Quarantine Agency (S-2158) / Avian Influenza Research & Diagnostic Division) |
| EPI_ISL_408794 | A/duck/Korea/H1757/2014 | A / H5N8 | Asia / Korea, Republic of | Yu-Na Lee (Animal and Plant Quarantine Agency (S-2158) / Avian Influenza Research & Diagnostic Division) |
| EPI_ISL_408795 | A/duck/Korea/H1759/2014 | A / H5N8 | Asia / Korea, Republic of | Yu-Na Lee (Animal and Plant Quarantine Agency (S-2158) / Avian Influenza Research & Diagnostic Division) |
| EPI_ISL_408796 | A/duck/Korea/H1760/2014 | A / H5N8 | Asia / Korea, Republic of | Yu-Na Lee (Animal and Plant Quarantine Agency (S-2158) / Avian Influenza Research & Diagnostic Division) |
| EPI_ISL_408797 | A/duck/Korea/H1764/2014 | A / H5N8 | Asia / Korea, Republic of | Yu-Na Lee (Animal and Plant Quarantine Agency (S-2158) / Avian Influenza Research & Diagnostic Division) |
| EPI_ISL_408798 | A/duck/Korea/H1765/2014 | A / H5N8 | Asia / Korea, Republic of | Yu-Na Lee (Animal and Plant Quarantine Agency (S-2158) / Avian Influenza Research & Diagnostic Division) |
| EPI_ISL_408799 | A/duck/Korea/H1780/2014 | A / H5N8 | Asia / Korea, Republic of | Yu-Na Lee (Animal and Plant Quarantine Agency (S-2158) / Avian Influenza Research & Diagnostic Division) |
| EPI_ISL_408800 | A/duck/Korea/H1798/2014 | A / H5N8 | Asia / Korea, Republic of | Yu-Na Lee (Animal and Plant Quarantine Agency (S-2158) / Avian Influenza Research & Diagnostic Division) |
| EPI_ISL_408802 | A/Korean_native_chicken/Korea/H1804/2014 | A / H5N8 | Asia / Korea, Republic of | Yu-Na Lee (Animal and Plant Quarantine Agency (S-2158) / Avian Influenza Research & Diagnostic Division) |
| EPI_ISL_408803 | A/duck/Korea/H1806/2014 | A / H5N8 | Asia / Korea, Republic of | Yu-Na Lee (Animal and Plant Quarantine Agency (S-2158) / Avian Influenza Research & Diagnostic Division) |

| | | | | |
|---|---|---|---|---|
| EPI_ISL_408804 | A/duck/Korea/H1813/2014 | A / H5N8 | Asia / Korea, Republic of | Yu-Na Lee (Animal and Plant Quarantine Agency (S-2158) / Avian Influenza Research & Diagnostic Division) |
| EPI_ISL_408809 | A/duck/Korea/H1880/2014 | A / H5N8 | Asia / Korea, Republic of | Yu-Na Lee (Animal and Plant Quarantine Agency (S-2158) / Avian Influenza Research & Diagnostic Division) |
| EPI_ISL_408811 | A/Korean_native_chicken/Korea/H1964/2014 | A / H5N8 | Asia / Korea, Republic of | Yu-Na Lee (Animal and Plant Quarantine Agency (S-2158) / Avian Influenza Research & Diagnostic Division) |
| EPI_ISL_408812 | A/Korean_native_chicken/Korea/H2028/2014 | A / H5N8 | Asia / Korea, Republic of | Yu-Na Lee (Animal and Plant Quarantine Agency (S-2158) / Avian Influenza Research & Diagnostic Division) |
| EPI_ISL_408813 | A/Korean_native_chicken/Korea/H2031/2014 | A / H5N8 | Asia / Korea, Republic of | Yu-Na Lee (Animal and Plant Quarantine Agency (S-2158) / Avian Influenza Research & Diagnostic Division) |
| EPI_ISL_408816 | A/Korean_native_chicken/Korea/H2036/2014 | A / H5N8 | Asia / Korea, Republic of | Yu-Na Lee (Animal and Plant Quarantine Agency (S-2158) / Avian Influenza Research & Diagnostic Division) |
| EPI_ISL_408817 | A/Korean_native_chicken/Korea/H2037/2014 | A / H5N8 | Asia / Korea, Republic of | Yu-Na Lee (Animal and Plant Quarantine Agency (S-2158) / Avian Influenza Research & Diagnostic Division) |
| EPI_ISL_408818 | A/Korean_native_chicken/Korea/H2038/2014 | A / H5N8 | Asia / Korea, Republic of | Yu-Na Lee (Animal and Plant Quarantine Agency (S-2158) / Avian Influenza Research & Diagnostic Division) |
| EPI_ISL_408819 | A/Korean_native_chicken/Korea/H2039/2014 | A / H5N8 | Asia / Korea, Republic of | Yu-Na Lee (Animal and Plant Quarantine Agency (S-2158) / Avian Influenza Research & Diagnostic Division) |
| EPI_ISL_408820 | A/duck/Korea/H2075/2015 | A / H5N8 | Asia / Korea, Republic of | Yu-Na Lee (Animal and Plant Quarantine Agency (S-2158) / Avian Influenza Research & Diagnostic Division) |
| EPI_ISL_408821 | A/environment/Korea/H2081/2015 | A / H5N8 | Asia / Korea, Republic of | Yu-Na Lee (Animal and Plant Quarantine Agency (S-2158) / Avian Influenza Research & Diagnostic Division) |
| EPI_ISL_408823 | A/duck/Korea/H2088/2015 | A / H5N8 | Asia / Korea, Republic of | Yu-Na Lee (Animal and Plant Quarantine Agency (S-2158) / Avian Influenza Research & Diagnostic Division) |
| EPI_ISL_408824 | A/environment/Korea/H2093/2015 | A / H5N8 | Asia / Korea, Republic of | Yu-Na Lee (Animal and Plant Quarantine Agency (S-2158) / Avian Influenza Research & Diagnostic Division) |
| EPI_ISL_408825 | A/goose/Korea/H2095/2015 | A / H5N8 | Asia / Korea, Republic of | Yu-Na Lee (Animal and Plant Quarantine Agency (S-2158) / Avian Influenza Research & Diagnostic Division) |
| EPI_ISL_408826 | A/Korean_native_chicken/Korea/H2098/2015 | A / H5N8 | Asia / Korea, Republic of | Yu-Na Lee (Animal and Plant Quarantine Agency (S-2158) / Avian Influenza Research & Diagnostic Division) |
| EPI_ISL_408827 | A/duck/Korea/H2100/2015 | A / H5N8 | Asia / Korea, Republic of | Yu-Na Lee (Animal and Plant Quarantine Agency (S-2158) / Avian Influenza Research & Diagnostic Division) |
| EPI_ISL_408828 | A/chicken/Korea/H2103/2015 | A / H5N8 | Asia / Korea, Republic of | Yu-Na Lee (Animal and Plant Quarantine Agency (S-2158) / Avian Influenza Research & Diagnostic Division) |
| EPI_ISL_408829 | A/duck/Korea/H2104/2015 | A / H5N8 | Asia / Korea, Republic of | Yu-Na Lee (Animal and Plant Quarantine Agency (S-2158) / Avian Influenza Research & Diagnostic Division) |
| EPI_ISL_408831 | A/duck/Korea/H2117/2015 | A / H5N8 | Asia / Korea, Republic of | Yu-Na Lee (Animal and Plant Quarantine Agency (S-2158) / Avian Influenza Research & Diagnostic Division) |
| EPI_ISL_408832 | A/duck/Korea/H2118/2015 | A / H5N8 | Asia / Korea, Republic of | Yu-Na Lee (Animal and Plant Quarantine Agency (S-2158) / Avian Influenza Research & Diagnostic Division) |
| EPI_ISL_408833 | A/grey_heron/Korea/H2119/2015 | A / H5N8 | Asia / Korea, Republic of | Yu-Na Lee (Animal and Plant Quarantine Agency (S-2158) / Avian Influenza Research & Diagnostic Division) |
| EPI_ISL_408834 | A/spot_billed_duck/Korea/H2126/2015 | A / H5N8 | Asia / Korea, Republic of | Yu-Na Lee (Animal and Plant Quarantine Agency (S-2158) / Avian Influenza Research & Diagnostic Division) |
| EPI_ISL_408835 | A/duck/Korea/H2131/2015 | A / H5N8 | Asia / Korea, Republic of | Yu-Na Lee (Animal and Plant Quarantine Agency (S-2158) / Avian Influenza Research & Diagnostic Division) |
| EPI_ISL_408836 | A/duck/Korea/H2147/2015 | A / H5N8 | Asia / Korea, Republic of | Yu-Na Lee (Animal and Plant Quarantine Agency (S-2158) / Avian Influenza Research & Diagnostic Division) |
| EPI_ISL_408837 | A/duck/Korea/H2148/2015 | A / H5N8 | Asia / Korea, Republic of | Yu-Na Lee (Animal and Plant Quarantine Agency (S-2158) / Avian Influenza Research & Diagnostic Division) |
| EPI_ISL_408838 | A/duck/Korea/H2161/2015 | A / H5N8 | Asia / Korea, Republic of | Yu-Na Lee (Animal and Plant Quarantine Agency (S-2158) / Avian Influenza Research & Diagnostic Division) |
| EPI_ISL_408839 | A/chicken/Korea/H2181/2015 | A / H5N8 | Asia / Korea, Republic of | Yu-Na Lee (Animal and Plant Quarantine Agency (S-2158) / Avian Influenza Research & Diagnostic Division) |
| EPI_ISL_408840 | A/duck/Korea/H2190/2015 | A / H5N8 | Asia / Korea, Republic of | Yu-Na Lee (Animal and Plant Quarantine Agency (S-2158) / Avian Influenza Research & Diagnostic Division) |
| EPI_ISL_408841 | A/chicken/Korea/H2192/2015 | A / H5N8 | Asia / Korea, Republic of | Yu-Na Lee (Animal and Plant Quarantine Agency (S-2158) / Avian Influenza Research & Diagnostic Division) |
| EPI_ISL_408843 | A/chicken/Korea/H2205/2015 | A / H5N8 | Asia / Korea, Republic of | Yu-Na Lee (Animal and Plant Quarantine Agency (S-2158) / Avian Influenza Research & Diagnostic Division) |
| EPI_ISL_408844 | A/duck/Korea/H2206/2015 | A / H5N8 | Asia / Korea, Republic of | Yu-Na Lee (Animal and Plant Quarantine Agency (S-2158) / Avian Influenza Research & Diagnostic Division) |
| EPI_ISL_408845 | A/gadwall/Korea/H2209/2015 | A / H5N8 | Asia / Korea, Republic of | Yu-Na Lee (Animal and Plant Quarantine Agency (S-2158) / Avian Influenza Research & Diagnostic Division) |
| EPI_ISL_408846 | A/Eurasian_wigeon/Korea/H2210/2015 | A / H5N8 | Asia / Korea, Republic of | Yu-Na Lee (Animal and Plant Quarantine Agency (S-2158) / Avian Influenza Research & Diagnostic Division) |
| EPI_ISL_408847 | A/bean_goose/Korea/H2211/2015 | A / H5N8 | Asia / Korea, Republic of | Yu-Na Lee (Animal and Plant Quarantine Agency (S-2158) / Avian Influenza Research & Diagnostic Division) |
| EPI_ISL_408848 | A/duck/Korea/H2230/2015 | A / H5N8 | Asia / Korea, Republic of | Yu-Na Lee (Animal and Plant Quarantine Agency (S-2158) / Avian Influenza Research & Diagnostic Division) |
| EPI_ISL_408849 | A/duck/Korea/H2239/2015 | A / H5N8 | Asia / Korea, Republic of | Yu-Na Lee (Animal and Plant Quarantine Agency (S-2158) / Avian Influenza Research & Diagnostic Division) |
| EPI_ISL_408850 | A/duck/Korea/H2253/2015 | A / H5N8 | Asia / Korea, Republic of | Yu-Na Lee (Animal and Plant Quarantine Agency (S-2158) / Avian Influenza Research & Diagnostic Division) |
| EPI_ISL_408851 | A/common_teal/Korea/H2261/2015 | A / H5N8 | Asia / Korea, Republic of | Yu-Na Lee (Animal and Plant Quarantine Agency (S-2158) / Avian Influenza Research & Diagnostic Division) |
| EPI_ISL_408852 | A/duck/Korea/H2263/2015 | A / H5N8 | Asia / Korea, Republic of | Yu-Na Lee (Animal and Plant Quarantine Agency (S-2158) / Avian Influenza Research & Diagnostic Division) |
| EPI_ISL_408853 | A/duck/Korea/H2264/2015 | A / H5N8 | Asia / Korea, Republic of | Yu-Na Lee (Animal and Plant Quarantine Agency (S-2158) / Avian Influenza Research & Diagnostic Division) |
| EPI_ISL_408855 | A/duck/Korea/H2267/2015 | A / H5N8 | Asia / Korea, Republic of | Yu-Na Lee (Animal and Plant Quarantine Agency (S-2158) / Avian Influenza Research & Diagnostic Division) |
| EPI_ISL_408856 | A/duck/Korea/H2268/2015 | A / H5N8 | Asia / Korea, Republic of | Yu-Na Lee (Animal and Plant Quarantine Agency (S-2158) / Avian Influenza Research & Diagnostic Division) |
| EPI_ISL_408857 | A/duck/Korea/H2269/2015 | A / H5N8 | Asia / Korea, Republic of | Yu-Na Lee (Animal and Plant Quarantine Agency (S-2158) / Avian Influenza Research & Diagnostic Division) |
| EPI_ISL_408858 | A/duck/Korea/H2273/2015 | A / H5N8 | Asia / Korea, Republic of | Yu-Na Lee (Animal and Plant Quarantine Agency (S-2158) / Avian Influenza Research & Diagnostic Division) |
| EPI_ISL_408860 | A/duck/Korea/H2279/2015 | A / H5N8 | Asia / Korea, Republic of | Yu-Na Lee (Animal and Plant Quarantine Agency (S-2158) / Avian Influenza Research & Diagnostic Division) |
| EPI_ISL_408861 | A/duck/Korea/H2280/2015 | A / H5N8 | Asia / Korea, Republic of | Yu-Na Lee (Animal and Plant Quarantine Agency (S-2158) / Avian Influenza Research & Diagnostic Division) |
| EPI_ISL_408863 | A/duck/Korea/H2294/2015 | A / H5N8 | Asia / Korea, Republic of | Yu-Na Lee (Animal and Plant Quarantine Agency (S-2158) / Avian Influenza Research & Diagnostic Division) |
| EPI_ISL_408864 | A/duck/Korea/H2300/2015 | A / H5N8 | Asia / Korea, Republic of | Yu-Na Lee (Animal and Plant Quarantine Agency (S-2158) / Avian Influenza Research & Diagnostic Division) |
| EPI_ISL_408866 | A/environment/Korea/H2305/2015 | A / H5N8 | Asia / Korea, Republic of | Yu-Na Lee (Animal and Plant Quarantine Agency (S-2158) / Avian Influenza Research & Diagnostic Division) |
| EPI_ISL_408867 | A/duck/Korea/H2311/2015 | A / H5N8 | Asia / Korea, Republic of | Yu-Na Lee (Animal and Plant Quarantine Agency (S-2158) / Avian Influenza Research & Diagnostic Division) |
| EPI_ISL_408868 | A/duck/Korea/H2312/2015 | A / H5N8 | Asia / Korea, Republic of | Yu-Na Lee (Animal and Plant Quarantine Agency (S-2158) / Avian Influenza Research & Diagnostic Division) |
| EPI_ISL_408869 | A/duck/Korea/H2319/2015 | A / H5N8 | Asia / Korea, Republic of | Yu-Na Lee (Animal and Plant Quarantine Agency (S-2158) / Avian Influenza Research & Diagnostic Division) |
| EPI_ISL_408870 | A/duck/Korea/H2320/2015 | A / H5N8 | Asia / Korea, Republic of | Yu-Na Lee (Animal and Plant Quarantine Agency (S-2158) / Avian Influenza Research & Diagnostic Division) |
| EPI_ISL_408872 | A/chicken/Korea/H2357/2015 | A / H5N8 | Asia / Korea, Republic of | Yu-Na Lee (Animal and Plant Quarantine Agency (S-2158) / Avian Influenza Research & Diagnostic Division) |
| EPI_ISL_408873 | A/duck/Korea/H2358/2015 | A / H5N8 | Asia / Korea, Republic of | Yu-Na Lee (Animal and Plant Quarantine Agency (S-2158) / Avian Influenza Research & Diagnostic Division) |
| EPI_ISL_408874 | A/duck/Korea/H2363/2015 | A / H5N8 | Asia / Korea, Republic of | Yu-Na Lee (Animal and Plant Quarantine Agency (S-2158) / Avian Influenza Research & Diagnostic Division) |
| EPI_ISL_408875 | A/environment/Korea/H2369/2015 | A / H5N8 | Asia / Korea, Republic of | Yu-Na Lee (Animal and Plant Quarantine Agency (S-2158) / Avian Influenza Research & Diagnostic Division) |
| EPI_ISL_408877 | A/duck/Korea/H2388/2015 | A / H5N8 | Asia / Korea, Republic of | Yu-Na Lee (Animal and Plant Quarantine Agency (S-2158) / Avian Influenza Research & Diagnostic Division) |
| EPI_ISL_408878 | A/duck/Korea/H2391/2015 | A / H5N8 | Asia / Korea, Republic of | Yu-Na Lee (Animal and Plant Quarantine Agency (S-2158) / Avian Influenza Research & Diagnostic Division) |
| EPI_ISL_408879 | A/duck/Korea/H2392/2015 | A / H5N8 | Asia / Korea, Republic of | Yu-Na Lee (Animal and Plant Quarantine Agency (S-2158) / Avian Influenza Research & Diagnostic Division) |
| EPI_ISL_408881 | A/duck/Korea/H2394/2015 | A / H5N8 | Asia / Korea, Republic of | Yu-Na Lee (Animal and Plant Quarantine Agency (S-2158) / Avian Influenza Research & Diagnostic Division) |
| EPI_ISL_408882 | A/duck/Korea/H2395/2015 | A / H5N8 | Asia / Korea, Republic of | Yu-Na Lee (Animal and Plant Quarantine Agency (S-2158) / Avian Influenza Research & Diagnostic Division) |
| EPI_ISL_408883 | A/duck/Korea/H2397/2015 | A / H5N8 | Asia / Korea, Republic of | Yu-Na Lee (Animal and Plant Quarantine Agency (S-2158) / Avian Influenza Research & Diagnostic Division) |
| EPI_ISL_408884 | A/duck/Korea/H2399/2015 | A / H5N8 | Asia / Korea, Republic of | Yu-Na Lee (Animal and Plant Quarantine Agency (S-2158) / Avian Influenza Research & Diagnostic Division) |
| EPI_ISL_408886 | A/duck/Korea/H2401/2015 | A / H5N8 | Asia / Korea, Republic of | Yu-Na Lee (Animal and Plant Quarantine Agency (S-2158) / Avian Influenza Research & Diagnostic Division) |
| EPI_ISL_408887 | A/duck/Korea/H2402/2015 | A / H5N8 | Asia / Korea, Republic of | Yu-Na Lee (Animal and Plant Quarantine Agency (S-2158) / Avian Influenza Research & Diagnostic Division) |
| EPI_ISL_408889 | A/duck/Korea/H2410/2015 | A / H5N8 | Asia / Korea, Republic of | Yu-Na Lee (Animal and Plant Quarantine Agency (S-2158) / Avian Influenza Research & Diagnostic Division) |
| EPI_ISL_408890 | A/duck/Korea/H2411/2015 | A / H5N8 | Asia / Korea, Republic of | Yu-Na Lee (Animal and Plant Quarantine Agency (S-2158) / Avian Influenza Research & Diagnostic Division) |
| EPI_ISL_408891 | A/duck/Korea/H2412/2015 | A / H5N8 | Asia / Korea, Republic of | Yu-Na Lee (Animal and Plant Quarantine Agency (S-2158) / Avian Influenza Research & Diagnostic Division) |
| EPI_ISL_408892 | A/duck/Korea/H2426/2015 | A / H5N8 | Asia / Korea, Republic of | Yu-Na Lee (Animal and Plant Quarantine Agency (S-2158) / Avian Influenza Research & Diagnostic Division) |
| EPI_ISL_408893 | A/duck/Korea/H2435/2015 | A / H5N8 | Asia / Korea, Republic of | Yu-Na Lee (Animal and Plant Quarantine Agency (S-2158) / Avian Influenza Research & Diagnostic Division) |
| EPI_ISL_408894 | A/duck/Korea/H2436/2015 | A / H5N8 | Asia / Korea, Republic of | Yu-Na Lee (Animal and Plant Quarantine Agency (S-2158) / Avian Influenza Research & Diagnostic Division) |

| | | | | |
|---|---|---|---|---|
| EPI_ISL_408895 | A/duck/Korea/H2441/2015 | A / H5N8 | Asia / Korea, Republic of | Yu-Na Lee (Animal and Plant Quarantine Agency (S-2158) / Avian Influenza Research & Diagnostic Division) |
| EPI_ISL_408896 | A/duck/Korea/H2442/2015 | A / H5N8 | Asia / Korea, Republic of | Yu-Na Lee (Animal and Plant Quarantine Agency (S-2158) / Avian Influenza Research & Diagnostic Division) |
| EPI_ISL_408897 | A/duck/Korea/H2444/2015 | A / H5N8 | Asia / Korea, Republic of | Yu-Na Lee (Animal and Plant Quarantine Agency (S-2158) / Avian Influenza Research & Diagnostic Division) |
| EPI_ISL_408898 | A/duck/Korea/H2445/2015 | A / H5N8 | Asia / Korea, Republic of | Yu-Na Lee (Animal and Plant Quarantine Agency (S-2158) / Avian Influenza Research & Diagnostic Division) |
| EPI_ISL_408899 | A/duck/Korea/H2446/2015 | A / H5N8 | Asia / Korea, Republic of | Yu-Na Lee (Animal and Plant Quarantine Agency (S-2158) / Avian Influenza Research & Diagnostic Division) |
| EPI_ISL_408900 | A/duck/Korea/H2447/2015 | A / H5N8 | Asia / Korea, Republic of | Yu-Na Lee (Animal and Plant Quarantine Agency (S-2158) / Avian Influenza Research & Diagnostic Division) |
| EPI_ISL_408901 | A/duck/Korea/H2449/2015 | A / H5N8 | Asia / Korea, Republic of | Yu-Na Lee (Animal and Plant Quarantine Agency (S-2158) / Avian Influenza Research & Diagnostic Division) |
| EPI_ISL_408902 | A/duck/Korea/H2454/2015 | A / H5N8 | Asia / Korea, Republic of | Yu-Na Lee (Animal and Plant Quarantine Agency (S-2158) / Avian Influenza Research & Diagnostic Division) |
| EPI_ISL_408903 | A/duck/Korea/H2455/2015 | A / H5N8 | Asia / Korea, Republic of | Yu-Na Lee (Animal and Plant Quarantine Agency (S-2158) / Avian Influenza Research & Diagnostic Division) |
| EPI_ISL_408904 | A/duck/Korea/H2459/2015 | A / H5N8 | Asia / Korea, Republic of | Yu-Na Lee (Animal and Plant Quarantine Agency (S-2158) / Avian Influenza Research & Diagnostic Division) |
| EPI_ISL_408905 | A/duck/Korea/H2460/2015 | A / H5N8 | Asia / Korea, Republic of | Yu-Na Lee (Animal and Plant Quarantine Agency (S-2158) / Avian Influenza Research & Diagnostic Division) |
| EPI_ISL_408906 | A/duck/Korea/H2462/2015 | A / H5N8 | Asia / Korea, Republic of | Yu-Na Lee (Animal and Plant Quarantine Agency (S-2158) / Avian Influenza Research & Diagnostic Division) |
| EPI_ISL_408907 | A/duck/Korea/H2465/2015 | A / H5N8 | Asia / Korea, Republic of | Yu-Na Lee (Animal and Plant Quarantine Agency (S-2158) / Avian Influenza Research & Diagnostic Division) |
| EPI_ISL_408908 | A/duck/Korea/H2466/2015 | A / H5N8 | Asia / Korea, Republic of | Yu-Na Lee (Animal and Plant Quarantine Agency (S-2158) / Avian Influenza Research & Diagnostic Division) |
| EPI_ISL_408909 | A/duck/Korea/H2467/2015 | A / H5N8 | Asia / Korea, Republic of | Yu-Na Lee (Animal and Plant Quarantine Agency (S-2158) / Avian Influenza Research & Diagnostic Division) |
| EPI_ISL_408910 | A/duck/Korea/H2469/2015 | A / H5N8 | Asia / Korea, Republic of | Yu-Na Lee (Animal and Plant Quarantine Agency (S-2158) / Avian Influenza Research & Diagnostic Division) |
| EPI_ISL_408911 | A/duck/Korea/H2474/2015 | A / H5N8 | Asia / Korea, Republic of | Yu-Na Lee (Animal and Plant Quarantine Agency (S-2158) / Avian Influenza Research & Diagnostic Division) |
| EPI_ISL_408912 | A/duck/Korea/H2475/2015 | A / H5N8 | Asia / Korea, Republic of | Yu-Na Lee (Animal and Plant Quarantine Agency (S-2158) / Avian Influenza Research & Diagnostic Division) |
| EPI_ISL_408913 | A/duck/Korea/H2484/2015 | A / H5N8 | Asia / Korea, Republic of | Yu-Na Lee (Animal and Plant Quarantine Agency (S-2158) / Avian Influenza Research & Diagnostic Division) |
| EPI_ISL_408914 | A/duck/Korea/H2485/2015 | A / H5N8 | Asia / Korea, Republic of | Yu-Na Lee (Animal and Plant Quarantine Agency (S-2158) / Avian Influenza Research & Diagnostic Division) |
| EPI_ISL_408915 | A/chicken/Korea/H2486/2015 | A / H5N8 | Asia / Korea, Republic of | Yu-Na Lee (Animal and Plant Quarantine Agency (S-2158) / Avian Influenza Research & Diagnostic Division) |
| EPI_ISL_408917 | A/chicken/Korea/H2497/2015 | A / H5N8 | Asia / Korea, Republic of | Yu-Na Lee (Animal and Plant Quarantine Agency (S-2158) / Avian Influenza Research & Diagnostic Division) |
| EPI_ISL_408918 | A/duck/Korea/H2504/2015 | A / H5N8 | Asia / Korea, Republic of | Yu-Na Lee (Animal and Plant Quarantine Agency (S-2158) / Avian Influenza Research & Diagnostic Division) |
| EPI_ISL_408919 | A/chicken/Korea/H2513/2015 | A / H5N8 | Asia / Korea, Republic of | Yu-Na Lee (Animal and Plant Quarantine Agency (S-2158) / Avian Influenza Research & Diagnostic Division) |
| EPI_ISL_408920 | A/chicken/Korea/H2514/2015 | A / H5N8 | Asia / Korea, Republic of | Yu-Na Lee (Animal and Plant Quarantine Agency (S-2158) / Avian Influenza Research & Diagnostic Division) |
| EPI_ISL_408922 | A/chicken/Korea/H2519/2015 | A / H5N8 | Asia / Korea, Republic of | Yu-Na Lee (Animal and Plant Quarantine Agency (S-2158) / Avian Influenza Research & Diagnostic Division) |
| EPI_ISL_408923 | A/duck/Korea/H2529/2015 | A / H5N8 | Asia / Korea, Republic of | Yu-Na Lee (Animal and Plant Quarantine Agency (S-2158) / Avian Influenza Research & Diagnostic Division) |
| EPI_ISL_408925 | A/chicken/Korea/H2541/2015 | A / H5N8 | Asia / Korea, Republic of | Yu-Na Lee (Animal and Plant Quarantine Agency (S-2158) / Avian Influenza Research & Diagnostic Division) |
| EPI_ISL_408926 | A/chicken/Korea/H2551/2015 | A / H5N8 | Asia / Korea, Republic of | Yu-Na Lee (Animal and Plant Quarantine Agency (S-2158) / Avian Influenza Research & Diagnostic Division) |
| EPI_ISL_408928 | A/chicken/Korea/H2555/2015 | A / H5N8 | Asia / Korea, Republic of | Yu-Na Lee (Animal and Plant Quarantine Agency (S-2158) / Avian Influenza Research & Diagnostic Division) |
| EPI_ISL_408929 | A/chicken/Korea/H2560/2015 | A / H5N8 | Asia / Korea, Republic of | Yu-Na Lee (Animal and Plant Quarantine Agency (S-2158) / Avian Influenza Research & Diagnostic Division) |
| EPI_ISL_408930 | A/duck/Korea/H2564/2015 | A / H5N8 | Asia / Korea, Republic of | Yu-Na Lee (Animal and Plant Quarantine Agency (S-2158) / Avian Influenza Research & Diagnostic Division) |
| EPI_ISL_408932 | A/chicken/Korea/H2568/2015 | A / H5N8 | Asia / Korea, Republic of | Yu-Na Lee (Animal and Plant Quarantine Agency (S-2158) / Avian Influenza Research & Diagnostic Division) |
| EPI_ISL_408933 | A/chicken/Korea/H2570/2015 | A / H5N8 | Asia / Korea, Republic of | Yu-Na Lee (Animal and Plant Quarantine Agency (S-2158) / Avian Influenza Research & Diagnostic Division) |
| EPI_ISL_408934 | A/chicken/Korea/H2573/2015 | A / H5N8 | Asia / Korea, Republic of | Yu-Na Lee (Animal and Plant Quarantine Agency (S-2158) / Avian Influenza Research & Diagnostic Division) |
| EPI_ISL_408935 | A/chicken/Korea/H2580/2015 | A / H5N8 | Asia / Korea, Republic of | Yu-Na Lee (Animal and Plant Quarantine Agency (S-2158) / Avian Influenza Research & Diagnostic Division) |
| EPI_ISL_408936 | A/chicken/Korea/H2584/2015 | A / H5N8 | Asia / Korea, Republic of | Yu-Na Lee (Animal and Plant Quarantine Agency (S-2158) / Avian Influenza Research & Diagnostic Division) |
| EPI_ISL_408938 | A/chicken/Korea/H2596/2015 | A / H5N8 | Asia / Korea, Republic of | Yu-Na Lee (Animal and Plant Quarantine Agency (S-2158) / Avian Influenza Research & Diagnostic Division) |
| EPI_ISL_408940 | A/chicken/Korea/H2599/2015 | A / H5N8 | Asia / Korea, Republic of | Yu-Na Lee (Animal and Plant Quarantine Agency (S-2158) / Avian Influenza Research & Diagnostic Division) |
| EPI_ISL_408941 | A/chicken/Korea/H2600/2015 | A / H5N8 | Asia / Korea, Republic of | Yu-Na Lee (Animal and Plant Quarantine Agency (S-2158) / Avian Influenza Research & Diagnostic Division) |
| EPI_ISL_408943 | A/duck/Korea/H2613/2015 | A / H5N8 | Asia / Korea, Republic of | Yu-Na Lee (Animal and Plant Quarantine Agency (S-2158) / Avian Influenza Research & Diagnostic Division) |
| EPI_ISL_408947 | A/duck/Korea/H2629/2015 | A / H5N8 | Asia / Korea, Republic of | Yu-Na Lee (Animal and Plant Quarantine Agency (S-2158) / Avian Influenza Research & Diagnostic Division) |
| EPI_ISL_408948 | A/duck/Korea/H2631/2015 | A / H5N8 | Asia / Korea, Republic of | Yu-Na Lee (Animal and Plant Quarantine Agency (S-2158) / Avian Influenza Research & Diagnostic Division) |
| EPI_ISL_408955 | A/environment/Korea/H2833/2015 | A / H5N8 | Asia / Korea, Republic of | Yu-Na Lee (Animal and Plant Quarantine Agency (S-2158) / Avian Influenza Research & Diagnostic Division) |
| EPI_ISL_408962 | A/duck/Korea/H3105/2015 | A / H5N8 | Asia / Korea, Republic of | Yu-Na Lee (Animal and Plant Quarantine Agency (S-2158) / Avian Influenza Research & Diagnostic Division) |
| EPI_ISL_408966 | A/duck/Korea/H0088/2014 | A / H5N8 | Asia / Korea, Republic of | Yu-Na Lee (Animal and Plant Quarantine Agency (S-2158) / Avian Influenza Research & Diagnostic Division) |
| EPI_ISL_408967 | A/chicken/Korea/H0453/2014 | A / H5N8 | Asia / Korea, Republic of | Yu-Na Lee (Animal and Plant Quarantine Agency (S-2158) / Avian Influenza Research & Diagnostic Division) |
| EPI_ISL_408969 | A/duck/Korea/H1592/2014 | A / H5N8 | Asia / Korea, Republic of | Yu-Na Lee (Animal and Plant Quarantine Agency (S-2158) / Avian Influenza Research & Diagnostic Division) |
| EPI_ISL_408970 | A/duck/Korea/H1750/2014 | A / H5N8 | Asia / Korea, Republic of | Yu-Na Lee (Animal and Plant Quarantine Agency (S-2158) / Avian Influenza Research & Diagnostic Division) |
| EPI_ISL_408971 | A/duck/Korea/H1763/2014 | A / H5N8 | Asia / Korea, Republic of | Yu-Na Lee (Animal and Plant Quarantine Agency (S-2158) / Avian Influenza Research & Diagnostic Division) |
| EPI_ISL_408972 | A/duck/Korea/H2240/2015 | A / H5N8 | Asia / Korea, Republic of | Yu-Na Lee (Animal and Plant Quarantine Agency (S-2158) / Avian Influenza Research & Diagnostic Division) |
| EPI_ISL_408973 | A/chicken/Korea/H2295/2015 | A / H5N8 | Asia / Korea, Republic of | Yu-Na Lee (Animal and Plant Quarantine Agency (S-2158) / Avian Influenza Research & Diagnostic Division) |
| EPI_ISL_410213 | A/ostrich/Korea/H0829/2014 | A / H5N8 | Asia / Korea, Republic of | Yu-Na Lee (Animal and Plant Quarantine Agency (S-2158) / Avian Influenza Research & Diagnostic Division) |
| EPI_ISL_410291 | A/chicken/Germany-BW/AI00049/2020 | A / H5N8 | Europe / Germany / Baden-Wuerttemberg / Regierungsbezirk Karlsruhe / Hohenlohekreis | Jacqueline King (Friedrich-Loeffler-Institut) |
| EPI_ISL_412984 | A/chicken/Iran/17RS654-01/2016 | A / H5N8 | Asia / Iran, Islamic Republic of | Bianca Zecchin (Istituto Zooprofilattico Sperimentale Delle Venezie) |
| EPI_ISL_412986 | A/chicken/Iran/17RS654-03/2016 | A / H5N8 | Asia / Iran, Islamic Republic of | Bianca Zecchin (Istituto Zooprofilattico Sperimentale Delle Venezie) |
| EPI_ISL_412987 | A/chicken/Iran/17RS654-08/2016 | A / H5N8 | Asia / Iran, Islamic Republic of | Bianca Zecchin (Istituto Zooprofilattico Sperimentale Delle Venezie) |
| EPI_ISL_412988 | A/chicken/Iran/17RS654-15/2017 | A / H5N8 | Asia / Iran, Islamic Republic of | Bianca Zecchin (Istituto Zooprofilattico Sperimentale Delle Venezie) |
| EPI_ISL_412989 | A/chicken/Iran/17RS654-26/2017 | A / H5N8 | Asia / Iran, Islamic Republic of | Bianca Zecchin (Istituto Zooprofilattico Sperimentale Delle Venezie) |
| EPI_ISL_412990 | A/chicken/Iran/17RS654-28/2016 | A / H5N8 | Asia / Iran, Islamic Republic of | Bianca Zecchin (Istituto Zooprofilattico Sperimentale Delle Venezie) |
| EPI_ISL_412991 | A/chicken/Iran/17RS654-29/2017 | A / H5N8 | Asia / Iran, Islamic Republic of | Bianca Zecchin (Istituto Zooprofilattico Sperimentale Delle Venezie) |
| EPI_ISL_412992 | A/chicken/Iran/17RS654-34/2017 | A / H5N8 | Asia / Iran, Islamic Republic of | Bianca Zecchin (Istituto Zooprofilattico Sperimentale Delle Venezie) |
| EPI_ISL_412994 | A/chicken/Iran/18VIR2027-01/2017 | A / H5N8 | Asia / Iran, Islamic Republic of | Bianca Zecchin (Istituto Zooprofilattico Sperimentale Delle Venezie) |
| EPI_ISL_412995 | A/chicken/Iran/18VIR2027-02/2017 | A / H5N8 | Asia / Iran, Islamic Republic of | Bianca Zecchin (Istituto Zooprofilattico Sperimentale Delle Venezie) |
| EPI_ISL_412996 | A/chicken/Iran/18VIR2027-03/2017 | A / H5N8 | Asia / Iran, Islamic Republic of | Bianca Zecchin (Istituto Zooprofilattico Sperimentale Delle Venezie) |
| EPI_ISL_412997 | A/chicken/Iran/18VIR2027-04/2017 | A / H5N8 | Asia / Iran, Islamic Republic of | Bianca Zecchin (Istituto Zooprofilattico Sperimentale Delle Venezie) |
| EPI_ISL_412998 | A/chicken/Iran/18VIR2027-05/2017 | A / H5N8 | Asia / Iran, Islamic Republic of | Bianca Zecchin (Istituto Zooprofilattico Sperimentale Delle Venezie) |
| EPI_ISL_412999 | A/chicken/Iran/18VIR2027-06/2017 | A / H5N8 | Asia / Iran, Islamic Republic of | Bianca Zecchin (Istituto Zooprofilattico Sperimentale Delle Venezie) |
| EPI_ISL_413000 | A/chicken/Iran/18VIR2027-07/2017 | A / H5N8 | Asia / Iran, Islamic Republic of | Bianca Zecchin (Istituto Zooprofilattico Sperimentale Delle Venezie) |
| EPI_ISL_413001 | A/chicken/Iran/18VIR2027-08/2017 | A / H5N8 | Asia / Iran, Islamic Republic of | Bianca Zecchin (Istituto Zooprofilattico Sperimentale Delle Venezie) |
| EPI_ISL_413002 | A/chicken/Iran/18VIR2027-09/2017 | A / H5N8 | Asia / Iran, Islamic Republic of | Bianca Zecchin (Istituto Zooprofilattico Sperimentale Delle Venezie) |
| EPI_ISL_413003 | A/chicken/Iran/18VIR2027-10/2017 | A / H5N8 | Asia / Iran, Islamic Republic of | Bianca Zecchin (Istituto Zooprofilattico Sperimentale Delle Venezie) |

| | | | | |
|---|---|---|---|---|
| EPI_ISL_413005 | A/chicken/Iran/18VIR2027-14/2017 | A / H5N8 | Asia / Iran, Islamic Republic of | Bianca Zecchin (Istituto Zooprofilattico Sperimentale Delle Venezie) |
| EPI_ISL_413006 | A/chicken/Iran/18VIR2027-15/2017 | A / H5N8 | Asia / Iran, Islamic Republic of | Bianca Zecchin (Istituto Zooprofilattico Sperimentale Delle Venezie) |
| EPI_ISL_413008 | A/flamingo/Iran/17RS654-18/2016 | A / H5N8 | Asia / Iran, Islamic Republic of | Bianca Zecchin (Istituto Zooprofilattico Sperimentale Delle Venezie) |
| EPI_ISL_413009 | A/gadwall/Iran/18VIR2027-20/2018 | A / H5N6 | Asia / Iran, Islamic Republic of | Bianca Zecchin (Istituto Zooprofilattico Sperimentale Delle Venezie) |
| EPI_ISL_413010 | A/little_grebe/Iran/17RS654-10/2016 | A / H5N8 | Asia / Iran, Islamic Republic of | Bianca Zecchin (Istituto Zooprofilattico Sperimentale Delle Venezie) |
| EPI_ISL_413011 | A/wild_bird/Iran/17RS654-24/2016 | A / H5N8 | Asia / Iran, Islamic Republic of | Bianca Zecchin (Istituto Zooprofilattico Sperimentale Delle Venezie) |
| EPI_ISL_413225 | A/Mallard/Republic of Georgia/1/2015 | A / H5N8 | Asia / Georgia | Maria Johanna Poen (Erasmus Medical Center / Department of Virology) |
| EPI_ISL_413226 | A/Mallard/Republic of Georgia/2/2015 | A / H5N8 | Asia / Georgia | Maria Johanna Poen (Erasmus Medical Center / Department of Virology) |
| EPI_ISL_415197 | A/chicken/Germany-SN/AI00276/2020 | A / H5N8 | Europe / Germany / Saxony / Regierungsbezirk Leipzig / Leipzig | Jacqueline King (Friedrich-Loeffler-Institut) |
| EPI_ISL_417414 | A/buzzard/Germany-SN/AI00285/2020 | A / H5N8 | Europe / Germany / Saxony | Jacqueline King (Friedrich-Loeffler-Institut) |
| EPI_ISL_417415 | A/turkey/Germany-NI/AI00334/2020 | A / H5N8 | Europe / Germany / Lower Saxony / Aurich | Jacqueline King (Friedrich-Loeffler-Institut) |
| EPI_ISL_418169 | A/Whooper swan/Xinjiang/1/2020 | A / H5N6 | Asia / China / Xinjiang | Yanbing Li (Harbin Veterinary Research Institute (CAAS) / Ministry of Agriculture) |
| EPI_ISL_418170 | A/Whooper swan/Xinjiang/2/2020 | A / H5N6 | Asia / China / Xinjiang | Yanbing Li (Harbin Veterinary Research Institute (CAAS) / Ministry of Agriculture) |
| EPI_ISL_418171 | A/Whooper swan/Xinjiang/3/2020 | A / H5N6 | Asia / China / Xinjiang | Yanbing Li (Harbin Veterinary Research Institute (CAAS) / Ministry of Agriculture) |
| EPI_ISL_418172 | A/Mute swan/Xinjiang/4/2020 | A / H5N6 | Asia / China / Xinjiang | Yanbing Li (Harbin Veterinary Research Institute (CAAS) / Ministry of Agriculture) |
| EPI_ISL_418173 | A/Mute swan/Xinjiang/5/2020 | A / H5N6 | Asia / China / Xinjiang | Yanbing Li (Harbin Veterinary Research Institute (CAAS) / Ministry of Agriculture) |
| EPI_ISL_418174 | A/Whooper swan/Xinjiang/6/2020 | A / H5N6 | Asia / China / Xinjiang | Yanbing Li (Harbin Veterinary Research Institute (CAAS) / Ministry of Agriculture) |
| EPI_ISL_418175 | A/Whooper swan/Xinjiang/7/2020 | A / H5N6 | Asia / China / Xinjiang | Yanbing Li (Harbin Veterinary Research Institute (CAAS) / Ministry of Agriculture) |
| EPI_ISL_418176 | A/Whooper swan/Xinjiang/8/2020 | A / H5N6 | Asia / China / Xinjiang | Yanbing Li (Harbin Veterinary Research Institute (CAAS) / Ministry of Agriculture) |
| EPI_ISL_418177 | A/Whooper swan/Xinjiang/9/2020 | A / H5N6 | Asia / China / Xinjiang | Yanbing Li (Harbin Veterinary Research Institute (CAAS) / Ministry of Agriculture) |
| EPI_ISL_418178 | A/Whooper swan/Xinjiang/10/2020 | A / H5N6 | Asia / China / Xinjiang | Yanbing Li (Harbin Veterinary Research Institute (CAAS) / Ministry of Agriculture) |
| EPI_ISL_418179 | A/Whooper swan/Xinjiang/11/2020 | A / H5N6 | Asia / China / Xinjiang | Yanbing Li (Harbin Veterinary Research Institute (CAAS) / Ministry of Agriculture) |
| EPI_ISL_418180 | A/Whooper swan/Xinjiang/12/2020 | A / H5N6 | Asia / China / Xinjiang | Yanbing Li (Harbin Veterinary Research Institute (CAAS) / Ministry of Agriculture) |
| EPI_ISL_418181 | A/Whooper swan/Xinjiang/13/2020 | A / H5N6 | Asia / China / Xinjiang | Yanbing Li (Harbin Veterinary Research Institute (CAAS) / Ministry of Agriculture) |
| EPI_ISL_418266 | A/turkey/Czech Republic/3071/2020 | A / H5N8 | Europe / Czech Republic / Pardubicky kraj / Okres Pardubice / Slepotice, Moravany | Alexander Nagy (State Veterinary Institute Prague) |
| EPI_ISL_419212 | A/chicken/Bulgaria/Dobrich/12-1/2018 | A / H5N8 | Europe / Bulgaria / Oblast Dobrich | Divya Venkatesh (University of Cambridge / Zoology) |
| EPI_ISL_419220 | A/turkey/Hungary/1020_20VIR749-1/2020 | A / H5N8 | Europe / Hungary | Ambra Pastori (Istituto Zooprofilattico Sperimentale Delle Venezie) |
| EPI_ISL_419239 | A/duck/Hungary/1565_20VIR749-2/2020 | A / H5N8 | Europe / Hungary | Ambra Pastori (Istituto Zooprofilattico Sperimentale Delle Venezie) |
| EPI_ISL_419312 | A/steamer duck/Germany-SN/AI00346/2020 | A / H5N8 | Europe / Germany / Saxony / Zwickau | Jacqueline King (Friedrich-Loeffler-Institut) |
| EPI_ISL_419314 | A/turkey/Germany-ST/AI00352/2020 | A / H5N8 | Europe / Germany / Saxony-Anhalt / Börde | Jacqueline King (Friedrich-Loeffler-Institut) |
| EPI_ISL_419344 | A/chicken/Bulgaria/Dobrich/12-2/2018 | A / H5N8 | Europe / Bulgaria | Divya Venkatesh (University of Cambridge / Zoology) |
| EPI_ISL_419345 | A/chicken/Bulgaria/Plovdiv/224-1/2018 | A / H5N8 | Europe / Bulgaria | Divya Venkatesh (University of Cambridge / Zoology) |
| EPI_ISL_419346 | A/chicken/Bulgaria/Plovdiv/224-2/2018 | A / H5N8 | Europe / Bulgaria | Divya Venkatesh (University of Cambridge / Zoology) |
| EPI_ISL_419347 | A/chicken/Bulgaria/Plovdiv/224-3/2018 | A / H5N8 | Europe / Bulgaria | Divya Venkatesh (University of Cambridge / Zoology) |
| EPI_ISL_419348 | A/chicken/Bulgaria/Haskovo/411/2017 | A / H5N8 | Europe / Bulgaria | Divya Venkatesh (University of Cambridge / Zoology) |
| EPI_ISL_419349 | A/chicken/Bulgaria/Sliven/432/2017 | A / H5N8 | Europe / Bulgaria | Divya Venkatesh (University of Cambridge / Zoology) |
| EPI_ISL_419350 | A/duck/Bulgaria/Dobrich/407/2017 | A / H5N8 | Europe / Bulgaria | Divya Venkatesh (University of Cambridge / Zoology) |
| EPI_ISL_419351 | A/duck/Bulgaria/Yambol/436/2017 | A / H5N8 | Europe / Bulgaria | Divya Venkatesh (University of Cambridge / Zoology) |
| EPI_ISL_419352 | A/duck/Bulgaria/Stara-Zagora/623/2017 | A / H5N8 | Europe / Bulgaria | Divya Venkatesh (University of Cambridge / Zoology) |
| EPI_ISL_419353 | A/duck/Bulgaria/Yambol/35-1/2018 | A / H5N8 | Europe / Bulgaria | Divya Venkatesh (University of Cambridge / Zoology) |
| EPI_ISL_419355 | A/partridge/Bulgaria/Plovdiv/60-2/2018 | A / H5N8 | Europe / Bulgaria | Divya Venkatesh (University of Cambridge / Zoology) |
| EPI_ISL_419357 | A/duck/Bulgaria/Plovdiv/76-1/2018 | A / H5N8 | Europe / Bulgaria | Divya Venkatesh (University of Cambridge / Zoology) |
| EPI_ISL_419358 | A/duck/Bulgaria/Plovdiv/76-2/2018 | A / H5N8 | Europe / Bulgaria | Divya Venkatesh (University of Cambridge / Zoology) |
| EPI_ISL_419359 | A/chicken/Bulgaria/Dobrich/115/2018 | A / H5N8 | Europe / Bulgaria | Divya Venkatesh (University of Cambridge / Zoology) |
| EPI_ISL_419360 | A/chicken/Bulgaria/Dobrich/163-1/2018 | A / H5N8 | Europe / Bulgaria | Divya Venkatesh (University of Cambridge / Zoology) |
| EPI_ISL_419361 | A/chicken/Bulgaria/Dobrich/163-2/2018 | A / H5N8 | Europe / Bulgaria | Divya Venkatesh (University of Cambridge / Zoology) |
| EPI_ISL_419362 | A/chicken/Bulgaria/Haskovo/286/2018 | A / H5N8 | Europe / Bulgaria | Divya Venkatesh (University of Cambridge / Zoology) |
| EPI_ISL_419363 | A/chicken/Bulgaria/Plovdiv/295/2018 | A / H5N8 | Europe / Bulgaria | Divya Venkatesh (University of Cambridge / Zoology) |
| EPI_ISL_419364 | A/chicken/Bulgaria/Plovdiv/333/2018 | A / H5N8 | Europe / Bulgaria | Divya Venkatesh (University of Cambridge / Zoology) |
| EPI_ISL_419365 | A/turkey/Bulgaria/Haskovo/336/2018 | A / H5N8 | Europe / Bulgaria | Divya Venkatesh (University of Cambridge / Zoology) |
| EPI_ISL_419366 | A/duck/Bulgaria/Plovdiv/74-1/2018 | A / H5N8 | Europe / Bulgaria | Divya Venkatesh (University of Cambridge / Zoology) |
| EPI_ISL_419367 | A/chicken/Bulgaria/Vidin-Kosovo/550/2018 | A / H5N8 | Europe / Bulgaria | Divya Venkatesh (University of Cambridge / Zoology) |
| EPI_ISL_436133 | A/chicken/Germany-MV/AR9311-L02968/2016 | A / H5N8 | Europe / Germany / Mecklenburg-Vorpommern / Vorpommern-Rügen | Jacqueline King (Friedrich-Loeffler-Institut) |
| EPI_ISL_436134 | A/domestic duck/Germany-MV/AR9433-L02969/2016 | A / H5N8 | Europe / Germany / Mecklenburg-Vorpommern / Vorpommern-Greifswald | Jacqueline King (Friedrich-Loeffler-Institut) |
| EPI_ISL_436135 | A/chicken/Germany-MV/AR9528-L02970/2016 | A / H5N8 | Europe / Germany / Mecklenburg-Vorpommern / Vorpommern-Rügen | Jacqueline King (Friedrich-Loeffler-Institut) |
| EPI_ISL_436136 | A/chicken/Germany-MV/AR9738-L02971/2016 | A / H5N8 | Europe / Germany / Mecklenburg-Vorpommern / Vorpommern-Greifswald | Jacqueline King (Friedrich-Loeffler-Institut) |
| EPI_ISL_436142 | A/domestic duck/Germany-ST/AR11090-L02972/2016 | A / H5N8 | Europe / Germany / Saxony-Anhalt / Jerichower Land | Jacqueline King (Friedrich-Loeffler-Institut) |
| EPI_ISL_436143 | A/turkey/Germany-NW/AR11095-L02973/2016 | A / H5N8 | Europe / Germany / North Rhine-Westphalia / Soest | Jacqueline King (Friedrich-Loeffler-Institut) |
| EPI_ISL_436144 | A/domestic goose/Germany-NW/AR11222-L02974/2016 | A / H5N8 | Europe / Germany / North Rhine-Westphalia / Gütersloh | Jacqueline King (Friedrich-Loeffler-Institut) |
| EPI_ISL_436145 | A/turkey/Germany-NW/AR11455-L02975/2016 | A / H5N8 | Europe / Germany / North Rhine-Westphalia / Kleve | Jacqueline King (Friedrich-Loeffler-Institut) |
| EPI_ISL_436146 | A/chicken/Germany-ST/AR101-L02976/2017 | A / H5N8 | Europe / Germany / Saxony-Anhalt / Salzlandkreis | Jacqueline King (Friedrich-Loeffler-Institut) |
| EPI_ISL_436147 | A/turkey/Germany-NW/AR246-L02977/2017 | A / H5N8 | Europe / Germany / North Rhine-Westphalia / Wesel | Jacqueline King (Friedrich-Loeffler-Institut) |
| EPI_ISL_436148 | A/turkey/Germany-NW/AR258-L02994/2017 | A / H5N8 | Europe / Germany / North Rhine-Westphalia / Paderborn | Jacqueline King (Friedrich-Loeffler-Institut) |
| EPI_ISL_436149 | A/domestic duck/Germany-NI/AR350-L02995/2017 | A / H5N8 | Europe / Germany / Lower Saxony / Diepholz | Jacqueline King (Friedrich-Loeffler-Institut) |
| EPI_ISL_436150 | A/turkey/Germany-SH/AR495-L02996/2017 | A / H5N5 | Europe / Germany / Schleswig-Holstein / Steinburg | Jacqueline King (Friedrich-Loeffler-Institut) |
| EPI_ISL_436151 | A/turkey/Germany-SH/AR804-L02997/2017 | A / H5N5 | Europe / Germany / Schleswig-Holstein / Steinburg | Jacqueline King (Friedrich-Loeffler-Institut) |
| EPI_ISL_436152 | A/turkey/Germany-MV/AR622-L02998/2017 | A / H5N8 | Europe / Germany / Mecklenburg-Vorpommern / Grimmen | Jacqueline King (Friedrich-Loeffler-Institut) |
| EPI_ISL_436159 | A/chicken/Germany-BY/AR845-L02999/2017 | A / H5N8 | Europe / Germany / Bavaria / Hof | Jacqueline King (Friedrich-Loeffler-Institut) |
| EPI_ISL_436160 | A/chicken/Germany-MV/AR851-L03000/2017 | A / H5N8 | Europe / Germany / Mecklenburg-Vorpommern / Ludwigslust-Parchim | Jacqueline King (Friedrich-Loeffler-Institut) |
| EPI_ISL_436161 | A/domestic duck/Germany-BB/AR877-L03001/2017 | A / H5N8 | Europe / Germany / Brandenburg / Märkisch-Oderland | Jacqueline King (Friedrich-Loeffler-Institut) |
| EPI_ISL_436162 | A/turkey/Germany-SN/AR888-L03002/2017 | A / H5N8 | Europe / Germany / Saxony / Nordsachsen | Jacqueline King (Friedrich-Loeffler-Institut) |
| EPI_ISL_436163 | A/domestic duck/Germany-BB/AR906-L03003/2017 | A / H5N8 | Europe / Germany / Brandenburg / Märkisch-Oderland | Jacqueline King (Friedrich-Loeffler-Institut) |

| | | | | |
|---|---|---|---|---|
| EPI_ISL_436164 | A/turkey/Germany-BB/AR1015-L03026/2017 | A / H5N8 | Europe / Germany / Brandenburg / Märkisch-Oderland | Jacqueline King (Friedrich-Loeffler-Institut) |
| EPI_ISL_436165 | A/turkey/Germany-BY/AR1132-L03027/2017 | A / H5N8 | Europe / Germany / Bavaria / Straubing-Bogen | Jacqueline King (Friedrich-Loeffler-Institut) |
| EPI_ISL_436166 | A/turkey/Germany-BB/AR1223-L03028/2017 | A / H5N8 | Europe / Germany / Brandenburg / Ostprignitz-Ruppin | Jacqueline King (Friedrich-Loeffler-Institut) |
| EPI_ISL_436167 | A/domestic duck/Germany-BB/AR1341-L03029/2017 | A / H5N8 | Europe / Germany / Brandenburg / Märkisch-Oderland | Jacqueline King (Friedrich-Loeffler-Institut) |
| EPI_ISL_436168 | A/chicken/Germany-NW/AR1384-L03030/2017 | A / H5N8 | Europe / Germany / North Rhine-Westphalia / Paderborn | Jacqueline King (Friedrich-Loeffler-Institut) |
| EPI_ISL_436169 | A/chicken/Germany-MV/AR1398-L03031/2017 | A / H5N8 | Europe / Germany / Mecklenburg-Vorpommern / Rostock | Jacqueline King (Friedrich-Loeffler-Institut) |
| EPI_ISL_436170 | A/domestic duck/Germany-BB/AR1464-L03032/2017 | A / H5N8 | Europe / Germany / Brandenburg / Märkisch-Oderland | Jacqueline King (Friedrich-Loeffler-Institut) |
| EPI_ISL_436171 | A/chicken/Germany-TH/AR1489-L03033/2017 | A / H5N8 | Europe / Germany / Thuringia / Greiz | Jacqueline King (Friedrich-Loeffler-Institut) |
| EPI_ISL_436172 | A/chicken/Germany-TH/AR1638-L03034/2017 | A / H5N8 | Europe / Germany / Thuringia / Sömmerda | Jacqueline King (Friedrich-Loeffler-Institut) |
| EPI_ISL_436173 | A/chicken/Germany-RP/AR1774-L03035/2017 | A / H5N8 | Europe / Germany / Rhineland-Palatinate / Bad Dürkheim | Jacqueline King (Friedrich-Loeffler-Institut) |
| EPI_ISL_436174 | A/chicken/Germany-NW/AR3197-L03036/2017 | A / H5N8 | Europe / Germany / North Rhine-Westphalia / Lippe | Jacqueline King (Friedrich-Loeffler-Institut) |
| EPI_ISL_436175 | A/turkey/Germany-NI/AR1964-L02703/2017 | A / H5N8 | Europe / Germany / Lower Saxony / Cloppenburg | Jacqueline King (Friedrich-Loeffler-Institut) |
| EPI_ISL_436176 | A/turkey/Germany-NI/AR2024-L02704/2017 | A / H5N8 | Europe / Germany / Lower Saxony / Cloppenburg | Jacqueline King (Friedrich-Loeffler-Institut) |
| EPI_ISL_436177 | A/turkey/Germany-NI/AR2095-L02705/2017 | A / H5N8 | Europe / Germany / Lower Saxony / Cloppenburg | Jacqueline King (Friedrich-Loeffler-Institut) |
| EPI_ISL_436178 | A/turkey/Germany-NI/AR2142-L02706/2017 | A / H5N8 | Europe / Germany / Lower Saxony / Cloppenburg | Jacqueline King (Friedrich-Loeffler-Institut) |
| EPI_ISL_436179 | A/turkey/Germany-NI/AR2195-L02707/2017 | A / H5N8 | Europe / Germany / Lower Saxony / Cloppenburg | Jacqueline King (Friedrich-Loeffler-Institut) |
| EPI_ISL_436180 | A/turkey/Germany-NI/AR2248-L02708/2017 | A / H5N8 | Europe / Germany / Lower Saxony / Cloppenburg | Jacqueline King (Friedrich-Loeffler-Institut) |
| EPI_ISL_436181 | A/turkey/Germany-NI/AR2350-L02709/2017 | A / H5N8 | Europe / Germany / Lower Saxony / Cloppenburg | Jacqueline King (Friedrich-Loeffler-Institut) |
| EPI_ISL_436182 | A/turkey/Germany-NI/AR2374-L02710/2017 | A / H5N8 | Europe / Germany / Lower Saxony / Cloppenburg | Jacqueline King (Friedrich-Loeffler-Institut) |
| EPI_ISL_436183 | A/turkey/Germany-NI/AR2493-L02711/2017 | A / H5N8 | Europe / Germany / Lower Saxony / Cloppenburg | Jacqueline King (Friedrich-Loeffler-Institut) |
| EPI_ISL_436184 | A/chicken/Germany-NI/AR2541-L02712/2017 | A / H5N8 | Europe / Germany / Lower Saxony / Cloppenburg | Jacqueline King (Friedrich-Loeffler-Institut) |
| EPI_ISL_436185 | A/turkey/Germany-NI/AR2704-L02713/2017 | A / H5N8 | Europe / Germany / Lower Saxony / Cloppenburg | Jacqueline King (Friedrich-Loeffler-Institut) |
| EPI_ISL_436186 | A/turkey/Germany-NI/AR2552-L02714/2017 | A / H5N8 | Europe / Germany / Lower Saxony / Cloppenburg | Jacqueline King (Friedrich-Loeffler-Institut) |
| EPI_ISL_436187 | A/turkey/Germany-NI/AR2732-L02715/2017 | A / H5N8 | Europe / Germany / Lower Saxony / Cloppenburg | Jacqueline King (Friedrich-Loeffler-Institut) |
| EPI_ISL_436188 | A/turkey/Germany-NI/AR2769-L02716/2017 | A / H5N8 | Europe / Germany / Lower Saxony / Cloppenburg | Jacqueline King (Friedrich-Loeffler-Institut) |
| EPI_ISL_436189 | A/turkey/Germany-NI/AR2795-L02717/2017 | A / H5N8 | Europe / Germany / Lower Saxony / Cloppenburg | Jacqueline King (Friedrich-Loeffler-Institut) |
| EPI_ISL_436190 | A/chicken/Germany-NI/AR2921-L02720/2017 | A / H5N8 | Europe / Germany / Lower Saxony / Ammerland | Jacqueline King (Friedrich-Loeffler-Institut) |
| EPI_ISL_436191 | A/turkey/Germany-NI/AR2909-L02719/2017 | A / H5N8 | Europe / Germany / Lower Saxony / Cloppenburg | Jacqueline King (Friedrich-Loeffler-Institut) |
| EPI_ISL_436192 | A/turkey/Germany-NI/AR3012-L02721/2017 | A / H5N8 | Europe / Germany / Lower Saxony / Cloppenburg | Jacqueline King (Friedrich-Loeffler-Institut) |
| EPI_ISL_436193 | A/turkey/Germany-NI/AR3128-L02722/2017 | A / H5N8 | Europe / Germany / Lower Saxony / Cloppenburg | Jacqueline King (Friedrich-Loeffler-Institut) |
| EPI_ISL_436690 | A/turkey/Germany-SH/AR417-L03291/2017 | A / H5N5 | Europe / Germany / Schleswig-Holstein / Steinburg | Jacqueline King (Friedrich-Loeffler-Institut) |
| EPI_ISL_436692 | A/turkey/Germany-SH/AR420-L03292/2017 | A / H5N5 | Europe / Germany / Schleswig-Holstein / Steinburg | Jacqueline King (Friedrich-Loeffler-Institut) |
| EPI_ISL_436693 | A/turkey/Germany-SH/AR424-L03293/2017 | A / H5N5 | Europe / Germany / Schleswig-Holstein / Steinburg | Jacqueline King (Friedrich-Loeffler-Institut) |
| EPI_ISL_436694 | A/turkey/Germany-SH/AR425-L03294/2017 | A / H5N5 | Europe / Germany / Schleswig-Holstein / Steinburg | Jacqueline King (Friedrich-Loeffler-Institut) |
| EPI_ISL_436695 | A/turkey/Germany-SH/AR426-L03295/2017 | A / H5N5 | Europe / Germany / Schleswig-Holstein / Steinburg | Jacqueline King (Friedrich-Loeffler-Institut) |
| EPI_ISL_436696 | A/turkey/Germany-SH/AR494-L03296/2017 | A / H5N5 | Europe / Germany / Schleswig-Holstein / Steinburg | Jacqueline King (Friedrich-Loeffler-Institut) |
| EPI_ISL_436697 | A/turkey/Germany-SH/AR769-L03298/2017 | A / H5N5 | Europe / Germany / Schleswig-Holstein / Steinburg | Jacqueline King (Friedrich-Loeffler-Institut) |
| EPI_ISL_436698 | A/turkey/Germany-SH/AR780-L03299/2017 | A / H5N5 | Europe / Germany / Schleswig-Holstein / Steinburg | Jacqueline King (Friedrich-Loeffler-Institut) |
| EPI_ISL_436699 | A/turkey/Germany-SH/AR786-L03300/2017 | A / H5N5 | Europe / Germany / Schleswig-Holstein / Steinburg | Jacqueline King (Friedrich-Loeffler-Institut) |
| EPI_ISL_436700 | A/turkey/Germany-SH/AR793-L03301/2017 | A / H5N5 | Europe / Germany / Schleswig-Holstein / Steinburg | Jacqueline King (Friedrich-Loeffler-Institut) |
| EPI_ISL_436701 | A/turkey/Germany-SH/AR803-L03302/2017 | A / H5N5 | Europe / Germany / Schleswig-Holstein / Steinburg | Jacqueline King (Friedrich-Loeffler-Institut) |
| EPI_ISL_436702 | A/turkey/Germany-SH/AR806-L03303/2017 | A / H5N5 | Europe / Germany / Schleswig-Holstein / Steinburg | Jacqueline King (Friedrich-Loeffler-Institut) |
| EPI_ISL_436703 | A/turkey/Germany-SH/AR815-L03304/2017 | A / H5N5 | Europe / Germany / Schleswig-Holstein / Steinburg | Jacqueline King (Friedrich-Loeffler-Institut) |
| EPI_ISL_436704 | A/turkey/Germany-SH/AR822-L03305/2017 | A / H5N5 | Europe / Germany / Schleswig-Holstein / Steinburg | Jacqueline King (Friedrich-Loeffler-Institut) |
| EPI_ISL_436705 | A/turkey/Germany-SH/AR827-L03306/2017 | A / H5N5 | Europe / Germany / Schleswig-Holstein / Steinburg | Jacqueline King (Friedrich-Loeffler-Institut) |
| EPI_ISL_436707 | A/greylag goose/Germany-SH/AR1609-L03307/2017 | A / H5N5 | Europe / Germany / Schleswig-Holstein / Neumünster | Jacqueline King (Friedrich-Loeffler-Institut) |
| EPI_ISL_436708 | A/mallard/Germany-SH/AR1659-L03308/2017 | A / H5N5 | Europe / Germany / Schleswig-Holstein / Rendsburg-Eckernförde | Jacqueline King (Friedrich-Loeffler-Institut) |
| EPI_ISL_436709 | A/mute swan/Germany-SH/AR1802-L03309/2017 | A / H5N5 | Europe / Germany / Schleswig-Holstein / Rendsburg-Eckernförde | Jacqueline King (Friedrich-Loeffler-Institut) |
| EPI_ISL_436710 | A/greylag goose/Germany-SH/AR1803-L03310/2017 | A / H5N5 | Europe / Germany / Schleswig-Holstein / Neumünster | Jacqueline King (Friedrich-Loeffler-Institut) |
| EPI_ISL_436712 | A/common buzzard/Germany-SH/AR2172-L03311/2017 | A / H5N5 | Europe / Germany / Schleswig-Holstein / Segeberg | Jacqueline King (Friedrich-Loeffler-Institut) |
| EPI_ISL_4396748 | A/Meleagris gallopavo/Belgium/11293_001/2021 | A / H5N8 | Europe / Belgium / Provincie West-Vlaanderen | Steven Van Borm (Sciensano, Department of Animal Infectious Diseases / Animal Infectious Diseases) |
| EPI_ISL_4558108 | A/Sichuan/06689/2021 | A / H5N6 | Asia / China / Sichuan Province | Lei Yang (WHO Chinese National Influenza Center / Virology Institute, Chinese CDC) |
| EPI_ISL_4568642 | A/Hunan/09285/2021 | A / H5N6 | Asia / China / Hunan Province / Hunan | Lei Yang (WHO Chinese National Influenza Center / Virology Institute, Chinese CDC) |
| EPI_ISL_4568643 | A/Hunan/09911/2021 | A / H5N6 | Asia / China / Hunan Province / Hunan | Lei Yang (WHO Chinese National Influenza Center / Virology Institute, Chinese CDC) |
| EPI_ISL_4568644 | A/Chongqing/02/2021 | A / H5N6 | Asia / China / Chongqinq Municipality / Chongqing | Lei Yang (WHO Chinese National Influenza Center / Virology Institute, Chinese CDC) |
| EPI_ISL_463009 | A/duck/Bulgaria/Plovdiv/74-2/2018 | A / H5N2 | Europe / Bulgaria | Divya Venkatesh (University of Cambridge / Zoology) |
| EPI_ISL_4651963 | A/common pheasant /Sweden/SVA210923SZ0341/KN000365/ | A / H5N1 | Europe / Sweden / Skane Lan / Kristianstads Kommun | Siamak Zohari (National Veterinary Institute) |
| EPI_ISL_4653177 | A/common pheasant /Sweden/SVA210923SZ0341/KN000366/ | A / H5N1 | Europe / Sweden / Skane Lan / Kristianstads Kommun | Siamak Zohari (National Veterinary Institute) |
| EPI_ISL_4804850 | A/seal/Germany-SH/AI05373/2021 | A / H5N8 | Europe / Germany / Schleswig-Holstein / Nordfriesland | Jacqueline King (Friedrich-Loeffler-Institut) |
| EPI_ISL_4805852 | A/seal/Germany-SH/AI05377/2021 | A / H5N8 | Europe / Germany / Schleswig-Holstein / Dithmarschen | Jacqueline King (Friedrich-Loeffler-Institut) |
| EPI_ISL_4805936 | A/seal/Germany-SH/AI05379/2021 | A / H5N8 | Europe / Germany / Schleswig-Holstein / Nordfriesland | Jacqueline King (Friedrich-Loeffler-Institut) |
| EPI_ISL_486440 | recombinant A/tufted duck/Germany/AR8444/2016 | A / H5N8 | Europe / Germany | El-Sayed M. Abdelwhab (Friedrich-Loeffler-Institut) |
| EPI_ISL_486441 | recombinant A/turkey/Germany-MV/AR2487/2014 | A / H5N8 | Europe / Germany | El-Sayed M. Abdelwhab (Friedrich-Loeffler-Institut) |
| EPI_ISL_499181 | A/goose/China/GS39/2016 | A / H5N6 | Asia / China | |
| EPI_ISL_499182 | A/chicken/China/CK44/2016 | A / H5N6 | Asia / China | |
| EPI_ISL_499183 | A/duck/China/DK47/2016 | A / H5N6 | Asia / China | |
| EPI_ISL_499184 | A/chicken/China/CK74/2016 | A / H5N6 | Asia / China | |
| EPI_ISL_501644 | A/Duck/Viet Nam/HU12-981/2019 | A / H5N6 | Asia / Vietnam | |
| EPI_ISL_501645 | A/Duck/Viet Nam/HU12-1303/2019 | A / H5N6 | Asia / Vietnam | |
| EPI_ISL_501646 | A/Duck/Viet Nam/HU12-982/2019 | A / H5N6 | Asia / Vietnam | |
| EPI_ISL_501647 | A/Duck/Viet Nam/HU12-1310/2019 | A / H5N6 | Asia / Vietnam | |
| EPI_ISL_501648 | A/Duck/Viet Nam/HU12-1305/2019 | A / H5N6 | Asia / Vietnam | |

| | | | | |
|---|---|---|---|---|
| EPI_ISL_501649 | A/Duck/Viet Nam/HU12-1309/2019 | A / H5N6 | Asia / Vietnam | |
| EPI_ISL_501651 | A/Duck/Viet Nam/HU12-1467/2019 | A / H5N6 | Asia / Vietnam | |
| EPI_ISL_501652 | A/Duck/Viet Nam/HU12-1465/2019 | A / H5N6 | Asia / Vietnam | |
| EPI_ISL_501653 | A/Duck/Viet Nam/HU12-1471/2019 | A / H5N6 | Asia / Vietnam | |
| EPI_ISL_501654 | A/Duck/Viet Nam/HU12-1470/2019 | A / H5N6 | Asia / Vietnam | |
| EPI_ISL_501655 | A/Duck/Viet Nam/HU12-1472/2019 | A / H5N6 | Asia / Vietnam | |
| EPI_ISL_502069 | A/muscovy duck/Japan/AQ-HE30-77C1/2018 | A / H5N6 | Asia / Japan | |
| EPI_ISL_502070 | A/muscovy duck/Japan/AQ-HE30-77C2/2018 | A / H5N6 | Asia / Japan | |
| EPI_ISL_502201 | A/mink/Eastern China/006/2018 | A / H5N6 | Asia / China | |
| EPI_ISL_502202 | A/mink/Eastern China/032/2018 | A / H5N6 | Asia / China | |
| EPI_ISL_502203 | A/mink/Northern China/110/2018 | A / H5N6 | Asia / China | |
| EPI_ISL_502204 | A/mink/Eastern China/149/2018 | A / H5N6 | Asia / China | |
| EPI_ISL_502205 | A/mink/Eastern China/528/2018 | A / H5N6 | Asia / China | |
| EPI_ISL_502206 | A/mink/Eastern China/0824/2018 | A / H5N6 | Asia / China | |
| EPI_ISL_502207 | A/mink/Eastern China/0712/2018 | A / H5N1 | Asia / China | |
| EPI_ISL_502208 | A/mink/Eastern China/571/2018 | A / H5N6 | Asia / China | |
| EPI_ISL_502279 | A/chicken/Egypt/F15099/2018 | A / H5N8 | Africa / Egypt | |
| EPI_ISL_502280 | A/chicken/Egypt/N15173D/2018 | A / H5N8 | Africa / Egypt | |
| EPI_ISL_502393 | A/chicken/Taiwan/A3/2019 | A / H5N2 | Asia / Taiwan | |
| EPI_ISL_502394 | A/chicken/Taiwan/A4/2019 | A / H5N2 | Asia / Taiwan | |
| EPI_ISL_502395 | A/chicken/Taiwan/A5/2019 | A / H5N2 | Asia / Taiwan | |
| EPI_ISL_502396 | A/chicken/Taiwan/D2/2019 | A / H5N2 | Asia / Taiwan | |
| EPI_ISL_502397 | A/chicken/Taiwan/D9/2019 | A / H5N2 | Asia / Taiwan | |
| EPI_ISL_503010 | A/muscovy duck/Vietnam/LBM1021/2017 | A / H5N6 | Asia / Vietnam | |
| EPI_ISL_503012 | A/duck/Vietnam/LBM1099/2018 | A / H5N6 | Asia / Vietnam | |
| EPI_ISL_503013 | A/muscovy duck/Vietnam/LBM1101/2018 | A / H5N6 | Asia / Vietnam | |
| EPI_ISL_503015 | A/duck/Nha Trang/271/2018 | A / H5N6 | Asia / Vietnam / Tinh Khanh Hoa | |
| EPI_ISL_503016 | A/duck/Nha Trang/301/2018 | A / H5N6 | Asia / Vietnam / Tinh Khanh Hoa | |
| EPI_ISL_503017 | A/duck/Quang Ninh/371/2018 | A / H5N6 | Asia / Vietnam / Tinh Quang Ninh | |
| EPI_ISL_503018 | A/chicken/Japan/AQ-HE30-35C1/2018 | A / H5N2 | Asia / Japan | |
| EPI_ISL_503024 | A/duck/China/0936/2017 | A / H5N6 | Asia / China | |
| EPI_ISL_503031 | A/Anser fabalis/China/P563/2015 | A / H5N6 | Asia / China | |
| EPI_ISL_503036 | A/Anser cygnoides/China/P126/2015 | A / H5N1 | Asia / China | |
| EPI_ISL_503037 | A/swan/China/ST/2016 | A / H5N8 | Asia / China | |
| EPI_ISL_503058 | A/pigeon/Egypt/A15052/2018 | A / H5N8 | Africa / Egypt | |
| EPI_ISL_503150 | A/duck/Nigeria/SK28T_19VIR8424-2/2019 | A / H5N6 | Africa / Nigeria | |
| EPI_ISL_5034659 | A/turkey/Poland/100s3/2016 | A / H5N8 | Europe / Poland / Lubusz Voivodeship | Edyta ?wi?to? (National Veterinary Research Institut Poland, PIWet-PIB) |
| EPI_ISL_5034849 | A/domestic_duck/Poland/106/2017(H5N8) | A / H5N8 | Europe / Poland / Greater Poland | Edyta ?wi?to? (National Veterinary Research Institut Poland, PIWet-PIB) |
| EPI_ISL_5034850 | A/turkey/Poland/109s1/2016(H5N8) | A / H5N8 | Europe / Poland / Lubusz | Edyta ?wi?to? (National Veterinary Research Institut Poland, PIWet-PIB) |
| EPI_ISL_5034851 | A/chicken/Poland/115/2016(H5N8) | A / H5N8 | Europe / Poland / Lubusz | Edyta ?wi?to? (National Veterinary Research Institut Poland, PIWet-PIB) |
| EPI_ISL_5034852 | A/turkey/Poland/15s4/2017(H5N8) | A / H5N8 | Europe / Poland / Lubusz | Edyta ?wi?to? (National Veterinary Research Institut Poland, PIWet-PIB) |
| EPI_ISL_5034853 | A/domestic_goose/Poland/204/2017(H5N8) | A / H5N8 | Europe / Poland / Greater Poland | Edyta ?wi?to? (National Veterinary Research Institut Poland, PIWet-PIB) |
| EPI_ISL_5034854 | A/domestic_duck/Poland/238/2017(H5N8) | A / H5N8 | Europe / Poland / Greater Poland | Edyta ?wi?to? (National Veterinary Research Institut Poland, PIWet-PIB) |
| EPI_ISL_5034855 | A/turkey/Poland/78s1/2016(H5N8) | A / H5N8 | Europe / Poland / Lubusz | Edyta ?wi?to? (National Veterinary Research Institut Poland, PIWet-PIB) |
| EPI_ISL_5034856 | A/domestic_duck/Poland/90/2017(H5N8) | A / H5N8 | Europe / Poland / Greater Poland | Edyta ?wi?to? (National Veterinary Research Institut Poland, PIWet-PIB) |
| EPI_ISL_503594 | A/duck/Viet Nam/HU13-60/2019 | A / H5N6 | Asia / Vietnam | |
| EPI_ISL_503599 | A/duck/Viet Nam/HU12-970/2019 | A / H5N6 | Asia / Vietnam | |
| EPI_ISL_503600 | A/duck/Viet Nam/HU12-971/2019 | A / H5N6 | Asia / Vietnam | |
| EPI_ISL_503601 | A/duck/Viet Nam/HU12-972/2019 | A / H5N6 | Asia / Vietnam | |
| EPI_ISL_503602 | A/duck/Viet Nam/HU12-977/2019 | A / H5N6 | Asia / Vietnam | |
| EPI_ISL_503603 | A/duck/Viet Nam/HU12-978/2019 | A / H5N6 | Asia / Vietnam | |
| EPI_ISL_503604 | A/duck/Viet Nam/HU12-979/2019 | A / H5N6 | Asia / Vietnam | |
| EPI_ISL_503605 | A/duck/Viet Nam/HU12-980/2019 | A / H5N6 | Asia / Vietnam | |
| EPI_ISL_503606 | A/duck/Viet Nam/HU13-163/2019 | A / H5N6 | Asia / Vietnam | |
| EPI_ISL_503607 | A/duck/Viet Nam/HU13-164/2019 | A / H5N6 | Asia / Vietnam | |
| EPI_ISL_503608 | A/duck/Viet Nam/HU13-64/2019 | A / H5N6 | Asia / Vietnam | |
| EPI_ISL_503609 | A/duck/Viet Nam/HU13-65/2019 | A / H5N6 | Asia / Vietnam | |
| EPI_ISL_503610 | A/duck/Viet Nam/HU13-71/2019 | A / H5N6 | Asia / Vietnam | |
| EPI_ISL_503791 | A/Guinea fowl/Belgium/810/2017 | A / H5N8 | Europe / Belgium | |
| EPI_ISL_503819 | A/duck/Viet Nam/HU12-1473/2019 | A / H5N6 | Asia / Vietnam | |
| EPI_ISL_503935 | A/Teal/Dakahlia/VRLCU/2019 | A / H5N8 | Africa / Egypt / Dakahlia | |
| EPI_ISL_503968 | A/chicken/Bulgaria/217_20VIR1724-1/2020 | A / H5N8 | Europe / Bulgaria | |
| EPI_ISL_503969 | A/chicken/Bulgaria/77_20VIR1727/2020 | A / H5N2 | Europe / Bulgaria | |
| EPI_ISL_504004 | A/chicken/Quang Tri/MT11/2016 | A / H5N6 | Asia / Vietnam / Tinh Quang Tri | |
| EPI_ISL_504005 | A/duck/Ha Tinh/HT12/2014 | A / H5N6 | Asia / Vietnam | |
| EPI_ISL_504006 | A/duck/Ha Tinh/HT7/2014 | A / H5N6 | Asia / Vietnam | |
| EPI_ISL_504007 | A/chicken/Nha Trang/NT3/2017 | A / H5N6 | Asia / Vietnam | |
| EPI_ISL_5049590 | A/chicken/Luxembourg/21168413/2021 | A / H5N8 | Europe / Luxembourg / District de Grevenmacher | Chantal J. Snoeck (Luxembourg Institute of Health / Department of Infection and Immunity) |
| EPI_ISL_504984 | A/duck/Viet Nam/HU13-161/2019 | A / H5N6 | Asia / Vietnam | |
| EPI_ISL_504985 | A/duck/Viet Nam/HU13-162/2019 | A / H5N6 | Asia / Vietnam | |
| EPI_ISL_505073 | A/chicken/Sichuan/k141/2017 | A / H5N6 | Asia / China / Sichuan Province | |
| EPI_ISL_505396 | A/Duck/Giza/1814/2018 | A / H5N8 | Africa / Egypt / Giza | |

| | | | | |
|---|---|---|---|---|
| EPI_ISL_505397 | A/Duck/Al-Shaqiya/172AS/2017 | A / H5N8 | Africa / Egypt | |
| EPI_ISL_505398 | A/Duck/AL-Minia/1777FAO-S/2017 | A / H5N8 | Africa / Egypt | |
| EPI_ISL_505399 | A/chicken/Al-Sharqia/1822FM/2018 | A / H5N8 | Africa / Egypt | |
| EPI_ISL_505400 | A/chicken/Al-Minia/1785Fao-S/2017 | A / H5N8 | Africa / Egypt | |
| EPI_ISL_505401 | A/Duck/Ismalia/171Fao-SI/2017 | A / H5N8 | Africa / Egypt | |
| EPI_ISL_505402 | A/Duck/Ismalia/175Fao-SI/2017 | A / H5N8 | Africa / Egypt | |
| EPI_ISL_505403 | A/Duck/Ismalia/1719Fao-SI/2017 | A / H5N8 | Africa / Egypt | |
| EPI_ISL_505404 | A/chicken/Al Qaliobia/1755FAO-S/2017 | A / H5N8 | Africa / Egypt | |
| EPI_ISL_505405 | A/Duck/Ismalia/176FAO-SI/2017 | A / H5N8 | Africa / Egypt | |
| EPI_ISL_505406 | A/chicken/AL Suez/184/2018 | A / H5N8 | Africa / Egypt | |
| EPI_ISL_505407 | A/chicken/Cairo/1794FAO-S/2017 | A / H5N8 | Africa / Egypt / Cairo | |
| EPI_ISL_505408 | A/Duck/AL-Qaliobia/182Fao-s/2018 | A / H5N8 | Africa / Egypt | |
| EPI_ISL_505409 | A/Duck/Cairo/189Fao-S/2018 | A / H5N8 | Africa / Egypt / Cairo | |
| EPI_ISL_505410 | A/chicken/Al-Minia/183/2018 | A / H5N8 | Africa / Egypt | |
| EPI_ISL_505411 | A/Turkey/Beni Sueif/18296F/2018 | A / H5N8 | Africa / Egypt | |
| EPI_ISL_505412 | A/chicken/Giza/1836CAL/2017 | A / H5N8 | Africa / Egypt / Giza | |
| EPI_ISL_505413 | A/chicken/Albehra/18319F/2018 | A / H5N8 | Africa / Egypt | |
| EPI_ISL_505414 | A/chicken/Al- Minia/189CA/2018 | A / H5N8 | Africa / Egypt | |
| EPI_ISL_505415 | A/chicken/Giza/1810CA/2018 | A / H5N8 | Africa / Egypt / Giza | |
| EPI_ISL_505416 | A/Duck/Al-Monfia/1727/2017 | A / H5N8 | Africa / Egypt | |
| EPI_ISL_505417 | A/Duck/EL-WadiAlgidid/185/2018 | A / H5N8 | Africa / Egypt | |
| EPI_ISL_505418 | A/Duck/Cairo/187Fao-S/2018 | A / H5N8 | Africa / Egypt / Cairo | |
| EPI_ISL_505419 | A/Duck/Ismalia/1721Fao-SI/2017 | A / H5N8 | Africa / Egypt | |
| EPI_ISL_505420 | A/Duck/Al-Sharqiya/1733FM/2017 | A / H5N8 | Africa / Egypt | |
| EPI_ISL_505421 | A/Duck/Cairo/1895CA/2017 | A / H5N8 | Africa / Egypt / Cairo | |
| EPI_ISL_505422 | A/chicken/Al-Minia/188Fao-S/2018 | A / H5N8 | Africa / Egypt | |
| EPI_ISL_505423 | A/Duck/Giza/1754FAO-S/2017 | A / H5N8 | Africa / Egypt / Giza | |
| EPI_ISL_505431 | A/quail/South Africa/AI5930/2018 | A / H5N8 | Africa / South Africa | |
| EPI_ISL_505432 | A/chicken/South Africa/499723/2018 | A / H5N8 | Africa / South Africa | |
| EPI_ISL_505528 | A/duck/Saudi Arabia/KFU-HKU3616_2017/2017 | A / H5N8 | Asia / Saudi Arabia | |
| EPI_ISL_505529 | A/duck/Saudi Arabia/KFU-HKU3617_2017/2017 | A / H5N8 | Asia / Saudi Arabia | |
| EPI_ISL_505530 | A/duck/Saudi Arabia/KFU-HKU3618_2017/2017 | A / H5N8 | Asia / Saudi Arabia | |
| EPI_ISL_505531 | A/duck/Saudi Arabia/KFU-HKU3619_2017/2017 | A / H5N8 | Asia / Saudi Arabia | |
| EPI_ISL_505532 | A/ostrich/Saudi Arabia/KFU-HKU3590_2017/2017 | A / H5N8 | Asia / Saudi Arabia | |
| EPI_ISL_505533 | A/ostrich/Saudi Arabia/KFU-HKU3591_2017/2017 | A / H5N8 | Asia / Saudi Arabia | |
| EPI_ISL_505534 | A/ostrich/Saudi Arabia/KFU-HKU3592_2017/2017 | A / H5N8 | Asia / Saudi Arabia | |
| EPI_ISL_505535 | A/ostrich/Saudi Arabia/KFU-HKU3593_2017/2017 | A / H5N8 | Asia / Saudi Arabia | |
| EPI_ISL_505536 | A/ostrich/Saudi Arabia/KFU-HKU3594_2017/2017 | A / H5N8 | Asia / Saudi Arabia | |
| EPI_ISL_505537 | A/pigeon/Saudi Arabia/KFU-HKU3598_2017/2017 | A / H5N8 | Asia / Saudi Arabia | |
| EPI_ISL_505538 | A/pigeon/Saudi Arabia/KFU-HKU3599_2017/2017 | A / H5N8 | Asia / Saudi Arabia | |
| EPI_ISL_505539 | A/pigeon/Saudi Arabia/KFU-HKU3600_2017/2017 | A / H5N8 | Asia / Saudi Arabia | |
| EPI_ISL_5057844 | A/barnacle goose/Germany-SH/AI02168/2020 | A / H5N8 | Europe / Germany / Schleswig-Holstein / Nordfriesland | Jacqueline King (Friedrich-Loeffler-Institut) |
| EPI_ISL_5057856 | A/peregrine falcon/Germany-SH/AI02162/2020 | A / H5N8 | Europe / Germany / Schleswig-Holstein / Nordfriesland | Jacqueline King (Friedrich-Loeffler-Institut) |
| EPI_ISL_5057874 | A/barnacle goose/Germany-SH/AI02172/2020 | A / H5N8 | Europe / Germany / Schleswig-Holstein / Nordfriesland | Jacqueline King (Friedrich-Loeffler-Institut) |
| EPI_ISL_5057875 | A/Eurasian wigeon/Germany-SH/AI02176/2020 | A / H5N8 | Europe / Germany / Schleswig-Holstein / Nordfriesland | Jacqueline King (Friedrich-Loeffler-Institut) |
| EPI_ISL_5057876 | A/Eurasian wigeon/Germany-SH/AI02179/2020 | A / H5N8 | Europe / Germany / Schleswig-Holstein / Dithmarschen | Jacqueline King (Friedrich-Loeffler-Institut) |
| EPI_ISL_5057963 | A/barnacle goose/Germany-SH/AI02180/2020 | A / H5N8 | Europe / Germany / Schleswig-Holstein / Dithmarschen | Jacqueline King (Friedrich-Loeffler-Institut) |
| EPI_ISL_5058090 | A/herring gull/Germany-HH/AI02182/2020 | A / H5N8 | Europe / Germany / Hamburg / Hamburg | Jacqueline King (Friedrich-Loeffler-Institut) |
| EPI_ISL_5058262 | A/wild duck/Germany-SH/AI02189/2020 | A / H5N8 | Europe / Germany / Schleswig-Holstein / Nordfriesland | Jacqueline King (Friedrich-Loeffler-Institut) |
| EPI_ISL_5058432 | A/barnacle goose/Germany-SH/AI02190/2020 | A / H5N8 | Europe / Germany / Schleswig-Holstein / Nordfriesland | Jacqueline King (Friedrich-Loeffler-Institut) |
| EPI_ISL_5058630 | A/greylag goose/Germany-SH/AI02191/2020 | A / H5N8 | Europe / Germany / Schleswig-Holstein / Nordfriesland | Jacqueline King (Friedrich-Loeffler-Institut) |
| EPI_ISL_5058855 | A/wild goose/Germany-SH/AI02194/2020 | A / H5N8 | Europe / Germany / Schleswig-Holstein / Rendsburg-Eckernförde | Jacqueline King (Friedrich-Loeffler-Institut) |
| EPI_ISL_5060031 | A/barnacle goose/Germany-SH/AI02199/2020 | A / H5N8 | Europe / Germany / Schleswig-Holstein / Nordfriesland | Jacqueline King (Friedrich-Loeffler-Institut) |
| EPI_ISL_5061390 | A/greylag goose/Germany-SH/AI02207/2020 | A / H5N8 | Europe / Germany / Schleswig-Holstein / Nordfriesland | Jacqueline King (Friedrich-Loeffler-Institut) |
| EPI_ISL_5061617 | A/chicken/Germany-MV/AI02431/2020 | A / H5N5 | Europe / Germany / Schleswig-Holstein / Vorpommern-Rügen | Jacqueline King (Friedrich-Loeffler-Institut) |
| EPI_ISL_5061890 | A/domestic goose/Germany-SH/AI02884/2020 | A / H5N8 | Europe / Germany / Schleswig-Holstein / Nordfriesland | Jacqueline King (Friedrich-Loeffler-Institut) |
| EPI_ISL_5062078 | A/Eurasian oystercatcher/Germany-SH/AI02269/2020 | A / H5N8 | Europe / Germany / Schleswig-Holstein / Nordfriesland | Jacqueline King (Friedrich-Loeffler-Institut) |
| EPI_ISL_5062851 | A/herring gull/Germany-MV/AI02300/2020 | A / H5N8 | Europe / Germany / Mecklenburg-Vorpommern / Vorpommern-Rügen | Jacqueline King (Friedrich-Loeffler-Institut) |
| EPI_ISL_5063706 | A/barnacle goose/Germany-SH/AI02379/2020 | A / H5N8 | Europe / Germany / Schleswig-Holstein / Nordfriesland | Jacqueline King (Friedrich-Loeffler-Institut) |
| EPI_ISL_5064270 | A/turkey/Germany-NI/AI03453/2020 | A / H5N8 | Europe / Germany / Lower Saxony / Oldenburg | Jacqueline King (Friedrich-Loeffler-Institut) |
| EPI_ISL_5064509 | A/turkey/Germany-NI/AI03584/2020 | A / H5N8 | Europe / Germany / Lower Saxony / Cloppenburg | Jacqueline King (Friedrich-Loeffler-Institut) |
| EPI_ISL_5064707 | A/turkey/Germany-NI/AI03594/2020 | A / H5N8 | Europe / Germany / Lower Saxony / Cloppenburg | Jacqueline King (Friedrich-Loeffler-Institut) |
| EPI_ISL_5064825 | A/turkey/Germany-NI/AI03599/2020 | A / H5N8 | Europe / Germany / Lower Saxony / Cloppenburg | Jacqueline King (Friedrich-Loeffler-Institut) |
| EPI_ISL_5064922 | A/turkey/Germany-NI/AI03606/2020 | A / H5N8 | Europe / Germany / Lower Saxony / Cloppenburg | Jacqueline King (Friedrich-Loeffler-Institut) |
| EPI_ISL_5064971 | A/turkey/Germany-NI/AI03609/2020 | A / H5N8 | Europe / Germany / Lower Saxony / Cloppenburg | Jacqueline King (Friedrich-Loeffler-Institut) |
| EPI_ISL_5065068 | A/turkey/Germany-NI/AI00016/2021 | A / H5N8 | Europe / Germany / Lower Saxony / Cloppenburg | Jacqueline King (Friedrich-Loeffler-Institut) |
| EPI_ISL_5065144 | A/turkey/Germany-NI/AI00038/2021 | A / H5N8 | Europe / Germany / Lower Saxony / Cloppenburg | Jacqueline King (Friedrich-Loeffler-Institut) |
| EPI_ISL_5065276 | A/turkey/Germany-NI/AI00042/2021 | A / H5N8 | Europe / Germany / Lower Saxony / Cloppenburg | Jacqueline King (Friedrich-Loeffler-Institut) |
| EPI_ISL_5066981 | A/turkey/Germany-NI/AI03452/2020 | A / H5N8 | Europe / Germany / Lower Saxony / Oldenburg | Jacqueline King (Friedrich-Loeffler-Institut) |
| EPI_ISL_5066982 | A/turkey/Germany-NI/AI03654/2020 | A / H5N8 | Europe / Germany / Lower Saxony / Cloppenburg | Jacqueline King (Friedrich-Loeffler-Institut) |
| EPI_ISL_5066996 | A/turkey/Germany-NI/AI03432/2020 | A / H5N8 | Europe / Germany / Lower Saxony / Cloppenburg | Jacqueline King (Friedrich-Loeffler-Institut) |
| EPI_ISL_5067044 | A/turkey/Germany-NI/AI00063/2021 | A / H5N8 | Europe / Germany / Lower Saxony / Oldenburg | Jacqueline King (Friedrich-Loeffler-Institut) |

| | | | | |
|---|---|---|---|---|
| EPI_ISL_5067335 | A/turkey/Germany-NI/AI00072/2021 | A / H5N8 | Europe / Germany / Lower Saxony / Cloppenburg | Jacqueline King (Friedrich-Loeffler-Institut) |
| EPI_ISL_5067957 | A/domestic duck/Germany-NI/AI00079/2021 | A / H5N8 | Europe / Germany / Lower Saxony / Cloppenburg | Jacqueline King (Friedrich-Loeffler-Institut) |
| EPI_ISL_5095319 | A/turkey/Germany-NI/AI00402/2021 | A / H5N8 | Europe / Germany / Lower Saxony / Cloppenburg | Jacqueline King (Friedrich-Loeffler-Institut) |
| EPI_ISL_5095320 | A/turkey/Germany-NI/AI00406/2021 | A / H5N8 | Europe / Germany / Lower Saxony / Cloppenburg | Jacqueline King (Friedrich-Loeffler-Institut) |
| EPI_ISL_5095321 | A/turkey/Germany-NI/AI00429/2021 | A / H5N8 | Europe / Germany / Lower Saxony / Cloppenburg | Jacqueline King (Friedrich-Loeffler-Institut) |
| EPI_ISL_5095322 | A/turkey/Germany-NI/AI00439/2021 | A / H5N8 | Europe / Germany / Lower Saxony / Cloppenburg | Jacqueline King (Friedrich-Loeffler-Institut) |
| EPI_ISL_5095323 | A/wild goose/Germany-NI/AI03220/2020 | A / H5N8 | Europe / Germany / Lower Saxony / Friesland | Jacqueline King (Friedrich-Loeffler-Institut) |
| EPI_ISL_5095336 | A/wild goose/Germany-NI/AI03473/2020 | A / H5N8 | Europe / Germany / Lower Saxony / Wilhelmshaven, Stadt | Jacqueline King (Friedrich-Loeffler-Institut) |
| EPI_ISL_5095554 | A/chicken/Germany-MV/AI00539/2021 | A / H5N8 | Europe / Germany / Mecklenburg-Vorpommern / Landkreis Rostock | Jacqueline King (Friedrich-Loeffler-Institut) |
| EPI_ISL_5095555 | A/chicken/Germany-NI/AI00547/2021 | A / H5N8 | Europe / Germany / Lower Saxony / Oldenburg | Jacqueline King (Friedrich-Loeffler-Institut) |
| EPI_ISL_5095556 | A/turkey/Germany-NI/AI00589/2021 | A / H5N8 | Europe / Germany / Lower Saxony / Cuxhaven | Jacqueline King (Friedrich-Loeffler-Institut) |
| EPI_ISL_5095644 | A/turkey/Germany-BB/AI00868/2021 | A / H5N8 | Europe / Germany / Brandenburg / Prignitz | Jacqueline King (Friedrich-Loeffler-Institut) |
| EPI_ISL_5095645 | A/turkey/Germany-MV/AI00909/2021 | A / H5N8 | Europe / Germany / Mecklenburg-Vorpommern / Nordwestmecklenburg | Jacqueline King (Friedrich-Loeffler-Institut) |
| EPI_ISL_5095649 | A/turkey/Germany-MV/AI01015/2021 | A / H5N8 | Europe / Germany / Mecklenburg-Vorpommern / Ludwigslust-Parchim | Jacqueline King (Friedrich-Loeffler-Institut) |
| EPI_ISL_5095650 | A/turkey/Germany-BB/AI01023/2021 | A / H5N8 | Europe / Germany / Brandenburg / Uckermark | Jacqueline King (Friedrich-Loeffler-Institut) |
| EPI_ISL_5096070 | A/mallard/Germany-MV/AI00639/2021 | A / H5N8 | Europe / Germany / Mecklenburg-Vorpommern / Vorpommern-Greifswald | Jacqueline King (Friedrich-Loeffler-Institut) |
| EPI_ISL_5096104 | A/wild goose/Germany-NI/AI00855/2021 | A / H5N8 | Europe / Germany / Lower Saxony / Stade | Jacqueline King (Friedrich-Loeffler-Institut) |
| EPI_ISL_5096105 | A/domestic duck/Germany-NI/AI00646/2021 | A / H5N8 | Europe / Germany / Lower Saxony / Wittmund | Jacqueline King (Friedrich-Loeffler-Institut) |
| EPI_ISL_5096107 | A/turkey/Germany-NI/AI00647/2021 | A / H5N8 | Europe / Germany / Lower Saxony / Oldenburg | Jacqueline King (Friedrich-Loeffler-Institut) |
| EPI_ISL_5096629 | A/wild goose/Germany-NI/AI00625/2021 | A / H5N8 | Europe / Germany / Lower Saxony / Cuxhaven | Jacqueline King (Friedrich-Loeffler-Institut) |
| EPI_ISL_5097280 | A/turkey/Germany-NI/AI00621/2021 | A / H5N8 | Europe / Germany / Lower Saxony / Cuxhaven | Jacqueline King (Friedrich-Loeffler-Institut) |
| EPI_ISL_5098130 | A/turkey/Germany-NI/AI00612/2021 | A / H5N8 | Europe / Germany / Lower Saxony / Cloppenburg | Jacqueline King (Friedrich-Loeffler-Institut) |
| EPI_ISL_5098132 | A/turkey/Germany-NI/AI00616/2021 | A / H5N8 | Europe / Germany / Lower Saxony / Cloppenburg | Jacqueline King (Friedrich-Loeffler-Institut) |
| EPI_ISL_5098133 | A/turkey/Germany-NI/AI00591/2021 | A / H5N8 | Europe / Germany / Lower Saxony / Cloppenburg | Jacqueline King (Friedrich-Loeffler-Institut) |
| EPI_ISL_5098134 | A/wild duck/Germany-NI/AI03479/2020 | A / H5N8 | Europe / Germany / Lower Saxony / Lüneburg | Jacqueline King (Friedrich-Loeffler-Institut) |
| EPI_ISL_5098135 | A/kestrel/Germany-NI/AI03672/2020 | A / H5N8 | Europe / Germany / Lower Saxony / Friesland | Jacqueline King (Friedrich-Loeffler-Institut) |
| EPI_ISL_5098138 | A/wild goose/Germany-NI/AI00626/2021 | A / H5N8 | Europe / Germany / Lower Saxony / Harburg | Jacqueline King (Friedrich-Loeffler-Institut) |
| EPI_ISL_5098139 | A/wild goose/Germany-NI/AI00854/2021 | A / H5N8 | Europe / Germany / Lower Saxony / Lüchow-Dannenberg | Jacqueline King (Friedrich-Loeffler-Institut) |
| EPI_ISL_5098142 | A/Hawaiian goose/Germany-RP/AI00856/2021 | A / H5N8 | Europe / Germany / Rhineland-Palatinate / Rhein-Pfalz-Kreis | Jacqueline King (Friedrich-Loeffler-Institut) |
| EPI_ISL_5098143 | A/chicken/Germany-NI/AI00887/2021 | A / H5N8 | Europe / Germany / Lower Saxony / Cloppenburg | Jacqueline King (Friedrich-Loeffler-Institut) |
| EPI_ISL_5098144 | A/pomeranian goose/Germany-MV/AI01100/2021 | A / H5N8 | Europe / Germany / Mecklenburg-Vorpommern / Vorpommern-Rügen | Jacqueline King (Friedrich-Loeffler-Institut) |
| EPI_ISL_5098145 | A/turkey/Germany-BB/AI01120/2021 | A / H5N8 | Europe / Germany / Brandenburg / Uckermark | Jacqueline King (Friedrich-Loeffler-Institut) |
| EPI_ISL_5098146 | A/chicken/Germany-MV/AI01347/2021 | A / H5N8 | Europe / Germany / Mecklenburg-Vorpommern / Landkreis Rostock | Jacqueline King (Friedrich-Loeffler-Institut) |
| EPI_ISL_5098147 | A/wild goose/Germany-NI/AI03471/2020 | A / H5N8 | Europe / Germany / Lower Saxony / Friesland | Jacqueline King (Friedrich-Loeffler-Institut) |
| EPI_ISL_5098153 | A/wild goose/Germany-NI/AI03101/2020 | A / H5N8 | Europe / Germany / Lower Saxony / Hannover | Jacqueline King (Friedrich-Loeffler-Institut) |
| EPI_ISL_5098154 | A/turkey/Germany-MV/AI01127/2021 | A / H5N8 | Europe / Germany / Mecklenburg-Vorpommern / Vorpommern-Rügen | Jacqueline King (Friedrich-Loeffler-Institut) |
| EPI_ISL_5098158 | A/turkey/Germany-BB/AI01419/2021 | A / H5N8 | Europe / Germany / Brandenburg / Prignitz | Jacqueline King (Friedrich-Loeffler-Institut) |
| EPI_ISL_5098159 | A/domestic duck/Germany-BB/AI01423/2021 | A / H5N8 | Europe / Germany / Brandenburg / Märkisch-Oderland | Jacqueline King (Friedrich-Loeffler-Institut) |
| EPI_ISL_5098160 | A/domestic duck/Germany-BB/AI01437/2021 | A / H5N8 | Europe / Germany / Brandenburg / Uckermark | Jacqueline King (Friedrich-Loeffler-Institut) |
| EPI_ISL_5099356 | A/domestic duck/Germany-BB/AI01444/2021 | A / H5N8 | Europe / Germany / Brandenburg / Märkisch-Oderland | Jacqueline King (Friedrich-Loeffler-Institut) |
| EPI_ISL_5099367 | A/chicken/Germany-BY/AI01355/2021 | A / H5N8 | Europe / Germany / Bavaria / Weißenburg-Gunzenhausen | Jacqueline King (Friedrich-Loeffler-Institut) |
| EPI_ISL_5099368 | A/turkey/Germany-MV/AI01477/2021 | A / H5N8 | Europe / Germany / Mecklenburg-Vorpommern / Vorpommern-Rügen | Jacqueline King (Friedrich-Loeffler-Institut) |
| EPI_ISL_5099372 | A/Laridae/Germany-SH/AI01498/2021 | A / H5N4 | Europe / Germany / Schleswig-Holstein / Ostholstein | Jacqueline King (Friedrich-Loeffler-Institut) |
| EPI_ISL_5099373 | A/chicken/Germany-MV/AI01587/2021 | A / H5N8 | Europe / Germany / Mecklenburg-Vorpommern / Landkreis Rostock | Jacqueline King (Friedrich-Loeffler-Institut) |
| EPI_ISL_5099451 | A/chicken/Germany-NI/AI01599/2021 | A / H5N1 | Europe / Germany / Lower Saxony / Aurich | Jacqueline King (Friedrich-Loeffler-Institut) |
| EPI_ISL_5099456 | A/barnacle goose/Germany-NI/AI01605/2021 | A / H5N1 | Europe / Germany / Lower Saxony / Aurich | Jacqueline King (Friedrich-Loeffler-Institut) |
| EPI_ISL_5099462 | A/chicken/Germany-BY/AI01617/2021 | A / H5N8 | Europe / Germany / Bavaria / Würzburg | Jacqueline King (Friedrich-Loeffler-Institut) |
| EPI_ISL_5099463 | A/chicken/Germany-MV/AI01794/2021 | A / H5N8 | Europe / Germany / Mecklenburg-Vorpommern / Vorpommern-Greifswald | Jacqueline King (Friedrich-Loeffler-Institut) |
| EPI_ISL_5099468 | A/turkey/Germany-NI/AI01799/2021 | A / H5N8 | Europe / Germany / Lower Saxony / Cloppenburg | Jacqueline King (Friedrich-Loeffler-Institut) |
| EPI_ISL_5099474 | A/turkey/Germany-NI/AI01805/2021 | A / H5N8 | Europe / Germany / Lower Saxony / Cloppenburg | Jacqueline King (Friedrich-Loeffler-Institut) |
| EPI_ISL_5099476 | A/chicken/Germany-NI/AI01815/2021 | A / H5N8 | Europe / Germany / Lower Saxony / Vechta | Jacqueline King (Friedrich-Loeffler-Institut) |
| EPI_ISL_5099628 | A/domestic duck/Germany-BB/AI01443/2021 | A / H5N8 | Europe / Germany / Brandenburg / Märkisch-Oderland | Jacqueline King (Friedrich-Loeffler-Institut) |
| EPI_ISL_5100056 | A/turkey/Germany-NI/AI02013/2021 | A / H5N8 | Europe / Germany / Lower Saxony / Diepholz | Jacqueline King (Friedrich-Loeffler-Institut) |
| EPI_ISL_5100599 | A/turkey/Germany-NI/AI02025/2021 | A / H5N8 | Europe / Germany / Lower Saxony / Cloppenburg | Jacqueline King (Friedrich-Loeffler-Institut) |
| EPI_ISL_5102033 | A/domestic duck/Germany-NW/AI02049/2021 | A / H5N8 | Europe / Germany / North Rhine-Westphalia / Gütersloh | Jacqueline King (Friedrich-Loeffler-Institut) |
| EPI_ISL_5102130 | A/chicken/Germany-NW/AI02054/2021 | A / H5N8 | Europe / Germany / North Rhine-Westphalia / Paderborn | Jacqueline King (Friedrich-Loeffler-Institut) |
| EPI_ISL_5102132 | A/chicken/Germany-BY/AI02064/2021 | A / H5N8 | Europe / Germany / Bavaria / Würzburg | Jacqueline King (Friedrich-Loeffler-Institut) |
| EPI_ISL_5102136 | A/chicken/Germany-BY/AI02066/2021 | A / H5N8 | Europe / Germany / Bavaria / Schwandorf | Jacqueline King (Friedrich-Loeffler-Institut) |
| EPI_ISL_5102449 | A/white-tailed eagle/Germany-SH/AI02170/2020 | A / H5N8 | Europe / Germany / Schleswig-Holstein / Nordfriesland | Jacqueline King (Friedrich-Loeffler-Institut) |
| EPI_ISL_5102578 | A/chicken/Germany-MV/AI01407/2021 | A / H5N8 | Europe / Germany / Mecklenburg-Vorpommern / Mecklenburgische Seenplatte | Jacqueline King (Friedrich-Loeffler-Institut) |
| EPI_ISL_5115613 | A/chicken/Germany-MV/AI01965/2021 | A / H5N8 | Europe / Germany / Mecklenburg-Vorpommern / Mecklenburgische Seenplatte | Jacqueline King (Friedrich-Loeffler-Institut) |
| EPI_ISL_5115614 | A/chicken/Germany-BE/AI01974/2021 | A / H5N8 | Europe / Germany / Berlin / Berlin,Stadt | Jacqueline King (Friedrich-Loeffler-Institut) |
| EPI_ISL_5115615 | A/turkey/Germany-BB/AI01980/2021 | A / H5N8 | Europe / Germany / Brandenburg / Prignitz | Jacqueline King (Friedrich-Loeffler-Institut) |
| EPI_ISL_5115616 | A/domestic goose/Germany-SH/AI02100/2021 | A / H5N8 | Europe / Germany / Schleswig-Holstein / Rendsburg-Eckernförde | Jacqueline King (Friedrich-Loeffler-Institut) |
| EPI_ISL_5115617 | A/domestic goose/Germany-SH/AI02102/2021 | A / H5N8 | Europe / Germany / Schleswig-Holstein / Stormarn | Jacqueline King (Friedrich-Loeffler-Institut) |
| EPI_ISL_5115618 | A/turkey/Germany-BB/AI02117/2021 | A / H5N8 | Europe / Germany / Brandenburg / Märkisch-Oderland | Jacqueline King (Friedrich-Loeffler-Institut) |
| EPI_ISL_5115619 | A/turkey/Germany-NI/AI02122/2021 | A / H5N8 | Europe / Germany / Lower Saxony / Diepholz | Jacqueline King (Friedrich-Loeffler-Institut) |
| EPI_ISL_5115620 | A/turkey/Germany-NI/AI02128/2021 | A / H5N8 | Europe / Germany / Lower Saxony / Cloppenburg | Jacqueline King (Friedrich-Loeffler-Institut) |
| EPI_ISL_5115621 | A/chicken/Germany-MV/AI02141/2021 | A / H5N8 | Europe / Germany / Mecklenburg-Vorpommern / Vorpommern-Greifswald | Jacqueline King (Friedrich-Loeffler-Institut) |
| EPI_ISL_5115763 | A/turkey/Germany-NW/AI02290/2021 | A / H5N8 | Europe / Germany / North Rhine-Westphalia / Minden-Lübbecke | Jacqueline King (Friedrich-Loeffler-Institut) |
| EPI_ISL_5115955 | A/turkey/Germany-NI/AI02303/2021 | A / H5N8 | Europe / Germany / Lower Saxony / Cloppenburg | Jacqueline King (Friedrich-Loeffler-Institut) |
| EPI_ISL_5115956 | A/turkey/Germany-NI/AI02306/2021 | A / H5N8 | Europe / Germany / Lower Saxony / Cloppenburg | Jacqueline King (Friedrich-Loeffler-Institut) |

| | | | | |
|---|---|---|---|---|
| EPI_ISL_5115967 | A/chicken/Germany-SH/AI02312/2021 | A / H5N8 | Europe / Germany / Schleswig-Holstein / Vechta Plön | Jacqueline King (Friedrich-Loeffler-Institut) |
| EPI_ISL_5115968 | A/turkey/Germany-BB/AI02318/2021 | A / H5N8 | Europe / Germany / Brandenburg / Märkisch-Oderland | Jacqueline King (Friedrich-Loeffler-Institut) |
| EPI_ISL_5115969 | A/domestic goose/Germany-MV/AI02319/2021 | A / H5N8 | Europe / Germany / Mecklenburg-Vorpommern / Vorpommern-Rügen | Jacqueline King (Friedrich-Loeffler-Institut) |
| EPI_ISL_5115970 | A/chicken/Germany-NI/AI02328/2021 | A / H5N8 | Europe / Germany / Lower Saxony / Vechta | Jacqueline King (Friedrich-Loeffler-Institut) |
| EPI_ISL_5116063 | A/swan/Germany-HE/AI02335/2021 | A / H5N8 | Europe / Germany / Hessen / Offenbach a. Main | Jacqueline King (Friedrich-Loeffler-Institut) |
| EPI_ISL_5116064 | A/turkey/Germany-NI/AI02349/2021 | A / H5N8 | Europe / Germany / Lower Saxony / Cloppenburg | Jacqueline King (Friedrich-Loeffler-Institut) |
| EPI_ISL_5116065 | A/greater rhea/Germany-SN/AI02351/2021 | A / H5N8 | Europe / Germany / Saxony / Zwickau | Jacqueline King (Friedrich-Loeffler-Institut) |
| EPI_ISL_5116093 | A/chicken/Germany-SN/AI02353/2021 | A / H5N8 | Europe / Germany / Saxony / Mittelsachsen | Jacqueline King (Friedrich-Loeffler-Institut) |
| EPI_ISL_5116172 | A/mute swan/Germany-HE/AI02373/2021 | A / H5N8 | Europe / Germany / Hessen / Frankfurt a. Main | Jacqueline King (Friedrich-Loeffler-Institut) |
| EPI_ISL_5116326 | A/chicken/Germany-MV/AI02376/2021 | A / H5N8 | Europe / Germany / Mecklenburg-Vorpommern / Cloppenburg Vorpommern-Rügen | Jacqueline King (Friedrich-Loeffler-Institut) |
| EPI_ISL_5123578 | A/domestic goose/Germany-NI/AI03093/2021 | A / H5N8 | Europe / Germany / Lower Saxony / Vechta | Jacqueline King (Friedrich-Loeffler-Institut) |
| EPI_ISL_5123579 | A/domestic duck/Germany-NI/AI03099/2021 | A / H5N8 | Europe / Germany / Lower Saxony / Cloppenburg | Jacqueline King (Friedrich-Loeffler-Institut) |
| EPI_ISL_5123584 | A/turkey/Germany-NI/AI03126/2021 | A / H5N8 | Europe / Germany / Lower Saxony / Vechta | Jacqueline King (Friedrich-Loeffler-Institut) |
| EPI_ISL_5123589 | A/turkey/Germany-NI/AI02615/2021 | A / H5N8 | Europe / Germany / Lower Saxony / Cloppenburg | Jacqueline King (Friedrich-Loeffler-Institut) |
| EPI_ISL_5123590 | A/domestic duck/Germany-BB/AI02599/2021 | A / H5N8 | Europe / Germany / Brandenburg / Oder-Spree | Jacqueline King (Friedrich-Loeffler-Institut) |
| EPI_ISL_5123847 | A/chicken/Germany-SH/AI02562/2021 | A / H5N8 | Europe / Germany / Schleswig-Holstein / Plön | Jacqueline King (Friedrich-Loeffler-Institut) |
| EPI_ISL_5126214 | A/eagle owl/Germany-ST/AI02542/2021 | A / H5N8 | Europe / Germany / Saxony-Anhalt / Altmarkkreis Salzwedel | Jacqueline King (Friedrich-Loeffler-Institut) |
| EPI_ISL_5133724 | A/chicken/Germany-NI/AI02954/2021 | A / H5N8 | Europe / Germany / Lower Saxony / Diepholz | Jacqueline King (Friedrich-Loeffler-Institut) |
| EPI_ISL_5134186 | A/turkey/Germany-NI/AI02950/2021 | A / H5N8 | Europe / Germany / Lower Saxony / Vechta | Jacqueline King (Friedrich-Loeffler-Institut) |
| EPI_ISL_5134658 | A/turkey/Germany-NI/AI02943/2021 | A / H5N8 | Europe / Germany / Lower Saxony / Ammerland | Jacqueline King (Friedrich-Loeffler-Institut) |
| EPI_ISL_5135087 | A/chicken/Germany-NI/AI02942/2021 | A / H5N8 | Europe / Germany / Lower Saxony / Cloppenburg | Jacqueline King (Friedrich-Loeffler-Institut) |
| EPI_ISL_5136192 | A/turkey/Germany-NI/AI02933/2021 | A / H5N8 | Europe / Germany / Lower Saxony / Cloppenburg | Jacqueline King (Friedrich-Loeffler-Institut) |
| EPI_ISL_5136837 | A/turkey/Germany-NI/AI02671/2021 | A / H5N8 | Europe / Germany / Lower Saxony / Cloppenburg | Jacqueline King (Friedrich-Loeffler-Institut) |
| EPI_ISL_5137351 | A/turkey/Germany-NI/AI02678/2021 | A / H5N8 | Europe / Germany / Lower Saxony / Cloppenburg | Jacqueline King (Friedrich-Loeffler-Institut) |
| EPI_ISL_5137952 | A/turkey/Germany-NI/AI02674/2021 | A / H5N8 | Europe / Germany / Lower Saxony / Cloppenburg | Jacqueline King (Friedrich-Loeffler-Institut) |
| EPI_ISL_5139441 | A/turkey/Germany-NI/AI02663/2021 | A / H5N8 | Europe / Germany / Lower Saxony / Vechta | Jacqueline King (Friedrich-Loeffler-Institut) |
| EPI_ISL_5141340 | A/domestic duck/Germany-NI/AI02660/2021 | A / H5N8 | Europe / Germany / Lower Saxony / Vechta | Jacqueline King (Friedrich-Loeffler-Institut) |
| EPI_ISL_5142451 | A/turkey/Germany-NW/AI03104/2021 | A / H5N8 | Europe / Germany / North Rhine-Westphalia / Hochsauerlandkreis | Jacqueline King (Friedrich-Loeffler-Institut) |
| EPI_ISL_5142455 | A/turkey/Germany-NI/AI02991/2021 | A / H5N8 | Europe / Germany / Lower Saxony / Cloppenburg | Jacqueline King (Friedrich-Loeffler-Institut) |
| EPI_ISL_5142488 | A/chicken/Germany-ST/AI02967/2021 | A / H5N8 | Europe / Germany / Saxony-Anhalt / Altmarkkreis Salzwedel | Jacqueline King (Friedrich-Loeffler-Institut) |
| EPI_ISL_5142489 | A/turkey/Germany-NI/AI02553/2021 | A / H5N8 | Europe / Germany / Lower Saxony / Cloppenburg | Jacqueline King (Friedrich-Loeffler-Institut) |
| EPI_ISL_5142894 | A/chicken/Germany-NI/AI02543/2021 | A / H5N8 | Europe / Germany / Lower Saxony / Wesermarsch | Jacqueline King (Friedrich-Loeffler-Institut) |
| EPI_ISL_5142961 | A/turkey/Germany-MV/AI02458/2021 | A / H5N8 | Europe / Germany / Mecklenburg-Vorpommern / Vorpommern-Rügen | Jacqueline King (Friedrich-Loeffler-Institut) |
| EPI_ISL_5143100 | A/turkey/Germany-NI/AI02424/2021 | A / H5N8 | Europe / Germany / Lower Saxony / Cloppenburg | Jacqueline King (Friedrich-Loeffler-Institut) |
| EPI_ISL_5143300 | A/domestic goose/Germany-MV/AI02558/2021 | A / H5N8 | Europe / Germany / Mecklenburg-Vorpommern / Vorpommern-Rügen | Jacqueline King (Friedrich-Loeffler-Institut) |
| EPI_ISL_5143417 | A/turkey/Germany-BB/AI02434/2021 | A / H5N8 | Europe / Germany / Brandenburg / Märkisch-Oderland | Jacqueline King (Friedrich-Loeffler-Institut) |
| EPI_ISL_5143554 | A/chicken/Germany-NI/AI02412/2021 | A / H5N8 | Europe / Germany / Lower Saxony / Cuxhaven | Jacqueline King (Friedrich-Loeffler-Institut) |
| EPI_ISL_5143583 | A/chicken/Germany-TH/AI03513/2021 | A / H5N8 | Europe / Germany / Thuringia / Sömmerda | Jacqueline King (Friedrich-Loeffler-Institut) |
| EPI_ISL_5143584 | A/chicken/Germany-BY/AI03544/2021 | A / H5N8 | Europe / Germany / Bavaria / Ebersberg | Jacqueline King (Friedrich-Loeffler-Institut) |
| EPI_ISL_5143585 | A/turkey/Germany-MV/AI03535/2021 | A / H5N8 | Europe / Germany / Mecklenburg-Vorpommern / Landkreis Rostock | Jacqueline King (Friedrich-Loeffler-Institut) |
| EPI_ISL_5143586 | A/chicken/Germany-SN/AI03496/2021 | A / H5N8 | Europe / Germany / Saxony / Mittelsachsen | Jacqueline King (Friedrich-Loeffler-Institut) |
| EPI_ISL_5143587 | A/chicken/Germany-SN/AI03502/2021 | A / H5N8 | Europe / Germany / Saxony / Vogtlandkreis | Jacqueline King (Friedrich-Loeffler-Institut) |
| EPI_ISL_5143626 | A/chicken/Germany-TH/AI03193/2021 | A / H5N8 | Europe / Germany / Thuringia / Weimar-Land | Jacqueline King (Friedrich-Loeffler-Institut) |
| EPI_ISL_5143627 | A/turkey/Germany-NI/AI03209/2021 | A / H5N8 | Europe / Germany / Lower Saxony / Cloppenburg | Jacqueline King (Friedrich-Loeffler-Institut) |
| EPI_ISL_5143820 | A/chicken/Germany-BW/AI03205/2021 | A / H5N8 | Europe / Germany / Baden-Wuerttemberg / Breisgau-Hochschwarzwald | Jacqueline King (Friedrich-Loeffler-Institut) |
| EPI_ISL_5143995 | A/domestic duck/Germany-ST/AI03183/2021 | A / H5N8 | Europe / Germany / Saxony-Anhalt / Saalekreis | Jacqueline King (Friedrich-Loeffler-Institut) |
| EPI_ISL_5144149 | A/turkey/Germany-NI/AI03160/2021 | A / H5N8 | Europe / Germany / Lower Saxony / Cloppenburg | Jacqueline King (Friedrich-Loeffler-Institut) |
| EPI_ISL_5144341 | A/chicken/Germany-NW/AI03148/2021 | A / H5N8 | Europe / Germany / North Rhine-Westphalia / Warendorf | Jacqueline King (Friedrich-Loeffler-Institut) |
| EPI_ISL_5144368 | A/domestic duck/Germany-NI/AI03088/2021 | A / H5N8 | Europe / Germany / Lower Saxony / Cloppenburg | Jacqueline King (Friedrich-Loeffler-Institut) |
| EPI_ISL_5144407 | A/chicken/Germany-NW/AI03154/2021 | A / H5N8 | Europe / Germany / North Rhine-Westphalia / Paderborn | Jacqueline King (Friedrich-Loeffler-Institut) |
| EPI_ISL_5144529 | A/domestic goose/Germany-NI/AI03142/2021 | A / H5N8 | Europe / Germany / Lower Saxony / Vechta | Jacqueline King (Friedrich-Loeffler-Institut) |
| EPI_ISL_5144632 | A/turkey/Germany-NI/AI02393/2021 | A / H5N8 | Europe / Germany / Lower Saxony / Cloppenburg | Jacqueline King (Friedrich-Loeffler-Institut) |
| EPI_ISL_5144725 | A/domestic goose/Germany-NI/AI02980/2021 | A / H5N8 | Europe / Germany / Lower Saxony / Vechta | Jacqueline King (Friedrich-Loeffler-Institut) |
| EPI_ISL_5144778 | A/turkey/Germany-NI/AI01488/2021 | A / H5N8 | Europe / Germany / Lower Saxony / Cloppenburg | Jacqueline King (Friedrich-Loeffler-Institut) |
| EPI_ISL_5144920 | A/turkey/Germany-NI/AI01483/2021 | A / H5N8 | Europe / Germany / Lower Saxony / Diepholz | Jacqueline King (Friedrich-Loeffler-Institut) |
| EPI_ISL_5145223 | A/turkey/Germany-NI/AI04455/2021 | A / H5N1 | Europe / Germany / Lower Saxony / Emsland | Jacqueline King (Friedrich-Loeffler-Institut) |
| EPI_ISL_5145422 | A/barnacle goose/Germany-NI/AI03914/2021 | A / H5N1 | Europe / Germany / Lower Saxony / Leer | Jacqueline King (Friedrich-Loeffler-Institut) |
| EPI_ISL_5145613 | A/chicken/Germany-BY/AI02060/2021 | A / H5N8 | Europe / Germany / Bavaria / Würzburg | Jacqueline King (Friedrich-Loeffler-Institut) |
| EPI_ISL_5145744 | A/oystercatcher/Germany-NI/AI05047/2021 | A / H5N1 | Europe / Germany / Lower Saxony / Aurich | Jacqueline King (Friedrich-Loeffler-Institut) |
| EPI_ISL_5145796 | A/swan/Germany-NI/AI05045/2021 | A / H5N8 | Europe / Germany / Lower Saxony / Osnabrück | Jacqueline King (Friedrich-Loeffler-Institut) |
| EPI_ISL_5145858 | A/chicken/Germany-TH/AI03512/2021 | A / H5N8 | Europe / Germany / Thuringia / Saale-Holzland-Kreis | Jacqueline King (Friedrich-Loeffler-Institut) |
| EPI_ISL_5145921 | A/chicken/Germany-BW/AI03554/2021 | A / H5N8 | Europe / Germany / Baden-Wuerttemberg / Böblingen | Jacqueline King (Friedrich-Loeffler-Institut) |
| EPI_ISL_5146171 | A/chicken/Germany-BW/AI03634/2021 | A / H5N8 | Europe / Germany / Baden-Wuerttemberg / Rottweil | Jacqueline King (Friedrich-Loeffler-Institut) |
| EPI_ISL_5146288 | A/chicken/Germany-NW/AI03705/2021 | A / H5N8 | Europe / Germany / North Rhine-Westphalia / Märkischer Kreis | Jacqueline King (Friedrich-Loeffler-Institut) |
| EPI_ISL_5146291 | A/barnacle goose/Germany-NI/AI03980/2021 | A / H5N1 | Europe / Germany / Lower Saxony / Leer | Jacqueline King (Friedrich-Loeffler-Institut) |
| EPI_ISL_5146292 | A/turkey/Germany-NI/AI04373/2021 | A / H5N1 | Europe / Germany / Lower Saxony / Emsland | Jacqueline King (Friedrich-Loeffler-Institut) |
| EPI_ISL_5146481 | A/turkey/Germany-NI/AI04425/2021 | A / H5N1 | Europe / Germany / Lower Saxony / Emsland | Jacqueline King (Friedrich-Loeffler-Institut) |
| EPI_ISL_5159426 | A/Hunan/10117/2021 | A / H5N6 | Asia / China / Hunan | Lei Yang (WHO Chinese National Influenza Center / Virology Institute, Chinese CDC) |
| EPI_ISL_5234711 | A/duck/Laos/NL-2175410/2021 | A / H5N6 | Asia / Lao, People's Democratic Republic | Yunho Jang (Centers for Disease Control and Prevention / WHO Collaborating Center for Surveillance, Epidemiology and Control of Influ |
| EPI_ISL_5234724 | A/duck/Laos/NL-2175411/2021 | A / H5N6 | Asia / Lao, People's Democratic Republic | Yunho Jang (Centers for Disease Control and Prevention / WHO Collaborating Center for Surveillance, Epidemiology and Control of Influ |
| EPI_ISL_5234816 | A/duck/Laos/NL-2175412/2021 | A / H5N6 | Asia / Lao, People's Democratic Republic | Yunho Jang (Centers for Disease Control and Prevention / WHO Collaborating Center for Surveillance, Epidemiology and Control of Influ |
| EPI_ISL_525439 | A/laying_hen/Poland/002/2020 | A / H5N8 | Europe / Poland / Greater Poland Voivodeship | Edyta ?wi?to? (National Veterinary Research Institut Poland, PIWet-PIB) |

| | | | | |
|---|---|---|---|---|
| EPI_ISL_525440 | A/chicken/Poland/003/2020 | A / H5N8 | Europe / Poland / Lublin Voivodeship | Edyta ?wi?to? (National Veterinary Research Institut Poland, PIWet-PIB) |
| EPI_ISL_525441 | A/chicken/Poland/004/2020 | A / H5N8 | Europe / Poland / Lublin Voivodeship | Edyta ?wi?to? (National Veterinary Research Institut Poland, PIWet-PIB) |
| EPI_ISL_525442 | A/turkey/Poland/027/2020 | A / H5N8 | Europe / Poland | Edyta ?wi?to? (National Veterinary Research Institut Poland, PIWet-PIB) |
| EPI_ISL_525443 | A/domestic_goose/Poland/028/2020 | A / H5N8 | Europe / Poland | Edyta ?wi?to? (National Veterinary Research Institut Poland, PIWet-PIB) |
| EPI_ISL_525444 | A/chicken/Poland/054/2020 | A / H5N8 | Europe / Poland | Edyta ?wi?to? (National Veterinary Research Institut Poland, PIWet-PIB) |
| EPI_ISL_525445 | A/turkey/Poland/079/2020 | A / H5N8 | Europe / Poland | Edyta ?wi?to? (National Veterinary Research Institut Poland, PIWet-PIB) |
| EPI_ISL_525446 | A/laying_hen/Poland/095/2020 | A / H5N8 | Europe / Poland | Edyta ?wi?to? (National Veterinary Research Institut Poland, PIWet-PIB) |
| EPI_ISL_525447 | A/turkey/Poland/096/2020 | A / H5N8 | Europe / Poland | Edyta ?wi?to? (National Veterinary Research Institut Poland, PIWet-PIB) |
| EPI_ISL_525448 | A/turkey/Poland/182/2020 | A / H5N8 | Europe / Poland | Edyta ?wi?to? (National Veterinary Research Institut Poland, PIWet-PIB) |
| EPI_ISL_525449 | A/domestic_duck/Poland/219/2020 | A / H5N8 | Europe / Poland | Edyta ?wi?to? (National Veterinary Research Institut Poland, PIWet-PIB) |
| EPI_ISL_525450 | A/domestic_duck/Poland/221/2020 | A / H5N8 | Europe / Poland | Edyta ?wi?to? (National Veterinary Research Institut Poland, PIWet-PIB) |
| EPI_ISL_525451 | A/domestic_duck/Poland/222/2020 | A / H5N8 | Europe / Poland | Edyta ?wi?to? (National Veterinary Research Institut Poland, PIWet-PIB) |
| EPI_ISL_525452 | A/domestic_duck/Poland/223/2020 | A / H5N8 | Europe / Poland | Edyta ?wi?to? (National Veterinary Research Institut Poland, PIWet-PIB) |
| EPI_ISL_525453 | A/domestic_duck/Poland/229/2020 | A / H5N8 | Europe / Poland | Edyta ?wi?to? (National Veterinary Research Institut Poland, PIWet-PIB) |
| EPI_ISL_525454 | A/domestic_duck/Poland/230/2020 | A / H5N8 | Europe / Poland | Edyta ?wi?to? (National Veterinary Research Institut Poland, PIWet-PIB) |
| EPI_ISL_525455 | A/domestic_duck/Poland/237/2020 | A / H5N8 | Europe / Poland | Edyta ?wi?to? (National Veterinary Research Institut Poland, PIWet-PIB) |
| EPI_ISL_525459 | A/laying_hen/Poland/312/2020 | A / H5N8 | Europe / Poland | Edyta ?wi?to? (National Veterinary Research Institut Poland, PIWet-PIB) |
| EPI_ISL_525461 | A/turkey/Poland/366/2020 | A / H5N8 | Europe / Poland | Edyta ?wi?to? (National Veterinary Research Institut Poland, PIWet-PIB) |
| EPI_ISL_525462 | A/domestic_goose/Poland/274/2020 | A / H5N8 | Europe / Poland | Edyta ?wi?to? (National Veterinary Research Institut Poland, PIWet-PIB) |
| EPI_ISL_525463 | A/domestic_duck/Poland/285/2020 | A / H5N8 | Europe / Poland | Edyta ?wi?to? (National Veterinary Research Institut Poland, PIWet-PIB) |
| EPI_ISL_525464 | A/domestic_duck/Poland/263/2020 | A / H5N8 | Europe / Poland | Edyta ?wi?to? (National Veterinary Research Institut Poland, PIWet-PIB) |
| EPI_ISL_525465 | A/domestic_duck/Poland/271/2020 | A / H5N8 | Europe / Poland | Edyta ?wi?to? (National Veterinary Research Institut Poland, PIWet-PIB) |
| EPI_ISL_5260416 | A/black-backed gull/Shandong/SC189/2021 | A / H5N8 | Asia / China / Shandong | Pengfei Cui (Harbin Veterinary Research Institute (CAAS) / Ministry of Agriculture) |
| EPI_ISL_5260417 | A/brown-headed gull/Tibet/1-1/2021 | A / H5N8 | Asia / China / Tibet | Pengfei Cui (Harbin Veterinary Research Institute (CAAS) / Ministry of Agriculture) |
| EPI_ISL_5260418 | A/black swan/Beijing/1/2021 | A / H5N8 | Asia / China / Beijing | Pengfei Cui (Harbin Veterinary Research Institute (CAAS) / Ministry of Agriculture) |
| EPI_ISL_5260419 | A/duck/Guangdong/S1269/2021 | A / H5N8 | Asia / China / Guangdong | Pengfei Cui (Harbin Veterinary Research Institute (CAAS) / Ministry of Agriculture) |
| EPI_ISL_5260420 | A/duck/Guangxi/S10099/2021 | A / H5N8 | Asia / China / Guangxi | Pengfei Cui (Harbin Veterinary Research Institute (CAAS) / Ministry of Agriculture) |
| EPI_ISL_5260421 | A/duck/Guangxi/S10263/2021 | A / H5N8 | Asia / China / Guangxi | Pengfei Cui (Harbin Veterinary Research Institute (CAAS) / Ministry of Agriculture) |
| EPI_ISL_5260422 | A/duck/Guangxi/S11043/2021 | A / H5N8 | Asia / China / Guangxi | Pengfei Cui (Harbin Veterinary Research Institute (CAAS) / Ministry of Agriculture) |
| EPI_ISL_5260426 | A/duck/Guangxi/S21194/2021 | A / H5N8 | Asia / China / Guangxi | Pengfei Cui (Harbin Veterinary Research Institute (CAAS) / Ministry of Agriculture) |
| EPI_ISL_5260435 | A/duck/Henan/S1381/2021 | A / H5N8 | Asia / China / Henan | Pengfei Cui (Harbin Veterinary Research Institute (CAAS) / Ministry of Agriculture) |
| EPI_ISL_5260444 | A/duck/Jiangxi/S10252/2021 | A / H5N8 | Asia / China / Jiangxi | Pengfei Cui (Harbin Veterinary Research Institute (CAAS) / Ministry of Agriculture) |
| EPI_ISL_5260445 | A/duck/Hebei/S1070/2021 | A / H5N8 | Asia / China / Hebei | Pengfei Cui (Harbin Veterinary Research Institute (CAAS) / Ministry of Agriculture) |
| EPI_ISL_5260446 | A/goose/Guangxi/S20601/2021 | A / H5N8 | Asia / China / Guangxi | Pengfei Cui (Harbin Veterinary Research Institute (CAAS) / Ministry of Agriculture) |
| EPI_ISL_5260447 | A/goose/Henan/S1315/2021 | A / H5N8 | Asia / China / Henan | Pengfei Cui (Harbin Veterinary Research Institute (CAAS) / Ministry of Agriculture) |
| EPI_ISL_5260448 | A/goose/Hunan/S11288/2021 | A / H5N8 | Asia / China / Hunan | Pengfei Cui (Harbin Veterinary Research Institute (CAAS) / Ministry of Agriculture) |
| EPI_ISL_5260449 | A/goose/Jiangsu/S1385/2021 | A / H5N8 | Asia / China / Jiangsu | Pengfei Cui (Harbin Veterinary Research Institute (CAAS) / Ministry of Agriculture) |
| EPI_ISL_5260450 | A/goose/Zhejiang/S1266/2021 | A / H5N8 | Asia / China / Zhejiang | Pengfei Cui (Harbin Veterinary Research Institute (CAAS) / Ministry of Agriculture) |
| EPI_ISL_5260451 | A/goose/Liaoning/S1266/2021 | A / H5N8 | Asia / China / Liaoning | Pengfei Cui (Harbin Veterinary Research Institute (CAAS) / Ministry of Agriculture) |
| EPI_ISL_5260452 | A/green-winged teal/Guangdong/SD004/2021 | A / H5N8 | Asia / China / Guangdong | Pengfei Cui (Harbin Veterinary Research Institute (CAAS) / Ministry of Agriculture) |
| EPI_ISL_5260453 | A/mute swan/Shandong/1/2021 | A / H5N8 | Asia / China / Shandong | Pengfei Cui (Harbin Veterinary Research Institute (CAAS) / Ministry of Agriculture) |
| EPI_ISL_5260454 | A/wild duck/Shandong/SC177/2021 | A / H5N8 | Asia / China / Shandong | Pengfei Cui (Harbin Veterinary Research Institute (CAAS) / Ministry of Agriculture) |
| EPI_ISL_5260455 | A/whooper swan/Shandong/SC176/2021 | A / H5N8 | Asia / China / Shandong | Pengfei Cui (Harbin Veterinary Research Institute (CAAS) / Ministry of Agriculture) |
| EPI_ISL_5260456 | A/whooper swan/Shandong/SC185/2021 | A / H5N8 | Asia / China / Shandong | Pengfei Cui (Harbin Veterinary Research Institute (CAAS) / Ministry of Agriculture) |
| EPI_ISL_5260457 | A/whooper swan/Shandong/SC195/2021 | A / H5N8 | Asia / China / Shandong | Pengfei Cui (Harbin Veterinary Research Institute (CAAS) / Ministry of Agriculture) |
| EPI_ISL_5260458 | A/whooper swan/Shandong/SC198/2021 | A / H5N8 | Asia / China / Shandong | Pengfei Cui (Harbin Veterinary Research Institute (CAAS) / Ministry of Agriculture) |
| EPI_ISL_5260459 | A/whooper swan/Shandong/SC199/2021 | A / H5N8 | Asia / China / Shandong | Pengfei Cui (Harbin Veterinary Research Institute (CAAS) / Ministry of Agriculture) |
| EPI_ISL_5260460 | A/whooper swan/Shandong/SC200/2021 | A / H5N8 | Asia / China / Shandong | Pengfei Cui (Harbin Veterinary Research Institute (CAAS) / Ministry of Agriculture) |
| EPI_ISL_5260461 | A/whooper swan/Shandong/SC188/2021 | A / H5N8 | Asia / China / Shandong | Pengfei Cui (Harbin Veterinary Research Institute (CAAS) / Ministry of Agriculture) |
| EPI_ISL_5260462 | A/whooper swan/Shanxi/4-1/2020 | A / H5N8 | Asia / China / Shanxi | Pengfei Cui (Harbin Veterinary Research Institute (CAAS) / Ministry of Agriculture) |
| EPI_ISL_5260463 | A/whooper swan/Shanxi/4-2/2020 | A / H5N8 | Asia / China / Shanxi | Pengfei Cui (Harbin Veterinary Research Institute (CAAS) / Ministry of Agriculture) |
| EPI_ISL_5260464 | A/common coot/Shandong/SC197/2021 | A / H5N8 | Asia / China / Shandong | Pengfei Cui (Harbin Veterinary Research Institute (CAAS) / Ministry of Agriculture) |
| EPI_ISL_5260465 | A/egret/Jiangsu/SD021/2021 | A / H5N8 | Asia / China / Jiangsu | Pengfei Cui (Harbin Veterinary Research Institute (CAAS) / Ministry of Agriculture) |
| EPI_ISL_5260466 | A/grebe/Shandong/SC184/2021 | A / H5N8 | Asia / China / Shandong | Pengfei Cui (Harbin Veterinary Research Institute (CAAS) / Ministry of Agriculture) |
| EPI_ISL_5260467 | A/grebe/Shaanxi/SD001/2021 | A / H5N8 | Asia / China / Shaanxi | Pengfei Cui (Harbin Veterinary Research Institute (CAAS) / Ministry of Agriculture) |
| EPI_ISL_5260468 | A/grebe/Ningxia/SD001/2021 | A / H5N8 | Asia / China / Ningxia | Pengfei Cui (Harbin Veterinary Research Institute (CAAS) / Ministry of Agriculture) |
| EPI_ISL_5260469 | A/wild duck/Jiangsu/SD019/2021 | A / H5N8 | Asia / China / Jiangsu | Pengfei Cui (Harbin Veterinary Research Institute (CAAS) / Ministry of Agriculture) |
| EPI_ISL_5260470 | A/wild goose/Shandong/SC196/2021 | A / H5N8 | Asia / China / Shandong | Pengfei Cui (Harbin Veterinary Research Institute (CAAS) / Ministry of Agriculture) |
| EPI_ISL_529179 | A/duck/Netherlands/16014829-001005/2016 | A / H5N8 | Europe / Netherlands / Provincie Flevoland / Gemeente Dronten / Biddinghuizen | Rene Heutink (Wageningen Bioveterinary Research) |
| EPI_ISL_5323346 | A/goose/Czech Republic/18520-1/2021 | A / H5N1 | Europe / Czech Republic / Stredocesky Kraj / Okres Pribram / Trhove Dusniky; Zip Code:26101 | Alexander Nagy (State Veterinary Institute Prague) |
| EPI_ISL_5323347 | A/duck/Czech Republic/18520-2/2021 | A / H5N1 | Europe / Czech Republic / Stredocesky Kraj / Okres Pribram / Trhove Dusniky; Zip Code:26101 | Alexander Nagy (State Veterinary Institute Prague) |
| EPI_ISL_5403566 | A/Eurasian wigeon/Germany-SH/AI05948/2021 | A / H5N1 | Europe / Germany / Schleswig-Holstein / Nordfriesland | Jacqueline King (Friedrich-Loeffler-Institut) |
| EPI_ISL_5449305 | A/barnacle_goose/Denmark/17572-1.01/2021-03-01 | A / H5N1 | Europe / Denmark / Region Sjaland / Slagelse Kommune | Charlotte Kristiane Hjulsager (Statens Serum Institute / Microbiological Diagnostic and Virology) |
| EPI_ISL_5463793 | A/chicken/Tyumen/27-31V/2021 | A / H5N1 | Europe / Russian Federation / Tyumen Oblast / Tyumen | Natalia Goncharova (State Research Center of Virology and Biotechnology (VECTOR) / Emerging Zoonotic Diseases and Influenza) |
| EPI_ISL_5463794 | A/chicken/Tyumen/27-39V/2021 | A / H5N1 | Europe / Russian Federation / Tyumen Oblast / Tyumen | Natalia Goncharova (State Research Center of Virology and Biotechnology (VECTOR) / Emerging Zoonotic Diseases and Influenza) |
| EPI_ISL_5463795 | A/chicken/Tyumen/27-40V/2021 | A / H5N1 | Europe / Russian Federation / Tyumen Oblast / Tyumen | Natalia Goncharova (State Research Center of Virology and Biotechnology (VECTOR) / Emerging Zoonotic Diseases and Influenza) |
| EPI_ISL_5463796 | A/chicken/Tyumen/27-42V/2021 | A / H5N1 | Europe / Russian Federation / Tyumen Oblast / Tyumen | Natalia Goncharova (State Research Center of Virology and Biotechnology (VECTOR) / Emerging Zoonotic Diseases and Influenza) |
| EPI_ISL_5463797 | A/duck/Saratov/29-02V/2021 | A / H5N1 | Europe / Russian Federation / Saratov Oblast / Saratov | Natalia Goncharova (State Research Center of Virology and Biotechnology (VECTOR) / Emerging Zoonotic Diseases and Influenza) |
| EPI_ISL_5463798 | A/duck/Saratov/29-03V/2021 | A / H5N1 | Europe / Russian Federation / Saratov Oblast / Saratov | Natalia Goncharova (State Research Center of Virology and Biotechnology (VECTOR) / Emerging Zoonotic Diseases and Influenza) |
| EPI_ISL_5463799 | A/duck/Saratov/29-04V/2021 | A / H5N1 | Europe / Russian Federation / Saratov Oblast / Saratov | Natalia Goncharova (State Research Center of Virology and Biotechnology (VECTOR) / Emerging Zoonotic Diseases and Influenza) |
| EPI_ISL_5463800 | A/chicken/Saratov/29-06V/2021 | A / H5N1 | Europe / Russian Federation / Saratov Oblast / Saratov | Natalia Goncharova (State Research Center of Virology and Biotechnology (VECTOR) / Emerging Zoonotic Diseases and Influenza) |
| EPI_ISL_5463801 | A/chicken/Saratov/29-07V/2021 | A / H5N1 | Europe / Russian Federation / Saratov Oblast / Saratov | Natalia Goncharova (State Research Center of Virology and Biotechnology (VECTOR) / Emerging Zoonotic Diseases and Influenza) |

| | | | | |
|---|---|---|---|---|
| EPI_ISL_5463802 | A/duck/Saratov/29-08V/2021 | A / H5N1 | Europe / Russian Federation / Saratov Oblast / Saratov | Natalia Goncharova (State Research Center of Virology and Biotechnology (VECTOR) / Emerging Zoonotic Diseases and Influenza) |
| EPI_ISL_5463803 | A/duck/Saratov/29-11V/2021 | A / H5N1 | Europe / Russian Federation / Saratov Oblast / Saratov | Natalia Goncharova (State Research Center of Virology and Biotechnology (VECTOR) / Emerging Zoonotic Diseases and Influenza) |
| EPI_ISL_5463804 | A/chicken/Tyumen/33-45V/2021 | A / H5N1 | Europe / Russian Federation / Tyumen Oblast / Tyumen | Natalia Goncharova (State Research Center of Virology and Biotechnology (VECTOR) / Emerging Zoonotic Diseases and Influenza) |
| EPI_ISL_5463805 | A/goose/Tyumen/33-52V/2021 | A / H5N1 | Europe / Russian Federation / Tyumen Oblast / Tyumen | Natalia Goncharova (State Research Center of Virology and Biotechnology (VECTOR) / Emerging Zoonotic Diseases and Influenza) |
| EPI_ISL_5463806 | A/goose/Tyumen/33-53V/2021 | A / H5N1 | Europe / Russian Federation / Tyumen Oblast / Tyumen | Natalia Goncharova (State Research Center of Virology and Biotechnology (VECTOR) / Emerging Zoonotic Diseases and Influenza) |
| EPI_ISL_5524384 | A/turkey/Italy/21VIR8585-1/2021 | A / H5N1 | Europe / Italy / Veneto / Province of Verona / Ronco All'Adige | Bianca Zecchin (Istituto Zooprofilattico Sperimentale Delle Venezie) |
| EPI_ISL_5530613 | A/great_skua/Scotland/041672/2021 | A / H5N1 | Europe / United Kingdom / Western Isles / Flannan Isles | Alex Byrne (Animal and Plant Health Agency (APHA) / Virology Department) |
| EPI_ISL_5588100 | A/chicken/Netherlands/21037287-006010/2021 | A / H5N1 | Europe / Netherlands / Provincie Flevoland / Gemeente Zeewolde | Rene Heutink (Wageningen Bioveterinary Research) |
| EPI_ISL_5588106 | A/chicken/Netherlands/21037233-001/2021 | A / H5N1 | Europe / Netherlands / Provincie Noord-Holland | Rene Heutink (Wageningen Bioveterinary Research) |
| EPI_ISL_5804698 | A/chicken/Netherlands/21037708-006010/2021 | A / H5N1 | Europe / Netherlands / Provincie Noord-Holland | Rene Heutink (Wageningen Bioveterinary Research) |
| EPI_ISL_5804708 | A/mute_swan/England/053054/2021 | A / H5N1 | Europe / United Kingdom / Worcestershire | Alex Byrne (Animal and Plant Health Agency (APHA) / Virology Department) |
| EPI_ISL_5804788 | A/Mute swan/Netherlands/21037283-002/2021 | A / H5N1 | Europe / Netherlands / Provincie Overijssel | Rene Heutink (Wageningen Bioveterinary Research) |
| EPI_ISL_5804789 | A/great_skua/Scotland/B07779/2021 | A / H5N1 | Europe / United Kingdom / Western Isles | Alex Byrne (Animal and Plant Health Agency (APHA) / Virology Department) |
| EPI_ISL_5862398 | A/green-winged-teal/Georgia/DT-22810/2020 | A / H5N8 | Asia / Georgia | Nicola Lewis (Royal Veterinary College) |
| EPI_ISL_5863089 | A/mallard/Georgia/DT-22620/2020 | A / H5N8 | Asia / Georgia | Nicola Lewis (Royal Veterinary College) |
| EPI_ISL_5863358 | A/mallard/Georgia/DT-22360/2020 | A / H5N8 | Asia / Georgia | Nicola Lewis (Royal Veterinary College) |
| EPI_ISL_5863359 | A/mallard/Georgia/DT-23072/2020 | A / H5N8 | Asia / Georgia | Nicola Lewis (Royal Veterinary College) |
| EPI_ISL_5863447 | A/green-winged-teal/Georgia/DT-22332/2020 | A / H5N8 | Asia / Georgia | Nicola Lewis (Royal Veterinary College) |
| EPI_ISL_5863490 | A/green-winged-teal/Georgia/DT-22792/2020 | A / H5N8 | Asia / Georgia | Nicola Lewis (Royal Veterinary College) |
| EPI_ISL_5863491 | A/mallard/Georgia/DT-22456/2020 | A / H5N8 | Asia / Georgia | Nicola Lewis (Royal Veterinary College) |
| EPI_ISL_5863497 | A/green-winged-teal/Georgia/DT-23070/2020 | A / H5N8 | Asia / Georgia | Nicola Lewis (Royal Veterinary College) |
| EPI_ISL_5863557 | A/green-winged-teal/Georgia/DT-22978/2020 | A / H5N8 | Asia / Georgia | Nicola Lewis (Royal Veterinary College) |
| EPI_ISL_5863567 | A/green-winged-teal/Georgia/DT-22392/2020 | A / H5N8 | Asia / Georgia | Nicola Lewis (Royal Veterinary College) |
| EPI_ISL_5863615 | A/mallard/Georgia/DT-22862/2020 | A / H5N8 | Asia / Georgia | Nicola Lewis (Royal Veterinary College) |
| EPI_ISL_5863641 | A/mallard/Georgia/DT-22495/2020 | A / H5N8 | Asia / Georgia | Nicola Lewis (Royal Veterinary College) |
| EPI_ISL_5863650 | A/mallard/Georgia/DT-20222/2020 | A / H5N8 | Asia / Georgia | Nicola Lewis (Royal Veterinary College) |
| EPI_ISL_5863749 | A/green-winged-teal/Georgia/DT-22894/2020 | A / H5N3 | Asia / Georgia | Nicola Lewis (Royal Veterinary College) |
| EPI_ISL_5863929 | A/garganey/Georgia/DT-22572/2020 | A / H5N8 | Asia / Georgia | Nicola Lewis (Royal Veterinary College) |
| EPI_ISL_5865417 | A/domestic duck/Georgia/DT-22368/2020 | A / H5N8 | Asia / Georgia | Nicola Lewis (Royal Veterinary College) |
| EPI_ISL_5865550 | A/mallard/Georgia/DT-22362/2020 | A / H5N8 | Asia / Georgia | Nicola Lewis (Royal Veterinary College) |
| EPI_ISL_5865629 | A/garganey/Georgia/DT-22770/2020 | A / H5N8 | Asia / Georgia | Nicola Lewis (Royal Veterinary College) |
| EPI_ISL_5865682 | A/mallard/Georgia/DT-22356/2020 | A / H5N8 | Asia / Georgia | Nicola Lewis (Royal Veterinary College) |
| EPI_ISL_5865695 | A/green-winged-teal/Georgia/DT-22246/2020 | A / H5N8 | Asia / Georgia | Nicola Lewis (Royal Veterinary College) |
| EPI_ISL_591075 | A/mute swan/Netherlands/20015931-001/2020 | A / H5N8 | Europe / Netherlands / Provincie Utrecht / Kockengen | Rene Heutink (Wageningen Bioveterinary Research) |
| EPI_ISL_5941469 | A/duck/Netherlands/21037958-001005/2021 | A / H5N1 | Europe / Netherlands / Provincie Flevoland | Rene Heutink (Wageningen Bioveterinary Research) |
| EPI_ISL_5942524 | A/chicken/Netherlands/21037750-001005/2021 | A / H5N1 | Europe / Netherlands / Provincie Noord-Holland | Rene Heutink (Wageningen Bioveterinary Research) |
| EPI_ISL_5945412 | A/European Herring Gull/Netherlands/21037412-002/2021 | A / H5N1 | Europe / Netherlands / Provincie Friesland | Rene Heutink (Wageningen Bioveterinary Research) |
| EPI_ISL_6025676 | A/Barnacle goose/Netherlands/21037293-001/2021 | A / H5N1 | Europe / Netherlands / Provincie Friesland | Rene Heutink (Wageningen Bioveterinary Research) |
| EPI_ISL_6029360 | A/great_skua/Scotland/042505/2021 | A / H5N1 | Europe / United Kingdom / Western Isles | Alex Byrne (Animal and Plant Health Agency (APHA) / Virology Department) |
| EPI_ISL_603132 | A/chicken/Netherlands/20016597-026030/2020 | A / H5N8 | Europe / Netherlands / Provincie Gelderland / West Maas en Waal / Altforst | Rene Heutink (Wageningen Bioveterinary Research) |
| EPI_ISL_603133 | A/Eurasian Wigeon/Netherlands/1/2020 | A / H5N1 | Europe / Netherlands | Theo Bestebroer (Erasmus Medical Center / Department of Virology) |
| EPI_ISL_603134 | A/Eurasian Wigeon/Netherlands/4/2020 | A / H5N1 | Europe / Netherlands | Theo Bestebroer (Erasmus Medical Center / Department of Virology) |
| EPI_ISL_603135 | A/Eurasian Wigeon/Netherlands/5/2020 | A / H5N1 | Europe / Netherlands | Theo Bestebroer (Erasmus Medical Center / Department of Virology) |
| EPI_ISL_603136 | A/Eurasian Wigeon/Netherlands/7/2020 | A / H5N8 | Europe / Netherlands | Theo Bestebroer (Erasmus Medical Center / Department of Virology) |
| EPI_ISL_603226 | A/chicken/Bulgaria/136_19VIR3315-1/2019 | A / H5N8 | Europe / Bulgaria / Oblast Plovdiv / Krumovo | Bianca Zecchin (Istituto Zooprofilattico Sperimentale Delle Venezie) |
| EPI_ISL_603227 | A/chicken/Bulgaria/143_19VIR3315-2/2019 | A / H5N8 | Europe / Bulgaria / Oblast Plovdiv / Asenograd | Bianca Zecchin (Istituto Zooprofilattico Sperimentale Delle Venezie) |
| EPI_ISL_603228 | A/chicken/Bulgaria/148-01_19VIR3315-3/2019 | A / H5N8 | Europe / Bulgaria / Oblast Plovdiv / Asenograd | Bianca Zecchin (Istituto Zooprofilattico Sperimentale Delle Venezie) |
| EPI_ISL_603229 | A/chicken/Bulgaria/150-01_19VIR3315-5/2019 | A / H5N8 | Europe / Bulgaria / Oblast Plovdiv / Asenograd | Bianca Zecchin (Istituto Zooprofilattico Sperimentale Delle Venezie) |
| EPI_ISL_603230 | A/duck/Bulgaria/78-4t_20VIR1416-3/2020 | A / H5N8 | Europe / Bulgaria / Oblast Plovdiv / Padarsko | Bianca Zecchin (Istituto Zooprofilattico Sperimentale Delle Venezie) |
| EPI_ISL_603231 | A/chicken/Bulgaria/201_20VIR1723-1/2020 | A / H5N2 | Europe / Bulgaria / Oblast Plovdiv / Trilistnik | Bianca Zecchin (Istituto Zooprofilattico Sperimentale Delle Venezie) |
| EPI_ISL_603232 | A/chicken/Bulgaria/221_20VIR1725-1/2020 | A / H5N2 | Europe / Bulgaria / Oblast Plovdiv / Manole | Bianca Zecchin (Istituto Zooprofilattico Sperimentale Delle Venezie) |
| EPI_ISL_603233 | A/chicken/Bulgaria/380-1_20VIR3542-1/2020 | A / H5N8 | Europe / Bulgaria / Oblast Plovdiv / Asenograd | Bianca Zecchin (Istituto Zooprofilattico Sperimentale Delle Venezie) |
| EPI_ISL_603234 | A/mule_duck/Bulgaria/50-506_20VIR1414-1/2020 | A / H5N2 | Europe / Bulgaria / Oblast Plovdiv / Obshtina Rakovski | Bianca Zecchin (Istituto Zooprofilattico Sperimentale Delle Venezie) |
| EPI_ISL_603235 | A/mule_duck/Bulgaria/147_20VIR1721-1/2020 | A / H5N8 | Europe / Bulgaria / Oblast Plovdiv / Bolyarino | Bianca Zecchin (Istituto Zooprofilattico Sperimentale Delle Venezie) |
| EPI_ISL_603237 | A/duck/Bulgaria/130-1_19VIR3314-1/2019 | A / H5N8 | Europe / Bulgaria / Oblast Lovech / Yoglav | Bianca Zecchin (Istituto Zooprofilattico Sperimentale Delle Venezie) |
| EPI_ISL_6101847 | A/duck/Netherlands/21038151-001005/2021 | A / H5N1 | Europe / Netherlands / Provincie Flevoland | Rene Heutink (Wageningen Bioveterinary Research) |
| EPI_ISL_6101848 | A/chicken/Netherlands/21038165-006010/2021 | A / H5N1 | Europe / Netherlands / Provincie Groningen | Rene Heutink (Wageningen Bioveterinary Research) |
| EPI_ISL_6101868 | A/black-headed gull/Netherlands/21037589-002/2021 | A / H5N1 | Europe / Netherlands / Provincie Friesland | Rene Heutink (Wageningen Bioveterinary Research) |
| EPI_ISL_6101869 | A/goose/Netherlands/21037720-001/2021 | A / H5N1 | Europe / Netherlands / Provincie Noord-Holland | Rene Heutink (Wageningen Bioveterinary Research) |
| EPI_ISL_614399 | A/buzzard/Germany-MV/AI02166/2020 | A / H5N5 | Europe / Germany / Mecklenburg-Vorpommern | Jacqueline King (Friedrich-Loeffler-Institut) |
| EPI_ISL_614400 | A/barnacle goose/Germany-SH/AI02167/2020 | A / H5N8 | Europe / Germany / Schleswig-Holstein | Jacqueline King (Friedrich-Loeffler-Institut) |
| EPI_ISL_614401 | A/mute_swan/Kazakhstan/1-267-20-B/2020 | A / H5N8 | Asia / Kazakhstan / Akmola | Alex Byrne (Animal and Plant Health Agency (APHA) / Virology Department) |
| EPI_ISL_615065 | A/domestic_goose/Kazakhstan/1-261_2-20-B/2020 | A / H5N8 | Asia / Kazakhstan / Kostanay | Alex Byrne (Animal and Plant Health Agency (APHA) / Virology Department) |
| EPI_ISL_615068 | A/domestic_goose/Kazakhstan/1-248_2-20-B/2020 | A / H5N8 | Asia / Kazakhstan / Akmola | Alex Byrne (Animal and Plant Health Agency (APHA) / Virology Department) |
| EPI_ISL_615072 | A/domestic_duck/Kazakhstan/1-274-20-B/2020 | A / H5N8 | Asia / Kazakhstan / Kostanay | Alex Byrne (Animal and Plant Health Agency (APHA) / Virology Department) |
| EPI_ISL_615073 | A/domestic_goose/Kazakhstan/1-242_2-20-B/2020 | A / H5N8 | Asia / Kazakhstan / North Kazakhstan | Alex Byrne (Animal and Plant Health Agency (APHA) / Virology Department) |
| EPI_ISL_623074 | A/chicken/Iraq/1/2020 | A / H5N8 | Asia / Iraq / Muhafazat Ninawa | Alex Byrne (Animal and Plant Health Agency (APHA) / Virology Department) |
| EPI_ISL_623075 | A/chicken/Netherlands/20017138-016020/2020 | A / H5N8 | Europe / Netherlands / Provincie Gelderland / Gemeente Druten | Rene Heutink (Wageningen Bioveterinary Research) |
| EPI_ISL_625671 | A/whooper swan/Inner Mongolia/w1-1/2020 | A / H5N8 | Asia / China / Inner Mongolia Autonomous Region | Hongliang Chai (Northeast Forestry University / College of Wildlife Resources) |
| EPI_ISL_625672 | A/mute swan/Inner Mongolia/w2-1/2020 | A / H5N8 | Asia / China / Inner Mongolia Autonomous Region | Hongliang Chai (Northeast Forestry University / College of Wildlife Resources) |
| EPI_ISL_626647 | A/chicken/Russian_Federation/Omsk/1680-10/2020 | A / H5N5 | Europe / Russian Federation / Omsk Oblast | Alex Byrne (Animal and Plant Health Agency (APHA) / Virology Department) |
| EPI_ISL_626648 | A/goose/Russian_Federation/Omsk/1680-6/2020 | A / H5N5 | Europe / Russian Federation / Omsk Oblast | Alex Byrne (Animal and Plant Health Agency (APHA) / Virology Department) |

| | | | | |
|---|---|---|---|---|
| EPI_ISL_626649 | A/duck/Russian_Federation/Saratov/1578-2/2020 | A / H5N8 | Europe / Russian Federation / Saratov Oblast | Alex Byrne (Animal and Plant Health Agency (APHA) / Virology Department) |
| EPI_ISL_626650 | A/duck/Russian_Federation/Omsk/1328-2/2020 | A / H5N8 | Europe / Russian Federation / Omsk Oblast | Alex Byrne (Animal and Plant Health Agency (APHA) / Virology Department) |
| EPI_ISL_626651 | A/goose/Russian_Federation/Kurgan/1345-25/2020 | A / H5N8 | Europe / Russian Federation / Kurgan Oblast | Alex Byrne (Animal and Plant Health Agency (APHA) / Virology Department) |
| EPI_ISL_626652 | A/chicken/England/030720/2020 | A / H5N8 | Europe / United Kingdom / Cheshire | Alex Byrne (Animal and Plant Health Agency (APHA) / Virology Department) |
| EPI_ISL_631824 | A/Mandarin duck/Korea/H242/2020 | A / H5N8 | Asia / Korea, Republic of / Chungcheongbuk-do | Yu-Na Lee (Animal and Plant Quarantine Agency (S-2158) / Avian Influenza Research & Diagnostic Division) |
| EPI_ISL_632314 | A/greylag goose/Netherlands/20016582-004/2020 | A / H5N1 | Europe / Netherlands / Provincie Noord-Holland / Gemeente Purmerend | Rene Heutink (Wageningen Bioveterinary Research) |
| EPI_ISL_632315 | A/eurasian teal/Netherlands/20016896-013/2020 | A / H5N1 | Europe / Netherlands / Provincie Friesland / Holwerd | Rene Heutink (Wageningen Bioveterinary Research) |
| EPI_ISL_632316 | A/eurasian curlew/Netherlands/20016890-001/2020 | A / H5N1 | Europe / Netherlands / Provincie Friesland / Hallum | Rene Heutink (Wageningen Bioveterinary Research) |
| EPI_ISL_632317 | A/barnacle goose/Netherlands/20016935-002/2020 | A / H5N8 | Europe / Netherlands / Provincie Friesland / Holwerd | Rene Heutink (Wageningen Bioveterinary Research) |
| EPI_ISL_632318 | A/greylag goose/Netherlands/20016879-001/2020 | A / H5N8 | Europe / Netherlands / Provincie Friesland / Earnewald | Rene Heutink (Wageningen Bioveterinary Research) |
| EPI_ISL_632319 | A/chicken/Netherlands/20017460-011015/2020 | A / H5N8 | Europe / Netherlands / Provincie Groningen / Lutjegast | Rene Heutink (Wageningen Bioveterinary Research) |
| EPI_ISL_6328001 | A/greylag goose/Netherlands/21037497-001/2021 | A / H5N1 | Europe / Netherlands / Provincie Noord-Holland | Rene Heutink (Wageningen Bioveterinary Research) |
| EPI_ISL_6328009 | A/swan/Netherlands/21037791-001/2021 | A / H5N1 | Europe / Netherlands / Provincie Friesland | Rene Heutink (Wageningen Bioveterinary Research) |
| EPI_ISL_6328022 | A/chicken/Netherlands/21037907-006010/2021 | A / H5N1 | Europe / Netherlands / Provincie Noord-Holland | Rene Heutink (Wageningen Bioveterinary Research) |
| EPI_ISL_6328036 | A/chicken/Netherlands/21038675-001005/2021 | A / H5N1 | Europe / Netherlands / Provincie Friesland | Rene Heutink (Wageningen Bioveterinary Research) |
| EPI_ISL_6328409 | A/mute swan/Czech Republic/21312/2021 | A / H5N1 | Europe / Czech Republic / Jihocesky Kraj / Okres Tabor / 49°14'23.870''N, 14°42''46.680E | Alexander Nagy (State Veterinary Institute Prague) |
| EPI_ISL_637098 | A/duck/Chelyabinsk/1207-1/2020 | A / H5N8 | Europe / Russian Federation / Chelyabinsk Oblast | Nikolay Zinyakov (Federal Centre for Animal Health (ARRIAH) / OIE Regional Reference Laboratory) |
| EPI_ISL_6375134 | A/waterfowl/Netherlands/21037914-006/2021 | A / H5N1 | Europe / Netherlands / Provincie Friesland | Rene Heutink (Wageningen Bioveterinary Research) |
| EPI_ISL_641377 | A/chicken/Netherlands/20016978-001/2020 | A / H5N8 | Europe / Netherlands / Provincie Noord-Holland / Edam-Volendam | Rene Heutink (Wageningen Bioveterinary Research) |
| EPI_ISL_641378 | A/greater white-fronted goose/Netherlands/20017403-004/20 | A / H5N5 | Europe / Netherlands / South Holland / Gemeente Rijswijk | Rene Heutink (Wageningen Bioveterinary Research) |
| EPI_ISL_641394 | A/chicken/Netherlands/20017639-001/2020 | A / H5N8 | Europe / Netherlands / Provincie Friesland / Wons | Rene Heutink (Wageningen Bioveterinary Research) |
| EPI_ISL_641395 | A/chicken/Netherlands/20017694-004/2020 | A / H5N8 | Europe / Netherlands / Provincie Noord-Holland / Gemeente Purmerend | Rene Heutink (Wageningen Bioveterinary Research) |
| EPI_ISL_641518 | A/duck/Netherlands/20017868-016020/2020 | A / H5N8 | Europe / Netherlands / Provincie Gelderland / Gemeente Voorst | Rene Heutink (Wageningen Bioveterinary Research) |
| EPI_ISL_644121 | A/turkey/Omsk/0001/2020 | A / H5N8 | Europe / Russian Federation / Omsk Oblast / Omsk | Natalia Goncharova (State Research Center of Virology and Biotechnology (VECTOR) / Emerging Zoonotic Diseases and Influenza) |
| EPI_ISL_644122 | A/goose/Omsk/0002/2020 | A / H5N8 | Europe / Russian Federation / Omsk Oblast / Omsk region | Natalia Goncharova (State Research Center of Virology and Biotechnology (VECTOR) / Emerging Zoonotic Diseases and Influenza) |
| EPI_ISL_644123 | A/turkey/Omsk/0003/2020 | A / H5N8 | Europe / Russian Federation / Omsk Oblast / Omsk | Natalia Goncharova (State Research Center of Virology and Biotechnology (VECTOR) / Emerging Zoonotic Diseases and Influenza) |
| EPI_ISL_644124 | A/duck/Omsk/0004/2020 | A / H5N8 | Europe / Russian Federation / Omsk Oblast / Omsk | Natalia Goncharova (State Research Center of Virology and Biotechnology (VECTOR) / Emerging Zoonotic Diseases and Influenza) |
| EPI_ISL_644125 | A/goose/Omsk/0111/2020 | A / H5N8 | Europe / Russian Federation / Omsk Oblast / Omsk | Natalia Goncharova (State Research Center of Virology and Biotechnology (VECTOR) / Emerging Zoonotic Diseases and Influenza) |
| EPI_ISL_644126 | A/chicken/Omsk/0112/2020 | A / H5N8 | Europe / Russian Federation / Omsk Oblast / Omsk | Natalia Goncharova (State Research Center of Virology and Biotechnology (VECTOR) / Emerging Zoonotic Diseases and Influenza) |
| EPI_ISL_644127 | A/goose/Omsk/0113/2020 | A / H5N8 | Europe / Russian Federation / Omsk Oblast / Omsk | Natalia Goncharova (State Research Center of Virology and Biotechnology (VECTOR) / Emerging Zoonotic Diseases and Influenza) |
| EPI_ISL_644128 | A/goose/Omsk/0114/2020 | A / H5N8 | Europe / Russian Federation / Omsk Oblast / Omsk | Natalia Goncharova (State Research Center of Virology and Biotechnology (VECTOR) / Emerging Zoonotic Diseases and Influenza) |
| EPI_ISL_644129 | A/goose/Omsk/0115/2020 | A / H5N8 | Europe / Russian Federation / Omsk Oblast / Omsk | Natalia Goncharova (State Research Center of Virology and Biotechnology (VECTOR) / Emerging Zoonotic Diseases and Influenza) |
| EPI_ISL_644130 | A/goose/Omsk/01161/2020 | A / H5N8 | Europe / Russian Federation / Omsk Oblast / Omsk | Natalia Goncharova (State Research Center of Virology and Biotechnology (VECTOR) / Emerging Zoonotic Diseases and Influenza) |
| EPI_ISL_644132 | A/goose/Omsk/01171/2020 | A / H5N8 | Europe / Russian Federation / Omsk Oblast / Omsk | Natalia Goncharova (State Research Center of Virology and Biotechnology (VECTOR) / Emerging Zoonotic Diseases and Influenza) |
| EPI_ISL_644134 | A/chicken/Omsk/0118/2020 | A / H5N8 | Europe / Russian Federation / Omsk Oblast / Omsk | Natalia Goncharova (State Research Center of Virology and Biotechnology (VECTOR) / Emerging Zoonotic Diseases and Influenza) |
| EPI_ISL_644135 | A/chicken/Omsk/0119/2020 | A / H5N8 | Europe / Russian Federation / Omsk Oblast / Omsk | Natalia Goncharova (State Research Center of Virology and Biotechnology (VECTOR) / Emerging Zoonotic Diseases and Influenza) |
| EPI_ISL_644136 | A/goose/Omsk/011101/2020 | A / H5N8 | Europe / Russian Federation / Omsk Oblast / Omsk | Natalia Goncharova (State Research Center of Virology and Biotechnology (VECTOR) / Emerging Zoonotic Diseases and Influenza) |
| EPI_ISL_644138 | A/wild duck/Omsk/01111/2020 | A / H5N8 | Europe / Russian Federation / Omsk Oblast / Omsk | Natalia Goncharova (State Research Center of Virology and Biotechnology (VECTOR) / Emerging Zoonotic Diseases and Influenza) |
| EPI_ISL_644139 | A/goose/Omsk/0071/2020 | A / H5N8 | Europe / Russian Federation / Omsk Oblast / Omsk | Natalia Goncharova (State Research Center of Virology and Biotechnology (VECTOR) / Emerging Zoonotic Diseases and Influenza) |
| EPI_ISL_644140 | A/chicken/Omsk/0073/2020 | A / H5N8 | Europe / Russian Federation / Omsk Oblast / Omsk | Natalia Goncharova (State Research Center of Virology and Biotechnology (VECTOR) / Emerging Zoonotic Diseases and Influenza) |
| EPI_ISL_644141 | A/goose/Omsk/0074/2020 | A / H5N8 | Europe / Russian Federation / Omsk Oblast / Omsk | Natalia Goncharova (State Research Center of Virology and Biotechnology (VECTOR) / Emerging Zoonotic Diseases and Influenza) |
| EPI_ISL_644142 | A/duck/Omsk/0075/2020 | A / H5N8 | Europe / Russian Federation / Omsk Oblast / Omsk | Natalia Goncharova (State Research Center of Virology and Biotechnology (VECTOR) / Emerging Zoonotic Diseases and Influenza) |
| EPI_ISL_644143 | A/duck/Omsk/0076/2020 | A / H5N8 | Europe / Russian Federation / Omsk Oblast / Omsk | Natalia Goncharova (State Research Center of Virology and Biotechnology (VECTOR) / Emerging Zoonotic Diseases and Influenza) |
| EPI_ISL_644144 | A/duck/Omsk/0077/2020 | A / H5N8 | Europe / Russian Federation / Omsk Oblast / Omsk | Natalia Goncharova (State Research Center of Virology and Biotechnology (VECTOR) / Emerging Zoonotic Diseases and Influenza) |
| EPI_ISL_644145 | A/chicken/Chelyabinsk/401/2020 | A / H5N8 | Europe / Russian Federation / Chelyabinsk Oblast / Chelyabinsk | Natalia Goncharova (State Research Center of Virology and Biotechnology (VECTOR) / Emerging Zoonotic Diseases and Influenza) |
| EPI_ISL_644146 | A/chicken/Chelyabinsk/402/2020 | A / H5N8 | Europe / Russian Federation / Chelyabinsk Oblast / Chelyabinsk | Natalia Goncharova (State Research Center of Virology and Biotechnology (VECTOR) / Emerging Zoonotic Diseases and Influenza) |
| EPI_ISL_644147 | A/chicken/Chelyabinsk/403/2020 | A / H5N8 | Europe / Russian Federation / Chelyabinsk Oblast / Chelyabinsk | Natalia Goncharova (State Research Center of Virology and Biotechnology (VECTOR) / Emerging Zoonotic Diseases and Influenza) |
| EPI_ISL_644148 | A/chicken/Chelyabinsk/404/2020 | A / H5N8 | Europe / Russian Federation / Chelyabinsk Oblast / Chelyabinsk | Natalia Goncharova (State Research Center of Virology and Biotechnology (VECTOR) / Emerging Zoonotic Diseases and Influenza) |
| EPI_ISL_644735 | A/Cygnus_olor/Belgium/11956_001/2020 (H5N8) | A / H5N8 | Europe / Belgium / Provincie West-Vlaanderen / Oudenburg | Steven Van Borm (Sciensano, Department of Animal Infectious Diseases / Animal Infectious Diseases) |
| EPI_ISL_644737 | A/peregrine falcon/Denmark/13776-1/2020-10-30(H5N5) | A / H5N5 | Europe / Denmark / Region Sjaland / Lolland | Charlotte Kristiane Hjulsager (Statens Serum Institute / Microbiological Diagnostic and Virology) |
| EPI_ISL_644824 | A/barnacle goose/Denmark/14138-1/2020-11-04(H5N8) | A / H5N8 | Europe / Denmark / Region Syddanmark / Tonder Kommune | Charlotte Kristiane Hjulsager (Statens Serum Institute / Microbiological Diagnostic and Virology) |
| EPI_ISL_647969 | A/Turkey/Sweden/SVA201114SZ0001/20KN303106/2020 | A / H5N8 | Europe / Sweden / Skane Lan / Tomelilla Kommun | Siamak Zohari (National Veterinary Institute) |
| EPI_ISL_6507374 | A/mute swan/Croatia/100/2021 | A / H5N1 | Europe / Croatia / Sisacko-Moslavacka Zupanija / Muzilovcica (45° 23' 25.9'' N; 16° 40' 49.3'' E) | Vladimir Savi? (Croatian Veterinary Institute / Poultry Centre) |
| EPI_ISL_653918 | A/greylag goose/Netherlands/20016414-001/2020 | A / H5N8 | Europe / Netherlands / Provincie Utrecht / Gemeente Woerden | Rene Heutink (Wageningen Bioveterinary Research) |
| EPI_ISL_653919 | A/greylag goose/Netherlands/20016494-001/2020 | A / H5N8 | Europe / Netherlands / Provincie Noord-Holland / Gemeente Naarden | Rene Heutink (Wageningen Bioveterinary Research) |
| EPI_ISL_653920 | A/chicken/Netherlands/20018496-006010/2020 | A / H5N8 | Europe / Netherlands / Provincie Friesland / Witmarsum | Rene Heutink (Wageningen Bioveterinary Research) |
| EPI_ISL_653921 | A/chicken/Netherlands/20018523-001005/2020 | A / H5N8 | Europe / Netherlands / Provincie Utrecht / Gemeente Oudewater | Rene Heutink (Wageningen Bioveterinary Research) |
| EPI_ISL_654821 | A/chicken/Saratov/29801/2020 | A / H5N8 | Europe / Russian Federation / Saratov Oblast / Saratov | Natalia Goncharova (State Research Center of Virology and Biotechnology (VECTOR) / Emerging Zoonotic Diseases and Influenza) |
| EPI_ISL_654824 | A/duck/Saratov/29804/2020 | A / H5N8 | Europe / Russian Federation / Saratov Oblast / Saratov | Natalia Goncharova (State Research Center of Virology and Biotechnology (VECTOR) / Emerging Zoonotic Diseases and Influenza) |
| EPI_ISL_654826 | A/goose/Omsk/30001/2020 | A / H5N8 | Europe / Russian Federation / Omsk Oblast / Omsk | Natalia Goncharova (State Research Center of Virology and Biotechnology (VECTOR) / Emerging Zoonotic Diseases and Influenza) |
| EPI_ISL_654827 | A/goose/Omsk/30003/2020 | A / H5N8 | Europe / Russian Federation / Omsk Oblast / Omsk | Natalia Goncharova (State Research Center of Virology and Biotechnology (VECTOR) / Emerging Zoonotic Diseases and Influenza) |
| EPI_ISL_654828 | A/goose/Omsk/30004/2020 | A / H5N8 | Europe / Russian Federation / Omsk Oblast / Omsk | Natalia Goncharova (State Research Center of Virology and Biotechnology (VECTOR) / Emerging Zoonotic Diseases and Influenza) |
| EPI_ISL_654829 | A/goose/Omsk/30006/2020 | A / H5N8 | Europe / Russian Federation / Omsk Oblast / Omsk | Natalia Goncharova (State Research Center of Virology and Biotechnology (VECTOR) / Emerging Zoonotic Diseases and Influenza) |
| EPI_ISL_654830 | A/chicken/Omsk/30007/2020 | A / H5N8 | Europe / Russian Federation / Omsk Oblast / Omsk | Natalia Goncharova (State Research Center of Virology and Biotechnology (VECTOR) / Emerging Zoonotic Diseases and Influenza) |
| EPI_ISL_654831 | A/goose/Omsk/30009/2020 | A / H5N8 | Europe / Russian Federation / Omsk Oblast / Omsk | Natalia Goncharova (State Research Center of Virology and Biotechnology (VECTOR) / Emerging Zoonotic Diseases and Influenza) |
| EPI_ISL_654832 | A/goose/Omsk/30010/2020 | A / H5N8 | Europe / Russian Federation / Omsk Oblast / Omsk | Natalia Goncharova (State Research Center of Virology and Biotechnology (VECTOR) / Emerging Zoonotic Diseases and Influenza) |
| EPI_ISL_654833 | A/chicken/Chelyabinsk/201/2020 | A / H5N8 | Europe / Russian Federation / Chelyabinsk region | Natalia Goncharova (State Research Center of Virology and Biotechnology (VECTOR) / Emerging Zoonotic Diseases and Influenza) |
| EPI_ISL_654834 | A/chicken/Kurgan/1001/2020 | A / H5N8 | Europe / Russian Federation / Kurgan region | Natalia Goncharova (State Research Center of Virology and Biotechnology (VECTOR) / Emerging Zoonotic Diseases and Influenza) |
| EPI_ISL_654835 | A/chicken/Kurgan/1003/2020 | A / H5N8 | Europe / Russian Federation / Kurgan region | Natalia Goncharova (State Research Center of Virology and Biotechnology (VECTOR) / Emerging Zoonotic Diseases and Influenza) |
| EPI_ISL_654836 | A/chicken/Kurgan/1004/2020 | A / H5N8 | Europe / Russian Federation / Kurgan Oblast | Natalia Goncharova (State Research Center of Virology and Biotechnology (VECTOR) / Emerging Zoonotic Diseases and Influenza) |
| EPI_ISL_654837 | A/chicken/Kurgan/1005/2020 | A / H5N8 | Europe / Russian Federation / Kurgan region | Natalia Goncharova (State Research Center of Virology and Biotechnology (VECTOR) / Emerging Zoonotic Diseases and Influenza) |
| EPI_ISL_654838 | A/chicken/Kurgan/1010/2020 | A / H5N8 | Europe / Russian Federation / Kurgan region | Natalia Goncharova (State Research Center of Virology and Biotechnology (VECTOR) / Emerging Zoonotic Diseases and Influenza) |

| | | | | |
|---|---|---|---|---|
| EPI_ISL_654839 | A/goose/Kurgan/01/2020 | A / H5N8 | Europe / Russian Federation / Kurgan region | Natalia Goncharova (State Research Center of Virology and Biotechnology (VECTOR) / Emerging Zoonotic Diseases and Influenza) |
| EPI_ISL_654840 | A/duck/Thanh Hoa/676VTC/2020 | A / H5N6 | Asia / Vietnam / Tinh Thanh Hoa / Tan Tho | Natalia Goncharova (State Research Center of Virology and Biotechnology (VECTOR) / Emerging Zoonotic Diseases and Influenza) |
| EPI_ISL_654841 | A/duck/Nghe An/694VTC/2020 | A / H5N6 | Asia / Vietnam / Tinh Nghe An / Quynh Hung | Natalia Goncharova (State Research Center of Virology and Biotechnology (VECTOR) / Emerging Zoonotic Diseases and Influenza) |
| EPI_ISL_654842 | A/duck/Nghe An/695VTC/2020 | A / H5N6 | Asia / Vietnam / Tinh Nghe An / Quynh Hau | Natalia Goncharova (State Research Center of Virology and Biotechnology (VECTOR) / Emerging Zoonotic Diseases and Influenza) |
| EPI_ISL_654843 | A/duck/Thanh Hoa/722VTC/2020 | A / H5N6 | Asia / Vietnam / Tinh Thanh Hoa / Quang Hoa | Natalia Goncharova (State Research Center of Virology and Biotechnology (VECTOR) / Emerging Zoonotic Diseases and Influenza) |
| EPI_ISL_654844 | A/duck/Thanh Hoa/740VTC/2020 | A / H5N6 | Asia / Vietnam / Tinh Thanh Hoa / Vinh | Natalia Goncharova (State Research Center of Virology and Biotechnology (VECTOR) / Emerging Zoonotic Diseases and Influenza) |
| EPI_ISL_654845 | A/chicken/Thanh Hoa/VTC741/2020 | A / H5N6 | Asia / Vietnam / Tinh Thanh Hoa / Tan Binh | Natalia Goncharova (State Research Center of Virology and Biotechnology (VECTOR) / Emerging Zoonotic Diseases and Influenza) |
| EPI_ISL_654846 | A/duck/Thanh Hoa/752VTC/2020 | A / H5N6 | Asia / Vietnam / Tinh Thanh Hoa / Thanh Hoa | Natalia Goncharova (State Research Center of Virology and Biotechnology (VECTOR) / Emerging Zoonotic Diseases and Influenza) |
| EPI_ISL_654847 | A/duck/Thanh Hoa/808VTC/2020 | A / H5N6 | Asia / Vietnam / Tinh Thanh Hoa / Dong Cao | Natalia Goncharova (State Research Center of Virology and Biotechnology (VECTOR) / Emerging Zoonotic Diseases and Influenza) |
| EPI_ISL_654848 | A/chicken/Nghe An/842VTC/2020 | A / H5N6 | Asia / Vietnam / Tinh Nghe An / Xom Thuan | Natalia Goncharova (State Research Center of Virology and Biotechnology (VECTOR) / Emerging Zoonotic Diseases and Influenza) |
| EPI_ISL_654849 | A/chicken/Thanh Hoa/844VTC/2020 | A / H5N6 | Asia / Vietnam / Tinh Thanh Hoa / Van Son | Natalia Goncharova (State Research Center of Virology and Biotechnology (VECTOR) / Emerging Zoonotic Diseases and Influenza) |
| EPI_ISL_654850 | A/muscovy duck/Thanh Hoa/879VTC/2020 | A / H5N6 | Asia / Vietnam / Tinh Thanh Hoa / Quang Hop | Natalia Goncharova (State Research Center of Virology and Biotechnology (VECTOR) / Emerging Zoonotic Diseases and Influenza) |
| EPI_ISL_654851 | A/muscovy duck/Thanh Hoa/892VTC/2020 | A / H5N6 | Asia / Vietnam / Tinh Thanh Hoa / Hung Loc | Natalia Goncharova (State Research Center of Virology and Biotechnology (VECTOR) / Emerging Zoonotic Diseases and Influenza) |
| EPI_ISL_654852 | A/duck/Thanh Hoa/893VTC/2020 | A / H5N6 | Asia / Vietnam / Tinh Thanh Hoa / Thieu Hoa | Natalia Goncharova (State Research Center of Virology and Biotechnology (VECTOR) / Emerging Zoonotic Diseases and Influenza) |
| EPI_ISL_654853 | A/duck/Thanh Hoa/923VTC/2020 | A / H5N6 | Asia / Vietnam / Tinh Thanh Hoa / Dong Son | Natalia Goncharova (State Research Center of Virology and Biotechnology (VECTOR) / Emerging Zoonotic Diseases and Influenza) |
| EPI_ISL_654854 | A/duck/Thanh Hoa/945VTC/2020 | A / H5N6 | Asia / Vietnam / Tinh Thanh Hoa / Quang Trung | Natalia Goncharova (State Research Center of Virology and Biotechnology (VECTOR) / Emerging Zoonotic Diseases and Influenza) |
| EPI_ISL_654855 | A/chicken/Thanh Hoa/968VTC/2020 | A / H5N6 | Asia / Vietnam / Tinh Thanh Hoa / Phu Son | Natalia Goncharova (State Research Center of Virology and Biotechnology (VECTOR) / Emerging Zoonotic Diseases and Influenza) |
| EPI_ISL_654856 | A/chicken/Thanh Hoa/980VTC/2020 | A / H5N6 | Asia / Vietnam / Tinh Thanh Hoa / Mai Lam | Natalia Goncharova (State Research Center of Virology and Biotechnology (VECTOR) / Emerging Zoonotic Diseases and Influenza) |
| EPI_ISL_654857 | A/duck/Thanh Hoa/1107VTC/2020 | A / H5N6 | Asia / Vietnam / Tinh Thanh Hoa / Phu Son | Natalia Goncharova (State Research Center of Virology and Biotechnology (VECTOR) / Emerging Zoonotic Diseases and Influenza) |
| EPI_ISL_654858 | A/duck/Thanh Hoa/1151VTC/2020 | A / H5N6 | Asia / Vietnam / Tinh Thanh Hoa / Quang Trung | Natalia Goncharova (State Research Center of Virology and Biotechnology (VECTOR) / Emerging Zoonotic Diseases and Influenza) |
| EPI_ISL_654859 | A/chicken/Thanh Hoa/1152VTC/2020 | A / H5N6 | Asia / Vietnam / Tinh Thanh Hoa / Phu Son | Natalia Goncharova (State Research Center of Virology and Biotechnology (VECTOR) / Emerging Zoonotic Diseases and Influenza) |
| EPI_ISL_654860 | A/duck/Thanh Hoa/1180VTC/2020 | A / H5N6 | Asia / Vietnam / Tinh Thanh Hoa / Phu Son | Natalia Goncharova (State Research Center of Virology and Biotechnology (VECTOR) / Emerging Zoonotic Diseases and Influenza) |
| EPI_ISL_654861 | A/duck/Thanh Hoa/1182VTC/2020 | A / H5N6 | Asia / Vietnam / Tinh Thanh Hoa / Quang Trung | Natalia Goncharova (State Research Center of Virology and Biotechnology (VECTOR) / Emerging Zoonotic Diseases and Influenza) |
| EPI_ISL_654862 | A/duck/Nghe An/1693VTC/2020 | A / H5N6 | Asia / Vietnam / Tinh Nghe An / Quynh Vinh | Natalia Goncharova (State Research Center of Virology and Biotechnology (VECTOR) / Emerging Zoonotic Diseases and Influenza) |
| EPI_ISL_654863 | A/duck/Thanh Hoa/2202VTC/2020 | A / H5N6 | Asia / Vietnam / Tinh Thanh Hoa / Truong Lam | Natalia Goncharova (State Research Center of Virology and Biotechnology (VECTOR) / Emerging Zoonotic Diseases and Influenza) |
| EPI_ISL_654864 | A/chicken/Nghe An/14475VTC/2020 | A / H5N6 | Asia / Vietnam / Tinh Nghe An / Quynh Chau | Natalia Goncharova (State Research Center of Virology and Biotechnology (VECTOR) / Emerging Zoonotic Diseases and Influenza) |
| EPI_ISL_654865 | A/duck/Nghe An/14528VTC/2020 | A / H5N6 | Asia / Vietnam / Tinh Nghe An / Ngoc Son | Natalia Goncharova (State Research Center of Virology and Biotechnology (VECTOR) / Emerging Zoonotic Diseases and Influenza) |
| EPI_ISL_654958 | A/mallard/Italy/20VIR7139-73/2020 | A / H5N8 | Europe / Italy / Veneto / Province of Rovigo / Porto Tolle, Ca' Zuliani | Bianca Zecchin (Istituto Zooprofilattico Sperimentale Delle Venezie) |
| EPI_ISL_6590766 | A/swan/France/21P012384/2021 | A / H5N1 | Europe / France / Region Lorraine / Departement de la Meuse | Francois-Xavier Briand (ANSES Agence Nationale De Securite Sanitaire De L'alimentation / Laboratoire de Ploufragan-Plouzané) |
| EPI_ISL_6596216 | A/barnacle goose/Sweden/SVA211102SZ0402/FB004395/M-2( | A / H5N1 | Europe / Sweden / Skane Lan / Malmo Kommun | Siamak Zohari (National Veterinary Institute) |
| EPI_ISL_6599075 | A/greylag goose /Sweden/SVA211103SZ0398/FB004410/M-20 | A / H5N1 | Europe / Sweden / Skane Lan / Svedala Kommun | Siamak Zohari (National Veterinary Institute) |
| EPI_ISL_6599895 | A/common pheasant /Sweden/SVA211104SZ0320/FB004417/I | A / H5N1 | Europe / Sweden / Skane Lan / Svedala Kommun | Siamak Zohari (National Veterinary Institute) |
| EPI_ISL_6600745 | A/common pheasant /Sweden/SVA211104SZ0320/FB004418/I | A / H5N1 | Europe / Sweden / Skane Lan / Svedala Kommun | Siamak Zohari (National Veterinary Institute) |
| EPI_ISL_6600769 | A/common buzzard /Sweden/SVA211104SZ0320/FB004419/M | A / H5N1 | Europe / Sweden / Skane Lan / Svedala Kommun | Siamak Zohari (National Veterinary Institute) |
| EPI_ISL_6600943 | A/great black-backed gull/Sweden/SVA211109SZ0434/FB0044 | A / H5N1 | Europe / Sweden / Skane Lan / Landskrona Kommun | Siamak Zohari (National Veterinary Institute) |
| EPI_ISL_660264 | A/Gallus_gallus/Belgium/12168_002/2020 | A / H5N5 | Europe / Belgium / Provincie West-Vlaanderen / Menen | Steven Van Borm (Sciensano, Department of Animal Infectious Diseases / Animal Infectious Diseases) |
| EPI_ISL_661177 | A/chicken/Poland/448/2020 | A / H5N8 | Europe / Poland / Greater Poland Voivodeship | Edyta ?wi?to? (National Veterinary Research Institut Poland, PIWet-PIB) |
| EPI_ISL_661178 | A/swan/Tumen/1479-2/2020 | A / H5N8 | Europe / Russian Federation / Tyumen Oblast | Nikolay Zinyakov (Federal Centre for Animal Health (ARRIAH) / OIE Regional Reference Laboratory) |
| EPI_ISL_661313 | A/Anser_albifrons/Belgium/11956_005/2020 | A / H5N8 | Europe / Belgium / Provincie West-Vlaanderen / Bredene | Steven Van Borm (Sciensano, Department of Animal Infectious Diseases / Animal Infectious Diseases) |
| EPI_ISL_664102 | A/Numenius_arquata/Belgium/11956_003/2020 | A / H5N8 | Europe / Belgium / Provincie West-Vlaanderen / Wilskerke | Steven Van Borm (Sciensano, Department of Animal Infectious Diseases / Animal Infectious Diseases) |
| EPI_ISL_666687 | A/Madarin duck/Korea/K20-551-4/2020 | A / H5N8 | Asia / Korea, Republic of / Gyeonggi-do | Sol Jeong (Konkuk University / 302, School of Veterinary Medicine) |
| EPI_ISL_666890 | A/turkey/Croatia/104/2020 | A / H5N8 | Europe / Croatia / Koprivnicko-Krizevacka Zupanija / Delovi | Vladimir Savi? (Croatian Veterinary Institute / Poultry Centre) |
| EPI_ISL_6675733 | A/chicken/Netherlands/21039174-001005/2021 | A / H5N1 | Europe / Netherlands / Provincie Utrecht | Rene Heutink (Wageningen Bioveterinary Research) |
| EPI_ISL_6675739 | A/chicken/Netherlands/21039108-001005/2021 | A / H5N1 | Europe / Netherlands / Provincie Utrecht | Rene Heutink (Wageningen Bioveterinary Research) |
| EPI_ISL_6675744 | A/mute swan/Netherlands/21039008-003/2021 | A / H5N1 | Europe / Netherlands / Provincie Noord-Holland | Rene Heutink (Wageningen Bioveterinary Research) |
| EPI_ISL_6675750 | A/mute swan/Netherlands/21038412-002/2021 | A / H5N1 | Europe / Netherlands / South Holland | Rene Heutink (Wageningen Bioveterinary Research) |
| EPI_ISL_6675755 | A/goose/Netherlands/21038413-002/2021 | A / H5N1 | Europe / Netherlands / Provincie Friesland | Rene Heutink (Wageningen Bioveterinary Research) |
| EPI_ISL_667810 | A/chicken/France/20P016448/2020 | A / H5N8 | Europe / France / Corsica / Departement de la Haute-Corse | Francois-Xavier Briand (ANSES Agence Nationale De Securite Sanitaire De L'alimentation / Laboratoire de Ploufragan-Plouzané) |
| EPI_ISL_668456 | A/Peregrine falcon/Sweden/SVA201117SZ0467/20KN003345/2 | A / H5N8 | Europe / Sweden / Skane Lan / Lunds Kommun | Siamak Zohari (National Veterinary Institute) |
| EPI_ISL_668457 | A/barnacle goose/Sweden/SVA201117SZ0468/KN003355/202( | A / H5N8 | Europe / Sweden / Skane Lan / Lunds Kommun | Siamak Zohari (National Veterinary Institute) |
| EPI_ISL_6757629 | A/Duck/Guangdong/21314/2021(H5N6) | A / H5N6 | Asia / China / Guangdong Province | Jiahao Zhang (South China Agricultural University / College of Veterinary Medicine) |
| EPI_ISL_6757630 | A/Goose/Sichuan/21406-5/2021(H5N6) | A / H5N6 | Asia / China / Sichuan Province | Jiahao Zhang (South China Agricultural University / College of Veterinary Medicine) |
| EPI_ISL_6757631 | A/Duck/Sichuan/21406-8/2021(H5N6) | A / H5N6 | Asia / China / Sichuan Province | Jiahao Zhang (South China Agricultural University / College of Veterinary Medicine) |
| EPI_ISL_6758007 | A/Goose/Sichuan/21418-2/2021(H5N6) | A / H5N6 | Asia / China / Sichuan Province | Jiahao Zhang (South China Agricultural University / College of Veterinary Medicine) |
| EPI_ISL_6759148 | A/Goose/Sichuan/21765-8/2021(H5N6) | A / H5N6 | Asia / China / Sichuan Province | Jiahao Zhang (South China Agricultural University / College of Veterinary Medicine) |
| EPI_ISL_6760430 | A/Duck/Sichuan/21826-1/2021(H5N6) | A / H5N6 | Asia / China / Sichuan Province | Jiahao Zhang (South China Agricultural University / College of Veterinary Medicine) |
| EPI_ISL_6760679 | A/Duck/Sichuan/21826-5/2021(H5N6) | A / H5N6 | Asia / China / Sichuan Province | Jiahao Zhang (South China Agricultural University / College of Veterinary Medicine) |
| EPI_ISL_6760681 | A/Duck/Sichuan/21840-5/2021(H5N6) | A / H5N6 | Asia / China / Sichuan Province | Jiahao Zhang (South China Agricultural University / College of Veterinary Medicine) |
| EPI_ISL_6760682 | A/Chicken/Sichuan/21859-5/2021(H5N6) | A / H5N6 | Asia / China / Sichuan Province | Jiahao Zhang (South China Agricultural University / College of Veterinary Medicine) |
| EPI_ISL_6760683 | A/Duck/Sichuan/21898/2021(H5N6) | A / H5N6 | Asia / China / Sichuan Province | Jiahao Zhang (South China Agricultural University / College of Veterinary Medicine) |
| EPI_ISL_6760684 | A/Duck/Sichuan/21921-1/2021(H5N6) | A / H5N6 | Asia / China / Sichuan Province | Jiahao Zhang (South China Agricultural University / College of Veterinary Medicine) |
| EPI_ISL_6760685 | A/Duck/Sichuan/21957-1/2021(H5N6) | A / H5N6 | Asia / China / Sichuan Province | Jiahao Zhang (South China Agricultural University / College of Veterinary Medicine) |
| EPI_ISL_6760687 | A/Goose/Guangdong/211057/2021(H5N6) | A / H5N6 | Asia / China / Guangdong Province | Jiahao Zhang (South China Agricultural University / College of Veterinary Medicine) |
| EPI_ISL_6760999 | A/black-headed gull/Netherlands/21037590-002/2021 | A / H5N1 | Europe / Netherlands / Provincie Friesland | Rene Heutink (Wageningen Bioveterinary Research) |
| EPI_ISL_6761000 | A/greylag goose/Netherlands/21037809-001/2021 | A / H5N1 | Europe / Netherlands / South Holland | Rene Heutink (Wageningen Bioveterinary Research) |
| EPI_ISL_6761001 | A/mute swan/Netherlands/21038142-001/2021 | A / H5N1 | Europe / Netherlands / Provincie Gelderland | Rene Heutink (Wageningen Bioveterinary Research) |
| EPI_ISL_6761002 | A/mute swan/Netherlands/21038142-004/2021 | A / H5N1 | Europe / Netherlands / Provincie Gelderland | Rene Heutink (Wageningen Bioveterinary Research) |
| EPI_ISL_6761003 | A/mute swan/Netherlands/21038142-007/2021 | A / H5N1 | Europe / Netherlands / Provincie Gelderland | Rene Heutink (Wageningen Bioveterinary Research) |
| EPI_ISL_6761004 | A/greylag goose/Netherlands/21037883-001/2021 | A / H5N1 | Europe / Netherlands / Provincie Flevoland | Rene Heutink (Wageningen Bioveterinary Research) |
| EPI_ISL_6761005 | A/chicken/Netherlands/21038211-001/2021 | A / H5N1 | Europe / Netherlands / Provincie Friesland | Rene Heutink (Wageningen Bioveterinary Research) |
| EPI_ISL_6761006 | A/mute swan/Netherlands/21038362-001/2021 | A / H5N1 | Europe / Netherlands / Provincie Friesland | Rene Heutink (Wageningen Bioveterinary Research) |
| EPI_ISL_6761007 | A/mute swan/Netherlands/21038362-002/2021 | A / H5N1 | Europe / Netherlands / Provincie Friesland | Rene Heutink (Wageningen Bioveterinary Research) |

| | | | | |
|---|---|---|---|---|
| EPI_ISL_6761008 | A/greylag goose/Netherlands/21038036-002/2021 | A / H5N1 | Europe / Netherlands / Provincie Noord-Holland | Rene Heutink (Wageningen Bioveterinary Research) |
| EPI_ISL_6761009 | A/greylag goose/Netherlands/21038037-001/2021 | A / H5N1 | Europe / Netherlands / Provincie Noord-Holland | Rene Heutink (Wageningen Bioveterinary Research) |
| EPI_ISL_6761010 | A/mute swan/Netherlands/21038038-001/2021 | A / H5N1 | Europe / Netherlands / Provincie Groningen | Rene Heutink (Wageningen Bioveterinary Research) |
| EPI_ISL_6761011 | A/peregrine falcon/Netherlands/21038169-002/2021 | A / H5N1 | Europe / Netherlands / Provincie Noord-Holland | Rene Heutink (Wageningen Bioveterinary Research) |
| EPI_ISL_6761012 | A/barnacle goose/Netherlands/21038248-001/2021 | A / H5N1 | Europe / Netherlands / South Holland | Rene Heutink (Wageningen Bioveterinary Research) |
| EPI_ISL_6761013 | A/greylag goose/Netherlands/21038252-001/2021 | A / H5N1 | Europe / Netherlands / Provincie Utrecht | Rene Heutink (Wageningen Bioveterinary Research) |
| EPI_ISL_6761016 | A/greylag goose/Netherlands/21038253-001/2021 | A / H5N1 | Europe / Netherlands / Provincie Utrecht | Rene Heutink (Wageningen Bioveterinary Research) |
| EPI_ISL_6761017 | A/chicken/Netherlands/21038669-001005/2021 | A / H5N1 | Europe / Netherlands / Provincie Utrecht | Rene Heutink (Wageningen Bioveterinary Research) |
| EPI_ISL_6761018 | A/mute swan/Netherlands/21038537-004/2021 | A / H5N1 | Europe / Netherlands / Provincie Noord-Holland | Rene Heutink (Wageningen Bioveterinary Research) |
| EPI_ISL_6761019 | A/mute swan/Netherlands/21038706-002/2021 | A / H5N1 | Europe / Netherlands / South Holland | Rene Heutink (Wageningen Bioveterinary Research) |
| EPI_ISL_6761020 | A/mute swan/Netherlands/21038479-002/2021 | A / H5N1 | Europe / Netherlands / Provincie Friesland | Rene Heutink (Wageningen Bioveterinary Research) |
| EPI_ISL_6761021 | A/greylag goose/Netherlands/21038565-002/2021 | A / H5N1 | Europe / Netherlands / Provincie Utrecht | Rene Heutink (Wageningen Bioveterinary Research) |
| EPI_ISL_6761022 | A/greylag goose/Netherlands/21038567-002/2021 | A / H5N1 | Europe / Netherlands / Provincie Utrecht | Rene Heutink (Wageningen Bioveterinary Research) |
| EPI_ISL_6761023 | A/mallard/Netherlands/21038796-002/2021 | A / H5N1 | Europe / Netherlands / South Holland | Rene Heutink (Wageningen Bioveterinary Research) |
| EPI_ISL_6761024 | A/goose/Netherlands/21038799-002/2021 | A / H5N1 | Europe / Netherlands / North Brabant | Rene Heutink (Wageningen Bioveterinary Research) |
| EPI_ISL_6761101 | A/Branta_leucopsis/Belgium/14735_0001/2021 | A / H5N1 | Europe / Belgium / Provincie Antwerpen / Ekerne | Steven Van Borm (Sciensano, Department of Animal Infectious Diseases / Animal Infectious Diseases) |
| EPI_ISL_6772736 | A/Chicken/Guangdong/211064-1/2021(H5N6) | A / H5N6 | Asia / China / Guangdong Province | Jiahao Zhang (South China Agricultural University / College of Veterinary Medicine) |
| EPI_ISL_6772737 | A/Chicken/Guangdong/211064-2/2021(H5N6) | A / H5N6 | Asia / China / Guangdong Province | Jiahao Zhang (South China Agricultural University / College of Veterinary Medicine) |
| EPI_ISL_6772738 | A/Chicken/Guangdong/211064-3/2021(H5N6) | A / H5N6 | Asia / China / Guangdong Province | Jiahao Zhang (South China Agricultural University / College of Veterinary Medicine) |
| EPI_ISL_6772739 | A/Chicken/Guangdong/211064-4/2021(H5N6) | A / H5N6 | Asia / China / Guangdong Province | Jiahao Zhang (South China Agricultural University / College of Veterinary Medicine) |
| EPI_ISL_6772759 | A/Chicken/Guangdong/211064-5/2021(H5N6) | A / H5N6 | Asia / China / Guangdong Province | Jiahao Zhang (South China Agricultural University / College of Veterinary Medicine) |
| EPI_ISL_6772898 | A/Goose/Guangdong/211106-2/2021(H5N6) | A / H5N6 | Asia / China / Guangdong Province | Jiahao Zhang (South China Agricultural University / College of Veterinary Medicine) |
| EPI_ISL_6780544 | A/Muscovy duck/Quang Ninh/353/2017 | A / H5N6 | Asia / Vietnam / Tinh Quang Ninh | |
| EPI_ISL_6780643 | A/Duck/Luxor/51/2018 | A / H5N8 | Africa / Egypt | |
| EPI_ISL_6780654 | A/Duck/Luxor/62/2018 | A / H5N8 | Africa / Egypt | |
| EPI_ISL_6780663 | A/chicken/Luxor/103/2018 | A / H5N8 | Africa / Egypt | |
| EPI_ISL_6780673 | A/Turkey/Luxor/240/2018 | A / H5N8 | Africa / Egypt | |
| EPI_ISL_6780682 | A/Duck/Luxor/294/2019 | A / H5N8 | Africa / Egypt | |
| EPI_ISL_6780696 | A/Duck/Luxor/297/2019 | A / H5N8 | Africa / Egypt | |
| EPI_ISL_6780727 | A/Duck/Luxor/248/2018 | A / H5N8 | Africa / Egypt | |
| EPI_ISL_6780760 | A/chicken/Luxor/313/2019 | A / H5N8 | Africa / Egypt | |
| EPI_ISL_6781195 | A/Whooper swan/Mongolia/01/2020 | A / H5N6 | Asia / Mongolia | |
| EPI_ISL_6781200 | A/Whooper swan/Mongolia/03/2020 | A / H5N6 | Asia / Mongolia | |
| EPI_ISL_6781352 | A/goose/China/Wuhu01/2019 | A / H5N6 | Asia / China | |
| EPI_ISL_6781353 | A/goose/China/Yizheng11/2016 | A / H5N6 | Asia / China | |
| EPI_ISL_6781354 | A/goose/China/BaoYing09/2016 | A / H5N6 | Asia / China | |
| EPI_ISL_6781355 | A/chicken/China/HJ11/2017 | A / H5N6 | Asia / China | |
| EPI_ISL_6781356 | A/chicken/China/LA10/2017 | A / H5N6 | Asia / China | |
| EPI_ISL_6781405 | A/Whooper Swan/Khuvsgul/#1/2020 | A / H5N6 | Asia / Mongolia | |
| EPI_ISL_6781406 | A/Whooper Swan/Khuvsgul/#4/2020 | A / H5N6 | Asia / Mongolia | |
| EPI_ISL_6781491 | A/Turkey/Egypt/A2/2021 | A / H5N8 | Africa / Egypt | |
| EPI_ISL_6795229 | A/duck/China/FJ1807/2018 | A / H5N6 | Asia / China | |
| EPI_ISL_6795230 | A/duck/China/FJ1808/2018 | A / H5N6 | Asia / China | |
| EPI_ISL_6795231 | A/duck/China/FJ18105/2018 | A / H5N6 | Asia / China | |
| EPI_ISL_6795232 | A/duck/China/FJ18106/2018 | A / H5N6 | Asia / China | |
| EPI_ISL_6795233 | A/duck/China/FJ18107/2018 | A / H5N6 | Asia / China | |
| EPI_ISL_6795234 | A/duck/China/FJ18108/2018 | A / H5N6 | Asia / China | |
| EPI_ISL_6795235 | A/duck/China/FJ1813/2018 | A / H5N6 | Asia / China | |
| EPI_ISL_6795236 | A/duck/China/FJ18203/2018 | A / H5N6 | Asia / China | |
| EPI_ISL_6795237 | A/duck/China/FJ1543/2015 | A / H5N6 | Asia / China | |
| EPI_ISL_6795238 | A/duck/China/FJ18210/2018 | A / H5N6 | Asia / China | |
| EPI_ISL_6795239 | A/duck/China/FJ18212/2018 | A / H5N6 | Asia / China | |
| EPI_ISL_6795240 | A/duck/China/FJ18220/2018 | A / H5N6 | Asia / China | |
| EPI_ISL_6795241 | A/duck/China/FJ1519/2015 | A / H5N6 | Asia / China | |
| EPI_ISL_6795242 | A/duck/China/FJ1545/2015 | A / H5N6 | Asia / China | |
| EPI_ISL_6795243 | A/duck/China/FJ1563/2015 | A / H5N6 | Asia / China | |
| EPI_ISL_6795244 | A/duck/China/FJ1567/2016 | A / H5N6 | Asia / China | |
| EPI_ISL_6795245 | A/duck/China/FJ1602/2016 | A / H5N6 | Asia / China | |
| EPI_ISL_6795246 | A/duck/China/FJ1683/2016 | A / H5N6 | Asia / China | |
| EPI_ISL_6795247 | A/duck/China/FJ17139/2017 | A / H5N6 | Asia / China | |
| EPI_ISL_6795248 | A/duck/China/FJ17152/2017 | A / H5N6 | Asia / China | |
| EPI_ISL_6795249 | A/duck/China/FJ19179/2017 | A / H5N6 | Asia / China | |
| EPI_ISL_6795250 | A/duck/China/JX1519/2015 | A / H5N6 | Asia / China | |
| EPI_ISL_6795251 | A/duck/China/JX1545/2015 | A / H5N6 | Asia / China | |
| EPI_ISL_6795252 | A/duck/China/ZJ1604/2016 | A / H5N6 | Asia / China | |
| EPI_ISL_6795253 | A/duck/China/ZJ1504/2015 | A / H5N6 | Asia / China | |
| EPI_ISL_6795254 | A/duck/China/ZJ1532/2015 | A / H5N6 | Asia / China | |
| EPI_ISL_6795255 | A/duck/China/ZJ1606/2016 | A / H5N6 | Asia / China | |
| EPI_ISL_6795256 | A/duck/China/FJ18231/2019 | A / H5N6 | Asia / China | |
| EPI_ISL_6795257 | A/duck/China/FJ18248/2019 | A / H5N6 | Asia / China | |
| EPI_ISL_6795258 | A/duck/China/FJ18252/2019 | A / H5N6 | Asia / China | |
| EPI_ISL_6795259 | A/duck/China/FJ18262/2018 | A / H5N6 | Asia / China | |

| | | | | |
|---|---|---|---|---|
| EPI_ISL_6795260 | A/duck/China/FJ18271/2018 | A / H5N6 | Asia / China | |
| EPI_ISL_6795261 | A/duck/China/FJ1829/2019 | A / H5N6 | Asia / China | |
| EPI_ISL_6795262 | A/duck/China/FJ18299/2018 | A / H5N6 | Asia / China | |
| EPI_ISL_6795263 | A/duck/China/FJ1854/2018 | A / H5N6 | Asia / China | |
| EPI_ISL_6795264 | A/duck/China/FJ1904/2019 | A / H5N6 | Asia / China | |
| EPI_ISL_6795265 | A/duck/China/FJ1914/2019 | A / H5N6 | Asia / China | |
| EPI_ISL_6795266 | A/duck/China/FJ1915/2019 | A / H5N6 | Asia / China | |
| EPI_ISL_6795267 | A/duck/China/FJ1921/2019 | A / H5N6 | Asia / China | |
| EPI_ISL_6795268 | A/duck/China/FJ1922/2019 | A / H5N6 | Asia / China | |
| EPI_ISL_6795269 | A/duck/China/FJ1923/2019 | A / H5N6 | Asia / China | |
| EPI_ISL_6795270 | A/duck/China/FJ1931/2019 | A / H5N6 | Asia / China | |
| EPI_ISL_6795271 | A/duck/China/FJ19323/2019 | A / H5N6 | Asia / China | |
| EPI_ISL_6795272 | A/duck/China/FJ19332/2019 | A / H5N6 | Asia / China | |
| EPI_ISL_6795273 | A/duck/China/FJ1935/2019 | A / H5N6 | Asia / China | |
| EPI_ISL_681271 | A/chicken/Kagawa/F2T/2020 | A / H5N8 | Asia / Japan / Kagawa | Takehiko Saito (National Institute of Animal Health) |
| EPI_ISL_681272 | A/chicken/Kagawa/F3C/2020 | A / H5N8 | Asia / Japan / Kagawa | Takehiko Saito (National Institute of Animal Health) |
| EPI_ISL_681273 | A/chicken/Kagawa/G12C/2020 | A / H5N8 | Asia / Japan / Kagawa | Takehiko Saito (National Institute of Animal Health) |
| EPI_ISL_681274 | A/chicken/Kagawa/G4T/2020 | A / H5N8 | Asia / Japan / Kagawa | Takehiko Saito (National Institute of Animal Health) |
| EPI_ISL_681275 | A/chicken/Kagawa/H2T/2020 | A / H5N8 | Asia / Japan / Kagawa | Takehiko Saito (National Institute of Animal Health) |
| EPI_ISL_681276 | A/chicken/Kagawa/H8C/2020 | A / H5N8 | Asia / Japan / Kagawa | Takehiko Saito (National Institute of Animal Health) |
| EPI_ISL_681281 | A/swan/Niigata/151118/2020 | A / H5N8 | Asia / Japan / Niigata | Takehiko Saito (National Institute of Animal Health) |
| EPI_ISL_681282 | A/chicken/Fukuoka/T1/2020 | A / H5N8 | Asia / Japan / Fukuoka | Takehiko Saito (National Institute of Animal Health) |
| EPI_ISL_681283 | A/chicken/Fukuoka/T2/2020 | A / H5N8 | Asia / Japan / Fukuoka | Takehiko Saito (National Institute of Animal Health) |
| EPI_ISL_681284 | A/chicken/Hyogo/1T/2020 | A / H5N8 | Asia / Japan / Hyogo | Takehiko Saito (National Institute of Animal Health) |
| EPI_ISL_681285 | A/chicken/Hyogo/2T/2020 | A / H5N8 | Asia / Japan / Hyogo | Takehiko Saito (National Institute of Animal Health) |
| EPI_ISL_681286 | A/chicken/Kagawa/11C/2020 | A / H5N8 | Asia / Japan / Kagawa | Takehiko Saito (National Institute of Animal Health) |
| EPI_ISL_681287 | A/chicken/Kagawa/7C/2020 | A / H5N8 | Asia / Japan / Kagawa | Takehiko Saito (National Institute of Animal Health) |
| EPI_ISL_681288 | A/chicken/Kagawa/B10T/2020 | A / H5N8 | Asia / Japan / Kagawa | Takehiko Saito (National Institute of Animal Health) |
| EPI_ISL_681289 | A/chicken/Kagawa/B13C/2020 | A / H5N8 | Asia / Japan / Kagawa | Takehiko Saito (National Institute of Animal Health) |
| EPI_ISL_681290 | A/chicken/Kagawa/B8T/2020 | A / H5N8 | Asia / Japan / Kagawa | Takehiko Saito (National Institute of Animal Health) |
| EPI_ISL_681291 | A/chicken/Kagawa/C3C/2020 | A / H5N8 | Asia / Japan / Kagawa | Takehiko Saito (National Institute of Animal Health) |
| EPI_ISL_681292 | A/chicken/Kagawa/C3T/2020 | A / H5N8 | Asia / Japan / Kagawa | Takehiko Saito (National Institute of Animal Health) |
| EPI_ISL_681293 | A/chicken/Kagawa/C5T/2020 | A / H5N8 | Asia / Japan / Kagawa | Takehiko Saito (National Institute of Animal Health) |
| EPI_ISL_681294 | A/chicken/Kagawa/D2C/2020 | A / H5N8 | Asia / Japan / Kagawa | Takehiko Saito (National Institute of Animal Health) |
| EPI_ISL_681295 | A/chicken/Kagawa/D5T/2020 | A / H5N8 | Asia / Japan / Kagawa | Takehiko Saito (National Institute of Animal Health) |
| EPI_ISL_681296 | A/chicken/Kagawa/E13T/2020 | A / H5N8 | Asia / Japan / Kagawa | Takehiko Saito (National Institute of Animal Health) |
| EPI_ISL_681297 | A/chicken/Kagawa/E7T/2020 | A / H5N8 | Asia / Japan / Kagawa | Takehiko Saito (National Institute of Animal Health) |
| EPI_ISL_6816050 | A/Duck/Egypt/BEH2/2020 | A / H5N8 | Africa / Egypt | |
| EPI_ISL_6816051 | A/chicken/Egypt/GIZ4/2020 | A / H5N8 | Africa / Egypt | |
| EPI_ISL_6816052 | A/chicken/Egypt/MEN6/2020 | A / H5N8 | Africa / Egypt | |
| EPI_ISL_6816053 | A/chicken/Egypt/MEN10/2020 | A / H5N8 | Africa / Egypt | |
| EPI_ISL_6816054 | A/chicken/Egypt/CAI11/2020 | A / H5N8 | Africa / Egypt | |
| EPI_ISL_6816057 | A/environment sample/China/TZ001/2021 | A / H5N8 | Asia / China | |
| EPI_ISL_682297 | A/environment/Kagoshima/KU-ngr-J2/2020(H5N8) | A / H5N8 | Asia / Japan / Kagoshima | Ahmed Magdy Ahmed Khalil (Faculty of Veterinary Medicine - Zagazig University / Zoonoses) |
| EPI_ISL_6829497 | A/hooded crane/Kagoshima/KU-5T/2021 (H5N8) | A / H5N8 | Asia / Japan / Kagoshima | Makoto Ozawa (Kagoshima University ) |
| EPI_ISL_6829532 | A/chicken/Akita/7C/2021 | A / H5N8 | Asia / Japan / Akita | Takehiko Saito (National Institute of Animal Health) |
| EPI_ISL_6829533 | A/chicken/Kagoshima/21A6T/2021 | A / H5N1 | Asia / Japan / Kagoshima | Takehiko Saito (National Institute of Animal Health) |
| EPI_ISL_6829536 | A/chicken/Kagoshima/B3T/2021 | A / H5N8 | Asia / Japan / Kagoshima | Takehiko Saito (National Institute of Animal Health) |
| EPI_ISL_683592 | A/Eurasian_wigeon/Italy/20VIR7301-206/2020 | A / H5N1 | Europe / Italy / Veneto / Province of Venice / Jesolo | Bianca Zecchin (Istituto Zooprofilattico Sperimentale Delle Venezie) |
| EPI_ISL_683593 | A/Eurasian_wigeon/Italy/20VIR7139-121/2020 | A / H5N8 | Europe / Italy / Veneto / Province of Rovigo / Valle Ca' Zuliani | Bianca Zecchin (Istituto Zooprofilattico Sperimentale Delle Venezie) |
| EPI_ISL_683594 | A/mallard/Italy/20VIR7139-124_feather/2020 | A / H5N8 | Europe / Italy / Veneto / Province of Rovigo / Valle Ca' Zuliani | Bianca Zecchin (Istituto Zooprofilattico Sperimentale Delle Venezie) |
| EPI_ISL_683751 | A/Eurasian_wigeon/Italy/20VIR7301-31/2020 | A / H5N8 | Europe / Italy / Veneto / Province of Venice / Jesolo | Bianca Zecchin (Istituto Zooprofilattico Sperimentale Delle Venezie) |
| EPI_ISL_683752 | A/Eurasian_wigeon/Italy/20VIR7301-34/2020 | A / H5N8 | Europe / Italy / Veneto / Province of Venice / Jesolo | Bianca Zecchin (Istituto Zooprofilattico Sperimentale Delle Venezie) |
| EPI_ISL_683997 | A/mute swan/Wales/048069/2020 | A / H5N5 | Europe / United Kingdom | Alex Byrne (Animal and Plant Health Agency (APHA) / Virology Department) |
| EPI_ISL_683999 | A/mute swan/Wales/048068/2020 | A / H5N5 | Europe / United Kingdom | Alex Byrne (Animal and Plant Health Agency (APHA) / Virology Department) |
| EPI_ISL_684037 | A/goose/Tatarstan/1730-2/2020 | A / H5N8 | Europe / Russian Federation / Republic of Tatarstan | Nikolay Zinyakov (Federal Centre for Animal Health (ARRIAH) / OIE Regional Reference Laboratory) |
| EPI_ISL_6914131 | A/chicken/chongqing/H1/2021 | A / H5N6 | Asia / China / Chongqinq Municipality | Wenming Jiang (China Animal Health and Epidemiology Center (CAHEC)) |
| EPI_ISL_6929958 | A/turkey/Poland/H1910-T3/2021 | A / H5N1 | Europe / Poland / Masovian Voivodeship | Edyta ?wi?to? (National Veterinary Research Institut Poland, PIWet-PIB) |
| EPI_ISL_6929970 | A/turkey/Poland/H1911-N/2021 | A / H5N1 | Europe / Poland / Masovian Voivodeship | Edyta ?wi?to? (National Veterinary Research Institut Poland, PIWet-PIB) |
| EPI_ISL_6930238 | A/turkey/Poland/H1913-T1/2021 | A / H5N1 | Europe / Poland / Masovian Voivodeship | Edyta ?wi?to? (National Veterinary Research Institut Poland, PIWet-PIB) |
| EPI_ISL_6930564 | A/turkey/Poland/H1924-T1/2021 | A / H5N1 | Europe / Poland / Masovian Voivodeship | Edyta ?wi?to? (National Veterinary Research Institut Poland, PIWet-PIB) |
| EPI_ISL_6931008 | A/domestic_goose/Poland/H1931-T1/2021 | A / H5N1 | Europe / Poland / Lubusz Voivodeship | Edyta ?wi?to? (National Veterinary Research Institut Poland, PIWet-PIB) |
| EPI_ISL_6931288 | A/chicken/Poland/H1940-N/2021 | A / H5N1 | Europe / Poland / Silesian Voivodeship | Edyta ?wi?to? (National Veterinary Research Institut Poland, PIWet-PIB) |
| EPI_ISL_6934175 | A/domestic_duck/Poland/H1942-N/2021 | A / H5N1 | Europe / Poland / Lodz Voivodeship | Edyta ?wi?to? (National Veterinary Research Institut Poland, PIWet-PIB) |
| EPI_ISL_693515 | A/chicken/Netherlands/20019411-006010/2020 | A / H5N8 | Europe / Netherlands / South Holland / Midden-Delfland | Rene Heutink (Wageningen Bioveterinary Research) |
| EPI_ISL_6935584 | A/turkey/Poland/H1944-N/2021 | A / H5N1 | Europe / Poland / Masovian Voivodeship | Edyta ?wi?to? (National Veterinary Research Institut Poland, PIWet-PIB) |
| EPI_ISL_6937114 | A/mute_swan/Poland/MB490-L1/2021 | A / H5N1 | Europe / Poland / Greater Poland Voivodeship | Edyta ?wi?to? (National Veterinary Research Institut Poland, PIWet-PIB) |
| EPI_ISL_694042 | A/chicken/Vietnam/NCVD-17A505/2017 | A / H5N6 | Asia / Vietnam | Yunho Jang (Centers for Disease Control and Prevention / WHO Collaborating Center for Surveillance, Epidemiology and Control of Influ |
| EPI_ISL_6959592 | A/mandarin duck/Korea/WA585/2021 | A / H5N1 | Asia / Korea, Republic of / Chungcheongnam-do | Yu-Na Lee (Animal and Plant Quarantine Agency (S-2158) / Avian Influenza Research & Diagnostic Division) |
| EPI_ISL_6959593 | A/quail/Korea/H526/2021 | A / H5N1 | Asia / Korea, Republic of / Chungcheongbuk-do | Yu-Na Lee (Animal and Plant Quarantine Agency (S-2158) / Avian Influenza Research & Diagnostic Division) |
| EPI_ISL_696629 | A/chicken/Shanxi/04.28_TGRL001-O/2018 | A / H5N6 | Asia / China / Shanxi Province | |
| EPI_ISL_696630 | A/chicken/Shanxi/04.28_TGRL024-O/2018 | A / H5N6 | Asia / China / Shanxi Province | |

| | | | |
|---|---|---|---|
| EPI_ISL_696686 | A/duck/Jiangxi/11.29_NCNP22D3-OC/2017 | A / H5N6 | Asia / China / Jiangxi Province |
| EPI_ISL_696689 | A/duck/Hunan/03.07_YYGK25L3-OC/2018 | A / H5N6 | Asia / China / Hunan Province |
| EPI_ISL_696703 | A/duck/Jiangxi/11.29_NCNP39D3-OC/2017 | A / H5N6 | Asia / China / Jiangxi Province |
| EPI_ISL_696707 | A/duck/Hunan/7.21_YYGK41Y1-C/2016 | A / H5N6 | Asia / China / Hunan Province |
| EPI_ISL_696739 | A/chicken/Hunan/03.07_YYGK74L3-OC/2018 | A / H5N6 | Asia / China / Hunan Province |
| EPI_ISL_696740 | A/duck/Hunan/11.30_YYGK75E3-OC/2017 | A / H5N6 | Asia / China / Hunan Province |
| EPI_ISL_696807 | A/duck/Guizhou/7.27_ZYLJJ017-O/2018 | A / H5N6 | Asia / China / Guizhou Province |
| EPI_ISL_696813 | A/duck/Guizhou/8.26_ZYLJJ016-O/2018 | A / H5N6 | Asia / China / Guizhou Province |
| EPI_ISL_696836 | A/chicken/Guizhou/10.28_ZYLJJ008-C/2018 | A / H5N6 | Asia / China / Guizhou Province |
| EPI_ISL_696843 | A/pigeon/Hainan/1.14_HKPL006-C/2018 | A / H5N6 | Asia / China / Hainan Province |
| EPI_ISL_696943 | A/duck/Yunnan/9.27_DQXY003-O/2016 | A / H5N6 | Asia / China / Yunnan Province |
| EPI_ISL_696991 | A/goose/Fujian/3.15_FZHX0001-O/2018 | A / H5N6 | Asia / China / Fujian Province |
| EPI_ISL_696992 | A/goose/Fujian/3.15_FZHX0007-O/2018 | A / H5N6 | Asia / China / Fujian Province |
| EPI_ISL_696993 | A/goose/Fujian/3.15_FZHX0010-O/2018 | A / H5N6 | Asia / China / Fujian Province |
| EPI_ISL_696994 | A/goose/Fujian/3.15_FZHX0005-C/2018 | A / H5N6 | Asia / China / Fujian Province |
| EPI_ISL_696995 | A/goose/Fujian/3.15_FZHX0007-C/2018 | A / H5N6 | Asia / China / Fujian Province |
| EPI_ISL_697004 | A/duck/Guangdong/7.20_DGCP036-O/2017 | A / H5N6 | Asia / China / Guangdong Province |
| EPI_ISL_697077 | A/chicken/Shandong/8.28_TAWM017-O/2017 | A / H5N6 | Asia / China / Shandong Province |
| EPI_ISL_697143 | A/chicken/Fujian/11.23_FZHX0014-O/2017 | A / H5N6 | Asia / China / Fujian Province |
| EPI_ISL_697144 | A/chicken/Fujian/11.23_FZHX0015-O/2017 | A / H5N6 | Asia / China / Fujian Province |
| EPI_ISL_697188 | A/duck/Fujian/10.26_FZHX0034-C/2017 | A / H5N6 | Asia / China / Fujian Province |
| EPI_ISL_697189 | A/duck/Fujian/11.09_FZHX-O/2017 | A / H5N6 | Asia / China / Fujian Province |
| EPI_ISL_697210 | A/goose/Fujian/11.23_FZHX0008-O/2017 | A / H5N6 | Asia / China / Fujian Province |
| EPI_ISL_697211 | A/goose/Fujian/11.23_FZHX0009-O/2017 | A / H5N6 | Asia / China / Fujian Province |
| EPI_ISL_697214 | A/goose/Fujian/3.15_FZHX0005-O/2018 | A / H5N6 | Asia / China / Fujian Province |
| EPI_ISL_697216 | A/goose/Fujian/3.15_FZHX0008-O/2018 | A / H5N6 | Asia / China / Fujian Province |
| EPI_ISL_697697 | A/chicken/Egypt/ME-2018/2018 | A / H5N8 | Africa / Egypt |
| EPI_ISL_697771 | A/northern pintail/Hokkaido/M13/2020 | A / H5N8 | Asia / Japan / Hokkaido |
| EPI_ISL_697804 | A/chicken/Shandong/2.28_TAWM016-C/2017 | A / H5N6 | Asia / China / Shandong Province |
| EPI_ISL_697808 | A/chicken/Fujian/11.23_FZHX0006-O/2017 | A / H5N6 | Asia / China / Fujian Province |
| EPI_ISL_697819 | A/chicken/Guizhou/12.20_ZYLJJ017-O/2017 | A / H5N6 | Asia / China / Guizhou Province |
| EPI_ISL_697826 | A/chicken/Fujian/3.15_FZHX0009-C/2018 | A / H5N2 | Asia / China / Fujian Province |
| EPI_ISL_697829 | A/duck/Fujian/11.23_FZHX0010-C/2017 | A / H5N2 | Asia / China / Fujian Province |
| EPI_ISL_697830 | A/duck/Fujian/11.23_FZHX0018-O/2017 | A / H5N2 | Asia / China / Fujian Province |
| EPI_ISL_697831 | A/duck/Fujian/11.23_FZHX0020-O/2017 | A / H5N2 | Asia / China / Fujian Province |
| EPI_ISL_697833 | A/chicken/Guizhou/8.26_ZYLJJ014-O/2018 | A / H5N6 | Asia / China / Guizhou Province |
| EPI_ISL_697834 | A/chicken/Shandong/12.26_TAWL007-O/2018 | A / H5N6 | Asia / China / Shandong Province |
| EPI_ISL_697835 | A/chicken/Shandong/12.26_TAWL015-O/2018 | A / H5N6 | Asia / China / Shandong Province |
| EPI_ISL_697837 | A/duck/Henan/01.01_LYGL003-O/2017 | A / H5N6 | Asia / China / Henan Province |
| EPI_ISL_697839 | A/chicken/Shanxi/11.30_TGRL017-O/2018 | A / H5N6 | Asia / China / Shanxi Province |
| EPI_ISL_697840 | A/chicken/Shanxi/11.30_TGRL008-C/2018 | A / H5N6 | Asia / China / Shanxi Province |
| EPI_ISL_697841 | A/chicken/Shanxi/12.01_TGRL025-C/2018 | A / H5N6 | Asia / China / Shanxi Province |
| EPI_ISL_697842 | A/chicken/Henan/10.31_XXHM015-C/2018 | A / H5N6 | Asia / China / Henan Province |
| EPI_ISL_697846 | A/chicken/Henan/11.29_XXHM014-C/2018 | A / H5N6 | Asia / China / Henan Province |
| EPI_ISL_697847 | A/chicken/Shanxi/11.30_TGRL008-O/2018 | A / H5N6 | Asia / China / Shanxi Province |
| EPI_ISL_697848 | A/chicken/Shanxi/12.01_TGRL016-O/2018 | A / H5N6 | Asia / China / Shanxi Province |
| EPI_ISL_697851 | A/chicken/Hunan/12.27_YYGK19J2-C/2016 | A / H5N6 | Asia / China / Hunan Province |
| EPI_ISL_697884 | A/chicken/Hunan/03.07_YYGK60L3-OC/2018 | A / H5N2 | Asia / China / Hunan Province |
| EPI_ISL_697885 | A/chicken/Hunan/3.07_YYGK81L3-OC/2018 | A / H5N6 | Asia / China / Hunan Province |
| EPI_ISL_697888 | A/goose/Hunan/1.11_YYGK92E3-OC/2018 | A / H5N6 | Asia / China / Hunan Province |
| EPI_ISL_697891 | A/chicken/Anhui/9.29_YHZGS014-O/2018 | A / H5N6 | Asia / China / Anhui Province |
| EPI_ISL_697897 | A/environment/Anhui/01.20_YHZGS003-E/2019 | A / H5N6 | Asia / China / Anhui Province |
| EPI_ISL_697898 | A/environment/Anhui/01.20_YHZGS004-E/2019 | A / H5N2 | Asia / China / Anhui Province |
| EPI_ISL_697901 | A/chicken/Shandong/01.26_TAWL003-O/2019 | A / H5N6 | Asia / China / Shandong Province |
| EPI_ISL_697903 | A/chicken/Shandong/01.26_TAWL011-O/2019 | A / H5N6 | Asia / China / Shandong Province |
| EPI_ISL_697904 | A/chicken/Shandong/01.26_TAWL017-O/2019 | A / H5N6 | Asia / China / Shandong Province |
| EPI_ISL_697905 | A/chicken/Shandong/01.26_TAWL022-O/2019 | A / H5N2 | Asia / China / Shandong Province |
| EPI_ISL_697907 | A/chicken/Shandong/01.26_TAWL007-C/2019 | A / H5N6 | Asia / China / Shandong Province |
| EPI_ISL_697908 | A/chicken/Shandong/01.26_TAWL009-C/2019 | A / H5N6 | Asia / China / Shandong Province |
| EPI_ISL_697920 | A/duck/Zhejiang/8.25_HZBX003-O/2018 | A / H5 | Asia / China / Zhejiang Province |
| EPI_ISL_697941 | A/duck/Zhejiang/11.09_HZBX009-C/2018 | A / H5 | Asia / China / Zhejiang Province |
| EPI_ISL_697946 | A/duck/Zhejiang/12.28_HZBX012-O/2018 | A / H5 | Asia / China / Zhejiang Province |
| EPI_ISL_697963 | A/chicken/Guangxi/8.25_NNXH008-O/2018 | A / H5 | Asia / China / Guangxi Zhuang Autonomous Region |
| EPI_ISL_697983 | A/chicken/Yunnan/11.22_DQWGH005-O/2018 | A / H5 | Asia / China / Yunnan Province |
| EPI_ISL_697984 | A/environment/Yunnan/11.22_DQWGH002-E/2018 | A / H5N6 | Asia / China / Yunnan Province |
| EPI_ISL_697985 | A/chicken/Fujian/9.24_FZHX0067-O/2018 | A / H5N6 | Asia / China / Fujian Province |
| EPI_ISL_697986 | A/chicken/Fujian/9.24_FZHX0068-C/2018 | A / H5N6 | Asia / China / Fujian Province |
| EPI_ISL_697987 | A/chicken/Fujian/9.24_FZHX0071-O/2018 | A / H5 | Asia / China / Fujian Province |
| EPI_ISL_697989 | A/chicken/Fujian/9.24_FZHX0076-O/2018 | A / H5 | Asia / China / Fujian Province |
| EPI_ISL_697990 | A/duck/Fujian/1.25_FZHX0041-O/2018 | A / H5 | Asia / China / Fujian Province |
| EPI_ISL_697991 | A/chicken/Fujian/9.24_FZHX0088-C/2018 | A / H5N6 | Asia / China / Fujian Province |
| EPI_ISL_698002 | A/chicken/Jiangxi/4.25_NCDZT53Q3-OC/2018 | A / H5N6 | Asia / China / Jiangxi Province |

| | | | | |
|---|---|---|---|---|
| EPI_ISL_698004 | A/duck/Jiangxi/6.21_NCNP83S3-OC/2018 | A / H5N6 | Asia / China / Jiangxi Province | |
| EPI_ISL_698008 | A/duck/Jiangxi/12.17_NCNP71D4-O/2018 | A / H5N6 | Asia / China / Jiangxi Province | |
| EPI_ISL_698009 | A/duck/Hunan/04.26_YYGK75R3-OC/2018 | A / H5N6 | Asia / China / Hunan Province | |
| EPI_ISL_698010 | A/chicken/Hunan/04.26_YYGK19R3-OC/2018 | A / H5N6 | Asia / China / Hunan Province | |
| EPI_ISL_698011 | A/duck/Hunan/06.22_YYGK90T3-OC/2018 | A / H5N6 | Asia / China / Hunan Province | |
| EPI_ISL_698012 | A/chicken/Hunan/07.26_YYGK28V3-OC/2018 | A / H5N6 | Asia / China / Hunan Province | |
| EPI_ISL_698013 | A/chicken/Hunan/7.26_YYGK56V3-OC/2018 | A / H5N6 | Asia / China / Hunan Province | |
| EPI_ISL_698015 | A/chicken/Jiangxi/5.28_NCDZT44N3-OC/2018 | A / H5 | Asia / China / Jiangxi Province | |
| EPI_ISL_698017 | A/duck/Jiangxi/5.28_NCNP34N3-OC/2018 | A / H5 | Asia / China / Jiangxi Province | |
| EPI_ISL_698020 | A/duck/Jiangxi/05.28_NCNP30N3-OC/2018 | A / H5 | Asia / China / Jiangxi Province | |
| EPI_ISL_698022 | A/duck/Jiangxi/05.28_NCNP17N3-OC/2018 | A / H5 | Asia / China / Jiangxi Province | |
| EPI_ISL_698025 | A/duck/Jiangxi/5.28_NCNP12N3-OC/2018 | A / H5 | Asia / China / Jiangxi Province | |
| EPI_ISL_698031 | A/chicken/Jiangxi/09.22_NCNP6Z3-OC/2018 | A / H5 | Asia / China / Jiangxi Province | |
| EPI_ISL_698034 | A/duck/Jiangxi/9.22_NCDZT76Z3-OC/2018 | A / H5 | Asia / China / Jiangxi Province | |
| EPI_ISL_698045 | A/chicken/Hunan/4.26_YYGK62R3-OC/2018 | A / H5 | Asia / China / Hunan Province | |
| EPI_ISL_698047 | A/duck/Hunan/4.26_YYGK69R3-OC/2018 | A / H5 | Asia / China / Hunan Province | |
| EPI_ISL_698052 | A/chicken/Hunan/04.26_YYGK37R3-OC/2018 | A / H5 | Asia / China / Hunan Province | |
| EPI_ISL_698060 | A/duck/Hunan/09.22_YYGK26Y3-OC/2018 | A / H5 | Asia / China / Hunan Province | |
| EPI_ISL_698070 | A/chicken/Anhui/2.22_YHZGS003-O/2019 | A / H5N6 | Asia / China / Anhui Province | |
| EPI_ISL_698071 | A/chicken/Anhui/2.22_YHZGS007-O/2019 | A / H5N2 | Asia / China / Anhui Province | |
| EPI_ISL_699166 | A/chicken/Anhui/8.28_YHZGS017-O/2018 | A / H5N6 | Asia / China / Anhui Province | |
| EPI_ISL_699204 | A/chicken/Fujian/11.23_FZHX0007-O/2017 | A / H5N6 | Asia / China / Fujian Province | |
| EPI_ISL_699429 | A/chicken/Hunan/02.06_YYGK37J3-OC/2018 | A / H5N6 | Asia / China / Hunan Province | |
| EPI_ISL_699451 | A/chicken/Hunan/12.27_YYGK55J2-O/2016 | A / H5N6 | Asia / China / Hunan Province | |
| EPI_ISL_699454 | A/chicken/Hunan/12.27_YYGKK148-OC/2016 | A / H5N6 | Asia / China / Hunan Province | |
| EPI_ISL_7049600 | A/bean goose/Sweden/SVA211111SZ0372/FB004482/2021 | A / H5N1 | Europe / Sweden / Skane Lan / Simrishamns Kommun | Siamak Zohari (National Veterinary Institute) |
| EPI_ISL_7050532 | A/western jackdaw/Sweden/SVA211111SZ0376/FB004483/202 | A / H5N1 | Europe / Sweden / Skane Lan / Malmo Kommun | Siamak Zohari (National Veterinary Institute) |
| EPI_ISL_7053000 | A/common buzzard/Sweden/SVA211111SZ0376/FB004484/20 | A / H5N1 | Europe / Sweden / Skane Lan / Malmo Kommun | Siamak Zohari (National Veterinary Institute) |
| EPI_ISL_7053817 | A/barnacle goose/Sweden/SVA211111SZ0376/FB004496/2021 | A / H5N1 | Europe / Sweden / Skane Lan / Malmo Kommun | Siamak Zohari (National Veterinary Institute) |
| EPI_ISL_7054529 | A/greylag goose /Sweden/SVA211111SZ0376/FB004497/M-20 | A / H5N1 | Europe / Sweden / Skane Lan / Malmo Kommun | Siamak Zohari (National Veterinary Institute) |
| EPI_ISL_7054770 | A/European herring gull/Sweden/SVA211116SZ0432/FB00451 | A / H5N1 | Europe / Sweden / Hallands Lan / Hylte Kommun | Siamak Zohari (National Veterinary Institute) |
| EPI_ISL_7055384 | A/greylag goose /Sweden/SVA211118SZ0354/FB004497/I-202 | A / H5N1 | Europe / Sweden / Kalmar Lan / Kalmar Kommun | Siamak Zohari (National Veterinary Institute) |
| EPI_ISL_707015 | A/Whooper swan/Mongolia/24/2020 | A / H5N6 | Asia / Mongolia | |
| EPI_ISL_707016 | A/Whooper swan/Mongolia/25/2020 | A / H5N6 | Asia / Mongolia | |
| EPI_ISL_707363 | A/chicken/Shanxi/12.01_TGYL014-C/2016 | A / H5N6 | Asia / China / Shanxi Province | |
| EPI_ISL_707384 | A/duck/Fujian/1.17_FZHX0123-C/2017 | A / H5N6 | Asia / China / Fujian Province | |
| EPI_ISL_707414 | A/duck/Fujian/10.26_FZHX0002-O/2017 | A / H5N6 | Asia / China / Fujian Province | |
| EPI_ISL_707415 | A/duck/Fujian/10.26_FZHX0010-O/2017 | A / H5N6 | Asia / China / Fujian Province | |
| EPI_ISL_707417 | A/duck/Fujian/11.23_FZHX0009-C/2017 | A / H5N6 | Asia / China / Fujian Province | |
| EPI_ISL_707418 | A/duck/Fujian/11.23_FZHX0014-C/2017 | A / H5N6 | Asia / China / Fujian Province | |
| EPI_ISL_707419 | A/duck/Fujian/11.23_FZHX0015-C/2017 | A / H5N6 | Asia / China / Fujian Province | |
| EPI_ISL_707421 | A/duck/Fujian/11.3_FZHX1132-O/2016 | A / H5N6 | Asia / China / Fujian Province | |
| EPI_ISL_707422 | A/duck/Fujian/11.3_FZHX1141-C/2016 | A / H5N6 | Asia / China / Fujian Province | |
| EPI_ISL_707425 | A/duck/Fujian/12.7_FZHX1202-O/2016 | A / H5N6 | Asia / China / Fujian Province | |
| EPI_ISL_707426 | A/duck/Fujian/12.7_FZHX1218-C/2016 | A / H5N6 | Asia / China / Fujian Province | |
| EPI_ISL_707427 | A/duck/Fujian/12.7_FZHX1219-O/2016 | A / H5N6 | Asia / China / Fujian Province | |
| EPI_ISL_707428 | A/duck/Fujian/12.7_FZHX1221-C/2016 | A / H5N6 | Asia / China / Fujian Province | |
| EPI_ISL_707453 | A/duck/Guangdong/11.18_SZBJ003-C/2016 | A / H5N6 | Asia / China / Guangdong Province | |
| EPI_ISL_707454 | A/duck/Guangdong/11.18_SZBJ003-O/2016 | A / H5N6 | Asia / China / Guangdong Province | |
| EPI_ISL_707455 | A/duck/Guangdong/11.18_SZBJ007-O/2016 | A / H5N6 | Asia / China / Guangdong Province | |
| EPI_ISL_707465 | A/duck/Guizhou/8.26_ZYLJJ017-O/2018 | A / H5N6 | Asia / China / Guizhou Province | |
| EPI_ISL_707469 | A/duck/Hainan/12.29_HKPL002-C/2017 | A / H5N6 | Asia / China / Hainan Province | |
| EPI_ISL_707475 | A/duck/Hunan/01.12_YYGK82H3-OC/2018 | A / H5N6 | Asia / China / Hunan Province | |
| EPI_ISL_707477 | A/duck/Hunan/03.07_YYGK4L3-OC/2018 | A / H5N6 | Asia / China / Hunan Province | |
| EPI_ISL_707478 | A/duck/Hunan/03.24_YYGK174M2-C/2017 | A / H5N6 | Asia / China / Hunan Province | |
| EPI_ISL_707479 | A/duck/Hunan/03.24_YYGK178M2-C/2017 | A / H5N6 | Asia / China / Hunan Province | |
| EPI_ISL_707480 | A/duck/Hunan/03.24_YYGK18M2-O/2017 | A / H5N6 | Asia / China / Hunan Province | |
| EPI_ISL_707481 | A/duck/Hunan/03.24_YYGK21M2-O/2017 | A / H5N6 | Asia / China / Hunan Province | |
| EPI_ISL_707489 | A/duck/Hunan/1.12_YYGK68H3-OC/2018 | A / H5N6 | Asia / China / Hunan Province | |
| EPI_ISL_707490 | A/duck/Hunan/1.12_YYGK72H3-OC/2018 | A / H5N6 | Asia / China / Hunan Province | |
| EPI_ISL_707491 | A/duck/Hunan/1.12_YYGK74H3-OC/2018 | A / H5N6 | Asia / China / Hunan Province | |
| EPI_ISL_707493 | A/duck/Hunan/1.17_YYGKK80-OC/2017 | A / H5N6 | Asia / China / Hunan Province | |
| EPI_ISL_707500 | A/duck/Hunan/11.30_YYGK59E3-OC/2017 | A / H5N6 | Asia / China / Hunan Province | |
| EPI_ISL_707501 | A/duck/Hunan/11.30_YYGK62E3-OC/2017 | A / H5N6 | Asia / China / Hunan Province | |
| EPI_ISL_707502 | A/duck/Hunan/11.30_YYGK63E3-OC/2017 | A / H5N6 | Asia / China / Hunan Province | |
| EPI_ISL_707503 | A/duck/Hunan/11.30_YYGK74E3-OC/2017 | A / H5N8 | Asia / China / Hunan Province | |
| EPI_ISL_707504 | A/duck/Hunan/12.27_YYGK100J2-C/2016 | A / H5N6 | Asia / China / Hunan Province | |
| EPI_ISL_707506 | A/duck/Hunan/12.27_YYGK60J2-C/2016 | A / H5N6 | Asia / China / Hunan Province | |
| EPI_ISL_707507 | A/duck/Hunan/12.27_YYGK71J2-O/2016 | A / H5N6 | Asia / China / Hunan Province | |
| EPI_ISL_707508 | A/duck/Hunan/12.27_YYGK73J2-O/2016 | A / H5N6 | Asia / China / Hunan Province | |
| EPI_ISL_707510 | A/duck/Hunan/12.27_YYGK81J2-O/2016 | A / H5N6 | Asia / China / Hunan Province | |
| EPI_ISL_707512 | A/duck/Hunan/12.27_YYGK86J2-C/2016 | A / H5N6 | Asia / China / Hunan Province | |

| | | | | |
|---|---|---|---|---|
| EPI_ISL_707515 | A/duck/Hunan/12.27_YYGK89J2-O/2016 | A / H5N6 | Asia / China / Hunan Province | |
| EPI_ISL_707516 | A/duck/Hunan/12.27_YYGK91J2-O/2016 | A / H5N6 | Asia / China / Hunan Province | |
| EPI_ISL_707517 | A/duck/Hunan/12.27_YYGK93J2-O/2016 | A / H5N6 | Asia / China / Hunan Province | |
| EPI_ISL_707519 | A/duck/Hunan/12.27_YYGK95J2-O/2016 | A / H5N6 | Asia / China / Hunan Province | |
| EPI_ISL_707520 | A/duck/Hunan/12.27_YYGK96J2-O/2016 | A / H5N6 | Asia / China / Hunan Province | |
| EPI_ISL_707521 | A/duck/Hunan/12.27_YYGK97J2-O/2016 | A / H5N6 | Asia / China / Hunan Province | |
| EPI_ISL_707523 | A/duck/Hunan/2.06_YYGK77J3-OC/2018 | A / H5N6 | Asia / China / Hunan Province | |
| EPI_ISL_707524 | A/duck/Hunan/2.06_YYGK78J3-OC/2018 | A / H5N6 | Asia / China / Hunan Province | |
| EPI_ISL_707525 | A/duck/Hunan/2.06_YYGK84J3-OC/2018 | A / H5N6 | Asia / China / Hunan Province | |
| EPI_ISL_707527 | A/duck/Hunan/2.06_YYGK92J3-OC/2018 | A / H5N6 | Asia / China / Hunan Province | |
| EPI_ISL_707533 | A/duck/Jiangxi/01.11_NCNP101G2-OC/2017 | A / H5N6 | Asia / China / Jiangxi Province | |
| EPI_ISL_707543 | A/duck/Jiangxi/11.17_NCNP30Y2-O/2016 | A / H5N6 | Asia / China / Jiangxi Province | |
| EPI_ISL_707544 | A/duck/Jiangxi/11.17_NCNP48Y2-C/2016 | A / H5N6 | Asia / China / Jiangxi Province | |
| EPI_ISL_707546 | A/duck/Jiangxi/11.29_NCDZT71H2-O/2016 | A / H5N6 | Asia / China / Jiangxi Province | |
| EPI_ISL_707554 | A/duck/Jiangxi/2.28NCNP23K3-OC/2018 | A / H5N6 | Asia / China / Jiangxi Province | |
| EPI_ISL_707555 | A/duck/Jiangxi/2.28_NCNP25K3-OC/2018 | A / H5N6 | Asia / China / Jiangxi Province | |
| EPI_ISL_707570 | A/duck/Jiangxi/5.26_NCDZT80Q2-OC/2017 | A / H5N6 | Asia / China / Jiangxi Province | |
| EPI_ISL_707581 | A/duck/Xinjiang/12.18_WLMQXL001-C/2016 | A / H5N8 | Asia / China / Xinjiang Uyghur Autonomous Region | |
| EPI_ISL_707641 | A/goose/Fujian/10.11_FZHX1001-C/2016 | A / H5N6 | Asia / China / Fujian Province | |
| EPI_ISL_707642 | A/goose/Fujian/10.11_FZHX1002-O/2016 | A / H5N6 | Asia / China / Fujian Province | |
| EPI_ISL_707643 | A/goose/Fujian/10.26_FZHX0002-C/2017 | A / H5N6 | Asia / China / Fujian Province | |
| EPI_ISL_707644 | A/goose/Fujian/10.26_FZHX0002-O/2017 | A / H5N6 | Asia / China / Fujian Province | |
| EPI_ISL_707645 | A/goose/Fujian/10.26_FZHX0004-C/2017 | A / H5N6 | Asia / China / Fujian Province | |
| EPI_ISL_707646 | A/goose/Fujian/10.26_FZHX0004-O/2017 | A / H5N6 | Asia / China / Fujian Province | |
| EPI_ISL_707647 | A/goose/Fujian/10.26_FZHX0006-C/2017 | A / H5N6 | Asia / China / Fujian Province | |
| EPI_ISL_707648 | A/goose/Fujian/10.26_FZHX0006-O/2017 | A / H5N6 | Asia / China / Fujian Province | |
| EPI_ISL_707649 | A/goose/Fujian/10.26_FZHX0007-O/2017 | A / H5N6 | Asia / China / Fujian Province | |
| EPI_ISL_707650 | A/goose/Fujian/10.26_FZHX0008-C/2017 | A / H5N6 | Asia / China / Fujian Province | |
| EPI_ISL_707651 | A/goose/Fujian/10.26_FZHX0014-O/2017 | A / H5N6 | Asia / China / Fujian Province | |
| EPI_ISL_707652 | A/goose/Fujian/11.3_FZHX1102-C/2016 | A / H5N6 | Asia / China / Fujian Province | |
| EPI_ISL_707653 | A/goose/Fujian/3.15_FZHX0008-C/2018 | A / H5N6 | Asia / China / Fujian Province | |
| EPI_ISL_707658 | A/goose/Guangdong/7.20_DGCP010-C/2017 | A / H5N6 | Asia / China / Guangdong Province | |
| EPI_ISL_707659 | A/goose/Henan/01.01_LYGL001-O/2017 | A / H5N6 | Asia / China / Henan Province | |
| EPI_ISL_707660 | A/goose/Xinjiang/11.29_WLMQXL001-O/2017 | A / H5N6 | Asia / China / Xinjiang Uyghur Autonomous Region | |
| EPI_ISL_707661 | A/goose/Xinjiang/12.18_WLMQXL001-C/2016 | A / H5N6 | Asia / China / Xinjiang Uyghur Autonomous Region | |
| EPI_ISL_707662 | A/goose/Xinjiang/12.18_WLMQXL002-O/2016 | A / H5N6 | Asia / China / Xinjiang Uyghur Autonomous Region | |
| EPI_ISL_707663 | A/goose/Xinjiang/12.18_WLMQXL003-C/2016 | A / H5N8 | Asia / China / Xinjiang Uyghur Autonomous Region | |
| EPI_ISL_707664 | A/goose/Xinjiang/12.18_WLMQXL003-O/2016 | A / H5N8 | Asia / China / Xinjiang Uyghur Autonomous Region | |
| EPI_ISL_707665 | A/goose/Xinjiang/12.18_WLMQXL004-C/2016 | A / H5N8 | Asia / China / Xinjiang Uyghur Autonomous Region | |
| EPI_ISL_707666 | A/goose/Xinjiang/12.18_WLMQXL004-O/2016 | A / H5N8 | Asia / China / Xinjiang Uyghur Autonomous Region | |
| EPI_ISL_707676 | A/pigeon/Fujian/1.17_FZHX0104-O/2017 | A / H5N6 | Asia / China / Fujian Province | |
| EPI_ISL_708701 | A/chicken/Kagawa/I1C/2020 | A / H5N8 | Asia / Japan / Kagawa | Takehiko Saito (National Institute of Animal Health) |
| EPI_ISL_708702 | A/chicken/Kagawa/I1T/2020 | A / H5N8 | Asia / Japan / Kagawa | Takehiko Saito (National Institute of Animal Health) |
| EPI_ISL_708703 | A/chicken/Kagawa/I2C/2020 | A / H5N8 | Asia / Japan / Kagawa | Takehiko Saito (National Institute of Animal Health) |
| EPI_ISL_708704 | A/chicken/Kagawa/I2T/2020 | A / H5N8 | Asia / Japan / Kagawa | Takehiko Saito (National Institute of Animal Health) |
| EPI_ISL_708705 | A/chicken/Kagawa/I4C/2020 | A / H5N8 | Asia / Japan / Kagawa | Takehiko Saito (National Institute of Animal Health) |
| EPI_ISL_708706 | A/chicken/Kagawa/I4T/2020 | A / H5N8 | Asia / Japan / Kagawa | Takehiko Saito (National Institute of Animal Health) |
| EPI_ISL_708707 | A/chicken/Kagawa/I5C/2020 | A / H5N8 | Asia / Japan / Kagawa | Takehiko Saito (National Institute of Animal Health) |
| EPI_ISL_708708 | A/chicken/Kagawa/I5T/2020 | A / H5N8 | Asia / Japan / Kagawa | Takehiko Saito (National Institute of Animal Health) |
| EPI_ISL_708709 | A/chicken/Kagawa/I6C/2020 | A / H5N8 | Asia / Japan / Kagawa | Takehiko Saito (National Institute of Animal Health) |
| EPI_ISL_708710 | A/chicken/Kagawa/I6T/2020 | A / H5N8 | Asia / Japan / Kagawa | Takehiko Saito (National Institute of Animal Health) |
| EPI_ISL_708711 | A/chicken/Kagawa/J3C/2020 | A / H5N8 | Asia / Japan / Kagawa | Takehiko Saito (National Institute of Animal Health) |
| EPI_ISL_708712 | A/chicken/Kagawa/J3T/2020 | A / H5N8 | Asia / Japan / Kagawa | Takehiko Saito (National Institute of Animal Health) |
| EPI_ISL_708713 | A/chicken/Kagawa/J4C/2020 | A / H5N8 | Asia / Japan / Kagawa | Takehiko Saito (National Institute of Animal Health) |
| EPI_ISL_708714 | A/chicken/Kagawa/J4T/2020 | A / H5N8 | Asia / Japan / Kagawa | Takehiko Saito (National Institute of Animal Health) |
| EPI_ISL_708715 | A/chicken/Kagawa/J7C/2020 | A / H5N8 | Asia / Japan / Kagawa | Takehiko Saito (National Institute of Animal Health) |
| EPI_ISL_708716 | A/chicken/Kagawa/J7T/2020 | A / H5N8 | Asia / Japan / Kagawa | Takehiko Saito (National Institute of Animal Health) |
| EPI_ISL_708717 | A/chicken/Kagawa/J8C/2020 | A / H5N8 | Asia / Japan / Kagawa | Takehiko Saito (National Institute of Animal Health) |
| EPI_ISL_708718 | A/chicken/Kagawa/J8T/2020 | A / H5N8 | Asia / Japan / Kagawa | Takehiko Saito (National Institute of Animal Health) |
| EPI_ISL_708719 | A/chicken/Kagawa/J9C/2020 | A / H5N8 | Asia / Japan / Kagawa | Takehiko Saito (National Institute of Animal Health) |
| EPI_ISL_708720 | A/chicken/Kagawa/J9T/2020 | A / H5N8 | Asia / Japan / Kagawa | Takehiko Saito (National Institute of Animal Health) |
| EPI_ISL_710434 | A/chicken/Hiroshima/1T/2020 | A / H5N8 | Asia / Japan / Hiroshima | Takehiko Saito (National Institute of Animal Health) |
| EPI_ISL_710435 | A/chicken/Hiroshima/2T/2020 | A / H5N8 | Asia / Japan / Hiroshima | Takehiko Saito (National Institute of Animal Health) |
| EPI_ISL_710436 | A/chicken/Hiroshima/3T/2020 | A / H5N8 | Asia / Japan / Hiroshima | Takehiko Saito (National Institute of Animal Health) |
| EPI_ISL_710437 | A/chicken/Hiroshima/4T/2020 | A / H5N8 | Asia / Japan / Hiroshima | Takehiko Saito (National Institute of Animal Health) |
| EPI_ISL_710438 | A/chicken/Miyazaki/1T/2020 | A / H5N8 | Asia / Japan / Miyazaki | Takehiko Saito (National Institute of Animal Health) |
| EPI_ISL_710439 | A/chicken/Miyazaki/2T/2020 | A / H5N8 | Asia / Japan / Miyazaki | Takehiko Saito (National Institute of Animal Health) |
| EPI_ISL_710440 | A/chicken/Miyazaki/4T/2020 | A / H5N8 | Asia / Japan / Miyazaki | Takehiko Saito (National Institute of Animal Health) |
| EPI_ISL_710441 | A/chicken/Miyazaki/6T/2020 | A / H5N8 | Asia / Japan / Miyazaki | Takehiko Saito (National Institute of Animal Health) |
| EPI_ISL_710442 | A/chicken/Miyazaki/B10T/2020 | A / H5N8 | Asia / Japan / Miyazaki | Takehiko Saito (National Institute of Animal Health) |
| EPI_ISL_710443 | A/chicken/Miyazaki/B1T/2020 | A / H5N8 | Asia / Japan / Miyazaki | Takehiko Saito (National Institute of Animal Health) |
| EPI_ISL_710444 | A/chicken/Miyazaki/B2T/2020 | A / H5N8 | Asia / Japan / Miyazaki | Takehiko Saito (National Institute of Animal Health) |

| | | | | |
|---|---|---|---|---|
| EPI_ISL_710445 | A/chicken/Miyazaki/B6T/2020 | A / H5N8 | Asia / Japan / Miyazaki | Takehiko Saito (National Institute of Animal Health) |
| EPI_ISL_710446 | A/chicken/Miyazaki/C4T/2020 | A / H5N8 | Asia / Japan / Miyazaki | Takehiko Saito (National Institute of Animal Health) |
| EPI_ISL_710447 | A/chicken/Miyazaki/C6T/2020 | A / H5N8 | Asia / Japan / Miyazaki | Takehiko Saito (National Institute of Animal Health) |
| EPI_ISL_710448 | A/chicken/Miyazaki/C7T/2020 | A / H5N8 | Asia / Japan / Miyazaki | Takehiko Saito (National Institute of Animal Health) |
| EPI_ISL_710449 | A/chicken/Miyazaki/C8T/2020 | A / H5N8 | Asia / Japan / Miyazaki | Takehiko Saito (National Institute of Animal Health) |
| EPI_ISL_710450 | A/chicken/Nara/12T/2020 | A / H5N8 | Asia / Japan / Nara | Takehiko Saito (National Institute of Animal Health) |
| EPI_ISL_710451 | A/chicken/Nara/1C/2020 | A / H5N8 | Asia / Japan / Nara | Takehiko Saito (National Institute of Animal Health) |
| EPI_ISL_710452 | A/chicken/Nara/5T/2020 | A / H5N8 | Asia / Japan / Nara | Takehiko Saito (National Institute of Animal Health) |
| EPI_ISL_710453 | A/chicken/Nara/9T/2020 | A / H5N8 | Asia / Japan / Nara | Takehiko Saito (National Institute of Animal Health) |
| EPI_ISL_710504 | A/turkey/England/037784/2020 | A / H5N8 | Europe / United Kingdom / North Yorkshire | Alex Byrne (Animal and Plant Health Agency (APHA) / Virology Department) |
| EPI_ISL_710505 | A/turkey/England/038115/2020 | A / H5N8 | Europe / United Kingdom / North Yorkshire | Alex Byrne (Animal and Plant Health Agency (APHA) / Virology Department) |
| EPI_ISL_710506 | A/Canada_goose/England/032697/2020 | A / H5N8 | Europe / United Kingdom / Gloucestershire | Alex Byrne (Animal and Plant Health Agency (APHA) / Virology Department) |
| EPI_ISL_710507 | A/Greylag_goose/England/032698/2020 | A / H5N8 | Europe / United Kingdom / Gloucestershire | Alex Byrne (Animal and Plant Health Agency (APHA) / Virology Department) |
| EPI_ISL_710508 | A/Greylag_goose/England/033100/2020 | A / H5N8 | Europe / United Kingdom / Gloucestershire | Alex Byrne (Animal and Plant Health Agency (APHA) / Virology Department) |
| EPI_ISL_710509 | A/chicken/England/033708/2020 | A / H5N8 | Europe / United Kingdom / Hertfordshire | Alex Byrne (Animal and Plant Health Agency (APHA) / Virology Department) |
| EPI_ISL_710511 | A/chicken/England/037052/2020 | A / H5N8 | Europe / United Kingdom / Leicestershire | Alex Byrne (Animal and Plant Health Agency (APHA) / Virology Department) |
| EPI_ISL_710512 | A/whistling_duck/England/035643/2020 | A / H5N8 | Europe / United Kingdom / Gloucestershire | Alex Byrne (Animal and Plant Health Agency (APHA) / Virology Department) |
| EPI_ISL_710533 | A/chicken/Netherlands/20019237-001005/2020 | A / H5N8 | Europe / Netherlands / South Holland / Gemeente Brielle | Rene Heutink (Wageningen Bioveterinary Research) |
| EPI_ISL_710538 | A/chicken/Netherlands/20019226-001/2020 | A / H5N8 | Europe / Netherlands / Provincie Noord-Holland / Edam-Volendam | Rene Heutink (Wageningen Bioveterinary Research) |
| EPI_ISL_710539 | A/chicken/Netherlands/20019422-001005/2020 | A / H5N8 | Europe / Netherlands / Provincie Friesland / Sint Annaparochie | Rene Heutink (Wageningen Bioveterinary Research) |
| EPI_ISL_711055 | A/chicken/Netherlands/20019879-001005/2020 | A / H5N1 | Europe / Netherlands / Provincie Friesland / Achtkarspelen | Rene Heutink (Wageningen Bioveterinary Research) |
| EPI_ISL_711056 | A/mute swan/Netherlands/20019252-002/2020 | A / H5N8 | Europe / Netherlands / South Holland / Leidschendam-Voorburg | Rene Heutink (Wageningen Bioveterinary Research) |
| EPI_ISL_711058 | A/mute swan/Netherlands/20019255-002/2020 | A / H5N8 | Europe / Netherlands / Provincie Utrecht / Gemeente Woerden | Rene Heutink (Wageningen Bioveterinary Research) |
| EPI_ISL_711059 | A/greylag goose/Netherlands/20019685-002/2020 | A / H5N1 | Europe / Netherlands / Provincie Utrecht / Gemeente Woerden | Rene Heutink (Wageningen Bioveterinary Research) |
| EPI_ISL_718106 | A/chicken/Miyazaki/D5C/2020 | A / H5N8 | Asia / Japan / Miyazaki | Takehiko Saito (National Institute of Animal Health) |
| EPI_ISL_718107 | A/chicken/Miyazaki/D5T/2020 | A / H5N8 | Asia / Japan / Miyazaki | Takehiko Saito (National Institute of Animal Health) |
| EPI_ISL_718108 | A/chicken/Miyazaki/E1T/2020 | A / H5N8 | Asia / Japan / Miyazaki | Takehiko Saito (National Institute of Animal Health) |
| EPI_ISL_718109 | A/chicken/Miyazaki/E3T/2020 | A / H5N8 | Asia / Japan / Miyazaki | Takehiko Saito (National Institute of Animal Health) |
| EPI_ISL_718110 | A/chicken/Miyazaki/E6T/2020 | A / H5N8 | Asia / Japan / Miyazaki | Takehiko Saito (National Institute of Animal Health) |
| EPI_ISL_718111 | A/chicken/Miyazaki/E9T/2020 | A / H5N8 | Asia / Japan / Miyazaki | Takehiko Saito (National Institute of Animal Health) |
| EPI_ISL_718112 | A/chicken/Oita/27T/2020 | A / H5N8 | Asia / Japan / Oita | Takehiko Saito (National Institute of Animal Health) |
| EPI_ISL_718113 | A/chicken/Oita/2T/2020 | A / H5N8 | Asia / Japan / Oita | Takehiko Saito (National Institute of Animal Health) |
| EPI_ISL_718114 | A/chicken/Oita/3T/2020 | A / H5N8 | Asia / Japan / Oita | Takehiko Saito (National Institute of Animal Health) |
| EPI_ISL_718115 | A/chicken/Oita/5T/2020 | A / H5N8 | Asia / Japan / Oita | Takehiko Saito (National Institute of Animal Health) |
| EPI_ISL_718116 | A/chicken/Okayama/1T/2020 | A / H5N8 | Asia / Japan / Okayama | Takehiko Saito (National Institute of Animal Health) |
| EPI_ISL_718117 | A/chicken/Okayama/2T/2020 | A / H5N8 | Asia / Japan / Okayama | Takehiko Saito (National Institute of Animal Health) |
| EPI_ISL_718118 | A/chicken/Okayama/6T/2020 | A / H5N8 | Asia / Japan / Okayama | Takehiko Saito (National Institute of Animal Health) |
| EPI_ISL_718119 | A/chicken/Okayama/7T/2020 | A / H5N8 | Asia / Japan / Okayama | Takehiko Saito (National Institute of Animal Health) |
| EPI_ISL_718120 | A/chicken/Wakayama/1T/2020 | A / H5N8 | Asia / Japan / Wakayama | Takehiko Saito (National Institute of Animal Health) |
| EPI_ISL_718121 | A/chicken/Wakayama/2T/2020 | A / H5N8 | Asia / Japan / Wakayama | Takehiko Saito (National Institute of Animal Health) |
| EPI_ISL_718122 | A/chicken/Wakayama/3T/2020 | A / H5N8 | Asia / Japan / Wakayama | Takehiko Saito (National Institute of Animal Health) |
| EPI_ISL_718123 | A/chicken/Wakayama/4T/2020 | A / H5N8 | Asia / Japan / Wakayama | Takehiko Saito (National Institute of Animal Health) |
| EPI_ISL_718223 | A/barnacle goose/Sweden/SVA201125SZ0472/KN003488/202( | A / H5N8 | Europe / Sweden / Gotlands Lan / Gotlands Kommun | Siamak Zohari (National Veterinary Institute) |
| EPI_ISL_718225 | A/Jiangsu/1/2020 ?H5N6? | A / H5N6 | Asia / China / Jiangsu Provice | Xian Qi (Jiangsu Provincial Center for Disease Control & Prevention) |
| EPI_ISL_718266 | A/Jiangsu/1/2020(H5N6) | A / H5N6 | Asia / China / Jiangsu Provice / Nanjing | Min He (Nanjing Center for Disease Control and Prevention / Microbiological Testing) |
| EPI_ISL_7192735 | A/Duck/Sichuan/21022-4/2021(H5N8) | A / H5N8 | Asia / China / Sichuan Province | Jiahao Zhang (South China Agricultural University / College of Veterinary Medicine) |
| EPI_ISL_722176 | A/barnacle goose/Sweden/SVA201201SZ0353/KN003624/202( | A / H5N5 | Europe / Sweden / Skane Lan / Staffanstorps Kommun | Siamak Zohari (National Veterinary Institute) |
| EPI_ISL_7224437 | A/chicken/Czech Republic/22224-2T/2021 | A / H5N1 | Europe / Czech Republic / Jihocesky Kraj / Okres Jindrichuv Hradec / Straz nad Nezarkou, Dolni Ll | Alexander Nagy (State Veterinary Institute Prague) |
| EPI_ISL_7224454 | A/chicken/Czech Republic/22224-3K/2021 | A / H5N1 | Europe / Czech Republic / Jihocesky Kraj / Okres Jindrichuv Hradec / Straz nad Nezarkou, Dolni Ll | Alexander Nagy (State Veterinary Institute Prague) |
| EPI_ISL_7224463 | A/chicken/Czech Republic/22224-3T/2021 | A / H5N1 | Europe / Czech Republic / Jihocesky Kraj / Okres Jindrichuv Hradec / Straz nad Nezarkou, Dolni Ll | Alexander Nagy (State Veterinary Institute Prague) |
| EPI_ISL_7224472 | A/chicken/Czech Republic/22224-4T/2021 | A / H5N1 | Europe / Czech Republic / Jihocesky Kraj / Okres Jindrichuv Hradec / Straz nad Nezarkou, Dolni Ll | Alexander Nagy (State Veterinary Institute Prague) |
| EPI_ISL_7267243 | A/greylag goose/Netherlands/21038570-002/2021 | A / H5N1 | Europe / Netherlands / Provincie Friesland | Rene Heutink (Wageningen Bioveterinary Research) |
| EPI_ISL_7267244 | A/common buzzard/Netherlands/21038793-001/2021 | A / H5N1 | Europe / Netherlands / Provincie Friesland | Rene Heutink (Wageningen Bioveterinary Research) |
| EPI_ISL_7267245 | A/mute swan/Netherlands/21038797-001/2021 | A / H5N1 | Europe / Netherlands / South Holland | Rene Heutink (Wageningen Bioveterinary Research) |
| EPI_ISL_7267246 | A/mute swan/Netherlands/21039192-001/2021 | A / H5N1 | Europe / Netherlands / Provincie Friesland | Rene Heutink (Wageningen Bioveterinary Research) |
| EPI_ISL_7267247 | A/mute swan/Netherlands/21039202-001/2021 | A / H5N1 | Europe / Netherlands / South Holland | Rene Heutink (Wageningen Bioveterinary Research) |
| EPI_ISL_7267248 | A/goose/Netherlands/21038940-002/2021 | A / H5N1 | Europe / Netherlands / Provincie Flevoland | Rene Heutink (Wageningen Bioveterinary Research) |
| EPI_ISL_7267249 | A/grey heron/Netherlands/21038941-001/2021 | A / H5N1 | Europe / Netherlands / Provincie Gelderland | Rene Heutink (Wageningen Bioveterinary Research) |
| EPI_ISL_7267250 | A/greylag goose/Netherlands/21038942-001/2021 | A / H5N1 | Europe / Netherlands / Provincie Friesland | Rene Heutink (Wageningen Bioveterinary Research) |
| EPI_ISL_7267251 | A/greater canada goose /Netherlands/21038971-001/2021 | A / H5N1 | Europe / Netherlands / South Holland | Rene Heutink (Wageningen Bioveterinary Research) |
| EPI_ISL_7267252 | A/goose/Netherlands/21039029-001/2021 | A / H5N1 | Europe / Netherlands / Provincie Noord-Holland | Rene Heutink (Wageningen Bioveterinary Research) |
| EPI_ISL_7267253 | A/gull/Netherlands/21039180-002/2021 | A / H5N1 | Europe / Netherlands / Provincie Noord-Holland | Rene Heutink (Wageningen Bioveterinary Research) |
| EPI_ISL_7267254 | A/greylag goose/Netherlands/21039376-001/2021 | A / H5N1 | Europe / Netherlands / Provincie Utrecht | Rene Heutink (Wageningen Bioveterinary Research) |
| EPI_ISL_732927 | A/chicken/Kagawa/K2T/2020 | A / H5N8 | Asia / Japan / Kagawa | Takehiko Saito (National Institute of Animal Health) |
| EPI_ISL_732928 | A/chicken/Kagawa/K3T/2020 | A / H5N8 | Asia / Japan / Kagawa | Takehiko Saito (National Institute of Animal Health) |
| EPI_ISL_732929 | A/chicken/Kagawa/K4T/2020 | A / H5N8 | Asia / Japan / Kagawa | Takehiko Saito (National Institute of Animal Health) |
| EPI_ISL_732930 | A/chicken/Kagawa/K6T/2020 | A / H5N8 | Asia / Japan / Kagawa | Takehiko Saito (National Institute of Animal Health) |
| EPI_ISL_732931 | A/chicken/Kochi/4C/2020 | A / H5N8 | Asia / Japan / Kochi | Takehiko Saito (National Institute of Animal Health) |
| EPI_ISL_732932 | A/chicken/Kochi/5C/2020 | A / H5N8 | Asia / Japan / Kochi | Takehiko Saito (National Institute of Animal Health) |
| EPI_ISL_732933 | A/chicken/Kochi/6T/2020 | A / H5N8 | Asia / Japan / Kochi | Takehiko Saito (National Institute of Animal Health) |
| EPI_ISL_732934 | A/chicken/Kochi/7T/2020 | A / H5N8 | Asia / Japan / Kochi | Takehiko Saito (National Institute of Animal Health) |
| EPI_ISL_732935 | A/chicken/Miyazaki/F1T/2020 | A / H5N8 | Asia / Japan / Miyazaki | Takehiko Saito (National Institute of Animal Health) |

| | | | |
|---|---|---|---|
| EPI_ISL_732936 | A/chicken/Miyazaki/F4T/2020 | A / H5N8 | Asia / Japan / Miyazaki | Takehiko Saito (National Institute of Animal Health) |
| EPI_ISL_732937 | A/chicken/Miyazaki/F5T/2020 | A / H5N8 | Asia / Japan / Miyazaki | Takehiko Saito (National Institute of Animal Health) |
| EPI_ISL_732938 | A/chicken/Miyazaki/F7T/2020 | A / H5N8 | Asia / Japan / Miyazaki | Takehiko Saito (National Institute of Animal Health) |
| EPI_ISL_732939 | A/chicken/Miyazaki/G1T/2020 | A / H5N8 | Asia / Japan / Miyazaki | Takehiko Saito (National Institute of Animal Health) |
| EPI_ISL_732940 | A/chicken/Miyazaki/G3T/2020 | A / H5N8 | Asia / Japan / Miyazaki | Takehiko Saito (National Institute of Animal Health) |
| EPI_ISL_732941 | A/chicken/Miyazaki/G5T/2020 | A / H5N8 | Asia / Japan / Miyazaki | Takehiko Saito (National Institute of Animal Health) |
| EPI_ISL_732942 | A/chicken/Miyazaki/G9T/2020 | A / H5N8 | Asia / Japan / Miyazaki | Takehiko Saito (National Institute of Animal Health) |
| EPI_ISL_732943 | A/chicken/Shiga/4T/2020 | A / H5N8 | Asia / Japan / Shiga | Takehiko Saito (National Institute of Animal Health) |
| EPI_ISL_732944 | A/chicken/Shiga/5T/2020 | A / H5N8 | Asia / Japan / Shiga | Takehiko Saito (National Institute of Animal Health) |
| EPI_ISL_732945 | A/chicken/Shiga/6T/2020 | A / H5N8 | Asia / Japan / Shiga | Takehiko Saito (National Institute of Animal Health) |
| EPI_ISL_732946 | A/chicken/Shiga/7T/2020 | A / H5N8 | Asia / Japan / Shiga | Takehiko Saito (National Institute of Animal Health) |
| EPI_ISL_7349517 | A/Duck/Sichuan/21022-1/2021(H5N8) | A / H5N8 | Asia / China / Sichuan Province | Jiahao Zhang (South China Agricultural University / College of Veterinary Medicine) |
| EPI_ISL_7357590 | A/mute swan/Croatia/101/2021 | A / H5N1 | Europe / Croatia / Sisacko-Moslavacka Zupanija / Cigoc (45° 25' 16,9" N; 16° 37' 1,2" E) | Vladimir Savi? (Croatian Veterinary Institute / Poultry Centre) |
| EPI_ISL_7379463 | A/Goose/Henan/21028/2021(H5N8) | A / H5N8 | Asia / China / Henan Province | Jiahao Zhang (South China Agricultural University / College of Veterinary Medicine) |
| EPI_ISL_7379464 | A/Duck/Sichuan/21044-2/2021(H5N8) | A / H5N8 | Asia / China / Sichuan Province | Jiahao Zhang (South China Agricultural University / College of Veterinary Medicine) |
| EPI_ISL_7379465 | A/Goose/Henan/21056-2/2021(H5N8) | A / H5N8 | Asia / China / Henan Province | Jiahao Zhang (South China Agricultural University / College of Veterinary Medicine) |
| EPI_ISL_7379509 | A/Duck/Guangdong/21057/2021(H5N8) | A / H5N8 | Asia / China / Guangdong Province | Jiahao Zhang (South China Agricultural University / College of Veterinary Medicine) |
| EPI_ISL_7380253 | A/Goose/Jiangsu/21153-2/2021(H5N8) | A / H5N8 | Asia / China / Jiangsu Provice | Jiahao Zhang (South China Agricultural University / College of Veterinary Medicine) |
| EPI_ISL_738049 | A/chicken/Kagawa/L4T/2020 | A / H5N8 | Asia / Japan / Kagawa | Takehiko Saito (National Institute of Animal Health) |
| EPI_ISL_738050 | A/chicken/Kagawa/L6T/2020 | A / H5N8 | Asia / Japan / Kagawa | Takehiko Saito (National Institute of Animal Health) |
| EPI_ISL_7380500 | A/Goose/Shandong/21153-3/2021(H5N8) | A / H5N8 | Asia / China / Shandong Province | Jiahao Zhang (South China Agricultural University / College of Veterinary Medicine) |
| EPI_ISL_738051 | A/chicken/Kagawa/L7T/2020 | A / H5N8 | Asia / Japan / Kagawa | Takehiko Saito (National Institute of Animal Health) |
| EPI_ISL_738052 | A/chicken/Kagawa/L9T/2020 | A / H5N8 | Asia / Japan / Kagawa | Takehiko Saito (National Institute of Animal Health) |
| EPI_ISL_738053 | A/chicken/Miyazaki/H1T/2020 | A / H5N8 | Asia / Japan / Miyazaki | Takehiko Saito (National Institute of Animal Health) |
| EPI_ISL_738054 | A/chicken/Miyazaki/H3T/2020 | A / H5N8 | Asia / Japan / Miyazaki | Takehiko Saito (National Institute of Animal Health) |
| EPI_ISL_738055 | A/chicken/Miyazaki/H6T/2020 | A / H5N8 | Asia / Japan / Miyazaki | Takehiko Saito (National Institute of Animal Health) |
| EPI_ISL_738056 | A/chicken/Miyazaki/H9T/2020 | A / H5N8 | Asia / Japan / Miyazaki | Takehiko Saito (National Institute of Animal Health) |
| EPI_ISL_738057 | A/chicken/Tokushima/1T/20200 | A / H5N8 | Asia / Japan / Tokushima | Takehiko Saito (National Institute of Animal Health) |
| EPI_ISL_738058 | A/chicken/Tokushima/2T/2020 | A / H5N8 | Asia / Japan / Tokushima | Takehiko Saito (National Institute of Animal Health) |
| EPI_ISL_738059 | A/chicken/Tokushima/3T/2020 | A / H5N8 | Asia / Japan / Tokushima | Takehiko Saito (National Institute of Animal Health) |
| EPI_ISL_738060 | A/chicken/Tokushima/4T/2020 | A / H5N8 | Asia / Japan / Tokushima | Takehiko Saito (National Institute of Animal Health) |
| EPI_ISL_7380623 | A/Duck/Shandong/21232-5/2021(H5N8) | A / H5N8 | Asia / China / Shandong Province | Jiahao Zhang (South China Agricultural University / College of Veterinary Medicine) |
| EPI_ISL_7380813 | A/Duck/Guangdong/21316/2021(H5N8) | A / H5N8 | Asia / China / Guangdong Province | Jiahao Zhang (South China Agricultural University / College of Veterinary Medicine) |
| EPI_ISL_7381026 | A/Chicken/Liaoning/21346-2/2021(H5N8) | A / H5N8 | Asia / China / Liaoning Province | Jiahao Zhang (South China Agricultural University / College of Veterinary Medicine) |
| EPI_ISL_7381065 | A/Goose/Shandong/21369-4/2021(H5N8) | A / H5N8 | Asia / China / Shandong Province | Jiahao Zhang (South China Agricultural University / College of Veterinary Medicine) |
| EPI_ISL_7381103 | A/Goose/Shandong/21369-5/2021(H5N8) | A / H5N8 | Asia / China / Shandong Province | Jiahao Zhang (South China Agricultural University / College of Veterinary Medicine) |
| EPI_ISL_7381115 | A/Goose/Liaoning/21640/2021(H5N8) | A / H5N8 | Asia / China / Liaoning Province | Jiahao Zhang (South China Agricultural University / College of Veterinary Medicine) |
| EPI_ISL_7381183 | A/Duck/Shandong/21644-4/2021(H5N8) | A / H5N8 | Asia / China / Shandong Province | Jiahao Zhang (South China Agricultural University / College of Veterinary Medicine) |
| EPI_ISL_7381397 | A/Duck/Shandong/21644-7/2021(H5N8) | A / H5N8 | Asia / China / Shandong Province | Jiahao Zhang (South China Agricultural University / College of Veterinary Medicine) |
| EPI_ISL_7381423 | A/Duck/Shandong/21931-8/2021(H5N8) | A / H5N8 | Asia / China / Shandong Province | Jiahao Zhang (South China Agricultural University / College of Veterinary Medicine) |
| EPI_ISL_7381438 | A/Duck/Guangdong/21964/2021(H5N8) | A / H5N8 | Asia / China / Guangdong Province | Jiahao Zhang (South China Agricultural University / College of Veterinary Medicine) |
| EPI_ISL_7381448 | A/Chicken/Guangxi/21989-3/2021(H5N8) | A / H5N8 | Asia / China / Guangxi Zhuang Autonomous Region | Jiahao Zhang (South China Agricultural University / College of Veterinary Medicine) |
| EPI_ISL_7381695 | A/Goose/Guangdong/211030-1/2021(H5N8) | A / H5N8 | Asia / China / Guangdong Province | Jiahao Zhang (South China Agricultural University / College of Veterinary Medicine) |
| EPI_ISL_739684 | A/goose/Russia_Novosibirsk region/1-12/2020 | A / H5N8 | Europe / Russian Federation / Novosibirsk region | Ivan Sobolev (Research Institute of Experimental and Clinical Medicine) |
| EPI_ISL_739685 | A/goose/Russia_Omsk region/55-1/2020 | A / H5N8 | Europe / Russian Federation / Omsk region | Ivan Sobolev (Research Institute of Experimental and Clinical Medicine) |
| EPI_ISL_739686 | A/chicken/Kazakhstan/Kn-3/2020 | A / H5N8 | Asia / Kazakhstan / North Kazakhstan | Ivan Sobolev (Research Institute of Experimental and Clinical Medicine) |
| EPI_ISL_739687 | A/chicken/Kazakhstan/Kn-6/2020 | A / H5N8 | Asia / Kazakhstan / North Kazakhstan | Ivan Sobolev (Research Institute of Experimental and Clinical Medicine) |
| EPI_ISL_739688 | A/chicken/Russia_Novosibirsk region/1910-1/2020 | A / H5N8 | Europe / Russian Federation / Novosibirsk region | Ivan Sobolev (Research Institute of Experimental and Clinical Medicine) |
| EPI_ISL_739689 | A/chicken/Russia_Novosibirsk region/1910-2/2020 | A / H5N8 | Europe / Russian Federation / Novosibirsk region | Ivan Sobolev (Research Institute of Experimental and Clinical Medicine) |
| EPI_ISL_739690 | A/chicken/Russia_Novosibirsk region/3-1/2020 | A / H5N8 | Europe / Russian Federation / Novosibirsk region | Ivan Sobolev (Research Institute of Experimental and Clinical Medicine) |
| EPI_ISL_739691 | A/chicken/Russia_Novosibirsk region/3-15/2020 | A / H5N8 | Europe / Russian Federation / Novosibirsk region | Ivan Sobolev (Research Institute of Experimental and Clinical Medicine) |
| EPI_ISL_739692 | A/chicken/Russia_Novosibirsk region/3-29/2020 | A / H5N8 | Europe / Russian Federation / Novosibirsk Oblast / Novosibirsk region | Ivan Sobolev (Research Institute of Experimental and Clinical Medicine) |
| EPI_ISL_7452805 | A/Chicken/Sweden/SVA211130SZ0427/FB290424-IP-1/M-2021 | A / H5N1 | Europe / Sweden / Skane Lan / Skurups Kommun | Siamak Zohari (National Veterinary Institute) |
| EPI_ISL_7457654 | A/Chicken/Guangdong/211106-3/2021(H5N6) | A / H5N6 | Asia / China / Guangdong Province | Jiahao Zhang (South China Agricultural University / College of Veterinary Medicine) |
| EPI_ISL_7458702 | A/Duck/Shandong/21644-1/2021(H5N8) | A / H5N8 | Asia / China / Shandong Province | Jiahao Zhang (South China Agricultural University / College of Veterinary Medicine) |
| EPI_ISL_7570462 | A/mute swan/Croatia/104/2021 | A / H5N1 | Europe / Croatia / Medimurska Zupanija / Gornji Kursanec (46° 19' 27.3" N; 16° 22' 45" E) | Vladimir Savi? (Croatian Veterinary Institute / Poultry Centre) |
| EPI_ISL_7570634 | A/goose/Croatia/107/2021 | A / H5N1 | Europe / Croatia / Sisacko-Moslavacka Zupanija / Staro Pracno | Vladimir Savi? (Croatian Veterinary Institute / Poultry Centre) |
| EPI_ISL_7585764 | A/Aalopochen_Aegyptiaca/Belgium/2928_002/2021 | A / H5N8 | Europe / Belgium / Provincie Limburg / Genk | Steven Van Borm (Sciensano, Department of Animal Infectious Diseases / Animal Infectious Diseases) |
| EPI_ISL_7589566 | A/Gallus_gallus/Belgium/5107_002/2021 | A / H5N8 | Europe / Belgium / Province de Hainaut / Silly | Steven Van Borm (Sciensano, Department of Animal Infectious Diseases / Animal Infectious Diseases) |
| EPI_ISL_7590312 | A/Branta_canadensis/Belgium/500/2021 | A / H5N8 | Europe / Belgium / Province de Liege / Amay | Steven Van Borm (Sciensano, Department of Animal Infectious Diseases / Animal Infectious Diseases) |
| EPI_ISL_7591058 | A/Anas_platyrhynchos domestica/Belgium/5517/2021 | A / H5N8 | Europe / Belgium / Provincie West-Vlaanderen / Beveren | Steven Van Borm (Sciensano, Department of Animal Infectious Diseases / Animal Infectious Diseases) |
| EPI_ISL_7596571 | A/Pica_pica/Belgium/12100_005/2020 | A / H5N8 | Europe / Belgium / Provincie West-Vlaanderen / Oudenburg | Steven Van Borm (Sciensano, Department of Animal Infectious Diseases / Animal Infectious Diseases) |
| EPI_ISL_7597827 | A/Podiceps_cristatus/Belgium/12659_0015/2020 | A / H5N8 | Europe / Belgium / Provincie Oost-Vlaanderen / Gent (Blaarmeersen) | Steven Van Borm (Sciensano, Department of Animal Infectious Diseases / Animal Infectious Diseases) |
| EPI_ISL_759850 | A/chicken/Chiba/1T/2020 | A / H5N8 | Asia / Japan / Chiba | Takehiko Saito (National Institute of Animal Health) |
| EPI_ISL_759851 | A/chicken/Chiba/2T/2020 | A / H5N8 | Asia / Japan / Chiba | Takehiko Saito (National Institute of Animal Health) |
| EPI_ISL_759852 | A/chicken/Chiba/3T/2020 | A / H5N8 | Asia / Japan / Chiba | Takehiko Saito (National Institute of Animal Health) |
| EPI_ISL_759853 | A/chicken/Chiba/4T/2020 | A / H5N8 | Asia / Japan / Chiba | Takehiko Saito (National Institute of Animal Health) |
| EPI_ISL_759854 | A/chicken/Kagawa/M10T/2020 | A / H5N8 | Asia / Japan / Kagawa | Takehiko Saito (National Institute of Animal Health) |
| EPI_ISL_759855 | A/chicken/Kagawa/M11T/2020 | A / H5N8 | Asia / Japan / Kagawa | Takehiko Saito (National Institute of Animal Health) |
| EPI_ISL_759856 | A/chicken/Kagawa/M12T/2020 | A / H5N8 | Asia / Japan / Kagawa | Takehiko Saito (National Institute of Animal Health) |
| EPI_ISL_759857 | A/chicken/Kagawa/M2T/2020 | A / H5N8 | Asia / Japan / Kagawa | Takehiko Saito (National Institute of Animal Health) |
| EPI_ISL_7598788 | A/Anser_brachyrhynchus/Belgium/13275_0009/2020 | A / H5N8 | Europe / Belgium / Provincie West-Vlaanderen / De Panne | Steven Van Borm (Sciensano, Department of Animal Infectious Diseases / Animal Infectious Diseases) |

| | | | | |
|---|---|---|---|---|
| EPI_ISL_7622619 | A/Anser_Brachyrhynchus_Anser_Anser/Belgium/13846/2020 | A / H5N8 | Europe / Belgium / Provincie West-Vlaanderen / Brugge and Zwin | Steven Van Borm (Sciensano, Department of Animal Infectious Diseases / Animal Infectious Diseases) |
| EPI_ISL_7622862 | A/Anser_brachyrhynchus/Belgium/151/2020 | A / H5N8 | Europe / Belgium / Provincie West-Vlaanderen / Zedelgem | Steven Van Borm (Sciensano, Department of Animal Infectious Diseases / Animal Infectious Diseases) |
| EPI_ISL_7623043 | A/Larus_argentatus/Belgium/568/2021 | A / H5N8 | Europe / Belgium / Provincie Oost-Vlaanderen / Merelbeke | Steven Van Borm (Sciensano, Department of Animal Infectious Diseases / Animal Infectious Diseases) |
| EPI_ISL_7623299 | A/Columba_palumbus/Belgium/2928_004/2021 | A / H5N8 | Europe / Belgium / Provincie Oost-Vlaanderen / Erembodegem | Steven Van Borm (Sciensano, Department of Animal Infectious Diseases / Animal Infectious Diseases) |
| EPI_ISL_7623303 | A/Garrulus_glandarius/Belgium/2928_003/2021 | A / H5N8 | Europe / Belgium / Provincie Oost-Vlaanderen / Lokeren | Steven Van Borm (Sciensano, Department of Animal Infectious Diseases / Animal Infectious Diseases) |
| EPI_ISL_7623511 | A/Gallus_gallus/Belgium/7578_001/2021 | A / H5N8 | Europe / Belgium / Province de Hainaut / Le Roeulx | Steven Van Borm (Sciensano, Department of Animal Infectious Diseases / Animal Infectious Diseases) |
| EPI_ISL_7623789 | A/Gallus_gallus/Belgium/9247/2021 | A / H5N8 | Europe / Belgium / Provincie Oost-Vlaanderen / Moerzeke | Steven Van Borm (Sciensano, Department of Animal Infectious Diseases / Animal Infectious Diseases) |
| EPI_ISL_7624142 | A/Gallus_gallus/Belgium/11644/2021 | A / H5N8 | Europe / Belgium / Province de Luxembourg / Attert | Steven Van Borm (Sciensano, Department of Animal Infectious Diseases / Animal Infectious Diseases) |
| EPI_ISL_7626484 | A/chicken/Czech Republic/23404-2K/2021 | A / H5N1 | Europe / Czech Republic / South Moravian Region / Lednice; ZipCode:69103 | Alexander Nagy (State Veterinary Institute Prague) |
| EPI_ISL_7626506 | A/chicken/Czech Republic/23404-4K/2021 | A / H5N1 | Europe / Czech Republic / South Moravian Region / Okres Breclav / Lednice; ZipCode:69103 | Alexander Nagy (State Veterinary Institute Prague) |
| EPI_ISL_7626512 | A/chicken/Czech Republic/23404-2T/2021 | A / H5N1 | Europe / Czech Republic / South Moravian Region / Okres Breclav / Lednice; ZipCode:69103 | Alexander Nagy (State Veterinary Institute Prague) |
| EPI_ISL_7626513 | A/chicken/Czech Republic/23404-4T/2021 | A / H5N1 | Europe / Czech Republic / South Moravian Region / Okres Breclav / Lednice; ZipCode:69103 | Alexander Nagy (State Veterinary Institute Prague) |
| EPI_ISL_7626514 | A/duck/Czech Republic/23589-1T/2021 | A / H5N1 | Europe / Czech Republic / Plzensky Kraj / Okres Plzen-Jih / Zdirec-Myt; ZipCode:33601 | Alexander Nagy (State Veterinary Institute Prague) |
| EPI_ISL_7626526 | A/chicken/Czech Republic/23589-1T/2021 | A / H5N1 | Europe / Czech Republic / Plzensky Kraj / Okres Plzen-Jih / Zdirec-Myt; ZipCode:33601 | Alexander Nagy (State Veterinary Institute Prague) |
| EPI_ISL_7626532 | A/chicken/Czech Republic/23589-3/2021 | A / H5N1 | Europe / Czech Republic / Plzensky Kraj / Okres Plzen-Jih / Zdirec-Myt; ZipCode:33601 | Alexander Nagy (State Veterinary Institute Prague) |
| EPI_ISL_7626533 | A/chicken/Czech Republic/23589-4/2021 | A / H5N1 | Europe / Czech Republic / Plzensky Kraj / Okres Plzen-Jih / Zdirec-Myt; ZipCode:33601 | Alexander Nagy (State Veterinary Institute Prague) |
| EPI_ISL_7635907 | A/white-tailed eagle/Sweden/SVA211201SZ0380/FB004721/M | A / H5N1 | Europe / Sweden / Skane Lan / Svedala Kommun | Siamak Zohari (National Veterinary Institute) |
| EPI_ISL_766052 | A/turkey/England/038730/2020 | A / H5N8 | Europe / United Kingdom / Norfolk | Alex Byrne (Animal and Plant Health Agency (APHA) / Virology Department) |
| EPI_ISL_766053 | A/turkey/England/039352/2020 | A / H5N8 | Europe / United Kingdom / Norfolk | Alex Byrne (Animal and Plant Health Agency (APHA) / Virology Department) |
| EPI_ISL_766054 | A/turkey/England/039472/2020 | A / H5N8 | Europe / United Kingdom / Norfolk | Alex Byrne (Animal and Plant Health Agency (APHA) / Virology Department) |
| EPI_ISL_766056 | A/falcon/England/041976/2020 | A / H5N8 | Europe / United Kingdom / Derbyshire | Alex Byrne (Animal and Plant Health Agency (APHA) / Virology Department) |
| EPI_ISL_766876 | A/mute_swan/England/234255/2020 | A / H5N1 | Europe / United Kingdom / Essex | Alex Byrne (Animal and Plant Health Agency (APHA) / Virology Department) |
| EPI_ISL_7683077 | A/mute swan/Netherlands/21039407-001/2021 | A / H5N1 | Europe / Netherlands / Provincie Gelderland | Rene Heutink (Wageningen Bioveterinary Research) |
| EPI_ISL_7683079 | A/mute swan/Netherlands/21039627-002/2021 | A / H5N1 | Europe / Netherlands / South Holland | Rene Heutink (Wageningen Bioveterinary Research) |
| EPI_ISL_7683101 | A/common buzzard/Netherlands/21039607-002/2021 | A / H5N1 | Europe / Netherlands / Provincie Noord-Holland | Rene Heutink (Wageningen Bioveterinary Research) |
| EPI_ISL_7683118 | A/red fox/Netherlands/21040099-001/2021 | A / H5N1 | Europe / Netherlands / North Brabant | Rene Heutink (Wageningen Bioveterinary Research) |
| EPI_ISL_7683137 | A/red fox/Netherlands/21040099-006/2021 | A / H5N1 | Europe / Netherlands / North Brabant | Rene Heutink (Wageningen Bioveterinary Research) |
| EPI_ISL_7683138 | A/red fox/Netherlands/21040099-007/2021 | A / H5N1 | Europe / Netherlands / North Brabant | Rene Heutink (Wageningen Bioveterinary Research) |
| EPI_ISL_7733048 | A/black-headed gull/Germany-NI/AI05962/2021 | A / H5N1 | Europe / Germany / Lower Saxony / Aurich | Jacqueline King (Friedrich-Loeffler-Institut) |
| EPI_ISL_7733582 | A/chicken/Italy/21VIR10388/2021 | A / H5N1 | Europe / Italy / BS | Bianca Zecchin (Istituto Zooprofilattico Sperimentale Delle Venezie) |
| EPI_ISL_7733584 | A/duck/Italy/21VIR10447/2021 | A / H5N1 | Europe / Italy / BS | Bianca Zecchin (Istituto Zooprofilattico Sperimentale Delle Venezie) |
| EPI_ISL_7733585 | A/Eurasian_wigeon/Italy/21VIR8919-3/2021 | A / H5N1 | Europe / Italy / BS | Bianca Zecchin (Istituto Zooprofilattico Sperimentale Delle Venezie) |
| EPI_ISL_7733587 | A/turkey/Italy/21VIR9618-7/2021 | A / H5N1 | Europe / Italy / BS | Bianca Zecchin (Istituto Zooprofilattico Sperimentale Delle Venezie) |
| EPI_ISL_7733591 | A/mallard/Italy/21VIR8919-2/2021 | A / H5N1 | Europe / Italy / BS | Bianca Zecchin (Istituto Zooprofilattico Sperimentale Delle Venezie) |
| EPI_ISL_7733595 | A/chicken/Italy/21VIR10239/2021 | A / H5N1 | Europe / Italy / CR | Bianca Zecchin (Istituto Zooprofilattico Sperimentale Delle Venezie) |
| EPI_ISL_7733597 | A/turkey/Italy/21VIR10340/2021 | A / H5N1 | Europe / Italy / CR | Bianca Zecchin (Istituto Zooprofilattico Sperimentale Delle Venezie) |
| EPI_ISL_7733598 | A/turkey/Italy/21VIR9816-1/2021 | A / H5N1 | Europe / Italy / MN | Bianca Zecchin (Istituto Zooprofilattico Sperimentale Delle Venezie) |
| EPI_ISL_7733599 | A/chicken/Italy/21VIR10389/2021 | A / H5N1 | Europe / Italy / MN | Bianca Zecchin (Istituto Zooprofilattico Sperimentale Delle Venezie) |
| EPI_ISL_7733603 | A/chicken/Italy/21VIR9951-25/2021 | A / H5N1 | Europe / Italy / MN | Bianca Zecchin (Istituto Zooprofilattico Sperimentale Delle Venezie) |
| EPI_ISL_7733607 | A/chicken/Italy/21VIR10384/2021 | A / H5N1 | Europe / Italy / PD | Bianca Zecchin (Istituto Zooprofilattico Sperimentale Delle Venezie) |
| EPI_ISL_7733609 | A/chicken/Italy/21VIR9580-22/2021 | A / H5N1 | Europe / Italy / PD | Bianca Zecchin (Istituto Zooprofilattico Sperimentale Delle Venezie) |
| EPI_ISL_7733610 | A/chicken/Italy/21VIR10343/2021 | A / H5N1 | Europe / Italy / UD | Bianca Zecchin (Istituto Zooprofilattico Sperimentale Delle Venezie) |
| EPI_ISL_7733611 | A/Wild_goose/Italy/21VIR10193/2021 | A / H5N1 | Europe / Italy / VE | Bianca Zecchin (Istituto Zooprofilattico Sperimentale Delle Venezie) |
| EPI_ISL_7733612 | A/magpie/Italy/21VIR9487-2/2021 | A / H5N1 | Europe / Italy / UD | Bianca Zecchin (Istituto Zooprofilattico Sperimentale Delle Venezie) |
| EPI_ISL_7733613 | A/turkey/Italy/21VIR10456/2021 | A / H5N1 | Europe / Italy / VI | Bianca Zecchin (Istituto Zooprofilattico Sperimentale Delle Venezie) |
| EPI_ISL_7733617 | A/chicken/Italy/21VIR9133-20/2021 | A / H5N1 | Europe / Italy / VR | Bianca Zecchin (Istituto Zooprofilattico Sperimentale Delle Venezie) |
| EPI_ISL_7733621 | A/seagull/Italy/21VIR9432-2/2021 | A / H5N1 | Europe / Italy / VR | Bianca Zecchin (Istituto Zooprofilattico Sperimentale Delle Venezie) |
| EPI_ISL_7733624 | A/turkey/Italy/21VIR9767-3/2021 | A / H5N1 | Europe / Italy / VR | Bianca Zecchin (Istituto Zooprofilattico Sperimentale Delle Venezie) |
| EPI_ISL_7733628 | A/turkey/Italy/21VIR10251/2021 | A / H5N1 | Europe / Italy / VR | Bianca Zecchin (Istituto Zooprofilattico Sperimentale Delle Venezie) |
| EPI_ISL_7733629 | A/turkey/Italy/21VIR9652-2/2021 | A / H5N1 | Europe / Italy / VR | Bianca Zecchin (Istituto Zooprofilattico Sperimentale Delle Venezie) |
| EPI_ISL_7733630 | A/chicken/Italy/21VIR9691-2/2021 | A / H5N1 | Europe / Italy / VR | Bianca Zecchin (Istituto Zooprofilattico Sperimentale Delle Venezie) |
| EPI_ISL_7733632 | A/turkey/Italy/21VIR9512-1/2021 | A / H5N1 | Europe / Italy / VR | Bianca Zecchin (Istituto Zooprofilattico Sperimentale Delle Venezie) |
| EPI_ISL_7733633 | A/turkey/Italy/21VIR9143-2/2021 | A / H5N1 | Europe / Italy / VR | Bianca Zecchin (Istituto Zooprofilattico Sperimentale Delle Venezie) |
| EPI_ISL_7733634 | A/chicken/Italy/21VIR9765-12/2021 | A / H5N1 | Europe / Italy / VR | Bianca Zecchin (Istituto Zooprofilattico Sperimentale Delle Venezie) |
| EPI_ISL_7733635 | A/turkey/Italy/21VIR9510-1/2021 | A / H5N1 | Europe / Italy / VR | Bianca Zecchin (Istituto Zooprofilattico Sperimentale Delle Venezie) |
| EPI_ISL_7733638 | A/turkey/Italy/21VIR9768-8/2021 | A / H5N1 | Europe / Italy / VR | Bianca Zecchin (Istituto Zooprofilattico Sperimentale Delle Venezie) |
| EPI_ISL_7733641 | A/chicken/Italy/21VIR10352/2021 | A / H5N1 | Europe / Italy / VR | Bianca Zecchin (Istituto Zooprofilattico Sperimentale Delle Venezie) |
| EPI_ISL_7733642 | A/kestrel/Italy/21VIR10468/2021 | A / H5N1 | Europe / Italy / VR | Bianca Zecchin (Istituto Zooprofilattico Sperimentale Delle Venezie) |
| EPI_ISL_7733643 | A/turkey/Italy/21VIR9649-2/2021 | A / H5N1 | Europe / Italy / VR | Bianca Zecchin (Istituto Zooprofilattico Sperimentale Delle Venezie) |
| EPI_ISL_7733644 | A/chicken/Italy/IZSLT-122448_21VIR9218-1/2021 | A / H5N1 | Europe / Italy / Roma | Bianca Zecchin (Istituto Zooprofilattico Sperimentale Delle Venezie) |
| EPI_ISL_7733645 | A/goose/Italy/IZSLT-21VIR10273/2021 | A / H5N1 | Europe / Italy / Roma | Bianca Zecchin (Istituto Zooprofilattico Sperimentale Delle Venezie) |
| EPI_ISL_7733647 | A/turkey/Italy/21VIR9210-1/2021 | A / H5N1 | Europe / Italy / VR | Bianca Zecchin (Istituto Zooprofilattico Sperimentale Delle Venezie) |
| EPI_ISL_7733648 | A/turkey/Italy/21VIR8817-1/2021 | A / H5N1 | Europe / Italy / VR | Bianca Zecchin (Istituto Zooprofilattico Sperimentale Delle Venezie) |
| EPI_ISL_7748001 | A/Eurasian curlew/Germany-SH/AI05960/2021 | A / H5N1 | Europe / Germany / Schleswig-Holstein / Nordfriesland | Jacqueline King (Friedrich-Loeffler-Institut) |
| EPI_ISL_7748188 | A/Eurasian wigeon/Germany-SH/AI05950/2021 | A / H5N1 | Europe / Germany / Schleswig-Holstein / Nordfriesland | Jacqueline King (Friedrich-Loeffler-Institut) |
| EPI_ISL_7748262 | A/Eurasian wigeon/Germany-SH/AI05952/2021 | A / H5N1 | Europe / Germany / Schleswig-Holstein / Nordfriesland | Jacqueline King (Friedrich-Loeffler-Institut) |
| EPI_ISL_775248 | A/muscovy duck/Netherlands/20019914-001/2020 | A / H5N8 | Europe / Netherlands / South Holland / Gemeente alphen aan den Rijn | Rene Heutink (Wageningen Bioveterinary Research) |
| EPI_ISL_775251 | A/peregrine falcon/Netherlands/20020038-001/2020 | A / H5N8 | Europe / Netherlands / Provincie Limburg / Gemeente Nederweert | Rene Heutink (Wageningen Bioveterinary Research) |
| EPI_ISL_775253 | A/mute swan/Netherlands/20020133-001/2020 | A / H5N8 | Europe / Netherlands / Provincie Groningen / Gemeente Groningen | Rene Heutink (Wageningen Bioveterinary Research) |
| EPI_ISL_775266 | A/black-headed gull/Netherlands/20020162-002/2020 | A / H5N8 | Europe / Netherlands / Provincie Flevoland / Almere | Rene Heutink (Wageningen Bioveterinary Research) |
| EPI_ISL_775267 | A/turkey/Netherlands/21020942-001005/2021 | A / H5N8 | Europe / Netherlands / North Brabant / Gemeente Oisterwijk | Rene Heutink (Wageningen Bioveterinary Research) |
| EPI_ISL_7753130 | A/Eurasian wigeon/Germany-SH/AI05955/2021 | A / H5N1 | Europe / Germany / Schleswig-Holstein / Nordfriesland | Jacqueline King (Friedrich-Loeffler-Institut) |

| | | | | |
|---|---|---|---|---|
| EPI_ISL_7753152 | A/Eurasian wigeon/Germany-SH/AI05953/2021 | A / H5N1 | Europe / Germany / Schleswig-Holstein / Nordfriesland | Jacqueline King (Friedrich-Loeffler-Institut) |
| EPI_ISL_7753173 | A/Eurasian wigeon/Germany-SH/AI05956/2021 | A / H5N1 | Europe / Germany / Schleswig-Holstein / Nordfriesland | Jacqueline King (Friedrich-Loeffler-Institut) |
| EPI_ISL_7753193 | A/Eurasian wigeon/Germany-SH/AI05951/2021 | A / H5N1 | Europe / Germany / Schleswig-Holstein / Nordfriesland | Jacqueline King (Friedrich-Loeffler-Institut) |
| EPI_ISL_7753211 | A/Eurasian wigeon/Germany-SH/AI05954/2021 | A / H5N1 | Europe / Germany / Schleswig-Holstein / Nordfriesland | Jacqueline King (Friedrich-Loeffler-Institut) |
| EPI_ISL_7753230 | A/mallard/Germany-NI/AI06010/2021 | A / H5N1 | Europe / Germany / Lower Saxony / Harburg | Jacqueline King (Friedrich-Loeffler-Institut) |
| EPI_ISL_7753251 | A/barnacle goose/Germany-SH/AI06005/2021 | A / H5N1 | Europe / Germany / Schleswig-Holstein / Nordfriesland | Jacqueline King (Friedrich-Loeffler-Institut) |
| EPI_ISL_7753272 | A/white-tailed eagle/Germany-MV/AI05975/2021 | A / H5N1 | Europe / Germany / Mecklenburg-Vorpommern / Vorpommern-Greifswald | Jacqueline King (Friedrich-Loeffler-Institut) |
| EPI_ISL_7753290 | A/turkey/Germany-MV/AI06035/2021 | A / H5N1 | Europe / Germany / Mecklenburg-Vorpommern / Vorpommern-Greifswald | Jacqueline King (Friedrich-Loeffler-Institut) |
| EPI_ISL_7753309 | A/lesser white-fronted goose/Germany-MV/AI05973/2021 | A / H5N1 | Europe / Germany / Mecklenburg-Vorpommern / Vorpommern-Greifswald | Jacqueline King (Friedrich-Loeffler-Institut) |
| EPI_ISL_7753329 | A/white stork/Germany-MV/AI05979/2021 | A / H5N1 | Europe / Germany / Mecklenburg-Vorpommern / Vorpommern-Greifswald | Jacqueline King (Friedrich-Loeffler-Institut) |
| EPI_ISL_7753347 | A/herring gull/Germany-SH/AI06141/2021 | A / H5N1 | Europe / Germany / Schleswig-Holstein / Nordfriesland | Jacqueline King (Friedrich-Loeffler-Institut) |
| EPI_ISL_7753367 | A/Eurasian teal/Germany-BY/AI05977/2021 | A / H5N1 | Europe / Germany / Bayern | Jacqueline King (Friedrich-Loeffler-Institut) |
| EPI_ISL_7753387 | A/Eurasian wigeon/Germany-SH/AI06143/2021 | A / H5N1 | Europe / Germany / Schleswig-Holstein / Dithmarschen | Jacqueline King (Friedrich-Loeffler-Institut) |
| EPI_ISL_7753396 | A/domestic goose/Germany-SH/AI06147/2021 | A / H5N1 | Europe / Germany / Schleswig-Holstein / Dithmarschen | Jacqueline King (Friedrich-Loeffler-Institut) |
| EPI_ISL_7753399 | A/domestic goose/Germany-SH/AI06150/2021 | A / H5N1 | Europe / Germany / Schleswig-Holstein / Dithmarschen | Jacqueline King (Friedrich-Loeffler-Institut) |
| EPI_ISL_7753400 | A/domestic goose/Germany-SH/AI06024/2021 | A / H5N1 | Europe / Germany / Schleswig-Holstein / Dithmarschen | Jacqueline King (Friedrich-Loeffler-Institut) |
| EPI_ISL_7753405 | A/Eurasian wigeon/Germany-SH/AI06142/2021 | A / H5N1 | Europe / Germany / Schleswig-Holstein / Dithmarschen | Jacqueline King (Friedrich-Loeffler-Institut) |
| EPI_ISL_7753406 | A/greylag goose/Germany-SH/AI06144/2021 | A / H5N1 | Europe / Germany / Schleswig-Holstein / Dithmarschen | Jacqueline King (Friedrich-Loeffler-Institut) |
| EPI_ISL_7753407 | A/barnacle goose/Germany-SH/AI06145/2021 | A / H5N1 | Europe / Germany / Schleswig-Holstein / Dithmarschen | Jacqueline King (Friedrich-Loeffler-Institut) |
| EPI_ISL_7753425 | A/greylag goose/Germany-SH/AI06205/2021 | A / H5N1 | Europe / Germany / Schleswig-Holstein / Nordfriesland | Jacqueline King (Friedrich-Loeffler-Institut) |
| EPI_ISL_7753443 | A/buzzard/Germany-SH/AI06210/2021 | A / H5N1 | Europe / Germany / Schleswig-Holstein / Dithmarschen | Jacqueline King (Friedrich-Loeffler-Institut) |
| EPI_ISL_7778753 | A/Eagle_owl/Estonia/TA2122239_21VIR10433-10/2021 | A / H5N8 | Europe / Estonia | Bianca Zecchin (Istituto Zooprofilattico Sperimentale Delle Venezie) |
| EPI_ISL_7778754 | A/Withe-tiled_eagle/Estonia/TA2124126-1_21VIR10433-11/20 | A / H5N1 | Europe / Estonia | Bianca Zecchin (Istituto Zooprofilattico Sperimentale Delle Venezie) |
| EPI_ISL_7778755 | A/white-tailed_eagle/Estonia/TA2111864-2_21VIR7512-6/202 | A / H5N1 | Europe / Estonia | Bianca Zecchin (Istituto Zooprofilattico Sperimentale Delle Venezie) |
| EPI_ISL_7778756 | A/gull/Estonia/TA2113284-4_21VIR7512-8/2021 | A / H5N1 | Europe / Estonia | Bianca Zecchin (Istituto Zooprofilattico Sperimentale Delle Venezie) |
| EPI_ISL_7778757 | A/mute_swan/Estonia/TA2106111-2_21VIR7512-1/2021 | A / H5N8 | Europe / Estonia | Bianca Zecchin (Istituto Zooprofilattico Sperimentale Delle Venezie) |
| EPI_ISL_7778758 | A/mute_swan/Estonia/TA2106241_21VIR7512-2/2021 | A / H5N8 | Europe / Estonia | Bianca Zecchin (Istituto Zooprofilattico Sperimentale Delle Venezie) |
| EPI_ISL_7778759 | A/mute_swan/Estonia/TA2106419-1_21VIR7512-3/2021 | A / H5N8 | Europe / Estonia | Bianca Zecchin (Istituto Zooprofilattico Sperimentale Delle Venezie) |
| EPI_ISL_7778760 | A/mute_swan/Estonia/TA2106615_21VIR7512-4/2021 | A / H5N8 | Europe / Estonia | Bianca Zecchin (Istituto Zooprofilattico Sperimentale Delle Venezie) |
| EPI_ISL_7778761 | A/mute_swan/Estonia/TA2108545-1_21VIR7512-5/2021 | A / H5N8 | Europe / Estonia | Bianca Zecchin (Istituto Zooprofilattico Sperimentale Delle Venezie) |
| EPI_ISL_7778762 | A/guinea_fowl/Estonia/TA2104719_21VIR7512-9/2021 | A / H5N8 | Europe / Estonia | Bianca Zecchin (Istituto Zooprofilattico Sperimentale Delle Venezie) |
| EPI_ISL_7778763 | A/white-tailed_eagle/Finland/6984_21VIR7689-10/2021 | A / H5N1 | Europe / Finland | Bianca Zecchin (Istituto Zooprofilattico Sperimentale Delle Venezie) |
| EPI_ISL_7778764 | A/white-tailed_eagle/Finland/9257_21VIR7689-11/2021 | A / H5N8 | Europe / Finland | Bianca Zecchin (Istituto Zooprofilattico Sperimentale Delle Venezie) |
| EPI_ISL_7778765 | A/golden_eagle/Finland/9378_21VIR7689-12/2021 | A / H5N1 | Europe / Finland | Bianca Zecchin (Istituto Zooprofilattico Sperimentale Delle Venezie) |
| EPI_ISL_7778766 | A/European_herring_gull/Finland/9722_21VIR7689-13/2021 | A / H5N1 | Europe / Finland | Bianca Zecchin (Istituto Zooprofilattico Sperimentale Delle Venezie) |
| EPI_ISL_7778767 | A/whooper_swan/Finland/9906_21VIR7689-14/2021 | A / H5N8 | Europe / Finland | Bianca Zecchin (Istituto Zooprofilattico Sperimentale Delle Venezie) |
| EPI_ISL_7778768 | A/Eurasian_eagle-owl/Finland/10617_21VIR7689-15/2021 | A / H5N1 | Europe / Finland | Bianca Zecchin (Istituto Zooprofilattico Sperimentale Delle Venezie) |
| EPI_ISL_7778769 | A/pheasant/Finland/499_21VIR7689-1/2021 | A / H5N8 | Europe / Finland | Bianca Zecchin (Istituto Zooprofilattico Sperimentale Delle Venezie) |
| EPI_ISL_7778770 | A/mute_swan/Finland/1325_21VIR7689-2/2021 | A / H5N8 | Europe / Finland | Bianca Zecchin (Istituto Zooprofilattico Sperimentale Delle Venezie) |
| EPI_ISL_7778771 | A/pheasant/Finland/1589_21VIR7689-3/2021 | A / H5N8 | Europe / Finland | Bianca Zecchin (Istituto Zooprofilattico Sperimentale Delle Venezie) |
| EPI_ISL_7778773 | A/barnacle_goose/Finland/6247_21VIR7689-6/2021 | A / H5N1 | Europe / Finland | Bianca Zecchin (Istituto Zooprofilattico Sperimentale Delle Venezie) |
| EPI_ISL_7778774 | A/barnacle_goose/Finland//6378_21VIR7689-7/2021 | A / H5N1 | Europe / Finland | Bianca Zecchin (Istituto Zooprofilattico Sperimentale Delle Venezie) |
| EPI_ISL_7778775 | A/barnacle_goose/Finland/6955_21VIR7689-9/2021 | A / H5N1 | Europe / Finland | Bianca Zecchin (Istituto Zooprofilattico Sperimentale Delle Venezie) |
| EPI_ISL_7778776 | A/greylag_goose/Norway/V294_21VIR7634-1/2021 | A / H5N8 | Europe / Norway | Bianca Zecchin (Istituto Zooprofilattico Sperimentale Delle Venezie) |
| EPI_ISL_7778777 | A/mute_swan/Norway/FU452_21VIR7634-2/2021 | A / H5N8 | Europe / Norway | Bianca Zecchin (Istituto Zooprofilattico Sperimentale Delle Venezie) |
| EPI_ISL_7778778 | A/common_eider/Norway/FU453_21VIR7634-3/2021 | A / H5N8 | Europe / Norway | Bianca Zecchin (Istituto Zooprofilattico Sperimentale Delle Venezie) |
| EPI_ISL_7778779 | A/common_eider/Norway/FU458_21VIR7634-4/2021 | A / H5N8 | Europe / Norway | Bianca Zecchin (Istituto Zooprofilattico Sperimentale Delle Venezie) |
| EPI_ISL_7778780 | A/common_eider/Norway/FU474_21VIR7634-5/2021 | A / H5N8 | Europe / Norway | Bianca Zecchin (Istituto Zooprofilattico Sperimentale Delle Venezie) |
| EPI_ISL_7778880 | A/Red_fox/Estonia/TA2126820_21VIR10433-13/2021 | A / H5N1 | Europe / Estonia | Bianca Zecchin (Istituto Zooprofilattico Sperimentale Delle Venezie) |
| EPI_ISL_779129 | A/turkey/Poland/464/2020(H5N8) | A / H5N8 | Europe / Poland / Masovian Voivodeship | Edyta ?wi?to? (National Veterinary Research Institut Poland, PIWet-PIB) |
| EPI_ISL_780096 | A/Duck/Hungary/14788/2020 | A / H5N8 | Europe / Hungary / Bacs-Kiskun | Katalin Szentpáli-Gavallér (National Food Chain Safety Office, Hungary / Virology) |
| EPI_ISL_7880689 | A/Gallus_gallus/Belgium/16070_003/2021 | A / H5N1 | Europe / Belgium / Provincie Antwerpen / Weelde (Ravels) | Steven Van Borm (Sciensano, Department of Animal Infectious Diseases / Animal Infectious Diseases) |
| EPI_ISL_7880696 | A/Gallus_gallus/Belgium/15977/2021 | A / H5N1 | Europe / Belgium / Provincie West-Vlaanderen / Alveringem | Steven Van Borm (Sciensano, Department of Animal Infectious Diseases / Animal Infectious Diseases) |
| EPI_ISL_7892492 | A/grey heron/Croatia/132/2021 | A / H5N1 | Europe / Croatia / Osjecko-Baranjska Zupanija / Kopacki rit (45.606587° 18.801463°) | Vladimir Savi? (Croatian Veterinary Institute / Poultry Centre) |
| EPI_ISL_7952115 | A/mute swan/Netherlands/21039526-002/2021 | A / H5N1 | Europe / Netherlands / South Holland | Rene Heutink (Wageningen Bioveterinary Research) |
| EPI_ISL_7952116 | A/chicken/Netherlands/21039901-001-005/2021 | A / H5N1 | Europe / Netherlands / South Holland | Rene Heutink (Wageningen Bioveterinary Research) |
| EPI_ISL_7952117 | A/chicken/Netherlands/21040808-001-005/2021 | A / H5N1 | Europe / Netherlands / Provincie Overijssel | Rene Heutink (Wageningen Bioveterinary Research) |
| EPI_ISL_7952118 | A/chicken/Netherlands/21040811-001-005/2021 | A / H5N1 | Europe / Netherlands / Provincie Overijssel | Rene Heutink (Wageningen Bioveterinary Research) |
| EPI_ISL_7952119 | A/great black-backed gull/Netherlands/21039609-001-002/202 | A / H5N1 | Europe / Netherlands / Provincie Friesland | Rene Heutink (Wageningen Bioveterinary Research) |
| EPI_ISL_7952120 | A/greylag goose/Netherlands/21039525-001/2021 | A / H5N1 | Europe / Netherlands / South Holland | Rene Heutink (Wageningen Bioveterinary Research) |
| EPI_ISL_7952121 | A/sanderling/Netherlands/21039528-002/2021 | A / H5N1 | Europe / Netherlands / Provincie Friesland | Rene Heutink (Wageningen Bioveterinary Research) |
| EPI_ISL_7952122 | A/greylag goose/Netherlands/21039746-002/2021 | A / H5N1 | Europe / Netherlands / South Holland | Rene Heutink (Wageningen Bioveterinary Research) |
| EPI_ISL_7952123 | A/barnacle goose/Netherlands/21039750-002/2021 | A / H5N1 | Europe / Netherlands / Provincie Flevoland | Rene Heutink (Wageningen Bioveterinary Research) |
| EPI_ISL_7952124 | A/mute swan/Netherlands/21039841-002/2021 | A / H5N1 | Europe / Netherlands / Provincie Utrecht | Rene Heutink (Wageningen Bioveterinary Research) |
| EPI_ISL_7952125 | A/mute swan/Netherlands/21039824-002/2021 | A / H5N1 | Europe / Netherlands / Provincie Friesland | Rene Heutink (Wageningen Bioveterinary Research) |
| EPI_ISL_7952126 | A/greylag goose/Netherlands/21038417-001/2021 | A / H5N1 | Europe / Netherlands / South Holland | Rene Heutink (Wageningen Bioveterinary Research) |
| EPI_ISL_7952127 | A/goose/Netherlands/21039364-002/2021 | A / H5N1 | Europe / Netherlands / Provincie Friesland | Rene Heutink (Wageningen Bioveterinary Research) |
| EPI_ISL_7952128 | A/gadwall/Netherlands/21038566-002/2021 | A / H5N1 | Europe / Netherlands / South Holland | Rene Heutink (Wageningen Bioveterinary Research) |
| EPI_ISL_7952129 | A/western jackdaw/Netherlands/21039297-002/2021 | A / H5N1 | Europe / Netherlands / Provincie Flevoland | Rene Heutink (Wageningen Bioveterinary Research) |
| EPI_ISL_796011 | A/Goose/Hungary/15267/2020 | A / H5N8 | Europe / Hungary / Bacs-Kiskun | Katalin Szentpáli-Gavallér (National Food Chain Safety Office, Hungary / Virology) |
| EPI_ISL_7983813 | A/Hangzhou/01/2021 | A / H5N6 | Asia / China / Zhejiang Province / Hangzhou | Jun Li (Hangzhou Center for Disease Control and Prevention ) |
| EPI_ISL_7996368 | A/mute swan/Netherlands/21038568-005/2021 | A / H5N1 | Europe / Netherlands / Provincie Friesland | Rene Heutink (Wageningen Bioveterinary Research) |

| | | | | |
|---|---|---|---|---|
| EPI_ISL_7996369 | A/mute swan/Netherlands/21039291-002/2021 | A / H5N1 | Europe / Netherlands / South Holland | Rene Heutink (Wageningen Bioveterinary Research) |
| EPI_ISL_7996370 | A/western jackdaw/Netherlands/21039294-002/2021 | A / H5N1 | Europe / Netherlands / Provincie Flevoland | Rene Heutink (Wageningen Bioveterinary Research) |
| EPI_ISL_7996371 | A/turkey/Netherlands/21040980-001005/2021 | A / H5N1 | Europe / Netherlands / Provincie Limburg | Rene Heutink (Wageningen Bioveterinary Research) |
| EPI_ISL_804043 | A/Mallard_duck/Hungary/17319/2020 | A / H5N8 | Europe / Hungary / Bacs-Kiskun | Katalin Szentpáli-Gavallér (National Food Chain Safety Office, Hungary / Virology) |
| EPI_ISL_804045 | A/Duck/Hungary/17806/2020 | A / H5N8 | Europe / Hungary / Bacs-Kiskun | Katalin Szentpáli-Gavallér (National Food Chain Safety Office, Hungary / Virology) |
| EPI_ISL_804047 | A/Duck/Hungary/17957/2020 | A / H5N8 | Europe / Hungary / Csongrad megye | Katalin Szentpáli-Gavallér (National Food Chain Safety Office, Hungary / Virology) |
| EPI_ISL_808250 | A/Duck/Hungary/18358/2020 | A / H5N8 | Europe / Hungary / Bacs-Kiskun | Katalin Szentpáli-Gavallér (National Food Chain Safety Office, Hungary / Virology) |
| EPI_ISL_809658 | A/Goose/Hungary/18406/2020 | A / H5N8 | Europe / Hungary / Bacs-Kiskun | Katalin Szentpáli-Gavallér (National Food Chain Safety Office, Hungary / Virology) |
| EPI_ISL_810963 | A/Duck/Hungary/18444/2020 | A / H5N8 | Europe / Hungary / Bacs-Kiskun | Katalin Szentpáli-Gavallér (National Food Chain Safety Office, Hungary / Virology) |
| EPI_ISL_810970 | A/Chicken/Hungary/18466/2020 | A / H5N8 | Europe / Hungary / Bacs-Kiskun | Katalin Szentpáli-Gavallér (National Food Chain Safety Office, Hungary / Virology) |
| EPI_ISL_811128 | A/Chicken/Hungary/18467/2020 | A / H5N8 | Europe / Hungary / Bacs-Kiskun | Katalin Szentpáli-Gavallér (National Food Chain Safety Office, Hungary / Virology) |
| EPI_ISL_811134 | A/Chicken/Hungary/19776/2020 | A / H5N8 | Europe / Hungary / Csongrad megye | Katalin Szentpáli-Gavallér (National Food Chain Safety Office, Hungary / Virology) |
| EPI_ISL_811135 | A/Chicken/Hungary/20227/2020 | A / H5N8 | Europe / Hungary / Csongrad megye | Katalin Szentpáli-Gavallér (National Food Chain Safety Office, Hungary / Virology) |
| EPI_ISL_811144 | A/Goose/Hungary/21737/2020 | A / H5N8 | Europe / Hungary / Bekes megye | Katalin Szentpáli-Gavallér (National Food Chain Safety Office, Hungary / Virology) |
| EPI_ISL_811146 | A/Turkey/Hungary/21753/2020 | A / H5N8 | Europe / Hungary / Csongrad megye | Katalin Szentpáli-Gavallér (National Food Chain Safety Office, Hungary / Virology) |
| EPI_ISL_813599 | A/Goose/Hungary/22493/2020 | A / H5N8 | Europe / Hungary / Bacs-Kiskun | Katalin Szentpáli-Gavallér (National Food Chain Safety Office, Hungary / Virology) |
| EPI_ISL_813971 | A/Turkey/Hungary/22494/2020 | A / H5N8 | Europe / Hungary / Bekes megye | Katalin Szentpáli-Gavallér (National Food Chain Safety Office, Hungary / Virology) |
| EPI_ISL_813972 | A/Goose/Hungary/24021/2020 | A / H5N8 | Europe / Hungary / Bekes megye | Katalin Szentpáli-Gavallér (National Food Chain Safety Office, Hungary / Virology) |
| EPI_ISL_813979 | A/peregrine_falcon/Ireland/20VIR7872-1/2020 | A / H5N8 | Europe / Ireland | Bianca Zecchin (Istituto Zooprofilattico Sperimentale Delle Venezie) |
| EPI_ISL_8215653 | A/Bar-headed Goose/Tibet/XZQ5-1/2021 | A / H5N8 | Asia / China / Tibet Autonomous Region | Hongliang Chai (Northeast Forestry University / College of Wildlife Resources) |
| EPI_ISL_8215654 | A/Bar-headed Goose/Tibet/XZQ7/2021 | A / H5N8 | Asia / China / Tibet Autonomous Region | Hongliang Chai (Northeast Forestry University / College of Wildlife Resources) |
| EPI_ISL_8215655 | A/Bar-headed Goose/Tibet/XZQ8-1/2021 | A / H5N8 | Asia / China / Tibet Autonomous Region | Hongliang Chai (Northeast Forestry University / College of Wildlife Resources) |
| EPI_ISL_8215656 | A/Bar-headed Goose/Tibet/XZQ9-1/2021 | A / H5N8 | Asia / China / Tibet Autonomous Region | Hongliang Chai (Northeast Forestry University / College of Wildlife Resources) |
| EPI_ISL_8215657 | A/Bar-headed Goose/Tibet/XZQ10-1/2021 | A / H5N8 | Asia / China / Tibet Autonomous Region | Hongliang Chai (Northeast Forestry University / College of Wildlife Resources) |
| EPI_ISL_8215658 | A/Bar-headed Goose/Tibet/XZ6/2021 | A / H5N8 | Asia / China / Tibet Autonomous Region | Hongliang Chai (Northeast Forestry University / College of Wildlife Resources) |
| EPI_ISL_8215659 | A/Bar-headed Goose/Tibet/XZ71/2021 | A / H5N8 | Asia / China / Tibet Autonomous Region | Hongliang Chai (Northeast Forestry University / College of Wildlife Resources) |
| EPI_ISL_8215660 | A/Bar-headed Goose/Tibet/XZ81/2021 | A / H5N8 | Asia / China / Tibet Autonomous Region | Hongliang Chai (Northeast Forestry University / College of Wildlife Resources) |
| EPI_ISL_8215661 | A/Bar-headed Goose/Tibet/XZ181/2021 | A / H5N8 | Asia / China / Tibet Autonomous Region | Hongliang Chai (Northeast Forestry University / College of Wildlife Resources) |
| EPI_ISL_8215662 | A/Bar-headed Goose/Tibet/XZQ13-1/2021 | A / H5N8 | Asia / China / Tibet Autonomous Region | Hongliang Chai (Northeast Forestry University / College of Wildlife Resources) |
| EPI_ISL_8215663 | A/Brown-headed Gull/Tibet/XZQ15-2/2021 | A / H5N8 | Asia / China / Tibet Autonomous Region | Hongliang Chai (Northeast Forestry University / College of Wildlife Resources) |
| EPI_ISL_8215684 | A/Brown-headed Gull/Tibet/XZQ16-2/2021 | A / H5N8 | Asia / China / Tibet Autonomous Region | Hongliang Chai (Northeast Forestry University / College of Wildlife Resources) |
| EPI_ISL_8215685 | A/Bar-headed Goose/Tibet/XZQ17-1/2021 | A / H5N8 | Asia / China / Tibet Autonomous Region | Hongliang Chai (Northeast Forestry University / College of Wildlife Resources) |
| EPI_ISL_8215686 | A/Bar-headed Goose/Tibet/XZQ18-1/2021 | A / H5N8 | Asia / China / Tibet Autonomous Region | Hongliang Chai (Northeast Forestry University / College of Wildlife Resources) |
| EPI_ISL_8215687 | A/Brown-headed Gull/Tibet/XZ19/2021 | A / H5N8 | Asia / China / Tibet Autonomous Region | Hongliang Chai (Northeast Forestry University / College of Wildlife Resources) |
| EPI_ISL_8215688 | A/Bar-headed Goose/Tibet/XZ901/2021 | A / H5N1 | Asia / China / Tibet Autonomous Region | Hongliang Chai (Northeast Forestry University / College of Wildlife Resources) |
| EPI_ISL_8215689 | A/Bar-headed Goose/Tibet/XZ1131/2021 | A / H5N1 | Asia / China / Tibet Autonomous Region | Hongliang Chai (Northeast Forestry University / College of Wildlife Resources) |
| EPI_ISL_826283 | A/Chicken/Sweden/SVA201221SZ0027/KN342632-IP2/2020 | A / H5N8 | Europe / Sweden / Skane Lan / Svedala Kommun | Siamak Zohari (National Veterinary Institute) |
| EPI_ISL_826285 | A/barnacle goose/Sweden/SVA201215SZ0368/KN003768/202( | A / H5N8 | Europe / Sweden / Skane Lan / Malmo Kommun | Siamak Zohari (National Veterinary Institute) |
| EPI_ISL_826296 | A/barnacle goose/Sweden/SVA201221SZ0154/KN003828/202( | A / H5N8 | Europe / Sweden / Skane Lan / Skurups Kommun | Siamak Zohari (National Veterinary Institute) |
| EPI_ISL_826457 | A/Chicken/Sweden/SVA210102SZ0002/KN000002-IP1/2021 | A / H5N8 | Europe / Sweden / Skane Lan / Sjobo Kommun | Siamak Zohari (National Veterinary Institute) |
| EPI_ISL_832205 | A/Chicken/Sweden/SVA210111SZ0065/KN005298-IP1/2021 | A / H5N8 | Europe / Sweden / Skane Lan / Sjobo Kommun | Siamak Zohari (National Veterinary Institute) |
| EPI_ISL_832206 | A/Turkey/Sweden/SVA210114SZ0001/KN009555-IP2/2021 | A / H5N8 | Europe / Sweden / Skane Lan / Skurups Kommun | Siamak Zohari (National Veterinary Institute) |
| EPI_ISL_833248 | A/Muscovy duck/China/FJFZ21/H5N6/2020 | A / H5N6 | Asia / China / Fujian Province | Rui Zhang (Fujian Agriculture and Forestry University) |
| EPI_ISL_833482 | A/Goose/Hungary/18325/2020 | A / H5N8 | Europe / Hungary / Bacs-Kiskun | Katalin Szentpáli-Gavallér (National Food Chain Safety Office, Hungary / Virology) |
| EPI_ISL_833485 | A/Mallard_duck/Hungary/18410/2020 | A / H5N8 | Europe / Hungary / Csongrad megye | Katalin Szentpáli-Gavallér (National Food Chain Safety Office, Hungary / Virology) |
| EPI_ISL_833487 | A/Pheasant/Hungary/18731/2020 | A / H5N8 | Europe / Hungary / Bacs-Kiskun | Katalin Szentpáli-Gavallér (National Food Chain Safety Office, Hungary / Virology) |
| EPI_ISL_833488 | A/Turkey/Hungary/19338/2020 | A / H5N8 | Europe / Hungary / Bacs-Kiskun | Katalin Szentpáli-Gavallér (National Food Chain Safety Office, Hungary / Virology) |
| EPI_ISL_833579 | A/Duck/Hungary/18414/2020 | A / H5N8 | Europe / Hungary / Csongrad megye | Katalin Szentpáli-Gavallér (National Food Chain Safety Office, Hungary / Virology) |
| EPI_ISL_833580 | A/Turkey/Hungary/19394/2020 | A / H5N8 | Europe / Hungary / Bacs-Kiskun | Katalin Szentpáli-Gavallér (National Food Chain Safety Office, Hungary / Virology) |
| EPI_ISL_833581 | A/Goose/Hungary/19118/2020 | A / H5N8 | Europe / Hungary / Bacs-Kiskun | Katalin Szentpáli-Gavallér (National Food Chain Safety Office, Hungary / Virology) |
| EPI_ISL_833582 | A/Goose/Hungary/19953/2020 | A / H5N8 | Europe / Hungary / Bacs-Kiskun | Katalin Szentpáli-Gavallér (National Food Chain Safety Office, Hungary / Virology) |
| EPI_ISL_833583 | A/Goose/Hungary/19959/2020 | A / H5N8 | Europe / Hungary / Csongrad megye | Katalin Szentpáli-Gavallér (National Food Chain Safety Office, Hungary / Virology) |
| EPI_ISL_8338002 | A/Turkey/Sweden/SVA211212SZ0001/FB301013-IP-2/M-2021 | A / H5N1 | Europe / Sweden / Skane Lan / Skurups Kommun | Siamak Zohari (National Veterinary Institute) |
| EPI_ISL_8338456 | A/mute swan/Croatia/144/2021 | A / H5N1 | Europe / Croatia / Istarska Zupanija / the mouth of the river Mirna (45° 18´ 55?N, 13° 35´ 28?E) | Vladimir Savi? (Croatian Veterinary Institute / Poultry Centre) |
| EPI_ISL_8338502 | A/mute swan/Croatia/145/2021 | A / H5N1 | Europe / Croatia / Zagrebacka Zupanija / Rugvica - Siromaja lakes (45° 45´ 54?N, 16° 10´ 11?E) | Vladimir Savi? (Croatian Veterinary Institute / Poultry Centre) |
| EPI_ISL_8377056 | A/pelican/France/21P013720/2021 | A / H5N1 | Europe / France / Languedoc-Roussillon / Departement de l'Aude | Francois-Xavier Briand (ANSES Agence Nationale De Securite Sanitaire De L'alimentation / Laboratoire de Ploufragan-Plouzané) |
| EPI_ISL_8377254 | A/egret/France/21P013418/2021 | A / H5N1 | Europe / France / Rhone-Alpes / Departement de l'Ain | Francois-Xavier Briand (ANSES Agence Nationale De Securite Sanitaire De L'alimentation / Laboratoire de Ploufragan-Plouzané) |
| EPI_ISL_8377417 | A/chicken/France/21P013076/2021 | A / H5N1 | Europe / France / Region Nord-Pas-De Calais / Departement du Nord | Francois-Xavier Briand (ANSES Agence Nationale De Securite Sanitaire De L'alimentation / Laboratoire de Ploufragan-Plouzané) |
| EPI_ISL_8377418 | A/goose/France/21P013228/2021 | A / H5N1 | Europe / France / Brittany / Departement du Morbihan | Francois-Xavier Briand (ANSES Agence Nationale De Securite Sanitaire De L'alimentation / Laboratoire de Ploufragan-Plouzané) |
| EPI_ISL_8416806 | A/turkey/Lisbon/1/2021 | A / H5N1 | Europe / Portugal / Distrito de Setubal | Aryse Martins Melo (Instituto Nacional de Saúde Dr. Ricardo Jorge) |
| EPI_ISL_8439476 | A/mute swan/Croatia/146/2021 | A / H5N1 | Europe / Croatia / Vukovarsko-Srijemska Zupanija / Lipovac (45.06517, 19.07613) | Vladimir Savi? (Croatian Veterinary Institute / Poultry Centre) |
| EPI_ISL_8440175 | A/Cygnus_olor/Romania/16381_21VIR10306/2021 | A / H5N1 | Europe / Romania | Bianca Zecchin (Istituto Zooprofilattico Sperimentale Delle Venezie) |
| EPI_ISL_8440976 | A/mute_swan/Romania/16790_21VIR11355/2021 | A / H5N1 | Europe / Romania | Bianca Zecchin (Istituto Zooprofilattico Sperimentale Delle Venezie) |
| EPI_ISL_846583 | A/Chicken/Hungary/21379/2020 | A / H5N8 | Europe / Hungary / Bacs-Kiskun | Katalin Szentpáli-Gavallér (National Food Chain Safety Office, Hungary / Virology) |
| EPI_ISL_846584 | A/Goose/Hungary/19128/2020 | A / H5N8 | Europe / Hungary / Bacs-Kiskun | Katalin Szentpáli-Gavallér (National Food Chain Safety Office, Hungary / Virology) |
| EPI_ISL_846585 | A/Chicken/Hungary/24596/2020 | A / H5N8 | Europe / Hungary / Bacs-Kiskun | Katalin Szentpáli-Gavallér (National Food Chain Safety Office, Hungary / Virology) |
| EPI_ISL_846586 | A/Chicken/Hungary/29723/2020 | A / H5N8 | Europe / Hungary / Bacs-Kiskun | Katalin Szentpáli-Gavallér (National Food Chain Safety Office, Hungary / Virology) |
| EPI_ISL_846601 | A/chicken/Poland/474/2020(H5N8) | A / H5N8 | Europe / Poland / pomorskie | Edyta ?wi?to? (National Veterinary Research Institut Poland, PIWet-PIB) |
| EPI_ISL_846602 | A/turkey/Poland/475/2020(H5N8) | A / H5N8 | Europe / Poland / wielkopolskie | Edyta ?wi?to? (National Veterinary Research Institut Poland, PIWet-PIB) |
| EPI_ISL_846603 | A/chicken/Poland/476/2020(H5N8) | A / H5N8 | Europe / Poland / wielkopolskie | Edyta ?wi?to? (National Veterinary Research Institut Poland, PIWet-PIB) |
| EPI_ISL_846604 | A/turkey/Poland/477/2020(H5N8) | A / H5N8 | Europe / Poland / Masovian Voivodeship / mazowieckie | Edyta ?wi?to? (National Veterinary Research Institut Poland, PIWet-PIB) |
| EPI_ISL_846614 | A/turkey/Poland/542/2020(H5N8) | A / H5N8 | Europe / Poland / warminsko-mazurskie | Edyta ?wi?to? (National Veterinary Research Institut Poland, PIWet-PIB) |

| | | | | |
|---|---|---|---|---|
| EPI_ISL_846617 | A/chicken/Poland/565/2020(H5N8) | A / H5N8 | Europe / Poland / dolnoslaskie | Edyta ?wi?to? (National Veterinary Research Institut Poland, PIWet-PIB) |
| EPI_ISL_846623 | A/swan/Poland/MB141/2020(H5N8) | A / H5N8 | Europe / Poland / Pomeranian Voivodeship / pomorskie | Edyta ?wi?to? (National Veterinary Research Institut Poland, PIWet-PIB) |
| EPI_ISL_846624 | A/wild_goose/Poland/MB142/2020(H5N8) | A / H5N8 | Europe / Poland / pomorskie | Edyta ?wi?to? (National Veterinary Research Institut Poland, PIWet-PIB) |
| EPI_ISL_846625 | A/tundra_bean_goose/Poland/MB128/2020(H5N8) | A / H5N8 | Europe / Poland / wielkopolskie | Edyta ?wi?to? (National Veterinary Research Institut Poland, PIWet-PIB) |
| EPI_ISL_8515478 | A/goose/Czech_Republic/22608-1/2021 | A / H5N1 | Europe / Czech Republic / Jihocesky Kraj / Okres Ceske Budejovice / Nové Hrady, By?ov, ZipCode | Alexander Nagy (State Veterinary Institute Prague) |
| EPI_ISL_8515479 | A/goose/Czech_Republic/22608-2/2021 | A / H5N1 | Europe / Czech Republic / Jihocesky Kraj / Okres Ceske Budejovice / Nové Hrady, By?ov, ZipCode | Alexander Nagy (State Veterinary Institute Prague) |
| EPI_ISL_8515480 | A/chicken/Czech_Republic/23404/2021 | A / H5N1 | Europe / Czech Republic / South Moravian Region / Okres Breclav / Lednice, ZipCode: 69103 | Alexander Nagy (State Veterinary Institute Prague) |
| EPI_ISL_8515481 | A/grey heron/Czech_Republic/23608/2021 | A / H5N1 | Europe / Czech Republic / Liberecky Kraj / Okres Ceska Lipa / He?mani?ky u Dobranova, GPS:50.( | Alexander Nagy (State Veterinary Institute Prague) |
| EPI_ISL_8515482 | A/grey heron/Czech_Republic/23608-1K/2021 | A / H5N1 | Europe / Czech Republic / Liberecky Kraj / Okres Ceska Lipa / He?mani?ky u Dobranova, GPS: 50. | Alexander Nagy (State Veterinary Institute Prague) |
| EPI_ISL_8515483 | A/great egret/Czech_Republic/23609/2021 | A / H5N1 | Europe / Czech Republic / Liberecky Kraj / Okres Ceska Lipa / He?mani?ky u Dobranova, GPS:50.( | Alexander Nagy (State Veterinary Institute Prague) |
| EPI_ISL_8517996 | A/Environment/Hangzhou/E1147/2021 | A / H5N6 | Asia / China / Zhejiang | Jun Li (Hangzhou Center for Disease Control and Prevention ) |
| EPI_ISL_8517997 | A/Environment/Hangzhou/E1150/2021 | A / H5N6 | Asia / China / Zhejiang | Jun Li (Hangzhou Center for Disease Control and Prevention ) |
| EPI_ISL_8517998 | A/Environment/Hangzhou/E1151/2021 | A / H5N6 | Asia / China / Zhejiang | Jun Li (Hangzhou Center for Disease Control and Prevention ) |
| EPI_ISL_8517999 | A/Environment/Hangzhou/E1153/2021 | A / H5N6 | Asia / China / Zhejiang | Jun Li (Hangzhou Center for Disease Control and Prevention ) |
| EPI_ISL_8518000 | A/Environment/Hangzhou/E1154/2021 | A / H5N6 | Asia / China / Zhejiang | Jun Li (Hangzhou Center for Disease Control and Prevention ) |
| EPI_ISL_8518268 | A/Chicken/Hangzhou/E1149/2021 | A / H5N6 | Asia / China / Zhejiang | Jun Li (Hangzhou Center for Disease Control and Prevention ) |
| EPI_ISL_854595 | A/chicken/Chiba/B2T/2021 | A / H5N8 | Asia / Japan / Chiba | Takehiko Saito (National Institute of Animal Health) |
| EPI_ISL_854596 | A/chicken/Chiba/B3T/2021 | A / H5N8 | Asia / Japan / Chiba | Takehiko Saito (National Institute of Animal Health) |
| EPI_ISL_854597 | A/chicken/Chiba/B4T/2021 | A / H5N8 | Asia / Japan / Chiba | Takehiko Saito (National Institute of Animal Health) |
| EPI_ISL_854598 | A/chicken/Chiba/B5T/2021 | A / H5N8 | Asia / Japan / Chiba | Takehiko Saito (National Institute of Animal Health) |
| EPI_ISL_854599 | A/chicken/Gifu/10T/2021 | A / H5N8 | Asia / Japan / Gifu | Takehiko Saito (National Institute of Animal Health) |
| EPI_ISL_854600 | A/chicken/Gifu/11T/2021 | A / H5N8 | Asia / Japan / Gifu | Takehiko Saito (National Institute of Animal Health) |
| EPI_ISL_854601 | A/chicken/Gifu/2T/2021 | A / H5N8 | Asia / Japan / Gifu | Takehiko Saito (National Institute of Animal Health) |
| EPI_ISL_854602 | A/chicken/Gifu/3T/2021 | A / H5N8 | Asia / Japan / Gifu | Takehiko Saito (National Institute of Animal Health) |
| EPI_ISL_854603 | A/chicken/Gifu/6T/2021 | A / H5N8 | Asia / Japan / Gifu | Takehiko Saito (National Institute of Animal Health) |
| EPI_ISL_854604 | A/chicken/Gifu/7T/2021 | A / H5N8 | Asia / Japan / Gifu | Takehiko Saito (National Institute of Animal Health) |
| EPI_ISL_854605 | A/chicken/Kagoshima/1T/2021 | A / H5N8 | Asia / Japan / Kagoshima | Takehiko Saito (National Institute of Animal Health) |
| EPI_ISL_854606 | A/chicken/Kagoshima/3T/2021 | A / H5N8 | Asia / Japan / Kagoshima | Takehiko Saito (National Institute of Animal Health) |
| EPI_ISL_854607 | A/chicken/Kagoshima/5T/2021 | A / H5N8 | Asia / Japan / Kagoshima | Takehiko Saito (National Institute of Animal Health) |
| EPI_ISL_854608 | A/chicken/Kagoshima/6T/2021 | A / H5N8 | Asia / Japan / Kagoshima | Takehiko Saito (National Institute of Animal Health) |
| EPI_ISL_854609 | A/chicken/Miyazaki/I11T/2020 | A / H5N8 | Asia / Japan / Miyazaki | Takehiko Saito (National Institute of Animal Health) |
| EPI_ISL_854610 | A/chicken/Miyazaki/I12T/2020 | A / H5N8 | Asia / Japan / Miyazaki | Takehiko Saito (National Institute of Animal Health) |
| EPI_ISL_854611 | A/chicken/Miyazaki/I5T/2020 | A / H5N8 | Asia / Japan / Miyazaki | Takehiko Saito (National Institute of Animal Health) |
| EPI_ISL_854612 | A/chicken/Miyazaki/I9T/2020 | A / H5N8 | Asia / Japan / Miyazaki | Takehiko Saito (National Institute of Animal Health) |
| EPI_ISL_8568483 | A/gadwall/Croatia/108/2021 | A / H5N1 | Europe / Croatia / Splitsko-Dalmatinska Zupanija / Zaostrog, Podaca cove | Vladimir Savi? (Croatian Veterinary Institute / Poultry Centre) |
| EPI_ISL_8650947 | A/red fox/Netherlands/22000097-004/2022 | A / H5N1 | Europe / Netherlands / Provincie Noord-Holland | Rene Heutink (Wageningen Bioveterinary Research) |
| EPI_ISL_8650949 | A/red knot/Netherlands/21041030-008/2021 | A / H5N1 | Europe / Netherlands / Provincie Friesland | Rene Heutink (Wageningen Bioveterinary Research) |
| EPI_ISL_8694562 | A/mute swan/Croatia/6/2022 | A / H5N1 | Europe / Croatia / Karlovacka Zupanija / Karlovac | Vladimir Savi? (Croatian Veterinary Institute / Poultry Centre) |
| EPI_ISL_8694715 | A/chicken/Croatia/7/2022 | A / H5N1 | Europe / Croatia / Osjecko-Baranjska Zupanija / Branjin vrh | Vladimir Savi? (Croatian Veterinary Institute / Poultry Centre) |
| EPI_ISL_8768942 | A/turkey/Tyumen/15-1V/2021 | A / H5N1 | Europe / Russian Federation / Tyumen Oblast / Tyumen | Natalia Goncharova (State Research Center of Virology and Biotechnology (VECTOR) / Emerging Zoonotic Diseases and Influenza) |
| EPI_ISL_8768944 | A/turkey/Tyumen/15-2V/2021 | A / H5N1 | Europe / Russian Federation / Tyumen Oblast / Tyumen | Natalia Goncharova (State Research Center of Virology and Biotechnology (VECTOR) / Emerging Zoonotic Diseases and Influenza) |
| EPI_ISL_8768947 | A/turkey/Tyumen/15-3V/2021 | A / H5N1 | Europe / Russian Federation / Tyumen Oblast / Tyumen | Natalia Goncharova (State Research Center of Virology and Biotechnology (VECTOR) / Emerging Zoonotic Diseases and Influenza) |
| EPI_ISL_8768948 | A/turkey/Tyumen/15-4V/2021 | A / H5N1 | Europe / Russian Federation / Tyumen Oblast / Tyumen | Natalia Goncharova (State Research Center of Virology and Biotechnology (VECTOR) / Emerging Zoonotic Diseases and Influenza) |
| EPI_ISL_8768949 | A/turkey/Tyumen/15-6V/2021 | A / H5N1 | Europe / Russian Federation / Tyumen Oblast / Tyumen | Natalia Goncharova (State Research Center of Virology and Biotechnology (VECTOR) / Emerging Zoonotic Diseases and Influenza) |
| EPI_ISL_8768950 | A/turkey/Tyumen/15-9V/2021 | A / H5N1 | Europe / Russian Federation / Tyumen Oblast / Tyumen | Natalia Goncharova (State Research Center of Virology and Biotechnology (VECTOR) / Emerging Zoonotic Diseases and Influenza) |
| EPI_ISL_8768953 | A/turkey/Tyumen/15-14V/2021 | A / H5N1 | Europe / Russian Federation / Tyumen Oblast / Tyumen | Natalia Goncharova (State Research Center of Virology and Biotechnology (VECTOR) / Emerging Zoonotic Diseases and Influenza) |
| EPI_ISL_8768954 | A/duck/Omsk/19-1V/2021 | A / H5N1 | Europe / Russian Federation / Omsk Oblast / Omsk | Natalia Goncharova (State Research Center of Virology and Biotechnology (VECTOR) / Emerging Zoonotic Diseases and Influenza) |
| EPI_ISL_8768955 | A/duck/Omsk/19-2V/2021 | A / H5N1 | Europe / Russian Federation / Omsk Oblast / Omsk | Natalia Goncharova (State Research Center of Virology and Biotechnology (VECTOR) / Emerging Zoonotic Diseases and Influenza) |
| EPI_ISL_8768965 | A/duck/Omsk/19-3V/2021 | A / H5N1 | Europe / Russian Federation / Omsk Oblast / Omsk | Natalia Goncharova (State Research Center of Virology and Biotechnology (VECTOR) / Emerging Zoonotic Diseases and Influenza) |
| EPI_ISL_8768966 | A/duck/Omsk/19-5V/2021 | A / H5N1 | Europe / Russian Federation / Omsk Oblast / Omsk | Natalia Goncharova (State Research Center of Virology and Biotechnology (VECTOR) / Emerging Zoonotic Diseases and Influenza) |
| EPI_ISL_8768986 | A/goose/Rostov-on-Don/28-1V/2021 | A / H5N1 | Europe / Russian Federation / Rostov Oblast / Rostov-on-Don | Natalia Goncharova (State Research Center of Virology and Biotechnology (VECTOR) / Emerging Zoonotic Diseases and Influenza) |
| EPI_ISL_8768989 | A/duck/Saratov/29-2V/2021 | A / H5N1 | Europe / Russian Federation / Saratov Oblast / Saratov | Natalia Goncharova (State Research Center of Virology and Biotechnology (VECTOR) / Emerging Zoonotic Diseases and Influenza) |
| EPI_ISL_8769004 | A/chicken/Saratov/29-6V/2021 | A / H5N1 | Europe / Russian Federation / Saratov Oblast / Saratov | Natalia Goncharova (State Research Center of Virology and Biotechnology (VECTOR) / Emerging Zoonotic Diseases and Influenza) |
| EPI_ISL_8769010 | A/goose/Chelyabinsk/34-1V/2021 | A / H5N1 | Europe / Russian Federation / Chelyabinsk Oblast / Chelyabinsk | Natalia Goncharova (State Research Center of Virology and Biotechnology (VECTOR) / Emerging Zoonotic Diseases and Influenza) |
| EPI_ISL_8769012 | A/chicken/Orenburg/46-1V/2021 | A / H5N1 | Europe / Russian Federation / Orenburg Oblast / Orenburg | Natalia Goncharova (State Research Center of Virology and Biotechnology (VECTOR) / Emerging Zoonotic Diseases and Influenza) |
| EPI_ISL_8769014 | A/chicken/Orenburg/46-2V/2021 | A / H5N1 | Europe / Russian Federation / Orenburg Oblast / Orenburg | Natalia Goncharova (State Research Center of Virology and Biotechnology (VECTOR) / Emerging Zoonotic Diseases and Influenza) |
| EPI_ISL_8769016 | A/goose/Orenburg/46-3V/2021 | A / H5N1 | Europe / Russian Federation / Orenburg Oblast / Orenburg | Natalia Goncharova (State Research Center of Virology and Biotechnology (VECTOR) / Emerging Zoonotic Diseases and Influenza) |
| EPI_ISL_8769017 | A/chicken/Tyumen/47-66V/2021 | A / H5N1 | Europe / Russian Federation / Tyumen Oblast / Tyumen | Natalia Goncharova (State Research Center of Virology and Biotechnology (VECTOR) / Emerging Zoonotic Diseases and Influenza) |
| EPI_ISL_8769018 | A/chicken/Tyumen/47-79V/2021 | A / H5N1 | Europe / Russian Federation / Tyumen Oblast / Tyumen | Natalia Goncharova (State Research Center of Virology and Biotechnology (VECTOR) / Emerging Zoonotic Diseases and Influenza) |
| EPI_ISL_8769019 | A/chicken/Tyumen/47-85V/2021 | A / H5N1 | Europe / Russian Federation / Tyumen Oblast / Tyumen | Natalia Goncharova (State Research Center of Virology and Biotechnology (VECTOR) / Emerging Zoonotic Diseases and Influenza) |
| EPI_ISL_8769021 | A/chicken/Tyumen/47-88V/2021 | A / H5N1 | Europe / Russian Federation / Tyumen Oblast / Tyumen | Natalia Goncharova (State Research Center of Virology and Biotechnology (VECTOR) / Emerging Zoonotic Diseases and Influenza) |
| EPI_ISL_8769022 | A/chicken/Tyumen/47-95V/2021 | A / H5N1 | Europe / Russian Federation / Tyumen Oblast / Tyumen | Natalia Goncharova (State Research Center of Virology and Biotechnology (VECTOR) / Emerging Zoonotic Diseases and Influenza) |
| EPI_ISL_8769023 | A/chicken/Kirov/63-1V/2021 | A / H5N1 | Europe / Russian Federation / Kirov Oblast / Kirov | Natalia Goncharova (State Research Center of Virology and Biotechnology (VECTOR) / Emerging Zoonotic Diseases and Influenza) |
| EPI_ISL_8769024 | A/chicken/Kirov/63-2V/2021 | A / H5N1 | Europe / Russian Federation / Kirov Oblast / Kirov | Natalia Goncharova (State Research Center of Virology and Biotechnology (VECTOR) / Emerging Zoonotic Diseases and Influenza) |
| EPI_ISL_8769025 | A/chicken/Kirov/63-5V/2021 | A / H5N1 | Europe / Russian Federation / Kirov Oblast / Kirov | Natalia Goncharova (State Research Center of Virology and Biotechnology (VECTOR) / Emerging Zoonotic Diseases and Influenza) |
| EPI_ISL_8769026 | A/chicken/Kurgan/72-1V/2021 | A / H5N1 | Europe / Russian Federation / Kurgan Oblast / Kurgan | Natalia Goncharova (State Research Center of Virology and Biotechnology (VECTOR) / Emerging Zoonotic Diseases and Influenza) |
| EPI_ISL_8769028 | A/chicken/Orenburg/73-5V/2021 | A / H5N1 | Europe / Russian Federation / Orenburg Oblast / Orenburg | Natalia Goncharova (State Research Center of Virology and Biotechnology (VECTOR) / Emerging Zoonotic Diseases and Influenza) |
| EPI_ISL_8769029 | A/chicken/Orenburg/73-6V/2021 | A / H5N1 | Europe / Russian Federation / Orenburg Oblast / Orenburg | Natalia Goncharova (State Research Center of Virology and Biotechnology (VECTOR) / Emerging Zoonotic Diseases and Influenza) |
| EPI_ISL_8769030 | A/chicken/Orenburg/73-7V/2021 | A / H5N1 | Europe / Russian Federation / Orenburg Oblast / Orenburg | Natalia Goncharova (State Research Center of Virology and Biotechnology (VECTOR) / Emerging Zoonotic Diseases and Influenza) |
| EPI_ISL_8769031 | A/black-necked grebe/Kalmykia/78-1V/2021 | A / H5N5 | Europe / Russian Federation / Republic of Kalmykia | Natalia Goncharova (State Research Center of Virology and Biotechnology (VECTOR) / Emerging Zoonotic Diseases and Influenza) |
| EPI_ISL_8769033 | A/turkey/Tyumen/81-96V/2021 | A / H5N1 | Europe / Russian Federation / Tyumen Oblast / Tyumen | Natalia Goncharova (State Research Center of Virology and Biotechnology (VECTOR) / Emerging Zoonotic Diseases and Influenza) |
| EPI_ISL_8769034 | A/chicken/Tyumen/81-97V/2021 | A / H5N1 | Europe / Russian Federation / Tyumen Oblast / Tyumen | Natalia Goncharova (State Research Center of Virology and Biotechnology (VECTOR) / Emerging Zoonotic Diseases and Influenza) |

| | | | | |
|---|---|---|---|---|
| EPI_ISL_8769036 | A/chicken/Saratov/102-12V/2021 | A / H5N1 | Europe / Russian Federation / Saratov Oblast / Saratov | Natalia Goncharova (State Research Center of Virology and Biotechnology (VECTOR) / Emerging Zoonotic Diseases and Influenza) |
| EPI_ISL_8769038 | A/chicken/Saratov/102-15V/2021 | A / H5N1 | Europe / Russian Federation / Saratov Oblast / Saratov | Natalia Goncharova (State Research Center of Virology and Biotechnology (VECTOR) / Emerging Zoonotic Diseases and Influenza) |
| EPI_ISL_8799314 | A/red knot/Netherlands/21040956-003/2021 | A / H5N1 | Europe / Netherlands / Provincie Groningen | Rene Heutink (Wageningen Bioveterinary Research) |
| EPI_ISL_8799317 | A/seagull/Netherlands/21041370-001/2021 | A / H5N1 | Europe / Netherlands / Provincie Gelderland | Rene Heutink (Wageningen Bioveterinary Research) |
| EPI_ISL_8799318 | A/common buzzard/Netherlands/21041449-002/2021 | A / H5N1 | Europe / Netherlands / Provincie Zeeland | Rene Heutink (Wageningen Bioveterinary Research) |
| EPI_ISL_8799320 | A/polecat/Netherlands/22000516-003/2022 | A / H5N1 | Europe / Netherlands / South Holland | Rene Heutink (Wageningen Bioveterinary Research) |
| EPI_ISL_8799552 | A/England/215201407/2021 | A / H5N1 | Europe / United Kingdom | Steven Platt (UK Health Security Agency (UKHSA)) |
| EPI_ISL_8809153 | A/Muscovy duck/England/074477/2021 | A / H5N1 | Europe / United Kingdom / Devon | Alex Byrne (Animal and Plant Health Agency (APHA) / Virology Department) |
| EPI_ISL_8814146 | A/turkey/England/055251/2021 | A / H5N1 | Europe / United Kingdom / Warwickshire / England | Alex Byrne (Animal and Plant Health Agency (APHA) / Virology Department) |
| EPI_ISL_8814195 | A/turkey/England/056764/2021 | A / H5N1 | Europe / United Kingdom / Lancashire / England | Alex Byrne (Animal and Plant Health Agency (APHA) / Virology Department) |
| EPI_ISL_885144 | A/environment/Hong Kong/WCRB-01/2018 | A / H5N6 | Asia / Hong Kong (SAR) | Huyen Trang Bui (The University of Hong Kong) |
| EPI_ISL_885145 | A/Spoonbill/HK/17-18259/2017 | A / H5N6 | Asia / Hong Kong (SAR) / Mudflat of HK Wetland Park | Huyen Trang Bui (The University of Hong Kong) |
| EPI_ISL_885147 | A/Northern Pintail/HK/MP692/2016 | A / H5N6 | Asia / Hong Kong (SAR) / Mai Po Tower Hide | Huyen Trang Bui (The University of Hong Kong) |
| EPI_ISL_885148 | A/Chicken/Egypt/F1366A/2017 | A / H5N8 | Africa / Egypt | Huyen Trang Bui (The University of Hong Kong) |
| EPI_ISL_885149 | A/Grey-Headed Gull/Uganda/200144/2017 | A / H5N8 | Africa / Uganda | Huyen Trang Bui (The University of Hong Kong) |
| EPI_ISL_8882166 | A/turkey/Italy/21VIR11586-2/2021 | A / H5N1 | Europe / Italy | Giacomo Barbierato (Istituto Zooprofilattico Sperimentale delle Venezie / Ricerca e innovazione) |
| EPI_ISL_8882167 | A/turkey/Italy/21VIR11804-1/2021 | A / H5N1 | Europe / Italy | Giacomo Barbierato (Istituto Zooprofilattico Sperimentale delle Venezie / Ricerca e innovazione) |
| EPI_ISL_8882168 | A/chicken/Italy/21VIR10850/2021 | A / H5N1 | Europe / Italy | Giacomo Barbierato (Istituto Zooprofilattico Sperimentale delle Venezie / Ricerca e innovazione) |
| EPI_ISL_8882169 | A/chicken/Italy/21VIR10573-1/2021 | A / H5N1 | Europe / Italy | Giacomo Barbierato (Istituto Zooprofilattico Sperimentale delle Venezie / Ricerca e innovazione) |
| EPI_ISL_8882170 | A/turkey/Italy/21VIR11887-3/2021 | A / H5N1 | Europe / Italy | Giacomo Barbierato (Istituto Zooprofilattico Sperimentale delle Venezie / Ricerca e innovazione) |
| EPI_ISL_8882171 | A/turkey/Italy/21VIR11591-8/2021 | A / H5N1 | Europe / Italy | Giacomo Barbierato (Istituto Zooprofilattico Sperimentale delle Venezie / Ricerca e innovazione) |
| EPI_ISL_8882172 | A/turkey/Italy/21VIR10851/2021 | A / H5N1 | Europe / Italy | Giacomo Barbierato (Istituto Zooprofilattico Sperimentale delle Venezie / Ricerca e innovazione) |
| EPI_ISL_8882173 | A/laying_hen/Italy/21VIR11502/2021 | A / H5N1 | Europe / Italy | Giacomo Barbierato (Istituto Zooprofilattico Sperimentale delle Venezie / Ricerca e innovazione) |
| EPI_ISL_8882187 | A/turkey/Italy/21VIR11053-1/2021 | A / H5N1 | Europe / Italy | Giacomo Barbierato (Istituto Zooprofilattico Sperimentale delle Venezie / Ricerca e innovazione) |
| EPI_ISL_8882201 | A/turkey/Italy/21VIR11803-1/2021 | A / H5N1 | Europe / Italy | Giacomo Barbierato (Istituto Zooprofilattico Sperimentale delle Venezie / Ricerca e innovazione) |
| EPI_ISL_8882209 | A/little_owl/Italy/21VIR11382-1/2021 | A / H5N1 | Europe / Italy | Giacomo Barbierato (Istituto Zooprofilattico Sperimentale delle Venezie / Ricerca e innovazione) |
| EPI_ISL_8882210 | A/seagull/Italy/21VIR10481-2/2021 | A / H5N1 | Europe / Italy | Giacomo Barbierato (Istituto Zooprofilattico Sperimentale delle Venezie / Ricerca e innovazione) |
| EPI_ISL_8882211 | A/magpie/Italy/21VIR11656/2021 | A / H5N1 | Europe / Italy | Giacomo Barbierato (Istituto Zooprofilattico Sperimentale delle Venezie / Ricerca e innovazione) |
| EPI_ISL_8882212 | A/cignus_olor/Italy/IZSLT_21VIR10529-1/2021 | A / H5N1 | Europe / Italy | Giacomo Barbierato (Istituto Zooprofilattico Sperimentale delle Venezie / Ricerca e innovazione) |
| EPI_ISL_8882213 | A/seagull/Italy/21VIR11259-10/2021 | A / H5N1 | Europe / Italy | Giacomo Barbierato (Istituto Zooprofilattico Sperimentale delle Venezie / Ricerca e innovazione) |
| EPI_ISL_8882214 | A/heron/Italy/21VIR10998-1/2021 | A / H5N1 | Europe / Italy | Giacomo Barbierato (Istituto Zooprofilattico Sperimentale delle Venezie / Ricerca e innovazione) |
| EPI_ISL_8882215 | A/owl/Italy/21VIR11899-1/2021 | A / H5N1 | Europe / Italy | Giacomo Barbierato (Istituto Zooprofilattico Sperimentale delle Venezie / Ricerca e innovazione) |
| EPI_ISL_8882216 | A/buzzard/Italy/21VIR11899-5/2021 | A / H5N1 | Europe / Italy | Giacomo Barbierato (Istituto Zooprofilattico Sperimentale delle Venezie / Ricerca e innovazione) |
| EPI_ISL_8882217 | A/owl/Italy/21VIR11057/2021 | A / H5N1 | Europe / Italy | Giacomo Barbierato (Istituto Zooprofilattico Sperimentale delle Venezie / Ricerca e innovazione) |
| EPI_ISL_8882218 | A/seagull/Italy/21VIR11259-12/2021 | A / H5N1 | Europe / Italy | Giacomo Barbierato (Istituto Zooprofilattico Sperimentale delle Venezie / Ricerca e innovazione) |
| EPI_ISL_9009291 | A/chicken/Kursk/132-1V/2021 | A / H5N1 | Europe / Russian Federation / Kursk Oblast / Kursk | Natalia Goncharova (State Research Center of Virology and Biotechnology (VECTOR) / Emerging Zoonotic Diseases and Influenza) |
| EPI_ISL_9009294 | A/chicken/Stavropol/146-1V/2021 | A / H5N1 | Europe / Russian Federation / Stavropol Krai / Stavropol | Natalia Goncharova (State Research Center of Virology and Biotechnology (VECTOR) / Emerging Zoonotic Diseases and Influenza) |
| EPI_ISL_9009296 | A/chicken/Rostov-on-Don/159-1V/2021 | A / H5N1 | Europe / Russian Federation / Rostov Oblast / Rostov-on-Don | Natalia Goncharova (State Research Center of Virology and Biotechnology (VECTOR) / Emerging Zoonotic Diseases and Influenza) |
| EPI_ISL_9009297 | A/chicken/Rostov-on-Don/159-2V/2021 | A / H5N1 | Europe / Russian Federation / Rostov Oblast / Rostov-on-Don | Natalia Goncharova (State Research Center of Virology and Biotechnology (VECTOR) / Emerging Zoonotic Diseases and Influenza) |
| EPI_ISL_9009302 | A/chicken/Rostov-on-Don/159-3V/2021 | A / H5N1 | Europe / Russian Federation / Rostov Oblast / Rostov-on-Don | Natalia Goncharova (State Research Center of Virology and Biotechnology (VECTOR) / Emerging Zoonotic Diseases and Influenza) |
| EPI_ISL_9009303 | A/chicken/Rostov-on-Don/159-4V/2021 | A / H5N1 | Europe / Russian Federation / Rostov Oblast / Rostov-on-Don | Natalia Goncharova (State Research Center of Virology and Biotechnology (VECTOR) / Emerging Zoonotic Diseases and Influenza) |
| EPI_ISL_9009304 | A/chicken/Rostov-on-Don/159-5V/2021 | A / H5N1 | Europe / Russian Federation / Rostov Oblast / Rostov-on-Don | Natalia Goncharova (State Research Center of Virology and Biotechnology (VECTOR) / Emerging Zoonotic Diseases and Influenza) |
| EPI_ISL_9009309 | A/chicken/Rostov-on-Don/159-6V/2021 | A / H5N1 | Europe / Russian Federation / Rostov Oblast / Rostov-on-Don | Natalia Goncharova (State Research Center of Virology and Biotechnology (VECTOR) / Emerging Zoonotic Diseases and Influenza) |
| EPI_ISL_9009317 | A/chicken/Rostov-on-Don/159-7V/2021 | A / H5N1 | Europe / Russian Federation / Rostov Oblast / Rostov-on-Don | Natalia Goncharova (State Research Center of Virology and Biotechnology (VECTOR) / Emerging Zoonotic Diseases and Influenza) |
| EPI_ISL_9009472 | A/chicken/Quang Tri/V4S4VTC/2020 | A / H5N6 | Asia / Vietnam / Tinh Quang Tri | Natalia Goncharova (State Research Center of Virology and Biotechnology (VECTOR) / Emerging Zoonotic Diseases and Influenza) |
| EPI_ISL_9009479 | A/chicken/Thanh Hoa/V3S3VTC/2020 | A / H5N6 | Asia / Vietnam / Tinh Thanh Hoa | Natalia Goncharova (State Research Center of Virology and Biotechnology (VECTOR) / Emerging Zoonotic Diseases and Influenza) |
| EPI_ISL_9009481 | A/muscovy duck/Nghe An/6873VTC/2020 | A / H5N6 | Asia / Vietnam / Nhge An | Natalia Goncharova (State Research Center of Virology and Biotechnology (VECTOR) / Emerging Zoonotic Diseases and Influenza) |
| EPI_ISL_9009483 | A/chicken/Nghe An/7007VTC/2020 | A / H5N6 | Asia / Vietnam / Nhge An | Natalia Goncharova (State Research Center of Virology and Biotechnology (VECTOR) / Emerging Zoonotic Diseases and Influenza) |
| EPI_ISL_9009488 | A/muscovy duck/Nghe An/7006VTC/2020 | A / H5N6 | Asia / Vietnam / Nhge An | Natalia Goncharova (State Research Center of Virology and Biotechnology (VECTOR) / Emerging Zoonotic Diseases and Influenza) |
| EPI_ISL_9009499 | A/muscovy duck/Nghe An/259VTC/2021 | A / H5N6 | Asia / Vietnam / Nhge An | Natalia Goncharova (State Research Center of Virology and Biotechnology (VECTOR) / Emerging Zoonotic Diseases and Influenza) |
| EPI_ISL_9009515 | A/chicken/Thanh Hoa/1351VTC/2021 | A / H5N6 | Asia / Vietnam / Tinh Thanh Hoa / Tan Thanh | Natalia Goncharova (State Research Center of Virology and Biotechnology (VECTOR) / Emerging Zoonotic Diseases and Influenza) |
| EPI_ISL_9009531 | A/chicken/Thanh Hoa/6081VTC/2020 | A / H5N6 | Asia / Vietnam / Tinh Thanh Hoa / Quang Thanh | Natalia Goncharova (State Research Center of Virology and Biotechnology (VECTOR) / Emerging Zoonotic Diseases and Influenza) |
| EPI_ISL_9009546 | A/duck/Thanh Hoa/5331VTC/2020 | A / H5N6 | Asia / Vietnam / Tinh Thanh Hoa / Co Lung | Natalia Goncharova (State Research Center of Virology and Biotechnology (VECTOR) / Emerging Zoonotic Diseases and Influenza) |
| EPI_ISL_9009552 | A/chicken/Ha Tinh/514VTC/2021 | A / H5N6 | Asia / Vietnam / Nam Ha Tinh / Thach Van | Natalia Goncharova (State Research Center of Virology and Biotechnology (VECTOR) / Emerging Zoonotic Diseases and Influenza) |
| EPI_ISL_9009554 | A/duck/Thanh Hoa/4643VTC/2020 | A / H5N6 | Asia / Vietnam / Tinh Thanh Hoa / Tan Tho | Natalia Goncharova (State Research Center of Virology and Biotechnology (VECTOR) / Emerging Zoonotic Diseases and Influenza) |
| EPI_ISL_9012457 | A/chicken/England/053052/2021 | A / H5N1 | Europe / United Kingdom / Worcestershire / England | Alex Byrne (Animal and Plant Health Agency (APHA) / Virology Department) |
| EPI_ISL_9012572 | A/pheasant/Wales/385129/2021 | A / H5N1 | Europe / United Kingdom / County Borough of Wrexham / Wales | Alex Byrne (Animal and Plant Health Agency (APHA) / Virology Department) |
| EPI_ISL_9012574 | A/Herring Gull/Netherlands/1/2021 | A / H5N1 | Europe / Netherlands | Pascal Lexmond (Erasmus Medical Center / Viroscience) |
| EPI_ISL_9012577 | A/Mallard/Netherlands/13/2021 | A / H5N1 | Europe / Netherlands | Pascal Lexmond (Erasmus Medical Center / Viroscience) |
| EPI_ISL_9012578 | A/Mallard/Netherlands/14/2021 | A / H5N1 | Europe / Netherlands | Pascal Lexmond (Erasmus Medical Center / Viroscience) |
| EPI_ISL_9012580 | A/Mallard/Netherlands/15/2021 | A / H5N1 | Europe / Netherlands | Pascal Lexmond (Erasmus Medical Center / Viroscience) |
| EPI_ISL_9012618 | A/chicken/Wales/053969/2021 | A / H5N1 | Europe / United Kingdom / County Borough of Wrexham / Wales | Alex Byrne (Animal and Plant Health Agency (APHA) / Virology Department) |
| EPI_ISL_9012694 | A/guineafowl/Scotland/054471/2021 | A / H5N1 | Europe / United Kingdom / Angus / Scotland | Alex Byrne (Animal and Plant Health Agency (APHA) / Virology Department) |
| EPI_ISL_9012696 | A/chicken/Scotland/054477/2021 | A / H5N1 | Europe / United Kingdom / Angus / Scotland | Alex Byrne (Animal and Plant Health Agency (APHA) / Virology Department) |
| EPI_ISL_9012700 | A/domestic_duck/Scotland/054469/2021 | A / H5N1 | Europe / United Kingdom / Angus / Scotland | Alex Byrne (Animal and Plant Health Agency (APHA) / Virology Department) |
| EPI_ISL_9029956 | A/chicken/England/057314/2021 | A / H5N1 | Europe / United Kingdom / Lancashire / England | Alex Byrne (Animal and Plant Health Agency (APHA) / Virology Department) |
| EPI_ISL_9029957 | A/turkey/England/057679/2021 | A / H5N1 | Europe / United Kingdom / Derbyshire / England | Alex Byrne (Animal and Plant Health Agency (APHA) / Virology Department) |
| EPI_ISL_9029959 | A/domestic_duck/England/058612/2021 | A / H5N1 | Europe / United Kingdom / Dorset / England | Alex Byrne (Animal and Plant Health Agency (APHA) / Virology Department) |
| EPI_ISL_9029960 | A/mute_swan/England/385466/2021 | A / H5N1 | Europe / United Kingdom / Gloucestershire / England | Alex Byrne (Animal and Plant Health Agency (APHA) / Virology Department) |
| EPI_ISL_9029961 | A/Canada_goose/England/385250/2021 | A / H5N1 | Europe / United Kingdom / Stratford-upon-Avon | Alex Byrne (Animal and Plant Health Agency (APHA) / Virology Department) |
| EPI_ISL_9029962 | A/Whooper_swan/Scotland/056219/2021 | A / H5N1 | Europe / United Kingdom / Dumfries and Galloway / Scotland | Alex Byrne (Animal and Plant Health Agency (APHA) / Virology Department) |
| EPI_ISL_9029965 | A/Greylag_goose/England/054503/2021 | A / H5N1 | Europe / United Kingdom / Lancashire / England | Alex Byrne (Animal and Plant Health Agency (APHA) / Virology Department) |

| | | | | |
|---|---|---|---|---|
| EPI_ISL_9111095 | A/goose/Netherlands/22000487-001/2022 | A / H5N1 | Europe / Netherlands / Provincie Gelderland | Rene Heutink (Wageningen Bioveterinary Research) |
| EPI_ISL_9111104 | A/otter/Netherlands/22001014-005/2022 | A / H5N1 | Europe / Netherlands / Provincie Flevoland | Rene Heutink (Wageningen Bioveterinary Research) |
| EPI_ISL_9111105 | A/chicken/Netherlands/22001401-001005/2022 | A / H5N1 | Europe / Netherlands / Provincie Flevoland | Rene Heutink (Wageningen Bioveterinary Research) |
| EPI_ISL_9117228 | A/northern goshawk/Netherlands/22000305-002/2022 | A / H5N1 | Europe / Netherlands / Provincie Utrecht | Rene Heutink (Wageningen Bioveterinary Research) |
| EPI_ISL_9160206 | A/Anser_albifrons/Belgium/15465_0010/2021 | A / H5N1 | Europe / Belgium / Provincie West-Vlaanderen / Oudenburg | Steven Van Borm (Sciensano, Department of Animal Infectious Diseases / Animal Infectious Diseases) |
| EPI_ISL_9161618 | A/Gallus_gallus/Belgium/17100_0001/2021 | A / H5N1 | Europe / Belgium / Provincie West-Vlaanderen / Veurne | Steven Van Borm (Sciensano, Department of Animal Infectious Diseases / Animal Infectious Diseases) |
| EPI_ISL_9162331 | A/Phasianus_colchicus/Belgium/294/2022 | A / H5N1 | Europe / Belgium / Provincie Limburg / Bocholt | Steven Van Borm (Sciensano, Department of Animal Infectious Diseases / Animal Infectious Diseases) |
| EPI_ISL_9221411 | A/barnacle goose/Netherlands/21041450-001/2021 | A / H5N1 | Europe / Netherlands / South Holland | Rene Heutink (Wageningen Bioveterinary Research) |
| EPI_ISL_9221430 | A/common buzzard/Netherlands/21041451-002/2021 | A / H5N1 | Europe / Netherlands / South Holland | Rene Heutink (Wageningen Bioveterinary Research) |
| EPI_ISL_9221469 | A/lesser black-backed gull/Netherlands/21041570-001/2021 | A / H5N1 | Europe / Netherlands / Provincie Noord-Holland | Rene Heutink (Wageningen Bioveterinary Research) |
| EPI_ISL_9221519 | A/barnacle goose/Netherlands/22000419-002/2022 | A / H5N1 | Europe / Netherlands / Provincie Flevoland | Rene Heutink (Wageningen Bioveterinary Research) |
| EPI_ISL_9250718 | A/Gallus_gallus/Belgium/11372_0001/2021 | A / H5N8 | Europe / Belgium / Province de Luxembourg / Chassepierre | Steven Van Borm (Sciensano, Department of Animal Infectious Diseases / Animal Infectious Diseases) |
| EPI_ISL_9261740 | A/Mallard/Netherlands/17/2021 | A / H5N3 | Europe / Netherlands / Provincie Noord-Holland / Wieringen | Pascal Lexmond (Erasmus Medical Center / Viroscience) |
| EPI_ISL_9261741 | A/Barnacle goose/Netherlands/1/2022 | A / H5N1 | Europe / Netherlands / Provincie Gelderland / Stroe | Pascal Lexmond (Erasmus Medical Center / Viroscience) |
| EPI_ISL_9261742 | A/Great black-backed gull/1/2022 | A / H5N1 | Europe / Netherlands / Provincie Gelderland / Stroe | Pascal Lexmond (Erasmus Medical Center / Viroscience) |
| EPI_ISL_9261743 | A/Oystercatcher/Netherlands/1/2022 | A / H5N1 | Europe / Netherlands / Provincie Noord-Holland / Normerven | Pascal Lexmond (Erasmus Medical Center / Viroscience) |
| EPI_ISL_9261744 | A/Barnacle goose/Netherlands/2/2022 | A / H5N1 | Europe / Netherlands / Provincie Noord-Holland / Normerven | Pascal Lexmond (Erasmus Medical Center / Viroscience) |
| EPI_ISL_9261745 | A/Barnacle goose/Netherlands/3/2022 | A / H5N1 | Europe / Netherlands / Provincie Noord-Holland / Zuiderdijk | Pascal Lexmond (Erasmus Medical Center / Viroscience) |
| EPI_ISL_9261746 | A/Barnacle goose/Netherlands/4/2022 | A / H5N1 | Europe / Netherlands / Provincie Noord-Holland / Zuiderdijk | Pascal Lexmond (Erasmus Medical Center / Viroscience) |
| EPI_ISL_9261747 | A/Barnacle goose/Netherlands/5/2022 | A / H5N1 | Europe / Netherlands / Provincie Noord-Holland / De Kreupel | Pascal Lexmond (Erasmus Medical Center / Viroscience) |
| EPI_ISL_9261748 | A/Grey heron/Netherlands/1/2022 | A / H5N1 | Europe / Netherlands / Provincie Noord-Holland / De Kreupel | Pascal Lexmond (Erasmus Medical Center / Viroscience) |
| EPI_ISL_9261749 | A/Caspian gull/Netherlands/1/2022 | A / H5N1 | Europe / Netherlands / Provincie Noord-Holland / De Kreupel | Pascal Lexmond (Erasmus Medical Center / Viroscience) |
| EPI_ISL_9261750 | A/Black-headed gull/Netherlands/1/2022 | A / H5N1 | Europe / Netherlands / Provincie Noord-Holland / De Kreupel | Pascal Lexmond (Erasmus Medical Center / Viroscience) |
| EPI_ISL_9261751 | A/Barnacle goose/Netherlands/6/2022 | A / H5N1 | Europe / Netherlands / Provincie Noord-Holland / De Kreupel | Pascal Lexmond (Erasmus Medical Center / Viroscience) |
| EPI_ISL_9304992 | A/barnacle goose/Netherlands/22000551-002/2022 | A / H5N1 | Europe / Netherlands / Provincie Friesland | Rene Heutink (Wageningen Bioveterinary Research) |
| EPI_ISL_9377021 | A/goose/France/21P014207/2021 | A / H5N1 | Europe / France / Midi-Pyrenees / Departement des Hautes-Pyrenees | Francois-Xavier Briand (ANSES Agence Nationale De Securite Sanitaire De L'alimentation / Laboratoire de Ploufragan-Plouzané) |
| EPI_ISL_9401912 | A/white-fronted Goose/Croatia/16/2022 | A / H5N1 | Europe / Croatia / Osjecko-Baranjska Zupanija | Vladimir Savi? (Croatian Veterinary Institute / Poultry Centre) |
| EPI_ISL_94329 | A/duck/Eastern China/008/2008 | A / H5N5 | Asia / China | |
| EPI_ISL_94330 | A/duck/Eastern China/031/2009 | A / H5N5 | Asia / China | |
| EPI_ISL_94331 | A/duck/Eastern China/108/2008 | A / H5N1 | Asia / China | |
| EPI_ISL_943551 | A/Chicken/Sweden/SVA210121SZ0033/KN000273-IP3/2021 | A / H5N5 | Europe / Sweden / Kalmar Lan / Monsteras Kommun | Siamak Zohari (National Veterinary Institute) |
| EPI_ISL_943553 | A/Chicken/Sweden/SVA210117SZ0004/KN011326-IP3/2021 | A / H5N5 | Europe / Sweden / Kalmar Lan / Monsteras Kommun | Siamak Zohari (National Veterinary Institute) |
| EPI_ISL_9439410 | A/grey heron/Netherlands/22000049-002/2021 | A / H5N1 | Europe / Netherlands / Provincie Noord-Holland | Rene Heutink (Wageningen Bioveterinary Research) |
| EPI_ISL_956368 | A/mute swan/Czech Republic/1410-2/2021 | A / H5N8 | Europe / Czech Republic / Jihocesky Kraj / Okres Pisek / Zlivicky pond; GPS 49°21'26.626''N 14°6' | Alexander Nagy (State Veterinary Institute Prague) |
| EPI_ISL_956408 | A/common_teal/Italy/20VIR7608-73/2020 | A / H5N8 | Europe / Italy / Venezia | Bianca Zecchin (Istituto Zooprofilattico Sperimentale Delle Venezie) |
| EPI_ISL_956409 | A/common_teal/Italy/20VIR7439-190/2020 | A / H5N5 | Europe / Italy / Rovigo | Bianca Zecchin (Istituto Zooprofilattico Sperimentale Delle Venezie) |
| EPI_ISL_956411 | A/greylag_goose/Italy/20VIR7660-6/2020 | A / H5N8 | Europe / Italy / Venezia | Bianca Zecchin (Istituto Zooprofilattico Sperimentale Delle Venezie) |
| EPI_ISL_956412 | A/greater_white-fronted_goose/Italy/20VIR8073-4/2020 | A / H5N1 | Europe / Italy / Gorizia | Bianca Zecchin (Istituto Zooprofilattico Sperimentale Delle Venezie) |
| EPI_ISL_956414 | A/Eurasian_wigeon/Italy/20VIR7301-362/2020 | A / H5N8 | Europe / Italy / Venezia | Bianca Zecchin (Istituto Zooprofilattico Sperimentale Delle Venezie) |
| EPI_ISL_9572654 | A/duck/Nigeria/VRD19-068DC019T_19RS1081-12/2019 | A / H5N8 | Africa / Nigeria | |
| EPI_ISL_9572655 | A/duck/Nigeria/VRD19-068DC004T_19RS1081-10/2019 | A / H5N8 | Africa / Nigeria | |
| EPI_ISL_9572656 | A/chicken/Nigeria/VRD-19-023_19RS1081-1/2019 | A / H5N8 | Africa / Nigeria | |
| EPI_ISL_9572657 | A/chicken/Nigeria/VRD-19-055_19RS1081-3/2019 | A / H5N8 | Africa / Nigeria | |
| EPI_ISL_9572658 | A/chicken/Nigeria/VRD-19-069_19RS1081-5/2019 | A / H5N8 | Africa / Nigeria | |
| EPI_ISL_9572659 | A/chicken/Nigeria/VRD-19-073_19RS1081-6/2019 | A / H5N8 | Africa / Nigeria | |
| EPI_ISL_9572672 | A/chicken/Nigeria/VRD-18-NS10_18RS1971-29/2018 | A / H5N8 | Africa / Nigeria | |
| EPI_ISL_9572689 | A/chicken/Egypt/FN1/2020 | A / H5 | Africa / Egypt | |
| EPI_ISL_9572690 | A/chicken/Egypt/FN2/2020 | A / H5 | Africa / Egypt | |
| EPI_ISL_9572691 | A/chicken/Egypt/FN3/2020 | A / H5 | Africa / Egypt | |
| EPI_ISL_9572739 | A/Muscovy duck/Vietnam/HN6610/2020 | A / H5N6 | Asia / Vietnam | |
| EPI_ISL_9572741 | A/duck/Vietnam/HN6611/2020 | A / H5N6 | Asia / Vietnam | |
| EPI_ISL_9572745 | A/Muscovy duck/Vietnam/HN6609/2020 | A / H5N6 | Asia / Vietnam | |
| EPI_ISL_9572754 | A/Muscovy duck/Vietnam/HN6606/2020 | A / H5N6 | Asia / Vietnam | |
| EPI_ISL_9572759 | A/Muscovy duck/Vietnam/HN6607/2020 | A / H5N6 | Asia / Vietnam | |
| EPI_ISL_9572770 | A/Muscovy duck/Vietnam/HN6608/2020 | A / H5N6 | Asia / Vietnam | |
| EPI_ISL_9572787 | A/Muscovy duck/Vietnam/HN6113/2020 | A / H5N6 | Asia / Vietnam | |
| EPI_ISL_9572792 | A/Muscovy duck/Vietnam/HN6120/2020 | A / H5N6 | Asia / Vietnam | |
| EPI_ISL_9572794 | A/Muscovy duck/Vietnam/HN6115/2020 | A / H5N6 | Asia / Vietnam | |
| EPI_ISL_9572800 | A/Muscovy duck/Vietnam/HN6119/2020 | A / H5N6 | Asia / Vietnam | |
| EPI_ISL_9572808 | A/Muscovy duck/Vietnam/HN6114/2020 | A / H5N6 | Asia / Vietnam | |
| EPI_ISL_9572812 | A/Muscovy duck/Vietnam/HN6111/2020 | A / H5N6 | Asia / Vietnam | |
| EPI_ISL_9572816 | A/duck/Vietnam/QN6519/2020 | A / H5N6 | Asia / Vietnam | |
| EPI_ISL_9594290 | A/chicken/Egypt/A19670/2021 | A / H5N8 | Africa / Egypt | |
| EPI_ISL_9594292 | A/chicken/Egypt/N15174C/2018 | A / H5N8 | Africa / Egypt | |
| EPI_ISL_9594293 | A/chicken/Egypt/N15175C/2018 | A / H5N8 | Africa / Egypt | |
| EPI_ISL_9594294 | A/chicken/Egypt/N15173A/2018 | A / H5N8 | Africa / Egypt | |
| EPI_ISL_9594297 | A/chicken/Egypt/N15178B/2018 | A / H5N8 | Africa / Egypt | |
| EPI_ISL_9594298 | A/chicken/Egypt/N15176A/2018 | A / H5N8 | Africa / Egypt | |
| EPI_ISL_9594299 | A/chicken/Egypt/N15168C/2018 | A / H5N8 | Africa / Egypt | |
| EPI_ISL_9594301 | A/chicken/Egypt/N15174A/2018 | A / H5N8 | Africa / Egypt | |
| EPI_ISL_9594302 | A/chicken/Egypt/F15100/2018 | A / H5N8 | Africa / Egypt | |
| EPI_ISL_9594303 | A/chicken/Egypt/N15173C/2018 | A / H5N8 | Africa / Egypt | |

| | | | | |
|---|---|---|---|---|
| EPI_ISL_9594305 | A/chicken/Egypt/N15173B/2018 | A / H5N8 | Africa / Egypt | |
| EPI_ISL_9594306 | A/chicken/Egypt/S18182C/2020 | A / H5N8 | Africa / Egypt | |
| EPI_ISL_9594307 | A/chicken/Egypt/N15178D/2018 | A / H5N8 | Africa / Egypt | |
| EPI_ISL_9594308 | A/chicken/Egypt/N15178C/2018 | A / H5N8 | Africa / Egypt | |
| EPI_ISL_9594309 | A/chicken/Egypt/N15178A/2018 | A / H5N8 | Africa / Egypt | |
| EPI_ISL_9594310 | A/chicken/Egypt/N15176B/2018 | A / H5N8 | Africa / Egypt | |
| EPI_ISL_9594311 | A/chicken/Egypt/N15169A/2018 | A / H5N8 | Africa / Egypt | |
| EPI_ISL_9594312 | A/chicken/Egypt/N15177C/2018 | A / H5N8 | Africa / Egypt | |
| EPI_ISL_9603767 | A/goose/Czech Republic/23458-2T/2021 | A / H5N1 | Europe / Czech Republic / Jihocesky Kraj / Okres Ceske Budejovice / Nové Hrady, Štipto? | Alexander Nagy (State Veterinary Institute Prague) |
| EPI_ISL_9603769 | A/goose/Czech Republic/23458-5T/2021 | A / H5N1 | Europe / Czech Republic / Jihocesky Kraj / Okres Ceske Budejovice / Nové Hrady, Štipto? | Alexander Nagy (State Veterinary Institute Prague) |
| EPI_ISL_9603775 | A/goose/Czech Republic/23458-4T/2021 | A / H5N1 | Europe / Czech Republic / Jihocesky Kraj / Okres Ceske Budejovice / Nové Hrady, Štipto? | Alexander Nagy (State Veterinary Institute Prague) |
| EPI_ISL_9603780 | A/goose/Czech Republic/23458-1K/2021 | A / H5N1 | Europe / Czech Republic / Jihocesky Kraj / Okres Ceske Budejovice / Nové Hrady, Štipto? | Alexander Nagy (State Veterinary Institute Prague) |
| EPI_ISL_9603795 | A/goose/Czech Republic/22608-1T/2021 | A / H5N1 | Europe / Czech Republic / Nové Hrady, By?ov | Alexander Nagy (State Veterinary Institute Prague) |
| EPI_ISL_9603819 | A/goose/Czech Republic/22608-2T/2021 | A / H5N1 | Europe / Czech Republic / Jihocesky Kraj / Okres Ceske Budejovice / Nové Hrady, By?ov | Alexander Nagy (State Veterinary Institute Prague) |
| EPI_ISL_9603901 | A/goose/Czech Republic/22608-3T/2021 | A / H5N1 | Europe / Czech Republic / Jihocesky Kraj / Okres Ceske Budejovice / Nové Hrady, By?ov | Alexander Nagy (State Veterinary Institute Prague) |
| EPI_ISL_9603916 | A/mute swan/Czech Republic/22477-1/2021 | A / H5N1 | Europe / Czech Republic / Zlinsky Kraj / Okres Kromeriz / Doubravicky pond, GPS: 49°17'17.233'' | Alexander Nagy (State Veterinary Institute Prague) |
| EPI_ISL_9603918 | A/mute swan/Czech Republic/22477-2/2021 | A / H5N1 | Europe / Czech Republic / Zlinsky Kraj / Okres Kromeriz / Doubravicky pond, GPS: 49°17'17.233'' | Alexander Nagy (State Veterinary Institute Prague) |
| EPI_ISL_9603920 | A/mute swan/Czech Republic/22380/2021 | A / H5N1 | Europe / Czech Republic / Kralovehradecky Kraj / Okres Jicin / Ostruzensky pond, GPS: 50°26'33, | Alexander Nagy (State Veterinary Institute Prague) |
| EPI_ISL_9603922 | A/mute swan/Czech Republic/22684/2021 | A / H5N1 | Europe / Czech Republic / Olomoucky Kraj / Okres Prerov / Na hrazi pond, GPS: 49,3582233N, 17 | Alexander Nagy (State Veterinary Institute Prague) |
| EPI_ISL_9603924 | A/goose/Czech Republic/22750/2021 | A / H5N1 | Europe / Czech Republic / Jihocesky Kraj / Okres Ceske Budejovice / Nové Hrady, Velebil pond | Alexander Nagy (State Veterinary Institute Prague) |
| EPI_ISL_9603927 | A/mute swan/Czech Republic/25702-2/2021 | A / H5N1 | Europe / Czech Republic / Olomoucky Kraj / Okres Prostejov / Víceme?ice | Alexander Nagy (State Veterinary Institute Prague) |
| EPI_ISL_9603929 | A/chicken/Czech Republic/25690/2021 | A / H5N1 | Europe / Czech Republic / Ustecky Kraj / Okres Litomerice / Libotenice, GPS: 50°28'8.285''N, 14° | Alexander Nagy (State Veterinary Institute Prague) |
| EPI_ISL_9603931 | A/pheasant/Czech Republic/25827-1/2021 | A / H5N1 | Europe / Czech Republic / South Moravian Region / Okres Hodonin / Jarohn?vice-Mist?ín | Alexander Nagy (State Veterinary Institute Prague) |
| EPI_ISL_9603937 | A/pheasant/Czech Republic/25827-2/2021 | A / H5N1 | Europe / Czech Republic / South Moravian Region / Okres Hodonin / Jarohn?vice-Mist?ín | Alexander Nagy (State Veterinary Institute Prague) |
| EPI_ISL_9603939 | A/pheasant/Czech Republic/25827-3/2021 | A / H5N1 | Europe / Czech Republic / South Moravian Region / Okres Hodonin / Jarohn?vice-Mist?ín | Alexander Nagy (State Veterinary Institute Prague) |
| EPI_ISL_9603943 | A/mute swan/Czech Republic/785/2022 | A / H5N1 | Europe / Czech Republic / Stredocesky Kraj / Okres Praha-Zapad / Davle, GPS: 49.8909144N, 14. | Alexander Nagy (State Veterinary Institute Prague) |
| EPI_ISL_9616210 | A/gadwall/Netherlands/22000307-001/2022 | A / H5N1 | Europe / Netherlands / Provincie Noord-Holland | Rene Heutink (Wageningen Bioveterinary Research) |
| EPI_ISL_9616212 | A/red knot/Netherlands/22000409-002/2022 | A / H5N1 | Europe / Netherlands / Provincie Noord-Holland | Rene Heutink (Wageningen Bioveterinary Research) |
| EPI_ISL_9616263 | A/brant goose/Netherlands/22000648-002/2022 | A / H5N1 | Europe / Netherlands / Provincie Zeeland | Rene Heutink (Wageningen Bioveterinary Research) |
| EPI_ISL_9616265 | A/red fox/Netherlands/22001901-005/2022 | A / H5N1 | Europe / Netherlands / Provincie Utrecht | Rene Heutink (Wageningen Bioveterinary Research) |
| EPI_ISL_9616266 | A/polecat/Netherlands/22002047-001/2022 | A / H5N1 | Europe / Netherlands / Provincie Gelderland | Rene Heutink (Wageningen Bioveterinary Research) |
| EPI_ISL_9616271 | A/red fox/Netherlands/22002190-003/2022 | A / H5N1 | Europe / Netherlands / Provincie Flevoland | Rene Heutink (Wageningen Bioveterinary Research) |
| EPI_ISL_9616272 | A/red fox/Netherlands/22002191-003/2022 | A / H5N1 | Europe / Netherlands / Provincie Utrecht | Rene Heutink (Wageningen Bioveterinary Research) |
| EPI_ISL_9690841 | A/great black-backed gull/Netherlands/22000090-002/2022 | A / H5N1 | Europe / Netherlands / South Holland | Rene Heutink (Wageningen Bioveterinary Research) |
| EPI_ISL_9690956 | A/common buzzard/Netherlands/22000192-001/2022 | A / H5N1 | Europe / Netherlands / South Holland | Rene Heutink (Wageningen Bioveterinary Research) |
| EPI_ISL_977513 | A/chicken/Czech Republic/1566-1/2021 | A / H5N8 | Europe / Czech Republic / Jihocesky Kraj / Okres Tabor / Dlouha Lhota/Chynov; GPS 49.3561703 | Alexander Nagy (State Veterinary Institute Prague) |
| EPI_ISL_977582 | A/duck/Northern China/ZGL/2020(H5N8) | A / H5N8 | Asia / China / Shandong Province | Jiahao Zhang (South China Agricultural University / College of Veterinary Medicine) |
| EPI_ISL_977595 | A/duck/Southwestern China/B1904/2020(H5N8) | A / H5N8 | Asia / China / Sichuan Province | Jiahao Zhang (South China Agricultural University / College of Veterinary Medicine) |
| EPI_ISL_977599 | A/duck/Northern China/LSP/2020(H5N8) | A / H5N8 | Asia / China / Shandong Province | Jiahao Zhang (South China Agricultural University / College of Veterinary Medicine) |
| EPI_ISL_978890 | A/chicken/Chiba/E1T/2021 | A / H5N8 | Asia / Japan / Chiba | Takehiko Saito (National Institute of Animal Health) |
| EPI_ISL_978891 | A/chicken/Chiba/E2T/2021 | A / H5N8 | Asia / Japan / Chiba | Takehiko Saito (National Institute of Animal Health) |
| EPI_ISL_978892 | A/chicken/Chiba/E3T/2021 | A / H5N8 | Asia / Japan / Chiba | Takehiko Saito (National Institute of Animal Health) |
| EPI_ISL_978893 | A/chicken/Chiba/E4T/2021 | A / H5N8 | Asia / Japan / Chiba | Takehiko Saito (National Institute of Animal Health) |
| EPI_ISL_978894 | A/chicken/Chiba/F1T/2021 | A / H5N8 | Asia / Japan / Chiba | Takehiko Saito (National Institute of Animal Health) |
| EPI_ISL_978895 | A/chicken/Chiba/F2T/2021 | A / H5N8 | Asia / Japan / Chiba | Takehiko Saito (National Institute of Animal Health) |
| EPI_ISL_978896 | A/chicken/Chiba/F3C/2021 | A / H5N8 | Asia / Japan / Chiba | Takehiko Saito (National Institute of Animal Health) |
| EPI_ISL_978897 | A/chicken/Chiba/F4T/2021 | A / H5N8 | Asia / Japan / Chiba | Takehiko Saito (National Institute of Animal Health) |
| EPI_ISL_978898 | A/chicken/Ibaraki/1C/2021 | A / H5N8 | Asia / Japan / Ibaraki | Takehiko Saito (National Institute of Animal Health) |
| EPI_ISL_978899 | A/chicken/Ibaraki/3T/2021 | A / H5N8 | Asia / Japan / Ibaraki | Takehiko Saito (National Institute of Animal Health) |
| EPI_ISL_978900 | A/chicken/Ibaraki/7C/2021 | A / H5N8 | Asia / Japan / Ibaraki | Takehiko Saito (National Institute of Animal Health) |
| EPI_ISL_978901 | A/chicken/Ibaraki/8T/2021 | A / H5N8 | Asia / Japan / Ibaraki | Takehiko Saito (National Institute of Animal Health) |
| EPI_ISL_978902 | A/chicken/Miyazaki/J12T/2021 | A / H5N8 | Asia / Japan / Miyazaki | Takehiko Saito (National Institute of Animal Health) |
| EPI_ISL_978903 | A/chicken/Miyazaki/J3T/2021 | A / H5N8 | Asia / Japan / Miyazaki | Takehiko Saito (National Institute of Animal Health) |
| EPI_ISL_978904 | A/chicken/Miyazaki/J4T/2021 | A / H5N8 | Asia / Japan / Miyazaki | Takehiko Saito (National Institute of Animal Health) |
| EPI_ISL_978905 | A/chicken/Miyazaki/J6T/2021 | A / H5N8 | Asia / Japan / Miyazaki | Takehiko Saito (National Institute of Animal Health) |
| EPI_ISL_978906 | A/chicken/Toyama/1T/2021 | A / H5N8 | Asia / Japan / Toyama | Takehiko Saito (National Institute of Animal Health) |
| EPI_ISL_978907 | A/chicken/Toyama/2T/2021 | A / H5N8 | Asia / Japan / Toyama | Takehiko Saito (National Institute of Animal Health) |
| EPI_ISL_978908 | A/chicken/Toyama/3T/2021 | A / H5N8 | Asia / Japan / Toyama | Takehiko Saito (National Institute of Animal Health) |
| EPI_ISL_978909 | A/chicken/Toyama/4T/2021 | A / H5N8 | Asia / Japan / Toyama | Takehiko Saito (National Institute of Animal Health) |
| EPI_ISL_978910 | A/duck/Chiba/C1T/2021 | A / H5N8 | Asia / Japan / Chiba | Takehiko Saito (National Institute of Animal Health) |
| EPI_ISL_978911 | A/duck/Chiba/C2T/2021 | A / H5N8 | Asia / Japan / Chiba | Takehiko Saito (National Institute of Animal Health) |
| EPI_ISL_978912 | A/duck/Chiba/C3T/2021 | A / H5N8 | Asia / Japan / Chiba | Takehiko Saito (National Institute of Animal Health) |
| EPI_ISL_978913 | A/duck/Chiba/C5T/2021 | A / H5N8 | Asia / Japan / Chiba | Takehiko Saito (National Institute of Animal Health) |
| EPI_ISL_978914 | A/duck/Chiba/D1-3T/2021 | A / H5N8 | Asia / Japan / Chiba | Takehiko Saito (National Institute of Animal Health) |
| EPI_ISL_978915 | A/mute swan/Ibaraki/080203C/2021 | A / H5N8 | Asia / Japan / Ibaraki | Takehiko Saito (National Institute of Animal Health) |
| EPI_ISL_978916 | A/mute swan/Ibaraki/080203T/2021 | A / H5N8 | Asia / Japan / Ibaraki | Takehiko Saito (National Institute of Animal Health) |
| EPI_ISL_980839 | A/mute swan/Czech Republic/1410-1/2021 | A / H5N8 | Europe / Czech Republic / Jihocesky Kraj / Okres Pisek / Zlivicky pond; GPS 49°21'26.626''N 14°6' | Alexander Nagy (State Veterinary Institute Prague) |
| EPI_ISL_984670 | A/barnacle goose/Denmark/14139-1/2020 | A / H5N8 | Europe / Denmark / Region Syddanmark / Tonder Kommune | Charlotte Kristiane Hjulsager (Statens Serum Institute / Microbiological Diagnostic and Virology) |
| EPI_ISL_984673 | A/barnacle goose/Denmark/14139-2/2020 | A / H5N8 | Europe / Denmark / Region Syddanmark / Tonder Kommune | Charlotte Kristiane Hjulsager (Statens Serum Institute / Microbiological Diagnostic and Virology) |
| EPI_ISL_984675 | A/barnacle goose/Denmark/14139-3/2020 | A / H5N8 | Europe / Denmark / Region Syddanmark / Tonder Kommune | Charlotte Kristiane Hjulsager (Statens Serum Institute / Microbiological Diagnostic and Virology) |
| EPI_ISL_984680 | A/black-headed gull/Denmark/14139-4/2020 | A / H5N8 | Europe / Denmark / Region Syddanmark / Tonder Kommune | Charlotte Kristiane Hjulsager (Statens Serum Institute / Microbiological Diagnostic and Virology) |
| EPI_ISL_984681 | A/barnacle goose/Denmark/14534-1/2020 | A / H5N8 | Europe / Denmark / Region Syddanmark / Tonder Kommune | Charlotte Kristiane Hjulsager (Statens Serum Institute / Microbiological Diagnostic and Virology) |

| | | | | |
|---|---|---|---|---|
| EPI_ISL_984684 | A/barnacle goose/Denmark/14537-1/2020 | A / H5N8 | Europe / Denmark / Region Syddanmark / Vejle Kommune | Charlotte Kristiane Hjulsager (Statens Serum Institute / Microbiological Diagnostic and Virology) |
| EPI_ISL_984686 | A/barnacle goose/Denmark/14536-1/2020 | A / H5N8 | Europe / Denmark / Region Syddanmark / Abenra Kommune | Charlotte Kristiane Hjulsager (Statens Serum Institute / Microbiological Diagnostic and Virology) |
| EPI_ISL_984688 | A/barnacle goose/Denmark/14538-1/2020 | A / H5N8 | Europe / Denmark / Region Syddanmark / Sonderborg Kommune | Charlotte Kristiane Hjulsager (Statens Serum Institute / Microbiological Diagnostic and Virology) |
| EPI_ISL_984690 | A/peregrine falcon/Denmark/14596-1/2020 | A / H5N8 | Europe / Denmark / Region Syddanmark / Svendborg Kommune | Charlotte Kristiane Hjulsager (Statens Serum Institute / Microbiological Diagnostic and Virology) |
| EPI_ISL_984691 | A/common buzzard/Denmark/14600-1/2020 | A / H5N8 | Europe / Denmark / Region Syddanmark / Abenra Kommune | Charlotte Kristiane Hjulsager (Statens Serum Institute / Microbiological Diagnostic and Virology) |
| EPI_ISL_984692 | A/barnacle goose/Denmark/14600-2/2020 | A / H5N8 | Europe / Denmark / Region Syddanmark / Abenra Kommune | Charlotte Kristiane Hjulsager (Statens Serum Institute / Microbiological Diagnostic and Virology) |
| EPI_ISL_984693 | A/barnacle goose/Denmark/14599-1/2020 | A / H5N8 | Europe / Denmark / Region Sjaland / Kalundborg Kommune | Charlotte Kristiane Hjulsager (Statens Serum Institute / Microbiological Diagnostic and Virology) |
| EPI_ISL_984694 | A/graylag goose/Denmark/14535-1/2020 | A / H5N8 | Europe / Denmark / Region Nordjylland / Alborg | Charlotte Kristiane Hjulsager (Statens Serum Institute / Microbiological Diagnostic and Virology) |
| EPI_ISL_984695 | A/chicken/Denmark/14819-6/2020 | A / H5N8 | Europe / Denmark / Region Midtjylland / Randers Kommune | Charlotte Kristiane Hjulsager (Statens Serum Institute / Microbiological Diagnostic and Virology) |
| EPI_ISL_985179 | A/duck/Korea/H338/2020 | A / H5N8 | Asia / Korea, Republic of / JB | Yu-Na Lee (Animal and Plant Quarantine Agency (S-2158) / Avian Influenza Research & Diagnostic Division) |
| EPI_ISL_985180 | A/chicken/Korea/H365/2020 | A / H5N8 | Asia / Korea, Republic of / GB | Yu-Na Lee (Animal and Plant Quarantine Agency (S-2158) / Avian Influenza Research & Diagnostic Division) |
| EPI_ISL_985181 | A/duck/Korea/H385/2020 | A / H5N8 | Asia / Korea, Republic of / JN | Yu-Na Lee (Animal and Plant Quarantine Agency (S-2158) / Avian Influenza Research & Diagnostic Division) |
| EPI_ISL_985182 | A/chicken/Korea/H390/2020 | A / H5N8 | Asia / Korea, Republic of / GG | Yu-Na Lee (Animal and Plant Quarantine Agency (S-2158) / Avian Influenza Research & Diagnostic Division) |
| EPI_ISL_985183 | A/quail/Korea/H394/2020 | A / H5N8 | Asia / Korea, Republic of / CB | Yu-Na Lee (Animal and Plant Quarantine Agency (S-2158) / Avian Influenza Research & Diagnostic Division) |
| EPI_ISL_985184 | A/duck/Korea/H411/2020 | A / H5N8 | Asia / Korea, Republic of / JN | Yu-Na Lee (Animal and Plant Quarantine Agency (S-2158) / Avian Influenza Research & Diagnostic Division) |
| EPI_ISL_985185 | A/quail/Korea/H412/2020 | A / H5N8 | Asia / Korea, Republic of / GG | Yu-Na Lee (Animal and Plant Quarantine Agency (S-2158) / Avian Influenza Research & Diagnostic Division) |
| EPI_ISL_985186 | A/duck/Korea/H419/2020 | A / H5N8 | Asia / Korea, Republic of / JN | Yu-Na Lee (Animal and Plant Quarantine Agency (S-2158) / Avian Influenza Research & Diagnostic Division) |
| EPI_ISL_985187 | A/duck/Korea/H431/2020 | A / H5N8 | Asia / Korea, Republic of / JN | Yu-Na Lee (Animal and Plant Quarantine Agency (S-2158) / Avian Influenza Research & Diagnostic Division) |
| EPI_ISL_985188 | A/duck/Korea/H432/2020 | A / H5N8 | Asia / Korea, Republic of / JN | Yu-Na Lee (Animal and Plant Quarantine Agency (S-2158) / Avian Influenza Research & Diagnostic Division) |
| EPI_ISL_985189 | A/duck/Korea/H438/2020 | A / H5N8 | Asia / Korea, Republic of / JN | Yu-Na Lee (Animal and Plant Quarantine Agency (S-2158) / Avian Influenza Research & Diagnostic Division) |
| EPI_ISL_985190 | A/duck/Korea/H439/2020 | A / H5N8 | Asia / Korea, Republic of / JN | Yu-Na Lee (Animal and Plant Quarantine Agency (S-2158) / Avian Influenza Research & Diagnostic Division) |
| EPI_ISL_985191 | A/chicken/Korea/H440/2020 | A / H5N8 | Asia / Korea, Republic of / GG | Yu-Na Lee (Animal and Plant Quarantine Agency (S-2158) / Avian Influenza Research & Diagnostic Division) |
| EPI_ISL_985192 | A/chicken/Korea/H441/2020 | A / H5N8 | Asia / Korea, Republic of / JB | Yu-Na Lee (Animal and Plant Quarantine Agency (S-2158) / Avian Influenza Research & Diagnostic Division) |
| EPI_ISL_985193 | A/chicken/Korea/H450/2020 | A / H5N8 | Asia / Korea, Republic of / GB | Yu-Na Lee (Animal and Plant Quarantine Agency (S-2158) / Avian Influenza Research & Diagnostic Division) |
| EPI_ISL_985194 | A/goose/Korea/H449/2020 | A / H5N8 | Asia / Korea, Republic of / CN | Yu-Na Lee (Animal and Plant Quarantine Agency (S-2158) / Avian Influenza Research & Diagnostic Division) |
| EPI_ISL_985195 | A/chicken/Korea/H470/2020 | A / H5N8 | Asia / Korea, Republic of / GG | Yu-Na Lee (Animal and Plant Quarantine Agency (S-2158) / Avian Influenza Research & Diagnostic Division) |
| EPI_ISL_985196 | A/duck/Korea/H471/2020 | A / H5N8 | Asia / Korea, Republic of / JN | Yu-Na Lee (Animal and Plant Quarantine Agency (S-2158) / Avian Influenza Research & Diagnostic Division) |
| EPI_ISL_985197 | A/duck/Korea/H499/2020 | A / H5N8 | Asia / Korea, Republic of / GG | Yu-Na Lee (Animal and Plant Quarantine Agency (S-2158) / Avian Influenza Research & Diagnostic Division) |
| EPI_ISL_985198 | A/chicken/Korea/H491/2020 | A / H5N8 | Asia / Korea, Republic of / GG | Yu-Na Lee (Animal and Plant Quarantine Agency (S-2158) / Avian Influenza Research & Diagnostic Division) |
| EPI_ISL_985199 | A/chicken/Korea/H510/2020 | A / H5N8 | Asia / Korea, Republic of / GG | Yu-Na Lee (Animal and Plant Quarantine Agency (S-2158) / Avian Influenza Research & Diagnostic Division) |
| EPI_ISL_985200 | A/duck/Korea/H509/2020 | A / H5N8 | Asia / Korea, Republic of / CB | Yu-Na Lee (Animal and Plant Quarantine Agency (S-2158) / Avian Influenza Research & Diagnostic Division) |
| EPI_ISL_985201 | A/duck/Korea/H511/2020 | A / H5N8 | Asia / Korea, Republic of / JB | Yu-Na Lee (Animal and Plant Quarantine Agency (S-2158) / Avian Influenza Research & Diagnostic Division) |
| EPI_ISL_985202 | A/duck/Korea/H514/2020 | A / H5N8 | Asia / Korea, Republic of / JB | Yu-Na Lee (Animal and Plant Quarantine Agency (S-2158) / Avian Influenza Research & Diagnostic Division) |
| EPI_ISL_985203 | A/duck/Korea/H515/2020 | A / H5N8 | Asia / Korea, Republic of / JN | Yu-Na Lee (Animal and Plant Quarantine Agency (S-2158) / Avian Influenza Research & Diagnostic Division) |
| EPI_ISL_985204 | A/duck/Korea/H516/2020 | A / H5N8 | Asia / Korea, Republic of / JN | Yu-Na Lee (Animal and Plant Quarantine Agency (S-2158) / Avian Influenza Research & Diagnostic Division) |
| EPI_ISL_985205 | A/duck/Korea/H524/2020 | A / H5N8 | Asia / Korea, Republic of / CN | Yu-Na Lee (Animal and Plant Quarantine Agency (S-2158) / Avian Influenza Research & Diagnostic Division) |
| EPI_ISL_985206 | A/chicken/Korea/H525/2020 | A / H5N8 | Asia / Korea, Republic of / CN | Yu-Na Lee (Animal and Plant Quarantine Agency (S-2158) / Avian Influenza Research & Diagnostic Division) |
| EPI_ISL_985207 | A/chicken/Korea/H526/2020 | A / H5N8 | Asia / Korea, Republic of / GB | Yu-Na Lee (Animal and Plant Quarantine Agency (S-2158) / Avian Influenza Research & Diagnostic Division) |
| EPI_ISL_985208 | A/duck/Korea/H528/2020 | A / H5N8 | Asia / Korea, Republic of / JB | Yu-Na Lee (Animal and Plant Quarantine Agency (S-2158) / Avian Influenza Research & Diagnostic Division) |
| EPI_ISL_985209 | A/chicken/Korea/H531/2020 | A / H5N8 | Asia / Korea, Republic of / GG | Yu-Na Lee (Animal and Plant Quarantine Agency (S-2158) / Avian Influenza Research & Diagnostic Division) |
| EPI_ISL_985210 | A/chicken/Korea/H532/2020 | A / H5N8 | Asia / Korea, Republic of / GG | Yu-Na Lee (Animal and Plant Quarantine Agency (S-2158) / Avian Influenza Research & Diagnostic Division) |
| EPI_ISL_985211 | A/duck/Korea/H538/2020 | A / H5N8 | Asia / Korea, Republic of / JB | Yu-Na Lee (Animal and Plant Quarantine Agency (S-2158) / Avian Influenza Research & Diagnostic Division) |
| EPI_ISL_985220 | A/chicken/Chiba/G1C/2021 | A / H5N8 | Asia / Japan / Chiba | Takehiko Saito (National Institute of Animal Health) |
| EPI_ISL_985221 | A/chicken/Chiba/G1T/2021 | A / H5N8 | Asia / Japan / Chiba | Takehiko Saito (National Institute of Animal Health) |
| EPI_ISL_985222 | A/chicken/Chiba/G2C/2021 | A / H5N8 | Asia / Japan / Chiba | Takehiko Saito (National Institute of Animal Health) |
| EPI_ISL_985223 | A/chicken/Chiba/G2T/2021 | A / H5N8 | Asia / Japan / Chiba | Takehiko Saito (National Institute of Animal Health) |
| EPI_ISL_985224 | A/chicken/Chiba/H1C/2021 | A / H5N8 | Asia / Japan / Chiba | Takehiko Saito (National Institute of Animal Health) |
| EPI_ISL_985225 | A/chicken/Chiba/H1T/2021 | A / H5N8 | Asia / Japan / Chiba | Takehiko Saito (National Institute of Animal Health) |
| EPI_ISL_985226 | A/chicken/Chiba/H2C/2021 | A / H5N8 | Asia / Japan / Chiba | Takehiko Saito (National Institute of Animal Health) |
| EPI_ISL_985227 | A/chicken/Chiba/H2T/2021 | A / H5N8 | Asia / Japan / Chiba | Takehiko Saito (National Institute of Animal Health) |
| EPI_ISL_985228 | A/chicken/Miyazaki/K1T/2021 | A / H5N8 | Asia / Japan / Miyazaki | Takehiko Saito (National Institute of Animal Health) |
| EPI_ISL_985229 | A/chicken/Miyazaki/K3T/2021 | A / H5N8 | Asia / Japan / Miyazaki | Takehiko Saito (National Institute of Animal Health) |
| EPI_ISL_985230 | A/chicken/Miyazaki/K5T/2021 | A / H5N8 | Asia / Japan / Miyazaki | Takehiko Saito (National Institute of Animal Health) |
| EPI_ISL_985231 | A/chicken/Miyazaki/K8T/2021 | A / H5N8 | Asia / Japan / Miyazaki | Takehiko Saito (National Institute of Animal Health) |
| EPI_ISL_985232 | A/mallard duck/Niigata/150209C/2021 | A / H5N8 | Asia / Japan / Niigata | Takehiko Saito (National Institute of Animal Health) |
| EPI_ISL_985233 | A/mallard duck/Niigata/150209T/2021 | A / H5N8 | Asia / Japan / Niigata | Takehiko Saito (National Institute of Animal Health) |
| EPI_ISL_9869760 | A/American wigeon/South Carolina/AH0195145/2021 | A / H5N1 | North America / United States / South Carolina / Colleton County | Mary Lea Killian (National Veterinary Services Laboratories - USDA / Diagnostic Virology Laboratory) |
| EPI_ISL_9876777 | A/American blue-winged teal/South Carolina/AH0195150/202 | A / H5N1 | North America / United States / South Carolina / Colleton County | Mary Lea Killian (National Veterinary Services Laboratories - USDA / Diagnostic Virology Laboratory) |
| EPI_ISL_9880015 | A/American wigeon/North Carolina/AH0182517/2022 | A / H5N1 | North America / United States / North Carolina / Hyde | Mary Lea Killian (National Veterinary Services Laboratories - USDA / Diagnostic Virology Laboratory) |
| EPI_ISL_9880019 | A/mallard/North Carolina/AH0182886/2022 | A / H5N1 | North America / United States / North Carolina / Pamlico | Mary Lea Killian (National Veterinary Services Laboratories - USDA / Diagnostic Virology Laboratory) |
| EPI_ISL_9880021 | A/northern pintail/North Carolina/AH0182892/2022 | A / H5N1 | North America / United States / North Carolina / Pamlico | Mary Lea Killian (National Veterinary Services Laboratories - USDA / Diagnostic Virology Laboratory) |
| EPI_ISL_9880151 | A/gadwall/North Carolina/AH0182894/2022 | A / H5N1 | North America / United States / North Carolina / Pamlico | Mary Lea Killian (National Veterinary Services Laboratories - USDA / Diagnostic Virology Laboratory) |
| EPI_ISL_9880152 | A/northern shoveler/North Carolina/AH0182913/2022 | A / H5N1 | North America / United States / North Carolina / Pamlico | Mary Lea Killian (National Veterinary Services Laboratories - USDA / Diagnostic Virology Laboratory) |
| EPI_ISL_9880153 | A/American wigeon/North Carolina/AH0182954/2022 | A / H5N1 | North America / United States / North Carolina / Pamlico | Mary Lea Killian (National Veterinary Services Laboratories - USDA / Diagnostic Virology Laboratory) |
| EPI_ISL_9909371 | A/turkey/Indiana/22-003707-003/2022 | A / H5N1 | North America / United States / Indiana / Jasper County / Dubois | Mary Lea Killian (National Veterinary Services Laboratories - USDA / Diagnostic Virology Laboratory) |
| EPI_ISL_995172 | A/chicken/Northern_Ireland/2020-17671_21VIR113-11/2020 | A / H5N8 | Europe / United Kingdom / Northern Ireland | Bianca Zecchin (Istituto Zooprofilattico Sperimentale Delle Venezie) |
| EPI_ISL_996003 | A/chicken/Northern_Ireland/2021-000067_21VIR114-19/2021 | A / H5N8 | Europe / United Kingdom / Northern Ireland | Bianca Zecchin (Istituto Zooprofilattico Sperimentale Delle Venezie) |
