## Supplemental Materials and Methods for "Detection and Phylogenetic Analysis of Highly Pathogenic A/H5N1 Avian Influenza Clade 2.3.4.4b Virus in Chile, 2022"

### Sample collection, screening, and sequencing

Fresh environmental samples were collected using sterile flocked swabs (Copan, Italy) and places in vials containing 1mL of universal transport media (Copa, Italy). Samples were stored at 4 °C during sampling and until transport to the laboratory at the Faculty of Veterinary Medicine of the University of Chile and stored at -80 °C until analysis. Nucleic acid extraction was performed using the MagMax-96 AI/ND kit (Applied Biosystems) following manufacturer's instructions. Influenzas A M-gene screening was performed according to protocol describes elsewhere (Sui et al) using the Fast Virus master mix (Thermofisher, USA), with cycle threshold of cutoff value of 38 on a Stratagene m3000p (Agilent technologies, USA) Real-Time PCR thermocycler. Host species were identified using specific cytochrome oxidase I primers as described [1] and Sanger sequenced at St. Jude's. Influenza A virus positive samples were sequenced at the St. Jude Children's Hospital Harwell Center using the Nextera XT DNA-seq library preparation kit on an Illumina MiSeq genome sequencer (Illumina, USA), following the centers protocols. Raw sequencing reads were processed using Trimmomatic [2] and *de-novo* assembled with the SPADes package [3]. After de novo assembly, only samples with all eight genetic segments were kept for following analysis. Nucleic acid sequences were annotated using the Influenza Virus Sequence Annotation Tool from NCBI [4]. Amino acid sequences were downloaded for the above samples. Sequences for each gene were separated and multi-alignments were done in CLC Genomics Workbench (v7.5.1, Qiagen, Germany). Each gene was examined for sequence motives of interest and sequences were recorded and summarized. Host species were identified using primers designed to amplify a segment of the mitochondrial cytochrome-oxidase I (COI) as described [5].

### Phylogenetic analysis

All H5Nx sequences were obtained from the GenBank and GISAID databases (assessed on 2023-01-22. Additionally, for the HA and NA genes, we added the top 100 BLAST matches that were closest to the Chilean H5N1 sequences. Duplicates (based on strain name), laboratory derived, mixed subtype, and low coverage (<90% of full length) sequences were excluded from downstream analysis when possible. Sequences were aligned with MAFFT v.7.490 [6], trimmed using trimAL[7] and phylogenetic trees were constructed using FastTree using the generalized time-reversible (GTR) models of nucleotide evolution (PMID: 20224823) and IQ-TREE v.2.0.3 [8]. with the best-fit nucleotide substitution model (TPM2u+F+I chosen according to Bayesian Information Criterion (BIC)). Trees were visualized and annotated using FigTree v.1.4.4 (http://tree.bio.ed.ac.uk/software/figtree/). Clade nomenclature was adopted from the WHO antigenic and genetic characteristics of zoonotic influenza A viruses and development of candidate vaccine viruses for pandemic preparedness (https://apps.who.int/iris/handle/10665/363866).

**References**


1. Bravo-Vasquez, N., et al., *Equine-like H3 avian influenza viruses in wild birds, Chile.* Emerging infectious diseases, 2020. **26**(12): p. 2887.
2. Bolger, A.M., M. Lohse, and B. Usadel, *Trimmomatic: a flexible trimmer for Illumina sequence data.* Bioinformatics, 2014. **30**(15): p. 2114-2120.
3. Prjibelski, A., et al., *Using SPAdes de novo assembler.* Current protocols in bioinformatics, 2020. **70**(1): p. e102.
4. Bao, Y., et al., *The influenza virus resource at the National Center for Biotechnology Information.* Journal of virology, 2008. **82**(2): p. 596-601.
5. Cheung, P.P., et al., *Identifying the species-origin of faecal droppings used for avian influenza virus surveillance in wild-birds.* Journal of clinical virology, 2009. **46**(1): p. 90-93.
6. Katoh, K. and D.M. Standley, *MAFFT multiple sequence alignment software version 7: improvements in performance and usability.* Molecular biology and evolution, 2013. **30**(4): p. 772-780.
7. Capella-Gutiérrez, S., J.M. Silla-Martínez, and T. Gabaldón, *trimAl: a tool for automated alignment trimming in large-scale phylogenetic analyses.* Bioinformatics, 2009. **25**(15): p. 1972-1973.
8. Minh, B.Q., et al., *IQ-TREE 2: new models and efficient methods for phylogenetic inference in the genomic era.* Molecular biology and evolution, 2020. **37**(5): p. 1530-1534.
